# A cell state specific metabolic vulnerability to GPX4-dependent ferroptosis in glioblastoma

**DOI:** 10.1101/2023.02.22.529581

**Authors:** Matei A. Banu, Athanassios Dovas, Michael G. Argenziano, Wenting Zhao, Henar Cuervo Grajal, Dominique M.O. Higgins, Colin P. Sperring, Brianna Pereira, Ling F. Ye, Aayushi Mahajan, Nelson Humala, Julia L. Furnari, Pavan S. Upadhyayula, Fereshteh Zandkarimi, Trang T. T. Nguyen, Peter B. Wu, Li Hai, Charles Karan, Aida Razavilar, Markus D. Siegelin, Jan Kitajewski, Jeffrey N. Bruce, Brent R. Stockwell, Peter A. Sims, Peter D. Canoll

## Abstract

Glioma cells hijack developmental transcriptional programs to control cell state. During neural development, lineage trajectories rely on specialized metabolic pathways. However, the link between tumor cell state and metabolic programs is poorly understood in glioma. Here we uncover a glioma cell state-specific metabolic liability that can be leveraged therapeutically. To model cell state diversity, we generated genetically engineered murine gliomas, induced by deletion of p53 alone (p53) or with constitutively active Notch signaling (N1IC), a pathway critical in controlling cellular fate. N1IC tumors harbored quiescent astrocyte-like transformed cell states while p53 tumors were predominantly comprised of proliferating progenitor-like cell states. N1IC cells exhibit distinct metabolic alterations, with mitochondrial uncoupling and increased ROS production rendering them more sensitive to inhibition of the lipid hydroperoxidase GPX4 and induction of ferroptosis. Importantly, treating patient-derived organotypic slices with a GPX4 inhibitor induced selective depletion of quiescent astrocyte-like glioma cell populations with similar metabolic profiles.

## INTRODUCTION

Intratumoral heterogeneity remains a central therapeutic hurdle in glioblastoma(Neftel et al., 2019; Wang et al., 2022; Yuan et al., 2018; Zhao et al., 2021). Previous studies have identified four major glioma cell states in GBM(Johnson et al., 2021; Liu et al., 2022; Neftel et al., 2019; Yuan et al., 2018). These glioma states differ in their resemblance to neural or glial lineages as well as their proliferative status(Neftel et al., 2019; Xie et al., 2022). To drive a particular state, glioma cells hijack neurodevelopmental transcriptional programs and master regulators(Neftel et al., 2019). We recently discovered that these glioma states also differ in their therapeutic vulnerabilities, with proliferative populations demonstrating sensitivity to mitotic poisons(Zhao et al., 2021). These findings are consistent with clinical experience showing that cycling cells are effectively targeted by standard chemotherapy and radiation(Barthel et al., 2019; Liau et al., 2017; Spinazzi et al., 2022). In contrast, quiescent populations with mesenchymal or astrocytic features are relatively insensitive to standard forms of treatment and, while abundant in primary GBM, are even more abundant in recurrent GBM(Artegiani et al., 2017; Chen et al., 2012; Wang et al., 2022; Xie et al., 2022). Importantly, a subset of these persister tumor cells reenter the cell cycle even under cytotoxic pressure and repopulate the proliferating cell pool(Oren et al., 2021; Xie et al., 2022). Quiescent tumor cells are therefore the likely culprit for resistance and tumor recurrence in GBM following standard of care treatment(Couturier et al., 2020; Hoang-Minh et al., 2018; Xie et al., 2022). Targeting specific vulnerabilities in the quiescent population can potentially delay tumor relapse and lead to durable therapeutic responses(Hangauer et al., 2017; Xie et al., 2022). Cell state-specific druggable targets remain, however, an unmet need in GBM.

Here, we explored the link between metabolic dependencies and glioma cell states to identify state specific druggable vulnerabilities. We hypothesized that individual cell states depend on highly specialized metabolic programs. As an experimental system to explore the metabolic dependencies and therapeutic vulnerabilities of glioma states we used a p53-deleted, PDGFR-B overexpressing genetic murine glioma model with or without Notch activation and analyzed tumor cell lineage trajectories at single cell resolution. Notch signaling is a master regulator of cell fate decisions during neural development. Without Notch overexpression, transformed progenitor cells maintained a neural-progenitor cell (NPC) phenotype with high proliferative capacity, whereas constitutive Notch activation induced a slow-cycling subpopulation of glioma cells with a more astrocyte-like phenotype. We leveraged this model to perform functional and metabolomic studies and identified state specific metabolic programs. NPC-like glioma cells relied on amino acid metabolism, glycolysis, and high mitochondrial oxygen consumption. Astrocyte-like glioma cells preferentially used lipid peroxidation and fatty acid oxidation, with alterations in mitochondrial metabolism and increased ROS production. Based on these programs, we then identified an astrocytic state that is selectively vulnerable to inhibition of the lipid hydroperoxidase GPX4 and ferroptosis, a regulated non-necroptotic form of cell death driven by iron dependent lipid peroxidation. High sensitivity to GPX4 inhibition of astrocyte-like glioma cells was directly linked to dysfunctional mitochondrial activity. Treating acute slices generated from patient derived surgical samples of GBM selectively targeted quiescent astrocyte-like glioma cells, highlighting the potential clinical significance of our findings.

## RESULTS

### Activated Notch induces a slow-growing astrocytic phenotype in a PDGFB/p53-/- murine glioma model

While genetically engineered retrovirally induced murine glioma models recapitulate the key histopathological features of human GBM, they do not recapitulate the diversity of glioma states, limiting their utility in pre-clinical studies of cell state specific druggable targets(Couturier et al., 2020; Weng et al., 2019). Notch activation in neural (NPC) or oligodendrocyte (OPC) progenitor cells represses progenitor programs, initiating and maintaining astrocytic differentiation(Benner et al., 2013; Dray et al., 2021; Engler et al., 2018; Wang et al., 2020; Zamboni et al., 2020). Given its critical role in regulating proliferation and lineage trajectories in the CNS, we hypothesized that Notch activation could have similar effects on glioma cell phenotype. To test this, we generated two genetic murine glioma models by injecting an HA tagged PDGFB-IRES-Cre expressing retrovirus to target progenitor cells in the subcortical white matter of p53^fl/fl^ mice(Eyler et al., 2020), with one of the models harboring a constitutively-active form of Notch1 in transformed cells at tumor inception (**Figure 1A**). Compared to the p53 control model, the N1IC model showed significantly longer survival, ranging from 29 to 151 days post-injection (dpi), and longer latency in tumor formation (**Figure 1B**), as confirmed by weekly bioluminescence imaging (**Figure 1C**). Serial MRI scans also demonstrated aggressive tumors in the p53 model at approximately four weeks post-injection while N1IC mice developed radiographically detectable tumors by 60 dpi at the earliest (**Figure S1A**). Histologically, both models exhibited high grade features at end stage (**Figure S1B)**, however the N1IC model showed significantly lower proliferation in the transformed and recruited populations, as measured by Ki67 labeling index (**Figure 1D**). Quiescent cell populations with high Notch activity have been identified in primary, treatment naïve GBMs(Liau et al., 2017). Notably, slow cycling persister cells driven by Notch signaling also emerge in GBM after prolonged antiproliferative drug exposure(Liau et al., 2017).

**Figure 1:**
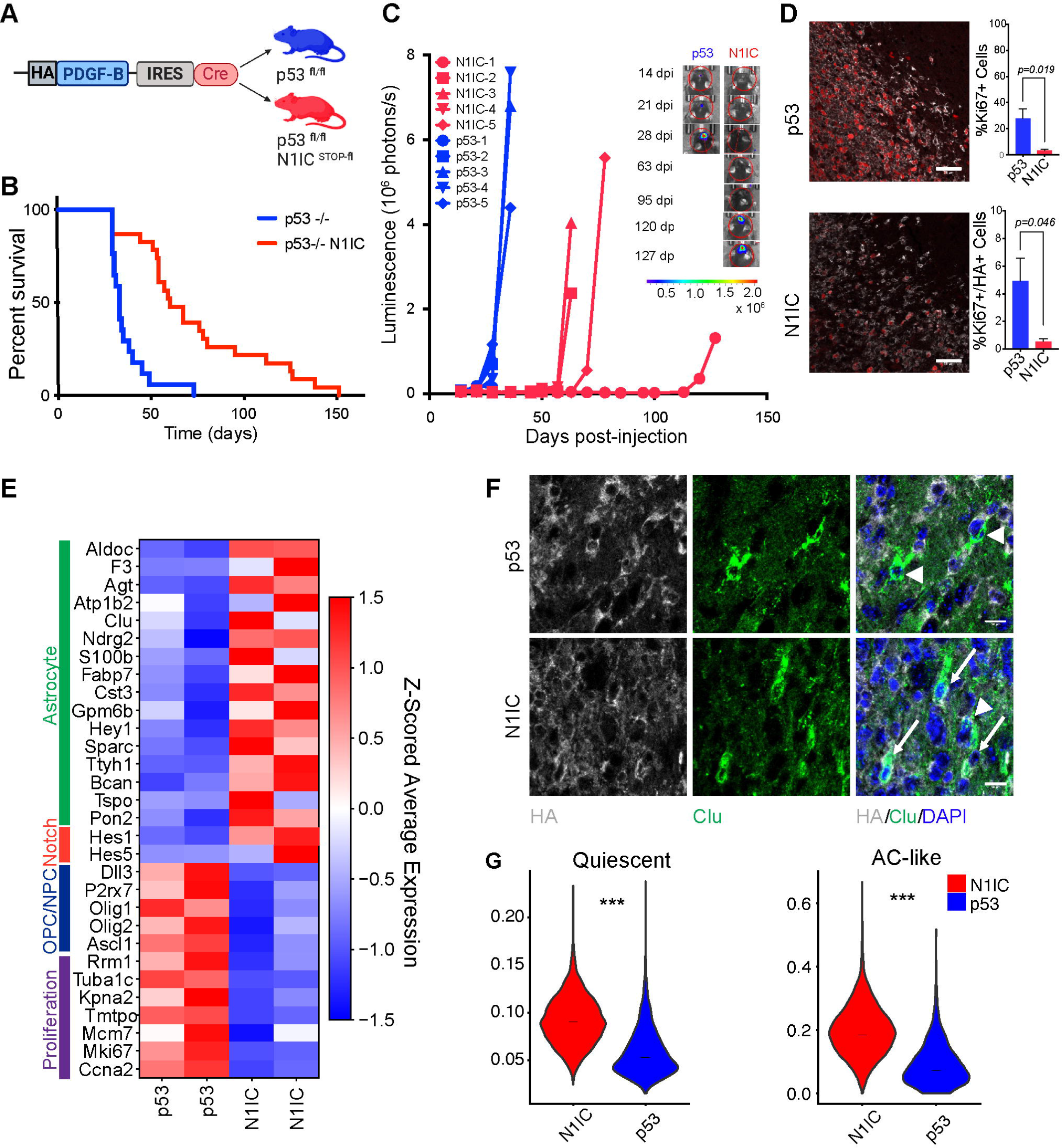
**Notch-driven murine glioma model of a quiescent astrocyte-like tumor cell state** A. Schematic depicting genetically engineered murine glioma models with or without Notch activation. PDGF-B IRES Cre retrovirus was injected in the corpus callosum of p53^fl/fl^ or p53^fl/fl^ N1IC^STOP-fl^ mice at 6 weeks. B. Survival curve for the two glioma models. Survival was significantly longer in p53 -/- N1IC mice compared to p53 -/- only mice (p<0.0001 by Mantel Cox Log rank test). Note the significant variability in survival for N1IC mice (29-151 days survival). C. Serial IVIS imaging demonstrating increased latency in signal detection and tumor formation in N1IC tumors compared to p53 tumors with subsequent sharp increase in bioluminescence and aggressive tumor growth. Inset demonstrates representative IVIS images for one mouse from each model. Also refer to **Supplementary Figure S1.** D. Representative immunofluorescence images of Ki67 and HA in N1IC and p53 endstage tumors. Quantification of Ki67+ cells and Ki67/HA double + cells demonstrating a decreased proliferative transformed population in N1IC tumors as well as an increased recruited proliferative population in p53 tumors. Bar graph shows mean proportions ± SEM. p=0.019 (%Ki67+ cells) and p=0.048 (%Ki67+/HA+ cells) by Welch’s t test, data pooled from n=6 p53/n=4 N1IC animals. Scale bar, 50 μm. E. Heatmap showing the expression of selected glioma cell state markers as well as Notch canonical downstream targets and proliferation genes in the p53 and N1IC transformed populations, as derived from scRNA-seq of the retrovirus induced tumors (n=2 from each model). Significantly differentially expressed genes are reported in **Table S1**. Also refer to **Supplementary Figure S1** and **Table S2** for direct N1ICD targets. F. Representative immunofluorescence of Clu/HA demonstrating double positive transformed astrocytic tumor cells only in the N1IC model (arrows) with presence of Clu+/HA- non-transformed tumor-associated astrocytes in both models (arrow heads). Scale bar, 10 μm. G. Violin plots of AUCell scores of the AC-like cell state and quiescence gene signatures in tumor cells from the two models. ***p<0.001 Welch’s t test

We next characterized the transcriptional states of the transformed tumor cells in the two models using scRNA-seq. N1IC glioma cells expressed higher levels of astrocytic markers while p53 transformed cells expressed higher levels of proliferation markers as well as NPC and OPC markers (**Figure 1E, Table S1**). Canonical Notch downstream targets were highly expressed in the transformed N1IC population, including *Hey1, Hes1* and *Hes5*. The inhibitory Notch ligand Dll3, critical in maintenance of undifferentiated neural progenitors(Zhao et al., 2009), was highly expressed in p53 tumor cells (**Figure 1E**). N1IC tumor cells activated Notch-dependent transcriptional programs regulating lineage identity in the persister cell population identified in human GBM (Liau et al., 2017) **(Figure S1C-D, Table S2**). Early NPC or OPC transcription factors were downregulated in N1IC tumors, including *Sox6, Olig1* and *Myt1*. In contrast, astrocytic master regulators such as *Id3* and *Runx3* were upregulated in the transformed N1IC population (**Figure 1E, S1D, Table S2**). Using immunofluorescence and taking advantage of the HA-tag that marks the transformed cells (Methods), we confirmed the presence of this Clu+/HA+ astrocytic glioma cell population in Notch tumors, while Clu was only seen in HA-tumor-associated, untransformed astrocytes in the p53 model (**Figure 1F)**. We lastly mapped out the landscape of tumor cell states based on proliferation status in the two models. While a significant proportion of the N1IC transformed tumor cells remained proliferative, the majority were quiescent and had AC-like features (**Figure 1G)**. Thus, even under constitutive Notch activation a subset of transformed glioma cells can re-enter the cell cycle and promote tumor growth, similar to subpopulations of persister cells identified in other cancer types(Oren et al., 2021). Based on these findings, we demonstrate that p53 and N1IC tumors are valid models to study diverse transcriptional tumor cell states in GBM.

### Multi-omic studies reveal cell state specific metabolic programs

In the CNS, lineage identity is tightly linked to specialized metabolic programs(Llorens-Bobadilla et al., 2015). Tumor cell populations with specific metabolic affinities and potentially unique therapeutic vulnerabilities have been recently identified in glioma(Garofano et al., 2021; Hoang-Minh et al., 2018; Rusu et al., 2019), but correlations between transcriptional cell state and metabolism have not been studied. We probed this potential link and first performed gene ontology analysis using the scRNA-seq data of the transformed cells from the two murine models. This revealed profound differences in metabolic programs at the transcriptional level, with N1IC glioma cells showing significant enrichment in genes associated with mitochondrial metabolism (electron transport chain/ETC, oxidative phosphorylation, tricyclic acid cycle/TCA), lipid catabolism and fatty acid β-oxidation (FAO). Notably, N1IC cells were also enriched in programs involved in oxidative stress responses, including taurine/hypotaurine metabolism and glutathione peroxidase activity. In contrast, p53 tumor cells demonstrated enrichment in amino acid metabolism (seleno-aminoacids, alanine, aspartate, glutamate) and nitrogen catabolism (**Figure 2A, Table S3**).

**Figure 2:**
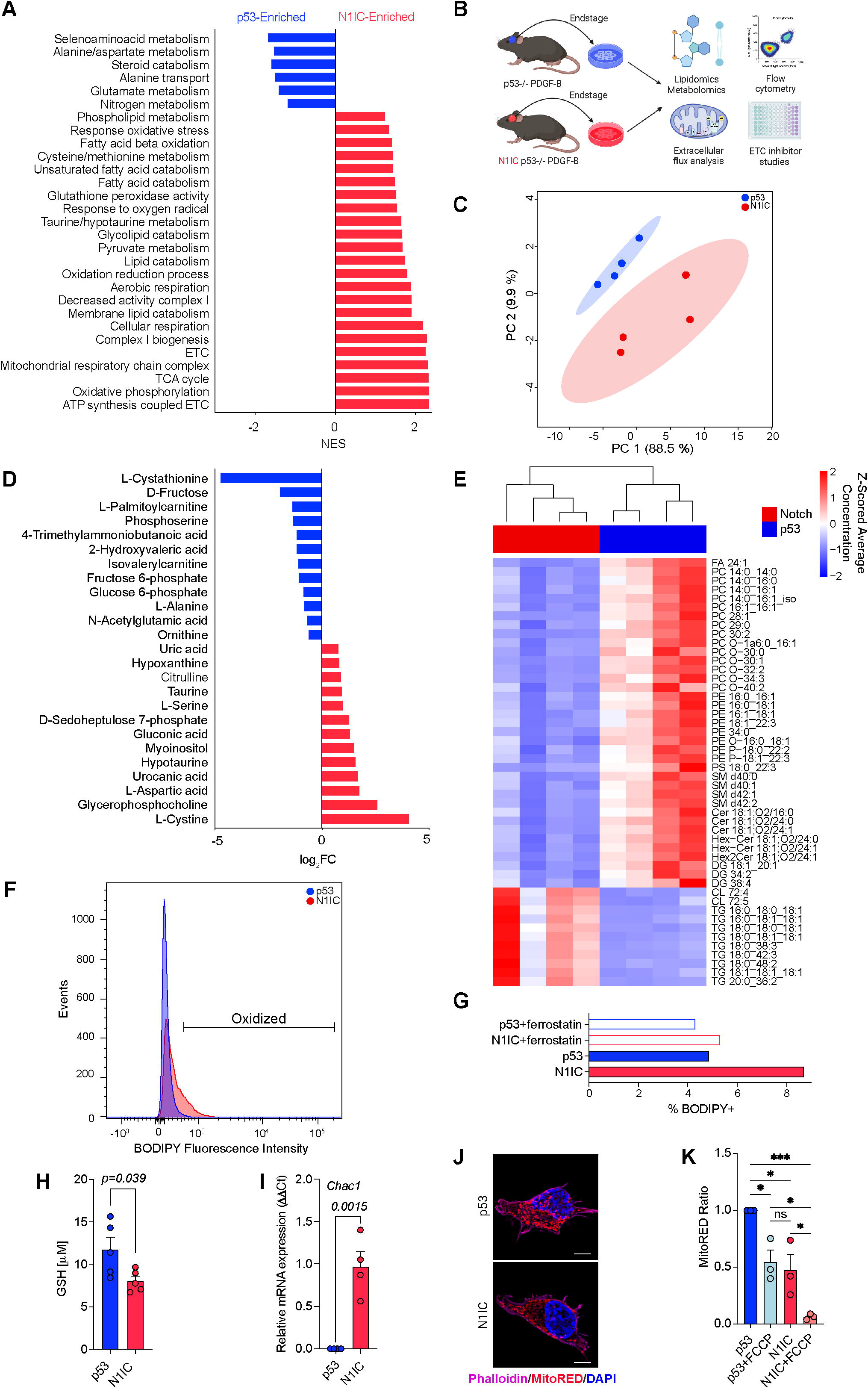
Multi-omic analysis reveals differences in metabolic programs between N1IC and p53 cells. A. Bar graph depicting significant metabolic gene ontologies in the two models via GSEA. NES – normalized enrichment score. Full list of transcriptional metabolic programs provided in **Table S3**. B. Schematic diagram depicting workflow for multi-omic analysis including functional studies, metabolic and lipidomic LC-MS analysis of cell lines isolated from the two models. Also refer to **Supplementary Figure S2** for further characterization of isolated cell lines. C. PCA analysis based on untargeted metabolomics performed on p53 and N1IC cells lines demonstrating separation of models based on metabolic profile. D. Bar graph demonstrating log_2_FC of significant differentially enriched metabolites in N1IC and p53 cells. P-values provided in **Table S4**. Also refer to **Supplementary Figure S3** for in depth description of N1IC metabolic pathways based on integrated transcriptomic and metabolomic data. E. Heatmap of differentially enriched lipid species in p53 and N1IC cells in both the positive and negative mode. Scale bar represents Z-scored average concentration of distinct lipid species normalized by protein concentration. FDR-corrected P-values and lipid ontology analysis provided in **Table S5** and **Supplementary Figure S3**. F. Flow cytometry of BODIPY-C11 fluorescence demonstrating higher lipid peroxidation in N1IC vs. p53 cells at baseline. Representative experiment of n=4 independent experiments. G. Percent BODIPY+ cells in p53 and N1IC cells and effects of ferroptosis inhibitor Ferrostatin-1 in reversing baseline lipid peroxidation. Representative experiment of n=3 independent experiments. H. Quantification of GSH levels in the same pair of p53 and N1IC cells by GSH fluorometric assay. Data pooled from n=4 independent experiments, p=0.039 by unpaired two tailed t-test. I. PCR of Chac1, ferroptosis marker involved in GSH degradation and Notch receptor inhibition, demonstrating higher expression in a N1IC cell line compared to a p53 cell line. Data pooled from n=3 independent experiments, p=0.0184 by unpaired two tailed t-test. J. Representative images acquired via SIM of MitoRED stained p53 and N1IC cells demonstrating differences in mitochondrial staining (red). DAPI stain (blue) labels nuclei and Phalloidin stain (magenta) labels filamentous actin. Scale bar, 5 μm. K. Bar plot showing mean ± SEM MitoRED ratios, data pooled from n=3 independent experiments, untreated p53 cells used as reference. p values calculated by Kruskal-Wallis test with Dunn’s multiple comparison test. *p < 0.05; **p < 0.01; ***p < 0.001. Also refer to **Supplementary Figures S4** and **S5** for characterization of the ETC and basal energetic metabolism in the two models.

To further dissect state-specific functional and metabolic differences, we next isolated primary cell cultures from end-stage tumors (**Figure 2B**). N1IC tumor cells retained the N1IC transgene and showed increased expression of Notch downstream targets Hey1 and Hes5, similar to the transformed population in the retroviral N1IC murine model (**Figure S2A-C**). The Notch1 intracellular domain (N1ICD) was highly expressed and exhibited nuclear translocation in cells isolated from the Notch model (**Figure S2C**). We performed liquid chromatography-mass spectrometry (LC-MS/MS) untargeted metabolic and lipidomic profiling on these primary cell cultures from the two models (**Figure 2 C-E**). Principal component analysis (PCA) of 203 quantified metabolites clearly separated the p53 and N1IC cell lines, with 25 metabolites differentially abundant between the two populations (**Figure 2C-D, Table S4**). Notably, metabolic ontology analysis (Methods) in N1IC cells revealed enrichment in oxidative stress response and redox balance pathways, such as cysteine/methionine metabolism (L-serine, L-cystine) and taurine/hypotaurine metabolism (**Figure 2D, S3A, Table S4**). N1IC cells were enriched in D-seduheptulose 7-phosphate and gluconic acid, metabolites in the pentose phosphate pathway, a pathway highly active in astrocytes driving NADPH regeneration for reductive recycling of glutathione (GSH) under oxidative stress(Vicente-Gutierrez et al., 2019) (**Figure 2D, S3A**). Joint transcriptomic-metabolomic pathway analysis in the N1IC model (Methods) revealed high mitochondrial activity (TCA cycle, pyruvate metabolism, oxidative phosphorylation) and ineffective cysteine/methionine metabolism, with degradation of cysteine to taurine/hypotaurine via upregulated cysteine sulfinic acid decarboxylase (*Csad*) instead of utilization for GSH synthesis (**Figure S3B-C, Table S4**). In contrast, p53 cells were enriched in glycolysis and galactose pathway metabolites (D-fructose, glucose 6-posphate, fructose 6-phosphate), programs shown to play a major role in the energy balance of rapidly proliferating glioma cells(Hoang-Minh et al., 2018). Furthermore, p53 cells had high levels of the L-cystathionine metabolite and increased expression of glutamate-cysteine ligase gene (*Gcl*), both involved in GSH synthesis via the transsulfuration pathway. In line with the transcriptomic ontologies, p53 cells demonstrated high levels of L-alanine, which has been shown to inhibit glial fate of NPCs in the CNS(Radu et al., 2019) (**Figure 2A, D, S3A**).

Lipid profiling also revealed differences in the two models, with p53 cells showing a higher abundance and variety of lipid species compared to the N1IC cells (**Figure 2E)**. Eleven lipid types were differentially enriched in N1IC cells compared to 37 species in p53 cells. The lipid profile segregated by model, in both the negative and positive mode (**Figure S3D**). Notably, p53 cells had higher levels of multiple functionally important lipid groups, including glycerophosphocholines (PC), glycerophosphoethanolamines (PE), ceramides (Cer) and sphingolipids (SM) which are important in stabilizing the membrane lipid bilayer (**Figure 2E, S3D-E, Table S5**). In contrast, the most abundant lipid species in N1IC cells were triacylglycerols (TG), with depletion of most other lipid species containing fatty acids. Interestingly, the only differentially abundant polyunsaturated fatty acid (PUFAs) species were in the p53 cell lines. Based on our transcriptomic and lipidomic data, we hypothesized that depletion of PUFAs is secondary to baseline lipid peroxidation in N1IC cells. To verify this, we used the fluorescent dye BODIPY-C11 and quantified baseline lipid peroxides in the two cell types via flow cytometry (**Figure 2F-G**). N1IC cells had an 8.7% oxidized fraction compared to only 4.8% in p53 cells. When using Ferrostatin-1, a radical scavenger and ferroptosis inhibitor, generation of lipid peroxides in N1IC cells was reduced close to baseline p53 levels (**Figure 2G**). Ferrostatin-1 had no effect on lipid peroxide generation in p53 cells. Furthermore, N1IC cells had lower levels of the antioxidant glutathione in its reduced form, and higher baseline levels of the ferroptosis marker *Chac1* (**Figure 2H-I**), suggesting that N1IC cells may be undergoing baseline low-level ferroptosis, a form of programed cell death based on iron-dependent oxidation of phospholipids with polyunsaturated fatty acyl tails.

Given the high redox imbalance noted in N1IC cells, we next examined the role of the electron transport chain (ETC) in the two cell models. High FAO and TCA cycle activity noted in N1IC cells will lead to increased oxidative phosphorylation (**Figure S4A**). Notably, inhibition of complex I (IACS-010759), complex III (Antimycin) or ATP synthase (Oligomycin) in the ETC led to variable effects on cell viability for p53 and N1IC cells, 80.6 vs. 77.4%, 30.8 vs. 31.9% and 57.6 vs. 64% respectively, but without significant differences in sensitivity between the two models (p=0.449, 0.61 and 0.122 respectively) (**Figure S4B-D**). However, p53 cells were significantly more sensitive to uncoupling of oxidative phosphorylation with the ionophore carbonyl cyanide-p-trifluoro-methoxyphenylhydrazone FCCP (47.3 vs. 59% cell viability, p=0.0005) (**Figure S4E**). Lipid-peroxide-induced mitochondrial damage can lead to uncoupling of oxidative phosphorylation in N1IC cells. Furthermore, translocator protein (Tspo), highly upregulated in the N1IC model (**Figure 1E**, **Table S1**), regulates mitochondrial respiration and generation of ROS. Indeed, when measuring baseline mitochondrial membrane potential ((ΔΨ_m_) in the two models using MitoRED mitochondrial dye, we found significant membrane depolarization and less dye uptake in N1IC cells. Addition of the uncoupling agent FCCP decreased MitoRED fluorescence in p53 cells to baseline N1IC levels, and even further reduced MitoRED fluorescence of N1IC cells (**Figure 2J-K**). This demonstrates that N1IC cells have lower baseline ΔΨ_m_ secondary to mitochondrial uncoupling. Even low levels of uncoupling can have dramatic metabolic effects, including induction of lipolysis and modulation of ROS production(Demine et al., 2019). Indeed, mirroring normal astrocytic metabolism(Vicente-Gutierrez et al., 2019), N1IC cells demonstrated significantly higher total ROS production compared to p53 cells (**Figure S4F**). Together, these findings suggest that mitochondrial dysfunction in N1IC cells leads to profound redox imbalance and increased generation of ROS.

Uncoupling profoundly impacts ATP production and cellular energetic metabolism. We therefore compared basal energy metabolism and mitochondrial function in the two glioma cell models. First, we measured oxygen consumption rate (OCR) and found significantly higher basal and maximal respiration as well as spare respiratory capacity in the p53 cells (**Figure S5A-C**). Notably, there were no significant differences between models in non-mitochondrial OCR. Therefore, p53 cells appear to have more efficient mitochondrial metabolism with higher OCR and increased mitochondrial adaptability. Seahorse extracellular acidification rate (ECAR) analysis suggested significantly higher glucose usage in the rapidly proliferating p53 NPC-like cell state, in line with recent studies (**Figure S5B**)(Garofano et al., 2021; Hoang-Minh et al., 2018). This is further supported by enrichment in glycolytic metabolites glucose 6-P and fructose 6-P in p53 cells (**Figure 2D**). The slow-cycling N1IC cells had decreased energetic requirements and lower ATP production (**Figure S5C)**. ROS, abundant and poorly buffered in N1IC cells, can lead to mitochondrial membrane damage and dysfunction. We used cysteine and methionine deprivation to mimic inefficient GSH synthesis noted in N1IC cells and observed significant decrease in p53 maximal respiration but minimal effect on basal OCR. Intriguingly, there were no effects of cysteine/methionine deprivation on energy metabolism in the Notch model (**Figure S5D-E**).

### Mitochondrial dependent differential sensitivity to Gpx4 inhibition and ferroptosis of the astrocyte-like N1IC cell state

We next investigated if these metabolic differences can be leveraged to design cell state-targeted therapies. As detailed above, N1IC cells show uncoupling of oxidative phosphorylation, increased production of ROS, and elevated levels of lipid peroxidation (**Figure 2**). Furthermore, high ROS levels and ineffective cysteine metabolism can lead to depletion of glutathione stores (**Figure 2H, S3C**). The selenocysteine enzyme glutathione peroxidase 4 (GPX4), using GSH as a cofactor, plays a central role in protecting cells from oxidative stress, particularly under thiol deprivation conditions(Dixon et al., 2012; Jiang et al., 2021). Given these metabolic differences, we compared the sensitivity of N1IC and p53 cells to induction of ferroptosis with a pharmacologic inhibitor of GPX4 (RSL3). *In vitro* viability assays showed that N1IC cell lines were preferentially sensitive to RSL3 compared to p53 cell lines (**Figure 3A**). We investigated if this differential sensitivity was secondary to ferroptosis or off-target effects of RSL3 using three independent modalities. First, cell death induced by GPX4 inhibition in N1IC cells was significantly reduced, although not completely abrogated, with addition of the ferroptosis rescue drug Ferrostatin-1 (**Figure S6A**). Ferrostatin-1 decreased efficacy of GPX4 inhibition in both N1IC and p53 cell lines (**Figure 3B**). In contrast, addition of the apoptosis inhibitor Z-VAD-FMK or the necroptosis inhibitor Necrostatin-1 did not significantly modify sensitivity to RSL3 in the N1IC cell line (**Figure S6A)**. Notably, treatment of a N1IC cell line with Ferrostatin-1 in the absence of RSL3 did not significantly affect cell viability, suggesting that low level baseline lipid peroxidation and ferroptosis encountered in this cell state do not lead to significant levels of cell death. As expected, Necrostatin-1 and Z-VAD-FMK alone did not affect cell viability (**Figure S6B**). Second, in both cell types RSL3 induced upregulation of canonical ferroptosis markers at a transcriptional level, including glutathione-specific γ-glutamylcyclotransferase 1 (*Chac1*), prostaglandin-endoperoxide synthase 2 (*Ptgs2*) and the system x_c_^-^ antiporter (*Slc7a11)* (**Figure 3C**). Importantly, all three markers are upregulated in vehicle-treated N1IC cells compared to p53 cells, further supporting low level baseline ferroptosis in the AC-like cell state (**Figure 3C**). Notably, *Chac1* catalyzes the cleavage of glutathione, likely leading to further depletion of glutathione stores in N1IC cells (**Figure S3C**). All three markers are significantly higher in RSL3-treated N1IC cells, demonstrating a higher ferroptotic response in the AC-like cell state. Third, exposure of N1IC cells to RSL3 led to a significant increase in BODIPY-C11 fluorescence, a marker of increased lipid peroxidation secondary to ferroptosis, starting at approximately 10 minutes after drug treatment (**Figure S7C**). Taken together, these results demonstrate that in both p53 and N1IC cell lines RSL3 inhibition of GPX4 induces ferroptosis, with the N1IC cells showing significantly higher sensitivity to ferroptosis.

**Figure 3:**
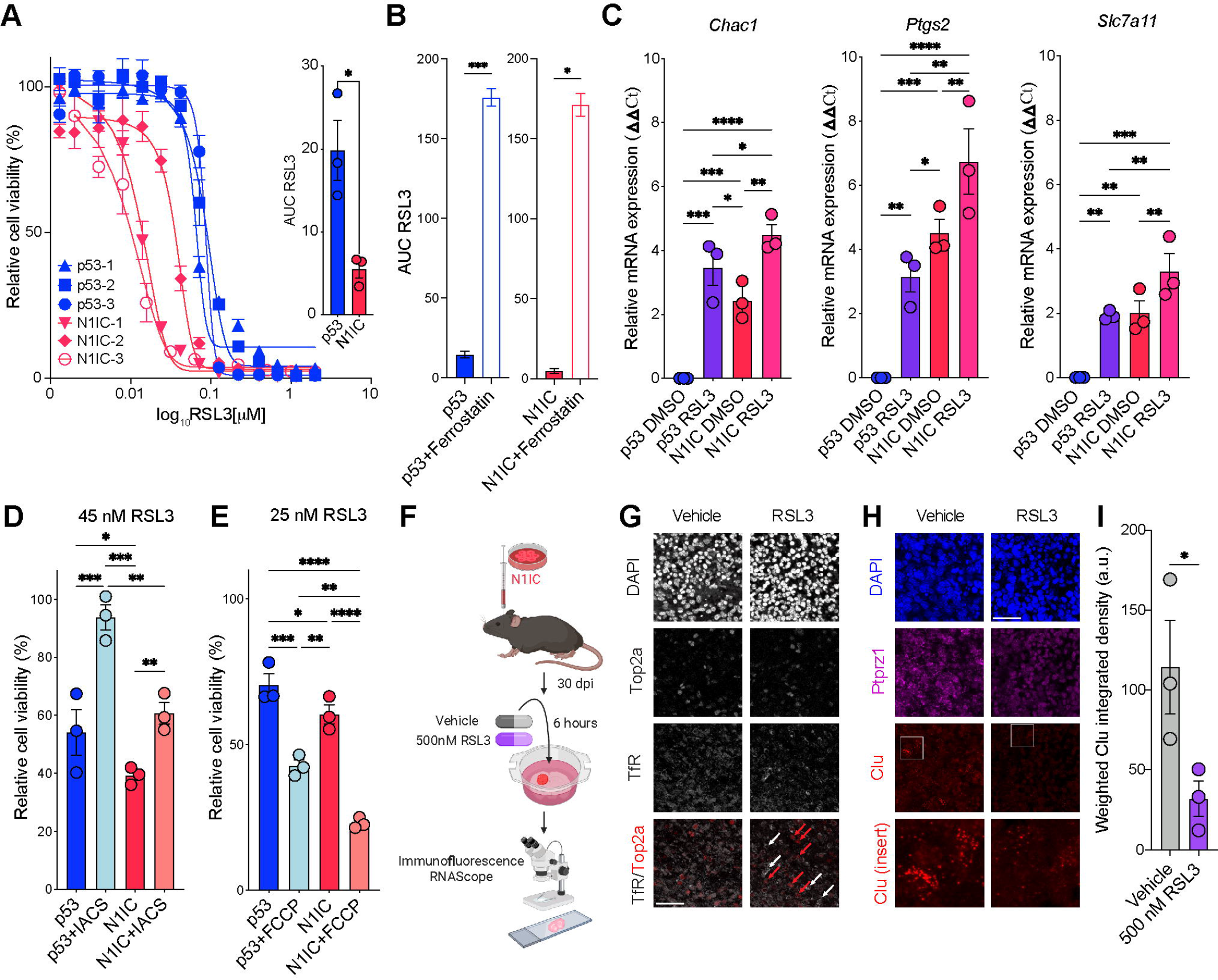
Mitochondrial dependent differential sensitivity to Gpx4 inhibition and ferroptosis of the quiescent AC-like cell state. A. Representative RSL3 drug screen on n=3 p53 and N1IC independent cell lines, error bars represent SEM from three technical replicates. Inset: Bar graph depicting mean area under the curve (AUC) ± SEM for panel A, comparison by Welch’s t test. *p< 0.05 B. AUC from RSL3 dose response curves with and without Ferrostatin-1 in n=3 p53 cell lines and n=2 N1IC cell lines. Also refer to Supplementary Figure S6. C. PCR ΔΔCT values of ferroptosis markers in p53 and N1IC treated vs. untreated cell lines, data pooled from n=3 independent experiments. Normalized to actin, p53 DMSO used as reference. D. Combined drug screen using RSL3 and complex I inhibitor IACS-010759 demonstrating increased resistance to RSL3 with complete inhibition of the ETC and oxidative phosphorylation. 45 nM RSL3, 2 μM IACS, normalized to IACS. n=3 technical replicates. Representative experiment from n=3 independent experiments. E. Combined drug screen using RSL3 and mitochondrial uncoupling agent FCCP demonstrating increased sensitivity to RSL3 with uncoupling of oxidative phosphorylation. 25 nM RSL3, 2 μM FCCP, normalized to FCCP. n=3 technical replicates. Representative experiment from n=4 independent experiments. C, D, E: P-values calculated by two-way ANOVA correcting for multiple comparisons by controlling the FDR using the 2-stage linear step-up procedure of Benjamini, Krieger, and Yekutieli. *p < 0.05; **p < 0.01; ***p < 0.001, ****p < 0.0001. F. Schematic representation of experimental setup of murine organotypic slice cultures. 3 slices per condition (technical replicates) were generated from 3 different tumor bearing mice (biological replicates). Mice were sacrificed at 30 dpi. Treatment with 500 nM RSL3 or DMSO vehicle for 6 hours. G. Double immunofluorescence of proliferation marker Top2a and ferroptosis marker transferrin receptor (TfR) demonstrating lack of TfR staining in vehicle treated slices and upregulation of TfR after 500 nM RSL3 in Top2a negative non-proliferating cells. Red arrows mark Top2a+ cells, white arrows marking TfR+ cells. Scale bar, 50 μm. H. RNAScope of Ptprz (tumor marker - magenta) and Clu (astrocytic marker – red) after 500 nM RSL3 or DMSO vehicle demonstrating depletion of the Clu+ transformed cell population. Scale bar, 50 μm, insets 40x40 μm. I. Quantification of weighted integrated density on n=3 independent slice cultures. Bar graph depicting mean ± SEM, p=0.0185 by paired t-test.

We sought to dissect mechanisms of differential sensitivity to ferroptosis in the two models. First, we asked if increased ferroptosis sensitivity in N1IC cells is specifically linked to GPX4 dependency. N1IC cell lines demonstrated significantly higher sensitivity and lower area under the curve (AUC) for both ML162, a chloroacetamide, and ML210, a nitroisoxazole (**Figure S6D, F**). ML162 and ML210 are GPX4 inhibitors with substantially different chemical structures, and, therefore, distinct off-target effects. Thus, differential sensitivity is directly linked to degree of GPX4 dependency. Importantly, inhibition of GPX4 with ML162 and ML210 induced ferroptosis, which was rescued with addition of Ferrostatin-1 (**Figure S7E, G**). Intriguingly, both cell models were resistant to IKE, a potent system x_c_^-/^Slc7a11 inhibitor and ferroptosis inducer. Doses of up to 20 mM were insufficient to achieve IC50 and addition of Ferrostatin-1 did not significantly alter AUC (**Figure S7H-I**). Therefore, inhibition of Slc7a11 did not induce ferroptosis in either N1IC or p53 cell lines, further demonstrating the specific sensitivity to inhibition of GPX4, particularly in the N1IC model. Mitochondria generate most of the energy and ROS in cells, playing a central role in programmed cell death. Therefore, we also investigated the effects of disrupting mitochondrial function on sensitivity to GPX4 inhibitors in the two models. Blocking complex I activity with the specific inhibitor IACS-010759 significantly increased resistance to RSL3 in both cell models, with higher impact in the p53 cell line (**Figure 3D**). In contrast, inducing uncoupling of the ETC with FCCP increased response to RSL3 in both cell models, and more so in N1IC cells (**Figure 3E**). This may be explained by the central role of complex I in generating ROS. Taken together, this data demonstrates increased sensitivity to GPX4-driven ferroptosis in the AC-like cell state and an important role for the ETC driven ROS production in modulating GPX4 dependency.

Tumor microenvironment and intercellular interactions can significantly impact ferroptosis(Wu et al., 2019). To test the effects of ferroptosis on N1IC tumor cells in the context of the complex cellular architecture of gliomas, we generated acute slice cultures from N1IC cell-transplanted murine tumors at 30 dpi, treated the slices with either vehicle or RSL3 and performed immunofluorescence and RNAscope studies (**Figure 3F**). GPX4-driven ferroptosis specifically targeted the non-cycling tumor cell states, with upregulation of the canonical ferroptosis marker TfR(Feng et al., 2020) solely in Top2a negative cells (**Figure 3G**). Furthermore, RSL3 specifically targeted the AC-like transformed population with significant depletion of Ptprz+ Clu+ cells after 6 hours of 500 nM RSL3 (**Figure 3H**). Overall, these findings support a cell state specific metabolic vulnerability to GPX4 inhibition and ferroptosis in quiescent AC-like glioma cell populations.

### RSL3 targets quiescent astrocyte-like transformed cell populations in acute slice cultures from human glioma

To test the clinical significance of our models, we next investigated the presence of N1IC-like tumor cell populations in human gliomas. We derived signatures from differentially up and downregulated genes in the N1IC tumor cell population and analyzed scRNA-seq profiles from IDH1-wild type GBM samples in recently published datasets (Methods, **Figure S7**)(Yuan et al., 2018). Quiescent tumor cell populations, predominantly adopting the AC-like cell state, were present in all eight patient samples (**Figure S7A-B**). We compiled Spearman correlations between glioma cell states(Neftel et al., 2019) and the two N1IC gene signatures (Methods). The “N1IC_up” signature most closely correlated with the AC and MES1/2 gene signatures, while the “N1IC_down” signature clustered with highly proliferative NPC-like states (**Figure S7C**). Notably, in a cohort of five patients undergoing chronic intratumoral delivery of topotecan, a mitotic poison, post-treatment tissue analysis revealed significant enrichment in the “N1IC_up” gene signature(Spinazzi et al., 2022) (**Figure S7D**). These findings demonstrate that the N1IC murine model recapitulates tumor cell populations present in treatment naïve GBM as well as populations expanded after exposure to chronic cytotoxic therapies. To explore cell-state specific metabolic programs in human glioma, we also calculated the Spearman correlation coefficient between metabolic pathways and cell state signatures using the transcriptomic data from eight human GBM samples(Yuan et al., 2018) (Methods). In line with our previous findings, the AC-like signature highly correlated with multiple metabolic pathways identified in the N1IC model, including lipid peroxidation, fatty acid (FA) metabolism, generation of ROS, and mitochondrial metabolism. Conversely, NPC-like and proliferating cell state scores negatively correlated with these metabolic programs but positively correlated with other pathways, such as amino acid metabolism (**Figure S7E, Table S6**). Thus, we demonstrate that quiescent AC-like cell states with specific metabolic programs identified in the N1IC murine model are ubiquitously present in human GBM samples.

Lastly, to test the potential translational impact of our findings, we applied our recently reported approach of treating acute slice cultures of human glioma surgical specimens with drugs and deconvolving cell type-specific responses with scRNA-seq(Zhao et al., 2021). We performed either DMSO (vehicle) or RSL3 treatments for 18 hours on organotypic slice cultures from six gliomas, including five primary GBMs and an IDH1-mutant anaplastic astrocytoma (**Figure 4A-F**, **Table S7**). Cells from the RSL3-treated slice cultures co-clustered with subsets of cells from vehicle-treated slices, suggesting that RSL3 selectively depletes specific subpopulations (**Figure 4A**). We used aneuploidies or other large copy number alterations, which cause significant increases or decreases in the relative expression of genes on amplified or deleted chromosomes, to identify transformed glioma cells(Yuan et al., 2018) (**Figure 4E-F,** Methods). For the five GBMs we used the relative expression of Chr. 7 (amplified) and Chr. 10 (deleted) (**Figure 4E)**, while for the IDH1 mutant glioma we used Chr. 1 (deleted) and Chr. 2 (unaltered) (**Figure 4F)**. This analysis allowed us to perform differential expression analysis between the vehicle- and RSL3-treated slices specifically for the transformed glioma cells from each patient (**Figure S8, Table S8**). Using the gene signatures for the four-state model of glioma cell phenotype (Neftel et al., 2019), we performed GSEA to show that the astrocyte-like glioma cell state was depleted across all six patient samples (**Figure 4G**), consistent with our findings in the animal model. Interestingly, this effect was independent of patient’s gender, MGMT methylation, p53 mutations or EGFR status. As we and others have shown, this astrocyte-like glioma cell subpopulation is largely quiescent and resistant to conventional anti-proliferative therapies (Zhao et al., 2021). For three of the six patients, we also co-treated slice cultures with RSL3 and Ferrostatin-1, ferroptosis inhibitor which we would expect to reverse the effects of the GPX4 inhibitor. Indeed, differential expression analysis and GSEA show an enrichment of the astrocyte-like signature in the transformed cells that were co-treated relative to those treated with RSL3 alone (**Figure 4H, Table S8**). These results suggest that RSL3 depletion of astrocyte-like glioma cells occurs primarily through ferroptosis. Notably, RSL3 induced widespread depletion of the astrocyte-like signature in transformed glioma cells across patient samples, including several genes with important metabolic functions, such as *FABP7* (fatty acid binding protein 7), involved in fatty acid and mitochondrial metabolism, *DBI* (Diazepam Binding Inhibitor, Acyl-CoA Binding Protein) involved in lipid metabolism, and *GPM6B* (glycoprotein M6B), membrane proteolipid with important roles in membrane trafficking (**Figure 4I, Table S8**).

**Figure 4:**
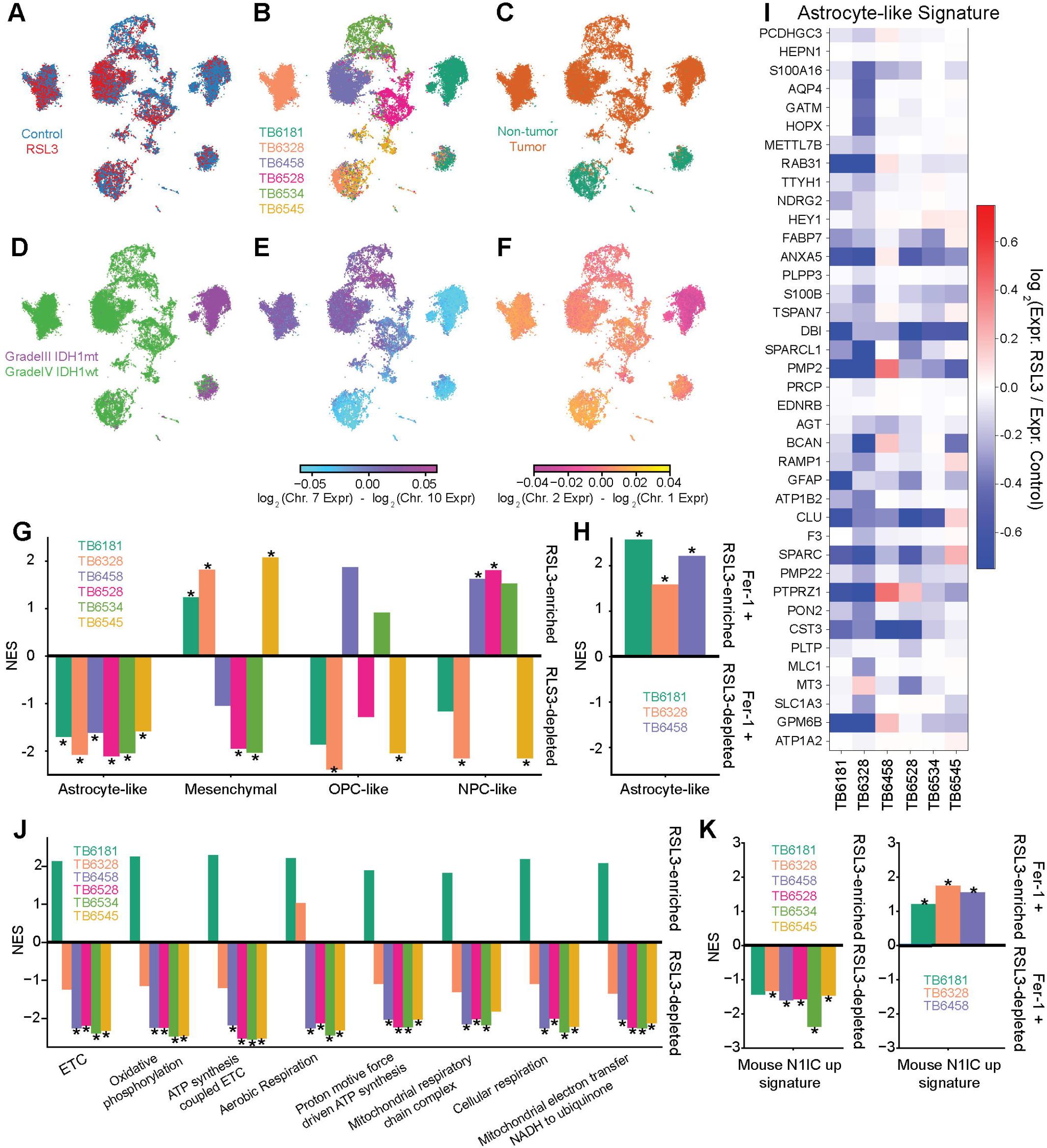
RSL3 targets quiescent AC-like transformed cell populations in acute slice cultures from human glioma. A-F. UMAP embedding of scRNA-seq data from vehicle- and RSL3-treated slice cultures from six gliomas, including five primary GBMs and one primary IDH1 mutant adult-type diffuse glioma. A shows the cells annotated by treatment; control (Blue), RSL3 (Red). B shows the cells annotated by tumor. C shows cells annotated as non-tumor (green) or tumor (orange). D shows cells annotated by IDHmt (purple) or IDHwt (green). E and F show cells annotated by chromosomal copy number alterations. Also refer to **Supplementary Figures S7** and **S8** and **Table S6-8**. A. G. Normalized enrichment scores (NES) for glioma cell state-specific gene signatures comparing the vehicle treated vs. RSL3 treated slices for all 6 cases. B. H. NES for the astrocyte-like gene signature, comparing RSL3 treated vs. RSL3+Ferrostatin-1 treated slices for 3 of the 6 cases. I. Heatmap of the fold-change for each gene in the astrocyte-like gene signature across all six patients comparing the vehicle and RSL3 treated slices. C. J. GSEA of mitochondrial metabolic signatures comparing the transformed glioma cells in the vehicle and RSL3 treated slices. D. K. Normalized enrichment scores for the “N1IC_up” gene signature derived from the murine model comparing the vehicle treated vs. RSL3 treated slices for all 6 cases. Right panel depicts the same gene signature comparing RSL3 treated vs. RSL3+Ferrostatin-1 treated slices for 3 of the 6 cases. G, H, J, K: Significant (FDR-corrected p<0.05) NES marked with asterisk (*).

While these results are highly concordant with our findings in the N1IC glioma cells, which harbor an astrocyte-like phenotype and are highly sensitive to RSL3-induced ferroptosis, we wanted to know whether the RSL3-sensitive cells in human gliomas bore a metabolic resemblance to N1IC glioma cells. Thus, we performed GSEA on the differential expression analysis comparing the transformed glioma cells in the vehicle- and RSL3-treated slices using mitochondrial metabolic signatures that were enriched in the N1IC model relative to the p53 model (**Figures 4J** and **2A**). Indeed, signatures related to mitochondrial respiration and oxidative phosphorylation were depleted by RSL3 treatment in five of the six gliomas. Interestingly, the IDH1 mutant glioma was the tumor that did not conform to this trend, suggesting that it has a distinct metabolic profile from the primary GBMs, but still harbors an astrocyte-like subpopulation that is selectively vulnerable to ferroptosis. Lastly, we found that RSL3 depletes the “N1IC_up” gene signature in all patient samples with the exception of the IDH1 mutant astrocytoma, an effect which is rescued with addition of Ferrostatin-1 (**Figure 4K**). These findings demonstrate that the murine derived N1IC gene signature can be used to label ferroptosis responsive tumor cell populations in human IDH1-wt GBM. This signature could be used as biomarker for patient selection and response monitoring in future clinical trials employing GPX4 inhibitors. Taken together, these experiments support the potential clinical applicability of GPX4 inhibitors to eradicate quiescent astrocyte-like tumor cells in human GBM by targeting a mitochondrial vulnerability of this persister-like population which is resistant to standard of care antiproliferative therapies.

## DISCUSSION

Using a combination of genetically engineered mouse glioma models, *in vitro* functional and metabolomic studies and *ex vivo* studies with organotypic slice cultures generated from patient-derived GBM samples, we show that Notch signaling can induce a quiescent astrocyte-like persister phenotype with a metabolic profile characterized by alterations in mitochondrial function leading to increased ROS and sensitivity to GPX4-driven ferroptosis. Our results are consistent with analysis of human GBM samples showing high Notch signaling in a subpopulation of slowly cycling glioma cells, present both in treatment naïve samples and enriched after exposure to cytotoxic therapies(Liau et al., 2017). Furthermore, the Notch-induced quiescent phenotype has been mechanistically linked to the upregulation of Notch targets *HES5* and *HEY1*(Liau et al., 2017), which are both significantly higher in the N1IC cells, and downregulation of *ASCL1*(Park et al., 2017; Sueda et al., 2019), a neural progenitor master regulator which is significantly downregulated in N1IC compared to p53 glioma cells. The importance of these findings extends beyond the specifics of Notch signaling in glioma cells and reveals a link between a quiescent AC-like glioma state, cellular metabolism, and therapeutic vulnerability.

### The clinical and therapeutic implications of cellular quiescence are context specific

Genes that promote a quiescent phenotype can slow tumor growth and prolong survival, such as in the N1IC mouse glioma model. Genetically deleting Notch induces faster tumor growth and significantly shortens survival(Giachino et al., 2015). Thus, Notch signaling can function as a tumor suppressor in mouse models of gliomagenesis. In our murine model, high constitutive Notch signaling in endstage N1IC tumors refutes a possible selection process of N1IC inactive clones as mechanism for driving tumor formation. Therefore, rather than acting as a pure tumor suppressor, Notch activation in our model leads to induction of a quiescent astrocyte-like cell state which recapitulates glioma cell populations present both in primary and recurrent GBM. Moreover, previous studies have shown that Notch-dependent quiescent glioma cells are resistant to cytotoxic/anti-proliferative therapies(Eyler et al., 2020; Jung et al., 2021; Liau et al., 2017), and analysis of patient samples has also shown that quiescent cells are insensitive to drugs that preferentially target proliferating cells(Zhao et al., 2021),(Spinazzi et al., 2022). Quiescent cells are therefore likely culprits for resistance and tumor recurrence in GBM following treatments that target proliferating populations(Couturier et al., 2020; Hoang-Minh et al., 2018; Xie et al., 2022). Most studies have considered quiescence through the conceptual framework of glioma stem cells. In contrast, our results highlight that quiescent glioma cells can have an astrocyte-like phenotype that is enriched in lineage restricted genes. It has been proposed that this is a more differentiated phenotype with relatively limited tumor propagation potential(Suva and Tirosh, 2020). However, analysis of the TCGA database revealed significantly shorter overall survival for patients whose tumors samples are enriched for the astrocyte-like signature(Levitin et al., 2019). Furthermore, scRNA-seq analysis of MRI-localized biopsies showed that astrocyte-like glioma cells are relatively abundant at the infiltrated margin, where they may escape surgical resection(Levitin et al., 2019), and due to their quiescent state may be resistant to standard of care radiation and chemotherapy. Importantly, a subset of quiescent tumor cells is capable of reentering the cell cycle and repopulating the proliferating cell pool, thereby reconstituting the heterogeneous cellular architecture of the tumor after cytotoxic treatment(Oren et al., 2021; Xie et al., 2022). Together, these findings highlight the need to develop more effective ways to target quiescent AC-like glioma cells.

### The link between glioma cell state and metabolic programs remains an important question in GBM

Nutrient and oxygen availability are important bottlenecks in tumor growth, modulating tumor phenotype(Barthel et al., 2019). Therefore, glioma metabolic status is another key determinant of tumor behavior and resistance to therapy (Nguyen et al., 2021; Rabe et al., 2020; Torrini et al., 2022). One recent study proposed a functional dichotomy in primary GBM, with glioma cells following either a metabolic or a neurodevelopmental axis during gliomagenesis(Garofano et al., 2021). Our results further refine the link between transcriptional cell states and highly specialized metabolic programs. In addition to the enrichment for transcriptional signatures for oxidative phosphorylation and ETC in the N1IC glioma model, several genes associated with the Notch-induced astrocytic-like phenotype have metabolic implications. Notably, two genes upregulated in N1IC cells, *Fabp7* and *Tspo*, have been implicated in alterations of mitochondrial function. Fabp7 is highly expressed in astrocytes with damaged mitochondria(Killoy et al., 2020) while Tspo is a translocator protein localized to the outer mitochondrial membrane that regulates mitochondrial respiration and generation of ROS by opening the mitochondrial permeability transition pore, which causes depolarization of the mitochondrial membrane and uncoupling of the ETC(Daugherty et al., 2016; Milenkovic et al., 2019; Rupprecht et al., 2010). Consistent with these transcriptional alterations, our metabolomic and functional analyses highlighted metabolic differences between N1IC and p53 cells. Specifically, we found higher glycolysis in the rapidly proliferating p53 model, in line with several recent studies (Garofano et al., 2021; Hoang-Minh et al., 2018). In contrast, N1IC glioma cells showed uncoupling of mitochondrial oxidative phosphorylation leading to increase in ROS and lipid peroxidation. Notably, noncanonical Notch activity has been recently shown to directly modulate complex I function in the electron respiratory chain(Ojha et al., 2022). Oxidative phosphorylation plays a critical role in glioma progression(Bonnay et al., 2020; Ojha et al., 2022) while uncoupling has been shown to drive therapeutic resistance in cancer(Demine et al., 2019). In our model, uncoupling leads to profound redox imbalance in the N1IC AC-like population, which, in conjunction with ineffective cysteine/methionine metabolism, leads to depletion of GSH stores. In contrast, p53 cells rely on the transsulfuration pathway, with more efficient production of reduced glutathione from L-cystathionine. Lipidomic signatures demonstrated enrichment of glycerolipids in the AC-like N1IC model, mirroring the lipid profile recently described in astrocytes(Fitzner et al., 2020) as well as profound depletion of PUFAs, likely secondary to baseline lipid peroxidation. Overall, ROS may play important roles in modulating the link between cell state and metabolic programs and reveal potential cell state specific therapeutic liabilities. Further studies are needed to investigate this link.

### Developing therapeutic strategies based on cell-state specific vulnerabilities remains an important goal in oncology

Tumor cell populations with specific metabolic affinities and potentially unique therapeutic vulnerabilities have been recently identified in glioma(Garofano et al., 2021; Hoang-Minh et al., 2018; Rusu et al., 2019). Long-term exposure of glioma cells to cytotoxic therapies such as temozolomide induced a quiescent cell state with mitochondrial metabolic reprogramming(Rabe et al., 2020). In line with these findings, we show that chronic intratumoral topotecan leads to expansion of a quiescent AC-like tumor population enriched in the N1IC metabolic gene signature. In other solid cancers, treatment-resistant persister-like cells demonstrate acquired GPX4 dependency and specific sensitivity to ferroptotic cell death(Hangauer et al., 2017; Oren et al., 2021; Viswanathan et al., 2017). In line with these studies, we found that mitochondrial dysfunction renders the N1IC cells more sensitive to GPX4 inhibitors and ferroptotic cell death (Yang et al., 2014). Intriguingly, modulating the electron flow in the ETC of glioma cells directly affected sensitivity to RSL3, a GPX4 inhibitor. In other cancer models inhibition of complex I attenuated sensitivity to cysteine-deprivation but did not affect GPX4 driven ferroptosis(Gao et al., 2019), suggesting a highly cell type specific and context dependent effect. Remarkably, treating acute slices of GBM tissue generated from surgical samples consistently with RSL3 depleted the quiescent AC-like glioma cell populations in all cases tested. Furthermore, RSL3 treatment caused a significant depletion in genes associated with mitochondrial respiration and oxidative phosphorylation in all five IDH-wild type tumors. Together these findings reveal a link between the quiescent AC-like glioma state and a specific metabolic/therapeutic vulnerability that translates from the N1IC mouse glioma model to human IDH-wild type GBM. Notably, the IDH-mutant glioma we tested also contained quiescent AC-like glioma cells that were selectively depleted by RSL3, although effects on the metabolic signatures were different, suggesting that sensitivity to ferroptosis is mechanistically distinct in IDH-mutant gliomas, possibly related to the effects of the IDH-mutation on cellular metabolism(Karpel-Massler et al., 2017) (Fack et al., 2017). Lastly, the dependency on GPX4 yet not SLC7A11 in both models may suggest that glioma states obtain cysteine independent of the system x_c_^-^ antiporter while still depending on robust GPX4 activity to eliminate lethal concentrations of lipid peroxides. NPC-like cells appear to rely on the transsulfuration pathway for cysteine and GSH synthesis. In contrast, AC-like cells have depleted GSH stores at baseline secondary to lipid peroxidation and use of cysteine in the taurine/hypotaurine pathway. Notably, GPX4 has been shown to modulate peroxide-induced ferroptosis through thiol independent selenolate based catalysis in some cell types of the developing brain(Ingold et al., 2018). The role of GPX4-independent mechanisms in governing ferroptosis susceptibility of various glioma cell states needs further investigation.

### Our translational studies also address the need for models that recapitulate the quiescent glioma population

One major barrier to developing new treatments that can effectively target quiescent tumor cells is the lack of experimental models that recapitulate the quiescent glioma populations. Both *in vitro* glioma cell culture models and *in vivo* cell transplantation glioma models commonly used to test drugs and other anti-glioma therapies are predominantly composed of proliferating tumor cells, and therefore are not well suited to study quiescent glioma cells(LeBlanc et al., 2022; Pine et al., 2020). In this respect, the acute slice culture model provides an important advantage. We previously reported that the slice model recapitulates the diversity of glioma cell states seen in GBM samples, and that scRNA-seq of slices treated *ex vivo* provides a way to assess cell type specific effects of drugs on the different glioma cell states(Zhao et al., 2021). In our previous studies we showed that treating acute GBM slices with the topoisomerase inhibitor etoposide selectively depletes subpopulations of proliferating glioma cells, consistent with its known mechanism of action(Hande, 1998; Zhao et al., 2021). Our results in the current study show that RSL3 can target quiescent astrocyte-like glioma cells, which are relatively resistant to etoposide(Zhao et al., 2021). Future studies could use the slice culture model to test if combinations of drugs, such as Etoposide and RSL3, can more effectively eliminate both proliferating and quiescent glioma cell populations in order to obtain a durable response in GBM.

## ACKNOWLEDGMENTS

We would like to thank all patients who generously donated tissue for these studies. We thank members of the Stockwell group and members in the High Throughput Screening Facility for excellent technical assistance, D. Teasley and O. Adeuyan for optimizing *in vitro* experiments, Dr. M. Chiriac for assistance with figure editing, Drs. X. Yang and G. Zhang from the Proteomics and Metabolomics Core Facility at Weill Cornell Medicine for LC-MS metabolomics analysis, Dr. M. Sisti. Dr. G. M. McKhann and Dr. B. Youngerman for assistance with tissue acquisition. M.A.B. was partially supported by the NREF&AANS/CNS Section on Tumors Research Fellowship Grant and the R38CA231577 StARR Kirschstein NRSA award.

J.N.B. was partially supported by William Rhodes and Louise Tilzer Rhodes Center for Glioblastoma, the Khatib Foundation and The Gary and Yael Fegel Foundation. This study was supported by the Emerson Health Collective Cancer Research Fund and the NIH/NCI/NINDS//NHLBI (J.N.B., P.A.S., P.C. – R01NS103473, J.N.B., P.C. – UL1TR001873, B.R.S. – P01CA87497, R35CA209896 and R61NS109407, J.K. – 5R01HL112626 N-SIM microscope – S100D014584, Nikon A1RMP confocal microscope – S10RR02568). This study used Cancer Center Flow and Confocal Core Facilities funded and the Genomics and High Throughput Screening Shared Resource in part funded by the NIH/NCI Cancer Center Support Grant P30CA013696.

## Author contributions

Conceptualization, M.A.B., A.D., H.C., J.K., P.A.S., P.C.; Methodology, M.A.B., A.D., M.G.A., W.Z., H.C., C.K., M.D.S., J.K., J.N.B., B.R.S., P.A.S., P.C.; Software, M.A.B., W.Z., M.G.A., C.K., P.A.S.; Investigation, M.A.B., A.D., M.G.A., W.Z., H.C. C.P.S., B.P., D.M.O.H., L.F.Y., A.M., N.H., P.S.U., F.Z., T.T.T.N., P.B.W., C.K., A.R., J.L.F. ; Writing – Original Draft, M.A.B., A.D., P.A.S., P.C.; Writing – Review & Editing, M.A.B., A.D., M.G.A., H.C., F.Z., M.D.S., J.K., J.N.B., B.R.S., P.A.S., P.C.; Funding Acquisition, M.A.B., J.K., J.N.B., B.R.S., P.A.S., P.C.; Resources, H.C., L.F.Y., A.M., N.H., F.Z., T.T.T.N., M.D.S., J.K., B.R.S., P.C.; Supervision, J.K., J.N.B., B.R.S., P.A.S., P.C.

## METHODS

### Murine glioma models

#### Animals

All procedures were reviewed and approved by the Columbia University Institutional Animal Care and Use committee (IACUC). C57BL/6 male and female mice were used as background for all experiments, including genetic and cell transplantation models. Wild-type C57BL/6 mice were obtained from Jackson Laboratories (strain #000664). Mice harboring a stop-flox N1IC construct(Buonamici et al., 2009) were crossed with *TP53*^fl/fl^ mice(Lei et al., 2011). The resulting mouse line was maintained by breeding of N1IC^+^/TP53^fl/fl^ male mice with TP53^fl/fl^ female mice. N1IC^+^/TP53^fl/fl^ mice were further crossed with TP53^fl/fl^/mCherry-Luciferase^stop-flox^ mice(Montgomery et al., 2020) to generate N1IC^+^/TP53^fl/fl^/mCherry-Luciferase^+^ mice. Mice were bred in house and housed under standard conditions in pathogen-free facilities at Columbia University Medical Center. Correct expression of transgenes was confirmed by genotyping from genomic DNA.

### Viral and orthotopic cell injections

Mice of both sexes (6-8 weeks old) were used for viral injections. 1 μL of PDGF-BB – IRES – Cre retrovirus (10^6^/mL titer) was injected into the subcortical white matter at the following coordinates (with bregma as reference): 2.2 mm lateral, 2.2 mm rostral and 2 mm deep, at a flow rate of 0.33 μL/minute. Tumor growth in mCherry-Luciferase^+^ mice was monitored by bioluminescent imaging on an IVIS Spectrum Optical Imaging System (Caliper) once weekly as previously described(Lei et al., 2011). Retrovirally induced tumors from 2 male N1IC^+^/TP53^fl/fl^ and 2 male TP53^fl/fl^ were dissected at end-stage for single cell RNA sequencing (scRNA-seq) analysis, as described below. For orthotopic cell transplantation experiments, 6-week-old C57BL/6 male mice were injected with 100,000 cells (p53-1 or N1IC-1, passage 10, isolated from littermates) resuspended in 2 μL PBS at the following coordinates (with bregma as reference): 2.5 mm lateral, 2 mm rostral and 2.5 mm deep, and at a flow rate of 0.25 μL/min. Mice were sacrificed either at 30 days post injection or when endstage criteria were met, as indicated in the corresponding Figures.

### Cell line isolation and culture

Primary murine glioma cells were isolated from virally induced end-stage tumors. Tumor tissue from the right hemisphere was minced and enzymatically dissociated in a cocktail of TrypLE and DNase at 37°C in a shaking water bath for 5 minutes. Tissue was triturated by passing through a flame-polished glass Pasteur pipette, filtered through a 70 μm mesh filter and neutralized with 50% FBS. Cells were collected by centrifugation and resuspended in BFP media, composed of DMEM, 10 ng/mL PDGF-AA (Peprotech), 10 ng/mL bFGF (Peprotech), N2 supplement (Gibco), antibiotic/antimycotic and 0.5% FBS. Cells were propagated in poly-L-lysine (PLL) coated tissue culture dishes in a humidified incubator (5% CO_2_) at 37°C. Studies were performed in litter and viral batch matched pairs. Cells were regularly checked for mycoplasma.

### Ferroptosis compounds and *in vitro* screening

Drug screens were performed in 384-well plates (Greiner Bio-One 781075) pre-coated with 10 μg/mL PLL for 60 minutes followed by 3ᵡ washes in PBS. Cells were plated at a density of 1,000 cells/well in 50 μL BFP medium and incubated overnight. Drugs were dispensed at the indicated concentrations using a HP Digital Dispenser. For drug combinations, wells were normalized by volume of DMSO added. Following a 24-hour (RSL3, ML162, IKE, IACS, Antimycin, Oligomycin, FCCP) or 48-hour (ML210) incubation, the plates were allowed to equilibrate to room temperature for 30 minutes and 25 μL of Cytotox Glo solution (Promega) was added to the wells using a Wellmate plate filler. Plates were read in a Tecan Infinite F200 multi-label plate reader. Data were exported to Microsoft Excel and GraphPad Prism for further analysis.

### BODIPY-C11 staining for flow cytometry and time-lapse confocal imaging

Cells were seeded in 6-well plates at a density of 25ᵡ10^4^ cells/well. After an 18-hour incubation, cells were labelled with 2 μM BODIPY-C11 dissolved in BFP media for 30 minutes. Cells were subsequently washed once with PBS, lifted with 10 mM EDTA in PBS (pH 7.4), and centrifuged for 5 minutes at 400g. The cell pellet was resuspended in flow buffer (0.1% w/v BSA, 1μg/ml DAPI in PBS) and filtered into polystyrene flow tubes. Flow cytometry was performed using a BD LSRFortessa Cell Analyzer using Pacific Blue, FITC and PE-Texas Red filters to detect DAPI, oxidized and reduced BODIPY-C11 signals, respectively. Data were processed using FlowJo v10.

For time-lapse imaging, 1.3ᵡ10^5^ N1IC-1 cells were plated on poly-L-lysine coated 35mm glass-bottom dishes (MatTek Corporation) for 24 hours. Cells were incubated for 30 minutes in BFP media containing 2 µM BODIPY-C11. Cells were washed once with warm PBS and media were replaced with fresh BFP. Cells were imaged on a Nikon A1RMP confocal microscope at 37°C in a humidified chamber with 5% CO_2_. Time-lapse images (512ᵡ512 pixels) were acquired using a 40ᵡ/1.3 NA oil immersion objective and focus was maintained using the Perfect Focus System. Excitation was achieved using 488 nm and 561 nm laser illumination; emission of the oxidized and reduced forms of BODIPY-C11 was captured using a 525/50 and a 595/50 filter, respectively. At time 0, RSL3 (500nM) was added and images were acquired every 30 seconds for a total of 30 minutes. Images were exported to ImageJ and ratio images of the 488 over the 568 channel were obtained using a pipeline detailed elsewhere(Kardash et al., 2011).

### MitoRED staining and flow cytometry

N1IC and p53 cells were seeded in 6-well plates at a density of 25ᵡ10^4^ cells/well. After an 18-hour incubation, cells were labelled with 25 nM MitoTracker Red CMXRos dissolved in BFP for 45 minutes. Cells were washed once with PBS and incubated with BFP media containing 2 μM FCCP or DMSO for 60 minutes. Cells were subsequently washed once with PBS, lifted with 10 mM EDTA in PBS (pH 7.4), and centrifuged for 5 minutes at 400g. The cell pellet was resuspended in flow buffer (0.1% w/v BSA, 1μg/ml DAPI in PBS) and filtered into polysterene flow tubes. Flow cytometry was performed using a BD LSRFortessa Cell Analyzer using Pacific Blue and PE filters to detect DAPI and MitoTracker signals, respectively, and data loaded into FlowJo v10 for further analysis. Cells were first gated by forward and side scatter areas (FSC-A and SSC-A) to remove debris, followed by filtering of multiplets using forward scatter area and height (FSC-A and FSC-H). Dead cells were filtered out by gating on low DAPI. MitoRED was visualized in the PE-A channel as a density distribution, and was gated at the median of the p53 condition treated with DMSO (which had the highest mean PE-A across the 4 conditions), and the percentage of live cells above this threshold were recorded (e.g. 50% for p53 cells in DMSO). The ratio of live cells above threshold between each condition with respect to the p53 control within each experiment was calculated, and technical triplicates for each condition were plotted using Prism.

### ROS measurement using H2DCFDA staining for flow cytometry

N1IC and p53 cells were seeded in 6-well plates at a density of 25ᵡ10^4^ cells/well. After an 18-hour incubation period, cells were labelled with 5 μM H2DCFDA reagent dissolved in BFP media for 30 minutes. Cells were washed, lifted, and prepared for flow cytometry exactly as described above. Flow cytometry was performed using a BD LSRFortessa Cell Analyzer using Pacific Blue and FITC filters to detect DAPI and ROS signals, respectively. Data were processed using FlowJo v10. Gating was carried out as described in the above MitoRED experiment, first by gating on cells (excluding debris), then singlets, then live cells. Quantification was performed in similar fashion to MitoRED. Briefly, H2DCFDA was visualized in the FITC-A channel as a density distribution, and was gated at the median of the p53 condition treated with DMSO (which had the highest mean FITC-A across the 2 cell lines), and the percentage of live cells above this threshold were recorded (e.g. 50% for p53 cells). The ratio of live cells above threshold between each condition with respect to p53 within each experiment was calculated, and technical triplicates for each condition were plotted using Prism.

### Liquid chromatography-mass spectrometry (LC-MS/MS) lipidomic profiling and analysis

Lipids were extracted using previously described methods (Ye et al., 2020). 2ᵡ10^6^ cells (p53 or N1IC of similar passage, n=4 technical replicates per condition) were grown in 10-cm dishes overnight, as detailed above. Cells were subsequently collected by scraping on dry ice with a microtip homogenizer and 132 μL of solution containing ice cold methanol and 0.01% (w/v) butylated hydroxyl toluene (BHT) were added to 100 μL of each cell pellet in a glass vial with a Teflon-lined cap. Samples were vortex mixed for 15 seconds. 668 μL of methyl tert-butyl ether (MTBE) was added and samples were again vortex mixed vigorously for 30 seconds. Samples were then incubated for 20 minutes in a cold room at 4^0^C on an orbital shaker and incubated for an additional 5 minutes on ice. The samples then underwent centrifugation at 3,000 rpm in a cold room at 4^0^C for 15-minutes for phase separation. The organic top layer was collected in a glass vial, dried under nitrogen stream, and stored at -80^0^C until used for analysis. Dried samples were reconstituted in 2-propanol/MeCN/H_2_O (4:3:1, v/v/v) for LC-MS analysis. 20 μL of each sample were used as quality control (QC). The protein pellet was used to measure protein concentration for normalization using the colorimetric Bradford assay (ThermoFisher Scientific).

Separation of extracted lipids was done at 55^0^C on ACQUITY UPLC HSS C18 Column over a 20-minute gradient elution (1.7 mM, 100 x 2.1 mm; Waters). The column was eluted isocratically for 1 minute with 80% mobile phase A (MeCN/water 60:40; v/v) with 10 mM ammonium acetate/0.1% acetic acid followed by 2-minute linear gradient 40% mobile phase B (2-propranol/MeCN/water 85:10:5; V/V/V) with 10 mM ammonium acetate/0.1% acetic acid followed by 70% at 2 minutes and 90% at 18 minutes. The flow rate was set to 400 µL/min and injection volumes were set at 6 µL e. Analysis was performed using a Synapt G2-Q-ToF mass spectrometrer (Water, Manchester, UK) in positive and negative electrospray ionization (ESI). Technical details were as follows: positive mode – +3 kV capillary voltage/32 V sampling cone voltage/source temperature 120^0^C/desolvation temperature 500^0^C; negative mode – -2 kV capillary voltage/30 V sampling cone voltage/source temperature 120^0^C/desolvation gas flow 900 L/hour. The traveling wave velocity was set to 650 m/s and wave height was 40 V. The helium gas flow in the helium cell region of the ion-mobility spectrometry cell was set to 180 mL/min to reduce the internal energy of the ions and minimize fragmentation. Nitrogen as the drift gas was held at a flow rate of 90 mL/min in the IMS cell. The low collision energy was set to 4 eV, and high collision energy was ramping from 25 to 65 eV in the transfer region of the T-Wave device to induce fragmentation of mobility separated precursor ions. All raw data were converted to netCDF using the MassLynx software v4.1. XCMS package in R was used for subsequent analysis. The CenWave algorithm was used for peak picking (CenWave algorithm, peak width window 2-25s) and peak grouping (bandwith 2s, m/z-width 0.01 Da). Peaks with variations larger than 30% in QCs were eliminated.

All the extracted features were normalized to measured protein concentrations measured by Bradford assay. Statistical analysis was performed in MetaboAnalyst. Group differences were calculated using Welch’s t-test with FDR-corrected p-value < 0.05Lipid identities were determined based on available online databases such as METLIN, Lipid MAPS, and HMDB and matching with MS^E^ fragments. Lipid ontology analysis was performed using the LION webtool (lipidontology.com), as previously described (Molenaar et al., 2019). The ranking mode was used for input and lipids were ranked based on log_2_FC (**Table S4**). One tailed-T-test was used to assess statistical significance of the comparison.

### Untargeted metabolomics by LC-MS/MS

Cells were processed for untargeted metabolomics as described elsewhere(Nguyen et al., 2021). Briefly, 2ᵡ10^6^ cells (p53 or N1IC of similar passage) were plated in 10 cm dishes overnight. The next day plates were washed twice with ice-cold PBS, placed on dry ice and 1mL 100% HPLC grade methanol was added to the dish. Cells were then scraped and transferred to cold Eppendorf tubes centrifuged at 14,000 x g for 20 minutes at 4°C and the supernatant was transferred to a fresh Eppendorf tube and dried using SpeedVac. The dried sample was reconstituted in water/acetonitrile (1:1; v/v) before LC-MS analysis. Protein was extracted from the pellets after centrifugation using cell extraction buffer with protease and phosphatase inhibitors. A colorimetric Bradford assay (ThermoFisher Scientific) was read at 740nm for evaluation of protein content.

LC-MS analyses were performed on a Q Exactive Orbitrap mass spectrometer (ThermoFisher Scientific) coupled to a Vanquish UPLC system (ThermoFisher Scientific). The Q Exactive operated in polarity-switching mode. A Sequant ZIC-HILIC column (2.1 mm i.d. × 150 mm, Merck) was used for separation of metabolites. Flow rate was set at 150 μL/min. Buffers consisted of 100% acetonitrile for mobile B, and 0.1% NH4OH/20 mM CH3COONH4 in water for mobile A. Gradient ran from 85% to 30% B in 20 min followed by a wash with 30% B and re-equilibration at 85% B. Initial data analysis was done using TraceFinder 4.1 (ThermoFisher Scientific). Metabolites were identified based on exact mass within 5 ppm and matching the retention times with the standards. Relative metabolite quantitation was performed based on peak area for each metabolite. Samples were normalized by protein content using a Bradford assay. MetaboAnalyst version 5.0 (metaboanalyst.ca) was used for differential assessment analysis, principal component analysis, metabolite set enrichment analysis (MSEA) and quantitative pathway analysis. Differentially enriched metabolites in N1IC or p53 cells were identified based on Welch’s t-test with a cutoff p<0.05 and no specific cutoff for the log_2_FC. MSEA analysis was performed using a KEGG based overrepresentation analysis (**Table S3**). For joint pathway analysis, we used differentially expressed genes with log_2_FC>0 based on the scRNA-seq analysis (**Table S1**) and differentially enriched metabolites in the N1IC cell line (**Table S3**). The pathway analysis was performed using a hypergeometric test with degree centrality and combined queries as integration method.

### Seahorse analysis of cellular respiration and extracellular acidification

Murine glioma cells (similar low passage N1IC-1/2 and p53-1/2) were seeded in PLL coated XFe24 cell culture microplates (Agilent TEchnologies) at 1.8ᵡ10^4^ cells per well in 250μL BFP (n=10 technical replicates for the baseline experiment and n=5 technical replicates for the cysteine/methionine deprivation experiment) and were allowed to attach overnight. For the cysteine/methionine deprivation (CMD) experiment, media was aspirated and replaced with either BFP or CMD BFP media after 4 hours and cells were grown in respective media for an additional 18 hours. Cells were then washed twice in PBS before changing to Seahorse XF base media (Agilent, 102353-100). Mitochondrial stress tests were run with the following concentrations of media: 10 mM glucose, 2 mM glutamine, and 1 mM pyruvate in assay medium, and 2 μM oligomycin, 2 μM trifluoromethoxy carbonylcyanide phenylhydrazone (FCCP), and 0.5 μM rotenone/antimycin A. The assay involved injection of glucose (10 mM), followed by oligomycin (1 μM), followed by 50 mM 2-deoxy-D-glucose. Cells were incubated at 37^0^C in a CO_2_-free atmosphere for 1 hour. Oxygen consumption rate (OCR) and extracellular acidification rate (ECAR) were analyzed using the XFe24 Extracellular Flux Analyzer (Agilent, Santa Clara, CA).

### Glutathione measurement

GSH concentration was calculated using the GSH Detection Assay Kit (Abcam; ab138881) as detailed in the manufacturer’s protocol. Briefly, 2.5ᵡ10^5^ cells (p53 or N1IC of similar passage) were plated per well of a six-well plate and grown for 24 hours as detailed above. Cells were washed with cold PBS and lysed with 100 μL ice-cold 1ᵡ Mammalian Lysis Buffer. Lysates were centrifuged at 21,000 g for 15 min. The supernatant was collected, mixed with 20 μL ice-cold trichloroacetic acid, incubated on ice for 10 minutes and centrifuged at 12,000 g for 5 minutes. The supernatant was neutralized by addition of NaHCO_3_ to a final pH of 6, centrifuged at 13,000 g for 5 minutes, and supernatants collected. Samples were corrected for the diluted volumes, mixed with GSH Assay Mixture and GSH was calculated by measuring in a microplate reader (Promega GloMax) using 490/520 nm excitation/emission filter and fitting into a GSH standard curve.

### qRT-PCR

2.5 µg of RNA were used for first strand cDNA synthesis using the SuperScript Vilo cDNA synthesis kit (ThermoFisher, 11754050). cDNA was diluted to 250 ng/µL and the RT-qPCR reactions were conducted using Thermo Scientific ABsolute Blue qPCR SYBR mix (ThermoFisher, AB4322B). Reactions were performed in triplicate samples per condition on an Applied Biosystems QuantStudio 3 qPCR instrument and all experiments were repeated 3 independent times. Samples were normalized to β-actin, used as reference, and fold change between conditions was calculated using the ΔΔCT method.

### Immunofluorescence and immunocytochemistry

Immunohistochemistry was performed on 4% PFA-fixed, paraffin-embedded sections or OCT-frozen tissue. Paraffin-embedded sections were first deparaffinized in xylene (35 min) and rehydrated with sequential incubations in 100 % ethanol (25 min), 95% ethanol (25 min) and 75% ethanol (15 min). Following a wash in distilled water, antigen retrieval was performed in 10 mM Citrate (pH 6.0) solution in a pressure cooker for 10 minutes. Slides were allowed to equilibrate to room temperature for 30 minutes, washed in PBS and incubated in blocking buffer (10% normal goat serum, 0.5% Triton X-100 in PBS) for 30 minutes at room temperature. For obtaining frozen sections, 4% PFA-fixed tissue was incubated in 30% sucrose solution in PBS followed by immersion and freezing in OCT. Frozen sections (12 μm thickness) were obtained in a cryostat (Leica), mounted on glass slides, and stored at -80°C. Sections were washed in PBS before incubation in blocking buffer for 30 minutes at room temperature. For immunocytochemistry, cells were plated at a density of 310^4^ cells on 12 mm glass coverslips in 24-well plates. Cells were fixed with 4% PFA for 15 minutes, permeabilized with 0.5% Triton X-100 in PBS for 5 minutes and blocked with blocking buffer (10% normal goat serum in PBS) for 30 minutes. Primary antibody incubations were performed at 4°C overnight. After three 10-minute washes in PBS, sections were incubated with the appropriate Alexa Fluor conjugated antibodies for 1 hour. Sections were washed extensively with PBS, incubated with DAPI for 10 minutes (Thermo Scientific, D1306, 0.5 μg/mL), and mounted using Fluoro-Gel with TES buffer (Electron Microscopy Sciences).

#### Confocal imaging

Confocal images (1024×1024 pixels) were acquired on a Zeiss LSM800 confocal microscope equipped with GaAsP point-detectors, using a Plan-Neofluar 40×/1.3 NA oil DIC objective. Fluorophores were excited using 405 nm, 488 nm, 561 nm and 639 nm wavelength lasers. Confocal stacks were acquired using 1 AU pinhole size and at 0.58 μm steps. Images were exported to ImageJ for further analysis. A minimum of three fields were obtained from each biological sample.

#### SIM imaging

Cells were plated on 1.5 glass coverslips and stained with 25 nM MitoTracker Red CMXRos for 1 hour. After a wash with warm PBS, cells were fixed with 4% PFA for 15 minutes, permeabilized with 0.2% Triton X-100 for 5 minutes and counterstained with DAPI and Alexa Fluor 488 conjugated Phalloidin for 30 minutes at room temperature. Coverslips were mounted using SlowFade Diamond Antifade mountant. Multichannel Structured Illumination Microscopy was performed on a Nikon N-SIM system using a 100ᵡ/1.49 NA objective. Fluorophores were excited using 405, 488 and 561 nm diode lasers and z-stack images (voxel size: 0.0634×0.0634×0.125 μm^3^) were acquired using an Andor iXon3 DU897 electron-multiplying CCD camera. Images were processed using Nikon Elements and ImageJ software. The resulting voxel size post-reconstruction was 0.0317×0.0317×0.125 μm^3^.

### *Ex vivo* organotypic murine slice experiments

6-week-old C57BL/6 male mice were injected with 100,000 cells (N1IC-1, passage 10) resuspended in 2 μL PBS at coordinates and rate described above. Mice were sacrificed at 30 days post injection, and tumor-bearing brains were dissected and placed into artificial CSF (aCSF). Murine tumor slices were generated and cultured as previously described(Zhao et al., 2021). Briefly, a McIlwain tissue chopper was used to generate 500 µm coronal sections of the tumor-bearing murine brains, starting at the anterior aspect of visible tumor. Slices were plated on Millicell cell culture inserts (catalog # PICM0RG50) in 6 well dishes and incubated in slice culture media (DMEM/F12, 1% N2 supplement, 1% anti-mycotic/anti-bacterial) with either DMSO control or 500nM RSL3 for 6 hours. After treatment was completed, slices were fixed in 4% PFA, embedded in paraffin, and cut into 5 μm-thick sections.

### Fluorescent *in situ* hybridization/RNAscope

*In situ* hybridization of murine Clu and Ptprz1 from paraffin-embedded 5 μm-thick sections was performed using the RNAscope Fluorescent v2 assay (ACDbio), as detailed in the manufacturer’s protocol. Of note, we used Ptprz1 as a cell-state agnostic tumor specific marker as it was not differentially expressed between the two murine models (**Table S1**). Briefly, sections were baked for 1 hour at 60°C, followed by deparaffinization in xylene and 100% ethanol. Slides were dried for 5 min at 60°C, followed by incubation with H_2_O_2_ for 10 min at room temperature. Antigen retrieval was performed in boiling 1 Target Retrieval reagent for 15 min. Slides were then washed in water, dehydrated in 100% ethanol, and incubated with Protease Plus for 30 min at 40°C. The C3 probe (Clu) was diluted 1:50 in the C1 probe (Ptprz1) and hybridized on the sections for 2 hours at 40°C. Ptprz1 was detected with TSA-Cy5 and Clu with TSA-Cy3. Nuclei were stained with DAPI and the slides were mounted with Fluoro-Gel with TES buffer (Electron Microscopy Sciences). Images were obtained from Ptprz1 high signal areas to identify tumor. For quantification of RNAscope images, nuclei were first segmented using the Huang intensity threshold followed by a watershed filter. For identification of Clu+ signals, images were thresholded using the “Sambhag white” filter to generate masks. Positive particles were identified using the “Analyze Particles” function and integrated density of Clu from the entire field was obtained. Clu intensity was expressed as weighted Integrated Density by dividing the sum of Clu integrated densities over the sum of cells per biological sample.

### Drug treatment of patient-derived tissue slices

Patient-derived tissue slices were generated and cultured as previously described(Zhao et al., 2021). Patient diagnosis information can be found in **Table S7**. From each individual patient tumor resection, we performed an 18-hour drug perturbation with 50 nM RSL3 or drug vehicle (DMSO) on spatially adjacent slices. For three patient tumor resections (TB6181, TB6328 and TB6458), we also co-treated slice cultures with 50 nM RSL3 and 10 μM Ferrostatin-1.

### scRNA-seq analysis

#### Tissue dissociation

Murine brain tumors and patient-derived tissue slices generated from TB6181, TB6328 and TB6458 were collected for dissociation using the Adult Brain Dissociation kit (Miltenyi Biotec) according to the manufacturer’s instructions. Patient-derived tissue slices generated from TB6528, TB6534 and TB6545 were collected for dissociation using Papain Dissociation System (Worthington) as previously described(Mizrak et al., 2020; Mizrak et al., 2019) with following modifications. Briefly, minced tissue was digested with papain (Worthington, 10 units per sample) in PIPES solution (120 mM NaCl, 5 mM KCl, 20 mM PIPES (Sigma), 0.45% glucose, DNase I (Worthington, 100 units per sample), 1ᵡ Antibiotic/Antimycotic (GIBCO, pH adjusted to 7.6) for 30 min at 37°C on gentleMACS™ Octo Dissociator with Heaters (Miltenyi Biotec, program 37C_ABDK). After digestion, the cell suspension was applied to a MACS SmartStrainer (70 μm, Miltenyi Biotec) and centrifuged at 300g for 10 minutes at 4°C. The cell pellets were then subjected to red blood cell removal (Red Blood Cell Lysis Solution (10ᵡ), Miltenyi Biotec) and debris removal (Debris Removal Solution, Miltenyi Biotec) according to the manufacturer’s instructions.

#### Microwell-based scRNA-seq

Dissociated cells from each mouse brain tumor specimen or patient-derived slice cultures were profiled using microwell-based single-cell RNA-seq(Yuan and Sims, 2016) as previously described(Zhao et al., 2021). Each cDNA library was barcoded with an Illumina sample index. Libraries with unique Illumina sample indices were pooled for sequencing on an Illumina Nextseq500/550 with an 8-base index read, a 26-base read 1 containing the cell barcode (CB) and unique molecular identifier (UMI), and a 58-base read 2 containing the transcript sequence (mouse brain tumors, TB6181, TB6328 and TB6458) or on an Illumina NovaSeq 6000 with an 8-base index read, a 26-base read 1 containing CB and UMI, and a 151-base read 2 containing the transcript sequence (TB6528, TB6534 and TB6545).

#### scRNA-seq data preprocessing

scRNA-seq data were preprocessed as described previously(Zhao et al., 2021). Briefly, raw sequencing data were trimmed and aligned (Levitin et al., 2019; Yuan et al., 2018). Data obtained from the Illumina NovaSeq 6000 were corrected for index swapping to avoid cross-talk between sample index sequences using the algorithm described by (Griffiths et al., 2018) before assigning read addresses for each sample. Reads with the same CB, UMI and aligned gene were collapsed and sequencing errors in the CB and UMI were corrected to generate a preliminary matrix of molecular counts for each cell. We applied the EmptyDrops algorithm(Lun et al., 2019) to recover cell-identifying barcodes in the digital gene expression matrix and filtered out CBs as described by (Zhao et al., 2021) to remove low quality cells.

#### Identification of neoplastic glioma cells and differential gene expression analysis in murine tumors

We identified transformed glioma cells from the murine scRNA-seq data by performing unsupervised clustering with Louvain community detection as implemented in Phenograph (Levine et al, Cell, 2015) followed by differential enrichment analysis using the binomial test to identify genes that are statistically enriched in each cluster as described previously(Shekhar et al., 2016). The pipeline for performing this analysis is described in (Levitin et al., 2019) and the corresponding code can be found at https://github.com/simslab/cluster_diffex2018. We defined the transformed glioma cells as cells in clusters that exhibit statistical enrichment of the Cre recombinase transcript. We performed differential expression analysis between transformed glioma cell populations in the two mouse glioma models using the Mann-Whitney U-test as described in (Zhao et al., 2021) after subsampling the relevant groups to the same cell numbers and the same average number of unique transcripts per cell and normalizing the resulting count data with scran (Lun et al., 2016). Genes were defined as differentially expressed between the two models if they log_2_FC>1 and FDR<0.05 in at least three of the four pairwise comparisons of the two sets of biological replicates for each model. Based on these parameters, these significantly upregulated and significantly downregulated genes therefore comprise the “N1IC_up” and “N1IC_down” signatures, respectively.

#### Gene set enrichment analysis (GSEA) in the murine model

Differential gene expression analysis was performed as described above and log_2_FC was then used to rank genes for Preranked GSEA. Preranked GSEA was performed in GSEA version 4.3.2 (Subramanian et al., 2005) using MSigDB C2 (KEGG, REACTOME, etc) and C5 (Gene Ontology) gene set libraries (Liberzon et al., 2011). Only statistically significant gene sets with a nominal p value<0.05 were depicted. Each tumor cell was also scored using the package AUCell in R (version 1.16.0) (Aibar et al., 2017) using the “Astrocyte-like” (AC) gene signature described in (Neftel et al., 2019), the “quiescence” gene signature described in (Shin et al., 2015), and the “persister” gene signature described in (Liau et al., 2017) and visualized as Violin plots using Seurat (version 4.1.0)(Hao et al., 2021).

#### Identification of neoplastic glioma cells and non-tumor cells in patient-derived slice cultures

The malignant glioma cells and non-tumor cells were identified as previously described(Zhao et al., 2021). Briefly, we first merged scRNA-seq data of all samples derived from the same patient for unsupervised clustering analysis. We used Louvain community detection as implemented in Phenograph for unsupervised clustering with k=20 for all k-nearest neighbor graphs. The marker genes were identified using the drop-out curve method as described in (Levitin et al., 2019)(https://github.com/simslab/cluster_diffex2018) for each individual sample, and we took the union of the resulting marker sets to cluster and embed the merged dataset. We defined putative malignant cells and non-tumor cells using the genes most specific to each cluster. Putative tumor-myeloid doublet clusters were removed first. Next, we computed the average gene expression of each somatic chromosome as described in (Yuan et al., 2018). For data obtained from IDH1-wild type glioma tissues (TB6328 and TB6458, TB6528, TB6534 and TB6545), we define the malignancy score to be the log-ratio of the average expression of Chr. 7 (amplified) genes to that of Chr. 10 (deleted) genes(Zhao et al., 2021). For data obtained from the IDH1-mutant glioma (TB6181), we define the malignancy score to be the log-ratio of the average expression of Chr. 2 (unaltered) genes to that of Chr. 1 (deleted) genes. We plotted the distribution of malignancy scores and fit a double Gaussian to the malignancy score distribution and established a threshold at 1.96 standard deviations below the mean of the Gaussian with the higher mean (i.e. 95% confidence interval). Putative malignant cells with malignancy scores below this threshold and putative non-tumor cells with malignancy scores above this threshold were discarded as non-malignant or potential multiplets (**Figure S8**).

#### Whole genome sequencing and analysis

Genomic DNA was extracted from a piece of fresh-frozen tissue from each patient tumor resection using the DNeasy Blood & Tissue Kit (Qiagen) according to the manufacturer’s instructions. Libraries for whole genome sequencing were constructed using the Nextera XT kit (Illumina) with unique i7 indices for each patient sample according to the manufacturer’s instructions.

Sequencing was performed with an Illumina NextSeq 500/550 using a 150-cycle High Output Kit (Illumina) with an 8-base index read, 80-base read 1, and 80-base read 2. Raw sequencing data were aligned to the human genome (GRCh38) using bwa mem as described previously(Yuan et al., 2018; Zhao et al., 2021). Briefly, we computed the number of de-duplicated reads that aligned to each chromosome for each patient and divided by the number of de-duplicated reads that aligned to each chromosome for a diploid germline sample from patient TB6545 (peripheral blood mononuclear cells) after normalization by total reads. We then normalized this ratio by the median ratio across all somatic chromosomes and multiplied by two to estimate the average copy number of each chromosome.

#### Cell-type specific differential expression analysis in patient derived slice cultures

We identified differentially expressed genes for RSL3-vs. vehicle-treated tumor cells as previously described (Zhao et al., 2021). Briefly, we first randomly sub-sampled number of cells in each comparison to have the same number of cells as the condition with fewer cells. Next, we subsampled the count matrices for the two conditions such that they had the same average number of molecules per cell and normalized the resulting count matrix using scran(Lun et al., 2016). We then conducted differential expression analysis for protein-coding genes using the two-sided Mann-Whitney U-test as implemented by the “mannwhitneyu” command in the Python module “scipy”. The resulting p-values were corrected for false discovery using the Benjamini-Hochberg procedure as implemented in the “mutlipletests” command in the Python module “statsmodels”. We used the same approach for the differential expression analysis for RSL3 and Ferrostatin-1 co-treated vs. RSL3-treated tumor cells.

#### Gene set enrichment analysis (GSEA) in patient derived slice cultures

GSEA was performed using GSEA version 4.3.2 by ranking differentially expressed genes by +/-log-ratio of q-values obtained from comparing the RSL3-vs. vehicle-treated tumor cells or the RSL3 and Ferrostatin-1 co-treated vs. RSL3-treated tumor cells. Gene signatures used for the four-state model of glioma cell phenotype were described by (Neftel et al., 2019). Gene signatures used for identifying the metabolic signatures were the gene sets from the GO Biological Process ontology.

#### Single cell hierarchical Poisson factorization (scHPF) analysis

The scHPF model was generated by combining scRNA-seq profiles from all patient slice culture data, randomly subsampling to an even number of cells from each patient, and randomly downsampling the resulting count matrix so that the average number of unique transcripts per cell was equal for each patient. We subjected the processed count matrix to probabilistic matrix factorization with scHPF (https://github.com/simslab/scHPF)(Levitin et al., 2019). To generate **Figure 4A-F**, we embedded the resulting cell score matrix in two dimensions with UMAP. The relative expression of chromosomes exhibiting aneuploidy shown in **Figure 4E-F** was computed using scHPF-imputed average expression values for the genes on each chromosome as described in Zhao et al(Zhao et al., 2021).

### Single-cell RNA sequencing analysis of published IDH1-wild type human glioma dataset

Single-cell RNA sequencing data was obtained from (Yuan et al., 2018), which is an annotated scRNA-seq dataset comprising tumors from 8 patients with IDH-wild-type glioblastoma. Cells that were annotated as “transformed” tumor cells were subsetted for each patient, marker genes were identified for each patient using the drop-out curve method as described in (Levitin et al., 2019) (www.github.com/simslab/cluster_diffex2018), and we took the union of the resulting marker sets to cluster and embed the merged 8-patient dataset into a UMAP. Merged scRNA-seq count data was loaded into R and normalized using the Seurat package.

Each tumor cell was scored using the package AUCell in R (version 1.16.0) (Aibar et al., 2017) for each of the six Neftel lineage gene sets: Mesenchymal 1 (MES1), Mesenchymal 2 (MES2), Astrocyte-like (AC), NPC-like 1 (NPC1), NPC-like 2 (NPC2), OPC-like (OPC). For each transformed cell, the highest of the 6 lineage scores via AUCell defined its lineage classification, and NPC1/2 and MES1/2 were then aggregated, resulting in 4 unique lineage classifications: MES, AC, NPC, OPC. Then, each cell also underwent cell cycle scoring using the “CellCycleScoring” function in Seurat(Butler et al., 2018) and the proliferation gene sets G1/S and G2/M described in (Neftel et al., 2019), which classified each tumor cell into “G0”, “G1/S”, and “G2/M” groups. Those assigned to G1/S or G2/M were defined as “proliferative”, while those assigned to G0 were defined as “quiescent”. These classifications were then projected on to the merged UMAP.

AUCell scoring for this dataset was also performed on each gene set in MSigDB C2 (KEGG, REACTOME, etc) and C5 (Gene Ontology) gene set libraries (Liberzon et al., 2011), as well as using the “N1IC_up” and “N1IC_down” gene signatures described above. AUCell scores for each of the 8 Neftel gene sets (6 lineage sets and 2 proliferation sets) across all patients underwent pairwise Spearman correlation analysis with AUCell scores for select C2 and C5 metabolic gene sets, as well as the “N1IC_up” and “N1IC_down” signatures, and these correlation results are depicted in heatmaps using the ComplexHeatmap (version 2.10.0) package in R (Gu et al., 2016).

### Bulk RNA sequencing analysis of convection-enhanced delivery clinical trial dataset

Transcript count data was obtained from (Spinazzi et al., 2022), which describes a clinical trial in recurrent glioblastoma in which 86 MRI-localized biopsies were taken before or after intratumoral delivery of topotecan across 5 patients. Differential gene expression analysis comparing pre- and post-topotecan biopsies was performed using the DESeq2 package in R and was done both on a patient-by-patient basis as well as across all patients. Genes were ranked using log_2_FC, and Preranked GSEA was performed using the GSEA application as described above using the “N1IC up” gene signature.

### Statistical Analysis

Statistical analysis and generation of graphs were performed using GraphPad Prism v9.0 and RStudio. P values were calculated using unpaired two-tailed t-tests with unequal variances or paired two-tailed t-tests, where appropriate. A P value of .05 was the cutoff for statistical significance (*p<0.05, **p<0.01, ***p<0.001). Log rank/Mantel-Cox test was used for analysis of survival in the mouse model. Welch’s t-test was used for simple comparisons. Kruskal Wallis with uncorrected Dunn’s *post hoc* test was used for multiple comparisons. Multiple hypothesis correction was performed using the Benjamini-Hochberg method or the 2-stage linear step-up procedure of Benjamini, Krieger, and Yekutieli, where indicated. Dose response curves for ferroptosis drugs were generated using sigmoidal 4 parameter nonlinear regression. Data are reported as mean ± standard error of the mean (SEM). Number of replicates and details of statistical tests used are described in the figure legends and text.

### Data and software availability

Datasets generated for this study including murine scRNA-seq data and the human slice culture scRNA-seq data are deposited in the Gene Expression Omnibus (GEO: GSE224727). The code used for analysis and generation of figures is available on Github ((https://github.com/simslab). All unique/stable reagents and mouse/cell lines generated in this study are available within reasonable request from the lead contact with a completed Materials Transfer Agreement. Further information and requests for resources and reagents should be directed to and will be fulfilled by the lead contact, P.C.

## Supplementary Figure Legends

**Figure S1 (associated with Figure 1):**
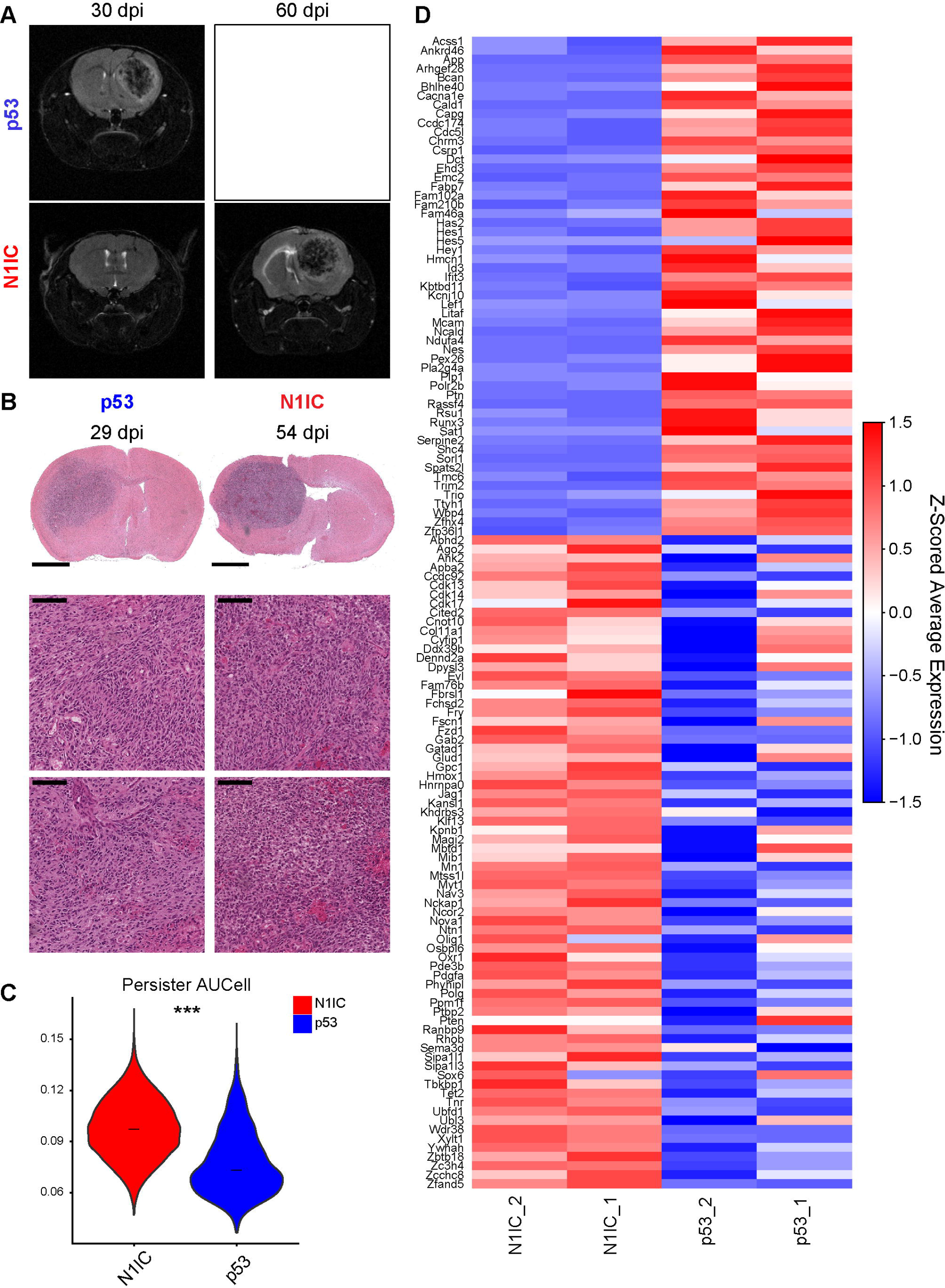
Radiographic, histopathologic, and transcriptomic features of the N1IC and p53 mouse glioma models. A. MRI demonstrates significantly slower tumor progression and increased latency in tumor formation in p53-/- N1IC mice with diffusely infiltrative tumors at 30 dpi and large necrotic and hemorrhagic tumors at 60 dpi. p53 tumors reach end stage around 30 dpi. B. Representative H&E staining of endstage tumors demonstrate high grade histological features, including pseudopalisading necrosis and neo-angiogenesis in both models. Scale bars, 2 mm for panoramic view and 100 μm for detailed inset. C. Violin plot of AUCell persister scores in the two models, ***p<0.001 Welch’s t test. Persister scores based on gene set described in (Liau et al., 2017). D. Heatmap of differentially expressed genes that are direct N1ICD targets in persister cells. Complete gene list also provided in **Table S2.**

**Figure S2 (Associated with Figure 2):**
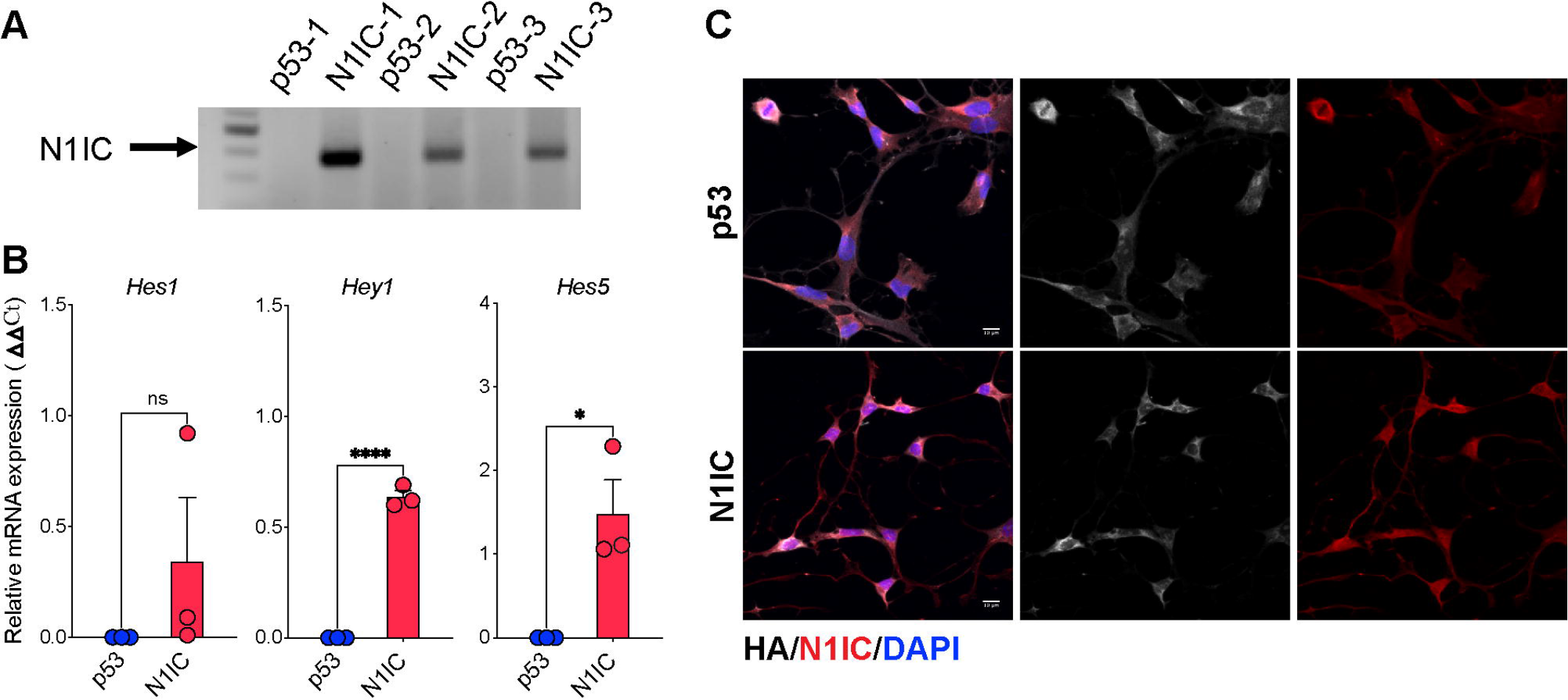
Differential activation of Notch signaling in glioma cells isolated from N1IC vs. p53 tumors. A. Genomic DNA PCR isolated from 6 different murine cell lines demonstrating persistence of the N1IC transgene in all three N1IC cell lines. B. Immunofluorescence of the N1IC and the tumor marker HA in a pair of p53/N1IC cells demonstrating nuclear localization of the intracellular portion of the Notch1 receptor in the N1IC cell line. All cells cultured *in vitro* expressed the HA marker. Scale bar, 10 μm. C. qRT-PCR of direct Notch downstream targets Hey1, Hes1 and Hes5 demonstrating upregulation of Hey1/Hes5 in the N1IC cell line. Data obtained on the same pair of N1IC/p53 cells used in panel B pooled from n=3 independent experiment. *p<0.05, unpaired t test.

**Figure S3 (Associated with Figure 2):**
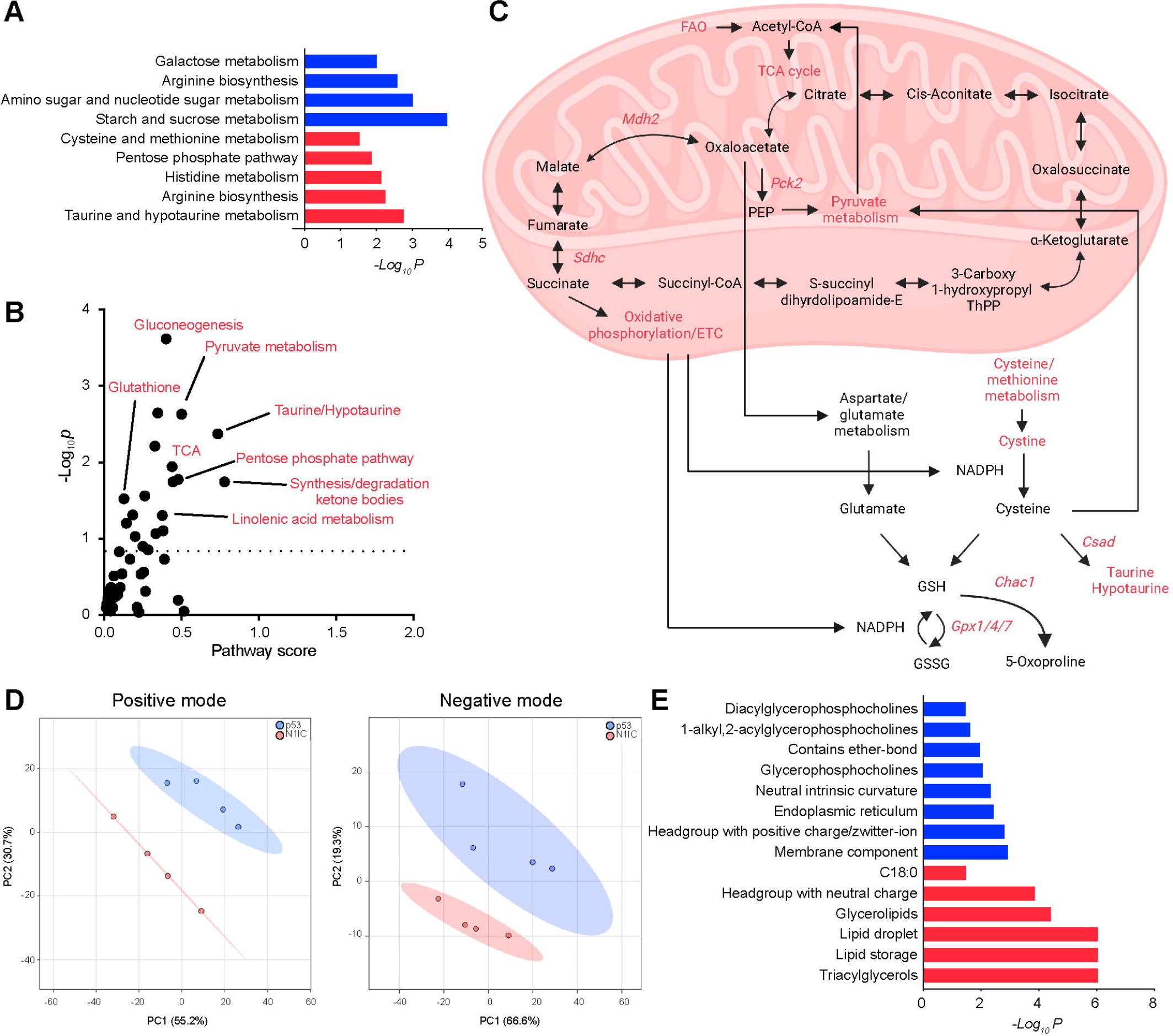
Metabolomic/lipidomic pathway analysis reveals differences between N1IC and p53 cells. A. Metabolite Set Enrichment Analysis (MSEA) depicting significant ontologies from untargeted metabolomic LC-MS on p53 and N1IC cell lines. B. Combined transcriptomic/metabolomic pathway analysis in the N1IC model demonstrating activation of taurine/hypotaurine, pyruvate and pentose phosphate pathways. Select significant pathways are highlighted in red. Punctate line indicates cutoff level of significance (p<0.05). C. Diagram depicting upregulated enzymes and metabolic pathways in the N1IC model demonstrating ineffective cysteine/methionine metabolism and highly active mitochondrial metabolic pathways including FAO, TCA cycle, pyruvate metabolism and oxidative phosphorylation. FAO – fatty acid oxidation, TCA – tricarboxylic acid. Select significant metabolites/enzymes/pathways upregulated in N1IC model are highlighted in red. D. PCA of lipidomics data shows that the lipid profile segregates by model, in both the negative and positive mode. E. Lipid ontology analysis demonstrating upregulation in TAG in the N1IC model with enrichment in lipids stabilizing the membrane in p53 cells. TAG – triacylglycerols.

**Figure S4 (Associated with Figure 2):**
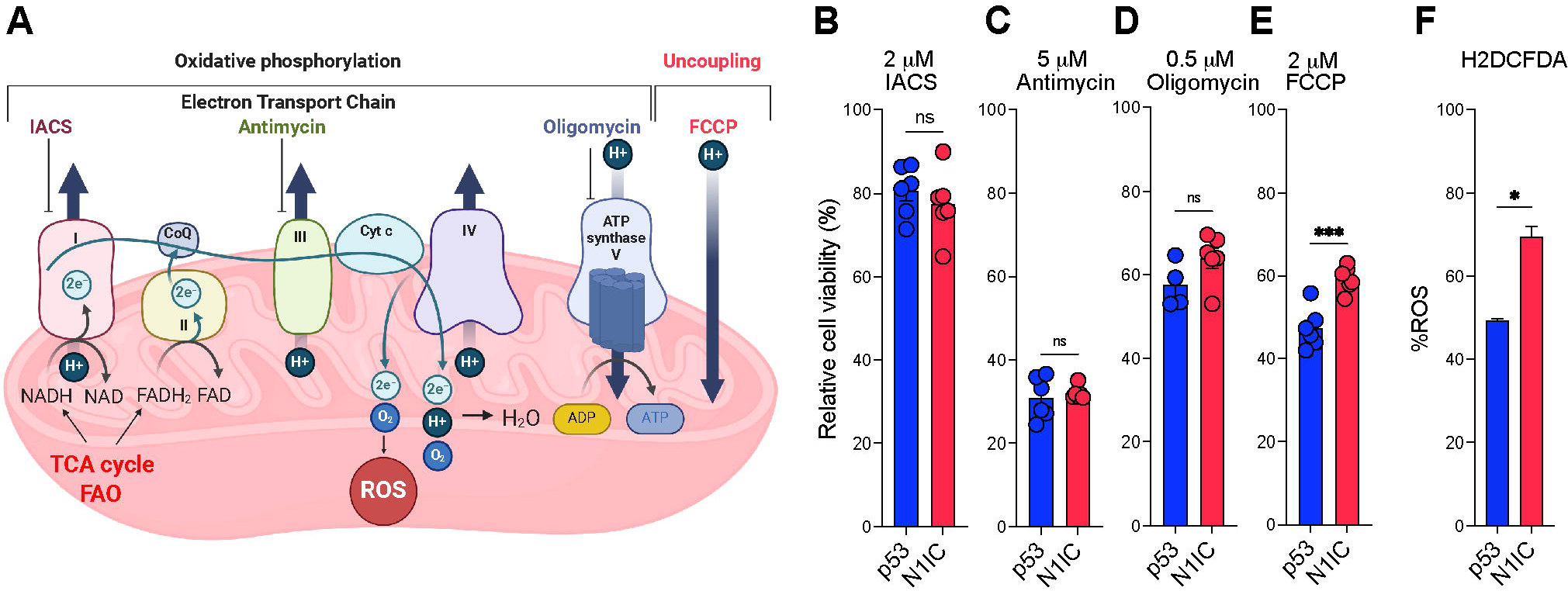
N1IC and p53 cells are differentially sensitive to mitochondrial uncoupling. A. Image summarizing the electron transport chain (ETC) and role of uncoupling in controlling ROS production. Image adapted in Biorender from (Shrestha et al., 2021). Pharmacologic ETC inhibitors used in panels B-E with their specific targets are also depicted. B-E. Response to mitochondrial ETC inhibitors demonstrating differential sensitivity to uncoupling agent FCCP (B) but no difference in response to complex I inhibitor IACS (C), complex III inhibitor Antimycin (D) or complex V/ATP synthase inhibitor Oligomycin (E) between p53/N1IC cells. Representative data from n=3 independent experiments. n=6 technical replicates per condition, ***p< 0.001, unpaired t test. A. F. % baseline ROS in a p53 and a N1IC cell line as assessed by H2DCFDA flow cytometry from n=2 independent experiments. p=0.0469 by Welch’s t test.

**Figure S5 (Associated with Figure 2):**
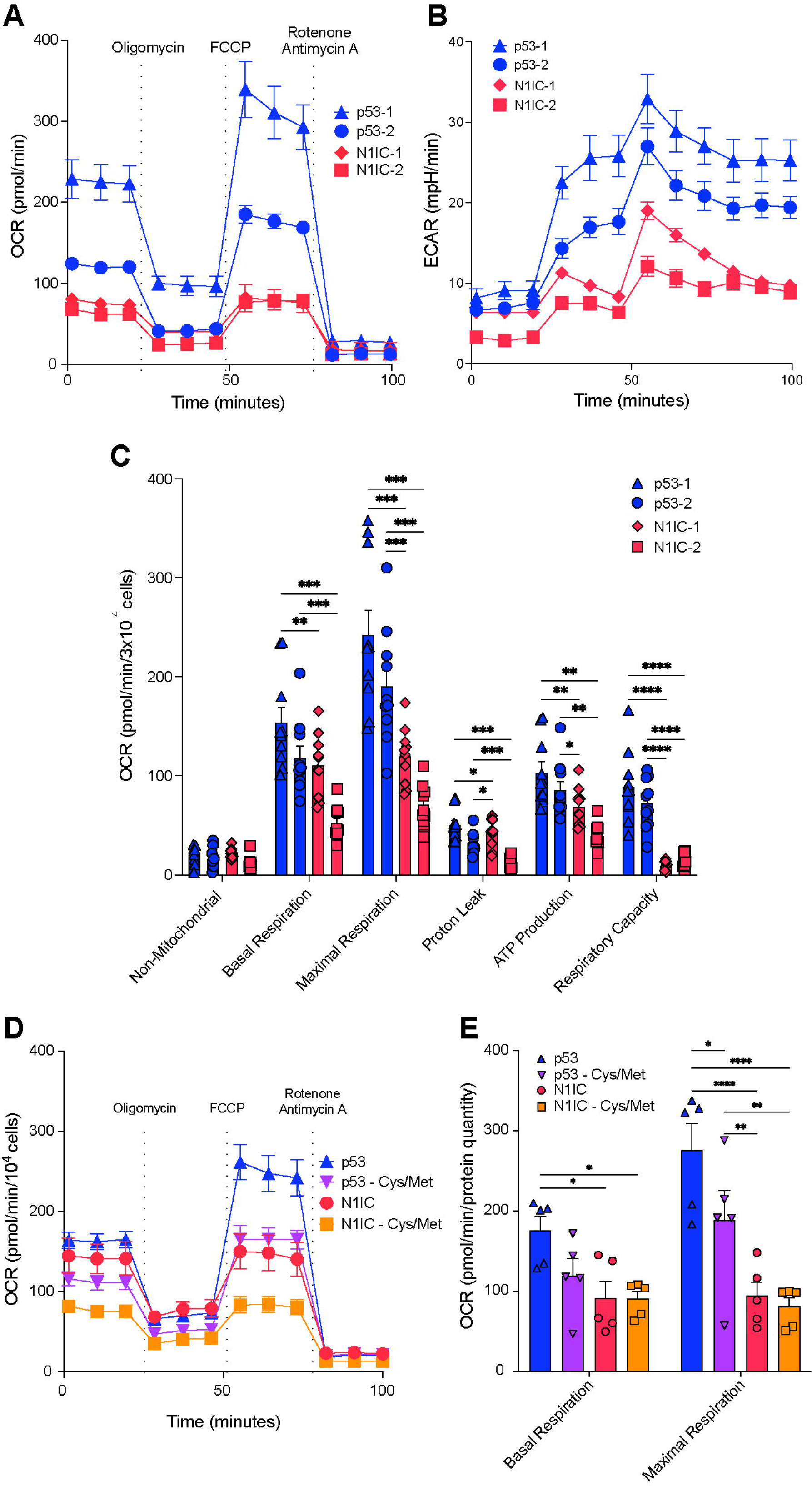
Differences in energetic metabolism between N1IC and p53 cells. A. Seahorse OCR analysis of two different p53 and N1IC cell lines demonstrating higher energetic metabolism in p53 tumor cells compared to N1IC tumor cells. Each point represents mean, error bars SEM of n=10 technical replicates. Representative experiment of n=4 independent experiments. B. Seahorse ECAR analysis suggesting increased glycolysis in p53 cells accounting for the higher OCR. Each point represents mean, error bars represent SEM of n=10 technical replicates. Representative experiment of n=4 independent experiments. C. Quantification of different Seahorse parameters demonstrating higher ATP production, higher respiratory capacity, and higher basal and maximal respiration in p53 tumor cells. Bar graph depicting means ± SEM. Normalized by protein quantification. Data from n=2 independent experiments, n=5 replicates per condition per experiment. D. Representative time course of Seahorse analysis of a p53/N1IC tumor cell pair with or without cysteine/methionine deprivation. Each point represents mean, errors bars represent SEM calculated from n=5 technical replicates. E. Quantification of Seahorse parameters demonstrating decreased basal respiration and maximal respiration after cysteine/methionine deprivation in p53 cells with less impact on N1IC cells. Normalized by protein quantification. Mean ± SEM from n=5 technical replicates. C, E P-values calculated by two-way ANOVA correcting for multiple comparisons by controlling the FDR using the 2-stage linear step-up procedure of Benjamini, Krieger, and Yekutieli. *p < 0.05; **p < 0.01; ***p < 0.001, ****p < 0.0001.

**Figure S6 (Associated with Figure 3):**
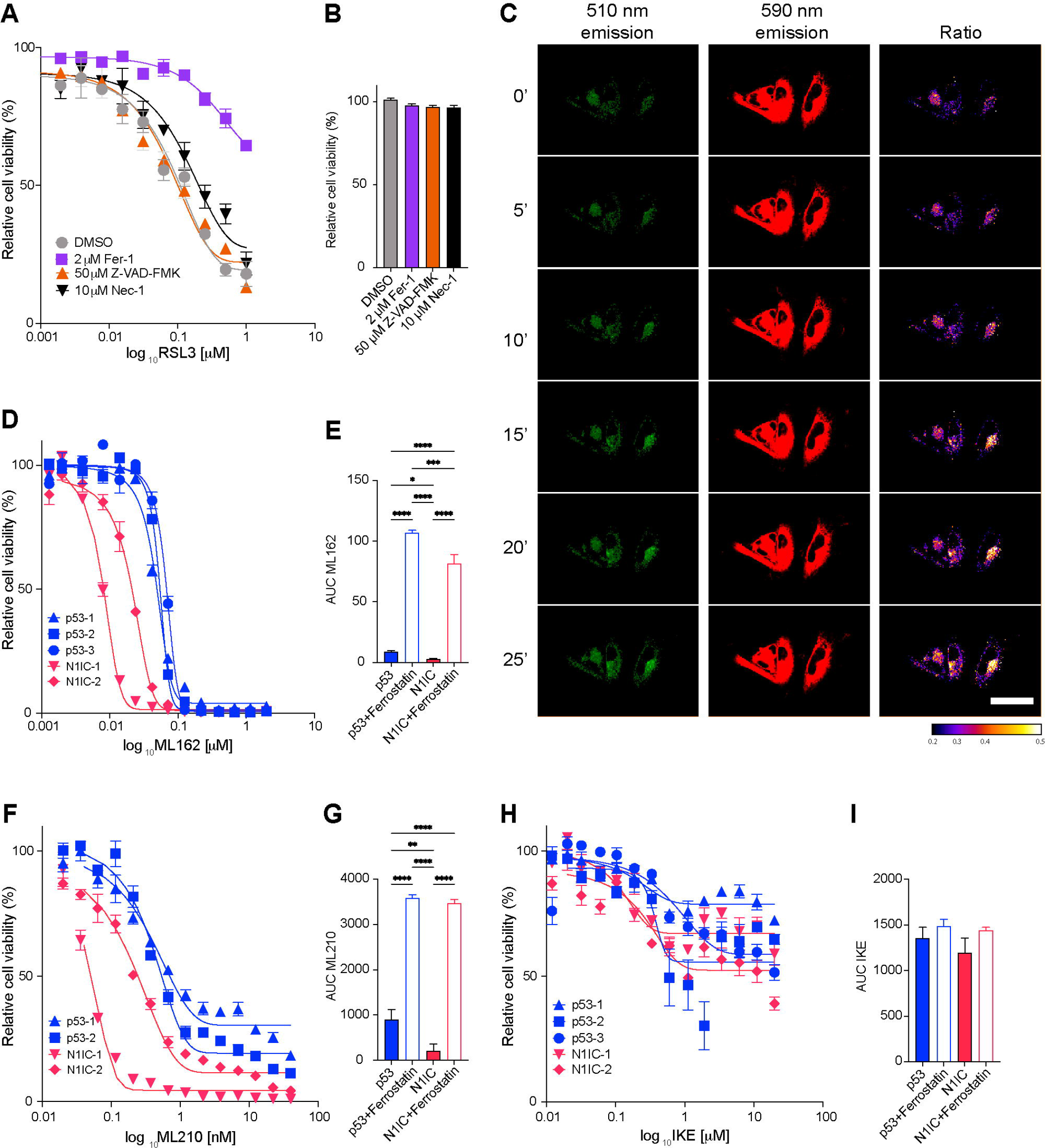
Increased sensitivity to Gpx4-driven but not Slc7a11-dependent ferroptosis of N1IC cells. A. Dose response curves to RSL3 in a N1IC cell line with ferroptosis, apoptosis and necroptosis inhibitors. Only the ferroptosis inhibitor Ferrostatin (Fer-1, 2 μm) induces partial rescue, while Z-VAD-FKM (50 μm) and Necrostatin-1 (Nec1, 10 μm) have no significant effect, demonstrating that RSL3 induces cell death via ferroptosis-specific mechanisms. B. Viability with ferroptosis, necroptosis and apoptosis inhibitors alone in the same N1IC cell line. Error bars represent mean ± SEM, n=3 technical replicates. Nonsignificant p-values not depicted. C. Live imaging with BODIPY-C11 stains demonstrating time course of 500 nM RSL3-induced ferroptosis in N1IC cell line. The 510/590 ratio, marker of increased lipid peroxidation secondary to ferroptosis, between 10 to 25 minutes after drug treatment. Representative experiment of n=3 independent experiments. Scale bar, 50 μm. D. Dose response curve for ML162, 24-hour treatment, normalized to DMSO. E. AUC without/with Ferrostatin, n=3 p53/n=2 N1IC cell lines. Error bars represent SEM. F. Dose response curves for ML210, 48-hour treatment, normalized to DMSO. G. AUC without/with Ferrostatin, n=2 p53/n=2 N1IC cell lines. Error bars represent SEM. H. Dose response curves for IKE, 24-hour treatment, normalized to DMSO. I. AUC without/with Ferrostatin, n=3 p53/n=2 N1IC cell lines. Error bars represent SEM. P-values in E, G, I calculated by ANOVA with FDR correction via Benjamini, Krieger, and Yekutieli. *Q < 0.05; **Q < 0.01; ***Q < 0.001, ****Q < 0.0001.

**Figure S7 (Associated with Figure 4):**
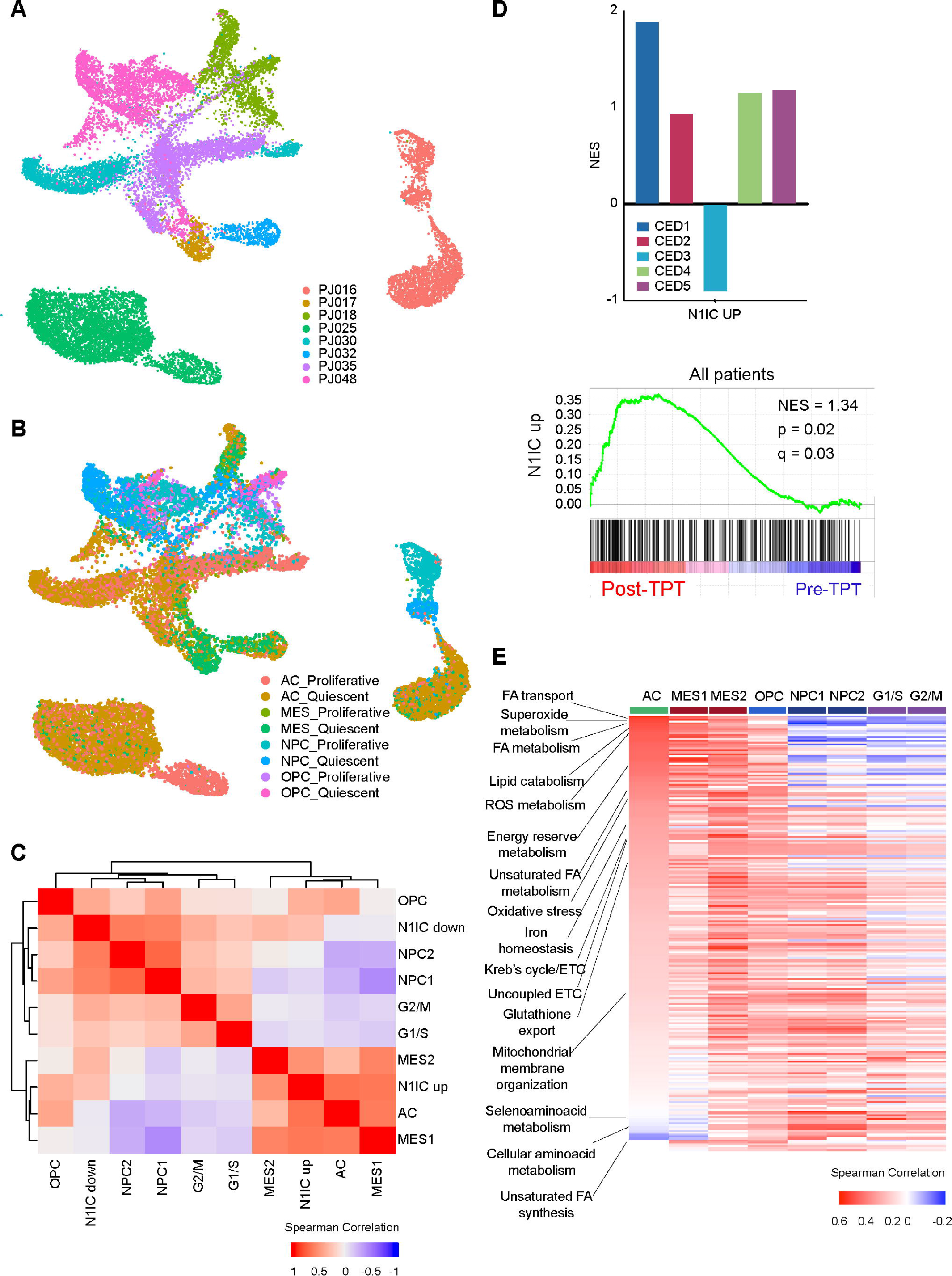
N1IC-like tumor populations in treatment-naïve and post-treatment human GBM. A-B. UMAP embedding of scRNAseq data from n=8 GBM patient samples described in (Yuan et al., 2018) colored by patient (A) and cell state + proliferation status (B). A. C. Heatmap of Spearman correlation index between the AUCell gene signatures generated from the Notch murine model (N1IC_up and N1IC_down) and Neftel cell state using the scRNAseq data in panels A-B. Note the clustering of the “N1IC_up” signature with the AC/MES signatures. The “N1IC_down” signature clusters with the NPC/OPC and proliferation signatures. B. D. Top: Bar plot demonstrating enrichment in the “N1IC_up” signature in 4 out of 5 patients undergoing chronic intratumoral delivery of topotecan in a recurrent GBM cohort (Spinazzi et al., 2022). Bottom: GSEA of the “N1IC_up” signature in all 5 patients combined. C. E. Heatmap of Spearman correlation between cell states and metabolic gene ontologies using the same scRNA-seq dataset used in panels A-C. Note divergent programs in the AC and NPC cell states, including upregulated mitochondrial programs in AC and enriched amino acid programs in NPC. Full list provided in **Table S6**.

**Figure S8 (Associated with Figure 4):**
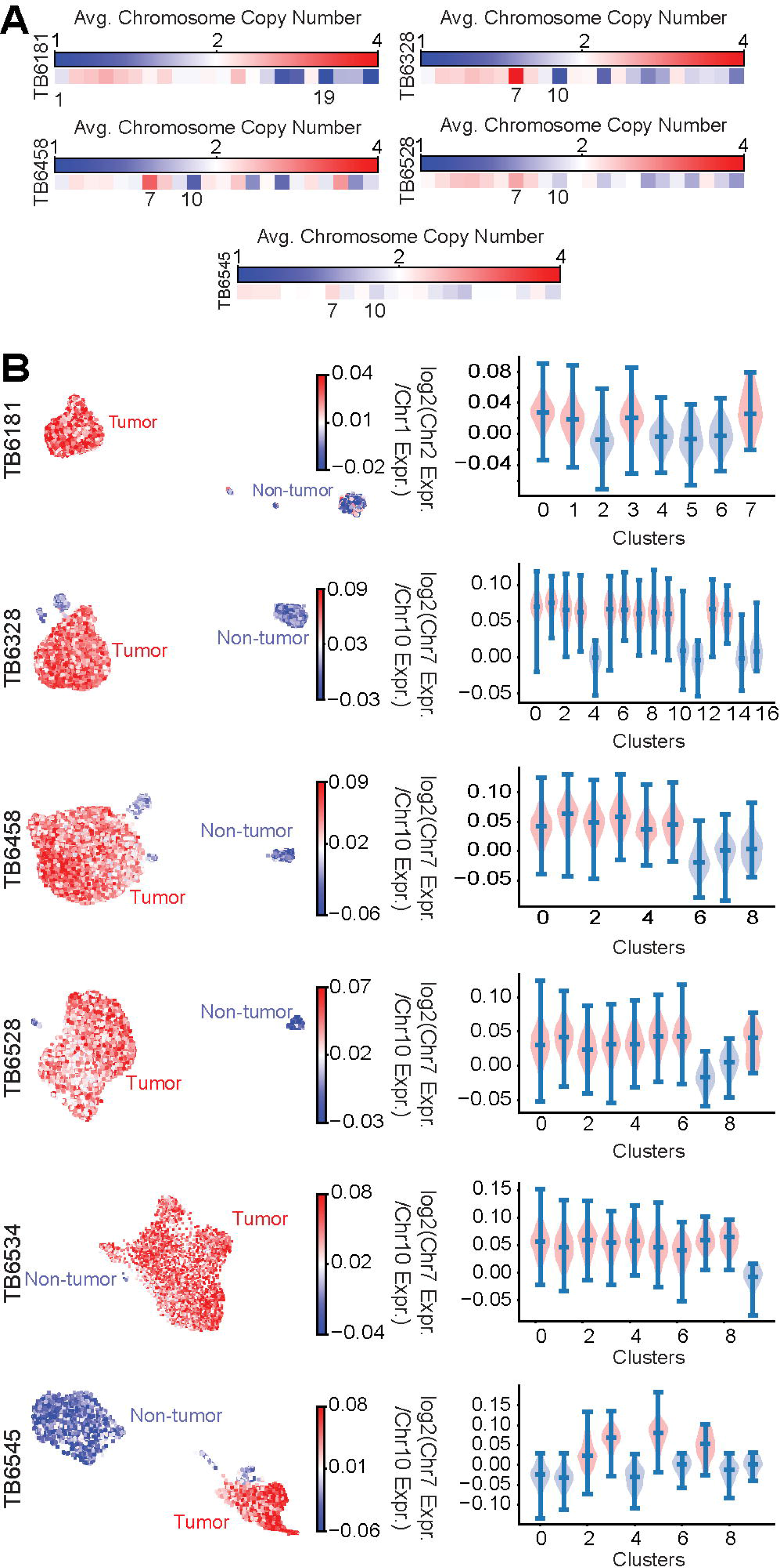
Identification of transformed populations in slice cultures via chromosome copy number alterations. A. Heatmaps showing the average chromosomal copy number from whole genome sequencing of patient samples used for slice cultures in **Figure 4**. B. Left: UMAP embeddings of scRNA-seq profiles from TB6181, TB6328, TB6458, TB6528, TB6534 and TB6545 slice cultures colored by the log-ratio of Chr. 2 to Chr. 1 average expression (TB6181) or by the log-ratio of Chr. 7 to Chr. 10 average expression (TB6328, TB6458, TB6528, TB6534 and TB6545). The high ratio (red) indicates malignant transformation. Right: Violin plots showing the distributions of log-ratios of Chr. 2 to Chr. 1 or Chr. 7 to Chr. 10 average expression for each Phenograph cluster identified for TB6181, TB6328, TB6458, TB6528, TB6534 and TB6545 slice cultures.

## Supplementary Tables

**Table S1:** Genetic murine glioma model scRNA-seq differentially expressed genes

**Table S2:** Common differentially expressed genes in persister cells and the N1IC model

**Table S3:** Murine model transcriptional metabolic programs

**Table S4:** Metabolic pathway analysis

**Table S5:** Lipid ontology analysis

**Table S6:** Correlations between cell state and metabolic programs in human GBM

**Table S7:** Clinical and pathologic patient characteristics of organotypic slice culture specimens

**Table S8:** Differentially expressed genes in treated acute slice cultures

