## Supplementary material for "A cell state specific metabolic vulnerability to GPX4-dependent ferroptosis in glioblastoma": Table S1

| **N1IC_up**  **(l2fc≥1)** | **N1IC_down (l2fc≤-1)** | **N1IC_up**  **(l2fc≥0.5)** | **N1IC_down**  **(l2fc≤-0.5)** | **N1IC_up**  **(l2fc>0)** | **N1IC_down**  **(l2fc<0)** |
| --- | --- | --- | --- | --- | --- |
| Abcg1 | Aard | 1500015O10Rik | 1700001O22Rik | 1110012L19Rik | 1700001O22Rik |
| Abracl | Abhd17c | 1700019D03Rik | 1810026J23Rik | 1500015O10Rik | 1810013L24Rik |
| Acot1 | Abhd2 | 1700025G04Rik | 2410131K14Rik | 1700019D03Rik | 1810026J23Rik |
| Acss1 | Adam22 | 2210013O21Rik | 2700081O15Rik | 1700025G04Rik | 2410131K14Rik |
| Agt | Agap3 | 2700029M09Rik | 3110035E14Rik | 1810037I17Rik | 2700081O15Rik |
| Aim1 | Ak5 | 2900011O08Rik | Aard | 2210013O21Rik | 3110035E14Rik |
| Akr1e1 | Akap7 | 44806 | Abce1 | 2700029M09Rik | Aak1 |
| Aldoc | Akt1s1 | Abcg1 | Abhd17c | 2900011O08Rik | Aard |
| Ankrd46 | Aldh1a3 | Abracl | Abhd2 | 44806 | Abce1 |
| Apbb1ip | Alyref | Acaa1a | Adam22 | 44811 | Abhd17c |
| Apobec1 | Amz1 | Aco2 | Adgrb2 | Abcf1 | Abhd2 |
| Arhgef28 | Anapc15 | Acot1 | Ado | Abcg1 | Acaca |
| Arpc5 | Anapc7 | Acss1 | Agap3 | Abracl | Adam22 |
| Atp10b | Ankib1 | Aebp1 | Ago2 | Acaa1a | Adgrb2 |
| B230118H07Rik | Ankle2 | Agt | Ajap1 | Aco2 | Ado |
| B2m | Ano8 | Aim1 | Ak5 | Acot1 | Agap3 |
| Bfsp2 | Ap1s1 | Akr1e1 | Akap7 | Acss1 | Ago2 |
| Bst2 | Aplp1 | Aldoc | Akirin2 | Actg1 | Ajap1 |
| Btbd17 | Arf6 | Ankrd46 | Akt1 | Aebp1 | Ak5 |
| C4b | Arhgap21 | Apbb1ip | Akt1s1 | Agt | Akap7 |
| Cacna1e | Arhgap33 | Apoa1bp | Aldh1a3 | Ahsa1 | Akirin2 |
| Cald1 | Arhgef11 | Apobec1 | Alyref | Aim1 | Akt1 |
| Capg | Arl8a | App | Amfr | Akr1e1 | Akt1s1 |
| Car8 | Armc10 | Arhgef28 | Amz1 | Aldoc | Aldh1a3 |
| Ccdc134 | Armc8 | Arpc2 | Anapc15 | Ankrd46 | Aldoa |
| Ccser2 | Armcx4 | Arpc5 | Anapc7 | Ap2m1 | Alyref |
| Cd200 | Arnt2 | Atp10b | Ank2 | Apbb1ip | Amfr |
| Chrm3 | Arpc3 | Atp1b2 | Ankib1 | Apoa1bp | Amz1 |
| Cited1 | Arrb1 | Atp6v1a | Ankle2 | Apobec1 | Anapc15 |
| Clvs2 | Arxes2 | Atp6v1e1 | Ankrd13a | App | Anapc7 |
| Cmtm5 | Asap2 | Atrx | Ankrd17 | Arhgef28 | Ank2 |
| Cnn2 | Ascl1 | Atxn10 | Ankrd28 | Arpc2 | Ankib1 |
| Commd7 | Asphd2 | B230118H07Rik | Ano8 | Arpc5 | Ankle2 |
| Coq3 | Aspscr1 | B2m | Ap1s1 | Atp10b | Ankrd13a |
| Cpped1 | Asxl1 | B9d1 | Ap2a1 | Atp1b2 | Ankrd17 |
| Cpq | Atp13a3 | Bcan | Ap2a2 | Atp5b | Ankrd28 |
| Creb3 | Atp2a2 | Bcas1 | Apba2 | Atp5c1 | Ano8 |
| Creg1 | Atp8b2 | Bfsp2 | Api5 | Atp5d | Ap1s1 |
| Cryab | Atxn2 | Bhlhe40 | Aplp1 | Atp5f1 | Ap2a1 |
| Csad | Bax | Blcap | Arf6 | Atp5j | Ap2a2 |
| Csrp1 | Bbc3 | Bst2 | Arhgap21 | Atp6v1a | Apba2 |
| Cst3 | BC030336 | Btbd17 | Arhgap33 | Atp6v1e1 | Api5 |
| Ctnnbip1 | Brdt | C4b | Arhgap35 | Atp6v1g1 | Aplp1 |
| Cyb5r3 | Btbd1 | Cacna1e | Arhgef11 | Atrx | Arf6 |
| Cyp2j9 | C1qb | Cald1 | Arl8a | Atxn10 | Arhgap21 |
| Cyr61 | C2cd3 | Capg | Armc10 | Atxn7l3b | Arhgap33 |
| Cystm1 | Cacna1c | Car8 | Armc8 | B230118H07Rik | Arhgap35 |
| Ddx58 | Cacna1h | Casp3 | Armcx4 | B2m | Arhgef11 |
| Dhrs1 | Cacng7 | Ccdc134 | Arnt2 | B9d1 | Arl8a |
| Dlc1 | Cadm4 | Ccdc174 | Arpc3 | Bcan | Armc10 |
| Dtx3l | Cand1 | Ccnd3 | Arrb1 | Bcas1 | Armc8 |
| Dusp15 | Cand2 | Ccser2 | Arxes2 | Bfsp2 | Armcx4 |
| Echdc2 | Cblb | Cd200 | Asap2 | Bhlhe40 | Arnt2 |
| Edf1 | Ccdc38 | Cdc42se2 | Ascl1 | Blcap | Arpc3 |
| Ehd3 | Ccl3 | Cdc5l | Asphd2 | Brk1 | Arrb1 |
| Eif2ak2 | Ccl4 | Cdk11b | Aspscr1 | Bst2 | Arxes2 |
| Eif2s3y | Ccna2 | Cetn2 | Asxl1 | Btbd17 | Asap2 |
| En1 | Ccnk | Cfap36 | Asxl2 | C4b | Ascl1 |
| Enpp2 | Ccz1 | Chrm3 | Atp13a3 | Cacna1e | Asphd2 |
| Erbb3 | Cd93 | Cited1 | Atp1b1 | Cald1 | Aspscr1 |
| Exosc4 | Cdk14 | Clu | Atp2a2 | Calm1 | Asxl1 |
| Fabp7 | Cdk17 | Clvs2 | Atp8b2 | Calm2 | Asxl2 |
| Fam102a | Cdk2ap1 | Cmtm5 | Atxn2 | Capg | Atp13a3 |
| Fam118a | Cdk5r1 | Cnih4 | Atxn2l | Car8 | Atp1b1 |
| Fam210b | Cdkn2a | Cnn2 | Axl | Casp3 | Atp2a2 |
| Fam46a | Cdkn2aip | Cnn3 | Bag6 | Cbx5 | Atp8b2 |
| Fermt2 | Cep89 | Cnp | Bax | Ccdc134 | Atxn2 |
| Fkbp9 | Chek2 | Cnpy2 | Baz1b | Ccdc174 | Atxn2l |
| Flt1 | Chka | Col16a1 | Bbc3 | Ccnd2 | Axl |
| Flywch1 | Chpf2 | Commd7 | BC030336 | Ccnd3 | B3gat1 |
| Fxyd2 | Chpt1 | Cope | Bcl7a | Ccser2 | Bag6 |
| Fzd2 | Chrna4 | Copz1 | Bex1 | Cd200 | Bax |
| Gbp2 | Chst11 | Coq3 | Brd1 | Cd302 | Baz1b |
| Gbp3 | Chst2 | Cpe | Brdt | Cd47 | Bbc3 |
| Gbp6 | Chuk | Cpped1 | Bri3bp | Cd63 | BC030336 |
| Gbp7 | Cic | Cpq | Bscl2 | Cd9 | Bcl7a |
| Gm266 | Cited2 | Creb3 | Btbd1 | Cdc42se2 | Bex1 |
| Gm43302 | Clcn4 | Creg1 | C1qb | Cdc5l | Birc6 |
| Gm4951 | Cldn12 | Crip2 | C2cd3 | Cdk11b | Brd1 |
| Gng11 | Clptm1 | Cryab | C330027C09Rik | Cdk4 | Brdt |
| Gnpda2 | Cnot11 | Csad | C530008M17Rik | Cetn2 | Bri3 |
| Gpt2 | Cnot3 | Csdc2 | Cacna1c | Cfap20 | Bri3bp |
| Gpx7 | Cnpy4 | Csrp1 | Cacna1h | Cfap36 | Bscl2 |
| H2-Ab1 | Cox16 | Cst3 | Cacng7 | Cfdp1 | Btbd1 |
| H2-D1 | Crebzf | Ctnnbip1 | Cadm4 | Chrm3 | C1qb |
| H2-K1 | Crls1 | Cyb5r3 | Calm3 | Cited1 | C1qc |
| H2-Q4 | Cse1l | Cyp2j9 | Camk2b | Clu | C2cd3 |
| H2-Q6 | Ctbp1 | Cyr61 | Camsap2 | Clvs2 | C330027C09Rik |
| H2-Q7 | Ctr9 | Cystm1 | Cand1 | Cmtm5 | C530008M17Rik |
| H2-T22 | Ctsd | Dct | Cand2 | Cnbp | Cab39 |
| H2-T23 | Cttn | Ddx26b | Casc4 | Cnih1 | Cacna1c |
| Has2 | Cux1 | Ddx58 | Cblb | Cnih4 | Cacna1h |
| Heg1 | Cxcl14 | Decr1 | Ccdc38 | Cnn2 | Cacng7 |
| Hes1 | Cyfip1 | Dek | Ccdc92 | Cnn3 | Cadm4 |
| Hes5 | D5Ertd579e | Dhrs1 | Ccl3 | Cnp | Calm3 |
| Hey1 | Daglb | Dlc1 | Ccl4 | Cnpy2 | Camk2b |
| Hmcn1 | Dcx | Dnal4 | Ccna2 | Col16a1 | Camsap2 |
| Hmgcl | Deaf1 | Dnttip2 | Ccnd1 | Commd7 | Cand1 |
| Hpgd | Dennd2a | Dpy30 | Ccni | Cope | Cand2 |
| Id3 | Dennd5a | Dscr3 | Ccnk | Copz1 | Cars |
| Ifi27l2a | Dennd5b | Dtx3l | Ccnt2 | Coq3 | Casc4 |
| Ifi35 | Dgat2 | Dusp15 | Ccz1 | Cotl1 | Cblb |
| Ifi47 | Dhcr7 | Dync1i2 | Cd2bp2 | Cox7b | Ccdc38 |
| Ifih1 | Dhx32 | Dynlrb1 | Cd93 | Cpe | Ccdc92 |
| Ifit1 | Dido1 | Ebpl | Cdk13 | Cpped1 | Ccl3 |
| Ifit3 | Dip2b | Echdc2 | Cdk14 | Cpq | Ccl4 |
| Ift20 | Dlg1 | Edf1 | Cdk17 | Creb3 | Ccna2 |
| Igfbp5 | Dlk1 | Ehd3 | Cdk2ap1 | Creg1 | Ccnd1 |
| Igtp | Dll3 | Eif2ak2 | Cdk5r1 | Crip2 | Ccni |
| Il11ra1 | Dmpk | Eif2s3y | Cdkn2a | Cryab | Ccnk |
| Il18 | Dmwd | Eif4a2 | Cdkn2aip | Csad | Ccnt2 |
| Irf7 | Dnajc13 | Eif5b | Cep89 | Csdc2 | Cct2 |
| Irgm1 | Dock1 | Emc2 | Chd2 | Csrp1 | Ccz1 |
| Irgm2 | Dock11 | Emc4 | Chek2 | Cst3 | Cd2bp2 |
| Kbtbd11 | Dyrk1a | Emid1 | Chka | Ctnnb1 | Cd93 |
| Kcnj16 | E2f5 | En1 | Chpf2 | Ctnnbip1 | Cdk13 |
| Keap1 | Edrf1 | Enpp2 | Chpt1 | Ctsl | Cdk14 |
| Kit | Eef1a2 | Erbb3 | Chrna4 | Cyb5r3 | Cdk17 |
| Ldhb | Efcc1 | Erlec1 | Chst11 | Cyp2j9 | Cdk2ap1 |
| Lgals3bp | Eid1 | Exosc4 | Chst2 | Cyr61 | Cdk5r1 |
| Lgals9 | Eid2 | Ezr | Chuk | Cystm1 | Cdkn2a |
| Lims2 | Eif2s3x | F3 | Churc1 | Dct | Cdkn2aip |
| Litaf | Eif3b | Fabp5 | Cic | Ddit4 | Cep89 |
| Lman1 | Eif3f | Fabp7 | Cited2 | Ddx26b | Chd2 |
| Lnx1 | Eif4g2 | Fam102a | Clcn4 | Ddx58 | Chek2 |
| Lpar4 | Elfn1 | Fam118a | Cldn12 | Decr1 | Chfr |
| Lrp4 | Emc10 | Fam210b | Clic1 | Dek | Chka |
| Lxn | Eme1 | Fam213a | Clip3 | Desi1 | Chpf2 |
| Map1lc3b | Ep400 | Fam213b | Clns1a | Dhrs1 | Chpt1 |
| Map7d2 | Ephx4 | Fam46a | Clptm1 | Dlc1 | Chrna4 |
| Mbnl2 | Epn1 | Fermt2 | Cltc | Dnaja1 | Chst11 |
| Mgll | Erf | Fez1 | Cnot10 | Dnajc8 | Chst2 |
| Mgst1 | Eri2 | Fkbp11 | Cnot11 | Dnal4 | Chuk |
| Mical1 | Ermp1 | Fkbp1a | Cnot3 | Dnttip2 | Churc1 |
| Mmp14 | Evl | Fkbp9 | Cnpy4 | Dpy30 | Cic |
| Mphosph8 | Fam101a | Flt1 | Col11a1 | Dscr3 | Cited2 |
| Mrpl55 | Fam166a | Flywch1 | Cox16 | Dtx3l | Clcn4 |
| Mxra8 | Fam174b | Fxyd2 | Cpeb4 | Dusp15 | Cldn11 |
| Myl9 | Fam177a | Fzd2 | Cpsf4 | Dusp6 | Cldn12 |
| Nbn | Fam179b | Gadd45gip1 | Crebzf | Dync1i2 | Clic1 |
| Ncald | Fam19a2 | Gbp2 | Crls1 | Dynll1 | Clip2 |
| Ndufaf2 | Fam208a | Gbp3 | Crtc3 | Dynlrb1 | Clip3 |
| Necap2 | Fam221b | Gbp6 | Cse1l | Ebpl | Clns1a |
| Nupr1 | Far1 | Gbp7 | Ctbp1 | Echdc2 | Clptm1 |
| Oas1a | Fasn | Gm266 | Ctr9 | Edf1 | Cltc |
| Oasl2 | Fau | Gm2a | Ctsd | Ehd3 | Cnot10 |
| Oxct1 | Fbl | Gm43302 | Cttn | Eif1 | Cnot11 |
| Palmd | Fbrsl1 | Gm4951 | Ctxn1 | Eif2ak2 | Cnot3 |
| Paqr4 | Fcer1g | Gng11 | Cul3 | Eif2s2 | Cnpy4 |
| Paqr7 | Fgfrl1 | Gnpda2 | Cux1 | Eif2s3y | Col11a1 |
| Parp14 | Fnbp4 | Golgb1 | Cxcl14 | Eif3d | Cox16 |
| Parp9 | Fndc4 | Gpm6b | Cyfip1 | Eif4a2 | Cpeb4 |
| Pcdhb14 | Fosl2 | Gpr180 | Cyth2 | Eif5b | Cpsf4 |
| Pcdhb6 | Fry | Gpr37l1 | D5Ertd579e | Emc2 | Crebzf |
| Pck2 | Fscn1 | Gpt2 | Daglb | Emc4 | Crls1 |
| Pcolce | Fstl3 | Gpx1 | Dazap1 | Emid1 | Crtc3 |
| Pdpn | Ftl1 | Gpx7 | Dcx | En1 | Cse1l |
| Pdrg1 | Fus | Gtf3c6 | Ddx17 | Enpp2 | Ctbp1 |
| Pex26 | Fzd1 | Guk1 | Ddx39b | Eny2 | Ctr9 |
| Pgm2 | Gab2 | H2-Ab1 | Ddx3x | Erbb3 | Ctsd |
| Pigk | Gabpa | H2-D1 | Deaf1 | Erlec1 | Cttn |
| Pla2g4a | Gadl1 | H2-K1 | Dennd2a | Exosc4 | Ctxn1 |
| Plin3 | Gas6 | H2-Q4 | Dennd5a | Ezr | Cul3 |
| Plp1 | Gcn1l1 | H2-Q6 | Dennd5b | F3 | Cux1 |
| Pmel | Get4 | H2-Q7 | Dgat2 | Fabp5 | Cxcl14 |
| Ppp1r14c | Gm10094 | H2-T22 | Dhcr7 | Fabp7 | Cyfip1 |
| Prss35 | Gm10116 | H2-T23 | Dhx32 | Fam102a | Cyth2 |
| Psma7 | Gm10131 | Has2 | Dido1 | Fam118a | D5Ertd579e |
| Psmb8 | Gm10260 | Heg1 | Dip2b | Fam162a | Daglb |
| Psmb9 | Gm10282 | Hes1 | Dlg1 | Fam210b | Dazap1 |
| Psme1 | Gm10320 | Hes5 | Dlk1 | Fam213a | Dcx |
| Rassf4 | Gm11273 | Hey1 | Dll3 | Fam213b | Ddx17 |
| Rfc1 | Gm12184 | Hint2 | Dmpk | Fam46a | Ddx39b |
| Rnf31 | Gm12355 | Hist1h1c | Dmwd | Fermt2 | Ddx3x |
| Rpa2 | Gm12728 | Hist1h2bc | Dnajb6 | Fez1 | Deaf1 |
| Rsrc1 | Gm13889 | Hmcn1 | Dnajc13 | Fkbp11 | Dennd2a |
| Rtp4 | Gm15013 | Hmg20b | Dock1 | Fkbp1a | Dennd5a |
| Runx3 | Gm17087 | Hmgcl | Dock11 | Fkbp9 | Dennd5b |
| S100a11 | Gm4707 | Hpgd | Dock3 | Flt1 | Dgat2 |
| S100b | Gm5148 | Hsbp1 | Dpysl2 | Flywch1 | Dhcr7 |
| Samd9l | Gm6525 | Id3 | Dpysl3 | Frg1 | Dhx32 |
| Sat1 | Gm9774 | Ier3ip1 | Dr1 | Fuca1 | Dido1 |
| Scamp2 | Gm9803 | Ifi27 | Dynlt1c | Fxyd2 | Dip2b |
| Scrg1 | Gm9833 | Ifi27l2a | Dyrk1a | Fzd2 | Dlg1 |
| Serpine2 | Gm9844 | Ifi35 | E130309D02Rik | Gadd45gip1 | Dlg4 |
| Sh3bgr | Gna12 | Ifi44 | E2f5 | Gbp2 | Dlk1 |
| Sh3bgrl3 | Golga3 | Ifi47 | Edrf1 | Gbp3 | Dll3 |
| Shc4 | Gpc1 | Ifih1 | Eed | Gbp6 | Dmpk |
| Slc26a2 | Gprc5b | Ifit1 | Eef1a1 | Gbp7 | Dmwd |
| Slc9a3r1 | Grin1 | Ifit3 | Eef1a2 | Glul | Dnajb6 |
| Sorl1 | Gsk3a | Ift20 | Efcc1 | Gm266 | Dnajc13 |
| Sparc | Gtf3c1 | Igfbp5 | Eid1 | Gm2a | Dock1 |
| Spats2l | Hcfc1 | Igtp | Eid2 | Gm43302 | Dock11 |
| Srek1ip1 | Hip1r | Il11ra1 | Eif2s3x | Gm4951 | Dock3 |
| Srpr | Hipk1 | Il18 | Eif3b | Gng11 | Dpysl2 |
| Stat1 | Hirip3 | Immp1l | Eif3f | Gng12 | Dpysl3 |
| Synpr | Hmga1 | Irf7 | Eif4a1 | Gnpda2 | Dr1 |
| Tap1 | Hmga1-rs1 | Irgm1 | Eif4g2 | Golgb1 | Dscam |
| Tap2 | Hmox1 | Irgm2 | Eif4h | Gpm6b | Dynlt1c |
| Tax1bp3 | Hnrnpa0 | Itm2b | Elavl1 | Gpr180 | Dyrk1a |
| Ten1 | Hnrnpd | Itm2c | Elavl3 | Gpr37l1 | E130309D02Rik |
| Tgtp2 | Hnrnpl | Kbtbd11 | Elfn1 | Gpt2 | E2f5 |
| Thbs3 | Hnrnpul1 | Kcnj10 | Emc10 | Gpx1 | Edrf1 |
| Timp1 | Hook2 | Kcnj16 | Eme1 | Gpx4 | Eed |
| Timp3 | Hsd17b11 | Keap1 | Emsy | Gpx7 | Eef1a1 |
| Tk1 | Hsph1 | Kit | Ep400 | Gtf3c6 | Eef1a2 |
| Tlr3 | Htra3 | Lage3 | Ephb2 | Guk1 | Efcc1 |
| Tmc6 | Htt | Lamp2 | Ephx4 | H2-Ab1 | Eid1 |
| Tmem100 | Hyou1 | Lamtor5 | Epn1 | H2-D1 | Eid2 |
| Tmem29 | Ick | Lbh | Erf | H2-K1 | Eif2ak1 |
| Tmem50a | Ier5l | Ldhb | Eri2 | H2-Q4 | Eif2s3x |
| Tnfaip2 | Iglon5 | Lef1 | Ermp1 | H2-Q6 | Eif3b |
| Tnni1 | Il1a | Leprot | Evl | H2-Q7 | Eif3f |
| Tpd52l1 | Ints1 | Lgals3bp | Fam101a | H2-T22 | Eif4a1 |
| Trf | Ipo7 | Lgals9 | Fam120a | H2-T23 | Eif4g2 |
| Trim2 | Ireb2 | Lims2 | Fam166a | H2afz | Eif4h |
| Tspan17 | Irf2bp1 | Litaf | Fam174b | H3f3b | Elavl1 |
| Tspo | Irgq | Lman1 | Fam177a | Has2 | Elavl3 |
| Txndc15 | Isg20l2 | Lnx1 | Fam179b | Heg1 | Elfn1 |
| Usp18 | Jag1 | Lpar4 | Fam19a2 | Hes1 | Emc10 |
| Vit | Jmjd1c | Lrp4 | Fam208a | Hes5 | Eme1 |
| Xaf1 | Kansl1 | Luc7l3 | Fam219a | Hey1 | Emsy |
| Zcchc17 | Kbtbd2 | Lxn | Fam221b | Hhipl2 | Ep400 |
| Zfhx4 | Kcnip1 | Mad2l2 | Fam76b | Hint2 | Epb41l2 |
| Zfp36l1 | Kctd5 | Maf1 | Fam91a1 | Hist1h1c | Ephb2 |
| Zfp87 | Kdm2b | Maged2 | Far1 | Hist1h2bc | Ephx4 |
| Zfyve21 | Kdm3b | Mageh1 | Fasn | Hmcn1 | Epn1 |
| 1500015O10Rik | Khsrp | Map1lc3b | Fau | Hmg20b | Erf |
| 1700019D03Rik | Kif14 | Map7d2 | Fbl | Hmgcl | Eri2 |
| 1700025G04Rik | Klf13 | Mbnl2 | Fbrsl1 | Hn1 | Ermp1 |
|  | Klhl15 | Mcam | Fbxl3 | Hpgd | Evl |
|  | Klhl9 | Mdh1 | Fcer1g | Hsbp1 | Ewsr1 |
|  | Kmt2d | Med19 | Fcho2 | Hsp90ab1 | Faf1 |
|  | Kmt5a | Mgll | Fchsd2 | Hsp90b1 | Fam101a |
|  | Kpna2 | Mgst1 | Fgfrl1 | Hspa5 | Fam120a |
|  | Kpna4 | Mical1 | Fmr1 | Id3 | Fam166a |
|  | Krit1 | Mmp14 | Fnbp4 | Ier3ip1 | Fam174b |
|  | Larp1 | Mocs2 | Fndc3a | Ifi27 | Fam177a |
|  | Lars2 | Mpc1 | Fndc4 | Ifi27l2a | Fam179b |
|  | Leng8 | Mphosph8 | Fosl2 | Ifi35 | Fam19a2 |
|  | Lilrb4a | Mrpl18 | Fry | Ifi44 | Fam208a |
|  | Lin54 | Mrpl19 | Fscn1 | Ifi47 | Fam219a |
|  | Lin7a | Mrpl40 | Fstl3 | Ifih1 | Fam221b |
|  | Lmnb1 | Mrpl55 | Ftl1 | Ifit1 | Fam76b |
|  | Lmtk2 | Mrps33 | Fus | Ifit3 | Fam91a1 |
|  | Lrp11 | Msrb1 | Fzd1 | Ift20 | Far1 |
|  | Lrp3 | Mxra8 | Gab2 | Igfbp5 | Fasn |
|  | Lrrc4b | Myl12b | Gabpa | Igsf3 | Fau |
|  | Lrrfip1 | Myl9 | Gadl1 | Igtp | Fbl |
|  | Lrwd1 | Nans | Gas6 | Il11ra1 | Fbrsl1 |
|  | Lsm14a | Nbn | Gatad1 | Il18 | Fbxl3 |
|  | Lsm7 | Ncald | Gatad2a | Immp1l | Fcer1g |
|  | Ltbp4 | Ndrg2 | Gclm | Irf7 | Fcho2 |
|  | Map1lc3a | Ndufa4 | Gcn1l1 | Irf9 | Fchsd2 |
|  | Map3k10 | Ndufa5 | Get4 | Irgm1 | Fgfrl1 |
|  | Map3k12 | Ndufaf2 | Ggt7 | Irgm2 | Fmr1 |
|  | Map4k5 | Ndufb10 | Gins1 | Itm2b | Fnbp4 |
|  | Map6 | Ndufb5 | Gls | Itm2c | Fndc3a |
|  | Mapk12 | Ndufs4 | Gltp | Kank1 | Fndc4 |
|  | Mapk6 | Ndufs5 | Glud1 | Kbtbd11 | Fosl2 |
|  | Mark4 | Necap2 | Gm10094 | Kcnj10 | Fry |
|  | Mast2 | Nes | Gm10116 | Kcnj16 | Fscn1 |
|  | Matn4 | Ngly1 | Gm10131 | Keap1 | Fstl3 |
|  | Maz | Nmi | Gm10260 | Kit | Ftl1 |
|  | Mcm7 | Nsrp1 | Gm10282 | Lage3 | Fubp1 |
|  | Med25 | Nupr1 | Gm10320 | Lamp2 | Fus |
|  | Mex3b | Nusap1 | Gm11273 | Lamtor5 | Fzd1 |
|  | Mib1 | Oas1a | Gm12184 | Lbh | Gab2 |
|  | Mki67 | Oasl2 | Gm12355 | Ldhb | Gabpa |
|  | Mlip | Oxct1 | Gm12728 | Lef1 | Gadl1 |
|  | Mmp17 | Palmd | Gm13889 | Leprot | Gas6 |
|  | Mms19 | Paqr4 | Gm15013 | Lgals3bp | Gatad1 |
|  | Mn1 | Paqr7 | Gm17087 | Lgals9 | Gatad2a |
|  | Morc4 | Parp14 | Gm2000 | Lims2 | Gclm |
|  | Mphosph9 | Parp9 | Gm4707 | Litaf | Gcn1l1 |
|  | Mrpl23 | Pbxip1 | Gm5148 | Lman1 | Get4 |
|  | Mta1 | Pcdhb14 | Gm6525 | Lmo4 | Ggt7 |
|  | Mthfd2l | Pcdhb6 | Gm9774 | Lnx1 | Gins1 |
|  | Myc | Pck2 | Gm9803 | Lpar4 | Gls |
|  | Myef2 | Pcolce | Gm9833 | Lrp4 | Gltp |
|  | Myt1 | Pdcd6 | Gm9844 | Luc7l3 | Glud1 |
|  | Nav2 | Pdpn | Gna12 | Lxn | Gm10094 |
|  | Nav3 | Pdrg1 | Gnb2 | Mad2l2 | Gm10116 |
|  | Ncor2 | Pex26 | Golga3 | Maf1 | Gm10131 |
|  | Nell2 | Pfdn5 | Gpc1 | Maged2 | Gm10260 |
|  | Nme2 | Pgm2 | Gprc5b | Mageh1 | Gm10282 |
|  | Nomo1 | Phax | Grin1 | Map1lc3b | Gm10320 |
|  | Nova1 | Pigk | Grtp1 | Map7d2 | Gm11273 |
|  | Nt5dc2 | Pik3ip1 | Gsk3a | Mbnl2 | Gm12184 |
|  | Ntn1 | Pla2g4a | Gsx1 | Mcam | Gm12355 |
|  | Ntrk3 | Plin3 | Gtf2h3 | Mdh1 | Gm12728 |
|  | Numbl | Plp1 | Gtf3c1 | Med19 | Gm13889 |
|  | Nup98 | Pmel | Gtf3c2 | Mgll | Gm15013 |
|  | Olfm2 | Pmvk | Hcfc1 | Mgst1 | Gm17087 |
|  | Oraov1 | Pnkd | Hdac2 | Mical1 | Gm2000 |
|  | Orc5 | Polr2b | Hdgf | Mmp14 | Gm4707 |
|  | Otud6b | Polr3h | Hdgfrp3 | Mocs2 | Gm5148 |
|  | P2rx7 | Polr3k | Hip1r | Morf4l1 | Gm6525 |
|  | Paf1 | Pon2 | Hipk1 | Mpc1 | Gm9774 |
|  | Paip1 | Ppp1r14c | Hirip3 | Mphosph8 | Gm9803 |
|  | Pan3 | Prmt2 | Hmbox1 | Mrpl13 | Gm9833 |
|  | Patz1 | Prss35 | Hmga1 | Mrpl15 | Gm9844 |
|  | Paxbp1 | Psma7 | Hmga1-rs1 | Mrpl18 | Gna12 |
|  | Paxip1 | Psmb1 | Hmga2 | Mrpl19 | Gnb2 |
|  | Pcf11 | Psmb10 | Hmgn2 | Mrpl28 | Golga3 |
|  | Pclo | Psmb2 | Hmox1 | Mrpl40 | Gpc1 |
|  | Pde3b | Psmb3 | Hnrnpa0 | Mrpl42 | Gprc5b |
|  | Pde8a | Psmb8 | Hnrnpa1 | Mrpl55 | Gramd1a |
|  | Pdgfa | Psmb9 | Hnrnpd | Mrps14 | Gria4 |
|  | Peak1 | Psmc3 | Hnrnpl | Mrps25 | Grin1 |
|  | Peg10 | Psmd10 | Hnrnpu | Mrps33 | Grtp1 |
|  | Phactr3 | Psmd12 | Hnrnpul1 | Mrps6 | Gsap |
|  | Phf2 | Psmd14 | Hook2 | Msrb1 | Gsk3a |
|  | Phyhipl | Psmd7 | Hras | Mthfd2 | Gspt1 |
|  | Phykpl | Psme1 | Hsd17b11 | Mxra8 | Gsx1 |
|  | Pianp | Ptn | Hsf2 | Mydgf | Gtf2h3 |
|  | Picalm | Rasa2 | Hspa4 | Myl12b | Gtf3c1 |
|  | Pih1d1 | Rassf4 | Hsph1 | Myl9 | Gtf3c2 |
|  | Pld3 | Rbm28 | Htra3 | Nans | H2afy |
|  | Plekha1 | Rfc1 | Htt | Nav1 | Hacd1 |
|  | Plekhm2 | Rhoc | Hyou1 | Nbn | Hcfc1 |
|  | Plk2 | Rnf31 | Ick | Ncald | Hdac2 |
|  | Plxna1 | Rpa2 | Ier5l | Ndrg2 | Hdgf |
|  | Pnmal1 | Rps19bp1 | Iglon5 | Ndufa1 | Hdgfrp3 |
|  | Podxl2 | Rps4x | Il1a | Ndufa13 | Hip1r |
|  | Pogz | Rsrc1 | Ino80e | Ndufa2 | Hipk1 |
|  | Ppard | Rsu1 | Inpp5f | Ndufa4 | Hirip3 |
|  | Ppfia1 | Rtp4 | Insig1 | Ndufa5 | Hmbox1 |
|  | Ppm1a | Runx3 | Ints1 | Ndufaf2 | Hmga1 |
|  | Ppm1d | S100a11 | Ipo7 | Ndufb10 | Hmga1-rs1 |
|  | Ppm1g | S100b | Ireb2 | Ndufb11 | Hmga2 |
|  | Ppp1cb | Samd9l | Irf2bp1 | Ndufb5 | Hmgn2 |
|  | Ppp1cc | Sat1 | Irgq | Ndufb7 | Hmox1 |
|  | Ppp1r15a | Scamp2 | Iscu | Ndufb8 | Hnrnpa0 |
|  | Ppp1r9a | Scg5 | Isg20l2 | Ndufc1 | Hnrnpa1 |
|  | Ppp2r2c | Scp2 | Jag1 | Ndufs2 | Hnrnpa2b1 |
|  | Ppp2r2d | Scrg1 | Jmjd1c | Ndufs4 | Hnrnpd |
|  | Ppp6r1 | Sdc4 | Kansl1 | Ndufs5 | Hnrnpl |
|  | Prex2 | Sdf4 | Kbtbd2 | Ndufs6 | Hnrnpu |
|  | Prkar2b | Sdhc | Kcnd3 | Necap2 | Hnrnpul1 |
|  | Prkcb | Sec11c | Kcnip1 | Nes | Hook2 |
|  | Prmt1 | Sec13 | Kctd5 | Nfib | Hras |
|  | Prmt3 | Sec61b | Kdelr1 | Nfu1 | Hsd17b11 |
|  | Prpf8 | Sec62 | Kdelr2 | Ngly1 | Hsf2 |
|  | Prr7 | Serf1 | Kdm1a | Nhp2l1 | Hspa4 |
|  | Prrc2a | Serpine2 | Kdm2a | Nim1k | Hsph1 |
|  | Psmd13 | Sf3a3 | Kdm2b | Nmi | Htra3 |
|  | Ptbp1 | Sh3bgr | Kdm3b | Nnat | Htt |
|  | Ptms | Sh3bgrl3 | Kdm6b | Npc2 | Hyou1 |
|  | Ptov1 | Shc4 | Khdrbs1 | Nsrp1 | Ick |
|  | Ptp4a1 | Shisa5 | Khdrbs3 | Nupr1 | Ier5l |
|  | Ptpn12 | Sin3b | Khsrp | Nusap1 | Igfbp2 |
|  | Rab35 | Slc25a4 | Kif14 | Oas1a | Iglon5 |
|  | Rab3b | Slc26a2 | Kif18b | Oasl2 | Il1a |
|  | Rab6a | Slc9a3r1 | Klf13 | Oxct1 | Ino80e |
|  | Rac1 | Sorl1 | Klhdc2 | Pabpc1 | Inpp5f |
|  | Rad23b | Sparc | Klhl15 | Palmd | Insig1 |
|  | Ralgapb | Sparcl1 | Klhl7 | Paqr4 | Ints1 |
|  | Raly | Spats2l | Klhl9 | Paqr7 | Ipo5 |
|  | Ranbp6 | Srek1ip1 | Kmt2b | Parp14 | Ipo7 |
|  | Ranbp9 | Srp9 | Kmt2c | Parp9 | Iqgap1 |
|  | Rasgrf1 | Srpr | Kmt2d | Pbxip1 | Ireb2 |
|  | Rasl2-9 | St13 | Kmt5a | Pcdhb14 | Irf2bp1 |
|  | Rbm15b | Stat1 | Kpna2 | Pcdhb6 | Irgq |
|  | Rbm33 | Sub1 | Kpna4 | Pck2 | Iscu |
|  | Rbm45 | Svbp | Kpnb1 | Pcna | Isg20l2 |
|  | Rbm8a2 | Synpr | Kras | Pcolce | Jag1 |
|  | Rcc1 | Tagln3 | Krit1 | Pdcd6 | Jmjd1c |
|  | Rdh13 | Tap1 | Larp1 | Pdgfra | Junb |
|  | Relb | Tap2 | Lars2 | Pdpn | Kansl1 |
|  | Reps1 | Tax1bp3 | Leng8 | Pdrg1 | Kbtbd2 |
|  | Rfwd2 | Tbc1d20 | Lgals3 | Pex26 | Kcnd3 |
|  | Rgs10 | Tcf25 | Lhfpl3 | Pfdn5 | Kcnip1 |
|  | Rnf10 | Ten1 | Lilrb4a | Pfn1 | Kctd5 |
|  | Rnf111 | Tgtp2 | Lin54 | Pgm2 | Kdelr1 |
|  | Rnf126 | Thap3 | Lin7a | Pgpep1 | Kdelr2 |
|  | Rnf19a | Thbs3 | Lmnb1 | Phax | Kdm1a |
|  | Rpgrip1 | Thoc7 | Lmtk2 | Phb | Kdm2a |
|  | Rpl10 | Timm22 | Lrp11 | Pigk | Kdm2b |
|  | Rpl10a | Timp1 | Lrp3 | Pik3ip1 | Kdm3b |
|  | Rpl11 | Timp3 | Lrrc4b | Pkm | Kdm6b |
|  | Rpl18 | Tk1 | Lrrc58 | Pla2g4a | Khdrbs1 |
|  | Rpl19 | Tlr3 | Lrrfip1 | Plin3 | Khdrbs3 |
|  | Rpl21 | Tm2d2 | Lrwd1 | Plp1 | Khsrp |
|  | Rpl27a | Tmc6 | Lsm14a | Pmel | Kif14 |
|  | Rpl28 | Tmem100 | Lsm14b | Pmvk | Kif18b |
|  | Rpl3 | Tmem234 | Lsm7 | Pnkd | Klf13 |
|  | Rpl36a-ps1 | Tmem29 | Ltbp4 | Polr2b | Klhdc2 |
|  | Rpl6 | Tmem50a | Luc7l2 | Polr3h | Klhl15 |
|  | Rpl7a | Tnfaip2 | Magi2 | Polr3k | Klhl7 |
|  | Rplp0 | Tnni1 | Map1lc3a | Pon2 | Klhl9 |
|  | Rps12 | Tpd52l1 | Map3k10 | Ppp1r14c | Kmt2b |
|  | Rps13 | Tpd52l2 | Map3k12 | Prdx1 | Kmt2c |
|  | Rps16 | Trf | Map4k5 | Prmt2 | Kmt2d |
|  | Rps17 | Trim2 | Map6 | Prss35 | Kmt5a |
|  | Rps23 | Trio | Map7d1 | Psap | Kpna2 |
|  | Rps27 | Tsn | Mapk12 | Psma2 | Kpna4 |
|  | Rps27rt | Tspan17 | Mapk6 | Psma4 | Kpnb1 |
|  | Rpsa | Tspo | Mark1 | Psma7 | Kras |
|  | Rrm1 | Ttyh1 | Mark2 | Psmb1 | Krit1 |
|  | Ruvbl2 | Tubb2a | Mark4 | Psmb10 | Larp1 |
|  | Sag | Twf1 | Mast2 | Psmb2 | Larp4 |
|  | Samd1 | Txn1 | Matn4 | Psmb3 | Lars2 |
|  | Sart3 | Txndc15 | Maz | Psmb6 | Leng8 |
|  | Satb1 | Txndc17 | Mbd6 | Psmb8 | Lgals3 |
|  | Sbf2 | Uqcc2 | Mbtd1 | Psmb9 | Lhfpl3 |
|  | Sbk1 | Uqcrb | Mcm7 | Psmc3 | Lilrb4a |
|  | Scaf1 | Usp18 | Med25 | Psmd10 | Lin54 |
|  | Sec24b | Usp46 | Mettl9 | Psmd12 | Lin7a |
|  | Serp2 | Vim | Mex3b | Psmd14 | Lmnb1 |
|  | Serpine1 | Vit | Mgat4b | Psmd4 | Lmtk2 |
|  | Setd1a | Wbp4 | Mib1 | Psmd7 | Lrp11 |
|  | Setd1b | Wdfy1 | Midn | Psme1 | Lrp3 |
|  | Sf1 | Wls | Mki67 | Ptn | Lrrc4b |
|  | Sfpq | Xaf1 | Mlip | Pttg1ip | Lrrc58 |
|  | Sh2b3 | Yae1d1 | Mmp17 | Rasa2 | Lrrfip1 |
|  | Sh3gl3 | Zcchc17 | Mms19 | Rassf4 | Lrwd1 |
|  | Shisa7 | Zcrb1 | Mn1 | Rbm28 | Lsm14a |
|  | Simc1 | Zfhx4 | Morc4 | Rbm39 | Lsm14b |
|  | Sipa1l1 | Zfp36l1 | Mphosph9 | Rfc1 | Lsm7 |
|  | Sipa1l3 | Zfp87 | Mras | Rhoc | Ltbp4 |
|  | Sirt1 | Zfyve21 | Mrpl23 | Rnf31 | Luc7l2 |
|  | Slc12a6 |  | Mrpl33 | Rpa2 | Maged1 |
|  | Slc16a1 |  | Msi1 | Rpl22l1 | Magi2 |
|  | Slc1a1 |  | Msn | Rpl26 | Map1lc3a |
|  | Slc20a1 |  | Mta1 | Rpl32 | Map3k10 |
|  | Slc30a4 |  | Mthfd2l | Rpl36a | Map3k12 |
|  | Slc35e3 |  | Mtss1l | Rpl37 | Map4k5 |
|  | Sorcs2 |  | Myc | Rps19bp1 | Map6 |
|  | Sox8 |  | Myef2 | Rps24 | Map7d1 |
|  | Sp3 |  | Myt1 | Rps4x | Mapk12 |
|  | Sppl3 |  | Nap1l4 | Rrbp1 | Mapk6 |
|  | Spred3 |  | Nav2 | Rrp1 | Mark1 |
|  | Srgn |  | Nav3 | Rsrc1 | Mark2 |
|  | Srsf1 |  | Nckap1 | Rsrp1 | Mark3 |
|  | Srsf9 |  | Ncoa6 | Rsu1 | Mark4 |
|  | Srxn1 |  | Ncor2 | Rtp4 | Mast2 |
|  | Stab1 |  | Negr1 | Runx3 | Matn4 |
|  | Stk11 |  | Nelfa | S100a11 | Maz |
|  | Stmn4 |  | Nell2 | S100a6 | Mbd6 |
|  | Strn4 |  | Neto2 | S100b | Mbtd1 |
|  | Suds3 |  | Nfatc2ip | Samd9l | Mcm7 |
|  | Sun1 |  | Nme2 | Saraf | Med25 |
|  | Sympk |  | Nomo1 | Sat1 | Mettl9 |
|  | Syn1 |  | Nova1 | Scamp2 | Mex3b |
|  | Syt6 |  | Nt5dc2 | Sccpdh | Mgat4b |
|  | Tada2b |  | Ntn1 | Scg5 | Mib1 |
|  | Taf10 |  | Ntrk3 | Scp2 | Midn |
|  | Taf4 |  | Nudt3 | Scrg1 | Mier1 |
|  | Tarsl2 |  | Numbl | Scrn1 | Mif |
|  | Tbc1d1 |  | Nup153 | Sdc4 | Mki67 |
|  | Tbkbp1 |  | Nup98 | Sdf4 | Mlip |
|  | Tceal5 |  | Olfm2 | Sdhc | Mmp17 |
|  | Tceal6 |  | Olig1 | Sec11c | Mms19 |
|  | Tgfa |  | Olig2 | Sec13 | Mn1 |
|  | Tjp1 |  | Oraov1 | Sec61b | Morc4 |
|  | Tlcd1 |  | Orc5 | Sec62 | Mphosph9 |
|  | Tmcc1 |  | Osbpl6 | Sepp1 | Mras |
|  | Tmed2 |  | Otud4 | Serbp1 | Mrpl23 |
|  | Tmem131 |  | Otud6b | Serf1 | Mrpl33 |
|  | Tmem144 |  | Oxr1 | Serhl | Msi1 |
|  | Tmem243 |  | P2rx7 | Serpine2 | Msl2 |
|  | Tmem245 |  | Paf1 | Sf3a3 | Msn |
|  | Tmem248 |  | Paip1 | Sh3bgr | Mt1 |
|  | Tmem80 |  | Pak1 | Sh3bgrl3 | Mt2 |
|  | Tmod3 |  | Pam16 | Shc4 | Mta1 |
|  | Tmpo |  | Pan3 | Shisa5 | Mthfd2l |
|  | Tnks2 |  | Pank3 | Sin3b | Mtss1l |
|  | Tnr |  | Patz1 | Slc25a4 | Myc |
|  | Tnrc18 |  | Paxbp1 | Slc25a5 | Myef2 |
|  | Tollip |  | Paxip1 | Slc26a2 | Myt1 |
|  | Tomm40 |  | Pcf11 | Slc9a3r1 | Nap1l4 |
|  | Topbp1 |  | Pcgf3 | Sltm | Nav2 |
|  | Tpm3-rs7 |  | Pclo | Slu7 | Nav3 |
|  | Tpst1 |  | Pde3b | Snrnp27 | Nckap1 |
|  | Traf4 |  | Pde8a | Snx1 | Ncoa6 |
|  | Trim28 |  | Pdgfa | Socs3 | Ncor2 |
|  | Tsc22d2 |  | Peak1 | Sorl1 | Negr1 |
|  | Ttyh3 |  | Peg10 | Sparc | Nelfa |
|  | Tuba1c |  | Phactr3 | Sparcl1 | Nell2 |
|  | Tuba4a |  | Phf2 | Spats2l | Neto2 |
|  | Tyrobp |  | Phldb1 | Spin1 | Nfatc2ip |
|  | U2af2 |  | Phrf1 | Srek1ip1 | Nme2 |
|  | Ubc |  | Phyhipl | Srp9 | Nomo1 |
|  | Ube2m |  | Phykpl | Srpr | Nova1 |
|  | Ube2s |  | Pianp | Ssb | Nt5dc2 |
|  | Ube3c |  | Picalm | Ssr2 | Ntn1 |
|  | Ubfd1 |  | Pigm | Ssrp1 | Ntrk3 |
|  | Ubqln4 |  | Pigt | St13 | Nudt3 |
|  | Ugcg |  | Pih1d1 | Stat1 | Numbl |
|  | Uhmk1 |  | Pithd1 | Stk25 | Nup153 |
|  | Uhrf2 |  | Pld3 | Sub1 | Nup98 |
|  | Ulk4 |  | Plekha1 | Svbp | Olfm1 |
|  | Upf1 |  | Plekhm2 | Synpr | Olfm2 |
|  | Usf2 |  | Plk2 | Tagln2 | Olig1 |
|  | Usp31 |  | Plxna1 | Tagln3 | Olig2 |
|  | Usp42 |  | Pnmal1 | Tap1 | Oraov1 |
|  | Usp50 |  | Podxl2 | Tap2 | Orc5 |
|  | Uspl1 |  | Pogz | Tapbp | Osbpl6 |
|  | Ust |  | Polg | Tax1bp3 | Otud4 |
|  | Utp20 |  | Ppard | Tbc1d20 | Otud6b |
|  | Vegfb |  | Ppfia1 | Tbca | Oxr1 |
|  | Vps37b |  | Ppm1a | Tcf25 | P2rx7 |
|  | Wdfy3 |  | Ppm1d | Tcf4 | Paf1 |
|  | Wdr38 |  | Ppm1f | Ten1 | Paip1 |
|  | Wdr89 |  | Ppm1g | Tfrc | Pak1 |
|  | Wipi2 |  | Ppp1cb | Tgtp2 | Pam16 |
|  | Wiz |  | Ppp1cc | Thap3 | Pan3 |
|  | Wsb2 |  | Ppp1r10 | Thbs3 | Pank3 |
|  | Wtap |  | Ppp1r15a | Thoc7 | Patz1 |
|  | Xpo6 |  | Ppp1r9a | Timm10 | Paxbp1 |
|  | Xylt1 |  | Ppp2r2c | Timm22 | Paxip1 |
|  | Ywhah |  | Ppp2r2d | Timp1 | Pcf11 |
|  | Zbed5 |  | Ppp4r2 | Timp3 | Pcgf3 |
|  | Zbtb10 |  | Ppp6r1 | Tk1 | Pclo |
|  | Zbtb18 |  | Prex2 | Tlr3 | Pde3b |
|  | Zc3h11a |  | Prkar2b | Tm2d2 | Pde8a |
|  | Zc3h4 |  | Prkcb | Tmc6 | Pdgfa |
|  | Zcchc8 |  | Prkg2 | Tmed1 | Peak1 |
|  | Zfand5 |  | Prkrir | Tmem100 | Peg10 |
|  | Zfp236 |  | Prmt1 | Tmem234 | Phactr3 |
|  | Zfp384 |  | Prmt3 | Tmem256 | Phf2 |
|  | Zfp664 |  | Prpf39 | Tmem29 | Phldb1 |
|  | Zfp787 |  | Prpf8 | Tmem50a | Phrf1 |
|  | Zfp821 |  | Prr7 | Tnfaip2 | Phyhipl |
|  | Zmiz2 |  | Prrc2a | Tnni1 | Phykpl |
|  | Zranb1 |  | Psenen | Tpd52l1 | Pianp |
|  | Zswim8 |  | Psmc4 | Tpd52l2 | Picalm |
|  | 1700001O22Rik |  | Psmd13 | Tpi1 | Pigm |
|  | 1810026J23Rik |  | Psmd8 | Trappc4 | Pigt |
|  | 2410131K14Rik |  | Ptbp1 | Trf | Pih1d1 |
|  |  |  | Ptbp2 | Trim2 | Pithd1 |
|  |  |  | Pten | Trio | Pld3 |
|  |  |  | Ptms | Tsn | Plekha1 |
|  |  |  | Ptov1 | Tspan17 | Plekhm2 |
|  |  |  | Ptp4a1 | Tspan6 | Plk2 |
|  |  |  | Ptpn12 | Tspo | Plxna1 |
|  |  |  | Pum2 | Ttyh1 | Pnmal1 |
|  |  |  | Qrich1 | Tuba1a | Podxl2 |
|  |  |  | R3hdm2 | Tubb2a | Pogz |
|  |  |  | Rab35 | Twf1 | Polg |
|  |  |  | Rab3b | Txn1 | Ppard |
|  |  |  | Rab6a | Txn2 | Ppfia1 |
|  |  |  | Rac1 | Txndc15 | Ppia |
|  |  |  | Rad23b | Txndc17 | Ppm1a |
|  |  |  | Ralgapb | Ugp2 | Ppm1d |
|  |  |  | Raly | Uqcc2 | Ppm1f |
|  |  |  | Ran | Uqcr11 | Ppm1g |
|  |  |  | Ranbp6 | Uqcrb | Ppp1cb |
|  |  |  | Ranbp9 | Uqcrq | Ppp1cc |
|  |  |  | Rasgrf1 | Usp18 | Ppp1r10 |
|  |  |  | Rasl2-9 | Usp46 | Ppp1r15a |
|  |  |  | Rbbp6 | Vim | Ppp1r9a |
|  |  |  | Rbm12b2 | Vit | Ppp2r2c |
|  |  |  | Rbm15b | Vmp1 | Ppp2r2d |
|  |  |  | Rbm33 | Vwa5a | Ppp4r2 |
|  |  |  | Rbm45 | Wbp4 | Ppp6r1 |
|  |  |  | Rbm8a2 | Wdfy1 | Prex2 |
|  |  |  | Rcc1 | Wls | Prkar2b |
|  |  |  | Rcn2 | Xaf1 | Prkcb |
|  |  |  | Rdh13 | Yae1d1 | Prkg2 |
|  |  |  | Relb | Ywhab | Prkrir |
|  |  |  | Reps1 | Zc3h15 | Prmt1 |
|  |  |  | Rfwd2 | Zcchc17 | Prmt3 |
|  |  |  | Rgs10 | Zcrb1 | Prpf39 |
|  |  |  | Rhob | Zfhx4 | Prpf8 |
|  |  |  | Rlim | Zfp251 | Prr7 |
|  |  |  | Rnf10 | Zfp36l1 | Prrc2a |
|  |  |  | Rnf111 | Zfp87 | Psenen |
|  |  |  | Rnf126 | Zfyve21 | Psmc4 |
|  |  |  | Rnf19a |  | Psmd13 |
|  |  |  | Rpgrip1 |  | Psmd8 |
|  |  |  | Rpl10 |  | Ptbp1 |
|  |  |  | Rpl10a |  | Ptbp2 |
|  |  |  | Rpl11 |  | Pten |
|  |  |  | Rpl12 |  | Ptms |
|  |  |  | Rpl13a |  | Ptov1 |
|  |  |  | Rpl15 |  | Ptp4a1 |
|  |  |  | Rpl18 |  | Ptpn12 |
|  |  |  | Rpl18a |  | Pum2 |
|  |  |  | Rpl19 |  | Qrich1 |
|  |  |  | Rpl21 |  | R3hdm2 |
|  |  |  | Rpl24 |  | Rab35 |
|  |  |  | Rpl27 |  | Rab3b |
|  |  |  | Rpl27a |  | Rab6a |
|  |  |  | Rpl28 |  | Rac1 |
|  |  |  | Rpl29 |  | Rack1 |
|  |  |  | Rpl3 |  | Rad23b |
|  |  |  | Rpl31 |  | Ralgapb |
|  |  |  | Rpl35 |  | Raly |
|  |  |  | Rpl36 |  | Ran |
|  |  |  | Rpl36a-ps1 |  | Ranbp6 |
|  |  |  | Rpl5 |  | Ranbp9 |
|  |  |  | Rpl6 |  | Rasgrf1 |
|  |  |  | Rpl7a |  | Rasl2-9 |
|  |  |  | Rplp0 |  | Rbbp6 |
|  |  |  | Rplp2 |  | Rbm12b2 |
|  |  |  | Rps11 |  | Rbm15b |
|  |  |  | Rps12 |  | Rbm33 |
|  |  |  | Rps13 |  | Rbm45 |
|  |  |  | Rps15a |  | Rbm8a2 |
|  |  |  | Rps16 |  | Rcc1 |
|  |  |  | Rps17 |  | Rcn2 |
|  |  |  | Rps19 |  | Rdh13 |
|  |  |  | Rps23 |  | Relb |
|  |  |  | Rps25 |  | Reps1 |
|  |  |  | Rps27 |  | Rfwd2 |
|  |  |  | Rps27a |  | Rgs10 |
|  |  |  | Rps27rt |  | Rhob |
|  |  |  | Rps6 |  | Rlim |
|  |  |  | Rps8 |  | Rnf10 |
|  |  |  | Rpsa |  | Rnf111 |
|  |  |  | Rrm1 |  | Rnf126 |
|  |  |  | Rsl24d1 |  | Rnf19a |
|  |  |  | Ruvbl2 |  | Rpgrip1 |
|  |  |  | Sae1 |  | Rpl10 |
|  |  |  | Sag |  | Rpl10a |
|  |  |  | Samd1 |  | Rpl11 |
|  |  |  | Sart3 |  | Rpl12 |
|  |  |  | Satb1 |  | Rpl13a |
|  |  |  | Sbf2 |  | Rpl15 |
|  |  |  | Sbk1 |  | Rpl18 |
|  |  |  | Scaf1 |  | Rpl18a |
|  |  |  | Sec24b |  | Rpl19 |
|  |  |  | Sec31a |  | Rpl21 |
|  |  |  | Sema3d |  | Rpl24 |
|  |  |  | Serp2 |  | Rpl27 |
|  |  |  | Serpine1 |  | Rpl27a |
|  |  |  | Set |  | Rpl28 |
|  |  |  | Setd1a |  | Rpl29 |
|  |  |  | Setd1b |  | Rpl3 |
|  |  |  | Setd2 |  | Rpl31 |
|  |  |  | Sf1 |  | Rpl35 |
|  |  |  | Sf3a2 |  | Rpl36 |
|  |  |  | Sfpq |  | Rpl36a-ps1 |
|  |  |  | Sh2b3 |  | Rpl36al |
|  |  |  | Sh3gl3 |  | Rpl5 |
|  |  |  | Shisa7 |  | Rpl6 |
|  |  |  | Sik3 |  | Rpl7a |
|  |  |  | Simc1 |  | Rplp0 |
|  |  |  | Sipa1l1 |  | Rplp2 |
|  |  |  | Sipa1l3 |  | Rps11 |
|  |  |  | Sirt1 |  | Rps12 |
|  |  |  | Skil |  | Rps13 |
|  |  |  | Slc12a6 |  | Rps15a |
|  |  |  | Slc16a1 |  | Rps16 |
|  |  |  | Slc1a1 |  | Rps17 |
|  |  |  | Slc20a1 |  | Rps18 |
|  |  |  | Slc24a5 |  | Rps19 |
|  |  |  | Slc30a4 |  | Rps23 |
|  |  |  | Slc35e2 |  | Rps25 |
|  |  |  | Slc35e3 |  | Rps27 |
|  |  |  | Slc39a1 |  | Rps27a |
|  |  |  | Slc7a11 |  | Rps27rt |
|  |  |  | Smarcad1 |  | Rps28 |
|  |  |  | Smarce1 |  | Rps29 |
|  |  |  | Smek1 |  | Rps5 |
|  |  |  | Smg7 |  | Rps6 |
|  |  |  | Snrnp70 |  | Rps8 |
|  |  |  | Snx17 |  | Rpsa |
|  |  |  | Snx5 |  | Rrm1 |
|  |  |  | Sorcs2 |  | Rsl24d1 |
|  |  |  | Sox6 |  | Ruvbl2 |
|  |  |  | Sox8 |  | Sae1 |
|  |  |  | Sp3 |  | Sag |
|  |  |  | Spon1 |  | Samd1 |
|  |  |  | Sppl3 |  | Sart3 |
|  |  |  | Spred3 |  | Satb1 |
|  |  |  | Srgn |  | Sbf2 |
|  |  |  | Srsf1 |  | Sbk1 |
|  |  |  | Srsf2 |  | Scaf1 |
|  |  |  | Srsf3 |  | Sdc3 |
|  |  |  | Srsf4 |  | Sec23ip |
|  |  |  | Srsf5 |  | Sec24b |
|  |  |  | Srsf6 |  | Sec31a |
|  |  |  | Srsf9 |  | Sema3d |
|  |  |  | Srxn1 |  | Serp2 |
|  |  |  | Ssh2 |  | Serpine1 |
|  |  |  | Stab1 |  | Set |
|  |  |  | Stk11 |  | Setd1a |
|  |  |  | Stmn4 |  | Setd1b |
|  |  |  | Strn4 |  | Setd2 |
|  |  |  | Stx16 |  | Sf1 |
|  |  |  | Suds3 |  | Sf3a2 |
|  |  |  | Sun1 |  | Sfpq |
|  |  |  | Sympk |  | Sfswap |
|  |  |  | Syn1 |  | Sh2b3 |
|  |  |  | Syne2 |  | Sh3gl3 |
|  |  |  | Syt6 |  | Shisa7 |
|  |  |  | Tada2b |  | Sik3 |
|  |  |  | Taf10 |  | Simc1 |
|  |  |  | Taf15 |  | Sipa1l1 |
|  |  |  | Taf4 |  | Sipa1l3 |
|  |  |  | Tarsl2 |  | Sirt1 |
|  |  |  | Tbc1d1 |  | Skil |
|  |  |  | Tbkbp1 |  | Slc12a6 |
|  |  |  | Tbrg4 |  | Slc16a1 |
|  |  |  | Tceal3 |  | Slc1a1 |
|  |  |  | Tceal5 |  | Slc20a1 |
|  |  |  | Tceal6 |  | Slc24a5 |
|  |  |  | Tdg |  | Slc25a36 |
|  |  |  | Tet2 |  | Slc30a4 |
|  |  |  | Tfdp1 |  | Slc35e2 |
|  |  |  | Tgfa |  | Slc35e3 |
|  |  |  | Tial1 |  | Slc39a1 |
|  |  |  | Timm50 |  | Slc7a11 |
|  |  |  | Tjp1 |  | Smarcad1 |
|  |  |  | Tlcd1 |  | Smarce1 |
|  |  |  | Tmcc1 |  | Smek1 |
|  |  |  | Tmed2 |  | Smg7 |
|  |  |  | Tmed7 |  | Snrnp70 |
|  |  |  | Tmem131 |  | Snx17 |
|  |  |  | Tmem144 |  | Snx5 |
|  |  |  | Tmem243 |  | Sorcs2 |
|  |  |  | Tmem245 |  | Sox2 |
|  |  |  | Tmem248 |  | Sox6 |
|  |  |  | Tmem41b |  | Sox8 |
|  |  |  | Tmem80 |  | Sp3 |
|  |  |  | Tmod3 |  | Spon1 |
|  |  |  | Tmpo |  | Spopl |
|  |  |  | Tnks2 |  | Sppl3 |
|  |  |  | Tnr |  | Spred3 |
|  |  |  | Tnrc18 |  | Srgn |
|  |  |  | Tollip |  | Srsf1 |
|  |  |  | Tomm40 |  | Srsf2 |
|  |  |  | Tomm70a |  | Srsf3 |
|  |  |  | Top2b |  | Srsf4 |
|  |  |  | Topbp1 |  | Srsf5 |
|  |  |  | Tpm3-rs7 |  | Srsf6 |
|  |  |  | Tpst1 |  | Srsf9 |
|  |  |  | Traf4 |  | Srxn1 |
|  |  |  | Trim28 |  | Ssh2 |
|  |  |  | Trim33 |  | Stab1 |
|  |  |  | Trim44 |  | Stk11 |
|  |  |  | Trim59 |  | Stmn4 |
|  |  |  | Tsc22d2 |  | Strn4 |
|  |  |  | Ttyh3 |  | Stx16 |
|  |  |  | Tub |  | Suds3 |
|  |  |  | Tuba1c |  | Sun1 |
|  |  |  | Tuba4a |  | Suz12 |
|  |  |  | Tubb4a |  | Sympk |
|  |  |  | Txnrd1 |  | Syn1 |
|  |  |  | Tyrobp |  | Syne2 |
|  |  |  | U2af2 |  | Syt6 |
|  |  |  | Uba1 |  | Tada2b |
|  |  |  | Uba2 |  | Taf10 |
|  |  |  | Ubc |  | Taf15 |
|  |  |  | Ube2m |  | Taf4 |
|  |  |  | Ube2s |  | Tarsl2 |
|  |  |  | Ube3c |  | Tbc1d1 |
|  |  |  | Ube4b |  | Tbc1d10b |
|  |  |  | Ubfd1 |  | Tbkbp1 |
|  |  |  | Ubl3 |  | Tbrg4 |
|  |  |  | Ubqln1 |  | Tceal3 |
|  |  |  | Ubqln2 |  | Tceal5 |
|  |  |  | Ubqln4 |  | Tceal6 |
|  |  |  | Ubr3 |  | Tdg |
|  |  |  | Ugcg |  | Tet2 |
|  |  |  | Uhmk1 |  | Tfdp1 |
|  |  |  | Uhrf2 |  | Tgfa |
|  |  |  | Ulk4 |  | Tial1 |
|  |  |  | Upf1 |  | Timm50 |
|  |  |  | Usf2 |  | Tjp1 |
|  |  |  | Uso1 |  | Tlcd1 |
|  |  |  | Usp24 |  | Tmcc1 |
|  |  |  | Usp31 |  | Tmed2 |
|  |  |  | Usp42 |  | Tmed7 |
|  |  |  | Usp50 |  | Tmem131 |
|  |  |  | Usp9x |  | Tmem144 |
|  |  |  | Uspl1 |  | Tmem243 |
|  |  |  | Ust |  | Tmem245 |
|  |  |  | Utp18 |  | Tmem248 |
|  |  |  | Utp20 |  | Tmem41b |
|  |  |  | Vapb |  | Tmem80 |
|  |  |  | Vars |  | Tmod3 |
|  |  |  | Vegfb |  | Tmpo |
|  |  |  | Vkorc1l1 |  | Tnks2 |
|  |  |  | Vps37b |  | Tnpo1 |
|  |  |  | Vrk1 |  | Tnr |
|  |  |  | Wdfy3 |  | Tnrc18 |
|  |  |  | Wdr1 |  | Tollip |
|  |  |  | Wdr38 |  | Tomm40 |
|  |  |  | Wdr82 |  | Tomm70a |
|  |  |  | Wdr89 |  | Top2b |
|  |  |  | Whsc1 |  | Topbp1 |
|  |  |  | Wipi2 |  | Tpm3-rs7 |
|  |  |  | Wiz |  | Tpst1 |
|  |  |  | Wrnip1 |  | Tpt1 |
|  |  |  | Wsb2 |  | Traf4 |
|  |  |  | Wtap |  | Trim28 |
|  |  |  | Xpo6 |  | Trim33 |
|  |  |  | Xylt1 |  | Trim44 |
|  |  |  | Ywhag |  | Trim59 |
|  |  |  | Ywhah |  | Tsc22d2 |
|  |  |  | Zbed5 |  | Ttyh3 |
|  |  |  | Zbtb10 |  | Tub |
|  |  |  | Zbtb18 |  | Tuba1c |
|  |  |  | Zc3h11a |  | Tuba4a |
|  |  |  | Zc3h14 |  | Tubb4a |
|  |  |  | Zc3h4 |  | Tubb5 |
|  |  |  | Zcchc8 |  | Txnrd1 |
|  |  |  | Zfand5 |  | Tyrobp |
|  |  |  | Zfp236 |  | U2af2 |
|  |  |  | Zfp384 |  | Uba1 |
|  |  |  | Zfp664 |  | Uba2 |
|  |  |  | Zfp68 |  | Ubc |
|  |  |  | Zfp787 |  | Ube2c |
|  |  |  | Zfp821 |  | Ube2m |
|  |  |  | Zmiz2 |  | Ube2s |
|  |  |  | Zranb1 |  | Ube3c |
|  |  |  | Zswim4 |  | Ube4b |
|  |  |  | Zswim8 |  | Ubfd1 |
|  |  |  |  |  | Ubl3 |
|  |  |  |  |  | Ubqln1 |
|  |  |  |  |  | Ubqln2 |
|  |  |  |  |  | Ubqln4 |
|  |  |  |  |  | Ubr3 |
|  |  |  |  |  | Ugcg |
|  |  |  |  |  | Uhmk1 |
|  |  |  |  |  | Uhrf2 |
|  |  |  |  |  | Ulk4 |
|  |  |  |  |  | Upf1 |
|  |  |  |  |  | Usf2 |
|  |  |  |  |  | Uso1 |
|  |  |  |  |  | Usp24 |
|  |  |  |  |  | Usp31 |
|  |  |  |  |  | Usp42 |
|  |  |  |  |  | Usp50 |
|  |  |  |  |  | Usp9x |
|  |  |  |  |  | Uspl1 |
|  |  |  |  |  | Ust |
|  |  |  |  |  | Utp18 |
|  |  |  |  |  | Utp20 |
|  |  |  |  |  | Vapb |
|  |  |  |  |  | Vars |
|  |  |  |  |  | Vcpip1 |
|  |  |  |  |  | Vegfb |
|  |  |  |  |  | Vkorc1l1 |
|  |  |  |  |  | Vps37b |
|  |  |  |  |  | Vrk1 |
|  |  |  |  |  | Vwa9 |
|  |  |  |  |  | Wdfy3 |
|  |  |  |  |  | Wdr1 |
|  |  |  |  |  | Wdr38 |
|  |  |  |  |  | Wdr82 |
|  |  |  |  |  | Wdr89 |
|  |  |  |  |  | Whsc1 |
|  |  |  |  |  | Wipi2 |
|  |  |  |  |  | Wiz |
|  |  |  |  |  | Wrnip1 |
|  |  |  |  |  | Wsb2 |
|  |  |  |  |  | Wtap |
|  |  |  |  |  | Xpo6 |
|  |  |  |  |  | Xylt1 |
|  |  |  |  |  | Ywhag |
|  |  |  |  |  | Ywhah |
|  |  |  |  |  | Zbed5 |
|  |  |  |  |  | Zbtb10 |
|  |  |  |  |  | Zbtb18 |
|  |  |  |  |  | Zbtb37 |
|  |  |  |  |  | Zc3h11a |
|  |  |  |  |  | Zc3h14 |
|  |  |  |  |  | Zc3h4 |
|  |  |  |  |  | Zcchc8 |
|  |  |  |  |  | Zfand5 |
|  |  |  |  |  | Zfp236 |
|  |  |  |  |  | Zfp384 |
|  |  |  |  |  | Zfp532 |
|  |  |  |  |  | Zfp664 |
|  |  |  |  |  | Zfp68 |
|  |  |  |  |  | Zfp787 |
|  |  |  |  |  | Zfp821 |
|  |  |  |  |  | Zmiz2 |
|  |  |  |  |  | Zranb1 |
|  |  |  |  |  | Zswim4 |
|  |  |  |  |  | Zswim8 |
