## Supplementary material for "A cell state specific metabolic vulnerability to GPX4-dependent ferroptosis in glioblastoma": Table S2

| **Persister**  **UP_N1IC**  **(l2fc≥1)** | **Persister UP_N1IC**  **(l2fc≥0.5)** | **Persister UP_N1IC**  **(l2fc>0)** | **Persister**  **DOWN_N1IC**  **(l2fc≤1)** | **Persister**  **DOWN_N1IC**  **(l2fc≤0.5)** | **Persister DOWN_N1IC**  **(l2fc<0)** |
| --- | --- | --- | --- | --- | --- |
| Acss1 | Acss1 | Acss1 | Abhd2 | Abhd2 | Abhd2 |
| Ankrd46 | Ankrd46 | Ankrd46 | Cdk14 | Ago2 | Acaca |
| Arhgef28 | App | App | Cdk17 | Ank2 | Ago2 |
| Cacna1e | Arhgef28 | Arhgef28 | Cited2 | Apba2 | Ank2 |
| Cald1 | Bcan | Bcan | Cyfip1 | Ccdc92 | Apba2 |
| Capg | Bhlhe40 | Bhlhe40 | Dennd2a | Cdk13 | B3gat1 |
| Chrm3 | Cacna1e | Cacna1e | Evl | Cdk14 | Cab39 |
| Csrp1 | Cald1 | Cald1 | Fbrsl1 | Cdk17 | Ccdc92 |
| Ehd3 | Capg | Calm1 | Fry | Cited2 | Cdk13 |
| Fabp7 | Ccdc174 | Capg | Fscn1 | Cnot10 | Cdk14 |
| Fam102a | Cdc5l | Ccdc174 | Fzd1 | Col11a1 | Cdk17 |
| Fam210b | Chrm3 | Ccnd2 | Gab2 | Cyfip1 | Cited2 |
| Fam46a | Csrp1 | Cd47 | Gpc1 | Ddx39b | Cldn11 |
| Has2 | Dct | Cdc5l | Hmox1 | Dennd2a | Cnot10 |
| Hes1 | Ehd3 | Chrm3 | Hnrnpa0 | Dpysl3 | Col11a1 |
| Hes5 | Emc2 | Csrp1 | Jag1 | Evl | Cyfip1 |
| Hey1 | Fabp7 | Ctnnb1 | Kansl1 | Fam76b | Ddx39b |
| Hmcn1 | Fam102a | Dct | Klf13 | Fbrsl1 | Dennd2a |
| Id3 | Fam210b | Ehd3 | Mib1 | Fchsd2 | Dpysl3 |
| Ifit3 | Fam46a | Emc2 | Mn1 | Fry | Epb41l2 |
| Kbtbd11 | Has2 | Fabp7 | Myt1 | Fscn1 | Evl |
| Litaf | Hes1 | Fam102a | Nav3 | Fzd1 | Fam76b |
| Ncald | Hes5 | Fam210b | Ncor2 | Gab2 | Fbrsl1 |
| Pex26 | Hey1 | Fam46a | Nova1 | Gatad1 | Fchsd2 |
| Pla2g4a | Hmcn1 | Frg1 | Ntn1 | Glud1 | Fry |
| Plp1 | Id3 | Gng12 | Pde3b | Gpc1 | Fscn1 |
| Rassf4 | Ifit3 | Has2 | Pdgfa | Hmox1 | Fzd1 |
| Runx3 | Kbtbd11 | Hes1 | Phyhipl | Hnrnpa0 | Gab2 |
| Sat1 | Kcnj10 | Hes5 | Ranbp9 | Jag1 | Gatad1 |
| Serpine2 | Lef1 | Hey1 | Sipa1l1 | Kansl1 | Glud1 |
| Shc4 | Litaf | Hmcn1 | Sipa1l3 | Khdrbs3 | Gpc1 |
| Sorl1 | Mcam | Id3 | Tbkbp1 | Klf13 | Gsap |
| Spats2l | Ncald | Ifit3 | Tnr | Kpnb1 | Hmox1 |
| Tmc6 | Ndufa4 | Igsf3 | Ubfd1 | Magi2 | Hnrnpa0 |
| Trim2 | Nes | Kank1 | Wdr38 | Mbtd1 | Igfbp2 |
| Zfhx4 | Pex26 | Kbtbd11 | Xylt1 | Mib1 | Jag1 |
| Zfp36l1 | Pla2g4a | Kcnj10 | Ywhah | Mn1 | Kansl1 |
|  | Plp1 | Lef1 | Zbtb18 | Mtss1l | Khdrbs3 |
|  | Polr2b | Litaf | Zc3h4 | Myt1 | Klf13 |
|  | Ptn | Lmo4 | Zcchc8 | Nav3 | Kpnb1 |
|  | Rassf4 | Mcam | Zfand5 | Nckap1 | Magi2 |
|  | Rsu1 | Ncald |  | Ncor2 | Mbtd1 |
|  | Runx3 | Ndufa4 |  | Nova1 | Mib1 |
|  | Sat1 | Nes |  | Ntn1 | Mn1 |
|  | Serpine2 | Pdgfra |  | Olig1 | Msl2 |
|  | Shc4 | Pex26 |  | Osbpl6 | Mtss1l |
|  | Sorl1 | Pla2g4a |  | Oxr1 | Myt1 |
|  | Spats2l | Plp1 |  | Pde3b | Nav3 |
|  | Tmc6 | Polr2b |  | Pdgfa | Nckap1 |
|  | Trim2 | Psap |  | Phyhipl | Ncor2 |
|  | Trio | Ptn |  | Polg | Nova1 |
|  | Ttyh1 | Rassf4 |  | Ppm1f | Ntn1 |
|  | Wbp4 | Rsu1 |  | Ptbp2 | Olig1 |
|  | Zfhx4 | Runx3 |  | Pten | Osbpl6 |
|  | Zfp36l1 | Sat1 |  | Ranbp9 | Oxr1 |
|  |  | Serpine2 |  | Rhob | Pde3b |
|  |  | Shc4 |  | Sema3d | Pdgfa |
|  |  | Slu7 |  | Sipa1l1 | Phyhipl |
|  |  | Sorl1 |  | Sipa1l3 | Polg |
|  |  | Spats2l |  | Sox6 | Ppm1f |
|  |  | Tbca |  | Tbkbp1 | Ptbp2 |
|  |  | Tcf4 |  | Tet2 | Pten |
|  |  | Tmc6 |  | Tnr | Ranbp9 |
|  |  | Trim2 |  | Ubfd1 | Rhob |
|  |  | Trio |  | Ubl3 | Sema3d |
|  |  | Ttyh1 |  | Wdr38 | Sipa1l1 |
|  |  | Ugp2 |  | Xylt1 | Sipa1l3 |
|  |  | Vmp1 |  | Ywhah | Sox2 |
|  |  | Wbp4 |  | Zbtb18 | Sox6 |
|  |  | Zc3h15 |  | Zc3h4 | Tbkbp1 |
|  |  | Zfhx4 |  | Zcchc8 | Tet2 |
|  |  | Zfp36l1 |  | Zfand5 | Tnr |
|  |  |  |  |  | Ubfd1 |
|  |  |  |  |  | Ubl3 |
|  |  |  |  |  | Wdr38 |
|  |  |  |  |  | Xylt1 |
|  |  |  |  |  | Ywhah |
|  |  |  |  |  | Zbtb18 |
|  |  |  |  |  | Zc3h4 |
|  |  |  |  |  | Zcchc8 |
|  |  |  |  |  | Zfand5 |
