## Supplementary material for "A cell state specific metabolic vulnerability to GPX4-dependent ferroptosis in glioblastoma": Table S3

| **GENE SET NAME** | **SIZE** | **ES** | **NES** | **NOM**  **p-val** | **FDR q-val** | **FWER p-val** | **RANK AT MAX** | **LEADING EDGE** |
| --- | --- | --- | --- | --- | --- | --- | --- | --- |
| KEGG_OXIDATIVE_PHOSPHORYLATION | 123 | 0.86258525 | 2.4507883 | 0 | 0 | 0 | 1761 | tags=58%, list=7%, signal=61% |
| REACTOME_RESPIRATORY_ELECTRON_TRANSPORT_ATP_SYNTHESIS_BY_CHEMIOSMOTIC_COUPLING_AND_HEAT_PRODUCTION_BY_UNCOUPLING_PROTEINS_ | 119 | 0.8536449 | 2.41815 | 0 | 0 | 0 | 1881 | tags=62%, list=7%, signal=67% |
| GO_INNER_MITOCHONDRIAL_MEMBRANE_PROTEIN_COMPLEX | 120 | 0.8397967 | 2.4117274 | 0 | 0 | 0 | 1761 | tags=58%, list=7%, signal=61% |
| REACTOME_RESPIRATORY_ELECTRON_TRANSPORT | 98 | 0.85887533 | 2.3999891 | 0 | 0 | 0 | 1881 | tags=64%, list=7%, signal=69% |
| KEGG_PARKINSONS_DISEASE | 115 | 0.8504379 | 2.3891587 | 0 | 0 | 0 | 1812 | tags=60%, list=7%, signal=64% |
| GO_RESPIRASOME | 90 | 0.8746036 | 2.3632152 | 0 | 0 | 0 | 1812 | tags=66%, list=7%, signal=70% |
| GO_ATP_SYNTHESIS_COUPLED_ELECTRON_TRANSPORT | 91 | 0.84505075 | 2.3305883 | 0 | 0 | 0 | 1244 | tags=55%, list=5%, signal=57% |
| GO_OXIDATIVE_PHOSPHORYLATION | 134 | 0.82442206 | 2.3295429 | 0 | 0 | 0 | 1812 | tags=53%, list=7%, signal=57% |
| REACTOME_THE_CITRIC_ACID_TCA_CYCLE_AND_RESPIRATORY_ELECTRON_TRANSPORT | 168 | 0.820501 | 2.3185546 | 0 | 0 | 0 | 2254 | tags=55%, list=8%, signal=59% |
| MOOTHA_VOXPHOS | 83 | 0.8420065 | 2.3109727 | 0 | 0 | 0 | 1812 | tags=61%, list=7%, signal=66% |
| GO_NADH_DEHYDROGENASE_ACTIVITY | 43 | 0.9098177 | 2.310499 | 0 | 0 | 0 | 978 | tags=70%, list=4%, signal=72% |
| GO_MITOCHONDRIAL_RESPIRATORY_CHAIN_COMPLEX_ASSEMBLY | 94 | 0.8295528 | 2.300705 | 0 | 0 | 0 | 2095 | tags=56%, list=8%, signal=61% |
| GO_RESPIRATORY_CHAIN_COMPLEX | 76 | 0.8848652 | 2.2874024 | 0 | 0 | 0 | 1088 | tags=62%, list=4%, signal=64% |
| REACTOME_COMPLEX_I_BIOGENESIS | 53 | 0.8790547 | 2.2793396 | 0 | 0 | 0 | 1881 | tags=72%, list=7%, signal=77% |
| WP_MITOCHONDRIAL_COMPLEX_I_ASSEMBLY_MODEL_OXPHOS_SYSTEM | 51 | 0.88031864 | 2.2788043 | 0 | 0 | 0 | 1881 | tags=69%, list=7%, signal=74% |
| WP_ELECTRON_TRANSPORT_CHAIN_OXPHOS_SYSTEM_IN_MITOCHONDRIA | 97 | 0.87567294 | 2.2714279 | 0 | 0 | 0 | 1251 | tags=62%, list=5%, signal=65% |
| GO_RESPIRATORY_ELECTRON_TRANSPORT_CHAIN | 108 | 0.8307688 | 2.2695172 | 0 | 0 | 0 | 1244 | tags=51%, list=5%, signal=53% |
| GO_MITOCHONDRIAL_PROTEIN_COMPLEX | 244 | 0.76442015 | 2.2679794 | 0 | 0 | 0 | 2410 | tags=52%, list=9%, signal=57% |
| GO_ELECTRON_TRANSPORT_CHAIN | 165 | 0.7767949 | 2.253811 | 0 | 0 | 0 | 2442 | tags=49%, list=9%, signal=54% |
| KEGG_HUNTINGTONS_DISEASE | 166 | 0.7541597 | 2.236705 | 0 | 0 | 0 | 2115 | tags=48%, list=8%, signal=51% |
| GO_OXIDOREDUCTASE_ACTIVITY_ACTING_ON_NAD_P_H_QUINONE_OR_SIMILAR_COMPOUND_AS_ACCEPTOR | 53 | 0.8893177 | 2.2213078 | 0 | 0 | 0 | 2259 | tags=74%, list=8%, signal=80% |
| HP_INCREASED_CSF_LACTATE | 82 | 0.81103826 | 2.2148 | 0 | 0 | 0 | 2442 | tags=55%, list=9%, signal=60% |
| GO_OXIDOREDUCTASE_COMPLEX | 99 | 0.81705207 | 2.1941977 | 0 | 0 | 0 | 1130 | tags=47%, list=4%, signal=49% |
| GO_CELLULAR_RESPIRATION | 173 | 0.7696814 | 2.1868882 | 0 | 0 | 0 | 2442 | tags=48%, list=9%, signal=52% |
| GO_MITOCHONDRIAL_ELECTRON_TRANSPORT_NADH_TO_UBIQUINONE | 51 | 0.8917118 | 2.1820028 | 0 | 0 | 0 | 978 | tags=65%, list=4%, signal=67% |
| HP_SCOTOMA | 58 | 0.8339778 | 2.1695547 | 0 | 0 | 0 | 1165 | tags=24%, list=4%, signal=25% |
| GO_NADH_DEHYDROGENASE_COMPLEX | 46 | 0.90862185 | 2.1677215 | 0 | 0 | 0 | 978 | tags=67%, list=4%, signal=70% |
| WP_OXIDATIVE_PHOSPHORYLATION | 56 | 0.88646775 | 2.1650333 | 0 | 0 | 0 | 1761 | tags=70%, list=7%, signal=74% |
| KEGG_ALZHEIMERS_DISEASE | 155 | 0.7444452 | 2.1631806 | 0 | 0 | 0 | 1862 | tags=44%, list=7%, signal=47% |
| HP_RENAL_TUBULAR_DYSFUNCTION | 95 | 0.7542126 | 2.154526 | 0 | 0 | 0 | 2254 | tags=38%, list=8%, signal=41% |
| GO_ORGANELLE_INNER_MEMBRANE | 481 | 0.65782046 | 2.1523092 | 0 | 0 | 0 | 2442 | tags=36%, list=9%, signal=39% |
| GO_ATP_METABOLIC_PROCESS | 282 | 0.6907384 | 2.1411192 | 0 | 0 | 0 | 2444 | tags=40%, list=9%, signal=43% |
| HP_ABNORMAL_ACTIVITY_OF_MITOCHONDRIAL_RESPIRATORY_CHAIN | 67 | 0.80984175 | 2.1409748 | 0 | 0 | 0 | 2400 | tags=57%, list=9%, signal=62% |
| GO_OXIDOREDUCTASE_ACTIVITY_ACTING_ON_NAD_P_H | 94 | 0.82070524 | 2.1288536 | 0 | 0 | 0 | 2259 | tags=48%, list=8%, signal=52% |
| JECHLINGER_EPITHELIAL_TO_MESENCHYMAL_TRANSITION_UP | 67 | 0.76150477 | 2.1144698 | 0 | 0 | 0 | 1920 | tags=36%, list=7%, signal=38% |
| HP_ABNORMAL_CSF_METABOLITE_LEVEL | 99 | 0.7948011 | 2.113559 | 0 | 0 | 0 | 2442 | tags=49%, list=9%, signal=54% |
| WONG_MITOCHONDRIA_GENE_MODULE | 213 | 0.75338227 | 2.1133087 | 0 | 0 | 0 | 2622 | tags=51%, list=10%, signal=56% |
| GO_NADH_DEHYDROGENASE_COMPLEX_ASSEMBLY | 59 | 0.8635111 | 2.1046586 | 0 | 0 | 0 | 1881 | tags=66%, list=7%, signal=71% |
| GO_PROTON_TRANSMEMBRANE_TRANSPORT | 136 | 0.7260804 | 2.0777953 | 0 | 0 | 0 | 1740 | tags=32%, list=6%, signal=34% |
| DER_IFN_ALPHA_RESPONSE_UP | 63 | 0.76286125 | 2.0681589 | 0 | 0 | 0 | 1880 | tags=40%, list=7%, signal=43% |
| HP_ABNORMALITY_OF_THE_MITOCHONDRION | 145 | 0.7134287 | 2.0674198 | 0 | 0 | 0 | 2400 | tags=43%, list=9%, signal=47% |
| YAO_TEMPORAL_RESPONSE_TO_PROGESTERONE_CLUSTER_13 | 164 | 0.70079577 | 2.0615132 | 0 | 0 | 0 | 2646 | tags=56%, list=10%, signal=62% |
| HP_BLURRED_VISION | 27 | 0.889763 | 2.0592983 | 0 | 0 | 0 | 1447 | tags=44%, list=5%, signal=47% |
| GO_THREONINE_TYPE_PEPTIDASE_ACTIVITY | 24 | 0.8594237 | 2.057166 | 0 | 0 | 0 | 1549 | tags=50%, list=6%, signal=53% |
| HP_INCREASED_SERUM_LACTATE | 153 | 0.7190617 | 2.053921 | 0 | 0 | 0 | 2442 | tags=40%, list=9%, signal=44% |
| WP_NONALCOHOLIC_FATTY_LIVER_DISEASE | 146 | 0.70806926 | 2.0537925 | 0 | 0 | 0 | 2132 | tags=43%, list=8%, signal=47% |
| HP_ABNORMAL_BASAL_GANGLIA_MRI_SIGNAL_INTENSITY | 44 | 0.83657324 | 2.0392911 | 0 | 0 | 0 | 2400 | tags=57%, list=9%, signal=62% |
| GO_ELECTRON_TRANSFER_ACTIVITY | 97 | 0.7501156 | 2.0354276 | 0 | 0 | 0 | 2442 | tags=47%, list=9%, signal=52% |
| WUNDER_INFLAMMATORY_RESPONSE_AND_CHOLESTEROL_UP | 53 | 0.8079293 | 2.033379 | 0 | 0 | 0 | 1639 | tags=42%, list=6%, signal=44% |
| HP_MITOCHONDRIAL_MYOPATHY | 49 | 0.8271732 | 2.0301068 | 0 | 0 | 0 | 1795 | tags=53%, list=7%, signal=57% |
| GO_ENERGY_DERIVATION_BY_OXIDATION_OF_ORGANIC_COMPOUNDS | 251 | 0.67121303 | 2.023343 | 0 | 0 | 0 | 2254 | tags=35%, list=8%, signal=38% |
| HP_PIGMENTARY_RETINOPATHY | 146 | 0.7012876 | 2.0212421 | 0 | 0 | 0 | 2970 | tags=38%, list=11%, signal=42% |
| HP_CENTRAL_SCOTOMA | 40 | 0.8745504 | 2.015303 | 0 | 0 | 0 | 854 | tags=30%, list=3%, signal=31% |
| ICHIBA_GRAFT_VERSUS_HOST_DISEASE_D7_UP | 107 | 0.73361063 | 2.0103927 | 0 | 0 | 0 | 2079 | tags=37%, list=8%, signal=40% |
| HP_PAROXYSMAL_INVOLUNTARY_EYE_MOVEMENTS | 35 | 0.8509353 | 2.0085695 | 0 | 4.44E-04 | 0.01 | 2254 | tags=60%, list=8%, signal=65% |
| HP_VOMITING | 192 | 0.6733133 | 2.000445 | 0 | 4.32E-04 | 0.01 | 3274 | tags=42%, list=12%, signal=47% |
| HP_DECREASED_ACTIVITY_OF_THE_PYRUVATE_DEHYDROGENASE_COMPLEX | 32 | 0.8250378 | 1.9859438 | 0 | 4.20E-04 | 0.01 | 2400 | tags=59%, list=9%, signal=65% |
| HP_LETHARGY | 125 | 0.68170756 | 1.9856639 | 0 | 4.10E-04 | 0.01 | 2254 | tags=33%, list=8%, signal=36% |
| GO_PROTON_TRANSPORTING_TWO_SECTOR_ATPASE_COMPLEX | 44 | 0.83339566 | 1.983624 | 0 | 3.99E-04 | 0.01 | 1488 | tags=45%, list=6%, signal=48% |
| BECKER_TAMOXIFEN_RESISTANCE_UP | 45 | 0.8260171 | 1.9779192 | 0 | 0 | 0 | 1874 | tags=49%, list=7%, signal=52% |
| HP_LACTIC_ACIDOSIS | 155 | 0.6839272 | 1.9773839 | 0 | 3.90E-04 | 0.01 | 2442 | tags=40%, list=9%, signal=44% |
| MOOTHA_HUMAN_MITODB_6_2002 | 409 | 0.6057701 | 1.9768381 | 0 | 0 | 0 | 2545 | tags=38%, list=9%, signal=41% |
| MOOTHA_MITOCHONDRIA | 420 | 0.6180763 | 1.9755096 | 0 | 0 | 0 | 3223 | tags=42%, list=12%, signal=47% |
| REACTOME_METABOLISM_OF_POLYAMINES | 56 | 0.75662607 | 1.9752252 | 0 | 0 | 0 | 910 | tags=39%, list=3%, signal=41% |
| GO_PROTON_TRANSMEMBRANE_TRANSPORTER_ACTIVITY | 110 | 0.6899556 | 1.975142 | 0 | 3.80E-04 | 0.01 | 1740 | tags=28%, list=6%, signal=30% |
| HP_PROXIMAL_TUBULOPATHY | 45 | 0.8366669 | 1.9702946 | 0 | 3.71E-04 | 0.01 | 2254 | tags=58%, list=8%, signal=63% |
| HP_HEMIPARESIS | 68 | 0.7373099 | 1.9692844 | 0 | 3.63E-04 | 0.01 | 2158 | tags=26%, list=8%, signal=29% |
| HP_ABNORMAL_MITOCHONDRIA_IN_MUSCLE_TISSUE | 38 | 0.8625573 | 1.9576699 | 0 | 3.55E-04 | 0.01 | 2400 | tags=68%, list=9%, signal=75% |
| GO_MITOCHONDRIAL_ENVELOPE | 683 | 0.6134189 | 1.9510378 | 0 | 6.92E-04 | 0.02 | 3134 | tags=37%, list=12%, signal=41% |
| HP_MUSCLE_ABNORMALITY_RELATED_TO_MITOCHONDRIAL_DYSFUNCTION | 62 | 0.8090341 | 1.9505315 | 0 | 6.77E-04 | 0.02 | 2442 | tags=56%, list=9%, signal=62% |
| HP_ABNORMAL_CELLULAR_PHENOTYPE | 435 | 0.5945327 | 1.9474093 | 0 | 6.63E-04 | 0.02 | 2434 | tags=26%, list=9%, signal=29% |
| HP_ABNORMAL_CNS_MYELINATION | 208 | 0.64892894 | 1.9456086 | 0 | 6.49E-04 | 0.02 | 2554 | tags=34%, list=9%, signal=37% |
| HP_VENTRICULAR_ARRHYTHMIA | 89 | 0.7149517 | 1.9416112 | 0 | 6.36E-04 | 0.02 | 954 | tags=18%, list=4%, signal=19% |
| HP_SLOW_DECREASE_IN_VISUAL_ACUITY | 27 | 0.89494675 | 1.9407775 | 0 | 6.24E-04 | 0.02 | 854 | tags=41%, list=3%, signal=42% |
| CASTELLANO_NRAS_TARGETS_UP | 65 | 0.7602493 | 1.9389981 | 0 | 0.00139104 | 0.02 | 1692 | tags=32%, list=6%, signal=34% |
| HP_LEUKODYSTROPHY | 96 | 0.744763 | 1.9366904 | 0 | 6.12E-04 | 0.02 | 2452 | tags=47%, list=9%, signal=51% |
| BURTON_ADIPOGENESIS_6 | 176 | 0.665905 | 1.932349 | 0 | 0.0013354 | 0.02 | 3660 | tags=49%, list=14%, signal=57% |
| REACTOME_MITOCHONDRIAL_TRANSLATION | 92 | 0.7163991 | 1.9274408 | 0 | 0.00128404 | 0.02 | 2348 | tags=51%, list=9%, signal=56% |
| DOANE_BREAST_CANCER_ESR1_DN | 37 | 0.79050446 | 1.9251989 | 0 | 0.00185517 | 0.03 | 2207 | tags=27%, list=8%, signal=29% |
| GO_METANEPHROS_MORPHOGENESIS | 31 | 0.8016135 | 1.9214334 | 0 | 0.00120888 | 0.04 | 267 | tags=19%, list=1%, signal=20% |
| COLINA_TARGETS_OF_4EBP1_AND_4EBP2 | 335 | 0.6151928 | 1.9185965 | 0 | 0.00178891 | 0.03 | 3081 | tags=38%, list=11%, signal=42% |
| HP_TELANGIECTASIA | 121 | 0.67563 | 1.9185822 | 0 | 0.00148243 | 0.05 | 3484 | tags=36%, list=13%, signal=42% |
| BENNETT_SYSTEMIC_LUPUS_ERYTHEMATOSUS | 21 | 0.87623477 | 1.9171402 | 0 | 0.00228314 | 0.03 | 2141 | tags=57%, list=8%, signal=62% |
| STARK_PREFRONTAL_CORTEX_22Q11_DELETION_DN | 491 | 0.59225094 | 1.9162159 | 0 | 0.00277703 | 0.04 | 3011 | tags=44%, list=11%, signal=48% |
| GO_REGULATION_OF_CELLULAR_AMINO_ACID_METABOLIC_PROCESS | 59 | 0.7337049 | 1.9153321 | 0 | 0.00145547 | 0.05 | 1634 | tags=42%, list=6%, signal=45% |
| HP_HYPERTROPHIC_CARDIOMYOPATHY | 231 | 0.6489546 | 1.9135591 | 0 | 0.00142948 | 0.05 | 3128 | tags=39%, list=12%, signal=44% |
| GO_ENDOPEPTIDASE_REGULATOR_ACTIVITY | 114 | 0.6764677 | 1.9120162 | 0 | 0.00140441 | 0.05 | 1824 | tags=21%, list=7%, signal=22% |
| XU_GH1_AUTOCRINE_TARGETS_UP | 146 | 0.6397851 | 1.9113344 | 0 | 0.00268745 | 0.04 | 3433 | tags=35%, list=13%, signal=40% |
| KARLSSON_TGFB1_TARGETS_DN | 188 | 0.661626 | 1.910716 | 0 | 0.00260347 | 0.04 | 3104 | tags=46%, list=12%, signal=52% |
| HP_ABNORMALITY_OF_ACID_BASE_HOMEOSTASIS | 312 | 0.61495525 | 1.9073695 | 0 | 0.00138019 | 0.05 | 3291 | tags=38%, list=12%, signal=42% |
| WP_TYPE_II_INTERFERON_SIGNALING_IFNG | 29 | 0.80830246 | 1.90425 | 0 | 0.00303377 | 0.05 | 1909 | tags=38%, list=7%, signal=41% |
| HP_TYPE_I_DIABETES_MELLITUS | 34 | 0.81271577 | 1.9041559 | 0 | 0.00162927 | 0.06 | 1197 | tags=29%, list=4%, signal=31% |
| HP_ABNORMAL_CIRCULATING_CARBOHYDRATE_CONCENTRATION | 50 | 0.7723193 | 1.9016536 | 0 | 0.00187 | 0.07 | 2254 | tags=48%, list=8%, signal=52% |
| HP_VASCULAR_TORTUOSITY | 19 | 0.8760545 | 1.9011331 | 0 | 0.00183934 | 0.07 | 1194 | tags=42%, list=4%, signal=44% |
| LIU_SMARCA4_TARGETS | 43 | 0.7766604 | 1.8986105 | 0 | 0.00294454 | 0.05 | 1388 | tags=23%, list=5%, signal=24% |
| GO_MEMBRANE_LIPID_CATABOLIC_PROCESS | 32 | 0.82723945 | 1.8968791 | 0 | 0.00180968 | 0.07 | 3612 | tags=59%, list=13%, signal=68% |
| GO_GENERATION_OF_PRECURSOR_METABOLITES_AND_ENERGY | 473 | 0.61043906 | 1.8963597 | 0 | 0.00178095 | 0.07 | 2444 | tags=32%, list=9%, signal=35% |
| HP_ABNORMAL_MUSCLE_FIBER_MORPHOLOGY | 186 | 0.6354691 | 1.8950937 | 0 | 0.00175313 | 0.07 | 2442 | tags=27%, list=9%, signal=30% |
| HP_DECREASED_ACTIVITY_OF_MITOCHONDRIAL_COMPLEX_I | 37 | 0.826472 | 1.895056 | 0 | 0.00172615 | 0.07 | 2322 | tags=62%, list=9%, signal=68% |
| ZHAN_MULTIPLE_MYELOMA_HP_DN | 43 | 0.7665734 | 1.8913004 | 0 | 0.0033513 | 0.06 | 2852 | tags=44%, list=11%, signal=49% |
| GO_ORGANELLAR_RIBOSOME | 85 | 0.71487045 | 1.8910097 | 0 | 0.0017 | 0.07 | 2348 | tags=49%, list=9%, signal=54% |
| GO_AEROBIC_RESPIRATION | 81 | 0.71473837 | 1.8896949 | 0 | 0.00191781 | 0.08 | 1883 | tags=41%, list=7%, signal=44% |
| HP_EPISODIC_VOMITING | 25 | 0.83282775 | 1.88539 | 0 | 0.00236086 | 0.09 | 1093 | tags=44%, list=4%, signal=46% |
| KIM_GLIS2_TARGETS_UP | 81 | 0.707273 | 1.8831614 | 0 | 0.00372281 | 0.07 | 2340 | tags=37%, list=9%, signal=40% |
| REACTOME_CROSS_PRESENTATION_OF_SOLUBLE_EXOGENOUS_ANTIGENS_ENDOSOMES_ | 46 | 0.7675965 | 1.8823026 | 0 | 0.00454442 | 0.09 | 805 | tags=41%, list=3%, signal=43% |
| GO_PROTEASOMAL_UBIQUITIN_INDEPENDENT_PROTEIN_CATABOLIC_PROCESS | 20 | 0.8577937 | 1.8807608 | 0 | 0.0030214 | 0.12 | 597 | tags=50%, list=2%, signal=51% |
| GO_REGULATION_OF_CELLULAR_AMINE_METABOLIC_PROCESS | 74 | 0.7166766 | 1.8769133 | 0 | 0.00343186 | 0.14 | 1634 | tags=35%, list=6%, signal=37% |
| KLEIN_PRIMARY_EFFUSION_LYMPHOMA_DN | 52 | 0.7435548 | 1.8741969 | 0 | 0.00531735 | 0.11 | 3146 | tags=40%, list=12%, signal=46% |
| HOFFMANN_PRE_BI_TO_LARGE_PRE_BII_LYMPHOCYTE_DN | 71 | 0.73587376 | 1.8713669 | 0 | 0.005181 | 0.11 | 2331 | tags=37%, list=9%, signal=40% |
| DER_IFN_GAMMA_RESPONSE_UP | 62 | 0.7289439 | 1.8704509 | 0 | 0.00547533 | 0.12 | 1880 | tags=40%, list=7%, signal=43% |
| GO_MITOCHONDRIAL_TRANSLATIONAL_TERMINATION | 87 | 0.71714395 | 1.8691226 | 0 | 0.00338352 | 0.14 | 2348 | tags=51%, list=9%, signal=55% |
| GO_MITOCHONDRION | 1427 | 0.5367474 | 1.8642639 | 0 | 0.00378043 | 0.16 | 3944 | tags=38%, list=15%, signal=43% |
| HP_POLYNEUROPATHY | 50 | 0.7597808 | 1.8638539 | 0 | 0.00372864 | 0.16 | 1938 | tags=32%, list=7%, signal=34% |
| HP_APHASIA | 46 | 0.75232124 | 1.8635584 | 0 | 0.00367826 | 0.16 | 1197 | tags=35%, list=4%, signal=36% |
| GO_PROTON_TRANSPORTING_TWO_SECTOR_ATPASE_COMPLEX_PROTON_TRANSPORTING_DOMAIN | 20 | 0.8481496 | 1.8628286 | 0 | 0.00362921 | 0.16 | 822 | tags=35%, list=3%, signal=36% |
| HP_RETINAL_TELANGIECTASIA | 16 | 0.9431773 | 1.860327 | 0 | 0.00400732 | 0.18 | 941 | tags=69%, list=3%, signal=71% |
| GRAESSMANN_RESPONSE_TO_MC_AND_SERUM_DEPRIVATION_UP | 194 | 0.6285914 | 1.8592473 | 0 | 0.00574835 | 0.13 | 2159 | tags=27%, list=8%, signal=29% |
| HP_MITOCHONDRIAL_INHERITANCE | 26 | 0.8689117 | 1.8590456 | 0 | 0.00416314 | 0.19 | 2622 | tags=73%, list=10%, signal=81% |
| KIM_LRRC3B_TARGETS | 24 | 0.8431871 | 1.8577707 | 0 | 0.00561149 | 0.13 | 1315 | tags=58%, list=5%, signal=61% |
| GO_MITOCHONDRIAL_LARGE_RIBOSOMAL_SUBUNIT | 55 | 0.70862514 | 1.857642 | 0 | 0.00431465 | 0.19 | 2252 | tags=53%, list=8%, signal=57% |
| HP_ABNORMAL_BRAINSTEM_MRI_SIGNAL_INTENSITY | 37 | 0.82060397 | 1.8566549 | 0 | 0.00446232 | 0.19 | 2254 | tags=62%, list=8%, signal=68% |
| REN_ALVEOLAR_RHABDOMYOSARCOMA_DN | 399 | 0.5804439 | 1.8565147 | 0 | 0.00587543 | 0.14 | 3236 | tags=37%, list=12%, signal=42% |
| BROWNE_INTERFERON_RESPONSIVE_GENES | 57 | 0.814297 | 1.8533218 | 0 | 0.00611251 | 0.15 | 3668 | tags=54%, list=14%, signal=63% |
| MOOTHA_PGC | 398 | 0.56725365 | 1.8517992 | 0 | 0.00635167 | 0.16 | 3234 | tags=43%, list=12%, signal=48% |
| GO_STRUCTURAL_CONSTITUENT_OF_EYE_LENS | 18 | 0.91425955 | 1.8492757 | 0 | 0.00480717 | 0.2 | 1550 | tags=22%, list=6%, signal=24% |
| MARKEY_RB1_ACUTE_LOF_DN | 220 | 0.6297987 | 1.8474476 | 0 | 0.00621359 | 0.16 | 1919 | tags=25%, list=7%, signal=27% |
| GO_ANTIGEN_PROCESSING_AND_PRESENTATION_OF_PEPTIDE_ANTIGEN_VIA_MHC_CLASS_I | 90 | 0.6939216 | 1.8472853 | 0 | 0.00474782 | 0.2 | 1624 | tags=33%, list=6%, signal=35% |
| HP_OPTIC_NEUROPATHY | 52 | 0.7378155 | 1.8471835 | 0 | 0.00468992 | 0.2 | 1830 | tags=54%, list=7%, signal=58% |
| YAO_TEMPORAL_RESPONSE_TO_PROGESTERONE_CLUSTER_10 | 65 | 0.6822451 | 1.8471053 | 0 | 0.00608139 | 0.16 | 3563 | tags=62%, list=13%, signal=71% |
| GO_MITOCHONDRION_ORGANIZATION | 500 | 0.5569051 | 1.8463552 | 0 | 0.00463342 | 0.2 | 2442 | tags=30%, list=9%, signal=32% |
| HP_REDUCED_CONSCIOUSNESS_CONFUSION | 277 | 0.6051903 | 1.8418531 | 0 | 0.00496037 | 0.22 | 2398 | tags=27%, list=9%, signal=29% |
| BERENJENO_TRANSFORMED_BY_RHOA_DN | 374 | 0.59181255 | 1.8414742 | 0 | 0.00595469 | 0.16 | 2836 | tags=38%, list=11%, signal=42% |
| GO_NEGATIVE_REGULATION_OF_ENDOTHELIAL_CELL_PROLIFERATION | 36 | 0.72300524 | 1.839739 | 0 | 0.00527571 | 0.24 | 2587 | tags=28%, list=10%, signal=31% |
| HP_VENTRICULAR_PREEXCITATION | 21 | 0.9050914 | 1.8390878 | 0 | 0.00521436 | 0.24 | 854 | tags=52%, list=3%, signal=54% |
| HP_SENSORIMOTOR_NEUROPATHY | 64 | 0.66253406 | 1.8353894 | 0 | 0.00588987 | 0.27 | 2329 | tags=28%, list=9%, signal=31% |
| GO_ANTIGEN_PROCESSING_AND_PRESENTATION_OF_EXOGENOUS_PEPTIDE_ANTIGEN_VIA_MHC_CLASS_I | 72 | 0.721722 | 1.834655 | 0 | 0.00618959 | 0.28 | 1624 | tags=38%, list=6%, signal=40% |
| PELLICCIOTTA_HDAC_IN_ANTIGEN_PRESENTATION_DN | 44 | 0.7541354 | 1.8269112 | 0 | 0.00754799 | 0.21 | 1624 | tags=48%, list=6%, signal=51% |
| REACTOME_ROS_AND_RNS_PRODUCTION_IN_PHAGOCYTES | 35 | 0.7688597 | 1.8260347 | 0 | 0.00806261 | 0.23 | 1206 | tags=23%, list=4%, signal=24% |
| HP_PANCREATITIS | 55 | 0.6986353 | 1.825106 | 0 | 0.00756276 | 0.34 | 2025 | tags=24%, list=8%, signal=26% |
| KEGG_PROTEASOME | 41 | 0.7646845 | 1.8244551 | 0 | 0.00790452 | 0.23 | 805 | tags=44%, list=3%, signal=45% |
| GRAHAM_CML_QUIESCENT_VS_NORMAL_QUIESCENT_DN | 39 | 0.7595135 | 1.8227918 | 0 | 0.00775251 | 0.23 | 1903 | tags=33%, list=7%, signal=36% |
| HP_PROGRESSIVE_SENSORINEURAL_HEARING_IMPAIRMENT | 30 | 0.82512313 | 1.8224071 | 0 | 0.00800993 | 0.36 | 1098 | tags=23%, list=4%, signal=24% |
| HP_DROWSINESS | 29 | 0.78294706 | 1.8212318 | 0 | 0.00792191 | 0.36 | 4033 | tags=45%, list=15%, signal=53% |
| TSENG_ADIPOGENIC_POTENTIAL_DN | 43 | 0.73254246 | 1.8204669 | 0 | 0.00791392 | 0.23 | 2881 | tags=33%, list=11%, signal=36% |
| GO_OXIDOREDUCTASE_ACTIVITY | 619 | 0.5814058 | 1.8195544 | 0 | 0.0080129 | 0.36 | 3236 | tags=32%, list=12%, signal=35% |
| LIU_VAV3_PROSTATE_CARCINOGENESIS_UP | 76 | 0.67611676 | 1.8194174 | 0 | 0.00776736 | 0.23 | 1774 | tags=26%, list=7%, signal=28% |
| GO_MUSCLE_FILAMENT_SLIDING | 31 | 0.7694615 | 1.8138844 | 0 | 0.00896116 | 0.39 | 1714 | tags=19%, list=6%, signal=21% |
| HP_INCREASED_SERUM_PYRUVATE | 40 | 0.7939875 | 1.8135761 | 0 | 0.00886583 | 0.39 | 2254 | tags=55%, list=8%, signal=60% |
| KRASNOSELSKAYA_ILF3_TARGETS_UP | 26 | 0.8490036 | 1.8132654 | 0 | 0.00822724 | 0.24 | 1086 | tags=35%, list=4%, signal=36% |
| GO_TRANSLATIONAL_ELONGATION | 128 | 0.61538976 | 1.8131328 | 0 | 0.00894072 | 0.4 | 2386 | tags=43%, list=9%, signal=47% |
| CROMER_TUMORIGENESIS_DN | 33 | 0.76284736 | 1.8122017 | 0 | 0.00808033 | 0.24 | 3318 | tags=33%, list=12%, signal=38% |
| HP_ABNORMAL_CIRCULATING_PYRUVATE_FAMILY_AMINO_ACID_CONCENTRATION | 28 | 0.8081306 | 1.8108538 | 0 | 0.00935144 | 0.43 | 1633 | tags=36%, list=6%, signal=38% |
| HP_PARAPLEGIA | 143 | 0.6067979 | 1.81068 | 0 | 0.00925503 | 0.43 | 2645 | tags=27%, list=10%, signal=30% |
| HP_STROKE_LIKE_EPISODE | 14 | 0.9004089 | 1.8088331 | 0 | 0.01014319 | 0.47 | 854 | tags=43%, list=3%, signal=44% |
| HP_LEFT_VENTRICULAR_DYSFUNCTION | 57 | 0.7190441 | 1.8081195 | 0 | 0.01004073 | 0.47 | 1194 | tags=18%, list=4%, signal=18% |
| HOLLMANN_APOPTOSIS_VIA_CD40_DN | 224 | 0.5957803 | 1.8072731 | 0 | 0.00911567 | 0.28 | 2313 | tags=29%, list=9%, signal=32% |
| HP_ABNORMAL_SPEECH_PROSODY | 19 | 0.85630214 | 1.8069096 | 0 | 0.01009787 | 0.48 | 854 | tags=37%, list=3%, signal=38% |
| HP_ERYTHEMA | 96 | 0.66724855 | 1.8029362 | 0 | 0.01125946 | 0.55 | 3498 | tags=28%, list=13%, signal=32% |
| GO_REGULATION_OF_STEM_CELL_DIFFERENTIATION | 109 | 0.6394793 | 1.8028544 | 0 | 0.01114908 | 0.55 | 1939 | tags=33%, list=7%, signal=35% |
| GO_CYTOCHROME_COMPLEX | 28 | 0.8309715 | 1.8004086 | 0 | 0.01104083 | 0.55 | 1088 | tags=46%, list=4%, signal=48% |
| HP_OPTIC_DISC_PALLOR | 111 | 0.62665707 | 1.799457 | 0 | 0.01093467 | 0.55 | 2341 | tags=28%, list=9%, signal=30% |
| HP_POOR_EYE_CONTACT | 68 | 0.70488846 | 1.7986454 | 0 | 0.01098596 | 0.56 | 2302 | tags=41%, list=9%, signal=45% |
| GO_MEMBRANE_PROTEIN_COMPLEX | 1033 | 0.5427339 | 1.7984732 | 0 | 0.01088231 | 0.56 | 2410 | tags=24%, list=9%, signal=25% |
| SMID_BREAST_CANCER_NORMAL_LIKE_UP | 398 | 0.5608508 | 1.7984385 | 0 | 0.01187323 | 0.35 | 2830 | tags=17%, list=11%, signal=19% |
| HP_PROGRESSIVE_VISUAL_LOSS | 91 | 0.6649727 | 1.7983342 | 0 | 0.01078061 | 0.56 | 2068 | tags=24%, list=8%, signal=26% |
| GO_PROTEASOME_CORE_COMPLEX | 18 | 0.88389915 | 1.7965301 | 0 | 0.01142352 | 0.58 | 597 | tags=56%, list=2%, signal=57% |
| HP_FETAL_DISTRESS | 36 | 0.8069366 | 1.796254 | 0 | 0.01131871 | 0.58 | 2254 | tags=61%, list=8%, signal=67% |
| MCBRYAN_PUBERTAL_BREAST_4_5WK_UP | 236 | 0.5834143 | 1.7956662 | 0 | 0.01254538 | 0.38 | 3364 | tags=30%, list=12%, signal=34% |
| GO_SCF_DEPENDENT_PROTEASOMAL_UBIQUITIN_DEPENDENT_PROTEIN_CATABOLIC_PROCESS | 88 | 0.66487914 | 1.7955775 | 0 | 0.01165296 | 0.59 | 2297 | tags=34%, list=9%, signal=37% |
| GO_PEPTIDASE_REGULATOR_ACTIVITY | 146 | 0.6362607 | 1.7953564 | 0 | 0.01169217 | 0.59 | 2661 | tags=24%, list=10%, signal=26% |
| HP_VITILIGO | 22 | 0.8223333 | 1.7931439 | 0 | 0.01216206 | 0.63 | 1487 | tags=27%, list=6%, signal=29% |
| WP_MITOCHONDRIAL_CIV_ASSEMBLY | 33 | 0.7826648 | 1.7921774 | 0 | 0.01373339 | 0.41 | 2442 | tags=52%, list=9%, signal=57% |
| HP_RAGGED_RED_MUSCLE_FIBERS | 40 | 0.79632366 | 1.7905824 | 0 | 0.01262333 | 0.64 | 1481 | tags=33%, list=6%, signal=34% |
| LAIHO_COLORECTAL_CANCER_SERRATED_UP | 106 | 0.67762196 | 1.7903961 | 0 | 0.01406165 | 0.41 | 2434 | tags=46%, list=9%, signal=51% |
| GO_OXIDATION_REDUCTION_PROCESS | 857 | 0.5438354 | 1.7898407 | 0 | 0.01279414 | 0.64 | 3236 | tags=30%, list=12%, signal=33% |
| BOQUEST_STEM_CELL_UP | 232 | 0.5997011 | 1.7895936 | 0 | 0.01465893 | 0.44 | 2800 | tags=24%, list=10%, signal=27% |
| HP_VISUAL_FIELD_DEFECT | 166 | 0.5800548 | 1.7895305 | 0 | 0.01282149 | 0.64 | 3665 | tags=26%, list=14%, signal=30% |
| LEE_BMP2_TARGETS_UP | 683 | 0.53795755 | 1.7884402 | 0 | 0.0146941 | 0.45 | 3305 | tags=27%, list=12%, signal=30% |
| HP_HEPATOMEGALY | 431 | 0.5590258 | 1.787437 | 0 | 0.01298852 | 0.65 | 3525 | tags=34%, list=13%, signal=38% |
| GO_MYOSIN_COMPLEX | 46 | 0.72048813 | 1.7871655 | 0 | 0.0130149 | 0.65 | 986 | tags=20%, list=4%, signal=20% |
| GO_THREONINE_TYPE_ENDOPEPTIDASE_ACTIVITY | 19 | 0.8769014 | 1.7871524 | 0 | 0.01290461 | 0.65 | 597 | tags=53%, list=2%, signal=54% |
| BAUS_TFF2_TARGETS_UP | 25 | 0.81344163 | 1.7853398 | 0 | 0.01525306 | 0.45 | 1097 | tags=20%, list=4%, signal=21% |
| HP_ABNORMAL_CIRCULATING_CARBOXYLIC_ACID_CONCENTRATION | 117 | 0.65395886 | 1.7852706 | 0 | 0.0134686 | 0.66 | 2493 | tags=28%, list=9%, signal=31% |
| KIM_ALL_DISORDERS_DURATION_CORR_DN | 140 | 0.62725365 | 1.7851816 | 0 | 0.0150184 | 0.45 | 2936 | tags=46%, list=11%, signal=52% |
| HP_BASAL_GANGLIA_CALCIFICATION | 30 | 0.7994997 | 1.7850521 | 0 | 0.01335636 | 0.66 | 981 | tags=30%, list=4%, signal=31% |
| COWLING_MYCN_TARGETS | 37 | 0.73494864 | 1.7850517 | 0 | 0.01479085 | 0.45 | 1550 | tags=35%, list=6%, signal=37% |
| GO_TRANSLATIONAL_TERMINATION | 101 | 0.66737574 | 1.784452 | 0 | 0.01337904 | 0.67 | 2373 | tags=49%, list=9%, signal=53% |
| GO_METANEPHRIC_NEPHRON_MORPHOGENESIS | 24 | 0.8314537 | 1.7833087 | 0 | 0.01353371 | 0.67 | 267 | tags=25%, list=1%, signal=25% |
| GILDEA_METASTASIS | 22 | 0.82636744 | 1.7814333 | 0 | 0.01658826 | 0.48 | 1882 | tags=41%, list=7%, signal=44% |
| MOSERLE_IFNA_RESPONSE | 24 | 0.8499557 | 1.7805896 | 0 | 0.01659269 | 0.48 | 2141 | tags=54%, list=8%, signal=59% |
| HP_ABNORMAL_BRAIN_LACTATE_LEVEL_BY_MRS | 13 | 0.89590555 | 1.7800885 | 0 | 0.01472313 | 0.72 | 854 | tags=54%, list=3%, signal=56% |
| BLANCO_MELO_COVID19_SARS_COV_2_INFECTION_A594_ACE2_EXPRESSING_CELLS_RUXOLITINIB_DN | 62 | 0.6676659 | 1.778822 | 0 | 0.01684073 | 0.48 | 2300 | tags=35%, list=9%, signal=39% |
| GO_ATP_SYNTHESIS_COUPLED_PROTON_TRANSPORT | 21 | 0.83222395 | 1.7787244 | 0 | 0.01486549 | 0.72 | 1663 | tags=62%, list=6%, signal=66% |
| HP_ACTION_TREMOR | 134 | 0.62965536 | 1.7786956 | 0 | 0.01474657 | 0.72 | 1776 | tags=23%, list=7%, signal=25% |
| HAN_JNK_SINGALING_UP | 30 | 0.753706 | 1.7786592 | 0 | 0.01709019 | 0.5 | 959 | tags=30%, list=4%, signal=31% |
| LEE_LIVER_CANCER_MYC_E2F1_UP | 50 | 0.7119173 | 1.7768228 | 0 | 0.01708655 | 0.51 | 3625 | tags=48%, list=13%, signal=55% |
| HP_IMPAIRED_SOCIAL_INTERACTIONS | 95 | 0.660918 | 1.7752922 | 0 | 0.01678613 | 0.75 | 2302 | tags=34%, list=9%, signal=37% |
| DER_IFN_BETA_RESPONSE_UP | 91 | 0.66336054 | 1.7733076 | 0 | 0.01780262 | 0.54 | 2141 | tags=36%, list=8%, signal=39% |
| HP_ABNORMAL_PATTERN_OF_RESPIRATION | 320 | 0.57574224 | 1.772516 | 0 | 0.01766509 | 0.79 | 2935 | tags=26%, list=11%, signal=28% |
| HP_TYPE_1_MUSCLE_FIBER_PREDOMINANCE | 41 | 0.7074437 | 1.7708902 | 0 | 0.01827985 | 0.82 | 4184 | tags=34%, list=16%, signal=40% |
| GO_CELLULAR_KETONE_METABOLIC_PROCESS | 218 | 0.589344 | 1.7699414 | 0 | 0.01900594 | 0.85 | 2476 | tags=27%, list=9%, signal=29% |
| HP_ABNORMAL_CSF_PROTEIN_LEVEL | 37 | 0.7676673 | 1.7695926 | 0 | 0.01935516 | 0.85 | 2158 | tags=38%, list=8%, signal=41% |
| HP_EXTERNAL_OPHTHALMOPLEGIA | 47 | 0.7392131 | 1.7672979 | 0 | 0.02042924 | 0.86 | 1382 | tags=30%, list=5%, signal=31% |
| YAO_TEMPORAL_RESPONSE_TO_PROGESTERONE_CLUSTER_16 | 72 | 0.6974245 | 1.7672466 | 0 | 0.01988685 | 0.57 | 2853 | tags=46%, list=11%, signal=51% |
| GO_CIS_TRANS_ISOMERASE_ACTIVITY | 32 | 0.77824575 | 1.7670164 | 0 | 0.02039535 | 0.86 | 3869 | tags=63%, list=14%, signal=73% |
| HP_ABNORMAL_CARDIAC_VENTRICULAR_FUNCTION | 69 | 0.6793448 | 1.7663114 | 0 | 0.02036216 | 0.86 | 1237 | tags=17%, list=5%, signal=18% |
| HP_ABNORMALITY_OF_PULMONARY_CIRCULATION | 124 | 0.62216973 | 1.7655835 | 0 | 0.02092442 | 0.88 | 2928 | tags=24%, list=11%, signal=27% |
| CHIARADONNA_NEOPLASTIC_TRANSFORMATION_CDC25_DN | 140 | 0.61411846 | 1.7652199 | 0 | 0.02076584 | 0.6 | 2484 | tags=33%, list=9%, signal=36% |
| GO_NEGATIVE_REGULATION_OF_ANIMAL_ORGAN_MORPHOGENESIS | 30 | 0.7902431 | 1.763587 | 0 | 0.02136656 | 0.89 | 1194 | tags=27%, list=4%, signal=28% |
| SANA_RESPONSE_TO_IFNG_UP | 59 | 0.69817066 | 1.7628207 | 0 | 0.0213967 | 0.64 | 2468 | tags=42%, list=9%, signal=47% |
| GO_MESENCHYMAL_TO_EPITHELIAL_TRANSITION | 18 | 0.8914748 | 1.7626015 | 0 | 0.02144384 | 0.89 | 158 | tags=17%, list=1%, signal=17% |
| HAN_JNK_SINGALING_DN | 36 | 0.74466926 | 1.7623574 | 0 | 0.02111516 | 0.64 | 3174 | tags=44%, list=12%, signal=50% |
| GO_PROTON_TRANSPORTING_ATP_SYNTHASE_COMPLEX | 16 | 0.8661537 | 1.7620364 | 0 | 0.02140485 | 0.89 | 1251 | tags=63%, list=5%, signal=66% |
| GO_METANEPHRIC_NEPHRON_DEVELOPMENT | 33 | 0.7296132 | 1.7619495 | 0 | 0.02124974 | 0.89 | 267 | tags=21%, list=1%, signal=21% |
| HP_GLOMERULOSCLEROSIS | 67 | 0.6677658 | 1.7598968 | 0 | 0.02155493 | 0.9 | 1674 | tags=24%, list=6%, signal=25% |
| NAKAJIMA_MAST_CELL | 37 | 0.7342132 | 1.7585418 | 0 | 0.02345839 | 0.65 | 1104 | tags=30%, list=4%, signal=31% |
| HP_DILATED_CARDIOMYOPATHY | 127 | 0.6477031 | 1.7582173 | 0 | 0.02231274 | 0.92 | 2506 | tags=29%, list=9%, signal=32% |
| GO_PROTON_CHANNEL_ACTIVITY | 21 | 0.84781224 | 1.7580931 | 0 | 0.02226775 | 0.92 | 1663 | tags=52%, list=6%, signal=56% |
| BUCKANOVICH_T_LYMPHOCYTE_HOMING_ON_TUMOR_UP | 15 | 0.837538 | 1.7572078 | 0 | 0.02403004 | 0.66 | 2544 | tags=53%, list=9%, signal=59% |
| LI_DCP2_BOUND_MRNA | 85 | 0.64286697 | 1.7570097 | 0 | 0.02394306 | 0.66 | 3020 | tags=53%, list=11%, signal=59% |
| GO_AMINE_METABOLIC_PROCESS | 118 | 0.6670472 | 1.756159 | 0 | 0.02368352 | 0.92 | 912 | tags=22%, list=3%, signal=23% |
| RADAEVA_RESPONSE_TO_IFNA1_UP | 39 | 0.72673976 | 1.7553195 | 0 | 0.02426828 | 0.66 | 1700 | tags=44%, list=6%, signal=46% |
| HP_ABNORMAL_CIRCULATING_AROMATIC_AMINO_ACID_CONCENTRATION | 12 | 0.8725082 | 1.7551755 | 0.02941177 | 0.02385247 | 0.92 | 1744 | tags=33%, list=6%, signal=36% |
| MCBRYAN_PUBERTAL_BREAST_3_4WK_DN | 34 | 0.7513469 | 1.7543725 | 0 | 0.02417717 | 0.67 | 2806 | tags=38%, list=10%, signal=43% |
| GO_AXON_ENSHEATHMENT_IN_CENTRAL_NERVOUS_SYSTEM | 20 | 0.8463368 | 1.7543132 | 0 | 0.02402241 | 0.93 | 161 | tags=20%, list=1%, signal=20% |
| MISSIAGLIA_REGULATED_BY_METHYLATION_UP | 104 | 0.64513135 | 1.7541653 | 0 | 0.02408821 | 0.67 | 1776 | tags=27%, list=7%, signal=29% |
| GO_ATPASE_ACTIVITY_COUPLED_TO_TRANSMEMBRANE_MOVEMENT_OF_IONS_ROTATIONAL_MECHANISM | 19 | 0.83449274 | 1.7533652 | 0 | 0.02440821 | 0.94 | 1206 | tags=37%, list=4%, signal=39% |
| GO_REGULATION_OF_CELLULAR_KETONE_METABOLIC_PROCESS | 165 | 0.5834504 | 1.753074 | 0 | 0.02456813 | 0.94 | 2476 | tags=28%, list=9%, signal=31% |
| GO_HEMATOPOIETIC_PROGENITOR_CELL_DIFFERENTIATION | 162 | 0.5935415 | 1.752452 | 0 | 0.02461846 | 0.94 | 2270 | tags=26%, list=8%, signal=28% |
| GO_ENVELOPE | 1081 | 0.5180195 | 1.7520249 | 0 | 0.02456141 | 0.95 | 2937 | tags=30%, list=11%, signal=33% |
| REACTOME_INSULIN_RECEPTOR_RECYCLING | 24 | 0.79041904 | 1.7520117 | 0 | 0.02482424 | 0.7 | 1206 | tags=33%, list=4%, signal=35% |
| GO_MITOCHONDRIAL_TRANSLATION | 132 | 0.6177352 | 1.7513901 | 0 | 0.02482664 | 0.96 | 2348 | tags=40%, list=9%, signal=44% |
| WESTON_VEGFA_TARGETS | 81 | 0.6500266 | 1.7503359 | 0 | 0.02493427 | 0.7 | 2614 | tags=25%, list=10%, signal=27% |
| HP_WEAKNESS_DUE_TO_UPPER_MOTOR_NEURON_DYSFUNCTION | 380 | 0.5554414 | 1.7496276 | 0 | 0.02551661 | 0.96 | 2421 | tags=23%, list=9%, signal=25% |
| BOWIE_RESPONSE_TO_EXTRACELLULAR_MATRIX | 15 | 0.9113811 | 1.749309 | 0 | 0.02523449 | 0.7 | 1289 | tags=60%, list=5%, signal=63% |
| GO_REGULATION_OF_HEMATOPOIETIC_PROGENITOR_CELL_DIFFERENTIATION | 82 | 0.6647401 | 1.7489533 | 0 | 0.02577021 | 0.97 | 1624 | tags=32%, list=6%, signal=34% |
| HP_LEFT_VENTRICULAR_HYPERTROPHY | 69 | 0.6533019 | 1.7474256 | 0 | 0.02644558 | 0.98 | 1994 | tags=22%, list=7%, signal=23% |
| LU_AGING_BRAIN_UP | 249 | 0.5748982 | 1.7470515 | 0 | 0.02611616 | 0.73 | 3053 | tags=38%, list=11%, signal=42% |
| HP_FOCAL_T2_HYPERINTENSE_BRAINSTEM_LESION | 30 | 0.84275293 | 1.7467333 | 0 | 0.02669142 | 0.98 | 2254 | tags=70%, list=8%, signal=76% |
| GO_PHAGOCYTIC_VESICLE_MEMBRANE | 68 | 0.6473756 | 1.7445737 | 0 | 0.0275599 | 0.98 | 2425 | tags=29%, list=9%, signal=32% |
| PELLICCIOTTA_HDAC_IN_ANTIGEN_PRESENTATION_UP | 61 | 0.7092997 | 1.7444412 | 0 | 0.0275665 | 0.74 | 1815 | tags=51%, list=7%, signal=54% |
| BYSTRYKH_HEMATOPOIESIS_STEM_CELL_QTL_CIS | 121 | 0.62915564 | 1.7436359 | 0 | 0.02744508 | 0.75 | 2819 | tags=41%, list=10%, signal=46% |
| HP_RECURRENT_PAROXYSMAL_HEADACHE | 25 | 0.8105673 | 1.7434254 | 0 | 0.02779557 | 0.98 | 1365 | tags=44%, list=5%, signal=46% |
| HP_ABNORMAL_MYOCARDIUM_MORPHOLOGY | 431 | 0.5582145 | 1.7434214 | 0 | 0.0276174 | 0.98 | 3528 | tags=33%, list=13%, signal=38% |
| GO_REGULATION_OF_INNATE_IMMUNE_RESPONSE | 258 | 0.543419 | 1.7432282 | 0 | 0.02744149 | 0.98 | 2354 | tags=23%, list=9%, signal=25% |
| HP_HYPOTHERMIA | 22 | 0.86146355 | 1.7422 | 0 | 0.02777367 | 0.98 | 1503 | tags=45%, list=6%, signal=48% |
| GO_MHC_PROTEIN_BINDING | 23 | 0.83007216 | 1.7414639 | 0 | 0.02810285 | 0.98 | 1765 | tags=48%, list=7%, signal=51% |
| KRIEG_KDM3A_TARGETS_NOT_HYPOXIA | 177 | 0.58718604 | 1.7406726 | 0 | 0.02902302 | 0.8 | 3482 | tags=42%, list=13%, signal=48% |
| GO_PEPTIDYL_PROLINE_MODIFICATION | 48 | 0.7077691 | 1.7400999 | 0 | 0.02932157 | 0.99 | 3386 | tags=48%, list=13%, signal=55% |
| GO_REGULATION_OF_GLYCOPROTEIN_METABOLIC_PROCESS | 40 | 0.7501892 | 1.7393459 | 0 | 0.02993498 | 1 | 1427 | tags=28%, list=5%, signal=29% |
| GO_TUMOR_NECROSIS_FACTOR_MEDIATED_SIGNALING_PATHWAY | 149 | 0.60476685 | 1.739128 | 0 | 0.03004648 | 1 | 3182 | tags=34%, list=12%, signal=38% |
| HP_MYOCLONIC_SEIZURE | 93 | 0.64413214 | 1.7385604 | 0 | 0.03045092 | 1 | 2466 | tags=27%, list=9%, signal=29% |
| REACTOME_ABC_TRANSPORTER_DISORDERS | 72 | 0.6487525 | 1.738069 | 0 | 0.03019583 | 0.83 | 2352 | tags=42%, list=9%, signal=46% |
| HP_FOCAL_SEGMENTAL_GLOMERULOSCLEROSIS | 40 | 0.73046595 | 1.7368853 | 0 | 0.03113982 | 1 | 1674 | tags=20%, list=6%, signal=21% |
| HP_OPHTHALMOPARESIS | 249 | 0.56186724 | 1.7365389 | 0 | 0.03104782 | 1 | 2442 | tags=22%, list=9%, signal=24% |
| YAN_ESCAPE_FROM_ANOIKIS | 21 | 0.8088129 | 1.735207 | 0 | 0.03171631 | 0.85 | 2141 | tags=57%, list=8%, signal=62% |
| LEIN_ASTROCYTE_MARKERS | 37 | 0.7085895 | 1.7330655 | 0 | 0.03228848 | 0.85 | 791 | tags=32%, list=3%, signal=33% |
| RICKMAN_HEAD_AND_NECK_CANCER_D | 17 | 0.8622827 | 1.7323475 | 0 | 0.03286126 | 0.85 | 1751 | tags=29%, list=7%, signal=31% |
| GO_HEMATOPOIETIC_STEM_CELL_DIFFERENTIATION | 83 | 0.6570553 | 1.7322837 | 0 | 0.03240253 | 1 | 1624 | tags=31%, list=6%, signal=33% |
| JISON_SICKLE_CELL_DISEASE_UP | 166 | 0.6105476 | 1.7317213 | 0 | 0.03287722 | 0.86 | 2488 | tags=37%, list=9%, signal=40% |
| BOWIE_RESPONSE_TO_TAMOXIFEN | 15 | 0.8800349 | 1.7311225 | 0 | 0.03271015 | 0.86 | 1289 | tags=47%, list=5%, signal=49% |
| HP_HEMIANOPIA | 37 | 0.7438678 | 1.7307675 | 0 | 0.03268703 | 1 | 854 | tags=16%, list=3%, signal=17% |
| GO_CELLULAR_LIPID_CATABOLIC_PROCESS | 197 | 0.5851814 | 1.7306286 | 0 | 0.03258725 | 1 | 3957 | tags=40%, list=15%, signal=47% |
| GRANDVAUX_IRF3_TARGETS_UP | 11 | 0.905434 | 1.7304685 | 0.02380952 | 0.03342309 | 0.87 | 1370 | tags=36%, list=5%, signal=38% |
| REACTOME_TRANSFERRIN_ENDOCYTOSIS_AND_RECYCLING | 30 | 0.8016803 | 1.7302804 | 0 | 0.03325473 | 0.87 | 1206 | tags=33%, list=4%, signal=35% |
| SMID_BREAST_CANCER_LUMINAL_B_DN | 467 | 0.53449625 | 1.7301497 | 0 | 0.03308996 | 0.87 | 1851 | tags=15%, list=7%, signal=16% |
| NIKOLSKY_BREAST_CANCER_8Q23_Q24_AMPLICON | 121 | 0.63942283 | 1.7301203 | 0 | 0.03275572 | 0.87 | 3094 | tags=36%, list=11%, signal=41% |
| GO_COPII_COATED_ER_TO_GOLGI_TRANSPORT_VESICLE | 75 | 0.64992243 | 1.7300284 | 0 | 0.03277275 | 1 | 2180 | tags=43%, list=8%, signal=46% |
| HP_ABNORMALITY_OF_RETINAL_PIGMENTATION | 289 | 0.56378585 | 1.7298557 | 0 | 0.03257997 | 1 | 3810 | tags=33%, list=14%, signal=37% |
| HP_ABSENCE_OF_SECONDARY_SEX_CHARACTERISTICS | 34 | 0.6987928 | 1.7298055 | 0 | 0.03248324 | 1 | 602 | tags=12%, list=2%, signal=12% |
| HP_HEMIPLEGIA_HEMIPARESIS | 176 | 0.5903207 | 1.7296607 | 0 | 0.03238866 | 1 | 2350 | tags=19%, list=9%, signal=21% |
| GO_AMINE_CATABOLIC_PROCESS | 19 | 0.82315105 | 1.7295916 | 0 | 0.03220144 | 1 | 2593 | tags=21%, list=10%, signal=23% |
| HP_ABNORMALITY_OF_THE_CEREBRAL_VASCULATURE | 246 | 0.5555992 | 1.7272882 | 0 | 0.03312291 | 1 | 3165 | tags=28%, list=12%, signal=31% |
| GO_MITOCHONDRIAL_ELECTRON_TRANSPORT_UBIQUINOL_TO_CYTOCHROME_C | 12 | 0.9034347 | 1.7268981 | 0 | 0.03302546 | 1 | 1812 | tags=67%, list=7%, signal=71% |
| GO_INNER_MITOCHONDRIAL_MEMBRANE_ORGANIZATION | 44 | 0.7209354 | 1.726311 | 0 | 0.03329474 | 1 | 3671 | tags=59%, list=14%, signal=68% |
| IGLESIAS_E2F_TARGETS_UP | 138 | 0.5964678 | 1.7253426 | 0 | 0.03496491 | 0.9 | 1064 | tags=27%, list=4%, signal=28% |
| REACTOME_DEFECTIVE_CFTR_CAUSES_CYSTIC_FIBROSIS | 58 | 0.6598353 | 1.7247083 | 0 | 0.03478587 | 0.9 | 2352 | tags=52%, list=9%, signal=57% |
| GO_ANTIGEN_PROCESSING_AND_PRESENTATION_OF_PEPTIDE_ANTIGEN | 170 | 0.59332436 | 1.7246355 | 0 | 0.03355805 | 1 | 2364 | tags=33%, list=9%, signal=36% |
| ZHAN_MULTIPLE_MYELOMA_MS_UP | 41 | 0.75144005 | 1.7242031 | 0 | 0.03461 | 0.9 | 2741 | tags=34%, list=10%, signal=38% |
| HP_APNEA | 224 | 0.57285434 | 1.7238526 | 0 | 0.03399876 | 1 | 3876 | tags=36%, list=14%, signal=42% |
| HP_ABNORMAL_URINE_PH | 168 | 0.58615386 | 1.723166 | 0 | 0.03408001 | 1 | 3597 | tags=38%, list=13%, signal=44% |
| HP_ABNORMAL_LIVER_MORPHOLOGY | 680 | 0.52696204 | 1.7228966 | 0 | 0.03424534 | 1 | 3528 | tags=30%, list=13%, signal=34% |
| HP_EXERCISE_INTOLERANCE | 70 | 0.69083 | 1.7228711 | 0 | 0.03405614 | 1 | 1088 | tags=24%, list=4%, signal=25% |
| ZHONG_SECRETOME_OF_LUNG_CANCER_AND_MACROPHAGE | 70 | 0.6651804 | 1.7227333 | 0 | 0.03543109 | 0.9 | 1919 | tags=43%, list=7%, signal=46% |
| GO_MONOVALENT_INORGANIC_CATION_HOMEOSTASIS | 131 | 0.58796704 | 1.7227086 | 0 | 0.03395738 | 1 | 1952 | tags=21%, list=7%, signal=23% |
| GO_AROMATIC_AMINO_ACID_FAMILY_METABOLIC_PROCESS | 33 | 0.7504233 | 1.722454 | 0 | 0.0338584 | 1 | 3426 | tags=24%, list=13%, signal=28% |
| HP_POSTURAL_TREMOR | 46 | 0.70577323 | 1.7214003 | 0 | 0.03410772 | 1 | 854 | tags=26%, list=3%, signal=27% |
| PLASARI_TGFB1_TARGETS_10HR_DN | 239 | 0.56623584 | 1.7213398 | 0 | 0.03606428 | 0.9 | 3271 | tags=32%, list=12%, signal=36% |
| GO_MIDBRAIN_DEVELOPMENT | 78 | 0.639223 | 1.7210382 | 0 | 0.03409637 | 1 | 2207 | tags=33%, list=8%, signal=36% |
| GO_ENDOPLASMIC_RETICULUM_LUMEN | 265 | 0.55057895 | 1.7205132 | 0 | 0.0343462 | 1 | 3452 | tags=29%, list=13%, signal=33% |
| GO_METANEPHRIC_RENAL_VESICLE_MORPHOGENESIS | 14 | 0.89132726 | 1.7195728 | 0 | 0.03475982 | 1 | 158 | tags=21%, list=1%, signal=22% |
| WP_THE_HUMAN_IMMUNE_RESPONSE_TO_TUBERCULOSIS | 22 | 0.82046485 | 1.7191437 | 0 | 0.03684982 | 0.91 | 2378 | tags=55%, list=9%, signal=60% |
| KEGG_CARDIAC_MUSCLE_CONTRACTION | 70 | 0.67754334 | 1.7188476 | 0 | 0.03682643 | 0.91 | 1740 | tags=30%, list=6%, signal=32% |
| HP_LOWER_LIMB_SPASTICITY | 172 | 0.57133657 | 1.7186911 | 0 | 0.03474395 | 1 | 2673 | tags=28%, list=10%, signal=31% |
| GO_ASTROCYTE_DIFFERENTIATION | 75 | 0.65945864 | 1.7186826 | 0 | 0.03456013 | 1 | 2318 | tags=27%, list=9%, signal=29% |
| JOHANSSON_BRAIN_CANCER_EARLY_VS_LATE_DN | 41 | 0.71346277 | 1.7175025 | 0 | 0.03759591 | 0.91 | 1600 | tags=34%, list=6%, signal=36% |
| HP_HYPERVENTILATION | 25 | 0.81237173 | 1.7173645 | 0 | 0.03522493 | 1 | 2293 | tags=40%, list=9%, signal=44% |
| HP_ABNORMALITY_OF_KREBS_CYCLE_METABOLISM | 14 | 0.88993394 | 1.7169623 | 0 | 0.03537253 | 1 | 2254 | tags=71%, list=8%, signal=78% |
| WP_MITOCHONDRIAL_CIII_ASSEMBLY | 15 | 0.9115473 | 1.7165939 | 0 | 0.03756251 | 0.91 | 1036 | tags=60%, list=4%, signal=62% |
| GO_POSITIVE_REGULATION_OF_LIPASE_ACTIVITY | 59 | 0.6638611 | 1.7156035 | 0 | 0.03602479 | 1 | 2728 | tags=29%, list=10%, signal=32% |
| YAO_TEMPORAL_RESPONSE_TO_PROGESTERONE_CLUSTER_9 | 69 | 0.65032053 | 1.715359 | 0.0212766 | 0.03783284 | 0.91 | 2994 | tags=43%, list=11%, signal=49% |
| HP_EPISODIC_RESPIRATORY_DISTRESS | 15 | 0.89748013 | 1.715276 | 0 | 0.03600496 | 1 | 854 | tags=40%, list=3%, signal=41% |
| LINDGREN_BLADDER_CANCER_CLUSTER_2B | 349 | 0.5325204 | 1.71453 | 0 | 0.03779705 | 0.92 | 2821 | tags=27%, list=10%, signal=29% |
| HP_SPASTIC_PARAPLEGIA | 119 | 0.6121215 | 1.714286 | 0 | 0.03615076 | 1 | 2645 | tags=31%, list=10%, signal=34% |
| OHGUCHI_LIVER_HNF4A_TARGETS_UP | 39 | 0.68542624 | 1.7142105 | 0 | 0.03776375 | 0.92 | 2625 | tags=28%, list=10%, signal=31% |
| GO_AROMATIC_AMINO_ACID_FAMILY_CATABOLIC_PROCESS | 23 | 0.801317 | 1.7133726 | 0 | 0.03653685 | 1 | 1291 | tags=13%, list=5%, signal=14% |
| TAKEDA_TARGETS_OF_NUP98_HOXA9_FUSION_10D_UP | 150 | 0.5971459 | 1.7133001 | 0 | 0.03788297 | 0.92 | 2544 | tags=25%, list=9%, signal=28% |
| GO_PROTEIN_PEPTIDYL_PROLYL_ISOMERIZATION | 33 | 0.74007905 | 1.7132117 | 0 | 0.03643389 | 1 | 3869 | tags=64%, list=14%, signal=74% |
| GO_MYELIN_SHEATH | 43 | 0.64214635 | 1.7130789 | 0 | 0.03624895 | 1 | 1181 | tags=21%, list=4%, signal=22% |
| PICCALUGA_ANGIOIMMUNOBLASTIC_LYMPHOMA_UP | 187 | 0.59301263 | 1.7128448 | 0 | 0.03799102 | 0.93 | 1978 | tags=24%, list=7%, signal=26% |
| GRAHAM_CML_DIVIDING_VS_NORMAL_DIVIDING_DN | 9 | 0.91979605 | 1.7109731 | 0 | 0.03825471 | 0.94 | 1903 | tags=44%, list=7%, signal=48% |
| SERVITJA_ISLET_HNF1A_TARGETS_UP | 157 | 0.5601144 | 1.7109303 | 0 | 0.03792206 | 0.94 | 2800 | tags=31%, list=10%, signal=34% |
| TAKEDA_TARGETS_OF_NUP98_HOXA9_FUSION_8D_UP | 123 | 0.58717936 | 1.7097359 | 0 | 0.03818494 | 0.95 | 2141 | tags=24%, list=8%, signal=25% |
| YAMAZAKI_TCEB3_TARGETS_UP | 154 | 0.5738703 | 1.7094984 | 0 | 0.03843477 | 0.96 | 2793 | tags=29%, list=10%, signal=32% |
| REACTOME_ABC_FAMILY_PROTEINS_MEDIATED_TRANSPORT | 95 | 0.6409422 | 1.7094115 | 0 | 0.03810905 | 0.96 | 1624 | tags=33%, list=6%, signal=35% |
| EINAV_INTERFERON_SIGNATURE_IN_CANCER | 23 | 0.8286199 | 1.709285 | 0 | 0.03807488 | 0.96 | 1371 | tags=39%, list=5%, signal=41% |
| HP_DYSTONIA | 374 | 0.53526545 | 1.7091688 | 0 | 0.03914231 | 1 | 3266 | tags=31%, list=12%, signal=35% |
| GO_PH_REDUCTION | 53 | 0.7097706 | 1.708931 | 0 | 0.03902711 | 1 | 1952 | tags=26%, list=7%, signal=28% |
| HP_NEPHROPATHY | 85 | 0.6298319 | 1.7087451 | 0.04347826 | 0.03891182 | 1 | 3535 | tags=29%, list=13%, signal=34% |
| HP_INCREASED_CIRCULATING_ANTIBODY_LEVEL | 65 | 0.64924437 | 1.708742 | 0 | 0.03871823 | 1 | 2427 | tags=18%, list=9%, signal=20% |
| ROZANOV_MMP14_TARGETS_UP | 233 | 0.57107776 | 1.7086241 | 0 | 0.03803753 | 0.96 | 3158 | tags=30%, list=12%, signal=33% |
| ZHAN_MULTIPLE_MYELOMA_LB_DN | 31 | 0.7760357 | 1.7085005 | 0 | 0.03799916 | 0.96 | 1599 | tags=45%, list=6%, signal=48% |
| HP_HEPATIC_NECROSIS | 15 | 0.88524055 | 1.7076051 | 0 | 0.03884222 | 1 | 2883 | tags=80%, list=11%, signal=90% |
| WESTON_VEGFA_TARGETS_3HR | 57 | 0.68277943 | 1.7073283 | 0 | 0.03810366 | 0.96 | 1449 | tags=21%, list=5%, signal=22% |
| GO_PEPTIDASE_COMPLEX | 85 | 0.6498535 | 1.707253 | 0 | 0.03865087 | 1 | 2316 | tags=40%, list=9%, signal=44% |
| GO_PLATELET_DEGRANULATION | 107 | 0.6151398 | 1.7072358 | 0 | 0.03853875 | 1 | 3369 | tags=36%, list=13%, signal=40% |
| CHIARADONNA_NEOPLASTIC_TRANSFORMATION_KRAS_DN | 134 | 0.5976175 | 1.7068083 | 0 | 0.03806647 | 0.96 | 2162 | tags=24%, list=8%, signal=26% |
| GO_INTEGRIN_BINDING | 127 | 0.58734185 | 1.7058234 | 0 | 0.03952206 | 1 | 2429 | tags=22%, list=9%, signal=24% |
| GO_MHC_CLASS_I_PROTEIN_BINDING | 16 | 0.84613615 | 1.7052343 | 0 | 0.03963878 | 1 | 1663 | tags=56%, list=6%, signal=60% |
| GO_AMYLOID_FIBRIL_FORMATION | 16 | 0.8726821 | 1.7039654 | 0 | 0.0406867 | 1 | 444 | tags=38%, list=2%, signal=38% |
| HP_PSYCHOSIS | 86 | 0.61763996 | 1.7038544 | 0 | 0.04049109 | 1 | 2645 | tags=24%, list=10%, signal=27% |
| HP_HYPERTRICHOSIS | 193 | 0.5732904 | 1.7037846 | 0 | 0.04029735 | 1 | 3100 | tags=34%, list=12%, signal=38% |
| HP_SEGMENTAL_PERIPHERAL_DEMYELINATION_REMYELINATION | 16 | 0.8463392 | 1.702793 | 0 | 0.04124527 | 1 | 923 | tags=56%, list=3%, signal=58% |
| NAKAYAMA_SOFT_TISSUE_TUMORS_PCA2_DN | 61 | 0.64084494 | 1.7026304 | 0 | 0.03995307 | 0.96 | 2916 | tags=23%, list=11%, signal=26% |
| REACTOME_RESPONSE_TO_ELEVATED_PLATELET_CYTOSOLIC_CA2_ | 113 | 0.61280537 | 1.7025204 | 0 | 0.03963345 | 0.96 | 3409 | tags=36%, list=13%, signal=41% |
| GO_MALE_GENITALIA_DEVELOPMENT | 18 | 0.79423225 | 1.702192 | 0 | 0.0412023 | 1 | 206 | tags=11%, list=1%, signal=11% |
| HP_STUTTERING | 10 | 0.92886394 | 1.700213 | 0 | 0.04350575 | 1 | 854 | tags=50%, list=3%, signal=52% |
| HP_RED_EYE | 81 | 0.607558 | 1.6998869 | 0 | 0.04352775 | 1 | 1967 | tags=19%, list=7%, signal=20% |
| BLALOCK_ALZHEIMERS_DISEASE_DN | 1180 | 0.507614 | 1.6998401 | 0 | 0.04120764 | 0.96 | 3057 | tags=34%, list=11%, signal=37% |
| GO_RESPIRATORY_CHAIN_COMPLEX_III | 11 | 0.944465 | 1.6996258 | 0 | 0.04347331 | 1 | 1036 | tags=73%, list=4%, signal=76% |
| YAO_TEMPORAL_RESPONSE_TO_PROGESTERONE_CLUSTER_1 | 61 | 0.6651136 | 1.6995716 | 0 | 0.0410101 | 0.96 | 2763 | tags=31%, list=10%, signal=35% |
| GO_ANTIBACTERIAL_HUMORAL_RESPONSE | 14 | 0.8151469 | 1.6981642 | 0 | 0.04475739 | 1 | 1272 | tags=21%, list=5%, signal=22% |
| GO_L_PHENYLALANINE_METABOLIC_PROCESS | 8 | 0.90396327 | 1.6979644 | 0 | 0.04484617 | 1 | 1291 | tags=25%, list=5%, signal=26% |
| HP_DEVELOPMENTAL_CATARACT | 85 | 0.64589274 | 1.6974101 | 0 | 0.04530246 | 1 | 2434 | tags=29%, list=9%, signal=32% |
| GO_OXIDOREDUCTASE_ACTIVITY_ACTING_ON_A_HEME_GROUP_OF_DONORS | 25 | 0.80062 | 1.6971289 | 0 | 0.04524165 | 1 | 1740 | tags=48%, list=6%, signal=51% |
| GO_REGULATION_OF_MORPHOGENESIS_OF_AN_EPITHELIUM | 168 | 0.56175107 | 1.6969733 | 0 | 0.04532713 | 1 | 2270 | tags=25%, list=8%, signal=27% |
| REACTOME_NEGATIVE_REGULATION_OF_NOTCH4_SIGNALING | 51 | 0.6681634 | 1.6968317 | 0 | 0.04279937 | 0.96 | 805 | tags=39%, list=3%, signal=40% |
| CHEBOTAEV_GR_TARGETS_DN | 113 | 0.627741 | 1.6961925 | 0 | 0.04337874 | 0.96 | 2340 | tags=31%, list=9%, signal=34% |
| ZHONG_SECRETOME_OF_LUNG_CANCER_AND_FIBROBLAST | 126 | 0.61179775 | 1.6959604 | 0 | 0.04304506 | 0.96 | 1919 | tags=37%, list=7%, signal=40% |
| SEITZ_NEOPLASTIC_TRANSFORMATION_BY_8P_DELETION_UP | 60 | 0.6810005 | 1.6949604 | 0 | 0.04322035 | 0.96 | 1692 | tags=27%, list=6%, signal=28% |
| HP_KERATOCONJUNCTIVITIS | 34 | 0.73117393 | 1.6943173 | 0 | 0.04679597 | 1 | 1967 | tags=24%, list=7%, signal=25% |
| HP_POOR_HEAD_CONTROL | 155 | 0.5924763 | 1.6934991 | 0 | 0.04716355 | 1 | 2400 | tags=32%, list=9%, signal=35% |
| GO_LEUKOCYTE_MIGRATION_INVOLVED_IN_INFLAMMATORY_RESPONSE | 12 | 0.910793 | 1.693483 | 0 | 0.0469511 | 1 | 1692 | tags=50%, list=6%, signal=53% |
| GO_PRIMARY_ACTIVE_TRANSMEMBRANE_TRANSPORTER_ACTIVITY | 97 | 0.5989727 | 1.6927488 | 0 | 0.0473869 | 1 | 1315 | tags=20%, list=5%, signal=21% |
| GO_PROTON_TRANSPORTING_ATP_SYNTHASE_ACTIVITY_ROTATIONAL_MECHANISM | 16 | 0.8687179 | 1.6925538 | 0 | 0.04717535 | 1 | 1251 | tags=63%, list=5%, signal=66% |
| AKL_HTLV1_INFECTION_UP | 24 | 0.7454713 | 1.6924806 | 0 | 0.04455759 | 0.97 | 2060 | tags=42%, list=8%, signal=45% |
| GO_PROTON_TRANSPORTING_TWO_SECTOR_ATPASE_COMPLEX_CATALYTIC_DOMAIN | 16 | 0.86002225 | 1.6923141 | 0 | 0.04724995 | 1 | 1488 | tags=63%, list=6%, signal=66% |
| HP_ENCEPHALOPATHY | 235 | 0.55967295 | 1.6921976 | 0 | 0.04711155 | 1 | 2466 | tags=30%, list=9%, signal=33% |
| HP_ABNORMAL_METABOLIC_BRAIN_IMAGING_BY_MRS | 25 | 0.8159782 | 1.6912551 | 0 | 0.04761154 | 1 | 1830 | tags=52%, list=7%, signal=56% |
| KEGG_AUTOIMMUNE_THYROID_DISEASE | 23 | 0.8025046 | 1.6903652 | 0.02439024 | 0.04561876 | 0.98 | 1347 | tags=22%, list=5%, signal=23% |
| GO_ACTIVATION_OF_PHOSPHOLIPASE_C_ACTIVITY | 27 | 0.7680844 | 1.6899946 | 0.0212766 | 0.04838152 | 1 | 2728 | tags=26%, list=10%, signal=29% |
| STEIN_ESRRA_TARGETS_UP | 358 | 0.5508237 | 1.6898032 | 0 | 0.04578338 | 0.98 | 3899 | tags=44%, list=14%, signal=51% |
| HP_BULBAR_SIGNS | 26 | 0.78870934 | 1.689779 | 0 | 0.04858999 | 1 | 1776 | tags=38%, list=7%, signal=41% |
| HP_RETINAL_ARTERIAL_TORTUOSITY | 9 | 0.9501691 | 1.6894034 | 0 | 0.04879597 | 1 | 854 | tags=67%, list=3%, signal=69% |
| HP_ABNORMALITY_OF_THE_BASAL_GANGLIA | 174 | 0.57752824 | 1.6885118 | 0 | 0.04969271 | 1 | 2443 | tags=28%, list=9%, signal=30% |
| GO_MITOCHONDRIAL_MATRIX | 439 | 0.53935176 | 1.6882529 | 0 | 0.04975608 | 1 | 3512 | tags=40%, list=13%, signal=45% |
| MILI_PSEUDOPODIA_HAPTOTAXIS_UP | 489 | 0.5107837 | 1.6878287 | 0 | 0.04683053 | 0.98 | 3206 | tags=41%, list=12%, signal=46% |
| MARTINEZ_RESPONSE_TO_TRABECTEDIN_UP | 67 | 0.64215285 | 1.6874819 | 0 | 0.04660948 | 0.98 | 1812 | tags=28%, list=7%, signal=30% |
| LEE_LIVER_CANCER_E2F1_UP | 55 | 0.6886056 | 1.6869308 | 0 | 0.04676083 | 0.98 | 1195 | tags=27%, list=4%, signal=28% |
| KIM_RESPONSE_TO_TSA_AND_DECITABINE_UP | 95 | 0.60552245 | 1.6868839 | 0 | 0.04642198 | 0.98 | 1615 | tags=16%, list=6%, signal=17% |
| REACTOME_REGULATION_OF_RUNX2_EXPRESSION_AND_ACTIVITY | 69 | 0.6604444 | 1.6865573 | 0 | 0.04620951 | 0.98 | 2396 | tags=43%, list=9%, signal=48% |
| HP_AXIAL_MUSCLE_WEAKNESS | 21 | 0.78705215 | 1.6861583 | 0 | 0.05105575 | 1 | 732 | tags=14%, list=3%, signal=15% |
| REACTOME_DISORDERS_OF_TRANSMEMBRANE_TRANSPORTERS | 154 | 0.55858487 | 1.6850916 | 0 | 0.04756551 | 0.98 | 2352 | tags=26%, list=9%, signal=28% |
| DE_YY1_TARGETS_UP | 11 | 0.9036816 | 1.6849622 | 0 | 0.04734704 | 0.98 | 565 | tags=18%, list=2%, signal=19% |
| GO_ISOMERASE_ACTIVITY | 134 | 0.58964056 | 1.6845208 | 0 | 0.05199938 | 1 | 3386 | tags=42%, list=13%, signal=48% |
| DELYS_THYROID_CANCER_DN | 204 | 0.5472442 | 1.6844202 | 0 | 0.04724919 | 0.98 | 3000 | tags=25%, list=11%, signal=27% |
| HP_PROGRESSIVE_SPASTIC_PARAPLEGIA | 53 | 0.70425993 | 1.6840447 | 0 | 0.05205042 | 1 | 2400 | tags=40%, list=9%, signal=43% |
| GO_NEGATIVE_REGULATION_OF_METANEPHROS_DEVELOPMENT | 7 | 0.9788015 | 1.6832035 | 0 | 0.05250869 | 1 | 158 | tags=43%, list=1%, signal=43% |
| JINESH_BLEBBISHIELD_VS_LIVE_CONTROL_DN | 239 | 0.5549434 | 1.6831613 | 0 | 0.04763629 | 0.98 | 3770 | tags=38%, list=14%, signal=44% |
| GO_CELLULAR_MONOVALENT_INORGANIC_CATION_HOMEOSTASIS | 99 | 0.60110545 | 1.6821916 | 0 | 0.05316438 | 1 | 1952 | tags=23%, list=7%, signal=25% |
| HP_KERATITIS | 60 | 0.64394784 | 1.6807593 | 0 | 0.05455731 | 1 | 1967 | tags=18%, list=7%, signal=20% |
| GRAHAM_CML_DIVIDING_VS_NORMAL_QUIESCENT_DN | 76 | 0.6226822 | 1.6805868 | 0 | 0.04965622 | 0.98 | 3203 | tags=29%, list=12%, signal=33% |
| GO_REGULATION_OF_PH | 89 | 0.6334252 | 1.680543 | 0 | 0.05459935 | 1 | 1527 | tags=22%, list=6%, signal=24% |
| LEE_LIVER_CANCER_DENA_DN | 59 | 0.65469426 | 1.6801335 | 0 | 0.04954904 | 0.98 | 4677 | tags=36%, list=17%, signal=43% |
| HP_GOITER | 39 | 0.6913183 | 1.6801004 | 0 | 0.05497021 | 1 | 906 | tags=21%, list=3%, signal=21% |
| WEBER_METHYLATED_IN_COLON_CANCER | 14 | 0.8143573 | 1.6789222 | 0 | 0.05001203 | 0.98 | 3195 | tags=50%, list=12%, signal=57% |
| HP_ABNORMAL_ABDOMEN_MORPHOLOGY | 585 | 0.51494205 | 1.6787201 | 0 | 0.05620791 | 1 | 3471 | tags=30%, list=13%, signal=34% |
| HP_FLUCTUATIONS_IN_CONSCIOUSNESS | 16 | 0.8777429 | 1.6785406 | 0 | 0.05604156 | 1 | 854 | tags=44%, list=3%, signal=45% |
| GO_VESICLE_LUMEN | 280 | 0.5383235 | 1.6780096 | 0 | 0.05620609 | 1 | 3045 | tags=31%, list=11%, signal=35% |
| HP_IMPAIRED_VISUOSPATIAL_CONSTRUCTIVE_COGNITION | 22 | 0.816215 | 1.6777864 | 0.02083333 | 0.05630546 | 1 | 854 | tags=27%, list=3%, signal=28% |
| LINDVALL_IMMORTALIZED_BY_TERT_DN | 67 | 0.6589424 | 1.6773331 | 0 | 0.05105395 | 0.99 | 1425 | tags=24%, list=5%, signal=25% |
| GRUETZMANN_PANCREATIC_CANCER_UP | 331 | 0.5385824 | 1.6772652 | 0 | 0.05070898 | 0.99 | 2346 | tags=29%, list=9%, signal=31% |
| HP_HEPATIC_FAILURE | 131 | 0.58904684 | 1.6772032 | 0 | 0.05679232 | 1 | 3528 | tags=37%, list=13%, signal=43% |
| WANG_MLL_TARGETS | 249 | 0.58743936 | 1.6770786 | 0 | 0.05036866 | 0.99 | 3139 | tags=22%, list=12%, signal=25% |
| GO_RESPONSE_TO_OXYGEN_LEVELS | 354 | 0.529509 | 1.6766908 | 0 | 0.0568225 | 1 | 2360 | tags=24%, list=9%, signal=26% |
| GO_MITOCHONDRIAL_GENE_EXPRESSION | 158 | 0.5885273 | 1.6763909 | 0 | 0.05665747 | 1 | 3909 | tags=47%, list=15%, signal=55% |
| HP_ABNORMALITY_OF_THE_CEREBROSPINAL_FLUID | 424 | 0.5225456 | 1.6763233 | 0 | 0.05649345 | 1 | 3291 | tags=29%, list=12%, signal=32% |
| GNATENKO_PLATELET_SIGNATURE | 34 | 0.7166574 | 1.676284 | 0.02702703 | 0.05048406 | 0.99 | 1006 | tags=44%, list=4%, signal=46% |
| REACTOME_INTERLEUKIN_1_FAMILY_SIGNALING | 127 | 0.5606654 | 1.6755548 | 0 | 0.05082554 | 0.99 | 2352 | tags=28%, list=9%, signal=30% |
| GO_PROTON_TRANSPORTING_V_TYPE_ATPASE_COMPLEX | 25 | 0.8049023 | 1.6755397 | 0 | 0.05690628 | 1 | 1206 | tags=32%, list=4%, signal=33% |
| KEGG_PYRUVATE_METABOLISM | 36 | 0.69397837 | 1.6750399 | 0 | 0.05071286 | 0.99 | 3769 | tags=53%, list=14%, signal=61% |
| HUANG_DASATINIB_RESISTANCE_UP | 71 | 0.6381 | 1.6746594 | 0 | 0.05092976 | 0.99 | 1376 | tags=18%, list=5%, signal=19% |
| HP_ABNORMALITY_OF_THE_COAGULATION_CASCADE | 101 | 0.58766186 | 1.674574 | 0 | 0.05725713 | 1 | 4526 | tags=43%, list=17%, signal=51% |
| BAE_BRCA1_TARGETS_UP | 64 | 0.65902734 | 1.673977 | 0 | 0.05103533 | 0.99 | 2262 | tags=36%, list=8%, signal=39% |
| KEGG_PROTEIN_EXPORT | 21 | 0.8089092 | 1.6739728 | 0 | 0.05070607 | 0.99 | 752 | tags=48%, list=3%, signal=49% |
| GO_REGULATION_OF_CELLULAR_PH | 82 | 0.6316595 | 1.6738219 | 0 | 0.05753628 | 1 | 1527 | tags=23%, list=6%, signal=24% |
| HP_INFANTILE_SPASMS | 51 | 0.67247206 | 1.6726813 | 0 | 0.0583282 | 1 | 3347 | tags=39%, list=12%, signal=45% |
| REACTOME_HEDGEHOG_LIGAND_BIOGENESIS | 61 | 0.6500529 | 1.672229 | 0 | 0.05243329 | 0.99 | 2352 | tags=48%, list=9%, signal=52% |
| GO_NIK_NF_KAPPAB_SIGNALING | 155 | 0.58640885 | 1.6709793 | 0 | 0.05955167 | 1 | 2775 | tags=30%, list=10%, signal=34% |
| HP_MITOCHONDRIAL_RESPIRATORY_CHAIN_DEFECTS | 17 | 0.88059026 | 1.6708691 | 0 | 0.05931721 | 1 | 854 | tags=53%, list=3%, signal=55% |
| GO_PLATELET_ALPHA_GRANULE_MEMBRANE | 16 | 0.84395015 | 1.6706547 | 0 | 0.05908459 | 1 | 807 | tags=31%, list=3%, signal=32% |
| REACTOME_STABILIZATION_OF_P53 | 53 | 0.64367163 | 1.6704481 | 0 | 0.05296242 | 0.99 | 805 | tags=40%, list=3%, signal=41% |
| LEIN_OLIGODENDROCYTE_MARKERS | 72 | 0.5819333 | 1.669626 | 0 | 0.05348507 | 0.99 | 2685 | tags=33%, list=10%, signal=37% |
| HP_ABNORMALITY_OF_SECONDARY_SEXUAL_HAIR | 48 | 0.6615471 | 1.6694922 | 0.02631579 | 0.05941913 | 1 | 2301 | tags=25%, list=9%, signal=27% |
| REACTOME_DEGRADATION_OF_GLI1_BY_THE_PROTEASOME | 54 | 0.6592117 | 1.6682357 | 0 | 0.05420926 | 0.99 | 805 | tags=37%, list=3%, signal=38% |
| WANG_CLASSIC_ADIPOGENIC_TARGETS_OF_PPARG | 21 | 0.79087925 | 1.6681567 | 0.02380952 | 0.0539765 | 0.99 | 327 | tags=19%, list=1%, signal=19% |
| HP_DISTAL_PERIPHERAL_SENSORY_NEUROPATHY | 12 | 0.8528228 | 1.6679585 | 0 | 0.0611796 | 1 | 854 | tags=42%, list=3%, signal=43% |
| HUANG_GATA2_TARGETS_UP | 145 | 0.5830531 | 1.6678841 | 0 | 0.05374597 | 0.99 | 1104 | tags=14%, list=4%, signal=15% |
| HP_ABNORMAL_MYELINATION | 430 | 0.53207076 | 1.6677568 | 0 | 0.06125195 | 1 | 3129 | tags=31%, list=12%, signal=35% |
| KEGG_ALLOGRAFT_REJECTION | 21 | 0.77046937 | 1.6670322 | 0 | 0.05457448 | 0.99 | 1347 | tags=24%, list=5%, signal=25% |
| HP_ABNORMAL_CIRCULATING_CITRULLINE_CONCENTRATION | 10 | 0.9152337 | 1.6664261 | 0 | 0.06268074 | 1 | 854 | tags=60%, list=3%, signal=62% |
| GO_COLLAGEN_CONTAINING_EXTRACELLULAR_MATRIX | 348 | 0.5271876 | 1.6656712 | 0 | 0.06330073 | 1 | 3097 | tags=22%, list=12%, signal=25% |
| REACTOME_SIGNALING_BY_NOTCH4 | 79 | 0.6268987 | 1.6653708 | 0 | 0.05538011 | 1 | 805 | tags=30%, list=3%, signal=31% |
| GO_CONTRACTILE_FIBER | 206 | 0.56268024 | 1.6652641 | 0 | 0.06361171 | 1 | 2948 | tags=24%, list=11%, signal=27% |
| GO_MITOCHONDRIAL_TRANSMEMBRANE_TRANSPORT | 91 | 0.6286422 | 1.6647756 | 0 | 0.06379735 | 1 | 2410 | tags=34%, list=9%, signal=37% |
| GO_GLYCOLIPID_CATABOLIC_PROCESS | 14 | 0.8489002 | 1.6641164 | 0 | 0.0644686 | 1 | 3410 | tags=71%, list=13%, signal=82% |
| YAO_TEMPORAL_RESPONSE_TO_PROGESTERONE_CLUSTER_12 | 76 | 0.6382383 | 1.6638883 | 0 | 0.05637644 | 1 | 2363 | tags=43%, list=9%, signal=47% |
| GO_NEGATIVE_REGULATION_OF_GLYCOPROTEIN_METABOLIC_PROCESS | 14 | 0.8669253 | 1.6632864 | 0 | 0.06544004 | 1 | 1112 | tags=36%, list=4%, signal=37% |
| REACTOME_ANTIGEN_PROCESSING_CROSS_PRESENTATION | 89 | 0.64379776 | 1.6625981 | 0 | 0.05736226 | 1 | 2352 | tags=38%, list=9%, signal=42% |
| GO_REGULATION_OF_HEMATOPOIETIC_STEM_CELL_DIFFERENTIATION | 70 | 0.6582559 | 1.6625602 | 0 | 0.06561212 | 1 | 1624 | tags=34%, list=6%, signal=36% |
| GO_MULTIVESICULAR_BODY | 48 | 0.69323546 | 1.6615099 | 0 | 0.06680689 | 1 | 2698 | tags=40%, list=10%, signal=44% |
| GO_CERAMIDE_CATABOLIC_PROCESS | 17 | 0.82768136 | 1.6614751 | 0 | 0.06655668 | 1 | 3410 | tags=65%, list=13%, signal=74% |
| REACTOME_INTERLEUKIN_1_SIGNALING | 96 | 0.59374994 | 1.6608211 | 0 | 0.05946239 | 1 | 805 | tags=24%, list=3%, signal=25% |
| GRUETZMANN_PANCREATIC_CANCER_DN | 176 | 0.5552722 | 1.6601837 | 0 | 0.05971051 | 1 | 1492 | tags=23%, list=6%, signal=24% |
| HP_WIDENED_SUBARACHNOID_SPACE | 39 | 0.69861096 | 1.6592114 | 0 | 0.06840075 | 1 | 2815 | tags=33%, list=10%, signal=37% |
| KIM_HYPOXIA | 16 | 0.7697358 | 1.6591173 | 0.025 | 0.06025853 | 1 | 2939 | tags=50%, list=11%, signal=56% |
| GO_PHAGOSOME_ACIDIFICATION | 27 | 0.8120954 | 1.659116 | 0 | 0.06826627 | 1 | 1206 | tags=30%, list=4%, signal=31% |
| GO_STEROL_BINDING | 53 | 0.63433015 | 1.6590179 | 0 | 0.068073 | 1 | 3311 | tags=40%, list=12%, signal=45% |
| HP_PROGRESSIVE_EXTERNAL_OPHTHALMOPLEGIA | 24 | 0.7651314 | 1.6581416 | 0 | 0.06889051 | 1 | 1139 | tags=33%, list=4%, signal=35% |
| HP_EPILEPTIC_SPASM | 98 | 0.57240725 | 1.6580635 | 0 | 0.06863723 | 1 | 3184 | tags=32%, list=12%, signal=36% |
| HP_LACTICACIDURIA | 8 | 0.9394709 | 1.6575766 | 0 | 0.068679 | 1 | 854 | tags=75%, list=3%, signal=77% |
| ZHENG_GLIOBLASTOMA_PLASTICITY_DN | 57 | 0.6475786 | 1.6574982 | 0 | 0.06139519 | 1 | 3224 | tags=46%, list=12%, signal=52% |
| RUAN_RESPONSE_TO_TROGLITAZONE_UP | 19 | 0.8171682 | 1.6572635 | 0 | 0.06103405 | 1 | 2659 | tags=58%, list=10%, signal=64% |
| HP_HETEROGENEOUS | 248 | 0.52227765 | 1.6569958 | 0 | 0.06953607 | 1 | 2639 | tags=18%, list=10%, signal=19% |
| LEE_AGING_NEOCORTEX_UP | 70 | 0.632681 | 1.656971 | 0 | 0.06097061 | 1 | 923 | tags=27%, list=3%, signal=28% |
| BOQUEST_STEM_CELL_DN | 193 | 0.5643806 | 1.6568514 | 0 | 0.06070986 | 1 | 3085 | tags=23%, list=11%, signal=26% |
| GO_MESENCHYMAL_TO_EPITHELIAL_TRANSITION_INVOLVED_IN_METANEPHROS_MORPHOGENESIS | 11 | 0.92674774 | 1.6559453 | 0.02564103 | 0.070852 | 1 | 158 | tags=27%, list=1%, signal=27% |
| HP_NAUSEA_AND_VOMITING | 359 | 0.5283283 | 1.6556926 | 0 | 0.07088444 | 1 | 3228 | tags=32%, list=12%, signal=36% |
| HP_ELEVATED_HEPATIC_TRANSAMINASE | 202 | 0.5438391 | 1.655304 | 0 | 0.07132059 | 1 | 3528 | tags=33%, list=13%, signal=38% |
| GO_PHENOL_CONTAINING_COMPOUND_METABOLIC_PROCESS | 85 | 0.6230829 | 1.6537709 | 0 | 0.07296994 | 1 | 1634 | tags=18%, list=6%, signal=19% |
| GO_TRANSFERRIN_TRANSPORT | 35 | 0.70119786 | 1.6536008 | 0 | 0.07288103 | 1 | 1488 | tags=31%, list=6%, signal=33% |
| HP_SEVERE_GLOBAL_DEVELOPMENTAL_DELAY | 119 | 0.5796652 | 1.653194 | 0 | 0.07342247 | 1 | 2023 | tags=27%, list=8%, signal=29% |
| GO_ATP_BIOSYNTHETIC_PROCESS | 46 | 0.67173773 | 1.653149 | 0 | 0.07316118 | 1 | 1939 | tags=41%, list=7%, signal=44% |
| HP_ABNORMAL_CIRCULATING_METABOLITE_CONCENTRATION | 880 | 0.48766443 | 1.6529056 | 0 | 0.07301503 | 1 | 3537 | tags=28%, list=13%, signal=31% |
| REACTOME_CRISTAE_FORMATION | 26 | 0.74957514 | 1.6528698 | 0.04347826 | 0.06447657 | 1 | 3113 | tags=65%, list=12%, signal=74% |
| HP_ABNORMALITY_OF_THE_VASCULATURE_OF_THE_EYE | 276 | 0.54494923 | 1.651742 | 0 | 0.07445711 | 1 | 3471 | tags=27%, list=13%, signal=31% |
| GO_REGULATION_OF_PHOSPHOLIPASE_ACTIVITY | 58 | 0.6421253 | 1.6513282 | 0 | 0.07487098 | 1 | 2728 | tags=28%, list=10%, signal=31% |
| GO_POSITIVE_REGULATION_OF_VIRAL_TRANSCRIPTION | 40 | 0.66197383 | 1.6499456 | 0 | 0.07612836 | 1 | 2782 | tags=63%, list=10%, signal=70% |
| GO_CELLULAR_PROTEIN_COMPLEX_DISASSEMBLY | 204 | 0.5468478 | 1.6498438 | 0 | 0.07608613 | 1 | 2471 | tags=34%, list=9%, signal=37% |
| GO_AEROBIC_ELECTRON_TRANSPORT_CHAIN | 19 | 0.79618174 | 1.6495209 | 0 | 0.07660642 | 1 | 1812 | tags=53%, list=7%, signal=56% |
| HP_ABNORMALITY_OF_BLOOD_CIRCULATION | 244 | 0.5414129 | 1.6486367 | 0 | 0.07761543 | 1 | 3413 | tags=27%, list=13%, signal=31% |
| LIANG_SILENCED_BY_METHYLATION_UP | 23 | 0.7713021 | 1.6485469 | 0 | 0.06905466 | 1 | 411 | tags=17%, list=2%, signal=18% |
| BOSCO_EPITHELIAL_DIFFERENTIATION_MODULE | 39 | 0.63789165 | 1.6485121 | 0 | 0.06866006 | 1 | 3174 | tags=18%, list=12%, signal=20% |
| GO_IRON_ION_TRANSPORT | 66 | 0.64132464 | 1.6479 | 0 | 0.07812087 | 1 | 1488 | tags=26%, list=6%, signal=27% |
| REACTOME_DETOXIFICATION_OF_REACTIVE_OXYGEN_SPECIES | 33 | 0.7002546 | 1.647809 | 0 | 0.0693309 | 1 | 2604 | tags=48%, list=10%, signal=54% |
| GO_NEGATIVE_REGULATION_OF_WNT_SIGNALING_PATHWAY | 195 | 0.56883174 | 1.6473929 | 0 | 0.07834795 | 1 | 2270 | tags=26%, list=8%, signal=28% |
| KAAB_HEART_ATRIUM_VS_VENTRICLE_UP | 233 | 0.54743433 | 1.6473672 | 0 | 0.06922514 | 1 | 3424 | tags=33%, list=13%, signal=38% |
| HP_HYPOKINESIA | 36 | 0.7125634 | 1.6467215 | 0.02439024 | 0.07857471 | 1 | 2158 | tags=28%, list=8%, signal=30% |
| YU_MYC_TARGETS_DN | 51 | 0.6448866 | 1.6460595 | 0 | 0.06998505 | 1 | 2915 | tags=33%, list=11%, signal=37% |
| LU_EZH2_TARGETS_UP | 234 | 0.5613961 | 1.6457744 | 0 | 0.06978358 | 1 | 3404 | tags=36%, list=13%, signal=41% |
| GO_CATENIN_COMPLEX | 25 | 0.7436012 | 1.645657 | 0.02439024 | 0.07978512 | 1 | 3105 | tags=28%, list=12%, signal=32% |
| GO_HEXOSAMINIDASE_ACTIVITY | 10 | 0.87271863 | 1.6446507 | 0 | 0.0807155 | 1 | 3080 | tags=60%, list=11%, signal=68% |
| GO_MONOVALENT_INORGANIC_CATION_TRANSPORT | 459 | 0.5133163 | 1.644634 | 0 | 0.08049613 | 1 | 2442 | tags=17%, list=9%, signal=18% |
| HP_WEAK_CRY | 35 | 0.7115954 | 1.6442735 | 0 | 0.08066016 | 1 | 2421 | tags=23%, list=9%, signal=25% |
| GO_RENAL_VESICLE_DEVELOPMENT | 17 | 0.7886228 | 1.6441748 | 0 | 0.08044165 | 1 | 267 | tags=24%, list=1%, signal=24% |
| SWEET_LUNG_CANCER_KRAS_DN | 359 | 0.53093606 | 1.6438948 | 0 | 0.07136913 | 1 | 2458 | tags=22%, list=9%, signal=24% |
| REACTOME_INCRETIN_SYNTHESIS_SECRETION_AND_INACTIVATION | 14 | 0.83462703 | 1.6438675 | 0.02439024 | 0.07106689 | 1 | 351 | tags=21%, list=1%, signal=22% |
| GO_CRISTAE_FORMATION | 31 | 0.71150887 | 1.6437215 | 0 | 0.08060131 | 1 | 3671 | tags=58%, list=14%, signal=67% |
| HP_LEBER_OPTIC_ATROPHY | 10 | 0.9546889 | 1.6431755 | 0 | 0.08103102 | 1 | 854 | tags=70%, list=3%, signal=72% |
| GO_GOLGI_ASSOCIATED_VESICLE | 155 | 0.57571816 | 1.6427962 | 0 | 0.08135021 | 1 | 2180 | tags=32%, list=8%, signal=35% |
| ALONSO_METASTASIS_UP | 177 | 0.57107264 | 1.6425271 | 0 | 0.07170499 | 1 | 2655 | tags=39%, list=10%, signal=43% |
| GO_REGULATION_OF_RESPONSE_TO_INTERFERON_GAMMA | 23 | 0.80071807 | 1.6423113 | 0 | 0.08139928 | 1 | 2468 | tags=43%, list=9%, signal=48% |
| GO_COPPER_ION_BINDING | 50 | 0.62716395 | 1.6421635 | 0 | 0.08118153 | 1 | 839 | tags=18%, list=3%, signal=19% |
| KEGG_TAURINE_AND_HYPOTAURINE_METABOLISM | 8 | 0.9355428 | 1.6414466 | 0 | 0.07214043 | 1 | 384 | tags=25%, list=1%, signal=25% |
| WP_PATHOGENIC_ESCHERICHIA_COLI_INFECTION | 49 | 0.62041616 | 1.6410513 | 0 | 0.07221574 | 1 | 2819 | tags=41%, list=10%, signal=46% |
| TOMLINS_PROSTATE_CANCER_DN | 35 | 0.668919 | 1.6409198 | 0 | 0.07209609 | 1 | 2678 | tags=40%, list=10%, signal=44% |
| KIM_BIPOLAR_DISORDER_OLIGODENDROCYTE_DENSITY_CORR_UP | 646 | 0.4965766 | 1.640897 | 0 | 0.07179924 | 1 | 2447 | tags=35%, list=9%, signal=38% |
| RHEIN_ALL_GLUCOCORTICOID_THERAPY_DN | 351 | 0.5098637 | 1.6407924 | 0 | 0.07168737 | 1 | 2587 | tags=35%, list=10%, signal=38% |
| HP_VERTIGO | 89 | 0.6064128 | 1.6403939 | 0 | 0.0834595 | 1 | 3347 | tags=33%, list=12%, signal=37% |
| SANSOM_WNT_PATHWAY_REQUIRE_MYC | 58 | 0.6836152 | 1.6400671 | 0 | 0.07166835 | 1 | 1671 | tags=28%, list=6%, signal=29% |
| GO_NEGATIVE_REGULATION_OF_ACTIVATED_T_CELL_PROLIFERATION | 12 | 0.78255147 | 1.6392282 | 0.025 | 0.08471321 | 1 | 2158 | tags=33%, list=8%, signal=36% |
| HILLION_HMGA1_TARGETS | 75 | 0.619668 | 1.6389776 | 0 | 0.07244777 | 1 | 1939 | tags=39%, list=7%, signal=42% |
| HP_LOW_PLASMA_CITRULLINE | 7 | 0.9461146 | 1.6389397 | 0 | 0.08501667 | 1 | 854 | tags=86%, list=3%, signal=88% |
| GO_PLATELET_ALPHA_GRANULE | 74 | 0.61507094 | 1.6385286 | 0 | 0.08526228 | 1 | 1194 | tags=19%, list=4%, signal=20% |
| HP_POLAR_CATARACT | 9 | 0.91043454 | 1.6382529 | 0 | 0.08529908 | 1 | 104 | tags=33%, list=0%, signal=33% |
| FAELT_B_CLL_WITH_VH3_21_DN | 42 | 0.6607116 | 1.6380599 | 0 | 0.07313859 | 1 | 2714 | tags=45%, list=10%, signal=50% |
| HP_PULMONARY_ARTERIAL_HYPERTENSION | 94 | 0.6195832 | 1.6376411 | 0 | 0.0860658 | 1 | 3866 | tags=34%, list=14%, signal=40% |
| GO_PHENOL_CONTAINING_COMPOUND_BIOSYNTHETIC_PROCESS | 37 | 0.6929576 | 1.6375122 | 0.02777778 | 0.08599311 | 1 | 1194 | tags=22%, list=4%, signal=23% |
| GO_POSITIVE_REGULATION_OF_CANONICAL_WNT_SIGNALING_PATHWAY | 141 | 0.55949754 | 1.6374056 | 0 | 0.08586964 | 1 | 2717 | tags=28%, list=10%, signal=31% |
| GO_PHAGOCYTIC_VESICLE | 129 | 0.58142465 | 1.637255 | 0 | 0.08574677 | 1 | 2512 | tags=23%, list=9%, signal=26% |
| GO_POSITIVE_REGULATION_OF_KERATINOCYTE_PROLIFERATION | 9 | 0.9170329 | 1.6370382 | 0 | 0.0858837 | 1 | 366 | tags=22%, list=1%, signal=23% |
| EBAUER_MYOGENIC_TARGETS_OF_PAX3_FOXO1_FUSION | 43 | 0.6990822 | 1.6368352 | 0 | 0.07399521 | 1 | 1170 | tags=19%, list=4%, signal=19% |
| GO_NEGATIVE_REGULATION_OF_EPITHELIAL_CELL_PROLIFERATION | 115 | 0.59805673 | 1.6368132 | 0 | 0.08612296 | 1 | 1663 | tags=17%, list=6%, signal=18% |
| YAO_HOXA10_TARGETS_VIA_PROGESTERONE_UP | 67 | 0.6431969 | 1.6367815 | 0 | 0.07360981 | 1 | 2344 | tags=28%, list=9%, signal=31% |
| HP_FUNCTIONAL_RESPIRATORY_ABNORMALITY | 1023 | 0.47160655 | 1.6357788 | 0 | 0.08728381 | 1 | 3354 | tags=26%, list=12%, signal=28% |
| PLASARI_TGFB1_SIGNALING_VIA_NFIC_10HR_UP | 49 | 0.6569644 | 1.6351206 | 0 | 0.07471637 | 1 | 2157 | tags=27%, list=8%, signal=29% |
| GO_SECONDARY_METABOLITE_BIOSYNTHETIC_PROCESS | 20 | 0.75422144 | 1.6346235 | 0 | 0.08807886 | 1 | 1067 | tags=30%, list=4%, signal=31% |
| REACTOME_REGULATION_OF_IFNG_SIGNALING | 13 | 0.8084188 | 1.6341729 | 0.02564103 | 0.07503006 | 1 | 2232 | tags=54%, list=8%, signal=59% |
| HELLER_SILENCED_BY_METHYLATION_UP | 229 | 0.56510425 | 1.6336619 | 0 | 0.07551639 | 1 | 2173 | tags=21%, list=8%, signal=23% |
| HP_PERIPHERAL_NEUROPATHY | 515 | 0.48829708 | 1.6331612 | 0 | 0.0894732 | 1 | 3150 | tags=26%, list=12%, signal=28% |
| LIU_PROSTATE_CANCER_DN | 427 | 0.52003336 | 1.633108 | 0 | 0.07564496 | 1 | 2434 | tags=22%, list=9%, signal=23% |
| HOLLERN_MICROACINAR_BREAST_TUMOR_UP | 40 | 0.6529165 | 1.6330715 | 0 | 0.07526097 | 1 | 2704 | tags=38%, list=10%, signal=42% |
| GO_POST_TRANSLATIONAL_PROTEIN_MODIFICATION | 327 | 0.52664036 | 1.6328908 | 0 | 0.08964503 | 1 | 2857 | tags=27%, list=11%, signal=30% |
| WONG_ADULT_TISSUE_STEM_MODULE | 672 | 0.5041111 | 1.6325266 | 0 | 0.07513522 | 1 | 3205 | tags=31%, list=12%, signal=34% |
| HUPER_BREAST_BASAL_VS_LUMINAL_DN | 43 | 0.6888402 | 1.6319 | 0.04081633 | 0.07551937 | 1 | 1714 | tags=26%, list=6%, signal=27% |
| HP_ABNORMAL_RENAL_CORTEX_MORPHOLOGY | 156 | 0.5666778 | 1.6316968 | 0 | 0.09082509 | 1 | 3471 | tags=30%, list=13%, signal=34% |
| REACTOME_INTERLEUKIN_20_FAMILY_SIGNALING | 15 | 0.8074509 | 1.6314702 | 0.02439024 | 0.07522614 | 1 | 1939 | tags=40%, list=7%, signal=43% |
| HP_LEUKOENCEPHALOPATHY | 156 | 0.58373183 | 1.6313583 | 0 | 0.0909439 | 1 | 2622 | tags=36%, list=10%, signal=40% |
| GO_ADULT_FEEDING_BEHAVIOR | 6 | 0.9162999 | 1.6309826 | 0 | 0.0913648 | 1 | 790 | tags=33%, list=3%, signal=34% |
| REACTOME_PYRUVATE_METABOLISM_AND_CITRIC_ACID_TCA_CYCLE | 52 | 0.675268 | 1.6301931 | 0.02777778 | 0.07610659 | 1 | 2982 | tags=38%, list=11%, signal=43% |
| GO_EMBRYONIC_SKELETAL_JOINT_DEVELOPMENT | 9 | 0.895695 | 1.6301743 | 0 | 0.09182773 | 1 | 627 | tags=22%, list=2%, signal=23% |
| HAMAI_APOPTOSIS_VIA_TRAIL_UP | 611 | 0.49760163 | 1.6299893 | 0 | 0.07614097 | 1 | 3254 | tags=36%, list=12%, signal=40% |
| MORI_PLASMA_CELL_UP | 47 | 0.65013283 | 1.6291832 | 0 | 0.07734992 | 1 | 2593 | tags=51%, list=10%, signal=56% |
| WP_PPAR_SIGNALING_PATHWAY | 53 | 0.6765203 | 1.6285416 | 0 | 0.07805221 | 1 | 3030 | tags=26%, list=11%, signal=30% |
| REACTOME_FORMATION_OF_ATP_BY_CHEMIOSMOTIC_COUPLING | 15 | 0.87099886 | 1.627996 | 0 | 0.07833248 | 1 | 1251 | tags=67%, list=5%, signal=70% |
| CHARAFE_BREAST_CANCER_LUMINAL_VS_BASAL_DN | 391 | 0.5006885 | 1.6274612 | 0 | 0.07836329 | 1 | 3240 | tags=28%, list=12%, signal=31% |
| HP_ABNORMAL_LEFT_VENTRICLE_MORPHOLOGY | 85 | 0.6082225 | 1.6274474 | 0 | 0.09533573 | 1 | 1994 | tags=19%, list=7%, signal=20% |
| PAPASPYRIDONOS_UNSTABLE_ATEROSCLEROTIC_PLAQUE_UP | 51 | 0.6584252 | 1.6271477 | 0 | 0.07814701 | 1 | 2115 | tags=33%, list=8%, signal=36% |
| GOBERT_OLIGODENDROCYTE_DIFFERENTIATION_DN | 1020 | 0.46848676 | 1.6269306 | 0 | 0.07818183 | 1 | 3821 | tags=36%, list=14%, signal=41% |
| GO_DNA_REPLICATION_FACTOR_A_COMPLEX | 13 | 0.80645895 | 1.626521 | 0 | 0.09623417 | 1 | 1715 | tags=62%, list=6%, signal=66% |
| GO_COATED_VESICLE | 259 | 0.5257397 | 1.6264421 | 0 | 0.09593623 | 1 | 2425 | tags=29%, list=9%, signal=32% |
| REACTOME_DOWNSTREAM_SIGNALING_EVENTS_OF_B_CELL_RECEPTOR_BCR_ | 77 | 0.6044863 | 1.6262956 | 0 | 0.07861733 | 1 | 805 | tags=29%, list=3%, signal=29% |
| GRADE_COLON_AND_RECTAL_CANCER_DN | 93 | 0.5923275 | 1.6257174 | 0 | 0.07904493 | 1 | 3710 | tags=33%, list=14%, signal=39% |
| ABBUD_LIF_SIGNALING_1_UP | 35 | 0.70386434 | 1.6257098 | 0 | 0.07867031 | 1 | 2204 | tags=34%, list=8%, signal=37% |
| WESTON_VEGFA_TARGETS_12HR | 21 | 0.7242106 | 1.6256114 | 0.02777778 | 0.07853611 | 1 | 1133 | tags=19%, list=4%, signal=20% |
| GO_ORGANELLE_ENVELOPE_LUMEN | 84 | 0.603481 | 1.6252633 | 0 | 0.09716979 | 1 | 3173 | tags=43%, list=12%, signal=48% |
| RAY_TUMORIGENESIS_BY_ERBB2_CDC25A_DN | 244 | 0.53990155 | 1.6251668 | 0 | 0.07864492 | 1 | 3073 | tags=27%, list=11%, signal=30% |
| SANSOM_APC_TARGETS_DN | 312 | 0.53648305 | 1.6247433 | 0 | 0.07874887 | 1 | 2916 | tags=22%, list=11%, signal=24% |
| GO_POSITIVE_REGULATION_OF_ALPHA_BETA_T_CELL_PROLIFERATION | 18 | 0.76106536 | 1.6243067 | 0.02173913 | 0.09800164 | 1 | 1765 | tags=22%, list=7%, signal=24% |
| HP_ABNORMAL_CRY | 44 | 0.6558976 | 1.6241564 | 0 | 0.09804559 | 1 | 2421 | tags=20%, list=9%, signal=22% |
| HP_INFLAMMATORY_ABNORMALITY_OF_THE_EYE | 159 | 0.5369042 | 1.6241266 | 0 | 0.09784394 | 1 | 2174 | tags=17%, list=8%, signal=18% |
| GO_UTERUS_DEVELOPMENT | 17 | 0.8347186 | 1.6241196 | 0 | 0.09754564 | 1 | 3413 | tags=35%, list=13%, signal=40% |
| GO_ENDOPEPTIDASE_COMPLEX | 65 | 0.6381292 | 1.6241012 | 0 | 0.09724915 | 1 | 1624 | tags=35%, list=6%, signal=38% |
| LEE_LIVER_CANCER_MYC_TGFA_DN | 53 | 0.59524804 | 1.6228296 | 0 | 0.08027372 | 1 | 4564 | tags=34%, list=17%, signal=41% |
| HP_CENTROCECAL_SCOTOMA | 11 | 0.95233715 | 1.6220906 | 0 | 0.10016212 | 1 | 854 | tags=73%, list=3%, signal=75% |
| GO_EXTRACELLULAR_MATRIX | 460 | 0.5013057 | 1.6214526 | 0 | 0.10072833 | 1 | 2752 | tags=18%, list=10%, signal=20% |
| GO_NEGATIVE_REGULATION_OF_CELL_KILLING | 17 | 0.78151476 | 1.6209475 | 0 | 0.1014343 | 1 | 2389 | tags=29%, list=9%, signal=32% |
| NABA_ECM_REGULATORS | 168 | 0.5649978 | 1.6208097 | 0 | 0.08241522 | 1 | 2686 | tags=17%, list=10%, signal=19% |
| GO_NEGATIVE_REGULATION_OF_MULTI_ORGANISM_PROCESS | 18 | 0.81933 | 1.6201451 | 0 | 0.10257389 | 1 | 474 | tags=22%, list=2%, signal=23% |
| GO_POLYOL_BIOSYNTHETIC_PROCESS | 52 | 0.6391551 | 1.6201277 | 0.02941177 | 0.10226678 | 1 | 4324 | tags=44%, list=16%, signal=53% |
| NUYTTEN_NIPP1_TARGETS_UP | 642 | 0.4892983 | 1.6193286 | 0 | 0.08360742 | 1 | 3762 | tags=34%, list=14%, signal=39% |
| GO_TERTIARY_GRANULE_LUMEN | 46 | 0.66247016 | 1.6192214 | 0 | 0.10368105 | 1 | 1348 | tags=24%, list=5%, signal=25% |
| HP_RESPIRATORY_DISTRESS | 183 | 0.54790676 | 1.6190214 | 0 | 0.10361035 | 1 | 3049 | tags=25%, list=11%, signal=28% |
| GO_PROTON_TRANSPORTING_ATP_SYNTHASE_COMPLEX_COUPLING_FACTOR_F_O | 10 | 0.8439158 | 1.6189266 | 0.04347826 | 0.10349334 | 1 | 822 | tags=50%, list=3%, signal=52% |
| GO_REGULATION_OF_PEPTIDASE_ACTIVITY | 337 | 0.51536465 | 1.618688 | 0 | 0.10351826 | 1 | 2874 | tags=24%, list=11%, signal=26% |
| YANG_BCL3_TARGETS_UP | 314 | 0.5241581 | 1.618292 | 0 | 0.08423075 | 1 | 2688 | tags=26%, list=10%, signal=28% |
| BOQUEST_STEM_CELL_CULTURED_VS_FRESH_DN | 22 | 0.76511717 | 1.6180571 | 0 | 0.08392622 | 1 | 2090 | tags=50%, list=8%, signal=54% |
| REACTOME_HEDGEHOG_ON_STATE | 81 | 0.60297686 | 1.6179826 | 0 | 0.08362275 | 1 | 2352 | tags=36%, list=9%, signal=39% |
| GO_INTRINSIC_COMPONENT_OF_MITOCHONDRIAL_INNER_MEMBRANE | 42 | 0.6625223 | 1.6173798 | 0 | 0.10465364 | 1 | 3709 | tags=45%, list=14%, signal=52% |
| HP_MUSCLE_WEAKNESS | 782 | 0.48621994 | 1.61719 | 0 | 0.10467466 | 1 | 3537 | tags=30%, list=13%, signal=33% |
| GO_RESPONSE_TO_INTERFERON_BETA | 24 | 0.76402295 | 1.6171513 | 0 | 0.10441566 | 1 | 2374 | tags=46%, list=9%, signal=50% |
| WALLACE_PROSTATE_CANCER_RACE_UP | 246 | 0.5141141 | 1.6170844 | 0 | 0.08408041 | 1 | 2144 | tags=17%, list=8%, signal=19% |
| GO_PEPTIDYL_ASPARAGINE_MODIFICATION | 28 | 0.70863694 | 1.6169604 | 0 | 0.10443848 | 1 | 3715 | tags=57%, list=14%, signal=66% |
| WP_GLYCOLYSIS_AND_GLUCONEOGENESIS | 41 | 0.6810143 | 1.6160231 | 0 | 0.08438584 | 1 | 3141 | tags=39%, list=12%, signal=44% |
| DELPUECH_FOXO3_TARGETS_UP | 59 | 0.6456993 | 1.6158987 | 0 | 0.08408496 | 1 | 827 | tags=15%, list=3%, signal=16% |
| GAUSSMANN_MLL_AF4_FUSION_TARGETS_F_UP | 171 | 0.54633164 | 1.6154542 | 0 | 0.08423399 | 1 | 2983 | tags=26%, list=11%, signal=29% |
| HP_DEMYELINATING_PERIPHERAL_NEUROPATHY | 36 | 0.7214745 | 1.615241 | 0 | 0.10641731 | 1 | 1529 | tags=31%, list=6%, signal=32% |
| HP_FEVER | 221 | 0.54073733 | 1.6148938 | 0 | 0.10657064 | 1 | 2915 | tags=24%, list=11%, signal=26% |
| HP_CARDIAC_CONDUCTION_ABNORMALITY | 99 | 0.6299356 | 1.6140573 | 0 | 0.10741638 | 1 | 1736 | tags=24%, list=6%, signal=26% |
| GO_MHC_CLASS_II_PROTEIN_COMPLEX | 7 | 0.90011996 | 1.6140049 | 0.02941177 | 0.10719891 | 1 | 1347 | tags=57%, list=5%, signal=60% |
| GAUSSMANN_MLL_AF4_FUSION_TARGETS_E_UP | 92 | 0.61467874 | 1.6138151 | 0 | 0.08551655 | 1 | 2993 | tags=27%, list=11%, signal=30% |
| HP_ABNORMALITY_OF_THE_OPTIC_DISC | 558 | 0.49911392 | 1.613542 | 0 | 0.1074452 | 1 | 3524 | tags=30%, list=13%, signal=34% |
| GO_EXTERNAL_SIDE_OF_PLASMA_MEMBRANE | 272 | 0.5200359 | 1.6132479 | 0 | 0.10764226 | 1 | 3703 | tags=18%, list=14%, signal=21% |
| NATSUME_RESPONSE_TO_INTERFERON_BETA_DN | 45 | 0.63965887 | 1.6131921 | 0 | 0.08580979 | 1 | 1096 | tags=36%, list=4%, signal=37% |
| GO_T_CELL_RECEPTOR_SIGNALING_PATHWAY | 170 | 0.55788386 | 1.6129043 | 0 | 0.10762287 | 1 | 2510 | tags=26%, list=9%, signal=29% |
| HP_ABNORMALITY_OF_THE_PUBIC_HAIR | 35 | 0.7231331 | 1.612732 | 0 | 0.10759228 | 1 | 2301 | tags=26%, list=9%, signal=28% |
| GO_POSITIVE_REGULATION_OF_ENDOTHELIAL_CELL_DIFFERENTIATION | 15 | 0.8160371 | 1.6126492 | 0 | 0.10737784 | 1 | 1428 | tags=27%, list=5%, signal=28% |
| HP_VASCULAR_SKIN_ABNORMALITY | 419 | 0.5048237 | 1.6113497 | 0 | 0.1094314 | 1 | 3997 | tags=29%, list=15%, signal=34% |
| SABATES_COLORECTAL_ADENOMA_DN | 226 | 0.5493802 | 1.6105623 | 0 | 0.08840919 | 1 | 4149 | tags=21%, list=15%, signal=25% |
| SCHAEFFER_PROSTATE_DEVELOPMENT_48HR_UP | 384 | 0.50813276 | 1.6104597 | 0 | 0.08809487 | 1 | 3158 | tags=25%, list=12%, signal=28% |
| HP_PSYCHOTIC_MENTATION | 9 | 0.919687 | 1.6103247 | 0 | 0.11092138 | 1 | 854 | tags=56%, list=3%, signal=57% |
| HP_CEREBRAL_HEMORRHAGE | 53 | 0.6271408 | 1.6102486 | 0 | 0.11065572 | 1 | 2400 | tags=28%, list=9%, signal=31% |
| WP_NOCGMPPKG_MEDIATED_NEUROPROTECTION | 40 | 0.6440341 | 1.6101612 | 0 | 0.08822703 | 1 | 1812 | tags=23%, list=7%, signal=24% |
| GO_CHOLESTEROL_BINDING | 44 | 0.6811849 | 1.6096516 | 0 | 0.11115176 | 1 | 2434 | tags=34%, list=9%, signal=37% |
| GO_FC_RECEPTOR_SIGNALING_PATHWAY | 172 | 0.55550814 | 1.609512 | 0 | 0.11115348 | 1 | 2478 | tags=29%, list=9%, signal=32% |
| NIKOLSKY_BREAST_CANCER_8Q12_Q22_AMPLICON | 115 | 0.56575525 | 1.609109 | 0 | 0.0889413 | 1 | 2980 | tags=25%, list=11%, signal=28% |
| LIANG_HEMATOPOIESIS_STEM_CELL_NUMBER_LARGE_VS_TINY_UP | 39 | 0.69531286 | 1.6087877 | 0.02857143 | 0.08877542 | 1 | 2076 | tags=36%, list=8%, signal=39% |
| HP_MULTIPLE_GLOMERULAR_CYSTS | 8 | 0.9436655 | 1.6079412 | 0 | 0.11285256 | 1 | 854 | tags=75%, list=3%, signal=77% |
| HP_ABNORMAL_SUBARACHNOID_SPACE_MORPHOLOGY | 70 | 0.6366873 | 1.6076767 | 0 | 0.11334018 | 1 | 2815 | tags=23%, list=10%, signal=25% |
| HP_TACHYCARDIA | 85 | 0.6137691 | 1.607462 | 0 | 0.11329347 | 1 | 1628 | tags=19%, list=6%, signal=20% |
| HP_FACIAL_DIPLEGIA | 33 | 0.68565667 | 1.6074015 | 0 | 0.11311325 | 1 | 2009 | tags=27%, list=7%, signal=29% |
| LIM_MAMMARY_STEM_CELL_UP | 448 | 0.4783278 | 1.6067419 | 0 | 0.09173244 | 1 | 2318 | tags=18%, list=9%, signal=19% |
| HP_ABNORMAL_SOCIAL_BEHAVIOR | 145 | 0.5514763 | 1.6064739 | 0 | 0.11407726 | 1 | 2360 | tags=30%, list=9%, signal=32% |
| GO_LONG_CHAIN_FATTY_ACID_BIOSYNTHETIC_PROCESS | 25 | 0.75483906 | 1.6062301 | 0 | 0.11393973 | 1 | 1267 | tags=24%, list=5%, signal=25% |
| HP_BLINDNESS | 233 | 0.5424012 | 1.6060485 | 0 | 0.11411084 | 1 | 2500 | tags=24%, list=9%, signal=26% |
| HP_DYSPHAGIA | 317 | 0.5090407 | 1.6059281 | 0 | 0.11388634 | 1 | 3152 | tags=27%, list=12%, signal=31% |
| GO_METALLOEXOPEPTIDASE_ACTIVITY | 48 | 0.63566273 | 1.6058638 | 0 | 0.11362053 | 1 | 3898 | tags=38%, list=14%, signal=44% |
| HP_OSTEOLYSIS_INVOLVING_BONES_OF_THE_UPPER_LIMBS | 17 | 0.7616348 | 1.6057503 | 0 | 0.11348604 | 1 | 2030 | tags=41%, list=8%, signal=45% |
| VANDESLUIS_COMMD1_TARGETS_GROUP_3_DN | 32 | 0.72776467 | 1.6057042 | 0 | 0.09278572 | 1 | 2450 | tags=41%, list=9%, signal=45% |
| REACTOME_REGULATION_OF_RAS_BY_GAPS | 65 | 0.622278 | 1.6056094 | 0 | 0.09261158 | 1 | 2352 | tags=45%, list=9%, signal=49% |
| HP_CEREBRAL_CALCIFICATION | 106 | 0.6116172 | 1.6055864 | 0 | 0.11347964 | 1 | 2068 | tags=27%, list=8%, signal=30% |
| GO_CELL_DIFFERENTIATION_INVOLVED_IN_METANEPHROS_DEVELOPMENT | 21 | 0.7573819 | 1.6054394 | 0.02631579 | 0.11334648 | 1 | 1528 | tags=24%, list=6%, signal=25% |
| LEE_CALORIE_RESTRICTION_MUSCLE_UP | 36 | 0.7105818 | 1.6053951 | 0 | 0.09257891 | 1 | 1406 | tags=28%, list=5%, signal=29% |
| GO_PLATELET_DENSE_GRANULE | 16 | 0.8511054 | 1.6053693 | 0 | 0.11312749 | 1 | 2870 | tags=50%, list=11%, signal=56% |
| PEDERSEN_METASTASIS_BY_ERBB2_ISOFORM_4 | 90 | 0.576013 | 1.6053337 | 0 | 0.09233043 | 1 | 3308 | tags=28%, list=12%, signal=32% |
| HECKER_IFNB1_TARGETS | 75 | 0.5953726 | 1.6053118 | 0 | 0.09194248 | 1 | 1599 | tags=32%, list=6%, signal=34% |
| SMID_BREAST_CANCER_RELAPSE_IN_BONE_DN | 256 | 0.5447655 | 1.6052153 | 0 | 0.09176938 | 1 | 2394 | tags=22%, list=9%, signal=24% |
| GO_REGULATION_OF_ANIMAL_ORGAN_MORPHOGENESIS | 229 | 0.52628505 | 1.6051239 | 0 | 0.11320996 | 1 | 2270 | tags=21%, list=8%, signal=23% |
| BEIER_GLIOMA_STEM_CELL_DN | 61 | 0.64317876 | 1.6049594 | 0 | 0.09194942 | 1 | 3211 | tags=38%, list=12%, signal=43% |
| HP_WORMIAN_BONES | 42 | 0.64057845 | 1.6047254 | 0 | 0.11371969 | 1 | 4288 | tags=43%, list=16%, signal=51% |
| GO_SITE_OF_DNA_DAMAGE | 71 | 0.575688 | 1.6046629 | 0.02564103 | 0.11345862 | 1 | 3457 | tags=41%, list=13%, signal=47% |
| HP_ABNORMALITY_OF_HEPATOBILIARY_SYSTEM_PHYSIOLOGY | 177 | 0.55919915 | 1.6046457 | 0 | 0.11315686 | 1 | 3528 | tags=37%, list=13%, signal=43% |
| NABA_MATRISOME | 736 | 0.47561225 | 1.60464 | 0 | 0.09184898 | 1 | 3097 | tags=17%, list=12%, signal=19% |
| GO_POSITIVE_REGULATION_OF_INNATE_IMMUNE_RESPONSE | 182 | 0.5362319 | 1.6044322 | 0 | 0.11306756 | 1 | 1624 | tags=18%, list=6%, signal=19% |
| BROWNE_HCMV_INFECTION_24HR_DN | 130 | 0.57040757 | 1.6041069 | 0 | 0.09202904 | 1 | 2945 | tags=29%, list=11%, signal=33% |
| HP_GLUCOSE_INTOLERANCE | 337 | 0.5127571 | 1.6040298 | 0 | 0.11336235 | 1 | 2329 | tags=21%, list=9%, signal=22% |
| GO_CARBOHYDRATE_DERIVATIVE_CATABOLIC_PROCESS | 161 | 0.559924 | 1.6038166 | 0 | 0.11340053 | 1 | 3454 | tags=34%, list=13%, signal=39% |
| GO_EXTRACELLULAR_MATRIX_DISASSEMBLY | 65 | 0.6132942 | 1.6037364 | 0 | 0.11335601 | 1 | 875 | tags=15%, list=3%, signal=16% |
| REACTOME_RNA_POLYMERASE_III_TRANSCRIPTION_TERMINATION | 22 | 0.7754065 | 1.6037351 | 0.02325581 | 0.0921384 | 1 | 2452 | tags=55%, list=9%, signal=60% |
| GO_PROTEASOME_CORE_COMPLEX_BETA_SUBUNIT_COMPLEX | 9 | 0.93841964 | 1.6035028 | 0 | 0.11339532 | 1 | 597 | tags=78%, list=2%, signal=80% |
| REACTOME_REGULATION_OF_IFNA_SIGNALING | 12 | 0.8854017 | 1.6032598 | 0.02439024 | 0.09238143 | 1 | 2378 | tags=75%, list=9%, signal=82% |
| OUELLET_OVARIAN_CANCER_INVASIVE_VS_LMP_UP | 111 | 0.55798745 | 1.60083 | 0 | 0.09476733 | 1 | 1848 | tags=35%, list=7%, signal=38% |
| GO_WATER_TRANSMEMBRANE_TRANSPORTER_ACTIVITY | 9 | 0.8484848 | 1.6008066 | 0.02631579 | 0.11754464 | 1 | 2751 | tags=22%, list=10%, signal=25% |
| GO_REGULATION_OF_DEVELOPMENT_HETEROCHRONIC | 13 | 0.84491193 | 1.6004932 | 0 | 0.11778156 | 1 | 2315 | tags=46%, list=9%, signal=50% |
| MARSON_FOXP3_TARGETS_UP | 64 | 0.63139844 | 1.6002783 | 0 | 0.09479745 | 1 | 3091 | tags=41%, list=11%, signal=46% |
| GO_METALLOENDOPEPTIDASE_INHIBITOR_ACTIVITY | 15 | 0.88097715 | 1.5999568 | 0 | 0.11855608 | 1 | 1774 | tags=33%, list=7%, signal=36% |
| GO_LONG_CHAIN_FATTY_ACID_METABOLIC_PROCESS | 85 | 0.614213 | 1.5998076 | 0 | 0.11870467 | 1 | 1854 | tags=21%, list=7%, signal=23% |
| REACTOME_THE_ROLE_OF_GTSE1_IN_G2_M_PROGRESSION_AFTER_G2_CHECKPOINT | 70 | 0.6429806 | 1.5996151 | 0 | 0.09502661 | 1 | 805 | tags=34%, list=3%, signal=35% |
| WP_PROTEASOME_DEGRADATION | 54 | 0.6576195 | 1.5992126 | 0 | 0.09505387 | 1 | 2352 | tags=52%, list=9%, signal=57% |
| BURTON_ADIPOGENESIS_5 | 106 | 0.5845039 | 1.5991926 | 0 | 0.09467212 | 1 | 2744 | tags=42%, list=10%, signal=46% |
| GO_STEROID_BINDING | 89 | 0.5818964 | 1.5991359 | 0 | 0.11943156 | 1 | 3311 | tags=29%, list=12%, signal=33% |
| GO_NEGATIVE_REGULATION_OF_PROTEOLYSIS | 255 | 0.51814604 | 1.5985619 | 0 | 0.11999492 | 1 | 2879 | tags=24%, list=11%, signal=26% |
| GO_MONOVALENT_INORGANIC_CATION_TRANSMEMBRANE_TRANSPORTER_ACTIVITY | 314 | 0.50622797 | 1.5984483 | 0 | 0.11976894 | 1 | 1740 | tags=13%, list=6%, signal=14% |
| GO_REGULATION_OF_HEMOPOIESIS | 412 | 0.49636424 | 1.597534 | 0 | 0.12128116 | 1 | 2604 | tags=22%, list=10%, signal=24% |
| GO_POSITIVE_REGULATION_OF_T_CELL_APOPTOTIC_PROCESS | 10 | 0.9304696 | 1.5972415 | 0 | 0.12142202 | 1 | 673 | tags=40%, list=3%, signal=41% |
| WIELAND_UP_BY_HBV_INFECTION | 85 | 0.60305476 | 1.5970798 | 0 | 0.0972034 | 1 | 2146 | tags=28%, list=8%, signal=31% |
| GO_NEGATIVE_REGULATION_OF_PEPTIDASE_ACTIVITY | 170 | 0.5625943 | 1.5969884 | 0 | 0.12144041 | 1 | 1824 | tags=18%, list=7%, signal=19% |
| WONG_PROTEASOME_GENE_MODULE | 48 | 0.65164375 | 1.596604 | 0.02564103 | 0.09703761 | 1 | 1811 | tags=42%, list=7%, signal=45% |
| KEGG_GLYCOSPHINGOLIPID_BIOSYNTHESIS_GLOBO_SERIES | 11 | 0.8487163 | 1.596301 | 0 | 0.0969876 | 1 | 2998 | tags=36%, list=11%, signal=41% |
| GO_MAINTENANCE_OF_GASTROINTESTINAL_EPITHELIUM | 14 | 0.8095269 | 1.595948 | 0 | 0.12296937 | 1 | 267 | tags=7%, list=1%, signal=7% |
| CERVERA_SDHB_TARGETS_2 | 99 | 0.5653238 | 1.5957977 | 0 | 0.09713761 | 1 | 1964 | tags=20%, list=7%, signal=22% |
| CHESLER_BRAIN_QTL_CIS | 66 | 0.60862684 | 1.5956235 | 0 | 0.09728867 | 1 | 2789 | tags=41%, list=10%, signal=46% |
| HP_ABNORMAL_CONJUNCTIVA_MORPHOLOGY | 116 | 0.5824507 | 1.5955881 | 0 | 0.12306486 | 1 | 1994 | tags=21%, list=7%, signal=22% |
| REACTOME_SIGNALING_BY_THE_B_CELL_RECEPTOR_BCR_ | 106 | 0.57367283 | 1.595587 | 0 | 0.09690864 | 1 | 2389 | tags=31%, list=9%, signal=34% |
| REACTOME_FCERI_MEDIATED_NF_KB_ACTIVATION | 76 | 0.5945598 | 1.5952243 | 0 | 0.09719487 | 1 | 805 | tags=26%, list=3%, signal=27% |
| GO_RESPONSE_TO_INTERFERON_GAMMA | 162 | 0.53279656 | 1.5951182 | 0 | 0.12401212 | 1 | 1909 | tags=22%, list=7%, signal=24% |
| WP_DEGRADATION_PATHWAY_OF_SPHINGOLIPIDS_INCLUDING_DISEASES | 8 | 0.8658722 | 1.5949736 | 0.04347826 | 0.09746928 | 1 | 2998 | tags=63%, list=11%, signal=70% |
| WINZEN_DEGRADED_VIA_KHSRP | 88 | 0.59849775 | 1.5949321 | 0 | 0.09715745 | 1 | 1115 | tags=16%, list=4%, signal=17% |
| HP_ABNORMAL_URINE_POTASSIUM_CONCENTRATION | 6 | 0.94168514 | 1.5948104 | 0 | 0.12418295 | 1 | 161 | tags=17%, list=1%, signal=17% |
| GO_ANTIGEN_RECEPTOR_MEDIATED_SIGNALING_PATHWAY | 214 | 0.5236443 | 1.5948037 | 0 | 0.12390941 | 1 | 2587 | tags=24%, list=10%, signal=27% |
| GO_SECONDARY_METABOLIC_PROCESS | 43 | 0.6782009 | 1.5946579 | 0.02941177 | 0.12383906 | 1 | 2003 | tags=28%, list=7%, signal=30% |
| GO_FLUID_TRANSPORT | 21 | 0.76504564 | 1.5945253 | 0.02439024 | 0.12368862 | 1 | 2751 | tags=29%, list=10%, signal=32% |
| HP_APLASIA_HYPOPLASIA_OF_THE_CEREBRAL_WHITE_MATTER | 15 | 0.81708944 | 1.5944961 | 0.04878049 | 0.12341904 | 1 | 854 | tags=33%, list=3%, signal=34% |
| HP_LIPODYSTROPHY | 96 | 0.5607202 | 1.5944508 | 0 | 0.12322962 | 1 | 4375 | tags=39%, list=16%, signal=46% |
| GO_EPITHELIAL_CELL_PROLIFERATION | 350 | 0.48980594 | 1.594065 | 0 | 0.12344205 | 1 | 1694 | tags=15%, list=6%, signal=16% |
| GO_NEGATIVE_REGULATION_OF_CELL_CYCLE_G2_M_PHASE_TRANSITION | 96 | 0.61596215 | 1.5937248 | 0 | 0.12369496 | 1 | 2518 | tags=34%, list=9%, signal=38% |
| GO_ANTIGEN_PROCESSING_AND_PRESENTATION | 201 | 0.54674786 | 1.5934631 | 0 | 0.12354758 | 1 | 2364 | tags=29%, list=9%, signal=32% |
| MILI_PSEUDOPODIA | 46 | 0.67256004 | 1.5932055 | 0 | 0.09970503 | 1 | 3894 | tags=65%, list=14%, signal=76% |
| GO_MHC_PROTEIN_COMPLEX | 11 | 0.86167896 | 1.5925418 | 0 | 0.12518689 | 1 | 1347 | tags=55%, list=5%, signal=57% |
| GO_NEGATIVE_REGULATION_OF_ANTIGEN_RECEPTOR_MEDIATED_SIGNALING_PATHWAY | 26 | 0.684185 | 1.5925181 | 0.04651163 | 0.12499734 | 1 | 3025 | tags=38%, list=11%, signal=43% |
| REACTOME_INNATE_IMMUNE_SYSTEM | 867 | 0.49805638 | 1.5922887 | 0 | 0.10061578 | 1 | 2478 | tags=22%, list=9%, signal=23% |
| GO_NOTOCHORD_DEVELOPMENT | 17 | 0.79033834 | 1.591143 | 0 | 0.12638302 | 1 | 1528 | tags=18%, list=6%, signal=19% |
| GO_KETONE_CATABOLIC_PROCESS | 9 | 0.9006701 | 1.5910276 | 0 | 0.12626882 | 1 | 443 | tags=33%, list=2%, signal=34% |
| ST_TYPE_I_INTERFERON_PATHWAY | 7 | 0.9215958 | 1.590926 | 0 | 0.10171357 | 1 | 1909 | tags=57%, list=7%, signal=61% |
| HP_INTRACRANIAL_HEMORRHAGE | 106 | 0.6065874 | 1.590812 | 0 | 0.12631148 | 1 | 3767 | tags=35%, list=14%, signal=40% |
| GO_MITOCHONDRIAL_SMALL_RIBOSOMAL_SUBUNIT | 27 | 0.7597686 | 1.5907397 | 0 | 0.12608105 | 1 | 2348 | tags=48%, list=9%, signal=53% |
| WP_ALLOGRAFT_REJECTION | 63 | 0.60487914 | 1.5906997 | 0 | 0.10204059 | 1 | 1347 | tags=16%, list=5%, signal=17% |
| LI_WILMS_TUMOR_VS_FETAL_KIDNEY_1_UP | 173 | 0.55118483 | 1.5905945 | 0 | 0.10191191 | 1 | 2665 | tags=27%, list=10%, signal=30% |
| YAO_TEMPORAL_RESPONSE_TO_PROGESTERONE_CLUSTER_17 | 175 | 0.539484 | 1.5899904 | 0 | 0.10242358 | 1 | 2751 | tags=47%, list=10%, signal=52% |
| STEIN_ESRRA_TARGETS | 490 | 0.5031874 | 1.5895498 | 0 | 0.10267852 | 1 | 3899 | tags=40%, list=14%, signal=45% |
| GO_T_CELL_RECEPTOR_COMPLEX | 16 | 0.7722757 | 1.589045 | 0 | 0.12854795 | 1 | 2539 | tags=19%, list=9%, signal=21% |
| HP_RESPIRATORY_INSUFFICIENCY | 443 | 0.48780042 | 1.5890068 | 0 | 0.12831329 | 1 | 3524 | tags=32%, list=13%, signal=36% |
| GO_METANEPHROS_DEVELOPMENT | 77 | 0.58562624 | 1.5889078 | 0 | 0.1282746 | 1 | 1528 | tags=17%, list=6%, signal=18% |
| DAVICIONI_MOLECULAR_ARMS_VS_ERMS_DN | 166 | 0.5537581 | 1.5879586 | 0 | 0.10432693 | 1 | 3586 | tags=33%, list=13%, signal=38% |
| HOSHIDA_LIVER_CANCER_LATE_RECURRENCE_UP | 55 | 0.597248 | 1.587136 | 0 | 0.10514282 | 1 | 1918 | tags=31%, list=7%, signal=33% |
| HP_RETINAL_VASCULAR_TORTUOSITY | 38 | 0.7008447 | 1.5870144 | 0.04545455 | 0.13141339 | 1 | 854 | tags=24%, list=3%, signal=24% |
| BIOCARTA_MHC_PATHWAY | 8 | 0.9485856 | 1.586815 | 0 | 0.10544252 | 1 | 1237 | tags=88%, list=5%, signal=92% |
| YAO_TEMPORAL_RESPONSE_TO_PROGESTERONE_CLUSTER_6 | 63 | 0.6311339 | 1.5867877 | 0.02222222 | 0.10505199 | 1 | 2927 | tags=30%, list=11%, signal=34% |
| REACTOME_DEGRADATION_OF_DVL | 54 | 0.6699848 | 1.5866331 | 0 | 0.10510518 | 1 | 805 | tags=37%, list=3%, signal=38% |
| AFFAR_YY1_TARGETS_UP | 176 | 0.5518218 | 1.5866218 | 0 | 0.10471878 | 1 | 4054 | tags=35%, list=15%, signal=41% |
| HP_ABNORMALITY_OF_HEAD_BLOOD_VESSEL | 58 | 0.60312724 | 1.5865177 | 0 | 0.13179292 | 1 | 3197 | tags=34%, list=12%, signal=39% |
| HP_ABNORMALITY_OF_THE_LIVER | 824 | 0.47960022 | 1.5864992 | 0 | 0.13147534 | 1 | 3528 | tags=28%, list=13%, signal=32% |
| NABA_MATRISOME_ASSOCIATED | 514 | 0.4860181 | 1.586198 | 0 | 0.10501075 | 1 | 2783 | tags=14%, list=10%, signal=16% |
| HP_ABNORMALITY_OF_PANCREAS_PHYSIOLOGY | 103 | 0.57744247 | 1.5861217 | 0 | 0.13173804 | 1 | 3473 | tags=32%, list=13%, signal=37% |
| GO_SUCCINATE_DEHYDROGENASE_ACTIVITY | 4 | 0.97298807 | 1.5854762 | 0 | 0.13257314 | 1 | 730 | tags=100%, list=3%, signal=103% |
| HP_ABNORMALITY_OF_COAGULATION | 162 | 0.5557931 | 1.585318 | 0 | 0.13264045 | 1 | 3558 | tags=36%, list=13%, signal=41% |
| ASTON_MAJOR_DEPRESSIVE_DISORDER_DN | 149 | 0.5367439 | 1.5853052 | 0 | 0.10592914 | 1 | 2586 | tags=27%, list=10%, signal=30% |
| HP_HYPERHIDROSIS | 114 | 0.5466923 | 1.5852199 | 0 | 0.13244025 | 1 | 3407 | tags=32%, list=13%, signal=36% |
| GO_MICROVILLUS | 80 | 0.5957214 | 1.5849899 | 0.02857143 | 0.13269523 | 1 | 2372 | tags=23%, list=9%, signal=25% |
| GO_CYTOCHROME_COMPLEX_ASSEMBLY | 33 | 0.69112825 | 1.5849186 | 0.02222222 | 0.13245542 | 1 | 2622 | tags=39%, list=10%, signal=44% |
| GO_PURINE_NUCLEOSIDE_TRIPHOSPHATE_METABOLIC_PROCESS | 77 | 0.5935678 | 1.5848279 | 0 | 0.13233227 | 1 | 3276 | tags=43%, list=12%, signal=49% |
| HP_CONGESTIVE_HEART_FAILURE | 172 | 0.55516464 | 1.5846032 | 0 | 0.13243437 | 1 | 2434 | tags=27%, list=9%, signal=29% |
| JOHNSTONE_PARVB_TARGETS_2_UP | 131 | 0.56130135 | 1.584432 | 0 | 0.10683428 | 1 | 2870 | tags=33%, list=11%, signal=37% |
| GO_SENSORY_PERCEPTION_OF_BITTER_TASTE | 15 | 0.7781413 | 1.5843067 | 0.02325581 | 0.13272752 | 1 | 1662 | tags=13%, list=6%, signal=14% |
| WU_HBX_TARGETS_1_UP | 15 | 0.7933253 | 1.5842493 | 0 | 0.10662736 | 1 | 1195 | tags=33%, list=4%, signal=35% |
| KEGG_SYSTEMIC_LUPUS_ERYTHEMATOSUS | 37 | 0.6761397 | 1.58419 | 0.025 | 0.10624243 | 1 | 1638 | tags=30%, list=6%, signal=32% |
| HP_ABNORMAL_URINE_PROTEIN_LEVEL | 166 | 0.54422116 | 1.5841386 | 0 | 0.13275458 | 1 | 3535 | tags=30%, list=13%, signal=34% |
| HP_VENTRICULAR_HYPERTROPHY | 100 | 0.58533555 | 1.5840049 | 0 | 0.13266882 | 1 | 2581 | tags=23%, list=10%, signal=25% |
| HP_VENTRICULAR_TACHYCARDIA | 43 | 0.673032 | 1.5838996 | 0 | 0.13254464 | 1 | 1628 | tags=19%, list=6%, signal=20% |
| GO_T_CELL_LINEAGE_COMMITMENT | 24 | 0.7461317 | 1.5821196 | 0.025 | 0.13568044 | 1 | 2089 | tags=25%, list=8%, signal=27% |
| HP_ELEVATED_ERYTHROCYTE_SEDIMENTATION_RATE | 28 | 0.72236603 | 1.5819829 | 0 | 0.13562562 | 1 | 1785 | tags=25%, list=7%, signal=27% |
| GO_NEGATIVE_REGULATION_OF_PROTEIN_CATABOLIC_PROCESS | 123 | 0.561349 | 1.5819714 | 0 | 0.1353475 | 1 | 2304 | tags=30%, list=9%, signal=33% |
| HP_PSYCHOMOTOR_RETARDATION | 84 | 0.6044417 | 1.5816202 | 0 | 0.13570292 | 1 | 3266 | tags=40%, list=12%, signal=46% |
| GO_HINDBRAIN_DEVELOPMENT | 140 | 0.54245913 | 1.5815965 | 0 | 0.13542634 | 1 | 2371 | tags=23%, list=9%, signal=25% |
| HP_METABOLIC_ALKALOSIS | 10 | 0.8406907 | 1.5809486 | 0.02564103 | 0.13633502 | 1 | 1370 | tags=20%, list=5%, signal=21% |
| BIOCARTA_RNA_PATHWAY | 9 | 0.85005313 | 1.5804372 | 0 | 0.11085535 | 1 | 337 | tags=44%, list=1%, signal=45% |
| UROSEVIC_RESPONSE_TO_IMIQUIMOD | 17 | 0.83067 | 1.5803763 | 0 | 0.11069974 | 1 | 2141 | tags=29%, list=8%, signal=32% |
| KIM_MYCN_AMPLIFICATION_TARGETS_UP | 76 | 0.57847625 | 1.5803549 | 0 | 0.11030439 | 1 | 4247 | tags=45%, list=16%, signal=53% |
| GO_POSITIVE_REGULATION_OF_WNT_SIGNALING_PATHWAY | 171 | 0.52769506 | 1.5801857 | 0 | 0.13705534 | 1 | 2717 | tags=26%, list=10%, signal=28% |
| JINESH_BLEBBISHIELD_TO_IMMUNE_CELL_FUSION_PBSHMS_DN | 316 | 0.50422734 | 1.5798622 | 0 | 0.1109952 | 1 | 3925 | tags=35%, list=15%, signal=41% |
| REACTOME_NUCLEAR_SIGNALING_BY_ERBB4 | 31 | 0.7085155 | 1.579741 | 0 | 0.1106016 | 1 | 3053 | tags=35%, list=11%, signal=40% |
| HP_ROTARY_NYSTAGMUS | 8 | 0.9035516 | 1.5797269 | 0.01923077 | 0.13769692 | 1 | 1197 | tags=38%, list=4%, signal=39% |
| GO_DETECTION_OF_STIMULUS_INVOLVED_IN_SENSORY_PERCEPTION | 74 | 0.6021185 | 1.5796788 | 0 | 0.13745378 | 1 | 1729 | tags=11%, list=6%, signal=12% |
| GO_ANTIGEN_PROCESSING_AND_PRESENTATION_OF_ENDOGENOUS_PEPTIDE_ANTIGEN | 10 | 0.889475 | 1.5794708 | 0 | 0.13747104 | 1 | 1967 | tags=60%, list=7%, signal=65% |
| BURTON_ADIPOGENESIS_PEAK_AT_0HR | 57 | 0.6515031 | 1.579379 | 0 | 0.11057051 | 1 | 2539 | tags=32%, list=9%, signal=35% |
| GO_INNATE_IMMUNE_RESPONSE_ACTIVATING_SIGNAL_TRANSDUCTION | 99 | 0.59677935 | 1.5791255 | 0 | 0.13785096 | 1 | 1624 | tags=23%, list=6%, signal=25% |
| CHYLA_CBFA2T3_TARGETS_UP | 337 | 0.5158101 | 1.5790865 | 0 | 0.11041802 | 1 | 2889 | tags=19%, list=11%, signal=21% |
| HP_MUSCLE_HYPERTROPHY_OF_THE_LOWER_EXTREMITIES | 38 | 0.6414363 | 1.5786973 | 0.04651163 | 0.13837628 | 1 | 2302 | tags=26%, list=9%, signal=29% |
| HP_ABNORMAL_LYMPHOCYTE_PHYSIOLOGY | 204 | 0.531237 | 1.5781614 | 0 | 0.13937293 | 1 | 2876 | tags=22%, list=11%, signal=25% |
| REACTOME_PHENYLALANINE_AND_TYROSINE_METABOLISM | 8 | 0.90396327 | 1.5780771 | 0 | 0.11145417 | 1 | 1291 | tags=25%, list=5%, signal=26% |
| WANG_SMARCE1_TARGETS_UP | 258 | 0.5074139 | 1.5776688 | 0 | 0.11177555 | 1 | 3139 | tags=27%, list=12%, signal=30% |
| FLECHNER_BIOPSY_KIDNEY_TRANSPLANT_REJECTED_VS_OK_DN | 511 | 0.5005716 | 1.5776329 | 0 | 0.11144519 | 1 | 3025 | tags=29%, list=11%, signal=32% |
| REACTOME_DEGRADATION_OF_BETA_CATENIN_BY_THE_DESTRUCTION_COMPLEX | 80 | 0.58783835 | 1.5774444 | 0.0212766 | 0.11117494 | 1 | 2352 | tags=39%, list=9%, signal=42% |
| ANASTASSIOU_MULTICANCER_INVASIVENESS_SIGNATURE | 60 | 0.64028674 | 1.5772878 | 0 | 0.11114438 | 1 | 3049 | tags=27%, list=11%, signal=30% |
| SWEET_KRAS_TARGETS_UP | 74 | 0.59500045 | 1.5764751 | 0 | 0.11169602 | 1 | 2435 | tags=28%, list=9%, signal=31% |
| GO_RESPONSE_TO_TYPE_I_INTERFERON | 71 | 0.6182781 | 1.5763186 | 0 | 0.14261737 | 1 | 2468 | tags=46%, list=9%, signal=51% |
| HP_PSYCHOMOTOR_DETERIORATION | 12 | 0.86435246 | 1.5761027 | 0 | 0.14276613 | 1 | 1776 | tags=50%, list=7%, signal=54% |
| BRUINS_UVC_RESPONSE_LATE | 1050 | 0.47688034 | 1.5757728 | 0 | 0.11236356 | 1 | 4249 | tags=39%, list=16%, signal=45% |
| NIKOLSKY_BREAST_CANCER_17Q21_Q25_AMPLICON | 274 | 0.50950974 | 1.575635 | 0 | 0.11209565 | 1 | 3903 | tags=32%, list=14%, signal=37% |
| JIANG_VHL_TARGETS | 78 | 0.55288535 | 1.5751607 | 0 | 0.11235144 | 1 | 3564 | tags=47%, list=13%, signal=55% |
| HP_ABNORMALITY_OF_BRAINSTEM_MORPHOLOGY | 175 | 0.5324261 | 1.5749671 | 0 | 0.14512146 | 1 | 3274 | tags=33%, list=12%, signal=37% |
| HP_HYPOPARATHYROIDISM | 32 | 0.6630898 | 1.5749518 | 0.02702703 | 0.1447946 | 1 | 2295 | tags=28%, list=9%, signal=31% |
| KIM_LIVER_CANCER_POOR_SURVIVAL_DN | 33 | 0.6859996 | 1.5749265 | 0 | 0.11219866 | 1 | 2691 | tags=18%, list=10%, signal=20% |
| GO_MITOCHONDRIAL_TRANSPORT | 242 | 0.51697433 | 1.5748261 | 0 | 0.14472307 | 1 | 2410 | tags=28%, list=9%, signal=31% |
| HP_ABNORMALITY_OF_THE_THORACIC_SPINE | 72 | 0.60855263 | 1.5747204 | 0 | 0.14443436 | 1 | 2221 | tags=28%, list=8%, signal=30% |
| HP_TACHYPNEA | 96 | 0.5671331 | 1.5746988 | 0 | 0.14418267 | 1 | 3788 | tags=29%, list=14%, signal=34% |
| WU_SILENCED_BY_METHYLATION_IN_BLADDER_CANCER | 42 | 0.6311319 | 1.5745013 | 0.025 | 0.11285188 | 1 | 2459 | tags=21%, list=9%, signal=24% |
| GO_CALCIUM_DEPENDENT_PROTEIN_BINDING | 62 | 0.60748726 | 1.5743145 | 0 | 0.1443601 | 1 | 2131 | tags=24%, list=8%, signal=26% |
| GO_ACYL_COA_DEHYDROGENASE_ACTIVITY | 10 | 0.81795913 | 1.5739493 | 0.05 | 0.14464633 | 1 | 4505 | tags=90%, list=17%, signal=108% |
| GO_S_SHAPED_BODY_MORPHOGENESIS | 7 | 0.93753594 | 1.5737603 | 0 | 0.144538 | 1 | 167 | tags=29%, list=1%, signal=29% |
| GO_RNA_POLYMERASE_III_COMPLEX | 17 | 0.7569254 | 1.5737067 | 0.02631579 | 0.1443233 | 1 | 2452 | tags=53%, list=9%, signal=58% |
| GO_MULTICELLULAR_ORGANISM_AGING | 28 | 0.70260817 | 1.5729649 | 0.03225806 | 0.14531273 | 1 | 2350 | tags=32%, list=9%, signal=35% |
| GO_REGULATION_OF_GTP_BINDING | 10 | 0.8281144 | 1.5729567 | 0 | 0.14499265 | 1 | 3140 | tags=60%, list=12%, signal=68% |
| GO_INNATE_IMMUNE_RESPONSE | 665 | 0.47618738 | 1.5726248 | 0 | 0.14534336 | 1 | 2301 | tags=19%, list=9%, signal=21% |
| GO_TRIGLYCERIDE_CATABOLIC_PROCESS | 24 | 0.7240759 | 1.572518 | 0.02631579 | 0.14513053 | 1 | 2476 | tags=25%, list=9%, signal=28% |
| HP_INFLAMMATION_OF_THE_LARGE_INTESTINE | 44 | 0.6480901 | 1.5724622 | 0 | 0.14491808 | 1 | 1788 | tags=18%, list=7%, signal=19% |
| HP_ABNORMAL_RETINAL_ARTERY_MORPHOLOGY | 47 | 0.6429675 | 1.5724614 | 0 | 0.14460167 | 1 | 966 | tags=19%, list=4%, signal=20% |
| REACTOME_MITOCHONDRIAL_PROTEIN_IMPORT | 62 | 0.59692764 | 1.571986 | 0.03448276 | 0.1157849 | 1 | 2883 | tags=44%, list=11%, signal=49% |
| MEBARKI_HCC_PROGENITOR_WNT_UP_CTNNB1_DEPENDENT | 65 | 0.5707061 | 1.5719815 | 0 | 0.11539505 | 1 | 1896 | tags=20%, list=7%, signal=21% |
| GO_ORGANIC_ACID_METABOLIC_PROCESS | 985 | 0.46428433 | 1.5715929 | 0 | 0.14627689 | 1 | 3228 | tags=25%, list=12%, signal=28% |
| GO_HYPEROSMOTIC_RESPONSE | 25 | 0.7192889 | 1.5713503 | 0 | 0.14634341 | 1 | 2118 | tags=24%, list=8%, signal=26% |
| GO_FC_EPSILON_RECEPTOR_SIGNALING_PATHWAY | 106 | 0.560817 | 1.5710906 | 0 | 0.14657956 | 1 | 2389 | tags=28%, list=9%, signal=31% |
| NADLER_OBESITY_DN | 44 | 0.65414107 | 1.5706102 | 0 | 0.11687961 | 1 | 1609 | tags=25%, list=6%, signal=27% |
| REACTOME_AMINO_ACIDS_REGULATE_MTORC1 | 51 | 0.6260198 | 1.5705608 | 0.02631579 | 0.11660371 | 1 | 1933 | tags=27%, list=7%, signal=30% |
| RUTELLA_RESPONSE_TO_HGF_VS_CSF2RB_AND_IL4_UP | 378 | 0.50453335 | 1.5701519 | 0 | 0.11667208 | 1 | 3031 | tags=31%, list=11%, signal=34% |
| CHOW_RASSF1_TARGETS_DN | 28 | 0.73999345 | 1.5696155 | 0 | 0.11701371 | 1 | 4084 | tags=71%, list=15%, signal=84% |
| HP_MOTOR_SEIZURE | 280 | 0.51113147 | 1.5688449 | 0 | 0.1510043 | 1 | 3184 | tags=25%, list=12%, signal=28% |
| HP_CHEST_PAIN | 76 | 0.5873475 | 1.5686307 | 0 | 0.15098995 | 1 | 2400 | tags=28%, list=9%, signal=30% |
| GO_PHAGOSOME_MATURATION | 47 | 0.68692744 | 1.5686142 | 0.02777778 | 0.15069954 | 1 | 1206 | tags=26%, list=4%, signal=27% |
| GO_NEGATIVE_REGULATION_OF_INTRACELLULAR_TRANSPORT | 53 | 0.65372896 | 1.5682838 | 0.02702703 | 0.15099418 | 1 | 2747 | tags=36%, list=10%, signal=40% |
| GO_PROTEIN_TARGETING_TO_MITOCHONDRION | 92 | 0.53775036 | 1.5681134 | 0 | 0.15114985 | 1 | 2410 | tags=33%, list=9%, signal=36% |
| GO_REGULATION_OF_RESPONSE_TO_BIOTIC_STIMULUS | 338 | 0.5020801 | 1.5680637 | 0 | 0.15082689 | 1 | 2479 | tags=22%, list=9%, signal=23% |
| HP_BRAIN_IMAGING_ABNORMALITY | 63 | 0.6207047 | 1.568006 | 0 | 0.15070882 | 1 | 2068 | tags=35%, list=8%, signal=38% |
| DAUER_STAT3_TARGETS_DN | 41 | 0.6876253 | 1.5674446 | 0 | 0.11915144 | 1 | 1536 | tags=44%, list=6%, signal=46% |
| WEINMANN_ADAPTATION_TO_HYPOXIA_DN | 36 | 0.6567854 | 1.567403 | 0.02222222 | 0.1188695 | 1 | 2115 | tags=28%, list=8%, signal=30% |
| KEGG_ANTIGEN_PROCESSING_AND_PRESENTATION | 45 | 0.6472854 | 1.5671781 | 0 | 0.11892724 | 1 | 1557 | tags=31%, list=6%, signal=33% |
| GO_ENDOPEPTIDASE_ACTIVATOR_ACTIVITY | 7 | 0.91992813 | 1.567153 | 0 | 0.1526678 | 1 | 657 | tags=43%, list=2%, signal=44% |
| GO_CELLULAR_RESPONSE_TO_INTERFERON_BETA | 16 | 0.77785254 | 1.5670042 | 0.025 | 0.15278661 | 1 | 1289 | tags=38%, list=5%, signal=39% |
| REACTOME_DEGRADATION_OF_AXIN | 52 | 0.638086 | 1.5666718 | 0 | 0.11941958 | 1 | 805 | tags=37%, list=3%, signal=38% |
| GO_PLATELET_DENSE_GRANULE_LUMEN | 9 | 0.8623844 | 1.566173 | 0 | 0.15439418 | 1 | 2870 | tags=56%, list=11%, signal=62% |
| HP_GENERALIZED_JOINT_LAXITY | 36 | 0.66119146 | 1.565823 | 0 | 0.15455423 | 1 | 660 | tags=14%, list=2%, signal=14% |
| REACTOME_DECTIN_1_MEDIATED_NONCANONICAL_NF_KB_SIGNALING | 58 | 0.6141451 | 1.5654709 | 0 | 0.12052914 | 1 | 805 | tags=33%, list=3%, signal=34% |
| FOURNIER_ACINAR_DEVELOPMENT_LATE_UP | 9 | 0.8840569 | 1.5651678 | 0 | 0.12068909 | 1 | 269 | tags=22%, list=1%, signal=22% |
| GO_SECRETORY_GRANULE | 718 | 0.4784494 | 1.5650841 | 0 | 0.15567522 | 1 | 3182 | tags=24%, list=12%, signal=27% |
| REACTOME_CELLULAR_RESPONSE_TO_HYPOXIA | 69 | 0.5987395 | 1.5650092 | 0.02380952 | 0.12051883 | 1 | 805 | tags=29%, list=3%, signal=30% |
| FERREIRA_EWINGS_SARCOMA_UNSTABLE_VS_STABLE_DN | 84 | 0.60004896 | 1.5650033 | 0 | 0.12013006 | 1 | 2770 | tags=32%, list=10%, signal=36% |
| REACTOME_PYRUVATE_METABOLISM | 28 | 0.6704085 | 1.5649546 | 0 | 0.11974379 | 1 | 2246 | tags=29%, list=8%, signal=31% |
| HP_GASTROINTESTINAL_DYSMOTILITY | 57 | 0.61693954 | 1.5648816 | 0 | 0.15565212 | 1 | 2050 | tags=28%, list=8%, signal=30% |
| GO_OXIDOREDUCTASE_ACTIVITY_ACTING_ON_PEROXIDE_AS_ACCEPTOR | 44 | 0.6373945 | 1.5647933 | 0 | 0.15552825 | 1 | 2604 | tags=32%, list=10%, signal=35% |
| WP_STRIATED_MUSCLE_CONTRACTION_PATHWAY | 31 | 0.7237295 | 1.5645274 | 0 | 0.12039506 | 1 | 1714 | tags=23%, list=6%, signal=24% |
| GO_EXTRINSIC_COMPONENT_OF_EXTERNAL_SIDE_OF_PLASMA_MEMBRANE | 5 | 0.98691297 | 1.5644741 | 0 | 0.15563743 | 1 | 42 | tags=40%, list=0%, signal=40% |
| NIKOLSKY_BREAST_CANCER_8P12_P11_AMPLICON | 47 | 0.6295224 | 1.5644386 | 0 | 0.12011962 | 1 | 4127 | tags=45%, list=15%, signal=53% |
| SHETH_LIVER_CANCER_VS_TXNIP_LOSS_PAM4 | 240 | 0.53441936 | 1.5638645 | 0 | 0.12065312 | 1 | 3714 | tags=29%, list=14%, signal=33% |
| GO_EXTRACELLULAR_MATRIX_STRUCTURAL_CONSTITUENT | 138 | 0.53572124 | 1.5637361 | 0 | 0.15688299 | 1 | 4609 | tags=33%, list=17%, signal=40% |
| MOHANKUMAR_HOXA1_TARGETS_DN | 137 | 0.5307992 | 1.5631905 | 0 | 0.12155443 | 1 | 1609 | tags=15%, list=6%, signal=16% |
| HP_ABNORMALITY_OF_TEMPERATURE_REGULATION | 285 | 0.52304894 | 1.5630687 | 0 | 0.15792365 | 1 | 3989 | tags=30%, list=15%, signal=35% |
| GO_LATE_ENDOSOME | 239 | 0.5003047 | 1.5629053 | 0 | 0.15782921 | 1 | 2713 | tags=30%, list=10%, signal=33% |
| GO_LIPID_OXIDATION | 100 | 0.56037515 | 1.5626419 | 0 | 0.15799813 | 1 | 3779 | tags=40%, list=14%, signal=46% |
| SASAI_TARGETS_OF_CXCR6_AND_PTCH1_DN | 5 | 0.92312926 | 1.5626097 | 0 | 0.1222404 | 1 | 165 | tags=40%, list=1%, signal=40% |
| WINNEPENNINCKX_MELANOMA_METASTASIS_DN | 35 | 0.71563613 | 1.5624483 | 0 | 0.12238871 | 1 | 268 | tags=17%, list=1%, signal=17% |
| GO_ALCOHOL_BINDING | 74 | 0.58840054 | 1.5623614 | 0 | 0.15813576 | 1 | 2434 | tags=26%, list=9%, signal=28% |
| HP_THROMBOEMBOLISM | 21 | 0.7423272 | 1.5622137 | 0 | 0.15810585 | 1 | 4257 | tags=43%, list=16%, signal=51% |
| BURTON_ADIPOGENESIS_9 | 84 | 0.6055679 | 1.5619855 | 0 | 0.12264722 | 1 | 1622 | tags=30%, list=6%, signal=32% |
| TIEN_INTESTINE_PROBIOTICS_24HR_UP | 533 | 0.4900178 | 1.5617061 | 0 | 0.12290271 | 1 | 3034 | tags=38%, list=11%, signal=42% |
| HP_WHEEZING | 6 | 0.94690835 | 1.5616308 | 0 | 0.15913399 | 1 | 1108 | tags=33%, list=4%, signal=35% |
| GOTZMANN_EPITHELIAL_TO_MESENCHYMAL_TRANSITION_DN | 191 | 0.52649534 | 1.5615225 | 0 | 0.12288594 | 1 | 2154 | tags=34%, list=8%, signal=36% |
| GO_T_CELL_DIFFERENTIATION_IN_THYMUS | 68 | 0.602004 | 1.5608114 | 0 | 0.16062306 | 1 | 1679 | tags=16%, list=6%, signal=17% |
| KAAB_HEART_ATRIUM_VS_VENTRICLE_DN | 242 | 0.52379525 | 1.560767 | 0 | 0.12408862 | 1 | 3440 | tags=32%, list=13%, signal=36% |
| HP_VARIABLE_EXPRESSIVITY | 284 | 0.51363766 | 1.5606368 | 0 | 0.16068856 | 1 | 3437 | tags=27%, list=13%, signal=31% |
| HP_PSYCHOGENIC_NON_EPILEPTIC_SEIZURE | 32 | 0.7049354 | 1.5606318 | 0.02380952 | 0.16035928 | 1 | 4101 | tags=63%, list=15%, signal=74% |
| HP_ABNORMAL_RENAL_TUBULE_MORPHOLOGY | 72 | 0.58146715 | 1.5605462 | 0 | 0.16042373 | 1 | 3259 | tags=31%, list=12%, signal=35% |
| REACTOME_CELL_JUNCTION_ORGANIZATION | 75 | 0.5653679 | 1.560258 | 0 | 0.12491429 | 1 | 4872 | tags=40%, list=18%, signal=49% |
| GO_NEPHRON_MORPHOGENESIS | 66 | 0.60601354 | 1.5602101 | 0 | 0.16088158 | 1 | 1694 | tags=18%, list=6%, signal=19% |
| HP_ABNORMALITY_OF_COORDINATION | 856 | 0.4824684 | 1.5601332 | 0 | 0.16068533 | 1 | 3473 | tags=30%, list=13%, signal=34% |
| INGRAM_SHH_TARGETS_UP | 110 | 0.56493986 | 1.5600702 | 0 | 0.12489166 | 1 | 2853 | tags=33%, list=11%, signal=36% |
| HP_GENERALIZED_ONSET_MOTOR_SEIZURE | 111 | 0.56059444 | 1.5599936 | 0 | 0.16068402 | 1 | 2466 | tags=22%, list=9%, signal=24% |
| MCBRYAN_PUBERTAL_TGFB1_TARGETS_UP | 162 | 0.5408661 | 1.5595201 | 0 | 0.12549406 | 1 | 3085 | tags=37%, list=11%, signal=42% |
| GO_POSITIVE_REGULATION_OF_TYROSINE_PHOSPHORYLATION_OF_STAT_PROTEIN | 48 | 0.5952036 | 1.5594842 | 0 | 0.16123715 | 1 | 1939 | tags=19%, list=7%, signal=20% |
| VECCHI_GASTRIC_CANCER_EARLY_DN | 289 | 0.50560653 | 1.5590391 | 0 | 0.1255245 | 1 | 4168 | tags=26%, list=15%, signal=31% |
| CHANDRAN_METASTASIS_DN | 283 | 0.49184176 | 1.5586982 | 0 | 0.12560658 | 1 | 3588 | tags=29%, list=13%, signal=33% |
| GO_REGULATION_OF_IMMUNE_RESPONSE | 721 | 0.47402993 | 1.5583389 | 0 | 0.1633112 | 1 | 2510 | tags=19%, list=9%, signal=21% |
| HP_GOWERS_SIGN | 50 | 0.63381207 | 1.5577601 | 0 | 0.16408417 | 1 | 4184 | tags=42%, list=16%, signal=50% |
| HP_ELEVATED_TISSUE_NON_SPECIFIC_ALKALINE_PHOSPHATASE | 9 | 0.9037852 | 1.5571893 | 0 | 0.16503435 | 1 | 2008 | tags=33%, list=7%, signal=36% |
| REACTOME_DIGESTION_AND_ABSORPTION | 14 | 0.7738455 | 1.5569004 | 0 | 0.12815966 | 1 | 398 | tags=7%, list=1%, signal=7% |
| HP_ABNORMAL_CIRCULATING_IGG_LEVEL | 43 | 0.675899 | 1.5563359 | 0.02564103 | 0.16682765 | 1 | 2833 | tags=23%, list=11%, signal=26% |
| GO_BLOOD_MICROPARTICLE | 78 | 0.57728344 | 1.5562192 | 0 | 0.16690995 | 1 | 1646 | tags=15%, list=6%, signal=16% |
| HP_INTESTINAL_PSEUDO_OBSTRUCTION | 20 | 0.7388565 | 1.5561674 | 0.02173913 | 0.16667268 | 1 | 854 | tags=30%, list=3%, signal=31% |
| GO_PURINE_NUCLEOBASE_TRANSPORT | 6 | 0.94502836 | 1.5556933 | 0 | 0.1673996 | 1 | 171 | tags=33%, list=1%, signal=34% |
| GO_PEPTIDE_BINDING | 247 | 0.5029515 | 1.5552477 | 0 | 0.1679435 | 1 | 2650 | tags=21%, list=10%, signal=24% |
| GO_ESTABLISHMENT_OF_TISSUE_POLARITY | 118 | 0.5594079 | 1.5548657 | 0 | 0.16856341 | 1 | 2270 | tags=28%, list=8%, signal=30% |
| HP_APLASIA_HYPOPLASIA_OF_THE_LENS | 8 | 0.92606217 | 1.554765 | 0.02040816 | 0.1685473 | 1 | 18 | tags=13%, list=0%, signal=13% |
| FIGUEROA_AML_METHYLATION_CLUSTER_1_UP | 103 | 0.55964595 | 1.5547336 | 0 | 0.13095717 | 1 | 3780 | tags=34%, list=14%, signal=39% |
| GO_MESENCHYMAL_CELL_PROLIFERATION | 39 | 0.65041655 | 1.5547152 | 0 | 0.16840376 | 1 | 627 | tags=18%, list=2%, signal=18% |
| GO_ACTIN_FILAMENT_BUNDLE | 65 | 0.60149467 | 1.5546662 | 0.02439024 | 0.16819699 | 1 | 3368 | tags=34%, list=13%, signal=39% |
| GO_TRANSITION_METAL_ION_TRANSPORT | 111 | 0.5518925 | 1.5542616 | 0 | 0.16849592 | 1 | 1766 | tags=20%, list=7%, signal=21% |
| LUI_THYROID_CANCER_CLUSTER_4 | 11 | 0.8035776 | 1.5541091 | 0.04347826 | 0.13204615 | 1 | 1237 | tags=36%, list=5%, signal=38% |
| HP_FATIGABLE_WEAKNESS_OF_DISTAL_LIMB_MUSCLES | 7 | 0.9056237 | 1.5539435 | 0 | 0.16882555 | 1 | 427 | tags=29%, list=2%, signal=29% |
| BOSCO_INTERFERON_INDUCED_ANTIVIRAL_MODULE | 65 | 0.5845645 | 1.5537434 | 0 | 0.13220741 | 1 | 2374 | tags=37%, list=9%, signal=40% |
| HP_DEMENTIA | 153 | 0.5630444 | 1.553388 | 0 | 0.16953063 | 1 | 1153 | tags=20%, list=4%, signal=21% |
| SWEET_LUNG_CANCER_KRAS_UP | 431 | 0.47480264 | 1.5532386 | 0 | 0.13252978 | 1 | 3067 | tags=33%, list=11%, signal=37% |
| GALLUZZI_PREVENT_MITOCHONDIAL_PERMEABILIZATION | 13 | 0.8577117 | 1.5526493 | 0 | 0.13279435 | 1 | 1181 | tags=46%, list=4%, signal=48% |
| GO_REGULATION_OF_MESENCHYMAL_CELL_PROLIFERATION | 30 | 0.65919816 | 1.5525024 | 0.04255319 | 0.17117128 | 1 | 2066 | tags=27%, list=8%, signal=29% |
| BRUECKNER_TARGETS_OF_MIRLET7A3_UP | 98 | 0.58201253 | 1.5523558 | 0.02777778 | 0.13269651 | 1 | 3644 | tags=38%, list=14%, signal=44% |
| HP_APLASTIC_ANEMIA | 16 | 0.7937525 | 1.5522599 | 0 | 0.1714009 | 1 | 2156 | tags=38%, list=8%, signal=41% |
| REACTOME_INTERFERON_ALPHA_BETA_SIGNALING | 46 | 0.65853536 | 1.5521123 | 0 | 0.13244909 | 1 | 2378 | tags=54%, list=9%, signal=60% |
| HP_PROGRESSIVE_SPASTIC_QUADRIPLEGIA | 9 | 0.8853776 | 1.5519111 | 0.02564103 | 0.17159867 | 1 | 70 | tags=22%, list=0%, signal=22% |
| HP_NAUSEA | 53 | 0.63147783 | 1.5516635 | 0 | 0.17182627 | 1 | 4093 | tags=47%, list=15%, signal=56% |
| MOROSETTI_FACIOSCAPULOHUMERAL_MUSCULAR_DISTROPHY_UP | 16 | 0.7677379 | 1.551636 | 0.02173913 | 0.132912 | 1 | 411 | tags=25%, list=2%, signal=25% |
| GO_INTEGRIN_MEDIATED_SIGNALING_PATHWAY | 97 | 0.5860346 | 1.5515403 | 0 | 0.17177363 | 1 | 2346 | tags=21%, list=9%, signal=23% |
| REACTOME_TNFR2_NON_CANONICAL_NF_KB_PATHWAY | 90 | 0.5760446 | 1.5514241 | 0 | 0.13317175 | 1 | 805 | tags=21%, list=3%, signal=22% |
| REACTOME_ASYMMETRIC_LOCALIZATION_OF_PCP_PROTEINS | 61 | 0.62379473 | 1.5513908 | 0 | 0.13277659 | 1 | 1624 | tags=36%, list=6%, signal=38% |
| HP_ABNORMAL_BLEEDING | 376 | 0.48500317 | 1.5511816 | 0 | 0.17246316 | 1 | 3815 | tags=28%, list=14%, signal=32% |
| GO_CEREBELLAR_GRANULAR_LAYER_DEVELOPMENT | 12 | 0.8121586 | 1.5508935 | 0.02631579 | 0.1728422 | 1 | 165 | tags=17%, list=1%, signal=17% |
| HP_ABNORMALITY_OF_THE_ABDOMINAL_ORGANS | 970 | 0.46688575 | 1.5504438 | 0 | 0.17358965 | 1 | 3528 | tags=27%, list=13%, signal=30% |
| GO_GENITALIA_MORPHOGENESIS | 11 | 0.79330593 | 1.550207 | 0.02380952 | 0.17374866 | 1 | 531 | tags=18%, list=2%, signal=19% |
| GO_SUBSTANTIA_NIGRA_DEVELOPMENT | 37 | 0.69713765 | 1.5498335 | 0 | 0.17452385 | 1 | 2089 | tags=49%, list=8%, signal=53% |
| TORCHIA_TARGETS_OF_EWSR1_FLI1_FUSION_TOP20_DN | 16 | 0.7792793 | 1.5493193 | 0 | 0.13573769 | 1 | 4313 | tags=63%, list=16%, signal=74% |
| POOLA_INVASIVE_BREAST_CANCER_DN | 115 | 0.56161904 | 1.5493107 | 0 | 0.13538684 | 1 | 2639 | tags=20%, list=10%, signal=22% |
| GO_GLYCOSIDE_METABOLIC_PROCESS | 13 | 0.8251231 | 1.5492872 | 0 | 0.17557396 | 1 | 3410 | tags=69%, list=13%, signal=79% |
| HP_INFANTILE_MUSCULAR_HYPOTONIA | 134 | 0.52387446 | 1.5490634 | 0 | 0.1756655 | 1 | 1963 | tags=19%, list=7%, signal=20% |
| MASSARWEH_TAMOXIFEN_RESISTANCE_UP | 505 | 0.4716963 | 1.5488179 | 0 | 0.13518937 | 1 | 3693 | tags=29%, list=14%, signal=33% |
| HP_GASTROINTESTINAL_HEMORRHAGE | 114 | 0.56643504 | 1.5486656 | 0 | 0.17603366 | 1 | 2400 | tags=23%, list=9%, signal=25% |
| REACTOME_REGULATION_OF_INSULIN_LIKE_GROWTH_FACTOR_IGF_TRANSPORT_AND_UPTAKE_BY_INSULIN_LIKE_GROWTH_FACTOR_BINDING_PROTEINS_IGFBPS_ | 97 | 0.579166 | 1.5485657 | 0 | 0.13528718 | 1 | 1485 | tags=20%, list=6%, signal=21% |
| GO_TRICARBOXYLIC_ACID_CYCLE | 32 | 0.6850118 | 1.548373 | 0 | 0.17633864 | 1 | 2982 | tags=50%, list=11%, signal=56% |
| HP_APLASIA_HYPOPLASIA_OF_THE_OVARY | 57 | 0.6045039 | 1.548371 | 0 | 0.17600276 | 1 | 4790 | tags=42%, list=18%, signal=51% |
| HP_GYNECOMASTIA | 73 | 0.5561167 | 1.5478373 | 0 | 0.17700876 | 1 | 2338 | tags=22%, list=9%, signal=24% |
| REACTOME_TCR_SIGNALING | 106 | 0.566389 | 1.5477381 | 0 | 0.13631769 | 1 | 2352 | tags=30%, list=9%, signal=33% |
| HP_ABNORMALITY_OF_HUMORAL_IMMUNITY | 200 | 0.5232099 | 1.5474455 | 0 | 0.17749192 | 1 | 2876 | tags=21%, list=11%, signal=23% |
| GO_RESPONSE_TO_PAIN | 24 | 0.68403125 | 1.5471395 | 0.025 | 0.1778532 | 1 | 1634 | tags=13%, list=6%, signal=13% |
| GO_PIGMENT_BIOSYNTHETIC_PROCESS | 54 | 0.6205868 | 1.5469686 | 0 | 0.17794147 | 1 | 1501 | tags=22%, list=6%, signal=23% |
| GO_REGULATION_OF_TRANSCRIPTION_FROM_RNA_POLYMERASE_II_PROMOTER_IN_RESPONSE_TO_HYPOXIA | 72 | 0.5967157 | 1.5467899 | 0 | 0.17799918 | 1 | 805 | tags=29%, list=3%, signal=30% |
| WANG_NEOPLASTIC_TRANSFORMATION_BY_CCND1_MYC | 20 | 0.7794333 | 1.5467414 | 0.0212766 | 0.13794622 | 1 | 1012 | tags=35%, list=4%, signal=36% |
| GO_POSITIVE_REGULATION_OF_VOLTAGE_GATED_CALCIUM_CHANNEL_ACTIVITY | 9 | 0.87105864 | 1.54673 | 0 | 0.17778374 | 1 | 313 | tags=22%, list=1%, signal=22% |
| REACTOME_RUNX1_REGULATES_TRANSCRIPTION_OF_GENES_INVOLVED_IN_DIFFERENTIATION_OF_HSCS | 68 | 0.603108 | 1.5466466 | 0 | 0.13764334 | 1 | 805 | tags=29%, list=3%, signal=30% |
| GO_REGULATION_OF_LIPASE_ACTIVITY | 78 | 0.59631664 | 1.5465189 | 0 | 0.17787114 | 1 | 2728 | tags=24%, list=10%, signal=27% |
| GO_REGULATION_OF_WNT_SIGNALING_PATHWAY | 337 | 0.49107844 | 1.5464149 | 0 | 0.17768823 | 1 | 2804 | tags=24%, list=10%, signal=27% |
| HP_FAILURE_TO_THRIVE | 758 | 0.47894567 | 1.5462894 | 0 | 0.177716 | 1 | 3535 | tags=28%, list=13%, signal=32% |
| GO_MULTIVESICULAR_BODY_MEMBRANE | 11 | 0.7922434 | 1.5461593 | 0 | 0.17771237 | 1 | 2698 | tags=64%, list=10%, signal=71% |
| GO_SECRETORY_VESICLE | 861 | 0.46134108 | 1.5454947 | 0 | 0.17881641 | 1 | 3199 | tags=24%, list=12%, signal=26% |
| GO_AMIDE_BINDING | 310 | 0.49585044 | 1.5453172 | 0 | 0.17890264 | 1 | 2650 | tags=22%, list=10%, signal=24% |
| REACTOME_FGFR2_MUTANT_RECEPTOR_ACTIVATION | 27 | 0.6402865 | 1.544921 | 0.03225806 | 0.14061266 | 1 | 2758 | tags=41%, list=10%, signal=45% |
| GO_NEGATIVE_REGULATION_OF_CANONICAL_WNT_SIGNALING_PATHWAY | 163 | 0.5541904 | 1.5446893 | 0 | 0.17957221 | 1 | 1972 | tags=26%, list=7%, signal=28% |
| NUTT_GBM_VS_AO_GLIOMA_UP | 43 | 0.6626146 | 1.5445881 | 0.05 | 0.14083469 | 1 | 3250 | tags=49%, list=12%, signal=55% |
| FUNG_IL2_SIGNALING_1 | 9 | 0.88831174 | 1.544436 | 0 | 0.14052813 | 1 | 2591 | tags=67%, list=10%, signal=74% |
| GO_POSITIVE_REGULATION_OF_PHOSPHOLIPASE_ACTIVITY | 49 | 0.64986455 | 1.5443255 | 0 | 0.18001102 | 1 | 2728 | tags=29%, list=10%, signal=32% |
| FLECHNER_BIOPSY_KIDNEY_TRANSPLANT_REJECTED_VS_OK_UP | 82 | 0.53928894 | 1.5441029 | 0 | 0.1406124 | 1 | 2006 | tags=22%, list=7%, signal=24% |
| HP_MACULAR_DYSTROPHY | 11 | 0.8330792 | 1.5439934 | 0.02325581 | 0.18047933 | 1 | 640 | tags=18%, list=2%, signal=19% |
| REACTOME_CLEC7A_DECTIN_1_SIGNALING | 96 | 0.5605714 | 1.5436445 | 0 | 0.14132732 | 1 | 2389 | tags=31%, list=9%, signal=34% |
| GO_EPIDERMIS_MORPHOGENESIS | 26 | 0.76504564 | 1.543554 | 0.02564103 | 0.18144912 | 1 | 1194 | tags=15%, list=4%, signal=16% |
| FUNG_IL2_TARGETS_WITH_STAT5_BINDING_SITES | 6 | 0.88202184 | 1.5435374 | 0 | 0.14111637 | 1 | 2268 | tags=50%, list=8%, signal=55% |
| GO_CELLULAR_RESPONSE_TO_OXYGEN_LEVELS | 207 | 0.49288353 | 1.5434077 | 0 | 0.18141001 | 1 | 1786 | tags=23%, list=7%, signal=24% |
| REACTOME_NOTCH3_INTRACELLULAR_DOMAIN_REGULATES_TRANSCRIPTION | 24 | 0.7503463 | 1.5430937 | 0 | 0.14144087 | 1 | 333 | tags=25%, list=1%, signal=25% |
| KOBAYASHI_EGFR_SIGNALING_24HR_UP | 80 | 0.55962807 | 1.542381 | 0 | 0.1418071 | 1 | 1532 | tags=25%, list=6%, signal=26% |
| GO_CELL_SURFACE | 673 | 0.47627673 | 1.5422891 | 0 | 0.18396106 | 1 | 3514 | tags=19%, list=13%, signal=22% |
| HP_CONFUSION | 48 | 0.59635794 | 1.5422697 | 0 | 0.18365304 | 1 | 2360 | tags=27%, list=9%, signal=30% |
| TAKEDA_TARGETS_OF_NUP98_HOXA9_FUSION_3D_UP | 146 | 0.5472332 | 1.5418997 | 0 | 0.14230798 | 1 | 2703 | tags=28%, list=10%, signal=31% |
| KEGG_AMINO_SUGAR_AND_NUCLEOTIDE_SUGAR_METABOLISM | 40 | 0.6776639 | 1.5417961 | 0.025 | 0.14214666 | 1 | 3318 | tags=55%, list=12%, signal=63% |
| GO_DETECTION_OF_CHEMICAL_STIMULUS_INVOLVED_IN_SENSORY_PERCEPTION_OF_TASTE | 14 | 0.86265177 | 1.5417875 | 0 | 0.18446024 | 1 | 1662 | tags=14%, list=6%, signal=15% |
| GO_REGULATION_OF_RESPONSE_TO_EXTRACELLULAR_STIMULUS | 22 | 0.7225929 | 1.5414581 | 0.02564103 | 0.18491212 | 1 | 333 | tags=9%, list=1%, signal=9% |
| GO_PROTEIN_BINDING_INVOLVED_IN_HETEROTYPIC_CELL_CELL_ADHESION | 11 | 0.79191643 | 1.541429 | 0.02564103 | 0.18463333 | 1 | 893 | tags=18%, list=3%, signal=19% |
| GARCIA_TARGETS_OF_FLI1_AND_DAX1_UP | 46 | 0.64083076 | 1.5408512 | 0.02702703 | 0.14326459 | 1 | 4832 | tags=63%, list=18%, signal=77% |
| NAKAMURA_TUMOR_ZONE_PERIPHERAL_VS_CENTRAL_UP | 257 | 0.48854452 | 1.5408486 | 0 | 0.14286329 | 1 | 2938 | tags=30%, list=11%, signal=34% |
| HP_ABNORMALITY_OF_THE_AXILLARY_HAIR | 38 | 0.6577249 | 1.5407238 | 0 | 0.18616392 | 1 | 2807 | tags=26%, list=10%, signal=29% |
| GO_REGULATION_OF_PHOSPHOLIPASE_C_ACTIVITY | 40 | 0.6701753 | 1.5405408 | 0.02777778 | 0.18626127 | 1 | 2728 | tags=30%, list=10%, signal=33% |
| GO_STRUCTURAL_CONSTITUENT_OF_MUSCLE | 33 | 0.6933672 | 1.5402019 | 0.0212766 | 0.18691169 | 1 | 3292 | tags=33%, list=12%, signal=38% |
| REACTOME_AUF1_HNRNP_D0_BINDS_AND_DESTABILIZES_MRNA | 52 | 0.6164962 | 1.5401702 | 0 | 0.14368953 | 1 | 805 | tags=38%, list=3%, signal=40% |
| GO_POSITIVE_REGULATION_OF_ACTIVATION_OF_JANUS_KINASE_ACTIVITY | 4 | 0.9680952 | 1.5401651 | 0 | 0.18668935 | 1 | 665 | tags=25%, list=2%, signal=26% |
| HP_ABNORMAL_THROMBOCYTE_MORPHOLOGY | 308 | 0.48205638 | 1.5398633 | 0 | 0.18688363 | 1 | 3535 | tags=30%, list=13%, signal=34% |
| WP_ONE_CARBON_METABOLISM_AND_RELATED_PATHWAYS | 48 | 0.6100405 | 1.5395528 | 0.02564103 | 0.14433043 | 1 | 2750 | tags=31%, list=10%, signal=35% |
| GO_HEAD_DEVELOPMENT | 723 | 0.46610454 | 1.539525 | 0 | 0.18732508 | 1 | 2371 | tags=19%, list=9%, signal=21% |
| CREIGHTON_ENDOCRINE_THERAPY_RESISTANCE_5 | 432 | 0.47762266 | 1.5395228 | 0 | 0.14397681 | 1 | 3409 | tags=30%, list=13%, signal=34% |
| HP_RENAL_TUBULAR_ACIDOSIS | 40 | 0.6182271 | 1.5392098 | 0.02325581 | 0.18767884 | 1 | 2883 | tags=33%, list=11%, signal=36% |
| MCGARVEY_SILENCED_BY_METHYLATION_IN_COLON_CANCER | 33 | 0.6611031 | 1.5391027 | 0.02439024 | 0.14418182 | 1 | 4000 | tags=39%, list=15%, signal=46% |
| DURCHDEWALD_SKIN_CARCINOGENESIS_UP | 69 | 0.6040602 | 1.5389944 | 0.03846154 | 0.14387618 | 1 | 3433 | tags=39%, list=13%, signal=45% |
| COLDREN_GEFITINIB_RESISTANCE_DN | 180 | 0.52036285 | 1.538901 | 0 | 0.1437135 | 1 | 4414 | tags=28%, list=16%, signal=33% |
| KONDO_EZH2_TARGETS | 186 | 0.48851955 | 1.5387784 | 0 | 0.14355259 | 1 | 4475 | tags=37%, list=17%, signal=44% |
| HP_DECREASED_BODY_WEIGHT | 1063 | 0.45686296 | 1.5385425 | 0 | 0.18889692 | 1 | 3535 | tags=26%, list=13%, signal=29% |
| GO_REGULATION_OF_MYOBLAST_DIFFERENTIATION | 47 | 0.6598832 | 1.5382831 | 0.02702703 | 0.18927631 | 1 | 2587 | tags=26%, list=10%, signal=28% |
| HP_TITUBATION | 11 | 0.81117743 | 1.5382477 | 0.02325581 | 0.18902415 | 1 | 900 | tags=18%, list=3%, signal=19% |
| NELSON_RESPONSE_TO_ANDROGEN_UP | 73 | 0.57667804 | 1.5380694 | 0 | 0.1439003 | 1 | 2438 | tags=32%, list=9%, signal=35% |
| GO_SPHINGOMYELIN_METABOLIC_PROCESS | 12 | 0.77805054 | 1.5379189 | 0 | 0.18951662 | 1 | 5336 | tags=83%, list=20%, signal=104% |
| GO_AV_NODE_CELL_TO_BUNDLE_OF_HIS_CELL_COMMUNICATION | 8 | 0.90283847 | 1.5378605 | 0 | 0.18929325 | 1 | 234 | tags=13%, list=1%, signal=13% |
| TSENG_IRS1_TARGETS_DN | 121 | 0.5472344 | 1.5378602 | 0.03448276 | 0.14373985 | 1 | 2915 | tags=32%, list=11%, signal=36% |
| NAKAMURA_ADIPOGENESIS_EARLY_DN | 32 | 0.6786673 | 1.5377693 | 0 | 0.14357868 | 1 | 2882 | tags=31%, list=11%, signal=35% |
| BLANCO_MELO_MERS_COV_INFECTION_MCR5_CELLS_DN | 25 | 0.7227197 | 1.5377357 | 0.04255319 | 0.14323379 | 1 | 1026 | tags=28%, list=4%, signal=29% |
| GO_U1_SNRNP_BINDING | 4 | 0.9585346 | 1.5375948 | 0.04166667 | 0.1892416 | 1 | 1119 | tags=100%, list=4%, signal=104% |
| GO_PROTEIN_CONTAINING_COMPLEX_DISASSEMBLY | 301 | 0.5191197 | 1.5375568 | 0 | 0.1890201 | 1 | 2481 | tags=32%, list=9%, signal=35% |
| HP_DIARRHEA | 258 | 0.5075667 | 1.537359 | 0 | 0.18908285 | 1 | 3151 | tags=22%, list=12%, signal=25% |
| GO_HEART_TRABECULA_FORMATION | 11 | 0.838733 | 1.5371128 | 0.04444445 | 0.18925758 | 1 | 249 | tags=18%, list=1%, signal=18% |
| HP_MYOPATHY | 262 | 0.496456 | 1.5366805 | 0 | 0.19022419 | 1 | 2434 | tags=20%, list=9%, signal=22% |
| DURAND_STROMA_S_UP | 280 | 0.4836148 | 1.5361729 | 0 | 0.14472102 | 1 | 3003 | tags=24%, list=11%, signal=26% |
| GO_HYALURONAN_CATABOLIC_PROCESS | 13 | 0.8259015 | 1.5360621 | 0 | 0.19141147 | 1 | 2998 | tags=54%, list=11%, signal=61% |
| HP_MUSCLE_FIBER_INCLUSION_BODIES | 26 | 0.6909888 | 1.536061 | 0 | 0.19107448 | 1 | 1829 | tags=23%, list=7%, signal=25% |
| HP_SHOULDER_GIRDLE_MUSCLE_ATROPHY | 10 | 0.8816364 | 1.5359497 | 0 | 0.19090842 | 1 | 1829 | tags=60%, list=7%, signal=64% |
| GO_RESPONSE_TO_OXYGEN_RADICAL | 27 | 0.67741215 | 1.5357435 | 0.02173913 | 0.19099435 | 1 | 2750 | tags=37%, list=10%, signal=41% |
| HP_UPPER_MOTOR_NEURON_DYSFUNCTION | 1023 | 0.460408 | 1.5356251 | 0 | 0.19091225 | 1 | 4106 | tags=34%, list=15%, signal=39% |
| GO_REGULATION_OF_DEFENSE_RESPONSE | 571 | 0.4631856 | 1.5354793 | 0 | 0.19097096 | 1 | 2479 | tags=19%, list=9%, signal=21% |
| GO_REGULATION_OF_INTEGRIN_MEDIATED_SIGNALING_PATHWAY | 13 | 0.8105374 | 1.5354515 | 0.01923077 | 0.19072196 | 1 | 2030 | tags=54%, list=8%, signal=58% |
| GO_NEGATIVE_REGULATION_OF_INTRACELLULAR_PROTEIN_TRANSPORT | 41 | 0.6512502 | 1.53542 | 0 | 0.1905576 | 1 | 2317 | tags=32%, list=9%, signal=35% |
| HILLION_HMGA1B_TARGETS | 80 | 0.5746921 | 1.5353124 | 0 | 0.14578961 | 1 | 1939 | tags=36%, list=7%, signal=39% |
| GO_REGULATION_OF_APPETITE | 18 | 0.7715699 | 1.5348517 | 0 | 0.19187173 | 1 | 1 | tags=6%, list=0%, signal=6% |
| GO_NEGATIVE_REGULATION_OF_PROTEIN_LOCALIZATION_TO_NUCLEUS | 27 | 0.6870405 | 1.5348109 | 0.03225806 | 0.19165097 | 1 | 2857 | tags=37%, list=11%, signal=41% |
| LANDIS_ERBB2_BREAST_TUMORS_324_UP | 134 | 0.5628994 | 1.5347869 | 0 | 0.14635509 | 1 | 2438 | tags=34%, list=9%, signal=38% |
| GO_RESPONSE_TO_DRUG | 346 | 0.4932418 | 1.5344391 | 0 | 0.19220759 | 1 | 2729 | tags=21%, list=10%, signal=23% |
| WP_PROSTAGLANDIN_SYNTHESIS_AND_REGULATION | 37 | 0.669109 | 1.5341462 | 0 | 0.14669096 | 1 | 2768 | tags=27%, list=10%, signal=30% |
| GO_KIDNEY_MORPHOGENESIS | 83 | 0.5822778 | 1.5340806 | 0 | 0.192345 | 1 | 2160 | tags=18%, list=8%, signal=20% |
| GO_CALCIUM_ION_BINDING | 591 | 0.4704319 | 1.5340749 | 0 | 0.19201337 | 1 | 3905 | tags=25%, list=15%, signal=28% |
| REACTOME_TRANSCRIPTIONAL_REGULATION_BY_RUNX2 | 115 | 0.56143194 | 1.534064 | 0 | 0.14643222 | 1 | 2396 | tags=32%, list=9%, signal=35% |
| GO_QUINONE_BINDING | 15 | 0.81867504 | 1.5339341 | 0 | 0.19201407 | 1 | 707 | tags=33%, list=3%, signal=34% |
| CUI_TCF21_TARGETS_UP | 32 | 0.71433157 | 1.5336423 | 0.03125 | 0.14663105 | 1 | 802 | tags=25%, list=3%, signal=26% |
| GO_THREONINE_METABOLIC_PROCESS | 5 | 0.9245555 | 1.5335327 | 0.04651163 | 0.19264738 | 1 | 1126 | tags=40%, list=4%, signal=42% |
| MIKKELSEN_PLURIPOTENT_STATE_DN | 7 | 0.92736405 | 1.5333775 | 0 | 0.14655739 | 1 | 156 | tags=29%, list=1%, signal=29% |
| HP_ABNORMAL_ERYTHROCYTE_MORPHOLOGY | 497 | 0.47635806 | 1.5332946 | 0 | 0.19283761 | 1 | 3151 | tags=27%, list=12%, signal=30% |
| GO_TISSUE_REGENERATION | 57 | 0.61975104 | 1.533154 | 0 | 0.19289234 | 1 | 3281 | tags=32%, list=12%, signal=36% |
| SPIELMAN_LYMPHOBLAST_EUROPEAN_VS_ASIAN_DN | 562 | 0.47301105 | 1.5328488 | 0 | 0.14666776 | 1 | 3002 | tags=32%, list=11%, signal=35% |
| OUELLET_CULTURED_OVARIAN_CANCER_INVASIVE_VS_LMP_UP | 62 | 0.5816631 | 1.532656 | 0.03030303 | 0.14672948 | 1 | 2515 | tags=37%, list=9%, signal=41% |
| GO_POSITIVE_REGULATION_OF_RESPONSE_TO_EXTRACELLULAR_STIMULUS | 4 | 0.94370586 | 1.5325506 | 0 | 0.1940413 | 1 | 333 | tags=50%, list=1%, signal=51% |
| SATO_SILENCED_BY_METHYLATION_IN_PANCREATIC_CANCER_1 | 324 | 0.48852342 | 1.5324875 | 0 | 0.14683515 | 1 | 2141 | tags=16%, list=8%, signal=17% |
| HP_CONJUNCTIVAL_HAMARTOMA | 9 | 0.78357315 | 1.5323958 | 0 | 0.19414718 | 1 | 906 | tags=44%, list=3%, signal=46% |
| DEMAGALHAES_AGING_UP | 51 | 0.6237404 | 1.5322863 | 0 | 0.1467159 | 1 | 2204 | tags=31%, list=8%, signal=34% |
| GUILLAUMOND_KLF10_TARGETS_UP | 41 | 0.6422599 | 1.5322734 | 0.02702703 | 0.14632979 | 1 | 3432 | tags=44%, list=13%, signal=50% |
| REACTOME_SCF_SKP2_MEDIATED_DEGRADATION_OF_P27_P21 | 57 | 0.6097829 | 1.5322415 | 0 | 0.14599104 | 1 | 2352 | tags=47%, list=9%, signal=52% |
| GO_TRANSMEMBRANE_TRANSPORT | 1344 | 0.43895373 | 1.5319802 | 0 | 0.19480109 | 1 | 3440 | tags=21%, list=13%, signal=23% |
| GO_DEFENSE_RESPONSE_TO_OTHER_ORGANISM | 812 | 0.46326157 | 1.5319233 | 0 | 0.19460616 | 1 | 2378 | tags=19%, list=9%, signal=21% |
| GO_POSITIVE_REGULATION_OF_PHOSPHOLIPID_METABOLIC_PROCESS | 44 | 0.6459524 | 1.5317262 | 0 | 0.19492866 | 1 | 2476 | tags=23%, list=9%, signal=25% |
| HP_ABNORMAL_BREATH_SOUND | 42 | 0.6530053 | 1.5316005 | 0 | 0.19478814 | 1 | 2302 | tags=17%, list=9%, signal=18% |
| GO_NEGATIVE_REGULATION_OF_STEM_CELL_PROLIFERATION | 9 | 0.89155424 | 1.5312337 | 0.02380952 | 0.19535276 | 1 | 84 | tags=22%, list=0%, signal=22% |
| GO_LATE_ENDOSOME_TO_VACUOLE_TRANSPORT_VIA_MULTIVESICULAR_BODY_SORTING_PATHWAY | 13 | 0.8071733 | 1.5312275 | 0.04081633 | 0.19502278 | 1 | 1230 | tags=46%, list=5%, signal=48% |
| YOSHIMURA_MAPK8_TARGETS_DN | 345 | 0.48958576 | 1.5311253 | 0 | 0.14688596 | 1 | 2187 | tags=26%, list=8%, signal=28% |
| HP_PARAPLEGIA_PARAPARESIS | 194 | 0.53441536 | 1.5308415 | 0 | 0.19564043 | 1 | 2673 | tags=26%, list=10%, signal=28% |
| HIRSCH_CELLULAR_TRANSFORMATION_SIGNATURE_UP | 228 | 0.49658912 | 1.5305935 | 0 | 0.14760542 | 1 | 2470 | tags=25%, list=9%, signal=28% |
| GO_SPHERICAL_HIGH_DENSITY_LIPOPROTEIN_PARTICLE | 4 | 0.97257835 | 1.5305338 | 0.02040816 | 0.19590384 | 1 | 60 | tags=25%, list=0%, signal=25% |
| KEGG_RNA_POLYMERASE | 26 | 0.7166671 | 1.5302365 | 0.04 | 0.14779115 | 1 | 2758 | tags=54%, list=10%, signal=60% |
| GO_ANTIGEN_PROCESSING_AND_PRESENTATION_OF_ENDOGENOUS_ANTIGEN | 13 | 0.86479455 | 1.5298558 | 0 | 0.197407 | 1 | 2332 | tags=62%, list=9%, signal=67% |
| HP_VESTIBULAR_DYSFUNCTION | 110 | 0.55579877 | 1.52979 | 0 | 0.19721094 | 1 | 1999 | tags=21%, list=7%, signal=22% |
| HP_ABNORMALITY_OF_THE_URINARY_SYSTEM_PHYSIOLOGY | 789 | 0.46195182 | 1.5297688 | 0 | 0.19690795 | 1 | 3535 | tags=27%, list=13%, signal=30% |
| HP_ABNORMALITY_OF_THE_INNER_EAR | 776 | 0.45463023 | 1.5296463 | 0 | 0.19700691 | 1 | 3484 | tags=26%, list=13%, signal=29% |
| BLANCO_MELO_HUMAN_PARAINFLUENZA_VIRUS_3_INFECTION_A594_CELLS_UP | 167 | 0.51155573 | 1.5296168 | 0 | 0.14846082 | 1 | 2629 | tags=27%, list=10%, signal=30% |
| GO_TRNA_PROCESSING | 118 | 0.5364153 | 1.5295309 | 0 | 0.19694564 | 1 | 4137 | tags=48%, list=15%, signal=57% |
| HOLLERN_SOLID_NODULAR_BREAST_TUMOR_DN | 29 | 0.6774677 | 1.5294017 | 0.02631579 | 0.14847162 | 1 | 1012 | tags=17%, list=4%, signal=18% |
| GO_CEREBELLAR_GRANULAR_LAYER_MORPHOGENESIS | 8 | 0.8844376 | 1.5291402 | 0.04166667 | 0.19736129 | 1 | 1945 | tags=25%, list=7%, signal=27% |
| GO_ENDOSOME_LUMEN | 24 | 0.75090766 | 1.5289584 | 0.02040816 | 0.19762027 | 1 | 2661 | tags=38%, list=10%, signal=42% |
| RODWELL_AGING_KIDNEY_NO_BLOOD_DN | 128 | 0.5396341 | 1.5287547 | 0 | 0.14944603 | 1 | 2353 | tags=25%, list=9%, signal=27% |
| GO_REGULATION_OF_T_CELL_MIGRATION | 35 | 0.67878866 | 1.528397 | 0.03030303 | 0.19872819 | 1 | 3703 | tags=34%, list=14%, signal=40% |
| GO_NEGATIVE_REGULATION_OF_PROTEIN_CONTAINING_COMPLEX_ASSEMBLY | 118 | 0.527265 | 1.5281721 | 0 | 0.19932903 | 1 | 2283 | tags=25%, list=8%, signal=27% |
| LENAOUR_DENDRITIC_CELL_MATURATION_UP | 101 | 0.5502284 | 1.5280871 | 0 | 0.15023729 | 1 | 1108 | tags=19%, list=4%, signal=20% |
| HP_ABNORMAL_GLYCOSYLATION | 36 | 0.636654 | 1.5280353 | 0 | 0.199451 | 1 | 3527 | tags=44%, list=13%, signal=51% |
| GO_NON_CANONICAL_WNT_SIGNALING_PATHWAY | 147 | 0.54618543 | 1.5280259 | 0 | 0.199148 | 1 | 2746 | tags=27%, list=10%, signal=30% |
| HP_RECURRENT_PANCREATITIS | 9 | 0.92613626 | 1.527969 | 0.02702703 | 0.19895187 | 1 | 854 | tags=56%, list=3%, signal=57% |
| DELASERNA_TARGETS_OF_MYOD_AND_SMARCA4 | 9 | 0.8765215 | 1.5278263 | 0.02631579 | 0.15015402 | 1 | 732 | tags=22%, list=3%, signal=23% |
| GO_RESPIRATORY_CHAIN_COMPLEX_IV | 17 | 0.78304535 | 1.5277182 | 0.02173913 | 0.19928385 | 1 | 1740 | tags=41%, list=6%, signal=44% |
| HP_ABNORMALITY_OF_THE_OPTIC_NERVE | 648 | 0.46948114 | 1.5275542 | 0 | 0.19937937 | 1 | 3526 | tags=28%, list=13%, signal=32% |
| SENGUPTA_NASOPHARYNGEAL_CARCINOMA_WITH_LMP1_DN | 111 | 0.55998355 | 1.5273768 | 0 | 0.15037563 | 1 | 1796 | tags=9%, list=7%, signal=10% |
| GO_INTERLEUKIN_1_MEDIATED_SIGNALING_PATHWAY | 96 | 0.5394869 | 1.5273029 | 0 | 0.19948602 | 1 | 805 | tags=22%, list=3%, signal=22% |
| ZHAN_V2_LATE_DIFFERENTIATION_GENES | 41 | 0.6577213 | 1.5273002 | 0 | 0.15007931 | 1 | 4189 | tags=56%, list=16%, signal=66% |
| MARIADASON_RESPONSE_TO_BUTYRATE_CURCUMIN_SULINDAC_TSA_1 | 5 | 0.916367 | 1.5272161 | 0.02222222 | 0.14991072 | 1 | 300 | tags=40%, list=1%, signal=40% |
| WHITEHURST_PACLITAXEL_SENSITIVITY | 32 | 0.6929202 | 1.5271558 | 0 | 0.14961472 | 1 | 1302 | tags=19%, list=5%, signal=20% |
| GO_OXIDOREDUCTASE_ACTIVITY_ACTING_ON_DIPHENOLS_AND_RELATED_SUBSTANCES_AS_DONORS | 7 | 0.95968485 | 1.5270795 | 0 | 0.19968456 | 1 | 1036 | tags=86%, list=4%, signal=89% |
| GENTILE_UV_HIGH_DOSE_UP | 17 | 0.78863114 | 1.527006 | 0.02222222 | 0.14911233 | 1 | 855 | tags=24%, list=3%, signal=24% |
| CHIANG_LIVER_CANCER_SUBCLASS_INTERFERON_UP | 20 | 0.7569856 | 1.5269752 | 0.04651163 | 0.14873579 | 1 | 1751 | tags=30%, list=7%, signal=32% |
| GO_IMMUNE_EFFECTOR_PROCESS | 986 | 0.454929 | 1.5269167 | 0 | 0.19985527 | 1 | 2662 | tags=22%, list=10%, signal=23% |
| HP_CEREBRAL_VISUAL_IMPAIRMENT | 126 | 0.5304684 | 1.5265703 | 0 | 0.20028764 | 1 | 2316 | tags=21%, list=9%, signal=22% |
| GO_TRNA_MODIFICATION | 79 | 0.5478495 | 1.5261644 | 0 | 0.20066607 | 1 | 4134 | tags=48%, list=15%, signal=57% |
| ACEVEDO_LIVER_CANCER_UP | 886 | 0.46137246 | 1.5258802 | 0 | 0.14989409 | 1 | 3555 | tags=35%, list=13%, signal=39% |
| GO_CELL_DEATH_IN_RESPONSE_TO_HYDROGEN_PEROXIDE | 24 | 0.685044 | 1.5258375 | 0 | 0.20130232 | 1 | 1821 | tags=25%, list=7%, signal=27% |
| GO_NEGATIVE_REGULATION_OF_ATP_METABOLIC_PROCESS | 25 | 0.71532494 | 1.5256723 | 0 | 0.20146835 | 1 | 2089 | tags=32%, list=8%, signal=35% |
| CADWELL_ATG16L1_TARGETS_UP | 79 | 0.5696005 | 1.5254611 | 0 | 0.15032549 | 1 | 645 | tags=10%, list=2%, signal=10% |
| GO_MORPHOGENESIS_OF_AN_EPITHELIAL_BUD | 13 | 0.80424607 | 1.5254606 | 0.04347826 | 0.20166272 | 1 | 627 | tags=15%, list=2%, signal=16% |
| GO_DEFENSE_RESPONSE | 1257 | 0.44842118 | 1.5252224 | 0 | 0.20206262 | 1 | 2494 | tags=18%, list=9%, signal=18% |
| GO_M_BAND | 20 | 0.7415393 | 1.5251925 | 0 | 0.20186503 | 1 | 1912 | tags=20%, list=7%, signal=22% |
| GO_TYPE_I_INTERFERON_PRODUCTION | 113 | 0.54241323 | 1.525184 | 0 | 0.20159161 | 1 | 2468 | tags=31%, list=9%, signal=34% |
| LEI_MYB_TARGETS | 289 | 0.5024794 | 1.5250583 | 0 | 0.15049817 | 1 | 2450 | tags=28%, list=9%, signal=31% |
| GO_CELLULAR_RESPONSE_TO_OXYGEN_RADICAL | 25 | 0.6767746 | 1.5249228 | 0.02083333 | 0.20183414 | 1 | 2750 | tags=36%, list=10%, signal=40% |
| REACTOME_ORC1_REMOVAL_FROM_CHROMATIN | 67 | 0.60110605 | 1.52488 | 0 | 0.15054353 | 1 | 2352 | tags=42%, list=9%, signal=46% |
| HP_ABNORMALITY_OF_BLOOD_AND_BLOOD_FORMING_TISSUES | 1044 | 0.45713046 | 1.5248461 | 0 | 0.20169103 | 1 | 3558 | tags=27%, list=13%, signal=30% |
| HP_DIMINISHED_MOVEMENT | 53 | 0.61082184 | 1.524675 | 0 | 0.20172556 | 1 | 2158 | tags=23%, list=8%, signal=25% |
| HP_NECK_MUSCLE_WEAKNESS | 58 | 0.5985423 | 1.524542 | 0.03030303 | 0.20181341 | 1 | 4184 | tags=31%, list=16%, signal=37% |
| BLANCO_MELO_INFLUENZA_A_INFECTION_A594_CELLS_UP | 34 | 0.6283546 | 1.5245203 | 0 | 0.15067227 | 1 | 2757 | tags=38%, list=10%, signal=43% |
| GO_PIGMENTATION | 91 | 0.51502055 | 1.5242546 | 0 | 0.20218465 | 1 | 2779 | tags=22%, list=10%, signal=24% |
| REACTOME_DNA_REPLICATION_PRE_INITIATION | 81 | 0.5805303 | 1.5237143 | 0 | 0.15151371 | 1 | 2352 | tags=38%, list=9%, signal=42% |
| GO_ATPASE_COUPLED_ION_TRANSMEMBRANE_TRANSPORTER_ACTIVITY | 55 | 0.6018213 | 1.5235866 | 0 | 0.20362754 | 1 | 1206 | tags=22%, list=4%, signal=23% |
| GO_STEM_CELL_DIFFERENTIATION | 233 | 0.51389205 | 1.5234728 | 0 | 0.20355794 | 1 | 2677 | tags=25%, list=10%, signal=27% |
| GO_RESPONSE_TO_ETHANOL | 102 | 0.53177947 | 1.5233247 | 0 | 0.20324247 | 1 | 2729 | tags=24%, list=10%, signal=26% |
| GO_BIOLOGICAL_PHASE | 6 | 0.90244615 | 1.5232358 | 0.02702703 | 0.20314865 | 1 | 2396 | tags=50%, list=9%, signal=55% |
| GO_FATTY_ACID_BETA_OXIDATION_USING_ACYL_COA_DEHYDROGENASE | 9 | 0.8229568 | 1.523108 | 0 | 0.20310518 | 1 | 3506 | tags=78%, list=13%, signal=89% |
| GO_RESPONSE_TO_BIOTIC_STIMULUS | 1120 | 0.45015085 | 1.5229411 | 0 | 0.20281711 | 1 | 2378 | tags=19%, list=9%, signal=20% |
| FRASOR_RESPONSE_TO_ESTRADIOL_DN | 69 | 0.5758253 | 1.5229138 | 0.03030303 | 0.1522742 | 1 | 2999 | tags=30%, list=11%, signal=34% |
| REACTOME_INTERLEUKIN_35_SIGNALLING | 11 | 0.8401379 | 1.5223628 | 0.02380952 | 0.15277512 | 1 | 1939 | tags=27%, list=7%, signal=29% |
| GO_MHC_CLASS_I_PEPTIDE_LOADING_COMPLEX | 6 | 0.9297288 | 1.5223544 | 0 | 0.20421547 | 1 | 1315 | tags=67%, list=5%, signal=70% |
| HELLEBREKERS_SILENCED_DURING_TUMOR_ANGIOGENESIS | 70 | 0.634912 | 1.5222045 | 0 | 0.15260528 | 1 | 2685 | tags=33%, list=10%, signal=36% |
| HP_ABNORMAL_CORNEAL_EPITHELIUM_MORPHOLOGY | 80 | 0.58662724 | 1.5221288 | 0.03448276 | 0.20464887 | 1 | 1967 | tags=15%, list=7%, signal=16% |
| GO_NEGATIVE_REGULATION_OF_PLATELET_AGGREGATION | 9 | 0.8393525 | 1.5219344 | 0 | 0.20465448 | 1 | 312 | tags=22%, list=1%, signal=22% |
| OISHI_CHOLANGIOMA_STEM_CELL_LIKE_UP | 302 | 0.4955352 | 1.5218759 | 0 | 0.15285535 | 1 | 3876 | tags=36%, list=14%, signal=41% |
| REACTOME_SIGNALING_BY_HEDGEHOG | 140 | 0.53458506 | 1.521826 | 0 | 0.1525644 | 1 | 2352 | tags=28%, list=9%, signal=30% |
| GO_THYROID_HORMONE_METABOLIC_PROCESS | 20 | 0.77805 | 1.5216013 | 0 | 0.20488583 | 1 | 1197 | tags=15%, list=4%, signal=16% |
| GO_FATTY_ACID_METABOLIC_PROCESS | 316 | 0.4801365 | 1.5210911 | 0 | 0.20602065 | 1 | 3918 | tags=32%, list=15%, signal=37% |
| GO_INNER_EAR_RECEPTOR_CELL_STEREOCILIUM_ORGANIZATION | 26 | 0.6905406 | 1.5204498 | 0.02325581 | 0.20695107 | 1 | 1601 | tags=23%, list=6%, signal=25% |
| HP_ABNORMAL_PERIPHERAL_NERVOUS_SYSTEM_MORPHOLOGY | 746 | 0.47069418 | 1.5201505 | 0 | 0.20755453 | 1 | 3165 | tags=26%, list=12%, signal=28% |
| SCHAEFFER_SOX9_TARGETS_IN_PROSTATE_DEVELOPMENT_UP | 16 | 0.7742927 | 1.5199213 | 0.04347826 | 0.15541609 | 1 | 1094 | tags=19%, list=4%, signal=20% |
| HP_ELEVATED_DIASTOLIC_BLOOD_PRESSURE | 10 | 0.8172825 | 1.5192178 | 0 | 0.20945582 | 1 | 2530 | tags=30%, list=9%, signal=33% |
| GO_METANEPHRIC_TUBULE_MORPHOGENESIS | 9 | 0.8081129 | 1.5192018 | 0.04255319 | 0.20915467 | 1 | 267 | tags=33%, list=1%, signal=34% |
| BLANCO_MELO_BETA_INTERFERON_TREATED_BRONCHIAL_EPITHELIAL_CELLS_UP | 322 | 0.48618767 | 1.5188472 | 0 | 0.15644157 | 1 | 3924 | tags=31%, list=15%, signal=36% |
| GO_POSITIVE_REGULATION_OF_DEFENSE_RESPONSE | 316 | 0.48679954 | 1.5188082 | 0 | 0.20985378 | 1 | 2629 | tags=20%, list=10%, signal=21% |
| NATSUME_RESPONSE_TO_INTERFERON_BETA_UP | 64 | 0.60432094 | 1.5187864 | 0 | 0.15626624 | 1 | 1700 | tags=28%, list=6%, signal=30% |
| KIM_ALL_DISORDERS_OLIGODENDROCYTE_NUMBER_CORR_UP | 723 | 0.46496883 | 1.5187067 | 0 | 0.15609255 | 1 | 2447 | tags=30%, list=9%, signal=32% |
| GO_INSEMINATION | 7 | 0.92494744 | 1.5186 | 0 | 0.21019807 | 1 | 17 | tags=14%, list=0%, signal=14% |
| GO_REGULATION_OF_MESENCHYMAL_TO_EPITHELIAL_TRANSITION_INVOLVED_IN_METANEPHROS_MORPHOGENESIS | 6 | 0.9559761 | 1.5183849 | 0 | 0.21049105 | 1 | 85 | tags=33%, list=0%, signal=33% |
| LINDSTEDT_DENDRITIC_CELL_MATURATION_C | 58 | 0.6191581 | 1.5183712 | 0 | 0.15624467 | 1 | 2940 | tags=34%, list=11%, signal=39% |
| HP_CEREBRAL_DYSMYELINATION | 13 | 0.8185368 | 1.518266 | 0.02040816 | 0.21056193 | 1 | 1776 | tags=38%, list=7%, signal=41% |
| GO_POSITIVE_REGULATION_OF_IMMUNE_RESPONSE | 532 | 0.47666937 | 1.5182287 | 0 | 0.21028554 | 1 | 2510 | tags=20%, list=9%, signal=21% |
| KUMAR_TARGETS_OF_MLL_AF9_FUSION | 356 | 0.50306225 | 1.5180424 | 0 | 0.15644129 | 1 | 2589 | tags=24%, list=10%, signal=26% |
| GO_ENDOPLASMIC_RETICULUM | 1219 | 0.44289523 | 1.517746 | 0 | 0.2107233 | 1 | 3627 | tags=30%, list=13%, signal=33% |
| GO_ENDOCYTIC_VESICLE | 273 | 0.49445036 | 1.5172397 | 0 | 0.21170598 | 1 | 2586 | tags=25%, list=10%, signal=27% |
| LIANG_SILENCED_BY_METHYLATION_2 | 35 | 0.70556563 | 1.5171748 | 0 | 0.15711623 | 1 | 2141 | tags=34%, list=8%, signal=37% |
| GO_REGULATION_OF_LYSOSOMAL_LUMEN_PH | 12 | 0.7968278 | 1.51696 | 0.04878049 | 0.21224117 | 1 | 1952 | tags=42%, list=7%, signal=45% |
| TONKS_TARGETS_OF_RUNX1_RUNX1T1_FUSION_MONOCYTE_UP | 192 | 0.51543707 | 1.5169207 | 0 | 0.15718555 | 1 | 3069 | tags=34%, list=11%, signal=38% |
| GO_STRUCTURAL_CONSTITUENT_OF_MYELIN_SHEATH | 9 | 0.8632535 | 1.5167054 | 0.025 | 0.21260275 | 1 | 0 | tags=11%, list=0%, signal=11% |
| HP_ABNORMAL_METABOLISM | 95 | 0.5581658 | 1.5162935 | 0 | 0.2135265 | 1 | 3779 | tags=37%, list=14%, signal=43% |
| HUANG_FOXA2_TARGETS_DN | 32 | 0.6851578 | 1.5162055 | 0 | 0.1579434 | 1 | 3027 | tags=34%, list=11%, signal=39% |
| ELLWOOD_MYC_TARGETS_DN | 38 | 0.64199746 | 1.5161643 | 0 | 0.15764698 | 1 | 1841 | tags=34%, list=7%, signal=37% |
| HP_HEAD_TREMOR | 24 | 0.74130535 | 1.5157089 | 0.04761905 | 0.21471843 | 1 | 2461 | tags=33%, list=9%, signal=37% |
| GO_RIBONUCLEOSIDE_TRIPHOSPHATE_BIOSYNTHETIC_PROCESS | 61 | 0.6074429 | 1.5156374 | 0.025 | 0.214587 | 1 | 1939 | tags=33%, list=7%, signal=35% |
| GO_PROTEIN_INSERTION_INTO_MITOCHONDRIAL_INNER_MEMBRANE | 6 | 0.94306725 | 1.5155977 | 0 | 0.214333 | 1 | 1016 | tags=83%, list=4%, signal=87% |
| HP_BRONCHITIS | 71 | 0.5791174 | 1.5155821 | 0 | 0.21400675 | 1 | 4398 | tags=24%, list=16%, signal=29% |
| GO_RESPONSE_TO_TUMOR_NECROSIS_FACTOR | 262 | 0.5012262 | 1.5155425 | 0 | 0.21380389 | 1 | 3224 | tags=28%, list=12%, signal=31% |
| NADERI_BREAST_CANCER_PROGNOSIS_DN | 14 | 0.74138623 | 1.515424 | 0.04761905 | 0.15875897 | 1 | 2493 | tags=43%, list=9%, signal=47% |
| HP_GENERALIZED_ABNORMALITY_OF_SKIN | 726 | 0.46301478 | 1.515423 | 0 | 0.21379621 | 1 | 3524 | tags=24%, list=13%, signal=27% |
| REACTOME_PROGRAMMED_CELL_DEATH | 167 | 0.5040064 | 1.515047 | 0 | 0.15858309 | 1 | 2006 | tags=28%, list=7%, signal=30% |
| GO_POSITIVE_REGULATION_OF_RESPONSE_TO_BIOTIC_STIMULUS | 215 | 0.5123914 | 1.5149103 | 0 | 0.21471108 | 1 | 1624 | tags=16%, list=6%, signal=17% |
| GO_COATED_VESICLE_MEMBRANE | 154 | 0.52298707 | 1.5148507 | 0 | 0.2146287 | 1 | 2301 | tags=28%, list=9%, signal=30% |
| GO_COA_HYDROLASE_ACTIVITY | 18 | 0.76381606 | 1.5148448 | 0.02631579 | 0.21432865 | 1 | 4954 | tags=56%, list=18%, signal=68% |
| HP_NEMALINE_BODIES | 14 | 0.7928971 | 1.5147831 | 0.0212766 | 0.21415044 | 1 | 1580 | tags=21%, list=6%, signal=23% |
| WAMUNYOKOLI_OVARIAN_CANCER_LMP_UP | 248 | 0.50044936 | 1.5146247 | 0 | 0.1592092 | 1 | 3318 | tags=31%, list=12%, signal=35% |
| GO_COPULATION | 13 | 0.7993026 | 1.5145906 | 0.04347826 | 0.21438365 | 1 | 592 | tags=15%, list=2%, signal=16% |
| HP_THORACIC_SCOLIOSIS | 37 | 0.68491465 | 1.5142848 | 0.05 | 0.21507198 | 1 | 1927 | tags=30%, list=7%, signal=32% |
| HP_IMPAIRED_OROPHARYNGEAL_SWALLOW_RESPONSE | 4 | 0.9392329 | 1.5141379 | 0 | 0.21503736 | 1 | 1419 | tags=75%, list=5%, signal=79% |
| SHIN_B_CELL_LYMPHOMA_CLUSTER_6 | 8 | 0.83876705 | 1.5139886 | 0.02631579 | 0.15952775 | 1 | 1236 | tags=25%, list=5%, signal=26% |
| GO_GAMMA_AMINOBUTYRIC_ACID_METABOLIC_PROCESS | 4 | 0.9546802 | 1.5135624 | 0 | 0.21632223 | 1 | 445 | tags=50%, list=2%, signal=51% |
| CHEN_HOXA5_TARGETS_6HR_UP | 7 | 0.87630427 | 1.5134015 | 0 | 0.16002901 | 1 | 2330 | tags=43%, list=9%, signal=47% |
| GO_REGULATION_OF_RESPONSE_TO_FOOD | 16 | 0.76352334 | 1.5130584 | 0 | 0.21724588 | 1 | 1 | tags=6%, list=0%, signal=6% |
| BOYAULT_LIVER_CANCER_SUBCLASS_G23_DN | 8 | 0.8936053 | 1.5130498 | 0.02857143 | 0.16016915 | 1 | 28 | tags=13%, list=0%, signal=13% |
| HP_BILATERAL_SENSORINEURAL_HEARING_IMPAIRMENT | 55 | 0.59849745 | 1.5130416 | 0 | 0.21699333 | 1 | 1888 | tags=24%, list=7%, signal=25% |
| REACTOME_INTERLEUKIN_6_SIGNALING | 10 | 0.8926085 | 1.5128766 | 0 | 0.16003357 | 1 | 1939 | tags=40%, list=7%, signal=43% |
| MILICIC_FAMILIAL_ADENOMATOUS_POLYPOSIS_DN | 7 | 0.90154785 | 1.5126454 | 0.02222222 | 0.16009496 | 1 | 2555 | tags=57%, list=9%, signal=63% |
| PID_INTEGRIN2_PATHWAY | 21 | 0.7436762 | 1.5115342 | 0.02083333 | 0.16161554 | 1 | 723 | tags=14%, list=3%, signal=15% |
| GO_PROTON_TRANSPORTING_V_TYPE_ATPASE_V0_DOMAIN | 9 | 0.8670922 | 1.5114949 | 0.02439024 | 0.22095518 | 1 | 625 | tags=22%, list=2%, signal=23% |
| GO_CELLULAR_RESPONSE_TO_NITROSATIVE_STRESS | 5 | 0.94147265 | 1.5113384 | 0 | 0.22105598 | 1 | 749 | tags=60%, list=3%, signal=62% |
| HAHTOLA_SEZARY_SYNDROM_UP | 87 | 0.5715868 | 1.5111352 | 0 | 0.16218859 | 1 | 2993 | tags=33%, list=11%, signal=37% |
| DACOSTA_UV_RESPONSE_VIA_ERCC3_UP | 288 | 0.4853788 | 1.511058 | 0 | 0.16189009 | 1 | 3088 | tags=35%, list=11%, signal=40% |
| HP_IRRITABILITY | 131 | 0.5322577 | 1.5109202 | 0 | 0.22177553 | 1 | 4320 | tags=42%, list=16%, signal=50% |
| GO_GLUTATHIONE_PEROXIDASE_ACTIVITY | 17 | 0.75945336 | 1.5109136 | 0.04761905 | 0.22147068 | 1 | 1997 | tags=47%, list=7%, signal=51% |
| REACTOME_APC_C_CDH1_MEDIATED_DEGRADATION_OF_CDC20_AND_OTHER_APC_C_CDH1_TARGETED_PROTEINS_IN_LATE_MITOSIS_EARLY_G1 | 71 | 0.5736279 | 1.5104012 | 0.02857143 | 0.16273773 | 1 | 2375 | tags=46%, list=9%, signal=51% |
| HP_PTERYGIUM | 31 | 0.6823915 | 1.5101104 | 0 | 0.22313258 | 1 | 4549 | tags=48%, list=17%, signal=58% |
| NUYTTEN_EZH2_TARGETS_UP | 934 | 0.4619442 | 1.5097643 | 0 | 0.16318095 | 1 | 3668 | tags=30%, list=14%, signal=34% |
| INGRAM_SHH_TARGETS_DN | 61 | 0.5910768 | 1.5091628 | 0 | 0.16420482 | 1 | 1167 | tags=16%, list=4%, signal=17% |
| HP_PAPILLEDEMA | 20 | 0.71093047 | 1.508878 | 0.04545455 | 0.22654192 | 1 | 2068 | tags=35%, list=8%, signal=38% |
| MARKS_HDAC_TARGETS_UP | 19 | 0.69129926 | 1.508867 | 0.02702703 | 0.16429543 | 1 | 1095 | tags=21%, list=4%, signal=22% |
| GALLUZZI_PERMEABILIZE_MITOCHONDRIA | 38 | 0.63552904 | 1.5087907 | 0.03448276 | 0.16403401 | 1 | 2267 | tags=37%, list=8%, signal=40% |
| HP_HYPOKALEMIC_METABOLIC_ALKALOSIS | 4 | 0.967771 | 1.5087079 | 0 | 0.22658557 | 1 | 161 | tags=25%, list=1%, signal=25% |
| GO_MITOCHONDRIAL_CYTOCHROME_C_OXIDASE_ASSEMBLY | 18 | 0.7099156 | 1.5080945 | 0.02272727 | 0.22790329 | 1 | 2622 | tags=44%, list=10%, signal=49% |
| ISSAEVA_MLL2_TARGETS | 54 | 0.59242713 | 1.5078537 | 0.03225806 | 0.16541415 | 1 | 3010 | tags=26%, list=11%, signal=29% |
| GO_RENAL_TUBULE_DEVELOPMENT | 83 | 0.56120193 | 1.5077968 | 0 | 0.22837017 | 1 | 2587 | tags=20%, list=10%, signal=23% |
| RUAN_RESPONSE_TO_TNF_DN | 71 | 0.5687534 | 1.5076712 | 0 | 0.16526766 | 1 | 3734 | tags=44%, list=14%, signal=51% |
| KEGG_CITRATE_CYCLE_TCA_CYCLE | 27 | 0.7200938 | 1.5076227 | 0.025 | 0.16489118 | 1 | 2982 | tags=59%, list=11%, signal=67% |
| GO_INTRAMOLECULAR_OXIDOREDUCTASE_ACTIVITY | 45 | 0.6260426 | 1.5076199 | 0.03333334 | 0.22841118 | 1 | 2617 | tags=42%, list=10%, signal=47% |
| GO_POSITIVE_REGULATION_BY_HOST_OF_VIRAL_TRANSCRIPTION | 16 | 0.725308 | 1.5074128 | 0.04878049 | 0.22866392 | 1 | 2610 | tags=63%, list=10%, signal=69% |
| GO_GOLGI_ASSOCIATED_VESICLE_MEMBRANE | 90 | 0.56556815 | 1.5073798 | 0 | 0.22835262 | 1 | 1237 | tags=27%, list=5%, signal=28% |
| GO_DEFENSE_RESPONSE_TO_VIRUS | 194 | 0.5005172 | 1.5072671 | 0 | 0.22851163 | 1 | 2479 | tags=28%, list=9%, signal=31% |
| REACTOME_REGULATION_OF_TLR_BY_ENDOGENOUS_LIGAND | 13 | 0.7966374 | 1.5071968 | 0.02439024 | 0.16525106 | 1 | 2006 | tags=31%, list=7%, signal=33% |
| KEGG_PATHOGENIC_ESCHERICHIA_COLI_INFECTION | 49 | 0.62041616 | 1.506984 | 0.0212766 | 0.16545013 | 1 | 2819 | tags=41%, list=10%, signal=46% |
| HP_VISUAL_LOSS | 185 | 0.499129 | 1.5063137 | 0 | 0.23074982 | 1 | 3128 | tags=24%, list=12%, signal=27% |
| GO_POSITIVE_REGULATION_OF_CELL_SUBSTRATE_ADHESION | 107 | 0.5445101 | 1.5061033 | 0 | 0.23097299 | 1 | 3120 | tags=30%, list=12%, signal=34% |
| GO_GALACTOSIDE_BINDING | 5 | 0.96993893 | 1.5060986 | 0 | 0.23063678 | 1 | 383 | tags=40%, list=1%, signal=41% |
| HP_PROGRESSIVE_SPASTICITY | 21 | 0.7442687 | 1.5060977 | 0.04545455 | 0.23030156 | 1 | 1776 | tags=24%, list=7%, signal=25% |
| KINSEY_TARGETS_OF_EWSR1_FLII_FUSION_DN | 302 | 0.4844997 | 1.5059452 | 0 | 0.16653013 | 1 | 3236 | tags=28%, list=12%, signal=31% |
| GO_NEGATIVE_REGULATION_OF_CELL_POPULATION_PROLIFERATION | 606 | 0.46149537 | 1.5059059 | 0 | 0.23047642 | 1 | 2587 | tags=20%, list=10%, signal=21% |
| GO_SECRETION | 1371 | 0.44154707 | 1.5056896 | 0 | 0.23104903 | 1 | 2889 | tags=20%, list=11%, signal=22% |
| GO_PROTEIN_N_LINKED_GLYCOSYLATION | 68 | 0.61313146 | 1.505317 | 0 | 0.23168795 | 1 | 3527 | tags=49%, list=13%, signal=56% |
| PETRETTO_HEART_MASS_QTL_CIS_UP | 29 | 0.6808932 | 1.5053089 | 0 | 0.16669175 | 1 | 2994 | tags=34%, list=11%, signal=39% |
| GO_EXOCYTOSIS | 816 | 0.45029423 | 1.5049899 | 0 | 0.232394 | 1 | 2451 | tags=20%, list=9%, signal=22% |
| GO_OXIDOREDUCTASE_ACTIVITY_ACTING_ON_SINGLE_DONORS_WITH_INCORPORATION_OF_MOLECULAR_OXYGEN | 20 | 0.7435168 | 1.5049855 | 0.02380952 | 0.23208152 | 1 | 666 | tags=10%, list=2%, signal=10% |
| HARRIS_BRAIN_CANCER_PROGENITORS | 41 | 0.63247156 | 1.5048109 | 0 | 0.1675675 | 1 | 2947 | tags=34%, list=11%, signal=38% |
| HP_THORACIC_KYPHOSIS | 34 | 0.60303825 | 1.5045829 | 0.02857143 | 0.23282947 | 1 | 3879 | tags=44%, list=14%, signal=51% |
| GO_NEGATIVE_REGULATION_OF_KIDNEY_DEVELOPMENT | 13 | 0.8646576 | 1.5045395 | 0 | 0.2325174 | 1 | 627 | tags=31%, list=2%, signal=31% |
| HP_PHENOTYPIC_VARIABILITY | 382 | 0.49079186 | 1.5038753 | 0 | 0.23415972 | 1 | 3347 | tags=25%, list=12%, signal=28% |
| TORCHIA_TARGETS_OF_EWSR1_FLI1_FUSION_DN | 296 | 0.4827379 | 1.503622 | 0 | 0.16979446 | 1 | 4168 | tags=32%, list=15%, signal=38% |
| GRAESSMANN_APOPTOSIS_BY_SERUM_DEPRIVATION_UP | 502 | 0.46583286 | 1.5035802 | 0 | 0.16949168 | 1 | 3152 | tags=26%, list=12%, signal=29% |
| HP_MEDULLOBLASTOMA | 14 | 0.7508761 | 1.5034962 | 0.04166667 | 0.23476404 | 1 | 2350 | tags=36%, list=9%, signal=39% |
| GO_AMYLOID_BETA_BINDING | 74 | 0.5566366 | 1.50319 | 0 | 0.23530017 | 1 | 2372 | tags=23%, list=9%, signal=25% |
| GO_GLIAL_CELL_DIFFERENTIATION | 215 | 0.49103722 | 1.5028644 | 0 | 0.23574333 | 1 | 1260 | tags=16%, list=5%, signal=16% |
| HP_TREMOR_BY_ANATOMICAL_SITE | 53 | 0.5855249 | 1.5023483 | 0 | 0.23673365 | 1 | 2461 | tags=23%, list=9%, signal=25% |
| GO_CELLULAR_AMINO_ACID_METABOLIC_PROCESS | 307 | 0.47973666 | 1.5023309 | 0 | 0.2364642 | 1 | 3194 | tags=24%, list=12%, signal=27% |
| HP_STROKE | 111 | 0.5370124 | 1.5022416 | 0 | 0.23637858 | 1 | 2833 | tags=24%, list=11%, signal=27% |
| GO_I_BAND | 121 | 0.5414714 | 1.5017827 | 0 | 0.23756862 | 1 | 2942 | tags=21%, list=11%, signal=23% |
| DACOSTA_UV_RESPONSE_VIA_ERCC3_TTD_UP | 51 | 0.57950705 | 1.5017483 | 0 | 0.17290634 | 1 | 3151 | tags=43%, list=12%, signal=49% |
| GO_IMMUNE_RESPONSE_REGULATING_SIGNALING_PATHWAY | 342 | 0.4760928 | 1.5017146 | 0 | 0.23734522 | 1 | 2510 | tags=22%, list=9%, signal=24% |
| FLECHNER_BIOPSY_KIDNEY_TRANSPLANT_OK_VS_DONOR_UP | 528 | 0.46858713 | 1.5016619 | 0 | 0.1727849 | 1 | 2812 | tags=31%, list=10%, signal=34% |
| GO_REGULATION_OF_HOMOTYPIC_CELL_CELL_ADHESION | 26 | 0.68739283 | 1.5016077 | 0.03225806 | 0.23739587 | 1 | 1909 | tags=27%, list=7%, signal=29% |
| HP_ABNORMAL_RETINAL_VASCULAR_MORPHOLOGY | 182 | 0.5187913 | 1.501436 | 0 | 0.23771758 | 1 | 3197 | tags=25%, list=12%, signal=28% |
| GO_CELL_REDOX_HOMEOSTASIS | 54 | 0.61662173 | 1.5014014 | 0 | 0.23747234 | 1 | 2604 | tags=30%, list=10%, signal=33% |
| RUTELLA_RESPONSE_TO_HGF_UP | 386 | 0.48617113 | 1.5011317 | 0 | 0.17303355 | 1 | 3088 | tags=31%, list=11%, signal=35% |
| HP_TREMOR | 451 | 0.46147597 | 1.50105 | 0 | 0.23806684 | 1 | 2809 | tags=22%, list=10%, signal=24% |
| HP_AMINOACIDURIA | 121 | 0.54119813 | 1.5008203 | 0 | 0.23845409 | 1 | 4549 | tags=45%, list=17%, signal=54% |
| GO_NEGATIVE_REGULATION_OF_GLIOGENESIS | 35 | 0.6413769 | 1.5005105 | 0 | 0.23879541 | 1 | 1003 | tags=23%, list=4%, signal=24% |
| REACTOME_DIGESTION | 10 | 0.79434085 | 1.5003855 | 0.04081633 | 0.1739555 | 1 | 398 | tags=10%, list=1%, signal=10% |
| GO_ENTRY_OF_BACTERIUM_INTO_HOST_CELL | 12 | 0.82097757 | 1.5001152 | 0 | 0.23929484 | 1 | 3903 | tags=67%, list=14%, signal=78% |
| REACTOME_BETA_CATENIN_INDEPENDENT_WNT_SIGNALING | 142 | 0.5177621 | 1.5000609 | 0 | 0.17420942 | 1 | 2389 | tags=29%, list=9%, signal=32% |
| GO_PROTEIN_OXIDATION | 13 | 0.840461 | 1.4999169 | 0.04878049 | 0.23988202 | 1 | 3590 | tags=54%, list=13%, signal=62% |
| HP_ELEVATED_ALKALINE_PHOSPHATASE_OF_BONE_ORIGIN | 6 | 0.9192843 | 1.4997146 | 0.02272727 | 0.24012733 | 1 | 2008 | tags=33%, list=7%, signal=36% |
| VERHAAK_GLIOBLASTOMA_NEURAL | 117 | 0.5349735 | 1.4996878 | 0 | 0.17460851 | 1 | 4140 | tags=38%, list=15%, signal=44% |
| HP_INCOMPLETE_PENETRANCE | 133 | 0.51515055 | 1.4996561 | 0 | 0.23994817 | 1 | 1037 | tags=12%, list=4%, signal=12% |
| GO_RESPONSE_TO_SALT_STRESS | 21 | 0.71867144 | 1.4995724 | 0.02439024 | 0.23979227 | 1 | 2118 | tags=29%, list=8%, signal=31% |
| HP_IMMUNE_DYSREGULATION | 5 | 0.9533788 | 1.4995567 | 0 | 0.23954736 | 1 | 72 | tags=20%, list=0%, signal=20% |
| PANGAS_TUMOR_SUPPRESSION_BY_SMAD1_AND_SMAD5_UP | 122 | 0.50956917 | 1.4994675 | 0 | 0.17493354 | 1 | 2704 | tags=25%, list=10%, signal=28% |
| GO_CHANNEL_INHIBITOR_ACTIVITY | 33 | 0.6596491 | 1.4991862 | 0.02325581 | 0.24006121 | 1 | 3014 | tags=42%, list=11%, signal=48% |
| GO_CARDIAC_MUSCLE_CELL_CARDIAC_MUSCLE_CELL_ADHESION | 6 | 0.9139465 | 1.4990317 | 0 | 0.24028465 | 1 | 234 | tags=17%, list=1%, signal=17% |
| HP_CHERRY_RED_SPOT_OF_THE_MACULA | 10 | 0.84325033 | 1.4989119 | 0.05 | 0.24037284 | 1 | 2999 | tags=60%, list=11%, signal=67% |
| LIEN_BREAST_CARCINOMA_METAPLASTIC | 31 | 0.67477405 | 1.4983686 | 0.02439024 | 0.17598635 | 1 | 2260 | tags=35%, list=8%, signal=39% |
| GO_NUCLEOSIDE_TRIPHOSPHATE_BIOSYNTHETIC_PROCESS | 72 | 0.5709875 | 1.4983507 | 0.03333334 | 0.24130404 | 1 | 3276 | tags=38%, list=12%, signal=43% |
| KEGG_GRAFT_VERSUS_HOST_DISEASE | 20 | 0.71056974 | 1.4982833 | 0.02222222 | 0.17578551 | 1 | 1347 | tags=25%, list=5%, signal=26% |
| FULCHER_INFLAMMATORY_RESPONSE_LECTIN_VS_LPS_UP | 512 | 0.47454223 | 1.4982537 | 0 | 0.17540419 | 1 | 3052 | tags=27%, list=11%, signal=29% |
| HP_MUSCLE_FIBER_SPLITTING | 12 | 0.8472014 | 1.4976724 | 0 | 0.2425583 | 1 | 16 | tags=8%, list=0%, signal=8% |
| CHEN_LUNG_CANCER_SURVIVAL | 24 | 0.6896923 | 1.4974933 | 0.02564103 | 0.1763011 | 1 | 3409 | tags=58%, list=13%, signal=67% |
| GO_PSEUDOPODIUM_ORGANIZATION | 12 | 0.80787593 | 1.4973648 | 0.03921569 | 0.24332711 | 1 | 4435 | tags=67%, list=16%, signal=80% |
| REACTOME_PCP_CE_PATHWAY | 88 | 0.5666708 | 1.4972987 | 0 | 0.1762854 | 1 | 1624 | tags=31%, list=6%, signal=33% |
| SANSOM_APC_TARGETS | 181 | 0.4863909 | 1.4970754 | 0 | 0.17623325 | 1 | 4158 | tags=38%, list=15%, signal=45% |
| REACTOME_PROTEIN_FOLDING | 96 | 0.5463299 | 1.4970156 | 0 | 0.1758902 | 1 | 2857 | tags=32%, list=11%, signal=36% |
| GO_PYROPHOSPHATE_HYDROLYSIS_DRIVEN_PROTON_TRANSMEMBRANE_TRANSPORTER_ACTIVITY | 29 | 0.70456517 | 1.4969498 | 0 | 0.24320567 | 1 | 1206 | tags=31%, list=4%, signal=32% |
| GO_INTERFERON_GAMMA_MEDIATED_SIGNALING_PATHWAY | 71 | 0.5825412 | 1.4969312 | 0 | 0.24291663 | 1 | 3336 | tags=42%, list=12%, signal=48% |
| HP_GENERALIZED_AMINOACIDURIA | 10 | 0.86949855 | 1.496821 | 0.02439024 | 0.24286997 | 1 | 2247 | tags=40%, list=8%, signal=44% |
| GO_LONG_CHAIN_FATTY_ACID_TRANSPORTER_ACTIVITY | 15 | 0.7614647 | 1.4965731 | 0.02380952 | 0.24332559 | 1 | 669 | tags=20%, list=2%, signal=20% |
| BLANCO_MELO_BRONCHIAL_EPITHELIAL_CELLS_INFLUENZA_A_INFECTION_UP | 110 | 0.51504165 | 1.4965 | 0.03125 | 0.17678596 | 1 | 3419 | tags=31%, list=13%, signal=35% |
| GO_INTERLEUKIN_27_MEDIATED_SIGNALING_PATHWAY | 9 | 0.86314565 | 1.496141 | 0 | 0.2441299 | 1 | 1939 | tags=33%, list=7%, signal=36% |
| GO_MONOCARBOXYLIC_ACID_METABOLIC_PROCESS | 550 | 0.45548645 | 1.4958606 | 0 | 0.24495353 | 1 | 3561 | tags=29%, list=13%, signal=32% |
| REACTOME_SYNTHESIS_SECRETION_AND_INACTIVATION_OF_GLUCOSE_DEPENDENT_INSULINOTROPIC_POLYPEPTIDE_GIP_ | 7 | 0.85922617 | 1.4951602 | 0.02439024 | 0.17915943 | 1 | 2196 | tags=43%, list=8%, signal=47% |
| GO_BIOACTIVE_LIPID_RECEPTOR_ACTIVITY | 13 | 0.7636128 | 1.4951575 | 0.05 | 0.24679977 | 1 | 3139 | tags=38%, list=12%, signal=44% |
| ONDER_CDH1_TARGETS_3_DN | 40 | 0.6366361 | 1.494872 | 0.02702703 | 0.17928237 | 1 | 498 | tags=10%, list=2%, signal=10% |
| HP_ABNORMAL_CERVICAL_SPINE_MORPHOLOGY | 12 | 0.80577976 | 1.4946135 | 0.0212766 | 0.24839787 | 1 | 2009 | tags=33%, list=7%, signal=36% |
| BOYAULT_LIVER_CANCER_SUBCLASS_G123_DN | 43 | 0.6533795 | 1.4946036 | 0.02564103 | 0.1795518 | 1 | 3596 | tags=33%, list=13%, signal=38% |
| GO_NEPHRON_TUBULE_EPITHELIAL_CELL_DIFFERENTIATION | 13 | 0.83499295 | 1.4944415 | 0.04651163 | 0.24842946 | 1 | 3104 | tags=38%, list=12%, signal=43% |
| GO_SLEEP | 25 | 0.6902208 | 1.4943241 | 0 | 0.24837568 | 1 | 1729 | tags=20%, list=6%, signal=21% |
| GO_AZUROPHIL_GRANULE | 131 | 0.52685684 | 1.4936031 | 0 | 0.25018716 | 1 | 2836 | tags=33%, list=11%, signal=37% |
| FURUKAWA_DUSP6_TARGETS_PCI35_UP | 51 | 0.6088697 | 1.493571 | 0.02702703 | 0.18102495 | 1 | 1626 | tags=22%, list=6%, signal=23% |
| WAKABAYASHI_ADIPOGENESIS_PPARG_RXRA_BOUND_8D | 786 | 0.45297417 | 1.4930439 | 0 | 0.18161184 | 1 | 3228 | tags=30%, list=12%, signal=33% |
| GO_REGULATION_OF_EPIDERMIS_DEVELOPMENT | 70 | 0.58230925 | 1.4929811 | 0 | 0.2521412 | 1 | 3651 | tags=31%, list=14%, signal=36% |
| HP_POSTERIOR_POLAR_CATARACT | 6 | 0.9301845 | 1.492903 | 0 | 0.2521052 | 1 | 104 | tags=50%, list=0%, signal=50% |
| FRIDMAN_SENESCENCE_UP | 70 | 0.5869251 | 1.4927894 | 0.02941177 | 0.1816925 | 1 | 1097 | tags=29%, list=4%, signal=30% |
| HP_HYPOREFLEXIA | 313 | 0.47045237 | 1.4926097 | 0 | 0.25271645 | 1 | 2465 | tags=20%, list=9%, signal=22% |
| BIOCARTA_PROTEASOME_PATHWAY | 18 | 0.7559515 | 1.4924976 | 0.04255319 | 0.18194675 | 1 | 1857 | tags=61%, list=7%, signal=66% |
| ONO_AML1_TARGETS_DN | 34 | 0.63072616 | 1.4924268 | 0 | 0.1817773 | 1 | 1765 | tags=24%, list=7%, signal=25% |
| GO_COMMA_SHAPED_BODY_MORPHOGENESIS | 5 | 0.9878784 | 1.4923218 | 0 | 0.2533249 | 1 | 167 | tags=40%, list=1%, signal=40% |
| HP_HYPOVENTILATION | 36 | 0.6409945 | 1.4921906 | 0 | 0.25337183 | 1 | 1736 | tags=25%, list=6%, signal=27% |
| HP_HYPERTONIA | 828 | 0.44362563 | 1.4914417 | 0 | 0.25503147 | 1 | 4106 | tags=35%, list=15%, signal=40% |
| GO_SPERMATOPROTEASOME_COMPLEX | 4 | 0.94104815 | 1.4909 | 0 | 0.2561485 | 1 | 357 | tags=75%, list=1%, signal=76% |
| TORCHIA_TARGETS_OF_EWSR1_FLI1_FUSION_TOP20_UP | 17 | 0.6964695 | 1.490853 | 0.04878049 | 0.1837813 | 1 | 237 | tags=18%, list=1%, signal=18% |
| HP_MODERATE_GLOBAL_DEVELOPMENTAL_DELAY | 16 | 0.7707012 | 1.4905379 | 0.03125 | 0.25681382 | 1 | 3144 | tags=50%, list=12%, signal=57% |
| SATO_SILENCED_BY_METHYLATION_IN_PANCREATIC_CANCER_2 | 37 | 0.63043046 | 1.4905211 | 0.02941177 | 0.18336841 | 1 | 1945 | tags=19%, list=7%, signal=20% |
| GO_REGULATION_OF_CELL_CYCLE_G2_M_PHASE_TRANSITION | 199 | 0.50235134 | 1.4904648 | 0 | 0.2566194 | 1 | 2547 | tags=27%, list=9%, signal=30% |
| HP_ABNORMAL_NEPHRON_MORPHOLOGY | 179 | 0.51282436 | 1.4903812 | 0 | 0.25670433 | 1 | 3471 | tags=28%, list=13%, signal=32% |
| GO_LIPID_CATABOLIC_PROCESS | 273 | 0.4922444 | 1.4903693 | 0 | 0.2563832 | 1 | 3990 | tags=34%, list=15%, signal=40% |
| GO_RESPONSE_TO_AUDITORY_STIMULUS | 24 | 0.6986271 | 1.4903283 | 0 | 0.25627506 | 1 | 574 | tags=17%, list=2%, signal=17% |
| HP_PROGRESSIVE_HEARING_IMPAIRMENT | 38 | 0.65871406 | 1.4902679 | 0 | 0.25623327 | 1 | 1098 | tags=18%, list=4%, signal=19% |
| GO_PURINE_DEOXYRIBONUCLEOTIDE_METABOLIC_PROCESS | 11 | 0.7620347 | 1.4901844 | 0 | 0.25616848 | 1 | 2792 | tags=45%, list=10%, signal=51% |
| MOOTHA_GLUCONEOGENESIS | 28 | 0.6926459 | 1.490162 | 0.02173913 | 0.18362026 | 1 | 3141 | tags=43%, list=12%, signal=48% |
| GO_CILIARY_TIP | 43 | 0.6222728 | 1.4901485 | 0.025 | 0.25595635 | 1 | 4017 | tags=42%, list=15%, signal=49% |
| CAIRO_LIVER_DEVELOPMENT_UP | 156 | 0.51774234 | 1.489879 | 0 | 0.18372974 | 1 | 2065 | tags=28%, list=8%, signal=30% |
| TONKS_TARGETS_OF_RUNX1_RUNX1T1_FUSION_ERYTHROCYTE_UP | 139 | 0.51150954 | 1.4896612 | 0 | 0.18384154 | 1 | 1639 | tags=19%, list=6%, signal=21% |
| REACTOME_FATTY_ACID_METABOLISM | 146 | 0.50406265 | 1.4893924 | 0 | 0.18405537 | 1 | 4331 | tags=40%, list=16%, signal=48% |
| GO_DIOL_METABOLIC_PROCESS | 22 | 0.72241026 | 1.4891512 | 0.02272727 | 0.25813943 | 1 | 3272 | tags=55%, list=12%, signal=62% |
| GO_PROTEIN_EXIT_FROM_ENDOPLASMIC_RETICULUM | 40 | 0.59369934 | 1.4889259 | 0 | 0.25849584 | 1 | 2176 | tags=40%, list=8%, signal=43% |
| HP_EXTRAPYRAMIDAL_MUSCULAR_RIGIDITY | 12 | 0.7752905 | 1.4888487 | 0.02631579 | 0.25847027 | 1 | 4269 | tags=67%, list=16%, signal=79% |
| HP_COGNITIVE_IMPAIRMENT | 540 | 0.4643369 | 1.4887599 | 0 | 0.25842524 | 1 | 3552 | tags=28%, list=13%, signal=31% |
| HP_ANHIDROSIS | 18 | 0.73899376 | 1.4886336 | 0.04761905 | 0.25838038 | 1 | 1273 | tags=28%, list=5%, signal=29% |
| HP_ABNORMAL_LARGE_INTESTINE_PHYSIOLOGY | 40 | 0.655197 | 1.4884479 | 0.02777778 | 0.25818753 | 1 | 2295 | tags=30%, list=9%, signal=33% |
| HP_ESOPHAGEAL_STRICTURE | 7 | 0.9006549 | 1.4884064 | 0.02631579 | 0.25797507 | 1 | 1800 | tags=43%, list=7%, signal=46% |
| HP_HEADACHE | 261 | 0.479798 | 1.4883243 | 0 | 0.257973 | 1 | 2400 | tags=18%, list=9%, signal=20% |
| GO_N_ACETYLNEURAMINATE_METABOLIC_PROCESS | 10 | 0.84752023 | 1.4882387 | 0.04 | 0.25786558 | 1 | 1829 | tags=60%, list=7%, signal=64% |
| REACTOME_TRANSPORT_OF_VITAMINS_NUCLEOSIDES_AND_RELATED_MOLECULES | 35 | 0.6307214 | 1.4880047 | 0.02702703 | 0.18566541 | 1 | 1354 | tags=17%, list=5%, signal=18% |
| SIMBULAN_PARP1_TARGETS_UP | 29 | 0.6709475 | 1.4878082 | 0.02272727 | 0.18570137 | 1 | 2434 | tags=34%, list=9%, signal=38% |
| LEE_TARGETS_OF_PTCH1_AND_SUFU_UP | 52 | 0.5543868 | 1.4877673 | 0 | 0.18549518 | 1 | 1962 | tags=29%, list=7%, signal=31% |
| REACTOME_ASSEMBLY_OF_THE_PRE_REPLICATIVE_COMPLEX | 65 | 0.6112595 | 1.4877272 | 0 | 0.18522002 | 1 | 2352 | tags=42%, list=9%, signal=45% |
| GOLDRATH_HOMEOSTATIC_PROLIFERATION | 139 | 0.5304587 | 1.4877015 | 0.03846154 | 0.18487592 | 1 | 2377 | tags=39%, list=9%, signal=42% |
| HP_PROMINENT_VEINS_ON_TRUNK | 4 | 0.93124014 | 1.4873692 | 0.02040816 | 0.26033172 | 1 | 344 | tags=50%, list=1%, signal=51% |
| GO_NEGATIVE_REGULATION_OF_NEUROBLAST_PROLIFERATION | 5 | 0.96563584 | 1.4869732 | 0 | 0.2611623 | 1 | 84 | tags=40%, list=0%, signal=40% |
| BASSO_B_LYMPHOCYTE_NETWORK | 138 | 0.5300951 | 1.486808 | 0.02941177 | 0.18643156 | 1 | 3150 | tags=44%, list=12%, signal=50% |
| GO_PLATELET_DENSE_GRANULE_MEMBRANE | 5 | 0.8927564 | 1.4867728 | 0 | 0.26132327 | 1 | 2353 | tags=60%, list=9%, signal=66% |
| GO_UNFOLDED_PROTEIN_BINDING | 97 | 0.5520203 | 1.4867727 | 0 | 0.2609839 | 1 | 3736 | tags=40%, list=14%, signal=47% |
| GO_METANEPHRIC_NEPHRON_EPITHELIUM_DEVELOPMENT | 18 | 0.74345124 | 1.4866538 | 0.02380952 | 0.26095742 | 1 | 1528 | tags=33%, list=6%, signal=35% |
| HP_GLYCOSURIA | 37 | 0.6508317 | 1.486379 | 0.02857143 | 0.2614263 | 1 | 3199 | tags=35%, list=12%, signal=40% |
| HP_CONCENTRIC_HYPERTROPHIC_CARDIOMYOPATHY | 9 | 0.9285754 | 1.4862921 | 0 | 0.26142022 | 1 | 1281 | tags=67%, list=5%, signal=70% |
| HP_RECURRENT_MYCOBACTERIAL_INFECTIONS | 8 | 0.90102524 | 1.4861147 | 0.01923077 | 0.26157874 | 1 | 72 | tags=25%, list=0%, signal=25% |
| BIOCARTA_ETC_PATHWAY | 8 | 0.92253876 | 1.4859774 | 0 | 0.18763426 | 1 | 1130 | tags=88%, list=4%, signal=91% |
| GO_NEUROTRANSMITTER_UPTAKE | 33 | 0.647338 | 1.4858778 | 0 | 0.2619234 | 1 | 1426 | tags=24%, list=5%, signal=26% |
| ZHU_CMV_ALL_UP | 99 | 0.5277675 | 1.485745 | 0 | 0.18780355 | 1 | 3595 | tags=40%, list=13%, signal=46% |
| REACTOME_G1_S_DNA_DAMAGE_CHECKPOINTS | 64 | 0.6004931 | 1.4851108 | 0.02564103 | 0.18862619 | 1 | 1759 | tags=39%, list=7%, signal=42% |
| GO_REGULATION_OF_STRESS_GRANULE_ASSEMBLY | 4 | 0.91685826 | 1.4848595 | 0.025 | 0.26451743 | 1 | 1124 | tags=25%, list=4%, signal=26% |
| WP_CYTOSOLIC_DNASENSING_PATHWAY | 53 | 0.597148 | 1.484736 | 0 | 0.18879132 | 1 | 2452 | tags=34%, list=9%, signal=37% |
| GO_ORGAN_OR_TISSUE_SPECIFIC_IMMUNE_RESPONSE | 14 | 0.77901185 | 1.4842949 | 0.04761905 | 0.26578316 | 1 | 1328 | tags=14%, list=5%, signal=15% |
| GO_NEGATIVE_REGULATION_OF_ISOTYPE_SWITCHING | 4 | 0.90068 | 1.4842519 | 0.04651163 | 0.26566824 | 1 | 1772 | tags=75%, list=7%, signal=80% |
| GO_NEGATIVE_REGULATION_OF_PROTEIN_IMPORT | 11 | 0.83340317 | 1.484198 | 0.02702703 | 0.26547194 | 1 | 2747 | tags=55%, list=10%, signal=61% |
| GO_PURINE_CONTAINING_COMPOUND_BIOSYNTHETIC_PROCESS | 183 | 0.49185473 | 1.4835461 | 0 | 0.2663153 | 1 | 3573 | tags=32%, list=13%, signal=36% |
| GO_INORGANIC_ION_TRANSMEMBRANE_TRANSPORT | 725 | 0.45134765 | 1.4834499 | 0 | 0.26628178 | 1 | 3471 | tags=20%, list=13%, signal=22% |
| GRANDVAUX_IFN_RESPONSE_NOT_VIA_IRF3 | 11 | 0.88317156 | 1.4834305 | 0 | 0.19089782 | 1 | 2141 | tags=82%, list=8%, signal=89% |
| HP_EMG_DECREMENTAL_RESPONSE_OF_COMPOUND_MUSCLE_ACTION_POTENTIAL_TO_REPETITIVE_NERVE_STIMULATION | 22 | 0.69203067 | 1.4830204 | 0.02272727 | 0.26739204 | 1 | 4184 | tags=27%, list=16%, signal=32% |
| REACTOME_SYNTHESIS_OF_12_EICOSATETRAENOIC_ACID_DERIVATIVES | 4 | 0.9758374 | 1.4828501 | 0 | 0.19129214 | 1 | 593 | tags=50%, list=2%, signal=51% |
| CHEN_METABOLIC_SYNDROM_NETWORK | 1155 | 0.4318104 | 1.4827952 | 0 | 0.19059227 | 1 | 3585 | tags=24%, list=13%, signal=26% |
| ONDER_CDH1_TARGETS_2_UP | 243 | 0.4724968 | 1.4825523 | 0 | 0.1904825 | 1 | 2871 | tags=27%, list=11%, signal=30% |
| RODWELL_AGING_KIDNEY_NO_BLOOD_UP | 190 | 0.5068719 | 1.4823419 | 0 | 0.19054353 | 1 | 1904 | tags=21%, list=7%, signal=22% |
| ONDER_CDH1_SIGNALING_VIA_CTNNB1 | 80 | 0.5757516 | 1.4817132 | 0 | 0.19154528 | 1 | 3432 | tags=38%, list=13%, signal=43% |
| GO_REGULATION_OF_POSTSYNAPTIC_CYTOSOLIC_CALCIUM_ION_CONCENTRATION | 12 | 0.80309767 | 1.481644 | 0.04444445 | 0.27082494 | 1 | 3355 | tags=33%, list=12%, signal=38% |
| GO_INTRAMOLECULAR_OXIDOREDUCTASE_ACTIVITY_TRANSPOSING_C_C_BONDS | 12 | 0.7742171 | 1.4815133 | 0.02222222 | 0.27084607 | 1 | 2290 | tags=50%, list=9%, signal=55% |
| REACTOME_RNA_POLYMERASE_III_TRANSCRIPTION | 40 | 0.6301435 | 1.48133 | 0.04444445 | 0.19187148 | 1 | 2627 | tags=43%, list=10%, signal=47% |
| HP_ABNORMAL_RENAL_PHYSIOLOGY | 524 | 0.4644184 | 1.4809643 | 0 | 0.2719484 | 1 | 3497 | tags=26%, list=13%, signal=29% |
| GO_MORPHOGENESIS_OF_A_POLARIZED_EPITHELIUM | 139 | 0.53367186 | 1.4804618 | 0 | 0.27326927 | 1 | 2270 | tags=26%, list=8%, signal=28% |
| GO_GLYCOSYLCERAMIDE_METABOLIC_PROCESS | 16 | 0.76477706 | 1.4802392 | 0 | 0.27350873 | 1 | 4699 | tags=75%, list=17%, signal=91% |
| GO_FATTY_ACID_CATABOLIC_PROCESS | 102 | 0.5440178 | 1.479915 | 0.03225806 | 0.27397394 | 1 | 3918 | tags=44%, list=15%, signal=51% |
| GO_ER_OVERLOAD_RESPONSE | 10 | 0.814136 | 1.4798551 | 0.02564103 | 0.27376974 | 1 | 2023 | tags=60%, list=8%, signal=65% |
| REACTOME_HOST_INTERACTIONS_OF_HIV_FACTORS | 119 | 0.5174976 | 1.4796891 | 0 | 0.19390798 | 1 | 2352 | tags=32%, list=9%, signal=35% |
| RICKMAN_TUMOR_DIFFERENTIATED_WELL_VS_POORLY_DN | 332 | 0.4714759 | 1.4795951 | 0 | 0.1938258 | 1 | 2884 | tags=24%, list=11%, signal=27% |
| HP_HYPOKALEMIA | 41 | 0.63412076 | 1.4793932 | 0.03030303 | 0.2747177 | 1 | 1628 | tags=17%, list=6%, signal=18% |
| HP_ABNORMAL_TENDON_MORPHOLOGY | 597 | 0.44713557 | 1.4793702 | 0 | 0.27445224 | 1 | 4006 | tags=29%, list=15%, signal=34% |
| GO_BASEMENT_MEMBRANE | 94 | 0.5401024 | 1.479353 | 0 | 0.27416712 | 1 | 3097 | tags=22%, list=12%, signal=25% |
| GO_RAGE_RECEPTOR_BINDING | 7 | 0.9263744 | 1.4792442 | 0 | 0.27384004 | 1 | 1777 | tags=86%, list=7%, signal=92% |
| GO_MIDGUT_DEVELOPMENT | 10 | 0.7742047 | 1.4792374 | 0.04651163 | 0.2735171 | 1 | 1003 | tags=20%, list=4%, signal=21% |
| BEGUM_TARGETS_OF_PAX3_FOXO1_FUSION_DN | 44 | 0.6045991 | 1.4791647 | 0.03030303 | 0.19392744 | 1 | 2330 | tags=27%, list=9%, signal=30% |
| HP_INCREASED_MUSCLE_LIPID_CONTENT | 31 | 0.6746127 | 1.4791172 | 0 | 0.2735363 | 1 | 2622 | tags=32%, list=10%, signal=36% |
| GO_INHIBITION_OF_NEUROEPITHELIAL_CELL_DIFFERENTIATION | 4 | 0.9738489 | 1.479098 | 0.02777778 | 0.2732942 | 1 | 167 | tags=50%, list=1%, signal=50% |
| GO_HOMOTYPIC_CELL_CELL_ADHESION | 74 | 0.5733799 | 1.4789542 | 0.025 | 0.27339283 | 1 | 2031 | tags=26%, list=8%, signal=28% |
| SMID_BREAST_CANCER_RELAPSE_IN_BRAIN_UP | 32 | 0.6858819 | 1.4788578 | 0.02439024 | 0.19369948 | 1 | 4326 | tags=41%, list=16%, signal=48% |
| GOZGIT_ESR1_TARGETS_DN | 641 | 0.45759133 | 1.4788461 | 0 | 0.19338703 | 1 | 3308 | tags=24%, list=12%, signal=27% |
| GO_REGULATION_OF_EPITHELIAL_CELL_DIFFERENTIATION_INVOLVED_IN_KIDNEY_DEVELOPMENT | 16 | 0.7995111 | 1.4786284 | 0.02702703 | 0.27401087 | 1 | 1528 | tags=31%, list=6%, signal=33% |
| GO_VACUOLAR_MEMBRANE | 382 | 0.4591779 | 1.4784852 | 0 | 0.2740872 | 1 | 3987 | tags=34%, list=15%, signal=39% |
| HP_ABNORMAL_VASCULAR_PHYSIOLOGY | 149 | 0.5137991 | 1.4783183 | 0 | 0.27408516 | 1 | 2928 | tags=23%, list=11%, signal=26% |
| REACTOME_GLUCURONIDATION | 5 | 0.9281087 | 1.4779117 | 0.02 | 0.19475205 | 1 | 1478 | tags=40%, list=5%, signal=42% |
| PUIFFE_INVASION_INHIBITED_BY_ASCITES_DN | 134 | 0.53477556 | 1.4779058 | 0 | 0.19440535 | 1 | 3975 | tags=46%, list=15%, signal=54% |
| HP_DECREASED_SERUM_IRON | 4 | 0.90971243 | 1.4777014 | 0.04444445 | 0.27541572 | 1 | 2433 | tags=100%, list=9%, signal=110% |
| ZHAN_MULTIPLE_MYELOMA_MF_UP | 42 | 0.6350181 | 1.4774969 | 0.02222222 | 0.19490011 | 1 | 1099 | tags=21%, list=4%, signal=22% |
| GO_PEPTIDASE_ACTIVATOR_ACTIVITY | 36 | 0.6391455 | 1.4774249 | 0.04347826 | 0.27594817 | 1 | 2661 | tags=33%, list=10%, signal=37% |
| HOSHIDA_LIVER_CANCER_SURVIVAL_UP | 64 | 0.5822453 | 1.4773831 | 0.02564103 | 0.1947197 | 1 | 2383 | tags=28%, list=9%, signal=31% |
| REACTOME_ALPHA_LINOLENIC_OMEGA3_AND_LINOLEIC_OMEGA6_ACID_METABOLISM | 12 | 0.8031034 | 1.4770552 | 0.02631579 | 0.19483368 | 1 | 1854 | tags=42%, list=7%, signal=45% |
| ONGUSAHA_TP53_TARGETS | 35 | 0.6707519 | 1.4769981 | 0.02325581 | 0.19449072 | 1 | 2917 | tags=46%, list=11%, signal=51% |
| HP_ABNORMAL_HOMEOSTASIS | 1113 | 0.44271302 | 1.4769835 | 0 | 0.27687177 | 1 | 3535 | tags=25%, list=13%, signal=28% |
| ICHIBA_GRAFT_VERSUS_HOST_DISEASE_35D_UP | 133 | 0.50366306 | 1.476908 | 0 | 0.19427906 | 1 | 1777 | tags=21%, list=7%, signal=22% |
| BOUDOUKHA_BOUND_BY_IGF2BP2 | 100 | 0.5457308 | 1.4768143 | 0.03448276 | 0.19410002 | 1 | 2058 | tags=41%, list=8%, signal=44% |
| GO_RESPONSE_TO_BACTERIUM | 463 | 0.46258494 | 1.4767998 | 0 | 0.27720457 | 1 | 2604 | tags=17%, list=10%, signal=19% |
| GO_FATTY_ACID_ALPHA_OXIDATION | 8 | 0.86167467 | 1.4767314 | 0.04166667 | 0.27700135 | 1 | 3647 | tags=75%, list=14%, signal=87% |
| HP_PERIPHERAL_AXONAL_NEUROPATHY | 119 | 0.5308694 | 1.4767265 | 0 | 0.2766598 | 1 | 2009 | tags=21%, list=7%, signal=23% |
| GO_POSITIVE_REGULATION_OF_BONE_MINERALIZATION | 34 | 0.6316099 | 1.4765066 | 0 | 0.2772667 | 1 | 1334 | tags=18%, list=5%, signal=19% |
| GO_RESPIRATORY_CHAIN_COMPLEX_II | 4 | 0.97298807 | 1.4764581 | 0 | 0.27706334 | 1 | 730 | tags=100%, list=3%, signal=103% |
| REACTOME_PEROXISOMAL_LIPID_METABOLISM | 27 | 0.70679665 | 1.4762783 | 0.02857143 | 0.19473718 | 1 | 4331 | tags=48%, list=16%, signal=57% |
| GO_NEGATIVE_REGULATION_OF_CHONDROCYTE_DIFFERENTIATION | 18 | 0.7700905 | 1.4761777 | 0.02857143 | 0.277804 | 1 | 1295 | tags=22%, list=5%, signal=23% |
| HP_LIMB_GIRDLE_MUSCLE_WEAKNESS | 72 | 0.61079985 | 1.4760596 | 0.03846154 | 0.2779347 | 1 | 2338 | tags=24%, list=9%, signal=26% |
| SENESE_HDAC1_AND_HDAC2_TARGETS_DN | 196 | 0.49643385 | 1.4759872 | 0 | 0.19504383 | 1 | 4057 | tags=35%, list=15%, signal=41% |
| GO_POSITIVE_REGULATION_OF_RESPONSE_TO_WOUNDING | 57 | 0.5956926 | 1.4759005 | 0 | 0.2781453 | 1 | 1334 | tags=18%, list=5%, signal=18% |
| HP_SPASTICITY | 694 | 0.4412229 | 1.4758433 | 0 | 0.27794158 | 1 | 3441 | tags=29%, list=13%, signal=33% |
| GO_TRNA_METHYLATION | 34 | 0.63216966 | 1.4757963 | 0.02272727 | 0.277778 | 1 | 4654 | tags=56%, list=17%, signal=67% |
| GO_RESPONSE_TO_OSMOTIC_STRESS | 78 | 0.5665658 | 1.4756595 | 0.04761905 | 0.2779884 | 1 | 2768 | tags=26%, list=10%, signal=28% |
| WP_REGULATION_OF_WNTBCATENIN_SIGNALING_BY_SMALL_MOLECULE_COMPOUNDS | 15 | 0.7287052 | 1.4753621 | 0.04347826 | 0.1959326 | 1 | 1972 | tags=33%, list=7%, signal=36% |
| GO_PROTEIN_FOLDING | 197 | 0.5115556 | 1.475254 | 0 | 0.27859813 | 1 | 2898 | tags=32%, list=11%, signal=36% |
| GO_CRANIAL_NERVE_DEVELOPMENT | 43 | 0.5992704 | 1.4752483 | 0.04347826 | 0.2782592 | 1 | 499 | tags=12%, list=2%, signal=12% |
| FOSTER_KDM1A_TARGETS_UP | 195 | 0.4906428 | 1.4751705 | 0.04166667 | 0.19600777 | 1 | 4039 | tags=25%, list=15%, signal=29% |
| COLIN_PILOCYTIC_ASTROCYTOMA_VS_GLIOBLASTOMA_UP | 30 | 0.65427953 | 1.4747442 | 0.02702703 | 0.19559999 | 1 | 848 | tags=30%, list=3%, signal=31% |
| WP_SMALL_LIGAND_GPCRS | 13 | 0.7850418 | 1.4745401 | 0 | 0.19555044 | 1 | 3139 | tags=23%, list=12%, signal=26% |
| NIKOLSKY_BREAST_CANCER_5P15_AMPLICON | 21 | 0.7612235 | 1.4741242 | 0.02439024 | 0.19613399 | 1 | 2400 | tags=33%, list=9%, signal=37% |
| HP_ABNORMAL_PLATELET_COUNT | 278 | 0.47917888 | 1.4740916 | 0 | 0.28191206 | 1 | 3535 | tags=29%, list=13%, signal=33% |
| GO_ENDOCARDIAL_CUSHION_MORPHOGENESIS | 31 | 0.64438134 | 1.4740393 | 0.04761905 | 0.28172508 | 1 | 2396 | tags=23%, list=9%, signal=25% |
| GO_SMAD_PROTEIN_SIGNAL_TRANSDUCTION | 63 | 0.5674437 | 1.4738827 | 0.02272727 | 0.2818504 | 1 | 2065 | tags=14%, list=8%, signal=15% |
| ACEVEDO_LIVER_TUMOR_VS_NORMAL_ADJACENT_TISSUE_UP | 789 | 0.44911936 | 1.4738716 | 0 | 0.19614601 | 1 | 3193 | tags=34%, list=12%, signal=38% |
| SENESE_HDAC2_TARGETS_DN | 109 | 0.53143984 | 1.4735857 | 0 | 0.19610609 | 1 | 2800 | tags=22%, list=10%, signal=24% |
| GO_NEGATIVE_REGULATION_OF_INSULIN_LIKE_GROWTH_FACTOR_RECEPTOR_SIGNALING_PATHWAY | 4 | 0.9895138 | 1.4733267 | 0 | 0.2826604 | 1 | 142 | tags=25%, list=1%, signal=25% |
| GO_VACUOLE | 705 | 0.4547649 | 1.4732907 | 0 | 0.28241563 | 1 | 3987 | tags=32%, list=15%, signal=36% |
| GO_CELLULAR_COMPONENT_DISASSEMBLY | 514 | 0.46851963 | 1.4731258 | 0 | 0.28242275 | 1 | 2481 | tags=26%, list=9%, signal=28% |
| GRAHAM_CML_QUIESCENT_VS_NORMAL_DIVIDING_DN | 10 | 0.866938 | 1.473031 | 0.04545455 | 0.1969426 | 1 | 1234 | tags=40%, list=5%, signal=42% |
| GO_NEGATIVE_REGULATION_OF_NATURAL_KILLER_CELL_MEDIATED_IMMUNITY | 10 | 0.8382824 | 1.4728879 | 0.025 | 0.2829307 | 1 | 383 | tags=20%, list=1%, signal=20% |
| GO_MACROPHAGE_MIGRATION | 45 | 0.56547815 | 1.4725801 | 0 | 0.28353426 | 1 | 2465 | tags=29%, list=9%, signal=32% |
| GO_ANTIOXIDANT_ACTIVITY | 68 | 0.5603698 | 1.47256 | 0 | 0.28321266 | 1 | 2604 | tags=28%, list=10%, signal=31% |
| HP_GLUTARIC_ACIDURIA | 7 | 0.84860075 | 1.4723636 | 0.04 | 0.28352764 | 1 | 3165 | tags=71%, list=12%, signal=81% |
| GO_REGULATION_OF_RESPONSE_TO_EXTERNAL_STIMULUS | 865 | 0.44400108 | 1.4721925 | 0 | 0.2837249 | 1 | 2479 | tags=17%, list=9%, signal=18% |
| KEGG_GLYCEROLIPID_METABOLISM | 42 | 0.5886327 | 1.4721224 | 0 | 0.19785562 | 1 | 4069 | tags=38%, list=15%, signal=45% |
| GINESTIER_BREAST_CANCER_20Q13_AMPLIFICATION_UP | 93 | 0.5384081 | 1.4720682 | 0 | 0.1976773 | 1 | 2575 | tags=33%, list=10%, signal=37% |
| GO_ACTIN_BASED_CELL_PROJECTION | 197 | 0.4837068 | 1.472015 | 0 | 0.28388572 | 1 | 2372 | tags=18%, list=9%, signal=19% |
| GO_POSITIVE_REGULATION_OF_TYPE_I_INTERFERON_PRODUCTION | 69 | 0.5644046 | 1.47183 | 0 | 0.2839874 | 1 | 2452 | tags=33%, list=9%, signal=37% |
| HP_PULMONARY_EMBOLISM | 27 | 0.6823364 | 1.4716583 | 0.02777778 | 0.28416374 | 1 | 1967 | tags=19%, list=7%, signal=20% |
| HP_TELANGIECTASIA_OF_THE_SKIN | 64 | 0.5779475 | 1.4716386 | 0 | 0.2838437 | 1 | 4619 | tags=47%, list=17%, signal=56% |
| HP_ABNORMALITY_OF_URINE_HOMEOSTASIS | 503 | 0.46426678 | 1.4713851 | 0 | 0.28426933 | 1 | 3535 | tags=28%, list=13%, signal=31% |
| PID_IL23_PATHWAY | 32 | 0.6817519 | 1.4713775 | 0.02380952 | 0.19856733 | 1 | 1939 | tags=19%, list=7%, signal=20% |
| GO_EMBRYONIC_SKELETAL_JOINT_MORPHOGENESIS | 5 | 0.9121285 | 1.4713469 | 0.02380952 | 0.28410336 | 1 | 627 | tags=40%, list=2%, signal=41% |
| GAVIN_FOXP3_TARGETS_CLUSTER_T7 | 98 | 0.55199945 | 1.4712511 | 0.02941177 | 0.19851077 | 1 | 2791 | tags=38%, list=10%, signal=42% |
| YAMASHITA_METHYLATED_IN_PROSTATE_CANCER | 46 | 0.6118939 | 1.4712244 | 0.02439024 | 0.19817446 | 1 | 2816 | tags=26%, list=10%, signal=29% |
| MCCLUNG_CREB1_TARGETS_UP | 95 | 0.5383606 | 1.4711965 | 0 | 0.19784042 | 1 | 2936 | tags=32%, list=11%, signal=35% |
| KATSANOU_ELAVL1_TARGETS_UP | 156 | 0.51627564 | 1.4711096 | 0 | 0.1975376 | 1 | 2661 | tags=22%, list=10%, signal=24% |
| GO_CARD_DOMAIN_BINDING | 12 | 0.7915214 | 1.4710171 | 0.02439024 | 0.28481424 | 1 | 1195 | tags=17%, list=4%, signal=17% |
| HP_ABNORMAL_RETINAL_MORPHOLOGY | 731 | 0.45644513 | 1.4709748 | 0 | 0.28466743 | 1 | 3535 | tags=27%, list=13%, signal=30% |
| REACTOME_PLATELET_ACTIVATION_SIGNALING_AND_AGGREGATION | 232 | 0.4721939 | 1.4708424 | 0 | 0.19754839 | 1 | 2353 | tags=22%, list=9%, signal=24% |
| GO_TAURINE_METABOLIC_PROCESS | 6 | 0.9086396 | 1.470773 | 0.02222222 | 0.2849956 | 1 | 445 | tags=50%, list=2%, signal=51% |
| RIGGINS_TAMOXIFEN_RESISTANCE_DN | 200 | 0.5014597 | 1.4705511 | 0 | 0.19758928 | 1 | 3588 | tags=35%, list=13%, signal=40% |
| HP_EXERTIONAL_DYSPNEA | 79 | 0.5525939 | 1.4705015 | 0 | 0.2853988 | 1 | 3709 | tags=29%, list=14%, signal=34% |
| GO_RESPONSE_TO_VIRUS | 272 | 0.46386555 | 1.4704822 | 0 | 0.28509864 | 1 | 2479 | tags=26%, list=9%, signal=29% |
| GO_SULFURTRANSFERASE_ACTIVITY | 9 | 0.8138987 | 1.4703828 | 0.04 | 0.2849129 | 1 | 2994 | tags=67%, list=11%, signal=75% |
| GO_L_LACTATE_DEHYDROGENASE_ACTIVITY | 3 | 0.9960923 | 1.4701531 | 0 | 0.28542867 | 1 | 44 | tags=33%, list=0%, signal=33% |
| ONKEN_UVEAL_MELANOMA_UP | 736 | 0.4604237 | 1.4699823 | 0 | 0.19785464 | 1 | 2663 | tags=29%, list=10%, signal=31% |
| GO_HETEROTYPIC_CELL_CELL_ADHESION | 53 | 0.60372424 | 1.469939 | 0 | 0.28562364 | 1 | 893 | tags=13%, list=3%, signal=14% |
| GO_MITOCHONDRIAL_RESPIRASOME_ASSEMBLY | 5 | 0.8733833 | 1.4699326 | 0 | 0.2852876 | 1 | 1244 | tags=60%, list=5%, signal=63% |
| HP_PERIPHERAL_DEMYELINATION | 42 | 0.6338823 | 1.4698371 | 0 | 0.28531086 | 1 | 2480 | tags=40%, list=9%, signal=45% |
| HP_HYPOMETRIC_SACCADES | 12 | 0.7594069 | 1.4697018 | 0.02222222 | 0.28525674 | 1 | 679 | tags=17%, list=3%, signal=17% |
| FARMER_BREAST_CANCER_APOCRINE_VS_LUMINAL | 284 | 0.47019437 | 1.4692733 | 0 | 0.19878829 | 1 | 3594 | tags=33%, list=13%, signal=37% |
| GO_HINDBRAIN_MORPHOGENESIS | 40 | 0.60330933 | 1.4692708 | 0.02439024 | 0.28689736 | 1 | 2315 | tags=25%, list=9%, signal=27% |
| HP_PTOSIS | 552 | 0.45811623 | 1.4690086 | 0 | 0.28729233 | 1 | 3537 | tags=30%, list=13%, signal=33% |
| REACTOME_FORMATION_OF_FIBRIN_CLOT_CLOTTING_CASCADE_ | 27 | 0.69531345 | 1.4690081 | 0 | 0.19861989 | 1 | 1086 | tags=11%, list=4%, signal=12% |
| HP_MANIA | 31 | 0.67704433 | 1.4688683 | 0.04545455 | 0.28746215 | 1 | 854 | tags=19%, list=3%, signal=20% |
| REACTOME_TRANSPORT_OF_SMALL_MOLECULES | 634 | 0.43955567 | 1.4682671 | 0 | 0.19970693 | 1 | 3199 | tags=21%, list=12%, signal=23% |
| BIOCARTA_RECK_PATHWAY | 8 | 0.83596855 | 1.4682255 | 0.04545455 | 0.19943577 | 1 | 1774 | tags=63%, list=7%, signal=67% |
| EBAUER_TARGETS_OF_PAX3_FOXO1_FUSION_UP | 173 | 0.511998 | 1.4681522 | 0 | 0.19913672 | 1 | 2892 | tags=22%, list=11%, signal=24% |
| HP_FEMALE_HYPOGONADISM | 21 | 0.7380935 | 1.468024 | 0.02631579 | 0.28981742 | 1 | 533 | tags=10%, list=2%, signal=10% |
| GO_NEURAL_NUCLEUS_DEVELOPMENT | 53 | 0.576351 | 1.4675121 | 0 | 0.29098925 | 1 | 2089 | tags=38%, list=8%, signal=41% |
| HP_ABNORMAL_HEART_MORPHOLOGY | 1064 | 0.43101254 | 1.467262 | 0 | 0.29137704 | 1 | 3528 | tags=26%, list=13%, signal=29% |
| HP_SLOW_PROGRESSION | 153 | 0.5072122 | 1.4671072 | 0 | 0.29146567 | 1 | 3437 | tags=27%, list=13%, signal=31% |
| GO_LACTATE_DEHYDROGENASE_ACTIVITY | 4 | 0.9588898 | 1.4670151 | 0.02564103 | 0.29135087 | 1 | 44 | tags=25%, list=0%, signal=25% |
| GO_EXOCRINE_SYSTEM_DEVELOPMENT | 47 | 0.61058 | 1.4661018 | 0.025 | 0.29381257 | 1 | 3317 | tags=30%, list=12%, signal=34% |
| HP_HYPOSMIA | 43 | 0.62809706 | 1.4655837 | 0.025 | 0.2943866 | 1 | 2338 | tags=23%, list=9%, signal=25% |
| HP_ABNORMALITY_OF_VISION | 1021 | 0.43727237 | 1.4655712 | 0 | 0.294084 | 1 | 4103 | tags=29%, list=15%, signal=33% |
| WAKABAYASHI_ADIPOGENESIS_PPARG_RXRA_BOUND_WITH_H4K20ME1_MARK | 136 | 0.51338917 | 1.4655468 | 0 | 0.20398545 | 1 | 3911 | tags=46%, list=15%, signal=53% |
| GO_NUCLEOSIDE_TRIPHOSPHATE_METABOLIC_PROCESS | 99 | 0.5544293 | 1.4653794 | 0 | 0.29426312 | 1 | 3276 | tags=37%, list=12%, signal=42% |
| BERNARD_PPAPDC1B_TARGETS_UP | 35 | 0.6006102 | 1.4652461 | 0.02777778 | 0.20432228 | 1 | 3202 | tags=46%, list=12%, signal=52% |
| GO_ER_TO_GOLGI_TRANSPORT_VESICLE_MEMBRANE | 46 | 0.6018483 | 1.4651346 | 0.025 | 0.29477543 | 1 | 2116 | tags=37%, list=8%, signal=40% |
| VANTVEER_BREAST_CANCER_POOR_PROGNOSIS | 45 | 0.58423567 | 1.4651288 | 0 | 0.20435135 | 1 | 2518 | tags=27%, list=9%, signal=29% |
| GO_SIGNAL_RECOGNITION_PARTICLE | 7 | 0.9013309 | 1.465081 | 0 | 0.2946569 | 1 | 2612 | tags=86%, list=10%, signal=95% |
| WP_PROXIMAL_TUBULE_TRANSPORT | 48 | 0.594774 | 1.4650387 | 0 | 0.20413733 | 1 | 1716 | tags=17%, list=6%, signal=18% |
| MORI_MATURE_B_LYMPHOCYTE_DN | 69 | 0.5660237 | 1.4645705 | 0.03571429 | 0.20483172 | 1 | 2792 | tags=42%, list=10%, signal=47% |
| GO_REGULATION_OF_EXTRACELLULAR_MATRIX_ORGANIZATION | 36 | 0.60597426 | 1.4644134 | 0.04545455 | 0.29620713 | 1 | 643 | tags=19%, list=2%, signal=20% |
| GO_7S_RNA_BINDING | 5 | 0.9158725 | 1.4643538 | 0 | 0.29599625 | 1 | 752 | tags=80%, list=3%, signal=82% |
| GO_POSITIVE_REGULATION_OF_INTEGRIN_MEDIATED_SIGNALING_PATHWAY | 6 | 0.8688761 | 1.4642471 | 0.02439024 | 0.29595137 | 1 | 2030 | tags=67%, list=8%, signal=72% |
| REACTOME_UCH_PROTEINASES | 82 | 0.5581025 | 1.4642385 | 0.02325581 | 0.20501105 | 1 | 900 | tags=26%, list=3%, signal=26% |
| HP_DIPLOPIA | 68 | 0.5659727 | 1.4641532 | 0.02777778 | 0.2959417 | 1 | 2302 | tags=22%, list=9%, signal=24% |
| GO_CATION_TRANSMEMBRANE_TRANSPORT | 751 | 0.438738 | 1.4637012 | 0 | 0.2965997 | 1 | 3463 | tags=19%, list=13%, signal=21% |
| GO_ANTIGEN_PROCESSING_AND_PRESENTATION_OF_PEPTIDE_ANTIGEN_VIA_MHC_CLASS_IB | 4 | 0.96236503 | 1.4634461 | 0 | 0.29693818 | 1 | 292 | tags=75%, list=1%, signal=76% |
| VALK_AML_WITH_FLT3_ITD | 30 | 0.63676995 | 1.4632992 | 0.02272727 | 0.2060628 | 1 | 1449 | tags=20%, list=5%, signal=21% |
| GO_EPITHELIAL_STRUCTURE_MAINTENANCE | 20 | 0.76284194 | 1.4629991 | 0.02941177 | 0.29793182 | 1 | 267 | tags=10%, list=1%, signal=10% |
| BYSTRYKH_HEMATOPOIESIS_STEM_CELL_AND_BRAIN_QTL_CIS | 63 | 0.58614224 | 1.4628022 | 0 | 0.2067524 | 1 | 2789 | tags=51%, list=10%, signal=57% |
| SENGUPTA_NASOPHARYNGEAL_CARCINOMA_DN | 269 | 0.48345223 | 1.4623966 | 0 | 0.20680234 | 1 | 5412 | tags=30%, list=20%, signal=37% |
| GO_POSITIVE_REGULATION_OF_CHONDROCYTE_PROLIFERATION | 4 | 0.98306787 | 1.4623746 | 0 | 0.29950187 | 1 | 267 | tags=25%, list=1%, signal=25% |
| GO_CATALYTIC_COMPLEX | 1282 | 0.42412537 | 1.4623097 | 0 | 0.29941693 | 1 | 3260 | tags=29%, list=12%, signal=32% |
| HP_RENAL_POTASSIUM_WASTING | 2 | 0.9863736 | 1.462251 | 0.03846154 | 0.29929543 | 1 | 161 | tags=50%, list=1%, signal=50% |
| GO_CATION_TRANSPORT | 990 | 0.4392152 | 1.4622025 | 0 | 0.2991018 | 1 | 2737 | tags=16%, list=10%, signal=17% |
| HP_ABNORMAL_TISSUE_METABOLITE_CONCENTRATION | 28 | 0.6700947 | 1.4621748 | 0 | 0.2988541 | 1 | 3524 | tags=50%, list=13%, signal=57% |
| GO_GLYCOSIDE_CATABOLIC_PROCESS | 7 | 0.87350905 | 1.4621034 | 0.02325581 | 0.29871497 | 1 | 3410 | tags=100%, list=13%, signal=114% |
| GO_DETERMINATION_OF_ADULT_LIFESPAN | 13 | 0.7490174 | 1.4620409 | 0.04761905 | 0.29874063 | 1 | 3392 | tags=38%, list=13%, signal=44% |
| SMID_BREAST_CANCER_BASAL_UP | 561 | 0.4550417 | 1.46177 | 0 | 0.20787789 | 1 | 2531 | tags=22%, list=9%, signal=23% |
| GO_REGULATION_OF_LYMPHOID_PROGENITOR_CELL_DIFFERENTIATION | 7 | 0.8541843 | 1.4617542 | 0.02272727 | 0.29945096 | 1 | 167 | tags=29%, list=1%, signal=29% |
| HP_EMG_IMPAIRED_NEUROMUSCULAR_TRANSMISSION | 29 | 0.6179281 | 1.4616048 | 0.02173913 | 0.29903552 | 1 | 4184 | tags=28%, list=16%, signal=33% |
| GO_REGULATION_OF_ASTROCYTE_DIFFERENTIATION | 25 | 0.716327 | 1.4613994 | 0 | 0.29947406 | 1 | 2318 | tags=36%, list=9%, signal=39% |
| GO_REGULATION_OF_IMMUNE_SYSTEM_PROCESS | 1213 | 0.4257047 | 1.461373 | 0 | 0.2992821 | 1 | 2604 | tags=19%, list=10%, signal=20% |
| WP_ENDOCHONDRAL_OSSIFICATION_WITH_SKELETAL_DYSPLASIAS | 60 | 0.5540544 | 1.4612663 | 0 | 0.20817155 | 1 | 2686 | tags=33%, list=10%, signal=37% |
| MORI_EMU_MYC_LYMPHOMA_BY_ONSET_TIME_DN | 16 | 0.813243 | 1.4606622 | 0.03030303 | 0.20857346 | 1 | 3088 | tags=44%, list=11%, signal=49% |
| KEGG_EPITHELIAL_CELL_SIGNALING_IN_HELICOBACTER_PYLORI_INFECTION | 67 | 0.5701497 | 1.4605448 | 0.02439024 | 0.20844544 | 1 | 1488 | tags=24%, list=6%, signal=25% |
| BIOCARTA_IFNA_PATHWAY | 6 | 0.91742337 | 1.4602759 | 0.02040816 | 0.20830514 | 1 | 2202 | tags=83%, list=8%, signal=91% |
| GO_REGULATION_OF_TYPE_I_INTERFERON_MEDIATED_SIGNALING_PATHWAY | 31 | 0.6400108 | 1.4602548 | 0 | 0.3028395 | 1 | 2468 | tags=42%, list=9%, signal=46% |
| GO_DIOL_BIOSYNTHETIC_PROCESS | 17 | 0.8010137 | 1.4601259 | 0.04761905 | 0.30303672 | 1 | 3272 | tags=59%, list=12%, signal=67% |
| GO_NEGATIVE_REGULATION_OF_MESENCHYMAL_CELL_PROLIFERATION | 7 | 0.87018365 | 1.4599057 | 0.02040816 | 0.30304188 | 1 | 1507 | tags=57%, list=6%, signal=61% |
| BLANCO_MELO_BETA_INTERFERON_TREATED_BRONCHIAL_EPITHELIAL_CELLS_DN | 141 | 0.5046755 | 1.4594871 | 0 | 0.20968461 | 1 | 3928 | tags=25%, list=15%, signal=29% |
| HP_RETINAL_DYSTROPHY | 248 | 0.5033639 | 1.4594828 | 0 | 0.30381534 | 1 | 3524 | tags=27%, list=13%, signal=31% |
| GO_PROTEIN_CONTAINING_COMPLEX_SUBUNIT_ORGANIZATION | 1686 | 0.4215497 | 1.4592584 | 0 | 0.3042076 | 1 | 2481 | tags=24%, list=9%, signal=24% |
| HP_MOTOR_DELAY | 509 | 0.4506623 | 1.4592582 | 0 | 0.30386958 | 1 | 3189 | tags=24%, list=12%, signal=27% |
| WEST_ADRENOCORTICAL_TUMOR_DN | 482 | 0.44975284 | 1.459149 | 0 | 0.20997329 | 1 | 3305 | tags=26%, list=12%, signal=29% |
| LIM_MAMMARY_LUMINAL_MATURE_DN | 92 | 0.55428547 | 1.4590919 | 0 | 0.209637 | 1 | 2830 | tags=35%, list=11%, signal=39% |
| HP_SPLENOMEGALY | 295 | 0.4718808 | 1.4589986 | 0 | 0.30431515 | 1 | 3464 | tags=26%, list=13%, signal=30% |
| HP_VISUAL_IMPAIRMENT | 837 | 0.44105604 | 1.4588181 | 0 | 0.3046189 | 1 | 3907 | tags=28%, list=15%, signal=31% |
| NAKAMURA_ADIPOGENESIS_LATE_DN | 32 | 0.66189754 | 1.4585264 | 0 | 0.21012731 | 1 | 2882 | tags=31%, list=11%, signal=35% |
| MILI_PSEUDOPODIA_CHEMOTAXIS_UP | 81 | 0.5769218 | 1.4582586 | 0 | 0.21049711 | 1 | 3894 | tags=53%, list=14%, signal=62% |
| HP_WEAKNESS_OF_MUSCLES_OF_RESPIRATION | 95 | 0.54357684 | 1.4581555 | 0 | 0.306284 | 1 | 3741 | tags=35%, list=14%, signal=40% |
| HP_MENTAL_DETERIORATION | 282 | 0.4860415 | 1.4581178 | 0 | 0.30608603 | 1 | 2421 | tags=22%, list=9%, signal=24% |
| MYLLYKANGAS_AMPLIFICATION_HOT_SPOT_7 | 6 | 0.8615366 | 1.4579225 | 0.02439024 | 0.21081011 | 1 | 2761 | tags=50%, list=10%, signal=56% |
| VART_KSHV_INFECTION_ANGIOGENIC_MARKERS_DN | 116 | 0.5417628 | 1.4577318 | 0 | 0.21073805 | 1 | 2846 | tags=19%, list=11%, signal=21% |
| GO_EPITHELIAL_CELL_DIFFERENTIATION | 530 | 0.44974855 | 1.4576262 | 0 | 0.3074286 | 1 | 3281 | tags=21%, list=12%, signal=24% |
| GO_REGULATION_OF_COAGULATION | 71 | 0.55303967 | 1.4575045 | 0.02702703 | 0.30746123 | 1 | 1346 | tags=15%, list=5%, signal=16% |
| GO_ACTIVATION_OF_INNATE_IMMUNE_RESPONSE | 123 | 0.53161234 | 1.4575034 | 0 | 0.30712265 | 1 | 2281 | tags=24%, list=8%, signal=26% |
| GO_SERINE_FAMILY_AMINO_ACID_CATABOLIC_PROCESS | 10 | 0.75637066 | 1.4572003 | 0.02222222 | 0.30791178 | 1 | 1863 | tags=30%, list=7%, signal=32% |
| GO_FASCIA_ADHERENS | 9 | 0.79941237 | 1.457155 | 0.04444445 | 0.30781978 | 1 | 2334 | tags=33%, list=9%, signal=36% |
| SHEPARD_BMYB_MORPHOLINO_UP | 200 | 0.49142638 | 1.457128 | 0 | 0.21163541 | 1 | 2524 | tags=29%, list=9%, signal=31% |
| HP_ABNORMAL_INVOLUNTARY_EYE_MOVEMENTS | 850 | 0.43324006 | 1.4570208 | 0 | 0.30820188 | 1 | 3535 | tags=26%, list=13%, signal=29% |
| KONDO_PROSTATE_CANCER_WITH_H3K27ME3 | 112 | 0.51691306 | 1.4569772 | 0 | 0.21141666 | 1 | 2751 | tags=13%, list=10%, signal=15% |
| REACTOME_TCF_DEPENDENT_SIGNALING_IN_RESPONSE_TO_WNT | 165 | 0.5001262 | 1.4569752 | 0 | 0.21108124 | 1 | 1624 | tags=21%, list=6%, signal=22% |
| ENK_UV_RESPONSE_KERATINOCYTE_UP | 473 | 0.46198103 | 1.4563485 | 0 | 0.21186872 | 1 | 3067 | tags=33%, list=11%, signal=36% |
| GO_FATTY_ACID_BINDING | 33 | 0.62495226 | 1.4557908 | 0.05 | 0.31109935 | 1 | 1678 | tags=18%, list=6%, signal=19% |
| HP_DYSGERMINOMA | 6 | 0.8506376 | 1.4557077 | 0.04255319 | 0.3109688 | 1 | 85 | tags=17%, list=0%, signal=17% |
| HP_ABNORMALITY_OF_THE_PROTEIN_C_ANTICOAGULANT_PATHWAY | 4 | 0.9158754 | 1.4553173 | 0.04761905 | 0.31188914 | 1 | 1273 | tags=50%, list=5%, signal=52% |
| HP_CHOREA | 184 | 0.49639857 | 1.4552895 | 0 | 0.31160173 | 1 | 1795 | tags=21%, list=7%, signal=23% |
| RODWELL_AGING_KIDNEY_UP | 423 | 0.4728483 | 1.4552039 | 0 | 0.21417326 | 1 | 2076 | tags=20%, list=8%, signal=21% |
| LIM_MAMMARY_STEM_CELL_DN | 379 | 0.46949658 | 1.4550248 | 0 | 0.21403708 | 1 | 4519 | tags=33%, list=17%, signal=39% |
| TURASHVILI_BREAST_DUCTAL_CARCINOMA_VS_DUCTAL_NORMAL_DN | 160 | 0.48642033 | 1.4549785 | 0 | 0.2138172 | 1 | 2115 | tags=14%, list=8%, signal=16% |
| REACTOME_SYNTHESIS_SECRETION_AND_INACTIVATION_OF_GLUCAGON_LIKE_PEPTIDE_1_GLP_1_ | 13 | 0.83514 | 1.4547882 | 0.04761905 | 0.21346352 | 1 | 351 | tags=23%, list=1%, signal=23% |
| GO_PURINE_NUCLEOBASE_TRANSMEMBRANE_TRANSPORTER_ACTIVITY | 4 | 0.97473097 | 1.4546716 | 0 | 0.31267336 | 1 | 171 | tags=50%, list=1%, signal=50% |
| HP_AMYLOIDOSIS | 20 | 0.72004014 | 1.4543842 | 0.02380952 | 0.31331006 | 1 | 444 | tags=20%, list=2%, signal=20% |
| GO_LYMPHOID_PROGENITOR_CELL_DIFFERENTIATION | 18 | 0.7451405 | 1.454271 | 0.02222222 | 0.31340522 | 1 | 1181 | tags=22%, list=4%, signal=23% |
| HP_MIXED_DEMYELINATING_AND_AXONAL_POLYNEUROPATHY | 8 | 0.9041941 | 1.4541112 | 0.02325581 | 0.3135504 | 1 | 854 | tags=63%, list=3%, signal=65% |
| GO_NATURAL_KILLER_CELL_LECTIN_LIKE_RECEPTOR_BINDING | 4 | 0.95712215 | 1.4540857 | 0.02222222 | 0.31324568 | 1 | 228 | tags=25%, list=1%, signal=25% |
| GO_RESPONSE_TO_INTERLEUKIN_6 | 38 | 0.6487516 | 1.454008 | 0.025 | 0.31318438 | 1 | 2693 | tags=26%, list=10%, signal=29% |
| GO_CENTRAL_NERVOUS_SYSTEM_DEVELOPMENT | 934 | 0.44773972 | 1.4539956 | 0 | 0.3129331 | 1 | 2371 | tags=18%, list=9%, signal=19% |
| GO_PROTEIN_DISULFIDE_ISOMERASE_ACTIVITY | 15 | 0.7567126 | 1.4538053 | 0.02222222 | 0.31318358 | 1 | 3293 | tags=60%, list=12%, signal=68% |
| RODRIGUES_THYROID_CARCINOMA_ANAPLASTIC_DN | 473 | 0.46224427 | 1.4534401 | 0 | 0.2159392 | 1 | 3980 | tags=34%, list=15%, signal=40% |
| GO_GLYCOSYLCERAMIDE_CATABOLIC_PROCESS | 6 | 0.87347656 | 1.4532257 | 0 | 0.31433663 | 1 | 3410 | tags=100%, list=13%, signal=114% |
| HP_ABNORMAL_CIRCULATING_PHENYLALANINE_CONCENTRATION | 6 | 0.9312065 | 1.4531796 | 0 | 0.3141877 | 1 | 1744 | tags=67%, list=6%, signal=71% |
| GO_SYNAPTIC_GROWTH_AT_NEUROMUSCULAR_JUNCTION | 9 | 0.76857996 | 1.4531791 | 0.04166667 | 0.31385025 | 1 | 570 | tags=22%, list=2%, signal=23% |
| GO_REGULATION_OF_BLOOD_VESSEL_REMODELING | 6 | 0.9237897 | 1.4531003 | 0.04651163 | 0.3137197 | 1 | 2 | tags=17%, list=0%, signal=17% |
| GO_NEGATIVE_REGULATION_OF_AMYLOID_BETA_FORMATION | 13 | 0.745143 | 1.4530343 | 0.03448276 | 0.31353715 | 1 | 2158 | tags=54%, list=8%, signal=59% |
| GO_GERM_CELL_PROLIFERATION | 4 | 0.935895 | 1.4529797 | 0 | 0.3132698 | 1 | 270 | tags=50%, list=1%, signal=50% |
| GO_METANEPHRIC_NEPHRON_TUBULE_EPITHELIAL_CELL_DIFFERENTIATION | 6 | 0.86789894 | 1.4529637 | 0.04347826 | 0.31300306 | 1 | 3104 | tags=50%, list=12%, signal=57% |
| GRAESSMANN_APOPTOSIS_BY_DOXORUBICIN_UP | 1077 | 0.44170335 | 1.4525765 | 0 | 0.21720919 | 1 | 3633 | tags=30%, list=13%, signal=33% |
| MARTINEZ_RB1_TARGETS_UP | 625 | 0.43749046 | 1.4523576 | 0 | 0.21690375 | 1 | 3527 | tags=29%, list=13%, signal=33% |
| CHEOK_RESPONSE_TO_MERCAPTOPURINE_DN | 18 | 0.741097 | 1.4518062 | 0.04761905 | 0.21787779 | 1 | 3608 | tags=44%, list=13%, signal=51% |
| HP_ARTERIAL_THROMBOSIS | 22 | 0.70230526 | 1.4516858 | 0.02439024 | 0.31637934 | 1 | 1967 | tags=27%, list=7%, signal=29% |
| GO_TETRAHYDROBIOPTERIN_METABOLIC_PROCESS | 6 | 0.89772105 | 1.4516854 | 0.04255319 | 0.31604275 | 1 | 2720 | tags=83%, list=10%, signal=93% |
| HP_ABNORMALITY_OF_RENIN_ANGIOTENSIN_SYSTEM | 12 | 0.7998718 | 1.4513737 | 0.04081633 | 0.3163747 | 1 | 173 | tags=17%, list=1%, signal=17% |
| HP_BILATERAL_TONIC_CLONIC_SEIZURE | 176 | 0.5048984 | 1.451334 | 0 | 0.3161912 | 1 | 1790 | tags=16%, list=7%, signal=17% |
| REACTOME_CITRIC_ACID_CYCLE_TCA_CYCLE_ | 21 | 0.69334245 | 1.4512992 | 0.02380952 | 0.21856356 | 1 | 2982 | tags=52%, list=11%, signal=59% |
| SASAI_RESISTANCE_TO_NEOPLASTIC_TRANSFROMATION | 45 | 0.59083915 | 1.4512594 | 0 | 0.21830836 | 1 | 2531 | tags=31%, list=9%, signal=34% |
| BLANCO_MELO_BRONCHIAL_EPITHELIAL_CELLS_INFLUENZA_A_DEL_NS1_INFECTION_UP | 531 | 0.45410633 | 1.4511768 | 0 | 0.21808572 | 1 | 4169 | tags=30%, list=15%, signal=35% |
| BRUINS_UVC_RESPONSE_VIA_TP53_GROUP_B | 496 | 0.44768742 | 1.4508681 | 0 | 0.21851316 | 1 | 3420 | tags=26%, list=13%, signal=29% |
| GO_REGULATION_OF_ATPASE_ACTIVITY | 71 | 0.54841477 | 1.4506909 | 0 | 0.31785908 | 1 | 3095 | tags=35%, list=11%, signal=40% |
| HP_RECURRENT_INFECTION_OF_THE_GASTROINTESTINAL_TRACT | 26 | 0.6821332 | 1.4506263 | 0.04651163 | 0.3178448 | 1 | 2687 | tags=23%, list=10%, signal=26% |
| GO_INTEGRIN_ACTIVATION | 23 | 0.66350865 | 1.4502158 | 0.02631579 | 0.31916848 | 1 | 2555 | tags=30%, list=9%, signal=34% |
| REACTOME_SIGNALING_BY_INSULIN_RECEPTOR | 69 | 0.5462556 | 1.4498475 | 0 | 0.21936391 | 1 | 2162 | tags=25%, list=8%, signal=27% |
| GO_SMALL_MOLECULE_CATABOLIC_PROCESS | 383 | 0.45137575 | 1.4497579 | 0 | 0.32050267 | 1 | 4189 | tags=35%, list=16%, signal=41% |
| HP_SEVERE_MUSCULAR_HYPOTONIA | 66 | 0.5694678 | 1.4496803 | 0.02272727 | 0.32033363 | 1 | 3524 | tags=35%, list=13%, signal=40% |
| RUAN_RESPONSE_TO_TNF_UP | 8 | 0.8877028 | 1.4495974 | 0 | 0.21961999 | 1 | 1639 | tags=50%, list=6%, signal=53% |
| GO_S_ADENOSYLMETHIONINE_METABOLIC_PROCESS | 14 | 0.76755387 | 1.4494725 | 0.0212766 | 0.32087526 | 1 | 4257 | tags=57%, list=16%, signal=68% |
| TOOKER_GEMCITABINE_RESISTANCE_UP | 70 | 0.5281117 | 1.4491897 | 0 | 0.22001368 | 1 | 2703 | tags=36%, list=10%, signal=40% |
| HP_MALE_PSEUDOHERMAPHRODITISM | 33 | 0.6472754 | 1.4489932 | 0 | 0.32174915 | 1 | 4359 | tags=42%, list=16%, signal=51% |
| GO_CELL_ACTIVATION_INVOLVED_IN_IMMUNE_RESPONSE | 605 | 0.43701014 | 1.4489647 | 0 | 0.3215128 | 1 | 2693 | tags=21%, list=10%, signal=23% |
| GO_LEUKOCYTE_CHEMOTAXIS_INVOLVED_IN_INFLAMMATORY_RESPONSE | 4 | 0.96685743 | 1.4487997 | 0 | 0.32167938 | 1 | 165 | tags=50%, list=1%, signal=50% |
| HP_REDUCED_TENDON_REFLEXES | 458 | 0.45190778 | 1.4487131 | 0 | 0.32172775 | 1 | 3537 | tags=27%, list=13%, signal=30% |
| GO_REGULATION_OF_FOREBRAIN_NEURON_DIFFERENTIATION | 4 | 0.98152196 | 1.4486862 | 0.0212766 | 0.32140833 | 1 | 167 | tags=50%, list=1%, signal=50% |
| GO_REGULATION_OF_CHONDROCYTE_DIFFERENTIATION | 42 | 0.6021724 | 1.4484607 | 0 | 0.32175753 | 1 | 1773 | tags=19%, list=7%, signal=20% |
| PASTURAL_RIZ1_TARGETS_UP | 7 | 0.8341771 | 1.4483601 | 0.04545455 | 0.22101942 | 1 | 2457 | tags=57%, list=9%, signal=63% |
| HP_PROXIMAL_RENAL_TUBULAR_ACIDOSIS | 5 | 0.9059668 | 1.448166 | 0 | 0.3222901 | 1 | 73 | tags=20%, list=0%, signal=20% |
| LUCAS_HNF4A_TARGETS_UP | 55 | 0.5794904 | 1.4480239 | 0.04878049 | 0.22143884 | 1 | 4084 | tags=35%, list=15%, signal=41% |
| GO_CARBOHYDRATE_DERIVATIVE_METABOLIC_PROCESS | 967 | 0.4319187 | 1.4479449 | 0 | 0.32273817 | 1 | 3623 | tags=28%, list=13%, signal=31% |
| HP_DICARBOXYLIC_ACIDURIA | 36 | 0.607725 | 1.447768 | 0.04761905 | 0.3226674 | 1 | 4505 | tags=61%, list=17%, signal=73% |
| MORI_PRE_BI_LYMPHOCYTE_DN | 72 | 0.5448338 | 1.4476033 | 0 | 0.22202174 | 1 | 3224 | tags=31%, list=12%, signal=35% |
| GO_PI_BODY | 5 | 0.850558 | 1.4475789 | 0.04761905 | 0.32249662 | 1 | 2872 | tags=20%, list=11%, signal=22% |
| HP_ABNORMALITY_OF_PROTHROMBIN | 28 | 0.6508056 | 1.4474275 | 0.04255319 | 0.32261023 | 1 | 3222 | tags=39%, list=12%, signal=45% |
| GO_CELLULAR_OXIDANT_DETOXIFICATION | 83 | 0.5423891 | 1.4472877 | 0 | 0.32207382 | 1 | 2750 | tags=27%, list=10%, signal=29% |
| HAN_SATB1_TARGETS_DN | 341 | 0.46658757 | 1.4471836 | 0 | 0.22240949 | 1 | 3099 | tags=26%, list=12%, signal=29% |
| HP_HYPOPROTEINEMIA | 11 | 0.8302087 | 1.4471304 | 0.025 | 0.32202104 | 1 | 6 | tags=9%, list=0%, signal=9% |
| GO_PROTEIN_PROCESSING | 183 | 0.49045387 | 1.4468561 | 0 | 0.32213038 | 1 | 3061 | tags=25%, list=11%, signal=28% |
| GO_VERY_LONG_CHAIN_FATTY_ACID_METABOLIC_PROCESS | 31 | 0.6795357 | 1.4467571 | 0.05 | 0.32222787 | 1 | 2167 | tags=26%, list=8%, signal=28% |
| GO_DIPEPTIDASE_ACTIVITY | 11 | 0.7368354 | 1.4466617 | 0.02777778 | 0.3220782 | 1 | 424 | tags=18%, list=2%, signal=18% |
| REACTOME_HIV_INFECTION | 213 | 0.48459777 | 1.446565 | 0 | 0.22299346 | 1 | 2395 | tags=31%, list=9%, signal=34% |
| ZHAN_LATE_DIFFERENTIATION_GENES_UP | 31 | 0.5989474 | 1.4465371 | 0 | 0.22265406 | 1 | 2998 | tags=39%, list=11%, signal=44% |
| GO_MYELOID_LEUKOCYTE_ACTIVATION | 575 | 0.44466788 | 1.4463006 | 0 | 0.32302898 | 1 | 2465 | tags=21%, list=9%, signal=22% |
| HP_RECURRENT_ASPERGILLUS_INFECTIONS | 4 | 0.8941559 | 1.4461814 | 0.02631579 | 0.32312396 | 1 | 72 | tags=25%, list=0%, signal=25% |
| WP_SARS_CORONAVIRUS_AND_INNATE_IMMUNITY | 16 | 0.762381 | 1.4461582 | 0.02564103 | 0.22301362 | 1 | 2378 | tags=56%, list=9%, signal=62% |
| HP_DYSPAREUNIA | 30 | 0.5924788 | 1.4456375 | 0 | 0.32436368 | 1 | 4982 | tags=40%, list=19%, signal=49% |
| GO_NEGATIVE_REGULATION_OF_MITOCHONDRION_ORGANIZATION | 56 | 0.5884495 | 1.4455982 | 0.02857143 | 0.32413092 | 1 | 804 | tags=18%, list=3%, signal=18% |
| GO_HUMORAL_IMMUNE_RESPONSE | 142 | 0.5014466 | 1.4455936 | 0 | 0.3238005 | 1 | 2587 | tags=18%, list=10%, signal=19% |
| ALFANO_MYC_TARGETS | 224 | 0.4801152 | 1.4455734 | 0 | 0.22406428 | 1 | 3724 | tags=40%, list=14%, signal=46% |
| NABA_SECRETED_FACTORS | 230 | 0.47179192 | 1.445336 | 0 | 0.22411136 | 1 | 3796 | tags=18%, list=14%, signal=21% |
| ACEVEDO_NORMAL_TISSUE_ADJACENT_TO_LIVER_TUMOR_UP | 148 | 0.47981188 | 1.4453117 | 0 | 0.22380027 | 1 | 2780 | tags=34%, list=10%, signal=38% |
| CHUNG_BLISTER_CYTOTOXICITY_UP | 119 | 0.5250549 | 1.4452593 | 0.03125 | 0.22349079 | 1 | 3369 | tags=39%, list=13%, signal=45% |
| GO_RETINA_HOMEOSTASIS | 60 | 0.56794256 | 1.445211 | 0.02325581 | 0.32451382 | 1 | 613 | tags=8%, list=2%, signal=9% |
| REACTOME_HEDGEHOG_OFF_STATE | 105 | 0.5635939 | 1.4450204 | 0 | 0.22365138 | 1 | 805 | tags=22%, list=3%, signal=22% |
| GARGALOVIC_RESPONSE_TO_OXIDIZED_PHOSPHOLIPIDS_PINK_DN | 29 | 0.6276893 | 1.445015 | 0 | 0.22328772 | 1 | 4882 | tags=52%, list=18%, signal=63% |
| GO_ENSHEATHMENT_OF_NEURONS | 134 | 0.49500838 | 1.4448086 | 0 | 0.3248765 | 1 | 1295 | tags=15%, list=5%, signal=16% |
| GO_INTERLEUKIN_35_MEDIATED_SIGNALING_PATHWAY | 10 | 0.84121764 | 1.44474 | 0 | 0.32447797 | 1 | 1939 | tags=30%, list=7%, signal=32% |
| GO_ENDOPEPTIDASE_ACTIVITY | 324 | 0.45541286 | 1.4447172 | 0 | 0.32421398 | 1 | 3436 | tags=18%, list=13%, signal=21% |
| REACTOME_FATTY_ACIDS | 6 | 0.8738067 | 1.444655 | 0.04347826 | 0.22366539 | 1 | 694 | tags=17%, list=3%, signal=17% |
| HP_ABNORMALITY_OF_THE_PARATHYROID_GLAND | 63 | 0.5585154 | 1.4445632 | 0.02631579 | 0.32437205 | 1 | 3197 | tags=27%, list=12%, signal=31% |
| GO_PROTEASE_BINDING | 109 | 0.52314764 | 1.4440007 | 0 | 0.32572097 | 1 | 2232 | tags=28%, list=8%, signal=31% |
| GO_REGULATION_OF_COLLAGEN_METABOLIC_PROCESS | 36 | 0.64732337 | 1.4439287 | 0 | 0.32516143 | 1 | 316 | tags=8%, list=1%, signal=8% |
| HP_PARONYCHIA | 8 | 0.8518666 | 1.4439156 | 0.04545455 | 0.32486662 | 1 | 1939 | tags=25%, list=7%, signal=27% |
| GO_REGULATION_OF_EPIDERMAL_CELL_DIFFERENTIATION | 49 | 0.58891875 | 1.4438004 | 0.03225806 | 0.3249421 | 1 | 3514 | tags=33%, list=13%, signal=37% |
| KEGG_OTHER_GLYCAN_DEGRADATION | 14 | 0.78962624 | 1.4434125 | 0.025 | 0.22598572 | 1 | 3112 | tags=50%, list=12%, signal=57% |
| GO_REGULATION_OF_INFLAMMATORY_RESPONSE_TO_ANTIGENIC_STIMULUS | 21 | 0.7170069 | 1.4434104 | 0.02 | 0.3255109 | 1 | 1765 | tags=33%, list=7%, signal=36% |
| BURTON_ADIPOGENESIS_8 | 76 | 0.5409725 | 1.4432988 | 0 | 0.2258941 | 1 | 3667 | tags=43%, list=14%, signal=50% |
| GO_INTERLEUKIN_23_MEDIATED_SIGNALING_PATHWAY | 6 | 0.8477723 | 1.4432813 | 0.04166667 | 0.32547367 | 1 | 1939 | tags=33%, list=7%, signal=36% |
| GO_FEEDING_BEHAVIOR | 80 | 0.5567063 | 1.4432374 | 0 | 0.32538858 | 1 | 2530 | tags=16%, list=9%, signal=18% |
| GO_ACTIN_CYTOSKELETON | 445 | 0.4499683 | 1.4431022 | 0 | 0.32512277 | 1 | 3390 | tags=25%, list=13%, signal=29% |
| HP_PROLONGED_BLEEDING_FOLLOWING_PROCEDURE | 14 | 0.73714256 | 1.4430214 | 0.04166667 | 0.32507017 | 1 | 4480 | tags=43%, list=17%, signal=51% |
| GO_POSITIVE_REGULATION_OF_RESPONSE_TO_EXTERNAL_STIMULUS | 434 | 0.4598466 | 1.4426941 | 0 | 0.32596025 | 1 | 2555 | tags=18%, list=9%, signal=20% |
| BAELDE_DIABETIC_NEPHROPATHY_DN | 416 | 0.45895752 | 1.44239 | 0 | 0.22639482 | 1 | 2350 | tags=24%, list=9%, signal=26% |
| BOCHKIS_FOXA2_TARGETS | 368 | 0.46514255 | 1.4422393 | 0 | 0.22630244 | 1 | 4009 | tags=30%, list=15%, signal=35% |
| GO_IMMUNE_SYSTEM_DEVELOPMENT | 858 | 0.4267253 | 1.4416232 | 0 | 0.3288739 | 1 | 2692 | tags=20%, list=10%, signal=21% |
| WANG_LMO4_TARGETS_DN | 330 | 0.46935365 | 1.4413227 | 0 | 0.2276022 | 1 | 3670 | tags=37%, list=14%, signal=43% |
| HP_GONADAL_DYSGENESIS | 26 | 0.64268064 | 1.441065 | 0.03921569 | 0.3304763 | 1 | 2301 | tags=31%, list=9%, signal=34% |
| HP_ECTOPIC_CALCIFICATION | 166 | 0.5105599 | 1.4409784 | 0 | 0.33048218 | 1 | 3128 | tags=30%, list=12%, signal=34% |
| MONNIER_POSTRADIATION_TUMOR_ESCAPE_DN | 349 | 0.46107972 | 1.4407611 | 0 | 0.22836006 | 1 | 3306 | tags=35%, list=12%, signal=39% |
| JAEGER_METASTASIS_DN | 192 | 0.48512256 | 1.440671 | 0 | 0.22821312 | 1 | 3467 | tags=18%, list=13%, signal=20% |
| JAZAG_TGFB1_SIGNALING_UP | 93 | 0.5129946 | 1.4404786 | 0 | 0.2281177 | 1 | 3095 | tags=27%, list=11%, signal=30% |
| GO_NEGATIVE_REGULATION_OF_LEUKOCYTE_MEDIATED_CYTOTOXICITY | 14 | 0.7771176 | 1.4401965 | 0.04651163 | 0.3326586 | 1 | 2389 | tags=29%, list=9%, signal=31% |
| WP_SYNTHESIS_AND_DEGRADATION_OF_KETONE_BODIES | 4 | 0.9069832 | 1.4400182 | 0.0212766 | 0.22871916 | 1 | 541 | tags=50%, list=2%, signal=51% |
| VICENT_METASTASIS_UP | 12 | 0.76191384 | 1.4399117 | 0.05 | 0.22859678 | 1 | 2941 | tags=50%, list=11%, signal=56% |
| HP_RECURRENT_BRONCHITIS | 26 | 0.65601474 | 1.4399033 | 0.02941177 | 0.3326819 | 1 | 4144 | tags=35%, list=15%, signal=41% |
| KEGG_CYSTEINE_AND_METHIONINE_METABOLISM | 29 | 0.66092443 | 1.4397218 | 0.04651163 | 0.22850242 | 1 | 1089 | tags=14%, list=4%, signal=14% |
| HOSHIDA_LIVER_CANCER_SURVIVAL_DN | 101 | 0.5363316 | 1.439664 | 0 | 0.22786093 | 1 | 3672 | tags=32%, list=14%, signal=37% |
| CHEN_HOXA5_TARGETS_9HR_DN | 38 | 0.6290102 | 1.4393619 | 0 | 0.22824799 | 1 | 2392 | tags=37%, list=9%, signal=40% |
| HP_PROGRESSIVE_NEUROLOGIC_DETERIORATION | 41 | 0.620636 | 1.439265 | 0 | 0.3342533 | 1 | 3441 | tags=49%, list=13%, signal=56% |
| REACTOME_REGULATION_OF_RUNX3_EXPRESSION_AND_ACTIVITY | 53 | 0.610649 | 1.439196 | 0 | 0.22800994 | 1 | 1025 | tags=38%, list=4%, signal=39% |
| GO_UNSATURATED_FATTY_ACID_METABOLIC_PROCESS | 82 | 0.5411587 | 1.4389547 | 0.03225806 | 0.3346996 | 1 | 2191 | tags=26%, list=8%, signal=28% |
| HP_ANISOCYTOSIS | 12 | 0.7932809 | 1.4389178 | 0.04545455 | 0.3344488 | 1 | 2364 | tags=42%, list=9%, signal=46% |
| GO_ENDOPLASMIC_RETICULUM_GOLGI_INTERMEDIATE_COMPARTMENT_MEMBRANE | 66 | 0.58215475 | 1.4387147 | 0.02941177 | 0.33448562 | 1 | 2423 | tags=41%, list=9%, signal=45% |
| HP_PERIPHERAL_AXONAL_DEGENERATION | 147 | 0.51315606 | 1.4385784 | 0 | 0.3346612 | 1 | 2076 | tags=22%, list=8%, signal=24% |
| GO_TRANSMEMBRANE_RECEPTOR_PROTEIN_TYROSINE_PHOSPHATASE_SIGNALING_PATHWAY | 5 | 0.89865136 | 1.4384902 | 0 | 0.33461544 | 1 | 150 | tags=40%, list=1%, signal=40% |
| LIEN_BREAST_CARCINOMA_METAPLASTIC_VS_DUCTAL_DN | 80 | 0.54363614 | 1.4384694 | 0 | 0.22893848 | 1 | 2852 | tags=16%, list=11%, signal=18% |
| REACTOME_C_TYPE_LECTIN_RECEPTORS_CLRS_ | 127 | 0.5044662 | 1.4382557 | 0 | 0.22864972 | 1 | 2389 | tags=25%, list=9%, signal=28% |
| GO_POSITIVE_REGULATION_BY_SYMBIONT_OF_ENTRY_INTO_HOST | 7 | 0.87180537 | 1.4377265 | 0.02272727 | 0.33661208 | 1 | 3029 | tags=71%, list=11%, signal=80% |
| GO_POSITIVE_REGULATION_OF_CEREBELLAR_GRANULE_CELL_PRECURSOR_PROLIFERATION | 6 | 0.88949287 | 1.4376091 | 0 | 0.33665913 | 1 | 377 | tags=17%, list=1%, signal=17% |
| GO_RESPONSE_TO_CYTOKINE | 1001 | 0.4216618 | 1.4374547 | 0 | 0.33679953 | 1 | 3182 | tags=24%, list=12%, signal=26% |
| HP_WEAKNESS_OF_FACIAL_MUSCULATURE | 205 | 0.46924025 | 1.4367957 | 0 | 0.3387232 | 1 | 3776 | tags=28%, list=14%, signal=33% |
| IVANOVA_HEMATOPOIESIS_INTERMEDIATE_PROGENITOR | 125 | 0.49043593 | 1.4366038 | 0 | 0.23236805 | 1 | 3786 | tags=38%, list=14%, signal=44% |
| ZHONG_RESPONSE_TO_AZACITIDINE_AND_TSA_UP | 146 | 0.4890084 | 1.4365163 | 0 | 0.23180476 | 1 | 3667 | tags=31%, list=14%, signal=35% |
| APRELIKOVA_BRCA1_TARGETS | 47 | 0.58067155 | 1.4364061 | 0 | 0.23176003 | 1 | 2938 | tags=32%, list=11%, signal=36% |
| WEI_MYCN_TARGETS_WITH_E_BOX | 734 | 0.43527135 | 1.4362206 | 0 | 0.23187311 | 1 | 3214 | tags=32%, list=12%, signal=35% |
| HP_CONGENITAL_BLINDNESS | 7 | 0.9035884 | 1.4361633 | 0.02173913 | 0.33958825 | 1 | 912 | tags=29%, list=3%, signal=30% |
| CERIBELLI_GENES_INACTIVE_AND_BOUND_BY_NFY | 23 | 0.7009617 | 1.4360745 | 0 | 0.23172244 | 1 | 2515 | tags=26%, list=9%, signal=29% |
| PLASARI_NFIC_TARGETS_BASAL_UP | 24 | 0.67000514 | 1.4359341 | 0.04651163 | 0.23159969 | 1 | 3997 | tags=38%, list=15%, signal=44% |
| GO_PROTEIN_MATURATION | 244 | 0.47647545 | 1.4359266 | 0 | 0.34022543 | 1 | 2874 | tags=26%, list=11%, signal=29% |
| TAKEDA_TARGETS_OF_NUP98_HOXA9_FUSION_10D_DN | 114 | 0.5240181 | 1.4354395 | 0.02857143 | 0.23203145 | 1 | 2274 | tags=19%, list=8%, signal=21% |
| HOLLMANN_APOPTOSIS_VIA_CD40_UP | 188 | 0.49070916 | 1.4352195 | 0 | 0.2314607 | 1 | 2942 | tags=33%, list=11%, signal=37% |
| GO_LIPID_TRANSPORTER_ACTIVITY | 111 | 0.5328725 | 1.4350033 | 0 | 0.34176037 | 1 | 3795 | tags=24%, list=14%, signal=28% |
| GO_PROTEIN_N_TERMINUS_BINDING | 99 | 0.5121647 | 1.4348097 | 0 | 0.34201834 | 1 | 3524 | tags=42%, list=13%, signal=49% |
| TONKS_TARGETS_OF_RUNX1_RUNX1T1_FUSION_SUSTAINDED_IN_ERYTHROCYTE_UP | 40 | 0.61610943 | 1.4345307 | 0 | 0.23255014 | 1 | 934 | tags=23%, list=3%, signal=23% |
| GO_MITOCHONDRIAL_RESPIRATORY_CHAIN_COMPLEX_IV | 11 | 0.83169454 | 1.43429 | 0 | 0.34278888 | 1 | 1740 | tags=45%, list=6%, signal=49% |
| BOSCO_TH1_CYTOTOXIC_MODULE | 92 | 0.5001406 | 1.4338527 | 0.02702703 | 0.23351151 | 1 | 2225 | tags=16%, list=8%, signal=18% |
| KEGG_BASAL_CELL_CARCINOMA | 48 | 0.5820902 | 1.4338405 | 0.03030303 | 0.23315555 | 1 | 1945 | tags=13%, list=7%, signal=13% |
| GO_NEGATIVE_REGULATION_OF_RELEASE_OF_CYTOCHROME_C_FROM_MITOCHONDRIA | 18 | 0.6888579 | 1.4337922 | 0 | 0.34394026 | 1 | 804 | tags=22%, list=3%, signal=23% |
| HOOI_ST7_TARGETS_DN | 95 | 0.54123354 | 1.4337082 | 0.02564103 | 0.23300792 | 1 | 2993 | tags=21%, list=11%, signal=24% |
| GO_FATTY_ACID_DERIVATIVE_BINDING | 26 | 0.62683386 | 1.4336804 | 0.02040816 | 0.3439639 | 1 | 3481 | tags=42%, list=13%, signal=49% |
| BIOCARTA_EOSINOPHILS_PATHWAY | 3 | 0.9443294 | 1.4336544 | 0.04444445 | 0.23278187 | 1 | 1237 | tags=67%, list=5%, signal=70% |
| HP_ABNORMALITY_OF_THE_NASOLABIAL_REGION | 9 | 0.8157469 | 1.4331367 | 0.04761905 | 0.34483033 | 1 | 1613 | tags=33%, list=6%, signal=35% |
| GO_MYELOID_LEUKOCYTE_MEDIATED_IMMUNITY | 490 | 0.44506162 | 1.4329885 | 0 | 0.34495988 | 1 | 2451 | tags=21%, list=9%, signal=23% |
| MULLIGHAN_MLL_SIGNATURE_1_UP | 350 | 0.44874683 | 1.4329858 | 0 | 0.23366596 | 1 | 3029 | tags=26%, list=11%, signal=29% |
| WELCSH_BRCA1_TARGETS_UP | 183 | 0.47800565 | 1.4329233 | 0 | 0.2334661 | 1 | 2425 | tags=33%, list=9%, signal=36% |
| GO_CEREBELLAR_CORTEX_DEVELOPMENT | 45 | 0.57245964 | 1.4326761 | 0.05 | 0.3454662 | 1 | 2315 | tags=20%, list=9%, signal=22% |
| GO_BEHAVIORAL_RESPONSE_TO_PAIN | 11 | 0.84586906 | 1.432613 | 0.02702703 | 0.34541103 | 1 | 29 | tags=9%, list=0%, signal=9% |
| HP_RIDGED_NAIL | 20 | 0.6805091 | 1.4325057 | 0.04761905 | 0.34546477 | 1 | 2807 | tags=35%, list=10%, signal=39% |
| GO_POSITIVE_REGULATION_OF_BIOMINERALIZATION | 39 | 0.598088 | 1.4324995 | 0.05 | 0.3451351 | 1 | 1334 | tags=15%, list=5%, signal=16% |
| GO_RESPONSE_TO_SELENIUM_ION | 5 | 0.9433303 | 1.4324237 | 0.04878049 | 0.34503496 | 1 | 419 | tags=40%, list=2%, signal=41% |
| CHEBOTAEV_GR_TARGETS_UP | 72 | 0.5548335 | 1.4322364 | 0 | 0.23429038 | 1 | 2008 | tags=22%, list=7%, signal=24% |
| BLANCO_MELO_COVID19_SARS_COV_2_INFECTION_A594_ACE2_EXPRESSING_CELLS_DN | 75 | 0.55270517 | 1.4321709 | 0.02702703 | 0.23416862 | 1 | 2247 | tags=33%, list=8%, signal=36% |
| WP_PATHWAYS_OF_NUCLEIC_ACID_METABOLISM_AND_INNATE_IMMUNE_SENSING | 11 | 0.7749434 | 1.4321666 | 0.04878049 | 0.23381595 | 1 | 4269 | tags=73%, list=16%, signal=86% |
| GAUSSMANN_MLL_AF4_FUSION_TARGETS_D_UP | 32 | 0.6414521 | 1.431228 | 0.02702703 | 0.23598063 | 1 | 1510 | tags=19%, list=6%, signal=20% |
| URS_ADIPOCYTE_DIFFERENTIATION_UP | 56 | 0.58136374 | 1.4311917 | 0.02941177 | 0.23565212 | 1 | 3608 | tags=32%, list=13%, signal=37% |
| REACTOME_RNA_POLYMERASE_III_TRANSCRIPTION_INITIATION_FROM_TYPE_3_PROMOTER | 27 | 0.6522911 | 1.4310185 | 0.02222222 | 0.23570497 | 1 | 2452 | tags=44%, list=9%, signal=49% |
| HOSHIDA_LIVER_CANCER_SUBCLASS_S1 | 227 | 0.48234877 | 1.4304148 | 0 | 0.23608223 | 1 | 2421 | tags=26%, list=9%, signal=28% |
| GO_OLIGODENDROCYTE_DEVELOPMENT | 45 | 0.6132762 | 1.430367 | 0.0212766 | 0.35027152 | 1 | 496 | tags=13%, list=2%, signal=14% |
| GO_ORGANIC_ACID_CATABOLIC_PROCESS | 248 | 0.465572 | 1.4303516 | 0 | 0.35001528 | 1 | 3940 | tags=33%, list=15%, signal=39% |
| CLAUS_PGR_POSITIVE_MENINGIOMA_DN | 11 | 0.81175303 | 1.4302329 | 0.02325581 | 0.23600665 | 1 | 2640 | tags=45%, list=10%, signal=50% |
| LE_EGR2_TARGETS_DN | 99 | 0.5088083 | 1.430171 | 0 | 0.23575643 | 1 | 3058 | tags=30%, list=11%, signal=34% |
| HOEBEKE_LYMPHOID_STEM_CELL_DN | 86 | 0.4975586 | 1.430047 | 0 | 0.23580994 | 1 | 2604 | tags=34%, list=10%, signal=37% |
| GO_ENDOPLASMIC_RETICULUM_GOLGI_INTERMEDIATE_COMPARTMENT | 118 | 0.5305342 | 1.430023 | 0 | 0.35095665 | 1 | 2709 | tags=36%, list=10%, signal=40% |
| BONOME_OVARIAN_CANCER_SURVIVAL_OPTIMAL_DEBULKING | 220 | 0.4849555 | 1.43002 | 0 | 0.23553543 | 1 | 2836 | tags=21%, list=11%, signal=24% |
| CHICAS_RB1_TARGETS_CONFLUENT | 498 | 0.46204105 | 1.4298738 | 0 | 0.23551212 | 1 | 3117 | tags=28%, list=12%, signal=31% |
| GO_INSULIN_RECEPTOR_SIGNALING_PATHWAY | 128 | 0.5077712 | 1.4294666 | 0 | 0.35239807 | 1 | 2614 | tags=30%, list=10%, signal=33% |
| HP_MEMORY_IMPAIRMENT | 106 | 0.5212923 | 1.429438 | 0.03030303 | 0.35218576 | 1 | 3100 | tags=28%, list=12%, signal=32% |
| GO_METALLOCHAPERONE_ACTIVITY | 5 | 0.9314043 | 1.4293087 | 0 | 0.35239762 | 1 | 284 | tags=40%, list=1%, signal=40% |
| OISHI_CHOLANGIOMA_STEM_CELL_LIKE_DN | 245 | 0.47014192 | 1.4290315 | 0 | 0.23619331 | 1 | 3345 | tags=25%, list=12%, signal=29% |
| HP_LATE_ONSET_PROXIMAL_MUSCLE_WEAKNESS | 5 | 0.978814 | 1.4286046 | 0 | 0.35457057 | 1 | 16 | tags=20%, list=0%, signal=20% |
| FEVR_CTNNB1_TARGETS_UP | 608 | 0.44416136 | 1.4284838 | 0 | 0.23703752 | 1 | 3398 | tags=26%, list=13%, signal=29% |
| GO_RESPONSE_TO_OXYGEN_CONTAINING_COMPOUND | 1454 | 0.4108242 | 1.428231 | 0 | 0.35492766 | 1 | 2614 | tags=18%, list=10%, signal=19% |
| LEE_LIVER_CANCER_MYC_TGFA_UP | 56 | 0.5618739 | 1.4278166 | 0 | 0.23810947 | 1 | 3929 | tags=45%, list=15%, signal=52% |
| CROMER_METASTASIS_UP | 66 | 0.5571829 | 1.4275889 | 0 | 0.23796096 | 1 | 1998 | tags=29%, list=7%, signal=31% |
| QI_PLASMACYTOMA_UP | 245 | 0.4837485 | 1.4271406 | 0 | 0.23868512 | 1 | 2108 | tags=18%, list=8%, signal=19% |
| TURASHVILI_BREAST_DUCTAL_CARCINOMA_VS_LOBULAR_NORMAL_DN | 50 | 0.5483633 | 1.4269829 | 0.02272727 | 0.23878244 | 1 | 2434 | tags=22%, list=9%, signal=24% |
| BROWNE_HCMV_INFECTION_48HR_DN | 416 | 0.4430658 | 1.4269667 | 0 | 0.23845755 | 1 | 3250 | tags=26%, list=12%, signal=30% |
| HP_ABNORMALITY_OF_THE_PANCREAS | 219 | 0.47230953 | 1.4266679 | 0 | 0.3587206 | 1 | 3473 | tags=27%, list=13%, signal=31% |
| GROSS_ELK3_TARGETS_UP | 25 | 0.6619768 | 1.4262762 | 0.04166667 | 0.23983964 | 1 | 2704 | tags=36%, list=10%, signal=40% |
| GO_FORELIMB_MORPHOGENESIS | 30 | 0.6617675 | 1.4261026 | 0 | 0.36011767 | 1 | 2396 | tags=23%, list=9%, signal=26% |
| WOO_LIVER_CANCER_RECURRENCE_UP | 101 | 0.5263362 | 1.4260798 | 0 | 0.23995857 | 1 | 3304 | tags=32%, list=12%, signal=36% |
| REACTOME_ACTIVATION_OF_C3_AND_C5 | 5 | 0.921584 | 1.4257926 | 0 | 0.23994872 | 1 | 69 | tags=20%, list=0%, signal=20% |
| GO_ION_CHANNEL_REGULATOR_ACTIVITY | 98 | 0.5083396 | 1.4257666 | 0 | 0.3611008 | 1 | 3160 | tags=28%, list=12%, signal=31% |
| GO_SITE_OF_DOUBLE_STRAND_BREAK | 52 | 0.5821662 | 1.4257559 | 0 | 0.3608084 | 1 | 2838 | tags=37%, list=11%, signal=41% |
| GO_OLIGODENDROCYTE_DIFFERENTIATION | 99 | 0.5186765 | 1.4256619 | 0 | 0.36084664 | 1 | 504 | tags=13%, list=2%, signal=13% |
| CONRAD_GERMLINE_STEM_CELL | 9 | 0.84468997 | 1.4255881 | 0.02439024 | 0.23994362 | 1 | 1376 | tags=22%, list=5%, signal=23% |
| HP_CEREBRAL_AMYLOID_ANGIOPATHY | 4 | 0.94750786 | 1.425581 | 0 | 0.36070466 | 1 | 444 | tags=75%, list=2%, signal=76% |
| HP_SPASTIC_PARAPARESIS | 45 | 0.58477545 | 1.4255176 | 0 | 0.36054772 | 1 | 2673 | tags=29%, list=10%, signal=32% |
| GO_ORGANONITROGEN_COMPOUND_CATABOLIC_PROCESS | 1173 | 0.42107978 | 1.4252267 | 0 | 0.36043653 | 1 | 3338 | tags=28%, list=12%, signal=30% |
| HP_HYPERALDOSTERONISM | 11 | 0.7993393 | 1.4250472 | 0.04878049 | 0.36080015 | 1 | 161 | tags=18%, list=1%, signal=18% |
| APPEL_IMATINIB_RESPONSE | 32 | 0.613562 | 1.4247241 | 0.02325581 | 0.24163249 | 1 | 2998 | tags=41%, list=11%, signal=46% |
| GO_PUTRESCINE_METABOLIC_PROCESS | 6 | 0.86355186 | 1.4247171 | 0 | 0.36059317 | 1 | 910 | tags=33%, list=3%, signal=34% |
| GO_CELL_ADHESION_MOLECULE_BINDING | 490 | 0.43641007 | 1.4246668 | 0 | 0.3604677 | 1 | 3139 | tags=29%, list=12%, signal=32% |
| GO_INTRINSIC_COMPONENT_OF_ORGANELLE_MEMBRANE | 354 | 0.4597339 | 1.4245666 | 0 | 0.3605043 | 1 | 3484 | tags=32%, list=13%, signal=37% |
| WP_ENDOTHELIN_PATHWAYS | 32 | 0.63713926 | 1.424303 | 0.02083333 | 0.24131735 | 1 | 1184 | tags=19%, list=4%, signal=20% |
| GO_CELLULAR_RESPONSE_TO_PEPTIDE | 349 | 0.4370116 | 1.424235 | 0 | 0.3615916 | 1 | 2535 | tags=22%, list=9%, signal=24% |
| GO_LEUKOCYTE_MEDIATED_IMMUNITY | 662 | 0.4286111 | 1.4239609 | 0 | 0.36194304 | 1 | 2451 | tags=19%, list=9%, signal=20% |
| GO_ACTIVATION_OF_IMMUNE_RESPONSE | 374 | 0.46637264 | 1.4239045 | 0 | 0.36183175 | 1 | 2510 | tags=21%, list=9%, signal=23% |
| VECCHI_GASTRIC_CANCER_ADVANCED_VS_EARLY_DN | 99 | 0.5408609 | 1.4237841 | 0 | 0.24189378 | 1 | 3395 | tags=26%, list=13%, signal=30% |
| GO_RIBONUCLEOSIDE_TRIPHOSPHATE_METABOLIC_PROCESS | 75 | 0.5415633 | 1.4237703 | 0.03225806 | 0.3616967 | 1 | 2089 | tags=29%, list=8%, signal=32% |
| BROWNE_HCMV_INFECTION_20HR_DN | 87 | 0.5256219 | 1.4237407 | 0 | 0.24159461 | 1 | 3056 | tags=28%, list=11%, signal=31% |
| SWEET_KRAS_ONCOGENIC_SIGNATURE | 81 | 0.54454285 | 1.423591 | 0 | 0.24149066 | 1 | 1998 | tags=32%, list=7%, signal=35% |
| REACTOME_EXTRINSIC_PATHWAY_OF_FIBRIN_CLOT_FORMATION | 4 | 0.95904267 | 1.4234165 | 0.02325581 | 0.2414353 | 1 | 188 | tags=25%, list=1%, signal=25% |
| HP_ABNORMAL_CARDIOVASCULAR_SYSTEM_PHYSIOLOGY | 977 | 0.42770943 | 1.4234072 | 0 | 0.3615765 | 1 | 3822 | tags=26%, list=14%, signal=29% |
| LEE_SP4_THYMOCYTE | 13 | 0.7915312 | 1.4231753 | 0.02272727 | 0.24145208 | 1 | 1599 | tags=31%, list=6%, signal=33% |
| HP_COLOR_VISION_DEFECT | 80 | 0.5130911 | 1.4231013 | 0.02631579 | 0.36252174 | 1 | 4353 | tags=30%, list=16%, signal=36% |
| GO_GLYCOSYL_COMPOUND_CATABOLIC_PROCESS | 32 | 0.65821236 | 1.4228514 | 0.02857143 | 0.36308187 | 1 | 4092 | tags=50%, list=15%, signal=59% |
| MOOTHA_TCA | 14 | 0.75145996 | 1.4227209 | 0.04444445 | 0.24212542 | 1 | 2744 | tags=64%, list=10%, signal=72% |
| HP_CNS_HYPOMYELINATION | 56 | 0.5596419 | 1.4225639 | 0.04761905 | 0.36365342 | 1 | 1607 | tags=21%, list=6%, signal=23% |
| CREIGHTON_ENDOCRINE_THERAPY_RESISTANCE_3 | 616 | 0.43351483 | 1.4218252 | 0 | 0.24368563 | 1 | 4134 | tags=31%, list=15%, signal=36% |
| GO_REGULATION_OF_AMYLOID_FIBRIL_FORMATION | 7 | 0.8738756 | 1.4217395 | 0 | 0.36550102 | 1 | 444 | tags=43%, list=2%, signal=44% |
| GO_CARBOHYDRATE_DERIVATIVE_BIOSYNTHETIC_PROCESS | 586 | 0.43240675 | 1.4216963 | 0 | 0.36529955 | 1 | 4526 | tags=34%, list=17%, signal=40% |
| HP_GENERALIZED_AMYLOID_DEPOSITION | 4 | 0.9098396 | 1.4216741 | 0.05 | 0.3649974 | 1 | 6 | tags=50%, list=0%, signal=50% |
| HP_METABOLIC_ACIDOSIS | 116 | 0.5086146 | 1.4213144 | 0.03571429 | 0.36572823 | 1 | 3709 | tags=37%, list=14%, signal=43% |
| GO_TITIN_BINDING | 11 | 0.7612949 | 1.4212322 | 0.04761905 | 0.36571538 | 1 | 573 | tags=18%, list=2%, signal=19% |
| IIZUKA_LIVER_CANCER_PROGRESSION_G2_G3_DN | 8 | 0.8251332 | 1.4211746 | 0.05 | 0.24500494 | 1 | 604 | tags=38%, list=2%, signal=38% |
| GO_NEGATIVE_REGULATION_OF_HOMOTYPIC_CELL_CELL_ADHESION | 12 | 0.75794417 | 1.4210427 | 0.04545455 | 0.36626863 | 1 | 1865 | tags=25%, list=7%, signal=27% |
| WP_MATRIX_METALLOPROTEINASES | 24 | 0.637699 | 1.421004 | 0 | 0.24506418 | 1 | 1774 | tags=21%, list=7%, signal=22% |
| PARK_APL_PATHOGENESIS_UP | 13 | 0.7628163 | 1.420874 | 0.04255319 | 0.24497968 | 1 | 2587 | tags=38%, list=10%, signal=43% |
| GO_REGULATION_OF_CELL_SUBSTRATE_ADHESION | 190 | 0.47394866 | 1.4205875 | 0.03571429 | 0.36696753 | 1 | 3120 | tags=28%, list=12%, signal=31% |
| GO_PIGMENT_METABOLIC_PROCESS | 63 | 0.53616107 | 1.420559 | 0.03225806 | 0.3667089 | 1 | 1501 | tags=22%, list=6%, signal=23% |
| GO_NUCLEOSIDE_PHOSPHATE_BIOSYNTHETIC_PROCESS | 239 | 0.46182692 | 1.4197296 | 0 | 0.36841342 | 1 | 3623 | tags=29%, list=13%, signal=34% |
| DANG_REGULATED_BY_MYC_DN | 232 | 0.4690048 | 1.4191954 | 0 | 0.2484388 | 1 | 2894 | tags=29%, list=11%, signal=32% |
| GO_NEGATIVE_REGULATION_OF_HYDROLASE_ACTIVITY | 348 | 0.4543581 | 1.4191356 | 0 | 0.36970845 | 1 | 2632 | tags=20%, list=10%, signal=21% |
| GO_REGULATION_OF_EPITHELIAL_CELL_DIFFERENTIATION | 130 | 0.4800541 | 1.4189243 | 0 | 0.36993796 | 1 | 3281 | tags=29%, list=12%, signal=33% |
| GO_SIGNAL_TRANSDUCTION_BY_PROTEIN_PHOSPHORYLATION | 799 | 0.4261961 | 1.4187602 | 0 | 0.36950427 | 1 | 3128 | tags=22%, list=12%, signal=25% |
| GO_ENDOCYTIC_VESICLE_MEMBRANE | 140 | 0.49862078 | 1.4186124 | 0 | 0.3695755 | 1 | 2434 | tags=27%, list=9%, signal=30% |
| GO_POSITIVE_REGULATION_OF_MYOTUBE_DIFFERENTIATION | 26 | 0.6504762 | 1.4185224 | 0.04347826 | 0.36960298 | 1 | 1181 | tags=15%, list=4%, signal=16% |
| GO_GOLGI_LUMEN | 74 | 0.5494019 | 1.4184166 | 0 | 0.36954528 | 1 | 2948 | tags=27%, list=11%, signal=30% |
| GO_NEGATIVE_REGULATION_OF_COAGULATION | 39 | 0.6282277 | 1.4184079 | 0.02941177 | 0.36930192 | 1 | 1086 | tags=15%, list=4%, signal=16% |
| HP_ABNORMAL_ADIPOSE_TISSUE_MORPHOLOGY | 171 | 0.47457612 | 1.4183822 | 0 | 0.3690872 | 1 | 2434 | tags=18%, list=9%, signal=20% |
| HP_FUNCTIONAL_ABNORMALITY_OF_THE_GASTROINTESTINAL_TRACT | 723 | 0.43854195 | 1.418249 | 0 | 0.3692152 | 1 | 3152 | tags=23%, list=12%, signal=25% |
| GO_POSITIVE_REGULATION_OF_EPITHELIAL_CELL_DIFFERENTIATION | 53 | 0.5529776 | 1.4181 | 0.02 | 0.36947235 | 1 | 2160 | tags=17%, list=8%, signal=18% |
| HP_DECREASED_CIRCULATING_ANTIBODY_LEVEL | 132 | 0.5010797 | 1.4177047 | 0.04761905 | 0.37020847 | 1 | 2876 | tags=23%, list=11%, signal=25% |
| TONKS_TARGETS_OF_RUNX1_RUNX1T1_FUSION_MONOCYTE_DN | 43 | 0.5795756 | 1.4171478 | 0.02564103 | 0.25231653 | 1 | 714 | tags=14%, list=3%, signal=14% |
| HP_HEMOLYTIC_UREMIC_SYNDROME | 10 | 0.7632711 | 1.417116 | 0.04 | 0.37137955 | 1 | 4033 | tags=40%, list=15%, signal=47% |
| GO_OTIC_VESICLE_FORMATION | 6 | 0.8775499 | 1.4170187 | 0.02564103 | 0.37113506 | 1 | 267 | tags=17%, list=1%, signal=17% |
| HP_INCREASED_INTRACRANIAL_PRESSURE | 66 | 0.57183784 | 1.4166996 | 0 | 0.3716654 | 1 | 2443 | tags=29%, list=9%, signal=32% |
| CHICAS_RB1_TARGETS_SENESCENT | 491 | 0.4537522 | 1.4166435 | 0 | 0.25305587 | 1 | 3491 | tags=32%, list=13%, signal=37% |
| ZHU_CMV_24_HR_UP | 77 | 0.5130387 | 1.4165419 | 0 | 0.25301054 | 1 | 2960 | tags=36%, list=11%, signal=41% |
| GO_INFLAMMATORY_RESPONSE_TO_ANTIGENIC_STIMULUS | 42 | 0.57326597 | 1.4162065 | 0.02857143 | 0.37255657 | 1 | 1765 | tags=19%, list=7%, signal=20% |
| CHICAS_RB1_TARGETS_LOW_SERUM | 74 | 0.52919716 | 1.4159902 | 0.03125 | 0.25408772 | 1 | 1621 | tags=27%, list=6%, signal=29% |
| DELYS_THYROID_CANCER_UP | 396 | 0.44138032 | 1.4158288 | 0 | 0.25358707 | 1 | 3500 | tags=26%, list=13%, signal=30% |
| GO_ENERGY_COUPLED_PROTON_TRANSMEMBRANE_TRANSPORT_AGAINST_ELECTROCHEMICAL_GRADIENT | 4 | 0.983243 | 1.4157574 | 0.02439024 | 0.37408513 | 1 | 454 | tags=100%, list=2%, signal=102% |
| HOSHIDA_LIVER_CANCER_SUBCLASS_S3 | 226 | 0.47396684 | 1.415152 | 0 | 0.25453526 | 1 | 3024 | tags=28%, list=11%, signal=31% |
| GO_MOLECULAR_CARRIER_ACTIVITY | 9 | 0.82105184 | 1.4151055 | 0.02941177 | 0.37574872 | 1 | 1965 | tags=44%, list=7%, signal=48% |
| HP_ABSENCE_OF_PUBERTAL_DEVELOPMENT | 19 | 0.6857422 | 1.4149826 | 0.04651163 | 0.37579736 | 1 | 1927 | tags=16%, list=7%, signal=17% |
| HP_ABNORMAL_CIRCULATING_ANDROGEN_LEVEL | 53 | 0.56293005 | 1.4146867 | 0.02857143 | 0.3765488 | 1 | 2301 | tags=19%, list=9%, signal=21% |
| REACTOME_DNA_REPLICATION | 123 | 0.5147524 | 1.4142069 | 0 | 0.25583702 | 1 | 2375 | tags=35%, list=9%, signal=38% |
| HP_ABNORMAL_HAIR_QUANTITY | 624 | 0.42988876 | 1.4139228 | 0 | 0.37836695 | 1 | 4004 | tags=29%, list=15%, signal=33% |
| BUYTAERT_PHOTODYNAMIC_THERAPY_STRESS_DN | 598 | 0.44526625 | 1.4137894 | 0 | 0.25656876 | 1 | 3471 | tags=31%, list=13%, signal=35% |
| GO_POLYAMINE_BIOSYNTHETIC_PROCESS | 12 | 0.7514355 | 1.4136127 | 0.04347826 | 0.37852144 | 1 | 2403 | tags=42%, list=9%, signal=46% |
| GO_SIDE_OF_MEMBRANE | 454 | 0.44249034 | 1.4135344 | 0 | 0.378198 | 1 | 2346 | tags=15%, list=9%, signal=16% |
| GO_CARBON_CARBON_LYASE_ACTIVITY | 45 | 0.5860485 | 1.413434 | 0 | 0.37830073 | 1 | 2960 | tags=33%, list=11%, signal=37% |
| HP_RENAL_MALROTATION | 5 | 0.9160779 | 1.4132853 | 0 | 0.3780609 | 1 | 2031 | tags=40%, list=8%, signal=43% |
| VERHAAK_GLIOBLASTOMA_MESENCHYMAL | 200 | 0.47265083 | 1.413251 | 0 | 0.25724646 | 1 | 2163 | tags=15%, list=8%, signal=16% |
| HP_ABNORMAL_GLUCOSE_HOMEOSTASIS | 483 | 0.44952974 | 1.4132442 | 0 | 0.377872 | 1 | 2329 | tags=18%, list=9%, signal=20% |
| GO_CELL_CELL_SIGNALING_BY_WNT | 474 | 0.4340878 | 1.4126872 | 0 | 0.3794876 | 1 | 2863 | tags=23%, list=11%, signal=25% |
| HP_BRONCHIECTASIS | 95 | 0.5325415 | 1.412563 | 0.02631579 | 0.3797409 | 1 | 3987 | tags=19%, list=15%, signal=22% |
| GO_REGULATION_OF_CELL_POPULATION_PROLIFERATION | 1384 | 0.419854 | 1.4123919 | 0 | 0.37996525 | 1 | 2874 | tags=19%, list=11%, signal=20% |
| HP_ABNORMALITY_OF_THE_UPPER_URINARY_TRACT | 1028 | 0.41726422 | 1.4121947 | 0 | 0.38046747 | 1 | 3535 | tags=25%, list=13%, signal=27% |
| GO_OLFACTORY_RECEPTOR_BINDING | 3 | 0.99333143 | 1.4118223 | 0 | 0.380974 | 1 | 131 | tags=33%, list=0%, signal=33% |
| WP_GENE_REGULATORY_NETWORK_MODELLING_SOMITOGENESIS | 10 | 0.8010464 | 1.4115853 | 0.02325581 | 0.25927168 | 1 | 2567 | tags=30%, list=10%, signal=33% |
| GO_CUL5_RING_UBIQUITIN_LIGASE_COMPLEX | 4 | 0.9337004 | 1.4115196 | 0 | 0.38118756 | 1 | 1425 | tags=75%, list=5%, signal=79% |
| GO_PHOTORECEPTOR_CELL_OUTER_SEGMENT_ORGANIZATION | 10 | 0.8287339 | 1.411472 | 0.02 | 0.38075352 | 1 | 669 | tags=20%, list=2%, signal=21% |
| HOQUE_METHYLATED_IN_CANCER | 48 | 0.57300556 | 1.4113712 | 0.02702703 | 0.2595913 | 1 | 1291 | tags=17%, list=5%, signal=17% |
| GO_DUCTUS_ARTERIOSUS_CLOSURE | 3 | 0.9853551 | 1.4113479 | 0 | 0.3808523 | 1 | 128 | tags=33%, list=0%, signal=33% |
| HP_RETINOSCHISIS | 3 | 0.97449505 | 1.4111938 | 0.025 | 0.38101915 | 1 | 434 | tags=33%, list=2%, signal=34% |
| MEBARKI_HCC_PROGENITOR_WNT_UP_CTNNB1_DEPENDENT_BLOCKED_BY_FZD8CRD | 36 | 0.6211991 | 1.4111714 | 0.02941177 | 0.2596739 | 1 | 1823 | tags=28%, list=7%, signal=30% |
| BLANCO_MELO_COVID19_SARS_COV_2_INFECTION_CALU3_CELLS_UP | 263 | 0.47157133 | 1.4111048 | 0 | 0.25940895 | 1 | 2488 | tags=21%, list=9%, signal=23% |
| HP_INCREASED_CIRCULATING_GONADOTROPIN_LEVEL | 48 | 0.580114 | 1.4110156 | 0.0212766 | 0.3808744 | 1 | 1422 | tags=15%, list=5%, signal=15% |
| REACTOME_INTERFERON_SIGNALING | 160 | 0.48378047 | 1.4106642 | 0 | 0.25971857 | 1 | 2378 | tags=30%, list=9%, signal=33% |
| KORKOLA_EMBRYONAL_CARCINOMA_UP | 38 | 0.61676383 | 1.4104666 | 0.025 | 0.25980356 | 1 | 2516 | tags=29%, list=9%, signal=32% |
| KOKKINAKIS_METHIONINE_DEPRIVATION_96HR_DN | 69 | 0.52786416 | 1.410466 | 0 | 0.25944668 | 1 | 3738 | tags=42%, list=14%, signal=49% |
| REACTOME_MITOCHONDRIAL_FATTY_ACID_BETA_OXIDATION | 33 | 0.5977214 | 1.4104297 | 0.04347826 | 0.25922957 | 1 | 3722 | tags=48%, list=14%, signal=56% |
| GO_REGULATION_OF_PROTEOLYSIS | 592 | 0.4424248 | 1.4103154 | 0 | 0.3826167 | 1 | 2896 | tags=24%, list=11%, signal=26% |
| DUTERTRE_ESTRADIOL_RESPONSE_24HR_DN | 473 | 0.44692412 | 1.4102923 | 0 | 0.25922027 | 1 | 3566 | tags=26%, list=13%, signal=29% |
| GO_ADENOSINE_TO_INOSINE_EDITING | 8 | 0.8432262 | 1.4102359 | 0.04545455 | 0.38265795 | 1 | 3995 | tags=63%, list=15%, signal=73% |
| HP_CONTIGUOUS_GENE_SYNDROME | 4 | 0.96159583 | 1.410042 | 0 | 0.3829624 | 1 | 286 | tags=25%, list=1%, signal=25% |
| HP_REDUCED_VISUAL_ACUITY | 401 | 0.45802546 | 1.4099562 | 0 | 0.38309944 | 1 | 2527 | tags=20%, list=9%, signal=21% |
| GO_CANONICAL_WNT_SIGNALING_PATHWAY | 305 | 0.47585407 | 1.4099268 | 0 | 0.38291055 | 1 | 2754 | tags=24%, list=10%, signal=26% |
| GO_EXTRACELLULAR_STRUCTURE_ORGANIZATION | 356 | 0.45008704 | 1.4096982 | 0 | 0.38288647 | 1 | 3316 | tags=19%, list=12%, signal=21% |
| GO_11_CIS_RETINAL_BINDING | 2 | 0.98046875 | 1.4096841 | 0.0212766 | 0.38258973 | 1 | 257 | tags=50%, list=1%, signal=50% |
| GO_NUCLEAR_OUTER_MEMBRANE_ENDOPLASMIC_RETICULUM_MEMBRANE_NETWORK | 954 | 0.4219776 | 1.4096231 | 0 | 0.38246807 | 1 | 3792 | tags=31%, list=14%, signal=35% |
| TIEN_INTESTINE_PROBIOTICS_6HR_DN | 162 | 0.4842732 | 1.4096122 | 0 | 0.25998384 | 1 | 3104 | tags=38%, list=12%, signal=43% |
| GO_REGULATION_OF_SENSORY_PERCEPTION | 28 | 0.64785457 | 1.4095765 | 0.02173913 | 0.38229284 | 1 | 1729 | tags=21%, list=6%, signal=23% |
| GO_GLYCOPROTEIN_CATABOLIC_PROCESS | 21 | 0.7106304 | 1.409562 | 0.03571429 | 0.3820107 | 1 | 1876 | tags=24%, list=7%, signal=26% |
| BASSO_HAIRY_CELL_LEUKEMIA_UP | 75 | 0.53658617 | 1.4095488 | 0.02702703 | 0.25972202 | 1 | 2189 | tags=28%, list=8%, signal=30% |
| GO_INTRINSIC_COMPONENT_OF_MITOCHONDRIAL_MEMBRANE | 73 | 0.55395085 | 1.4092759 | 0.04761905 | 0.38245982 | 1 | 4103 | tags=40%, list=15%, signal=47% |
| PASINI_SUZ12_TARGETS_DN | 296 | 0.46443415 | 1.4092112 | 0 | 0.2599101 | 1 | 3172 | tags=29%, list=12%, signal=33% |
| GO_INTERLEUKIN_1_ALPHA_PRODUCTION | 3 | 0.99112415 | 1.4091868 | 0 | 0.3824206 | 1 | 140 | tags=33%, list=1%, signal=34% |
| HP_KINETIC_TREMOR | 71 | 0.5389153 | 1.4091161 | 0.0212766 | 0.3820731 | 1 | 2531 | tags=28%, list=9%, signal=31% |
| GO_ACYL_COA_OXIDASE_ACTIVITY | 6 | 0.8583455 | 1.4088212 | 0 | 0.3826017 | 1 | 1180 | tags=33%, list=4%, signal=35% |
| LOPEZ_TRANSLATION_VIA_FN1_SIGNALING | 33 | 0.6395487 | 1.4088155 | 0.04761905 | 0.2602127 | 1 | 1618 | tags=39%, list=6%, signal=42% |
| GO_EPITHELIUM_DEVELOPMENT | 1005 | 0.42254037 | 1.4087741 | 0 | 0.3821479 | 1 | 3383 | tags=21%, list=13%, signal=23% |
| GO_B_CELL_MEDIATED_IMMUNITY | 98 | 0.47739378 | 1.4085456 | 0 | 0.38255197 | 1 | 2016 | tags=16%, list=7%, signal=18% |
| REACTOME_SYNTHESIS_OF_SUBSTRATES_IN_N_GLYCAN_BIOSYTHESIS | 59 | 0.52314407 | 1.4083031 | 0 | 0.26119328 | 1 | 4512 | tags=37%, list=17%, signal=45% |
| PROVENZANI_METASTASIS_UP | 183 | 0.47114572 | 1.408196 | 0 | 0.26106793 | 1 | 3625 | tags=35%, list=13%, signal=40% |
| HP_OVARIAN_PAPILLARY_ADENOCARCINOMA | 5 | 0.89152426 | 1.4077176 | 0.04347826 | 0.38442805 | 1 | 85 | tags=20%, list=0%, signal=20% |
| HP_ABNORMAL_ENZYME_COENZYME_ACTIVITY | 178 | 0.47891095 | 1.4071038 | 0 | 0.38597384 | 1 | 4127 | tags=33%, list=15%, signal=38% |
| GO_GLIAL_CELL_DEVELOPMENT | 115 | 0.5169688 | 1.406993 | 0 | 0.38605273 | 1 | 3035 | tags=24%, list=11%, signal=27% |
| ACOSTA_PROLIFERATION_INDEPENDENT_MYC_TARGETS_UP | 78 | 0.50829 | 1.406758 | 0.03333334 | 0.2632518 | 1 | 3358 | tags=33%, list=12%, signal=38% |
| HP_ELEVATED_CIRCULATING_FOLLICLE_STIMULATING_HORMONE_LEVEL | 21 | 0.67767626 | 1.4064748 | 0.04545455 | 0.38587373 | 1 | 2301 | tags=24%, list=9%, signal=26% |
| GO_FATTY_ACID_DERIVATIVE_CATABOLIC_PROCESS | 10 | 0.8117877 | 1.4063245 | 0.02272727 | 0.38588417 | 1 | 832 | tags=30%, list=3%, signal=31% |
| HP_SKIN_RASH | 85 | 0.54337925 | 1.4060775 | 0 | 0.38646334 | 1 | 2008 | tags=18%, list=7%, signal=19% |
| GO_POSITIVE_REGULATION_OF_RESPONSE_TO_INTERFERON_GAMMA | 5 | 0.91189754 | 1.4060758 | 0.04444445 | 0.3861448 | 1 | 278 | tags=40%, list=1%, signal=40% |
| GO_AZUROPHIL_GRANULE_LUMEN | 73 | 0.54908854 | 1.4057925 | 0.04255319 | 0.38645878 | 1 | 2998 | tags=36%, list=11%, signal=40% |
| GO_REGULATION_OF_CEREBELLAR_GRANULE_CELL_PRECURSOR_PROLIFERATION | 7 | 0.87476635 | 1.4055823 | 0.02941177 | 0.38657498 | 1 | 377 | tags=14%, list=1%, signal=14% |
| LEE_AGING_MUSCLE_DN | 37 | 0.6143498 | 1.4055089 | 0.02941177 | 0.26556644 | 1 | 1811 | tags=22%, list=7%, signal=23% |
| HP_FEMALE_SEXUAL_DYSFUNCTION | 45 | 0.5366099 | 1.4054604 | 0 | 0.38633448 | 1 | 4353 | tags=33%, list=16%, signal=40% |
| HP_SCANNING_SPEECH | 12 | 0.7754328 | 1.4052963 | 0 | 0.38676584 | 1 | 1003 | tags=33%, list=4%, signal=35% |
| MIKKELSEN_MEF_LCP_WITH_H3K4ME3 | 119 | 0.5025933 | 1.4051778 | 0 | 0.26591483 | 1 | 3380 | tags=28%, list=13%, signal=32% |
| GO_REGULATION_OF_INFLAMMATORY_RESPONSE_TO_WOUNDING | 3 | 0.9878083 | 1.4050139 | 0.04444445 | 0.38677248 | 1 | 300 | tags=67%, list=1%, signal=67% |
| REACTOME_REGULATION_OF_PTEN_STABILITY_AND_ACTIVITY | 66 | 0.5760566 | 1.4049323 | 0.02777778 | 0.26612702 | 1 | 805 | tags=30%, list=3%, signal=31% |
| REACTOME_SWITCHING_OF_ORIGINS_TO_A_POST_REPLICATIVE_STATE | 87 | 0.54925716 | 1.4046096 | 0.025 | 0.2664969 | 1 | 2375 | tags=39%, list=9%, signal=43% |
| HP_ABNORMAL_REFLEX | 992 | 0.4197445 | 1.4046016 | 0 | 0.38791168 | 1 | 4059 | tags=30%, list=15%, signal=34% |
| GO_SENSORY_PERCEPTION | 443 | 0.443142 | 1.4044206 | 0 | 0.38794467 | 1 | 2172 | tags=11%, list=8%, signal=12% |
| KEGG_VIBRIO_CHOLERAE_INFECTION | 51 | 0.63955975 | 1.4043386 | 0 | 0.26668143 | 1 | 1709 | tags=29%, list=6%, signal=31% |
| GO_FATTY_ACID_BETA_OXIDATION | 69 | 0.5312091 | 1.4043175 | 0 | 0.38798067 | 1 | 3779 | tags=41%, list=14%, signal=47% |
| GO_ATPASE_COUPLED_AMIDE_TRANSPORTER_ACTIVITY | 4 | 0.9381854 | 1.4041793 | 0.02631579 | 0.38812196 | 1 | 1315 | tags=75%, list=5%, signal=79% |
| GO_BLOOD_VESSEL_MORPHOGENESIS | 543 | 0.4364465 | 1.4038229 | 0 | 0.3887546 | 1 | 3307 | tags=22%, list=12%, signal=25% |
| GO_INTERCELLULAR_CANALICULUS | 5 | 0.9155455 | 1.4037864 | 0.04347826 | 0.38865992 | 1 | 68 | tags=20%, list=0%, signal=20% |
| GO_MACROLIDE_BINDING | 9 | 0.82207507 | 1.4037347 | 0.02222222 | 0.3886178 | 1 | 3869 | tags=78%, list=14%, signal=91% |
| ISHIKAWA_STING_SIGNALING | 5 | 0.9305168 | 1.4036726 | 0 | 0.2678588 | 1 | 1874 | tags=100%, list=7%, signal=107% |
| REACTOME_TRANSCRIPTIONAL_REGULATION_BY_RUNX3 | 93 | 0.5378721 | 1.4036186 | 0 | 0.26763535 | 1 | 1624 | tags=30%, list=6%, signal=32% |
| MARTINEZ_RB1_AND_TP53_TARGETS_UP | 556 | 0.43727055 | 1.4036152 | 0 | 0.267279 | 1 | 3275 | tags=26%, list=12%, signal=29% |
| GO_DETOXIFICATION | 106 | 0.5049166 | 1.4034231 | 0.03225806 | 0.3894978 | 1 | 2750 | tags=27%, list=10%, signal=30% |
| REACTOME_ADHERENS_JUNCTIONS_INTERACTIONS | 28 | 0.6527926 | 1.4031016 | 0.02173913 | 0.26733682 | 1 | 4174 | tags=39%, list=16%, signal=46% |
| HSIAO_LIVER_SPECIFIC_GENES | 178 | 0.48069718 | 1.4025989 | 0 | 0.2677712 | 1 | 2596 | tags=16%, list=10%, signal=18% |
| GO_LATE_ENDOSOME_MEMBRANE | 121 | 0.51671165 | 1.4025657 | 0.03030303 | 0.39092883 | 1 | 2331 | tags=31%, list=9%, signal=34% |
| JIANG_AGING_CEREBRAL_CORTEX_UP | 28 | 0.6538732 | 1.402467 | 0.02857143 | 0.26761767 | 1 | 3013 | tags=50%, list=11%, signal=56% |
| WAKABAYASHI_ADIPOGENESIS_PPARG_BOUND_8D | 602 | 0.4239413 | 1.4024587 | 0 | 0.26728684 | 1 | 3750 | tags=32%, list=14%, signal=36% |
| WANG_CISPLATIN_RESPONSE_AND_XPC_UP | 156 | 0.47367394 | 1.4021825 | 0 | 0.26742423 | 1 | 1748 | tags=20%, list=6%, signal=21% |
| BIOCARTA_MALATEX_PATHWAY | 7 | 0.8647048 | 1.4021497 | 0.02272727 | 0.267161 | 1 | 2302 | tags=71%, list=9%, signal=78% |
| ZHANG_BREAST_CANCER_PROGENITORS_DN | 133 | 0.4813858 | 1.4020274 | 0 | 0.26718655 | 1 | 3336 | tags=35%, list=12%, signal=40% |
| GO_NERVE_DEVELOPMENT | 69 | 0.54842776 | 1.4019086 | 0.025 | 0.39278927 | 1 | 875 | tags=10%, list=3%, signal=10% |
| GO_SOMATIC_STEM_CELL_POPULATION_MAINTENANCE | 65 | 0.55526996 | 1.4017389 | 0.02631579 | 0.39270994 | 1 | 3771 | tags=40%, list=14%, signal=46% |
| GO_MITOTIC_CHROMOSOME_CONDENSATION | 14 | 0.74811137 | 1.4015633 | 0.04347826 | 0.39314196 | 1 | 3923 | tags=64%, list=15%, signal=75% |
| FLECHNER_PBL_KIDNEY_TRANSPLANT_REJECTED_VS_OK_UP | 61 | 0.5968372 | 1.4015403 | 0.02777778 | 0.26770136 | 1 | 2452 | tags=30%, list=9%, signal=32% |
| GO_VACUOLAR_LUMEN | 145 | 0.49624145 | 1.4008377 | 0.03703704 | 0.3955094 | 1 | 3112 | tags=34%, list=12%, signal=38% |
| GO_RIBOSE_PHOSPHATE_BIOSYNTHETIC_PROCESS | 176 | 0.47480088 | 1.4007013 | 0 | 0.39541352 | 1 | 3623 | tags=32%, list=13%, signal=37% |
| HP_EPIGASTRIC_PAIN | 4 | 0.94557554 | 1.4006928 | 0.04545455 | 0.39513424 | 1 | 85 | tags=25%, list=0%, signal=25% |
| REACTOME_SIGNALING_BY_MEMBRANE_TETHERED_FUSIONS_OF_PDGFRA_OR_PDGFRB | 4 | 0.95498705 | 1.4006442 | 0.04878049 | 0.2689455 | 1 | 746 | tags=50%, list=3%, signal=51% |
| MIKKELSEN_NPC_ICP_WITH_H3K4ME3 | 420 | 0.4434533 | 1.4006178 | 0 | 0.26861706 | 1 | 4114 | tags=35%, list=15%, signal=41% |
| HORIUCHI_WTAP_TARGETS_UP | 266 | 0.4666181 | 1.4004464 | 0 | 0.2685537 | 1 | 2884 | tags=27%, list=11%, signal=30% |
| GO_GROWTH_FACTOR_COMPLEX | 3 | 0.9785209 | 1.4003508 | 0.02272727 | 0.39599937 | 1 | 453 | tags=67%, list=2%, signal=68% |
| GO_ORGANOPHOSPHATE_METABOLIC_PROCESS | 932 | 0.41231483 | 1.4001772 | 0 | 0.3960194 | 1 | 4002 | tags=28%, list=15%, signal=32% |
| HAN_SATB1_TARGETS_UP | 383 | 0.4373293 | 1.4000319 | 0 | 0.26930344 | 1 | 2994 | tags=26%, list=11%, signal=28% |
| CROONQUIST_NRAS_VS_STROMAL_STIMULATION_UP | 38 | 0.56938535 | 1.3998783 | 0.02272727 | 0.26934975 | 1 | 3617 | tags=29%, list=13%, signal=33% |
| GO_SENSORY_PERCEPTION_OF_MECHANICAL_STIMULUS | 150 | 0.48409557 | 1.3996979 | 0 | 0.3969233 | 1 | 4105 | tags=25%, list=15%, signal=29% |
| SCHUETZ_BREAST_CANCER_DUCTAL_INVASIVE_UP | 321 | 0.4469601 | 1.3993187 | 0 | 0.2700325 | 1 | 3104 | tags=25%, list=12%, signal=28% |
| REACTOME_CLASS_A_1_RHODOPSIN_LIKE_RECEPTORS_ | 213 | 0.4486259 | 1.3992021 | 0 | 0.2699016 | 1 | 3383 | tags=12%, list=13%, signal=13% |
| GO_PHOTORECEPTOR_CELL_DIFFERENTIATION | 55 | 0.5334585 | 1.3991048 | 0.03125 | 0.398516 | 1 | 1939 | tags=15%, list=7%, signal=16% |
| GO_INTESTINAL_EPITHELIAL_CELL_DIFFERENTIATION | 18 | 0.6627495 | 1.3979219 | 0.02380952 | 0.40040886 | 1 | 2675 | tags=33%, list=10%, signal=37% |
| HP_ECTROPION | 46 | 0.5861256 | 1.3978478 | 0.025 | 0.40029427 | 1 | 4620 | tags=37%, list=17%, signal=45% |
| REACTOME_HEMOSTASIS | 509 | 0.44058517 | 1.3977509 | 0 | 0.2721094 | 1 | 3122 | tags=24%, list=12%, signal=27% |
| HP_INSPIRATORY_STRIDOR | 4 | 0.98468506 | 1.397624 | 0 | 0.40033403 | 1 | 0 | tags=25%, list=0%, signal=25% |
| GO_WHOLE_MEMBRANE | 1541 | 0.42037255 | 1.3975884 | 0 | 0.3998911 | 1 | 3306 | tags=25%, list=12%, signal=27% |
| REACTOME_SIGNALING_BY_WNT | 257 | 0.44903535 | 1.3969667 | 0 | 0.27325684 | 1 | 2434 | tags=23%, list=9%, signal=25% |
| KIM_ALL_DISORDERS_CALB1_CORR_UP | 526 | 0.43113506 | 1.3960614 | 0 | 0.27514148 | 1 | 2403 | tags=29%, list=9%, signal=31% |
| GO_INTRINSIC_COMPONENT_OF_ENDOPLASMIC_RETICULUM_MEMBRANE | 143 | 0.5083811 | 1.3957139 | 0 | 0.40558514 | 1 | 2632 | tags=33%, list=10%, signal=36% |
| REACTOME_MAPK6_MAPK4_SIGNALING | 85 | 0.5173891 | 1.3956966 | 0 | 0.2751946 | 1 | 1624 | tags=29%, list=6%, signal=31% |
| LEIN_CHOROID_PLEXUS_MARKERS | 92 | 0.5174365 | 1.3953363 | 0 | 0.2757292 | 1 | 2995 | tags=22%, list=11%, signal=24% |
| HP_FINGER_SWELLING | 4 | 0.9724415 | 1.3951249 | 0.03921569 | 0.4065803 | 1 | 365 | tags=75%, list=1%, signal=76% |
| SENESE_HDAC1_TARGETS_DN | 231 | 0.45553297 | 1.3949136 | 0 | 0.27624017 | 1 | 2318 | tags=25%, list=9%, signal=27% |
| REACTOME_NEUTROPHIL_DEGRANULATION | 420 | 0.4572323 | 1.3947499 | 0 | 0.27627736 | 1 | 2451 | tags=22%, list=9%, signal=24% |
| REACTOME_HYALURONAN_UPTAKE_AND_DEGRADATION | 10 | 0.8479179 | 1.3940593 | 0.02380952 | 0.2774754 | 1 | 2998 | tags=60%, list=11%, signal=67% |
| SUNG_METASTASIS_STROMA_UP | 101 | 0.5092871 | 1.39378 | 0.02857143 | 0.2779423 | 1 | 1947 | tags=28%, list=7%, signal=30% |
| ELVIDGE_HYPOXIA_UP | 158 | 0.47188225 | 1.3937613 | 0 | 0.27758914 | 1 | 2550 | tags=24%, list=9%, signal=26% |
| KEGG_GLYOXYLATE_AND_DICARBOXYLATE_METABOLISM | 14 | 0.7394797 | 1.3935251 | 0.03773585 | 0.27766684 | 1 | 3194 | tags=50%, list=12%, signal=57% |
| GO_OPIOID_RECEPTOR_BINDING | 6 | 0.8514214 | 1.3933831 | 0.02439024 | 0.41177073 | 1 | 514 | tags=17%, list=2%, signal=17% |
| HP_SPONDYLOEPIMETAPHYSEAL_DYSPLASIA | 15 | 0.69944394 | 1.3933825 | 0 | 0.41144902 | 1 | 1934 | tags=33%, list=7%, signal=36% |
| PECE_MAMMARY_STEM_CELL_DN | 130 | 0.5026448 | 1.393252 | 0.04347826 | 0.27787483 | 1 | 2263 | tags=28%, list=8%, signal=31% |
| HP_GENERALIZED_ONSET_SEIZURE | 226 | 0.4627882 | 1.3931245 | 0 | 0.4116921 | 1 | 3156 | tags=23%, list=12%, signal=25% |
| GO_NEGATIVE_REGULATION_OF_MEMBRANE_PROTEIN_ECTODOMAIN_PROTEOLYSIS | 7 | 0.8475776 | 1.3930066 | 0.02272727 | 0.41168255 | 1 | 1774 | tags=71%, list=7%, signal=76% |
| GO_CELL_SUBSTRATE_ADHESION | 322 | 0.45169353 | 1.3929682 | 0 | 0.41149962 | 1 | 3373 | tags=25%, list=13%, signal=28% |
| GO_GLIOGENESIS | 280 | 0.44957688 | 1.3927993 | 0 | 0.41185263 | 1 | 3067 | tags=23%, list=11%, signal=26% |
| GO_HOMOPHILIC_CELL_ADHESION_VIA_PLASMA_MEMBRANE_ADHESION_MOLECULES | 145 | 0.49769792 | 1.3926692 | 0.03448276 | 0.41198224 | 1 | 4920 | tags=32%, list=18%, signal=39% |
| HP_MIGRAINE | 117 | 0.48872972 | 1.3926156 | 0.03125 | 0.411848 | 1 | 2129 | tags=19%, list=8%, signal=20% |
| WANG_SMARCE1_TARGETS_DN | 343 | 0.44262072 | 1.3925811 | 0 | 0.2786162 | 1 | 3107 | tags=31%, list=12%, signal=35% |
| WP_ULTRACONSERVED_REGION_339_MODULATION_OF_TUMOR_SUPPRESSOR_MICRORNAS_IN_CANCER | 1 | 0.9968793 | 1.3925778 | 0 | 0.27828565 | 1 | 84 | tags=100%, list=0%, signal=100% |
| MIKKELSEN_ES_ICP_WITH_H3K4ME3 | 628 | 0.42554438 | 1.3923528 | 0 | 0.27838528 | 1 | 4119 | tags=25%, list=15%, signal=29% |
| GO_METANEPHRIC_MESENCHYMAL_CELL_DIFFERENTIATION | 3 | 0.9771701 | 1.3920276 | 0.02272727 | 0.41258097 | 1 | 72 | tags=33%, list=0%, signal=33% |
| GO_REGULATION_OF_IRON_ION_TRANSPORT | 4 | 0.92577195 | 1.3920262 | 0.02325581 | 0.41227385 | 1 | 42 | tags=50%, list=0%, signal=50% |
| SENGUPTA_NASOPHARYNGEAL_CARCINOMA_WITH_LMP1_UP | 345 | 0.44849065 | 1.3918501 | 0.04166667 | 0.27872968 | 1 | 2524 | tags=23%, list=9%, signal=25% |
| MCBRYAN_PUBERTAL_BREAST_3_4WK_UP | 183 | 0.4787514 | 1.3918347 | 0 | 0.27842185 | 1 | 3066 | tags=23%, list=11%, signal=26% |
| HP_APLASIA_HYPOPLASIA_OF_THE_CEREBRUM | 1162 | 0.4023329 | 1.3916377 | 0 | 0.4134546 | 1 | 3535 | tags=26%, list=13%, signal=29% |
| WANG_TUMOR_INVASIVENESS_DN | 199 | 0.46658298 | 1.3915256 | 0 | 0.27873042 | 1 | 1785 | tags=29%, list=7%, signal=30% |
| TAVAZOIE_METASTASIS | 71 | 0.5269895 | 1.3909642 | 0.02222222 | 0.27930102 | 1 | 206 | tags=7%, list=1%, signal=7% |
| CAIRO_LIVER_DEVELOPMENT_DN | 179 | 0.4523268 | 1.39094 | 0 | 0.2790589 | 1 | 3667 | tags=28%, list=14%, signal=32% |
| REACTOME_NOTCH4_INTRACELLULAR_DOMAIN_REGULATES_TRANSCRIPTION | 19 | 0.6734397 | 1.390636 | 0 | 0.2793418 | 1 | 333 | tags=21%, list=1%, signal=21% |
| SHETH_LIVER_CANCER_VS_TXNIP_LOSS_PAM3 | 66 | 0.55399144 | 1.3905545 | 0 | 0.27885783 | 1 | 3174 | tags=30%, list=12%, signal=34% |
| AKL_HTLV1_INFECTION_DN | 64 | 0.54123044 | 1.3903656 | 0 | 0.27875254 | 1 | 2842 | tags=27%, list=11%, signal=30% |
| CONCANNON_APOPTOSIS_BY_EPOXOMICIN_DN | 155 | 0.46219397 | 1.3901925 | 0 | 0.2788473 | 1 | 3057 | tags=30%, list=11%, signal=34% |
| GO_MESENCHYMAL_CELL_DIFFERENTIATION_INVOLVED_IN_KIDNEY_DEVELOPMENT | 5 | 0.94693303 | 1.3901776 | 0 | 0.41700596 | 1 | 72 | tags=20%, list=0%, signal=20% |
| GO_EXOPEPTIDASE_ACTIVITY | 76 | 0.5377473 | 1.3901563 | 0.02702703 | 0.41683316 | 1 | 3419 | tags=26%, list=13%, signal=30% |
| QI_PLASMACYTOMA_DN | 92 | 0.5173974 | 1.3900219 | 0.025 | 0.2788982 | 1 | 1963 | tags=26%, list=7%, signal=28% |
| GO_CELLULAR_LIPID_METABOLIC_PROCESS | 902 | 0.41583902 | 1.3898859 | 0 | 0.41763458 | 1 | 3924 | tags=27%, list=15%, signal=31% |
| HP_ABNORMALITY_OF_FACIAL_MUSCULATURE | 269 | 0.4423441 | 1.389665 | 0 | 0.4180285 | 1 | 2819 | tags=21%, list=10%, signal=23% |
| SCHLOSSER_SERUM_RESPONSE_DN | 648 | 0.41917437 | 1.3896054 | 0 | 0.27968127 | 1 | 2407 | tags=26%, list=9%, signal=28% |
| SHETH_LIVER_CANCER_VS_TXNIP_LOSS_PAM2 | 152 | 0.48837274 | 1.3895549 | 0.03030303 | 0.2793993 | 1 | 2011 | tags=24%, list=7%, signal=25% |
| JIANG_AGING_HYPOTHALAMUS_UP | 44 | 0.5734589 | 1.3894242 | 0.04347826 | 0.27928334 | 1 | 3013 | tags=43%, list=11%, signal=49% |
| NOUSHMEHR_GBM_SILENCED_BY_METHYLATION | 45 | 0.55634034 | 1.3891908 | 0 | 0.2795217 | 1 | 5057 | tags=49%, list=19%, signal=60% |
| GO_REACTIVE_OXYGEN_SPECIES_METABOLIC_PROCESS | 225 | 0.45201388 | 1.3890084 | 0 | 0.41968322 | 1 | 2750 | tags=20%, list=10%, signal=22% |
| FERRANDO_T_ALL_WITH_MLL_ENL_FUSION_UP | 78 | 0.53027743 | 1.388768 | 0.02941177 | 0.27968764 | 1 | 2661 | tags=26%, list=10%, signal=28% |
| RUTELLA_RESPONSE_TO_CSF2RB_AND_IL4_UP | 311 | 0.4453304 | 1.388728 | 0 | 0.2794897 | 1 | 3088 | tags=29%, list=11%, signal=32% |
| KEGG_TYPE_I_DIABETES_MELLITUS | 28 | 0.6236812 | 1.3886427 | 0.02702703 | 0.27905345 | 1 | 2140 | tags=25%, list=8%, signal=27% |
| MIKKELSEN_MCV6_LCP_WITH_H3K4ME3 | 140 | 0.49030712 | 1.3886181 | 0.03448276 | 0.27879512 | 1 | 2976 | tags=18%, list=11%, signal=20% |
| HP_INVOLUNTARY_MOVEMENTS | 835 | 0.42719024 | 1.3884183 | 0 | 0.4209012 | 1 | 4106 | tags=33%, list=15%, signal=38% |
| GO_SMOOTHENED_SIGNALING_PATHWAY | 127 | 0.5166793 | 1.3883287 | 0 | 0.420873 | 1 | 4118 | tags=31%, list=15%, signal=36% |
| GO_NEGATIVE_REGULATION_OF_CELL_CYCLE_PHASE_TRANSITION | 227 | 0.44541138 | 1.3878886 | 0 | 0.42202368 | 1 | 3146 | tags=30%, list=12%, signal=34% |
| GO_REGULATION_OF_NEUTROPHIL_EXTRAVASATION | 4 | 0.9372369 | 1.3876499 | 0 | 0.42223528 | 1 | 1547 | tags=50%, list=6%, signal=53% |
| HP_ABNORMALITY_OF_VITAMIN_K_METABOLISM | 4 | 0.9486674 | 1.387622 | 0 | 0.42198676 | 1 | 1367 | tags=75%, list=5%, signal=79% |
| REACTOME_TRANSCRIPTIONAL_REGULATION_BY_RUNX1 | 167 | 0.4745264 | 1.3874241 | 0.02777778 | 0.2803785 | 1 | 2352 | tags=26%, list=9%, signal=29% |
| ZHAN_MULTIPLE_MYELOMA_SUBGROUPS | 29 | 0.59049803 | 1.3874224 | 0.03225806 | 0.28005812 | 1 | 3281 | tags=62%, list=12%, signal=71% |
| GO_POSITIVE_REGULATION_OF_SIGNALING | 1620 | 0.3981381 | 1.3873719 | 0 | 0.42216116 | 1 | 3128 | tags=21%, list=12%, signal=22% |
| GO_RESPONSE_TO_GROWTH_HORMONE | 31 | 0.5906248 | 1.3873405 | 0.04761905 | 0.4217275 | 1 | 2374 | tags=26%, list=9%, signal=28% |
| GO_BRANCHING_INVOLVED_IN_BLOOD_VESSEL_MORPHOGENESIS | 30 | 0.6622504 | 1.3872024 | 0.04878049 | 0.42210075 | 1 | 2044 | tags=20%, list=8%, signal=22% |
| GO_ACTIN_FILAMENT_BASED_MOVEMENT | 132 | 0.47931296 | 1.3871957 | 0 | 0.42178097 | 1 | 3268 | tags=23%, list=12%, signal=26% |
| KOYAMA_SEMA3B_TARGETS_DN | 323 | 0.4614327 | 1.3871429 | 0 | 0.28013757 | 1 | 3410 | tags=31%, list=13%, signal=35% |
| WP_METHYLATION_PATHWAYS | 6 | 0.85331494 | 1.387091 | 0.04761905 | 0.2799019 | 1 | 2799 | tags=50%, list=10%, signal=56% |
| GO_POSITIVE_REGULATION_OF_BINDING | 159 | 0.47014582 | 1.3869969 | 0 | 0.42229855 | 1 | 2271 | tags=25%, list=8%, signal=27% |
| WP_NEUROINFLAMMATION | 12 | 0.69587743 | 1.3864534 | 0.04878049 | 0.28081954 | 1 | 323 | tags=17%, list=1%, signal=17% |
| HP_BONE_MARROW_HYPOCELLULARITY | 46 | 0.58168536 | 1.3862307 | 0.02325581 | 0.42371973 | 1 | 3421 | tags=39%, list=13%, signal=45% |
| GO_INTRACILIARY_TRANSPORT_INVOLVED_IN_CILIUM_ASSEMBLY | 39 | 0.57791775 | 1.3859223 | 0.02325581 | 0.42449793 | 1 | 4017 | tags=46%, list=15%, signal=54% |
| HP_DYSPNEA | 345 | 0.44400248 | 1.3858235 | 0 | 0.4241844 | 1 | 3049 | tags=22%, list=11%, signal=25% |
| GO_ENDOPLASMIC_RETICULUM_TO_GOLGI_VESICLE_MEDIATED_TRANSPORT | 184 | 0.4776348 | 1.3857728 | 0.03125 | 0.42405772 | 1 | 2779 | tags=32%, list=10%, signal=36% |
| HSIAO_HOUSEKEEPING_GENES | 378 | 0.43786848 | 1.3857584 | 0 | 0.2818051 | 1 | 1857 | tags=33%, list=7%, signal=34% |
| GO_CELLULAR_RESPONSE_TO_TOXIC_SUBSTANCE | 100 | 0.517756 | 1.3856238 | 0 | 0.42438814 | 1 | 2750 | tags=25%, list=10%, signal=28% |
| HP_HYPOCHLOREMIA | 3 | 0.9855251 | 1.3855543 | 0 | 0.42408732 | 1 | 173 | tags=33%, list=1%, signal=34% |
| BOYAULT_LIVER_CANCER_SUBCLASS_G3_DN | 46 | 0.5677629 | 1.3854645 | 0.02777778 | 0.28223968 | 1 | 2994 | tags=28%, list=11%, signal=32% |
| JOHNSTONE_PARVB_TARGETS_3_UP | 384 | 0.43365017 | 1.3850309 | 0 | 0.28286532 | 1 | 3080 | tags=28%, list=11%, signal=31% |
| STEGER_ADIPOGENESIS_UP | 15 | 0.76820666 | 1.3847395 | 0.02857143 | 0.28307417 | 1 | 1646 | tags=20%, list=6%, signal=21% |
| GO_ENZYME_INHIBITOR_ACTIVITY | 279 | 0.4443944 | 1.3846712 | 0 | 0.4267235 | 1 | 2669 | tags=22%, list=10%, signal=24% |
| GO_POSITIVE_REGULATION_OF_IMMUNE_SYSTEM_PROCESS | 818 | 0.42092726 | 1.3846307 | 0 | 0.4267034 | 1 | 2587 | tags=18%, list=10%, signal=19% |
| HP_NEUROLOGICAL_SPEECH_IMPAIRMENT | 960 | 0.42060077 | 1.3844178 | 0 | 0.42692956 | 1 | 3510 | tags=27%, list=13%, signal=30% |
| HUTTMANN_B_CLL_POOR_SURVIVAL_UP | 244 | 0.45466784 | 1.3839334 | 0 | 0.28437153 | 1 | 3199 | tags=24%, list=12%, signal=27% |
| GO_CELL_CYCLE_G2_M_PHASE_TRANSITION | 253 | 0.45340365 | 1.3837217 | 0 | 0.42816517 | 1 | 2547 | tags=25%, list=9%, signal=27% |
| HIRSCH_CELLULAR_TRANSFORMATION_SIGNATURE_DN | 96 | 0.5116363 | 1.3835245 | 0.03225806 | 0.2843673 | 1 | 2454 | tags=30%, list=9%, signal=33% |
| BLANCO_MELO_RESPIRATORY_SYNCYTIAL_VIRUS_INFECTION_A594_CELLS_UP | 248 | 0.4575534 | 1.3832241 | 0.03333334 | 0.28493455 | 1 | 2659 | tags=21%, list=10%, signal=23% |
| GO_INNER_EAR_RECEPTOR_CELL_DEVELOPMENT | 39 | 0.59798276 | 1.3830822 | 0.04878049 | 0.42995074 | 1 | 4982 | tags=54%, list=19%, signal=66% |
| GO_NEGATIVE_REGULATION_OF_INTRACELLULAR_SIGNAL_TRANSDUCTION | 451 | 0.423384 | 1.383035 | 0 | 0.42985734 | 1 | 3025 | tags=25%, list=11%, signal=28% |
| GO_LYSOSOMAL_LUMEN | 85 | 0.51940614 | 1.3829708 | 0.03030303 | 0.42974052 | 1 | 3112 | tags=36%, list=12%, signal=41% |
| ZHU_CMV_8_HR_UP | 35 | 0.6309256 | 1.3829241 | 0.02857143 | 0.28529587 | 1 | 2267 | tags=29%, list=8%, signal=31% |
| WOO_LIVER_CANCER_RECURRENCE_DN | 66 | 0.55598116 | 1.3825737 | 0.03448276 | 0.28598455 | 1 | 3826 | tags=29%, list=14%, signal=33% |
| HP_HYPERREFLEXIA | 603 | 0.4335849 | 1.382443 | 0 | 0.43085533 | 1 | 3524 | tags=26%, list=13%, signal=30% |
| GO_SULFUR_COMPOUND_BIOSYNTHETIC_PROCESS | 169 | 0.4596384 | 1.3819789 | 0.02941177 | 0.43216518 | 1 | 4290 | tags=36%, list=16%, signal=43% |
| GO_PROTEIN_HOMODIMERIZATION_ACTIVITY | 568 | 0.42482543 | 1.381769 | 0 | 0.43234497 | 1 | 3561 | tags=25%, list=13%, signal=28% |
| DURAND_STROMA_NS_UP | 158 | 0.47794253 | 1.3817128 | 0 | 0.28649822 | 1 | 3588 | tags=27%, list=13%, signal=30% |
| GO_MUSCLE_CELL_PROLIFERATION | 170 | 0.46289337 | 1.3816752 | 0 | 0.43234578 | 1 | 2814 | tags=24%, list=10%, signal=27% |
| HP_ABNORMAL_ELECTROPHYSIOLOGY_OF_SINOATRIAL_NODE_ORIGIN | 56 | 0.5336872 | 1.3816319 | 0.02857143 | 0.43219247 | 1 | 3268 | tags=27%, list=12%, signal=30% |
| GO_RESPONSE_TO_INTERLEUKIN_1 | 176 | 0.47458804 | 1.381608 | 0.02631579 | 0.43199065 | 1 | 2352 | tags=24%, list=9%, signal=26% |
| REACTOME_TP53_REGULATES_METABOLIC_GENES | 83 | 0.5122718 | 1.3812301 | 0.025 | 0.28698283 | 1 | 1836 | tags=27%, list=7%, signal=28% |
| HP_ABNORMAL_INFLAMMATORY_RESPONSE | 734 | 0.4154512 | 1.381138 | 0 | 0.4327361 | 1 | 3526 | tags=21%, list=13%, signal=24% |
| HP_HYPOKALEMIC_ALKALOSIS | 4 | 0.967771 | 1.3808907 | 0.02040816 | 0.43330312 | 1 | 161 | tags=25%, list=1%, signal=25% |
| VISALA_AGING_LYMPHOCYTE_DN | 16 | 0.7330536 | 1.3795238 | 0.02325581 | 0.2890087 | 1 | 2457 | tags=38%, list=9%, signal=41% |
| GO_MORPHOGENESIS_OF_AN_EPITHELIUM | 496 | 0.42452604 | 1.379079 | 0 | 0.4370021 | 1 | 2661 | tags=18%, list=10%, signal=20% |
| HP_ABNORMAL_CIRCULATING_ESTROGEN_LEVEL | 33 | 0.5842999 | 1.3790398 | 0 | 0.43656293 | 1 | 2301 | tags=27%, list=9%, signal=30% |
| HP_ABNORMALITY_OF_THE_ENDOCRINE_SYSTEM | 973 | 0.39843905 | 1.3789964 | 0 | 0.43577144 | 1 | 3144 | tags=21%, list=12%, signal=23% |
| GO_TRANSPORT_VESICLE | 352 | 0.43072775 | 1.3789465 | 0 | 0.43561724 | 1 | 3199 | tags=29%, list=12%, signal=32% |
| WP_INTRAFLAGELLAR_TRANSPORT_PROTEINS_BINDING_TO_DYNEIN | 27 | 0.6354088 | 1.3789302 | 0.02173913 | 0.28987518 | 1 | 3477 | tags=44%, list=13%, signal=51% |
| GO_ENDOSOME | 852 | 0.42260742 | 1.3787488 | 0 | 0.43578824 | 1 | 3205 | tags=25%, list=12%, signal=28% |
| GO_PROTEIN_LOCALIZATION_TO_MITOCHONDRION | 132 | 0.49153584 | 1.3787152 | 0.03030303 | 0.43557665 | 1 | 2410 | tags=32%, list=9%, signal=35% |
| HP_ABNORMAL_BLOOD_GLUCOSE_CONCENTRATION | 198 | 0.47274938 | 1.3784865 | 0 | 0.43580472 | 1 | 3528 | tags=29%, list=13%, signal=33% |
| HP_ARTHRITIS | 159 | 0.49626708 | 1.3783529 | 0 | 0.4359419 | 1 | 3471 | tags=28%, list=13%, signal=32% |
| GO_TISSUE_REMODELING | 145 | 0.48041686 | 1.3781419 | 0.03703704 | 0.43601957 | 1 | 2931 | tags=21%, list=11%, signal=23% |
| REACTOME_APC_C_MEDIATED_DEGRADATION_OF_CELL_CYCLE_PROTEINS | 84 | 0.52019566 | 1.3774756 | 0.03571429 | 0.29195458 | 1 | 2375 | tags=39%, list=9%, signal=43% |
| HP_ABNORMAL_CRANIAL_NERVE_PHYSIOLOGY | 257 | 0.45056236 | 1.3774009 | 0 | 0.43790448 | 1 | 3597 | tags=26%, list=13%, signal=29% |
| GO_CHANNEL_REGULATOR_ACTIVITY | 124 | 0.47558075 | 1.3773077 | 0 | 0.43785506 | 1 | 3358 | tags=27%, list=12%, signal=30% |
| KAYO_AGING_MUSCLE_DN | 114 | 0.49615523 | 1.3772414 | 0 | 0.2920474 | 1 | 3182 | tags=36%, list=12%, signal=41% |
| GO_UBIQUINONE_BINDING | 6 | 0.902101 | 1.3769405 | 0.02631579 | 0.43755805 | 1 | 707 | tags=50%, list=3%, signal=51% |
| GO_PEPTIDASE_ACTIVITY | 475 | 0.42962676 | 1.3763753 | 0 | 0.4372193 | 1 | 3436 | tags=21%, list=13%, signal=24% |
| HP_ABNORMAL_NASAL_MUCUS_SECRETION | 4 | 0.9466478 | 1.375958 | 0.0212766 | 0.43778032 | 1 | 707 | tags=25%, list=3%, signal=26% |
| GO_MUSCLE_CONTRACTION | 289 | 0.44927034 | 1.375943 | 0 | 0.4375365 | 1 | 2874 | tags=16%, list=11%, signal=18% |
| REACTOME_FC_EPSILON_RECEPTOR_FCERI_SIGNALING | 125 | 0.49873698 | 1.3757402 | 0.02941177 | 0.2944605 | 1 | 2389 | tags=28%, list=9%, signal=31% |
| GO_METENCEPHALON_DEVELOPMENT | 104 | 0.49114242 | 1.3756279 | 0.02564103 | 0.43849292 | 1 | 2371 | tags=19%, list=9%, signal=21% |
| GO_NADH_METABOLIC_PROCESS | 38 | 0.58424616 | 1.3756012 | 0.05 | 0.43822664 | 1 | 3141 | tags=42%, list=12%, signal=48% |
| YOSHIMURA_MAPK8_TARGETS_UP | 989 | 0.41157326 | 1.3755934 | 0 | 0.29451013 | 1 | 3056 | tags=18%, list=11%, signal=19% |
| GO_DIGESTIVE_SYSTEM_DEVELOPMENT | 117 | 0.49473518 | 1.3754873 | 0.02380952 | 0.4383935 | 1 | 2675 | tags=17%, list=10%, signal=19% |
| VANTVEER_BREAST_CANCER_METASTASIS_DN | 102 | 0.5099729 | 1.3751794 | 0.02380952 | 0.29450408 | 1 | 2228 | tags=25%, list=8%, signal=27% |
| TURASHVILI_BREAST_LOBULAR_CARCINOMA_VS_LOBULAR_NORMAL_UP | 82 | 0.50564814 | 1.374172 | 0 | 0.29652953 | 1 | 3160 | tags=24%, list=12%, signal=28% |
| GO_REGULATION_OF_OSSIFICATION | 173 | 0.4628358 | 1.3739433 | 0 | 0.44288737 | 1 | 2008 | tags=15%, list=7%, signal=16% |
| LI_INDUCED_T_TO_NATURAL_KILLER_UP | 290 | 0.4571943 | 1.3736347 | 0 | 0.29726854 | 1 | 2598 | tags=21%, list=10%, signal=23% |
| GO_PHOSPHATIDYLCHOLINE_METABOLIC_PROCESS | 71 | 0.53020287 | 1.3731074 | 0.02702703 | 0.44528267 | 1 | 3911 | tags=30%, list=15%, signal=35% |
| GO_ICOSANOID_CATABOLIC_PROCESS | 5 | 0.9407716 | 1.3729712 | 0.01886793 | 0.445444 | 1 | 832 | tags=40%, list=3%, signal=41% |
| GO_CD4_POSITIVE_CD25_POSITIVE_ALPHA_BETA_REGULATORY_T_CELL_DIFFERENTIATION | 4 | 0.91784656 | 1.3729627 | 0.04761905 | 0.44517463 | 1 | 383 | tags=25%, list=1%, signal=25% |
| GO_ION_TRANSMEMBRANE_TRANSPORT | 987 | 0.40667564 | 1.372756 | 0 | 0.44518077 | 1 | 3471 | tags=19%, list=13%, signal=21% |
| GO_ORGANIC_HYDROXY_COMPOUND_METABOLIC_PROCESS | 459 | 0.4322412 | 1.3723359 | 0 | 0.446258 | 1 | 3628 | tags=25%, list=13%, signal=28% |
| PILON_KLF1_TARGETS_UP | 434 | 0.42388338 | 1.372324 | 0 | 0.29903427 | 1 | 3103 | tags=25%, list=12%, signal=27% |
| ROYLANCE_BREAST_CANCER_16Q_COPY_NUMBER_UP | 52 | 0.56422323 | 1.3721331 | 0.02631579 | 0.29925382 | 1 | 4332 | tags=38%, list=16%, signal=46% |
| GO_RNA_POLYMERASE_I_COMPLEX | 12 | 0.7288941 | 1.3719217 | 0.04444445 | 0.44667903 | 1 | 2452 | tags=58%, list=9%, signal=64% |
| GO_NEGATIVE_REGULATION_OF_ESTABLISHMENT_OF_PROTEIN_LOCALIZATION | 131 | 0.5031894 | 1.371805 | 0.04347826 | 0.4468036 | 1 | 3215 | tags=28%, list=12%, signal=32% |
| KRIGE_RESPONSE_TO_TOSEDOSTAT_6HR_DN | 841 | 0.4162765 | 1.3717355 | 0 | 0.29968783 | 1 | 3181 | tags=29%, list=12%, signal=32% |
| BASSO_CD40_SIGNALING_UP | 95 | 0.5022815 | 1.3716062 | 0.03333334 | 0.29952016 | 1 | 1703 | tags=21%, list=6%, signal=22% |
| BLUM_RESPONSE_TO_SALIRASIB_UP | 236 | 0.467526 | 1.3706993 | 0.04761905 | 0.3003009 | 1 | 3485 | tags=36%, list=13%, signal=41% |
| MARTINEZ_TP53_TARGETS_DN | 532 | 0.4204758 | 1.3705761 | 0 | 0.30013162 | 1 | 2784 | tags=26%, list=10%, signal=29% |
| REACTOME_CLASS_I_MHC_MEDIATED_ANTIGEN_PROCESSING_PRESENTATION | 339 | 0.43532342 | 1.3703134 | 0 | 0.3004818 | 1 | 2375 | tags=22%, list=9%, signal=24% |
| GO_TRANSEPITHELIAL_TRANSPORT | 23 | 0.63784236 | 1.3702403 | 0.03571429 | 0.4513377 | 1 | 2353 | tags=26%, list=9%, signal=29% |
| ZWANG_EGF_INTERVAL_DN | 181 | 0.47865102 | 1.3701284 | 0.03225806 | 0.30009022 | 1 | 3756 | tags=30%, list=14%, signal=35% |
| GO_PROTON_TRANSPORTING_TWO_SECTOR_ATPASE_COMPLEX_ASSEMBLY | 9 | 0.82134014 | 1.3700944 | 0.04878049 | 0.45135605 | 1 | 4364 | tags=89%, list=16%, signal=106% |
| MANNE_COVID19_NONICU_VS_HEALTHY_DONOR_PLATELETS_UP | 125 | 0.4824849 | 1.3700743 | 0.03333334 | 0.29992443 | 1 | 2212 | tags=27%, list=8%, signal=29% |
| REACTOME_TERMINAL_PATHWAY_OF_COMPLEMENT | 3 | 0.97631824 | 1.3700674 | 0 | 0.29958552 | 1 | 60 | tags=33%, list=0%, signal=33% |
| SPIELMAN_LYMPHOBLAST_EUROPEAN_VS_ASIAN_UP | 460 | 0.4355828 | 1.3700391 | 0 | 0.2993232 | 1 | 3095 | tags=31%, list=11%, signal=35% |
| GO_ION_TRANSPORT | 1405 | 0.40680042 | 1.3695936 | 0 | 0.4510308 | 1 | 3440 | tags=19%, list=13%, signal=20% |
| GO_NEGATIVE_REGULATION_OF_SIGNALING | 1234 | 0.40748525 | 1.3691285 | 0 | 0.45240915 | 1 | 3348 | tags=24%, list=12%, signal=26% |
| GO_PROTON_TRANSPORTING_ATP_SYNTHASE_COMPLEX_CATALYTIC_CORE_F_1 | 5 | 0.96366066 | 1.3689696 | 0.025 | 0.45270705 | 1 | 982 | tags=100%, list=4%, signal=104% |
| GO_VESICLE_MEMBRANE | 710 | 0.41935915 | 1.3687639 | 0 | 0.45320588 | 1 | 3171 | tags=23%, list=12%, signal=25% |
| HP_ABNORMAL_PROTEIN_GLYCOSYLATION | 30 | 0.65383697 | 1.3685318 | 0.04166667 | 0.45370176 | 1 | 3527 | tags=47%, list=13%, signal=54% |
| HP_MYOSITIS | 24 | 0.66748136 | 1.3682586 | 0.05 | 0.45411548 | 1 | 1967 | tags=29%, list=7%, signal=31% |
| GO_MULTICELLULAR_ORGANISMAL_IRON_ION_HOMEOSTASIS | 7 | 0.79742104 | 1.368197 | 0.04444445 | 0.45403403 | 1 | 5119 | tags=57%, list=19%, signal=71% |
| GAVIN_FOXP3_TARGETS_CLUSTER_P2 | 70 | 0.5350928 | 1.3679826 | 0.04 | 0.302351 | 1 | 3060 | tags=30%, list=11%, signal=34% |
| HP_RENAL_AGENESIS | 121 | 0.4748516 | 1.3679206 | 0.03703704 | 0.45444784 | 1 | 949 | tags=12%, list=4%, signal=12% |
| ZHU_CMV_ALL_DN | 107 | 0.48369431 | 1.3671361 | 0.05 | 0.30398095 | 1 | 2812 | tags=32%, list=10%, signal=35% |
| GO_POSITIVE_REGULATION_OF_OSSIFICATION | 79 | 0.5279599 | 1.3667935 | 0.04761905 | 0.45687672 | 1 | 2647 | tags=20%, list=10%, signal=22% |
| GO_POSITIVE_REGULATION_OF_RECEPTOR_SIGNALING_PATHWAY_VIA_STAT | 66 | 0.5154439 | 1.3666073 | 0.02857143 | 0.45713368 | 1 | 2768 | tags=20%, list=10%, signal=22% |
| GAUSSMANN_MLL_AF4_FUSION_TARGETS_A_DN | 80 | 0.50945735 | 1.3660427 | 0.02564103 | 0.30585197 | 1 | 1449 | tags=16%, list=5%, signal=17% |
| GO_REGULATION_OF_PROTEIN_CATABOLIC_PROCESS | 370 | 0.4479717 | 1.365998 | 0 | 0.45877472 | 1 | 2654 | tags=26%, list=10%, signal=29% |
| KYNG_DNA_DAMAGE_DN | 169 | 0.4733693 | 1.3658688 | 0 | 0.30611375 | 1 | 2582 | tags=31%, list=10%, signal=34% |
| HP_FUNCTIONAL_MOTOR_DEFICIT | 369 | 0.44256663 | 1.3658262 | 0 | 0.45880425 | 1 | 2694 | tags=21%, list=10%, signal=23% |
| GO_CELLULAR_RESPONSE_TO_EXOGENOUS_DSRNA | 13 | 0.7229053 | 1.3656834 | 0.04347826 | 0.45884544 | 1 | 1446 | tags=38%, list=5%, signal=41% |
| GO_REGULATION_OF_SMALL_MOLECULE_METABOLIC_PROCESS | 399 | 0.4430419 | 1.3648492 | 0 | 0.4610073 | 1 | 2623 | tags=23%, list=10%, signal=25% |
| GO_REGULATION_OF_INSULIN_SECRETION | 151 | 0.44549328 | 1.3646907 | 0 | 0.4613278 | 1 | 2389 | tags=17%, list=9%, signal=19% |
| LENAOUR_DENDRITIC_CELL_MATURATION_DN | 118 | 0.485764 | 1.3645277 | 0.03571429 | 0.3082272 | 1 | 2353 | tags=25%, list=9%, signal=27% |
| GO_LYASE_ACTIVITY | 164 | 0.4622937 | 1.3644952 | 0 | 0.4616898 | 1 | 2726 | tags=24%, list=10%, signal=27% |
| HP_TYPE_2_MUSCLE_FIBER_ATROPHY | 11 | 0.7472904 | 1.3644222 | 0.04347826 | 0.46165937 | 1 | 4184 | tags=55%, list=16%, signal=65% |
| HOFFMANN_SMALL_PRE_BII_TO_IMMATURE_B_LYMPHOCYTE_UP | 58 | 0.5178208 | 1.363976 | 0.02941177 | 0.30931187 | 1 | 4106 | tags=36%, list=15%, signal=43% |
| MARTORIATI_MDM4_TARGETS_FETAL_LIVER_UP | 215 | 0.4747899 | 1.3639098 | 0 | 0.30911762 | 1 | 2235 | tags=25%, list=8%, signal=27% |
| WU_CELL_MIGRATION | 158 | 0.4601821 | 1.3636672 | 0.02857143 | 0.30939442 | 1 | 2889 | tags=23%, list=11%, signal=26% |
| WP_MICRORNA_NETWORK_ASSOCIATED_WITH_CHRONIC_LYMPHOCYTIC_LEUKEMIA | 3 | 0.92746705 | 1.3634733 | 0.02083333 | 0.30950305 | 1 | 84 | tags=33%, list=0%, signal=33% |
| IVANOVA_HEMATOPOIESIS_LATE_PROGENITOR | 471 | 0.43966046 | 1.3634213 | 0 | 0.30936715 | 1 | 3611 | tags=29%, list=13%, signal=33% |
| DAVICIONI_TARGETS_OF_PAX_FOXO1_FUSIONS_UP | 234 | 0.44332117 | 1.3631525 | 0 | 0.30954608 | 1 | 3599 | tags=31%, list=13%, signal=35% |
| GO_RESPONSE_TO_PEPTIDE | 470 | 0.43432307 | 1.3630619 | 0 | 0.46486613 | 1 | 2535 | tags=21%, list=9%, signal=23% |
| GO_NUCLEOBASE_CONTAINING_SMALL_MOLECULE_METABOLIC_PROCESS | 546 | 0.42098358 | 1.3630259 | 0 | 0.46462715 | 1 | 3223 | tags=25%, list=12%, signal=28% |
| WANG_LMO4_TARGETS_UP | 315 | 0.43882763 | 1.3629862 | 0 | 0.30956084 | 1 | 3607 | tags=33%, list=13%, signal=38% |
| MATSUDA_NATURAL_KILLER_DIFFERENTIATION | 446 | 0.4287235 | 1.362861 | 0 | 0.3096843 | 1 | 3401 | tags=28%, list=13%, signal=31% |
| HP_SKELETAL_MUSCLE_ATROPHY | 429 | 0.4260764 | 1.3626815 | 0 | 0.46523494 | 1 | 3817 | tags=28%, list=14%, signal=33% |
| HP_ATROPHY_DEGENERATION_AFFECTING_THE_CENTRAL_NERVOUS_SYSTEM | 669 | 0.40082625 | 1.3623234 | 0 | 0.46605617 | 1 | 4103 | tags=32%, list=15%, signal=37% |
| HP_MYOCLONUS | 261 | 0.43422034 | 1.3618617 | 0 | 0.4662666 | 1 | 4106 | tags=34%, list=15%, signal=39% |
| HP_DELAYED_ABILITY_TO_SIT | 7 | 0.86134046 | 1.3617648 | 0 | 0.4662442 | 1 | 1888 | tags=71%, list=7%, signal=77% |
| REACTOME_G2_M_CHECKPOINTS | 129 | 0.49341837 | 1.3614228 | 0.02941177 | 0.31244785 | 1 | 3138 | tags=35%, list=12%, signal=39% |
| HP_ABNORMAL_THYROID_MORPHOLOGY | 109 | 0.48509377 | 1.3612964 | 0.02702703 | 0.46672317 | 1 | 3352 | tags=25%, list=12%, signal=28% |
| HP_OPISTHOTONUS | 32 | 0.61746544 | 1.3608879 | 0.04166667 | 0.46743906 | 1 | 4283 | tags=44%, list=16%, signal=52% |
| HP_POSTNATAL_MICROCEPHALY | 129 | 0.46716422 | 1.3608764 | 0 | 0.46714762 | 1 | 2500 | tags=22%, list=9%, signal=25% |
| WAKABAYASHI_ADIPOGENESIS_PPARG_RXRA_BOUND_36HR | 139 | 0.4740174 | 1.3608524 | 0 | 0.31290996 | 1 | 3586 | tags=35%, list=13%, signal=40% |
| GO_MALE_SEX_DIFFERENTIATION | 139 | 0.49202028 | 1.3607585 | 0.03703704 | 0.46698433 | 1 | 2845 | tags=23%, list=11%, signal=26% |
| NAKAMURA_METASTASIS | 44 | 0.56352746 | 1.3605705 | 0.05 | 0.31325075 | 1 | 1709 | tags=27%, list=6%, signal=29% |
| GO_CYTOKINE_MEDIATED_SIGNALING_PATHWAY | 637 | 0.42212498 | 1.3605268 | 0 | 0.46681878 | 1 | 3182 | tags=24%, list=12%, signal=27% |
| GO_ENDOSOME_MEMBRANE | 446 | 0.4274107 | 1.3605051 | 0 | 0.46661425 | 1 | 2759 | tags=25%, list=10%, signal=28% |
| HP_ABNORMALITY_OF_IMMUNE_SYSTEM_PHYSIOLOGY | 1112 | 0.4202891 | 1.3604906 | 0 | 0.466378 | 1 | 3197 | tags=21%, list=12%, signal=23% |
| GO_FATTY_ACID_BIOSYNTHETIC_PROCESS | 136 | 0.4752188 | 1.3604223 | 0 | 0.46636695 | 1 | 3643 | tags=29%, list=14%, signal=34% |
| HP_PLATYSPONDYLY | 98 | 0.49159554 | 1.3602333 | 0 | 0.46635327 | 1 | 3956 | tags=35%, list=15%, signal=41% |
| GO_REGULATION_OF_GLIAL_CELL_DIFFERENTIATION | 67 | 0.53237987 | 1.360227 | 0.02702703 | 0.4660541 | 1 | 1003 | tags=15%, list=4%, signal=15% |
| GO_SPERM_PLASMA_MEMBRANE | 4 | 0.94801176 | 1.3597845 | 0.02380952 | 0.4671245 | 1 | 376 | tags=25%, list=1%, signal=25% |
| REACTOME_INTERLEUKIN_27_SIGNALING | 9 | 0.86314565 | 1.3596925 | 0.04166667 | 0.3146672 | 1 | 1939 | tags=33%, list=7%, signal=36% |
| HP_DYSARTHRIA | 448 | 0.42269453 | 1.3590226 | 0 | 0.4683586 | 1 | 3839 | tags=30%, list=14%, signal=34% |
| HP_ABNORMAL_FEAR_ANXIETY_RELATED_BEHAVIOR | 270 | 0.44155997 | 1.3588488 | 0 | 0.46870387 | 1 | 2446 | tags=19%, list=9%, signal=21% |
| GO_RESPONSE_TO_ABIOTIC_STIMULUS | 1099 | 0.41554734 | 1.3588414 | 0 | 0.46843687 | 1 | 2530 | tags=18%, list=9%, signal=19% |
| GO_CIRCULATORY_SYSTEM_DEVELOPMENT | 962 | 0.39539942 | 1.3587358 | 0 | 0.46845517 | 1 | 3322 | tags=20%, list=12%, signal=22% |
| GO_RESPONSE_TO_MOLECULE_OF_BACTERIAL_ORIGIN | 284 | 0.44773608 | 1.3586568 | 0 | 0.46839064 | 1 | 2604 | tags=17%, list=10%, signal=19% |
| GO_REGULATION_OF_HYDROLASE_ACTIVITY | 1082 | 0.39850754 | 1.35824 | 0 | 0.46951345 | 1 | 2968 | tags=21%, list=11%, signal=22% |
| REACTOME_REGULATION_OF_MRNA_STABILITY_BY_PROTEINS_THAT_BIND_AU_RICH_ELEMENTS | 82 | 0.51760703 | 1.3580415 | 0 | 0.3172183 | 1 | 805 | tags=27%, list=3%, signal=28% |
| GO_NEGATIVE_REGULATION_OF_MITOTIC_CELL_CYCLE | 297 | 0.42081752 | 1.357902 | 0 | 0.46944872 | 1 | 3170 | tags=28%, list=12%, signal=31% |
| HP_ABNORMALITY_OF_FACIAL_SOFT_TISSUE | 358 | 0.43046057 | 1.3577653 | 0 | 0.46974915 | 1 | 4051 | tags=31%, list=15%, signal=36% |
| GO_RESPONSE_TO_VITAMIN | 76 | 0.5048922 | 1.3575965 | 0.02702703 | 0.46947604 | 1 | 627 | tags=9%, list=2%, signal=9% |
| HP_UNUSUAL_INFECTION_BY_ANATOMICAL_SITE | 70 | 0.508879 | 1.357555 | 0 | 0.46941012 | 1 | 1967 | tags=16%, list=7%, signal=17% |
| ONKEN_UVEAL_MELANOMA_DN | 503 | 0.41273662 | 1.3572509 | 0 | 0.3183103 | 1 | 3406 | tags=32%, list=13%, signal=36% |
| HP_MYALGIA | 117 | 0.47198355 | 1.3571005 | 0.02941177 | 0.47034472 | 1 | 4252 | tags=34%, list=16%, signal=40% |
| GO_GLYCOPROTEIN_METABOLIC_PROCESS | 349 | 0.43022665 | 1.3570707 | 0 | 0.47020465 | 1 | 3547 | tags=24%, list=13%, signal=28% |
| REACTOME_UB_SPECIFIC_PROCESSING_PROTEASES | 159 | 0.46853244 | 1.3570623 | 0 | 0.3183688 | 1 | 2352 | tags=28%, list=9%, signal=30% |
| CHICAS_RB1_TARGETS_GROWING | 226 | 0.46113324 | 1.3569635 | 0 | 0.3178259 | 1 | 2552 | tags=25%, list=9%, signal=28% |
| HP_INCREASED_BLOOD_PRESSURE | 299 | 0.4392159 | 1.3568333 | 0 | 0.47072124 | 1 | 3497 | tags=23%, list=13%, signal=27% |
| GO_TERTIARY_GRANULE | 140 | 0.4930465 | 1.3567692 | 0.03448276 | 0.47067568 | 1 | 2094 | tags=19%, list=8%, signal=21% |
| RUIZ_TNC_TARGETS_UP | 144 | 0.48246968 | 1.3566585 | 0.02631579 | 0.31832275 | 1 | 4002 | tags=37%, list=15%, signal=43% |
| REACTOME_CYCLIN_A_CDK2_ASSOCIATED_EVENTS_AT_S_PHASE_ENTRY | 82 | 0.51355165 | 1.356409 | 0.05 | 0.31845176 | 1 | 2352 | tags=33%, list=9%, signal=36% |
| GO_EPIDERMAL_CELL_DIFFERENTIATION | 175 | 0.45105395 | 1.356277 | 0 | 0.4711446 | 1 | 3651 | tags=21%, list=14%, signal=24% |
| GO_PIGMENT_GRANULE | 101 | 0.5054481 | 1.3562746 | 0.03030303 | 0.47083884 | 1 | 2257 | tags=30%, list=8%, signal=32% |
| HP_ABNORMALITY_OF_THE_THYROID_GLAND | 301 | 0.4476943 | 1.3562094 | 0 | 0.47050884 | 1 | 3128 | tags=22%, list=12%, signal=24% |
| GO_UROGENITAL_SYSTEM_DEVELOPMENT | 295 | 0.44893664 | 1.3562081 | 0 | 0.4702041 | 1 | 3127 | tags=20%, list=12%, signal=23% |
| HP_ABNORMALITY_OF_CARDIOVASCULAR_SYSTEM_ELECTROPHYSIOLOGY | 365 | 0.43728545 | 1.3558875 | 0 | 0.47126874 | 1 | 2025 | tags=19%, list=8%, signal=20% |
| HP_ABNORMALITY_OF_RENAL_EXCRETION | 36 | 0.5926283 | 1.3556813 | 0 | 0.47098595 | 1 | 2876 | tags=19%, list=11%, signal=22% |
| GO_NEGATIVE_REGULATION_OF_CELL_CYCLE_PROCESS | 306 | 0.42537555 | 1.3554536 | 0 | 0.471328 | 1 | 3307 | tags=29%, list=12%, signal=33% |
| GO_TISSUE_MORPHOGENESIS | 585 | 0.41268831 | 1.3553842 | 0 | 0.47137606 | 1 | 2661 | tags=17%, list=10%, signal=18% |
| GO_CELLULAR_METABOLIC_COMPOUND_SALVAGE | 30 | 0.59309244 | 1.3552674 | 0.04761905 | 0.47154644 | 1 | 5007 | tags=53%, list=19%, signal=65% |
| GO_REGULATION_OF_PROTEIN_TYROSINE_KINASE_ACTIVITY | 87 | 0.5041865 | 1.3550646 | 0 | 0.47144082 | 1 | 3067 | tags=26%, list=11%, signal=30% |
| HP_ABNORMAL_SYSTEMIC_ARTERIAL_MORPHOLOGY | 355 | 0.42883044 | 1.3549982 | 0 | 0.4714477 | 1 | 2999 | tags=23%, list=11%, signal=25% |
| KAN_RESPONSE_TO_ARSENIC_TRIOXIDE | 104 | 0.47713837 | 1.3545537 | 0 | 0.32122374 | 1 | 923 | tags=19%, list=3%, signal=20% |
| GO_CELL_DIFFERENTIATION_INVOLVED_IN_KIDNEY_DEVELOPMENT | 51 | 0.5281412 | 1.3544332 | 0.03125 | 0.47208002 | 1 | 1528 | tags=14%, list=6%, signal=15% |
| GO_STEROL_TRANSPORT | 83 | 0.50451565 | 1.3542589 | 0.02564103 | 0.47222292 | 1 | 2434 | tags=22%, list=9%, signal=24% |
| GO_AZUROPHIL_GRANULE_MEMBRANE | 51 | 0.506285 | 1.3538655 | 0.04761905 | 0.47261783 | 1 | 3112 | tags=33%, list=12%, signal=38% |
| GO_HYDRO_LYASE_ACTIVITY | 49 | 0.5619536 | 1.3537847 | 0.02564103 | 0.47259307 | 1 | 3318 | tags=37%, list=12%, signal=42% |
| GO_RNA_POLYMERASE_ACTIVITY | 38 | 0.6024684 | 1.353699 | 0.04651163 | 0.47261858 | 1 | 2758 | tags=39%, list=10%, signal=44% |
| RICKMAN_METASTASIS_DN | 232 | 0.4521199 | 1.3536712 | 0.03225806 | 0.3229657 | 1 | 3052 | tags=22%, list=11%, signal=24% |
| GO_RESPONSE_TO_XENOBIOTIC_STIMULUS | 80 | 0.5056198 | 1.3534319 | 0.02941177 | 0.47331864 | 1 | 2691 | tags=23%, list=10%, signal=25% |
| VANDESLUIS_COMMD1_TARGETS_GROUP_3_UP | 69 | 0.55632263 | 1.3531758 | 0 | 0.3230526 | 1 | 2420 | tags=25%, list=9%, signal=27% |
| HP_ARACHNOID_HEMANGIOMATOSIS | 15 | 0.7049555 | 1.3531102 | 0.05 | 0.4744135 | 1 | 2744 | tags=53%, list=10%, signal=59% |
| GO_INTERLEUKIN_4_PRODUCTION | 27 | 0.640127 | 1.3530807 | 0.02857143 | 0.47429466 | 1 | 1346 | tags=26%, list=5%, signal=27% |
| HP_MICROCYTIC_ANEMIA | 27 | 0.63400763 | 1.3528305 | 0.04545455 | 0.47472945 | 1 | 1511 | tags=22%, list=6%, signal=24% |
| GO_PURINE_CONTAINING_COMPOUND_METABOLIC_PROCESS | 411 | 0.41976586 | 1.35244 | 0 | 0.47465706 | 1 | 3223 | tags=25%, list=12%, signal=28% |
| WP_BREAST_CANCER_PATHWAY | 142 | 0.45746827 | 1.3524282 | 0.03030303 | 0.32411247 | 1 | 2330 | tags=19%, list=9%, signal=21% |
| WOOD_EBV_EBNA1_TARGETS_DN | 41 | 0.57130307 | 1.3524196 | 0.04878049 | 0.32380345 | 1 | 1592 | tags=29%, list=6%, signal=31% |
| SASAKI_ADULT_T_CELL_LEUKEMIA | 156 | 0.47824925 | 1.3523976 | 0 | 0.32353166 | 1 | 2305 | tags=25%, list=9%, signal=27% |
| GO_TUBE_DEVELOPMENT | 928 | 0.40347564 | 1.3520758 | 0 | 0.47542694 | 1 | 3862 | tags=24%, list=14%, signal=27% |
| GO_NEGATIVE_REGULATION_OF_MOLECULAR_FUNCTION | 942 | 0.39950672 | 1.3520647 | 0 | 0.47518688 | 1 | 3185 | tags=22%, list=12%, signal=25% |
| GO_LIPID_BINDING | 656 | 0.40531856 | 1.3518528 | 0 | 0.47548652 | 1 | 3167 | tags=21%, list=12%, signal=23% |
| HP_BRAIN_ATROPHY | 591 | 0.4067334 | 1.3517901 | 0 | 0.4754398 | 1 | 4103 | tags=32%, list=15%, signal=37% |
| GO_CELLULAR_RESPONSE_TO_OSMOTIC_STRESS | 38 | 0.54079014 | 1.3512402 | 0.02564103 | 0.47663078 | 1 | 2438 | tags=24%, list=9%, signal=26% |
| PASQUALUCCI_LYMPHOMA_BY_GC_STAGE_UP | 264 | 0.4275542 | 1.3512385 | 0 | 0.3233292 | 1 | 3944 | tags=36%, list=15%, signal=41% |
| GO_SUPRAMOLECULAR_POLYMER | 788 | 0.4097499 | 1.3511864 | 0 | 0.4765126 | 1 | 3292 | tags=23%, list=12%, signal=26% |
| HP_RESPIRATORY_TRACT_INFECTION | 434 | 0.43390855 | 1.351178 | 0 | 0.47625348 | 1 | 3549 | tags=24%, list=13%, signal=27% |
| HP_CLINICAL_COURSE | 1271 | 0.3935462 | 1.3510125 | 0 | 0.47630957 | 1 | 4106 | tags=29%, list=15%, signal=33% |
| ZHU_CMV_24_HR_DN | 75 | 0.484991 | 1.3509473 | 0.03333334 | 0.32375565 | 1 | 2812 | tags=32%, list=10%, signal=36% |
| WP_EUKARYOTIC_TRANSCRIPTION_INITIATION | 40 | 0.62315977 | 1.3506831 | 0.03225806 | 0.3237344 | 1 | 3095 | tags=45%, list=11%, signal=51% |
| GO_CATION_TRANSMEMBRANE_TRANSPORTER_ACTIVITY | 534 | 0.42107776 | 1.3506497 | 0 | 0.47661784 | 1 | 3463 | tags=17%, list=13%, signal=20% |
| GO_RESPONSE_TO_ORGANIC_CYCLIC_COMPOUND | 805 | 0.4080074 | 1.3506483 | 0 | 0.47632924 | 1 | 2530 | tags=17%, list=9%, signal=19% |
| HP_OSTEOPENIA | 191 | 0.46509635 | 1.3505144 | 0 | 0.47661218 | 1 | 3151 | tags=25%, list=12%, signal=28% |
| GO_NEURAL_PRECURSOR_CELL_PROLIFERATION | 136 | 0.48166034 | 1.3503819 | 0 | 0.47676605 | 1 | 2148 | tags=21%, list=8%, signal=22% |
| REACTOME_COLLAGEN_BIOSYNTHESIS_AND_MODIFYING_ENZYMES | 64 | 0.5344712 | 1.3500975 | 0.02777778 | 0.3247038 | 1 | 2882 | tags=25%, list=11%, signal=28% |
| GO_REGULATION_OF_RESPONSE_TO_STRESS | 1264 | 0.3901992 | 1.3498307 | 0 | 0.47731975 | 1 | 2947 | tags=21%, list=11%, signal=22% |
| REACTOME_BIOLOGICAL_OXIDATIONS | 133 | 0.46638104 | 1.349785 | 0 | 0.32505074 | 1 | 2900 | tags=23%, list=11%, signal=26% |
| GO_PRECATALYTIC_SPLICEOSOME | 50 | 0.56481063 | 1.3497841 | 0.02325581 | 0.47710276 | 1 | 2138 | tags=44%, list=8%, signal=48% |
| NABA_CORE_MATRISOME | 221 | 0.45808077 | 1.349705 | 0.02702703 | 0.32496068 | 1 | 3097 | tags=20%, list=12%, signal=23% |
| GO_CELLULAR_PROTEIN_CONTAINING_COMPLEX_ASSEMBLY | 915 | 0.4099235 | 1.3496073 | 0 | 0.47685942 | 1 | 2448 | tags=24%, list=9%, signal=26% |
| HP_INFANTILE_ONSET | 407 | 0.42242584 | 1.3495243 | 0 | 0.47695106 | 1 | 3746 | tags=28%, list=14%, signal=32% |
| HP_CHRONIC_SINUSITIS | 46 | 0.51470435 | 1.3493752 | 0.04878049 | 0.47705615 | 1 | 5223 | tags=28%, list=19%, signal=35% |
| LIU_COMMON_CANCER_GENES | 54 | 0.5495049 | 1.3493575 | 0.02222222 | 0.3249309 | 1 | 3287 | tags=33%, list=12%, signal=38% |
| HP_INTELLECTUAL_DISABILITY_SEVERE | 313 | 0.42920616 | 1.3491795 | 0 | 0.47723863 | 1 | 2095 | tags=21%, list=8%, signal=23% |
| ZHENG_GLIOBLASTOMA_PLASTICITY_UP | 238 | 0.44313142 | 1.3491486 | 0 | 0.3248 | 1 | 2098 | tags=26%, list=8%, signal=28% |
| GO_CELLULAR_RESPONSE_TO_OXYGEN_CONTAINING_COMPOUND | 1032 | 0.413862 | 1.3487729 | 0 | 0.47835463 | 1 | 2614 | tags=19%, list=10%, signal=20% |
| GO_AUTOPHAGOSOME_MATURATION | 35 | 0.60156727 | 1.348362 | 0 | 0.47947648 | 1 | 2189 | tags=34%, list=8%, signal=37% |
| GO_RECEPTOR_SIGNALING_PATHWAY_VIA_STAT | 114 | 0.48139927 | 1.3482562 | 0 | 0.4791319 | 1 | 2784 | tags=25%, list=10%, signal=28% |
| GO_TRANSMEMBRANE_RECEPTOR_PROTEIN_TYROSINE_KINASE_SIGNALING_PATHWAY | 654 | 0.40758726 | 1.3481832 | 0 | 0.4791721 | 1 | 2775 | tags=22%, list=10%, signal=24% |
| KAAB_FAILED_HEART_ATRIUM_DN | 141 | 0.4739459 | 1.348025 | 0 | 0.32689807 | 1 | 2448 | tags=26%, list=9%, signal=29% |
| MOROSETTI_FACIOSCAPULOHUMERAL_MUSCULAR_DISTROPHY_DN | 11 | 0.7050455 | 1.3479766 | 0.05 | 0.326612 | 1 | 2626 | tags=27%, list=10%, signal=30% |
| GO_NEGATIVE_REGULATION_OF_HYDROGEN_PEROXIDE_MEDIATED_PROGRAMMED_CELL_DEATH | 4 | 0.8902931 | 1.347776 | 0.02439024 | 0.4806755 | 1 | 284 | tags=25%, list=1%, signal=25% |
| HP_ABNORMALITY_OF_MUSCLE_SIZE | 532 | 0.41075322 | 1.3474579 | 0 | 0.48171005 | 1 | 3000 | tags=23%, list=11%, signal=25% |
| GO_POSITIVE_REGULATION_OF_ATPASE_ACTIVITY | 50 | 0.5480727 | 1.3470902 | 0.03030303 | 0.48183456 | 1 | 3727 | tags=42%, list=14%, signal=49% |
| GO_CELL_BODY | 555 | 0.41137466 | 1.3469557 | 0 | 0.4821284 | 1 | 1854 | tags=14%, list=7%, signal=15% |
| GO_TETRAPYRROLE_BIOSYNTHETIC_PROCESS | 25 | 0.6030629 | 1.3468226 | 0.05 | 0.48208693 | 1 | 3741 | tags=40%, list=14%, signal=46% |
| PURBEY_TARGETS_OF_CTBP1_NOT_SATB1_DN | 342 | 0.41754192 | 1.3465313 | 0 | 0.32820433 | 1 | 3144 | tags=24%, list=12%, signal=27% |
| KYNG_ENVIRONMENTAL_STRESS_RESPONSE_NOT_BY_4NQO_IN_WS | 35 | 0.53077954 | 1.346194 | 0.02702703 | 0.32851028 | 1 | 1454 | tags=31%, list=5%, signal=33% |
| HP_GENERALIZED_HYPOTONIA | 876 | 0.41675273 | 1.3460928 | 0 | 0.4831808 | 1 | 3281 | tags=28%, list=12%, signal=31% |
| REACTOME_PEPTIDE_LIGAND_BINDING_RECEPTORS | 123 | 0.4793099 | 1.3460487 | 0 | 0.32851925 | 1 | 2555 | tags=11%, list=9%, signal=13% |
| WINTER_HYPOXIA_METAGENE | 216 | 0.46063113 | 1.3460177 | 0 | 0.32825133 | 1 | 1962 | tags=22%, list=7%, signal=23% |
| GO_CENTRAL_NERVOUS_SYSTEM_NEURON_DIFFERENTIATION | 169 | 0.47952452 | 1.345936 | 0 | 0.48353294 | 1 | 2352 | tags=15%, list=9%, signal=17% |
| GO_U2_TYPE_SPLICEOSOMAL_COMPLEX | 88 | 0.51029074 | 1.3456111 | 0.03448276 | 0.4843735 | 1 | 2161 | tags=39%, list=8%, signal=42% |
| KRIGE_RESPONSE_TO_TOSEDOSTAT_24HR_DN | 930 | 0.40501282 | 1.3453128 | 0 | 0.328802 | 1 | 2771 | tags=27%, list=10%, signal=29% |
| GO_MECHANORECEPTOR_DIFFERENTIATION | 57 | 0.5183493 | 1.3452946 | 0 | 0.48484617 | 1 | 4765 | tags=42%, list=18%, signal=51% |
| HP_ABNORMALITY_OF_THYROID_PHYSIOLOGY | 225 | 0.4412638 | 1.3452219 | 0 | 0.4845189 | 1 | 2315 | tags=16%, list=9%, signal=18% |
| HP_KYPHOSCOLIOSIS | 148 | 0.45668036 | 1.3449274 | 0 | 0.48542482 | 1 | 3968 | tags=32%, list=15%, signal=37% |
| HAHTOLA_MYCOSIS_FUNGOIDES_SKIN_UP | 169 | 0.4709727 | 1.3448875 | 0 | 0.32926095 | 1 | 2697 | tags=31%, list=10%, signal=35% |
| GO_ADULT_BEHAVIOR | 115 | 0.47366142 | 1.3443389 | 0 | 0.48653647 | 1 | 2331 | tags=23%, list=9%, signal=25% |
| REACTOME_DIGESTION_OF_DIETARY_LIPID | 2 | 0.98469365 | 1.3441312 | 0.02173913 | 0.32990745 | 1 | 398 | tags=50%, list=1%, signal=51% |
| FARMER_BREAST_CANCER_APOCRINE_VS_BASAL | 292 | 0.42958674 | 1.3441274 | 0 | 0.32958832 | 1 | 2864 | tags=26%, list=11%, signal=29% |
| MARTIN_VIRAL_GPCR_SIGNALING_DN | 45 | 0.5578207 | 1.3439126 | 0.05 | 0.32966688 | 1 | 2040 | tags=22%, list=8%, signal=24% |
| SHEDDEN_LUNG_CANCER_GOOD_SURVIVAL_A12 | 218 | 0.441728 | 1.3436886 | 0 | 0.32933122 | 1 | 3744 | tags=24%, list=14%, signal=28% |
| GO_FATTY_ACID_DERIVATIVE_METABOLIC_PROCESS | 122 | 0.4660564 | 1.3436437 | 0.02702703 | 0.48758954 | 1 | 2617 | tags=23%, list=10%, signal=25% |
| HP_RECURRENT_ACUTE_RESPIRATORY_TRACT_INFECTION | 10 | 0.75917494 | 1.3435702 | 0.05 | 0.4872716 | 1 | 1315 | tags=30%, list=5%, signal=32% |
| ZHOU_TNF_SIGNALING_4HR | 46 | 0.5665072 | 1.3435625 | 0 | 0.32902327 | 1 | 2290 | tags=39%, list=9%, signal=43% |
| WANG_RESPONSE_TO_GSK3_INHIBITOR_SB216763_UP | 298 | 0.43859857 | 1.3434249 | 0 | 0.32908425 | 1 | 3782 | tags=29%, list=14%, signal=33% |
| REACTOME_S_PHASE | 158 | 0.45285064 | 1.3433336 | 0 | 0.3289221 | 1 | 2375 | tags=28%, list=9%, signal=31% |
| HP_DECREASED_PROPORTION_OF_CD8_POSITIVE_T_CELLS | 6 | 0.81277895 | 1.3431488 | 0.03030303 | 0.48795635 | 1 | 1125 | tags=17%, list=4%, signal=17% |
| KYNG_DNA_DAMAGE_BY_UV | 52 | 0.5317487 | 1.3430178 | 0.02941177 | 0.3290593 | 1 | 3608 | tags=31%, list=13%, signal=35% |
| GO_NEPHRON_DEVELOPMENT | 126 | 0.46843988 | 1.3427463 | 0 | 0.48855186 | 1 | 4500 | tags=32%, list=17%, signal=38% |
| GO_RESPONSE_TO_LIPID | 768 | 0.40597662 | 1.34206 | 0 | 0.4902429 | 1 | 2614 | tags=18%, list=10%, signal=19% |
| GO_SMALL_MOLECULE_BIOSYNTHETIC_PROCESS | 618 | 0.4123068 | 1.3420436 | 0 | 0.48998708 | 1 | 3108 | tags=22%, list=12%, signal=24% |
| GO_VACUOLE_ORGANIZATION | 160 | 0.4521883 | 1.341998 | 0.03448276 | 0.48992324 | 1 | 4245 | tags=41%, list=16%, signal=49% |
| GO_SUPRAMOLECULAR_FIBER_ORGANIZATION | 635 | 0.4144747 | 1.3419608 | 0 | 0.48974526 | 1 | 3413 | tags=24%, list=13%, signal=27% |
| GO_TAP_BINDING | 3 | 0.9512167 | 1.3419511 | 0.02631579 | 0.48949975 | 1 | 1315 | tags=100%, list=5%, signal=105% |
| HP_MUSCULAR_HYPOTONIA_OF_THE_TRUNK | 146 | 0.45192996 | 1.3418566 | 0 | 0.48957178 | 1 | 4106 | tags=38%, list=15%, signal=44% |
| TONKS_TARGETS_OF_RUNX1_RUNX1T1_FUSION_HSC_UP | 162 | 0.4852869 | 1.3418128 | 0.02564103 | 0.330394 | 1 | 3656 | tags=34%, list=14%, signal=39% |
| HP_STRIDOR | 24 | 0.6534163 | 1.3417903 | 0.05 | 0.48919678 | 1 | 2302 | tags=17%, list=9%, signal=18% |
| HP_ABNORMALITY_OF_THE_CEREBRAL_SUBCORTEX | 829 | 0.40371615 | 1.3414991 | 0 | 0.49003386 | 1 | 4140 | tags=31%, list=15%, signal=36% |
| REACTOME_EXTRACELLULAR_MATRIX_ORGANIZATION | 277 | 0.45505294 | 1.3414755 | 0 | 0.33049232 | 1 | 2756 | tags=19%, list=10%, signal=21% |
| LOPEZ_MBD_TARGETS | 874 | 0.4055751 | 1.3410736 | 0 | 0.33107412 | 1 | 3156 | tags=30%, list=12%, signal=32% |
| DIAZ_CHRONIC_MEYLOGENOUS_LEUKEMIA_UP | 1335 | 0.404993 | 1.3406811 | 0 | 0.33071783 | 1 | 3564 | tags=34%, list=13%, signal=37% |
| GO_REGULATION_OF_PEPTIDYL_TYROSINE_PHOSPHORYLATION | 219 | 0.43625206 | 1.3405259 | 0 | 0.49162933 | 1 | 2808 | tags=19%, list=10%, signal=21% |
| LEE_LIVER_CANCER_SURVIVAL_UP | 146 | 0.45546877 | 1.340487 | 0 | 0.33094501 | 1 | 2912 | tags=18%, list=11%, signal=21% |
| GO_REGULATION_OF_METAL_ION_TRANSPORT | 324 | 0.4168954 | 1.3401612 | 0 | 0.4922327 | 1 | 2728 | tags=16%, list=10%, signal=17% |
| GO_CLATHRIN_COATED_VESICLE | 167 | 0.47430798 | 1.3395383 | 0 | 0.4937007 | 1 | 3171 | tags=30%, list=12%, signal=34% |
| GO_MALATE_DEHYDROGENASE_ACTIVITY | 7 | 0.87533784 | 1.339018 | 0.02272727 | 0.49508068 | 1 | 2744 | tags=71%, list=10%, signal=80% |
| GO_NEGATIVE_REGULATION_OF_IMMUNE_SYSTEM_PROCESS | 364 | 0.41251627 | 1.3389674 | 0 | 0.49495813 | 1 | 2604 | tags=18%, list=10%, signal=20% |
| HP_SENSORY_NEUROPATHY | 164 | 0.47838363 | 1.3388337 | 0 | 0.49479997 | 1 | 3100 | tags=21%, list=12%, signal=24% |
| REACTOME_PHENYLALANINE_METABOLISM | 3 | 0.94808894 | 1.3388115 | 0.01960784 | 0.33314732 | 1 | 1291 | tags=67%, list=5%, signal=70% |
| HP_PEDIATRIC_ONSET | 512 | 0.414967 | 1.338527 | 0 | 0.49566296 | 1 | 4184 | tags=31%, list=16%, signal=36% |
| CUI_TCF21_TARGETS_2_DN | 793 | 0.41035044 | 1.3384712 | 0 | 0.33365437 | 1 | 3515 | tags=26%, list=13%, signal=29% |
| GO_ELASTIN_CATABOLIC_PROCESS | 3 | 0.953528 | 1.338355 | 0.02439024 | 0.49565473 | 1 | 875 | tags=67%, list=3%, signal=69% |
| GO_VASCULATURE_DEVELOPMENT | 645 | 0.407174 | 1.3381164 | 0 | 0.49642974 | 1 | 3069 | tags=21%, list=11%, signal=23% |
| GO_RNA_POLYMERASE_COMPLEX | 96 | 0.48690215 | 1.33789 | 0 | 0.4969101 | 1 | 2758 | tags=33%, list=10%, signal=37% |
| GO_GOLGI_APPARATUS | 1383 | 0.40017286 | 1.3377445 | 0 | 0.4968813 | 1 | 3672 | tags=26%, list=14%, signal=29% |
| HP_HYPOPLASIA_OF_THE_CORPUS_CALLOSUM | 366 | 0.4290759 | 1.3377261 | 0 | 0.49669272 | 1 | 3526 | tags=28%, list=13%, signal=32% |
| VANOEVELEN_MYOGENESIS_SIN3A_TARGETS | 207 | 0.4396318 | 1.3374511 | 0 | 0.3334519 | 1 | 2833 | tags=28%, list=11%, signal=31% |
| BASAKI_YBX1_TARGETS_DN | 342 | 0.43255448 | 1.337435 | 0 | 0.33317426 | 1 | 2665 | tags=20%, list=10%, signal=22% |
| GO_ANTERIOR_POSTERIOR_PATTERN_SPECIFICATION | 168 | 0.44965658 | 1.3374132 | 0 | 0.49714413 | 1 | 2567 | tags=13%, list=10%, signal=14% |
| GO_REGULATION_OF_CARDIAC_CONDUCTION | 63 | 0.5237928 | 1.3365577 | 0 | 0.49800962 | 1 | 3268 | tags=30%, list=12%, signal=34% |
| GO_REGULATION_OF_CELL_ADHESION | 627 | 0.4011673 | 1.3363217 | 0 | 0.49839178 | 1 | 3120 | tags=20%, list=12%, signal=23% |
| GO_PROTEIN_DIMERIZATION_ACTIVITY | 808 | 0.39784935 | 1.3362939 | 0 | 0.49816668 | 1 | 2591 | tags=19%, list=10%, signal=20% |
| REACTOME_FERTILIZATION | 14 | 0.6894165 | 1.3362017 | 0.02222222 | 0.33424228 | 1 | 312 | tags=7%, list=1%, signal=7% |
| HP_FEEDING_DIFFICULTIES | 740 | 0.40007937 | 1.3361546 | 0 | 0.49795032 | 1 | 3535 | tags=28%, list=13%, signal=31% |
| VART_KSHV_INFECTION_ANGIOGENIC_MARKERS_UP | 142 | 0.46406558 | 1.3360773 | 0.03571429 | 0.3341618 | 1 | 1750 | tags=15%, list=7%, signal=16% |
| GO_LIPID_METABOLIC_PROCESS | 1180 | 0.39495352 | 1.3357071 | 0 | 0.49837345 | 1 | 4093 | tags=27%, list=15%, signal=30% |
| HP_OSTEOPOROSIS | 219 | 0.45902795 | 1.3355796 | 0 | 0.49838084 | 1 | 3198 | tags=23%, list=12%, signal=26% |
| GO_RESPONSE_TO_OXIDATIVE_STRESS | 407 | 0.41099855 | 1.3355601 | 0 | 0.49818465 | 1 | 2938 | tags=22%, list=11%, signal=24% |
| GO_TRANSCRIPTION_BY_RNA_POLYMERASE_I | 58 | 0.5323499 | 1.3353246 | 0.02564103 | 0.4977734 | 1 | 3872 | tags=43%, list=14%, signal=50% |
| RODRIGUES_DCC_TARGETS_DN | 112 | 0.46177208 | 1.3351163 | 0 | 0.33423534 | 1 | 4077 | tags=36%, list=15%, signal=42% |
| GO_MONOSACCHARIDE_CATABOLIC_PROCESS | 58 | 0.49870235 | 1.334804 | 0.02439024 | 0.49936405 | 1 | 1962 | tags=22%, list=7%, signal=24% |
| GO_EPIDERMIS_DEVELOPMENT | 269 | 0.4332487 | 1.3347414 | 0 | 0.49910188 | 1 | 3948 | tags=23%, list=15%, signal=27% |
| HP_SPECIFIC_LEARNING_DISABILITY | 187 | 0.4392213 | 1.3347365 | 0 | 0.4988325 | 1 | 3100 | tags=27%, list=12%, signal=30% |
| GO_HUMORAL_IMMUNE_RESPONSE_MEDIATED_BY_CIRCULATING_IMMUNOGLOBULIN | 41 | 0.54388744 | 1.33444 | 0.02777778 | 0.49977443 | 1 | 2016 | tags=17%, list=7%, signal=18% |
| GO_NEGATIVE_REGULATION_OF_FIBROBLAST_GROWTH_FACTOR_PRODUCTION | 2 | 0.97577155 | 1.334376 | 0.04878049 | 0.49969864 | 1 | 316 | tags=50%, list=1%, signal=51% |
| GO_MITOCHONDRIAL_MEMBRANE_ORGANIZATION | 124 | 0.48308885 | 1.3341396 | 0.04347826 | 0.5002796 | 1 | 2358 | tags=29%, list=9%, signal=32% |
| GO_NEGATIVE_REGULATION_OF_RESPONSE_TO_STIMULUS | 1430 | 0.3985953 | 1.3335345 | 0 | 0.50140387 | 1 | 3379 | tags=23%, list=13%, signal=25% |
| GO_MULTIVESICULAR_BODY_SORTING_PATHWAY | 36 | 0.5845109 | 1.3335273 | 0.02380952 | 0.5011159 | 1 | 2395 | tags=33%, list=9%, signal=37% |
| GAURNIER_PSMD4_TARGETS | 51 | 0.51703995 | 1.3332113 | 0.025 | 0.33804336 | 1 | 1347 | tags=14%, list=5%, signal=14% |
| GO_T_CELL_DIFFERENTIATION | 223 | 0.44767407 | 1.3328533 | 0.03703704 | 0.5018124 | 1 | 2604 | tags=16%, list=10%, signal=18% |
| GO_NEGATIVE_REGULATION_OF_PROTEIN_METABOLIC_PROCESS | 942 | 0.40369627 | 1.3324419 | 0 | 0.5024813 | 1 | 3087 | tags=24%, list=11%, signal=26% |
| GO_ISG15_PROTEIN_CONJUGATION | 5 | 0.8774481 | 1.3318028 | 0.04651163 | 0.5042688 | 1 | 19 | tags=20%, list=0%, signal=20% |
| CHYLA_CBFA2T3_TARGETS_DN | 207 | 0.44563994 | 1.3315556 | 0 | 0.33998394 | 1 | 4335 | tags=28%, list=16%, signal=33% |
| HP_ONSET | 907 | 0.39610347 | 1.331188 | 0 | 0.50497186 | 1 | 4106 | tags=30%, list=15%, signal=35% |
| SPIRA_SMOKERS_LUNG_CANCER_UP | 33 | 0.59022874 | 1.3307352 | 0.04651163 | 0.34113097 | 1 | 1148 | tags=33%, list=4%, signal=35% |
| HP_MULTIPLE_JOINT_CONTRACTURES | 51 | 0.5103477 | 1.3307158 | 0.04878049 | 0.50625515 | 1 | 2833 | tags=25%, list=11%, signal=28% |
| MORI_LARGE_PRE_BII_LYMPHOCYTE_DN | 53 | 0.51563615 | 1.3307114 | 0.05 | 0.3408364 | 1 | 2910 | tags=28%, list=11%, signal=32% |
| WP_CILIARY_LANDSCAPE | 211 | 0.44183454 | 1.3304569 | 0 | 0.34055418 | 1 | 4093 | tags=37%, list=15%, signal=43% |
| REACTOME_SIGNALING_BY_NOTCH | 173 | 0.47254723 | 1.3303422 | 0 | 0.33983648 | 1 | 805 | tags=17%, list=3%, signal=17% |
| MIKKELSEN_IPS_WITH_HCP_H3K27ME3 | 66 | 0.5272968 | 1.3302939 | 0.04444445 | 0.33970726 | 1 | 3455 | tags=18%, list=13%, signal=21% |
| GO_ICOSANOID_METABOLIC_PROCESS | 76 | 0.5009829 | 1.3299661 | 0.04545455 | 0.50756615 | 1 | 2191 | tags=21%, list=8%, signal=23% |
| GO_UBIQUITIN_LIKE_PROTEIN_LIGASE_BINDING | 291 | 0.4162223 | 1.329918 | 0 | 0.5074966 | 1 | 2831 | tags=27%, list=11%, signal=30% |
| GO_REGULATION_OF_CELL_DEATH | 1442 | 0.3833694 | 1.3294045 | 0 | 0.5078816 | 1 | 2963 | tags=21%, list=11%, signal=23% |
| HP_ELBOW_FLEXION_CONTRACTURE | 60 | 0.5283642 | 1.3292378 | 0.04651163 | 0.5080502 | 1 | 3956 | tags=33%, list=15%, signal=39% |
| NABA_ECM_GLYCOPROTEINS | 152 | 0.46139917 | 1.3291894 | 0 | 0.34199804 | 1 | 3097 | tags=22%, list=12%, signal=24% |
| HP_ABNORMAL_CIRCULATING_PROTEIN_LEVEL | 388 | 0.4206456 | 1.3290384 | 0 | 0.5082078 | 1 | 2948 | tags=22%, list=11%, signal=25% |
| GO_PERIPHERAL_NERVOUS_SYSTEM_AXON_REGENERATION | 6 | 0.8761631 | 1.3290358 | 0.04545455 | 0.5079218 | 1 | 29 | tags=33%, list=0%, signal=33% |
| GO_APOPTOTIC_PROCESS | 1676 | 0.38770714 | 1.3287678 | 0 | 0.50836825 | 1 | 3174 | tags=23%, list=12%, signal=25% |
| HP_IMPAIRMENT_IN_PERSONALITY_FUNCTIONING | 510 | 0.41764495 | 1.3285588 | 0 | 0.5090628 | 1 | 2894 | tags=22%, list=11%, signal=24% |
| GO_TRNA_METHYLTRANSFERASE_ACTIVITY | 29 | 0.6460364 | 1.3285204 | 0.02857143 | 0.50896615 | 1 | 4654 | tags=59%, list=17%, signal=71% |
| GO_CARBON_OXYGEN_LYASE_ACTIVITY | 64 | 0.5171188 | 1.3282332 | 0.025 | 0.50978744 | 1 | 3318 | tags=33%, list=12%, signal=37% |
| HP_CONGENITAL_ICHTHYOSIFORM_ERYTHRODERMA | 20 | 0.66741306 | 1.3281865 | 0.04878049 | 0.5096724 | 1 | 3948 | tags=30%, list=15%, signal=35% |
| GO_PROTEIN_CATABOLIC_PROCESS | 858 | 0.4056309 | 1.3277768 | 0 | 0.5111919 | 1 | 2813 | tags=25%, list=10%, signal=27% |
| HP_DEVELOPMENTAL_REGRESSION | 292 | 0.42742217 | 1.3276 | 0 | 0.51106876 | 1 | 4046 | tags=33%, list=15%, signal=39% |
| PETROVA_ENDOTHELIUM_LYMPHATIC_VS_BLOOD_DN | 149 | 0.44920936 | 1.3273772 | 0.02702703 | 0.34519905 | 1 | 2629 | tags=21%, list=10%, signal=24% |
| GO_KIDNEY_EPITHELIUM_DEVELOPMENT | 127 | 0.47210285 | 1.3272303 | 0 | 0.5121095 | 1 | 2587 | tags=17%, list=10%, signal=18% |
| HOLLERN_EMT_BREAST_TUMOR_UP | 135 | 0.4636547 | 1.3267434 | 0 | 0.34606776 | 1 | 2076 | tags=20%, list=8%, signal=22% |
| ELVIDGE_HYPOXIA_BY_DMOG_UP | 124 | 0.49377054 | 1.3263236 | 0.04 | 0.34685874 | 1 | 2531 | tags=23%, list=9%, signal=26% |
| MARTINEZ_TP53_TARGETS_UP | 567 | 0.4073276 | 1.3262138 | 0 | 0.34670746 | 1 | 3288 | tags=26%, list=12%, signal=29% |
| GO_NEGATIVE_REGULATION_OF_CERAMIDE_BIOSYNTHETIC_PROCESS | 4 | 0.86271286 | 1.3261399 | 0 | 0.5139242 | 1 | 2077 | tags=50%, list=8%, signal=54% |
| GO_NEGATIVE_REGULATION_OF_CELL_DEATH | 840 | 0.40186906 | 1.3257778 | 0 | 0.51503706 | 1 | 2803 | tags=22%, list=10%, signal=24% |
| GO_APICAL_PART_OF_CELL | 359 | 0.42650267 | 1.3257352 | 0 | 0.51494753 | 1 | 2743 | tags=16%, list=10%, signal=18% |
| GO_NEURAL_RETINA_DEVELOPMENT | 63 | 0.50487214 | 1.3257055 | 0.03571429 | 0.51450855 | 1 | 1666 | tags=14%, list=6%, signal=15% |
| GO_CELLULAR_RESPONSE_TO_INSULIN_STIMULUS | 205 | 0.44826752 | 1.3255045 | 0 | 0.51470155 | 1 | 2614 | tags=26%, list=10%, signal=29% |
| GO_REGULATION_OF_CALCIUM_ION_TRANSMEMBRANE_TRANSPORTER_ACTIVITY | 71 | 0.507274 | 1.3253018 | 0.025 | 0.51429045 | 1 | 677 | tags=11%, list=3%, signal=12% |
| VANTVEER_BREAST_CANCER_ESR1_UP | 136 | 0.4627516 | 1.3250881 | 0 | 0.3490702 | 1 | 2970 | tags=25%, list=11%, signal=28% |
| GO_MONOOXYGENASE_ACTIVITY | 58 | 0.5005699 | 1.3248816 | 0 | 0.51546866 | 1 | 2720 | tags=12%, list=10%, signal=13% |
| GO_POSITIVE_REGULATION_OF_I_KAPPAB_KINASE_NF_KAPPAB_SIGNALING | 172 | 0.45906845 | 1.3247812 | 0 | 0.5152972 | 1 | 3077 | tags=26%, list=11%, signal=29% |
| GO_REGULATION_OF_PRODUCTION_OF_MOLECULAR_MEDIATOR_OF_IMMUNE_RESPONSE | 114 | 0.47140545 | 1.3243468 | 0.02439024 | 0.5162838 | 1 | 2465 | tags=18%, list=9%, signal=20% |
| HP_ABNORMALITY_OF_THE_LENS | 611 | 0.40257987 | 1.3242185 | 0 | 0.5164833 | 1 | 3524 | tags=23%, list=13%, signal=26% |
| RASHI_RESPONSE_TO_IONIZING_RADIATION_2 | 112 | 0.4730762 | 1.3241904 | 0.03571429 | 0.34996584 | 1 | 2085 | tags=26%, list=8%, signal=28% |
| GO_REGULATION_OF_CATABOLIC_PROCESS | 950 | 0.39581972 | 1.323909 | 0 | 0.516972 | 1 | 2925 | tags=25%, list=11%, signal=27% |
| WANG_CLIM2_TARGETS_DN | 164 | 0.45734075 | 1.3238149 | 0.03571429 | 0.35026148 | 1 | 2810 | tags=31%, list=10%, signal=35% |
| VECCHI_GASTRIC_CANCER_ADVANCED_VS_EARLY_UP | 154 | 0.45089972 | 1.3237269 | 0.03225806 | 0.35018605 | 1 | 3104 | tags=21%, list=12%, signal=23% |
| HP_CHRONIC_CONSTIPATION | 32 | 0.5985964 | 1.3236121 | 0.02631579 | 0.51672417 | 1 | 1853 | tags=22%, list=7%, signal=23% |
| KEGG_ASCORBATE_AND_ALDARATE_METABOLISM | 7 | 0.827006 | 1.3231483 | 0.05 | 0.35113937 | 1 | 3769 | tags=57%, list=14%, signal=66% |
| RODRIGUES_THYROID_CARCINOMA_POORLY_DIFFERENTIATED_DN | 692 | 0.40512556 | 1.3229667 | 0 | 0.35125187 | 1 | 3799 | tags=29%, list=14%, signal=33% |
| WANG_TUMOR_INVASIVENESS_UP | 366 | 0.41357297 | 1.3225706 | 0 | 0.35160616 | 1 | 2400 | tags=34%, list=9%, signal=37% |
| HP_UNUSUAL_INFECTION | 654 | 0.41551572 | 1.3225424 | 0 | 0.51896757 | 1 | 3471 | tags=24%, list=13%, signal=27% |
| MCBRYAN_PUBERTAL_TGFB1_TARGETS_DN | 59 | 0.5021714 | 1.3223159 | 0.02564103 | 0.35167575 | 1 | 4000 | tags=37%, list=15%, signal=44% |
| SANSOM_APC_TARGETS_UP | 114 | 0.47994366 | 1.3222735 | 0.02777778 | 0.35149083 | 1 | 3675 | tags=39%, list=14%, signal=45% |
| REACTOME_FLT3_SIGNALING | 275 | 0.422425 | 1.3222393 | 0 | 0.35127378 | 1 | 2352 | tags=20%, list=9%, signal=22% |
| GO_ONE_CARBON_METABOLIC_PROCESS | 35 | 0.5972664 | 1.3221401 | 0.02777778 | 0.518522 | 1 | 3194 | tags=29%, list=12%, signal=32% |
| GO_POSITIVE_REGULATION_OF_PEPTIDYL_TYROSINE_PHOSPHORYLATION | 163 | 0.47252962 | 1.3215737 | 0 | 0.5203063 | 1 | 2566 | tags=18%, list=10%, signal=20% |
| SHIPP_DLBCL_VS_FOLLICULAR_LYMPHOMA_UP | 44 | 0.56941915 | 1.3215162 | 0.04347826 | 0.3524589 | 1 | 2270 | tags=41%, list=8%, signal=45% |
| GO_IRON_ION_BINDING | 105 | 0.4669033 | 1.3211318 | 0.03703704 | 0.5205461 | 1 | 2644 | tags=16%, list=10%, signal=18% |
| GO_NEPHRON_EPITHELIUM_DEVELOPMENT | 96 | 0.4973346 | 1.3208873 | 0.03030303 | 0.52088517 | 1 | 2587 | tags=18%, list=10%, signal=20% |
| ACOSTA_PROLIFERATION_INDEPENDENT_MYC_TARGETS_DN | 105 | 0.47319835 | 1.3204939 | 0.02941177 | 0.3543808 | 1 | 1939 | tags=16%, list=7%, signal=17% |
| MATTIOLI_MGUS_VS_PCL | 92 | 0.47156453 | 1.3201877 | 0.02702703 | 0.35474357 | 1 | 1832 | tags=26%, list=7%, signal=28% |
| REACTOME_NEDDYLATION | 217 | 0.46549752 | 1.3199896 | 0.03448276 | 0.3548543 | 1 | 2909 | tags=28%, list=11%, signal=31% |
| GO_REGULATION_OF_DNA_TEMPLATED_TRANSCRIPTION_IN_RESPONSE_TO_STRESS | 108 | 0.4873574 | 1.31992 | 0 | 0.52311087 | 1 | 805 | tags=20%, list=3%, signal=21% |
| GO_PERINUCLEAR_REGION_OF_CYTOPLASM | 662 | 0.41073382 | 1.3193372 | 0 | 0.524659 | 1 | 2399 | tags=20%, list=9%, signal=22% |
| GO_GOLGI_MEMBRANE | 661 | 0.3934924 | 1.3192291 | 0 | 0.524636 | 1 | 3236 | tags=25%, list=12%, signal=28% |
| HP_ABNORMALITY_OF_SKULL_SIZE | 1205 | 0.38298076 | 1.3191624 | 0 | 0.52458656 | 1 | 3281 | tags=25%, list=12%, signal=27% |
| HP_CYANOSIS | 64 | 0.4783706 | 1.3181806 | 0.03125 | 0.52550846 | 1 | 2302 | tags=16%, list=9%, signal=17% |
| JOHNSTONE_PARVB_TARGETS_3_DN | 784 | 0.3940494 | 1.3177894 | 0 | 0.35824797 | 1 | 3970 | tags=36%, list=15%, signal=42% |
| HESS_TARGETS_OF_HOXA9_AND_MEIS1_DN | 80 | 0.44078258 | 1.3174306 | 0.03225806 | 0.3585539 | 1 | 3588 | tags=26%, list=13%, signal=30% |
| GO_PLATELET_AGGREGATION | 53 | 0.53804016 | 1.3172431 | 0.03030303 | 0.52661127 | 1 | 2334 | tags=26%, list=9%, signal=29% |
| GO_RESPONSE_TO_PEPTIDE_HORMONE | 393 | 0.42740563 | 1.3171686 | 0 | 0.52660376 | 1 | 2535 | tags=21%, list=9%, signal=23% |
| MCBRYAN_PUBERTAL_BREAST_6_7WK_DN | 73 | 0.49968076 | 1.3171418 | 0.04444445 | 0.35906297 | 1 | 2205 | tags=25%, list=8%, signal=27% |
| MEBARKI_HCC_PROGENITOR_FZD8CRD_DN | 333 | 0.4288976 | 1.3166525 | 0 | 0.3596623 | 1 | 3878 | tags=19%, list=14%, signal=22% |
| HP_GONOSOMAL_INHERITANCE | 218 | 0.43524012 | 1.3165256 | 0 | 0.52813655 | 1 | 2815 | tags=22%, list=10%, signal=24% |
| GO_POSITIVE_REGULATION_OF_PRODUCTION_OF_MOLECULAR_MEDIATOR_OF_IMMUNE_RESPONSE | 80 | 0.48453948 | 1.3164535 | 0.02631579 | 0.5281361 | 1 | 2465 | tags=15%, list=9%, signal=16% |
| LEE_NEURAL_CREST_STEM_CELL_UP | 131 | 0.4664933 | 1.3155754 | 0 | 0.36022753 | 1 | 2587 | tags=22%, list=10%, signal=24% |
| REACTOME_DISEASES_OF_SIGNAL_TRANSDUCTION_BY_GROWTH_FACTOR_RECEPTORS_AND_SECOND_MESSENGERS | 369 | 0.41817296 | 1.3153203 | 0 | 0.3605174 | 1 | 2758 | tags=25%, list=10%, signal=27% |
| GO_ENZYME_LINKED_RECEPTOR_PROTEIN_SIGNALING_PATHWAY | 938 | 0.39326444 | 1.3150219 | 0 | 0.53049916 | 1 | 2775 | tags=19%, list=10%, signal=21% |
| GO_MESONEPHROS_DEVELOPMENT | 90 | 0.46726322 | 1.3144976 | 0 | 0.53085756 | 1 | 2160 | tags=13%, list=8%, signal=14% |
| GO_REGULATION_OF_CELL_SHAPE | 146 | 0.46086675 | 1.3144897 | 0.04166667 | 0.53057736 | 1 | 3076 | tags=25%, list=11%, signal=28% |
| REACTOME_POST_TRANSLATIONAL_PROTEIN_MODIFICATION | 1231 | 0.38285357 | 1.314221 | 0 | 0.36224326 | 1 | 3318 | tags=25%, list=12%, signal=28% |
| HP_ABNORMAL_SPERMATOGENESIS | 96 | 0.4788814 | 1.3140458 | 0.025 | 0.53176725 | 1 | 4023 | tags=21%, list=15%, signal=24% |
| GAVIN_FOXP3_TARGETS_CLUSTER_P7 | 84 | 0.5016321 | 1.3137715 | 0.02777778 | 0.36265972 | 1 | 2906 | tags=27%, list=11%, signal=31% |
| HP_HEPATIC_FIBROSIS | 104 | 0.45863536 | 1.313698 | 0 | 0.5320039 | 1 | 4790 | tags=45%, list=18%, signal=55% |
| REACTOME_CHROMOSOME_MAINTENANCE | 88 | 0.4822022 | 1.3133949 | 0.02439024 | 0.36271626 | 1 | 3325 | tags=39%, list=12%, signal=44% |
| LANDIS_ERBB2_BREAST_TUMORS_324_DN | 138 | 0.46233687 | 1.3128566 | 0 | 0.36318475 | 1 | 3794 | tags=37%, list=14%, signal=43% |
| GO_CALCIUM_CHANNEL_REGULATOR_ACTIVITY | 39 | 0.56287503 | 1.3120942 | 0.03125 | 0.53453827 | 1 | 639 | tags=13%, list=2%, signal=13% |
| PECE_MAMMARY_STEM_CELL_UP | 117 | 0.4854816 | 1.3118844 | 0.03030303 | 0.36505347 | 1 | 1373 | tags=32%, list=5%, signal=34% |
| HP_ABNORMALITY_OF_BONE_MARROW_CELL_MORPHOLOGY | 140 | 0.4440843 | 1.3118365 | 0 | 0.53473735 | 1 | 3535 | tags=31%, list=13%, signal=36% |
| HP_EMG_MYOPATHIC_ABNORMALITIES | 98 | 0.4693541 | 1.311678 | 0.03571429 | 0.5348562 | 1 | 2315 | tags=18%, list=9%, signal=20% |
| GO_SIGNALING_RECEPTOR_BINDING | 1253 | 0.3800544 | 1.311616 | 0 | 0.53456646 | 1 | 3083 | tags=18%, list=11%, signal=20% |
| GO_GOLGI_VESICLE_TRANSPORT | 341 | 0.42963663 | 1.311594 | 0 | 0.5343963 | 1 | 3724 | tags=35%, list=14%, signal=40% |
| RIGGI_EWING_SARCOMA_PROGENITOR_UP | 378 | 0.40612867 | 1.3113494 | 0 | 0.36584994 | 1 | 3997 | tags=28%, list=15%, signal=32% |
| FULCHER_INFLAMMATORY_RESPONSE_LECTIN_VS_LPS_DN | 385 | 0.40573472 | 1.3113409 | 0 | 0.3655525 | 1 | 3799 | tags=28%, list=14%, signal=32% |
| HP_LIPID_ACCUMULATION_IN_HEPATOCYTES | 108 | 0.48335505 | 1.3111148 | 0.04 | 0.53467834 | 1 | 2916 | tags=32%, list=11%, signal=36% |
| GO_TUBE_MORPHOGENESIS | 759 | 0.4057666 | 1.3109274 | 0 | 0.5345947 | 1 | 3822 | tags=25%, list=14%, signal=28% |
| GO_FOREBRAIN_DEVELOPMENT | 349 | 0.4091253 | 1.3107815 | 0 | 0.5340531 | 1 | 2371 | tags=17%, list=9%, signal=19% |
| GO_POSITIVE_REGULATION_OF_INTERLEUKIN_8_PRODUCTION | 44 | 0.5420577 | 1.3106483 | 0.04545455 | 0.5341735 | 1 | 2264 | tags=27%, list=8%, signal=30% |
| GO_MUSCLE_STRUCTURE_DEVELOPMENT | 565 | 0.39954522 | 1.3105519 | 0 | 0.5342534 | 1 | 2675 | tags=17%, list=10%, signal=19% |
| HP_SKELETAL_MUSCLE_HYPERTROPHY | 147 | 0.44950232 | 1.3104597 | 0 | 0.5338154 | 1 | 3079 | tags=26%, list=11%, signal=29% |
| YAGI_AML_WITH_T_8_21_TRANSLOCATION | 324 | 0.42682874 | 1.310426 | 0 | 0.36660227 | 1 | 3151 | tags=29%, list=12%, signal=32% |
| BENPORATH_PROLIFERATION | 138 | 0.4752494 | 1.3104043 | 0 | 0.36630565 | 1 | 2842 | tags=34%, list=11%, signal=38% |
| FOSTER_TOLERANT_MACROPHAGE_UP | 155 | 0.4577884 | 1.3101426 | 0.03030303 | 0.36665645 | 1 | 1568 | tags=13%, list=6%, signal=14% |
| GO_BASOLATERAL_PLASMA_MEMBRANE | 210 | 0.4393627 | 1.3094745 | 0.04545455 | 0.53513026 | 1 | 2542 | tags=16%, list=9%, signal=18% |
| BOYLAN_MULTIPLE_MYELOMA_C_D_DN | 246 | 0.4302432 | 1.3091558 | 0.03703704 | 0.3673227 | 1 | 3855 | tags=22%, list=14%, signal=26% |
| GO_TRNA_METABOLIC_PROCESS | 173 | 0.43259627 | 1.3090534 | 0.02777778 | 0.53527755 | 1 | 4137 | tags=42%, list=15%, signal=49% |
| GO_ORGANONITROGEN_COMPOUND_BIOSYNTHETIC_PROCESS | 1556 | 0.37418744 | 1.308462 | 0 | 0.5363203 | 1 | 3633 | tags=28%, list=13%, signal=31% |
| MIKKELSEN_IPS_ICP_WITH_H3K4ME3_AND_H327ME3 | 99 | 0.47898832 | 1.3082348 | 0.025 | 0.36805665 | 1 | 4469 | tags=17%, list=17%, signal=21% |
| GO_REGULATION_OF_GLIOGENESIS | 111 | 0.4719486 | 1.3080335 | 0 | 0.5376511 | 1 | 1003 | tags=13%, list=4%, signal=13% |
| DODD_NASOPHARYNGEAL_CARCINOMA_UP | 1374 | 0.38120684 | 1.3067633 | 0 | 0.3691053 | 1 | 4518 | tags=23%, list=17%, signal=27% |
| HP_ABNORMAL_CONJUGATE_EYE_MOVEMENT | 813 | 0.39367986 | 1.3060348 | 0 | 0.5424075 | 1 | 3526 | tags=27%, list=13%, signal=30% |
| GO_MONOSACCHARIDE_METABOLIC_PROCESS | 248 | 0.4164436 | 1.3059486 | 0 | 0.5423678 | 1 | 2984 | tags=24%, list=11%, signal=27% |
| REACTOME_SEPARATION_OF_SISTER_CHROMATIDS | 177 | 0.4586753 | 1.3058418 | 0.02941177 | 0.36877078 | 1 | 2471 | tags=29%, list=9%, signal=32% |
| GO_SKIN_DEVELOPMENT | 233 | 0.42459658 | 1.3058325 | 0 | 0.5422565 | 1 | 3651 | tags=18%, list=14%, signal=21% |
| HP_CHOREOATHETOSIS | 79 | 0.49005154 | 1.3056142 | 0.03225806 | 0.5425681 | 1 | 1795 | tags=23%, list=7%, signal=24% |
| HP_PROGRESSIVE_CEREBELLAR_ATAXIA | 79 | 0.51921135 | 1.305085 | 0.03225806 | 0.54229707 | 1 | 2158 | tags=28%, list=8%, signal=30% |
| BLALOCK_ALZHEIMERS_DISEASE_INCIPIENT_UP | 345 | 0.4208816 | 1.3050683 | 0 | 0.3688447 | 1 | 3405 | tags=30%, list=13%, signal=34% |
| GO_CELL_ACTIVATION | 1191 | 0.39956492 | 1.3050096 | 0 | 0.5420008 | 1 | 2693 | tags=18%, list=10%, signal=19% |
| HP_COMPENSATED_HYPOTHYROIDISM | 3 | 0.94171906 | 1.3049077 | 0.04255319 | 0.5418423 | 1 | 161 | tags=33%, list=1%, signal=34% |
| TAKEDA_TARGETS_OF_NUP98_HOXA9_FUSION_16D_UP | 138 | 0.46009788 | 1.3044461 | 0 | 0.36992258 | 1 | 2141 | tags=16%, list=8%, signal=17% |
| GO_PROTEIN_CONTAINING_COMPLEX_BINDING | 1105 | 0.3887979 | 1.3044438 | 0 | 0.5424607 | 1 | 3293 | tags=23%, list=12%, signal=25% |
| TARTE_PLASMA_CELL_VS_PLASMABLAST_UP | 307 | 0.4330516 | 1.3044164 | 0 | 0.36971778 | 1 | 2164 | tags=20%, list=8%, signal=21% |
| YAGI_AML_WITH_11Q23_REARRANGED | 307 | 0.41402033 | 1.304402 | 0 | 0.36941227 | 1 | 3556 | tags=29%, list=13%, signal=33% |
| MARTINEZ_RB1_AND_TP53_TARGETS_DN | 538 | 0.39991564 | 1.3043072 | 0 | 0.36906525 | 1 | 3413 | tags=31%, list=13%, signal=34% |
| GO_SENSORY_ORGAN_DEVELOPMENT | 511 | 0.3968718 | 1.3041486 | 0 | 0.54248077 | 1 | 2338 | tags=14%, list=9%, signal=15% |
| GO_RESPONSE_TO_ESTRADIOL | 126 | 0.45172358 | 1.3041302 | 0.03333334 | 0.54227304 | 1 | 2530 | tags=16%, list=9%, signal=17% |
| GO_CELL_CELL_SIGNALING | 1538 | 0.3827969 | 1.3039956 | 0 | 0.5425553 | 1 | 2804 | tags=17%, list=10%, signal=17% |
| GO_VACUOLAR_TRANSPORT | 140 | 0.4534442 | 1.3038813 | 0.03225806 | 0.54247 | 1 | 4245 | tags=42%, list=16%, signal=50% |
| GO_LIPOSACCHARIDE_METABOLIC_PROCESS | 101 | 0.50753886 | 1.3036789 | 0 | 0.5422103 | 1 | 4723 | tags=44%, list=18%, signal=53% |
| BANDRES_RESPONSE_TO_CARMUSTIN_MGMT_48HR_DN | 134 | 0.44307175 | 1.3035696 | 0 | 0.36963126 | 1 | 1718 | tags=20%, list=6%, signal=21% |
| GO_CELL_LEADING_EDGE | 402 | 0.40821978 | 1.3034463 | 0 | 0.54293984 | 1 | 3387 | tags=24%, list=13%, signal=27% |
| HP_ROD_CONE_DYSTROPHY | 151 | 0.4527386 | 1.3034452 | 0.03571429 | 0.54266894 | 1 | 3817 | tags=26%, list=14%, signal=30% |
| LEE_NEURAL_CREST_STEM_CELL_DN | 109 | 0.4677652 | 1.30305 | 0.02777778 | 0.36966905 | 1 | 2061 | tags=17%, list=8%, signal=18% |
| SENESE_HDAC3_TARGETS_DN | 481 | 0.39419064 | 1.3025402 | 0 | 0.3697033 | 1 | 3435 | tags=29%, list=13%, signal=33% |
| MCLACHLAN_DENTAL_CARIES_UP | 215 | 0.44574612 | 1.30244 | 0 | 0.36959022 | 1 | 2910 | tags=20%, list=11%, signal=22% |
| WP_NEURAL_CREST_DIFFERENTIATION | 94 | 0.47143814 | 1.3024138 | 0.03225806 | 0.36871576 | 1 | 2459 | tags=19%, list=9%, signal=21% |
| SHEN_SMARCA2_TARGETS_UP | 409 | 0.41559985 | 1.3023815 | 0 | 0.36848807 | 1 | 3774 | tags=39%, list=14%, signal=45% |
| LINDGREN_BLADDER_CANCER_CLUSTER_1_DN | 349 | 0.40255192 | 1.302042 | 0 | 0.3690805 | 1 | 3625 | tags=30%, list=13%, signal=34% |
| HP_CEREBRAL_CORTICAL_ATROPHY | 251 | 0.43231592 | 1.3019062 | 0 | 0.5452822 | 1 | 2398 | tags=19%, list=9%, signal=21% |
| GO_RESPONSE_TO_HORMONE | 782 | 0.39435822 | 1.3018974 | 0 | 0.54507506 | 1 | 2535 | tags=20%, list=9%, signal=21% |
| HP_ABNORMALITY_OF_THE_PARANASAL_SINUSES | 113 | 0.48409143 | 1.3018597 | 0.03333334 | 0.5449562 | 1 | 3514 | tags=19%, list=13%, signal=22% |
| HP_CEREBELLAR_ATROPHY | 293 | 0.42362112 | 1.3015404 | 0 | 0.5450575 | 1 | 3447 | tags=27%, list=13%, signal=30% |
| GO_GLAND_MORPHOGENESIS | 110 | 0.48236498 | 1.3015232 | 0.03448276 | 0.5448593 | 1 | 3283 | tags=25%, list=12%, signal=29% |
| GARY_CD5_TARGETS_DN | 410 | 0.41601202 | 1.301361 | 0 | 0.3698784 | 1 | 3160 | tags=33%, list=12%, signal=37% |
| HP_HYPOGONADOTROPIC_HYPOGONADISM | 114 | 0.47621295 | 1.3011057 | 0 | 0.5457833 | 1 | 2068 | tags=16%, list=8%, signal=17% |
| GO_IDENTICAL_PROTEIN_BINDING | 1659 | 0.39242655 | 1.3002357 | 0 | 0.5468177 | 1 | 3720 | tags=25%, list=14%, signal=28% |
| GO_CYTOKINE_PRODUCTION | 644 | 0.39885968 | 1.3001069 | 0 | 0.5465161 | 1 | 2530 | tags=17%, list=9%, signal=19% |
| GO_CELL_CELL_ADHESION | 752 | 0.39039603 | 1.3000748 | 0 | 0.5463578 | 1 | 4087 | tags=25%, list=15%, signal=28% |
| HP_ABNORMAL_OVARIAN_MORPHOLOGY | 112 | 0.47373405 | 1.299851 | 0.02702703 | 0.54633975 | 1 | 1601 | tags=14%, list=6%, signal=15% |
| GO_RESPONSE_TO_NITROGEN_COMPOUND | 976 | 0.38828942 | 1.2996721 | 0 | 0.54628193 | 1 | 2535 | tags=18%, list=9%, signal=19% |
| GO_SECRETORY_GRANULE_MEMBRANE | 266 | 0.42231256 | 1.2996193 | 0 | 0.5456843 | 1 | 2445 | tags=18%, list=9%, signal=20% |
| KEGG_PYRIMIDINE_METABOLISM | 90 | 0.47574493 | 1.2994336 | 0 | 0.3724887 | 1 | 4159 | tags=42%, list=15%, signal=50% |
| ONDER_CDH1_TARGETS_2_DN | 394 | 0.40456823 | 1.2990927 | 0 | 0.37290335 | 1 | 4100 | tags=23%, list=15%, signal=26% |
| SCHLINGEMANN_SKIN_CARCINOGENESIS_TPA_DN | 22 | 0.6373798 | 1.2987956 | 0.04545455 | 0.37293068 | 1 | 2054 | tags=32%, list=8%, signal=34% |
| HELLER_HDAC_TARGETS_SILENCED_BY_METHYLATION_UP | 382 | 0.40915576 | 1.2987372 | 0 | 0.37271553 | 1 | 3298 | tags=25%, list=12%, signal=28% |
| HP_EPISODIC_TACHYPNEA | 28 | 0.5665953 | 1.2983605 | 0.04545455 | 0.54839844 | 1 | 4058 | tags=54%, list=15%, signal=63% |
| GO_LIPID_TRANSPORT_ACROSS_BLOOD_BRAIN_BARRIER | 5 | 0.91197866 | 1.2982944 | 0.04545455 | 0.5483651 | 1 | 669 | tags=40%, list=2%, signal=41% |
| MEISSNER_BRAIN_HCP_WITH_H3K4ME3_AND_H3K27ME3 | 978 | 0.38212034 | 1.2982855 | 0 | 0.37278485 | 1 | 3158 | tags=15%, list=12%, signal=17% |
| LOCKWOOD_AMPLIFIED_IN_LUNG_CANCER | 199 | 0.43121842 | 1.2980745 | 0.03846154 | 0.37296847 | 1 | 3089 | tags=30%, list=11%, signal=33% |
| ACEVEDO_LIVER_CANCER_DN | 455 | 0.40404195 | 1.2979443 | 0 | 0.37295315 | 1 | 2965 | tags=24%, list=11%, signal=26% |
| REACTOME_COPI_DEPENDENT_GOLGI_TO_ER_RETROGRADE_TRAFFIC | 89 | 0.43870217 | 1.297199 | 0.03125 | 0.3736859 | 1 | 2992 | tags=29%, list=11%, signal=33% |
| GO_REGULATION_OF_GLIAL_CELL_PROLIFERATION | 28 | 0.5590798 | 1.2970481 | 0.04761905 | 0.5508935 | 1 | 735 | tags=14%, list=3%, signal=15% |
| REACTOME_ADAPTIVE_IMMUNE_SYSTEM | 651 | 0.39912337 | 1.29685 | 0 | 0.3733417 | 1 | 2869 | tags=22%, list=11%, signal=24% |
| GO_HEXOSE_CATABOLIC_PROCESS | 49 | 0.5299816 | 1.296654 | 0.04761905 | 0.55163443 | 1 | 1962 | tags=24%, list=7%, signal=26% |
| GO_MYELOID_CELL_DIFFERENTIATION | 354 | 0.40893608 | 1.2962576 | 0.03030303 | 0.55222076 | 1 | 3013 | tags=25%, list=11%, signal=28% |
| CERVERA_SDHB_TARGETS_1_UP | 99 | 0.4714859 | 1.296057 | 0.02564103 | 0.37389544 | 1 | 1935 | tags=18%, list=7%, signal=20% |
| HP_SUPERNUMERARY_BONES_OF_THE_AXIAL_SKELETON | 22 | 0.5849867 | 1.2960484 | 0.05 | 0.5524452 | 1 | 4295 | tags=45%, list=16%, signal=54% |
| GO_REGULATION_OF_ACTIN_FILAMENT_BUNDLE_ASSEMBLY | 95 | 0.45011553 | 1.2955501 | 0 | 0.5527958 | 1 | 2616 | tags=24%, list=10%, signal=27% |
| SCHAEFFER_PROSTATE_DEVELOPMENT_6HR_UP | 143 | 0.44875646 | 1.2955478 | 0 | 0.37404594 | 1 | 3491 | tags=29%, list=13%, signal=33% |
| GO_MAINTENANCE_OF_CELL_NUMBER | 151 | 0.46161458 | 1.294551 | 0 | 0.55427396 | 1 | 3641 | tags=34%, list=14%, signal=40% |
| HP_ABNORMALITY_OF_ESOPHAGUS_PHYSIOLOGY | 505 | 0.40948185 | 1.294532 | 0 | 0.55406123 | 1 | 3265 | tags=24%, list=12%, signal=27% |
| GO_NUCLEOSIDE_BISPHOSPHATE_METABOLIC_PROCESS | 115 | 0.45666546 | 1.2944417 | 0.02702703 | 0.5539526 | 1 | 3557 | tags=32%, list=13%, signal=37% |
| GO_SENSORY_SYSTEM_DEVELOPMENT | 349 | 0.41459823 | 1.2942371 | 0 | 0.5539828 | 1 | 2315 | tags=14%, list=9%, signal=15% |
| HAMAI_APOPTOSIS_VIA_TRAIL_DN | 167 | 0.4305849 | 1.2934985 | 0 | 0.37786773 | 1 | 2598 | tags=14%, list=10%, signal=16% |
| GO_ENDOTHELIAL_CELL_PROLIFERATION | 131 | 0.4605618 | 1.2926537 | 0 | 0.5570724 | 1 | 2870 | tags=23%, list=11%, signal=26% |
| GO_ACTIN_FILAMENT_BINDING | 183 | 0.44673893 | 1.292639 | 0 | 0.55687565 | 1 | 3435 | tags=25%, list=13%, signal=29% |
| ST_WNT_CA2_CYCLIC_GMP_PATHWAY | 18 | 0.645907 | 1.2926216 | 0.05 | 0.37924522 | 1 | 2948 | tags=39%, list=11%, signal=44% |
| REACTOME_MUSCLE_CONTRACTION | 171 | 0.45193902 | 1.2923725 | 0.03030303 | 0.37872484 | 1 | 3437 | tags=22%, list=13%, signal=25% |
| GO_SULFUR_COMPOUND_BINDING | 216 | 0.4343624 | 1.2922723 | 0 | 0.5576514 | 1 | 3561 | tags=24%, list=13%, signal=27% |
| GO_NEGATIVE_REGULATION_OF_CATALYTIC_ACTIVITY | 651 | 0.39594275 | 1.292233 | 0 | 0.5575383 | 1 | 3140 | tags=22%, list=12%, signal=25% |
| CASORELLI_ACUTE_PROMYELOCYTIC_LEUKEMIA_DN | 620 | 0.39160267 | 1.2909392 | 0 | 0.38050663 | 1 | 3103 | tags=31%, list=12%, signal=35% |
| GO_SULFUR_COMPOUND_METABOLIC_PROCESS | 316 | 0.4348069 | 1.2904619 | 0.04 | 0.56170917 | 1 | 3194 | tags=27%, list=12%, signal=30% |
| WIERENGA_STAT5A_TARGETS_UP | 183 | 0.4171484 | 1.2902181 | 0.04545455 | 0.38112494 | 1 | 3383 | tags=23%, list=13%, signal=26% |
| HP_DERMATOLOGICAL_MANIFESTATIONS_OF_SYSTEMIC_DISORDERS | 201 | 0.45059323 | 1.2901788 | 0.02777778 | 0.56172913 | 1 | 3524 | tags=25%, list=13%, signal=29% |
| GO_STRESS_FIBER_ASSEMBLY | 98 | 0.45806316 | 1.2899405 | 0.03030303 | 0.5623265 | 1 | 2616 | tags=27%, list=10%, signal=29% |
| GO_HIPPOCAMPUS_DEVELOPMENT | 74 | 0.48699698 | 1.2896641 | 0.02857143 | 0.5625349 | 1 | 3736 | tags=34%, list=14%, signal=39% |
| HP_FOCAL_ONSET_SEIZURE | 175 | 0.43732336 | 1.2896166 | 0.03225806 | 0.5624887 | 1 | 2080 | tags=15%, list=8%, signal=16% |
| YAGI_AML_WITH_INV_16_TRANSLOCATION | 358 | 0.40338933 | 1.2892822 | 0 | 0.3818352 | 1 | 2817 | tags=26%, list=10%, signal=28% |
| GO_ACTIN_FILAMENT_BASED_PROCESS | 712 | 0.39694747 | 1.28903 | 0 | 0.56407297 | 1 | 3413 | tags=23%, list=13%, signal=26% |
| GO_REGULATION_OF_PHOSPHATIDYLINOSITOL_3_KINASE_SIGNALING | 105 | 0.47601262 | 1.2883794 | 0.03571429 | 0.5649692 | 1 | 2952 | tags=29%, list=11%, signal=32% |
| GO_DETECTION_OF_STIMULUS | 215 | 0.42969635 | 1.2883626 | 0.03571429 | 0.5644761 | 1 | 2601 | tags=15%, list=10%, signal=16% |
| HP_ABNORMALITY_OF_THE_MUSCULATURE_OF_THE_LIMBS | 381 | 0.39576533 | 1.288312 | 0 | 0.564179 | 1 | 4044 | tags=30%, list=15%, signal=34% |
| MANNE_COVID19_NONICU_VS_HEALTHY_DONOR_PLATELETS_DN | 172 | 0.42047858 | 1.2882458 | 0 | 0.38331383 | 1 | 3782 | tags=22%, list=14%, signal=25% |
| GO_DETECTION_OF_ABIOTIC_STIMULUS | 111 | 0.4722191 | 1.288094 | 0.03703704 | 0.56417567 | 1 | 1894 | tags=11%, list=7%, signal=12% |
| GO_PROTEOLYSIS | 1470 | 0.38851318 | 1.2879521 | 0 | 0.5639152 | 1 | 2896 | tags=21%, list=11%, signal=22% |
| MANNE_COVID19_COMBINED_COHORT_VS_HEALTHY_DONOR_PLATELETS_DN | 181 | 0.4277145 | 1.287379 | 0.03125 | 0.3847898 | 1 | 3868 | tags=24%, list=14%, signal=28% |
| HP_DECREASED_HEAD_CIRCUMFERENCE | 979 | 0.3870862 | 1.2873784 | 0 | 0.56475085 | 1 | 3535 | tags=26%, list=13%, signal=29% |
| IVANOVA_HEMATOPOIESIS_STEM_CELL | 168 | 0.4321539 | 1.2870188 | 0 | 0.38545087 | 1 | 2789 | tags=29%, list=10%, signal=32% |
| GO_CELLULAR_PROTEIN_CATABOLIC_PROCESS | 721 | 0.38689727 | 1.2870138 | 0 | 0.5654091 | 1 | 2896 | tags=25%, list=11%, signal=27% |
| BOQUEST_STEM_CELL_CULTURED_VS_FRESH_UP | 391 | 0.4008107 | 1.2865862 | 0 | 0.38537633 | 1 | 2459 | tags=20%, list=9%, signal=22% |
| HP_PACE_OF_PROGRESSION | 411 | 0.40727165 | 1.2865318 | 0 | 0.56604946 | 1 | 3839 | tags=28%, list=14%, signal=32% |
| BIOCARTA_PEPI_PATHWAY | 3 | 0.9452916 | 1.2865103 | 0.04761905 | 0.38487777 | 1 | 300 | tags=33%, list=1%, signal=34% |
| GO_REGULATION_OF_INFLAMMATORY_RESPONSE | 303 | 0.4189697 | 1.2864991 | 0 | 0.5657087 | 1 | 2947 | tags=20%, list=11%, signal=22% |
| GO_MEMBRANE_FUSION | 140 | 0.44695407 | 1.2863343 | 0.03333334 | 0.56550825 | 1 | 2535 | tags=25%, list=9%, signal=27% |
| GO_NEGATIVE_REGULATION_OF_CELLULAR_PROTEIN_LOCALIZATION | 106 | 0.4656751 | 1.2861019 | 0.03703704 | 0.56506026 | 1 | 3075 | tags=29%, list=11%, signal=33% |
| GO_ORGANIC_ACID_BIOSYNTHETIC_PROCESS | 308 | 0.43237108 | 1.2857913 | 0.04347826 | 0.5652459 | 1 | 3108 | tags=21%, list=12%, signal=24% |
| GO_CELL_SURFACE_RECEPTOR_SIGNALING_PATHWAY_INVOLVED_IN_CELL_CELL_SIGNALING | 563 | 0.4049188 | 1.2855963 | 0 | 0.56456995 | 1 | 2754 | tags=20%, list=10%, signal=22% |
| GO_METAL_ION_TRANSPORT | 758 | 0.39443868 | 1.2852864 | 0 | 0.5646143 | 1 | 3083 | tags=16%, list=11%, signal=17% |
| REACTOME_TELOMERE_MAINTENANCE | 64 | 0.47144628 | 1.2843437 | 0.03333334 | 0.3873385 | 1 | 3325 | tags=41%, list=12%, signal=46% |
| HP_ABNORMALITY_OF_BONE_MINERAL_DENSITY | 438 | 0.40024015 | 1.2841501 | 0 | 0.5663983 | 1 | 3199 | tags=23%, list=12%, signal=26% |
| GO_REGULATION_OF_CELL_DIFFERENTIATION | 1631 | 0.36977395 | 1.2839876 | 0 | 0.5667403 | 1 | 2678 | tags=18%, list=10%, signal=18% |
| GO_RESPONSE_TO_LEUKEMIA_INHIBITORY_FACTOR | 93 | 0.47670832 | 1.2834265 | 0 | 0.5674084 | 1 | 3287 | tags=33%, list=12%, signal=38% |
| GO_REGULATION_OF_NIK_NF_KAPPAB_SIGNALING | 98 | 0.4695971 | 1.2832497 | 0.02702703 | 0.56753474 | 1 | 3182 | tags=29%, list=12%, signal=32% |
| GO_GLYCOPROTEIN_BIOSYNTHETIC_PROCESS | 286 | 0.41396916 | 1.2832476 | 0 | 0.5670385 | 1 | 3547 | tags=23%, list=13%, signal=27% |
| GO_BIOLOGICAL_ADHESION | 1250 | 0.37512225 | 1.2831196 | 0 | 0.5665675 | 1 | 3606 | tags=21%, list=13%, signal=23% |
| KIM_MYC_AMPLIFICATION_TARGETS_UP | 180 | 0.4353764 | 1.2831068 | 0.03333334 | 0.38926965 | 1 | 2948 | tags=33%, list=11%, signal=37% |
| GO_RECYCLING_ENDOSOME | 168 | 0.43726766 | 1.2829152 | 0 | 0.56655186 | 1 | 2566 | tags=21%, list=10%, signal=24% |
| GO_POSITIVE_REGULATION_OF_SYNAPTIC_TRANSMISSION | 159 | 0.4270203 | 1.2825482 | 0 | 0.5670606 | 1 | 927 | tags=11%, list=3%, signal=11% |
| REACTOME_G_ALPHA_I_SIGNALLING_EVENTS | 301 | 0.41615647 | 1.2818509 | 0.05 | 0.391395 | 1 | 3208 | tags=18%, list=12%, signal=20% |
| GO_RNA_MODIFICATION | 143 | 0.44141817 | 1.2817755 | 0.03703704 | 0.56858826 | 1 | 4153 | tags=41%, list=15%, signal=48% |
| GO_ANIMAL_ORGAN_MORPHOGENESIS | 927 | 0.38424197 | 1.2812788 | 0 | 0.569659 | 1 | 4151 | tags=24%, list=15%, signal=27% |
| GO_RESPONSE_TO_WOUNDING | 531 | 0.4144391 | 1.2809759 | 0 | 0.56981385 | 1 | 2446 | tags=18%, list=9%, signal=19% |
| CREIGHTON_ENDOCRINE_THERAPY_RESISTANCE_1 | 479 | 0.39283568 | 1.2803315 | 0 | 0.39436758 | 1 | 3703 | tags=34%, list=14%, signal=39% |
| GRAESSMANN_RESPONSE_TO_MC_AND_DOXORUBICIN_UP | 572 | 0.4008806 | 1.279318 | 0 | 0.39637104 | 1 | 3633 | tags=26%, list=13%, signal=30% |
| GO_PEPTIDYL_TYROSINE_MODIFICATION | 325 | 0.40674502 | 1.2792077 | 0.03225806 | 0.57135886 | 1 | 2846 | tags=18%, list=11%, signal=20% |
| GO_CADHERIN_BINDING | 301 | 0.41577116 | 1.2792072 | 0 | 0.57110274 | 1 | 3117 | tags=34%, list=12%, signal=38% |
| GO_REGULATION_OF_TRANSPORT | 1575 | 0.36934212 | 1.2788229 | 0 | 0.5722911 | 1 | 2747 | tags=18%, list=10%, signal=18% |
| REACTOME_SIGNALING_BY_INTERLEUKINS | 397 | 0.40963835 | 1.2783879 | 0 | 0.3985726 | 1 | 2621 | tags=20%, list=10%, signal=21% |
| REACTOME_CYTOKINE_SIGNALING_IN_IMMUNE_SYSTEM | 757 | 0.38822517 | 1.2779545 | 0 | 0.39891818 | 1 | 2621 | tags=19%, list=10%, signal=21% |
| BENPORATH_NANOG_TARGETS | 905 | 0.38248938 | 1.2778774 | 0 | 0.39873677 | 1 | 2718 | tags=24%, list=10%, signal=25% |
| REACTOME_ER_TO_GOLGI_ANTEROGRADE_TRANSPORT | 140 | 0.456484 | 1.2777822 | 0.03125 | 0.39863443 | 1 | 2819 | tags=30%, list=10%, signal=33% |
| GO_AGING | 265 | 0.4323685 | 1.2775348 | 0 | 0.574319 | 1 | 2158 | tags=16%, list=8%, signal=17% |
| GO_LIPID_MODIFICATION | 236 | 0.4168617 | 1.277527 | 0.04347826 | 0.57409143 | 1 | 3798 | tags=30%, list=14%, signal=34% |
| GO_REGULATION_OF_NEURAL_PRECURSOR_CELL_PROLIFERATION | 79 | 0.4847839 | 1.2773726 | 0.03448276 | 0.5740798 | 1 | 2148 | tags=22%, list=8%, signal=23% |
| GO_POSITIVE_REGULATION_OF_CELL_POPULATION_PROLIFERATION | 762 | 0.38830334 | 1.277275 | 0 | 0.57311094 | 1 | 2870 | tags=19%, list=11%, signal=20% |
| WAMUNYOKOLI_OVARIAN_CANCER_LMP_DN | 184 | 0.41675913 | 1.2771041 | 0 | 0.3997263 | 1 | 2512 | tags=22%, list=9%, signal=24% |
| HP_HYPERPITUITARISM | 81 | 0.5146488 | 1.2768736 | 0.02941177 | 0.57310754 | 1 | 3128 | tags=22%, list=12%, signal=25% |
| MCBRYAN_PUBERTAL_BREAST_6_7WK_UP | 183 | 0.42937586 | 1.2767965 | 0 | 0.39962295 | 1 | 3085 | tags=31%, list=11%, signal=35% |
| GO_EARLY_ENDOSOME | 332 | 0.41268182 | 1.2762544 | 0 | 0.5735499 | 1 | 2582 | tags=22%, list=10%, signal=24% |
| GO_RIBOSE_PHOSPHATE_METABOLIC_PROCESS | 378 | 0.41102406 | 1.2761201 | 0 | 0.573581 | 1 | 3223 | tags=26%, list=12%, signal=30% |
| GO_REGULATION_OF_LIPID_METABOLIC_PROCESS | 368 | 0.4004183 | 1.2759923 | 0 | 0.5735203 | 1 | 2729 | tags=22%, list=10%, signal=24% |
| HP_OSTEOARTHRITIS | 52 | 0.4979701 | 1.2756697 | 0.02941177 | 0.57407296 | 1 | 1927 | tags=19%, list=7%, signal=21% |
| HP_ABNORMAL_RENAL_MORPHOLOGY | 771 | 0.37657532 | 1.2756385 | 0 | 0.57395405 | 1 | 3524 | tags=22%, list=13%, signal=25% |
| GO_TRANSPORT_VESICLE_MEMBRANE | 174 | 0.44020358 | 1.2752515 | 0 | 0.5738129 | 1 | 2417 | tags=24%, list=9%, signal=26% |
| HP_ABNORMALITY_OF_THE_LYMPHATIC_SYSTEM | 524 | 0.40447864 | 1.2752273 | 0 | 0.57363886 | 1 | 3468 | tags=23%, list=13%, signal=26% |
| GO_CELL_CHEMOTAXIS | 243 | 0.41945037 | 1.2747648 | 0 | 0.57365453 | 1 | 2555 | tags=19%, list=9%, signal=20% |
| GO_NEGATIVE_REGULATION_OF_CELL_ADHESION | 244 | 0.4108903 | 1.273905 | 0 | 0.57273835 | 1 | 2281 | tags=16%, list=8%, signal=17% |
| MULLIGHAN_NPM1_MUTATED_SIGNATURE_1_UP | 228 | 0.41259828 | 1.273374 | 0 | 0.40529418 | 1 | 3970 | tags=32%, list=15%, signal=37% |
| GO_PROCESS_UTILIZING_AUTOPHAGIC_MECHANISM | 497 | 0.3864417 | 1.2725405 | 0 | 0.5729846 | 1 | 2527 | tags=22%, list=9%, signal=24% |
| DARWICHE_SQUAMOUS_CELL_CARCINOMA_DN | 156 | 0.4495812 | 1.2722049 | 0.04545455 | 0.40689075 | 1 | 3675 | tags=33%, list=14%, signal=38% |
| GO_ION_TRANSMEMBRANE_TRANSPORTER_ACTIVITY | 726 | 0.38991868 | 1.2720618 | 0 | 0.57370913 | 1 | 3471 | tags=16%, list=13%, signal=18% |
| HP_ABNORMALITY_OF_JOINT_MOBILITY | 931 | 0.3771875 | 1.2715507 | 0 | 0.5744392 | 1 | 4136 | tags=28%, list=15%, signal=32% |
| FORTSCHEGGER_PHF8_TARGETS_DN | 705 | 0.3963994 | 1.2710433 | 0 | 0.4074087 | 1 | 4024 | tags=30%, list=15%, signal=34% |
| GAUSSMANN_MLL_AF4_FUSION_TARGETS_G_UP | 214 | 0.433456 | 1.2709749 | 0.02857143 | 0.40726236 | 1 | 3210 | tags=24%, list=12%, signal=27% |
| HP_ABNORMAL_INTESTINE_MORPHOLOGY | 582 | 0.3963237 | 1.2698965 | 0 | 0.5765599 | 1 | 2876 | tags=20%, list=11%, signal=21% |
| HP_ABNORMALITY_OF_THE_SPLEEN | 428 | 0.4114499 | 1.2694842 | 0 | 0.5762871 | 1 | 3468 | tags=23%, list=13%, signal=26% |
| GO_REGULATION_OF_CELL_CYCLE_PHASE_TRANSITION | 413 | 0.39723468 | 1.2693119 | 0 | 0.57641685 | 1 | 3146 | tags=27%, list=12%, signal=30% |
| GO_CIRCULATORY_SYSTEM_PROCESS | 465 | 0.3992926 | 1.2691479 | 0 | 0.5768381 | 1 | 3438 | tags=18%, list=13%, signal=20% |
| GOLDRATH_ANTIGEN_RESPONSE | 332 | 0.40098473 | 1.2686625 | 0 | 0.40767533 | 1 | 2598 | tags=26%, list=10%, signal=29% |
| GO_REGULATION_OF_PROTEIN_STABILITY | 267 | 0.40360856 | 1.2676951 | 0 | 0.5789574 | 1 | 2973 | tags=26%, list=11%, signal=29% |
| BERTUCCI_MEDULLARY_VS_DUCTAL_BREAST_CANCER_DN | 154 | 0.4437502 | 1.267188 | 0.03571429 | 0.4099174 | 1 | 4457 | tags=35%, list=17%, signal=42% |
| DARWICHE_PAPILLOMA_RISK_HIGH_DN | 161 | 0.4252985 | 1.2669072 | 0 | 0.41024882 | 1 | 3675 | tags=32%, list=14%, signal=37% |
| HP_X_LINKED_RECESSIVE_INHERITANCE | 157 | 0.44185546 | 1.2667118 | 0 | 0.58029795 | 1 | 2815 | tags=22%, list=10%, signal=25% |
| DARWICHE_PAPILLOMA_RISK_LOW_UP | 137 | 0.4309007 | 1.2662033 | 0 | 0.41084892 | 1 | 3269 | tags=34%, list=12%, signal=38% |
| WIERENGA_STAT5A_TARGETS_GROUP1 | 116 | 0.47275573 | 1.2660943 | 0 | 0.4107933 | 1 | 3364 | tags=22%, list=12%, signal=25% |
| GO_INFLAMMATORY_RESPONSE | 599 | 0.3955949 | 1.2660285 | 0 | 0.5812438 | 1 | 2947 | tags=17%, list=11%, signal=18% |
| GO_PROTEASOMAL_PROTEIN_CATABOLIC_PROCESS | 450 | 0.39101475 | 1.2654938 | 0 | 0.58260685 | 1 | 2896 | tags=26%, list=11%, signal=28% |
| HP_INTRAUTERINE_GROWTH_RETARDATION | 426 | 0.39559352 | 1.2648891 | 0 | 0.58303326 | 1 | 2972 | tags=22%, list=11%, signal=24% |
| PARENT_MTOR_SIGNALING_UP | 516 | 0.40142903 | 1.2648833 | 0 | 0.41204053 | 1 | 3900 | tags=32%, list=14%, signal=36% |
| GO_CELLULAR_RESPONSE_TO_BIOTIC_STIMULUS | 195 | 0.44375175 | 1.2643604 | 0 | 0.58381534 | 1 | 2604 | tags=19%, list=10%, signal=21% |
| GO_CARBOHYDRATE_METABOLIC_PROCESS | 531 | 0.39966154 | 1.264356 | 0 | 0.58357865 | 1 | 3141 | tags=23%, list=12%, signal=25% |
| GO_REGULATION_OF_WOUND_HEALING | 118 | 0.43673432 | 1.2643037 | 0 | 0.58350873 | 1 | 2205 | tags=16%, list=8%, signal=17% |
| GO_REGULATION_OF_BRANCHING_INVOLVED_IN_LUNG_MORPHOGENESIS | 4 | 0.9034984 | 1.2638464 | 0.04651163 | 0.5844472 | 1 | 267 | tags=50%, list=1%, signal=50% |
| GO_T_CELL_ACTIVATION | 397 | 0.39231426 | 1.263734 | 0 | 0.58442044 | 1 | 2604 | tags=17%, list=10%, signal=18% |
| GO_CELLULAR_AMIDE_METABOLIC_PROCESS | 984 | 0.37315607 | 1.2635168 | 0 | 0.5847201 | 1 | 2801 | tags=27%, list=10%, signal=29% |
| GO_POSITIVE_REGULATION_OF_PROTEIN_KINASE_B_SIGNALING | 145 | 0.44552985 | 1.263424 | 0.02857143 | 0.5845537 | 1 | 2357 | tags=17%, list=9%, signal=19% |
| GO_REGULATION_OF_VIRAL_TRANSCRIPTION | 63 | 0.51020956 | 1.2633731 | 0.02777778 | 0.584431 | 1 | 2782 | tags=48%, list=10%, signal=53% |
| ZHANG_BREAST_CANCER_PROGENITORS_UP | 413 | 0.39658874 | 1.263372 | 0 | 0.41445783 | 1 | 2452 | tags=25%, list=9%, signal=27% |
| GO_CELL_CELL_JUNCTION_ORGANIZATION | 179 | 0.4286024 | 1.2631814 | 0.04166667 | 0.5847927 | 1 | 3638 | tags=24%, list=14%, signal=28% |
| HP_ABNORMAL_ATRIOVENTRICULAR_VALVE_PHYSIOLOGY | 126 | 0.44504794 | 1.2630212 | 0.03125 | 0.5849115 | 1 | 2928 | tags=27%, list=11%, signal=30% |
| GO_NEGATIVE_REGULATION_OF_TRANSPORT | 385 | 0.39996585 | 1.2630053 | 0 | 0.58447355 | 1 | 2847 | tags=21%, list=11%, signal=23% |
| GO_RESPONSE_TO_ENDOGENOUS_STIMULUS | 1412 | 0.36472622 | 1.2626083 | 0 | 0.58513427 | 1 | 2729 | tags=19%, list=10%, signal=20% |
| GO_REGULATION_OF_INTRINSIC_APOPTOTIC_SIGNALING_PATHWAY | 146 | 0.4371302 | 1.2625475 | 0 | 0.5848692 | 1 | 2434 | tags=29%, list=9%, signal=31% |
| GO_MACROAUTOPHAGY | 292 | 0.39887056 | 1.2624711 | 0 | 0.58470374 | 1 | 2407 | tags=23%, list=9%, signal=25% |
| GO_PEPTIDE_SECRETION | 439 | 0.3968787 | 1.2622614 | 0 | 0.5853093 | 1 | 2458 | tags=17%, list=9%, signal=18% |
| GO_REGULATION_OF_CELLULAR_CATABOLIC_PROCESS | 803 | 0.38450664 | 1.2621868 | 0 | 0.58486307 | 1 | 2925 | tags=24%, list=11%, signal=27% |
| MANNE_COVID19_ICU_VS_HEALTHY_DONOR_PLATELETS_DN | 125 | 0.44319814 | 1.2613319 | 0 | 0.4169866 | 1 | 3868 | tags=26%, list=14%, signal=31% |
| GO_REGULATION_OF_VASCULATURE_DEVELOPMENT | 297 | 0.41416976 | 1.2612268 | 0 | 0.58500403 | 1 | 3430 | tags=26%, list=13%, signal=29% |
| HP_NEOPLASM_OF_THE_SKIN | 160 | 0.43317258 | 1.2609607 | 0.03030303 | 0.5854853 | 1 | 3484 | tags=24%, list=13%, signal=28% |
| DANG_BOUND_BY_MYC | 971 | 0.3838055 | 1.260221 | 0 | 0.4185387 | 1 | 3473 | tags=32%, list=13%, signal=35% |
| JIANG_HYPOXIA_NORMAL | 192 | 0.4337458 | 1.2601472 | 0.03030303 | 0.41837928 | 1 | 2924 | tags=31%, list=11%, signal=35% |
| GO_POSITIVE_REGULATION_OF_CELL_ADHESION | 374 | 0.4041223 | 1.2600844 | 0 | 0.58509076 | 1 | 3120 | tags=21%, list=12%, signal=23% |
| GO_APICAL_PLASMA_MEMBRANE | 296 | 0.39360172 | 1.260023 | 0 | 0.5850934 | 1 | 2743 | tags=14%, list=10%, signal=16% |
| BRUINS_UVC_RESPONSE_VIA_TP53_GROUP_D | 235 | 0.41476312 | 1.2599955 | 0.03448276 | 0.4180829 | 1 | 2876 | tags=19%, list=11%, signal=21% |
| HP_GAIT_ATAXIA | 161 | 0.438418 | 1.2597054 | 0.03125 | 0.5853011 | 1 | 2681 | tags=20%, list=10%, signal=22% |
| GO_POSITIVE_REGULATION_OF_CELLULAR_COMPONENT_MOVEMENT | 493 | 0.3931463 | 1.2594092 | 0 | 0.5858205 | 1 | 2600 | tags=19%, list=10%, signal=20% |
| GO_CELLULAR_RESPONSE_TO_NITROGEN_COMPOUND | 609 | 0.38792247 | 1.2593975 | 0 | 0.5856089 | 1 | 2614 | tags=18%, list=10%, signal=20% |
| GAUSSMANN_MLL_AF4_FUSION_TARGETS_C_UP | 156 | 0.43251076 | 1.2588359 | 0 | 0.418642 | 1 | 3444 | tags=31%, list=13%, signal=35% |
| IVANOVA_HEMATOPOIESIS_EARLY_PROGENITOR | 410 | 0.38705713 | 1.2584289 | 0 | 0.41909593 | 1 | 3569 | tags=27%, list=13%, signal=31% |
| HP_AUTISTIC_BEHAVIOR | 418 | 0.38516828 | 1.258412 | 0 | 0.58669096 | 1 | 2302 | tags=19%, list=9%, signal=21% |
| GO_CELLULAR_RESPONSE_TO_PEPTIDE_HORMONE_STIMULUS | 290 | 0.40574992 | 1.2582448 | 0 | 0.5867545 | 1 | 2614 | tags=22%, list=10%, signal=24% |
| GO_PROTEIN_POLYUBIQUITINATION | 315 | 0.40445527 | 1.258182 | 0.03846154 | 0.5867118 | 1 | 2857 | tags=26%, list=11%, signal=29% |
| GO_REGULATION_OF_ANATOMICAL_STRUCTURE_MORPHOGENESIS | 990 | 0.38467017 | 1.2579001 | 0 | 0.5867003 | 1 | 2887 | tags=19%, list=11%, signal=21% |
| HP_DEATH_IN_INFANCY | 138 | 0.4242133 | 1.2567102 | 0.03030303 | 0.58754003 | 1 | 4642 | tags=36%, list=17%, signal=43% |
| GO_REGULATION_OF_PEPTIDE_SECRETION | 340 | 0.39597443 | 1.2561561 | 0 | 0.5880465 | 1 | 2458 | tags=17%, list=9%, signal=19% |
| HP_ABNORMALITY_OF_SKIN_PHYSIOLOGY | 343 | 0.39941773 | 1.2557127 | 0 | 0.5879387 | 1 | 3197 | tags=20%, list=12%, signal=22% |
| HP_CONSTIPATION | 248 | 0.4143286 | 1.25527 | 0 | 0.5882861 | 1 | 2459 | tags=19%, list=9%, signal=20% |
| HP_ABNORMAL_EMOTION_AFFECT_BEHAVIOR | 391 | 0.39083058 | 1.2547956 | 0 | 0.58829355 | 1 | 2421 | tags=20%, list=9%, signal=22% |
| GO_REGULATION_OF_PEPTIDE_TRANSPORT | 602 | 0.38391563 | 1.253448 | 0 | 0.5908136 | 1 | 2747 | tags=21%, list=10%, signal=23% |
| GO_REGULATION_OF_DNA_BINDING | 111 | 0.43991956 | 1.2530236 | 0.02941177 | 0.591443 | 1 | 2396 | tags=23%, list=9%, signal=26% |
| HP_DEPRESSIVITY | 283 | 0.40301204 | 1.2524998 | 0 | 0.59273434 | 1 | 3241 | tags=22%, list=12%, signal=24% |
| SHEPARD_BMYB_MORPHOLINO_DN | 182 | 0.4242076 | 1.2519646 | 0.03703704 | 0.42705312 | 1 | 2480 | tags=23%, list=9%, signal=25% |
| HP_ABNORMAL_SYSTEMIC_BLOOD_PRESSURE | 353 | 0.3981871 | 1.251919 | 0 | 0.59344214 | 1 | 3497 | tags=22%, list=13%, signal=25% |
| GO_POSITIVE_REGULATION_OF_INFLAMMATORY_RESPONSE | 122 | 0.43968478 | 1.2507124 | 0.03448276 | 0.59399724 | 1 | 2899 | tags=23%, list=11%, signal=26% |
| GO_LEUKOCYTE_PROLIFERATION | 252 | 0.40576193 | 1.2502353 | 0.03333334 | 0.5946724 | 1 | 2604 | tags=16%, list=10%, signal=18% |
| GO_NCRNA_PROCESSING | 349 | 0.3918078 | 1.2499322 | 0 | 0.59541094 | 1 | 4015 | tags=42%, list=15%, signal=48% |
| REACTOME_ASPARAGINE_N_LINKED_GLYCOSYLATION | 281 | 0.41329652 | 1.2493821 | 0 | 0.43105394 | 1 | 2833 | tags=27%, list=11%, signal=30% |
| GO_TRANSITION_METAL_ION_BINDING | 870 | 0.3840188 | 1.2493652 | 0 | 0.59642214 | 1 | 3898 | tags=26%, list=14%, signal=29% |
| CHARAFE_BREAST_CANCER_LUMINAL_VS_MESENCHYMAL_UP | 379 | 0.39411166 | 1.2490805 | 0 | 0.4314504 | 1 | 3955 | tags=24%, list=15%, signal=28% |
| GRAESSMANN_APOPTOSIS_BY_DOXORUBICIN_DN | 1693 | 0.36584082 | 1.2489673 | 0 | 0.43137044 | 1 | 3242 | tags=28%, list=12%, signal=30% |
| BERTUCCI_MEDULLARY_VS_DUCTAL_BREAST_CANCER_UP | 175 | 0.42859375 | 1.2487766 | 0.02439024 | 0.4314609 | 1 | 2268 | tags=21%, list=8%, signal=23% |
| CHIANG_LIVER_CANCER_SUBCLASS_PROLIFERATION_DN | 134 | 0.43798527 | 1.2487037 | 0.05 | 0.4313462 | 1 | 4495 | tags=30%, list=17%, signal=36% |
| GO_LEUKOCYTE_DIFFERENTIATION | 454 | 0.37294662 | 1.2486757 | 0 | 0.5975249 | 1 | 3119 | tags=19%, list=12%, signal=21% |
| GO_REGULATION_OF_CYSTEINE_TYPE_ENDOPEPTIDASE_ACTIVITY | 206 | 0.40046248 | 1.2483945 | 0.03846154 | 0.5968973 | 1 | 2874 | tags=20%, list=11%, signal=23% |
| GO_CELL_MOTILITY | 1409 | 0.36929756 | 1.248038 | 0 | 0.5971409 | 1 | 3475 | tags=21%, list=13%, signal=23% |
| GO_APOPTOTIC_SIGNALING_PATHWAY | 546 | 0.39053568 | 1.2478342 | 0 | 0.5976236 | 1 | 2952 | tags=25%, list=11%, signal=27% |
| RAO_BOUND_BY_SALL4_ISOFORM_A | 145 | 0.4275011 | 1.2478325 | 0 | 0.4325486 | 1 | 3600 | tags=23%, list=13%, signal=26% |
| GO_BEHAVIOR | 524 | 0.3771351 | 1.2474773 | 0 | 0.5984894 | 1 | 2331 | tags=14%, list=9%, signal=15% |
| WP_IL18_SIGNALING_PATHWAY | 242 | 0.41396266 | 1.2472798 | 0.04878049 | 0.43287086 | 1 | 2267 | tags=21%, list=8%, signal=23% |
| PHONG_TNF_RESPONSE_NOT_VIA_P38 | 317 | 0.4077434 | 1.2470257 | 0.04166667 | 0.43291903 | 1 | 3536 | tags=29%, list=13%, signal=33% |
| GO_NUCLEOBASE_CONTAINING_SMALL_MOLECULE_BIOSYNTHETIC_PROCESS | 95 | 0.45890364 | 1.2464873 | 0.03225806 | 0.6003692 | 1 | 3623 | tags=28%, list=13%, signal=33% |
| GO_STRUCTURAL_MOLECULE_ACTIVITY | 553 | 0.38987786 | 1.2458802 | 0.03846154 | 0.6016401 | 1 | 2261 | tags=22%, list=8%, signal=24% |
| GO_ORGANOPHOSPHATE_BIOSYNTHETIC_PROCESS | 535 | 0.39205888 | 1.24577 | 0 | 0.60174185 | 1 | 3931 | tags=27%, list=15%, signal=31% |
| REACTOME_EPH_EPHRIN_SIGNALING | 89 | 0.46758637 | 1.2457646 | 0.03448276 | 0.4347776 | 1 | 2859 | tags=33%, list=11%, signal=36% |
| HP_ABNORMALITY_OF_THE_VOICE | 358 | 0.39512375 | 1.2456492 | 0 | 0.6019305 | 1 | 3128 | tags=22%, list=12%, signal=24% |
| HP_ABNORMALITY_OF_SKIN_ADNEXA_MORPHOLOGY | 1036 | 0.3655483 | 1.245509 | 0 | 0.6020811 | 1 | 3527 | tags=23%, list=13%, signal=25% |
| GO_NEGATIVE_REGULATION_OF_ION_TRANSPORT | 115 | 0.45137414 | 1.2454437 | 0.03225806 | 0.6021084 | 1 | 3152 | tags=21%, list=12%, signal=24% |
| HP_JAUNDICE | 139 | 0.45095688 | 1.2445679 | 0.03333334 | 0.60354865 | 1 | 4062 | tags=32%, list=15%, signal=38% |
| REACTOME_ORGANELLE_BIOGENESIS_AND_MAINTENANCE | 279 | 0.4095264 | 1.2439163 | 0 | 0.43638992 | 1 | 3907 | tags=35%, list=15%, signal=40% |
| GO_REGULATION_OF_DNA_BINDING_TRANSCRIPTION_FACTOR_ACTIVITY | 388 | 0.39884847 | 1.2438235 | 0 | 0.6043317 | 1 | 2604 | tags=21%, list=10%, signal=23% |
| GO_RESPONSE_TO_STEROID_HORMONE | 300 | 0.40676743 | 1.2433388 | 0 | 0.60406566 | 1 | 2488 | tags=21%, list=9%, signal=23% |
| REACTOME_GPCR_LIGAND_BINDING | 305 | 0.4184616 | 1.2431496 | 0.04761905 | 0.43753776 | 1 | 2555 | tags=9%, list=9%, signal=10% |
| BOYAULT_LIVER_CANCER_SUBCLASS_G3_UP | 183 | 0.42989072 | 1.2428827 | 0 | 0.4373158 | 1 | 3384 | tags=34%, list=13%, signal=38% |
| KOKKINAKIS_METHIONINE_DEPRIVATION_48HR_UP | 124 | 0.4367192 | 1.2427385 | 0.03333334 | 0.4373419 | 1 | 3146 | tags=25%, list=12%, signal=28% |
| GO_LIPID_LOCALIZATION | 343 | 0.4079411 | 1.2426975 | 0.04761905 | 0.60416585 | 1 | 3504 | tags=20%, list=13%, signal=23% |
| GO_ORGANOPHOSPHATE_CATABOLIC_PROCESS | 126 | 0.4432105 | 1.2423494 | 0.04347826 | 0.60412 | 1 | 4086 | tags=33%, list=15%, signal=38% |
| GO_REGULATION_OF_INTRACELLULAR_SIGNAL_TRANSDUCTION | 1601 | 0.36004636 | 1.2422264 | 0 | 0.6043098 | 1 | 3089 | tags=21%, list=11%, signal=22% |
| MILI_PSEUDOPODIA_CHEMOTAXIS_DN | 412 | 0.3859382 | 1.2422086 | 0.04347826 | 0.43759453 | 1 | 3091 | tags=27%, list=11%, signal=30% |
| GO_INTRACELLULAR_TRANSPORT | 1594 | 0.3515557 | 1.241707 | 0 | 0.60509413 | 1 | 2789 | tags=25%, list=10%, signal=26% |
| LASTOWSKA_NEUROBLASTOMA_COPY_NUMBER_UP | 167 | 0.42876562 | 1.2414567 | 0.05 | 0.43905437 | 1 | 3323 | tags=32%, list=12%, signal=36% |
| HP_ABNORMAL_CRANIAL_NERVE_MORPHOLOGY | 234 | 0.4150921 | 1.2413812 | 0 | 0.6054393 | 1 | 3636 | tags=26%, list=14%, signal=30% |
| SHETH_LIVER_CANCER_VS_TXNIP_LOSS_PAM1 | 237 | 0.41235724 | 1.2408667 | 0 | 0.4397171 | 1 | 3394 | tags=27%, list=13%, signal=31% |
| GO_NEGATIVE_REGULATION_OF_CYSTEINE_TYPE_ENDOPEPTIDASE_ACTIVITY | 80 | 0.44614688 | 1.2408563 | 0.03225806 | 0.6042826 | 1 | 2780 | tags=19%, list=10%, signal=21% |
| GRABARCZYK_BCL11B_TARGETS_UP | 66 | 0.4726147 | 1.2406415 | 0.03030303 | 0.439092 | 1 | 3029 | tags=29%, list=11%, signal=32% |
| GO_MONOCARBOXYLIC_ACID_BIOSYNTHETIC_PROCESS | 206 | 0.43938455 | 1.240554 | 0.03448276 | 0.6049368 | 1 | 3108 | tags=24%, list=12%, signal=27% |
| DELACROIX_RARG_BOUND_MEF | 329 | 0.39442912 | 1.2404678 | 0 | 0.4390114 | 1 | 1971 | tags=18%, list=7%, signal=19% |
| HP_LIMITATION_OF_JOINT_MOBILITY | 308 | 0.4001324 | 1.2401947 | 0 | 0.60541445 | 1 | 3441 | tags=21%, list=13%, signal=24% |
| GO_NERVOUS_SYSTEM_PROCESS | 859 | 0.36529446 | 1.2398001 | 0 | 0.6060253 | 1 | 3824 | tags=19%, list=14%, signal=21% |
| GO_MOLECULAR_TRANSDUCER_ACTIVITY | 796 | 0.36883208 | 1.2394116 | 0 | 0.605964 | 1 | 3472 | tags=13%, list=13%, signal=15% |
| GO_REGULATION_OF_BINDING | 329 | 0.39594033 | 1.2390281 | 0.04166667 | 0.60621583 | 1 | 2281 | tags=21%, list=8%, signal=23% |
| GO_HEART_DEVELOPMENT | 499 | 0.37830427 | 1.2384503 | 0 | 0.60684407 | 1 | 3322 | tags=18%, list=12%, signal=20% |
| GO_POLYMERIC_CYTOSKELETAL_FIBER | 576 | 0.36790925 | 1.2381736 | 0 | 0.6068399 | 1 | 3277 | tags=24%, list=12%, signal=27% |
| GO_ORGANELLE_DISASSEMBLY | 103 | 0.44703496 | 1.2380464 | 0.02702703 | 0.6063407 | 1 | 2357 | tags=23%, list=9%, signal=25% |
| DAZARD_RESPONSE_TO_UV_NHEK_UP | 185 | 0.4286408 | 1.2378479 | 0.02380952 | 0.44199446 | 1 | 3127 | tags=34%, list=12%, signal=38% |
| GO_ION_HOMEOSTASIS | 663 | 0.3776126 | 1.2377524 | 0 | 0.60587615 | 1 | 2810 | tags=16%, list=10%, signal=18% |
| YAGI_AML_FAB_MARKERS | 176 | 0.41496912 | 1.2372056 | 0.03448276 | 0.44281176 | 1 | 3776 | tags=32%, list=14%, signal=37% |
| GO_SUPRAMOLECULAR_COMPLEX | 1073 | 0.36701137 | 1.2371277 | 0 | 0.6064715 | 1 | 3117 | tags=22%, list=12%, signal=23% |
| GO_PHOSPHOLIPID_METABOLIC_PROCESS | 400 | 0.38336974 | 1.2369297 | 0 | 0.6066384 | 1 | 3957 | tags=27%, list=15%, signal=31% |
| HP_CHOLESTASIS | 185 | 0.42278978 | 1.2369125 | 0 | 0.606485 | 1 | 3524 | tags=26%, list=13%, signal=30% |
| KEGG_LYSOSOME | 113 | 0.44071177 | 1.2365962 | 0.02941177 | 0.44330522 | 1 | 3112 | tags=28%, list=12%, signal=32% |
| GO_ALCOHOL_METABOLIC_PROCESS | 315 | 0.3992851 | 1.2364122 | 0 | 0.6070811 | 1 | 4009 | tags=30%, list=15%, signal=35% |
| GO_EARLY_ENDOSOME_MEMBRANE | 137 | 0.42967686 | 1.2350026 | 0.03703704 | 0.60881346 | 1 | 2566 | tags=20%, list=10%, signal=22% |
| HP_MALABSORPTION | 173 | 0.42561728 | 1.2349151 | 0.02941177 | 0.6086135 | 1 | 3524 | tags=26%, list=13%, signal=30% |
| GO_TRANSPORTER_ACTIVITY | 989 | 0.36303377 | 1.2342346 | 0 | 0.6097586 | 1 | 3471 | tags=17%, list=13%, signal=19% |
| GO_ANATOMICAL_STRUCTURE_FORMATION_INVOLVED_IN_MORPHOGENESIS | 974 | 0.36338606 | 1.2337749 | 0 | 0.6100207 | 1 | 3430 | tags=20%, list=13%, signal=22% |
| HP_KYPHOSIS | 358 | 0.4037361 | 1.233634 | 0.04347826 | 0.6103311 | 1 | 3128 | tags=24%, list=12%, signal=27% |
| GO_REGULATION_OF_CELLULAR_COMPONENT_MOVEMENT | 897 | 0.37595102 | 1.233617 | 0 | 0.61013114 | 1 | 3083 | tags=20%, list=11%, signal=22% |
| KEGG_PURINE_METABOLISM | 143 | 0.43069974 | 1.2332978 | 0.03125 | 0.44796887 | 1 | 3206 | tags=27%, list=12%, signal=30% |
| SMID_BREAST_CANCER_BASAL_DN | 590 | 0.38378185 | 1.2331909 | 0 | 0.4479052 | 1 | 3903 | tags=23%, list=14%, signal=27% |
| MARTORIATI_MDM4_TARGETS_FETAL_LIVER_DN | 476 | 0.38095635 | 1.2322015 | 0 | 0.44909713 | 1 | 2714 | tags=26%, list=10%, signal=28% |
| HP_AGENESIS_OF_CORPUS_CALLOSUM | 244 | 0.4105668 | 1.2321322 | 0 | 0.61106133 | 1 | 3202 | tags=20%, list=12%, signal=22% |
| UEDA_PERIFERAL_CLOCK | 157 | 0.42048752 | 1.231788 | 0.03125 | 0.4496667 | 1 | 2360 | tags=25%, list=9%, signal=28% |
| IVANOVA_HEMATOPOIESIS_STEM_CELL_LONG_TERM | 198 | 0.4316887 | 1.2314463 | 0.03703704 | 0.45005116 | 1 | 3402 | tags=26%, list=13%, signal=30% |
| GO_CELLULAR_HOMEOSTASIS | 802 | 0.38105193 | 1.2313184 | 0 | 0.61213696 | 1 | 2947 | tags=19%, list=11%, signal=21% |
| HP_ABNORMALITY_OF_HINDBRAIN_MORPHOLOGY | 706 | 0.3718162 | 1.231317 | 0 | 0.61191493 | 1 | 3510 | tags=25%, list=13%, signal=27% |
| GO_EPITHELIAL_CELL_DEVELOPMENT | 195 | 0.4092126 | 1.2312815 | 0.03333334 | 0.61177427 | 1 | 3361 | tags=22%, list=12%, signal=25% |
| GO_ORGANIC_HYDROXY_COMPOUND_BIOSYNTHETIC_PROCESS | 218 | 0.4186493 | 1.2312388 | 0 | 0.6116861 | 1 | 3519 | tags=24%, list=13%, signal=27% |
| GO_NEGATIVE_REGULATION_OF_MULTICELLULAR_ORGANISMAL_PROCESS | 1042 | 0.37212062 | 1.2311314 | 0 | 0.61180645 | 1 | 2678 | tags=17%, list=10%, signal=18% |
| GO_REGULATION_OF_HORMONE_SECRETION | 218 | 0.4151016 | 1.2308998 | 0 | 0.61190784 | 1 | 2389 | tags=15%, list=9%, signal=16% |
| GO_NEUROGENESIS | 1519 | 0.34889892 | 1.2307931 | 0 | 0.61156905 | 1 | 3102 | tags=18%, list=12%, signal=19% |
| MARTENS_TRETINOIN_RESPONSE_DN | 693 | 0.37265217 | 1.2306143 | 0 | 0.45049992 | 1 | 2707 | tags=27%, list=10%, signal=29% |
| HELLER_HDAC_TARGETS_UP | 269 | 0.40710703 | 1.2305697 | 0 | 0.45034304 | 1 | 3198 | tags=26%, list=12%, signal=29% |
| GO_GLYCOSAMINOGLYCAN_BINDING | 187 | 0.41731998 | 1.2302421 | 0.04 | 0.6117395 | 1 | 2752 | tags=17%, list=10%, signal=18% |
| HP_OCULAR_ANTERIOR_SEGMENT_DYSGENESIS | 229 | 0.40999654 | 1.2301971 | 0 | 0.61147755 | 1 | 4269 | tags=30%, list=16%, signal=36% |
| GO_G_PROTEIN_COUPLED_RECEPTOR_ACTIVITY | 297 | 0.39931542 | 1.2296773 | 0.04347826 | 0.6127443 | 1 | 3472 | tags=9%, list=13%, signal=11% |
| HP_ABNORMALITY_OF_PRENATAL_DEVELOPMENT_OR_BIRTH | 686 | 0.37650892 | 1.2292603 | 0 | 0.61309886 | 1 | 3524 | tags=22%, list=13%, signal=25% |
| GO_ACTIN_BINDING | 383 | 0.38770425 | 1.2285551 | 0 | 0.61379284 | 1 | 3475 | tags=23%, list=13%, signal=26% |
| GO_CYTOSKELETAL_PROTEIN_BINDING | 885 | 0.3648037 | 1.2285136 | 0 | 0.61370504 | 1 | 3502 | tags=24%, list=13%, signal=27% |
| MILI_PSEUDOPODIA_HAPTOTAXIS_DN | 648 | 0.3758309 | 1.228512 | 0 | 0.45164785 | 1 | 3463 | tags=28%, list=13%, signal=31% |
| GO_CELLULAR_RESPONSE_TO_LIPID | 492 | 0.38227183 | 1.2284017 | 0 | 0.61358094 | 1 | 2614 | tags=19%, list=10%, signal=21% |
| GO_PROTEIN_KINASE_B_SIGNALING | 217 | 0.40750152 | 1.2280042 | 0.02777778 | 0.6137149 | 1 | 2357 | tags=17%, list=9%, signal=18% |
| GO_NUCLEOLUS | 842 | 0.37846613 | 1.2275187 | 0 | 0.6128177 | 1 | 3050 | tags=29%, list=11%, signal=32% |
| SCHAEFFER_PROSTATE_DEVELOPMENT_48HR_DN | 376 | 0.39575386 | 1.2271011 | 0 | 0.45301 | 1 | 4136 | tags=27%, list=15%, signal=32% |
| GO_CHEMICAL_HOMEOSTASIS | 969 | 0.36610904 | 1.2268908 | 0 | 0.6138153 | 1 | 2810 | tags=17%, list=10%, signal=18% |
| LINDGREN_BLADDER_CANCER_CLUSTER_3_UP | 299 | 0.39303696 | 1.2268744 | 0 | 0.4533184 | 1 | 3001 | tags=31%, list=11%, signal=35% |
| PEDRIOLI_MIR31_TARGETS_DN | 336 | 0.38552195 | 1.2268217 | 0 | 0.4531399 | 1 | 3980 | tags=26%, list=15%, signal=30% |
| REACTOME_INFECTIOUS_DISEASE | 675 | 0.38402963 | 1.2266546 | 0 | 0.4533334 | 1 | 2395 | tags=24%, list=9%, signal=26% |
| GO_I_KAPPAB_KINASE_NF_KAPPAB_SIGNALING | 257 | 0.39263925 | 1.226565 | 0 | 0.6143491 | 1 | 2900 | tags=23%, list=11%, signal=25% |
| HP_ARTHRALGIA | 133 | 0.4323453 | 1.2264037 | 0.03225806 | 0.6147043 | 1 | 2434 | tags=20%, list=9%, signal=22% |
| GO_MICROBODY | 126 | 0.4349782 | 1.226179 | 0.03448276 | 0.61453736 | 1 | 4079 | tags=40%, list=15%, signal=47% |
| GO_REGULATION_OF_PEPTIDE_HORMONE_SECRETION | 173 | 0.43121916 | 1.2257066 | 0.03125 | 0.6147804 | 1 | 2389 | tags=16%, list=9%, signal=17% |
| TAKEDA_TARGETS_OF_NUP98_HOXA9_FUSION_8D_DN | 156 | 0.41227385 | 1.225678 | 0.03030303 | 0.45563447 | 1 | 2685 | tags=19%, list=10%, signal=21% |
| HP_ABNORMAL_LEUKOCYTE_COUNT | 292 | 0.39426968 | 1.2254114 | 0.03125 | 0.61525744 | 1 | 3549 | tags=26%, list=13%, signal=30% |
| GO_PROTEIN_DOMAIN_SPECIFIC_BINDING | 650 | 0.3687995 | 1.2249031 | 0 | 0.61609536 | 1 | 2729 | tags=20%, list=10%, signal=22% |
| RAGHAVACHARI_PLATELET_SPECIFIC_GENES | 62 | 0.51139796 | 1.2237624 | 0.03846154 | 0.4569725 | 1 | 2556 | tags=29%, list=9%, signal=32% |
| REACTOME_DISEASE | 1318 | 0.35752895 | 1.2234569 | 0 | 0.4573647 | 1 | 2837 | tags=22%, list=11%, signal=24% |
| BENPORATH_MYC_MAX_TARGETS | 715 | 0.37199774 | 1.2229735 | 0 | 0.45773262 | 1 | 3419 | tags=33%, list=13%, signal=37% |
| HP_ABNORMALITY_OF_FLUID_REGULATION | 404 | 0.3828436 | 1.2228798 | 0 | 0.61911845 | 1 | 3128 | tags=19%, list=12%, signal=21% |
| LI_INDUCED_T_TO_NATURAL_KILLER_DN | 128 | 0.43261704 | 1.2212965 | 0.03571429 | 0.45935878 | 1 | 2814 | tags=23%, list=10%, signal=25% |
| GO_ENZYME_REGULATOR_ACTIVITY | 893 | 0.36004695 | 1.221053 | 0 | 0.62206143 | 1 | 2968 | tags=21%, list=11%, signal=23% |
| GO_T_CELL_PROLIFERATION | 159 | 0.42489177 | 1.2209702 | 0.02777778 | 0.6218796 | 1 | 2604 | tags=16%, list=10%, signal=18% |
| GABRIELY_MIR21_TARGETS | 276 | 0.38145453 | 1.2208564 | 0.05 | 0.46001145 | 1 | 4049 | tags=35%, list=15%, signal=41% |
| GO_REGULATION_OF_ION_TRANSPORT | 588 | 0.37203786 | 1.2199019 | 0 | 0.62312216 | 1 | 3440 | tags=17%, list=13%, signal=19% |
| GO_STEM_CELL_PROLIFERATION | 109 | 0.41811737 | 1.2195964 | 0.03125 | 0.6228429 | 1 | 2318 | tags=19%, list=9%, signal=21% |
| HP_IMPAIRMENT_OF_ACTIVITIES_OF_DAILY_LIVING | 150 | 0.40710422 | 1.219583 | 0.02857143 | 0.62267715 | 1 | 3410 | tags=21%, list=13%, signal=24% |
| BYSTRYKH_HEMATOPOIESIS_STEM_CELL_QTL_TRANS | 703 | 0.37688783 | 1.219531 | 0 | 0.46253884 | 1 | 3392 | tags=27%, list=13%, signal=30% |
| GO_REGULATION_OF_PROTEIN_CONTAINING_COMPLEX_ASSEMBLY | 390 | 0.383214 | 1.2194656 | 0 | 0.62217015 | 1 | 3421 | tags=26%, list=13%, signal=29% |
| HP_HYPERKERATOSIS | 204 | 0.4018607 | 1.2192286 | 0.03333334 | 0.62210125 | 1 | 3490 | tags=24%, list=13%, signal=27% |
| BLALOCK_ALZHEIMERS_DISEASE_UP | 1508 | 0.36634764 | 1.218918 | 0 | 0.46247834 | 1 | 3420 | tags=26%, list=13%, signal=29% |
| GROSS_HYPOXIA_VIA_HIF1A_DN | 104 | 0.42825708 | 1.2184284 | 0.03030303 | 0.46300116 | 1 | 3932 | tags=32%, list=15%, signal=37% |
| GO_TELENCEPHALON_DEVELOPMENT | 235 | 0.39721677 | 1.2184013 | 0.03125 | 0.6236714 | 1 | 2089 | tags=17%, list=8%, signal=19% |
| GO_MEMBRANE_ORGANIZATION | 783 | 0.3636362 | 1.2183684 | 0 | 0.62355036 | 1 | 3113 | tags=25%, list=12%, signal=27% |
| MARTINEZ_RB1_TARGETS_DN | 494 | 0.38494048 | 1.2170261 | 0 | 0.46472597 | 1 | 2784 | tags=25%, list=10%, signal=27% |
| HP_ABNORMAL_VASCULAR_MORPHOLOGY | 642 | 0.37449378 | 1.216445 | 0 | 0.62481076 | 1 | 3197 | tags=22%, list=12%, signal=24% |
| KYNG_DNA_DAMAGE_UP | 188 | 0.40955073 | 1.215904 | 0.04347826 | 0.46685165 | 1 | 2758 | tags=20%, list=10%, signal=22% |
| GO_GLUCOSE_METABOLIC_PROCESS | 185 | 0.4160751 | 1.2154632 | 0.04347826 | 0.62548816 | 1 | 2984 | tags=25%, list=11%, signal=28% |
| MULLIGHAN_MLL_SIGNATURE_1_DN | 208 | 0.40748206 | 1.215099 | 0.02857143 | 0.46804455 | 1 | 3684 | tags=27%, list=14%, signal=32% |
| GO_POSTTRANSCRIPTIONAL_REGULATION_OF_GENE_EXPRESSION | 552 | 0.3712849 | 1.2150544 | 0 | 0.62564296 | 1 | 2772 | tags=25%, list=10%, signal=27% |
| WANG_ESOPHAGUS_CANCER_VS_NORMAL_UP | 108 | 0.44523412 | 1.2149038 | 0.04255319 | 0.46832106 | 1 | 3586 | tags=34%, list=13%, signal=39% |
| HP_LIMB_MUSCLE_WEAKNESS | 205 | 0.4055289 | 1.2146522 | 0.04166667 | 0.6258651 | 1 | 3746 | tags=26%, list=14%, signal=30% |
| GO_RESPONSE_TO_INORGANIC_SUBSTANCE | 487 | 0.38820672 | 1.2146345 | 0 | 0.6257073 | 1 | 2938 | tags=18%, list=11%, signal=20% |
| GO_LAMELLIPODIUM | 192 | 0.4006145 | 1.213859 | 0.03125 | 0.6259435 | 1 | 3368 | tags=26%, list=13%, signal=30% |
| GO_NEGATIVE_REGULATION_OF_PHOSPHORYLATION | 404 | 0.3754482 | 1.2137423 | 0.04545455 | 0.6257319 | 1 | 3104 | tags=27%, list=12%, signal=30% |
| GO_MUSCLE_ORGAN_DEVELOPMENT | 331 | 0.40146092 | 1.2135913 | 0.04166667 | 0.6260231 | 1 | 2675 | tags=16%, list=10%, signal=17% |
| GO_CENTROSOME | 538 | 0.3764772 | 1.2135671 | 0 | 0.6259308 | 1 | 4158 | tags=34%, list=15%, signal=39% |
| GO_REPRODUCTIVE_SYSTEM_DEVELOPMENT | 380 | 0.37824246 | 1.2134075 | 0 | 0.62584394 | 1 | 2352 | tags=15%, list=9%, signal=16% |
| HP_CONGENITAL_ONSET | 264 | 0.39337778 | 1.2130222 | 0.04761905 | 0.6255015 | 1 | 4127 | tags=31%, list=15%, signal=36% |
| TONKS_TARGETS_OF_RUNX1_RUNX1T1_FUSION_HSC_DN | 169 | 0.40393344 | 1.2130103 | 0.04166667 | 0.46915781 | 1 | 4144 | tags=26%, list=15%, signal=31% |
| GO_RESPONSE_TO_TOXIC_SUBSTANCE | 210 | 0.4272742 | 1.212834 | 0.04166667 | 0.62560964 | 1 | 2750 | tags=22%, list=10%, signal=25% |
| NAKAMURA_TUMOR_ZONE_PERIPHERAL_VS_CENTRAL_DN | 548 | 0.3755922 | 1.2124684 | 0.04347826 | 0.46894622 | 1 | 3381 | tags=27%, list=13%, signal=30% |
| HP_SHORT_LONG_BONE | 165 | 0.42005026 | 1.2121723 | 0.03125 | 0.62647814 | 1 | 3484 | tags=25%, list=13%, signal=29% |
| GO_PEPTIDE_HORMONE_SECRETION | 211 | 0.4022539 | 1.2121582 | 0.03448276 | 0.626096 | 1 | 1331 | tags=12%, list=5%, signal=12% |
| BROWN_MYELOID_CELL_DEVELOPMENT_UP | 153 | 0.40803885 | 1.2118042 | 0.02941177 | 0.46970195 | 1 | 3236 | tags=20%, list=12%, signal=23% |
| GO_CLUSTER_OF_ACTIN_BASED_CELL_PROJECTIONS | 136 | 0.43631053 | 1.2091047 | 0.02777778 | 0.6291776 | 1 | 2041 | tags=12%, list=8%, signal=13% |
| HP_POLYMICROGYRIA | 164 | 0.39080083 | 1.2081132 | 0.03125 | 0.62959945 | 1 | 4204 | tags=35%, list=16%, signal=41% |
| HP_ABNORMALITY_OF_THE_CURVATURE_OF_THE_VERTEBRAL_COLUMN | 910 | 0.3479279 | 1.2071426 | 0 | 0.629672 | 1 | 3537 | tags=23%, list=13%, signal=26% |
| GRYDER_PAX3FOXO1_TOP_ENHANCERS | 422 | 0.37051025 | 1.2067857 | 0 | 0.47721934 | 1 | 3938 | tags=35%, list=15%, signal=40% |
| HP_ABNORMAL_BONE_STRUCTURE | 586 | 0.38541386 | 1.2066711 | 0 | 0.629924 | 1 | 3258 | tags=23%, list=12%, signal=26% |
| GO_POSITIVE_REGULATION_OF_TRANSPORT | 819 | 0.3664088 | 1.2060091 | 0 | 0.6304353 | 1 | 2530 | tags=18%, list=9%, signal=19% |
| GO_VESICLE_LOCALIZATION | 215 | 0.39870986 | 1.2057692 | 0.03846154 | 0.6306872 | 1 | 3871 | tags=35%, list=14%, signal=40% |
| HP_ABNORMALITY_OF_CONNECTIVE_TISSUE | 1124 | 0.35880542 | 1.2050427 | 0 | 0.63110757 | 1 | 3510 | tags=21%, list=13%, signal=23% |
| GO_POSITIVE_REGULATION_OF_MOLECULAR_FUNCTION | 1588 | 0.35478905 | 1.2042766 | 0 | 0.63085735 | 1 | 2948 | tags=19%, list=11%, signal=20% |
| VERHAAK_GLIOBLASTOMA_CLASSICAL | 145 | 0.4028137 | 1.2035306 | 0.02777778 | 0.48063594 | 1 | 4123 | tags=28%, list=15%, signal=32% |
| GO_SENSORY_PERCEPTION_OF_LIGHT_STIMULUS | 179 | 0.40148577 | 1.2027124 | 0.03703704 | 0.6331892 | 1 | 4294 | tags=17%, list=16%, signal=20% |
| LINSLEY_MIR16_TARGETS | 198 | 0.37750933 | 1.2026577 | 0.03571429 | 0.48091942 | 1 | 2950 | tags=26%, list=11%, signal=29% |
| REACTOME_MAPK_FAMILY_SIGNALING_CASCADES | 305 | 0.39427125 | 1.2025454 | 0 | 0.4805935 | 1 | 2841 | tags=22%, list=11%, signal=24% |
| REACTOME_ANTIGEN_PROCESSING_UBIQUITINATION_PROTEASOME_DEGRADATION | 285 | 0.3973857 | 1.201978 | 0 | 0.4813256 | 1 | 2857 | tags=24%, list=11%, signal=27% |
| GO_POSITIVE_REGULATION_OF_DNA_BINDING_TRANSCRIPTION_FACTOR_ACTIVITY | 239 | 0.3862241 | 1.2017792 | 0 | 0.63339585 | 1 | 2530 | tags=19%, list=9%, signal=21% |
| HP_ABNORMAL_RESPIRATORY_SYSTEM_MORPHOLOGY | 911 | 0.35977453 | 1.2017663 | 0 | 0.633025 | 1 | 3197 | tags=21%, list=12%, signal=23% |
| HP_ABNORMAL_CORPUS_CALLOSUM_MORPHOLOGY | 649 | 0.36821556 | 1.2013423 | 0 | 0.6333607 | 1 | 4140 | tags=30%, list=15%, signal=34% |
| HP_MORTALITY_AGING | 222 | 0.4079754 | 1.2009459 | 0.03030303 | 0.63368034 | 1 | 2350 | tags=17%, list=9%, signal=18% |
| GO_REGULATION_OF_PROTEIN_LOCALIZATION | 870 | 0.3683365 | 1.1999236 | 0 | 0.63431656 | 1 | 2647 | tags=21%, list=10%, signal=23% |
| BROWNE_HCMV_INFECTION_14HR_DN | 254 | 0.3993982 | 1.1998626 | 0.04 | 0.48501772 | 1 | 3718 | tags=27%, list=14%, signal=31% |
| REACTOME_METABOLISM_OF_VITAMINS_AND_COFACTORS | 161 | 0.41094747 | 1.1998161 | 0.03333334 | 0.4844987 | 1 | 4295 | tags=30%, list=16%, signal=35% |
| GO_NEGATIVE_REGULATION_OF_CELL_ACTIVATION | 174 | 0.40119615 | 1.1993905 | 0.02941177 | 0.63431287 | 1 | 2158 | tags=14%, list=8%, signal=16% |
| DODD_NASOPHARYNGEAL_CARCINOMA_DN | 1258 | 0.34756085 | 1.1986221 | 0 | 0.484776 | 1 | 3396 | tags=29%, list=13%, signal=31% |
| HP_ABNORMAL_FACIAL_EXPRESSION | 118 | 0.4211075 | 1.1980661 | 0.03571429 | 0.63522273 | 1 | 2353 | tags=19%, list=9%, signal=21% |
| GO_POSITIVE_REGULATION_OF_INTRACELLULAR_SIGNAL_TRANSDUCTION | 896 | 0.36119533 | 1.1976191 | 0 | 0.63520384 | 1 | 3077 | tags=20%, list=11%, signal=22% |
| MARSON_BOUND_BY_FOXP3_UNSTIMULATED | 1106 | 0.34067142 | 1.1969415 | 0 | 0.48534727 | 1 | 3124 | tags=23%, list=12%, signal=25% |
| GO_GUANYL_NUCLEOTIDE_BINDING | 334 | 0.3699247 | 1.196042 | 0 | 0.6357719 | 1 | 2819 | tags=22%, list=10%, signal=24% |
| ODONNELL_TFRC_TARGETS_UP | 286 | 0.4000515 | 1.194922 | 0.04761905 | 0.4879737 | 1 | 3467 | tags=19%, list=13%, signal=22% |
| VECCHI_GASTRIC_CANCER_EARLY_UP | 376 | 0.37869227 | 1.1943799 | 0.03846154 | 0.48855355 | 1 | 3281 | tags=25%, list=12%, signal=28% |
| GO_RESPONSE_TO_INSULIN | 253 | 0.38924563 | 1.1938987 | 0.04761905 | 0.6370978 | 1 | 2614 | tags=23%, list=10%, signal=25% |
| HP_GAIT_DISTURBANCE | 857 | 0.37763414 | 1.1937573 | 0 | 0.63719404 | 1 | 4059 | tags=28%, list=15%, signal=32% |
| GO_MODIFICATION_DEPENDENT_MACROMOLECULE_CATABOLIC_PROCESS | 588 | 0.37196782 | 1.1927133 | 0 | 0.63835615 | 1 | 2896 | tags=24%, list=11%, signal=27% |
| HP_ABNORMAL_JOINT_MORPHOLOGY | 847 | 0.3660133 | 1.1908454 | 0 | 0.6400805 | 1 | 3526 | tags=24%, list=13%, signal=27% |
| GO_ZINC_ION_BINDING | 666 | 0.36302638 | 1.1908138 | 0 | 0.63999677 | 1 | 3331 | tags=23%, list=12%, signal=25% |
| GO_LYMPHOCYTE_ACTIVATION_INVOLVED_IN_IMMUNE_RESPONSE | 139 | 0.3999897 | 1.1906646 | 0.03225806 | 0.63989854 | 1 | 2651 | tags=17%, list=10%, signal=18% |
| REACTOME_CELL_CYCLE | 602 | 0.36172792 | 1.1905086 | 0 | 0.49213046 | 1 | 3040 | tags=27%, list=11%, signal=30% |
| GO_REGULATION_OF_LEUKOCYTE_DIFFERENTIATION | 247 | 0.38530096 | 1.1903653 | 0.04545455 | 0.6400172 | 1 | 3102 | tags=19%, list=12%, signal=21% |
| KIM_WT1_TARGETS_12HR_DN | 200 | 0.40051275 | 1.1902026 | 0.03448276 | 0.49237105 | 1 | 3058 | tags=31%, list=11%, signal=34% |
| GO_RESPONSE_TO_ENDOPLASMIC_RETICULUM_STRESS | 265 | 0.3822652 | 1.1894045 | 0.03571429 | 0.6405094 | 1 | 2938 | tags=27%, list=11%, signal=30% |
| REACTOME_SIGNALING_BY_GPCR | 618 | 0.3569289 | 1.1890744 | 0 | 0.49251068 | 1 | 2971 | tags=13%, list=11%, signal=15% |
| HP_HYPOGONADISM | 330 | 0.4018239 | 1.1889575 | 0 | 0.6411108 | 1 | 3128 | tags=18%, list=12%, signal=20% |
| GO_POSITIVE_REGULATION_OF_CATALYTIC_ACTIVITY | 1283 | 0.34601986 | 1.1889318 | 0 | 0.64099306 | 1 | 2948 | tags=19%, list=11%, signal=21% |
| GO_HYDROLASE_ACTIVITY_ACTING_ON_ACID_ANHYDRIDES | 762 | 0.35510263 | 1.1868465 | 0 | 0.6450233 | 1 | 3289 | tags=23%, list=12%, signal=25% |
| GO_CELLULAR_ION_HOMEOSTASIS | 543 | 0.36982253 | 1.1864 | 0.04166667 | 0.64560133 | 1 | 2810 | tags=16%, list=10%, signal=18% |
| GO_REGULATION_OF_CELL_CYCLE_PROCESS | 676 | 0.3592976 | 1.1860245 | 0 | 0.64606494 | 1 | 3325 | tags=26%, list=12%, signal=29% |
| GO_REGULATION_OF_INTRACELLULAR_TRANSPORT | 329 | 0.38605148 | 1.1856769 | 0.03571429 | 0.646413 | 1 | 2747 | tags=25%, list=10%, signal=27% |
| MULLIGHAN_MLL_SIGNATURE_2_DN | 245 | 0.40490416 | 1.1856157 | 0.04347826 | 0.49687 | 1 | 1919 | tags=16%, list=7%, signal=17% |
| THUM_SYSTOLIC_HEART_FAILURE_UP | 376 | 0.37404338 | 1.1855904 | 0 | 0.4966454 | 1 | 3750 | tags=30%, list=14%, signal=34% |
| GO_VESICLE_ORGANIZATION | 309 | 0.3870673 | 1.1853198 | 0 | 0.64609504 | 1 | 2535 | tags=24%, list=9%, signal=26% |
| GO_REGULATION_OF_CELL_CELL_ADHESION | 366 | 0.37975013 | 1.1838741 | 0.05 | 0.64706475 | 1 | 2604 | tags=17%, list=10%, signal=18% |
| HP_ABNORMALITY_OF_DIGESTIVE_SYSTEM_MORPHOLOGY | 766 | 0.35757864 | 1.1828635 | 0 | 0.6472218 | 1 | 3524 | tags=21%, list=13%, signal=24% |
| HP_SHORT_STATURE | 1039 | 0.35330448 | 1.1825163 | 0 | 0.64746344 | 1 | 3984 | tags=27%, list=15%, signal=30% |
| HP_ABNORMAL_ANTERIOR_EYE_SEGMENT_MORPHOLOGY | 988 | 0.36434454 | 1.1823361 | 0 | 0.6477729 | 1 | 3526 | tags=21%, list=13%, signal=23% |
| GO_NEGATIVE_REGULATION_OF_DEVELOPMENTAL_PROCESS | 837 | 0.3562416 | 1.1801668 | 0 | 0.6505048 | 1 | 2678 | tags=17%, list=10%, signal=18% |
| GO_ORGANELLE_SUBCOMPARTMENT | 325 | 0.36896732 | 1.1800517 | 0.03703704 | 0.6505417 | 1 | 3205 | tags=27%, list=12%, signal=30% |
| FEVR_CTNNB1_TARGETS_DN | 523 | 0.36019 | 1.1798848 | 0 | 0.5015742 | 1 | 3541 | tags=31%, list=13%, signal=35% |
| GO_NEGATIVE_REGULATION_OF_RESPONSE_TO_EXTERNAL_STIMULUS | 309 | 0.3980284 | 1.1798295 | 0 | 0.65084493 | 1 | 1729 | tags=13%, list=6%, signal=13% |
| GO_PATTERN_SPECIFICATION_PROCESS | 376 | 0.36598155 | 1.1790674 | 0 | 0.65190595 | 1 | 4003 | tags=21%, list=15%, signal=24% |
| GO_RESPONSE_TO_MECHANICAL_STIMULUS | 190 | 0.39381796 | 1.1784797 | 0 | 0.65251684 | 1 | 2530 | tags=14%, list=9%, signal=16% |
| REACTOME_LEISHMANIA_INFECTION | 207 | 0.4010279 | 1.178454 | 0.03571429 | 0.50303674 | 1 | 2478 | tags=16%, list=9%, signal=18% |
| GO_REGULATION_OF_MITOTIC_CELL_CYCLE | 584 | 0.3708508 | 1.1780468 | 0 | 0.6527554 | 1 | 3307 | tags=27%, list=12%, signal=31% |
| BONOME_OVARIAN_CANCER_SURVIVAL_SUBOPTIMAL_DEBULKING | 454 | 0.36678213 | 1.1766208 | 0.04347826 | 0.5056218 | 1 | 3240 | tags=23%, list=12%, signal=26% |
| REACTOME_METABOLISM_OF_LIPIDS | 638 | 0.37127665 | 1.1760395 | 0 | 0.5068652 | 1 | 3506 | tags=25%, list=13%, signal=28% |
| GO_RESPONSE_TO_METAL_ION | 314 | 0.38185614 | 1.1759173 | 0.04166667 | 0.65460837 | 1 | 3010 | tags=19%, list=11%, signal=21% |
| CAIRO_HEPATOBLASTOMA_DN | 216 | 0.39556572 | 1.1755514 | 0.04166667 | 0.50753707 | 1 | 2925 | tags=21%, list=11%, signal=23% |
| GO_HOMEOSTATIC_PROCESS | 1619 | 0.34396887 | 1.17477 | 0 | 0.6552584 | 1 | 2676 | tags=16%, list=10%, signal=17% |
| GO_REGULATION_OF_PHOSPHORYLATION | 1387 | 0.3439897 | 1.1742866 | 0 | 0.6553509 | 1 | 3299 | tags=21%, list=12%, signal=23% |
| GO_REGULATION_OF_CALCIUM_ION_TRANSPORT | 206 | 0.38989142 | 1.1741779 | 0.03125 | 0.6552721 | 1 | 2810 | tags=15%, list=10%, signal=17% |
| RATTENBACHER_BOUND_BY_CELF1 | 344 | 0.37357435 | 1.1735799 | 0 | 0.50977814 | 1 | 2943 | tags=21%, list=11%, signal=23% |
| MARSON_BOUND_BY_E2F4_UNSTIMULATED | 637 | 0.35380307 | 1.1717649 | 0 | 0.50951034 | 1 | 3996 | tags=32%, list=15%, signal=37% |
| GO_REGULATION_OF_CELLULAR_PROTEIN_CATABOLIC_PROCESS | 242 | 0.3961772 | 1.1717367 | 0.03571429 | 0.6555758 | 1 | 2481 | tags=24%, list=9%, signal=26% |
| GO_EXTRINSIC_COMPONENT_OF_MEMBRANE | 274 | 0.3861382 | 1.171081 | 0.03333334 | 0.65694904 | 1 | 3283 | tags=21%, list=12%, signal=24% |
| CHARAFE_BREAST_CANCER_LUMINAL_VS_MESENCHYMAL_DN | 435 | 0.35976148 | 1.1707015 | 0 | 0.51027817 | 1 | 3173 | tags=24%, list=12%, signal=27% |
| GO_POSITIVE_REGULATION_OF_MULTICELLULAR_ORGANISMAL_PROCESS | 1533 | 0.3516637 | 1.1702024 | 0 | 0.6582052 | 1 | 2676 | tags=16%, list=10%, signal=17% |
| GO_MACROMOLECULE_CATABOLIC_PROCESS | 1287 | 0.34487325 | 1.1696193 | 0 | 0.6584844 | 1 | 2837 | tags=24%, list=11%, signal=26% |
| GO_OSSIFICATION | 359 | 0.38375974 | 1.1685729 | 0 | 0.65889573 | 1 | 3127 | tags=19%, list=12%, signal=21% |
| GO_CELL_PROJECTION_ASSEMBLY | 540 | 0.36077714 | 1.1682547 | 0 | 0.65832865 | 1 | 3497 | tags=25%, list=13%, signal=28% |
| GO_REGULATION_OF_CELLULAR_PROTEIN_LOCALIZATION | 521 | 0.3528174 | 1.1677508 | 0 | 0.658038 | 1 | 2647 | tags=23%, list=10%, signal=25% |
| REACTOME_SIGNALING_BY_RHO_GTPASES | 376 | 0.35763887 | 1.1675369 | 0 | 0.5132391 | 1 | 2616 | tags=20%, list=10%, signal=22% |
| GO_METAL_ION_HOMEOSTASIS | 522 | 0.35844293 | 1.1667833 | 0 | 0.65776473 | 1 | 2810 | tags=15%, list=10%, signal=17% |
| GO_HORMONE_TRANSPORT | 265 | 0.37495893 | 1.1660686 | 0 | 0.65754485 | 1 | 2389 | tags=14%, list=9%, signal=15% |
| GO_CELLULAR_RESPONSE_TO_ENDOGENOUS_STIMULUS | 1190 | 0.34592563 | 1.1656951 | 0 | 0.65787876 | 1 | 2801 | tags=19%, list=10%, signal=20% |
| HP_SUBCUTANEOUS_HEMORRHAGE | 177 | 0.41232076 | 1.1655407 | 0.02941177 | 0.6581715 | 1 | 4531 | tags=30%, list=17%, signal=36% |
| GO_CARBOHYDRATE_CATABOLIC_PROCESS | 176 | 0.3931164 | 1.1648729 | 0.02941177 | 0.6581762 | 1 | 3141 | tags=26%, list=12%, signal=29% |
| CUI_TCF21_TARGETS_2_UP | 400 | 0.36425754 | 1.1647502 | 0 | 0.5142082 | 1 | 3317 | tags=26%, list=12%, signal=30% |
| GO_GROWTH_FACTOR_ACTIVITY | 119 | 0.4240627 | 1.1645211 | 0.03448276 | 0.6581875 | 1 | 2160 | tags=12%, list=8%, signal=13% |
| GOBERT_OLIGODENDROCYTE_DIFFERENTIATION_UP | 542 | 0.34869796 | 1.1641202 | 0 | 0.51531106 | 1 | 3392 | tags=28%, list=13%, signal=31% |
| GO_POSITIVE_REGULATION_OF_PHOSPHORUS_METABOLIC_PROCESS | 975 | 0.3464283 | 1.1633576 | 0 | 0.6590058 | 1 | 2604 | tags=17%, list=10%, signal=18% |
| MANALO_HYPOXIA_UP | 192 | 0.3916779 | 1.1620821 | 0.03571429 | 0.51722366 | 1 | 2531 | tags=19%, list=9%, signal=21% |
| GO_NEGATIVE_REGULATION_OF_PHOSPHORUS_METABOLIC_PROCESS | 508 | 0.3637274 | 1.1605744 | 0 | 0.66089416 | 1 | 3104 | tags=26%, list=12%, signal=29% |
| GO_CELL_PROJECTION_ORGANIZATION | 1462 | 0.33690915 | 1.1599092 | 0 | 0.661671 | 1 | 3604 | tags=22%, list=13%, signal=24% |
| HP_ABNORMALITY_OF_THE_HYPOTHALAMUS_PITUITARY_AXIS | 281 | 0.38228604 | 1.1591396 | 0.03571429 | 0.66216123 | 1 | 4069 | tags=24%, list=15%, signal=28% |
| HP_ABNORMAL_CARDIAC_VENTRICLE_MORPHOLOGY | 454 | 0.37675965 | 1.1589074 | 0.05 | 0.66179085 | 1 | 2434 | tags=18%, list=9%, signal=20% |
| GO_RNA_BINDING | 1504 | 0.3487821 | 1.1588277 | 0 | 0.66160315 | 1 | 2824 | tags=29%, list=10%, signal=31% |
| GO_POSITIVE_REGULATION_OF_PROTEIN_MODIFICATION_PROCESS | 1049 | 0.34484464 | 1.1584834 | 0 | 0.66194946 | 1 | 2604 | tags=18%, list=10%, signal=19% |
| GO_REGULATION_OF_CELLULAR_LOCALIZATION | 914 | 0.35175288 | 1.1575662 | 0 | 0.6619937 | 1 | 2747 | tags=21%, list=10%, signal=22% |
| GO_SIGNAL_RELEASE | 507 | 0.35989866 | 1.1574433 | 0.04761905 | 0.66181904 | 1 | 2389 | tags=14%, list=9%, signal=15% |
| GO_NEGATIVE_REGULATION_OF_CELL_DIFFERENTIATION | 614 | 0.35651296 | 1.1573957 | 0 | 0.6617391 | 1 | 2678 | tags=18%, list=10%, signal=19% |
| GO_PLASMA_MEMBRANE_REGION | 1073 | 0.34930748 | 1.1573678 | 0 | 0.66145676 | 1 | 2552 | tags=14%, list=9%, signal=15% |
| REACTOME_RNA_POLYMERASE_II_TRANSCRIPTION | 1031 | 0.3508897 | 1.1562138 | 0 | 0.5242416 | 1 | 2854 | tags=21%, list=11%, signal=23% |
| GO_LYMPHOCYTE_ACTIVATION | 582 | 0.35102013 | 1.1551648 | 0 | 0.66380596 | 1 | 2687 | tags=16%, list=10%, signal=17% |
| GO_NEGATIVE_REGULATION_OF_CELL_CYCLE | 557 | 0.35021037 | 1.1546389 | 0 | 0.6643471 | 1 | 3170 | tags=24%, list=12%, signal=27% |
| GO_INTRINSIC_COMPONENT_OF_PLASMA_MEMBRANE | 1370 | 0.34435862 | 1.153455 | 0 | 0.66476023 | 1 | 3496 | tags=15%, list=13%, signal=17% |
| GO_ANCHORING_JUNCTION | 763 | 0.34513465 | 1.1532079 | 0 | 0.6651058 | 1 | 2800 | tags=21%, list=10%, signal=22% |
| GO_POSITIVE_REGULATION_OF_PROTEIN_METABOLIC_PROCESS | 1462 | 0.34793213 | 1.1524634 | 0 | 0.6659188 | 1 | 2747 | tags=19%, list=10%, signal=20% |
| GO_REGULATION_OF_PROTEIN_MODIFICATION_PROCESS | 1604 | 0.34349155 | 1.1524011 | 0 | 0.66594243 | 1 | 3089 | tags=21%, list=11%, signal=22% |
| GO_REGIONALIZATION | 286 | 0.37142807 | 1.1521533 | 0.03448276 | 0.66583115 | 1 | 3899 | tags=19%, list=14%, signal=22% |
| GO_SYNAPSE | 1239 | 0.33700746 | 1.1518924 | 0 | 0.6654554 | 1 | 2481 | tags=16%, list=9%, signal=17% |
| GO_REGULATION_OF_CELLULAR_COMPONENT_BIOGENESIS | 877 | 0.3502958 | 1.1518742 | 0 | 0.6653286 | 1 | 3440 | tags=24%, list=13%, signal=26% |
| GO_POSITIVE_REGULATION_OF_CELL_DEATH | 634 | 0.35445347 | 1.1512755 | 0 | 0.66590697 | 1 | 2938 | tags=21%, list=11%, signal=23% |
| GO_REGULATION_OF_PHOSPHORUS_METABOLIC_PROCESS | 1561 | 0.34196943 | 1.1511222 | 0 | 0.665488 | 1 | 3299 | tags=21%, list=12%, signal=23% |
| GO_PEPTIDYL_AMINO_ACID_MODIFICATION | 1121 | 0.3396354 | 1.1507767 | 0 | 0.66525406 | 1 | 3413 | tags=23%, list=13%, signal=25% |
| ACEVEDO_METHYLATED_IN_LIVER_CANCER_DN | 579 | 0.35483602 | 1.150613 | 0 | 0.5292279 | 1 | 3847 | tags=20%, list=14%, signal=22% |
| HP_PROGRESSIVE | 246 | 0.37904176 | 1.1500194 | 0.04761905 | 0.6666174 | 1 | 4320 | tags=35%, list=16%, signal=41% |
| GO_CELL_CELL_JUNCTION | 435 | 0.36500257 | 1.1498674 | 0 | 0.6666682 | 1 | 3355 | tags=21%, list=12%, signal=24% |
| GO_PEPTIDE_METABOLIC_PROCESS | 743 | 0.35408652 | 1.1491278 | 0 | 0.66706103 | 1 | 2778 | tags=28%, list=10%, signal=31% |
| GO_NEURON_DIFFERENTIATION | 1278 | 0.3403443 | 1.1470934 | 0 | 0.668046 | 1 | 3102 | tags=18%, list=12%, signal=19% |
| ACEVEDO_LIVER_CANCER_WITH_H3K27ME3_UP | 209 | 0.38709065 | 1.1449803 | 0.04166667 | 0.5330678 | 1 | 3371 | tags=17%, list=13%, signal=20% |
| GO_RNA_PROCESSING | 860 | 0.34726697 | 1.1444324 | 0.04761905 | 0.6718742 | 1 | 3123 | tags=31%, list=12%, signal=33% |
| GRYDER_PAX3FOXO1_ENHANCERS_IN_TADS | 972 | 0.34160042 | 1.143918 | 0 | 0.5347459 | 1 | 2745 | tags=22%, list=10%, signal=24% |
| GO_STRESS_ACTIVATED_PROTEIN_KINASE_SIGNALING_CASCADE | 265 | 0.36712882 | 1.1428572 | 0.04761905 | 0.673105 | 1 | 3069 | tags=24%, list=11%, signal=27% |
| REACTOME_MEMBRANE_TRAFFICKING | 581 | 0.35109022 | 1.1414231 | 0 | 0.5368647 | 1 | 3085 | tags=27%, list=11%, signal=29% |
| GO_REGULATION_OF_SECRETION | 615 | 0.35453105 | 1.1412734 | 0 | 0.6750688 | 1 | 2665 | tags=16%, list=10%, signal=17% |
| GO_DIVALENT_INORGANIC_CATION_TRANSPORT | 404 | 0.3557508 | 1.1407557 | 0.04347826 | 0.67544526 | 1 | 3083 | tags=16%, list=11%, signal=18% |
| GO_GLAND_DEVELOPMENT | 381 | 0.35998368 | 1.1370856 | 0.04 | 0.678948 | 1 | 3437 | tags=22%, list=13%, signal=24% |
| HP_ABNORMAL_FORM_OF_THE_VERTEBRAL_BODIES | 259 | 0.35541674 | 1.134302 | 0.04166667 | 0.68226 | 1 | 3956 | tags=29%, list=15%, signal=34% |
| HP_EMG_ABNORMALITY | 215 | 0.37224877 | 1.1325725 | 0.03333334 | 0.68466836 | 1 | 4030 | tags=27%, list=15%, signal=31% |
| HP_ABNORMALITY_OF_THE_CEREBRAL_CORTEX | 312 | 0.3568987 | 1.1301069 | 0.03448276 | 0.6886132 | 1 | 4136 | tags=31%, list=15%, signal=36% |
| GO_MICROTUBULE_CYTOSKELETON | 1122 | 0.3357529 | 1.12424 | 0 | 0.695065 | 1 | 4061 | tags=28%, list=15%, signal=32% |
| RICKMAN_METASTASIS_UP | 307 | 0.3787162 | 1.1195829 | 0.05 | 0.55996895 | 1 | 3232 | tags=27%, list=12%, signal=30% |
| GO_AMEBOIDAL_TYPE_CELL_MIGRATION | 366 | 0.34466472 | 1.118677 | 0 | 0.6992994 | 1 | 3438 | tags=24%, list=13%, signal=27% |
| GO_MICROTUBULE_ORGANIZING_CENTER | 706 | 0.34135148 | 1.1171465 | 0.04761905 | 0.6999785 | 1 | 4083 | tags=30%, list=15%, signal=34% |
| HP_ABNORMALITY_OF_THE_CEREBRAL_VENTRICLES | 661 | 0.34035647 | 1.1170783 | 0 | 0.70001495 | 1 | 4106 | tags=27%, list=15%, signal=31% |
| ZHENG_BOUND_BY_FOXP3 | 462 | 0.35468593 | 1.1152202 | 0.04166667 | 0.56514305 | 1 | 3710 | tags=25%, list=14%, signal=28% |
| RODRIGUES_THYROID_CARCINOMA_POORLY_DIFFERENTIATED_UP | 599 | 0.3427724 | 1.1124597 | 0.05 | 0.56598884 | 1 | 3454 | tags=30%, list=13%, signal=33% |
| REACTOME_VESICLE_MEDIATED_TRANSPORT | 617 | 0.34800372 | 1.1109885 | 0 | 0.5669993 | 1 | 3085 | tags=25%, list=11%, signal=28% |
| GO_CELLULAR_MACROMOLECULE_LOCALIZATION | 1811 | 0.3288395 | 1.1104726 | 0 | 0.7053845 | 1 | 2887 | tags=23%, list=11%, signal=24% |
| HP_ABNORMALITY_OF_THE_ORBITAL_REGION | 1146 | 0.32355168 | 1.1097747 | 0 | 0.70562834 | 1 | 3527 | tags=23%, list=13%, signal=26% |
| GO_NEGATIVE_REGULATION_OF_BIOSYNTHETIC_PROCESS | 1347 | 0.3193478 | 1.1085303 | 0 | 0.7056196 | 1 | 3316 | tags=21%, list=12%, signal=23% |
| GO_POSITIVE_REGULATION_OF_CELLULAR_COMPONENT_ORGANIZATION | 1108 | 0.31846836 | 1.1085103 | 0 | 0.70552397 | 1 | 3440 | tags=23%, list=13%, signal=25% |
| GO_PROTEIN_MODIFICATION_BY_SMALL_PROTEIN_CONJUGATION | 820 | 0.3305031 | 1.1067306 | 0 | 0.70821613 | 1 | 2917 | tags=22%, list=11%, signal=24% |
| KOINUMA_TARGETS_OF_SMAD2_OR_SMAD3 | 756 | 0.33665302 | 1.1049501 | 0 | 0.5741883 | 1 | 3419 | tags=26%, list=13%, signal=29% |
| GO_INTRACELLULAR_PROTEIN_TRANSPORT | 1080 | 0.32377264 | 1.1037357 | 0 | 0.71042657 | 1 | 2899 | tags=25%, list=11%, signal=27% |
| PATIL_LIVER_CANCER | 602 | 0.3499065 | 1.0998476 | 0 | 0.57836163 | 1 | 3352 | tags=27%, list=12%, signal=31% |
| GO_NEURON_PROJECTION | 1223 | 0.3229002 | 1.0998014 | 0 | 0.71315587 | 1 | 2470 | tags=14%, list=9%, signal=15% |
| GO_CYTOSKELETON_ORGANIZATION | 1237 | 0.3244447 | 1.0989537 | 0 | 0.71383876 | 1 | 3443 | tags=22%, list=13%, signal=24% |
| GO_SOMATODENDRITIC_COMPARTMENT | 791 | 0.3367539 | 1.0953269 | 0 | 0.7166126 | 1 | 2360 | tags=14%, list=9%, signal=15% |
| HP_ABNORMAL_FOOT_MORPHOLOGY | 1049 | 0.33203438 | 1.0871973 | 0 | 0.7211776 | 1 | 3537 | tags=22%, list=13%, signal=25% |
| KINSEY_TARGETS_OF_EWSR1_FLII_FUSION_UP | 1191 | 0.3209983 | 1.0870049 | 0 | 0.59114224 | 1 | 3476 | tags=28%, list=13%, signal=30% |
| GO_NEURON_DEVELOPMENT | 1053 | 0.3302204 | 1.0722506 | 0 | 0.73139817 | 1 | 3090 | tags=18%, list=11%, signal=19% |
| GO_POSITIVE_REGULATION_OF_CELLULAR_BIOSYNTHETIC_PROCESS | 1693 | 0.3129755 | 1.0702195 | 0 | 0.7318674 | 1 | 3451 | tags=23%, list=13%, signal=24% |
| GO_NEGATIVE_REGULATION_OF_RNA_BIOSYNTHETIC_PROCESS | 1075 | 0.31366432 | 1.0627393 | 0 | 0.73648196 | 1 | 3316 | tags=21%, list=12%, signal=23% |
| GO_CELL_CYCLE_PROCESS | 1219 | 0.31257734 | 1.0609087 | 0 | 0.7360851 | 1 | 2552 | tags=19%, list=9%, signal=20% |
| GO_REPRODUCTION | 1156 | 0.3101562 | 1.0544796 | 0 | 0.7410809 | 1 | 3072 | tags=15%, list=11%, signal=16% |
| GO_CELLULAR_NITROGEN_COMPOUND_CATABOLIC_PROCESS | 533 | -0.4100489 | -1.1844522 | 0.02469136 | 1 | 1 | 2915 | tags=26%, list=11%, signal=28% |
| GO_MOTOR_NEURON_APOPTOTIC_PROCESS | 7 | -0.7935962 | -1.2147592 | 0.04081633 | 1 | 1 | 3784 | tags=43%, list=14%, signal=50% |
| GO_RIBOSOMAL_SUBUNIT | 178 | -0.4409977 | -1.2194948 | 0.04545455 | 1 | 1 | 1647 | tags=28%, list=6%, signal=29% |
| GO_PROTEIN_SERINE_THREONINE_KINASE_ACTIVITY | 398 | -0.4221173 | -1.2243685 | 0.01298701 | 1 | 1 | 5348 | tags=36%, list=20%, signal=44% |
| PYEON_CANCER_HEAD_AND_NECK_VS_CERVICAL_UP | 166 | -0.4518232 | -1.2247323 | 0.04918033 | 1 | 1 | 3293 | tags=27%, list=12%, signal=31% |
| BIDUS_METASTASIS_UP | 202 | -0.456815 | -1.2385414 | 0.04477612 | 1 | 1 | 3480 | tags=34%, list=13%, signal=39% |
| GO_COVALENT_CHROMATIN_MODIFICATION | 439 | -0.4239354 | -1.2399583 | 0.01298701 | 1 | 1 | 4095 | tags=33%, list=15%, signal=39% |
| GO_ORGANIC_CYCLIC_COMPOUND_CATABOLIC_PROCESS | 568 | -0.4183804 | -1.2414944 | 0 | 1 | 1 | 2915 | tags=25%, list=11%, signal=28% |
| GO_STRUCTURAL_CONSTITUENT_OF_RIBOSOME | 152 | -0.4680692 | -1.2544777 | 0.02816901 | 1 | 1 | 1647 | tags=32%, list=6%, signal=33% |
| GO_RNA_CATABOLIC_PROCESS | 378 | -0.4364077 | -1.2640532 | 0.02439024 | 1 | 1 | 2915 | tags=31%, list=11%, signal=35% |
| GO_NEGATIVE_REGULATION_OF_LOW_DENSITY_LIPOPROTEIN_PARTICLE_CLEARANCE | 4 | -0.8957225 | -1.2656416 | 0.03389831 | 1 | 1 | 1239 | tags=50%, list=5%, signal=52% |
| GO_CYTOPLASMIC_SIDE_OF_ROUGH_ENDOPLASMIC_RETICULUM_MEMBRANE | 4 | -0.8490005 | -1.2711558 | 0.03703704 | 1 | 1 | 4069 | tags=100%, list=15%, signal=118% |
| GO_REGULATION_OF_MICROTUBULE_CYTOSKELETON_ORGANIZATION | 177 | -0.4656807 | -1.2738751 | 0 | 1 | 1 | 3105 | tags=31%, list=12%, signal=34% |
| HP_DILATATION_OF_THE_BLADDER | 3 | -0.9265946 | -1.283596 | 0.03773585 | 1 | 1 | 215 | tags=33%, list=1%, signal=34% |
| GO_CIA_COMPLEX | 1 | -0.9846566 | -1.2898091 | 0 | 1 | 1 | 415 | tags=100%, list=2%, signal=102% |
| GO_MEDIUM_CHAIN_FATTY_ACID_CATABOLIC_PROCESS | 3 | -0.9394753 | -1.29354 | 0.03389831 | 1 | 1 | 171 | tags=33%, list=1%, signal=34% |
| GO_PIRNA_BIOSYNTHETIC_PROCESS | 3 | -0.9293456 | -1.2967926 | 0.03508772 | 1 | 1 | 1583 | tags=33%, list=6%, signal=35% |
| ROSTY_CERVICAL_CANCER_PROLIFERATION_CLUSTER | 131 | -0.4825459 | -1.2977862 | 0 | 1 | 1 | 3051 | tags=33%, list=11%, signal=37% |
| GO_ESTABLISHMENT_OF_PROTEIN_LOCALIZATION_TO_MEMBRANE | 311 | -0.4529586 | -1.299653 | 0.02739726 | 1 | 1 | 1855 | tags=24%, list=7%, signal=26% |
| STANHILL_HRAS_TRANSFROMATION_UP | 5 | -0.8619046 | -1.3063667 | 0.04761905 | 1 | 1 | 1995 | tags=80%, list=7%, signal=86% |
| REACTOME_HUR_ELAVL1_BINDS_AND_STABILIZES_MRNA | 6 | -0.888418 | -1.3073117 | 0.03846154 | 1 | 1 | 1674 | tags=67%, list=6%, signal=71% |
| WP_MRNA_PROCESSING | 122 | -0.5099845 | -1.3096195 | 0.03448276 | 1 | 1 | 3262 | tags=42%, list=12%, signal=47% |
| GINESTIER_BREAST_CANCER_20Q13_AMPLIFICATION_DN | 164 | -0.4932313 | -1.3112761 | 0.02777778 | 1 | 1 | 5584 | tags=48%, list=21%, signal=60% |
| PID_MAPK_TRK_PATHWAY | 32 | -0.6206496 | -1.3225646 | 0.046875 | 1 | 1 | 4110 | tags=44%, list=15%, signal=52% |
| HP_HEMERALOPIA | 3 | -0.9833894 | -1.3233556 | 0.01694915 | 1 | 1 | 413 | tags=33%, list=2%, signal=34% |
| SMID_BREAST_CANCER_NORMAL_LIKE_DN | 4 | -0.9621154 | -1.334892 | 0.01754386 | 1 | 1 | 525 | tags=25%, list=2%, signal=25% |
| GO_ENDODERMAL_CELL_FATE_SPECIFICATION | 4 | -0.9642248 | -1.3350312 | 0 | 1 | 1 | 377 | tags=25%, list=1%, signal=25% |
| GO_PHOSPHOLIPASE_D_ACTIVITY | 4 | -0.9233193 | -1.335739 | 0.04477612 | 1 | 1 | 501 | tags=50%, list=2%, signal=51% |
| GO_RIBOSOME_ASSEMBLY | 60 | -0.5893483 | -1.3362095 | 0.03278688 | 1 | 1 | 1552 | tags=28%, list=6%, signal=30% |
| PID_E2F_PATHWAY | 68 | -0.5588933 | -1.3433759 | 0.03030303 | 1 | 1 | 4487 | tags=41%, list=17%, signal=49% |
| REACTOME_CIRCADIAN_CLOCK | 67 | -0.539909 | -1.3439696 | 0.01538462 | 1 | 1 | 4038 | tags=43%, list=15%, signal=51% |
| BIOCARTA_FXR_PATHWAY | 4 | -0.8820651 | -1.344305 | 0.04 | 1 | 1 | 1324 | tags=25%, list=5%, signal=26% |
| GO_POSITIVE_REGULATION_OF_OSTEOCLAST_DEVELOPMENT | 4 | -0.9476054 | -1.3446127 | 0.03508772 | 1 | 1 | 779 | tags=25%, list=3%, signal=26% |
| HP_EARLY_ONSET_OF_SEXUAL_MATURATION | 92 | -0.5418536 | -1.3484572 | 0.04411765 | 1 | 1 | 4579 | tags=39%, list=17%, signal=47% |
| GO_MYOSIN_FILAMENT_ORGANIZATION | 4 | -0.9234842 | -1.3524047 | 0.03921569 | 1 | 1 | 684 | tags=25%, list=3%, signal=26% |
| REACTOME_RRNA_PROCESSING | 186 | -0.4940305 | -1.3551751 | 0.01388889 | 1 | 1 | 1977 | tags=27%, list=7%, signal=29% |
| BIOCARTA_WNT_LRP6_PATHWAY | 4 | -0.9271977 | -1.356196 | 0.02941177 | 1 | 1 | 1062 | tags=25%, list=4%, signal=26% |
| GO_ALTERNATIVE_MRNA_SPLICING_VIA_SPLICEOSOME | 82 | -0.5329491 | -1.3562988 | 0.046875 | 1 | 1 | 2984 | tags=40%, list=11%, signal=45% |
| GO_TENDON_DEVELOPMENT | 4 | -0.9428808 | -1.3607091 | 0.02040816 | 1 | 1 | 525 | tags=25%, list=2%, signal=25% |
| BIOCARTA_CREB_PATHWAY | 21 | -0.7398577 | -1.3615994 | 0.03225806 | 1 | 1 | 3934 | tags=62%, list=15%, signal=72% |
| GO_CELLULAR_RESPONSE_TO_ESTRADIOL_STIMULUS | 34 | -0.6135053 | -1.3682019 | 0.04615385 | 1 | 1 | 4406 | tags=26%, list=16%, signal=32% |
| REACTOME_HEME_DEGRADATION | 11 | -0.782193 | -1.3683871 | 0.03571429 | 1 | 1 | 1215 | tags=18%, list=5%, signal=19% |
| REACTOME_ACTIVATION_OF_THE_TFAP2_AP_2_FAMILY_OF_TRANSCRIPTION_FACTORS | 9 | -0.8325867 | -1.371549 | 0.03773585 | 1 | 1 | 2063 | tags=44%, list=8%, signal=48% |
| GO_REGULATION_OF_GENETIC_IMPRINTING | 3 | -0.9628584 | -1.3718219 | 0.01886793 | 1 | 1 | 253 | tags=33%, list=1%, signal=34% |
| HP_ABNORMAL_ERYTHROCYTE_ENZYME_LEVEL | 4 | -0.9469143 | -1.3721542 | 0.01724138 | 1 | 1 | 1421 | tags=75%, list=5%, signal=79% |
| BYSTROEM_CORRELATED_WITH_IL5_DN | 55 | -0.600409 | -1.3730417 | 0.01666667 | 1 | 1 | 4597 | tags=45%, list=17%, signal=55% |
| GO_NEUROFIBRILLARY_TANGLE | 4 | -0.9307365 | -1.3739866 | 0.01886793 | 1 | 1 | 334 | tags=50%, list=1%, signal=51% |
| GO_REGULATION_OF_METANEPHRIC_GLOMERULUS_DEVELOPMENT | 4 | -0.9248325 | -1.3772964 | 0.03703704 | 1 | 1 | 392 | tags=25%, list=1%, signal=25% |
| GO_RETINOIC_ACID_METABOLIC_PROCESS | 15 | -0.748997 | -1.3780645 | 0.03636364 | 1 | 1 | 3104 | tags=20%, list=12%, signal=23% |
| GO_VENTRICULAR_CARDIAC_MUSCLE_CELL_MEMBRANE_REPOLARIZATION | 20 | -0.6840172 | -1.3832995 | 0.03225806 | 1 | 1 | 1689 | tags=20%, list=6%, signal=21% |
| GO_MITOTIC_SPINDLE_MIDZONE | 12 | -0.8032761 | -1.3877869 | 0.05 | 1 | 1 | 2766 | tags=67%, list=10%, signal=74% |
| GO_REGULATION_OF_RNA_SPLICING | 145 | -0.5144236 | -1.3883481 | 0.01515152 | 1 | 1 | 3263 | tags=39%, list=12%, signal=44% |
| HP_PSEUDOEPIPHYSES | 14 | -0.7907894 | -1.3919216 | 0 | 1 | 1 | 2797 | tags=36%, list=10%, signal=40% |
| GO_NUCLEAR_EXPORT_SIGNAL_RECEPTOR_ACTIVITY | 10 | -0.8088028 | -1.3934175 | 0 | 1 | 1 | 3508 | tags=50%, list=13%, signal=57% |
| GO_MICROTUBULE_MINUS_END | 7 | -0.8940748 | -1.3941224 | 0.01818182 | 1 | 1 | 1627 | tags=57%, list=6%, signal=61% |
| TESAR_ALK_TARGETS_HUMAN_ES_4D_DN | 4 | -0.9727817 | -1.3948714 | 0.01754386 | 1 | 1 | 533 | tags=25%, list=2%, signal=26% |
| GO_RESPONSE_TO_LITHIUM_ION | 21 | -0.7032012 | -1.3969393 | 0.046875 | 1 | 1 | 2729 | tags=29%, list=10%, signal=32% |
| GO_NEGATIVE_REGULATION_OF_MEGAKARYOCYTE_DIFFERENTIATION | 3 | -0.9314932 | -1.3977772 | 0.01754386 | 1 | 1 | 1670 | tags=67%, list=6%, signal=71% |
| FLOTHO_PEDIATRIC_ALL_THERAPY_RESPONSE_UP | 48 | -0.6123522 | -1.399563 | 0.03571429 | 1 | 1 | 2155 | tags=35%, list=8%, signal=38% |
| REACTOME_NOTCH3_ACTIVATION_AND_TRANSMISSION_OF_SIGNAL_TO_THE_NUCLEUS | 24 | -0.6688184 | -1.3996891 | 0.04918033 | 1 | 1 | 3949 | tags=42%, list=15%, signal=49% |
| REACTOME_SRP_DEPENDENT_COTRANSLATIONAL_PROTEIN_TARGETING_TO_MEMBRANE | 107 | -0.5628597 | -1.4000064 | 0.01449275 | 1 | 1 | 1647 | tags=44%, list=6%, signal=47% |
| HP_MACRODACTYLY | 4 | -0.9021555 | -1.4004415 | 0.0483871 | 1 | 1 | 2258 | tags=50%, list=8%, signal=55% |
| IRITANI_MAD1_TARGETS_DN | 42 | -0.6357324 | -1.4008296 | 0.03333334 | 1 | 1 | 4119 | tags=55%, list=15%, signal=65% |
| REACTOME_PTK6_REGULATES_RTKS_AND_THEIR_EFFECTORS_AKT1_AND_DOK1 | 8 | -0.8190334 | -1.4015422 | 0.01612903 | 1 | 1 | 1168 | tags=38%, list=4%, signal=39% |
| GO_PROTEIN_SERINE_THREONINE_PHOSPHATASE_ACTIVITY | 68 | -0.5783939 | -1.4025987 | 0 | 1 | 1 | 4774 | tags=41%, list=18%, signal=50% |
| GO_REGULATION_OF_GLUTAMATE_SECRETION | 10 | -0.8028041 | -1.4028279 | 0.03278688 | 1 | 1 | 2800 | tags=30%, list=10%, signal=33% |
| TESAR_ALK_TARGETS_HUMAN_ES_5D_DN | 4 | -0.9727817 | -1.4030329 | 0.01785714 | 1 | 1 | 533 | tags=25%, list=2%, signal=26% |
| HP_EXAGGERATED_CUPID_S_BOW | 7 | -0.8597853 | -1.4036033 | 0.01851852 | 1 | 1 | 1524 | tags=57%, list=6%, signal=61% |
| BIOCARTA_NPC_PATHWAY | 10 | -0.8549341 | -1.4037323 | 0 | 1 | 1 | 1577 | tags=60%, list=6%, signal=64% |
| GO_RNA_DEPENDENT_ATPASE_ACTIVITY | 4 | -0.9346113 | -1.4045845 | 0 | 1 | 1 | 957 | tags=50%, list=4%, signal=52% |
| GO_NEGATIVE_REGULATION_OF_RNA_CATABOLIC_PROCESS | 54 | -0.6263433 | -1.4049952 | 0.01515152 | 1 | 1 | 3690 | tags=39%, list=14%, signal=45% |
| NIKOLSKY_BREAST_CANCER_11Q12_Q14_AMPLICON | 129 | -0.5493577 | -1.4055934 | 0 | 1 | 1 | 6122 | tags=42%, list=23%, signal=54% |
| GO_ESTRADIOL_17_BETA_DEHYDROGENASE_ACTIVITY | 9 | -0.8538384 | -1.4061418 | 0.01724138 | 1 | 1 | 28 | tags=11%, list=0%, signal=11% |
| GO_PROTEIN_TARGETING_TO_MEMBRANE | 187 | -0.5240763 | -1.406606 | 0.01538462 | 1 | 1 | 1830 | tags=30%, list=7%, signal=32% |
| GO_POSITIVE_REGULATION_OF_MICROTUBULE_POLYMERIZATION_OR_DEPOLYMERIZATION | 29 | -0.6550586 | -1.4100478 | 0.03278688 | 1 | 1 | 2588 | tags=45%, list=10%, signal=50% |
| KEGG_FC_EPSILON_RI_SIGNALING_PATHWAY | 66 | -0.571163 | -1.4118251 | 0 | 1 | 1 | 4653 | tags=39%, list=17%, signal=48% |
| WP_MICROGLIA_PATHOGEN_PHAGOCYTOSIS_PATHWAY | 39 | -0.6511584 | -1.4134912 | 0.03703704 | 1 | 1 | 4695 | tags=41%, list=17%, signal=50% |
| JISON_SICKLE_CELL_DISEASE_DN | 166 | -0.5446096 | -1.415373 | 0.01369863 | 1 | 1 | 2288 | tags=31%, list=8%, signal=34% |
| HP_GRANULOCYTOPENIA | 7 | -0.8890197 | -1.4176164 | 0 | 1 | 1 | 1164 | tags=57%, list=4%, signal=60% |
| YAMASHITA_LIVER_CANCER_WITH_EPCAM_UP | 50 | -0.6451539 | -1.4181231 | 0.01492537 | 1 | 1 | 1376 | tags=40%, list=5%, signal=42% |
| GO_NLS_BEARING_PROTEIN_IMPORT_INTO_NUCLEUS | 11 | -0.805006 | -1.4193162 | 0.03333334 | 1 | 1 | 2837 | tags=64%, list=11%, signal=71% |
| REACTOME_TRANSCRIPTIONAL_ACTIVITY_OF_SMAD2_SMAD3_SMAD4_HETEROTRIMER | 43 | -0.6084636 | -1.4195427 | 0.03571429 | 1 | 1 | 3949 | tags=51%, list=15%, signal=60% |
| GO_ATP_GATED_ION_CHANNEL_ACTIVITY | 6 | -0.9471717 | -1.420131 | 0.03278688 | 1 | 1 | 243 | tags=17%, list=1%, signal=17% |
| GO_NUCLEOTIDE_RECEPTOR_ACTIVITY | 15 | -0.7621692 | -1.4222754 | 0.03125 | 1 | 1 | 243 | tags=7%, list=1%, signal=7% |
| GO_VIRAL_GENE_EXPRESSION | 189 | -0.5164642 | -1.423086 | 0 | 1 | 1 | 1908 | tags=34%, list=7%, signal=37% |
| PID_FOXM1_PATHWAY | 38 | -0.6507682 | -1.4240286 | 0.03225806 | 1 | 1 | 3089 | tags=47%, list=11%, signal=53% |
| GO_REGULATION_OF_CHAPERONE_MEDIATED_AUTOPHAGY | 7 | -0.8940032 | -1.4240375 | 0.01785714 | 1 | 1 | 450 | tags=43%, list=2%, signal=44% |
| GO_REGULATION_OF_BONE_DEVELOPMENT | 19 | -0.7348049 | -1.4254489 | 0.03636364 | 1 | 1 | 5308 | tags=42%, list=20%, signal=52% |
| PID_RHODOPSIN_PATHWAY | 20 | -0.7270585 | -1.4311703 | 0.03389831 | 1 | 1 | 1152 | tags=10%, list=4%, signal=10% |
| GO_HISTONE_METHYLTRANSFERASE_ACTIVITY | 53 | -0.582833 | -1.4313889 | 0.03174603 | 1 | 1 | 3988 | tags=38%, list=15%, signal=44% |
| GO_RESPONSE_TO_ARSENIC_CONTAINING_SUBSTANCE | 32 | -0.6561987 | -1.4315014 | 0.04285714 | 1 | 1 | 4882 | tags=59%, list=18%, signal=72% |
| GO_ESTABLISHMENT_OF_PROTEIN_LOCALIZATION_TO_ENDOPLASMIC_RETICULUM | 107 | -0.5800624 | -1.4329901 | 0.01492537 | 1 | 1 | 1647 | tags=44%, list=6%, signal=47% |
| GO_HISTONE_H3_DEACETYLATION | 19 | -0.7307317 | -1.4331164 | 0 | 1 | 1 | 4095 | tags=58%, list=15%, signal=68% |
| GO_GAS_TRANSPORT | 12 | -0.8335788 | -1.433167 | 0.03571429 | 1 | 1 | 2240 | tags=33%, list=8%, signal=36% |
| HP_PRIMARY_CONGENITAL_GLAUCOMA | 4 | -0.9516948 | -1.433929 | 0.01515152 | 1 | 1 | 572 | tags=25%, list=2%, signal=26% |
| GO_RESPONSE_TO_CISPLATIN | 7 | -0.8607677 | -1.4340919 | 0.01851852 | 1 | 1 | 1025 | tags=29%, list=4%, signal=30% |
| HP_SUPRAVALVULAR_AORTIC_STENOSIS | 14 | -0.8052554 | -1.4344482 | 0.046875 | 1 | 1 | 3021 | tags=50%, list=11%, signal=56% |
| BILANGES_SERUM_RESPONSE_TRANSLATION | 29 | -0.6861808 | -1.4384594 | 0 | 1 | 1 | 1302 | tags=45%, list=5%, signal=47% |
| GO_TRANSLATIONAL_INITIATION | 178 | -0.5416728 | -1.4401153 | 0.01515152 | 1 | 1 | 1830 | tags=37%, list=7%, signal=39% |
| GO_NEGATIVE_REGULATION_OF_DENDRITE_DEVELOPMENT | 24 | -0.7045907 | -1.4425297 | 0.05 | 1 | 1 | 4070 | tags=50%, list=15%, signal=59% |
| NIKOLSKY_BREAST_CANCER_12Q24_AMPLICON | 13 | -0.7811024 | -1.4449201 | 0.01694915 | 1 | 1 | 4271 | tags=77%, list=16%, signal=91% |
| BIOCARTA_VDR_PATHWAY | 23 | -0.6762025 | -1.4451 | 0.01587302 | 1 | 1 | 4274 | tags=65%, list=16%, signal=77% |
| GO_REGULATION_OF_DNA_DEMETHYLATION | 6 | -0.9112681 | -1.4455221 | 0 | 1 | 1 | 1874 | tags=67%, list=7%, signal=72% |
| GO_NUCLEAR_TRANSCRIBED_MRNA_CATABOLIC_PROCESS | 195 | -0.5342745 | -1.4484316 | 0 | 1 | 1 | 2764 | tags=37%, list=10%, signal=41% |
| GINESTIER_BREAST_CANCER_ZNF217_AMPLIFIED_DN | 295 | -0.5137137 | -1.4487983 | 0.0131579 | 1 | 1 | 4051 | tags=36%, list=15%, signal=42% |
| GO_NEGATIVE_REGULATION_OF_AXON_EXTENSION | 42 | -0.6790264 | -1.4503129 | 0.03703704 | 1 | 1 | 5273 | tags=48%, list=20%, signal=59% |
| PID_PI3KCI_PATHWAY | 45 | -0.616617 | -1.4505129 | 0 | 1 | 1 | 3934 | tags=40%, list=15%, signal=47% |
| HP_DEVIATION_OF_THE_HALLUX | 48 | -0.630965 | -1.4532263 | 0 | 1 | 1 | 3364 | tags=40%, list=12%, signal=45% |
| NADELLA_PRKAR1A_TARGETS_UP | 7 | -0.9507344 | -1.4559771 | 0 | 1 | 1 | 948 | tags=43%, list=4%, signal=44% |
| GARGALOVIC_RESPONSE_TO_OXIDIZED_PHOSPHOLIPIDS_CYAN_UP | 15 | -0.8147742 | -1.4580159 | 0 | 1 | 1 | 4367 | tags=73%, list=16%, signal=87% |
| GO_PROTEIN_LOCALIZATION_TO_ENDOPLASMIC_RETICULUM | 134 | -0.5703958 | -1.4580317 | 0 | 1 | 1 | 1830 | tags=43%, list=7%, signal=45% |
| HP_OBSESSIVE_COMPULSIVE_TRAIT | 19 | -0.7425914 | -1.4606047 | 0.01587302 | 1 | 1 | 4255 | tags=47%, list=16%, signal=56% |
| GO_RNA_STABILIZATION | 44 | -0.6457348 | -1.46272 | 0.01538462 | 1 | 1 | 3690 | tags=41%, list=14%, signal=47% |
| GO_CELLULAR_RESPONSE_TO_CADMIUM_ION | 27 | -0.7166207 | -1.462865 | 0 | 1 | 1 | 3077 | tags=44%, list=11%, signal=50% |
| GO_RIBOSOMAL_LARGE_SUBUNIT_ASSEMBLY | 27 | -0.7059823 | -1.4655294 | 0.04761905 | 1 | 1 | 1552 | tags=37%, list=6%, signal=39% |
| GROSS_HYPOXIA_VIA_ELK3_ONLY_UP | 33 | -0.6737377 | -1.4666933 | 0.01886793 | 1 | 1 | 5169 | tags=52%, list=19%, signal=64% |
| GO_ALANINE_TRANSMEMBRANE_TRANSPORTER_ACTIVITY | 10 | -0.8375365 | -1.4669573 | 0.01818182 | 1 | 1 | 3287 | tags=60%, list=12%, signal=68% |
| HP_RENAL_DUPLICATION | 23 | -0.7393004 | -1.4687966 | 0.02 | 1 | 1 | 3351 | tags=43%, list=12%, signal=50% |
| GO_AGGREPHAGY | 7 | -0.9067756 | -1.472659 | 0 | 1 | 1 | 1987 | tags=86%, list=7%, signal=93% |
| REACTOME_ACTIVATION_OF_THE_PHOTOTRANSDUCTION_CASCADE | 10 | -0.8750426 | -1.4764524 | 0 | 1 | 1 | 1152 | tags=20%, list=4%, signal=21% |
| NIKOLSKY_BREAST_CANCER_7P22_AMPLICON | 34 | -0.6723628 | -1.477189 | 0.01754386 | 1 | 1 | 1502 | tags=29%, list=6%, signal=31% |
| REACTOME_ACTIVATED_NOTCH1_TRANSMITS_SIGNAL_TO_THE_NUCLEUS | 30 | -0.7036167 | -1.477556 | 0 | 1 | 1 | 4380 | tags=53%, list=16%, signal=64% |
| GO_NEGATIVE_REGULATION_OF_MRNA_SPLICING_VIA_SPLICEOSOME | 20 | -0.7474439 | -1.4783099 | 0.02898551 | 1 | 1 | 2689 | tags=60%, list=10%, signal=67% |
| GO_INTERMEDIATE_FILAMENT_BUNDLE_ASSEMBLY | 5 | -0.9386836 | -1.4784442 | 0.01724138 | 1 | 1 | 334 | tags=20%, list=1%, signal=20% |
| GO_RESPONSE_TO_ISOQUINOLINE_ALKALOID | 27 | -0.7073346 | -1.4785142 | 0.01724138 | 1 | 1 | 5494 | tags=44%, list=20%, signal=56% |
| GO_RAN_GTPASE_BINDING | 33 | -0.7443116 | -1.4791327 | 0 | 1 | 1 | 3651 | tags=61%, list=14%, signal=70% |
| GO_NUCLEAR_IMPORT_SIGNAL_RECEPTOR_ACTIVITY | 18 | -0.7724078 | -1.4846712 | 0.02 | 1 | 1 | 2886 | tags=56%, list=11%, signal=62% |
| BILANGES_SERUM_AND_RAPAMYCIN_SENSITIVE_GENES | 66 | -0.6237451 | -1.4850353 | 0.03571429 | 1 | 1 | 1647 | tags=44%, list=6%, signal=47% |
| WP_NICOTINE_ACTIVITY_ON_DOPAMINERGIC_NEURONS | 15 | -0.8132888 | -1.4886715 | 0.03333334 | 1 | 1 | 4075 | tags=47%, list=15%, signal=55% |
| GO_ALANINE_TRANSPORT | 13 | -0.8122803 | -1.4888629 | 0 | 1 | 1 | 3855 | tags=62%, list=14%, signal=72% |
| HP_SEVERE_FAILURE_TO_THRIVE | 10 | -0.8215586 | -1.4921188 | 0.03225806 | 1 | 1 | 3360 | tags=60%, list=12%, signal=69% |
| REACTOME_ESTROGEN_BIOSYNTHESIS | 5 | -0.9854672 | -1.4961882 | 0 | 0.9709233 | 1 | 28 | tags=20%, list=0%, signal=20% |
| GO_POSITIVE_REGULATION_OF_METANEPHROS_DEVELOPMENT | 12 | -0.8398864 | -1.5116912 | 0 | 1 | 1 | 533 | tags=25%, list=2%, signal=25% |
| WP_PREIMPLANTATION_EMBRYO | 40 | -0.6811491 | -1.5128955 | 0.01515152 | 0.8039714 | 1 | 2275 | tags=30%, list=8%, signal=33% |
| HOLLEMAN_ASPARAGINASE_RESISTANCE_ALL_UP | 18 | -0.7681698 | -1.5138824 | 0 | 0.82488835 | 1 | 2395 | tags=56%, list=9%, signal=61% |
| MEISSNER_BRAIN_HCP_WITH_H3_UNMETHYLATED | 24 | -0.7318395 | -1.5178982 | 0 | 0.81400615 | 1 | 2034 | tags=13%, list=8%, signal=14% |
| IVANOVA_HEMATOPOIESIS_STEM_CELL_SHORT_TERM | 16 | -0.7962878 | -1.5233328 | 0.01754386 | 0.782753 | 1 | 2039 | tags=44%, list=8%, signal=47% |
| GO_ESTROGEN_BIOSYNTHETIC_PROCESS | 11 | -0.8409035 | -1.5248766 | 0.01923077 | 1 | 1 | 1650 | tags=18%, list=6%, signal=19% |
| WP_ALANINE_AND_ASPARTATE_METABOLISM | 11 | -0.8725429 | -1.5256636 | 0.01587302 | 0.79511523 | 1 | 5 | tags=9%, list=0%, signal=9% |
| REACTOME_DOWNREGULATION_OF_TGF_BETA_RECEPTOR_SIGNALING | 25 | -0.7220904 | -1.5260456 | 0.01587302 | 0.82934886 | 1 | 3949 | tags=44%, list=15%, signal=52% |
| REACTOME_EUKARYOTIC_TRANSLATION_INITIATION | 115 | -0.6016311 | -1.5307786 | 0 | 0.8152073 | 1 | 1830 | tags=47%, list=7%, signal=50% |
| REACTOME_INFLUENZA_INFECTION | 149 | -0.5691263 | -1.531072 | 0 | 0.8569491 | 1 | 1647 | tags=40%, list=6%, signal=43% |
| GO_NEGATIVE_REGULATION_OF_MRNA_PROCESSING | 29 | -0.7239965 | -1.5313977 | 0 | 1 | 1 | 2689 | tags=62%, list=10%, signal=69% |
| GO_ANDROGEN_METABOLIC_PROCESS | 21 | -0.7298737 | -1.5346049 | 0.01818182 | 1 | 1 | 1508 | tags=14%, list=6%, signal=15% |
| GO_NEGATIVE_REGULATION_OF_RNA_SPLICING | 24 | -0.7492119 | -1.5367498 | 0.03125 | 1 | 1 | 2689 | tags=58%, list=10%, signal=65% |
| REACTOME_RESPONSE_OF_MTB_TO_PHAGOCYTOSIS | 21 | -0.7634594 | -1.5394546 | 0 | 0.80245763 | 1 | 3949 | tags=67%, list=15%, signal=78% |
| HP_HYPOPLASTIC_TOENAILS | 54 | -0.6477348 | -1.5411717 | 0.01694915 | 1 | 1 | 5623 | tags=52%, list=21%, signal=65% |
| NAKAYAMA_FRA2_TARGETS | 37 | -0.6765255 | -1.5447619 | 0 | 0.7828688 | 1 | 3712 | tags=46%, list=14%, signal=53% |
| HP_CHORIORETINAL_ATROPHY | 18 | -0.7897995 | -1.5455925 | 0.01666667 | 1 | 1 | 3604 | tags=44%, list=13%, signal=51% |
| GO_COTRANSLATIONAL_PROTEIN_TARGETING_TO_MEMBRANE | 95 | -0.5975645 | -1.5505084 | 0 | 1 | 1 | 1647 | tags=48%, list=6%, signal=51% |
| FIGUEROA_AML_METHYLATION_CLUSTER_6_DN | 25 | -0.7475264 | -1.5538584 | 0 | 0.72623116 | 1 | 3196 | tags=28%, list=12%, signal=32% |
| KUUSELO_PANCREATIC_CANCER_19Q13_AMPLIFICATION | 21 | -0.8056273 | -1.5561262 | 0 | 0.74866664 | 1 | 2898 | tags=57%, list=11%, signal=64% |
| ZHENG_RESPONSE_TO_ARSENITE_UP | 8 | -0.9145843 | -1.575688 | 0.01724138 | 0.57827485 | 1 | 1621 | tags=75%, list=6%, signal=80% |
| LY_AGING_MIDDLE_DN | 14 | -0.8305092 | -1.5802056 | 0.01515152 | 0.58283615 | 1 | 1950 | tags=57%, list=7%, signal=62% |
| HOLLEMAN_ASPARAGINASE_RESISTANCE_B_ALL_UP | 25 | -0.7623396 | -1.5811721 | 0 | 0.6295088 | 1 | 2464 | tags=56%, list=9%, signal=62% |
| GO_STEROID_CATABOLIC_PROCESS | 19 | -0.7835944 | -1.5836442 | 0.03636364 | 1 | 1 | 1355 | tags=26%, list=5%, signal=28% |
| GO_ACID_SECRETION | 32 | -0.7077997 | -1.5934876 | 0 | 1 | 1 | 3352 | tags=19%, list=12%, signal=21% |
| REACTOME_NONSENSE_MEDIATED_DECAY_NMD_ | 111 | -0.6369427 | -1.5988094 | 0 | 0.4920955 | 1 | 1830 | tags=48%, list=7%, signal=51% |
| CHNG_MULTIPLE_MYELOMA_HYPERPLOID_UP | 51 | -0.7104101 | -1.5995511 | 0 | 0.5390178 | 1 | 1922 | tags=55%, list=7%, signal=59% |
| KEGG_RIBOSOME | 83 | -0.6753436 | -1.6046773 | 0 | 0.5621041 | 1 | 1647 | tags=55%, list=6%, signal=59% |
| WP_DISRUPTION_OF_POSTSYNAPTIC_SIGNALLING_BY_CNV | 31 | -0.7415741 | -1.6064427 | 0 | 0.6233715 | 1 | 3080 | tags=42%, list=11%, signal=47% |
| PID_FRA_PATHWAY | 33 | -0.7781711 | -1.6231017 | 0 | 0.54125875 | 1 | 3419 | tags=45%, list=13%, signal=52% |
| REACTOME_RESPONSE_OF_EIF2AK4_GCN2_TO_AMINO_ACID_DEFICIENCY | 96 | -0.6491418 | -1.6233639 | 0 | 0.6460482 | 1 | 1845 | tags=52%, list=7%, signal=56% |
| GO_CYTOSOLIC_RIBOSOME | 102 | -0.6238155 | -1.6269062 | 0 | 1 | 1 | 1647 | tags=46%, list=6%, signal=49% |
| GARGALOVIC_RESPONSE_TO_OXIDIZED_PHOSPHOLIPIDS_YELLOW_UP | 28 | -0.7307686 | -1.6303718 | 0.01639344 | 0.71895474 | 1 | 3111 | tags=54%, list=12%, signal=61% |
| GO_NUCLEAR_TRANSCRIBED_MRNA_CATABOLIC_PROCESS_NONSENSE_MEDIATED_DECAY | 117 | -0.6270835 | -1.6328609 | 0 | 1 | 1 | 1830 | tags=44%, list=7%, signal=47% |
| GO_NUCLEOCYTOPLASMIC_CARRIER_ACTIVITY | 28 | -0.7701641 | -1.6411302 | 0.01818182 | 1 | 1 | 3508 | tags=54%, list=13%, signal=62% |
| REACTOME_SELENOAMINO_ACID_METABOLISM | 110 | -0.6384841 | -1.6735749 | 0 | 0.4259952 | 0.9 | 1647 | tags=43%, list=6%, signal=45% |
| REACTOME_EUKARYOTIC_TRANSLATION_ELONGATION | 89 | -0.6720671 | -1.6754417 | 0 | 0.60996455 | 0.89 | 1647 | tags=53%, list=6%, signal=56% |
| GO_CYTOSOLIC_LARGE_RIBOSOMAL_SUBUNIT | 53 | -0.7237657 | -1.6773794 | 0 | 1 | 0.97 | 1647 | tags=55%, list=6%, signal=58% |
| GO_NEGATIVE_REGULATION_OF_MRNA_METABOLIC_PROCESS | 74 | -0.7009368 | -1.7370231 | 0 | 0.47590187 | 0.55 | 2908 | tags=49%, list=11%, signal=54% |
| WP_CYTOPLASMIC_RIBOSOMAL_PROTEINS | 85 | -0.6803463 | -1.7515233 | 0 | 0.2090369 | 0.32 | 1830 | tags=54%, list=7%, signal=58% |
