## Supplementary material for "A cell state specific metabolic vulnerability to GPX4-dependent ferroptosis in glioblastoma": Table S4

**Table 4a: Metaboanalyst differential metabolite analysis**

| **Metabolite** | **FC** | **log2(FC)** | **raw.pval** | **(-LOG10(p))** |
| --- | --- | --- | --- | --- |
| L-Cystine | 16.536 | 4.0476 | 0.025646 | 1.591 |
| Glycerophosphocholine | 5.9037 | 2.5616 | 0.001083 | 2.9653 |
| L-Aspartic acid | 3.3253 | 1.7335 | 0.030928 | 1.5096 |
| Urocanic acid | 3.166 | 1.6626 | 0.011656 | 1.9335 |
| Hypotaurine | 2.9577 | 1.5645 | 0.035305 | 1.4522 |
| Myoinositol | 2.7739 | 1.4719 | 0.002271 | 2.6439 |
| Gluconic acid | 2.4597 | 1.2985 | 0.028316 | 1.548 |
| D-Sedoheptulose 7-phosphate | 2.4063 | 1.2668 | 0.028508 | 1.545 |
| L-Serine | 1.9457 | 0.96025 | 0.00451 | 2.3459 |
| Taurine | 1.8787 | 0.90974 | 0.032291 | 1.4909 |
| Citrulline | 1.8295 | 0.87148 | 0.006163 | 2.2102 |
| Hypoxanthine | 1.7356 | 0.79547 | 0.034369 | 1.4638 |
| Uric acid | 1.6854 | 0.75309 | 0.001157 | 2.9367 |
| Ornithine | 0.65489 | -0.61068 | 0.013647 | 1.865 |
| N-Acetylglutamic acid | 0.61602 | -0.69895 | 0.040823 | 1.3891 |
| L-Alanine | 0.56767 | -0.81689 | 0.040566 | 1.3918 |
| Glucose 6-phosphate | 0.55538 | -0.84847 | 0.042153 | 1.3752 |
| Fructose 6-phosphate | 0.47462 | -1.0751 | 0.003093 | 2.5097 |
| Isovalerylcarnitine | 0.46893 | -1.0926 | 0.026096 | 1.5834 |
| 2-Hydroxyvaleric acid | 0.44325 | -1.1738 | 0.00089 | 3.0505 |
| 4-Trimethylammoniobutanoic acid | 0.44006 | -1.1842 | 0.003452 | 2.4619 |
| Phosphoserine | 0.3949 | -1.3404 | 0.000547 | 3.262 |
| L-Palmitoylcarnitine | 0.3877 | -1.367 | 0.016845 | 1.7735 |
| D-Fructose | 0.25873 | -1.9505 | 3.83E-05 | 4.4173 |
| L-Cystathionine | 0.037631 | -4.7319 | 1.79E-05 | 4.7481 |

**Table 4b: Pathway analysis UP in N1IC**

| **Metabolic Program** | **total** | **expected** | **hits** | **Raw p** | **Holm p** | **FDR** |
| --- | --- | --- | --- | --- | --- | --- |
| Taurine and hypotaurine metabolism | 8 | 0.0677 | 2 | 0.0018 | 0.151 | 0.151 |
| Arginine biosynthesis | 14 | 0.118 | 2 | 0.00568 | 0.472 | 0.208 |
| Histidine metabolism | 16 | 0.135 | 2 | 0.00742 | 0.609 | 0.208 |
| Pentose phosphate pathway | 22 | 0.186 | 2 | 0.0139 | 1 | 0.292 |
| Cysteine and methionine metabolism | 33 | 0.279 | 2 | 0.0301 | 1 | 0.506 |
| Aminoacyl-tRNA biosynthesis | 48 | 0.406 | 2 | 0.0599 | 1 | 0.791 |
| Ascorbate and aldarate metabolism | 8 | 0.0677 | 1 | 0.0659 | 1 | 0.791 |
| Purine metabolism | 65 | 0.55 | 2 | 0.102 | 1 | 1 |
| Nicotinate and nicotinamide metabolism | 15 | 0.127 | 1 | 0.12 | 1 | 1 |
| Pantothenate and CoA biosynthesis | 19 | 0.161 | 1 | 0.15 | 1 | 1 |
| Ether lipid metabolism | 20 | 0.169 | 1 | 0.157 | 1 | 1 |
| beta-Alanine metabolism | 21 | 0.178 | 1 | 0.164 | 1 | 1 |
| Sphingolipid metabolism | 21 | 0.178 | 1 | 0.164 | 1 | 1 |
| Galactose metabolism | 27 | 0.229 | 1 | 0.207 | 1 | 1 |
| Alanine, aspartate and glutamate metabolism | 28 | 0.237 | 1 | 0.213 | 1 | 1 |
| Phosphatidylinositol signaling system | 28 | 0.237 | 1 | 0.213 | 1 | 1 |
| Inositol phosphate metabolism | 30 | 0.254 | 1 | 0.227 | 1 | 1 |
| Glyoxylate and dicarboxylate metabolism | 32 | 0.271 | 1 | 0.24 | 1 | 1 |
| Glycine, serine and threonine metabolism | 33 | 0.279 | 1 | 0.247 | 1 | 1 |
| Glycerophospholipid metabolism | 36 | 0.305 | 1 | 0.266 | 1 | 1 |
| Primary bile acid biosynthesis | 46 | 0.389 | 1 | 0.328 | 1 | 1 |

**Table 4c: Pathway analysis DOWN in N1IC**

| **Metabolic Program** | **total** | **expected** | **hits** | **Raw p** | **Holm p** | **FDR** |
| --- | --- | --- | --- | --- | --- | --- |
| Starch and sucrose metabolism | 18 | 0.105 | 3 | 0.000109 | 0.0091 | 0.00914 |
| Amino sugar and nucleotide sugar metabolism | 37 | 0.217 | 3 | 0.000979 | 0.0813 | 0.0411 |
| Arginine biosynthesis | 14 | 0.082 | 2 | 0.00268 | 0.22 | 0.075 |
| Galactose metabolism | 27 | 0.158 | 2 | 0.00993 | 0.804 | 0.196 |
| Neomycin, kanamycin and gentamicin biosynthesis | 2 | 0.0117 | 1 | 0.0117 | 0.935 | 0.196 |
| Fructose and mannose metabolism | 20 | 0.117 | 1 | 0.112 | 1 | 0.997 |
| Selenocompound metabolism | 20 | 0.117 | 1 | 0.112 | 1 | 0.997 |
| Pentose phosphate pathway | 22 | 0.129 | 1 | 0.122 | 1 | 0.997 |
| Lysine degradation | 25 | 0.146 | 1 | 0.138 | 1 | 0.997 |
| Glycolysis / Gluconeogenesis | 26 | 0.152 | 1 | 0.143 | 1 | 0.997 |
| Alanine, aspartate and glutamate metabolism | 28 | 0.164 | 1 | 0.153 | 1 | 0.997 |
| Glutathione metabolism | 28 | 0.164 | 1 | 0.153 | 1 | 0.997 |
| Inositol phosphate metabolism | 30 | 0.176 | 1 | 0.163 | 1 | 0.997 |
| Glycine, serine and threonine metabolism | 33 | 0.193 | 1 | 0.178 | 1 | 0.997 |
| Cysteine and methionine metabolism | 33 | 0.193 | 1 | 0.178 | 1 | 0.997 |
| Arginine and proline metabolism | 38 | 0.223 | 1 | 0.202 | 1 | 1 |
| Fatty acid degradation | 39 | 0.229 | 1 | 0.207 | 1 | 1 |
| Aminoacyl-tRNA biosynthesis | 48 | 0.281 | 1 | 0.249 | 1 | 1 |

**Table 4d: Integrated scRNAseq-LC-MS UP in N1IC**

| **Metabolic**  **Program** | **Total** | **Expected** | **Hits** | **Raw p** | **(-LOG10(p))** | **Holm adjust** | **FDR** | **Impact** |
| --- | --- | --- | --- | --- | --- | --- | --- | --- |
| Glycolysis or Gluconeogenesis | 61 | 1.3044 | 7 | 0.0002 | 3.6165 | 0.020311 | 0.02 | 0.4 |
| Arginine biosynthesis | 27 | 0.57736 | 4 | 0.0023 | 2.645 | 0.18797 | 0.07 | 0.346 |
| Pyruvate metabolism | 45 | 0.96226 | 5 | 0.0023 | 2.6299 | 0.19228 | 0.07 | 0.5 |
| Taurine and hypotaurine metabolism | 16 | 0.34214 | 3 | 0.0042 | 2.3726 | 0.34346 | 0.09 | 0.733 |
| Glyoxylate and dicarboxylate metabolism | 56 | 1.1975 | 5 | 0.0061 | 2.2118 | 0.49121 | 0.1 | 0.327 |
| Citrate cycle (TCA cycle) | 42 | 0.89811 | 4 | 0.0114 | 1.9429 | 0.90108 | 0.16 | 0.439 |
| Pentose phosphate pathway | 47 | 1.005 | 4 | 0.0168 | 1.7745 | 1 | 0.17 | 0.478 |
| Nitrogen metabolism | 10 | 0.21384 | 2 | 0.0181 | 1.7428 | 1 | 0.17 | 0.444 |
| Synthesis and degradation of ketone bodies | 10 | 0.21384 | 2 | 0.0181 | 1.7428 | 1 | 0.17 | 0.778 |
| Amino sugar and nucleotide sugar metabolism | 81 | 1.7321 | 5 | 0.0276 | 1.5595 | 1 | 0.23 | 0.263 |
| Glutathione metabolism | 56 | 1.1975 | 4 | 0.0301 | 1.5219 | 1 | 0.23 | 0.127 |
| Ether lipid metabolism | 39 | 0.83396 | 3 | 0.0491 | 1.3092 | 1 | 0.32 | 0.184 |
| Linoleic acid metabolism | 17 | 0.36352 | 2 | 0.0497 | 1.3037 | 1 | 0.32 | 0.375 |
| Cysteine and methionine metabolism | 71 | 1.5182 | 4 | 0.063 | 1.2005 | 1 | 0.38 | 0.143 |
| alpha-Linolenic acid metabolism | 22 | 0.47044 | 2 | 0.0789 | 1.1029 | 1 | 0.44 | 0.381 |
| Arachidonic acid metabolism | 79 | 1.6893 | 4 | 0.0862 | 1.0647 | 1 | 0.45 | 0.333 |
| Galactose metabolism | 51 | 1.0906 | 3 | 0.0937 | 1.0283 | 1 | 0.46 | 0.2 |
| Butanoate metabolism | 29 | 0.62013 | 2 | 0.1264 | 0.89844 | 1 | 0.59 | 0.25 |
| Alanine, aspartate and glutamate metabolism | 61 | 1.3044 | 3 | 0.14 | 0.85396 | 1 | 0.62 | 0.283 |
| Histidine metabolism | 32 | 0.68428 | 2 | 0.1484 | 0.8287 | 1 | 0.62 | 0.097 |
| Starch and sucrose metabolism | 37 | 0.79119 | 2 | 0.1866 | 0.7292 | 1 | 0.71 | 0.167 |
| Fructose and mannose metabolism | 37 | 0.79119 | 2 | 0.1866 | 0.7292 | 1 | 0.71 | 0.389 |
| Propanoate metabolism | 48 | 1.0264 | 2 | 0.2741 | 0.56216 | 1 | 0.98 | 0.255 |
| Valine, leucine and isoleucine degradation | 88 | 1.8818 | 3 | 0.2903 | 0.53723 | 1 | 0.98 | 0.115 |
| Purine metabolism | 169 | 3.6138 | 5 | 0.2924 | 0.53409 | 1 | 0.98 | 0.238 |
| Ascorbate and aldarate metabolism | 17 | 0.36352 | 1 | 0.3084 | 0.51092 | 1 | 1 | 0.063 |
| Drug metabolism - other enzymes | 69 | 1.4755 | 2 | 0.4379 | 0.35859 | 1 | 1 | 0.044 |
| Inositol phosphate metabolism | 69 | 1.4755 | 2 | 0.4379 | 0.35859 | 1 | 1 | 0.103 |
| Aminoacyl-tRNA biosynthesis | 74 | 1.5824 | 2 | 0.4743 | 0.32393 | 1 | 1 | 0.041 |
| One carbon pool by folate | 31 | 0.66289 | 1 | 0.4905 | 0.30934 | 1 | 1 | 0.267 |
| Pentose and glucuronate interconversions | 32 | 0.68428 | 1 | 0.5016 | 0.29967 | 1 | 1 | 0.065 |
| Pantothenate and CoA biosynthesis | 34 | 0.72704 | 1 | 0.5229 | 0.28155 | 1 | 1 | 0.03 |
| Glycerolipid metabolism | 35 | 0.74843 | 1 | 0.5333 | 0.27304 | 1 | 1 | 0.088 |
| Terpenoid backbone biosynthesis | 36 | 0.76981 | 1 | 0.5434 | 0.26486 | 1 | 1 | 0.057 |
| Glycerophospholipid metabolism | 86 | 1.839 | 2 | 0.5558 | 0.25511 | 1 | 1 | 0.082 |
| Drug metabolism - cytochrome P450 | 39 | 0.83396 | 1 | 0.5725 | 0.2422 | 1 | 1 | 0.053 |
| Primary bile acid biosynthesis | 90 | 1.9245 | 2 | 0.5809 | 0.23588 | 1 | 1 | 0.067 |
| Nicotinate and nicotinamide metabolism | 43 | 0.9195 | 1 | 0.6085 | 0.21572 | 1 | 1 | 0.024 |
| beta-Alanine metabolism | 44 | 0.94088 | 1 | 0.6171 | 0.20968 | 1 | 1 | 0.047 |
| Biosynthesis of unsaturated fatty acids | 47 | 1.005 | 1 | 0.6416 | 0.19277 | 1 | 1 | 0.478 |
| Sphingolipid metabolism | 58 | 1.2403 | 1 | 0.7189 | 0.14332 | 1 | 1 | 0.018 |
| Folate biosynthesis | 60 | 1.283 | 1 | 0.7311 | 0.13602 | 1 | 1 | 0.034 |
| Glycine, serine and threonine metabolism | 72 | 1.5396 | 1 | 0.7941 | 0.10014 | 1 | 1 | 0.211 |
| Phosphatidylinositol signaling system | 74 | 1.5824 | 1 | 0.8031 | 0.095255 | 1 | 1 | 0.055 |
| Fatty acid elongation | 75 | 1.6038 | 1 | 0.8074 | 0.092911 | 1 | 1 | 0.014 |
| Arginine and proline metabolism | 78 | 1.6679 | 1 | 0.8199 | 0.086248 | 1 | 1 | 0.026 |
| Tyrosine metabolism | 88 | 1.8818 | 1 | 0.856 | 0.067512 | 1 | 1 | 0.023 |
| Pyrimidine metabolism | 99 | 2.117 | 1 | 0.8876 | 0.051784 | 1 | 1 | 0.041 |
| Fatty acid degradation | 102 | 2.1811 | 1 | 0.895 | 0.048198 | 1 | 1 | 0.515 |
| Metabolism of xenobiotics by cytochrome P450 | 117 | 2.5019 | 1 | 0.9252 | 0.033751 | 1 | 1 | 0.224 |

**Table 4e: Integrated scRNAseq-LC-MS DOWN in N1IC**

| **Metabolic**  **Program** | **Total** | **Expected** | **Hits** | **Raw p** | **(-LOG10(p))** | **Holm adjust** | **FDR** | **Impact** |
| --- | --- | --- | --- | --- | --- | --- | --- | --- |
| Lysine degradation | 49 | 1.1916 | 9 | 1.52E-06 | 5.8177 | 0.000128 | 0.000128 | 0.22917 |
| Arginine biosynthesis | 27 | 0.6566 | 4 | 0.003642 | 2.4387 | 0.30225 | 0.15295 | 0.38462 |
| Taurine and hypotaurine metabolism | 16 | 0.3891 | 3 | 0.006105 | 2.2144 | 0.50057 | 0.17093 | 0.4 |
| D-Glutamine and D-glutamate metabolism | 10 | 0.24319 | 2 | 0.023077 | 1.6368 | 1 | 0.48463 | 0.88889 |
| Aminoacyl-tRNA biosynthesis | 74 | 1.7996 | 5 | 0.032002 | 1.4948 | 1 | 0.53763 | 0.12329 |
| Glutathione metabolism | 56 | 1.3618 | 4 | 0.045265 | 1.3442 | 1 | 0.63371 | 0.32727 |
| Glycosaminoglycan biosynthesis - chondroitin sulfate / dermatan sulfate | 18 | 0.43774 | 2 | 0.069327 | 1.1591 | 1 | 0.83192 | 0.23529 |
| Neomycin, kanamycin and gentamicin biosynthesis | 4 | 0.097275 | 1 | 0.09384 | 1.0276 | 1 | 0.98532 | 0.66667 |
| Phenylalanine metabolism | 24 | 0.58365 | 2 | 0.11411 | 0.94268 | 1 | 1 | 0.34783 |
| Glycerophospholipid metabolism | 86 | 2.0914 | 4 | 0.15432 | 0.81157 | 1 | 1 | 0.16471 |
| Alanine, aspartate and glutamate metabolism | 61 | 1.4834 | 3 | 0.18361 | 0.73611 | 1 | 1 | 0.2 |
| Pantothenate and CoA biosynthesis | 34 | 0.82683 | 2 | 0.19956 | 0.69993 | 1 | 1 | 0.24242 |
| Selenocompound metabolism | 35 | 0.85115 | 2 | 0.20853 | 0.68083 | 1 | 1 | 0.14706 |
| Phosphonate and phosphinate metabolism | 10 | 0.24319 | 1 | 0.2186 | 0.66036 | 1 | 1 | 0.22222 |
| Nitrogen metabolism | 10 | 0.24319 | 1 | 0.2186 | 0.66036 | 1 | 1 | 0.44444 |
| Starch and sucrose metabolism | 37 | 0.89979 | 2 | 0.2266 | 0.64475 | 1 | 1 | 0.27778 |
| Inositol phosphate metabolism | 69 | 1.678 | 3 | 0.23453 | 0.6298 | 1 | 1 | 0.073529 |
| Ether lipid metabolism | 39 | 0.94843 | 2 | 0.24478 | 0.61122 | 1 | 1 | 0.15789 |
| Cysteine and methionine metabolism | 71 | 1.7266 | 3 | 0.24767 | 0.60612 | 1 | 1 | 0.11429 |
| beta-Alanine metabolism | 44 | 1.07 | 2 | 0.29043 | 0.53696 | 1 | 1 | 0.093023 |
| Thiamine metabolism | 14 | 0.34046 | 1 | 0.29222 | 0.53429 | 1 | 1 | 0.15385 |
| Ubiquinone and other terpenoid-quinone biosynthesis | 17 | 0.41342 | 1 | 0.34292 | 0.4648 | 1 | 1 | 0.25 |
| Purine metabolism | 169 | 4.1099 | 5 | 0.39342 | 0.40514 | 1 | 1 | 0.25 |
| Glycolysis or Gluconeogenesis | 61 | 1.4834 | 2 | 0.44064 | 0.35591 | 1 | 1 | 0.16667 |
| Drug metabolism - other enzymes | 69 | 1.678 | 2 | 0.50555 | 0.29623 | 1 | 1 | 0.088235 |
| Glycine, serine and threonine metabolism | 72 | 1.7509 | 2 | 0.52865 | 0.27683 | 1 | 1 | 0.084507 |
| Glycosylphosphatidylinositol (GPI)-anchor biosynthesis | 31 | 0.75388 | 1 | 0.53611 | 0.27075 | 1 | 1 | 0.1 |
| One carbon pool by folate | 31 | 0.75388 | 1 | 0.53611 | 0.27075 | 1 | 1 | 0.26667 |
| Phosphatidylinositol signaling system | 74 | 1.7996 | 2 | 0.54365 | 0.26468 | 1 | 1 | 0.082192 |
| Histidine metabolism | 32 | 0.7782 | 1 | 0.54754 | 0.26159 | 1 | 1 | 0.064516 |
| Glycerolipid metabolism | 35 | 0.85115 | 1 | 0.58019 | 0.23643 | 1 | 1 | 0.088235 |
| Fructose and mannose metabolism | 37 | 0.89979 | 1 | 0.60066 | 0.22137 | 1 | 1 | 0.27778 |
| Drug metabolism - cytochrome P450 | 39 | 0.94843 | 1 | 0.62015 | 0.2075 | 1 | 1 | 0.10526 |
| Tyrosine metabolism | 88 | 2.14 | 2 | 0.63944 | 0.1942 | 1 | 1 | 0.22989 |
| Nicotinate and nicotinamide metabolism | 43 | 1.0457 | 1 | 0.65637 | 0.18285 | 1 | 1 | 0.047619 |
| Pyruvate metabolism | 45 | 1.0943 | 1 | 0.67318 | 0.17187 | 1 | 1 | 0.045455 |
| Pentose phosphate pathway | 47 | 1.143 | 1 | 0.68918 | 0.16167 | 1 | 1 | 0.086957 |
| Propanoate metabolism | 48 | 1.1673 | 1 | 0.69689 | 0.15683 | 1 | 1 | 0.042553 |
| Pyrimidine metabolism | 99 | 2.4075 | 2 | 0.70335 | 0.15283 | 1 | 1 | 0.26531 |
| Porphyrin and chlorophyll metabolism | 53 | 1.2889 | 1 | 0.73271 | 0.13507 | 1 | 1 | 0.11538 |
| Sphingolipid metabolism | 58 | 1.4105 | 1 | 0.76437 | 0.1167 | 1 | 1 | 0.035088 |
| Fatty acid biosynthesis | 129 | 3.1371 | 2 | 0.83191 | 0.079922 | 1 | 1 | 1.0391 |
| Fatty acid elongation | 75 | 1.8239 | 1 | 0.84681 | 0.072215 | 1 | 1 | 0.027027 |
| N-Glycan biosynthesis | 77 | 1.8725 | 1 | 0.85441 | 0.068335 | 1 | 1 | 0.026316 |
| Arginine and proline metabolism | 78 | 1.8969 | 1 | 0.85807 | 0.06648 | 1 | 1 | 0.051948 |
| Amino sugar and nucleotide sugar metabolism | 81 | 1.9698 | 1 | 0.86851 | 0.061226 | 1 | 1 | 0.0125 |
| Steroid biosynthesis | 82 | 1.9941 | 1 | 0.87182 | 0.059574 | 1 | 1 | 0.074074 |
| Fatty acid degradation | 102 | 2.4805 | 1 | 0.9232 | 0.034703 | 1 | 1 | 0.029703 |
| Metabolism of xenobiotics by cytochrome P450 | 117 | 2.8453 | 1 | 0.94786 | 0.023256 | 1 | 1 | 0.017241 |
