## Supplementary material for "A cell state specific metabolic vulnerability to GPX4-dependent ferroptosis in glioblastoma": Table S5

**Table 5a: Significantly positively/negatively enriched lipids in N1IC:**

| **Name** | **Fold change**  **(N1IC/p53)** | **log2(FC)** | **FDR-corrected p-value** | **N1IC_1** | **N1IC_2** | **N1IC_3** | **N1IC_4** | **p53_1** | **p53_2** | **p53_3** | **p53_4** |
| --- | --- | --- | --- | --- | --- | --- | --- | --- | --- | --- | --- |
| FA 24:1 | 0.16675 | -2.5842413334775 | 0.0052942 | 2.08404881 | 2.35476107 | 3.80524506 | 3.19812525 | 19.9690674 | 21.4112412 | 12.1271382 | 15.1120782 |
| PC 14:0_14:0 | 0.31922 | -1.64737705173277 | 0.020032 | 178.52734 | 308.747449 | 411.290188 | 252.423216 | 1038.53146 | 1190.40219 | 660.230072 | 716.476655 |
| PC 14:0_16:0 | 0.32972 | -1.60068669491062 | 0.03147 | 1426.83274 | 2461.18036 | 3646.37574 | 2090.09384 | 8908.14279 | 9784.22478 | 4639.97468 | 5857.92226 |
| PC 14:0_16:1 | 0.33026 | -1.59832584913975 | 0.024753 | 543.276598 | 1012.61284 | 1239.09589 | 710.260836 | 3040.54893 | 3630.44523 | 1996.86609 | 1945.63966 |
| PC 14:0_16:1_iso | 0.25205 | -1.98821814054009 | 0.016625 | 163.820964 | 228.647997 | 361.040832 | 234.048245 | 1043.12775 | 1345.21969 | 722.993544 | 806.738339 |
| PC 16:1_16:1 | 0.31597 | -1.66214050770596 | 0.020032 | 278.827601 | 417.520067 | 625.913965 | 335.940534 | 1524.95302 | 1737.27488 | 950.261262 | 1035.40981 |
| PC 28:1 | 0.16375 | -2.61043318823727 | 0.014368 | 14.5164493 | 16.0268049 | 19.1364553 | 18.18346 | 114.845467 | 153.81195 | 69.2419263 | 76.5244136 |
| PC 29:0 | 0.32869 | -1.60520053007202 | 0.033571 | 13.0892491 | 29.7195801 | 37.0486638 | 27.2030404 | 94.8279617 | 105.838266 | 52.5856818 | 72.466672 |
| PC 30:2 | 0.17017 | -2.55495137435048 | 0.014368 | 25.7970355 | 32.1063148 | 46.5536348 | 34.7512195 | 222.515561 | 267.890346 | 158.515188 | 169.156381 |
| PC O-16:0_16:1 | 0.30466 | -1.71472799948985 | 0.021883 | 350.969879 | 652.106773 | 826.081491 | 639.012137 | 2447.04956 | 2660.73636 | 1628.4369 | 1365.13241 |
| PC O-30:0 | 0.27595 | -1.85752120935775 | 0.017287 | 44.294303 | 84.6112824 | 89.4681996 | 79.8041093 | 375.280463 | 284.68222 | 245.716528 | 174.874771 |
| PC O-30:1 | 0.11825 | -3.08008791132269 | 0.01642 | 8.59528415 | 19.9296382 | 24.1691337 | 15.7270415 | 169.212709 | 200.076538 | 103.165554 | 106.17286 |
| PC O-32:2 | 0.22338 | -2.16242807286414 | 0.01642 | 18.6911878 | 31.397359 | 39.1236852 | 27.3528365 | 159.743725 | 163.56091 | 96.0142551 | 102.500098 |
| PC O-34:3 | 0.18809 | -2.41050494636604 | 0.016625 | 26.3997643 | 47.3336581 | 63.9764945 | 46.5977966 | 264.364772 | 357.145542 | 210.087384 | 148.280447 |
| PC O-40:2 | 0.43086 | -1.21470892653951 | 0.022314 | 36.7203058 | 53.3455526 | 71.6045497 | 45.7938338 | 160.846653 | 112.339394 | 102.205883 | 106.124107 |
| PE 16:0_16:1 | 0.29875 | -1.74298938179398 | 0.03006 | 47.145853 | 105.350742 | 116.686182 | 77.6247737 | 344.170815 | 414.387857 | 194.544291 | 207.77419 |
| PE 16:0_18:1 | 0.36043 | -1.47220899758175 | 0.021883 | 49.6711231 | 78.0684998 | 97.6504063 | 66.3841751 | 216.30383 | 285.350572 | 151.466541 | 156.388264 |
| PE 16:1_18:1 | 0.37034 | -1.43307771312007 | 0.029279 | 90.1211244 | 109.29603 | 182.328264 | 132.97629 | 408.767982 | 458.916879 | 221.479352 | 300.710358 |
| PE 18:1_22:3 | 0.32888 | -1.60436681787287 | 0.03742 | 37.3020905 | 56.6282271 | 101.595135 | 63.6790778 | 235.708651 | 276.377477 | 120.343458 | 155.702192 |
| PE 34:0 | 0.24001 | -2.0588335780125 | 0.016625 | 36.9432867 | 66.7157372 | 87.235875 | 51.7054325 | 303.914436 | 323.034852 | 188.11529 | 195.734834 |
| PE O-16:0_18:1 | 0.30491 | -1.71354462832511 | 0.022578 | 10.5358157 | 15.0960452 | 27.631257 | 10.2059883 | 56.7338866 | 67.1904142 | 42.9289839 | 41.3044791 |
| PE P-18:0_22:2 | 0.47411 | -1.07670627195961 | 0.040566 | 45.6771767 | 74.5265277 | 87.4710757 | 61.8737081 | 180.186273 | 166.758861 | 103.078377 | 118.518007 |
| PE P-18:1_22:3 | 0.36313 | -1.461441971515 | 0.027872 | 260.327671 | 396.559122 | 553.360449 | 354.328496 | 1310.51134 | 1418.603 | 740.046182 | 839.461511 |
| PS 18:0_22:3 | 0.25708 | -1.95971071740019 | 0.04894 | 25.1316601 | 39.9360051 | 30.4919348 | 39.9688914 | 141.812295 | 212.669198 | 85.6597042 | 87.0396211 |
| SM d40:0 | 0.26412 | -1.9207345438241 | 0.020351 | 12.3665153 | 17.3705751 | 36.8129542 | 19.865417 | 109.116196 | 95.3872663 | 62.3740272 | 60.3039471 |
| SM d40:1 | 0.34763 | -1.5243755047269 | 0.023967 | 35.5991845 | 58.9973241 | 78.3889717 | 42.9352685 | 185.450464 | 198.658561 | 108.779987 | 128.230662 |
| SM d42:1 | 0.31374 | -1.67235861886881 | 0.020032 | 21.2833371 | 37.6165894 | 44.291633 | 35.3874373 | 140.325471 | 138.234445 | 73.0157631 | 90.1206518 |
| SM d42:2 | 0.31621 | -1.66104510188341 | 0.017287 | 312.729874 | 535.676846 | 658.604486 | 474.479347 | 1847.93196 | 1998.3755 | 1124.42758 | 1295.71752 |
| Cer 18:1;O2/16:0 | 0.40688 | -1.29732472774671 | 0.039594 | 39.6810647 | 58.3270298 | 86.5733677 | 53.6920697 | 168.115321 | 203.530884 | 101.1693 | 112.800422 |
| Cer 18:1;O2/24:0 | 0.3165 | -1.65972259523375 | 0.016625 | 4.96692346 | 8.09282501 | 9.8233388 | 7.54660734 | 26.1714542 | 29.9518099 | 18.4010221 | 21.6213752 |
| Cer 18:1;O2/24:1 | 0.31497 | -1.66671367234576 | 0.020032 | 41.8570808 | 66.3246597 | 96.3055379 | 59.7065084 | 237.196333 | 282.541274 | 152.736387 | 166.313881 |
| Hex-Cer 18:1;O2/24:0 | 0.31137 | -1.68329814512738 | 0.022578 | 7.80567301 | 16.8798476 | 21.1284561 | 11.8970503 | 50.4149319 | 60.2554974 | 34.0029838 | 40.6748591 |
| Hex-Cer 18:1;O2/24:1 | 0.31365 | -1.67277253223871 | 0.020032 | 31.4575468 | 57.9192754 | 74.769619 | 48.2628746 | 199.885776 | 217.381941 | 119.588673 | 140.36488 |
| Hex2Cer 18:1;O2/24:1 | 0.28066 | -1.8331046310146 | 0.016625 | 10.1000024 | 16.8641401 | 19.5132424 | 12.4103069 | 58.9493088 | 70.5105314 | 39.195927 | 41.1626765 |
| DG 18:1_20:1 | 0.22287 | -2.16572566296472 | 0.021883 | 6.57784699 | 15.4183787 | 25.9896998 | 13.7223089 | 94.9577846 | 75.9156369 | 48.309082 | 57.7033893 |
| DG 34:2 | 0.43358 | -1.20562988470454 | 0.03006 | 18.8488306 | 21.1337853 | 31.1663395 | 17.0966295 | 66.8119279 | 57.9503837 | 37.2508444 | 41.5156116 |
| DG 38:4 | 0.36744 | -1.44441940621587 | 0.049263 | 6.44690786 | 10.0313794 | 14.9237153 | 6.83243792 | 25.5769135 | 40.9874214 | 16.506532 | 20.9867732 |
| CL 72:4 | 3.6608 | 1.87215895722894 | 0.023976 | 75.7025937 | 140.868014 | 175.18396 | 130.754802 | 36.8124007 | 50.0920274 | 34.8515549 | 41.3722745 |
| CL 72:5 | 2.1399 | 1.09754337939512 | 0.035205 | 42.8349751 | 70.2186114 | 89.3366784 | 65.076851 | 28.9331162 | 40.8519008 | 28.058518 | 27.1442018 |
| TG 16:0_18:0_18:1 | 3.7938 | 1.9236436253457 | 0.016625 | 300.200228 | 514.841922 | 800.916807 | 448.31959 | 131.191403 | 164.932501 | 125.872597 | 122.119304 |
| TG 16:0_18:1_18:1 | 2.5433 | 1.34670164802286 | 0.039594 | 1014.60814 | 1945.61064 | 2770.84645 | 1575.82134 | 728.247671 | 878.432069 | 627.663056 | 638.630359 |
| TG 18:0_18:0_18:1 | 4.2204 | 2.07737974079701 | 0.01642 | 71.7565346 | 112.1335 | 177.839411 | 104.758731 | 23.0212488 | 32.189505 | 27.6968501 | 27.6240361 |
| TG 18:0_18:1_18:1 | 4.0687 | 2.02456790919802 | 0.016625 | 506.854458 | 946.820606 | 1387.53741 | 767.641757 | 222.144403 | 271.406509 | 186.652183 | 206.775307 |
| TG 18:0_38:3 | 3.0824 | 1.6240540911101 | 0.024581 | 232.98417 | 441.550707 | 651.925855 | 343.617938 | 134.224462 | 162.611749 | 116.731232 | 128.241292 |
| TG 18:0_42:3 | 2.659 | 1.41088377719558 | 0.022578 | 18.2859525 | 31.6025718 | 45.0893214 | 26.6598007 | 11.2106193 | 12.855543 | 10.7880143 | 10.8914106 |
| TG 18:0_48:2 | 2.1494 | 1.10393399105623 | 0.03147 | 12.674809 | 19.2616438 | 28.2907522 | 16.6154513 | 9.04704998 | 8.79521048 | 8.43812372 | 9.46970811 |
| TG 18:1_18:1_18:1 | 2.7212 | 1.44424299422248 | 0.032103 | 682.234757 | 1288.37187 | 1844.82375 | 1045.61478 | 455.660569 | 551.063885 | 378.352701 | 401.25305 |
| TG 20:0_36:2 | 4.1586 | 2.05609792428003 | 0.016625 | 102.777762 | 202.751543 | 277.917137 | 153.786047 | 45.0977163 | 49.4388408 | 42.3836026 | 40.3604411 |

**Table 5b: LION results positive enriched in N1IC:**

| **Term ID** | **Description** | **Annotated** | **p-value** | **FDR q-value** |
| --- | --- | --- | --- | --- |
| LION:0000622 | triacylglycerols [GL0301] | 9 | 9.30E-07 | 1.67E-05 |
| [#0040] TG 16:0_18:0_18:1 |  |  |  |  |
| [#0041] TG 16:0_18:1_18:1 |  |  |  |  |
| [#0042] TG 18:0_18:0_18:1 |  |  |  |  |
| [#0043] TG 18:0_18:1_18:1 |  |  |  |  |
| [#0044] TG 18:0_38:3 |  |  |  |  |
| [#0045] TG 18:0_42:3 |  |  |  |  |
| [#0046] TG 18:0_48:2 |  |  |  |  |
| [#0047] TG 18:1_18:1_18:1 |  |  |  |  |
| [#0048] TG 20:0_36:2 |  |  |  |  |
| LION:0012011 | lipid storage | 9 | 9.30E-07 | 1.67E-05 |
| [#0040] TG 16:0_18:0_18:1 |  |  |  |  |
| [#0041] TG 16:0_18:1_18:1 |  |  |  |  |
| [#0042] TG 18:0_18:0_18:1 |  |  |  |  |
| [#0043] TG 18:0_18:1_18:1 |  |  |  |  |
| [#0044] TG 18:0_38:3 |  |  |  |  |
| [#0045] TG 18:0_42:3 |  |  |  |  |
| [#0046] TG 18:0_48:2 |  |  |  |  |
| [#0047] TG 18:1_18:1_18:1 |  |  |  |  |
| [#0048] TG 20:0_36:2 |  |  |  |  |
| LION:0012084 | lipid droplet | 9 | 9.30E-07 | 1.67E-05 |
| [#0040] TG 16:0_18:0_18:1 |  |  |  |  |
| [#0041] TG 16:0_18:1_18:1 |  |  |  |  |
| [#0042] TG 18:0_18:0_18:1 |  |  |  |  |
| [#0043] TG 18:0_18:1_18:1 |  |  |  |  |
| [#0044] TG 18:0_38:3 |  |  |  |  |
| [#0045] TG 18:0_42:3 |  |  |  |  |
| [#0046] TG 18:0_48:2 |  |  |  |  |
| [#0047] TG 18:1_18:1_18:1 |  |  |  |  |
| [#0048] TG 20:0_36:2 |  |  |  |  |
| LION:0000002 | glycerolipids [GL] | 12 | 4.00E-05 | 5.40E-04 |
| [#0035] DG 18:1_20:1 |  |  |  |  |
| [#0036] DG 34:2 |  |  |  |  |
| [#0037] DG 38:4 |  |  |  |  |
| [#0040] TG 16:0_18:0_18:1 |  |  |  |  |
| [#0041] TG 16:0_18:1_18:1 |  |  |  |  |
| [#0042] TG 18:0_18:0_18:1 |  |  |  |  |
| [#0043] TG 18:0_18:1_18:1 |  |  |  |  |
| [#0044] TG 18:0_38:3 |  |  |  |  |
| [#0045] TG 18:0_42:3 |  |  |  |  |
| [#0046] TG 18:0_48:2 |  |  |  |  |
| [#0047] TG 18:1_18:1_18:1 |  |  |  |  |
| [#0048] TG 20:0_36:2 |  |  |  |  |
| LION:0000094 | headgroup with neutral charge | 13 | 0.00014 | 1.51E-03 |
| [#0034] Hex2Cer 18:1;O2/24:1 | |  |  |  |
| [#0035] DG 18:1_20:1 |  |  |  |  |
| [#0036] DG 34:2 |  |  |  |  |
| [#0037] DG 38:4 |  |  |  |  |
| [#0040] TG 16:0_18:0_18:1 |  |  |  |  |
| [#0041] TG 16:0_18:1_18:1 |  |  |  |  |
| [#0042] TG 18:0_18:0_18:1 |  |  |  |  |
| [#0043] TG 18:0_18:1_18:1 |  |  |  |  |
| [#0044] TG 18:0_38:3 |  |  |  |  |
| [#0045] TG 18:0_42:3 |  |  |  |  |
| [#0046] TG 18:0_48:2 |  |  |  |  |
| [#0047] TG 18:1_18:1_18:1 |  |  |  |  |
| [#0048] TG 20:0_36:2 |  |  |  |  |
| LION:0002921 | C18:0 | 5 | 0.03416 | 3.07E-01 |
| [#0022] PE P-18:0_22:2 |  |  |  |  |
| [#0024] PS 18:0_22:3 |  |  |  |  |
| [#0040] TG 16:0_18:0_18:1 |  |  |  |  |
| [#0042] TG 18:0_18:0_18:1 |  |  |  |  |
| [#0043] TG 18:0_18:1_18:1 |  |  |  |  |
| LION:0000100 | fatty acid with 18 carbons or less | 26 | 0.12797 | 9.15E-01 |
| [#0002] PC 14:0_14:0 |  |  |  |  |
| [#0003] PC 14:0_16:0 |  |  |  |  |
| [#0004] PC 14:0_16:1 |  |  |  |  |
| [#0005] PC 14:0_16:1_iso |  |  |  |  |
| [#0006] PC 16:1_16:1 |  |  |  |  |
| [#0010] PC O-16:0_16:1 |  |  |  |  |
| [#0016] PE 16:0_16:1 |  |  |  |  |
| [#0017] PE 16:0_18:1 |  |  |  |  |
| [#0018] PE 16:1_18:1 |  |  |  |  |
| [#0019] PE 18:1_22:3 |  |  |  |  |
| [#0021] PE O-16:0_18:1 |  |  |  |  |
| [#0022] PE P-18:0_22:2 |  |  |  |  |
| [#0023] PE P-18:1_22:3 |  |  |  |  |
| [#0024] PS 18:0_22:3 |  |  |  |  |
| [#0029] Cer 18:1;O2/16:0 |  |  |  |  |
| [#0030] Cer 18:1;O2/24:0 |  |  |  |  |
| [#0031] Cer 18:1;O2/24:1 |  |  |  |  |
| [#0032] Hex-Cer 18:1;O2/24:0 | |  |  |  |
| [#0033] Hex-Cer 18:1;O2/24:1 | |  |  |  |
| [#0034] Hex2Cer 18:1;O2/24:1 | |  |  |  |
| [#0035] DG 18:1_20:1 |  |  |  |  |
| [#0040] TG 16:0_18:0_18:1 |  |  |  |  |
| [#0041] TG 16:0_18:1_18:1 |  |  |  |  |
| [#0042] TG 18:0_18:0_18:1 |  |  |  |  |
| [#0043] TG 18:0_18:1_18:1 |  |  |  |  |
| [#0047] TG 18:1_18:1_18:1 |  |  |  |  |
| LION:0002922 | C18:1 | 17 | 0.1355 | 9.15E-01 |
| [#0017] PE 16:0_18:1 |  |  |  |  |
| [#0018] PE 16:1_18:1 |  |  |  |  |
| [#0019] PE 18:1_22:3 |  |  |  |  |
| [#0021] PE O-16:0_18:1 |  |  |  |  |
| [#0023] PE P-18:1_22:3 |  |  |  |  |
| [#0029] Cer 18:1;O2/16:0 |  |  |  |  |
| [#0030] Cer 18:1;O2/24:0 |  |  |  |  |
| [#0031] Cer 18:1;O2/24:1 |  |  |  |  |
| [#0032] Hex-Cer 18:1;O2/24:0 | |  |  |  |
| [#0033] Hex-Cer 18:1;O2/24:1 | |  |  |  |
| [#0034] Hex2Cer 18:1;O2/24:1 | |  |  |  |
| [#0035] DG 18:1_20:1 |  |  |  |  |
| [#0040] TG 16:0_18:0_18:1 |  |  |  |  |
| [#0041] TG 16:0_18:1_18:1 |  |  |  |  |
| [#0042] TG 18:0_18:0_18:1 |  |  |  |  |
| [#0043] TG 18:0_18:1_18:1 |  |  |  |  |
| [#0047] TG 18:1_18:1_18:1 |  |  |  |  |
| LION:0002948 | fatty acid with 16-18 carbons | 25 | 0.17145 | 9.55E-01 |
| [#0003] PC 14:0_16:0 |  |  |  |  |
| [#0004] PC 14:0_16:1 |  |  |  |  |
| [#0005] PC 14:0_16:1_iso |  |  |  |  |
| [#0006] PC 16:1_16:1 |  |  |  |  |
| [#0010] PC O-16:0_16:1 |  |  |  |  |
| [#0016] PE 16:0_16:1 |  |  |  |  |
| [#0017] PE 16:0_18:1 |  |  |  |  |
| [#0018] PE 16:1_18:1 |  |  |  |  |
| [#0019] PE 18:1_22:3 |  |  |  |  |
| [#0021] PE O-16:0_18:1 |  |  |  |  |
| [#0022] PE P-18:0_22:2 |  |  |  |  |
| [#0023] PE P-18:1_22:3 |  |  |  |  |
| [#0024] PS 18:0_22:3 |  |  |  |  |
| [#0029] Cer 18:1;O2/16:0 |  |  |  |  |
| [#0030] Cer 18:1;O2/24:0 |  |  |  |  |
| [#0031] Cer 18:1;O2/24:1 |  |  |  |  |
| [#0032] Hex-Cer 18:1;O2/24:0 | |  |  |  |
| [#0033] Hex-Cer 18:1;O2/24:1 | |  |  |  |
| [#0034] Hex2Cer 18:1;O2/24:1 | |  |  |  |
| [#0035] DG 18:1_20:1 |  |  |  |  |
| [#0040] TG 16:0_18:0_18:1 |  |  |  |  |
| [#0041] TG 16:0_18:1_18:1 |  |  |  |  |
| [#0042] TG 18:0_18:0_18:1 |  |  |  |  |
| [#0043] TG 18:0_18:1_18:1 |  |  |  |  |
| [#0047] TG 18:1_18:1_18:1 |  |  |  |  |
| LION:0080969 | low bilayer thickness | 4 | 0.21977 | 9.55E-01 |
| [#0003] PC 14:0_16:0 |  |  |  |  |
| [#0004] PC 14:0_16:1 |  |  |  |  |
| [#0006] PC 16:1_16:1 |  |  |  |  |
| [#0008] PC 29:0 |  |  |  |  |
| LION:0002957 | fatty acid with 18 carbons | 19 | 0.22237 | 9.55E-01 |
| [#0017] PE 16:0_18:1 |  |  |  |  |
| [#0018] PE 16:1_18:1 |  |  |  |  |
| [#0019] PE 18:1_22:3 |  |  |  |  |
| [#0021] PE O-16:0_18:1 |  |  |  |  |
| [#0022] PE P-18:0_22:2 |  |  |  |  |
| [#0023] PE P-18:1_22:3 |  |  |  |  |
| [#0024] PS 18:0_22:3 |  |  |  |  |
| [#0029] Cer 18:1;O2/16:0 |  |  |  |  |
| [#0030] Cer 18:1;O2/24:0 |  |  |  |  |
| [#0031] Cer 18:1;O2/24:1 |  |  |  |  |
| [#0032] Hex-Cer 18:1;O2/24:0 | |  |  |  |
| [#0033] Hex-Cer 18:1;O2/24:1 | |  |  |  |
| [#0034] Hex2Cer 18:1;O2/24:1 | |  |  |  |
| [#0035] DG 18:1_20:1 |  |  |  |  |
| [#0040] TG 16:0_18:0_18:1 |  |  |  |  |
| [#0041] TG 16:0_18:1_18:1 |  |  |  |  |
| [#0042] TG 18:0_18:0_18:1 |  |  |  |  |
| [#0043] TG 18:0_18:1_18:1 |  |  |  |  |
| [#0047] TG 18:1_18:1_18:1 |  |  |  |  |
| LION:0080978 | average lateral diffusion | 6 | 0.26203 | 9.55E-01 |
| [#0002] PC 14:0_14:0 |  |  |  |  |
| [#0003] PC 14:0_16:0 |  |  |  |  |
| [#0008] PC 29:0 |  |  |  |  |
| [#0016] PE 16:0_16:1 |  |  |  |  |
| [#0018] PE 16:1_18:1 |  |  |  |  |
| [#0019] PE 18:1_22:3 |  |  |  |  |
| LION:0000464 | negative intrinsic curvature | 16 | 0.2636 | 9.55E-01 |
| [#0016] PE 16:0_16:1 |  |  |  |  |
| [#0017] PE 16:0_18:1 |  |  |  |  |
| [#0018] PE 16:1_18:1 |  |  |  |  |
| [#0019] PE 18:1_22:3 |  |  |  |  |
| [#0020] PE 34:0 |  |  |  |  |
| [#0021] PE O-16:0_18:1 |  |  |  |  |
| [#0022] PE P-18:0_22:2 |  |  |  |  |
| [#0023] PE P-18:1_22:3 |  |  |  |  |
| [#0029] Cer 18:1;O2/16:0 |  |  |  |  |
| [#0030] Cer 18:1;O2/24:0 |  |  |  |  |
| [#0031] Cer 18:1;O2/24:1 |  |  |  |  |
| [#0035] DG 18:1_20:1 |  |  |  |  |
| [#0036] DG 34:2 |  |  |  |  |
| [#0037] DG 38:4 |  |  |  |  |
| [#0038] CL 72:4 |  |  |  |  |
| [#0039] CL 72:5 |  |  |  |  |
| LION:0002966 | fatty acid with less than 2 double bonds | 27 | 0.27792 | 9.55E-01 |
| [#0001] FA 24:1 |  |  |  |  |
| [#0002] PC 14:0_14:0 |  |  |  |  |
| [#0003] PC 14:0_16:0 |  |  |  |  |
| [#0004] PC 14:0_16:1 |  |  |  |  |
| [#0005] PC 14:0_16:1_iso |  |  |  |  |
| [#0006] PC 16:1_16:1 |  |  |  |  |
| [#0010] PC O-16:0_16:1 |  |  |  |  |
| [#0016] PE 16:0_16:1 |  |  |  |  |
| [#0017] PE 16:0_18:1 |  |  |  |  |
| [#0018] PE 16:1_18:1 |  |  |  |  |
| [#0019] PE 18:1_22:3 |  |  |  |  |
| [#0021] PE O-16:0_18:1 |  |  |  |  |
| [#0022] PE P-18:0_22:2 |  |  |  |  |
| [#0023] PE P-18:1_22:3 |  |  |  |  |
| [#0024] PS 18:0_22:3 |  |  |  |  |
| [#0029] Cer 18:1;O2/16:0 |  |  |  |  |
| [#0030] Cer 18:1;O2/24:0 |  |  |  |  |
| [#0031] Cer 18:1;O2/24:1 |  |  |  |  |
| [#0032] Hex-Cer 18:1;O2/24:0 | |  |  |  |
| [#0033] Hex-Cer 18:1;O2/24:1 | |  |  |  |
| [#0034] Hex2Cer 18:1;O2/24:1 | |  |  |  |
| [#0035] DG 18:1_20:1 |  |  |  |  |
| [#0040] TG 16:0_18:0_18:1 |  |  |  |  |
| [#0041] TG 16:0_18:1_18:1 |  |  |  |  |
| [#0042] TG 18:0_18:0_18:1 |  |  |  |  |
| [#0043] TG 18:0_18:1_18:1 |  |  |  |  |
| [#0047] TG 18:1_18:1_18:1 |  |  |  |  |
| LION:0002969 | monounsaturated fatty acid | 23 | 0.28292 | 9.55E-01 |
| [#0001] FA 24:1 |  |  |  |  |
| [#0004] PC 14:0_16:1 |  |  |  |  |
| [#0005] PC 14:0_16:1_iso |  |  |  |  |
| [#0006] PC 16:1_16:1 |  |  |  |  |
| [#0010] PC O-16:0_16:1 |  |  |  |  |
| [#0016] PE 16:0_16:1 |  |  |  |  |
| [#0017] PE 16:0_18:1 |  |  |  |  |
| [#0018] PE 16:1_18:1 |  |  |  |  |
| [#0019] PE 18:1_22:3 |  |  |  |  |
| [#0021] PE O-16:0_18:1 |  |  |  |  |
| [#0023] PE P-18:1_22:3 |  |  |  |  |
| [#0029] Cer 18:1;O2/16:0 |  |  |  |  |
| [#0030] Cer 18:1;O2/24:0 |  |  |  |  |
| [#0031] Cer 18:1;O2/24:1 |  |  |  |  |
| [#0032] Hex-Cer 18:1;O2/24:0 | |  |  |  |
| [#0033] Hex-Cer 18:1;O2/24:1 | |  |  |  |
| [#0034] Hex2Cer 18:1;O2/24:1 | |  |  |  |
| [#0035] DG 18:1_20:1 |  |  |  |  |
| [#0040] TG 16:0_18:0_18:1 |  |  |  |  |
| [#0041] TG 16:0_18:1_18:1 |  |  |  |  |
| [#0042] TG 18:0_18:0_18:1 |  |  |  |  |
| [#0043] TG 18:0_18:1_18:1 |  |  |  |  |
| [#0047] TG 18:1_18:1_18:1 |  |  |  |  |
| LION:0012081 | mitochondrion | 10 | 0.2917 | 9.55E-01 |
| [#0016] PE 16:0_16:1 |  |  |  |  |
| [#0017] PE 16:0_18:1 |  |  |  |  |
| [#0018] PE 16:1_18:1 |  |  |  |  |
| [#0019] PE 18:1_22:3 |  |  |  |  |
| [#0020] PE 34:0 |  |  |  |  |
| [#0021] PE O-16:0_18:1 |  |  |  |  |
| [#0022] PE P-18:0_22:2 |  |  |  |  |
| [#0023] PE P-18:1_22:3 |  |  |  |  |
| [#0038] CL 72:4 |  |  |  |  |
| [#0039] CL 72:5 |  |  |  |  |
| LION:0002882 | C16:0 | 7 | 0.30052 | 9.55E-01 |
| [#0003] PC 14:0_16:0 |  |  |  |  |
| [#0010] PC O-16:0_16:1 |  |  |  |  |
| [#0016] PE 16:0_16:1 |  |  |  |  |
| [#0017] PE 16:0_18:1 |  |  |  |  |
| [#0021] PE O-16:0_18:1 |  |  |  |  |
| [#0040] TG 16:0_18:0_18:1 |  |  |  |  |
| [#0041] TG 16:0_18:1_18:1 |  |  |  |  |
| LION:0002968 | saturated fatty acid | 14 | 0.33352 | 1.00E+00 |
| [#0002] PC 14:0_14:0 |  |  |  |  |
| [#0003] PC 14:0_16:0 |  |  |  |  |
| [#0004] PC 14:0_16:1 |  |  |  |  |
| [#0005] PC 14:0_16:1_iso |  |  |  |  |
| [#0010] PC O-16:0_16:1 |  |  |  |  |
| [#0016] PE 16:0_16:1 |  |  |  |  |
| [#0017] PE 16:0_18:1 |  |  |  |  |
| [#0021] PE O-16:0_18:1 |  |  |  |  |
| [#0022] PE P-18:0_22:2 |  |  |  |  |
| [#0024] PS 18:0_22:3 |  |  |  |  |
| [#0040] TG 16:0_18:0_18:1 |  |  |  |  |
| [#0041] TG 16:0_18:1_18:1 |  |  |  |  |
| [#0042] TG 18:0_18:0_18:1 |  |  |  |  |
| [#0043] TG 18:0_18:1_18:1 |  |  |  |  |
| LION:0012441 | N-acylsphingosines (ceramides) [SP0201] | 3 | 0.36686 | 1.00E+00 |
| [#0029] Cer 18:1;O2/16:0 |  |  |  |  |
| [#0030] Cer 18:1;O2/24:0 |  |  |  |  |
| [#0031] Cer 18:1;O2/24:1 |  |  |  |  |
| LION:0002955 | fatty acid with 16 carbons | 11 | 0.39695 | 1.00E+00 |
| [#0003] PC 14:0_16:0 |  |  |  |  |
| [#0004] PC 14:0_16:1 |  |  |  |  |
| [#0005] PC 14:0_16:1_iso |  |  |  |  |
| [#0006] PC 16:1_16:1 |  |  |  |  |
| [#0010] PC O-16:0_16:1 |  |  |  |  |
| [#0016] PE 16:0_16:1 |  |  |  |  |
| [#0017] PE 16:0_18:1 |  |  |  |  |
| [#0018] PE 16:1_18:1 |  |  |  |  |
| [#0021] PE O-16:0_18:1 |  |  |  |  |
| [#0040] TG 16:0_18:0_18:1 |  |  |  |  |
| [#0041] TG 16:0_18:1_18:1 |  |  |  |  |
| LION:0000004 | sphingolipids [SP] | 8 | 0.4346 | 1.00E+00 |
| [#0025] SM d40:0 |  |  |  |  |
| [#0026] SM d40:1 |  |  |  |  |
| [#0027] SM d42:1 |  |  |  |  |
| [#0028] SM d42:2 |  |  |  |  |
| [#0029] Cer 18:1;O2/16:0 |  |  |  |  |
| [#0030] Cer 18:1;O2/24:0 |  |  |  |  |
| [#0031] Cer 18:1;O2/24:1 |  |  |  |  |
| [#0034] Hex2Cer 18:1;O2/24:1 | |  |  |  |
| LION:0012009 | lipid-mediated signalling | 6 | 0.48664 | 1.00E+00 |
| [#0029] Cer 18:1;O2/16:0 |  |  |  |  |
| [#0030] Cer 18:1;O2/24:0 |  |  |  |  |
| [#0031] Cer 18:1;O2/24:1 |  |  |  |  |
| [#0035] DG 18:1_20:1 |  |  |  |  |
| [#0036] DG 34:2 |  |  |  |  |
| [#0037] DG 38:4 |  |  |  |  |
| LION:0012082 | plasma membrane | 8 | 0.50916 | 1.00E+00 |
| [#0024] PS 18:0_22:3 |  |  |  |  |
| [#0025] SM d40:0 |  |  |  |  |
| [#0026] SM d40:1 |  |  |  |  |
| [#0027] SM d42:1 |  |  |  |  |
| [#0028] SM d42:2 |  |  |  |  |
| [#0029] Cer 18:1;O2/16:0 |  |  |  |  |
| [#0030] Cer 18:1;O2/24:0 |  |  |  |  |
| [#0031] Cer 18:1;O2/24:1 |  |  |  |  |
| LION:0000093 | headgroup with negative charge | 4 | 0.52721 | 1.00E+00 |
| [#0001] FA 24:1 |  |  |  |  |
| [#0024] PS 18:0_22:3 |  |  |  |  |
| [#0038] CL 72:4 |  |  |  |  |
| [#0039] CL 72:5 |  |  |  |  |
| LION:0002961 | fatty acid with 22 carbons | 4 | 0.52721 | 1.00E+00 |
| [#0019] PE 18:1_22:3 |  |  |  |  |
| [#0022] PE P-18:0_22:2 |  |  |  |  |
| [#0023] PE P-18:1_22:3 |  |  |  |  |
| [#0024] PS 18:0_22:3 |  |  |  |  |
| LION:0002967 | polyunsaturated fatty acid | 4 | 0.52721 | 1.00E+00 |
| [#0019] PE 18:1_22:3 |  |  |  |  |
| [#0022] PE P-18:0_22:2 |  |  |  |  |
| [#0023] PE P-18:1_22:3 |  |  |  |  |
| [#0024] PS 18:0_22:3 |  |  |  |  |
| LION:0000607 | diacylglycerols [GL0201] | 3 | 0.53526 | 1.00E+00 |
| [#0035] DG 18:1_20:1 |  |  |  |  |
| [#0036] DG 34:2 |  |  |  |  |
| [#0037] DG 38:4 |  |  |  |  |
| LION:0000259 | C14:0 | 4 | 0.63234 | 1.00E+00 |
| [#0002] PC 14:0_14:0 |  |  |  |  |
| [#0003] PC 14:0_16:0 |  |  |  |  |
| [#0004] PC 14:0_16:1 |  |  |  |  |
| [#0005] PC 14:0_16:1_iso |  |  |  |  |
| LION:0000011 | glycerophosphoethanolamines [GP02] | 8 | 0.66476 | 1.00E+00 |
| [#0016] PE 16:0_16:1 |  |  |  |  |
| [#0017] PE 16:0_18:1 |  |  |  |  |
| [#0018] PE 16:1_18:1 |  |  |  |  |
| [#0019] PE 18:1_22:3 |  |  |  |  |
| [#0020] PE 34:0 |  |  |  |  |
| [#0021] PE O-16:0_18:1 |  |  |  |  |
| [#0022] PE P-18:0_22:2 |  |  |  |  |
| [#0023] PE P-18:1_22:3 |  |  |  |  |
| LION:0002900 | C16:1 | 6 | 0.68321 | 1.00E+00 |
| [#0004] PC 14:0_16:1 |  |  |  |  |
| [#0005] PC 14:0_16:1_iso |  |  |  |  |
| [#0006] PC 16:1_16:1 |  |  |  |  |
| [#0010] PC O-16:0_16:1 |  |  |  |  |
| [#0016] PE 16:0_16:1 |  |  |  |  |
| [#0018] PE 16:1_18:1 |  |  |  |  |
| LION:0000084 | ceramide phosphocholines (sphingomyelins) [SP0301] | 4 | 0.68469 | 1.00E+00 |
| [#0025] SM d40:0 |  |  |  |  |
| [#0026] SM d40:1 |  |  |  |  |
| [#0027] SM d42:1 |  |  |  |  |
| [#0028] SM d42:2 |  |  |  |  |
| LION:0001740 | above average transition temperature | 7 | 0.70569 | 1.00E+00 |
| [#0003] PC 14:0_16:0 |  |  |  |  |
| [#0017] PE 16:0_18:1 |  |  |  |  |
| [#0020] PE 34:0 |  |  |  |  |
| [#0025] SM d40:0 |  |  |  |  |
| [#0026] SM d40:1 |  |  |  |  |
| [#0027] SM d42:1 |  |  |  |  |
| [#0028] SM d42:2 |  |  |  |  |
| LION:0080973 | below average bilayer thickness | 7 | 0.75498 | 1.00E+00 |
| [#0002] PC 14:0_14:0 |  |  |  |  |
| [#0003] PC 14:0_16:0 |  |  |  |  |
| [#0004] PC 14:0_16:1 |  |  |  |  |
| [#0006] PC 16:1_16:1 |  |  |  |  |
| [#0007] PC 28:1 |  |  |  |  |
| [#0008] PC 29:0 |  |  |  |  |
| [#0009] PC 30:2 |  |  |  |  |
| LION:0000038 | diacylglycerophosphoethanolamines [GP0201] | 5 | 0.78867 | 1.00E+00 |
| [#0016] PE 16:0_16:1 |  |  |  |  |
| [#0017] PE 16:0_18:1 |  |  |  |  |
| [#0018] PE 16:1_18:1 |  |  |  |  |
| [#0019] PE 18:1_22:3 |  |  |  |  |
| [#0020] PE 34:0 |  |  |  |  |
| LION:0001739 | very high transition temperature | 5 | 0.78867 | 1.00E+00 |
| [#0020] PE 34:0 |  |  |  |  |
| [#0025] SM d40:0 |  |  |  |  |
| [#0026] SM d40:1 |  |  |  |  |
| [#0027] SM d42:1 |  |  |  |  |
| [#0028] SM d42:2 |  |  |  |  |
| LION:0002934 | C22:3 | 3 | 0.79852 | 1.00E+00 |
| [#0019] PE 18:1_22:3 |  |  |  |  |
| [#0023] PE P-18:1_22:3 |  |  |  |  |
| [#0024] PS 18:0_22:3 |  |  |  |  |
| LION:0080977 | low lateral diffusion | 3 | 0.79852 | 1.00E+00 |
| [#0016] PE 16:0_16:1 |  |  |  |  |
| [#0017] PE 16:0_18:1 |  |  |  |  |
| [#0024] PS 18:0_22:3 |  |  |  |  |
| LION:0080971 | high bilayer thickness | 4 | 0.8306 | 1.00E+00 |
| [#0017] PE 16:0_18:1 |  |  |  |  |
| [#0019] PE 18:1_22:3 |  |  |  |  |
| [#0020] PE 34:0 |  |  |  |  |
| [#0024] PS 18:0_22:3 |  |  |  |  |
| LION:0080981 | below average lateral diffusion | 4 | 0.8306 | 1.00E+00 |
| [#0016] PE 16:0_16:1 |  |  |  |  |
| [#0017] PE 16:0_18:1 |  |  |  |  |
| [#0020] PE 34:0 |  |  |  |  |
| [#0024] PS 18:0_22:3 |  |  |  |  |
| LION:0002950 | fatty acid with 22-24 carbons | 5 | 0.8496 | 1.00E+00 |
| [#0001] FA 24:1 |  |  |  |  |
| [#0019] PE 18:1_22:3 |  |  |  |  |
| [#0022] PE P-18:0_22:2 |  |  |  |  |
| [#0023] PE P-18:1_22:3 |  |  |  |  |
| [#0024] PS 18:0_22:3 |  |  |  |  |
| LION:0001736 | low transition temperature | 3 | 0.95653 | 1.00E+00 |
| [#0006] PC 16:1_16:1 |  |  |  |  |
| [#0009] PC 30:2 |  |  |  |  |
| [#0018] PE 16:1_18:1 |  |  |  |  |
| LION:0001737 | average transition temperature | 7 | 0.95891 | 1.00E+00 |
| [#0002] PC 14:0_14:0 |  |  |  |  |
| [#0004] PC 14:0_16:1 |  |  |  |  |
| [#0007] PC 28:1 |  |  |  |  |
| [#0008] PC 29:0 |  |  |  |  |
| [#0016] PE 16:0_16:1 |  |  |  |  |
| [#0019] PE 18:1_22:3 |  |  |  |  |
| [#0024] PS 18:0_22:3 |  |  |  |  |
| LION:0000030 | diacylglycerophosphocholines [GP0101] | 8 | 0.96722 | 1.00E+00 |
| [#0002] PC 14:0_14:0 |  |  |  |  |
| [#0003] PC 14:0_16:0 |  |  |  |  |
| [#0004] PC 14:0_16:1 |  |  |  |  |
| [#0005] PC 14:0_16:1_iso |  |  |  |  |
| [#0006] PC 16:1_16:1 |  |  |  |  |
| [#0007] PC 28:1 |  |  |  |  |
| [#0008] PC 29:0 |  |  |  |  |
| [#0009] PC 30:2 |  |  |  |  |
| LION:0002945 | fatty acid with more than 18 carbons | 6 | 0.97647 | 1.00E+00 |
| [#0001] FA 24:1 |  |  |  |  |
| [#0019] PE 18:1_22:3 |  |  |  |  |
| [#0022] PE P-18:0_22:2 |  |  |  |  |
| [#0023] PE P-18:1_22:3 |  |  |  |  |
| [#0024] PS 18:0_22:3 |  |  |  |  |
| [#0035] DG 18:1_20:1 |  |  |  |  |
| LION:0080979 | high lateral diffusion | 5 | 0.99517 | 1.00E+00 |
| [#0002] PC 14:0_14:0 |  |  |  |  |
| [#0004] PC 14:0_16:1 |  |  |  |  |
| [#0006] PC 16:1_16:1 |  |  |  |  |
| [#0007] PC 28:1 |  |  |  |  |
| [#0009] PC 30:2 |  |  |  |  |
| LION:0080968 | very low bilayer thickness | 4 | 0.99622 | 1.00E+00 |
| [#0002] PC 14:0_14:0 |  |  |  |  |
| [#0004] PC 14:0_16:1 |  |  |  |  |
| [#0007] PC 28:1 |  |  |  |  |
| [#0009] PC 30:2 |  |  |  |  |
| LION:0012010 | membrane component | 33 | 0.99924 | 1.00E+00 |
| [#0002] PC 14:0_14:0 |  |  |  |  |
| [#0003] PC 14:0_16:0 |  |  |  |  |
| [#0004] PC 14:0_16:1 |  |  |  |  |
| [#0005] PC 14:0_16:1_iso |  |  |  |  |
| [#0006] PC 16:1_16:1 |  |  |  |  |
| [#0007] PC 28:1 |  |  |  |  |
| [#0008] PC 29:0 |  |  |  |  |
| [#0009] PC 30:2 |  |  |  |  |
| [#0010] PC O-16:0_16:1 |  |  |  |  |
| [#0011] PC O-30:0 |  |  |  |  |
| [#0012] PC O-30:1 |  |  |  |  |
| [#0013] PC O-32:2 |  |  |  |  |
| [#0014] PC O-34:3 |  |  |  |  |
| [#0015] PC O-40:2 |  |  |  |  |
| [#0016] PE 16:0_16:1 |  |  |  |  |
| [#0017] PE 16:0_18:1 |  |  |  |  |
| [#0018] PE 16:1_18:1 |  |  |  |  |
| [#0019] PE 18:1_22:3 |  |  |  |  |
| [#0020] PE 34:0 |  |  |  |  |
| [#0021] PE O-16:0_18:1 |  |  |  |  |
| [#0022] PE P-18:0_22:2 |  |  |  |  |
| [#0023] PE P-18:1_22:3 |  |  |  |  |
| [#0024] PS 18:0_22:3 |  |  |  |  |
| [#0025] SM d40:0 |  |  |  |  |
| [#0026] SM d40:1 |  |  |  |  |
| [#0027] SM d42:1 |  |  |  |  |
| [#0028] SM d42:2 |  |  |  |  |
| [#0034] Hex2Cer 18:1;O2/24:1 | |  |  |  |
| [#0035] DG 18:1_20:1 |  |  |  |  |
| [#0036] DG 34:2 |  |  |  |  |
| [#0037] DG 38:4 |  |  |  |  |
| [#0038] CL 72:4 |  |  |  |  |
| [#0039] CL 72:5 |  |  |  |  |
| LION:0000003 | glycerophospholipids [GP] | 25 | 1 | 1.00E+00 |
| [#0002] PC 14:0_14:0 |  |  |  |  |
| [#0003] PC 14:0_16:0 |  |  |  |  |
| [#0004] PC 14:0_16:1 |  |  |  |  |
| [#0005] PC 14:0_16:1_iso |  |  |  |  |
| [#0006] PC 16:1_16:1 |  |  |  |  |
| [#0007] PC 28:1 |  |  |  |  |
| [#0008] PC 29:0 |  |  |  |  |
| [#0009] PC 30:2 |  |  |  |  |
| [#0010] PC O-16:0_16:1 |  |  |  |  |
| [#0011] PC O-30:0 |  |  |  |  |
| [#0012] PC O-30:1 |  |  |  |  |
| [#0013] PC O-32:2 |  |  |  |  |
| [#0014] PC O-34:3 |  |  |  |  |
| [#0015] PC O-40:2 |  |  |  |  |
| [#0016] PE 16:0_16:1 |  |  |  |  |
| [#0017] PE 16:0_18:1 |  |  |  |  |
| [#0018] PE 16:1_18:1 |  |  |  |  |
| [#0019] PE 18:1_22:3 |  |  |  |  |
| [#0020] PE 34:0 |  |  |  |  |
| [#0021] PE O-16:0_18:1 |  |  |  |  |
| [#0022] PE P-18:0_22:2 |  |  |  |  |
| [#0023] PE P-18:1_22:3 |  |  |  |  |
| [#0024] PS 18:0_22:3 |  |  |  |  |
| [#0038] CL 72:4 |  |  |  |  |
| [#0039] CL 72:5 |  |  |  |  |
| LION:0000010 | glycerophosphocholines [GP01] | 14 | 1 | 1.00E+00 |
| [#0002] PC 14:0_14:0 |  |  |  |  |
| [#0003] PC 14:0_16:0 |  |  |  |  |
| [#0004] PC 14:0_16:1 |  |  |  |  |
| [#0005] PC 14:0_16:1_iso |  |  |  |  |
| [#0006] PC 16:1_16:1 |  |  |  |  |
| [#0007] PC 28:1 |  |  |  |  |
| [#0008] PC 29:0 |  |  |  |  |
| [#0009] PC 30:2 |  |  |  |  |
| [#0010] PC O-16:0_16:1 |  |  |  |  |
| [#0011] PC O-30:0 |  |  |  |  |
| [#0012] PC O-30:1 |  |  |  |  |
| [#0013] PC O-32:2 |  |  |  |  |
| [#0014] PC O-34:3 |  |  |  |  |
| [#0015] PC O-40:2 |  |  |  |  |
| LION:0000031 | 1-alkyl,2-acylglycerophosphocholines [GP0102] | 6 | 1 | 1.00E+00 |
| [#0010] PC O-16:0_16:1 |  |  |  |  |
| [#0011] PC O-30:0 |  |  |  |  |
| [#0012] PC O-30:1 |  |  |  |  |
| [#0013] PC O-32:2 |  |  |  |  |
| [#0014] PC O-34:3 |  |  |  |  |
| [#0015] PC O-40:2 |  |  |  |  |
| LION:0000095 | headgroup with positive charge / zwitter-ion | 26 | 1 | 1.00E+00 |
| [#0002] PC 14:0_14:0 |  |  |  |  |
| [#0003] PC 14:0_16:0 |  |  |  |  |
| [#0004] PC 14:0_16:1 |  |  |  |  |
| [#0005] PC 14:0_16:1_iso |  |  |  |  |
| [#0006] PC 16:1_16:1 |  |  |  |  |
| [#0007] PC 28:1 |  |  |  |  |
| [#0008] PC 29:0 |  |  |  |  |
| [#0009] PC 30:2 |  |  |  |  |
| [#0010] PC O-16:0_16:1 |  |  |  |  |
| [#0011] PC O-30:0 |  |  |  |  |
| [#0012] PC O-30:1 |  |  |  |  |
| [#0013] PC O-32:2 |  |  |  |  |
| [#0014] PC O-34:3 |  |  |  |  |
| [#0015] PC O-40:2 |  |  |  |  |
| [#0016] PE 16:0_16:1 |  |  |  |  |
| [#0017] PE 16:0_18:1 |  |  |  |  |
| [#0018] PE 16:1_18:1 |  |  |  |  |
| [#0019] PE 18:1_22:3 |  |  |  |  |
| [#0020] PE 34:0 |  |  |  |  |
| [#0021] PE O-16:0_18:1 |  |  |  |  |
| [#0022] PE P-18:0_22:2 |  |  |  |  |
| [#0023] PE P-18:1_22:3 |  |  |  |  |
| [#0025] SM d40:0 |  |  |  |  |
| [#0026] SM d40:1 |  |  |  |  |
| [#0027] SM d42:1 |  |  |  |  |
| [#0028] SM d42:2 |  |  |  |  |
| LION:0000465 | neutral intrinsic curvature | 15 | 1 | 1.00E+00 |
| [#0002] PC 14:0_14:0 |  |  |  |  |
| [#0003] PC 14:0_16:0 |  |  |  |  |
| [#0004] PC 14:0_16:1 |  |  |  |  |
| [#0005] PC 14:0_16:1_iso |  |  |  |  |
| [#0006] PC 16:1_16:1 |  |  |  |  |
| [#0007] PC 28:1 |  |  |  |  |
| [#0008] PC 29:0 |  |  |  |  |
| [#0009] PC 30:2 |  |  |  |  |
| [#0010] PC O-16:0_16:1 |  |  |  |  |
| [#0011] PC O-30:0 |  |  |  |  |
| [#0012] PC O-30:1 |  |  |  |  |
| [#0013] PC O-32:2 |  |  |  |  |
| [#0014] PC O-34:3 |  |  |  |  |
| [#0015] PC O-40:2 |  |  |  |  |
| [#0024] PS 18:0_22:3 |  |  |  |  |
| LION:0000467 | contains ether-bond | 7 | 1 | 1.00E+00 |
| [#0010] PC O-16:0_16:1 |  |  |  |  |
| [#0011] PC O-30:0 |  |  |  |  |
| [#0012] PC O-30:1 |  |  |  |  |
| [#0013] PC O-32:2 |  |  |  |  |
| [#0014] PC O-34:3 |  |  |  |  |
| [#0015] PC O-40:2 |  |  |  |  |
| [#0021] PE O-16:0_18:1 |  |  |  |  |
| LION:0012080 | endoplasmic reticulum (ER) | 25 | 1 | 1.00E+00 |
| [#0002] PC 14:0_14:0 |  |  |  |  |
| [#0003] PC 14:0_16:0 |  |  |  |  |
| [#0004] PC 14:0_16:1 |  |  |  |  |
| [#0005] PC 14:0_16:1_iso |  |  |  |  |
| [#0006] PC 16:1_16:1 |  |  |  |  |
| [#0007] PC 28:1 |  |  |  |  |
| [#0008] PC 29:0 |  |  |  |  |
| [#0009] PC 30:2 |  |  |  |  |
| [#0010] PC O-16:0_16:1 |  |  |  |  |
| [#0011] PC O-30:0 |  |  |  |  |
| [#0012] PC O-30:1 |  |  |  |  |
| [#0013] PC O-32:2 |  |  |  |  |
| [#0014] PC O-34:3 |  |  |  |  |
| [#0015] PC O-40:2 |  |  |  |  |
| [#0016] PE 16:0_16:1 |  |  |  |  |
| [#0017] PE 16:0_18:1 |  |  |  |  |
| [#0018] PE 16:1_18:1 |  |  |  |  |
| [#0019] PE 18:1_22:3 |  |  |  |  |
| [#0020] PE 34:0 |  |  |  |  |
| [#0021] PE O-16:0_18:1 |  |  |  |  |
| [#0022] PE P-18:0_22:2 |  |  |  |  |
| [#0023] PE P-18:1_22:3 |  |  |  |  |
| [#0029] Cer 18:1;O2/16:0 |  |  |  |  |
| [#0030] Cer 18:1;O2/24:0 |  |  |  |  |
| [#0031] Cer 18:1;O2/24:1 |  |  |  |  |

**Table 5c: LION results negative enriched in N1IC:**

| **Term ID** | **Description** | **Annotated** | **p-value** | **FDR q-value** |
| --- | --- | --- | --- | --- |
| LION:0012010 | membrane component | 33 | 0.0012 | 0.0432 |
| [#0002] PC 14:0_14:0 |  |  |  |  |
| [#0003] PC 14:0_16:0 |  |  |  |  |
| [#0004] PC 14:0_16:1 |  |  |  |  |
| [#0005] PC 14:0_16:1_iso |  |  |  |  |
| [#0006] PC 16:1_16:1 |  |  |  |  |
| [#0007] PC 28:1 |  |  |  |  |
| [#0008] PC 29:0 |  |  |  |  |
| [#0009] PC 30:2 |  |  |  |  |
| [#0010] PC O-16:0_16:1 |  |  |  |  |
| [#0011] PC O-30:0 |  |  |  |  |
| [#0012] PC O-30:1 |  |  |  |  |
| [#0013] PC O-32:2 |  |  |  |  |
| [#0014] PC O-34:3 |  |  |  |  |
| [#0015] PC O-40:2 |  |  |  |  |
| [#0016] PE 16:0_16:1 |  |  |  |  |
| [#0017] PE 16:0_18:1 |  |  |  |  |
| [#0018] PE 16:1_18:1 |  |  |  |  |
| [#0019] PE 18:1_22:3 |  |  |  |  |
| [#0020] PE 34:0 |  |  |  |  |
| [#0021] PE O-16:0_18:1 |  |  |  |  |
| [#0022] PE P-18:0_22:2 |  |  |  |  |
| [#0023] PE P-18:1_22:3 |  |  |  |  |
| [#0024] PS 18:0_22:3 |  |  |  |  |
| [#0025] SM d40:0 |  |  |  |  |
| [#0026] SM d40:1 |  |  |  |  |
| [#0027] SM d42:1 |  |  |  |  |
| [#0028] SM d42:2 |  |  |  |  |
| [#0034] Hex2Cer 18:1;O2/24:1 | |  |  |  |
| [#0035] DG 18:1_20:1 |  |  |  |  |
| [#0036] DG 34:2 |  |  |  |  |
| [#0037] DG 38:4 |  |  |  |  |
| [#0038] CL 72:4 |  |  |  |  |
| [#0039] CL 72:5 |  |  |  |  |
| LION:0000095 | headgroup with positive charge / zwitter-ion | 26 | 0.0016 | 0.0432 |
| [#0002] PC 14:0_14:0 |  |  |  |  |
| [#0003] PC 14:0_16:0 |  |  |  |  |
| [#0004] PC 14:0_16:1 |  |  |  |  |
| [#0005] PC 14:0_16:1_iso |  |  |  |  |
| [#0006] PC 16:1_16:1 |  |  |  |  |
| [#0007] PC 28:1 |  |  |  |  |
| [#0008] PC 29:0 |  |  |  |  |
| [#0009] PC 30:2 |  |  |  |  |
| [#0010] PC O-16:0_16:1 |  |  |  |  |
| [#0011] PC O-30:0 |  |  |  |  |
| [#0012] PC O-30:1 |  |  |  |  |
| [#0013] PC O-32:2 |  |  |  |  |
| [#0014] PC O-34:3 |  |  |  |  |
| [#0015] PC O-40:2 |  |  |  |  |
| [#0016] PE 16:0_16:1 |  |  |  |  |
| [#0017] PE 16:0_18:1 |  |  |  |  |
| [#0018] PE 16:1_18:1 |  |  |  |  |
| [#0019] PE 18:1_22:3 |  |  |  |  |
| [#0020] PE 34:0 |  |  |  |  |
| [#0021] PE O-16:0_18:1 |  |  |  |  |
| [#0022] PE P-18:0_22:2 |  |  |  |  |
| [#0023] PE P-18:1_22:3 |  |  |  |  |
| [#0025] SM d40:0 |  |  |  |  |
| [#0026] SM d40:1 |  |  |  |  |
| [#0027] SM d42:1 |  |  |  |  |
| [#0028] SM d42:2 |  |  |  |  |
| LION:0012080 | endoplasmic reticulum (ER) | 25 | 0.0038 | 0.0648 |
| [#0002] PC 14:0_14:0 |  |  |  |  |
| [#0003] PC 14:0_16:0 |  |  |  |  |
| [#0004] PC 14:0_16:1 |  |  |  |  |
| [#0005] PC 14:0_16:1_iso |  |  |  |  |
| [#0006] PC 16:1_16:1 |  |  |  |  |
| [#0007] PC 28:1 |  |  |  |  |
| [#0008] PC 29:0 |  |  |  |  |
| [#0009] PC 30:2 |  |  |  |  |
| [#0010] PC O-16:0_16:1 |  |  |  |  |
| [#0011] PC O-30:0 |  |  |  |  |
| [#0012] PC O-30:1 |  |  |  |  |
| [#0013] PC O-32:2 |  |  |  |  |
| [#0014] PC O-34:3 |  |  |  |  |
| [#0015] PC O-40:2 |  |  |  |  |
| [#0016] PE 16:0_16:1 |  |  |  |  |
| [#0017] PE 16:0_18:1 |  |  |  |  |
| [#0018] PE 16:1_18:1 |  |  |  |  |
| [#0019] PE 18:1_22:3 |  |  |  |  |
| [#0020] PE 34:0 |  |  |  |  |
| [#0021] PE O-16:0_18:1 |  |  |  |  |
| [#0022] PE P-18:0_22:2 |  |  |  |  |
| [#0023] PE P-18:1_22:3 |  |  |  |  |
| [#0029] Cer 18:1;O2/16:0 |  |  |  |  |
| [#0030] Cer 18:1;O2/24:0 |  |  |  |  |
| [#0031] Cer 18:1;O2/24:1 |  |  |  |  |
| LION:0000465 | neutral intrinsic curvature | 15 | 0.0048 | 0.0648 |
| [#0002] PC 14:0_14:0 |  |  |  |  |
| [#0003] PC 14:0_16:0 |  |  |  |  |
| [#0004] PC 14:0_16:1 |  |  |  |  |
| [#0005] PC 14:0_16:1_iso |  |  |  |  |
| [#0006] PC 16:1_16:1 |  |  |  |  |
| [#0007] PC 28:1 |  |  |  |  |
| [#0008] PC 29:0 |  |  |  |  |
| [#0009] PC 30:2 |  |  |  |  |
| [#0010] PC O-16:0_16:1 |  |  |  |  |
| [#0011] PC O-30:0 |  |  |  |  |
| [#0012] PC O-30:1 |  |  |  |  |
| [#0013] PC O-32:2 |  |  |  |  |
| [#0014] PC O-34:3 |  |  |  |  |
| [#0015] PC O-40:2 |  |  |  |  |
| [#0024] PS 18:0_22:3 |  |  |  |  |
| LION:0000010 | glycerophosphocholines [GP01] | 14 | 0.009 | 0.0972 |
| [#0002] PC 14:0_14:0 |  |  |  |  |
| [#0003] PC 14:0_16:0 |  |  |  |  |
| [#0004] PC 14:0_16:1 |  |  |  |  |
| [#0005] PC 14:0_16:1_iso |  |  |  |  |
| [#0006] PC 16:1_16:1 |  |  |  |  |
| [#0007] PC 28:1 |  |  |  |  |
| [#0008] PC 29:0 |  |  |  |  |
| [#0009] PC 30:2 |  |  |  |  |
| [#0010] PC O-16:0_16:1 |  |  |  |  |
| [#0011] PC O-30:0 |  |  |  |  |
| [#0012] PC O-30:1 |  |  |  |  |
| [#0013] PC O-32:2 |  |  |  |  |
| [#0014] PC O-34:3 |  |  |  |  |
| [#0015] PC O-40:2 |  |  |  |  |
| LION:0000467 | contains ether-bond | 7 | 0.0111 | 0.0999 |
| [#0010] PC O-16:0_16:1 |  |  |  |  |
| [#0011] PC O-30:0 |  |  |  |  |
| [#0012] PC O-30:1 |  |  |  |  |
| [#0013] PC O-32:2 |  |  |  |  |
| [#0014] PC O-34:3 |  |  |  |  |
| [#0015] PC O-40:2 |  |  |  |  |
| [#0021] PE O-16:0_18:1 |  |  |  |  |
| LION:0000031 | 1-alkyl,2-acylglycerophosphocholines [GP0102] | 6 | 0.0242 | 0.1867 |
| [#0010] PC O-16:0_16:1 |  |  |  |  |
| [#0011] PC O-30:0 |  |  |  |  |
| [#0012] PC O-30:1 |  |  |  |  |
| [#0013] PC O-32:2 |  |  |  |  |
| [#0014] PC O-34:3 |  |  |  |  |
| [#0015] PC O-40:2 |  |  |  |  |
| LION:0000030 | diacylglycerophosphocholines [GP0101] | 8 | 0.0357 | 0.241 |
| [#0002] PC 14:0_14:0 |  |  |  |  |
| [#0003] PC 14:0_16:0 |  |  |  |  |
| [#0004] PC 14:0_16:1 |  |  |  |  |
| [#0005] PC 14:0_16:1_iso |  |  |  |  |
| [#0006] PC 16:1_16:1 |  |  |  |  |
| [#0007] PC 28:1 |  |  |  |  |
| [#0008] PC 29:0 |  |  |  |  |
| [#0009] PC 30:2 |  |  |  |  |
| LION:0000003 | glycerophospholipids [GP] | 25 | 0.052 | 0.2852 |
| [#0002] PC 14:0_14:0 |  |  |  |  |
| [#0003] PC 14:0_16:0 |  |  |  |  |
| [#0004] PC 14:0_16:1 |  |  |  |  |
| [#0005] PC 14:0_16:1_iso |  |  |  |  |
| [#0006] PC 16:1_16:1 |  |  |  |  |
| [#0007] PC 28:1 |  |  |  |  |
| [#0008] PC 29:0 |  |  |  |  |
| [#0009] PC 30:2 |  |  |  |  |
| [#0010] PC O-16:0_16:1 |  |  |  |  |
| [#0011] PC O-30:0 |  |  |  |  |
| [#0012] PC O-30:1 |  |  |  |  |
| [#0013] PC O-32:2 |  |  |  |  |
| [#0014] PC O-34:3 |  |  |  |  |
| [#0015] PC O-40:2 |  |  |  |  |
| [#0016] PE 16:0_16:1 |  |  |  |  |
| [#0017] PE 16:0_18:1 |  |  |  |  |
| [#0018] PE 16:1_18:1 |  |  |  |  |
| [#0019] PE 18:1_22:3 |  |  |  |  |
| [#0020] PE 34:0 |  |  |  |  |
| [#0021] PE O-16:0_18:1 |  |  |  |  |
| [#0022] PE P-18:0_22:2 |  |  |  |  |
| [#0023] PE P-18:1_22:3 |  |  |  |  |
| [#0024] PS 18:0_22:3 |  |  |  |  |
| [#0038] CL 72:4 |  |  |  |  |
| [#0039] CL 72:5 |  |  |  |  |
| LION:0001737 | average transition temperature | 7 | 0.0581 | 0.2852 |
| [#0002] PC 14:0_14:0 |  |  |  |  |
| [#0004] PC 14:0_16:1 |  |  |  |  |
| [#0007] PC 28:1 |  |  |  |  |
| [#0008] PC 29:0 |  |  |  |  |
| [#0016] PE 16:0_16:1 |  |  |  |  |
| [#0019] PE 18:1_22:3 |  |  |  |  |
| [#0024] PS 18:0_22:3 |  |  |  |  |
| LION:0080973 | below average bilayer thickness | 7 | 0.0581 | 0.2852 |
| [#0002] PC 14:0_14:0 |  |  |  |  |
| [#0003] PC 14:0_16:0 |  |  |  |  |
| [#0004] PC 14:0_16:1 |  |  |  |  |
| [#0006] PC 16:1_16:1 |  |  |  |  |
| [#0007] PC 28:1 |  |  |  |  |
| [#0008] PC 29:0 |  |  |  |  |
| [#0009] PC 30:2 |  |  |  |  |
| LION:0001740 | above average transition temperature | 7 | 0.0998 | 0.4491 |
| [#0003] PC 14:0_16:0 |  |  |  |  |
| [#0017] PE 16:0_18:1 |  |  |  |  |
| [#0020] PE 34:0 |  |  |  |  |
| [#0025] SM d40:0 |  |  |  |  |
| [#0026] SM d40:1 |  |  |  |  |
| [#0027] SM d42:1 |  |  |  |  |
| [#0028] SM d42:2 |  |  |  |  |
| LION:0080979 | high lateral diffusion | 5 | 0.144 | 0.5935 |
| [#0002] PC 14:0_14:0 |  |  |  |  |
| [#0004] PC 14:0_16:1 |  |  |  |  |
| [#0006] PC 16:1_16:1 |  |  |  |  |
| [#0007] PC 28:1 |  |  |  |  |
| [#0009] PC 30:2 |  |  |  |  |
| LION:0080981 | below average lateral diffusion | 4 | 0.1599 | 0.5935 |
| [#0016] PE 16:0_16:1 |  |  |  |  |
| [#0017] PE 16:0_18:1 |  |  |  |  |
| [#0020] PE 34:0 |  |  |  |  |
| [#0024] PS 18:0_22:3 |  |  |  |  |
| LION:0001739 | very high transition temperature | 5 | 0.1739 | 0.5935 |
| [#0020] PE 34:0 |  |  |  |  |
| [#0025] SM d40:0 |  |  |  |  |
| [#0026] SM d40:1 |  |  |  |  |
| [#0027] SM d42:1 |  |  |  |  |
| [#0028] SM d42:2 |  |  |  |  |
| LION:0000004 | sphingolipids [SP] | 8 | 0.1953 | 0.5935 |
| [#0025] SM d40:0 |  |  |  |  |
| [#0026] SM d40:1 |  |  |  |  |
| [#0027] SM d42:1 |  |  |  |  |
| [#0028] SM d42:2 |  |  |  |  |
| [#0029] Cer 18:1;O2/16:0 |  |  |  |  |
| [#0030] Cer 18:1;O2/24:0 |  |  |  |  |
| [#0031] Cer 18:1;O2/24:1 |  |  |  |  |
| [#0034] Hex2Cer 18:1;O2/24:1 | |  |  |  |
| LION:0012082 | plasma membrane | 8 | 0.1953 | 0.5935 |
| [#0024] PS 18:0_22:3 |  |  |  |  |
| [#0025] SM d40:0 |  |  |  |  |
| [#0026] SM d40:1 |  |  |  |  |
| [#0027] SM d42:1 |  |  |  |  |
| [#0028] SM d42:2 |  |  |  |  |
| [#0029] Cer 18:1;O2/16:0 |  |  |  |  |
| [#0030] Cer 18:1;O2/24:0 |  |  |  |  |
| [#0031] Cer 18:1;O2/24:1 |  |  |  |  |
| LION:0000259 | C14:0 | 4 | 0.2198 | 0.5935 |
| [#0002] PC 14:0_14:0 |  |  |  |  |
| [#0003] PC 14:0_16:0 |  |  |  |  |
| [#0004] PC 14:0_16:1 |  |  |  |  |
| [#0005] PC 14:0_16:1_iso |  |  |  |  |
| LION:0080968 | very low bilayer thickness | 4 | 0.2198 | 0.5935 |
| [#0002] PC 14:0_14:0 |  |  |  |  |
| [#0004] PC 14:0_16:1 |  |  |  |  |
| [#0007] PC 28:1 |  |  |  |  |
| [#0009] PC 30:2 |  |  |  |  |
| LION:0080969 | low bilayer thickness | 4 | 0.2198 | 0.5935 |
| [#0003] PC 14:0_16:0 |  |  |  |  |
| [#0004] PC 14:0_16:1 |  |  |  |  |
| [#0006] PC 16:1_16:1 |  |  |  |  |
| [#0008] PC 29:0 |  |  |  |  |
| LION:0000084 | ceramide phosphocholines (sphingomyelins) [SP0301] | 4 | 0.2548 | 0.6151 |
| [#0025] SM d40:0 |  |  |  |  |
| [#0026] SM d40:1 |  |  |  |  |
| [#0027] SM d42:1 |  |  |  |  |
| [#0028] SM d42:2 |  |  |  |  |
| LION:0002900 | C16:1 | 6 | 0.262 | 0.6151 |
| [#0004] PC 14:0_16:1 |  |  |  |  |
| [#0005] PC 14:0_16:1_iso |  |  |  |  |
| [#0006] PC 16:1_16:1 |  |  |  |  |
| [#0010] PC O-16:0_16:1 |  |  |  |  |
| [#0016] PE 16:0_16:1 |  |  |  |  |
| [#0018] PE 16:1_18:1 |  |  |  |  |
| LION:0080978 | average lateral diffusion | 6 | 0.262 | 0.6151 |
| [#0002] PC 14:0_14:0 |  |  |  |  |
| [#0003] PC 14:0_16:0 |  |  |  |  |
| [#0008] PC 29:0 |  |  |  |  |
| [#0016] PE 16:0_16:1 |  |  |  |  |
| [#0018] PE 16:1_18:1 |  |  |  |  |
| [#0019] PE 18:1_22:3 |  |  |  |  |
| LION:0080971 | high bilayer thickness | 4 | 0.2931 | 0.6595 |
| [#0017] PE 16:0_18:1 |  |  |  |  |
| [#0019] PE 18:1_22:3 |  |  |  |  |
| [#0020] PE 34:0 |  |  |  |  |
| [#0024] PS 18:0_22:3 |  |  |  |  |
| LION:0002945 | fatty acid with more than 18 carbons | 6 | 0.3114 | 0.6726 |
| [#0001] FA 24:1 |  |  |  |  |
| [#0019] PE 18:1_22:3 |  |  |  |  |
| [#0022] PE P-18:0_22:2 |  |  |  |  |
| [#0023] PE P-18:1_22:3 |  |  |  |  |
| [#0024] PS 18:0_22:3 |  |  |  |  |
| [#0035] DG 18:1_20:1 |  |  |  |  |
| LION:0000038 | diacylglycerophosphoethanolamines [GP0201] | 5 | 0.3362 | 0.6983 |
| [#0016] PE 16:0_16:1 |  |  |  |  |
| [#0017] PE 16:0_18:1 |  |  |  |  |
| [#0018] PE 16:1_18:1 |  |  |  |  |
| [#0019] PE 18:1_22:3 |  |  |  |  |
| [#0020] PE 34:0 |  |  |  |  |
| LION:0000464 | negative intrinsic curvature | 16 | 0.3603 | 0.7035 |
| [#0016] PE 16:0_16:1 |  |  |  |  |
| [#0017] PE 16:0_18:1 |  |  |  |  |
| [#0018] PE 16:1_18:1 |  |  |  |  |
| [#0019] PE 18:1_22:3 |  |  |  |  |
| [#0020] PE 34:0 |  |  |  |  |
| [#0021] PE O-16:0_18:1 |  |  |  |  |
| [#0022] PE P-18:0_22:2 |  |  |  |  |
| [#0023] PE P-18:1_22:3 |  |  |  |  |
| [#0029] Cer 18:1;O2/16:0 |  |  |  |  |
| [#0030] Cer 18:1;O2/24:0 |  |  |  |  |
| [#0031] Cer 18:1;O2/24:1 |  |  |  |  |
| [#0035] DG 18:1_20:1 |  |  |  |  |
| [#0036] DG 34:2 |  |  |  |  |
| [#0037] DG 38:4 |  |  |  |  |
| [#0038] CL 72:4 |  |  |  |  |
| [#0039] CL 72:5 |  |  |  |  |
| LION:0000011 | glycerophosphoethanolamines [GP02] | 8 | 0.3648 | 0.7035 |
| [#0016] PE 16:0_16:1 |  |  |  |  |
| [#0017] PE 16:0_18:1 |  |  |  |  |
| [#0018] PE 16:1_18:1 |  |  |  |  |
| [#0019] PE 18:1_22:3 |  |  |  |  |
| [#0020] PE 34:0 |  |  |  |  |
| [#0021] PE O-16:0_18:1 |  |  |  |  |
| [#0022] PE P-18:0_22:2 |  |  |  |  |
| [#0023] PE P-18:1_22:3 |  |  |  |  |
| LION:0080977 | low lateral diffusion | 3 | 0.4066 | 0.7571 |
| [#0016] PE 16:0_16:1 |  |  |  |  |
| [#0017] PE 16:0_18:1 |  |  |  |  |
| [#0024] PS 18:0_22:3 |  |  |  |  |
| LION:0012009 | lipid-mediated signalling | 6 | 0.4244 | 0.7639 |
| [#0029] Cer 18:1;O2/16:0 |  |  |  |  |
| [#0030] Cer 18:1;O2/24:0 |  |  |  |  |
| [#0031] Cer 18:1;O2/24:1 |  |  |  |  |
| [#0035] DG 18:1_20:1 |  |  |  |  |
| [#0036] DG 34:2 |  |  |  |  |
| [#0037] DG 38:4 |  |  |  |  |
| LION:0002934 | C22:3 | 3 | 0.4481 | 0.7806 |
| [#0019] PE 18:1_22:3 |  |  |  |  |
| [#0023] PE P-18:1_22:3 |  |  |  |  |
| [#0024] PS 18:0_22:3 |  |  |  |  |
| LION:0000093 | headgroup with negative charge | 4 | 0.476 | 0.8033 |
| [#0001] FA 24:1 |  |  |  |  |
| [#0024] PS 18:0_22:3 |  |  |  |  |
| [#0038] CL 72:4 |  |  |  |  |
| [#0039] CL 72:5 |  |  |  |  |
| LION:0001736 | low transition temperature | 3 | 0.5353 | 0.8536 |
| [#0006] PC 16:1_16:1 |  |  |  |  |
| [#0009] PC 30:2 |  |  |  |  |
| [#0018] PE 16:1_18:1 |  |  |  |  |
| LION:0002966 | fatty acid with less than 2 double bonds | 27 | 0.5514 | 0.8536 |
| [#0001] FA 24:1 |  |  |  |  |
| [#0002] PC 14:0_14:0 |  |  |  |  |
| [#0003] PC 14:0_16:0 |  |  |  |  |
| [#0004] PC 14:0_16:1 |  |  |  |  |
| [#0005] PC 14:0_16:1_iso |  |  |  |  |
| [#0006] PC 16:1_16:1 |  |  |  |  |
| [#0010] PC O-16:0_16:1 |  |  |  |  |
| [#0016] PE 16:0_16:1 |  |  |  |  |
| [#0017] PE 16:0_18:1 |  |  |  |  |
| [#0018] PE 16:1_18:1 |  |  |  |  |
| [#0019] PE 18:1_22:3 |  |  |  |  |
| [#0021] PE O-16:0_18:1 |  |  |  |  |
| [#0022] PE P-18:0_22:2 |  |  |  |  |
| [#0023] PE P-18:1_22:3 |  |  |  |  |
| [#0024] PS 18:0_22:3 |  |  |  |  |
| [#0029] Cer 18:1;O2/16:0 |  |  |  |  |
| [#0030] Cer 18:1;O2/24:0 |  |  |  |  |
| [#0031] Cer 18:1;O2/24:1 |  |  |  |  |
| [#0032] Hex-Cer 18:1;O2/24:0 |  |  |  |  |
| [#0033] Hex-Cer 18:1;O2/24:1 |  |  |  |  |
| [#0034] Hex2Cer 18:1;O2/24:1 | |  |  |  |
| [#0035] DG 18:1_20:1 |  |  |  |  |
| [#0040] TG 16:0_18:0_18:1 |  |  |  |  |
| [#0041] TG 16:0_18:1_18:1 |  |  |  |  |
| [#0042] TG 18:0_18:0_18:1 |  |  |  |  |
| [#0043] TG 18:0_18:1_18:1 |  |  |  |  |
| [#0047] TG 18:1_18:1_18:1 |  |  |  |  |
| LION:0002950 | fatty acid with 22-24 carbons | 5 | 0.5564 | 0.8536 |
| [#0001] FA 24:1 |  |  |  |  |
| [#0019] PE 18:1_22:3 |  |  |  |  |
| [#0022] PE P-18:0_22:2 |  |  |  |  |
| [#0023] PE P-18:1_22:3 |  |  |  |  |
| [#0024] PS 18:0_22:3 |  |  |  |  |
| LION:0012441 | N-acylsphingosines (ceramides) [SP0201] | 3 | 0.5802 | 0.8536 |
| [#0029] Cer 18:1;O2/16:0 |  |  |  |  |
| [#0030] Cer 18:1;O2/24:0 |  |  |  |  |
| [#0031] Cer 18:1;O2/24:1 |  |  |  |  |
| LION:0002955 | fatty acid with 16 carbons | 11 | 0.6142 | 0.8536 |
| [#0003] PC 14:0_16:0 |  |  |  |  |
| [#0004] PC 14:0_16:1 |  |  |  |  |
| [#0005] PC 14:0_16:1_iso |  |  |  |  |
| [#0006] PC 16:1_16:1 |  |  |  |  |
| [#0010] PC O-16:0_16:1 |  |  |  |  |
| [#0016] PE 16:0_16:1 |  |  |  |  |
| [#0017] PE 16:0_18:1 |  |  |  |  |
| [#0018] PE 16:1_18:1 |  |  |  |  |
| [#0021] PE O-16:0_18:1 |  |  |  |  |
| [#0040] TG 16:0_18:0_18:1 |  |  |  |  |
| [#0041] TG 16:0_18:1_18:1 |  |  |  |  |
| LION:0002969 | monounsaturated fatty acid | 23 | 0.6143 | 0.8536 |
| [#0001] FA 24:1 |  |  |  |  |
| [#0004] PC 14:0_16:1 |  |  |  |  |
| [#0005] PC 14:0_16:1_iso |  |  |  |  |
| [#0006] PC 16:1_16:1 |  |  |  |  |
| [#0010] PC O-16:0_16:1 |  |  |  |  |
| [#0016] PE 16:0_16:1 |  |  |  |  |
| [#0017] PE 16:0_18:1 |  |  |  |  |
| [#0018] PE 16:1_18:1 |  |  |  |  |
| [#0019] PE 18:1_22:3 |  |  |  |  |
| [#0021] PE O-16:0_18:1 |  |  |  |  |
| [#0023] PE P-18:1_22:3 |  |  |  |  |
| [#0029] Cer 18:1;O2/16:0 |  |  |  |  |
| [#0030] Cer 18:1;O2/24:0 |  |  |  |  |
| [#0031] Cer 18:1;O2/24:1 |  |  |  |  |
| [#0032] Hex-Cer 18:1;O2/24:0 |  |  |  |  |
| [#0033] Hex-Cer 18:1;O2/24:1 |  |  |  |  |
| [#0034] Hex2Cer 18:1;O2/24:1 | |  |  |  |
| [#0035] DG 18:1_20:1 |  |  |  |  |
| [#0040] TG 16:0_18:0_18:1 |  |  |  |  |
| [#0041] TG 16:0_18:1_18:1 |  |  |  |  |
| [#0042] TG 18:0_18:0_18:1 |  |  |  |  |
| [#0043] TG 18:0_18:1_18:1 |  |  |  |  |
| [#0047] TG 18:1_18:1_18:1 |  |  |  |  |
| LION:0002961 | fatty acid with 22 carbons | 4 | 0.6323 | 0.8536 |
| [#0019] PE 18:1_22:3 |  |  |  |  |
| [#0022] PE P-18:0_22:2 |  |  |  |  |
| [#0023] PE P-18:1_22:3 |  |  |  |  |
| [#0024] PS 18:0_22:3 |  |  |  |  |
| LION:0002967 | polyunsaturated fatty acid | 4 | 0.6323 | 0.8536 |
| [#0019] PE 18:1_22:3 |  |  |  |  |
| [#0022] PE P-18:0_22:2 |  |  |  |  |
| [#0023] PE P-18:1_22:3 |  |  |  |  |
| [#0024] PS 18:0_22:3 |  |  |  |  |
| LION:0000100 | fatty acid with 18 carbons or less | 26 | 0.6566 | 0.8618 |
| [#0002] PC 14:0_14:0 |  |  |  |  |
| [#0003] PC 14:0_16:0 |  |  |  |  |
| [#0004] PC 14:0_16:1 |  |  |  |  |
| [#0005] PC 14:0_16:1_iso |  |  |  |  |
| [#0006] PC 16:1_16:1 |  |  |  |  |
| [#0010] PC O-16:0_16:1 |  |  |  |  |
| [#0016] PE 16:0_16:1 |  |  |  |  |
| [#0017] PE 16:0_18:1 |  |  |  |  |
| [#0018] PE 16:1_18:1 |  |  |  |  |
| [#0019] PE 18:1_22:3 |  |  |  |  |
| [#0021] PE O-16:0_18:1 |  |  |  |  |
| [#0022] PE P-18:0_22:2 |  |  |  |  |
| [#0023] PE P-18:1_22:3 |  |  |  |  |
| [#0024] PS 18:0_22:3 |  |  |  |  |
| [#0029] Cer 18:1;O2/16:0 |  |  |  |  |
| [#0030] Cer 18:1;O2/24:0 |  |  |  |  |
| [#0031] Cer 18:1;O2/24:1 |  |  |  |  |
| [#0032] Hex-Cer 18:1;O2/24:0 |  |  |  |  |
| [#0033] Hex-Cer 18:1;O2/24:1 |  |  |  |  |
| [#0034] Hex2Cer 18:1;O2/24:1 | |  |  |  |
| [#0035] DG 18:1_20:1 |  |  |  |  |
| [#0040] TG 16:0_18:0_18:1 |  |  |  |  |
| [#0041] TG 16:0_18:1_18:1 |  |  |  |  |
| [#0042] TG 18:0_18:0_18:1 |  |  |  |  |
| [#0043] TG 18:0_18:1_18:1 |  |  |  |  |
| [#0047] TG 18:1_18:1_18:1 |  |  |  |  |
| LION:0000607 | diacylglycerols [GL0201] | 3 | 0.6703 | 0.8618 |
| [#0035] DG 18:1_20:1 |  |  |  |  |
| [#0036] DG 34:2 |  |  |  |  |
| [#0037] DG 38:4 |  |  |  |  |
| LION:0012081 | mitochondrion | 10 | 0.7434 | 0.9289 |
| [#0016] PE 16:0_16:1 |  |  |  |  |
| [#0017] PE 16:0_18:1 |  |  |  |  |
| [#0018] PE 16:1_18:1 |  |  |  |  |
| [#0019] PE 18:1_22:3 |  |  |  |  |
| [#0020] PE 34:0 |  |  |  |  |
| [#0021] PE O-16:0_18:1 |  |  |  |  |
| [#0022] PE P-18:0_22:2 |  |  |  |  |
| [#0023] PE P-18:1_22:3 |  |  |  |  |
| [#0038] CL 72:4 |  |  |  |  |
| [#0039] CL 72:5 |  |  |  |  |
| LION:0002948 | fatty acid with 16-18 carbons | 25 | 0.7569 | 0.9289 |
| [#0003] PC 14:0_16:0 |  |  |  |  |
| [#0004] PC 14:0_16:1 |  |  |  |  |
| [#0005] PC 14:0_16:1_iso |  |  |  |  |
| [#0006] PC 16:1_16:1 |  |  |  |  |
| [#0010] PC O-16:0_16:1 |  |  |  |  |
| [#0016] PE 16:0_16:1 |  |  |  |  |
| [#0017] PE 16:0_18:1 |  |  |  |  |
| [#0018] PE 16:1_18:1 |  |  |  |  |
| [#0019] PE 18:1_22:3 |  |  |  |  |
| [#0021] PE O-16:0_18:1 |  |  |  |  |
| [#0022] PE P-18:0_22:2 |  |  |  |  |
| [#0023] PE P-18:1_22:3 |  |  |  |  |
| [#0024] PS 18:0_22:3 |  |  |  |  |
| [#0029] Cer 18:1;O2/16:0 |  |  |  |  |
| [#0030] Cer 18:1;O2/24:0 |  |  |  |  |
| [#0031] Cer 18:1;O2/24:1 |  |  |  |  |
| [#0032] Hex-Cer 18:1;O2/24:0 |  |  |  |  |
| [#0033] Hex-Cer 18:1;O2/24:1 |  |  |  |  |
| [#0034] Hex2Cer 18:1;O2/24:1 | |  |  |  |
| [#0035] DG 18:1_20:1 |  |  |  |  |
| [#0040] TG 16:0_18:0_18:1 |  |  |  |  |
| [#0041] TG 16:0_18:1_18:1 |  |  |  |  |
| [#0042] TG 18:0_18:0_18:1 |  |  |  |  |
| [#0043] TG 18:0_18:1_18:1 |  |  |  |  |
| [#0047] TG 18:1_18:1_18:1 |  |  |  |  |
| LION:0002882 | C16:0 | 7 | 0.8619 | 1 |
| [#0003] PC 14:0_16:0 |  |  |  |  |
| [#0010] PC O-16:0_16:1 |  |  |  |  |
| [#0016] PE 16:0_16:1 |  |  |  |  |
| [#0017] PE 16:0_18:1 |  |  |  |  |
| [#0021] PE O-16:0_18:1 |  |  |  |  |
| [#0040] TG 16:0_18:0_18:1 |  |  |  |  |
| [#0041] TG 16:0_18:1_18:1 |  |  |  |  |
| LION:0002968 | saturated fatty acid | 14 | 0.9778 | 1 |
| [#0002] PC 14:0_14:0 |  |  |  |  |
| [#0003] PC 14:0_16:0 |  |  |  |  |
| [#0004] PC 14:0_16:1 |  |  |  |  |
| [#0005] PC 14:0_16:1_iso |  |  |  |  |
| [#0010] PC O-16:0_16:1 |  |  |  |  |
| [#0016] PE 16:0_16:1 |  |  |  |  |
| [#0017] PE 16:0_18:1 |  |  |  |  |
| [#0021] PE O-16:0_18:1 |  |  |  |  |
| [#0022] PE P-18:0_22:2 |  |  |  |  |
| [#0024] PS 18:0_22:3 |  |  |  |  |
| [#0040] TG 16:0_18:0_18:1 |  |  |  |  |
| [#0041] TG 16:0_18:1_18:1 |  |  |  |  |
| [#0042] TG 18:0_18:0_18:1 |  |  |  |  |
| [#0043] TG 18:0_18:1_18:1 |  |  |  |  |
| LION:0000002 | glycerolipids [GL] | 12 | 1 | 1 |
| [#0035] DG 18:1_20:1 |  |  |  |  |
| [#0036] DG 34:2 |  |  |  |  |
| [#0037] DG 38:4 |  |  |  |  |
| [#0040] TG 16:0_18:0_18:1 |  |  |  |  |
| [#0041] TG 16:0_18:1_18:1 |  |  |  |  |
| [#0042] TG 18:0_18:0_18:1 |  |  |  |  |
| [#0043] TG 18:0_18:1_18:1 |  |  |  |  |
| [#0044] TG 18:0_38:3 |  |  |  |  |
| [#0045] TG 18:0_42:3 |  |  |  |  |
| [#0046] TG 18:0_48:2 |  |  |  |  |
| [#0047] TG 18:1_18:1_18:1 |  |  |  |  |
| [#0048] TG 20:0_36:2 |  |  |  |  |
| LION:0000094 | headgroup with neutral charge | 13 | 1 | 1 |
| [#0034] Hex2Cer 18:1;O2/24:1 | |  |  |  |
| [#0035] DG 18:1_20:1 |  |  |  |  |
| [#0036] DG 34:2 |  |  |  |  |
| [#0037] DG 38:4 |  |  |  |  |
| [#0040] TG 16:0_18:0_18:1 |  |  |  |  |
| [#0041] TG 16:0_18:1_18:1 |  |  |  |  |
| [#0042] TG 18:0_18:0_18:1 |  |  |  |  |
| [#0043] TG 18:0_18:1_18:1 |  |  |  |  |
| [#0044] TG 18:0_38:3 |  |  |  |  |
| [#0045] TG 18:0_42:3 |  |  |  |  |
| [#0046] TG 18:0_48:2 |  |  |  |  |
| [#0047] TG 18:1_18:1_18:1 |  |  |  |  |
| [#0048] TG 20:0_36:2 |  |  |  |  |
| LION:0000622 | triacylglycerols [GL0301] | 9 | 1 | 1 |
| [#0040] TG 16:0_18:0_18:1 |  |  |  |  |
| [#0041] TG 16:0_18:1_18:1 |  |  |  |  |
| [#0042] TG 18:0_18:0_18:1 |  |  |  |  |
| [#0043] TG 18:0_18:1_18:1 |  |  |  |  |
| [#0044] TG 18:0_38:3 |  |  |  |  |
| [#0045] TG 18:0_42:3 |  |  |  |  |
| [#0046] TG 18:0_48:2 |  |  |  |  |
| [#0047] TG 18:1_18:1_18:1 |  |  |  |  |
| [#0048] TG 20:0_36:2 |  |  |  |  |
| LION:0002921 | C18:0 | 5 | 1 | 1 |
| [#0022] PE P-18:0_22:2 |  |  |  |  |
| [#0024] PS 18:0_22:3 |  |  |  |  |
| [#0040] TG 16:0_18:0_18:1 |  |  |  |  |
| [#0042] TG 18:0_18:0_18:1 |  |  |  |  |
| [#0043] TG 18:0_18:1_18:1 |  |  |  |  |
| LION:0002922 | C18:1 | 17 | 1 | 1 |
| [#0017] PE 16:0_18:1 |  |  |  |  |
| [#0018] PE 16:1_18:1 |  |  |  |  |
| [#0019] PE 18:1_22:3 |  |  |  |  |
| [#0021] PE O-16:0_18:1 |  |  |  |  |
| [#0023] PE P-18:1_22:3 |  |  |  |  |
| [#0029] Cer 18:1;O2/16:0 |  |  |  |  |
| [#0030] Cer 18:1;O2/24:0 |  |  |  |  |
| [#0031] Cer 18:1;O2/24:1 |  |  |  |  |
| [#0032] Hex-Cer 18:1;O2/24:0 |  |  |  |  |
| [#0033] Hex-Cer 18:1;O2/24:1 |  |  |  |  |
| [#0034] Hex2Cer 18:1;O2/24:1 | |  |  |  |
| [#0035] DG 18:1_20:1 |  |  |  |  |
| [#0040] TG 16:0_18:0_18:1 |  |  |  |  |
| [#0041] TG 16:0_18:1_18:1 |  |  |  |  |
| [#0042] TG 18:0_18:0_18:1 |  |  |  |  |
| [#0043] TG 18:0_18:1_18:1 |  |  |  |  |
| [#0047] TG 18:1_18:1_18:1 |  |  |  |  |
| LION:0002957 | fatty acid with 18 carbons | 19 | 1 | 1 |
| [#0017] PE 16:0_18:1 |  |  |  |  |
| [#0018] PE 16:1_18:1 |  |  |  |  |
| [#0019] PE 18:1_22:3 |  |  |  |  |
| [#0021] PE O-16:0_18:1 |  |  |  |  |
| [#0022] PE P-18:0_22:2 |  |  |  |  |
| [#0023] PE P-18:1_22:3 |  |  |  |  |
| [#0024] PS 18:0_22:3 |  |  |  |  |
| [#0029] Cer 18:1;O2/16:0 |  |  |  |  |
| [#0030] Cer 18:1;O2/24:0 |  |  |  |  |
| [#0031] Cer 18:1;O2/24:1 |  |  |  |  |
| [#0032] Hex-Cer 18:1;O2/24:0 |  |  |  |  |
| [#0033] Hex-Cer 18:1;O2/24:1 |  |  |  |  |
| [#0034] Hex2Cer 18:1;O2/24:1 | |  |  |  |
| [#0035] DG 18:1_20:1 |  |  |  |  |
| [#0040] TG 16:0_18:0_18:1 |  |  |  |  |
| [#0041] TG 16:0_18:1_18:1 |  |  |  |  |
| [#0042] TG 18:0_18:0_18:1 |  |  |  |  |
| [#0043] TG 18:0_18:1_18:1 |  |  |  |  |
| [#0047] TG 18:1_18:1_18:1 |  |  |  |  |
| LION:0012011 | lipid storage | 9 | 1 | 1 |
| [#0040] TG 16:0_18:0_18:1 |  |  |  |  |
| [#0041] TG 16:0_18:1_18:1 |  |  |  |  |
| [#0042] TG 18:0_18:0_18:1 |  |  |  |  |
| [#0043] TG 18:0_18:1_18:1 |  |  |  |  |
| [#0044] TG 18:0_38:3 |  |  |  |  |
| [#0045] TG 18:0_42:3 |  |  |  |  |
| [#0046] TG 18:0_48:2 |  |  |  |  |
| [#0047] TG 18:1_18:1_18:1 |  |  |  |  |
| [#0048] TG 20:0_36:2 |  |  |  |  |
| LION:0012084 | lipid droplet | 9 | 1 | 1 |
| [#0040] TG 16:0_18:0_18:1 |  |  |  |  |
| [#0041] TG 16:0_18:1_18:1 |  |  |  |  |
| [#0042] TG 18:0_18:0_18:1 |  |  |  |  |
| [#0043] TG 18:0_18:1_18:1 |  |  |  |  |
| [#0044] TG 18:0_38:3 |  |  |  |  |
| [#0045] TG 18:0_42:3 |  |  |  |  |
| [#0046] TG 18:0_48:2 |  |  |  |  |
| [#0047] TG 18:1_18:1_18:1 |  |  |  |  |
| [#0048] TG 20:0_36:2 |  |  |  |  |

**Table 5d: LION lipid associations:**

| LION-term | LION-name | [#0001] FA 24:1 | [#0002] PC 14:0_14:0 | [#0003] PC 14:0_16:0 | [#0004] PC 14:0_16:1 | [#0005] PC 14:0_16:1_iso | [#0006] PC 16:1_16:1 | [#0007] PC 28:1 | [#0008] PC 29:0 | [#0009] PC 30:2 | [#0010] PC O-16:0_16:1 | [#0011] PC O-30:0 | [#0012] PC O-30:1 | [#0013] PC O-32:2 | [#0014] PC O-34:3 | [#0015] PC O-40:2 | [#0016] PE 16:0_16:1 | [#0017] PE 16:0_18:1 | [#0018] PE 16:1_18:1 | [#0019] PE 18:1_22:3 | [#0020] PE 34:0 | [#0021] PE O-16:0_18:1 | [#0022] PE P-18:0_22:2 | [#0023] PE P-18:1_22:3 | [#0024] PS 18:0_22:3 | [#0025] SM d40:0 | [#0026] SM d40:1 | [#0027] SM d42:1 | [#0028] SM d42:2 | [#0029] Cer 18:1;O2/16:0 | [#0030] Cer 18:1;O2/24:0 | [#0031] Cer 18:1;O2/24:1 | [#0032] Hex-Cer 18:1;O2/24:0 | [#0033] Hex-Cer 18:1;O2/24:1 | [#0034] Hex2Cer 18:1;O2/24:1 | [#0035] DG 18:1_20:1 | [#0036] DG 34:2 | [#0037] DG 38:4 | [#0038] CL 72:4 | [#0039] CL 72:5 | [#0040] TG 16:0_18:0_18:1 | [#0041] TG 16:0_18:1_18:1 | [#0042] TG 18:0_18:0_18:1 | [#0043] TG 18:0_18:1_18:1 | [#0044] TG 18:0_38:3 | [#0045] TG 18:0_42:3 | [#0046] TG 18:0_48:2 | [#0047] TG 18:1_18:1_18:1 | [#0048] TG 20:0_36:2 |
| --- | --- | --- | --- | --- | --- | --- | --- | --- | --- | --- | --- | --- | --- | --- | --- | --- | --- | --- | --- | --- | --- | --- | --- | --- | --- | --- | --- | --- | --- | --- | --- | --- | --- | --- | --- | --- | --- | --- | --- | --- | --- | --- | --- | --- | --- | --- | --- | --- | --- |
| LION:0000001 | fatty acids [FA] | x |  |  |  |  |  |  |  |  |  |  |  |  |  |  |  |  |  |  |  |  |  |  |  |  |  |  |  |  |  |  |  |  |  |  |  |  |  |  |  |  |  |  |  |  |  |  |  |
| LION:0000002 | glycerolipids [GL] |  |  |  |  |  |  |  |  |  |  |  |  |  |  |  |  |  |  |  |  |  |  |  |  |  |  |  |  |  |  |  |  |  |  | x | x | x |  |  | x | x | x | x | x | x | x | x | x |
| LION:0000003 | glycerophospholipids [GP] |  | x | x | x | x | x | x | x | x | x | x | x | x | x | x | x | x | x | x | x | x | x | x | x |  |  |  |  |  |  |  |  |  |  |  |  |  | x | x |  |  |  |  |  |  |  |  |  |
| LION:0000004 | sphingolipids [SP] |  |  |  |  |  |  |  |  |  |  |  |  |  |  |  |  |  |  |  |  |  |  |  |  | x | x | x | x | x | x | x |  |  | x |  |  |  |  |  |  |  |  |  |  |  |  |  |  |
| LION:0000010 | glycerophosphocholines [GP01] |  | x | x | x | x | x | x | x | x | x | x | x | x | x | x |  |  |  |  |  |  |  |  |  |  |  |  |  |  |  |  |  |  |  |  |  |  |  |  |  |  |  |  |  |  |  |  |  |
| LION:0000011 | glycerophosphoethanolamines [GP02] |  |  |  |  |  |  |  |  |  |  |  |  |  |  |  | x | x | x | x | x | x | x | x |  |  |  |  |  |  |  |  |  |  |  |  |  |  |  |  |  |  |  |  |  |  |  |  |  |
| LION:0000013 | glycerophosphoserines [GP03] |  |  |  |  |  |  |  |  |  |  |  |  |  |  |  |  |  |  |  |  |  |  |  | x |  |  |  |  |  |  |  |  |  |  |  |  |  |  |  |  |  |  |  |  |  |  |  |  |
| LION:0000021 | glycerophosphoglycerophosphoglycerols [GP12] |  |  |  |  |  |  |  |  |  |  |  |  |  |  |  |  |  |  |  |  |  |  |  |  |  |  |  |  |  |  |  |  |  |  |  |  |  | x | x |  |  |  |  |  |  |  |  |  |
| LION:0000030 | diacylglycerophosphocholines [GP0101] |  | x | x | x | x | x | x | x | x |  |  |  |  |  |  |  |  |  |  |  |  |  |  |  |  |  |  |  |  |  |  |  |  |  |  |  |  |  |  |  |  |  |  |  |  |  |  |  |
| LION:0000031 | 1-alkyl,2-acylglycerophosphocholines [GP0102] |  |  |  |  |  |  |  |  |  | x | x | x | x | x | x |  |  |  |  |  |  |  |  |  |  |  |  |  |  |  |  |  |  |  |  |  |  |  |  |  |  |  |  |  |  |  |  |  |
| LION:0000038 | diacylglycerophosphoethanolamines [GP0201] |  |  |  |  |  |  |  |  |  |  |  |  |  |  |  | x | x | x | x | x |  |  |  |  |  |  |  |  |  |  |  |  |  |  |  |  |  |  |  |  |  |  |  |  |  |  |  |  |
| LION:0000039 | 1-alkyl,2-acylglycerophosphoethanolamines [GP0202] |  |  |  |  |  |  |  |  |  |  |  |  |  |  |  |  |  |  |  |  | x |  |  |  |  |  |  |  |  |  |  |  |  |  |  |  |  |  |  |  |  |  |  |  |  |  |  |  |
| LION:0000040 | 1-(1z-alkenyl),2-acylglycerophosphoethanolamines [GP0203] |  |  |  |  |  |  |  |  |  |  |  |  |  |  |  |  |  |  |  |  |  | x | x |  |  |  |  |  |  |  |  |  |  |  |  |  |  |  |  |  |  |  |  |  |  |  |  |  |
| LION:0000053 | diacylglycerophosphoserines [GP0301] |  |  |  |  |  |  |  |  |  |  |  |  |  |  |  |  |  |  |  |  |  |  |  | x |  |  |  |  |  |  |  |  |  |  |  |  |  |  |  |  |  |  |  |  |  |  |  |  |
| LION:0000077 | ceramides [SP02] |  |  |  |  |  |  |  |  |  |  |  |  |  |  |  |  |  |  |  |  |  |  |  |  |  |  |  |  | x | x | x |  |  |  |  |  |  |  |  |  |  |  |  |  |  |  |  |  |
| LION:0000078 | phosphosphingolipids [SP03] |  |  |  |  |  |  |  |  |  |  |  |  |  |  |  |  |  |  |  |  |  |  |  |  | x | x | x | x |  |  |  |  |  |  |  |  |  |  |  |  |  |  |  |  |  |  |  |  |
| LION:0000080 | neutral glycosphingolipids [SP05] |  |  |  |  |  |  |  |  |  |  |  |  |  |  |  |  |  |  |  |  |  |  |  |  |  |  |  |  |  |  |  |  |  | x |  |  |  |  |  |  |  |  |  |  |  |  |  |  |
| LION:0000084 | ceramide phosphocholines (sphingomyelins) [SP0301] |  |  |  |  |  |  |  |  |  |  |  |  |  |  |  |  |  |  |  |  |  |  |  |  | x | x | x | x |  |  |  |  |  |  |  |  |  |  |  |  |  |  |  |  |  |  |  |  |
| LION:0000093 | headgroup with negative charge | x |  |  |  |  |  |  |  |  |  |  |  |  |  |  |  |  |  |  |  |  |  |  | x |  |  |  |  |  |  |  |  |  |  |  |  |  | x | x |  |  |  |  |  |  |  |  |  |
| LION:0000094 | headgroup with neutral charge |  |  |  |  |  |  |  |  |  |  |  |  |  |  |  |  |  |  |  |  |  |  |  |  |  |  |  |  |  |  |  |  |  | x | x | x | x |  |  | x | x | x | x | x | x | x | x | x |
| LION:0000095 | headgroup with positive charge / zwitter-ion |  | x | x | x | x | x | x | x | x | x | x | x | x | x | x | x | x | x | x | x | x | x | x |  | x | x | x | x |  |  |  |  |  |  |  |  |  |  |  |  |  |  |  |  |  |  |  |  |
| LION:0000100 | fatty acid with 18 carbons or less |  | x | x | x | x | x |  |  |  | x |  |  |  |  |  | x | x | x | x |  | x | x | x | x |  |  |  |  | x | x | x | x | x | x | x |  |  |  |  | x | x | x | x |  |  |  | x |  |
| LION:0000104 | PC(O-16:0/16:1) |  |  |  |  |  |  |  |  |  | x |  |  |  |  |  |  |  |  |  |  |  |  |  |  |  |  |  |  |  |  |  |  |  |  |  |  |  |  |  |  |  |  |  |  |  |  |  |  |
| LION:0000140 | PC(28:0) |  | x |  |  |  |  |  |  |  |  |  |  |  |  |  |  |  |  |  |  |  |  |  |  |  |  |  |  |  |  |  |  |  |  |  |  |  |  |  |  |  |  |  |  |  |  |  |  |
| LION:0000143 | PC(O-30:0) |  |  |  |  |  |  |  |  |  |  | x |  |  |  |  |  |  |  |  |  |  |  |  |  |  |  |  |  |  |  |  |  |  |  |  |  |  |  |  |  |  |  |  |  |  |  |  |  |
| LION:0000145 | PC(30:2) |  |  |  |  |  |  |  |  | x |  |  |  |  |  |  |  |  |  |  |  |  |  |  |  |  |  |  |  |  |  |  |  |  |  |  |  |  |  |  |  |  |  |  |  |  |  |  |  |
| LION:0000146 | PC(30:1) |  |  |  | x |  |  |  |  |  |  |  |  |  |  |  |  |  |  |  |  |  |  |  |  |  |  |  |  |  |  |  |  |  |  |  |  |  |  |  |  |  |  |  |  |  |  |  |  |
| LION:0000147 | PC(30:0) |  |  | x |  |  |  |  |  |  |  |  |  |  |  |  |  |  |  |  |  |  |  |  |  |  |  |  |  |  |  |  |  |  |  |  |  |  |  |  |  |  |  |  |  |  |  |  |  |
| LION:0000151 | PC(O-32:2) |  |  |  |  |  |  |  |  |  |  |  |  | x |  |  |  |  |  |  |  |  |  |  |  |  |  |  |  |  |  |  |  |  |  |  |  |  |  |  |  |  |  |  |  |  |  |  |  |
| LION:0000152 | PC(O-32:1) |  |  |  |  |  |  |  |  |  | x |  |  |  |  |  |  |  |  |  |  |  |  |  |  |  |  |  |  |  |  |  |  |  |  |  |  |  |  |  |  |  |  |  |  |  |  |  |  |
| LION:0000157 | PC(32:2) |  |  |  |  |  | x |  |  |  |  |  |  |  |  |  |  |  |  |  |  |  |  |  |  |  |  |  |  |  |  |  |  |  |  |  |  |  |  |  |  |  |  |  |  |  |  |  |  |
| LION:0000163 | PC(O-34:3) |  |  |  |  |  |  |  |  |  |  |  |  |  | x |  |  |  |  |  |  |  |  |  |  |  |  |  |  |  |  |  |  |  |  |  |  |  |  |  |  |  |  |  |  |  |  |  |  |
| LION:0000259 | C14:0 |  | x | x | x | x |  |  |  |  |  |  |  |  |  |  |  |  |  |  |  |  |  |  |  |  |  |  |  |  |  |  |  |  |  |  |  |  |  |  |  |  |  |  |  |  |  |  |  |
| LION:0000324 | PE(32:1) |  |  |  |  |  |  |  |  |  |  |  |  |  |  |  | x |  |  |  |  |  |  |  |  |  |  |  |  |  |  |  |  |  |  |  |  |  |  |  |  |  |  |  |  |  |  |  |  |
| LION:0000331 | PE(34:2) |  |  |  |  |  |  |  |  |  |  |  |  |  |  |  |  |  | x |  |  |  |  |  |  |  |  |  |  |  |  |  |  |  |  |  |  |  |  |  |  |  |  |  |  |  |  |  |  |
| LION:0000332 | PE(34:1) |  |  |  |  |  |  |  |  |  |  |  |  |  |  |  |  | x |  |  |  |  |  |  |  |  |  |  |  |  |  |  |  |  |  |  |  |  |  |  |  |  |  |  |  |  |  |  |  |
| LION:0000333 | PE(34:0) |  |  |  |  |  |  |  |  |  |  |  |  |  |  |  |  |  |  |  | x |  |  |  |  |  |  |  |  |  |  |  |  |  |  |  |  |  |  |  |  |  |  |  |  |  |  |  |  |
| LION:0000378 | PS(40:3) |  |  |  |  |  |  |  |  |  |  |  |  |  |  |  |  |  |  |  |  |  |  |  | x |  |  |  |  |  |  |  |  |  |  |  |  |  |  |  |  |  |  |  |  |  |  |  |  |
| LION:0000409 | SM(40:1) |  |  |  |  |  |  |  |  |  |  |  |  |  |  |  |  |  |  |  |  |  |  |  |  |  | x |  |  |  |  |  |  |  |  |  |  |  |  |  |  |  |  |  |  |  |  |  |  |
| LION:0000410 | SM(40:0) |  |  |  |  |  |  |  |  |  |  |  |  |  |  |  |  |  |  |  |  |  |  |  |  | x |  |  |  |  |  |  |  |  |  |  |  |  |  |  |  |  |  |  |  |  |  |  |  |
| LION:0000414 | SM(42:2) |  |  |  |  |  |  |  |  |  |  |  |  |  |  |  |  |  |  |  |  |  |  |  |  |  |  |  | x |  |  |  |  |  |  |  |  |  |  |  |  |  |  |  |  |  |  |  |  |
| LION:0000415 | SM(42:1) |  |  |  |  |  |  |  |  |  |  |  |  |  |  |  |  |  |  |  |  |  |  |  |  |  |  | x |  |  |  |  |  |  |  |  |  |  |  |  |  |  |  |  |  |  |  |  |  |
| LION:0000453 | simple glc series [SP0501] |  |  |  |  |  |  |  |  |  |  |  |  |  |  |  |  |  |  |  |  |  |  |  |  |  |  |  |  |  |  |  |  |  | x |  |  |  |  |  |  |  |  |  |  |  |  |  |  |
| LION:0000464 | negative intrinsic curvature |  |  |  |  |  |  |  |  |  |  |  |  |  |  |  | x | x | x | x | x | x | x | x |  |  |  |  |  | x | x | x |  |  |  | x | x | x | x | x |  |  |  |  |  |  |  |  |  |
| LION:0000465 | neutral intrinsic curvature |  | x | x | x | x | x | x | x | x | x | x | x | x | x | x |  |  |  |  |  |  |  |  | x |  |  |  |  |  |  |  |  |  |  |  |  |  |  |  |  |  |  |  |  |  |  |  |  |
| LION:0000467 | contains ether-bond |  |  |  |  |  |  |  |  |  | x | x | x | x | x | x |  |  |  |  |  | x |  |  |  |  |  |  |  |  |  |  |  |  |  |  |  |  |  |  |  |  |  |  |  |  |  |  |  |
| LION:0000603 | diradylglycerols [GL02] |  |  |  |  |  |  |  |  |  |  |  |  |  |  |  |  |  |  |  |  |  |  |  |  |  |  |  |  |  |  |  |  |  |  | x | x | x |  |  |  |  |  |  |  |  |  |  |  |
| LION:0000604 | triradylglycerols [GL03] |  |  |  |  |  |  |  |  |  |  |  |  |  |  |  |  |  |  |  |  |  |  |  |  |  |  |  |  |  |  |  |  |  |  |  |  |  |  |  | x | x | x | x | x | x | x | x | x |
| LION:0000607 | diacylglycerols [GL0201] |  |  |  |  |  |  |  |  |  |  |  |  |  |  |  |  |  |  |  |  |  |  |  |  |  |  |  |  |  |  |  |  |  |  | x | x | x |  |  |  |  |  |  |  |  |  |  |  |
| LION:0000622 | triacylglycerols [GL0301] |  |  |  |  |  |  |  |  |  |  |  |  |  |  |  |  |  |  |  |  |  |  |  |  |  |  |  |  |  |  |  |  |  |  |  |  |  |  |  | x | x | x | x | x | x | x | x | x |
| LION:0000640 | DG(34:2) |  |  |  |  |  |  |  |  |  |  |  |  |  |  |  |  |  |  |  |  |  |  |  |  |  |  |  |  |  |  |  |  |  |  |  | x |  |  |  |  |  |  |  |  |  |  |  |  |
| LION:0000655 | DG(38:2) |  |  |  |  |  |  |  |  |  |  |  |  |  |  |  |  |  |  |  |  |  |  |  |  |  |  |  |  |  |  |  |  |  |  | x |  |  |  |  |  |  |  |  |  |  |  |  |  |
| LION:0000657 | DG(38:4) |  |  |  |  |  |  |  |  |  |  |  |  |  |  |  |  |  |  |  |  |  |  |  |  |  |  |  |  |  |  |  |  |  |  |  |  | x |  |  |  |  |  |  |  |  |  |  |  |
| LION:0000728 | diacylglycerophosphoglycerophosphodiradylglycerols [GP1201] |  |  |  |  |  |  |  |  |  |  |  |  |  |  |  |  |  |  |  |  |  |  |  |  |  |  |  |  |  |  |  |  |  |  |  |  |  | x | x |  |  |  |  |  |  |  |  |  |
| LION:0000760 | TG(52:1) |  |  |  |  |  |  |  |  |  |  |  |  |  |  |  |  |  |  |  |  |  |  |  |  |  |  |  |  |  |  |  |  |  |  |  |  |  |  |  | x |  |  |  |  |  |  |  |  |
| LION:0000761 | TG(52:2) |  |  |  |  |  |  |  |  |  |  |  |  |  |  |  |  |  |  |  |  |  |  |  |  |  |  |  |  |  |  |  |  |  |  |  |  |  |  |  |  | x |  |  |  |  |  |  |  |
| LION:0000770 | TG(54:1) |  |  |  |  |  |  |  |  |  |  |  |  |  |  |  |  |  |  |  |  |  |  |  |  |  |  |  |  |  |  |  |  |  |  |  |  |  |  |  |  |  | x |  |  |  |  |  |  |
| LION:0000773 | TG(54:2) |  |  |  |  |  |  |  |  |  |  |  |  |  |  |  |  |  |  |  |  |  |  |  |  |  |  |  |  |  |  |  |  |  |  |  |  |  |  |  |  |  |  | x |  |  |  |  |  |
| LION:0000774 | TG(54:3) |  |  |  |  |  |  |  |  |  |  |  |  |  |  |  |  |  |  |  |  |  |  |  |  |  |  |  |  |  |  |  |  |  |  |  |  |  |  |  |  |  |  |  |  |  |  | x |  |
| LION:0001006 | CL(72:4) |  |  |  |  |  |  |  |  |  |  |  |  |  |  |  |  |  |  |  |  |  |  |  |  |  |  |  |  |  |  |  |  |  |  |  |  |  | x |  |  |  |  |  |  |  |  |  |  |
| LION:0001007 | CL(72:5) |  |  |  |  |  |  |  |  |  |  |  |  |  |  |  |  |  |  |  |  |  |  |  |  |  |  |  |  |  |  |  |  |  |  |  |  |  |  | x |  |  |  |  |  |  |  |  |  |
| LION:0001285 | PE(40:4) |  |  |  |  |  |  |  |  |  |  |  |  |  |  |  |  |  |  | x |  |  |  |  |  |  |  |  |  |  |  |  |  |  |  |  |  |  |  |  |  |  |  |  |  |  |  |  |  |
| LION:0001320 | PC(O-30:1) |  |  |  |  |  |  |  |  |  |  |  | x |  |  |  |  |  |  |  |  |  |  |  |  |  |  |  |  |  |  |  |  |  |  |  |  |  |  |  |  |  |  |  |  |  |  |  |  |
| LION:0001333 | PC(O-40:2) |  |  |  |  |  |  |  |  |  |  |  |  |  |  | x |  |  |  |  |  |  |  |  |  |  |  |  |  |  |  |  |  |  |  |  |  |  |  |  |  |  |  |  |  |  |  |  |  |
| LION:0001355 | PE(O-34:1) |  |  |  |  |  |  |  |  |  |  |  |  |  |  |  |  |  |  |  |  | x |  |  |  |  |  |  |  |  |  |  |  |  |  |  |  |  |  |  |  |  |  |  |  |  |  |  |  |
| LION:0001734 | contains vinyl ether bond (plasmalogen) |  |  |  |  |  |  |  |  |  |  |  |  |  |  |  |  |  |  |  |  |  | x | x |  |  |  |  |  |  |  |  |  |  |  |  |  |  |  |  |  |  |  |  |  |  |  |  |  |
| LION:0001736 | low transition temperature |  |  |  |  |  | x |  |  | x |  |  |  |  |  |  |  |  | x |  |  |  |  |  |  |  |  |  |  |  |  |  |  |  |  |  |  |  |  |  |  |  |  |  |  |  |  |  |  |
| LION:0001737 | average transition temperature |  | x |  | x |  |  | x | x |  |  |  |  |  |  |  | x |  |  | x |  |  |  |  | x |  |  |  |  |  |  |  |  |  |  |  |  |  |  |  |  |  |  |  |  |  |  |  |  |
| LION:0001738 | high transition temperature |  |  | x |  |  |  |  |  |  |  |  |  |  |  |  |  | x |  |  |  |  |  |  |  |  |  |  |  |  |  |  |  |  |  |  |  |  |  |  |  |  |  |  |  |  |  |  |  |
| LION:0001739 | very high transition temperature |  |  |  |  |  |  |  |  |  |  |  |  |  |  |  |  |  |  |  | x |  |  |  |  | x | x | x | x |  |  |  |  |  |  |  |  |  |  |  |  |  |  |  |  |  |  |  |  |
| LION:0001740 | above average transition temperature |  |  | x |  |  |  |  |  |  |  |  |  |  |  |  |  | x |  |  | x |  |  |  |  | x | x | x | x |  |  |  |  |  |  |  |  |  |  |  |  |  |  |  |  |  |  |  |  |
| LION:0001741 | below average transition temperature |  |  |  |  |  | x |  |  | x |  |  |  |  |  |  |  |  | x |  |  |  |  |  |  |  |  |  |  |  |  |  |  |  |  |  |  |  |  |  |  |  |  |  |  |  |  |  |  |
| LION:0001742 | fatty acids and conjugates [FA01] | x |  |  |  |  |  |  |  |  |  |  |  |  |  |  |  |  |  |  |  |  |  |  |  |  |  |  |  |  |  |  |  |  |  |  |  |  |  |  |  |  |  |  |  |  |  |  |  |
| LION:0001767 | FFA(24:1) | x |  |  |  |  |  |  |  |  |  |  |  |  |  |  |  |  |  |  |  |  |  |  |  |  |  |  |  |  |  |  |  |  |  |  |  |  |  |  |  |  |  |  |  |  |  |  |  |
| LION:0002562 | PC(28:1) |  |  |  |  |  |  | x |  |  |  |  |  |  |  |  |  |  |  |  |  |  |  |  |  |  |  |  |  |  |  |  |  |  |  |  |  |  |  |  |  |  |  |  |  |  |  |  |  |
| LION:0002882 | C16:0 |  |  | x |  |  |  |  |  |  | x |  |  |  |  |  | x | x |  |  |  | x |  |  |  |  |  |  |  |  |  |  |  |  |  |  |  |  |  |  | x | x |  |  |  |  |  |  |  |
| LION:0002900 | C16:1 |  |  |  | x | x | x |  |  |  | x |  |  |  |  |  | x |  | x |  |  |  |  |  |  |  |  |  |  |  |  |  |  |  |  |  |  |  |  |  |  |  |  |  |  |  |  |  |  |
| LION:0002921 | C18:0 |  |  |  |  |  |  |  |  |  |  |  |  |  |  |  |  |  |  |  |  |  | x |  | x |  |  |  |  |  |  |  |  |  |  |  |  |  |  |  | x |  | x | x |  |  |  |  |  |
| LION:0002922 | C18:1 |  |  |  |  |  |  |  |  |  |  |  |  |  |  |  |  | x | x | x |  | x |  | x |  |  |  |  |  | x | x | x | x | x | x | x |  |  |  |  | x | x | x | x |  |  |  | x |  |
| LION:0002926 | C20:1 |  |  |  |  |  |  |  |  |  |  |  |  |  |  |  |  |  |  |  |  |  |  |  |  |  |  |  |  |  |  |  |  |  |  | x |  |  |  |  |  |  |  |  |  |  |  |  |  |
| LION:0002933 | C22:2 |  |  |  |  |  |  |  |  |  |  |  |  |  |  |  |  |  |  |  |  |  | x |  |  |  |  |  |  |  |  |  |  |  |  |  |  |  |  |  |  |  |  |  |  |  |  |  |  |
| LION:0002934 | C22:3 |  |  |  |  |  |  |  |  |  |  |  |  |  |  |  |  |  |  | x |  |  |  | x | x |  |  |  |  |  |  |  |  |  |  |  |  |  |  |  |  |  |  |  |  |  |  |  |  |
| LION:0002939 | C24:1 | x |  |  |  |  |  |  |  |  |  |  |  |  |  |  |  |  |  |  |  |  |  |  |  |  |  |  |  |  |  |  |  |  |  |  |  |  |  |  |  |  |  |  |  |  |  |  |  |
| LION:0002945 | fatty acid with more than 18 carbons | x |  |  |  |  |  |  |  |  |  |  |  |  |  |  |  |  |  | x |  |  | x | x | x |  |  |  |  |  |  |  |  |  |  | x |  |  |  |  |  |  |  |  |  |  |  |  |  |
| LION:0002947 | fatty acid with 13-15 carbons |  | x | x | x | x |  |  |  |  |  |  |  |  |  |  |  |  |  |  |  |  |  |  |  |  |  |  |  |  |  |  |  |  |  |  |  |  |  |  |  |  |  |  |  |  |  |  |  |
| LION:0002948 | fatty acid with 16-18 carbons |  |  | x | x | x | x |  |  |  | x |  |  |  |  |  | x | x | x | x |  | x | x | x | x |  |  |  |  | x | x | x | x | x | x | x |  |  |  |  | x | x | x | x |  |  |  | x |  |
| LION:0002949 | fatty acid with 19-21 carbons |  |  |  |  |  |  |  |  |  |  |  |  |  |  |  |  |  |  |  |  |  |  |  |  |  |  |  |  |  |  |  |  |  |  | x |  |  |  |  |  |  |  |  |  |  |  |  |  |
| LION:0002950 | fatty acid with 22-24 carbons | x |  |  |  |  |  |  |  |  |  |  |  |  |  |  |  |  |  | x |  |  | x | x | x |  |  |  |  |  |  |  |  |  |  |  |  |  |  |  |  |  |  |  |  |  |  |  |  |
| LION:0002953 | fatty acid with 14 carbons |  | x | x | x | x |  |  |  |  |  |  |  |  |  |  |  |  |  |  |  |  |  |  |  |  |  |  |  |  |  |  |  |  |  |  |  |  |  |  |  |  |  |  |  |  |  |  |  |
| LION:0002955 | fatty acid with 16 carbons |  |  | x | x | x | x |  |  |  | x |  |  |  |  |  | x | x | x |  |  | x |  |  |  |  |  |  |  |  |  |  |  |  |  |  |  |  |  |  | x | x |  |  |  |  |  |  |  |
| LION:0002957 | fatty acid with 18 carbons |  |  |  |  |  |  |  |  |  |  |  |  |  |  |  |  | x | x | x |  | x | x | x | x |  |  |  |  | x | x | x | x | x | x | x |  |  |  |  | x | x | x | x |  |  |  | x |  |
| LION:0002959 | fatty acid with 20 carbons |  |  |  |  |  |  |  |  |  |  |  |  |  |  |  |  |  |  |  |  |  |  |  |  |  |  |  |  |  |  |  |  |  |  | x |  |  |  |  |  |  |  |  |  |  |  |  |  |
| LION:0002961 | fatty acid with 22 carbons |  |  |  |  |  |  |  |  |  |  |  |  |  |  |  |  |  |  | x |  |  | x | x | x |  |  |  |  |  |  |  |  |  |  |  |  |  |  |  |  |  |  |  |  |  |  |  |  |
| LION:0002963 | fatty acid with 24 carbons | x |  |  |  |  |  |  |  |  |  |  |  |  |  |  |  |  |  |  |  |  |  |  |  |  |  |  |  |  |  |  |  |  |  |  |  |  |  |  |  |  |  |  |  |  |  |  |  |
| LION:0002966 | fatty acid with less than 2 double bonds | x | x | x | x | x | x |  |  |  | x |  |  |  |  |  | x | x | x | x |  | x | x | x | x |  |  |  |  | x | x | x | x | x | x | x |  |  |  |  | x | x | x | x |  |  |  | x |  |
| LION:0002967 | polyunsaturated fatty acid |  |  |  |  |  |  |  |  |  |  |  |  |  |  |  |  |  |  | x |  |  | x | x | x |  |  |  |  |  |  |  |  |  |  |  |  |  |  |  |  |  |  |  |  |  |  |  |  |
| LION:0002968 | saturated fatty acid |  | x | x | x | x |  |  |  |  | x |  |  |  |  |  | x | x |  |  |  | x | x |  | x |  |  |  |  |  |  |  |  |  |  |  |  |  |  |  | x | x | x | x |  |  |  |  |  |
| LION:0002969 | monounsaturated fatty acid | x |  |  | x | x | x |  |  |  | x |  |  |  |  |  | x | x | x | x |  | x |  | x |  |  |  |  |  | x | x | x | x | x | x | x |  |  |  |  | x | x | x | x |  |  |  | x |  |
| LION:0002970 | fatty acid with 2 double bonds |  |  |  |  |  |  |  |  |  |  |  |  |  |  |  |  |  |  |  |  |  | x |  |  |  |  |  |  |  |  |  |  |  |  |  |  |  |  |  |  |  |  |  |  |  |  |  |  |
| LION:0002971 | fatty acid with 3 double bonds |  |  |  |  |  |  |  |  |  |  |  |  |  |  |  |  |  |  | x |  |  |  | x | x |  |  |  |  |  |  |  |  |  |  |  |  |  |  |  |  |  |  |  |  |  |  |  |  |
| LION:0002976 | fatty acid with more than 3 double bonds |  |  |  |  |  |  |  |  |  |  |  |  |  |  |  |  |  |  | x |  |  |  | x | x |  |  |  |  |  |  |  |  |  |  |  |  |  |  |  |  |  |  |  |  |  |  |  |  |
| LION:0002977 | fatty acid with 3-5 double bonds |  |  |  |  |  |  |  |  |  |  |  |  |  |  |  |  |  |  | x |  |  |  | x | x |  |  |  |  |  |  |  |  |  |  |  |  |  |  |  |  |  |  |  |  |  |  |  |  |
| LION:0002993 | PC(14:0/14:0) |  | x |  |  |  |  |  |  |  |  |  |  |  |  |  |  |  |  |  |  |  |  |  |  |  |  |  |  |  |  |  |  |  |  |  |  |  |  |  |  |  |  |  |  |  |  |  |  |
| LION:0002996 | PC(14:0/16:1) |  |  |  | x |  |  |  |  |  |  |  |  |  |  |  |  |  |  |  |  |  |  |  |  |  |  |  |  |  |  |  |  |  |  |  |  |  |  |  |  |  |  |  |  |  |  |  |  |
| LION:0002998 | PC(14:0/16:0) |  |  | x |  |  |  |  |  |  |  |  |  |  |  |  |  |  |  |  |  |  |  |  |  |  |  |  |  |  |  |  |  |  |  |  |  |  |  |  |  |  |  |  |  |  |  |  |  |
| LION:0003004 | PC(16:1/16:1) |  |  |  |  |  | x |  |  |  |  |  |  |  |  |  |  |  |  |  |  |  |  |  |  |  |  |  |  |  |  |  |  |  |  |  |  |  |  |  |  |  |  |  |  |  |  |  |  |
| LION:0003542 | PE(16:0/16:1) |  |  |  |  |  |  |  |  |  |  |  |  |  |  |  | x |  |  |  |  |  |  |  |  |  |  |  |  |  |  |  |  |  |  |  |  |  |  |  |  |  |  |  |  |  |  |  |  |
| LION:0003551 | PE(16:1/18:1) |  |  |  |  |  |  |  |  |  |  |  |  |  |  |  |  |  | x |  |  |  |  |  |  |  |  |  |  |  |  |  |  |  |  |  |  |  |  |  |  |  |  |  |  |  |  |  |  |
| LION:0003554 | PE(16:0/18:1) |  |  |  |  |  |  |  |  |  |  |  |  |  |  |  |  | x |  |  |  |  |  |  |  |  |  |  |  |  |  |  |  |  |  |  |  |  |  |  |  |  |  |  |  |  |  |  |  |
| LION:0003741 | PS(18:0/22:3) |  |  |  |  |  |  |  |  |  |  |  |  |  |  |  |  |  |  |  |  |  |  |  | x |  |  |  |  |  |  |  |  |  |  |  |  |  |  |  |  |  |  |  |  |  |  |  |  |
| LION:0004460 | DG(18:1/20:1) |  |  |  |  |  |  |  |  |  |  |  |  |  |  |  |  |  |  |  |  |  |  |  |  |  |  |  |  |  |  |  |  |  |  | x |  |  |  |  |  |  |  |  |  |  |  |  |  |
| LION:0005133 | TG(16:0/18:0/18:1) |  |  |  |  |  |  |  |  |  |  |  |  |  |  |  |  |  |  |  |  |  |  |  |  |  |  |  |  |  |  |  |  |  |  |  |  |  |  |  | x |  |  |  |  |  |  |  |  |
| LION:0005152 | TG(16:0/18:1/18:1) |  |  |  |  |  |  |  |  |  |  |  |  |  |  |  |  |  |  |  |  |  |  |  |  |  |  |  |  |  |  |  |  |  |  |  |  |  |  |  |  | x |  |  |  |  |  |  |  |
| LION:0005290 | TG(18:0/18:0/18:1) |  |  |  |  |  |  |  |  |  |  |  |  |  |  |  |  |  |  |  |  |  |  |  |  |  |  |  |  |  |  |  |  |  |  |  |  |  |  |  |  |  | x |  |  |  |  |  |  |
| LION:0005320 | TG(18:0/18:1/18:1) |  |  |  |  |  |  |  |  |  |  |  |  |  |  |  |  |  |  |  |  |  |  |  |  |  |  |  |  |  |  |  |  |  |  |  |  |  |  |  |  |  |  | x |  |  |  |  |  |
| LION:0005353 | TG(18:1/18:1/18:1) |  |  |  |  |  |  |  |  |  |  |  |  |  |  |  |  |  |  |  |  |  |  |  |  |  |  |  |  |  |  |  |  |  |  |  |  |  |  |  |  |  |  |  |  |  |  | x |  |
| LION:0012009 | lipid-mediated signalling |  |  |  |  |  |  |  |  |  |  |  |  |  |  |  |  |  |  |  |  |  |  |  |  |  |  |  |  | x | x | x |  |  |  | x | x | x |  |  |  |  |  |  |  |  |  |  |  |
| LION:0012010 | membrane component |  | x | x | x | x | x | x | x | x | x | x | x | x | x | x | x | x | x | x | x | x | x | x | x | x | x | x | x |  |  |  |  |  | x | x | x | x | x | x |  |  |  |  |  |  |  |  |  |
| LION:0012011 | lipid storage |  |  |  |  |  |  |  |  |  |  |  |  |  |  |  |  |  |  |  |  |  |  |  |  |  |  |  |  |  |  |  |  |  |  |  |  |  |  |  | x | x | x | x | x | x | x | x | x |
| LION:0012018 | dihexosylceramides |  |  |  |  |  |  |  |  |  |  |  |  |  |  |  |  |  |  |  |  |  |  |  |  |  |  |  |  |  |  |  |  |  | x |  |  |  |  |  |  |  |  |  |  |  |  |  |  |
| LION:0012080 | endoplasmic reticulum (ER) |  | x | x | x | x | x | x | x | x | x | x | x | x | x | x | x | x | x | x | x | x | x | x |  |  |  |  |  | x | x | x |  |  |  |  |  |  |  |  |  |  |  |  |  |  |  |  |  |
| LION:0012081 | mitochondrion |  |  |  |  |  |  |  |  |  |  |  |  |  |  |  | x | x | x | x | x | x | x | x |  |  |  |  |  |  |  |  |  |  |  |  |  |  | x | x |  |  |  |  |  |  |  |  |  |
| LION:0012082 | plasma membrane |  |  |  |  |  |  |  |  |  |  |  |  |  |  |  |  |  |  |  |  |  |  |  | x | x | x | x | x | x | x | x |  |  |  |  |  |  |  |  |  |  |  |  |  |  |  |  |  |
| LION:0012084 | lipid droplet |  |  |  |  |  |  |  |  |  |  |  |  |  |  |  |  |  |  |  |  |  |  |  |  |  |  |  |  |  |  |  |  |  |  |  |  |  |  |  | x | x | x | x | x | x | x | x | x |
| LION:0012085 | golgi apparatus |  |  |  |  |  |  |  |  |  |  |  |  |  |  |  |  |  |  |  |  |  |  |  |  | x | x | x | x |  |  |  |  |  |  |  |  |  |  |  |  |  |  |  |  |  |  |  |  |
| LION:0012086 | endosome/lysosome |  |  |  |  |  |  |  |  |  |  |  |  |  |  |  |  |  |  |  |  |  |  |  |  | x | x | x | x |  |  |  |  |  |  |  |  |  |  |  |  |  |  |  |  |  |  |  |  |
| LION:0012441 | N-acylsphingosines (ceramides) [SP0201] |  |  |  |  |  |  |  |  |  |  |  |  |  |  |  |  |  |  |  |  |  |  |  |  |  |  |  |  | x | x | x |  |  |  |  |  |  |  |  |  |  |  |  |  |  |  |  |  |
| LION:0016003 | PE(P-40:2) |  |  |  |  |  |  |  |  |  |  |  |  |  |  |  |  |  |  |  |  |  | x |  |  |  |  |  |  |  |  |  |  |  |  |  |  |  |  |  |  |  |  |  |  |  |  |  |  |
| LION:0016009 | PE(P-40:4) |  |  |  |  |  |  |  |  |  |  |  |  |  |  |  |  |  |  |  |  |  |  | x |  |  |  |  |  |  |  |  |  |  |  |  |  |  |  |  |  |  |  |  |  |  |  |  |  |
| LION:0017306 | PE(O-16:0/18:1) |  |  |  |  |  |  |  |  |  |  |  |  |  |  |  |  |  |  |  |  | x |  |  |  |  |  |  |  |  |  |  |  |  |  |  |  |  |  |  |  |  |  |  |  |  |  |  |  |
| LION:0019464 | PE(P-18:0/22:2) |  |  |  |  |  |  |  |  |  |  |  |  |  |  |  |  |  |  |  |  |  | x |  |  |  |  |  |  |  |  |  |  |  |  |  |  |  |  |  |  |  |  |  |  |  |  |  |  |
| LION:0019470 | PE(P-18:1/22:3) |  |  |  |  |  |  |  |  |  |  |  |  |  |  |  |  |  |  |  |  |  |  | x |  |  |  |  |  |  |  |  |  |  |  |  |  |  |  |  |  |  |  |  |  |  |  |  |  |
| LION:0041232 | PC(29:0) |  |  |  |  |  |  |  | x |  |  |  |  |  |  |  |  |  |  |  |  |  |  |  |  |  |  |  |  |  |  |  |  |  |  |  |  |  |  |  |  |  |  |  |  |  |  |  |  |
| LION:0080664 | PE(18:1/22:3) |  |  |  |  |  |  |  |  |  |  |  |  |  |  |  |  |  |  | x |  |  |  |  |  |  |  |  |  |  |  |  |  |  |  |  |  |  |  |  |  |  |  |  |  |  |  |  |  |
| LION:0080968 | very low bilayer thickness |  | x |  | x |  |  | x |  | x |  |  |  |  |  |  |  |  |  |  |  |  |  |  |  |  |  |  |  |  |  |  |  |  |  |  |  |  |  |  |  |  |  |  |  |  |  |  |  |
| LION:0080969 | low bilayer thickness |  |  | x | x |  | x |  | x |  |  |  |  |  |  |  |  |  |  |  |  |  |  |  |  |  |  |  |  |  |  |  |  |  |  |  |  |  |  |  |  |  |  |  |  |  |  |  |  |
| LION:0080970 | average bilayer thickness |  |  |  |  |  |  |  |  |  |  |  |  |  |  |  | x |  | x |  |  |  |  |  |  |  |  |  |  |  |  |  |  |  |  |  |  |  |  |  |  |  |  |  |  |  |  |  |  |
| LION:0080971 | high bilayer thickness |  |  |  |  |  |  |  |  |  |  |  |  |  |  |  |  | x |  | x | x |  |  |  | x |  |  |  |  |  |  |  |  |  |  |  |  |  |  |  |  |  |  |  |  |  |  |  |  |
| LION:0080973 | below average bilayer thickness |  | x | x | x |  | x | x | x | x |  |  |  |  |  |  |  |  |  |  |  |  |  |  |  |  |  |  |  |  |  |  |  |  |  |  |  |  |  |  |  |  |  |  |  |  |  |  |  |
| LION:0080974 | above average bilayer thickness |  |  |  |  |  |  |  |  |  |  |  |  |  |  |  |  | x |  | x | x |  |  |  | x |  |  |  |  |  |  |  |  |  |  |  |  |  |  |  |  |  |  |  |  |  |  |  |  |
| LION:0080976 | very low lateral diffusion |  |  |  |  |  |  |  |  |  |  |  |  |  |  |  |  |  |  |  | x |  |  |  |  |  |  |  |  |  |  |  |  |  |  |  |  |  |  |  |  |  |  |  |  |  |  |  |  |
| LION:0080977 | low lateral diffusion |  |  |  |  |  |  |  |  |  |  |  |  |  |  |  | x | x |  |  |  |  |  |  | x |  |  |  |  |  |  |  |  |  |  |  |  |  |  |  |  |  |  |  |  |  |  |  |  |
| LION:0080978 | average lateral diffusion |  | x | x |  |  |  |  | x |  |  |  |  |  |  |  | x |  | x | x |  |  |  |  |  |  |  |  |  |  |  |  |  |  |  |  |  |  |  |  |  |  |  |  |  |  |  |  |  |
| LION:0080979 | high lateral diffusion |  | x |  | x |  | x | x |  | x |  |  |  |  |  |  |  |  |  |  |  |  |  |  |  |  |  |  |  |  |  |  |  |  |  |  |  |  |  |  |  |  |  |  |  |  |  |  |  |
| LION:0080980 | very high lateral diffusion |  |  |  |  |  |  |  |  | x |  |  |  |  |  |  |  |  |  |  |  |  |  |  |  |  |  |  |  |  |  |  |  |  |  |  |  |  |  |  |  |  |  |  |  |  |  |  |  |
| LION:0080981 | below average lateral diffusion |  |  |  |  |  |  |  |  |  |  |  |  |  |  |  | x | x |  |  | x |  |  |  | x |  |  |  |  |  |  |  |  |  |  |  |  |  |  |  |  |  |  |  |  |  |  |  |  |
| LION:0080982 | above average lateral diffusion |  | x |  | x |  | x | x |  | x |  |  |  |  |  |  |  |  |  |  |  |  |  |  |  |  |  |  |  |  |  |  |  |  |  |  |  |  |  |  |  |  |  |  |  |  |  |  |  |

Table 5e: LION term associations:

| LION-term | LION-name | nr_lipids | lipid identifiers |
| --- | --- | --- | --- |
| all | all | 48 | FA 24:1; PC 14:0_14:0; PC 14:0_16:0; PC 14:0_16:1; PC 14:0_16:1_iso; PC 16:1_16:1; PC 28:1; PC 29:0; PC 30:2; PC O-16:0_16:1; PC O-30:0; PC O-30:1; PC O-32:2; PC O-34:3; PC O-40:2; PE 16:0_16:1; PE 16:0_18:1; PE 16:1_18:1; PE 18:1_22:3; PE 34:0; PE O-16:0_18:1; PE P-18:0_22:2; PE P-18:1_22:3; PS 18:0_22:3; SM d40:0; SM d40:1; SM d42:1; SM d42:2; Cer 18:1;O2/16:0; Cer 18:1;O2/24:0; Cer 18:1;O2/24:1; Hex-Cer 18:1;O2/24:0; Hex-Cer 18:1;O2/24:1; Hex2Cer 18:1;O2/24:1; DG 18:1_20:1; DG 34:2; DG 38:4; CL 72:4; CL 72:5; TG 16:0_18:0_18:1; TG 16:0_18:1_18:1; TG 18:0_18:0_18:1; TG 18:0_18:1_18:1; TG 18:0_38:3; TG 18:0_42:3; TG 18:0_48:2; TG 18:1_18:1_18:1; TG 20:0_36:2 |
| CAT:0000000 | lipid classification | 46 | FA 24:1; PC 14:0_14:0; PC 14:0_16:0; PC 14:0_16:1; PC 14:0_16:1_iso; PC 16:1_16:1; PC 28:1; PC 29:0; PC 30:2; PC O-16:0_16:1; PC O-30:0; PC O-30:1; PC O-32:2; PC O-34:3; PC O-40:2; PE 16:0_16:1; PE 16:0_18:1; PE 16:1_18:1; PE 18:1_22:3; PE 34:0; PE O-16:0_18:1; PE P-18:0_22:2; PE P-18:1_22:3; PS 18:0_22:3; SM d40:0; SM d40:1; SM d42:1; SM d42:2; Cer 18:1;O2/16:0; Cer 18:1;O2/24:0; Cer 18:1;O2/24:1; Hex2Cer 18:1;O2/24:1; DG 18:1_20:1; DG 34:2; DG 38:4; CL 72:4; CL 72:5; TG 16:0_18:0_18:1; TG 16:0_18:1_18:1; TG 18:0_18:0_18:1; TG 18:0_18:1_18:1; TG 18:0_38:3; TG 18:0_42:3; TG 18:0_48:2; TG 18:1_18:1_18:1; TG 20:0_36:2 |
| CAT:0000091 | physical or chemical properties | 48 | FA 24:1; PC 14:0_14:0; PC 14:0_16:0; PC 14:0_16:1; PC 14:0_16:1_iso; PC 16:1_16:1; PC 28:1; PC 29:0; PC 30:2; PC O-16:0_16:1; PC O-30:0; PC O-30:1; PC O-32:2; PC O-34:3; PC O-40:2; PE 16:0_16:1; PE 16:0_18:1; PE 16:1_18:1; PE 18:1_22:3; PE 34:0; PE O-16:0_18:1; PE P-18:0_22:2; PE P-18:1_22:3; PS 18:0_22:3; SM d40:0; SM d40:1; SM d42:1; SM d42:2; Cer 18:1;O2/16:0; Cer 18:1;O2/24:0; Cer 18:1;O2/24:1; Hex-Cer 18:1;O2/24:0; Hex-Cer 18:1;O2/24:1; Hex2Cer 18:1;O2/24:1; DG 18:1_20:1; DG 34:2; DG 38:4; CL 72:4; CL 72:5; TG 16:0_18:0_18:1; TG 16:0_18:1_18:1; TG 18:0_18:0_18:1; TG 18:0_18:1_18:1; TG 18:0_38:3; TG 18:0_42:3; TG 18:0_48:2; TG 18:1_18:1_18:1; TG 20:0_36:2 |
| CAT:0000092 | charge headgroup | 43 | FA 24:1; PC 14:0_14:0; PC 14:0_16:0; PC 14:0_16:1; PC 14:0_16:1_iso; PC 16:1_16:1; PC 28:1; PC 29:0; PC 30:2; PC O-16:0_16:1; PC O-30:0; PC O-30:1; PC O-32:2; PC O-34:3; PC O-40:2; PE 16:0_16:1; PE 16:0_18:1; PE 16:1_18:1; PE 18:1_22:3; PE 34:0; PE O-16:0_18:1; PE P-18:0_22:2; PE P-18:1_22:3; PS 18:0_22:3; SM d40:0; SM d40:1; SM d42:1; SM d42:2; Hex2Cer 18:1;O2/24:1; DG 18:1_20:1; DG 34:2; DG 38:4; CL 72:4; CL 72:5; TG 16:0_18:0_18:1; TG 16:0_18:1_18:1; TG 18:0_18:0_18:1; TG 18:0_18:1_18:1; TG 18:0_38:3; TG 18:0_42:3; TG 18:0_48:2; TG 18:1_18:1_18:1; TG 20:0_36:2 |
| CAT:0000100 | contains fatty acid | 27 | FA 24:1; PC 14:0_14:0; PC 14:0_16:0; PC 14:0_16:1; PC 14:0_16:1_iso; PC 16:1_16:1; PC O-16:0_16:1; PE 16:0_16:1; PE 16:0_18:1; PE 16:1_18:1; PE 18:1_22:3; PE O-16:0_18:1; PE P-18:0_22:2; PE P-18:1_22:3; PS 18:0_22:3; Cer 18:1;O2/16:0; Cer 18:1;O2/24:0; Cer 18:1;O2/24:1; Hex-Cer 18:1;O2/24:0; Hex-Cer 18:1;O2/24:1; Hex2Cer 18:1;O2/24:1; DG 18:1_20:1; TG 16:0_18:0_18:1; TG 16:0_18:1_18:1; TG 18:0_18:0_18:1; TG 18:0_18:1_18:1; TG 18:1_18:1_18:1 |
| CAT:0000123 | type by bond | 9 | PC O-16:0_16:1; PC O-30:0; PC O-30:1; PC O-32:2; PC O-34:3; PC O-40:2; PE O-16:0_18:1; PE P-18:0_22:2; PE P-18:1_22:3 |
| CAT:0000463 | intrinsic curvature | 31 | PC 14:0_14:0; PC 14:0_16:0; PC 14:0_16:1; PC 14:0_16:1_iso; PC 16:1_16:1; PC 28:1; PC 29:0; PC 30:2; PC O-16:0_16:1; PC O-30:0; PC O-30:1; PC O-32:2; PC O-34:3; PC O-40:2; PE 16:0_16:1; PE 16:0_18:1; PE 16:1_18:1; PE 18:1_22:3; PE 34:0; PE O-16:0_18:1; PE P-18:0_22:2; PE P-18:1_22:3; PS 18:0_22:3; Cer 18:1;O2/16:0; Cer 18:1;O2/24:0; Cer 18:1;O2/24:1; DG 18:1_20:1; DG 34:2; DG 38:4; CL 72:4; CL 72:5 |
| CAT:0001734 | chain-melting transition temperature | 17 | PC 14:0_14:0; PC 14:0_16:0; PC 14:0_16:1; PC 16:1_16:1; PC 28:1; PC 29:0; PC 30:2; PE 16:0_16:1; PE 16:0_18:1; PE 16:1_18:1; PE 18:1_22:3; PE 34:0; PS 18:0_22:3; SM d40:0; SM d40:1; SM d42:1; SM d42:2 |
| CAT:0002945 | fatty acid unsaturation | 27 | FA 24:1; PC 14:0_14:0; PC 14:0_16:0; PC 14:0_16:1; PC 14:0_16:1_iso; PC 16:1_16:1; PC O-16:0_16:1; PE 16:0_16:1; PE 16:0_18:1; PE 16:1_18:1; PE 18:1_22:3; PE O-16:0_18:1; PE P-18:0_22:2; PE P-18:1_22:3; PS 18:0_22:3; Cer 18:1;O2/16:0; Cer 18:1;O2/24:0; Cer 18:1;O2/24:1; Hex-Cer 18:1;O2/24:0; Hex-Cer 18:1;O2/24:1; Hex2Cer 18:1;O2/24:1; DG 18:1_20:1; TG 16:0_18:0_18:1; TG 16:0_18:1_18:1; TG 18:0_18:0_18:1; TG 18:0_18:1_18:1; TG 18:1_18:1_18:1 |
| CAT:0002946 | fatty acid chain length | 27 | FA 24:1; PC 14:0_14:0; PC 14:0_16:0; PC 14:0_16:1; PC 14:0_16:1_iso; PC 16:1_16:1; PC O-16:0_16:1; PE 16:0_16:1; PE 16:0_18:1; PE 16:1_18:1; PE 18:1_22:3; PE O-16:0_18:1; PE P-18:0_22:2; PE P-18:1_22:3; PS 18:0_22:3; Cer 18:1;O2/16:0; Cer 18:1;O2/24:0; Cer 18:1;O2/24:1; Hex-Cer 18:1;O2/24:0; Hex-Cer 18:1;O2/24:1; Hex2Cer 18:1;O2/24:1; DG 18:1_20:1; TG 16:0_18:0_18:1; TG 16:0_18:1_18:1; TG 18:0_18:0_18:1; TG 18:0_18:1_18:1; TG 18:1_18:1_18:1 |
| CAT:0012007 | function | 45 | PC 14:0_14:0; PC 14:0_16:0; PC 14:0_16:1; PC 14:0_16:1_iso; PC 16:1_16:1; PC 28:1; PC 29:0; PC 30:2; PC O-16:0_16:1; PC O-30:0; PC O-30:1; PC O-32:2; PC O-34:3; PC O-40:2; PE 16:0_16:1; PE 16:0_18:1; PE 16:1_18:1; PE 18:1_22:3; PE 34:0; PE O-16:0_18:1; PE P-18:0_22:2; PE P-18:1_22:3; PS 18:0_22:3; SM d40:0; SM d40:1; SM d42:1; SM d42:2; Cer 18:1;O2/16:0; Cer 18:1;O2/24:0; Cer 18:1;O2/24:1; Hex2Cer 18:1;O2/24:1; DG 18:1_20:1; DG 34:2; DG 38:4; CL 72:4; CL 72:5; TG 16:0_18:0_18:1; TG 16:0_18:1_18:1; TG 18:0_18:0_18:1; TG 18:0_18:1_18:1; TG 18:0_38:3; TG 18:0_42:3; TG 18:0_48:2; TG 18:1_18:1_18:1; TG 20:0_36:2 |
| CAT:0012008 | cellular component | 41 | PC 14:0_14:0; PC 14:0_16:0; PC 14:0_16:1; PC 14:0_16:1_iso; PC 16:1_16:1; PC 28:1; PC 29:0; PC 30:2; PC O-16:0_16:1; PC O-30:0; PC O-30:1; PC O-32:2; PC O-34:3; PC O-40:2; PE 16:0_16:1; PE 16:0_18:1; PE 16:1_18:1; PE 18:1_22:3; PE 34:0; PE O-16:0_18:1; PE P-18:0_22:2; PE P-18:1_22:3; PS 18:0_22:3; SM d40:0; SM d40:1; SM d42:1; SM d42:2; Cer 18:1;O2/16:0; Cer 18:1;O2/24:0; Cer 18:1;O2/24:1; CL 72:4; CL 72:5; TG 16:0_18:0_18:1; TG 16:0_18:1_18:1; TG 18:0_18:0_18:1; TG 18:0_18:1_18:1; TG 18:0_38:3; TG 18:0_42:3; TG 18:0_48:2; TG 18:1_18:1_18:1; TG 20:0_36:2 |
| CAT:0080950 | lateral diffusion | 13 | PC 14:0_14:0; PC 14:0_16:0; PC 14:0_16:1; PC 16:1_16:1; PC 28:1; PC 29:0; PC 30:2; PE 16:0_16:1; PE 16:0_18:1; PE 16:1_18:1; PE 18:1_22:3; PE 34:0; PS 18:0_22:3 |
| CAT:0080951 | bilayer thickness | 13 | PC 14:0_14:0; PC 14:0_16:0; PC 14:0_16:1; PC 16:1_16:1; PC 28:1; PC 29:0; PC 30:2; PE 16:0_16:1; PE 16:0_18:1; PE 16:1_18:1; PE 18:1_22:3; PE 34:0; PS 18:0_22:3 |
| LION:0000001 | fatty acids [FA] | 1 | FA 24:1 |
| LION:0000002 | glycerolipids [GL] | 12 | DG 18:1_20:1; DG 34:2; DG 38:4; TG 16:0_18:0_18:1; TG 16:0_18:1_18:1; TG 18:0_18:0_18:1; TG 18:0_18:1_18:1; TG 18:0_38:3; TG 18:0_42:3; TG 18:0_48:2; TG 18:1_18:1_18:1; TG 20:0_36:2 |
| LION:0000003 | glycerophospholipids [GP] | 25 | PC 14:0_14:0; PC 14:0_16:0; PC 14:0_16:1; PC 14:0_16:1_iso; PC 16:1_16:1; PC 28:1; PC 29:0; PC 30:2; PC O-16:0_16:1; PC O-30:0; PC O-30:1; PC O-32:2; PC O-34:3; PC O-40:2; PE 16:0_16:1; PE 16:0_18:1; PE 16:1_18:1; PE 18:1_22:3; PE 34:0; PE O-16:0_18:1; PE P-18:0_22:2; PE P-18:1_22:3; PS 18:0_22:3; CL 72:4; CL 72:5 |
| LION:0000004 | sphingolipids [SP] | 8 | SM d40:0; SM d40:1; SM d42:1; SM d42:2; Cer 18:1;O2/16:0; Cer 18:1;O2/24:0; Cer 18:1;O2/24:1; Hex2Cer 18:1;O2/24:1 |
| LION:0000010 | glycerophosphocholines [GP01] | 14 | PC 14:0_14:0; PC 14:0_16:0; PC 14:0_16:1; PC 14:0_16:1_iso; PC 16:1_16:1; PC 28:1; PC 29:0; PC 30:2; PC O-16:0_16:1; PC O-30:0; PC O-30:1; PC O-32:2; PC O-34:3; PC O-40:2 |
| LION:0000011 | glycerophosphoethanolamines [GP02] | 8 | PE 16:0_16:1; PE 16:0_18:1; PE 16:1_18:1; PE 18:1_22:3; PE 34:0; PE O-16:0_18:1; PE P-18:0_22:2; PE P-18:1_22:3 |
| LION:0000013 | glycerophosphoserines [GP03] | 1 | PS 18:0_22:3 |
| LION:0000021 | glycerophosphoglycerophosphoglycerols [GP12] | 2 | CL 72:4; CL 72:5 |
| LION:0000030 | diacylglycerophosphocholines [GP0101] | 8 | PC 14:0_14:0; PC 14:0_16:0; PC 14:0_16:1; PC 14:0_16:1_iso; PC 16:1_16:1; PC 28:1; PC 29:0; PC 30:2 |
| LION:0000031 | 1-alkyl,2-acylglycerophosphocholines [GP0102] | 6 | PC O-16:0_16:1; PC O-30:0; PC O-30:1; PC O-32:2; PC O-34:3; PC O-40:2 |
| LION:0000038 | diacylglycerophosphoethanolamines [GP0201] | 5 | PE 16:0_16:1; PE 16:0_18:1; PE 16:1_18:1; PE 18:1_22:3; PE 34:0 |
| LION:0000039 | 1-alkyl,2-acylglycerophosphoethanolamines [GP0202] | 1 | PE O-16:0_18:1 |
| LION:0000040 | 1-(1z-alkenyl),2-acylglycerophosphoethanolamines [GP0203] | 2 | PE P-18:0_22:2; PE P-18:1_22:3 |
| LION:0000053 | diacylglycerophosphoserines [GP0301] | 1 | PS 18:0_22:3 |
| LION:0000077 | ceramides [SP02] | 3 | Cer 18:1;O2/16:0; Cer 18:1;O2/24:0; Cer 18:1;O2/24:1 |
| LION:0000078 | phosphosphingolipids [SP03] | 4 | SM d40:0; SM d40:1; SM d42:1; SM d42:2 |
| LION:0000080 | neutral glycosphingolipids [SP05] | 1 | Hex2Cer 18:1;O2/24:1 |
| LION:0000084 | ceramide phosphocholines (sphingomyelins) [SP0301] | 4 | SM d40:0; SM d40:1; SM d42:1; SM d42:2 |
| LION:0000093 | headgroup with negative charge | 4 | FA 24:1; PS 18:0_22:3; CL 72:4; CL 72:5 |
| LION:0000094 | headgroup with neutral charge | 13 | Hex2Cer 18:1;O2/24:1; DG 18:1_20:1; DG 34:2; DG 38:4; TG 16:0_18:0_18:1; TG 16:0_18:1_18:1; TG 18:0_18:0_18:1; TG 18:0_18:1_18:1; TG 18:0_38:3; TG 18:0_42:3; TG 18:0_48:2; TG 18:1_18:1_18:1; TG 20:0_36:2 |
| LION:0000095 | headgroup with positive charge / zwitter-ion | 26 | PC 14:0_14:0; PC 14:0_16:0; PC 14:0_16:1; PC 14:0_16:1_iso; PC 16:1_16:1; PC 28:1; PC 29:0; PC 30:2; PC O-16:0_16:1; PC O-30:0; PC O-30:1; PC O-32:2; PC O-34:3; PC O-40:2; PE 16:0_16:1; PE 16:0_18:1; PE 16:1_18:1; PE 18:1_22:3; PE 34:0; PE O-16:0_18:1; PE P-18:0_22:2; PE P-18:1_22:3; SM d40:0; SM d40:1; SM d42:1; SM d42:2 |
| LION:0000100 | fatty acid with 18 carbons or less | 26 | PC 14:0_14:0; PC 14:0_16:0; PC 14:0_16:1; PC 14:0_16:1_iso; PC 16:1_16:1; PC O-16:0_16:1; PE 16:0_16:1; PE 16:0_18:1; PE 16:1_18:1; PE 18:1_22:3; PE O-16:0_18:1; PE P-18:0_22:2; PE P-18:1_22:3; PS 18:0_22:3; Cer 18:1;O2/16:0; Cer 18:1;O2/24:0; Cer 18:1;O2/24:1; Hex-Cer 18:1;O2/24:0; Hex-Cer 18:1;O2/24:1; Hex2Cer 18:1;O2/24:1; DG 18:1_20:1; TG 16:0_18:0_18:1; TG 16:0_18:1_18:1; TG 18:0_18:0_18:1; TG 18:0_18:1_18:1; TG 18:1_18:1_18:1 |
| LION:0000104 | PC(O-16:0/16:1) | 1 | PC O-16:0_16:1 |
| LION:0000140 | PC(28:0) | 1 | PC 14:0_14:0 |
| LION:0000143 | PC(O-30:0) | 1 | PC O-30:0 |
| LION:0000145 | PC(30:2) | 1 | PC 30:2 |
| LION:0000146 | PC(30:1) | 1 | PC 14:0_16:1 |
| LION:0000147 | PC(30:0) | 1 | PC 14:0_16:0 |
| LION:0000151 | PC(O-32:2) | 1 | PC O-32:2 |
| LION:0000152 | PC(O-32:1) | 1 | PC O-16:0_16:1 |
| LION:0000157 | PC(32:2) | 1 | PC 16:1_16:1 |
| LION:0000163 | PC(O-34:3) | 1 | PC O-34:3 |
| LION:0000259 | C14:0 | 4 | PC 14:0_14:0; PC 14:0_16:0; PC 14:0_16:1; PC 14:0_16:1_iso |
| LION:0000324 | PE(32:1) | 1 | PE 16:0_16:1 |
| LION:0000331 | PE(34:2) | 1 | PE 16:1_18:1 |
| LION:0000332 | PE(34:1) | 1 | PE 16:0_18:1 |
| LION:0000333 | PE(34:0) | 1 | PE 34:0 |
| LION:0000378 | PS(40:3) | 1 | PS 18:0_22:3 |
| LION:0000409 | SM(40:1) | 1 | SM d40:1 |
| LION:0000410 | SM(40:0) | 1 | SM d40:0 |
| LION:0000414 | SM(42:2) | 1 | SM d42:2 |
| LION:0000415 | SM(42:1) | 1 | SM d42:1 |
| LION:0000453 | simple glc series [SP0501] | 1 | Hex2Cer 18:1;O2/24:1 |
| LION:0000464 | negative intrinsic curvature | 16 | PE 16:0_16:1; PE 16:0_18:1; PE 16:1_18:1; PE 18:1_22:3; PE 34:0; PE O-16:0_18:1; PE P-18:0_22:2; PE P-18:1_22:3; Cer 18:1;O2/16:0; Cer 18:1;O2/24:0; Cer 18:1;O2/24:1; DG 18:1_20:1; DG 34:2; DG 38:4; CL 72:4; CL 72:5 |
| LION:0000465 | neutral intrinsic curvature | 15 | PC 14:0_14:0; PC 14:0_16:0; PC 14:0_16:1; PC 14:0_16:1_iso; PC 16:1_16:1; PC 28:1; PC 29:0; PC 30:2; PC O-16:0_16:1; PC O-30:0; PC O-30:1; PC O-32:2; PC O-34:3; PC O-40:2; PS 18:0_22:3 |
| LION:0000467 | contains ether-bond | 7 | PC O-16:0_16:1; PC O-30:0; PC O-30:1; PC O-32:2; PC O-34:3; PC O-40:2; PE O-16:0_18:1 |
| LION:0000603 | diradylglycerols [GL02] | 3 | DG 18:1_20:1; DG 34:2; DG 38:4 |
| LION:0000604 | triradylglycerols [GL03] | 9 | TG 16:0_18:0_18:1; TG 16:0_18:1_18:1; TG 18:0_18:0_18:1; TG 18:0_18:1_18:1; TG 18:0_38:3; TG 18:0_42:3; TG 18:0_48:2; TG 18:1_18:1_18:1; TG 20:0_36:2 |
| LION:0000607 | diacylglycerols [GL0201] | 3 | DG 18:1_20:1; DG 34:2; DG 38:4 |
| LION:0000622 | triacylglycerols [GL0301] | 9 | TG 16:0_18:0_18:1; TG 16:0_18:1_18:1; TG 18:0_18:0_18:1; TG 18:0_18:1_18:1; TG 18:0_38:3; TG 18:0_42:3; TG 18:0_48:2; TG 18:1_18:1_18:1; TG 20:0_36:2 |
| LION:0000640 | DG(34:2) | 1 | DG 34:2 |
| LION:0000655 | DG(38:2) | 1 | DG 18:1_20:1 |
| LION:0000657 | DG(38:4) | 1 | DG 38:4 |
| LION:0000728 | diacylglycerophosphoglycerophosphodiradylglycerols [GP1201] | 2 | CL 72:4; CL 72:5 |
| LION:0000760 | TG(52:1) | 1 | TG 16:0_18:0_18:1 |
| LION:0000761 | TG(52:2) | 1 | TG 16:0_18:1_18:1 |
| LION:0000770 | TG(54:1) | 1 | TG 18:0_18:0_18:1 |
| LION:0000773 | TG(54:2) | 1 | TG 18:0_18:1_18:1 |
| LION:0000774 | TG(54:3) | 1 | TG 18:1_18:1_18:1 |
| LION:0001006 | CL(72:4) | 1 | CL 72:4 |
| LION:0001007 | CL(72:5) | 1 | CL 72:5 |
| LION:0001285 | PE(40:4) | 1 | PE 18:1_22:3 |
| LION:0001320 | PC(O-30:1) | 1 | PC O-30:1 |
| LION:0001333 | PC(O-40:2) | 1 | PC O-40:2 |
| LION:0001355 | PE(O-34:1) | 1 | PE O-16:0_18:1 |
| LION:0001734 | contains vinyl ether bond (plasmalogen) | 2 | PE P-18:0_22:2; PE P-18:1_22:3 |
| LION:0001736 | low transition temperature | 3 | PC 16:1_16:1; PC 30:2; PE 16:1_18:1 |
| LION:0001737 | average transition temperature | 7 | PC 14:0_14:0; PC 14:0_16:1; PC 28:1; PC 29:0; PE 16:0_16:1; PE 18:1_22:3; PS 18:0_22:3 |
| LION:0001738 | high transition temperature | 2 | PC 14:0_16:0; PE 16:0_18:1 |
| LION:0001739 | very high transition temperature | 5 | PE 34:0; SM d40:0; SM d40:1; SM d42:1; SM d42:2 |
| LION:0001740 | above average transition temperature | 7 | PC 14:0_16:0; PE 16:0_18:1; PE 34:0; SM d40:0; SM d40:1; SM d42:1; SM d42:2 |
| LION:0001741 | below average transition temperature | 3 | PC 16:1_16:1; PC 30:2; PE 16:1_18:1 |
| LION:0001742 | fatty acids and conjugates [FA01] | 1 | FA 24:1 |
| LION:0001767 | FFA(24:1) | 1 | FA 24:1 |
| LION:0002562 | PC(28:1) | 1 | PC 28:1 |
| LION:0002882 | C16:0 | 7 | PC 14:0_16:0; PC O-16:0_16:1; PE 16:0_16:1; PE 16:0_18:1; PE O-16:0_18:1; TG 16:0_18:0_18:1; TG 16:0_18:1_18:1 |
| LION:0002900 | C16:1 | 6 | PC 14:0_16:1; PC 14:0_16:1_iso; PC 16:1_16:1; PC O-16:0_16:1; PE 16:0_16:1; PE 16:1_18:1 |
| LION:0002921 | C18:0 | 5 | PE P-18:0_22:2; PS 18:0_22:3; TG 16:0_18:0_18:1; TG 18:0_18:0_18:1; TG 18:0_18:1_18:1 |
| LION:0002922 | C18:1 | 17 | PE 16:0_18:1; PE 16:1_18:1; PE 18:1_22:3; PE O-16:0_18:1; PE P-18:1_22:3; Cer 18:1;O2/16:0; Cer 18:1;O2/24:0; Cer 18:1;O2/24:1; Hex-Cer 18:1;O2/24:0; Hex-Cer 18:1;O2/24:1; Hex2Cer 18:1;O2/24:1; DG 18:1_20:1; TG 16:0_18:0_18:1; TG 16:0_18:1_18:1; TG 18:0_18:0_18:1; TG 18:0_18:1_18:1; TG 18:1_18:1_18:1 |
| LION:0002926 | C20:1 | 1 | DG 18:1_20:1 |
| LION:0002933 | C22:2 | 1 | PE P-18:0_22:2 |
| LION:0002934 | C22:3 | 3 | PE 18:1_22:3; PE P-18:1_22:3; PS 18:0_22:3 |
| LION:0002939 | C24:1 | 1 | FA 24:1 |
| LION:0002945 | fatty acid with more than 18 carbons | 6 | FA 24:1; PE 18:1_22:3; PE P-18:0_22:2; PE P-18:1_22:3; PS 18:0_22:3; DG 18:1_20:1 |
| LION:0002947 | fatty acid with 13-15 carbons | 4 | PC 14:0_14:0; PC 14:0_16:0; PC 14:0_16:1; PC 14:0_16:1_iso |
| LION:0002948 | fatty acid with 16-18 carbons | 25 | PC 14:0_16:0; PC 14:0_16:1; PC 14:0_16:1_iso; PC 16:1_16:1; PC O-16:0_16:1; PE 16:0_16:1; PE 16:0_18:1; PE 16:1_18:1; PE 18:1_22:3; PE O-16:0_18:1; PE P-18:0_22:2; PE P-18:1_22:3; PS 18:0_22:3; Cer 18:1;O2/16:0; Cer 18:1;O2/24:0; Cer 18:1;O2/24:1; Hex-Cer 18:1;O2/24:0; Hex-Cer 18:1;O2/24:1; Hex2Cer 18:1;O2/24:1; DG 18:1_20:1; TG 16:0_18:0_18:1; TG 16:0_18:1_18:1; TG 18:0_18:0_18:1; TG 18:0_18:1_18:1; TG 18:1_18:1_18:1 |
| LION:0002949 | fatty acid with 19-21 carbons | 1 | DG 18:1_20:1 |
| LION:0002950 | fatty acid with 22-24 carbons | 5 | FA 24:1; PE 18:1_22:3; PE P-18:0_22:2; PE P-18:1_22:3; PS 18:0_22:3 |
| LION:0002953 | fatty acid with 14 carbons | 4 | PC 14:0_14:0; PC 14:0_16:0; PC 14:0_16:1; PC 14:0_16:1_iso |
| LION:0002955 | fatty acid with 16 carbons | 11 | PC 14:0_16:0; PC 14:0_16:1; PC 14:0_16:1_iso; PC 16:1_16:1; PC O-16:0_16:1; PE 16:0_16:1; PE 16:0_18:1; PE 16:1_18:1; PE O-16:0_18:1; TG 16:0_18:0_18:1; TG 16:0_18:1_18:1 |
| LION:0002957 | fatty acid with 18 carbons | 19 | PE 16:0_18:1; PE 16:1_18:1; PE 18:1_22:3; PE O-16:0_18:1; PE P-18:0_22:2; PE P-18:1_22:3; PS 18:0_22:3; Cer 18:1;O2/16:0; Cer 18:1;O2/24:0; Cer 18:1;O2/24:1; Hex-Cer 18:1;O2/24:0; Hex-Cer 18:1;O2/24:1; Hex2Cer 18:1;O2/24:1; DG 18:1_20:1; TG 16:0_18:0_18:1; TG 16:0_18:1_18:1; TG 18:0_18:0_18:1; TG 18:0_18:1_18:1; TG 18:1_18:1_18:1 |
| LION:0002959 | fatty acid with 20 carbons | 1 | DG 18:1_20:1 |
| LION:0002961 | fatty acid with 22 carbons | 4 | PE 18:1_22:3; PE P-18:0_22:2; PE P-18:1_22:3; PS 18:0_22:3 |
| LION:0002963 | fatty acid with 24 carbons | 1 | FA 24:1 |
| LION:0002966 | fatty acid with less than 2 double bonds | 27 | FA 24:1; PC 14:0_14:0; PC 14:0_16:0; PC 14:0_16:1; PC 14:0_16:1_iso; PC 16:1_16:1; PC O-16:0_16:1; PE 16:0_16:1; PE 16:0_18:1; PE 16:1_18:1; PE 18:1_22:3; PE O-16:0_18:1; PE P-18:0_22:2; PE P-18:1_22:3; PS 18:0_22:3; Cer 18:1;O2/16:0; Cer 18:1;O2/24:0; Cer 18:1;O2/24:1; Hex-Cer 18:1;O2/24:0; Hex-Cer 18:1;O2/24:1; Hex2Cer 18:1;O2/24:1; DG 18:1_20:1; TG 16:0_18:0_18:1; TG 16:0_18:1_18:1; TG 18:0_18:0_18:1; TG 18:0_18:1_18:1; TG 18:1_18:1_18:1 |
| LION:0002967 | polyunsaturated fatty acid | 4 | PE 18:1_22:3; PE P-18:0_22:2; PE P-18:1_22:3; PS 18:0_22:3 |
| LION:0002968 | saturated fatty acid | 14 | PC 14:0_14:0; PC 14:0_16:0; PC 14:0_16:1; PC 14:0_16:1_iso; PC O-16:0_16:1; PE 16:0_16:1; PE 16:0_18:1; PE O-16:0_18:1; PE P-18:0_22:2; PS 18:0_22:3; TG 16:0_18:0_18:1; TG 16:0_18:1_18:1; TG 18:0_18:0_18:1; TG 18:0_18:1_18:1 |
| LION:0002969 | monounsaturated fatty acid | 23 | FA 24:1; PC 14:0_16:1; PC 14:0_16:1_iso; PC 16:1_16:1; PC O-16:0_16:1; PE 16:0_16:1; PE 16:0_18:1; PE 16:1_18:1; PE 18:1_22:3; PE O-16:0_18:1; PE P-18:1_22:3; Cer 18:1;O2/16:0; Cer 18:1;O2/24:0; Cer 18:1;O2/24:1; Hex-Cer 18:1;O2/24:0; Hex-Cer 18:1;O2/24:1; Hex2Cer 18:1;O2/24:1; DG 18:1_20:1; TG 16:0_18:0_18:1; TG 16:0_18:1_18:1; TG 18:0_18:0_18:1; TG 18:0_18:1_18:1; TG 18:1_18:1_18:1 |
| LION:0002970 | fatty acid with 2 double bonds | 1 | PE P-18:0_22:2 |
| LION:0002971 | fatty acid with 3 double bonds | 3 | PE 18:1_22:3; PE P-18:1_22:3; PS 18:0_22:3 |
| LION:0002976 | fatty acid with more than 3 double bonds | 3 | PE 18:1_22:3; PE P-18:1_22:3; PS 18:0_22:3 |
| LION:0002977 | fatty acid with 3-5 double bonds | 3 | PE 18:1_22:3; PE P-18:1_22:3; PS 18:0_22:3 |
| LION:0002993 | PC(14:0/14:0) | 1 | PC 14:0_14:0 |
| LION:0002996 | PC(14:0/16:1) | 1 | PC 14:0_16:1 |
| LION:0002998 | PC(14:0/16:0) | 1 | PC 14:0_16:0 |
| LION:0003004 | PC(16:1/16:1) | 1 | PC 16:1_16:1 |
| LION:0003542 | PE(16:0/16:1) | 1 | PE 16:0_16:1 |
| LION:0003551 | PE(16:1/18:1) | 1 | PE 16:1_18:1 |
| LION:0003554 | PE(16:0/18:1) | 1 | PE 16:0_18:1 |
| LION:0003741 | PS(18:0/22:3) | 1 | PS 18:0_22:3 |
| LION:0004460 | DG(18:1/20:1) | 1 | DG 18:1_20:1 |
| LION:0005133 | TG(16:0/18:0/18:1) | 1 | TG 16:0_18:0_18:1 |
| LION:0005152 | TG(16:0/18:1/18:1) | 1 | TG 16:0_18:1_18:1 |
| LION:0005290 | TG(18:0/18:0/18:1) | 1 | TG 18:0_18:0_18:1 |
| LION:0005320 | TG(18:0/18:1/18:1) | 1 | TG 18:0_18:1_18:1 |
| LION:0005353 | TG(18:1/18:1/18:1) | 1 | TG 18:1_18:1_18:1 |
| LION:0012009 | lipid-mediated signalling | 6 | Cer 18:1;O2/16:0; Cer 18:1;O2/24:0; Cer 18:1;O2/24:1; DG 18:1_20:1; DG 34:2; DG 38:4 |
| LION:0012010 | membrane component | 33 | PC 14:0_14:0; PC 14:0_16:0; PC 14:0_16:1; PC 14:0_16:1_iso; PC 16:1_16:1; PC 28:1; PC 29:0; PC 30:2; PC O-16:0_16:1; PC O-30:0; PC O-30:1; PC O-32:2; PC O-34:3; PC O-40:2; PE 16:0_16:1; PE 16:0_18:1; PE 16:1_18:1; PE 18:1_22:3; PE 34:0; PE O-16:0_18:1; PE P-18:0_22:2; PE P-18:1_22:3; PS 18:0_22:3; SM d40:0; SM d40:1; SM d42:1; SM d42:2; Hex2Cer 18:1;O2/24:1; DG 18:1_20:1; DG 34:2; DG 38:4; CL 72:4; CL 72:5 |
| LION:0012011 | lipid storage | 9 | TG 16:0_18:0_18:1; TG 16:0_18:1_18:1; TG 18:0_18:0_18:1; TG 18:0_18:1_18:1; TG 18:0_38:3; TG 18:0_42:3; TG 18:0_48:2; TG 18:1_18:1_18:1; TG 20:0_36:2 |
| LION:0012018 | dihexosylceramides | 1 | Hex2Cer 18:1;O2/24:1 |
| LION:0012080 | endoplasmic reticulum (ER) | 25 | PC 14:0_14:0; PC 14:0_16:0; PC 14:0_16:1; PC 14:0_16:1_iso; PC 16:1_16:1; PC 28:1; PC 29:0; PC 30:2; PC O-16:0_16:1; PC O-30:0; PC O-30:1; PC O-32:2; PC O-34:3; PC O-40:2; PE 16:0_16:1; PE 16:0_18:1; PE 16:1_18:1; PE 18:1_22:3; PE 34:0; PE O-16:0_18:1; PE P-18:0_22:2; PE P-18:1_22:3; Cer 18:1;O2/16:0; Cer 18:1;O2/24:0; Cer 18:1;O2/24:1 |
| LION:0012081 | mitochondrion | 10 | PE 16:0_16:1; PE 16:0_18:1; PE 16:1_18:1; PE 18:1_22:3; PE 34:0; PE O-16:0_18:1; PE P-18:0_22:2; PE P-18:1_22:3; CL 72:4; CL 72:5 |
| LION:0012082 | plasma membrane | 8 | PS 18:0_22:3; SM d40:0; SM d40:1; SM d42:1; SM d42:2; Cer 18:1;O2/16:0; Cer 18:1;O2/24:0; Cer 18:1;O2/24:1 |
| LION:0012084 | lipid droplet | 9 | TG 16:0_18:0_18:1; TG 16:0_18:1_18:1; TG 18:0_18:0_18:1; TG 18:0_18:1_18:1; TG 18:0_38:3; TG 18:0_42:3; TG 18:0_48:2; TG 18:1_18:1_18:1; TG 20:0_36:2 |
| LION:0012085 | golgi apparatus | 4 | SM d40:0; SM d40:1; SM d42:1; SM d42:2 |
| LION:0012086 | endosome/lysosome | 4 | SM d40:0; SM d40:1; SM d42:1; SM d42:2 |
| LION:0012441 | N-acylsphingosines (ceramides) [SP0201] | 3 | Cer 18:1;O2/16:0; Cer 18:1;O2/24:0; Cer 18:1;O2/24:1 |
| LION:0016003 | PE(P-40:2) | 1 | PE P-18:0_22:2 |
| LION:0016009 | PE(P-40:4) | 1 | PE P-18:1_22:3 |
| LION:0017306 | PE(O-16:0/18:1) | 1 | PE O-16:0_18:1 |
| LION:0019464 | PE(P-18:0/22:2) | 1 | PE P-18:0_22:2 |
| LION:0019470 | PE(P-18:1/22:3) | 1 | PE P-18:1_22:3 |
| LION:0041232 | PC(29:0) | 1 | PC 29:0 |
| LION:0080664 | PE(18:1/22:3) | 1 | PE 18:1_22:3 |
| LION:0080968 | very low bilayer thickness | 4 | PC 14:0_14:0; PC 14:0_16:1; PC 28:1; PC 30:2 |
| LION:0080969 | low bilayer thickness | 4 | PC 14:0_16:0; PC 14:0_16:1; PC 16:1_16:1; PC 29:0 |
| LION:0080970 | average bilayer thickness | 2 | PE 16:0_16:1; PE 16:1_18:1 |
| LION:0080971 | high bilayer thickness | 4 | PE 16:0_18:1; PE 18:1_22:3; PE 34:0; PS 18:0_22:3 |
| LION:0080973 | below average bilayer thickness | 7 | PC 14:0_14:0; PC 14:0_16:0; PC 14:0_16:1; PC 16:1_16:1; PC 28:1; PC 29:0; PC 30:2 |
| LION:0080974 | above average bilayer thickness | 4 | PE 16:0_18:1; PE 18:1_22:3; PE 34:0; PS 18:0_22:3 |
| LION:0080976 | very low lateral diffusion | 1 | PE 34:0 |
| LION:0080977 | low lateral diffusion | 3 | PE 16:0_16:1; PE 16:0_18:1; PS 18:0_22:3 |
| LION:0080978 | average lateral diffusion | 6 | PC 14:0_14:0; PC 14:0_16:0; PC 29:0; PE 16:0_16:1; PE 16:1_18:1; PE 18:1_22:3 |
| LION:0080979 | high lateral diffusion | 5 | PC 14:0_14:0; PC 14:0_16:1; PC 16:1_16:1; PC 28:1; PC 30:2 |
| LION:0080980 | very high lateral diffusion | 1 | PC 30:2 |
| LION:0080981 | below average lateral diffusion | 4 | PE 16:0_16:1; PE 16:0_18:1; PE 34:0; PS 18:0_22:3 |
| LION:0080982 | above average lateral diffusion | 5 | PC 14:0_14:0; PC 14:0_16:1; PC 16:1_16:1; PC 28:1; PC 30:2 |
