## Supplementary material for "A cell state specific metabolic vulnerability to GPX4-dependent ferroptosis in glioblastoma": Table S6

| Metabolic Gene Sets | AC | MES1 | MES2 | OPC | NPC1 | NPC2 | G1/S | G2/M |
| --- | --- | --- | --- | --- | --- | --- | --- | --- |
| GOBP_REGULATION_OF_GLYCOPROTEIN_METABOLIC_PROCESS | 0.633767 | 0.473886 | 0.315814 | 0.219108 | -0.20011 | -0.207863 | -0.151558 | -0.121049 |
| GOBP_FATTY_ACID_TRANSPORT | 0.534457 | 0.433565 | 0.279673 | 0.192772 | -0.14206 | -0.032892 | -0.205599 | -0.141684 |
| GOBP_REGULATION_OF_LIPID_LOCALIZATION | 0.520791 | 0.560606 | 0.408923 | 0.177001 | -0.169091 | -0.10106 | -0.193374 | -0.13097 |
| GOBP_PROTEIN_LIPID_COMPLEX_SUBUNIT_ORGANIZATION | 0.50977 |  | 0.328562 |  | -0.276981 | -0.227976 | -0.187962 | -0.191977 |
| GOBP_POSITIVE_REGULATION_OF_STEROID_METABOLIC_PROCESS | 0.497614 | 0.459183 | 0.247518 | 0.003216 | -0.355857 | -0.276725 | -0.240207 | -0.239623 |
| GOBP_SUPEROXIDE_METABOLIC_PROCESS | 0.495839 | 0.38933 | 0.236562 | 0.182647 | -0.166507 | -0.227389 | 0.018118 | -0.045049 |
| GOBP_REGULATION_OF_OXIDOREDUCTASE_ACTIVITY | 0.48685 | 0.400761 | 0.296561 | 0.171991 | -0.048477 |  | 0.025834 | 0.035896 |
| GOBP_FATTY_ACID_METABOLIC_PROCESS | 0.479732 | 0.387554 | 0.383107 | 0.36505 | 0.018041 | 0.036626 | -0.070404 | -0.074503 |
| GOBP_CELLULAR_LIPID_CATABOLIC_PROCESS | 0.464725 | 0.355928 | 0.315242 | 0.274324 | -0.088383 | -0.071292 | -0.131405 | -0.131748 |
| GOBP_REGULATION_OF_PHOSPHOLIPID_TRANSPORT | 0.456958 | 0.293705 | 0.188112 | 0.232046 | -0.062759 | -0.075979 | -0.038127 | -0.062623 |
| GOBP_POSITIVE_REGULATION_OF_REACTIVE_OXYGEN_SPECIES_METABOLIC_PROCESS | 0.453076 | 0.495099 | 0.445398 | 0.178293 | -0.032676 |  | -0.01681 | -0.011538 |
| GOBP_POSITIVE_REGULATION_OF_CHOLESTEROL_EFFLUX | 0.452636 | 0.510759 | 0.356346 | 0.053414 | -0.245458 | -0.133263 | -0.221761 | -0.205167 |
| KEGG_GLYCINE_SERINE_AND_THREONINE_METABOLISM | 0.452631 | 0.123651 | 0.097481 | 0.1261 | -0.205956 | -0.250337 | -0.008709 |  |
| GOBP_LONG_CHAIN_FATTY_ACID_TRANSPORT | 0.452515 | 0.494935 | 0.300308 | 0.053442 | -0.326203 | -0.246756 | -0.215216 | -0.210133 |
| GOBP_GLYCOPROTEIN_METABOLIC_PROCESS | 0.44411 | 0.383194 | 0.415047 | 0.378444 | 0.090749 | 0.065714 | -0.072175 |  |
| GOBP_REGULATION_OF_TRANSCRIPTION_FROM_RNA_POLYMERASE_II_PROMOTER_IN_RESPONSE_TO_OXIDATIVE_STRESS | 0.44063 | 0.428654 | 0.365396 | 0.054323 | -0.265816 | -0.290992 | -0.086958 | -0.075848 |
| GOBP_LIPID_CATABOLIC_PROCESS | 0.434727 | 0.379783 | 0.338629 | 0.260191 | -0.053089 | 0.010305 | -0.12445 | -0.117454 |
| REACTOME_FATTY_ACID_METABOLISM | 0.431888 | 0.240956 | 0.258255 | 0.373111 | 0.098029 | 0.088576 |  | -0.014464 |
| GOBP_RESPONSE_TO_LIPID | 0.428359 | 0.539857 | 0.525201 | 0.276929 | 0.094069 | 0.130855 | 0.065673 | 0.105032 |
| KEGG_ARGININE_AND_PROLINE_METABOLISM | 0.427255 | 0.289309 | 0.270666 | 0.199217 | 0.00846 | 0.039256 | -0.012696 | -0.010539 |
| GOBP_RESPONSE_TO_LIPID | 0.42693 | 0.540044 | 0.526512 | 0.276388 | 0.096484 | 0.134375 | 0.06483 | 0.103328 |
| GOBP_REGULATION_OF_CHOLESTEROL_EFFLUX | 0.425908 |  | 0.339939 | 0.065392 | -0.225273 | -0.123307 | -0.218629 | -0.210574 |
| GOBP_POSITIVE_REGULATION_OF_LIPID_LOCALIZATION | 0.424956 | 0.563236 | 0.412549 | 0.129153 | -0.167793 | -0.093812 | -0.169765 | -0.088002 |
| GOBP_POSITIVE_REGULATION_OF_LIPID_LOCALIZATION | 0.42118 | 0.560708 | 0.410396 | 0.130376 | -0.163616 |  | -0.168387 | -0.08589 |
| GOBP_POSITIVE_REGULATION_OF_LIPID_TRANSPORT | 0.412588 | 0.509463 | 0.370724 | 0.140879 | -0.125905 | -0.047404 | -0.150449 | -0.067236 |
| GOBP_FATTY_ACID_BIOSYNTHETIC_PROCESS | 0.406722 |  | 0.32659 | 0.421653 | 0.068002 | 0.025145 | -0.001871 | -0.011805 |
| GOBP_CELLULAR_RESPONSE_TO_LIPID | 0.403143 | 0.477493 | 0.474627 | 0.291701 | 0.128543 | 0.147818 | 0.075706 | 0.122571 |
| GOBP_ENERGY_RESERVE_METABOLIC_PROCESS | 0.39954 | 0.29552 | 0.358672 | 0.182243 | 0.018456 | 0.041231 | 0.051244 | 0.064139 |
| GOBP_PHOSPHOLIPID_TRANSPORT | 0.39899 | 0.348681 | 0.261014 | 0.210364 | -0.069973 | -0.022909 | -0.131201 | -0.124562 |
| GOBP_PROTEOGLYCAN_METABOLIC_PROCESS | 0.393534 | 0.240186 | 0.211183 | 0.38945 | 0.090404 | -0.031228 | -0.063132 | -0.069813 |
| GOBP_CHOLESTEROL_EFFLUX | 0.388389 | 0.474178 | 0.346535 | 0.056394 | -0.216672 | -0.099161 | -0.220977 | -0.209268 |
| GOBP_CARBOHYDRATE_DERIVATIVE_METABOLIC_PROCESS | 0.386424 | 0.345465 | 0.485854 | 0.409326 | 0.223179 | 0.226018 | 0.126813 | 0.069817 |
| REACTOME_GLYCOGEN_METABOLISM | 0.370954 | 0.287376 | 0.361287 | 0.145617 | -0.038394 | 0.006162 | 0.003895 | 0.019962 |
| GOBP_CELLULAR_MODIFIED_AMINO_ACID_METABOLIC_PROCESS | 0.365068 | 0.189131 | 0.197123 | 0.29375 | 0.089588 | 0.07455 | 0.127865 | 0.048168 |
| GOBP_NEGATIVE_REGULATION_OF_STEROID_METABOLIC_PROCESS | 0.36233 | 0.40643 | 0.314469 | 0.098112 | -0.114456 | 0.002718 |  |  |
| REACTOME_FATTY_ACYL_COA_BIOSYNTHESIS | 0.348459 | 0.157185 | 0.148987 | 0.342102 | 0.094775 |  |  | -0.002092 |
| GOBP_POSITIVE_REGULATION_OF_LIPID_BIOSYNTHETIC_PROCESS | 0.346831 | 0.490281 | 0.327558 | 0.03413 | -0.21158 | -0.109813 | -0.200142 | -0.162651 |
| REACTOME_METABOLISM_OF_CARBOHYDRATES | 0.34389 | 0.324264 |  | 0.358241 | 0.183516 | 0.187961 | 0.088617 | 0.076442 |
| GOBP_FATTY_ACYL_COA_BIOSYNTHETIC_PROCESS | 0.340404 | 0.142963 | 0.135188 | 0.349832 | 0.092365 | 0.024913 | 0.00346 | -0.000342 |
| GOBP_UNSATURATED_FATTY_ACID_METABOLIC_PROCESS | 0.338314 | 0.349261 | 0.30117 | 0.338534 | 0.020456 |  | -0.018901 | -0.028635 |
| GOBP_RESPONSE_TO_OXIDATIVE_STRESS | 0.335107 | 0.378475 | 0.506802 | 0.345454 | 0.23167 | 0.251032 | 0.170415 | 0.155789 |
| GOBP_ORGANIC_ACID_METABOLIC_PROCESS | 0.329848 | 0.262783 | 0.461582 | 0.38085 | 0.249322 | 0.278063 | 0.132643 | 0.109437 |
| GOBP_REGULATION_OF_LIPID_METABOLIC_PROCESS | 0.327047 | 0.400119 | 0.441911 | 0.283108 | 0.120086 | 0.211035 | -0.072531 | -0.045236 |
| REACTOME_ARACHIDONIC_ACID_METABOLISM | 0.32381 |  | 0.225476 | 0.156566 | -0.043029 | -0.004261 |  | -0.053765 |
| GOBP_POSITIVE_REGULATION_OF_PHOSPHORUS_METABOLIC_PROCESS | 0.31489 | 0.369885 | 0.484792 | 0.339377 | 0.273536 | 0.324397 | 0.097343 | 0.185168 |
| GOBP_SERINE_FAMILY_AMINO_ACID_METABOLIC_PROCESS | 0.303105 | 0.051548 | 0.125519 | 0.192938 | -0.025906 | -0.087816 | 0.127102 | 0.036035 |
| GOBP_FATTY_ACID_CATABOLIC_PROCESS | 0.295739 | 0.188893 | 0.23942 | 0.243327 | 0.062226 | 0.080217 | -0.019926 | -0.027161 |
| GOBP_REGULATION_OF_TRIGLYCERIDE_METABOLIC_PROCESS | 0.295466 | 0.398582 | 0.23832 | 0.026962 | -0.18554 | -0.088062 | -0.173491 | -0.162739 |
| GOBP_PURINE_CONTAINING_COMPOUND_METABOLIC_PROCESS | 0.289424 | 0.217989 | 0.44402 | 0.387585 | 0.277641 | 0.269836 | 0.168618 | 0.120235 |
| GOBP_GENERATION_OF_PRECURSOR_METABOLITES_AND_ENERGY | 0.287606 | 0.183176 | 0.414747 | 0.31172 | 0.264507 | 0.279209 | 0.203293 | 0.165068 |
| GOBP_NEGATIVE_REGULATION_OF_REACTIVE_OXYGEN_SPECIES_METABOLIC_PROCESS | 0.287214 | 0.332356 | 0.408497 | 0.28601 | 0.053895 | 0.04871 | -0.003389 | 0.005363 |
| GOBP_CELLULAR_GLUCAN_METABOLIC_PROCESS | 0.285797 | 0.234222 | 0.327903 | 0.150693 | 0.072246 | 0.126273 | 0.06251 | 0.079684 |
| GOBP_ENERGY_DERIVATION_BY_OXIDATION_OF_ORGANIC_COMPOUNDS | 0.285691 | 0.137601 | 0.343373 | 0.308974 | 0.271126 | 0.268839 | 0.210989 | 0.176167 |
| GOBP_REGULATION_OF_CHOLESTEROL_BIOSYNTHETIC_PROCESS | 0.281335 | 0.174444 | 0.203877 | 0.301125 | 0.144299 | 0.132673 | 0.057772 | 0.050459 |
| GOBP_REGULATION_OF_CHOLESTEROL_METABOLIC_PROCESS | 0.27435 | 0.184585 | 0.21744 | 0.285766 | 0.129799 | 0.12869 | 0.038037 | 0.037538 |
| GOBP_POLYSACCHARIDE_METABOLIC_PROCESS | 0.272956 |  | 0.32318 | 0.159369 | 0.072244 | 0.133078 | 0.033924 | 0.050043 |
| GOBP_ATP_METABOLIC_PROCESS | 0.272692 | 0.174001 | 0.402029 | 0.315532 | 0.27867 | 0.285582 |  | 0.168744 |
| GOBP_IRON_ION_HOMEOSTASIS | 0.271571 | 0.336696 | 0.431409 | 0.222375 | 0.15074 | 0.222459 | 0.021473 | 0.028295 |
| WP_UREA_CYCLE_AND_METABOLISM_OF_AMINO_GROUPS | 0.265726 | -0.054383 | -0.020457 | 0.209781 |  | 0.04796 |  |  |
| GOBP_FATTY_ACID_BETA_OXIDATION | 0.264402 | 0.19535 | 0.245703 | 0.22201 | 0.070604 | 0.101982 |  | -0.017106 |
| GOBP_STEROID_HORMONE_BIOSYNTHETIC_PROCESS | 0.264149 | 0.275841 | 0.296861 | 0.099109 | -0.066871 | -0.010524 | -0.095028 | -0.061139 |
| GOBP_LIPID_OXIDATION | 0.262236 | 0.20253 | 0.255862 | 0.245514 | 0.080879 | 0.104814 | -0.024288 | -0.023524 |
| GOBP_CARBOHYDRATE_METABOLIC_PROCESS | 0.259856 | 0.32382 | 0.542758 | 0.305844 |  | 0.250839 | 0.079445 | 0.075268 |
| GOBP_NEGATIVE_REGULATION_OF_PROTEIN_METABOLIC_PROCESS | 0.259167 | 0.310897 | 0.487646 | 0.362676 | 0.37372 | 0.417346 | 0.148771 | 0.201235 |
| GOBP_RESPONSE_TO_IRON_ION | 0.255564 | 0.304207 | 0.243873 | 0.137791 | -0.048856 | -0.038625 | -0.024581 | 0.091183 |
| WP_ELECTRON_TRANSPORT_CHAIN_OXPHOS_SYSTEM_IN_MITOCHONDRIA | 0.253393 |  | 0.296017 | 0.29231 | 0.269607 | 0.259428 | 0.207438 | 0.15806 |
| GOBP_SPHINGOLIPID_METABOLIC_PROCESS | 0.253322 | 0.251394 | 0.334602 | 0.270264 | 0.1357 | 0.192317 | -0.044474 | -0.045602 |
| REACTOME_THE_CITRIC_ACID_TCA_CYCLE_AND_RESPIRATORY_ELECTRON_TRANSPORT | 0.249913 | 0.10473 | 0.319542 | 0.310256 | 0.286054 | 0.276369 | 0.216255 | 0.167584 |
| REACTOME_RESPIRATORY_ELECTRON_TRANSPORT_ATP_SYNTHESIS_BY_CHEMIOSMOTIC_COUPLING_AND_HEAT_PRODUCTION_BY_UNCOUPLING_PROTEINS | 0.247541 | 0.103738 | 0.301134 | 0.293306 | 0.275622 | 0.268441 | 0.208266 | 0.160957 |
| GOBP_OXIDATIVE_PHOSPHORYLATION | 0.24672 | 0.108286 | 0.308395 | 0.287746 | 0.269396 | 0.26438 | 0.215351 | 0.173704 |
| REACTOME_RESPIRATORY_ELECTRON_TRANSPORT | 0.244432 | 0.101557 | 0.300797 | 0.292568 | 0.276481 | 0.271326 | 0.20946 | 0.162179 |
| KEGG_OXIDATIVE_PHOSPHORYLATION | 0.241157 | 0.114897 | 0.312838 | 0.286319 | 0.277877 | 0.281694 | 0.18859 | 0.14236 |
| WP_OXIDATIVE_PHOSPHORYLATION | 0.240173 | 0.113878 | 0.29113 | 0.273553 | 0.269886 | 0.26729 | 0.199497 | 0.151776 |
| GOBP_GLUTATHIONE_TRANSMEMBRANE_TRANSPORT | 0.237207 | 0.212833 | 0.049408 | 0.071517 | -0.085895 | -0.060227 | -0.112911 | -0.087748 |
| GOBP_NUCLEOBASE_CONTAINING_SMALL_MOLECULE_METABOLIC_PROCESS | 0.237061 | 0.206624 | 0.432655 | 0.383135 | 0.317825 | 0.310328 | 0.260966 | 0.170914 |
| GOBP_REGULATION_OF_HYDROGEN_PEROXIDE_BIOSYNTHETIC_PROCESS | 0.229113 | 0.343744 | 0.268071 | 0.181909 |  | -0.030305 | -0.03805 | -0.028928 |
| REACTOME_GLUTATHIONE_CONJUGATION | 0.228219 | 0.188659 | 0.162797 | 0.170013 | 0.022305 | 0.044828 | 0.045141 | 0.010283 |
| GOBP_POSITIVE_REGULATION_OF_HYDROGEN_PEROXIDE_BIOSYNTHETIC_PROCESS | 0.22663 | 0.398229 | 0.294392 | 0.06656 | -0.080465 | -0.074813 | -0.050721 | -0.048046 |
| GOBP_GLUTATHIONE_DERIVATIVE_METABOLIC_PROCESS | 0.225913 | 0.170722 | 0.151694 | 0.17988 | 0.038801 | 0.05939 | 0.033006 | 0.022376 |
| GOBP_POSITIVE_REGULATION_OF_HYDROGEN_PEROXIDE_METABOLIC_PROCESS | 0.22557 | 0.382281 | 0.282514 | 0.069904 | -0.07524 | -0.066902 | -0.044789 | -0.037975 |
| WP_GLYCOGEN_SYNTHESIS_AND_DEGRADATION | 0.224658 | 0.131249 | 0.258358 | 0.206645 | 0.178473 | 0.250036 | 0.086307 | 0.161941 |
| GOBP_REGULATION_OF_FATTY_ACID_OXIDATION | 0.219521 | 0.232241 | 0.232311 | 0.07212 | -0.027717 | 0.029145 | -0.085223 | -0.046423 |
| GOBP_CELLULAR_CARBOHYDRATE_METABOLIC_PROCESS | 0.21952 | 0.229456 | 0.411042 | 0.216078 | 0.176312 | 0.234564 | 0.05767 | 0.06554 |
| GOBP_POSITIVE_REGULATION_OF_MITOCHONDRIAL_FISSION | 0.219358 | 0.153234 | 0.306934 | 0.150491 | 0.053481 | 0.100288 | -0.014582 | -0.018938 |
| GOBP_NUCLEOSIDE_DIPHOSPHATE_METABOLIC_PROCESS | 0.216283 | 0.269959 | 0.532263 | 0.25912 | 0.18617 | 0.220322 | 0.13595 | 0.106087 |
| GOBP_REGULATION_OF_ATP_METABOLIC_PROCESS | 0.216134 | 0.189918 | 0.371475 | 0.272754 | 0.24288 | 0.24491 | 0.150172 | 0.120209 |
| GOBP_REGULATION_OF_NUCLEOTIDE_METABOLIC_PROCESS | 0.212508 | 0.21433 | 0.371013 | 0.228948 | 0.188787 | 0.193711 | 0.111264 | 0.094736 |
| GOBP_POSITIVE_REGULATION_OF_PROTEIN_METABOLIC_PROCESS | 0.211316 | 0.272809 | 0.473479 | 0.364205 | 0.403924 | 0.43122 | 0.195229 | 0.255081 |
| REACTOME_MITOCHONDRIAL_BIOGENESIS | 0.210216 |  | 0.2764 | 0.20573 | 0.223566 | 0.240437 | 0.136965 | 0.119912 |
| GOBP_REGULATION_OF_PHOSPHORUS_METABOLIC_PROCESS | 0.207994 | 0.282156 | 0.47001 | 0.356628 | 0.392762 | 0.436392 | 0.17074 | 0.242695 |
| GOBP_DICARBOXYLIC_ACID_METABOLIC_PROCESS | 0.200108 | 0.108719 | 0.213148 | 0.205388 | 0.122448 | 0.133742 | 0.10616 | 0.047997 |
| REACTOME_PYRUVATE_METABOLISM_AND_CITRIC_ACID_TCA_CYCLE | 0.195981 | 0.090947 | 0.307261 | 0.278587 | 0.210917 | 0.196557 | 0.160462 | 0.123199 |
| KEGG_PYRUVATE_METABOLISM | 0.195489 | 0.098473 | 0.276225 | 0.297322 | 0.220065 | 0.215468 | 0.176886 | 0.150164 |
| MOOTHA_VOXPHOS | 0.193249 | 0.061934 | 0.266443 | 0.27961 | 0.300449 | 0.310938 | 0.221364 | 0.175728 |
| GOBP_CELLULAR_IRON_ION_HOMEOSTASIS | 0.192185 | 0.258777 | 0.378186 | 0.219438 | 0.219066 | 0.295847 | 0.042005 | 0.043553 |
| GOBP_MITOCHONDRIAL_RESPIRATORY_CHAIN_COMPLEX_ASSEMBLY | 0.190979 | 0.087561 | 0.286604 | 0.263269 | 0.296792 | 0.308566 | 0.219961 | 0.168522 |
| WP_TCA_CYCLE_AKA_KREBS_OR_CITRIC_ACID_CYCLE | 0.190334 | 0.052463 | 0.169794 | 0.190278 | 0.146897 | 0.13962 |  |  |
| GOBP_MITOCHONDRIAL_RESPIRATORY_CHAIN_COMPLEX_ASSEMBLY | 0.189419 | 0.087593 | 0.28533 | 0.263579 | 0.294368 | 0.306944 | 0.22204 | 0.166072 |
| GOBP_INNER_MITOCHONDRIAL_MEMBRANE_ORGANIZATION | 0.186641 | 0.089761 | 0.26557 | 0.219591 | 0.216135 |  |  |  |
| GOBP_PYRUVATE_METABOLIC_PROCESS | 0.186273 |  | 0.536986 | 0.265629 | 0.205656 | 0.248577 | 0.11957 | 0.10807 |
| GOBP_MITOCHONDRIAL_TRANSMEMBRANE_TRANSPORT | 0.18498 | 0.071751 | 0.287638 | 0.278051 | 0.247964 | 0.267516 | 0.14804 | 0.08494 |
| REACTOME_CITRIC_ACID_CYCLE_TCA_CYCLE | 0.183391 | 0.047395 | 0.166902 | 0.193039 | 0.157065 | 0.155315 | 0.127524 | 0.06821 |
| GOBP_RESPONSE_TO_MITOCHONDRIAL_DEPOLARISATION | 0.179137 | 0.269816 | 0.373725 | 0.082999 | 0.010047 | 0.11808 | -0.078585 | -0.060706 |
| GOBP_RESPONSE_TO_MITOCHONDRIAL_DEPOLARISATION | 0.177669 | 0.270243 | 0.376071 | 0.080933 |  | 0.119708 | -0.079967 | -0.058232 |
| GOBP_LONG_CHAIN_FATTY_ACID_BIOSYNTHETIC_PROCESS | 0.174889 | 0.049104 | 0.082688 | 0.342412 | 0.155225 | 0.093359 |  | 0.065515 |
| REACTOME_PHOSPHOLIPID_METABOLISM | 0.174775 | 0.137593 | 0.250027 | 0.255771 | 0.198898 | 0.234341 | -0.046045 | -0.006845 |
| GOBP_POSITIVE_REGULATION_OF_FATTY_ACID_BETA_OXIDATION | 0.173619 | 0.149276 | 0.133003 | 0.034307 | -0.083045 | -0.083299 | -0.073419 | -0.04702 |
| GOBP_REGULATION_OF_MITOCHONDRION_ORGANIZATION | 0.169575 | 0.168965 | 0.416179 | 0.277474 | 0.330428 | 0.403434 | 0.117389 | 0.095687 |
| GOBP_ONE_CARBON_METABOLIC_PROCESS | 0.165784 | 0.157837 | 0.216729 | 0.147369 | 0.095689 | 0.097038 | 0.203223 | 0.090867 |
| GOBP_NEGATIVE_REGULATION_OF_PHOSPHORUS_METABOLIC_PROCESS | 0.164905 | 0.200624 | 0.400167 | 0.341812 | 0.407884 | 0.456012 | 0.157897 | 0.213275 |
| GOBP_MITOCHONDRIAL_FISSION | 0.16413 | 0.089165 | 0.241798 | 0.186357 | 0.148585 | 0.184473 |  |  |
| KEGG_CITRATE_CYCLE_TCA_CYCLE | 0.162509 | 0.03891 | 0.191586 | 0.228514 | 0.197855 | 0.19052 | 0.14267 | 0.100787 |
| KEGG_GLUTATHIONE_METABOLISM | 0.162409 | 0.140785 | 0.194933 | 0.212494 | 0.187752 | 0.18304 | 0.27253 | 0.179449 |
| GOBP_RIBONUCLEOSIDE_TRIPHOSPHATE_METABOLIC_PROCESS | 0.160142 | 0.146111 | 0.339404 | 0.291192 | 0.26282 | 0.212626 | 0.21095 | 0.152446 |
| REACTOME_IRON_UPTAKE_AND_TRANSPORT | 0.16008 | 0.217074 | 0.38849 | 0.239158 | 0.262764 | 0.347249 | 0.062748 | 0.058764 |
| GOBP_REGULATION_OF_CARBOHYDRATE_METABOLIC_PROCESS | 0.160022 | 0.180795 | 0.348949 | 0.253737 | 0.272941 | 0.286992 | 0.13055 | 0.143822 |
| GOBP_MONOSACCHARIDE_METABOLIC_PROCESS | 0.158404 | 0.274505 | 0.528316 | 0.280998 | 0.225544 | 0.264999 | 0.101706 | 0.10356 |
| GOBP_GLYCEROPHOSPHOLIPID_CATABOLIC_PROCESS | 0.15796 | 0.220283 | 0.144933 | 0.050119 | -0.101824 | -0.073854 | -0.073924 | -0.067922 |
| GOBP_TRICARBOXYLIC_ACID_CYCLE | 0.157292 | 0.031618 | 0.185467 | 0.223456 | 0.211336 | 0.207337 | 0.146465 | 0.108201 |
| GOBP_MITOCHONDRIAL_TRANSPORT | 0.156383 | 0.091376 | 0.363895 | 0.333616 | 0.394952 | 0.431224 | 0.199265 | 0.15936 |
| GOBP_GLUCOSE_METABOLIC_PROCESS | 0.152838 | 0.262972 | 0.52787 | 0.275987 | 0.228891 | 0.265454 | 0.103591 | 0.119227 |
| GOBP_NAD_METABOLIC_PROCESS | 0.152423 | 0.286913 | 0.569461 | 0.18808 | 0.086869 | 0.148576 | 0.050189 | 0.037892 |
| GOBP_REGULATION_OF_GENERATION_OF_PRECURSOR_METABOLITES_AND_ENERGY | 0.152411 | 0.139368 | 0.364694 | 0.249879 | 0.302665 | 0.321257 | 0.228114 | 0.237696 |
| REACTOME_GLYOXYLATE_METABOLISM_AND_GLYCINE_DEGRADATION | 0.14999 | 0.001308 | 0.099467 | 0.197295 | 0.126262 | 0.092791 | 0.105698 | 0.043529 |
| GOBP_TETRAPYRROLE_METABOLIC_PROCESS | 0.14828 | 0.136114 | 0.214341 | 0.178742 | 0.109983 | 0.154507 | 0.006841 | -0.029385 |
| GOBP_HEPARAN_SULFATE_PROTEOGLYCAN_METABOLIC_PROCESS | 0.14671 | 0.204489 | 0.238388 | 0.158885 | 0.066059 | 0.072623 | -0.04625 | -0.025039 |
| GOBP_PHOSPHOLIPID_CATABOLIC_PROCESS | 0.145648 | 0.187828 | 0.117818 | 0.064588 | -0.055259 | -0.027092 | -0.07963 | -0.063929 |
| GOBP_REGULATION_OF_OXIDATIVE_STRESS_INDUCED_CELL_DEATH | 0.144379 | 0.267203 | 0.402859 | 0.276139 | 0.260129 | 0.298944 | 0.121799 | 0.134924 |
| GOBP_REGULATION_OF_RESPONSE_TO_OXIDATIVE_STRESS | 0.144272 | 0.26259 | 0.383776 | 0.272855 | 0.261891 | 0.298103 | 0.14485 | 0.13734 |
| GOBP_REGULATION_OF_OXIDATIVE_PHOSPHORYLATION | 0.143516 | 0.142294 | 0.27181 | 0.115323 | 0.076816 | 0.143547 | 0.048831 | 0.025964 |
| GOBP_REGULATION_OF_RESPONSE_TO_OXIDATIVE_STRESS | 0.143205 | 0.263407 | 0.384617 | 0.2717 | 0.258892 | 0.299742 | 0.141948 | 0.139028 |
| GOBP_RIBONUCLEOSIDE_MONOPHOSPHATE_METABOLIC_PROCESS | 0.143176 | 0.100082 | 0.203381 | 0.16421 | 0.109032 | 0.119953 | 0.111839 | 0.030365 |
| GOBP_PORPHYRIN_CONTAINING_COMPOUND_METABOLIC_PROCESS | 0.141843 | 0.138475 | 0.223517 | 0.192938 | 0.122772 | 0.173487 | 0.009027 | -0.025746 |
| GOBP_GLUTAMINE_METABOLIC_PROCESS | 0.141203 | 0.154662 | 0.239688 | 0.092706 | 0.041968 | 0.072891 | 0.060387 | 0.02733 |
| GOBP_NADH_METABOLIC_PROCESS | 0.140009 | 0.245937 | 0.529278 | 0.201227 | 0.144634 | 0.207167 | 0.084184 | 0.056604 |
| GOBP_HEME_METABOLIC_PROCESS | 0.138532 | 0.145141 | 0.224709 | 0.188764 | 0.11744 | 0.177078 | 0.00626 | -0.027197 |
| GOBP_MEMBRANE_LIPID_CATABOLIC_PROCESS | 0.138413 | 0.199501 | 0.218232 | 0.090334 | -0.003632 | 0.061891 | -0.103165 | -0.099121 |
| GOBP_CELLULAR_AMIDE_METABOLIC_PROCESS | 0.137416 | 0.104211 | 0.390849 | 0.356557 | 0.422345 | 0.397142 | 0.254573 | 0.226251 |
| GOBP_MEMBRANE_LIPID_CATABOLIC_PROCESS | 0.136193 | 0.196725 | 0.218207 | 0.092565 |  | 0.066621 | -0.10708 | -0.101504 |
| GOBP_NUCLEOSIDE_MONOPHOSPHATE_METABOLIC_PROCESS | 0.13598 | 0.052834 | 0.169232 | 0.193552 | 0.173855 | 0.139189 | 0.362342 | 0.182503 |
| GOBP_PHOSPHATIDYLINOSITOL_METABOLIC_PROCESS | 0.135089 | 0.151012 | 0.240163 | 0.171727 | 0.15551 | 0.201408 | -0.059799 | -0.01932 |
| REACTOME_PYRUVATE_METABOLISM | 0.134013 | 0.100642 | 0.329655 | 0.252756 | 0.183835 |  | 0.136058 | 0.132603 |
| GOBP_MITOCHONDRIAL_MEMBRANE_ORGANIZATION | 0.133515 | 0.057494 | 0.31709 | 0.315643 | 0.409961 | 0.436555 | 0.242853 | 0.204996 |
| WP_GLUTATHIONE_METABOLISM | 0.132178 | 0.208578 | 0.211645 | 0.090354 | 0.041844 | 0.093532 | 0.015896 | 0.027437 |
| GOBP_REGULATION_OF_CELLULAR_CARBOHYDRATE_METABOLIC_PROCESS | 0.128216 | 0.15801 |  | 0.206783 | 0.223093 | 0.247509 | 0.074892 | 0.095002 |
| GOBP_NEGATIVE_REGULATION_OF_CELLULAR_AMIDE_METABOLIC_PROCESS | 0.125502 | 0.171537 | 0.36983 | 0.294151 | 0.415641 | 0.44042 | 0.22037 | 0.234406 |
| REACTOME_INTEGRATION_OF_ENERGY_METABOLISM | 0.124614 | 0.05302 | 0.213433 | 0.329789 | 0.356325 | 0.380814 | 0.140274 | 0.160775 |
| GOBP_CELL_DEATH_IN_RESPONSE_TO_OXIDATIVE_STRESS | 0.123644 | 0.245088 | 0.396553 | 0.285518 | 0.279125 | 0.32399 | 0.125654 | 0.13357 |
| GOBP_CELLULAR_AMINO_ACID_METABOLIC_PROCESS | 0.121137 | 0.057129 | 0.302522 | 0.319464 | 0.38079 | 0.384551 | 0.263029 | 0.23397 |
| GOBP_NUCLEOSIDE_TRIPHOSPHATE_METABOLIC_PROCESS | 0.116442 | 0.085336 | 0.281381 | 0.291136 | 0.309458 | 0.236999 |  | 0.251592 |
| GOBP_REGULATION_OF_OXIDATIVE_STRESS_INDUCED_INTRINSIC_APOPTOTIC_SIGNALING_PATHWAY | 0.116282 | 0.282264 | 0.393985 | 0.206512 | 0.239832 | 0.297227 | 0.139523 | 0.145086 |
| GOBP_ASPARTATE_FAMILY_AMINO_ACID_METABOLIC_PROCESS | 0.115665 |  | 0.239719 | 0.126109 | 0.127374 | 0.147873 | 0.135754 | 0.115612 |
| MOOTHA_GLYCOGEN_METABOLISM | 0.113673 | 0.179978 | 0.270354 | 0.14327 | 0.107724 | 0.140577 | 0.02646 | 0.042306 |
| KEGG_AMINO_SUGAR_AND_NUCLEOTIDE_SUGAR_METABOLISM | 0.113513 | 0.216142 | 0.303298 | 0.072483 | 0.008431 | 0.089153 | -0.017284 | -0.028398 |
| GOBP_GLYCOSYL_COMPOUND_METABOLIC_PROCESS | 0.113502 | 0.018145 | 0.176365 | 0.255146 | 0.270131 | 0.231992 | 0.28533 | 0.18529 |
| GOBP_PHOSPHATIDYLGLYCEROL_METABOLIC_PROCESS | 0.113452 | 0.104458 | 0.177805 | 0.128038 | 0.096572 | 0.120687 | 0.018292 | 0.008086 |
| MOOTHA_GLUCONEOGENESIS | 0.110934 | 0.17393 | 0.482972 | 0.222303 | 0.169202 | 0.213435 | 0.110664 | 0.096096 |
| GOBP_NUCLEOSIDE_METABOLIC_PROCESS | 0.109569 | 0.022263 | 0.176883 | 0.24993 | 0.261732 | 0.228356 | 0.28076 | 0.183172 |
| GOBP_PEPTIDE_METABOLIC_PROCESS | 0.10942 | 0.089609 | 0.380024 | 0.328433 | 0.419818 | 0.391279 |  | 0.242248 |
| KEGG_PURINE_METABOLISM | 0.105904 | 0.010498 | 0.191201 | 0.266875 | 0.296164 | 0.244325 | 0.33748 | 0.185275 |
| GOBP_CELLULAR_KETONE_METABOLIC_PROCESS | 0.099292 | 0.16791 | 0.374932 | 0.276278 | 0.34224 | 0.378569 | 0.215716 | 0.211275 |
| REACTOME_REGULATION_OF_CHOLESTEROL_BIOSYNTHESIS_BY_SREBP_SREBF | 0.099065 | -0.002286 | 0.142232 | 0.30091 | 0.300478 | 0.253588 | 0.12971 | 0.141811 |
| GOBP_REGULATION_OF_MITOCHONDRIAL_OUTER_MEMBRANE_PERMEABILIZATION_INVOLVED_IN_APOPTOTIC_SIGNALING_PATHWAY | 0.097673 | 0.042171 | 0.236145 | 0.267242 | 0.375788 | 0.408205 | 0.221778 | 0.192312 |
| GOBP_REGULATION_OF_CELLULAR_AMIDE_METABOLIC_PROCESS | 0.095261 | 0.135132 | 0.416263 | 0.334385 | 0.478135 | 0.482786 | 0.250726 |  |
| GOBP_REGULATION_OF_MITOCHONDRIAL_MEMBRANE_PERMEABILITY | 0.08956 | 0.040404 | 0.313382 | 0.307053 | 0.391961 | 0.424137 | 0.193464 | 0.156627 |
| GOBP_POSITIVE_REGULATION_OF_UBIQUITIN_DEPENDENT_PROTEIN_CATABOLIC_PROCESS | 0.086943 | 0.130533 | 0.324452 | 0.268234 | 0.340423 | 0.371217 | 0.116969 | 0.1929 |
| KEGG_GALACTOSE_METABOLISM | 0.085756 | 0.103969 | 0.184704 | 0.117288 | 0.060707 | 0.069962 | 0.031089 | 0.01682 |
| REACTOME_GLUCOSE_METABOLISM | 0.083473 | 0.181666 | 0.470412 | 0.227988 | 0.248533 | 0.294406 | 0.16883 | 0.148978 |
| KEGG_CYSTEINE_AND_METHIONINE_METABOLISM | 0.080287 | 0.100345 | 0.267305 | 0.210845 | 0.238283 | 0.245326 | 0.182918 | 0.16749 |
| GOBP_IRON_SULFUR_CLUSTER_ASSEMBLY | 0.076463 | 0.021535 | 0.199024 | 0.114994 | 0.147148 | 0.17348 | 0.102722 | 0.087114 |
| GOBP_POSITIVE_REGULATION_OF_MITOCHONDRIAL_DEPOLARIZATION | 0.076261 |  | 0.068872 | 0.141458 | 0.207331 | 0.201299 | 0.13605 | 0.068875 |
| REACTOME_METABOLISM_OF_AMINO_ACIDS_AND_DERIVATIVES | 0.07555 | 0.019139 | 0.301598 | 0.304782 | 0.364106 | 0.306233 | 0.258522 | 0.225936 |
| GOBP_REGULATION_OF_GLUCOSE_METABOLIC_PROCESS | 0.074737 | 0.100222 | 0.254789 | 0.199122 | 0.263946 | 0.265 | 0.117378 | 0.142243 |
| GOBP_GLUCOCORTICOID_METABOLIC_PROCESS | 0.073354 | 0.202673 | 0.236866 | 0.033495 | 0.031261 |  | -0.03737 | -0.03091 |
| GOBP_CELLULAR_METABOLIC_COMPOUND_SALVAGE | 0.07246 | 0.071131 | 0.151149 |  | 0.077767 | 0.118644 | 0.095853 | 0.013786 |
| GOBP_REGULATION_OF_MITOCHONDRIAL_ATP_SYNTHESIS_COUPLED_ELECTRON_TRANSPORT | 0.072015 | -0.024707 | 0.059273 | 0.103241 | 0.1304 | 0.095562 | 0.255099 | 0.405193 |
| GOBP_POSITIVE_REGULATION_OF_CELLULAR_AMIDE_METABOLIC_PROCESS | 0.067087 | 0.108523 | 0.36512 | 0.298009 | 0.434659 | 0.404078 | 0.251152 | 0.249358 |
| GOBP_NADP_METABOLIC_PROCESS | 0.065867 |  | 0.093804 | 0.193288 | 0.217195 | 0.172087 | 0.201294 | 0.176888 |
| GOBP_PYRIMIDINE_CONTAINING_COMPOUND_METABOLIC_PROCESS | 0.065039 | -0.045725 | 0.053477 | 0.213223 |  | 0.162949 | 0.462544 | 0.245191 |
| GOBP_TETRAHYDROFOLATE_METABOLIC_PROCESS | 0.064879 | -0.023204 | 0.048804 | 0.089852 | 0.088329 | 0.037528 | 0.369283 |  |
| GOBP_POSITIVE_REGULATION_OF_NUCLEOBASE_CONTAINING_COMPOUND_METABOLIC_PROCESS | 0.063252 | 0.115166 | 0.355705 | 0.323719 | 0.534649 | 0.516339 | 0.293978 | 0.344586 |
| GOBP_PURINE_NUCLEOBASE_METABOLIC_PROCESS | 0.062943 | 0.005421 | 0.056002 | 0.073973 | 0.048967 | 0.016271 | 0.122684 | 0.05226 |
| GOBP_MITOCHONDRIAL_CYTOCHROME_C_OXIDASE_ASSEMBLY | 0.062129 | 0.0544 | 0.183509 | 0.148748 | 0.207108 | 0.221774 | 0.152942 | 0.117438 |
| GOBP_SULFUR_AMINO_ACID_METABOLIC_PROCESS | 0.059557 | 0.027583 | 0.087665 | 0.086853 | 0.11473 | 0.128884 | 0.093853 | 0.064131 |
| KEGG_ETHER_LIPID_METABOLISM | 0.058787 | 0.09534 | 0.127297 | 0.136401 | 0.148914 | 0.218061 | 0.037403 | 0.042575 |
| GOBP_MITOCHONDRIAL_RNA_METABOLIC_PROCESS | 0.058657 | -0.000247 | 0.112896 | 0.11509 | 0.147693 | 0.152088 |  |  |
| GOBP_POSITIVE_REGULATION_OF_MITOCHONDRION_ORGANIZATION | 0.053131 | 0.043214 | 0.298936 | 0.262953 | 0.412015 |  | 0.188109 | 0.170639 |
| GOBP_PTERIDINE_CONTAINING_COMPOUND_METABOLIC_PROCESS | 0.050082 | -0.03724 | 0.043018 | 0.106344 | 0.127659 | 0.074869 | 0.363004 | 0.206476 |
| GOBP_GTP_METABOLIC_PROCESS | 0.047963 | 0.004882 | 0.157862 | 0.227634 | 0.246364 | 0.186999 | 0.21698 | 0.1589 |
| GOBP_GLUCOSE_6_PHOSPHATE_METABOLIC_PROCESS | 0.045605 | 0.046838 | 0.145544 | 0.147346 | 0.135655 | 0.113354 | 0.115105 | 0.057411 |
| WP_ENERGY_METABOLISM | 0.044356 | 0.00975 | 0.099916 | 0.105566 | 0.132979 | 0.129464 | -0.005013 | 0.027215 |
| GOBP_NUCLEOBASE_METABOLIC_PROCESS | 0.043489 | -0.030209 | 0.027563 | 0.128044 | 0.164562 | 0.088372 | 0.371896 | 0.205109 |
| REACTOME_SYNTHESIS_OF_PA | 0.041907 | 0.041786 | 0.016432 | 0.078533 | 0.014487 | 0.003837 | -0.03627 | -0.023367 |
| KEGG_GLYCOSPHINGOLIPID_BIOSYNTHESIS_LACTO_AND_NEOLACTO_SERIES | 0.041687 | 0.013192 | 0.015934 | 0.010954 | -0.02644 | -0.01603 | -0.030414 | -0.033809 |
| GOBP_PYRIMIDINE_NUCLEOTIDE_METABOLIC_PROCESS | 0.038792 | -0.03457 | 0.073617 | 0.201618 | 0.240861 | 0.154564 | 0.414974 | 0.211577 |
| GOBP_ESTABLISHMENT_OF_MITOCHONDRION_LOCALIZATION | 0.033804 | 0.039685 | 0.182512 | 0.219403 | 0.311154 | 0.449301 | 0.050311 | 0.088227 |
| GOBP_AMINE_METABOLIC_PROCESS | 0.032556 | 0.05387 | 0.280002 | 0.289598 | 0.422163 | 0.438048 | 0.263877 | 0.253987 |
| GOBP_NUCLEOTIDE_SUGAR_METABOLIC_PROCESS | 0.029187 | 0.099948 | 0.18695 | 0.060235 | 0.064633 | 0.106948 | 0.022735 | 0.026151 |
| WP_EICOSANOID_SYNTHESIS | 0.022928 | 0.191285 | 0.173689 | 0.014791 | -0.063347 | 0.025305 | -0.137414 | -0.116288 |
| GOBP_PYRIMIDINE_NUCLEOSIDE_METABOLIC_PROCESS | 0.021455 | -0.028061 | 0.014073 | 0.092059 | 0.138681 | 0.11013 | 0.264356 | 0.19735 |
| GOBP_RESPONSE_TO_FATTY_ACID | 0.020744 | 0.081596 | 0.154016 | 0.171973 | 0.240804 | 0.258875 | 0.141977 | 0.164901 |
| GOBP_PYRIMIDINE_NUCLEOSIDE_TRIPHOSPHATE_METABOLIC_PROCESS | 0.017014 | -0.076861 | 0.038811 | 0.205692 | 0.260662 | 0.152031 | 0.460501 | 0.252045 |
| REACTOME_PI_METABOLISM | 0.016684 | 0.018082 | 0.142513 | 0.175572 | 0.232731 | 0.246816 | -0.038705 | 0.023737 |
| REACTOME_SYNTHESIS_OF_SUBSTRATES_IN_N_GLYCAN_BIOSYTHESIS | 0.009348 | 0.058896 | 0.179604 | 0.134944 |  | 0.202453 | 0.017615 | -0.008218 |
| WP_LIPID_METABOLISM_PATHWAY | 0.00826 | 0.020106 | 0.161588 | 0.158969 | 0.2005 |  | 0.056757 | 0.076316 |
| GOBP_OLIGOSACCHARIDE_LIPID_INTERMEDIATE_BIOSYNTHETIC_PROCESS | -0.001983 | -0.03168 | 0.000157 | 0.080741 | 0.070322 | 0.023874 | 0.030445 | 0.01857 |
| GOBP_NEGATIVE_REGULATION_OF_NUCLEOBASE_CONTAINING_COMPOUND_METABOLIC_PROCESS | -0.004518 | 0.04148 | 0.313044 | 0.323249 | 0.586121 | 0.574589 | 0.315589 | 0.356336 |
| GOBP_REGULATION_OF_UBIQUITIN_DEPENDENT_PROTEIN_CATABOLIC_PROCESS | -0.004531 | 0.04653 | 0.331996 | 0.309065 | 0.469759 | 0.488826 | 0.19137 | 0.260539 |
| GOBP_PYRIMIDINE_DEOXYRIBONUCLEOTIDE_METABOLIC_PROCESS | -0.005173 | -0.041205 | 0.031476 | 0.120593 | 0.192773 | 0.132062 | 0.469859 | 0.253919 |
| REACTOME_SELENOAMINO_ACID_METABOLISM | -0.008668 | -0.023573 | 0.249051 | 0.241929 | 0.33452 | 0.261874 | 0.240336 | 0.211815 |
| GOBP_DEOXYRIBONUCLEOTIDE_METABOLIC_PROCESS | -0.016698 | -0.078442 | 0.041074 | 0.151862 | 0.263256 | 0.205727 | 0.566916 | 0.325226 |
| GOBP_UBIQUITIN_DEPENDENT_PROTEIN_CATABOLIC_PROCESS_VIA_THE_MULTIVESICULAR_BODY_SORTING_PATHWAY | -0.022297 | -0.034526 | 0.10887 | 0.151762 | 0.243709 | 0.243283 | 0.089067 | 0.110127 |
| GOBP_INTRACELLULAR_DISTRIBUTION_OF_MITOCHONDRIA | -0.028893 | -0.105375 |  | 0.132074 | 0.256408 | 0.250605 | 0.07617 | 0.085489 |
| GOBP_REGULATION_OF_CELLULAR_AMINE_METABOLIC_PROCESS | -0.032591 | -0.0132 | 0.234079 | 0.280771 | 0.456874 | 0.452387 | 0.312794 | 0.298713 |
| GOBP_REGULATION_OF_CELLULAR_AMINO_ACID_METABOLIC_PROCESS | -0.04837 | -0.028022 | 0.230125 | 0.281484 | 0.467929 | 0.460725 | 0.317207 | 0.307245 |
| REACTOME_OXIDATIVE_STRESS_INDUCED_SENESCENCE | -0.050835 | 0.028494 | 0.24422 | 0.23725 | 0.47046 | 0.491718 | 0.279652 | 0.288012 |
| GOBP_PEPTIDYL_GLUTAMIC_ACID_MODIFICATION | -0.056498 | -0.084601 | 0.003694 | 0.130272 | 0.185656 | 0.160857 | 0.17915 | 0.141176 |
| GOBP_CELLULAR_RESPONSE_TO_FATTY_ACID | -0.083059 | 0.02834 | 0.088253 | 0.129311 | 0.254428 | 0.281033 | 0.185324 | 0.209264 |
| WP_METABOLISM_OF_SPINGOLIPIDS_IN_ER_AND_GOLGI_APPARATUS | -0.170782 | -0.10816 | 0.003978 | 0.107777 | 0.255632 | 0.317765 |  |  |
| GOBP_MITOCHONDRION_DISTRIBUTION | -0.176058 | -0.17132 |  | 0.153389 | 0.349114 | 0.401893 | 0.05123 | 0.085 |
| GOBP_NEGATIVE_REGULATION_OF_UBIQUITIN_PROTEIN_TRANSFERASE_ACTIVITY | -0.189565 | -0.117502 | 0.161301 | 0.188783 | 0.41351 |  | 0.268146 | 0.276228 |
| GOBP_INOSITOL_LIPID_MEDIATED_SIGNALING |  | 0.343787 | 0.342424 | 0.286167 | 0.018739 | 0.008504 | -0.08965 |  |
| GOBP_UNSATURATED_FATTY_ACID_BIOSYNTHETIC_PROCESS |  | 0.322976 | 0.260334 | 0.342201 | 0.034309 | -0.008773 | -0.003546 | -0.010239 |
| GOBP_GLUTAMINE_FAMILY_AMINO_ACID_METABOLIC_PROCESS |  | 0.113451 | 0.188755 | 0.139445 | 0.096204 | 0.135084 | 0.053355 | 0.045155 |
| GOBP_ALPHA_AMINO_ACID_METABOLIC_PROCESS |  | 0.086184 |  | 0.264627 | 0.202305 | 0.205268 | 0.159407 | 0.11974 |
| GOBP_PYRIMIDINE_RIBONUCLEOTIDE_METABOLIC_PROCESS |  | 0.030639 | 0.122465 | 0.183006 | 0.164363 | 0.11356 | 0.151301 | 0.053297 |
| GOBP_GLYCEROL_ETHER_METABOLIC_PROCESS |  | -0.009895 | 0.130812 | 0.142825 | 0.224901 | 0.296006 | 0.069336 | 0.076917 |
