## Supplementary material for "A cell state specific metabolic vulnerability to GPX4-dependent ferroptosis in glioblastoma": Table S7

| **Tissue ID** | **Age range** | **Sex** | **Location** | **Diagnosis** | **IDH1 Status** | **EGFR status** |
| --- | --- | --- | --- | --- | --- | --- |
| **TB6181** | 30-39 | F | Frontal and temporal | Anaplastic Astrocytoma, WHO Grade III, IDH1 Mutant | Mutant | Unamplified |
| **TB6328** | 70-79 | F | Left frontal | Glioblastoma, WHO Grade IV | Wildtype | Amplified |
| **TB6458** | 50-59 | M | Right frontal | Glioblastoma, WHO Grade IV | Wildtype | Amplified |
| **TB6528** | 60-69 | F | Left temporal | Glioblastoma, WHO Grade IV | Wildtype | Unamplified |
| **TB6534** | 50-59 | F | Right temporal | Glioblastoma, WHO Grade IV | Wildtype | Unamplified |
| **TB6545** | 30-39 | M | Right Frontal | Glioblastoma, WHO Grade IV | Wildtype | Unamplified |
