## Supplementary material for "A cell state specific metabolic vulnerability to GPX4-dependent ferroptosis in glioblastoma": Table S8

| **GID** | **gene** | **TB6181_log2fc** | **TB6181_qval** | **TB6328_log2fc** | **TB6328_qval** | **TB6458_log2fc** | **TB6458_qval** | **TB6528_log2fc** | **TB6528_qval** | **TB6534_log2fc** | **TB6534_qval** | **TB6545_log2fc** | **TB6545_qval** |
| --- | --- | --- | --- | --- | --- | --- | --- | --- | --- | --- | --- | --- | --- |
| ENSG00000000003.14 | TSPAN6 | -0.989652 | 0.8867323 | 0.421091 | 0.8217443 | 0.550785 | 0.8753365 | 0.480859 | 0.7769736 | -1.386636 | 0.03192416 | 0.809346 | 0.9169962 |
| ENSG00000000419.12 | DPM1 | -2.080898 | 0.4406957 | 0.480441 | 0.638737 | -0.63569 | 0.1228759 | -0.105214 | 0.01683283 | -0.494827 | 0.179554 | 2.232364 | 0.6885065 |
| ENSG00000001167.14 | NFYA | -2.996422 | 1 | 0.789664 | 0.3596909 | 0.879479 | 0.7424 | 1.956041 | 0.000429593 | 2.100925 | 0.3974318 | -1.025204 | 1 |
| ENSG00000001629.9 | ANKIB1 | -0.311539 | 0.9753514 | -0.10489 | 0.6527736 | 0.956312 | 0.2492552 | 0.389258 | 0.02154744 | -0.153842 | 0.7449576 | 0.378 | 0.7767294 |
| ENSG00000002330.13 | BAD | -0.215489 | 1 | 0.222061 | 0.5012872 | -0.362561 | 0.002338449 | -0.672631 | 8.62E-05 | 0.308809 | 0.2880317 | -0.215501 | 0.6885065 |
| ENSG00000002834.17 | LASP1 | -0.358143 | 1 | -0.11383 | 0.7383998 | 0.884842 | 0.000163373 | -0.213189 | 0.859973 | 0.285006 | 0.691416 | 0.773733 | 0.6885065 |
| ENSG00000003056.7 | M6PR | -3.717379 | 0.7648042 | 0.170953 | 0.7613037 | 0.922134 | 0.01044833 | -0.014554 | 1 | 0.58371 | 0.8115805 | 0.315305 | 0.6885065 |
| ENSG00000003756.16 | RBM5 | -0.808736 | 1 | -0.474253 | 0.3225827 | 0.724058 | 0.01383094 | 0.981453 | 0.007865799 | 1.484438 | 0.05297091 | 0.654488 | 1 |
| ENSG00000004059.10 | ARF5 | 0.524837 | 0.9791227 | 1.684677 | 0.000556513 | -1.400135 | 0.000878597 | -0.579059 | 0.000160061 | -0.137167 | 0.6811951 | -1.123055 | 1 |
| ENSG00000004139.13 | SARM1 | -0.443216 | 0.6214382 | 0.433756 | 0.9114157 | 0.562273 | 1 | 1.400142 | 0.04278836 | 1.764143 | 0.002293693 | -0.426064 | 0.9576568 |
| ENSG00000004142.11 | POLDIP2 | -0.81166 | 0.529676 | -0.277781 | 0.1940102 | -0.382023 | 0.002309016 | 0.47054 | 1 | 0.288379 | 0.9768841 | 0.296441 | 0.8704093 |
| ENSG00000004455.16 | AK2 | 0.059778 | 0.9338818 | 0.73022 | 0.6527736 | -0.234133 | 0.02543885 | -0.286652 | 0.2047258 | -0.027783 | 0.1372704 | 0.509448 | 0.9787797 |
| ENSG00000004487.15 | KDM1A | -0.111866 | 0.9547114 | -0.357508 | 0.1141252 | -0.179421 | 0.9505825 | 0.885398 | 0.03460858 | 1.668311 | 0.000168448 | -0.903744 | 0.826754 |
| ENSG00000004534.14 | RBM6 | -0.258623 | 0.5063049 | 0.02501 | 0.6527736 | 0.336667 | 0.3018258 | 1.408778 | 7.79E-05 | 0.624605 | 0.23099 | -1.4724 | 0.6185816 |
| ENSG00000004700.15 | RECQL | -0.185466 | 0.8926671 | 0.60417 | 0.5884985 | 0.66454 | 0.005172759 | 0.329709 | 1 | 0.500454 | 0.2226201 | -0.97795 | 0.8056213 |
| ENSG00000004779.9 | NDUFAB1 | 0.789543 | 0.000182873 | -0.162037 | 0.3583128 | -0.325546 | 0.000737771 | -0.275445 | 0.006869538 | -0.060512 | 0.1399344 | -0.52662 | 0.6829816 |
| ENSG00000004838.13 | ZMYND10 | -1.195661 | 0.714717 | 1.821049 | 0.7585627 | 1.943325 | 0.01760987 | 2.545508 | 0.542299 | 3.06451 | 0.2363864 | inf | 0.6885065 |
| ENSG00000004864.13 | SLC25A13 | -1.413138 | 0.6603817 | -0.065744 | 0.6527736 | -0.928423 | 0.009393306 | 1.473049 | 0.7858185 | 0.838652 | 0.9510304 | 1.197682 | 0.6885065 |
| ENSG00000004866.18 | ST7 | 0.207489 | 0.8087883 | 0.173468 | 0.9104976 | 1.141675 | 0.01710394 | 0.051086 | 0.6711222 | -0.230854 | 0.676985 | 0.773354 | 0.6885065 |
| ENSG00000005007.12 | UPF1 | -1.819728 | 0.3166974 | -0.635934 | 0.1549389 | -0.743982 | 0.2636292 | -0.017454 | 0.211072 | 1.512424 | 0.02624132 | 0.773802 | 0.6829816 |
| ENSG00000005022.5 | SLC25A5 | 0.176156 | 0.02363633 | -0.08847 | 0.4114546 | -0.028006 | 0.6774982 | 0.28902 | 0.02584346 | -0.110005 | 0.000327706 | 0.225605 | 0.5841302 |
| ENSG00000005059.15 | CCDC109B | -0.227961 | 0.957965 | 0.049524 | 0.7062745 | -1.151044 | 0.002464869 | -0.172076 | 0.004713837 | 0.146812 | 0.8273528 | 0.425729 | 0.8479431 |
| ENSG00000005075.15 | POLR2J | -0.183789 | 0.7063838 | 0.054031 | 0.6077765 | -0.437674 | 5.63E-07 | 0.044933 | 0.8197402 | 0.168667 | 0.003418459 | -0.488335 | 0.6091918 |
| ENSG00000005100.12 | DHX33 | -0.101437 | 0.3946735 | -1.014533 | 0.05089515 | 1.134337 | 0.1998418 | 2.264027 | 0.000198323 | 1.290708 | 0.113469 | -2.540462 | 0.5246903 |
| ENSG00000005108.15 | THSD7A | -1.554149 | 0.6214382 | -0.483294 | 0.8363543 | 1.7799 | 0.4405949 | 2.332739 | 0.02786801 | 0.862937 | 0.01618785 | -3.891577 | 0.6861518 |
| ENSG00000005206.16 | SPPL2B | -2.62269 | 1 | -0.319487 | 0.592336 | -1.21951 | 0.8454005 | 1.989952 | 0.003218051 | 2.041009 | 0.3649717 | 0.655262 | 0.8856541 |
| ENSG00000005339.13 | CREBBP | -1.062334 | 1 | -0.239878 | 0.4886958 | 0.715085 | 0.05491754 | 1.246492 | 0.000276572 | 2.298457 | 0.000591261 | 0.257349 | 0.8489889 |
| ENSG00000005483.19 | KMT2E | -0.048647 | 0.8767328 | -0.11895 | 0.9839225 | 0.963281 | 1.11E-14 | 0.673525 | 0.1370833 | 0.643917 | 0.3974318 | -0.622621 | 0.6829816 |
| ENSG00000005513.9 | SOX8 | 0.124407 | 0.9858184 | -0.519518 | 0.09212669 | -0.359364 | 0.2754647 | 0.525443 | 0.330538 | 1.615238 | 0.00012503 | 0.622504 | 0.9780545 |
| ENSG00000005700.14 | IBTK | -0.622116 | 0.5355247 | -0.172993 | 0.1534359 | 0.836536 | 0.03002101 | 0.613292 | 0.000284065 | 1.560355 | 0.8965775 | -0.85932 | 1 |
| ENSG00000005801.16 | ZNF195 | 0.323022 | 0.6214382 | -0.070231 | 0.6527736 | 0.198256 | 0.2213291 | 0.355391 | 0.01816429 | 0.12508 | 0.7373849 | 2.102021 | 0.7501684 |
| ENSG00000005810.17 | MYCBP2 | 0.895756 | 0.6790677 | 0.56461 | 0.8689748 | 0.037319 | 0.09629423 | 2.619879 | 0.000284065 | 1.134231 | 0.001349185 | -1.544764 | 0.8479431 |
| ENSG00000005812.10 | FBXL3 | -0.527636 | 0.654369 | -0.322838 | 0.5245375 | -0.250331 | 0.5290845 | -0.089349 | 0.4762125 | 0.037517 | 0.04118451 | 1.797607 | 1 |
| ENSG00000005884.17 | ITGA3 | 0.113011 | 0.6214382 | -0.203393 | 0.6527736 | -0.32471 | 0.7437308 | 0.990552 | 0.01391062 | -0.590611 | 0.934012 | 0.808893 | 0.6576742 |
| ENSG00000005893.15 | LAMP2 | -1.233977 | 0.6931569 | -0.099972 | 0.8953649 | 0.433309 | 0.195658 | -1.491744 | 0.01045498 | 0.285224 | 0.920639 | 1.725779 | 0.5344421 |
| ENSG00000006007.11 | GDE1 | -0.417681 | 0.8480161 | -0.118186 | 0.592336 | -0.066169 | 0.6784963 | 1.472504 | 0.01271688 | 0.734226 | 0.9587693 | 0.202517 | 0.960853 |
| ENSG00000006015.17 | C19orf60 | -1.477751 | 0.4764974 | 0.010269 | 0.8678306 | -0.882788 | 0.08991471 | -0.197915 | 0.2754358 | -0.951126 | 0.008521047 | 0.950934 | 0.6885065 |
| ENSG00000006125.16 | AP2B1 | -0.154597 | 0.9706169 | -0.332374 | 0.7926154 | 0.425185 | 0.09230131 | 1.091094 | 0.01067607 | 0.304038 | 0.1688763 | 0.004031 | 0.6699469 |
| ENSG00000006327.13 | TNFRSF12A | -0.027132 | 0.6214382 | -0.222788 | 0.5894122 | -0.62313 | 1.01E-07 | -0.046378 | 0.08321889 | -0.087948 | 0.1119361 | -0.273333 | 0.3565625 |
| ENSG00000006451.7 | RALA | 0.060468 | 0.8006286 | 0.017595 | 0.7696508 | 0.15209 | 0.004077784 | 0.222763 | 0.9463108 | -0.268992 | 0.0129887 | 0.102344 | 0.6885065 |
| ENSG00000006468.13 | ETV1 | -0.324926 | 0.7117094 | -0.760123 | 0.000132809 | 1.348498 | 1.58E-11 | 0.058662 | 0.6075806 | -0.019869 | 0.01425771 | -0.945692 | 0.7045294 |
| ENSG00000006634.7 | DBF4 | -2.082774 | 0.6214382 | -0.443154 | 0.4961426 | 1.055511 | 0.000429525 | -0.524317 | 0.3508645 | 1.141697 | 0.7670514 | 0.735231 | 0.6885065 |
| ENSG00000006712.14 | PAF1 | -1.101534 | 0.4373461 | 0.123863 | 1 | -0.151187 | 0.5468728 | -0.171618 | 0.01799326 | -0.305874 | 0.01667537 | -0.15083 | 0.6885065 |
| ENSG00000006744.18 | ELAC2 | -0.941456 | 0.6364041 | 0.172263 | 0.8790165 | -0.910221 | 0.01286473 | 1.457248 | 0.000135252 | 1.58352 | 0.08170657 | -0.696959 | 0.4553016 |
| ENSG00000007080.10 | CCDC124 | -0.13126 | 0.9438474 | 0.355018 | 0.145165 | -0.465531 | 1.58E-05 | 0.81386 | 0.5668523 | -0.276245 | 0.1924287 | 1.740834 | 0.8863181 |
| ENSG00000007168.12 | PAFAH1B1 | -0.298182 | 0.714717 | -0.367692 | 0.2038521 | 0.816389 | 2.38E-07 | 0.477038 | 0.2982183 | 0.368386 | 0.5196843 | -0.827662 | 0.03724323 |
| ENSG00000007202.14 | KIAA0100 | -0.648708 | 0.3408865 | -0.113211 | 0.4922064 | -0.280794 | 0.4089974 | 2.175033 | 0.01870689 | 1.83787 | 0.07708632 | 1.213898 | 0.8580104 |
| ENSG00000007520.3 | TSR3 | -1.103715 | 0.6014343 | 0.119356 | 0.9854207 | -0.861699 | 0.000551725 | 0.578964 | 0.7704057 | -0.287181 | 0.178149 | 0.431233 | 0.6829816 |
| ENSG00000008083.13 | JARID2 | 1.546644 | 1 | 1.092054 | 0.4229072 | 2.06473 | 0.2000675 | 0.17897 | 0.002860427 | 1.052728 | 0.01550373 | 0.740277 | 0.6885065 |
| ENSG00000008128.22 | CDK11A | 0.211308 | 0.529676 | 0.648104 | 0.7549722 | -0.116753 | 0.2720913 | 2.995323 | 0.1698448 | 2.022569 | 0.6817312 |  |  |
| ENSG00000008130.15 | NADK | 0.608169 | 0.9183025 | -0.334262 | 0.6527736 | -1.014995 | 0.05669581 | 0.669687 | 0.006041353 | 0.96595 | 0.9795933 | 1.729503 | 0.6517664 |
| ENSG00000008256.15 | CYTH3 | 2.503947 | 0.8587992 | -0.344583 | 0.9790409 | 0.511672 | 0.5290845 | 1.869014 | 0.03640873 | 1.589452 | 0.02344635 | 0.545403 | 0.841154 |
| ENSG00000008277.14 | ADAM22 | 1.186734 | 0.6214382 | -0.556256 | 0.6834954 | 0.981603 | 0.08427854 | 0.747269 | 0.04248025 | 1.859824 | 0.000577827 | 0.372174 | 0.8949489 |
| ENSG00000008294.20 | SPAG9 | 0.081732 | 0.8867323 | -0.038404 | 0.6527736 | 0.567302 | 0.08175111 | 0.914884 | 0.000141599 | 1.276671 | 0.1403834 | -1.592509 | 0.6885065 |
| ENSG00000008300.14 | CELSR3 | 2.114747 | 0.4764974 | -0.836262 | 0.9104976 | 0.571342 | 1 | 4.211545 | 0.000271907 | 2.712421 | 0.004636219 | -0.405276 | 1 |
| ENSG00000008394.12 | MGST1 | -1.835278 | 0.529676 | 0.583736 | 0.6378755 | -1.324602 | 0.00452368 | 0.5266 | 0.2335397 | -2.004313 | 0.004641664 | 2.453354 | 0.5202568 |
| ENSG00000008405.11 | CRY1 | -1.808665 | 0.9047689 | 0.646577 | 0.6068884 | 0.389123 | 0.04799587 | 0.57726 | 0.2699839 | 0.971486 | 0.8080085 | -0.199807 | 0.9264343 |
| ENSG00000008441.16 | NFIX | -1.623939 | 0.008205756 | 0.225258 | 0.9507102 | 0.022082 | 0.6716532 | 1.192205 | 1.70E-06 | 1.067669 | 6.28E-06 | -0.494934 | 0.6885065 |
| ENSG00000008710.17 | PKD1 | 0.286927 | 0.8415359 | 0.999366 | 0.4723757 | -0.038997 | 0.7887553 | 1.662604 | 2.31E-05 | 2.254154 | 2.83E-07 | 0.407115 | 1 |
| ENSG00000008735.13 | MAPK8IP2 | -1.0867 | 0.7405237 | 0.539852 | 0.9452791 | -0.782774 | 0.1402812 | 1.560989 | 0.03537571 | 0.027049 | 0.3974318 | 2.985796 | 0.9147227 |
| ENSG00000008869.11 | HEATR5B | 0.625571 | 0.9198482 | -0.807359 | 0.6527736 | 0.117049 | 0.4787829 | 0.629644 | 0.03903959 | 1.859267 | 0.030568 | 0.585806 | 1 |
| ENSG00000008952.16 | SEC62 | -0.489496 | 0.9671207 | 0.249119 | 0.9156272 | 0.892401 | 1.99E-10 | -0.001378 | 0.3149163 | -0.152511 | 0.04675253 | -0.248545 | 0.6885065 |
| ENSG00000008988.9 | RPS20 | -1.001456 | 5.00E-08 | 0.080125 | 0.7418188 | 0.191779 | 0.0412862 | -0.172364 | 0.00013631 | -0.606401 | 6.15E-10 | -0.243882 | 0.04261865 |
| ENSG00000009307.15 | CSDE1 | 0.599397 | 0.8006286 | -0.185257 | 0.02222377 | 0.533362 | 2.92E-06 | 0.940767 | 0.000601717 | 0.42942 | 0.5196843 | -0.469828 | 0.2768456 |
| ENSG00000009335.17 | UBE3C | -1.606483 | 0.456765 | 0.256545 | 0.8263112 | 0.720734 | 0.1291104 | 2.222196 | 5.07E-08 | 0.335247 | 0.02215221 | 0.006084 | 1 |
| ENSG00000009413.15 | REV3L | -0.632017 | 0.8867323 | -0.451263 | 0.1940102 | 1.021432 | 0.000150194 | 1.742239 | 0.001282667 | 1.074257 | 0.03497153 | -0.358631 | 0.8708371 |
| ENSG00000009694.13 | TENM1 | 0.89615 | 0.8006286 | -0.641129 | 0.8263112 | 1.06429 | 0.001675839 | 0.68713 | 0.7144938 | 0.78255 | 0.4278631 | 0.231009 | 0.8949489 |
| ENSG00000009844.15 | VTA1 | 0.432796 | 0.563505 | 0.585734 | 0.521561 | 0.833646 | 0.3637026 | 1.701753 | 0.004197312 | 0.947096 | 0.06633533 | 0.361408 | 0.7577666 |
| ENSG00000009954.10 | BAZ1B | -0.019897 | 0.980017 | -0.081692 | 0.7056806 | 0.905998 | 5.90E-14 | 1.512637 | 1.36E-10 | 0.808957 | 0.2232993 | -0.402544 | 0.5344421 |
| ENSG00000010017.12 | RANBP9 | -0.641622 | 0.8006286 | -0.57852 | 0.3583128 | 0.073211 | 0.9579526 | 1.539103 | 0.02147156 | 0.171825 | 0.6626174 | 1.646788 | 0.8768029 |
| ENSG00000010165.19 | METTL13 | 0.518062 | 0.714717 | -0.543064 | 0.5010139 | -0.203895 | 0.5641573 | 1.504925 | 0.03084244 | 0.621619 | 0.5485224 | -2.596639 | 0.2856007 |
| ENSG00000010219.13 | DYRK4 | -0.282796 | 0.8867323 | -0.398804 | 0.3803801 | -0.187265 | 0.03835208 | 0.057849 | 0.01367005 | -0.701895 | 0.1102102 | -0.905016 | 0.6885065 |
| ENSG00000010244.16 | ZNF207 | 0.619248 | 0.7225091 | -0.483813 | 0.5509719 | 0.132364 | 0.6554376 | 1.046227 | 0.000345508 | 1.68472 | 0.003204401 | -0.108274 | 0.8711089 |
| ENSG00000010256.10 | UQCRC1 | 0.777002 | 0.4046488 | -0.04293 | 0.5736784 | -0.09938 | 0.09067289 | 0.14181 | 0.2541225 | -0.18307 | 0.02544541 | 0.838177 | 0.5203855 |
| ENSG00000010278.11 | CD9 | -1.350046 | 0.6214382 | 0.235547 | 0.5240506 | 0.255712 | 1 | -0.322079 | 1.36E-06 | -0.556341 | 0.01557237 | -0.359521 | 0.8178576 |
| ENSG00000010322.15 | NISCH | 0.829752 | 1 | -0.044002 | 0.9081785 | -0.215139 | 0.1489368 | 0.200062 | 0.02479946 | 1.139947 | 0.2306098 | -0.435628 | 0.7494343 |
| ENSG00000010404.17 | IDS | -0.13881 | 0.8006286 | 0.255349 | 1 | 0.432473 | 0.01953349 | 1.189205 | 0.000791326 | 0.939421 | 0.09215864 | -0.051322 | 0.9554184 |
| ENSG00000010810.17 | FYN | -1.303296 | 0.04338759 | -0.124364 | 0.6030937 | 0.863184 | 0.1344917 | 1.089098 | 0.01411413 | 1.046298 | 0.5805324 | -0.033451 | 0.6885065 |
| ENSG00000011007.12 | TCEB3 | -0.274361 | 0.8006286 | -0.023222 | 0.7156572 | 0.224869 | 0.003472498 | 1.064626 | 0.4919368 | 0.127937 | 1 | 1.197919 | 0.88512 |
| ENSG00000011021.21 | CLCN6 | -2.459488 | 0.6214382 | -0.102337 | 0.9791082 | -0.396673 | 0.7042635 | 2.842563 | 0.0202141 | 0.611157 | 0.3813186 | 2.472409 | 0.9737038 |
| ENSG00000011258.15 | MBTD1 | -0.716067 | 0.8006286 | -0.766298 | 0.5953732 | 1.444686 | 0.2343604 | 2.688296 | 0.01449893 | 0.760151 | 1 | 3.733611 | 0.6885065 |
| ENSG00000011275.18 | RNF216 | 0.846639 | 0.6214382 | 0.373395 | 0.6527736 | -0.276624 | 0.2101108 | 0.963161 | 0.03598227 | 1.362899 | 0.107751 | -2.722157 | 0.8479431 |
| ENSG00000011295.15 | TTC19 | 0.003485 | 0.6214382 | -0.266768 | 0.5953732 | 0.063806 | 0.9207295 | 1.176527 | 4.85E-06 | 1.384692 | 0.002293693 | 0.340152 | 0.2809502 |
| ENSG00000011304.17 | PTBP1 | -0.277379 | 0.992766 | -0.286536 | 0.4539559 | -0.054863 | 0.09455264 | 1.20234 | 1.04E-05 | 0.844291 | 0.5693759 | 0.353134 | 0.9147227 |
| ENSG00000011332.19 | DPF1 | 0.190072 | 0.9438446 | -0.43539 | 0.8263112 | -0.236113 | 0.4382561 | 2.336893 | 6.75E-07 | 1.458503 | 0.001495575 | -0.558794 | 0.5344421 |
| ENSG00000011451.17 | WIZ | -0.216799 | 0.9360557 | -0.241969 | 0.5010139 | 0.599646 | 0.9429584 | 1.411781 | 8.97E-06 | 0.783759 | 0.09116007 | -1.019282 | 0.7111448 |
| ENSG00000011454.16 | RABGAP1 | -0.41559 | 1 | 0.270339 | 0.9973179 | 0.665646 | 0.6600958 | 1.319552 | 0.01831499 | 0.482133 | 0.487611 | -1.87031 | 0.9568643 |
| ENSG00000011485.14 | PPP5C | -0.945687 | 0.6214382 | 0.070632 | 0.6527736 | -1.049849 | 0.02213292 | 1.800155 | 0.000793068 | 0.356443 | 0.7377262 | 0.066226 | 0.5733051 |
| ENSG00000012660.13 | ELOVL5 | 0.156821 | 0.8419179 | 0.004653 | 0.9727008 | 0.829462 | 0.02424566 | 0.428647 | 0.4479674 | -0.344798 | 0.8021966 | 0.840022 | 0.6885065 |
| ENSG00000013375.15 | PGM3 | -0.019018 | 0.9789948 | 0.269628 | 0.8842289 | 1.202833 | 1.02E-06 | 1.152215 | 0.7677345 | 0.769628 | 0.7494681 | -0.859214 | 0.8945638 |
| ENSG00000013441.15 | CLK1 | -1.111139 | 0.9506557 | 0.047706 | 0.9564082 | 0.169363 | 0.02280626 | 0.424918 | 0.03367078 | 0.722919 | 0.3831662 | 0.76101 | 0.9716102 |
| ENSG00000013561.17 | RNF14 | -0.319121 | 0.8426644 | -0.006482 | 0.7881421 | -0.374736 | 0.3276184 | 0.565186 | 0.02493694 | 1.068027 | 0.3212277 | -0.459713 | 0.8167193 |
| ENSG00000013573.16 | DDX11 | 1.210022 | 0.7736255 | -0.281943 | 0.8116276 | 0.177282 | 0.691147 | 0.680738 | 0.912238 | 2.638347 | 0.03658568 | 2.263686 | 0.8185572 |
| ENSG00000013583.8 | HEBP1 | -1.040481 | 1 | -0.472937 | 0.6822393 | 0.529918 | 0.896092 | -1.392974 | 0.000431472 | 2.048366 | 0.3974318 | -1.040135 | 0.8479431 |
| ENSG00000014123.9 | UFL1 | -1.056946 | 0.8006286 | 0.384852 | 1 | 0.63463 | 0.002841623 | 0.97008 | 0.8414154 | -0.238765 | 0.4951016 | 0.330387 | 0.8479431 |
| ENSG00000014641.17 | MDH1 | 0.619547 | 0.04338759 | 0.034637 | 0.9628891 | -0.424083 | 0.1663608 | -0.194657 | 0.001762572 | 0.583151 | 0.9629537 | 0.85326 | 1 |
| ENSG00000014824.13 | SLC30A9 | 0.101828 | 0.6616751 | -0.716465 | 0.3604204 | 0.431491 | 0.5290845 | 0.909859 | 0.03608197 | 0.840778 | 0.9642859 | 0.609132 | 0.6517664 |
| ENSG00000015475.18 | BID | 0.142232 | 0.8530607 | -0.270545 | 0.6527736 | -0.521942 | 0.000530924 | 0.952304 | 0.4399569 | 0.363548 | 0.8933383 | -0.375157 | 0.5620837 |
| ENSG00000015592.16 | STMN4 | -0.1727 | 0.9047689 | 0.084671 | 0.9420234 | 0.498083 | 2.37E-05 | 0.946015 | 1 | 0.356497 | 0.2244526 | -0.162042 | 0.8479431 |
| ENSG00000015676.17 | NUDCD3 | 0.038387 | 0.6214382 | 0.221958 | 0.8943231 | 0.265379 | 0.7437308 | 1.246731 | 0.000834282 | 0.400766 | 0.2465981 | 1.726146 | 0.950367 |
| ENSG00000017260.19 | ATP2C1 | -0.600372 | 1 | -0.15199 | 0.6527736 | 0.587669 | 0.2963013 | 1.432171 | 0.008003341 | 0.961029 | 0.260732 | 0.661059 | 0.9565287 |
| ENSG00000018189.12 | RUFY3 | 0.069879 | 0.88662 | -0.84235 | 0.00370122 | 0.391928 | 0.001234163 | 1.374338 | 0.01718177 | 0.04799 | 0.8978057 | -0.681097 | 0.4476145 |
| ENSG00000018236.14 | CNTN1 | 0.17037 | 0.8427546 | -0.477304 | 0.4947142 | -1.514692 | 1 | 0.590871 | 0.02237281 | 1.246315 | 0.0507771 | -1.948439 | 0.6885065 |
| ENSG00000018510.12 | AGPS | -1.035146 | 0.8587992 | 0.054776 | 0.7573724 | 0.430527 | 0.8308365 | 0.095539 | 0.8371191 | 0.034146 | 0.03539976 | 0.221648 | 0.9147227 |
| ENSG00000021645.18 | NRXN3 | -0.527066 | 0.714717 | 3.801505 | 0.8263112 | 2.208043 | 0.6295125 | 2.088455 | 0.009902535 | -0.275603 | 0.5520189 | -0.251623 | 1 |
| ENSG00000022267.16 | FHL1 | -0.21368 | 0.529676 | -0.149015 | 0.5283548 | 0.113453 | 0.4064599 | -0.563576 | 0.0136954 | -0.14912 | 0.7377262 | -0.05536 | 0.8829971 |
| ENSG00000023191.16 | RNH1 | -0.786837 | 0.9541095 | 0.133038 | 0.9860872 | -0.017137 | 0.3966431 | 0.358381 | 0.2129262 | 0.047197 | 0.0198219 | -0.670617 | 0.7695425 |
| ENSG00000023287.12 | RB1CC1 | -1.534016 | 0.4406957 | 0.213242 | 0.6527736 | 0.767608 | 2.64E-13 | 1.097576 | 0.006155218 | 0.418529 | 0.3759745 | -0.917236 | 0.6267144 |
| ENSG00000023572.8 | GLRX2 | -0.766054 | 0.7138283 | 0.026246 | 0.9194312 | -0.473321 | 0.1650103 | -0.358465 | 0.02835414 | 0.977036 | 0.6486434 | 0.440948 | 0.9919109 |
| ENSG00000023902.13 | PLEKHO1 | -0.537979 | 0.8006286 | 0.498064 | 0.7792763 | -0.111285 | 0.8599056 | 1.432365 | 0.02317381 | 0.274686 | 0.08151074 | 0.235368 | 1 |
| ENSG00000023909.9 | GCLM | -0.573265 | 0.6214382 | -0.080896 | 0.6527736 | -0.055742 | 0.1895736 | 2.225568 | 0.009475785 | 1.510837 | 0.3318854 | 0.423685 | 0.885753 |
| ENSG00000024048.10 | UBR2 | -2.484552 | 0.6214382 | -0.528473 | 0.9696336 | 0.217215 | 0.5290845 | 1.337666 | 0.009399882 | 2.009316 | 0.178378 | 1.718498 | 0.6576742 |
| ENSG00000025293.15 | PHF20 | -2.160493 | 0.7887824 | -0.41176 | 0.7952388 | 0.261821 | 0.1992941 | 0.739586 | 0.000139095 | 0.017684 | 0.6704453 | 0.641674 | 0.7750686 |
| ENSG00000025770.18 | NCAPH2 | -0.905926 | 0.8814146 | -0.28762 | 1 | -1.011733 | 0.03639066 | 0.270132 | 0.4919368 | 0.802568 | 0.4155911 | -0.472339 | 1 |
| ENSG00000025796.13 | SEC63 | 0.429873 | 0.4406884 | 0.13496 | 0.9104976 | 0.565112 | 0.000962523 | 1.060227 | 0.04008401 | -0.071121 | 0.2833571 | 0.516382 | 0.8383091 |
| ENSG00000026025.13 | VIM | -0.657227 | 0.2043253 | -0.207255 | 0.002276985 | -0.034999 | 0.1357519 | -0.071006 | 0.07237193 | -0.212085 | 7.78E-05 | 0.159486 | 0.9370627 |
| ENSG00000026297.15 | RNASET2 | -1.584683 | 0.8006286 | 0.532916 | 0.04760656 | -1.128888 | 0.05838784 | 0.05826 | 0.3527037 | 2.213232 | 0.3974318 | 1.794686 | 0.3746505 |
| ENSG00000026508.16 | CD44 | -0.371064 | 0.2423804 | 0.270444 | 0.005272355 | -0.769669 | 0.000371391 | 0.355615 | 0.3433335 | -0.097087 | 0.2228691 | 0.317327 | 0.9200004 |
| ENSG00000026652.13 | AGPAT4 | -0.027943 | 1 | -0.446398 | 0.6527736 | 0.316738 | 0.4383213 | 0.922485 | 0.0007863 | 1.308619 | 0.1329005 | 1.341819 | 0.9622177 |
| ENSG00000027847.13 | B4GALT7 | -1.87344 | 0.6014343 | 0.021707 | 0.8703522 | -1.057622 | 0.04639883 | -0.58027 | 0.2211193 | -0.464152 | 0.6720693 | -0.448787 | 0.9228374 |
| ENSG00000029153.14 | ARNTL2 | 2.658431 | 0.8006286 | -0.529538 | 0.06755823 | 0.056671 | 0.5039831 | -0.163726 | 0.01992068 | -0.49715 | 0.8983233 | -0.53329 | 0.8479431 |
| ENSG00000029363.15 | BCLAF1 | -0.479842 | 0.6214382 | 0.565255 | 0.5701602 | 1.207203 | 3.54E-10 | -0.083793 | 0.1652563 | 0.030972 | 0.7221921 | -0.487514 | 0.6885065 |
| ENSG00000029364.11 | SLC39A9 | -0.88461 | 0.9506557 | 0.470395 | 0.6706163 | -0.036595 | 1 | 1.998809 | 0.000337501 | 1.068772 | 0.3522585 | -0.693998 | 0.9532204 |
| ENSG00000030066.13 | NUP160 | 0.741083 | 1 | 0.026195 | 0.9583742 | -0.145205 | 0.7354314 | 0.531402 | 0.2970007 | 2.294791 | 0.04535071 | -1.039533 | 0.6516876 |
| ENSG00000030582.16 | GRN | 0.337697 | 0.714717 | 0.463222 | 0.6527736 | -1.070051 | 0.04448448 | 0.47011 | 0.8997082 | 1.504745 | 0.3974318 | 1.276654 | 0.6885065 |
| ENSG00000031698.12 | SARS | -0.271436 | 0.6014343 | 0.107434 | 0.9577957 | 0.505748 | 0.001163142 | 0.164179 | 0.3928563 | 0.077702 | 0.001282234 | 0.582349 | 0.6885065 |
| ENSG00000032219.18 | ARID4A | -0.09689 | 0.8006286 | 0.136934 | 1 | 0.443872 | 0.1400181 | 1.173239 | 0.000427487 | -0.48997 | 0.479149 | -0.81196 | 0.8479431 |
| ENSG00000032389.12 | TSSC1 | -0.227712 | 1 | -0.567798 | 0.5010139 | -0.451167 | 0.5017549 | -0.260724 | 0.01221979 | 0.037679 | 0.6908753 | 1.598071 | 0.7983408 |
| ENSG00000032444.15 | PNPLA6 | -0.312399 | 0.6859425 | 0.18058 | 0.770491 | 1.351199 | 1 | 0.081005 | 0.0201847 | 0.521082 | 0.5196843 | 1.27003 | 0.6885065 |
| ENSG00000033050.7 | ABCF2 | -0.031134 | 0.9178164 | -0.182627 | 0.9234066 | -0.467796 | 0.5290845 | 1.640923 | 0.01771888 | 0.44945 | 0.8768085 | -0.07684 | 0.7828689 |
| ENSG00000033100.14 | CHPF2 | 0.796658 | 0.6214382 | 0.760636 | 0.6514723 | -0.677758 | 0.01326601 | 1.241538 | 0.3760083 | 1.087951 | 0.5196843 | 0.786412 | 1 |
| ENSG00000033178.12 | UBA6 | -0.563043 | 0.6214382 | -0.285942 | 0.7062745 | 0.930548 | 0.02607316 | 1.34421 | 6.78E-05 | 1.15615 | 0.4535704 | -0.958743 | 0.6855855 |
| ENSG00000033327.12 | GAB2 | 4.199183 | 0.3946735 | -0.606609 | 0.535881 | 0.728156 | 0.6933858 | 1.234234 | 0.04291 | 1.267982 | 0.3232893 | -1.00467 | 0.8949489 |
| ENSG00000033627.14 | ATP6V0A1 | -0.270422 | 0.8867323 | 0.162278 | 0.9525045 | -0.419711 | 0.9076129 | 1.517172 | 0.000315064 | 1.052007 | 0.02163664 | -0.426404 | 0.6885065 |
| ENSG00000033867.16 | SLC4A7 | -0.669728 | 0.8587992 | -0.050341 | 0.6331189 | 0.791694 | 0.5087073 | 1.237318 | 0.01996536 | 1.451331 | 0.2312487 | 1.213991 | 0.6185816 |
| ENSG00000034053.14 | APBA2 | 1.097398 | 0.6214382 | 0.047248 | 0.7993746 | -0.486601 | 0.4319188 | 0.839885 | 0.003270086 | 1.721613 | 0.07401196 | -1.703152 | 0.8688812 |
| ENSG00000034152.18 | MAP2K3 | -0.548637 | 0.714717 | -0.156947 | 0.7368222 | -0.819658 | 0.08400047 | 1.828199 | 0.01052272 | 1.374255 | 0.3129511 | 0.054874 | 0.8856541 |
| ENSG00000034510.5 | TMSB10 | 0.303499 | 0.2261267 | 0.062156 | 0.6163022 | -0.221365 | 6.55E-07 | -0.011014 | 0.8818876 | -0.404749 | 2.02E-08 | 0.220757 | 0.6829816 |
| ENSG00000034713.7 | GABARAPL2 | -0.061931 | 0.3582875 | 0.271534 | 0.7834199 | -0.201894 | 0.7437308 | 0.241615 | 0.2918218 | 0.04993 | 0.007820647 | 0.105152 | 0.3469679 |
| ENSG00000035115.21 | SH3YL1 | 0.443282 | 0.8006286 | 0.016688 | 1 | 0.199798 | 0.3622474 | 1.479147 | 0.01645598 | 0.712622 | 0.1358132 | -0.470292 | 0.9029338 |
| ENSG00000035928.14 | RFC1 | -0.56552 | 0.6214382 | -0.014363 | 0.6527736 | 0.688073 | 0.001437818 | 0.539811 | 0.7967812 | 0.186934 | 0.7286448 | 0.094254 | 0.8479431 |
| ENSG00000036054.12 | TBC1D23 | -1.287642 | 0.4782461 | 0.074365 | 0.6527736 | 0.949305 | 0.002723788 | 0.206337 | 0.9527553 | 1.166444 | 0.5196843 | 0.938742 | 0.7862316 |
| ENSG00000036257.12 | CUL3 | 0.591511 | 0.6214382 | -0.068734 | 1 | 1.220568 | 0.0228706 | 1.454905 | 7.72E-05 | 1.60541 | 0.06340332 | 1.121793 | 0.6885065 |
| ENSG00000036549.12 | ZZZ3 | 0.319808 | 1 | 0.327658 | 0.9031929 | 0.56526 | 0.3767722 | 0.822981 | 0.3500508 | 2.464586 | 0.000168448 | -1.849593 | 0.7096009 |
| ENSG00000037474.14 | NSUN2 | -1.101154 | 0.3439898 | 0.241791 | 0.9612341 | 0.46448 | 0.5044352 | 2.617083 | 2.54E-05 | 0.86907 | 0.5360904 | -0.255571 | 0.8380376 |
| ENSG00000037637.10 | FBXO42 | 0.098169 | 0.714717 | 0.127469 | 0.8946072 | -0.259093 | 0.7558057 | 2.956169 | 0.0018434 | 2.707272 | 0.01398734 | 1.091976 | 0.9552941 |
| ENSG00000038002.8 | AGA | -0.077948 | 1 | 0.129772 | 0.7594702 | -1.1204 | 0.1611081 | 0.864482 | 0.02141464 | 1.536898 | 1 | 1.977619 | 0.8260537 |
| ENSG00000038210.12 | PI4K2B | -0.058796 | 0.714717 | -0.86157 | 0.592336 | 1.452916 | 0.7233932 | 2.347625 | 0.02639972 | 1.436093 | 0.5727095 | 2.950074 | 0.868568 |
| ENSG00000038219.12 | BOD1L1 | -0.066967 | 0.9703009 | 0.087246 | 0.8953649 | 0.824534 | 0.02390989 | 1.102593 | 7.13E-06 | 1.278347 | 0.000368774 | -0.972282 | 0.6885065 |
| ENSG00000038382.17 | TRIO | -0.542831 | 0.2585731 | 0.076678 | 0.9452791 | 0.299329 | 0.1610961 | 0.693043 | 3.89E-07 | 0.582594 | 0.1065267 | -1.885274 | 0.296307 |
| ENSG00000038427.15 | VCAN | 0.200087 | 0.529676 | 0.105088 | 0.7368405 | 1.686578 | 6.72E-19 | 1.586138 | 0.000782846 | 0.983365 | 0.1196747 | -0.032433 | 0.855268 |
| ENSG00000038532.14 | CLEC16A | 3.053286 | 0.4764974 | 1.088847 | 0.6670505 | 0.892576 | 0.4761297 | 1.339695 | 0.09332577 | 2.799833 | 0.0328027 | -0.321364 | 0.6885065 |
| ENSG00000039123.15 | SKIV2L2 | 0.369938 | 0.7952299 | -0.003355 | 0.6527736 | 0.604658 | 0.07986212 | 1.465607 | 0.000860514 | 0.716873 | 0.6915201 | -1.41374 | 0.6345741 |
| ENSG00000039523.17 | FAM65A | -0.035163 | 0.7780681 | 0.181471 | 0.9581211 | 1.100372 | 0.7488422 | 0.988528 | 0.03519259 | 1.558286 | 0.8345107 | 2.420283 | 0.8066053 |
| ENSG00000039560.13 | RAI14 | 1.700723 | 0.2032687 | -0.180693 | 0.9084242 | -0.303332 | 0.5290845 | 2.209835 | 0.02260367 | -0.333707 | 0.516599 | -0.364331 | 0.6829816 |
| ENSG00000040199.18 | PHLPP2 | -3.061813 | 1 | 1.065071 | 0.9590571 | 0.246751 | 0.4382561 | 0.85706 | 0.04364143 | 2.607802 | 0.08372708 | -1.234809 | 0.6928022 |
| ENSG00000040633.12 | PHF23 | 0.902364 | 0.6214382 | 0.027637 | 0.9653874 | 0.147237 | 0.942307 | 1.451594 | 0.000297795 | 0.072366 | 0.6813171 | -0.40493 | 0.9749564 |
| ENSG00000041357.15 | PSMA4 | -3.182548 | 0.2708189 | -0.193336 | 0.08175978 | 0.203246 | 0.538518 | 0.630139 | 0.2131424 | -0.481642 | 0.002111082 | -0.661483 | 0.6185816 |
| ENSG00000041802.10 | LSG1 | -1.27603 | 0.2373868 | 0.035192 | 0.5953732 | 0.238001 | 0.7283482 | 0.773484 | 0.03895965 | 1.296623 | 0.1137773 | -0.014099 | 0.6829816 |
| ENSG00000041982.15 | TNC | -0.942121 | 1 | -0.043479 | 0.69409 | 0.003723 | 0.3171226 | 1.292657 | 4.30E-06 | -0.514403 | 0.0117725 | 0.21955 | 0.9391913 |
| ENSG00000042753.11 | AP2S1 | -0.080974 | 0.9921711 | -0.155226 | 0.6774447 | -0.341013 | 0.000739632 | 0.070584 | 0.3059811 | 0.190076 | 0.2423939 | -0.48737 | 0.5066477 |
| ENSG00000043093.13 | DCUN1D1 | 0.463461 | 0.531287 | 0.211877 | 0.9924971 | 1.388841 | 0.006241239 | 2.585797 | 0.04764406 | 0.837674 | 1 | 0.431618 | 0.8949489 |
| ENSG00000044574.7 | HSPA5 | 0.793441 | 0.000196944 | 0.284492 | 7.88E-07 | 0.467827 | 6.60E-11 | 0.642201 | 0.2784124 | 0.124291 | 0.8624812 | 0.209301 | 0.7420459 |
| ENSG00000046651.14 | OFD1 | 0.283485 | 0.9539331 | 0.290009 | 0.8732309 | -0.338631 | 0.5033398 | 2.548076 | 0.006007256 | 0.043256 | 0.9669702 | 1.775259 | 0.6885065 |
| ENSG00000046653.14 | GPM6B | -0.569676 | 0.529676 | -0.191782 | 0.1120376 | 0.726921 | 8.99E-07 | 0.574206 | 0.4569212 | 0.060534 | 0.5196843 | -0.279507 | 0.7417751 |
| ENSG00000046889.18 | PREX2 | -1.309065 | 0.9183025 | -1.341233 | 0.02922882 | -0.848043 | 0.9343369 | 2.510001 | 0.006391833 | -3.144502 | 0.2357139 | 1.137032 | 0.826754 |
| ENSG00000047315.14 | POLR2B | -1.287022 | 0.00172503 | -0.052914 | 0.6527736 | 0.188715 | 0.03144921 | 0.80017 | 0.4134284 | 0.56097 | 0.2680669 | 1.355009 | 0.8853287 |
| ENSG00000047410.13 | TPR | -0.170105 | 0.6561323 | 0.0528 | 0.7374794 | 0.770042 | 1.70E-05 | 0.901744 | 0.4224573 | -0.292709 | 0.2532119 | 1.984974 | 0.9641669 |
| ENSG00000047662.4 | FAM184B | -0.061613 | 1 | -0.727725 | 0.6908049 | -0.922283 | 0.4129049 | 1.260202 | 0.02353059 | 1.584147 | 0.8595963 | -4.508575 | 0.5066477 |
| ENSG00000047849.21 | MAP4 | -0.441063 | 0.6214382 | 0.095492 | 0.9102749 | 0.779426 | 0.000441979 | 0.507474 | 0.4919368 | 1.093879 | 0.3810416 | -0.233118 | 0.8469357 |
| ENSG00000048028.11 | USP28 | 1.66451 | 0.6214382 | 0.075586 | 0.8513348 | 0.53524 | 0.8832504 | 2.012816 | 0.07274418 | 2.517246 | 0.0130078 | 0.575227 | 1 |
| ENSG00000048052.21 | HDAC9 | 3.244405 | 1 | 0.648595 | 0.7594721 | 0.104669 | 0.5701849 | 0.698538 | 0.03196063 | 0.653778 | 0.6487553 | -3.001333 | 0.826754 |
| ENSG00000048392.11 | RRM2B | 0.171831 | 1 | -0.161555 | 0.949856 | 0.437793 | 0.2180501 | 0.729055 | 0.01252251 | 0.995478 | 0.9418685 | -0.786051 | 0.9370888 |
| ENSG00000048649.13 | RSF1 | -0.595213 | 0.6792949 | -0.238952 | 0.4571222 | 1.584478 | 3.00E-20 | 1.572563 | 0.1619597 | 0.249359 | 0.7377262 | 1.397625 | 0.9205035 |
| ENSG00000048707.13 | VPS13D | 1.344339 | 1 | 0.544805 | 0.8917881 | 0.654824 | 0.9931698 | 1.39084 | 0.02535905 | 2.526597 | 0.1389793 | -1.103524 | 0.9062221 |
| ENSG00000048740.17 | CELF2 | 0.753413 | 0.8006286 | -0.547205 | 0.5159728 | 0.448559 | 0.01177253 | 0.370612 | 0.172332 | 0.5656 | 0.07142479 | -0.446555 | 0.7767294 |
| ENSG00000048828.16 | FAM120A | 0.827892 | 1 | -0.360614 | 0.5953732 | 1.014422 | 0.3271819 | 0.990864 | 2.84E-06 | 0.532164 | 0.7620165 | 0.032058 | 0.9589318 |
| ENSG00000048991.16 | R3HDM1 | 0.238465 | 0.8694227 | -0.907031 | 0.3121527 | 1.543904 | 0.005556955 | 0.99609 | 0.09883746 | -0.927869 | 0.8021966 | -0.844312 | 0.6885065 |
| ENSG00000049618.21 | ARID1B | -2.601335 | 0.937674 | -0.054714 | 0.7192354 | 1.194228 | 0.4064599 | 1.783351 | 0.001825432 | 1.328303 | 0.1515597 | 0.540839 | 0.9245437 |
| ENSG00000049656.13 | CLPTM1L | -1.059813 | 0.02746914 | 0.008686 | 0.9733961 | 0.201819 | 0.8774944 | 0.419686 | 0.2570203 | 0.29342 | 0.1704211 | 0.00878 | 0.6885065 |
| ENSG00000049759.16 | NEDD4L | 1.33682 | 0.8006286 | 0.159326 | 0.9452791 | 1.563014 | 0.7457949 | 2.463444 | 0.000334033 | 1.175528 | 0.2910219 | 2.23355 | 0.6829816 |
| ENSG00000049860.13 | HEXB | 0.459964 | 0.6214382 | 0.257953 | 0.4114546 | -0.156832 | 0.03960691 | -0.042062 | 0.8431309 | -0.494072 | 0.894525 | -0.019853 | 0.6885065 |
| ENSG00000050030.13 | KIAA2022 | 1.036081 | 0.6168416 | 1.032149 | 0.8528295 | 0.394908 | 0.01966866 | -0.695243 | 0.4919368 | 0.597663 | 0.4987279 | -3.402536 | 0.6885065 |
| ENSG00000050393.11 | MCUR1 | 0.719657 | 0.8222752 | -0.10685 | 0.7383998 | -1.237587 | 0.022069 | 0.096745 | 0.2998372 | 0.499015 | 0.7356185 | 0.608607 | 1 |
| ENSG00000050405.13 | LIMA1 | -0.342004 | 0.9678883 | -0.101001 | 0.3386108 | 0.465218 | 0.007296446 | 0.121239 | 0.9988044 | 0.07574 | 0.2653803 | 1.394136 | 1 |
| ENSG00000050820.16 | BCAR1 | -0.598126 | 0.6844928 | 0.133414 | 1 | -0.178023 | 0.645156 | 0.600064 | 0.009657092 | 0.586909 | 0.2534219 | 0.478293 | 0.9733288 |
| ENSG00000051108.14 | HERPUD1 | -2.128636 | 0.8006286 | -0.094341 | 0.5953732 | 0.387487 | 0.004579145 | 0.040815 | 0.9439697 | 0.569544 | 0.9812081 | 0.209356 | 0.8464135 |
| ENSG00000051128.18 | HOMER3 | -3.050003 | 0.08953158 | 0.504499 | 0.5967639 | -1.155599 | 0.000498639 | 0.722796 | 0.7551498 | -0.105456 | 0.5689995 | 0.196915 | 0.8716748 |
| ENSG00000052795.12 | FNIP2 | -0.866833 | 0.6792949 | -0.174857 | 0.7317746 | 1.778083 | 0.1669069 | 1.809236 | 0.02235605 | 1.022546 | 0.4883149 | 0.368683 | 0.9147227 |
| ENSG00000052802.12 | MSMO1 | 0.963393 | 0.000924656 | 0.162885 | 0.8532785 | 0.504893 | 0.0745984 | 0.193571 | 0.8697916 | 1.244512 | 0.5196843 | 0.356333 | 0.9646502 |
| ENSG00000052841.14 | TTC17 | 1.899647 | 0.07547825 | 0.299537 | 0.9222921 | 0.30125 | 0.004502289 | -0.280529 | 0.4008968 | 0.990091 | 0.3244978 | 1.542553 | 1 |
| ENSG00000053254.15 | FOXN3 | -2.197249 | 0.6214382 | 0.672026 | 0.9117805 | 1.121351 | 0.6521146 | 2.077338 | 0.003040752 | 0.085832 | 0.6845994 | 1.348585 | 0.6885065 |
| ENSG00000053371.12 | AKR7A2 | -1.646558 | 0.6792949 | -0.14078 | 0.5953732 | -2.491615 | 6.92E-06 | 0.450488 | 0.479858 | 2.611337 | 0.1434024 | -0.150225 | 0.8964771 |
| ENSG00000053900.10 | ANAPC4 | -1.684574 | 0.8006286 | -0.288149 | 0.8442222 | -0.24216 | 0.4383712 | 0.778731 | 0.03405717 | 1.132548 | 0.5196843 | -1.46962 | 1 |
| ENSG00000054116.11 | TRAPPC3 | 0.170389 | 0.6214382 | -0.234505 | 0.2097393 | 0.040129 | 0.8837341 | 0.731291 | 0.06156517 | -1.567678 | 0.01710408 | -0.441195 | 0.6885065 |
| ENSG00000054118.13 | THRAP3 | 0.543067 | 0.2335514 | 0.285323 | 0.8703522 | 1.158138 | 0.001019573 | 0.196238 | 0.6614105 | 0.260313 | 0.9022706 | 0.292905 | 0.7291058 |
| ENSG00000054148.17 | PHPT1 | 0.323865 | 0.2617323 | -0.249963 | 0.02488903 | -0.378258 | 4.95E-09 | -0.267496 | 0.000132784 | -0.301581 | 0.01129028 | 0.67331 | 0.9576568 |
| ENSG00000054267.20 | ARID4B | -0.285418 | 0.3946735 | 0.333233 | 0.6527736 | 0.640516 | 6.74E-06 | 0.934801 | 0.04504576 | 0.006868 | 0.7499404 | 0.066847 | 0.6623168 |
| ENSG00000054523.17 | KIF1B | 0.455207 | 0.8006286 | -0.228049 | 0.1004476 | 0.40497 | 0.3645882 | 1.219043 | 5.92E-09 | 1.637437 | 0.001515333 | -0.503441 | 0.8917781 |
| ENSG00000054654.15 | SYNE2 | 0.603814 | 0.8587992 | -0.517896 | 0.4905237 | 0.874379 | 0.02106069 | 1.076225 | 0.05691687 | 1.195196 | 0.1401461 | 1.791752 | 0.6185816 |
| ENSG00000054690.13 | PLEKHH1 | -1.826776 | 0.8587992 | 0.999556 | 0.4487482 | 1.336392 | 0.5290845 | 3.34558 | 0.01853051 | 2.752683 | 0.2363864 | -2.965723 | 0.6885065 |
| ENSG00000054965.10 | FAM168A | -0.450048 | 0.824666 | -0.359444 | 0.7436352 | -0.516776 | 0.1690356 | 0.581491 | 0.03861516 | 1.925299 | 0.006054633 | -0.045632 | 1 |
| ENSG00000055044.10 | NOP58 | 0.382995 | 0.6868924 | 0.058131 | 0.5953732 | 0.401724 | 0.001088591 | 0.342961 | 0.3467508 | 0.624194 | 0.8577401 | -0.555956 | 0.6044994 |
| ENSG00000055118.14 | KCNH2 | -0.109917 | 0.8587992 | 1.235873 | 1 | 0.373453 | 0.9287147 | 0.815438 | 0.5930297 | 1.580161 | 0.00086076 | -0.802855 | 0.6767422 |
| ENSG00000055163.18 | CYFIP2 | 0.92337 | 0.2506697 | -0.122019 | 0.5066068 | -0.389814 | 0.7236174 | 1.14555 | 0.0321729 | 0.632203 | 0.0171907 | 2.031771 | 0.7324591 |
| ENSG00000055208.17 | TAB2 | -0.687785 | 0.9047689 | -0.133963 | 0.4892267 | 0.87459 | 0.9316604 | 0.986262 | 0.09192941 | 1.863675 | 0.03706797 | -1.061379 | 0.6885065 |
| ENSG00000055332.16 | EIF2AK2 | -1.605414 | 0.01638188 | 0.193159 | 0.9859883 | 0.827166 | 0.004221161 | 0.565641 | 0.1577114 | 0.78603 | 0.8077499 | 0.417763 | 0.9428375 |
| ENSG00000055483.19 | USP36 | 0.285732 | 0.8861738 | -0.272332 | 0.9697133 | 0.034868 | 0.6049589 | 1.365015 | 0.02191916 | 0.677919 | 0.6382061 | 1.462527 | 0.6885065 |
| ENSG00000055609.17 | KMT2C | -1.090606 | 0.2643306 | 0.128325 | 0.5953732 | 0.778856 | 2.29E-06 | 0.910657 | 3.44E-07 | 1.204959 | 0.03622093 | -1.363549 | 0.6829816 |
| ENSG00000055917.15 | PUM2 | 1.393353 | 0.8867323 | -0.453006 | 0.492893 | 0.637445 | 0.2385992 | 1.556231 | 0.001533611 | 1.033052 | 0.003586819 | -0.118604 | 0.9264343 |
| ENSG00000056050.6 | C4orf27 | -1.653189 | 0.3982827 | 0.167959 | 0.8703522 | 0.057837 | 0.6064256 | -0.309544 | 0.04454587 | 0.176139 | 0.03056663 | 0.959363 | 0.7361208 |
| ENSG00000056097.15 | ZFR | -0.259768 | 0.8006286 | -0.233122 | 0.0597279 | 0.798042 | 0.000763356 | 0.771863 | 0.1031205 | 0.560497 | 0.937993 | -0.062208 | 0.863256 |
| ENSG00000056487.15 | PHF21B | 0.370741 | 0.8260297 | -0.265949 | 0.5953732 | -0.230703 | 0.9995062 | 1.546213 | 0.004777529 | -0.235863 | 0.6391587 | -0.126622 | 0.9532204 |
| ENSG00000056586.15 | RC3H2 | -0.724756 | 0.6493358 | -0.218962 | 0.9463739 | 0.712312 | 0.5290845 | 0.267397 | 0.000316394 | -0.077132 | 0.6913898 | 0.898859 | 0.9955898 |
| ENSG00000057019.15 | DCBLD2 | 0.183508 | 0.4764974 | -0.243512 | 0.6745155 | 0.324756 | 0.8305038 | 1.169415 | 2.75E-05 | 0.269681 | 0.9361679 | -0.405732 | 0.04261865 |
| ENSG00000057757.9 | PITHD1 | -0.41698 | 0.9004669 | 0.248891 | 0.8697084 | -0.008861 | 0.02196322 | 0.780539 | 0.62766 | 0.039143 | 0.007083031 | 0.484566 | 0.9351935 |
| ENSG00000058063.15 | ATP11B | -0.817324 | 0.7736255 | -0.093338 | 0.85618 | 1.536283 | 0.05283157 | 3.401986 | 0.000324536 | 1.308648 | 0.1565243 | 0.066572 | 0.9617345 |
| ENSG00000058091.16 | CDK14 | 0.51371 | 0.9183025 | -0.438835 | 0.4409757 | 0.555181 | 0.1808701 | 1.593036 | 0.02486581 | 1.890351 | 0.007905278 | -0.185804 | 0.793288 |
| ENSG00000058262.9 | SEC61A1 | -0.242947 | 0.3773249 | 0.138389 | 0.7107715 | -0.079304 | 0.8454005 | 0.877598 | 0.000543727 | 0.55948 | 0.7800472 | 0.215101 | 0.6885065 |
| ENSG00000058272.16 | PPP1R12A | -0.624069 | 0.8880907 | 0.013916 | 0.4539559 | 1.101494 | 7.92E-05 | 1.362805 | 0.1705746 | 0.433535 | 0.925632 | -0.099344 | 0.9657663 |
| ENSG00000058668.14 | ATP2B4 | -0.226225 | 0.9798794 | -0.094871 | 0.4207312 | 1.14181 | 0.4060104 | 0.218179 | 0.003953901 | 1.175694 | 0.08186725 | 0.715839 | 0.8949489 |
| ENSG00000058673.15 | ZC3H11A | 0.126212 | 0.6644434 | -0.200444 | 0.8106695 | 0.76921 | 0.000722601 | 0.533791 | 0.1589128 | 0.541972 | 0.5196843 | 1.322565 | 0.960662 |
| ENSG00000058866.14 | DGKG | 0.15999 | 0.9671207 | 0.713943 | 0.9575958 | 1.299868 | 0.3904243 | 1.09799 | 0.01826913 | 1.503319 | 0.7381046 | 1.6201 | 0.7595705 |
| ENSG00000059122.16 | FLYWCH1 | -0.835225 | 0.9288178 | 0.918548 | 0.8874017 | -0.844479 | 0.332994 | 0.305573 | 0.02914838 | 1.624408 | 0.07328082 | -1.264688 | 0.6885065 |
| ENSG00000059758.7 | CDK17 | -0.012447 | 0.5873352 | -0.368558 | 0.6527736 | 1.282941 | 0.3871766 | 0.63761 | 0.01378953 | 1.466742 | 0.2309279 | 0.181761 | 0.6185816 |
| ENSG00000060237.16 | WNK1 | 0.024764 | 0.7533508 | -0.199552 | 0.4011918 | 0.414643 | 0.02213292 | 1.649777 | 0.002666913 | 0.696389 | 0.03857745 | 0.074474 | 0.5344421 |
| ENSG00000060688.12 | SNRNP40 | 0.592543 | 0.8860308 | -0.737255 | 0.2713328 | -0.065509 | 0.8389806 | -0.33268 | 0.4926988 | -0.542821 | 0.02466882 | 1.126853 | 0.9782709 |
| ENSG00000060749.14 | QSER1 | -0.007679 | 0.9717201 | 0.18635 | 0.9828321 | 1.72028 | 0.000125894 | 1.316775 | 0.001230734 | 0.92896 | 0.3516806 | 2.517287 | 0.9995474 |
| ENSG00000060762.18 | MPC1 | -1.347815 | 0.1899814 | 0.367534 | 0.2217175 | -0.448232 | 0.02382767 | 0.251051 | 0.01679839 | -0.462998 | 0.08838444 | -0.925521 | 0.8549262 |
| ENSG00000060971.17 | ACAA1 | -0.530376 | 0.8847528 | 0.333444 | 0.8867595 | -0.144597 | 0.001345929 | -0.57205 | 0.4919368 | 0.05133 | 0.7326483 | 0.267997 | 0.8755007 |
| ENSG00000060982.14 | BCAT1 | 1.693294 | 0.6214382 | -0.32195 | 0.08732232 | 1.382379 | 5.74E-08 | 0.414676 | 0.01269998 | 0.690688 | 0.783109 | 0.259971 | 0.8949489 |
| ENSG00000061273.17 | HDAC7 | -1.996908 | 0.2445568 | -0.445599 | 0.9577957 | -1.476826 | 0.2872818 | 1.604018 | 0.0725107 | 2.217467 | 0.000298639 | 1.387975 | 0.6885065 |
| ENSG00000061676.14 | NCKAP1 | 0.219169 | 1 | 0.017295 | 0.8079089 | 0.658885 | 0.5177382 | 0.396524 | 0.002936828 | 0.402738 | 0.4366023 | -0.300571 | 0.8537209 |
| ENSG00000061794.12 | MRPS35 | -0.627067 | 0.6214382 | 0.291067 | 0.6413306 | -0.045619 | 0.8592855 | 0.303669 | 0.3467508 | -0.627305 | 0.001562471 | -0.657103 | 0.8479431 |
| ENSG00000061938.16 | TNK2 | -1.950592 | 0.6615279 | -0.198217 | 0.6200225 | -0.940423 | 0.1822174 | 0.587248 | 0.02882544 | 2.413121 | 0.0052154 | -0.846176 | 0.9726757 |
| ENSG00000062194.15 | GPBP1 | -0.165101 | 0.7134516 | -0.136989 | 0.7585627 | 0.567944 | 5.88E-05 | 0.523909 | 0.06578586 | 0.240658 | 1 | -0.065904 | 0.6885065 |
| ENSG00000062485.18 | CS | -0.416504 | 0.7979044 | -0.192696 | 0.3803801 | 0.323074 | 0.8306727 | 0.706305 | 0.003570292 | 1.765125 | 0.09570889 | -0.060931 | 0.6885065 |
| ENSG00000062598.17 | ELMO2 | 1.811246 | 0.9319271 | 0.073787 | 0.3891394 | -0.751363 | 1 | 0.897559 | 0.0214164 | 2.430965 | 0.01550373 | -0.01688 | 1 |
| ENSG00000062716.10 | VMP1 | 0.134013 | 0.5687266 | 0.100861 | 0.6527736 | -0.104837 | 0.2639048 | 2.053477 | 7.03E-05 | -0.325278 | 0.1526352 | 0.202682 | 0.9665677 |
| ENSG00000062725.9 | APPBP2 | -0.897324 | 0.6214382 | 0.134371 | 0.9827858 | 0.354994 | 0.9471555 | 1.362128 | 0.001106041 | 0.970359 | 0.08966305 | -0.719398 | 0.6576742 |
| ENSG00000063015.19 | SEZ6 | 0.483527 | 0.9438446 | 1.158649 | 0.9644556 | -1.660262 | 0.04753476 | 1.523293 | 0.4919368 | 0.59174 | 0.911559 | 0.978759 | 0.1350388 |
| ENSG00000063046.17 | EIF4B | -0.23742 | 0.6214382 | -0.013107 | 0.7300244 | 0.777806 | 8.26E-09 | 0.683294 | 0.7705399 | 0.940278 | 0.6602576 | -0.572387 | 1 |
| ENSG00000063169.10 | GLTSCR1 | 0.585848 | 0.6214382 | -1.662271 | 0.591903 | -0.309504 | 1 | 2.43242 | 0.18557 | 1.920217 | 0.04278906 | -1.138851 | 1 |
| ENSG00000063177.12 | RPL18 | -0.102271 | 0.8867323 | -0.15145 | 0.04870784 | -0.239929 | 1.85E-05 | 0.10287 | 0.6052147 | 0.098011 | 0.116144 | 0.099432 | 0.6530179 |
| ENSG00000063241.7 | ISOC2 | -0.750571 | 0.8860308 | 0.530944 | 0.0252253 | -0.452803 | 0.2189079 | 0.289067 | 0.5447452 | 0.363206 | 0.4739539 | 0.039974 | 0.6941888 |
| ENSG00000063245.14 | EPN1 | -0.980361 | 0.6214382 | 0.195768 | 0.9787982 | -0.838239 | 0.009166088 | 0.800049 | 0.1239499 | 0.018913 | 0.5196843 | -0.420593 | 0.8479431 |
| ENSG00000063587.13 | ZNF275 | -1.231811 | 0.8009748 | -0.063465 | 0.5953732 | -0.329709 | 0.1826859 | 1.215276 | 0.01011358 | 1.46332 | 0.642531 | -1.963612 | 0.6829816 |
| ENSG00000063601.16 | MTMR1 | 0.614337 | 0.9047689 | 0.316398 | 0.8262235 | 0.356762 | 0.7437308 | 1.233939 | 0.3493762 | 1.320511 | 0.01500431 | 3.05393 | 0.6185816 |
| ENSG00000063660.8 | GPC1 | -0.590576 | 0.5930856 | -0.268818 | 0.2562412 | -0.679492 | 0.001389931 | 1.30879 | 5.09E-06 | 1.004559 | 0.1627544 | -0.814669 | 1 |
| ENSG00000063854.12 | HAGH | 0.436284 | 0.7343911 | 0.320864 | 0.797408 | 0.137363 | 0.07810551 | -0.564762 | 0.03655283 | 0.629221 | 0.5574773 | -0.037988 | 0.9590434 |
| ENSG00000064042.17 | LIMCH1 | 1.334192 | 0.8006286 | 0.077137 | 0.4967252 | 0.912446 | 0.02360632 | 0.460184 | 0.954187 | 0.383312 | 0.673315 | 3.053165 | 0.6885065 |
| ENSG00000064300.8 | NGFR | -2.318831 | 0.6214382 | -0.111694 | 1 | -0.137206 | 0.6646119 | 1.442646 | 0.02378496 | 0.594389 | 0.7646967 | 0.396539 | 0.8479431 |
| ENSG00000064313.11 | TAF2 | 1.163541 | 0.5416712 | -0.033543 | 0.9911659 | -0.380668 | 0.3932734 | 0.502928 | 0.04115099 | 0.679267 | 0.7377262 | -1.288325 | 0.9510409 |
| ENSG00000064393.15 | HIPK2 | -0.390476 | 0.4318827 | -0.446421 | 0.1794587 | 0.867951 | 3.35E-05 | 1.737819 | 6.75E-11 | 0.680944 | 0.796751 | -0.205729 | 0.6517664 |
| ENSG00000064419.13 | TNPO3 | 0.639307 | 0.8006286 | -0.193986 | 0.6258774 | -0.055176 | 0.7437308 | 0.926809 | 0.000100462 | 1.116948 | 0.07221764 | -0.298875 | 0.6185816 |
| ENSG00000064607.16 | SUGP2 | -0.755341 | 0.09877095 | -0.060367 | 0.7079982 | 0.387751 | 0.2773118 | 1.187669 | 0.001122986 | 1.138629 | 0.0689338 | 0.534585 | 0.7871323 |
| ENSG00000064651.13 | SLC12A2 | -1.139245 | 0.6917716 | 0.116015 | 0.7545427 | 0.458072 | 0.4430409 | 0.790208 | 0.04055961 | -0.002003 | 0.72638 | 1.833337 | 0.9716684 |
| ENSG00000064666.14 | CNN2 | 0.813098 | 0.6014343 | 1.07796 | 0.32473 | 0.175407 | 0.7693698 | 5.72908 | 0.08788191 | -0.422664 | 0.04130846 | inf | 0.6885065 |
| ENSG00000064933.16 | PMS1 | -0.728956 | 0.6214382 | 0.184295 | 0.9903933 | 1.383572 | 0.008435237 | 2.698088 | 0.5866249 | 0.027924 | 0.7270816 | -0.22264 | 0.9169962 |
| ENSG00000064995.16 | TAF11 | -0.719699 | 0.8561029 | -0.082098 | 0.9973106 | -0.153319 | 0.04806286 | -0.237531 | 0.547158 | -0.114412 | 0.5196843 | 0.017755 | 0.6885065 |
| ENSG00000064999.13 | ANKS1A | 4.232355 | 0.563505 | -0.01745 | 0.8653838 | 0.274843 | 0.7437308 | 2.395346 | 0.00721147 | 0.566853 | 0.8427659 | 1.400933 | 0.8479431 |
| ENSG00000065000.15 | AP3D1 | -0.544466 | 0.636947 | -0.045201 | 0.370295 | 0.374005 | 0.04511416 | 0.686832 | 0.000719841 | -0.885455 | 1 | -0.127379 | 0.7515027 |
| ENSG00000065057.7 | NTHL1 | -0.222381 | 0.6014343 | -0.515599 | 0.6527736 | -0.670251 | 0.01208004 | -0.479489 | 0.3622213 | -0.685348 | 0.3638894 | -0.30551 | 0.8687124 |
| ENSG00000065060.16 | UHRF1BP1 | -0.980494 | 0.6214382 | -0.465679 | 0.5851668 | 0.140468 | 0.01290545 | 0.366681 | 0.1638517 | 0.403418 | 0.3181982 | -3.538395 | 0.5066477 |
| ENSG00000065135.8 | GNAI3 | 0.567855 | 0.6014343 | 0.067364 | 0.7735491 | 0.577485 | 0.002559654 | -0.028288 | 0.864147 | 0.685118 | 0.8753948 | 0.779997 | 0.3565625 |
| ENSG00000065154.11 | OAT | -0.973053 | 0.8932278 | 0.123656 | 0.9618421 | 1.375498 | 0.005854557 | -0.633679 | 0.314059 | 0.465509 | 0.7500517 | 0.711659 | 0.6885065 |
| ENSG00000065243.18 | PKN2 | 1.501939 | 0.3184862 | 0.014199 | 0.8893429 | 0.670654 | 0.007979364 | 0.717204 | 0.4207258 | 1.075989 | 0.4655627 | 0.014861 | 0.6885065 |
| ENSG00000065320.8 | NTN1 | -0.665718 | 0.8006286 | 0.011423 | 0.7881421 | 0.279222 | 0.08610771 | 1.586569 | 6.63E-06 | 1.433687 | 0.5196843 | 0.045507 | 0.715468 |
| ENSG00000065413.16 | ANKRD44 | -1.695054 | 0.6214382 | 0.074893 | 0.9754542 | 0.368281 | 0.8348198 | 3.840238 | 0.000105119 | 0.110328 | 0.7404912 | -0.206222 | 0.8479431 |
| ENSG00000065457.10 | ADAT1 | -0.018677 | 0.6792949 | 1.230555 | 0.8371079 | -0.58722 | 0.907132 | 1.716207 | 0.4919368 | 1.218383 | 0.04742931 | -2.828416 | 0.6885065 |
| ENSG00000065485.17 | PDIA5 | -0.982644 | 0.6214382 | 0.013776 | 0.9664473 | 0.47663 | 0.6877986 | 0.880576 | 0.1482986 | 3.270928 | 0.04278906 | 2.135451 | 1 |
| ENSG00000065518.7 | NDUFB4 | -0.487902 | 0.2617323 | -0.175939 | 0.5953732 | -0.40737 | 0.00010804 | -0.276508 | 8.34E-06 | 0.671129 | 0.8316598 | -0.726349 | 0.6829816 |
| ENSG00000065526.10 | SPEN | -0.691517 | 0.6014343 | 0.473484 | 0.5069871 | 0.949031 | 7.77E-06 | 0.57847 | 0.003228635 | 1.592287 | 0.003546663 | -1.20803 | 0.5638538 |
| ENSG00000065548.17 | ZC3H15 | -0.955343 | 0.1835496 | 0.131537 | 0.957632 | 0.964033 | 2.56E-07 | 0.835858 | 0.4060414 | -6.90E-05 | 0.1981203 | -1.144266 | 0.05044736 |
| ENSG00000065600.12 | TMEM206 | -0.97434 | 0.6084645 | 0.006291 | 0.5394427 | 0.384141 | 0.2469493 | 0.421631 | 1 | -0.245933 | 0.01412964 | -0.652984 | 0.6885065 |
| ENSG00000065802.11 | ASB1 | -0.76916 | 0.714717 | -0.038235 | 0.8703522 | 0.471519 | 0.9013136 | 1.002095 | 0.01539364 | 2.194442 | 0.09208386 | 0.020614 | 0.9526606 |
| ENSG00000065809.13 | FAM107B | -1.166093 | 0.6014343 | 0.2993 | 0.8989356 | 0.477816 | 0.5290845 | 1.422976 | 0.008249854 | 0.262655 | 0.08266705 | -1.145305 | 0.6885065 |
| ENSG00000065883.14 | CDK13 | -0.227336 | 0.8867323 | 0.226053 | 0.8033592 | 0.211396 | 0.06218784 | 1.407135 | 0.007565047 | 1.040538 | 0.2880317 | -0.129423 | 0.6885065 |
| ENSG00000065911.11 | MTHFD2 | -0.012138 | 0.8006286 | 0.275147 | 0.8863478 | 0.60471 | 0.003539928 | 0.677379 | 0.9597651 | 0.378795 | 0.5539249 | -0.942774 | 1 |
| ENSG00000065923.9 | SLC9A7 | -0.127406 | 0.8587992 | 0.176658 | 0.7871602 | 0.475584 | 0.5740698 | 1.913349 | 0.0307706 | 1.390434 | 0.2364543 | 1.972107 | 0.9704304 |
| ENSG00000065970.8 | FOXJ2 | 0.184439 | 1 | -0.161291 | 0.9793419 | 0.848883 | 0.01802117 | -0.412433 | 1 | 0.553693 | 0.8146399 | -0.012352 | 0.9684964 |
| ENSG00000066032.18 | CTNNA2 | 1.554007 | 0.6014343 | -2.208473 | 0.2815893 | -0.788756 | 0.8454005 | 0.64424 | 0.01635326 | 0.728727 | 0.8779042 | -0.18712 | 0.7041444 |
| ENSG00000066084.12 | DIP2B | 0.297656 | 0.6214382 | 0.062123 | 0.9981477 | 0.076031 | 0.7437308 | 0.550298 | 0.009064554 | -0.227371 | 0.7804595 | -0.673183 | 0.9204712 |
| ENSG00000066117.14 | SMARCD1 | 0.452276 | 0.4399567 | -0.454816 | 0.5289109 | 1.041968 | 0.09046443 | 1.407402 | 0.04372344 | 0.891256 | 0.4480624 | 0.683026 | 0.6185816 |
| ENSG00000066427.21 | ATXN3 | -0.476268 | 0.6214382 | 0.593082 | 0.8863806 | 1.045073 | 0.6932338 | 0.879121 | 0.1309039 | 2.830525 | 0.03306085 | 2.747347 | 0.6885065 |
| ENSG00000066654.12 | THUMPD1 | -0.867703 | 0.6014343 | -0.053446 | 0.8472233 | 0.401678 | 0.08713411 | 2.327028 | 0.002897201 | 0.524783 | 0.9056171 | -0.024731 | 0.7656733 |
| ENSG00000066739.11 | ATG2B | 2.472225 | 0.6214382 | -0.062563 | 0.8953649 | 1.115176 | 0.8485513 | -0.802467 | 0.02859072 | 2.396847 | 0.4874572 | -2.358339 | 0.6885065 |
| ENSG00000066777.8 | ARFGEF1 | -1.296981 | 0.6014343 | -0.099092 | 0.8263112 | 0.866062 | 0.004398576 | 1.673479 | 0.001023989 | 1.412796 | 0.01171107 | 1.52445 | 0.9077431 |
| ENSG00000066933.15 | MYO9A | -5.576517 | 0.65581 | 0.06104 | 0.7391331 | 2.129686 | 1.52E-05 | 1.921754 | 0.000510135 | -0.120812 | 0.5966036 | -3.381295 | 0.8041706 |
| ENSG00000067082.14 | KLF6 | 0.036953 | 0.6014343 | -0.006744 | 0.6527736 | 0.739665 | 2.53E-07 | 0.713904 | 0.6447903 | 1.265949 | 0.8910863 | -0.575896 | 0.8539279 |
| ENSG00000067141.16 | NEO1 | 0.32016 | 0.9047689 | -0.626083 | 0.5037981 | 0.268526 | 0.4674985 | 2.185527 | 0.125732 | 4.297389 | 0.0325649 | -0.058291 | 0.7308361 |
| ENSG00000067182.7 | TNFRSF1A | -5.505393 | 0.957965 | -0.174957 | 0.3494625 | -0.697501 | 7.39E-05 | 1.081457 | 0.03891837 | 0.878135 | 1 | -0.432841 | 0.6885065 |
| ENSG00000067208.14 | EVI5 | -0.75599 | 0.8006286 | 0.558647 | 0.9452791 | 0.659118 | 0.1182794 | -1.566916 | 0.3760083 | -2.412473 | 0.03104635 | 0.260308 | 0.7251306 |
| ENSG00000067248.9 | DHX29 | -0.29918 | 0.9016905 | 0.222343 | 0.9564082 | 1.111252 | 0.005370461 | 1.651381 | 0.4229371 | 1.14996 | 0.7003847 | 0.404877 | 0.6885065 |
| ENSG00000067369.13 | TP53BP1 | -1.556259 | 0.6374294 | 0.444799 | 0.7199683 | 0.311357 | 0.1710895 | 1.260541 | 0.01150992 | 1.520976 | 0.2089173 | 1.671811 | 0.7767294 |
| ENSG00000067560.10 | RHOA | 0.098687 | 0.6214382 | -0.038071 | 0.7443164 | 0.484181 | 0.01539792 | 0.083223 | 0.01929238 | 0.153973 | 0.0948816 | 0.306566 | 0.6885065 |
| ENSG00000067840.12 | PDZD4 | 1.09926 | 0.6214382 | 0.690186 | 0.5110299 | -0.101666 | 0.4070201 | 2.30766 | 6.59E-08 | 1.137899 | 0.01075306 | -0.610939 | 0.6421783 |
| ENSG00000067900.7 | ROCK1 | -0.409542 | 0.8006286 | -0.256266 | 0.5010139 | 1.406583 | 0.0123839 | 2.076912 | 5.19E-05 | 1.554816 | 0.006214297 | -0.486663 | 0.6948095 |
| ENSG00000068137.14 | PLEKHH3 | -0.912096 | 1 | -0.189652 | 0.7687946 | -2.019535 | 0.2808121 | 3.351786 | 0.09515288 | 1.704502 | 0.000369876 | -0.276467 | 0.8812173 |
| ENSG00000068305.17 | MEF2A | 0.355156 | 0.7000696 | 0.113017 | 0.6527736 | 0.407458 | 0.4044598 | 1.915654 | 0.001122986 | 0.421291 | 0.8975132 | 2.376904 | 0.6813272 |
| ENSG00000068323.16 | TFE3 | 4.27951 | 0.3098608 | -0.458246 | 0.9668554 | 1.391799 | 0.03662563 | 0.558228 | 0.4492037 | -0.2906 | 0.8327114 | -1.143583 | 0.6885065 |
| ENSG00000068394.10 | GPKOW | -0.15322 | 0.8006286 | 0.44136 | 0.6527736 | 0.196794 | 1 | -0.028929 | 0.02308096 | -0.486159 | 0.7574155 | -0.910583 | 0.967026 |
| ENSG00000068654.15 | POLR1A | -0.283189 | 0.9419384 | -0.686429 | 0.7132271 | 0.805654 | 0.5290845 | 1.362191 | 0.001302451 | 2.494395 | 0.098681 | 1.82747 | 0.8757616 |
| ENSG00000068697.6 | LAPTM4A | -0.511983 | 0.7757496 | 0.274792 | 0.4487482 | 0.302862 | 0.9619523 | -0.21174 | 0.01714385 | -0.246902 | 0.0171907 | 0.55829 | 0.7670687 |
| ENSG00000068796.16 | KIF2A | 0.161547 | 0.8867323 | 0.337349 | 1 | 0.668165 | 9.38E-05 | -0.108464 | 0.6733179 | -0.600462 | 0.009659558 | -0.536108 | 0.6885065 |
| ENSG00000068878.14 | PSME4 | -0.645936 | 0.9669959 | -0.43177 | 0.8314425 | 0.314743 | 0.3591176 | 0.607872 | 5.25E-05 | 1.982721 | 0.003261631 | 0.777106 | 0.926878 |
| ENSG00000068903.19 | SIRT2 | -0.690266 | 0.8587992 | 0.295179 | 0.7881421 | 0.07896 | 0.5290845 | 1.095353 | 0.004364008 | 1.397816 | 0.3956225 | -0.968221 | 0.9638718 |
| ENSG00000068912.13 | ERLEC1 | -0.301385 | 0.8587992 | 0.213863 | 0.6527736 | 0.673278 | 0.04947689 | -0.028934 | 0.06044308 | -0.313394 | 0.1213641 | 2.359234 | 0.5344421 |
| ENSG00000068971.13 | PPP2R5B | 0.858081 | 0.2774235 | 1.031623 | 0.2009498 | 0.033034 | 0.8384231 | 1.699567 | 0.01194267 | 0.315635 | 0.4821192 | -0.220724 | 0.7766607 |
| ENSG00000069248.11 | NUP133 | -1.303865 | 0.9047689 | 0.474495 | 0.9317444 | 0.759017 | 0.640899 | 2.798809 | 0.004386363 | 1.667767 | 0.7425244 | -0.166842 | 0.8949489 |
| ENSG00000069275.12 | NUCKS1 | -0.724388 | 0.06250334 | 0.03416 | 0.6657165 | 0.836739 | 3.16E-09 | 0.141073 | 0.03681075 | 0.401952 | 0.2626143 | -0.329369 | 0.2321124 |
| ENSG00000069329.15 | VPS35 | 0.267969 | 0.5006203 | -0.041464 | 0.5953732 | 1.265792 | 0.00013619 | 0.412946 | 0.7125861 | -0.129853 | 0.0722616 | -1.582307 | 0.9114828 |
| ENSG00000069345.11 | DNAJA2 | 0.107643 | 0.8006286 | -0.321719 | 0.5953732 | 0.573641 | 0.000904019 | 0.861653 | 0.9046903 | -0.086333 | 0.4671878 | 0.513536 | 0.958722 |
| ENSG00000069493.14 | CLEC2D | -1.160604 | 0.6214382 | -0.759638 | 0.8923146 | -0.018923 | 0.5840065 | 0.966768 | 0.00045875 | 0.725651 | 0.4838709 | -0.384303 | 0.6345741 |
| ENSG00000069849.10 | ATP1B3 | -0.60983 | 0.7558677 | -0.041281 | 0.7766756 | -0.299451 | 0.5510596 | 0.870916 | 0.06472093 | 1.412975 | 0.001955021 | -0.296239 | 0.4413458 |
| ENSG00000069869.15 | NEDD4 | 0.508904 | 0.8867323 | -0.535828 | 0.7955905 | 0.61076 | 0.2692975 | 1.670914 | 0.004511404 | 0.248945 | 0.9459937 | 0.391699 | 0.8278546 |
| ENSG00000069956.11 | MAPK6 | -0.054872 | 0.8712126 | 0.090257 | 0.659823 | 0.194078 | 0.04530346 | 1.194838 | 0.03118701 | 0.063261 | 0.8458207 | 1.529404 | 0.7815439 |
| ENSG00000069966.18 | GNB5 | -1.147119 | 0.6595756 | -0.55709 | 0.6527736 | 2.530233 | 0.006870424 | 1.570306 | 0.1089057 | 0.478062 | 0.7437199 | -0.524523 | 1 |
| ENSG00000070018.8 | LRP6 | -0.820828 | 0.8006286 | 0.554826 | 0.6527736 | 1.193466 | 0.005818383 | 3.291649 | 0.1022235 | 1.729833 | 0.008567685 | 0.611218 | 0.9663203 |
| ENSG00000070081.15 | NUCB2 | -1.23806 | 0.5182912 | 0.039519 | 0.5953732 | 0.470655 | 0.09399964 | 0.147897 | 0.001385729 | -0.489673 | 6.14E-06 | 2.113419 | 0.6517664 |
| ENSG00000070367.15 | EXOC5 | -0.256853 | 0.9183025 | -0.17674 | 0.5296018 | 0.859956 | 0.006321013 | 1.188214 | 0.1189105 | -0.485564 | 0.2713288 | 0.404344 | 0.9705209 |
| ENSG00000070444.14 | MNT | 0.936056 | 1 | -0.38478 | 0.6914453 | -0.212794 | 0.7925802 | 0.64295 | 0.02455892 | 0.796424 | 0.3812741 | 1.717684 | 0.7499106 |
| ENSG00000070669.16 | ASNS | -0.479579 | 0.977255 | 0.162291 | 0.7337542 | -0.208838 | 0.8599056 | 0.196601 | 1.83E-08 | 0.753349 | 0.001539519 | -0.298063 | 0.6885065 |
| ENSG00000070756.13 | PABPC1 | -0.830562 | 0.000369055 | 0.118022 | 1 | 1.035363 | 1.36E-29 | 0.032312 | 0.008823624 | 0.462601 | 0.8949249 | 0.532642 | 0.509185 |
| ENSG00000070770.8 | CSNK2A2 | 0.014768 | 0.8767328 | 0.110666 | 1 | 0.519124 | 0.05182881 | 1.778385 | 0.003258211 | 1.356234 | 0.1144781 | -0.627676 | 0.7491255 |
| ENSG00000070814.17 | TCOF1 | -0.437521 | 0.9085559 | -0.563387 | 0.5953732 | 0.070561 | 0.7735461 | 0.747498 | 0.02268262 | -0.223081 | 0.264938 | 0.372644 | 0.8479431 |
| ENSG00000070961.15 | ATP2B1 | 0.075163 | 0.6214382 | -0.041338 | 0.5953732 | 0.418574 | 0.22883 | 1.287543 | 0.000526142 | 0.11542 | 0.3570564 | 1.768997 | 0.6885065 |
| ENSG00000071054.16 | MAP4K4 | -0.425919 | 0.9305688 | -0.290891 | 0.5295559 | 0.874439 | 0.000890171 | 1.235426 | 1.56E-06 | 0.894526 | 0.006574444 | -1.145108 | 0.2892129 |
| ENSG00000071082.10 | RPL31 | -0.431939 | 0.6566178 | 0.036148 | 1 | -0.084948 | 0.0582643 | 0.033883 | 0.1898364 | -0.20505 | 3.20E-05 | 0.084475 | 0.6231061 |
| ENSG00000071246.10 | VASH1 | 1.588072 | 0.9047689 | -0.11322 | 0.964386 | 0.577672 | 0.7437308 | -0.622558 | 0.3153792 | 1.480923 | 0.005245519 | 1.161641 | 0.6885065 |
| ENSG00000071462.11 | WBSCR22 | -0.622198 | 0.2351592 | 0.219567 | 0.686184 | 0.201296 | 0.5290845 | -0.039712 | 0.000105119 | 0.119551 | 0.1253756 | -0.100897 | 0.3532606 |
| ENSG00000071537.13 | SEL1L | 0.198102 | 0.6214382 | 0.738048 | 0.1794587 | 0.596688 | 0.7437308 | 1.85576 | 0.000588925 | 1.309812 | 0.3676466 | -1.049147 | 0.8291878 |
| ENSG00000071553.16 | ATP6AP1 | 0.000263 | 0.9850641 | 0.318245 | 0.7828571 | -0.304723 | 0.03616489 | 0.6176 | 0.1124594 | 0.574178 | 0.9438088 | -0.327848 | 1 |
| ENSG00000071564.14 | TCF3 | -4.911502 | 0.9047689 | -0.158749 | 0.5953732 | -0.23219 | 0.335147 | 0.641634 | 0.01075895 | 0.786513 | 0.1318334 | -0.96516 | 1 |
| ENSG00000071575.11 | TRIB2 | -0.039445 | 1 | 0.002581 | 0.7759924 | 0.085388 | 0.1723306 | 0.773969 | 5.64E-05 | 0.943787 | 0.1532581 | -0.487843 | 0.7501684 |
| ENSG00000071626.16 | DAZAP1 | -2.43595 | 0.0318879 | -0.467868 | 0.8528295 | -0.03336 | 0.2894867 | 1.057493 | 0.005068064 | 1.084442 | 0.009101151 | -1.412335 | 0.5344421 |
| ENSG00000071655.17 | MBD3 | 0.216246 | 0.9057808 | 0.405161 | 0.8446275 | 0.40513 | 0.8592813 | 0.683713 | 9.84E-05 | 1.766413 | 0.01440053 | -0.812087 | 0.8479431 |
| ENSG00000071794.15 | HLTF | 0.871305 | 0.6214382 | 0.094474 | 0.704767 | 1.340952 | 0.03437177 | 0.37657 | 0.4062243 | -0.06249 | 0.5425935 | -0.137745 | 0.6885065 |
| ENSG00000071859.14 | FAM50A | -0.094341 | 0.8006286 | -0.088013 | 0.859273 | -1.207304 | 0.01907952 | -0.190253 | 0.4068061 | 1.29483 | 0.2363864 | 0.618129 | 0.3849244 |
| ENSG00000071994.10 | PDCD2 | -0.771746 | 0.1971316 | -0.281254 | 0.6527736 | 0.725611 | 0.2879524 | 1.083572 | 0.04561241 | 1.647565 | 0.03224226 | 1.761804 | 0.9963891 |
| ENSG00000072071.16 | ADGRL1 | -0.194799 | 0.8006286 | -0.438601 | 0.1905717 | -0.736445 | 0.7500241 | 0.138631 | 0.03580438 | 2.099958 | 0.00012052 | 0.715544 | 1 |
| ENSG00000072182.12 | ASIC4 | -1.07836 | 0.6214382 | -0.014959 | 0.7049761 | -0.223281 | 0.3276184 | 3.93667 | 0.002728072 | 0.991486 | 0.1835921 | -2.715678 | 0.9000979 |
| ENSG00000072364.12 | AFF4 | -1.136266 | 0.01740848 | -0.207201 | 0.6527736 | 0.664713 | 9.60E-06 | 0.421681 | 0.1190109 | 1.331345 | 0.02704086 | 0.659588 | 0.809678 |
| ENSG00000072501.17 | SMC1A | 0.381261 | 0.9687342 | -0.007753 | 0.7348129 | 0.248307 | 0.5020747 | 0.082982 | 0.000812686 | 0.733233 | 0.6447644 | -0.589158 | 0.9279511 |
| ENSG00000072506.12 | HSD17B10 | 0.537491 | 0.787986 | -0.001512 | 0.6527736 | -0.713084 | 8.91E-10 | 0.186656 | 0.4558202 | -0.360182 | 0.3907918 | 0.612079 | 0.3664193 |
| ENSG00000072518.20 | MARK2 | -1.24763 | 0.714717 | 0.454496 | 0.6527736 | 0.294906 | 0.5010887 | 1.508644 | 0.01439514 | 0.493306 | 0.2022966 | -1.451483 | 0.7614886 |
| ENSG00000072736.18 | NFATC3 | 0.051065 | 0.714717 | -0.140183 | 0.8263112 | -1.782889 | 0.9203772 | 1.833309 | 0.000289925 | 1.563401 | 0.002838476 | -3.584791 | 0.6185816 |
| ENSG00000072803.17 | FBXW11 | -1.274949 | 0.6214382 | -0.266095 | 0.9151764 | -0.412154 | 0.03844384 | 0.456698 | 0.1433935 | 1.086768 | 0.5912143 | -0.493519 | 0.7829412 |
| ENSG00000072832.14 | CRMP1 | 0.722681 | 0.367978 | -0.174533 | 0.6527736 | -0.275725 | 0.4555974 | 0.506741 | 0.05196313 | 0.930246 | 0.04590921 | 0.477563 | 0.3746505 |
| ENSG00000073008.14 | PVR | 0.461528 | 0.8587992 | -0.40874 | 0.5953732 | 0.026795 | 0.3897481 | 0.666839 | 0.01980018 | 0.328696 | 1 | -0.466846 | 0.6885065 |
| ENSG00000073169.13 | SELO | 0.837579 | 0.8006902 | -0.203562 | 1 | -0.218305 | 0.5290845 | 1.726718 | 0.02658249 | 3.069636 | 0.03056663 | -1.550442 | 0.8949489 |
| ENSG00000073417.14 | PDE8A | -1.308683 | 0.7736255 | -1.229093 | 0.04333419 | 0.544198 | 0.7669328 | 1.596702 | 0.009729605 | 1.531009 | 0.5196843 | 1.994679 | 0.8479431 |
| ENSG00000073614.11 | KDM5A | 1.19985 | 0.5546178 | 0.191107 | 0.8200822 | 0.382587 | 0.004950796 | 1.69604 | 0.002897201 | 0.512208 | 0.6185494 | -1.122633 | 0.916449 |
| ENSG00000073712.14 | FERMT2 | -0.040702 | 0.8491439 | -0.027244 | 0.5515692 | 0.333814 | 0.6983378 | 0.674476 | 0.7277058 | 2.686763 | 0.04995056 | 0.058316 | 0.6885065 |
| ENSG00000073921.17 | PICALM | -6.196297 | 0.5441969 | -0.123696 | 0.6369093 | 0.1543 | 0.7010938 | 0.62997 | 0.02375284 | 0.239127 | 0.4465705 | 1.673015 | 1 |
| ENSG00000074054.17 | CLASP1 | 0.607391 | 0.8408841 | 0.102342 | 0.9073435 | 1.011416 | 0.8026534 | 0.344116 | 0.3897483 | 1.375334 | 0.04635848 | -0.538663 | 0.8644876 |
| ENSG00000074071.13 | MRPS34 | 0.760978 | 0.529676 | 0.020626 | 0.5310556 | -0.423858 | 1.25E-06 | 0.149391 | 0.1708872 | 0.385866 | 0.006051963 | -1.002994 | 0.2995901 |
| ENSG00000074356.16 | NCBP3 | -0.517939 | 0.3946735 | -0.328904 | 0.9618421 | 0.721027 | 0.3427209 | 2.493197 | 1.01E-06 | 1.679093 | 0.01473534 | 1.371847 | 0.8479431 |
| ENSG00000074410.13 | CA12 | 0.80248 | 0.9972233 | 0.664945 | 0.008839116 | -0.167004 | 0.4973551 | 0.326737 | 0.3946292 | -0.642788 | 0.6244585 | 2.911915 | 0.5066477 |
| ENSG00000074603.18 | DPP8 | -0.440234 | 0.8806263 | -0.307081 | 0.7680378 | 0.6657 | 0.04939769 | 0.412473 | 0.00444805 | -0.304456 | 0.3766175 | 0.591451 | 0.6829816 |
| ENSG00000074657.13 | ZNF532 | -2.336563 | 0.6014343 | -0.474746 | 0.4562492 | 0.794536 | 0.3919757 | 1.099655 | 0.004012816 | 1.257256 | 0.1850728 | -0.159897 | 0.6185816 |
| ENSG00000074695.5 | LMAN1 | 0.143224 | 0.8006286 | -0.100077 | 0.7855215 | 1.22854 | 3.12E-14 | -0.218177 | 0.1680423 | -0.017178 | 0.3993666 | 0.992525 | 0.6829816 |
| ENSG00000074696.12 | HACD3 | 0.582249 | 0.6214382 | 0.051818 | 0.9906117 | 0.673404 | 0.02207811 | 0.445082 | 0.5328876 | -0.098188 | 0.5529159 | 0.585278 | 1 |
| ENSG00000074755.14 | ZZEF1 | -1.250723 | 0.6014343 | -0.012722 | 0.7581042 | -0.545822 | 0.6222102 | 1.870882 | 0.01487446 | 2.400382 | 1.34E-06 | 0.522055 | 0.9517616 |
| ENSG00000074800.13 | ENO1 | -0.461824 | 0.5104673 | 0.213895 | 0.005123222 | 0.349889 | 0.04711588 | -0.156082 | 0.003704982 | -0.509756 | 1.20E-08 | -0.178227 | 0.9728326 |
| ENSG00000074842.7 | MYDGF | -0.502945 | 0.862314 | 0.297528 | 0.6787205 | -0.553204 | 1.68E-08 | -0.272104 | 0.02665019 | 0.179251 | 0.005390303 | -0.234131 | 0.9270789 |
| ENSG00000074855.10 | ANO8 | 3.756335 | 0.6214382 | -0.291601 | 0.6527736 | 1.343731 | 0.7368441 | 0.700621 | 0.4778509 | 2.278892 | 0.0266455 | -0.388023 | 0.6885065 |
| ENSG00000075142.13 | SRI | 0.370896 | 0.714717 | -0.078913 | 0.3206709 | 0.232159 | 0.4722274 | -0.202094 | 3.49E-05 | -0.586927 | 1.53E-08 | -0.528119 | 0.6829816 |
| ENSG00000075151.20 | EIF4G3 | -0.330069 | 0.3098608 | -0.243976 | 0.06246231 | 0.777186 | 5.51E-05 | 0.850147 | 0.006549993 | 0.414456 | 0.6347671 | 0.698714 | 0.6829816 |
| ENSG00000075292.18 | ZNF638 | -0.857977 | 0.8921933 | 0.27736 | 0.6527736 | 0.599611 | 6.92E-06 | 1.005631 | 1.71E-07 | 1.618635 | 0.000145639 | -1.338823 | 0.5023959 |
| ENSG00000075407.17 | ZNF37A | -0.99657 | 0.529676 | -0.092116 | 0.7383998 | 0.787553 | 0.2938159 | 3.083874 | 0.01460212 | -0.030987 | 0.3628287 | -2.051856 | 0.9147227 |
| ENSG00000075413.17 | MARK3 | -0.722016 | 0.563505 | -0.207799 | 0.5953732 | 1.143958 | 0.2733328 | 0.478406 | 0.01539252 | 0.027644 | 0.8021966 | -0.650327 | 0.8479431 |
| ENSG00000075426.11 | FOSL2 | 0.527912 | 0.714717 | -0.073172 | 0.6527736 | -1.063943 | 0.1071577 | 1.671198 | 1.75E-05 | 1.810751 | 0.2707324 | 0.272868 | 0.7565106 |
| ENSG00000075461.5 | CACNG4 | 0.922287 | 1 | 0.546127 | 0.8653057 | 0.231006 | 0.5290845 | 1.162113 | 0.06861004 | 1.08771 | 0.03545956 | -1.245497 | 0.8949489 |
| ENSG00000075539.13 | FRYL | 0.878965 | 0.714717 | 0.211214 | 0.9410094 | 0.53552 | 0.00543407 | 0.672459 | 0.3878731 | 0.428418 | 0.5196843 | 1.036239 | 0.6231061 |
| ENSG00000075568.16 | TMEM131 | 1.186626 | 0.8006286 | -0.572281 | 0.01857467 | 0.214305 | 0.7437308 | 2.326945 | 0.000480157 | 1.262901 | 0.5206811 | 1.712605 | 0.5066477 |
| ENSG00000075618.17 | FSCN1 | 0.307255 | 0.8006286 | 0.148013 | 0.7994859 | -0.109548 | 0.0234506 | 1.00382 | 0.000276572 | 1.479043 | 4.14E-05 | -0.50131 | 0.7554837 |
| ENSG00000075624.13 | ACTB | 0.201733 | 3.60E-05 | 0.018691 | 0.9440556 | -0.211873 | 0.03439481 | 0.481826 | 1.86E-08 | 0.159865 | 0.8876055 | -0.03924 | 0.6885065 |
| ENSG00000075711.20 | DLG1 | 1.086445 | 0.8217227 | 0.066993 | 0.7255787 | -0.353516 | 0.4602617 | 0.529771 | 0.104499 | 1.672199 | 0.01777855 | 1.612567 | 0.6885065 |
| ENSG00000075945.12 | KIFAP3 | 0.888378 | 0.6169321 | 0.624519 | 0.5953732 | -0.435252 | 0.8583002 | 0.516695 | 0.912883 | -0.481143 | 0.01143583 | -0.250506 | 0.8992064 |
| ENSG00000075975.15 | MKRN2 | 2.226599 | 0.3946735 | -0.333455 | 0.5292164 | 0.083233 | 0.999519 | 1.58818 | 0.009399882 | 1.72401 | 0.02785769 | -1.055349 | 0.8479431 |
| ENSG00000076108.11 | BAZ2A | -0.889164 | 0.529676 | -0.193339 | 0.115387 | 0.5475 | 0.004447682 | 0.10753 | 0.4039579 | 0.990295 | 0.5539689 | 0.349033 | 0.359138 |
| ENSG00000076356.6 | PLXNA2 | 0.973139 | 0.714717 | -0.654441 | 0.6834954 | 0.011702 | 0.6233368 | 2.223058 | 0.007120975 | 1.96263 | 0.02745483 | 0.301988 | 0.9565287 |
| ENSG00000076513.16 | ANKRD13A | 0.920886 | 0.714717 | 0.064777 | 0.7426265 | -0.137747 | 0.7679107 | 1.209115 | 0.2564877 | 1.894635 | 0.04562023 | -1.19511 | 0.960853 |
| ENSG00000076706.14 | MCAM | -0.078283 | 0.9047689 | -0.188626 | 0.8263112 | -0.67438 | 0.9377685 | 1.257782 | 0.03629109 | 1.305184 | 0.7592987 | 0.808092 | 1 |
| ENSG00000076984.17 | MAP2K7 | 0.823649 | 0.7800288 | -0.048774 | 0.7599168 | -0.421176 | 0.2013354 | 1.730446 | 0.001697466 | 1.462817 | 0.02052479 | -0.27222 | 0.8273039 |
| ENSG00000077097.13 | TOP2B | 0.043132 | 0.9719553 | 0.150308 | 0.9068973 | 0.693054 | 0.2879524 | 1.458387 | 0.02384542 | 0.297573 | 0.1131988 | -0.100754 | 0.5344421 |
| ENSG00000077147.14 | TM9SF3 | -0.672339 | 0.8587992 | 0.830003 | 0.1236684 | -0.050408 | 0.6739221 | 0.60446 | 0.02280939 | 0.103039 | 0.8020128 | 0.102267 | 0.2281002 |
| ENSG00000077152.9 | UBE2T | -1.207093 | 0.7736255 | 0.223964 | 0.8309229 | 0.086217 | 0.5335698 | -0.640976 | 0.02764017 | -0.036929 | 0.1690216 | -1.231972 | 0.3133221 |
| ENSG00000077232.16 | DNAJC10 | 1.612335 | 0.529676 | 0.121929 | 0.8131911 | 0.072558 | 0.5690479 | 1.626696 | 0.000330905 | 1.253082 | 0.4954444 | -0.03505 | 0.9791298 |
| ENSG00000077235.17 | GTF3C1 | 2.505355 | 0.1336874 | -0.086919 | 0.7062745 | 0.049044 | 0.3556623 | 1.877675 | 0.01864847 | 0.166109 | 0.9958488 | -2.031296 | 0.8949489 |
| ENSG00000077254.14 | USP33 | -0.431544 | 0.8454066 | 0.364367 | 0.8496588 | 0.690872 | 0.1888762 | 0.411343 | 0.04923712 | 0.583021 | 0.9327454 | -0.437301 | 0.6885065 |
| ENSG00000077264.14 | PAK3 | -0.099809 | 0.957965 | -0.547934 | 0.6527736 | 0.276256 | 0.03711527 | 0.839946 | 0.009724107 | 1.412384 | 0.1401461 | -0.562947 | 0.8949489 |
| ENSG00000077279.16 | DCX | -1.112242 | 0.4808869 | -1.240433 | 0.02419363 | 0.560822 | 0.009443517 | 1.163957 | 0.01092796 | 0.257502 | 0.4728677 | 0.596476 | 1 |
| ENSG00000077348.8 | EXOSC5 | 0.304104 | 1 | -0.083373 | 0.8953649 | 0.226917 | 0.04532096 | 1.120264 | 0.3760083 | 0.8235 | 1 | 1.349953 | 0.6885065 |
| ENSG00000077380.15 | DYNC1I2 | -0.461913 | 0.9379425 | -0.159613 | 0.6527736 | 0.264336 | 0.3276184 | 0.603322 | 0.7621231 | -0.187945 | 2.24E-05 | 0.101388 | 0.6185816 |
| ENSG00000077684.15 | JADE1 | 1.071064 | 0.7678046 | -0.02369 | 0.8845769 | 0.793272 | 0.8200153 | 1.418261 | 4.79E-06 | 0.823941 | 0.4437504 | -2.132525 | 0.6231061 |
| ENSG00000077721.15 | UBE2A | -0.425545 | 0.9506557 | 0.215854 | 0.9100665 | 0.447086 | 0.9364284 | -0.413318 | 0.02317381 | 0.365474 | 0.6986649 | -0.568351 | 0.6885065 |
| ENSG00000078018.19 | MAP2 | -0.976889 | 0.002478701 | -0.447815 | 0.001065632 | 1.106458 | 3.39E-26 | 0.84168 | 0.001675881 | 0.282129 | 0.4193312 | -0.021925 | 0.4061635 |
| ENSG00000078177.13 | N4BP2 | 0.580783 | 0.7994453 | -0.538041 | 0.5835893 | 0.855711 | 0.004850934 | 1.219442 | 0.02310424 | 1.313585 | 0.6145671 | -1.83119 | 0.6829816 |
| ENSG00000078269.13 | SYNJ2 | 0.972789 | 1 | -0.746059 | 0.7201934 | 0.10787 | 0.1315283 | 1.065991 | 0.03793028 | 1.196283 | 0.4927467 | 1.267238 | 1 |
| ENSG00000078369.17 | GNB1 | -0.458612 | 0.6585586 | -0.173346 | 0.5110299 | -0.011939 | 0.4970176 | 0.745393 | 5.92E-06 | 0.422245 | 0.2363864 | 0.674942 | 0.8949489 |
| ENSG00000078549.14 | ADCYAP1R1 | -0.267033 | 0.6214382 | -0.241931 | 0.7607877 | -0.900006 | 0.9630279 | 2.015228 | 0.007348621 | 2.603666 | 0.2333487 | 3.989711 | 0.9147227 |
| ENSG00000078618.19 | NRD1 | 0.598668 | 0.6214382 | -0.006998 | 0.6527736 | 0.80642 | 7.97E-05 | 0.700563 | 0.3150683 | 0.33977 | 0.9361679 | -0.326248 | 0.6885065 |
| ENSG00000078674.17 | PCM1 | 0.246547 | 0.6014343 | -0.105143 | 0.6113936 | 0.715853 | 4.43E-07 | 0.425641 | 0.04638111 | 1.084816 | 0.04249027 | 1.592023 | 0.9510409 |
| ENSG00000078687.16 | TNRC6C | -2.060427 | 0.09356835 | 0.062522 | 1 | -0.235426 | 0.4321596 | 0.738319 | 0.009399882 | 1.318926 | 0.07956883 | 0.515655 | 0.8479431 |
| ENSG00000078699.21 | CBFA2T2 | 0.157318 | 1 | 0.248028 | 0.8639823 | -0.591046 | 0.8427291 | 1.468934 | 0.000695773 | 1.377101 | 0.00827723 | -3.266908 | 0.7313099 |
| ENSG00000078747.12 | ITCH | 0.047313 | 0.8867323 | -0.18089 | 0.8624047 | 1.182585 | 0.000995861 | 0.135068 | 0.9340378 | 0.537602 | 1 | 2.848496 | 0.9895571 |
| ENSG00000078804.12 | TP53INP2 | 1.678165 | 0.6214382 | -0.461269 | 0.9787499 | 0.019363 | 0.9558423 | 1.879685 | 0.002710251 | 3.105968 | 0.02745483 | -0.773963 | 0.8479431 |
| ENSG00000078808.16 | SDF4 | -0.186509 | 0.8587992 | -0.018377 | 0.4539559 | -1.879945 | 1.87E-21 | -0.000765 | 0.2410525 | 0.718996 | 0.5196843 | 0.965632 | 0.7420459 |
| ENSG00000078967.12 | UBE2D4 | 0.798218 | 0.4709738 | -0.096652 | 0.997768 | 0.133631 | 0.008142692 | 0.456629 | 0.8002793 | 0.872317 | 0.696089 | -2.148302 | 0.6939365 |
| ENSG00000079102.16 | RUNX1T1 | -1.486205 | 0.8006286 | -1.336089 | 0.7820217 | -0.051412 | 0.5290845 | -0.180415 | 0.03717626 | 1.359515 | 0.1536954 | -0.202061 | 1 |
| ENSG00000079150.17 | FKBP7 | 2.946138 | 0.714717 | 1.15126 | 0.8445455 | 0.148016 | 0.622671 | -0.637943 | 0.01524047 | -0.309971 | 0.8511053 | -1.643126 | 0.6885065 |
| ENSG00000079246.15 | XRCC5 | -0.085778 | 1 | -0.071118 | 0.5700931 | 0.791246 | 1.40E-05 | 0.608365 | 0.3634522 | 0.656682 | 0.5196843 | 0.310114 | 0.5066477 |
| ENSG00000079432.7 | CIC | 1.301175 | 0.8006286 | -0.387167 | 0.9444595 | -0.00359 | 0.9759919 | 1.186948 | 0.004269593 | 1.081731 | 0.3803925 | 0.43654 | 0.8949489 |
| ENSG00000079462.7 | PAFAH1B3 | -0.427854 | 0.6792949 | -0.302074 | 0.03619211 | -0.016318 | 0.7437308 | 0.2065 | 0.9733822 | -0.122155 | 0.06249556 | -0.424729 | 0.5733051 |
| ENSG00000080298.15 | RFX3 | 2.05256 | 0.563505 | 0.31402 | 0.9671539 | 1.113348 | 0.03540251 | 0.685762 | 0.8224647 | 2.406718 | 0.0006924 | -0.917389 | 0.6835249 |
| ENSG00000080345.17 | RIF1 | 0.432232 | 0.5687266 | -0.06141 | 0.9009117 | 0.647306 | 0.06923265 | 1.859791 | 0.000125539 | 1.105475 | 0.3387852 | -0.802086 | 0.8777121 |
| ENSG00000080371.5 | RAB21 | -0.022951 | 1 | 0.120549 | 0.9727008 | 0.353645 | 0.4382561 | 1.330994 | 0.001446231 | 1.326977 | 0.008344424 | 0.169923 | 0.6231061 |
| ENSG00000080802.18 | CNOT4 | -1.292584 | 0.714717 | -0.069366 | 0.6527736 | 0.596993 | 6.09E-09 | 1.330118 | 0.5361751 | 0.95998 | 0.9068628 | -1.973182 | 0.6885065 |
| ENSG00000080815.18 | PSEN1 | -0.734764 | 0.9503386 | -0.084815 | 0.77388 | 1.018564 | 0.01671052 | -0.060194 | 0.9279503 | 0.285804 | 0.8646421 | 1.499021 | 0.6885065 |
| ENSG00000080819.6 | CPOX | -0.720405 | 0.8587992 | -0.638211 | 0.4011918 | 0.559976 | 0.8432031 | 3.23278 | 0.02123146 | -0.411631 | 0.5768797 | -1.262059 | 0.9456372 |
| ENSG00000080824.18 | HSP90AA1 | -0.726144 | 0.08293861 | -0.316802 | 0.7160172 | 1.482039 | 4.60E-52 | 0.444439 | 0.000847916 | 0.699232 | 0.07334442 | -0.360531 | 0.8616307 |
| ENSG00000081019.13 | RSBN1 | -0.191905 | 1 | 0.740672 | 0.756107 | 0.29015 | 0.06202009 | 1.204168 | 0.013996 | 1.427097 | 0.5551535 | 2.54117 | 0.6885065 |
| ENSG00000081026.18 | MAGI3 | 1.201581 | 0.8006286 | -0.500685 | 0.4571222 | 1.229359 | 0.00240367 | 1.518472 | 0.04221781 | 0.966256 | 0.183027 | -1.119987 | 1 |
| ENSG00000081177.18 | EXD2 | -0.524838 | 0.7821237 | -0.230251 | 0.6527736 | 0.139177 | 0.4855596 | 0.690431 | 0.03838314 | 1.900787 | 0.5539249 | 0.869399 | 0.8479431 |
| ENSG00000081377.16 | CDC14B | -0.803643 | 0.9047689 | -0.341514 | 0.4002287 | 1.555752 | 0.2018778 | 0.93617 | 0.005541386 | 2.03203 | 0.006395008 | -0.108764 | 0.5066477 |
| ENSG00000081665.13 | ZNF506 | -2.829513 | 0.6214382 | -0.504278 | 0.9790409 | 0.528932 | 0.03905198 | 1.363764 | 0.01477387 | 0.560766 | 0.4863491 | 1.261586 | 0.7403741 |
| ENSG00000081870.11 | HSPB11 | -1.536176 | 0.6014343 | -0.181095 | 0.9859694 | 0.110904 | 0.02390989 | 0.52659 | 0.6244041 | 0.161859 | 0.1115558 | -1.722519 | 0.4149735 |
| ENSG00000082014.16 | SMARCD3 | 1.60408 | 0.3166974 | -0.30174 | 0.9577957 | -0.250353 | 0.3463677 | 0.267872 | 0.7830548 | 0.981267 | 0.02553981 | 1.104497 | 0.8322518 |
| ENSG00000082153.17 | BZW1 | 0.174452 | 0.6214382 | 0.519298 | 0.007219487 | 0.586896 | 0.000228539 | 0.337245 | 0.5566362 | -0.17051 | 0.006984314 | 0.060575 | 0.4081608 |
| ENSG00000082212.11 | ME2 | 0.102712 | 0.8821553 | -0.263504 | 0.7174098 | -0.333653 | 0.01994532 | 0.819855 | 0.00135868 | 1.69587 | 0.09831445 | 0.110867 | 0.6885065 |
| ENSG00000082258.12 | CCNT2 | -2.166686 | 1 | 0.117323 | 0.859711 | 0.314231 | 0.4721767 | 1.823221 | 0.03010423 | 2.519812 | 0.02150138 | 1.485104 | 0.6231583 |
| ENSG00000082458.11 | DLG3 | -0.789887 | 0.6014343 | -0.064026 | 0.7313293 | 1.895338 | 0.1996479 | 1.328933 | 0.02205499 | 1.736477 | 0.08729226 | 0.592101 | 0.9526606 |
| ENSG00000082515.16 | MRPL22 | -0.031016 | 0.8272675 | 0.296574 | 0.8703522 | -0.044685 | 0.0368713 | -0.864163 | 0.006869538 | 0.096915 | 0.1401461 | 0.337471 | 0.7020544 |
| ENSG00000082641.15 | NFE2L1 | 0.3815 | 0.9183025 | -0.040553 | 0.6527736 | -0.590131 | 0.1635361 | 0.738878 | 0.01616701 | 0.551645 | 0.7478583 | 1.50754 | 0.7845564 |
| ENSG00000082701.14 | GSK3B | -0.522998 | 0.7907283 | -0.15653 | 1 | -0.821673 | 0.3295893 | 1.17761 | 7.51E-06 | 1.206477 | 0.04616966 | -0.325679 | 0.8479431 |
| ENSG00000082898.16 | XPO1 | 0.483868 | 0.8867323 | -0.529594 | 0.04469129 | 0.247262 | 0.123838 | 0.326355 | 0.01090412 | 1.506211 | 0.003055278 | -0.003255 | 0.883407 |
| ENSG00000083097.14 | DOPEY1 | -2.071487 | 0.6214382 | 1.126477 | 0.5953732 | 1.00034 | 0.1569918 | 0.969317 | 0.03289865 | 1.285774 | 0.2452264 | -1.509789 | 0.916449 |
| ENSG00000083168.9 | KAT6A | 0.168776 | 0.737082 | -0.116597 | 0.7595553 | 0.325934 | 0.7284391 | 1.87919 | 0.00010121 | 1.282497 | 0.005649406 | 1.927469 | 0.9368176 |
| ENSG00000083223.17 | ZCCHC6 | 1.873423 | 1 | 0.060862 | 0.9889521 | -0.029343 | 0.05464236 | 0.945454 | 0.01279765 | 0.68952 | 0.5196843 | 0.133543 | 0.8479431 |
| ENSG00000083290.19 | ULK2 | -0.140453 | 1 | -0.317693 | 0.9028136 | -0.623438 | 0.3136618 | 0.265741 | 0.006916264 | 0.897498 | 0.3974318 | 1.045346 | 0.6885065 |
| ENSG00000083312.17 | TNPO1 | -0.514416 | 0.8006286 | 0.047991 | 0.8953649 | 0.935694 | 1.57E-05 | 1.156077 | 0.000321561 | 0.903039 | 0.2016786 | -1.731869 | 0.6885065 |
| ENSG00000083457.11 | ITGAE | 0.500373 | 0.6014343 | -0.158745 | 0.5284311 | -0.103617 | 0.1394359 | -0.581925 | 0.008110694 | 1.182079 | 0.9395667 | -2.011991 | 0.3532606 |
| ENSG00000083520.14 | DIS3 | 0.067491 | 0.6214382 | 0.07278 | 1 | 1.268253 | 3.14E-05 | 2.624303 | 0.1421118 | 1.770761 | 0.002293693 | -0.048397 | 1 |
| ENSG00000083635.7 | NUFIP1 | -1.839578 | 0.3254792 | -0.075319 | 0.8953649 | 1.328122 | 0.04753476 | 0.30237 | 0.9383784 | -0.119819 | 0.4638378 | -2.880207 | 0.9529516 |
| ENSG00000083642.18 | PDS5B | 0.573956 | 0.5711449 | -0.268421 | 1 | 0.892171 | 0.05361372 | 0.998541 | 0.005022021 | 0.656739 | 0.4099563 | -0.864649 | 0.6885065 |
| ENSG00000083799.17 | CYLD | -1.26064 | 1 | 0.308014 | 0.9188449 | 0.107206 | 0.001947133 | 1.488381 | 0.08709445 | 1.297361 | 0.3652046 | 1.355154 | 0.8917781 |
| ENSG00000083845.8 | RPS5 | -0.208774 | 0.9565296 | 0.007373 | 0.6677578 | 0.144615 | 0.5290845 | -0.256862 | 2.30E-07 | -0.181307 | 1.62E-05 | 0.131554 | 0.6784855 |
| ENSG00000083896.12 | YTHDC1 | -2.980726 | 1 | -0.101323 | 0.6565919 | 0.853946 | 0.000166866 | 0.760097 | 0.3395199 | 0.647874 | 0.4620424 | 0.323863 | 0.9576568 |
| ENSG00000084090.13 | STARD7 | 0.33094 | 0.4464112 | 0.049319 | 0.9249123 | 0.606016 | 0.7437308 | 1.47781 | 0.000960359 | 0.595295 | 0.963643 | -0.39521 | 0.6517664 |
| ENSG00000084093.15 | REST | -1.557865 | 0.7693197 | -0.196289 | 0.5953732 | 1.100189 | 0.0064626 | 1.507166 | 0.6720782 | 1.790857 | 0.8021966 | 1.233172 | 0.6885065 |
| ENSG00000084112.14 | SSH1 | -3.054644 | 0.3946735 | 0.464694 | 0.6782487 | -0.406281 | 0.9719251 | 1.812177 | 0.001191898 | 2.435526 | 0.08724925 | 0.874889 | 0.8868882 |
| ENSG00000084207.15 | GSTP1 | -0.760387 | 9.45E-05 | -0.084128 | 0.5953732 | -0.117805 | 0.001006486 | 0.13263 | 0.7157474 | -0.481643 | 1.73E-06 | -0.136056 | 0.01248611 |
| ENSG00000084234.16 | APLP2 | 0.498817 | 0.185057 | -0.018379 | 0.8953825 | 0.497464 | 9.02E-06 | 0.521607 | 0.4919368 | 0.039494 | 0.3013263 | 0.37857 | 0.9233554 |
| ENSG00000084444.13 | FAM234B | 1.891332 | 0.2654079 | -0.578912 | 0.6527736 | 0.708161 | 0.9346851 | 2.905197 | 0.002693974 | -0.003691 | 0.7987532 | 0.549479 | 0.7963477 |
| ENSG00000084463.7 | WBP11 | -0.144572 | 0.958326 | -0.066648 | 0.7118133 | 0.342903 | 0.3908542 | 0.979214 | 0.03607211 | 0.459638 | 0.2961378 | -0.403665 | 0.9380942 |
| ENSG00000084623.11 | EIF3I | 1.039892 | 0.001177428 | -0.184984 | 0.6925308 | 0.310338 | 0.1214755 | -0.283859 | 0.000470696 | -0.264158 | 0.001899548 | -0.120617 | 0.6185816 |
| ENSG00000084628.9 | NKAIN1 | 0.038105 | 0.7736255 | -1.569207 | 0.6527736 | 0.70433 | 0.5841679 | 0.281213 | 0.04055961 | 1.011854 | 0.5196843 | 0.088252 | 0.6979803 |
| ENSG00000084652.15 | TXLNA | 0.101704 | 0.7708086 | 0.071544 | 0.6527736 | 0.103868 | 1 | 1.110392 | 6.15E-05 | 1.644538 | 0.02392971 | -0.491105 | 0.9665677 |
| ENSG00000084676.15 | NCOA1 | -0.998874 | 0.5139066 | -0.015578 | 0.9056349 | -0.079744 | 0.04009954 | 0.401051 | 0.02090792 | 1.745302 | 0.03068444 | -0.106995 | 0.6829816 |
| ENSG00000084710.13 | EFR3B | -0.452845 | 0.6690771 | -0.093774 | 0.7383998 | -0.929264 | 0.005066113 | 1.349295 | 0.003301255 | 1.332977 | 0.3339985 | 1.22761 | 0.6885065 |
| ENSG00000084754.10 | HADHA | -0.187429 | 0.6014343 | 0.136305 | 0.8703522 | 0.311642 | 0.09396285 | -0.049137 | 0.04207541 | -0.734254 | 0.000616252 | 0.279539 | 0.6885065 |
| ENSG00000084774.13 | CAD | -0.621158 | 0.8245439 | -1.1776 | 0.1516841 | -1.25827 | 0.528724 | 2.93072 | 0.04318341 | 1.663895 | 0.4278631 | -0.128733 | 0.8479431 |
| ENSG00000085063.14 | CD59 | 0.343212 | 0.000286034 | -0.2764 | 0.1385302 | -0.271995 | 0.1450103 | 0.09182 | 0.9593911 | -0.345073 | 0.07383985 | 0.153945 | 0.6829816 |
| ENSG00000085224.20 | ATRX | 0.128568 | 0.6214382 | -0.325868 | 0.05090264 | 1.01445 | 4.87E-14 | 0.625924 | 4.54E-05 | -0.047988 | 0.6593794 | 0.357612 | 0.6517664 |
| ENSG00000085274.15 | MYNN | -1.671883 | 0.6214382 | 0.177479 | 0.9674023 | 0.916 | 0.002422785 | 0.097839 | 0.9802055 | -0.542765 | 0.290602 | -1.00391 | 0.7623154 |
| ENSG00000085433.15 | WDR47 | -0.835239 | 0.8587992 | 0.122096 | 0.9931449 | 0.783615 | 0.000367975 | 1.492518 | 0.1385281 | 1.499957 | 0.101823 | -0.099243 | 0.9786489 |
| ENSG00000085644.13 | ZNF213 | -0.239887 | 1 | -0.972095 | 0.8703522 | 0.917132 | 0.9687194 | 1.383833 | 0.211072 | 1.710926 | 0.01448391 | -2.002056 | 0.6829816 |
| ENSG00000085719.11 | CPNE3 | 0.633213 | 0.6574918 | 0.36631 | 0.8272197 | 0.824718 | 0.001745983 | 0.49062 | 0.6695085 | -0.136208 | 0.0957903 | -0.685448 | 0.5344421 |
| ENSG00000085721.12 | RRN3 | 0.800154 | 0.6214382 | 0.5988 | 0.9490229 | -0.115793 | 1 | 0.799082 | 0.001300776 | 1.021783 | 0.004672895 | -0.743615 | 0.9526606 |
| ENSG00000085832.16 | EPS15 | 3.09974 | 1 | 0.542521 | 0.9871396 | 0.973876 | 0.97505 | 2.051487 | 0.008357595 | 1.132727 | 0.7233645 | 1.155864 | 0.1954301 |
| ENSG00000085872.14 | CHERP | 0.381141 | 0.8264007 | -0.261889 | 0.470449 | -0.220806 | 0.4064599 | 0.883363 | 0.1866393 | 0.396877 | 0.007461499 | -1.584759 | 1 |
| ENSG00000086061.15 | DNAJA1 | 0.273308 | 0.07547825 | 0.012567 | 0.978688 | 0.361812 | 0.2543441 | 0.740418 | 4.19E-05 | 0.244326 | 0.7243765 | -0.225845 | 0.2305604 |
| ENSG00000086102.18 | NFX1 | -0.480294 | 0.4764974 | -0.927495 | 0.3202201 | -0.179087 | 0.9985944 | 1.379707 | 0.002864164 | 0.691115 | 0.9000411 | -1.233364 | 0.6885065 |
| ENSG00000086200.16 | IPO11 | 1.219276 | 0.9047689 | 0.443671 | 0.9856559 | -0.357339 | 0.9259471 | 0.81192 | 4.26E-05 | 1.044483 | 0.04488428 | 0.158579 | 1 |
| ENSG00000086475.14 | SEPHS1 | 0.740701 | 0.5711449 | -0.149003 | 0.6827803 | 1.441827 | 0.776456 | 0.490876 | 0.01015341 | 0.929389 | 0.5502401 | 1.862772 | 0.908391 |
| ENSG00000086598.10 | TMED2 | 0.994305 | 0.006263525 | -0.019548 | 0.5953732 | 0.229427 | 0.5654652 | 0.692799 | 0.01868866 | 0.448318 | 0.183027 | -1.263023 | 0.6829816 |
| ENSG00000086730.16 | LAT2 | -0.586727 | 0.8006286 | 0.339684 | 0.8703522 | 0.789668 | 0.7354314 | 4.204409 | 0.08787835 | 4.006627 | 0.02119726 | 1.617697 | 0.7767294 |
| ENSG00000086758.15 | HUWE1 | 0.780932 | 0.8787256 | -0.327527 | 0.2863181 | 0.101071 | 0.7437308 | 1.416289 | 7.70E-09 | 1.719573 | 0.00139047 | -0.366525 | 0.6829816 |
| ENSG00000086848.14 | ALG9 | 1.856761 | 0.714717 | -0.031146 | 0.9545506 | 3.920798 | 0.03552749 | 0.958032 | 0.1629094 | 3.055184 | 0.4480788 | 2.651342 | 0.8479431 |
| ENSG00000087077.11 | TRIP6 | -3.895415 | 0.4764974 | 0.028194 | 0.6048659 | -0.401092 | 0.03569965 | 0.357579 | 0.169916 | 0.810721 | 0.339022 | -0.071772 | 0.6993068 |
| ENSG00000087086.13 | FTL | -0.423893 | 0.6214382 | 0.284289 | 2.49E-16 | -0.302362 | 3.41E-05 | 0.24722 | 0.06128956 | -0.185671 | 0.008273711 | 0.736234 | 0.01867515 |
| ENSG00000087087.18 | SRRT | 0.211579 | 0.6136871 | 0.271635 | 0.5395105 | 0.08783 | 0.02483293 | 0.636064 | 0.1866915 | 0.437158 | 0.6598924 | -0.140423 | 0.3532606 |
| ENSG00000087088.19 | BAX | -2.499544 | 0.036921 | 0.4621 | 0.5010139 | -0.633928 | 0.000338866 | 0.900629 | 0.2346996 | 0.969765 | 1 | 1.129548 | 1 |
| ENSG00000087095.12 | NLK | 0.355362 | 0.6792949 | -0.193028 | 0.7806598 | 0.975016 | 0.5095143 | 1.825093 | 0.0004537 | 2.146999 | 0.4250235 | -1.072683 | 0.7419782 |
| ENSG00000087111.20 | PIGS | -0.376981 | 1 | 0.563467 | 0.9068973 | 0.672098 | 0.260758 | 0.37884 | 0.04148071 | 0.191404 | 0.7419008 | 2.37064 | 0.8479431 |
| ENSG00000087152.15 | ATXN7L3 | 0.709917 | 0.7683685 | 0.300717 | 0.9068973 | -0.917438 | 0.3389032 | 1.853081 | 0.01572337 | 1.211562 | 0.3974318 | -1.058862 | 0.9576568 |
| ENSG00000087245.12 | MMP2 | -1.060317 | 1 | -0.431178 | 0.8217443 | 0.112922 | 0.761562 | 1.163256 | 0.01960715 | -1.117321 | 0.5422031 | -0.557794 | 0.6967502 |
| ENSG00000087250.8 | MT3 | 0.191586 | 0.6704454 | -0.386639 | 0.3610905 | -0.391691 | 0.5290845 | -0.441813 | 9.54E-05 | 0.300317 | 0.1760064 | 0.22225 | 1 |
| ENSG00000087258.13 | GNAO1 | -0.656529 | 0.7138283 | -0.364072 | 0.8703522 | 1.392465 | 0.3288909 | -0.374111 | 0.07369839 | 1.152078 | 0.04521787 | -1.17939 | 0.9483075 |
| ENSG00000087274.16 | ADD1 | -0.006905 | 0.9305688 | 0.247817 | 0.8013363 | 0.261694 | 0.3695136 | 1.388592 | 0.02477329 | 1.651384 | 0.02061222 | -0.593 | 0.8479431 |
| ENSG00000087365.14 | SF3B2 | -0.61619 | 0.6214382 | -0.217406 | 0.7166706 | 0.357445 | 0.000674155 | 0.062688 | 0.2106491 | -0.24808 | 0.05319344 | 0.12503 | 0.8894718 |
| ENSG00000087460.23 | GNAS | -0.471567 | 0.4987686 | -0.418852 | 0.000832664 | 0.253845 | 0.01881054 | 0.853511 | 5.37E-10 | 0.999332 | 3.50E-10 | -0.239685 | 0.3946031 |
| ENSG00000087884.14 | AAMDC | 0.17366 | 0.7431767 | 0.182885 | 0.6527736 | -0.824684 | 0.0115343 | -1.127054 | 0.1080617 | 1.680779 | 0.1253902 | -0.923158 | 0.5344421 |
| ENSG00000088205.12 | DDX18 | -0.192255 | 0.9348915 | 0.34395 | 0.6527736 | 0.481528 | 0.00354589 | 0.735568 | 0.880362 | 0.338454 | 0.3974318 | -0.893957 | 0.6885065 |
| ENSG00000088247.15 | KHSRP | 0.143878 | 0.3946735 | -0.003418 | 0.9042506 | -0.202323 | 0.9166914 | 0.813751 | 0.001430378 | 1.36732 | 0.000281589 | 0.903973 | 0.8479431 |
| ENSG00000088256.8 | GNA11 | -0.96304 | 0.6214382 | 0.003428 | 0.704767 | 1.059872 | 0.3682205 | 1.346943 | 0.01284365 | 1.551004 | 0.01606565 | -1.118002 | 0.9539924 |
| ENSG00000088367.20 | EPB41L1 | 0.431425 | 0.8733861 | -0.843691 | 0.592336 | -0.290665 | 0.7235812 | 1.193788 | 0.005749726 | 0.085452 | 0.1864001 | -0.267872 | 0.501643 |
| ENSG00000088448.14 | ANKRD10 | -0.641909 | 0.4605085 | -0.479868 | 0.4264647 | -0.008421 | 0.5832822 | 0.999433 | 0.03967699 | 1.42466 | 0.001791395 | 0.886294 | 0.6885065 |
| ENSG00000088766.11 | CRLS1 | -0.693564 | 0.563505 | -0.262124 | 0.5953732 | -0.364946 | 0.0815126 | 0.820385 | 0.004484026 | 1.571922 | 0.2327404 | 0.177091 | 0.9264343 |
| ENSG00000088812.17 | ATRN | 0.531923 | 0.7736255 | -0.681744 | 0.9821403 | -0.277105 | 0.7437308 | 2.187562 | 0.000851278 | 2.080975 | 0.2307297 | 0.15636 | 0.6829816 |
| ENSG00000088832.14 | FKBP1A | 0.002651 | 0.6214382 | 0.083076 | 0.9974455 | -0.491635 | 0.01564226 | -0.050092 | 0.4016368 | 0.638139 | 0.6879887 | 0.326479 | 0.8384921 |
| ENSG00000088854.12 | C20orf194 | -1.464749 | 0.6169802 | 0.34709 | 0.6527736 | -0.751575 | 0.0618094 | 1.913344 | 0.008452094 | 1.450322 | 0.4594462 | 1.160183 | 0.789721 |
| ENSG00000088881.20 | EBF4 | -3.390246 | 0.563505 | -0.071436 | 0.6527736 | 0.347587 | 0.6734664 | 2.800934 | 0.003599989 | 1.591304 | 0.0597997 | 1.353993 | 0.8931285 |
| ENSG00000088930.7 | XRN2 | -1.056964 | 0.6214382 | -0.024238 | 0.6670949 | 0.84383 | 1.58E-07 | 0.363789 | 0.5869461 | -0.206379 | 0.01406336 | -0.541911 | 0.4578379 |
| ENSG00000088986.10 | DYNLL1 | -1.173867 | 0.7180972 | 0.262074 | 0.06493677 | -0.165004 | 9.19E-06 | -0.17143 | 0.001290919 | 0.105013 | 0.2363864 | 0.121342 | 0.5331811 |
| ENSG00000089009.15 | RPL6 | -0.764004 | 9.25E-06 | 0.216852 | 0.4812186 | 0.632781 | 8.28E-21 | 0.053985 | 0.450241 | -0.271777 | 2.32E-05 | 0.119903 | 0.5344421 |
| ENSG00000089048.14 | ESF1 | -0.678214 | 0.529676 | -0.480666 | 0.8703522 | 0.882087 | 2.56E-05 | 0.446541 | 0.3357428 | 0.176492 | 0.4582178 | -1.366328 | 0.7809378 |
| ENSG00000089057.14 | SLC23A2 | 1.490179 | 0.6214382 | 1.128129 | 0.1712543 | 0.237933 | 0.9262117 | 0.329511 | 0.005905424 | -2.008773 | 0.5196843 | 0.868369 | 1 |
| ENSG00000089094.16 | KDM2B | 0.878496 | 0.8575361 | -0.581833 | 0.8272197 | -0.38226 | 0.5290845 | 0.850881 | 0.2335397 | 1.523414 | 0.002548226 | 0.902388 | 0.6885065 |
| ENSG00000089123.15 | TASP1 | -2.815409 | 0.3946735 | 1.443751 | 0.4177445 | 0.490029 | 0.8218004 | 1.080467 | 0.01631219 | -0.040015 | 1 | -0.338552 | 0.8949489 |
| ENSG00000089154.10 | GCN1 | -0.529967 | 0.6214382 | -0.685453 | 0.1785916 | -1.040746 | 0.8979978 | 1.172561 | 0.02079597 | 1.139564 | 0.2880317 | 0.139187 | 0.8479431 |
| ENSG00000089157.15 | RPLP0 | 0.369825 | 1.66E-08 | 0.170751 | 0.2562412 | 0.272326 | 0.1041316 | -0.32391 | 1.89E-05 | 0.029654 | 0.3734275 | -0.201134 | 0.3746505 |
| ENSG00000089199.9 | CHGB | -0.116243 | 0.6353584 | -0.569408 | 1 | -0.893438 | 0.8057978 | 1.310716 | 0.004685378 | 1.327439 | 0.6857008 | -0.20351 | 0.8479431 |
| ENSG00000089220.4 | PEBP1 | -1.677013 | 0.1607609 | -0.005435 | 0.6527736 | 0.219675 | 0.9493498 | 0.390278 | 0.1273493 | -0.09098 | 0.001613143 | -0.354394 | 0.2635027 |
| ENSG00000089280.18 | FUS | -1.065639 | 1.11E-07 | -0.062632 | 0.7881421 | 0.151165 | 0.7212093 | 0.476704 | 0.01252251 | 0.357981 | 0.7734487 | 0.282286 | 0.4833616 |
| ENSG00000089597.16 | GANAB | 2.421723 | 0.02452349 | 0.163355 | 0.6527736 | -0.09816 | 0.1476253 | 1.621348 | 1.75E-05 | 0.96386 | 0.02339438 | -1.260123 | 0.5417688 |
| ENSG00000089682.16 | RBM41 | -3.254329 | 0.6214382 | 0.642909 | 0.9683845 | 0.181991 | 0.4791605 | 4.867845 | 0.001392502 | 1.628867 | 0.919153 | 1.236404 | 0.6885065 |
| ENSG00000089693.10 | MLF2 | 0.190583 | 0.1835496 | 0.44532 | 0.5375948 | -0.090116 | 0.03274039 | 0.30878 | 1 | 0.845208 | 0.8021966 | 0.244161 | 0.6885065 |
| ENSG00000089916.17 | GPATCH2L | -1.464167 | 0.2643306 | 0.221316 | 0.8263112 | 1.420597 | 0.09686449 | 0.437244 | 0.02571986 | 0.502767 | 0.3987518 | 2.972883 | 0.2256183 |
| ENSG00000090013.9 | BLVRB | 2.511963 | 0.6214382 | 0.76876 | 0.3387885 | 0.702801 | 0.05450271 | -0.562466 | 0.002463418 | 1.509049 | 0.8956005 | 0.370954 | 0.8949489 |
| ENSG00000090060.17 | PAPOLA | 0.007261 | 0.7383844 | -0.165747 | 0.4580322 | 0.659264 | 7.20E-05 | 0.225188 | 0.900835 | 0.50833 | 0.602879 | -1.011972 | 1 |
| ENSG00000090097.20 | PCBP4 | -0.330095 | 1 | 0.019895 | 0.7364864 | -0.137731 | 0.08191509 | 1.279249 | 0.01168837 | 0.264938 | 0.5118438 | -1.367891 | 0.6829816 |
| ENSG00000090266.12 | NDUFB2 | 0.634547 | 9.35E-06 | -0.170047 | 0.3580999 | -0.556436 | 1.45E-12 | -0.046952 | 0.003473137 | -0.286956 | 0.000567037 | -0.062816 | 0.05741837 |
| ENSG00000090273.13 | NUDC | -0.891582 | 0.529676 | -0.070072 | 0.7062105 | 0.31043 | 0.8884334 | 0.210909 | 0.01372311 | 0.034424 | 0.03752142 | -0.105284 | 0.2809502 |
| ENSG00000090372.14 | STRN4 | -1.294362 | 0.529676 | -0.110323 | 0.8703522 | -0.392244 | 0.5449454 | 1.957363 | 0.05921221 | 1.610011 | 0.001955021 | 0.929987 | 0.6939365 |
| ENSG00000090520.10 | DNAJB11 | 0.838524 | 0.1364817 | 0.130316 | 0.6526104 | -0.298302 | 0.006056379 | 0.496969 | 0.7890908 | 0.357019 | 0.9210594 | -0.313729 | 0.5246903 |
| ENSG00000090539.15 | CHRD | -1.932222 | 0.8006286 | -1.273833 | 0.5953732 | -1.742049 | 0.2348326 | 1.218483 | 0.6575769 | 1.372223 | 0.5196843 |  |  |
| ENSG00000090565.15 | RAB11FIP3 | -0.000976 | 0.7138283 | 0.274764 | 0.6527736 | 0.985911 | 0.8870699 | 1.016227 | 0.008527215 | 3.816687 | 0.000245861 | 0.086478 | 0.9606764 |
| ENSG00000090581.9 | GNPTG | -0.970835 | 0.6214382 | 0.237127 | 0.8849137 | -0.843507 | 0.001639474 | -0.291177 | 0.4919368 | 0.769578 | 0.480965 | -0.651837 | 0.8949489 |
| ENSG00000090612.20 | ZNF268 | -0.158815 | 0.6214382 | 0.147897 | 0.9839225 | -0.01694 | 0.9304971 | 1.131649 | 0.01292204 | 0.72175 | 0.1412962 | 1.264107 | 1 |
| ENSG00000090615.12 | GOLGA3 | -0.01076 | 0.6014343 | -0.16647 | 0.6527736 | 0.300039 | 0.1787542 | 1.634174 | 2.75E-05 | 1.352109 | 0.000288395 | -0.320593 | 0.705843 |
| ENSG00000090621.13 | PABPC4 | -0.190428 | 0.539139 | -0.127857 | 0.6527736 | 0.218993 | 0.8832504 | 0.927295 | 1.70E-05 | 0.92427 | 0.3974318 | -0.215689 | 0.8331363 |
| ENSG00000090686.15 | USP48 | -1.582722 | 0.6214382 | -0.254301 | 0.5953732 | 0.930939 | 0.3798808 | 2.092083 | 0.000586064 | 0.48469 | 0.005711494 | -1.119355 | 1 |
| ENSG00000090863.11 | GLG1 | 0.499718 | 0.8016328 | 0.178983 | 0.8263112 | 0.823191 | 0.0561467 | 1.212143 | 2.38E-05 | 0.775009 | 0.01345489 | -0.203845 | 0.7191877 |
| ENSG00000090905.17 | TNRC6A | -0.340518 | 0.529676 | 0.112325 | 0.7096117 | 0.969337 | 0.01755287 | 1.015693 | 0.01034389 | 2.010556 | 0.000215011 | -0.606759 | 0.9668212 |
| ENSG00000090924.14 | PLEKHG2 | 0.806837 | 1 | -0.951336 | 0.4691942 | -0.563414 | 0.2707921 | 2.54537 | 7.38E-06 | 1.817513 | 0.008780665 | 0.049454 | 0.8479431 |
| ENSG00000090932.10 | DLL3 | 0.238914 | 0.6214382 | -1.253691 | 0.04760656 | -0.018502 | 0.6625336 | 0.747006 | 0.3317899 | 0.725996 | 0.008907229 | 0.894016 | 0.8949489 |
| ENSG00000090975.12 | PITPNM2 | -0.065219 | 1 | 0.087782 | 0.9052551 | -0.712154 | 0.5290845 | 2.289251 | 0.004591457 | 3.195987 | 0.08655532 | -2.495921 | 0.9554184 |
| ENSG00000091039.16 | OSBPL8 | 0.145271 | 0.789972 | -0.25168 | 0.7697622 | 0.929911 | 0.000466473 | 0.636621 | 0.4175016 | 0.475464 | 0.5196843 | 1.283796 | 0.9824843 |
| ENSG00000091129.19 | NRCAM | -0.34905 | 0.8587992 | -0.316189 | 0.007339969 | 0.272011 | 0.1525822 | 0.625669 | 0.000752405 | 0.680411 | 0.7055534 | -0.212859 | 0.5344421 |
| ENSG00000091136.13 | LAMB1 | -4.508138 | 0.8006286 | 0.052013 | 0.6527736 | -0.438495 | 1 | 1.730285 | 4.07E-06 | -0.010633 | 0.953846 | 0.272878 | 0.9148375 |
| ENSG00000091140.12 | DLD | 1.164795 | 0.008781458 | 0.276511 | 0.8953649 | 0.095637 | 0.7530098 | 1.045935 | 0.0323983 | -0.013487 | 0.7136604 | 0.004264 | 0.8347311 |
| ENSG00000091157.13 | WDR7 | 0.641665 | 0.9183025 | 1.029988 | 0.6514723 | 2.947984 | 0.142087 | -0.089218 | 0.40476 | 2.543654 | 0.01313164 | -3.818464 | 1 |
| ENSG00000091164.12 | TXNL1 | -0.318537 | 0.9365286 | -0.291963 | 0.6818241 | 0.085682 | 0.5306219 | 0.935089 | 0.3647127 | -0.359488 | 0.004292528 | -1.061936 | 0.2093204 |
| ENSG00000091409.14 | ITGA6 | -0.181242 | 1 | -0.057115 | 0.6158356 | 0.283616 | 0.5290845 | 0.966352 | 0.01143362 | 1.091636 | 1 | 4.597642 | 0.0712442 |
| ENSG00000091527.15 | CDV3 | -0.840593 | 0.4764974 | 0.398679 | 0.4011918 | 0.559489 | 0.5290845 | 1.186604 | 1.98E-07 | 1.109542 | 0.09136799 | -0.553625 | 0.6829816 |
| ENSG00000091542.8 | ALKBH5 | 0.968371 | 0.3619888 | 0.191866 | 0.8350684 | 0.125858 | 0.5290845 | 1.227796 | 0.001044563 | 0.644012 | 0.3539308 | -0.170794 | 1 |
| ENSG00000091844.7 | RGS17 | 0.299882 | 0.6562396 | -0.413872 | 0.08794836 | 1.300429 | 0.000688818 | 0.729556 | 0.001676535 | 1.865197 | 0.3988814 | -0.193072 | 0.9617345 |
| ENSG00000091986.15 | CCDC80 | -1.517508 | 0.01669535 | 0.008626 | 0.6527736 | -0.033895 | 0.2017653 | 1.166513 | 0.4919368 | 0.136844 | 0.1196747 | 0.488535 | 0.6885065 |
| ENSG00000092010.14 | PSME1 | -0.465032 | 0.9506557 | 0.354815 | 0.8263112 | -0.477895 | 1.07E-05 | 0.01543 | 2.54E-05 | -0.322441 | 0.0507771 | -0.065052 | 0.6885065 |
| ENSG00000092051.16 | JPH4 | -0.702438 | 0.6214382 | 0.081433 | 0.9104976 | 0.604226 | 0.5290845 | 1.154356 | 0.672858 | 2.936975 | 0.007406492 | 0.123391 | 0.9174022 |
| ENSG00000092108.20 | SCFD1 | -0.001398 | 1 | -0.231283 | 0.4862625 | 0.98099 | 0.02037708 | 0.963054 | 0.3467508 | 0.585671 | 0.9501799 | 1.429493 | 0.6885065 |
| ENSG00000092199.17 | HNRNPC | -0.515407 | 0.05414451 | 0.066073 | 0.8703522 | 0.372899 | 5.87E-05 | 0.213089 | 0.4232343 | 0.084335 | 0.1199742 | -0.116829 | 0.5344421 |
| ENSG00000092201.9 | SUPT16H | 0.162132 | 0.6214382 | 0.162638 | 0.6527736 | 0.903966 | 0.000112326 | 0.786063 | 0.4828132 | -0.250221 | 0.6093143 | -0.489123 | 0.5344421 |
| ENSG00000092421.16 | SEMA6A | -1.855708 | 0.714717 | -0.886999 | 0.004581147 | 0.764844 | 0.4089878 | 0.071317 | 0.3315206 | 1.370784 | 0.5196843 | 2.555588 | 0.6261342 |
| ENSG00000092439.13 | TRPM7 | 0.4643 | 0.8477948 | -0.14134 | 0.7695049 | 1.179261 | 0.04334407 | 2.003354 | 0.01797105 | 0.371338 | 0.01285443 | 0.224815 | 0.8325085 |
| ENSG00000092445.11 | TYRO3 | 1.012185 | 0.6214382 | -1.024618 | 0.5953732 | 0.241387 | 0.7602311 | -0.098066 | 0.027258 | 1.467281 | 0.1380907 | -0.959439 | 0.9545867 |
| ENSG00000092621.11 | PHGDH | -0.750277 | 0.529676 | 0.426505 | 0.7869835 | 0.872223 | 0.001213578 | -0.430302 | 0.2492629 | -0.124559 | 1 | 0.431018 | 0.6885065 |
| ENSG00000092820.17 | EZR | -1.022171 | 0.8006286 | 0.248991 | 0.8263112 | -0.289246 | 0.3054882 | 1.390941 | 0.000251729 | 0.858714 | 0.9676088 | 1.813909 | 0.5733051 |
| ENSG00000092841.18 | MYL6 | -0.271102 | 0.6168416 | 0.204348 | 0.2417791 | -0.471254 | 8.79E-12 | 0.213486 | 0.9680918 | -0.266108 | 1.29E-06 | 0.12768 | 0.8479431 |
| ENSG00000092847.11 | AGO1 | -0.269666 | 0.8450993 | 0.082189 | 0.8132389 | 1.427326 | 0.04338503 | 0.68388 | 0.000191463 | 0.076294 | 0.9175623 | 0.163501 | 0.9590434 |
| ENSG00000092871.16 | RFFL | -1.212669 | 0.5711449 | -0.378834 | 0.5953732 | 0.241489 | 0.8095182 | 2.393848 | 0.00610922 | -0.036376 | 0.3271351 | 0.057821 | 0.9545867 |
| ENSG00000093000.18 | NUP50 | -1.282005 | 0.2998756 | -0.453159 | 0.3616583 | 0.635437 | 0.2996677 | 0.834999 | 0.001368015 | 0.961 | 0.8015013 | 0.661811 | 0.6984493 |
| ENSG00000093010.11 | COMT | 0.267562 | 0.6836146 | 0.405149 | 0.6527736 | -0.426199 | 0.001101424 | -0.328077 | 0.07264986 | 0.401761 | 0.1610264 | -0.191716 | 0.6885065 |
| ENSG00000094631.18 | HDAC6 | 2.221686 | 0.6014343 | 0.988923 | 0.8678408 | 0.21435 | 0.9372848 | 1.525981 | 0.08529855 | 2.300004 | 0.04012853 | 1.040372 | 0.6885065 |
| ENSG00000094880.10 | CDC23 | 1.493026 | 0.8404327 | 0.692577 | 0.6015027 | 0.764147 | 0.2239925 | -0.787337 | 0.02163847 | 0.828678 | 0.6096491 | 0.928949 | 0.8347311 |
| ENSG00000094916.13 | CBX5 | 0.032926 | 0.8006286 | -0.398784 | 0.1245072 | 0.878975 | 4.54E-07 | 0.541221 | 0.3472527 | 0.371148 | 0.9245349 | -0.341317 | 0.9147227 |
| ENSG00000094975.13 | SUCO | -0.034172 | 0.714717 | 0.388632 | 0.6527736 | 0.492911 | 0.007004779 | 1.299123 | 0.02918812 | 0.726262 | 0.5196843 | 0.184379 | 0.8759466 |
| ENSG00000095139.13 | ARCN1 | -1.55006 | 0.8142959 | -0.096359 | 0.6527736 | 0.629567 | 1.46E-05 | 0.603942 | 0.2914611 | 1.043191 | 0.1144644 | 0.376371 | 0.8949489 |
| ENSG00000095321.16 | CRAT | -0.312346 | 0.6214382 | -0.61504 | 0.04105643 | -0.473233 | 0.8100781 | 0.223363 | 0.9535748 | 1.53169 | 0.33052 | -1.374762 | 0.6829816 |
| ENSG00000095380.10 | NANS | -0.901533 | 0.6457414 | -0.060918 | 0.9227684 | 0.172587 | 0.7437308 | -0.541425 | 5.28E-05 | -0.283005 | 0.03956112 | 0.577378 | 0.6885065 |
| ENSG00000095637.21 | SORBS1 | 0.710368 | 0.543979 | -0.838381 | 0.1053945 | -0.703629 | 0.8691767 | 0.717334 | 0.01394664 | 1.681477 | 0.3974318 | 1.091373 | 0.9835231 |
| ENSG00000095752.6 | IL11 | -0.529197 | 0.4764974 | 0.585669 | 0.6527736 | -0.914074 | 0.00175392 | 1.14063 | 0.003399818 | 3.824473 | 0.6806131 | 1.561937 | 0.9214164 |
| ENSG00000095787.21 | WAC | -0.36498 | 0.7750198 | 0.288782 | 1 | 0.639469 | 0.000362535 | 0.136133 | 0.1133498 | 1.243756 | 0.1633949 | -1.856275 | 0.5344421 |
| ENSG00000095906.16 | NUBP2 | 0.509503 | 0.6214382 | -0.098815 | 0.9930962 | 0.211816 | 0.3841497 | 0.022001 | 0.6812015 | -0.372805 | 0.03321316 | 0.02923 | 0.9663203 |
| ENSG00000095951.16 | HIVEP1 | 4.361811 | 0.714717 | 0.387921 | 1 | 1.367138 | 0.02746102 | 1.112373 | 0.6241661 | 0.147186 | 0.1450246 | 2.602697 | 0.9539154 |
| ENSG00000096080.11 | MRPS18A | -0.461016 | 0.8856407 | -0.476021 | 0.6615455 | -0.720547 | 0.01553581 | -0.088655 | 1 | 0.406096 | 0.5196843 | -0.092799 | 0.7099033 |
| ENSG00000096384.19 | HSP90AB1 | 0.34884 | 3.65E-07 | -0.107938 | 0.6048501 | 0.874088 | 8.18E-53 | 0.61746 | 7.59E-05 | 0.43202 | 0.8008714 | 0.267654 | 0.6879794 |
| ENSG00000096401.7 | CDC5L | 0.386561 | 0.5149952 | -0.138246 | 0.8746358 | 1.09445 | 8.26E-09 | 0.1335 | 0.8477276 | -0.415779 | 0.05112503 | 0.918251 | 0.7972419 |
| ENSG00000096746.17 | HNRNPH3 | -3.700164 | 0.000333395 | -0.364105 | 0.5240506 | 0.276199 | 0.3095404 | 0.389615 | 0.8728987 | 0.895118 | 0.03056663 | -0.462118 | 0.5344421 |
| ENSG00000097007.17 | ABL1 | -0.941819 | 0.6014343 | -0.035515 | 0.6527736 | -0.503544 | 0.675525 | 1.923405 | 0.02205499 | 0.673319 | 0.5524766 | 0.646548 | 1 |
| ENSG00000097021.19 | ACOT7 | -1.202553 | 0.8958015 | -0.427223 | 0.9834088 | -0.252746 | 0.5290845 | 1.034861 | 4.64E-05 | 0.723366 | 0.2301528 | 0.085216 | 0.7219874 |
| ENSG00000099194.5 | SCD | 0.176005 | 0.6014343 | 0.224237 | 0.9969553 | 0.459492 | 0.4639633 | 0.890945 | 0.03048442 | 1.640061 | 0.3994316 | 0.531463 | 0.7812339 |
| ENSG00000099250.17 | NRP1 | -1.04945 | 0.8587992 | -0.431495 | 0.6527736 | -0.714121 | 0.182655 | 2.985148 | 3.39E-13 | 2.039765 | 0.09051777 | 1.182149 | 0.9351352 |
| ENSG00000099331.13 | MYO9B | 1.187224 | 0.7703009 | -0.002311 | 1 | -0.25453 | 0.7463765 | 0.873048 | 0.001516486 | 2.129508 | 0.004636219 | 2.047594 | 0.3532606 |
| ENSG00000099385.11 | BCL7C | -0.398266 | 0.6214382 | 0.602329 | 0.3242827 | -0.511472 | 0.0205573 | -0.897446 | 0.4919368 | 0.434018 | 0.7281781 | -0.980183 | 0.6861518 |
| ENSG00000099622.13 | CIRBP | -0.732131 | 0.04134193 | -0.032373 | 0.9690346 | 0.489361 | 0.05027396 | -0.144493 | 0.1619495 | 0.169143 | 0.5196843 | -0.071019 | 0.9708573 |
| ENSG00000099624.7 | ATP5D | -0.154778 | 1 | 0.301001 | 0.7383998 | -1.803075 | 3.99E-12 | -0.200499 | 0.4257214 | 0.312788 | 0.9984257 | -0.027286 | 0.7148199 |
| ENSG00000099625.12 | CBARP | 1.551574 | 0.714717 | 0.472424 | 0.8485564 | -2.533371 | 0.09046443 | 0.544398 | 0.1146644 | 0.789517 | 0.04057216 | 0.081015 | 0.960853 |
| ENSG00000099783.11 | HNRNPM | 0.539955 | 0.9047689 | 0.260091 | 0.8614527 | 0.11852 | 0.8427291 | 0.61899 | 0.2662722 | -0.020104 | 0.006214297 | -0.018502 | 0.2809502 |
| ENSG00000099795.6 | NDUFB7 | 0.158615 | 0.1640608 | 0.100354 | 0.8528295 | -0.397191 | 3.50E-05 | 0.187258 | 0.9650348 | 0.418237 | 0.363108 | 0.393883 | 0.7293549 |
| ENSG00000099797.11 | TECR | -1.405408 | 0.3350729 | 0.408971 | 0.2770455 | 0.146318 | 0.5290845 | -0.327898 | 0.004392971 | 0.220003 | 0.9264364 | -0.38072 | 0.7419782 |
| ENSG00000099800.7 | TIMM13 | 0.223108 | 0.7693197 | -0.180831 | 0.3893023 | -0.270349 | 3.03E-05 | -0.061132 | 0.09856702 | -0.887138 | 1.05E-08 | 0.006554 | 0.6185816 |
| ENSG00000099804.8 | CDC34 | -0.887416 | 0.8885732 | -0.003617 | 0.6527736 | -0.657349 | 0.212474 | 1.28373 | 0.01899838 | 1.688345 | 0.01557237 | -1.351293 | 0.84525 |
| ENSG00000099814.15 | CEP170B | 1.068959 | 0.8867323 | 1.057153 | 0.5240506 | 2.571959 | 0.8656249 | 1.1841 | 0.02915086 | 1.759144 | 0.1257281 | -0.928558 | 0.9449254 |
| ENSG00000099817.11 | POLR2E | -0.37892 | 0.8043486 | -0.079098 | 0.9078292 | -0.098606 | 0.1517379 | -0.461007 | 0.02879545 | 0.085783 | 0.5085946 | -0.997483 | 0.4331698 |
| ENSG00000099860.8 | GADD45B | -0.025151 | 0.7098524 | -0.280406 | 0.8623279 | -0.347816 | 0.4064599 | -0.382091 | 0.6663027 | 1.881927 | 0.006791043 | 0.439956 | 0.9867496 |
| ENSG00000099864.17 | PALM | 1.110193 | 0.536545 | -1.466177 | 0.9473701 | 0.544876 | 0.5290845 | 1.307381 | 0.004459652 | 1.362036 | 0.006425613 | -2.238759 | 0.9426376 |
| ENSG00000099875.14 | MKNK2 | -1.128367 | 0.8006286 | 0.117031 | 0.6527736 | 0.799797 | 0.7149658 | 0.600852 | 0.1075372 | 2.207464 | 0.01193561 | 1.028129 | 0.6829602 |
| ENSG00000099899.14 | TRMT2A | 1.015348 | 0.529676 | 0.40226 | 0.9146016 | -2.028682 | 0.005654245 | 0.428049 | 0.3467508 | -0.494663 | 0.7808339 | 1.270194 | 0.2314187 |
| ENSG00000099901.16 | RANBP1 | -0.050891 | 0.7736255 | -0.090676 | 0.9798987 | -0.952577 | 1.33E-14 | -0.067119 | 0.4644735 | 0.290417 | 0.5196843 | -0.078023 | 0.6108304 |
| ENSG00000099904.15 | ZDHHC8 | 1.946669 | 0.8587992 | 0.145893 | 0.826013 | -1.85132 | 0.1751648 | 2.384652 | 0.001509326 | 1.661445 | 0.1950713 | 0.127788 | 0.6885065 |
| ENSG00000099917.17 | MED15 | -0.753181 | 0.3610267 | -0.245065 | 0.2592088 | 0.997753 | 0.01546479 | 0.53413 | 0.966174 | 0.110944 | 0.827642 | -0.659602 | 0.913718 |
| ENSG00000099940.11 | SNAP29 | -0.291851 | 0.6214382 | 0.114134 | 0.6239827 | -1.610624 | 0.000789967 | 1.09927 | 0.2327125 | 0.649645 | 0.8021966 | 0.903636 | 0.996048 |
| ENSG00000099942.12 | CRKL | -0.409839 | 0.9495183 | 0.020938 | 0.9790409 | 0.001044 | 0.4442624 | 1.701328 | 0.001088311 | 1.245296 | 0.0003226 | 0.506698 | 0.6185816 |
| ENSG00000099949.18 | LZTR1 | -1.463814 | 0.6014343 | 0.054395 | 0.8136662 | 0.127167 | 0.4264835 | 1.593294 | 0.006550271 | 0.566694 | 0.3762104 | -1.297393 | 0.9007097 |
| ENSG00000099956.18 | SMARCB1 | 0.528285 | 0.529676 | 0.361675 | 0.9452791 | -0.919595 | 2.08E-07 | 0.522392 | 0.6274997 | 0.498844 | 0.6774748 | -1.516811 | 0.7096009 |
| ENSG00000099977.13 | DDT | 0.149157 | 0.8065323 | -0.116744 | 0.8263112 | -0.632031 | 0.000225524 | -0.20488 | 0.1483136 | -0.449182 | 0.165158 | 0.381438 | 0.8927477 |
| ENSG00000099991.16 | CABIN1 | -0.778341 | 1 | 0.240416 | 1 | -0.145725 | 0.9852492 | 1.833654 | 0.004146686 | 1.259982 | 0.05983326 | 0.678843 | 0.7915993 |
| ENSG00000100030.14 | MAPK1 | -0.73042 | 0.8006286 | -0.318864 | 0.3642382 | -0.068526 | 0.5935372 | 1.729065 | 3.33E-06 | 1.350782 | 0.0221427 | 0.141391 | 1 |
| ENSG00000100036.12 | SLC35E4 | -0.010339 | 0.7561952 | 0.016784 | 0.6527736 | -0.572043 | 0.5290845 | 0.925829 | 0.04454587 | 2.856064 | 0.0426606 | 1.786803 | 0.8829971 |
| ENSG00000100083.18 | GGA1 | -0.563606 | 0.8006286 | 0.619149 | 0.8434379 | 0.539435 | 0.8074554 | 1.665137 | 0.01052272 | 2.409961 | 0.04251491 | -1.365366 | 0.8949489 |
| ENSG00000100095.18 | SEZ6L | 0.964891 | 0.6214382 | -0.372663 | 0.6560795 | 0.683984 | 0.8154935 | 2.14803 | 0.000186827 | 1.156 | 0.4706961 | -2.536463 | 0.6829816 |
| ENSG00000100097.11 | LGALS1 | -2.451548 | 0.6214382 | -0.185268 | 0.4011918 | -0.627911 | 1.47E-08 | -0.494557 | 2.82E-06 | -0.578447 | 0.000160198 | 0.320296 | 0.8949489 |
| ENSG00000100105.17 | PATZ1 | -1.08678 | 0.7686507 | 1.131722 | 0.9780352 | -0.695921 | 0.6411711 | 0.79578 | 0.009832321 | 0.410081 | 0.4379425 | -2.05419 | 0.6885065 |
| ENSG00000100129.17 | EIF3L | -0.520036 | 0.9678883 | -0.15217 | 0.6527736 | 1.850953 | 0.001019573 | -0.506321 | 0.001116302 | 2.350736 | 0.006884015 | -0.446168 | 1 |
| ENSG00000100138.13 | SNU13 | 0.223005 | 0.6792949 | 0.081932 | 0.9360905 | -0.001849 | 0.05144653 | 0.241735 | 0.07723471 | -0.245206 | 0.003550296 | 0.284113 | 0.8479431 |
| ENSG00000100167.19 | 3-Sep | -0.303925 | 0.9865366 | -0.243849 | 0.8703522 | 0.923849 | 0.3258491 | 0.480754 | 0.005601583 | 0.764329 | 0.3974318 | -0.024895 | 0.6552711 |
| ENSG00000100201.19 | DDX17 | -0.134168 | 0.529676 | 0.167572 | 0.6601152 | 0.621398 | 3.11E-07 | 0.987593 | 0.000349241 | 0.870513 | 0.004704539 | 0.155759 | 0.9618742 |
| ENSG00000100209.9 | HSCB | 0.303169 | 0.9282145 | -0.216866 | 0.4324806 | -0.021476 | 0.8085825 | 0.434641 | 0.09977706 | -1.379298 | 0.02279401 | 2.258438 | 1 |
| ENSG00000100221.10 | JOSD1 | -1.73724 | 0.6214382 | 0.182059 | 0.6527736 | 0.12923 | 0.4064599 | 1.015076 | 0.01023944 | 1.366052 | 0.05691267 | 2.352012 | 0.5839815 |
| ENSG00000100227.17 | POLDIP3 | -0.603763 | 0.8006286 | 0.087896 | 0.4265132 | 0.684463 | 0.05663551 | 0.730493 | 0.08580736 | 1.220708 | 0.04567032 | 1.989989 | 0.5018621 |
| ENSG00000100234.11 | TIMP3 | -2.089692 | 0.02689108 | -0.27673 | 0.5394427 | -0.507715 | 0.7613083 | 1.518095 | 0.01075455 | -0.188048 | 1 | 2.201653 | 0.6885065 |
| ENSG00000100239.15 | PPP6R2 | 0.759034 | 0.3285896 | 0.44676 | 0.6527736 | -0.327383 | 0.04068304 | 0.052905 | 0.4687325 | 0.523646 | 0.7419008 | 1.263262 | 0.7767294 |
| ENSG00000100266.18 | PACSIN2 | 1.049379 | 0.9047689 | -0.512238 | 0.4333971 | -0.449549 | 0.5931059 | 0.221414 | 0.8259827 | 2.30612 | 0.04463757 | -1.303776 | 0.8479431 |
| ENSG00000100288.19 | CHKB | 0.957231 | 1 | 2.028522 | 0.1712543 | -0.273266 | 0.4064599 | 2.520157 | 0.02928659 | 0.492046 | 0.9169158 | -0.828402 | 1 |
| ENSG00000100292.16 | HMOX1 | 0.043059 | 0.733287 | 0.367281 | 9.29E-05 | -0.865879 | 1.44E-06 | 0.454057 | 0.2055888 | 0.666817 | 0.3974318 | -0.49541 | 0.9082795 |
| ENSG00000100294.12 | MCAT | -0.055904 | 0.9342632 | 0.466424 | 0.8110575 | -1.261257 | 1.50E-05 | 0.733557 | 0.4919368 | 0.571279 | 0.5360449 | 2.013253 | 0.7499106 |
| ENSG00000100296.13 | THOC5 | -1.288345 | 0.8006286 | 0.212391 | 0.7246758 | 0.35277 | 1 | 2.695017 | 0.01215412 | 0.891779 | 0.8021966 | 1.176786 | 0.6885065 |
| ENSG00000100300.17 | TSPO | -0.73347 | 1 | -0.207119 | 0.3776134 | -0.16997 | 0.01634596 | -0.189093 | 0.000548681 | -0.138879 | 0.02728334 | 1.182578 | 0.6885065 |
| ENSG00000100316.15 | RPL3 | -0.758274 | 0.01499139 | 0.159064 | 0.06246231 | 0.60874 | 6.82E-28 | -0.229145 | 7.15E-10 | -0.377427 | 1.15E-06 | -0.02159 | 0.4420445 |
| ENSG00000100320.22 | RBFOX2 | -0.590503 | 0.2617323 | -0.176162 | 0.4798292 | 0.768061 | 0.08735445 | 1.084552 | 3.81E-07 | 0.41434 | 0.6496523 | 1.611191 | 1 |
| ENSG00000100330.15 | MTMR3 | 0.444352 | 0.9338818 | 0.142192 | 0.6709033 | -0.141748 | 0.8310952 | 1.538811 | 0.02569154 | 1.740899 | 0.213263 | -1.503435 | 0.8949489 |
| ENSG00000100335.12 | MIEF1 | 0.182808 | 0.9183025 | 0.247534 | 0.9829027 | 0.318434 | 0.152402 | 1.078455 | 0.03894757 | 1.296952 | 0.05694888 | 0.118933 | 0.731863 |
| ENSG00000100345.20 | MYH9 | 1.008492 | 0.563505 | -0.011991 | 0.7828942 | 0.382215 | 0.772119 | 1.09124 | 0.003270086 | 1.805148 | 0.02670539 | -0.347953 | 0.8535547 |
| ENSG00000100348.9 | TXN2 | -0.328399 | 0.8006286 | 0.015642 | 1 | 0.478552 | 0.4738336 | -0.295013 | 0.133515 | -0.398151 | 0.000203276 | -0.805468 | 0.6185816 |
| ENSG00000100350.14 | FOXRED2 | 0.270208 | 0.6214239 | -0.447154 | 0.3044935 | -0.424528 | 0.980796 | 1.279882 | 0.000306426 | 0.639543 | 0.575616 | -0.956235 | 1 |
| ENSG00000100353.17 | EIF3D | -0.197349 | 0.9528265 | 0.177455 | 0.7439698 | 0.279622 | 0.02452832 | 0.143954 | 0.2729069 | 0.917657 | 1 | 1.220052 | 0.7595705 |
| ENSG00000100354.20 | TNRC6B | -0.793675 | 0.9520969 | -0.254129 | 0.6527736 | 0.521372 | 2.56E-07 | 1.552131 | 7.29E-05 | 1.060452 | 0.002249055 | -0.891864 | 0.6829816 |
| ENSG00000100364.18 | KIAA0930 | 0.09106 | 0.7736255 | -0.046382 | 1 | -0.476236 | 0.8610253 | 0.806804 | 0.105536 | 1.453435 | 0.04130846 | 0.671162 | 0.9278206 |
| ENSG00000100376.11 | FAM118A | -1.290803 | 0.8587992 | 0.311051 | 0.9061805 | 0.008065 | 0.7605385 | 1.685539 | 6.83E-05 | 1.220472 | 0.5196843 | -1.88225 | 0.6885065 |
| ENSG00000100379.17 | KCTD17 | -0.846862 | 0.6214382 | 0.084827 | 0.8593459 | -0.175136 | 0.2191854 | -0.056884 | 0.04919189 | -0.381898 | 0.4850443 | -2.000374 | 0.9576568 |
| ENSG00000100380.12 | ST13 | 0.351807 | 0.2600418 | 0.159699 | 0.7234382 | 0.572441 | 2.22E-06 | -0.104522 | 0.1189259 | -0.201351 | 0.08657607 | 0.995745 | 0.7473996 |
| ENSG00000100387.8 | RBX1 | -0.343466 | 0.8178953 | 0.072948 | 0.8394265 | -0.41827 | 0.001013746 | 0.187431 | 0.0616157 | 0.343649 | 0.007418885 | -0.552653 | 0.6885065 |
| ENSG00000100393.10 | EP300 | 1.847433 | 0.469053 | -0.700367 | 0.5059322 | 0.695959 | 0.1184711 | 1.461606 | 0.000107204 | 2.469295 | 0.001121532 | 0.875059 | 0.9370888 |
| ENSG00000100401.19 | RANGAP1 | 0.366107 | 0.9543887 | 0.057403 | 1 | 1.070593 | 0.3642925 | 0.788747 | 0.01753172 | 1.089219 | 0.2550605 | 0.169757 | 0.9646502 |
| ENSG00000100416.12 | TRMU | 0.8063 | 1 | -0.072213 | 0.9120034 | -0.543306 | 0.04447082 | 0.769171 | 0.284134 | 0.789491 | 0.3974318 | 1.52393 | 0.6885065 |
| ENSG00000100422.13 | CERK | 1.233486 | 1 | -0.232535 | 0.9768713 | 0.19977 | 0.7030899 | 2.088725 | 5.64E-06 | 1.402341 | 0.01512965 | 0.67886 | 0.6885065 |
| ENSG00000100425.18 | BRD1 | 0.2784 | 1 | 1.115172 | 0.4130975 | -0.212653 | 0.9578022 | 1.258573 | 0.005644152 | 1.466345 | 0.03469203 | -1.567674 | 0.8353756 |
| ENSG00000100426.6 | ZBED4 | 0.961208 | 0.9264933 | -1.038277 | 0.370295 | 0.159278 | 0.4495278 | 2.472826 | 0.001392121 | 2.814225 | 0.03068444 | 1.052282 | 0.7239287 |
| ENSG00000100427.15 | MLC1 | -2.856044 | 0.6214382 | -1.107906 | 0.00013882 | -0.080139 | 0.8880546 | -0.216275 | 0.2335397 | 0.151171 | 0.8021966 | 3.274909 | 0.6885065 |
| ENSG00000100442.10 | FKBP3 | -1.275508 | 0.01711078 | 0.170587 | 0.7062745 | 0.429093 | 0.7039722 | 0.521608 | 0.7308295 | 0.212537 | 0.05645591 | -0.15214 | 0.6839302 |
| ENSG00000100462.15 | PRMT5 | -0.241388 | 0.7641133 | 0.258249 | 0.9803143 | 0.451221 | 0.2685119 | 0.692502 | 0.03256381 | 0.439749 | 0.916949 | 1.244165 | 0.6829816 |
| ENSG00000100485.11 | SOS2 | -0.314368 | 0.8587992 | 1.021161 | 0.6527736 | -0.753412 | 0.8973482 | 1.738995 | 0.02101693 | 0.715029 | 0.4249407 | 1.572297 | 0.9565287 |
| ENSG00000100503.23 | NIN | 0.066646 | 0.6214382 | 0.011052 | 1 | 0.95922 | 0.01635334 | 1.486886 | 0.0829302 | 0.787569 | 0.9935049 | -1.883998 | 0.6885065 |
| ENSG00000100505.13 | TRIM9 | -0.222109 | 0.6214382 | -0.429478 | 0.01886794 | 0.94619 | 0.2811334 | 0.255489 | 0.07646181 | 0.520754 | 0.5425935 | -0.852332 | 0.7837542 |
| ENSG00000100523.14 | DDHD1 | -0.591537 | 0.6214382 | -0.229643 | 0.6527736 | 1.551062 | 0.233483 | 1.413058 | 0.2089529 | 3.11576 | 0.01361406 | -2.387559 | 0.4578379 |
| ENSG00000100578.14 | KIAA0586 | -1.367075 | 0.7379172 | 0.083789 | 0.4205387 | 1.098058 | 0.04267688 | 0.822269 | 0.5934132 | -0.887455 | 0.522515 | -1.003285 | 0.8755347 |
| ENSG00000100591.7 | AHSA1 | -0.871337 | 0.06605898 | -0.065423 | 0.5395105 | 0.561529 | 0.02823473 | 0.66128 | 0.6371775 | 0.361083 | 0.852599 | 0.188914 | 0.8479431 |
| ENSG00000100596.6 | SPTLC2 | 1.42718 | 0.7736255 | 0.508893 | 0.8905657 | 0.277381 | 0.2397283 | 1.532986 | 0.003599989 | -0.388313 | 0.2384222 | 1.89039 | 0.9627406 |
| ENSG00000100612.13 | DHRS7 | 0.247322 | 0.6214382 | -0.142503 | 0.3027683 | 0.506255 | 0.4388213 | 0.317056 | 0.3278696 | -0.861305 | 3.80E-05 | 1.163508 | 0.8479431 |
| ENSG00000100614.17 | PPM1A | 0.705444 | 0.7241575 | 0.016423 | 0.758504 | 2.164316 | 0.01834538 | 2.936812 | 0.002846276 | 1.269117 | 0.179792 | -0.359791 | 0.6885065 |
| ENSG00000100632.10 | ERH | 0.03718 | 0.529676 | 0.227988 | 0.9766519 | -0.329217 | 0.000634905 | 0.075463 | 0.2930841 | -0.151562 | 0.005199868 | -0.400383 | 0.01381583 |
| ENSG00000100644.16 | HIF1A | -0.099835 | 0.6763454 | -0.102921 | 0.1191613 | 1.461308 | 0.01363097 | 0.224449 | 0.4919368 | -0.025944 | 0.8887598 | -0.029923 | 0.6609803 |
| ENSG00000100650.15 | SRSF5 | -0.272997 | 0.8597832 | 0.482222 | 0.6527736 | -0.086349 | 0.1710895 | 1.356889 | 0.03434133 | 1.084883 | 0.1692823 | 1.064739 | 0.9513337 |
| ENSG00000100664.10 | EIF5 | 0.172274 | 0.6573292 | 0.472475 | 0.09707294 | 0.624532 | 0.1274122 | 0.789264 | 0.00761951 | 0.165152 | 0.5425982 | 1.020073 | 0.9381693 |
| ENSG00000100697.14 | DICER1 | -2.158681 | 0.7010877 | -0.149718 | 0.3121527 | 0.227612 | 0.4064599 | 2.585483 | 0.02260971 | 1.997868 | 0.1580162 | -2.760186 | 0.9023132 |
| ENSG00000100731.15 | PCNX | -0.898271 | 0.6014343 | -0.411066 | 0.4961426 | 0.598995 | 0.5290845 | 1.182317 | 0.001598398 | 0.6054 | 0.8021966 | -1.293167 | 0.6885065 |
| ENSG00000100764.13 | PSMC1 | -0.712364 | 0.2600418 | 0.22311 | 0.4951754 | 0.914419 | 0.0115343 | 0.381532 | 0.9373107 | -0.341845 | 0.002550804 | -0.134727 | 0.6829816 |
| ENSG00000100796.17 | PPP4R3A | 1.157991 | 0.9599095 | -0.135504 | 0.9054089 | 1.042479 | 0.4060104 | 0.261892 | 7.91E-06 | 0.361213 | 0.2058895 | 1.575723 | 0.7420459 |
| ENSG00000100804.18 | PSMB5 | 0.217152 | 0.6014343 | -0.018651 | 0.6527736 | -0.250175 | 0.004738414 | -0.107723 | 0.02095424 | 0.8348 | 0.3974318 | -1.16 | 0.6829816 |
| ENSG00000100811.10 | YY1 | -1.21208 | 0.3471857 | -0.225158 | 0.2388066 | -0.060418 | 0.9634096 | 1.270131 | 0.001195669 | 0.515792 | 0.004505662 | -0.240384 | 0.7565106 |
| ENSG00000100813.14 | ACIN1 | 0.236296 | 0.3946735 | -0.099506 | 0.8655457 | 0.492013 | 0.08679999 | 0.791658 | 0.01855834 | -0.009774 | 0.1026844 | -0.92991 | 0.6885065 |
| ENSG00000100815.12 | TRIP11 | 0.618013 | 0.7568783 | 0.266587 | 1 | 0.510132 | 0.3102224 | 1.358211 | 0.04666571 | 1.787434 | 0.09667266 | -3.191188 | 0.9759869 |
| ENSG00000100852.12 | ARHGAP5 | 0.460988 | 0.3585928 | -0.189601 | 0.6848565 | 0.837256 | 1.44E-10 | 0.974926 | 0.08459104 | 0.648308 | 0.5079513 | -0.213092 | 0.7780985 |
| ENSG00000100888.12 | CHD8 | 1.111107 | 0.6014343 | -0.334888 | 0.3921287 | 0.131338 | 0.3126857 | 1.801456 | 0.000255747 | 1.937957 | 0.01777855 | -0.37724 | 0.7333904 |
| ENSG00000100902.10 | PSMA6 | -0.580732 | 0.8964996 | 1.075599 | 0.5953732 | 0.706156 | 0.001559298 | -0.068557 | 0.7609387 | 1.588399 | 0.04473045 | 1.861443 | 0.6829816 |
| ENSG00000100906.10 | NFKBIA | -0.153535 | 0.7736255 | 0.242998 | 0.6030937 | -1.229501 | 4.55E-16 | 0.219937 | 0.9898756 | 1.376974 | 0.1826169 | -0.1478 | 0.5344421 |
| ENSG00000100911.13 | PSME2 | 0.357569 | 0.9282145 | 0.246618 | 0.7995795 | -0.664499 | 1.44E-10 | -0.00194 | 0.07266524 | 0.435689 | 0.8458207 | 0.235087 | 0.6885065 |
| ENSG00000100941.8 | PNN | -0.434011 | 0.6214382 | -0.051707 | 0.6140501 | -0.222712 | 0.9977776 | 0.871431 | 0.003528825 | 0.715039 | 0.9463565 | 0.210147 | 0.6829816 |
| ENSG00000100994.11 | PYGB | 1.499677 | 0.3589335 | -0.289141 | 0.8165715 | -0.074395 | 0.4796547 | 1.301792 | 0.000130593 | 0.652994 | 0.9372936 | -0.389567 | 0.6185816 |
| ENSG00000101040.19 | ZMYND8 | -1.984708 | 0.1607609 | -0.635139 | 0.5110299 | 0.146293 | 0.1049431 | 0.696908 | 8.64E-05 | 0.114171 | 0.7612996 | -0.628745 | 0.5344421 |
| ENSG00000101084.16 | C20orf24 | -0.834333 | 0.4061458 | 0.087457 | 0.8703522 | -0.722404 | 5.23E-13 | 0.499935 | 0.9508466 | -0.088232 | 0.5196843 | -0.21815 | 0.6885065 |
| ENSG00000101098.12 | RIMS4 | 0.064674 | 0.9426668 | 0.031392 | 1 | -0.205444 | 0.9604268 | 0.381772 | 0.0478576 | 0.268186 | 0.8754603 | 1.436801 | 0.868568 |
| ENSG00000101109.11 | STK4 | -1.635413 | 0.5084093 | 0.261016 | 0.8263112 | 1.21561 | 0.000710551 | 1.976586 | 0.000847916 | -0.117332 | 0.7913786 | -0.399424 | 0.9869001 |
| ENSG00000101126.15 | ADNP | 0.174064 | 0.6214382 | 0.108926 | 0.9353253 | 0.306269 | 0.5290845 | 0.992573 | 0.01524284 | 1.362012 | 0.000786243 | -0.443804 | 0.6885065 |
| ENSG00000101132.9 | PFDN4 | 0.151118 | 0.563505 | 0.203354 | 0.9577957 | -0.109509 | 0.0328671 | 0.212882 | 0.3003919 | -0.751533 | 0.000715496 | -0.175729 | 0.3746505 |
| ENSG00000101134.11 | DOK5 | -0.26741 | 0.9547805 | 0.355258 | 1 | -0.249885 | 0.1996959 | 0.026531 | 0.03189673 | -2.376117 | 0.04222895 | -3.071062 | 0.9420266 |
| ENSG00000101144.12 | BMP7 | 0.098645 | 0.6214382 | -1.182913 | 0.1277169 | 0.297352 | 0.9852492 | 1.413239 | 0.01046955 | 0.548806 | 1 | 0.686004 | 1 |
| ENSG00000101146.12 | RAE1 | 1.018324 | 0.714717 | 0.015014 | 1 | -0.361568 | 0.4064599 | 0.124292 | 0.02811515 | 0.930762 | 0.5767563 | 0.063813 | 0.874213 |
| ENSG00000101152.10 | DNAJC5 | 0.419304 | 0.8587992 | -0.348021 | 0.4605921 | -0.051021 | 0.5290845 | 0.593144 | 0.000284065 | 1.254069 | 0.4244936 | 0.184877 | 0.8949489 |
| ENSG00000101181.17 | MTG2 | -0.594185 | 0.9445494 | -0.064306 | 0.6527736 | -0.221112 | 0.1858002 | 0.96687 | 0.8412773 | 2.13353 | 0.02906571 | -2.68652 | 0.6885065 |
| ENSG00000101182.14 | PSMA7 | -0.305666 | 0.7736255 | 0.246189 | 0.5448012 | -0.16986 | 0.0122211 | -0.179289 | 0.000160243 | -0.393831 | 1.37E-08 | -0.286158 | 0.00077615 |
| ENSG00000101189.6 | MRGBP | -1.221297 | 0.8867323 | 0.555562 | 0.7199683 | -1.757318 | 0.02382767 | 0.842129 | 0.02693198 | 1.401495 | 0.4802509 | -0.54064 | 0.967979 |
| ENSG00000101191.16 | DIDO1 | 1.548295 | 0.563505 | -0.293421 | 0.5651089 | -0.109118 | 0.7285135 | 0.911537 | 0.003106508 | 0.569279 | 0.7377262 | -0.45302 | 0.7325363 |
| ENSG00000101199.12 | ARFGAP1 | -3.821203 | 0.003707355 | 0.092657 | 0.9696336 | 0.111488 | 0.7437308 | 0.876202 | 0.00136632 | 0.449743 | 0.898446 | 0.51796 | 0.6885065 |
| ENSG00000101203.16 | COL20A1 | 1.514347 | 1 | -0.971203 | 0.4011918 | 0.543652 | 0.4526189 | 0.955726 | 0.2568542 | 3.271487 | 0.01610973 | -3.897408 | 0.05357521 |
| ENSG00000101210.10 | EEF1A2 | -1.055661 | 0.714717 | 0.373858 | 0.6165309 | -1.926916 | 0.6010661 | 1.084598 | 6.03E-06 | 1.446257 | 0.001534519 | -0.755057 | 0.7745574 |
| ENSG00000101216.10 | GMEB2 | 0.921224 | 0.7736255 | -1.161349 | 0.6527736 | 1.125874 | 0.5290845 | 0.018881 | 0.7144938 | 3.079481 | 0.0191411 | -0.249893 | 0.6879794 |
| ENSG00000101220.17 | C20orf27 | 0.104594 | 0.6214382 | 0.197414 | 0.9643513 | -0.309088 | 0.000448866 | -0.42015 | 0.6051424 | -0.630457 | 0.4211066 | -0.656416 | 0.6185816 |
| ENSG00000101224.17 | CDC25B | -0.48029 | 0.8867323 | -0.530067 | 0.2553839 | -0.753801 | 0.03835208 | -0.161212 | 0.2374485 | 0.465807 | 0.1332756 | 0.564794 | 0.9147227 |
| ENSG00000101255.10 | TRIB3 | 1.225267 | 0.5711449 | 0.773035 | 0.03594505 | -0.085564 | 0.6827367 | 0.26015 | 1 | 0.692403 | 0.5079513 | 0.008082 | 0.9545062 |
| ENSG00000101265.15 | RASSF2 | -0.054024 | 1 | 0.072111 | 0.9028136 | 0.232901 | 0.02301863 | 0.747993 | 0.008380351 | 0.777452 | 0.9405498 | -1.237268 | 0.7648916 |
| ENSG00000101266.16 | CSNK2A1 | -0.121164 | 0.8740747 | 0.118677 | 0.8953649 | 0.52697 | 9.60E-06 | 0.208846 | 0.8448268 | -0.097402 | 0.1367421 | 0.618058 | 0.6885065 |
| ENSG00000101310.14 | SEC23B | -0.161393 | 0.9891705 | 0.152974 | 0.9856559 | 0.374161 | 0.5290845 | 0.501659 | 0.8472142 | -0.433649 | 0.02376642 | 1.082858 | 0.8479431 |
| ENSG00000101347.8 | SAMHD1 | 1.018161 | 0.9054014 | 0.964046 | 0.6527736 | 0.179186 | 0.9807333 | 0.183869 | 0.03396132 | -1.064588 | 0.621723 | 0.999029 | 0.9733288 |
| ENSG00000101350.7 | KIF3B | 0.278233 | 0.7107241 | -0.042007 | 0.8610208 | 0.945736 | 0.4580147 | 1.609973 | 0.009509884 | 1.42888 | 0.01544497 | -2.248315 | 0.6885065 |
| ENSG00000101361.14 | NOP56 | -0.133041 | 0.8867323 | 0.037229 | 0.9856559 | 0.049772 | 0.9545013 | 0.908329 | 8.99E-10 | 0.759929 | 0.0963637 | -0.268418 | 0.1987453 |
| ENSG00000101367.8 | MAPRE1 | 0.265519 | 0.5711449 | -0.392409 | 0.4886958 | 0.318429 | 0.5306026 | 0.7824 | 0.04071092 | 0.906094 | 0.05645591 | -1.355066 | 0.8479431 |
| ENSG00000101384.11 | JAG1 | 0.087243 | 0.4790285 | 0.249309 | 0.6414243 | 0.456439 | 0.005631247 | 1.160273 | 8.51E-08 | -0.09448 | 0.9690599 | 0.005272 | 0.6885065 |
| ENSG00000101421.3 | CHMP4B | -0.79985 | 0.59287 | -0.081558 | 0.5517607 | -0.432118 | 1.68E-08 | 0.893049 | 0.02840223 | -0.052294 | 0.2387576 | 0.562918 | 0.6829816 |
| ENSG00000101439.8 | CST3 | -0.362534 | 0.1035929 | -0.23485 | 0.1573875 | -2.532412 | 1.91E-148 | -0.530787 | 6.77E-07 | -0.95889 | 0.2011388 | 0.316567 | 0.8479431 |
| ENSG00000101442.9 | ACTR5 | -0.230661 | 0.9410335 | 0.316416 | 0.4518214 | -0.245033 | 0.7249136 | 0.553905 | 0.7205095 | 2.691133 | 0.0499092 | -2.247697 | 0.7990826 |
| ENSG00000101457.12 | DNTTIP1 | -1.618599 | 0.2132806 | 0.126982 | 0.7436238 | -0.394345 | 0.01098563 | 0.337399 | 0.9706579 | 0.344514 | 0.9498244 | -0.022935 | 0.6885065 |
| ENSG00000101460.12 | MAP1LC3A | 0.151399 | 0.7736255 | 0.658346 | 0.115361 | -0.791061 | 3.57E-10 | 0.006 | 0.01618006 | 0.469276 | 0.8972581 | 0.15289 | 0.557022 |
| ENSG00000101542.9 | CDH20 | 2.790586 | 0.6214382 | 0.591526 | 0.7881421 | 1.04471 | 0.8100781 | -0.029009 | 0.8764761 | -0.545736 | 0.643255 |  |  |
| ENSG00000101577.9 | LPIN2 | -1.474429 | 0.6214382 | 0.282838 | 0.9634109 | 0.358755 | 0.980796 | 2.001531 | 0.00224603 | 0.91511 | 0.7178266 | 1.784737 | 0.8489889 |
| ENSG00000101596.14 | SMCHD1 | -0.527983 | 0.6214382 | -0.105291 | 0.5953732 | 0.745878 | 0.03095238 | 1.692164 | 0.02173411 | 1.702141 | 0.04498667 | -0.205605 | 0.6885065 |
| ENSG00000101608.12 | MYL12A | -0.640222 | 0.8587992 | 0.483639 | 0.2785898 | -0.137351 | 0.03323509 | -0.028618 | 0.04446411 | -0.047822 | 0.2040744 | -0.393047 | 0.6885065 |
| ENSG00000101654.17 | RNMT | -0.504894 | 0.9798794 | 0.304638 | 0.8017884 | 0.481338 | 0.00238046 | 0.46222 | 0.8224647 | 0.489156 | 0.5196843 | -0.679724 | 0.8661501 |
| ENSG00000101745.15 | ANKRD12 | -1.155808 | 0.04500243 | 0.196799 | 0.9859694 | 1.528321 | 3.38E-13 | 0.929397 | 0.02787088 | 1.074774 | 0.3507718 | 0.779629 | 0.9581586 |
| ENSG00000101746.15 | NOL4 | -1.009645 | 0.6214382 | 0.480748 | 0.9726126 | 2.652941 | 0.000909714 | -0.113585 | 0.4566021 | 1.75387 | 0.03056663 | -0.186854 | 0.6885065 |
| ENSG00000101752.11 | MIB1 | -0.152299 | 0.8739908 | -0.27984 | 0.8263112 | 1.554381 | 0.000291694 | 1.815549 | 8.71E-05 | 1.523518 | 0.08379252 | -0.471688 | 0.9264343 |
| ENSG00000101782.14 | RIOK3 | 0.19062 | 0.8006286 | 0.14687 | 0.6527736 | 0.900852 | 0.00715996 | 1.381475 | 0.1142841 | 1.268422 | 0.8021966 | -0.002412 | 0.9147227 |
| ENSG00000101843.18 | PSMD10 | -0.504586 | 0.4925184 | 0.274979 | 0.6527736 | 0.212635 | 0.6717367 | -0.552802 | 0.07264986 | -1.120869 | 0.002294203 | 1.876549 | 0.6885065 |
| ENSG00000101966.12 | XIAP | -0.615755 | 0.7640133 | 0.401755 | 0.7427243 | 0.721116 | 0.01286473 | 1.382672 | 9.27E-07 | 0.82783 | 0.04159686 | 0.388988 | 0.8963036 |
| ENSG00000101972.18 | STAG2 | 1.758887 | 0.2909961 | -0.010295 | 0.8760416 | 0.801795 | 6.14E-06 | 1.879394 | 0.02610363 | -0.277137 | 0.2317816 | 0.212248 | 0.9147227 |
| ENSG00000102003.10 | SYP | -0.246121 | 0.8006286 | 1.868189 | 0.9104976 | 0.274895 | 0.1634946 | 0.361351 | 7.46E-05 | 0.469301 | 0.2154301 | -1.434039 | 0.6185816 |
| ENSG00000102007.10 | PLP2 | -4.037247 | 0.6214382 | 0.286917 | 0.6527736 | 0.370851 | 0.000144373 | -0.192635 | 0.1086815 | -0.026549 | 0.1778915 | 0.205703 | 0.8345878 |
| ENSG00000102024.17 | PLS3 | 1.298751 | 0.3946735 | -0.021974 | 0.6918607 | 0.342811 | 0.9807333 | -1.136963 | 0.3361528 | -1.749964 | 0.005809771 | 0.508366 | 0.3719182 |
| ENSG00000102030.15 | NAA10 | 0.354179 | 0.8006286 | 0.313782 | 0.6527736 | -0.521416 | 0.001612629 | 0.529105 | 0.4919368 | 0.518548 | 0.3812741 | 0.528253 | 0.8919564 |
| ENSG00000102038.15 | SMARCA1 | -0.604107 | 0.714717 | -0.071336 | 0.2570642 | 0.373183 | 0.007100439 | 0.667684 | 0.4611148 | -0.83596 | 0.3302713 | -0.901256 | 0.6108304 |
| ENSG00000102103.15 | PQBP1 | 0.051178 | 0.6214382 | 0.269967 | 0.6247401 | -0.356054 | 0.01616267 | -0.710715 | 0.02580419 | -0.043718 | 0.2376995 | -0.390112 | 1 |
| ENSG00000102109.8 | PCSK1N | -0.588557 | 0.714717 | 0.664731 | 0.02531857 | -2.614963 | 8.60E-13 | 0.571713 | 0.2064104 | 1.492805 | 0.001220683 | -0.212728 | 0.5344421 |
| ENSG00000102119.10 | EMD | -0.975113 | 1 | 0.412674 | 0.8613445 | -0.122461 | 0.03757911 | 0.068457 | 0.2335397 | 0.737533 | 0.8427757 | -0.4986 | 0.5215899 |
| ENSG00000102125.15 | TAZ | -1.29883 | 0.8867323 | 0.304224 | 0.9476887 | 0.409875 | 0.5290845 | 0.196435 | 0.04372344 | 0.887277 | 0.5196843 | 2.21389 | 0.6556863 |
| ENSG00000102144.13 | PGK1 | 0.318809 | 0.6214382 | 0.374504 | 0.005699963 | 0.154365 | 0.5290845 | 0.130454 | 0.3203369 | -0.16383 | 0.001611567 | -0.199825 | 0.5344421 |
| ENSG00000102172.15 | SMS | -0.479266 | 0.8787256 | -0.003824 | 0.9222921 | 0.390471 | 0.4696378 | 0.335778 | 0.5523657 | 0.026349 | 0.04311175 | 0.202896 | 0.8479431 |
| ENSG00000102181.20 | CD99L2 | -0.224194 | 0.9047689 | -0.152606 | 0.6184438 | 0.125105 | 0.8454005 | 0.37219 | 0.009224418 | 0.635431 | 0.573869 | -0.346143 | 0.7709617 |
| ENSG00000102189.16 | EEA1 | -2.958754 | 0.7621571 | -0.38241 | 0.1606333 | 1.23582 | 2.14E-08 | 0.844443 | 0.3164813 | 1.171632 | 0.9421969 | 0.007813 | 0.9626957 |
| ENSG00000102225.15 | CDK16 | -0.344218 | 0.9789948 | -0.31948 | 0.6527736 | -0.598833 | 0.07244964 | 1.76997 | 0.00168949 | 0.71112 | 0.04940493 | -0.156606 | 0.9219559 |
| ENSG00000102226.9 | USP11 | 0.212088 | 1 | 0.041321 | 0.9766519 | -0.274833 | 0.05669581 | 0.894092 | 0.004146686 | 0.237916 | 0.8297892 | 0.010407 | 0.6829816 |
| ENSG00000102265.11 | TIMP1 | -1.142032 | 0.9047689 | -0.048143 | 0.5010139 | -0.390081 | 9.39E-06 | -0.319216 | 0.000284065 | -0.047508 | 1.97E-06 | 0.809009 | 0.6421783 |
| ENSG00000102309.12 | PIN4 | 0.219898 | 0.9959253 | -0.028537 | 1 | -0.651187 | 0.005898071 | 0.141418 | 0.8635576 | 0.078371 | 0.4278631 | -0.086562 | 0.9627406 |
| ENSG00000102390.10 | PBDC1 | -0.358311 | 0.6595079 | -0.071252 | 0.6527736 | 0.791935 | 0.9642892 | 0.812725 | 0.1731044 | -0.874416 | 0.01297795 | 1.397459 | 0.9270789 |
| ENSG00000102547.18 | CAB39L | 0.221987 | 0.8467545 | -0.817235 | 0.7942243 | 0.423271 | 0.1694261 | -0.879853 | 0.03397274 | -0.578597 | 0.7253092 | -3.374703 | 0.7915993 |
| ENSG00000102606.17 | ARHGEF7 | -0.598591 | 0.5437972 | -0.181427 | 0.9797526 | 1.196389 | 0.01145515 | 0.593315 | 0.03626243 | 0.238443 | 0.6038912 | 2.312593 | 0.6517664 |
| ENSG00000102710.19 | SUPT20H | -0.56715 | 0.8006286 | -0.903747 | 0.2372836 | 0.839254 | 0.003743515 | 1.876715 | 0.1651369 | -1.426849 | 0.4909621 | -0.782852 | 0.7641741 |
| ENSG00000102763.15 | VWA8 | -2.553683 | 0.563505 | -0.056148 | 0.8874648 | 0.511879 | 0.9345978 | 1.032943 | 0.04248025 | 0.02495 | 0.7973092 | -1.623024 | 0.7767294 |
| ENSG00000102780.16 | DGKH | -0.191629 | 1 | -0.948871 | 1 | 1.368358 | 0.02427694 | 1.109275 | 0.5903915 | -0.004707 | 0.9535619 | -0.272479 | 0.9455597 |
| ENSG00000102786.14 | INTS6 | 1.596212 | 0.529676 | -0.219026 | 0.6527736 | 0.907416 | 0.02173691 | 2.06652 | 0.02353059 | 0.869903 | 0.2133723 | 1.880997 | 0.9147227 |
| ENSG00000102804.14 | TSC22D1 | 0.768226 | 0.01839876 | -0.383623 | 0.5046722 | 0.524255 | 0.08581726 | 0.780919 | 0.1417544 | 0.912331 | 0.3744595 | 0.377578 | 0.6440403 |
| ENSG00000102870.5 | ZNF629 | -2.763328 | 0.03629807 | -0.266969 | 0.9752186 | -0.460436 | 0.6608038 | 2.16593 | 0.01983441 | 2.11259 | 0.1908059 | -0.210319 | 0.9285229 |
| ENSG00000102893.15 | PHKB | -1.532989 | 0.6214382 | -0.682371 | 0.3121527 | 0.928851 | 0.01488258 | 2.078296 | 0.003314109 | 0.235497 | 0.8844564 | -0.60981 | 0.9536927 |
| ENSG00000102901.12 | CENPT | -0.463053 | 0.9513755 | -0.1705 | 0.7317746 | 0.058129 | 0.7437308 | 1.612684 | 0.03902563 | 2.153848 | 0.589803 | -1.027977 | 0.8479431 |
| ENSG00000102910.13 | LONP2 | -1.367666 | 0.6214382 | -0.099529 | 0.6527736 | 0.187145 | 0.02682175 | 0.311216 | 0.2744618 | 1.966317 | 0.1060143 | 0.427334 | 0.9696094 |
| ENSG00000102974.14 | CTCF | -1.206544 | 0.6214382 | -0.509224 | 0.3792317 | 0.230845 | 0.3744427 | 1.356119 | 0.001766262 | 0.224309 | 0.9566575 | 0.645086 | 0.7665307 |
| ENSG00000103035.10 | PSMD7 | -0.29894 | 0.8994202 | -0.04627 | 0.4487482 | 0.540355 | 0.000359112 | -0.565392 | 0.001221598 | 0.267722 | 0.07078934 | 0.870039 | 1 |
| ENSG00000103042.8 | SLC38A7 | 1.617818 | 0.3166974 | -0.157456 | 0.7955884 | -0.718635 | 0.5290845 | 3.139638 | 0.01275141 | 1.781796 | 0.5431796 | -2.266017 | 0.9315616 |
| ENSG00000103043.14 | VAC14 | 0.440806 | 0.9047689 | 0.349501 | 0.8106695 | -0.883119 | 0.5103391 | 2.769946 | 0.007914848 | 0.892017 | 0.4244936 | -2.357172 | 0.6885065 |
| ENSG00000103150.5 | MLYCD | -1.730434 | 0.6214382 | -1.122771 | 0.6200225 | -0.600252 | 0.1902429 | 0.556891 | 0.01264182 | -0.951997 | 0.4010034 | 2.994996 | 0.8949489 |
| ENSG00000103152.11 | MPG | -0.079845 | 0.8006286 | 0.190657 | 0.9114157 | -0.506408 | 0.003012856 | -0.161588 | 0.4050682 | 0.039298 | 0.3810728 | 0.343778 | 1 |
| ENSG00000103160.11 | HSDL1 | 0.601723 | 0.8235833 | -0.914221 | 0.6487262 | 0.312985 | 0.84816 | 1.271191 | 0.01595271 | -0.391831 | 0.5196843 | 1.245032 | 0.4498785 |
| ENSG00000103168.16 | TAF1C | 0.700153 | 0.5711449 | -0.007545 | 0.8517906 | -0.145029 | 0.5290845 | 2.904403 | 0.1250317 | 2.685297 | 0.007885052 | -2.389556 | 0.9653202 |
| ENSG00000103194.15 | USP10 | -1.105237 | 0.1540432 | 0.326598 | 0.6527736 | 0.529217 | 1.11E-06 | 0.378484 | 0.4919368 | -0.235902 | 0.621723 | -0.895456 | 0.7233657 |
| ENSG00000103202.12 | NME4 | -1.189535 | 0.02247068 | -0.290057 | 0.121014 | -1.471981 | 3.06E-14 | 0.121762 | 0.4919368 | 0.089172 | 0.3994662 | 0.227851 | 0.6829816 |
| ENSG00000103254.9 | FAM173A | -0.192567 | 0.6214382 | -0.291713 | 0.3121527 | -2.345125 | 0.000731729 | 0.18246 | 0.4369123 | -0.506973 | 0.3543796 | -1.824152 | 0.4553016 |
| ENSG00000103257.8 | SLC7A5 | -0.835435 | 0.6425861 | 0.40887 | 0.3778953 | -0.285096 | 0.5290845 | 1.451773 | 3.52E-05 | 1.250343 | 1.73E-06 | -0.765588 | 0.8816797 |
| ENSG00000103260.8 | METRN | -0.768536 | 0.5462034 | -0.575132 | 0.3620504 | -1.331441 | 0.000626055 | 1.288711 | 9.37E-07 | 1.659511 | 0.000428427 | -0.158552 | 0.2941125 |
| ENSG00000103266.10 | STUB1 | -0.69149 | 0.563505 | -0.220015 | 0.5222411 | -0.539778 | 0.000201068 | 0.517627 | 1 | 0.483145 | 0.8753948 | -0.003576 | 0.8479431 |
| ENSG00000103319.11 | EEF2K | 0.749072 | 0.6214382 | -0.904719 | 0.1284061 | -0.093532 | 0.8068696 | 0.13821 | 0.06459584 | 4.033895 | 0.0368586 | -0.211102 | 0.8979257 |
| ENSG00000103326.10 | CAPN15 | -3.885555 | 1 | 0.730362 | 0.6200225 | -1.431282 | 0.03552749 | 1.142049 | 0.00631271 | 1.510655 | 0.2372778 | -2.756463 | 0.4240407 |
| ENSG00000103335.19 | PIEZO1 | -0.896003 | 0.6214382 | -0.105722 | 1 | -0.840603 | 0.7250004 | 0.604164 | 0.002658559 | -0.059645 | 1 | -2.114538 | 0.9562356 |
| ENSG00000103342.12 | GSPT1 | -0.579148 | 0.8133722 | 0.014501 | 0.9561974 | -0.157393 | 0.7463765 | 0.276939 | 0.000896135 | 1.822048 | 0.0176653 | -0.759037 | 0.797124 |
| ENSG00000103363.14 | TCEB2 | -1.335769 | 5.45E-05 | -0.081418 | 0.5953732 | -0.466751 | 4.82E-08 | 0.224614 | 0.763492 | 0.350603 | 0.08221861 | -0.181021 | 0.2159565 |
| ENSG00000103429.10 | BFAR | -0.29429 | 0.8006286 | 0.12224 | 0.8217443 | 0.104603 | 0.8100781 | 2.077276 | 0.000110815 | 0.420201 | 1 | -0.361586 | 0.9213339 |
| ENSG00000103460.16 | TOX3 | -1.625292 | 0.6214382 | -5.001422 | 0.6200225 | 0.929884 | 0.02941896 | -0.023382 | 0.1377261 | 0.761949 | 0.5196843 | 1.831428 | 0.6885065 |
| ENSG00000103489.11 | XYLT1 | 0.478242 | 0.563505 | -1.08819 | 0.7647276 | 0.423204 | 0.007067549 | 3.618177 | 0.03028101 | 2.54718 | 0.03792058 | #NAME? | 0.5344421 |
| ENSG00000103495.13 | MAZ | 0.046197 | 0.8605883 | 0.856052 | 0.01578883 | -0.42273 | 0.01139218 | 1.179736 | 1.74E-05 | 0.986924 | 0.001848342 | -1.709437 | 0.09142162 |
| ENSG00000103496.14 | STX4 | -1.326701 | 0.0829426 | 0.655196 | 0.5953732 | -1.480991 | 0.1337577 | -0.17525 | 0.01135963 | 0.911026 | 0.4421603 | 0.239903 | 1 |
| ENSG00000103512.14 | NOMO1 | 1.231429 | 0.6214382 | 0.21064 | 0.95708 | 0.271699 | 0.8386403 | 1.522126 | 0.001657996 | -0.089577 | 0.1687325 | 0.282573 | 0.6778167 |
| ENSG00000103528.16 | SYT17 | 0.285778 | 0.8809848 | -1.102058 | 0.3776134 | -0.780567 | 0.3183629 | 1.328727 | 7.95E-05 | 1.147826 | 0.2265525 | 2.052803 | 0.8479431 |
| ENSG00000103540.16 | CCP110 | -1.485103 | 0.3683144 | -0.897459 | 0.05090264 | 1.235363 | 0.000308465 | 1.505154 | 0.001753928 | 1.638463 | 0.03247574 | 0.445447 | 0.6885065 |
| ENSG00000103647.12 | CORO2B | -0.273957 | 0.8006286 | 0.483341 | 0.3610905 | -0.455764 | 0.855815 | 1.460915 | 0.004266903 | 2.329249 | 0.1244284 | -0.227499 | 0.8145966 |
| ENSG00000103653.16 | CSK | 0.806938 | 0.3619888 | 0.711557 | 0.2968464 | -1.230905 | 0.6217702 | 2.17577 | 0.0002807 | 0.628053 | 0.04756918 | -0.88685 | 0.6885065 |
| ENSG00000103657.13 | HERC1 | 1.040408 | 0.8867323 | 0.488958 | 1 | -0.000581 | 0.3494524 | 1.499444 | 0.02658287 | 1.216866 | 0.1663076 | -2.460137 | 0.6885065 |
| ENSG00000103740.9 | ACSBG1 | -0.604064 | 0.8587992 | 0.196612 | 0.8962595 | -0.850674 | 0.6824517 | 4.707736 | 0.04008401 | -0.286456 | 1 | #NAME? | 0.6885065 |
| ENSG00000103855.17 | CD276 | -0.155754 | 0.6214382 | -0.290559 | 0.1834552 | -0.11503 | 0.8356299 | 1.138374 | 0.009744773 | 1.291999 | 0.04057216 | 1.102226 | 0.4149735 |
| ENSG00000103966.9 | EHD4 | 0.278813 | 1 | 0.043311 | 0.6532996 | 0.43969 | 0.9626429 | 2.047185 | 0.02147125 | 1.462998 | 0.3208953 | 1.01468 | 1 |
| ENSG00000103994.16 | ZNF106 | -0.776761 | 0.6214382 | 0.16214 | 0.6527736 | 0.651743 | 0.000481242 | 0.511436 | 0.08780156 | -0.427808 | 0.5440486 | 1.021945 | 0.9741465 |
| ENSG00000104093.13 | DMXL2 | -1.406256 | 0.6214382 | 0.204122 | 0.7605083 | 1.204599 | 0.04483046 | 2.303247 | 0.001646819 | 1.668397 | 0.1430508 | 3.291943 | 0.6556863 |
| ENSG00000104112.8 | SCG3 | 0.596648 | 0.2644645 | 0.239622 | 0.9653874 | 0.637956 | 0.002016951 | -0.153633 | 0.5690991 | 0.139705 | 0.2880317 | 0.213622 | 0.9415471 |
| ENSG00000104131.12 | EIF3J | -0.511244 | 0.8101751 | 0.048623 | 0.8955188 | 0.023397 | 0.4816459 | 0.155782 | 1 | 0.132392 | 0.009057514 | -0.223837 | 0.6829816 |
| ENSG00000104164.10 | BLOC1S6 | -0.684095 | 0.6595756 | 0.242481 | 0.9217775 | 0.162613 | 0.5614383 | 0.757399 | 0.009446349 | -0.126785 | 0.1772087 | 0.229242 | 0.9612903 |
| ENSG00000104177.17 | MYEF2 | -0.63018 | 0.6214382 | -0.268478 | 0.6200225 | 0.736636 | 0.01431886 | 0.96362 | 0.171072 | 0.167856 | 0.8968578 | -0.466073 | 0.9099979 |
| ENSG00000104218.14 | CSPP1 | -0.208165 | 1 | -0.062288 | 0.9080086 | 0.488731 | 0.4018207 | 0.51459 | 0.02549987 | 0.235293 | 0.7631604 | 1.780041 | 0.7788913 |
| ENSG00000104228.12 | TRIM35 | -0.549446 | 0.9282145 | 0.629137 | 0.9031929 | -1.202103 | 0.6295125 | 2.283392 | 0.007565047 | -0.884344 | 0.924393 | 1.047793 | 0.6885065 |
| ENSG00000104290.10 | FZD3 | -0.344343 | 0.6214382 | -0.129492 | 0.8647637 | 0.968885 | 0.000688818 | 0.922494 | 2.67E-05 | 0.903923 | 0.621723 | 0.214952 | 0.8250047 |
| ENSG00000104312.7 | RIPK2 | -1.775421 | 0.714717 | -0.068743 | 0.8110575 | 0.125795 | 0.8598754 | 0.17111 | 0.001794053 | 0.230079 | 0.8313245 | -0.952325 | 0.9719512 |
| ENSG00000104325.6 | DECR1 | -0.142013 | 1 | 0.006332 | 0.5010139 | -0.235714 | 0.632081 | -0.043328 | 0.01154793 | -0.493191 | 0.03666503 | 0.251381 | 0.6885065 |
| ENSG00000104331.8 | IMPAD1 | 0.0938 | 0.8075018 | -0.365727 | 0.5010139 | 0.287677 | 0.2879524 | 0.882059 | 0.002667715 | 1.182269 | 0.1505337 | -1.430325 | 0.7884953 |
| ENSG00000104341.16 | LAPTM4B | 0.483006 | 0.8509619 | -0.332072 | 0.0410984 | 0.329192 | 1 | 0.698326 | 0.008997755 | 0.042287 | 0.9566575 | 0.069059 | 0.6829816 |
| ENSG00000104356.10 | POP1 | -1.869504 | 0.714717 | -0.160283 | 0.978688 | 1.476293 | 0.03142041 | 1.625115 | 0.004091238 | 0.970314 | 0.5722804 | -2.671515 | 1 |
| ENSG00000104381.12 | GDAP1 | 0.205409 | 0.980017 | -0.987319 | 0.3583128 | -0.193537 | 0.782435 | 0.587695 | 0.004279695 | 2.093406 | 0.09456206 | 3.401035 | 0.6885065 |
| ENSG00000104408.9 | EIF3E | -0.06762 | 0.8971874 | 0.161335 | 0.9415457 | 0.69081 | 0.000599534 | 0.331834 | 0.1880764 | 1.157549 | 0.1420501 | -0.544574 | 0.4627089 |
| ENSG00000104412.7 | EMC2 | -1.701978 | 0.4552761 | -0.324657 | 0.7734014 | -0.662224 | 0.006978465 | 1.176914 | 0.462804 | 0.848811 | 0.5196843 | 0.608748 | 0.9169962 |
| ENSG00000104419.14 | NDRG1 | 2.798202 | 0.6214382 | 0.639656 | 0.003406272 | 0.31825 | 0.6937701 | 1.215452 | 0.2678339 | 1.295776 | 0.9907648 | 1.47174 | 0.6556863 |
| ENSG00000104435.13 | STMN2 | 1.314975 | 0.6214382 | -3.964469 | 0.6200225 | -1.797962 | 0.000418292 | 0.29608 | 0.8757984 | -0.074925 | 0.07005549 | 0.591265 | 0.6829816 |
| ENSG00000104447.11 | TRPS1 | -0.240475 | 0.8006286 | 0.609072 | 0.8532785 | 1.671957 | 0.04273058 | -0.634558 | 0.4919368 | -0.662215 | 0.9263598 | 3.854901 | 0.6267144 |
| ENSG00000104517.12 | UBR5 | -1.34459 | 0.5454431 | 0.048811 | 0.5953732 | 0.606234 | 0.000410193 | 1.015503 | 1.54E-06 | 1.914276 | 0.000254377 | -1.127953 | 0.8479431 |
| ENSG00000104529.17 | EEF1D | -1.35239 | 0.2213095 | 0.094393 | 0.7383998 | 0.237555 | 0.1065994 | 0.270412 | 0.4340365 | -0.101113 | 0.1229439 | -0.397928 | 0.02837516 |
| ENSG00000104635.13 | SLC39A14 | 1.033121 | 0.6014343 | 0.490776 | 0.338351 | -0.690645 | 0.5290845 | 1.790793 | 2.33E-05 | -0.213864 | 1 | 1.08969 | 0.9944299 |
| ENSG00000104643.9 | MTMR9 | -0.494853 | 0.8064015 | 0.056084 | 0.8045435 | 1.009326 | 0.4923004 | 1.18257 | 0.0102848 | 0.842404 | 0.7377262 | -0.495938 | 0.9048106 |
| ENSG00000104671.7 | DCTN6 | -0.236219 | 0.8689227 | -0.063316 | 0.5953732 | -0.021647 | 0.6140882 | -0.245504 | 0.05947855 | -1.019696 | 0.01236688 | -1.230844 | 0.6885065 |
| ENSG00000104687.12 | GSR | 0.099146 | 0.8006286 | 0.296929 | 0.6698464 | -1.637454 | 0.6529443 | 2.565571 | 0.01775402 | 1.582525 | 0.03189165 | -1.359576 | 0.8479431 |
| ENSG00000104695.12 | PPP2CB | -0.320309 | 0.6014343 | -0.011765 | 0.6048501 | 0.11777 | 0.8656249 | 1.313352 | 0.000357422 | 1.134121 | 0.00025243 | 0.295121 | 0.8479431 |
| ENSG00000104738.16 | MCM4 | 0.158817 | 0.9406686 | -0.854185 | 0.02312488 | 0.037597 | 0.2879524 | 0.831307 | 0.1189105 | 1.248914 | 0.2616371 | -0.328024 | 0.8102467 |
| ENSG00000104756.15 | KCTD9 | 0.696976 | 1 | 0.38363 | 0.6527736 | 1.002773 | 0.5290845 | 2.445818 | 0.004939588 | 0.736308 | 1 | -0.649368 | 0.9517616 |
| ENSG00000104765.14 | BNIP3L | -0.20881 | 0.8006286 | 0.531716 | 0.1885534 | 1.26434 | 0.000585923 | -0.301331 | 0.2587462 | -0.090023 | 0.5196843 | 1.774126 | 0.9429766 |
| ENSG00000104823.8 | ECH1 | 0.101974 | 0.6214382 | 0.173322 | 0.9194312 | -0.152228 | 0.0104491 | 0.277259 | 0.2171114 | 0.323847 | 0.2709989 | 0.576362 | 0.8782618 |
| ENSG00000104824.16 | HNRNPL | -0.265485 | 0.8743571 | -0.218181 | 0.1606176 | 0.093707 | 0.1252874 | 0.54574 | 0.000351389 | 1.133659 | 0.006268713 | -0.145934 | 0.5122384 |
| ENSG00000104852.14 | SNRNP70 | -5.73148 | 0.02033342 | 0.202109 | 0.8962595 | -0.614176 | 6.48E-05 | 0.28298 | 0.002965989 | 1.016929 | 0.1496273 | 0.565626 | 0.9147227 |
| ENSG00000104853.15 | CLPTM1 | -0.232666 | 0.9653624 | 0.425661 | 0.2009498 | -0.534292 | 0.0275606 | 0.657208 | 0.05345889 | 0.579446 | 0.2310724 | 0.767741 | 0.8479431 |
| ENSG00000104859.14 | CLASRP | -0.114614 | 0.824666 | 0.788496 | 0.5953732 | 0.206813 | 0.007695605 | 1.870678 | 0.06937481 | 0.604363 | 0.07739046 | -0.360241 | 0.6829816 |
| ENSG00000104866.10 | PPP1R37 | -1.25268 | 0.6798783 | 0.220297 | 1 | -0.550528 | 0.02105855 | 1.863998 | 4.51E-09 | 0.898439 | 0.3513722 | -0.495432 | 0.9716684 |
| ENSG00000104885.17 | DOT1L | -0.253713 | 0.8739578 | -0.358339 | 0.6887002 | -0.226455 | 1 | 1.6106 | 6.46E-06 | 1.762848 | 0.001077278 | 0.030333 | 1 |
| ENSG00000104897.9 | SF3A2 | -0.970939 | 0.3098608 | 0.571655 | 0.5953732 | -1.299455 | 2.23E-09 | 0.581213 | 0.5427147 | 1.489052 | 0.1411968 | 1.840301 | 0.5733051 |
| ENSG00000104904.12 | OAZ1 | -0.179538 | 0.6214382 | 0.335403 | 0.000806275 | 0.130145 | 1 | 0.116594 | 0.4863127 | 0.413161 | 0.621723 | 0.737576 | 0.6885065 |
| ENSG00000104936.17 | DMPK | 2.487744 | 0.8006286 | 0.63857 | 0.6617347 | -0.783059 | 0.870773 | 3.804306 | 0.000118843 | 3.849749 | 0.002340489 | 0.752953 | 1 |
| ENSG00000104946.10 | TBC1D17 | -2.256865 | 0.8587992 | -0.145945 | 0.9452791 | -1.887457 | 0.02012559 | -0.370189 | 0.5069474 | 2.701711 | 0.000106562 | 0.675857 | 0.6829816 |
| ENSG00000104960.15 | PTOV1 | -0.289502 | 0.6214382 | 0.080558 | 0.9590544 | -0.442144 | 0.001328081 | 0.189845 | 0.6399232 | 0.418906 | 0.8767837 | -0.016355 | 0.8479431 |
| ENSG00000104964.14 | AES | 0.048491 | 0.6938722 | -0.022524 | 0.6527736 | -0.393044 | 0.2716766 | 1.573499 | 1.32E-13 | 1.671245 | 3.79E-11 | -0.912569 | 0.6185816 |
| ENSG00000104967.6 | NOVA2 | -5.223642 | 0.2335514 | -1.093543 | 0.5953732 | -0.078015 | 0.4064599 | 1.296994 | 0.001892524 | 0.439781 | 0.3846513 | -0.789351 | 0.6885065 |
| ENSG00000104969.9 | SGTA | -0.219839 | 0.6759734 | 0.311754 | 0.9791082 | -0.227013 | 0.3443037 | 0.792231 | 4.37E-07 | 0.948878 | 0.009561275 | -0.475066 | 0.2273839 |
| ENSG00000104979.8 | C19orf53 | -0.014401 | 0.7736255 | -0.056192 | 0.3121527 | -0.324498 | 0.02066251 | -0.123301 | 0.1277032 | 0.052744 | 0.007470624 | 0.935875 | 0.6885065 |
| ENSG00000105058.11 | FAM32A | -0.830077 | 0.7736255 | 0.2078 | 0.9768713 | -0.533233 | 0.0198345 | 0.026135 | 0.7144938 | 0.053598 | 0.119277 | 0.095526 | 0.8479431 |
| ENSG00000105063.18 | PPP6R1 | 0.196608 | 0.8006286 | 0.926258 | 0.7150943 | 0.349366 | 0.5614808 | 3.436086 | 0.01413977 | 1.367496 | 0.7377262 | -0.188462 | 1 |
| ENSG00000105171.9 | POP4 | 0.047547 | 0.7814489 | 0.388295 | 0.859711 | -0.356086 | 0.002330498 | 0.841678 | 0.237157 | -0.678379 | 0.01947854 | -3.212748 | 0.3746505 |
| ENSG00000105176.17 | URI1 | -0.618904 | 0.9827503 | -0.061002 | 0.2401656 | 0.138684 | 0.6701871 | 0.705499 | 0.009883487 | 0.419884 | 0.6659625 | -1.355075 | 0.6829816 |
| ENSG00000105185.11 | PDCD5 | -0.182651 | 0.9282145 | -0.157179 | 0.6527736 | -0.313072 | 0.000342868 | -0.210574 | 0.1161075 | -0.374719 | 2.97E-05 | -0.725942 | 0.5733051 |
| ENSG00000105186.14 | ANKRD27 | 2.700749 | 0.8587992 | 0.663321 | 0.7383998 | 0.053665 | 0.5290845 | 2.025696 | 0.007812163 | 2.009416 | 0.5196843 | -0.840472 | 0.9270789 |
| ENSG00000105193.8 | RPS16 | -0.320252 | 0.6214382 | 0.074742 | 0.5010139 | 0.076828 | 0.1215462 | -0.121406 | 2.22E-06 | -0.29031 | 1.10E-06 | -0.052271 | 0.3565625 |
| ENSG00000105221.16 | AKT2 | -6.050636 | 1 | -0.239375 | 0.8263112 | -0.076037 | 0.6075097 | 1.433535 | 0.000203891 | 1.449778 | 0.2128722 | -1.20388 | 1 |
| ENSG00000105245.9 | NUMBL | -3.457332 | 0.6214382 | -0.473234 | 0.329874 | -0.512152 | 0.1825848 | 0.912185 | 0.006345602 | 2.045555 | 0.01299526 | 1.459821 | 0.6885065 |
| ENSG00000105251.10 | SHD | -1.8722 | 0.03954376 | -0.402278 | 0.8867595 | 0.469611 | 0.274519 | 1.070875 | 0.3610019 | 0.632619 | 1 | 1.590024 | 0.9547013 |
| ENSG00000105254.11 | TBCB | -0.916963 | 0.1835496 | -0.095687 | 0.6435797 | -0.606729 | 4.42E-09 | -0.087109 | 0.03385605 | 0.392739 | 0.0100807 | -0.475477 | 0.05538244 |
| ENSG00000105258.8 | POLR2I | -0.088289 | 0.8006286 | 0.012534 | 0.8263112 | -0.422007 | 6.39E-05 | 0.10222 | 0.4730254 | -0.615311 | 0.01127024 | -1.788907 | 0.6710264 |
| ENSG00000105290.11 | APLP1 | -0.172157 | 1 | 0.075967 | 0.9258417 | -0.440693 | 0.2112202 | 0.845183 | 0.04923712 | 0.816785 | 5.84E-05 | -0.484178 | 0.5841302 |
| ENSG00000105323.16 | HNRNPUL1 | 0.315781 | 0.903671 | -0.281452 | 0.1356646 | 0.384609 | 0.2113924 | 0.71774 | 9.13E-05 | 0.930655 | 0.000280062 | 1.048555 | 0.947968 |
| ENSG00000105364.13 | MRPL4 | 0.183452 | 0.6014343 | 0.066396 | 0.8953649 | -0.942613 | 0.001806849 | 0.567533 | 0.7962629 | 0.433126 | 0.2470944 | -0.089144 | 0.9533243 |
| ENSG00000105372.6 | RPS19 | -0.50915 | 1.91E-09 | -0.100158 | 0.06457881 | -0.138304 | 0.01921893 | -0.090764 | 0.05560288 | -0.173314 | 0.000452323 | -0.011577 | 0.5344421 |
| ENSG00000105373.18 | GLTSCR2 | 0.276348 | 0.714717 | 0.206279 | 0.912465 | -0.155423 | 0.0002759 | -0.004964 | 0.9642557 | 1.084294 | 0.7586399 | 0.697526 | 0.9919109 |
| ENSG00000105379.9 | ETFB | 0.141985 | 0.7345905 | 0.03157 | 0.8953649 | -0.199397 | 0.1595169 | -0.236828 | 0.008607601 | -0.080116 | 0.9783895 | -1.931594 | 0.9177601 |
| ENSG00000105404.10 | RABAC1 | 0.985428 | 0.9426668 | -0.002342 | 0.5981187 | -0.300041 | 2.47E-05 | -0.277328 | 0.000289925 | -0.38773 | 0.1386512 | 0.69975 | 1 |
| ENSG00000105409.15 | ATP1A3 | 1.96356 | 0.6214382 | 1.158316 | 1 | 1.017009 | 0.7437308 | 0.963734 | 0.001430378 | 0.929085 | 0.01231135 | -0.370105 | 0.3746505 |
| ENSG00000105419.17 | MEIS3 | 0.667503 | 0.8006286 | 0.201517 | 0.7548431 | 0.264448 | 0.6928639 | 2.075864 | 1.18E-06 | 0.932506 | 0.2341319 | -1.146578 | 0.9250166 |
| ENSG00000105426.14 | PTPRS | 1.019483 | 0.01246479 | -0.402797 | 0.09704869 | 0.228249 | 0.4450546 | 1.01906 | 5.37E-10 | 1.060606 | 0.000256485 | 0.310577 | 0.6885065 |
| ENSG00000105429.12 | MEGF8 | -3.264344 | 0.3098608 | -0.633386 | 0.4487482 | -0.505684 | 0.5412517 | 0.916563 | 0.001378031 | 1.084833 | 0.02943383 | 0.921487 | 0.929628 |
| ENSG00000105443.13 | CYTH2 | 1.325371 | 0.8901384 | -0.245171 | 0.8763548 | -0.408892 | 0.07373437 | 0.924022 | 0.03351087 | 1.143839 | 0.2206737 | 1.053374 | 1 |
| ENSG00000105464.3 | GRIN2D | -1.916556 | 0.6214382 | -2.254755 | 0.4158789 | -1.61116 | 0.5290845 | 0.696157 | 0.8919817 | 0.606047 | 0.003642399 | #NAME? | 0.5344421 |
| ENSG00000105472.12 | CLEC11A | 0.869996 | 0.714717 | -0.293266 | 0.5394427 | -0.966823 | 0.04639883 | -0.962901 | 0.2068989 | 1.499384 | 0.7302514 | 1.445862 | 0.7408601 |
| ENSG00000105486.13 | LIG1 | -0.561442 | 1 | -1.021553 | 0.8703522 | 0.151106 | 0.5302836 | 2.231992 | 0.001675881 | 1.099141 | 0.3363434 | 0.121897 | 0.9663203 |
| ENSG00000105518.13 | TMEM205 | -0.106966 | 0.8742761 | -0.008275 | 0.8278555 | -0.112694 | 0.04532096 | -0.347335 | 0.007120083 | -0.065605 | 0.5759119 | -0.48364 | 0.6885065 |
| ENSG00000105559.11 | PLEKHA4 | 0.263737 | 1 | -0.240761 | 0.2571098 | -0.776494 | 0.02102919 | 0.768718 | 0.4269893 | 1.128965 | 0.8883312 | 0.73406 | 0.6885065 |
| ENSG00000105568.17 | PPP2R1A | 0.667249 | 0.004886376 | -0.000746 | 0.6527736 | -0.299449 | 0.5825762 | 0.045954 | 0.384552 | 0.309413 | 0.8021966 | -0.544783 | 0.6829816 |
| ENSG00000105576.15 | TNPO2 | -0.130901 | 0.9744865 | 0.324015 | 0.7881421 | -0.027137 | 0.5581415 | 0.445927 | 0.001550612 | 1.183652 | 0.02780984 | -0.510249 | 0.6885065 |
| ENSG00000105583.9 | WDR83OS | 0.274741 | 0.1057507 | -0.040669 | 0.5886575 | -0.212699 | 0.01840642 | -0.218725 | 0.008171374 | 0.051033 | 0.00041383 | 0.001332 | 0.5344421 |
| ENSG00000105605.7 | CACNG7 | -1.930879 | 0.6214382 | 0.464699 | 0.7050684 | -0.323507 | 1.51E-05 | 0.985363 | 0.000688376 | 0.580254 | 0.1635289 | -1.341138 | 0.922577 |
| ENSG00000105617.3 | LENG1 | -0.832878 | 0.6214382 | 0.345884 | 1 | 0.248098 | 0.000633168 | -0.383714 | 0.4040131 | 0.168694 | 0.6192622 | -3.445951 | 0.8260537 |
| ENSG00000105640.12 | RPL18A | -1.160508 | 0.3535686 | 0.261843 | 0.000277555 | 0.034728 | 0.9711943 | 0.08638 | 0.3541189 | -0.031321 | 0.0464589 | 0.486168 | 0.7891857 |
| ENSG00000105647.14 | PIK3R2 | -2.565209 | 0.3946735 | 0.02211 | 0.9420234 | -1.170503 | 0.2430735 | 2.566842 | 0.001301183 | 1.986594 | 0.03622093 | -2.098952 | 0.8539279 |
| ENSG00000105649.9 | RAB3A | -1.201621 | 1 | -0.730256 | 0.2250307 | -1.262262 | 0.004396714 | 1.529339 | 0.1557388 | 0.505243 | 0.621723 | -0.27552 | 0.9147227 |
| ENSG00000105669.12 | COPE | 0.392746 | 0.9054014 | 0.649692 | 0.008657586 | -0.293029 | 2.78E-05 | 0.444176 | 0.5376676 | -0.051021 | 0.04615617 | -0.038556 | 0.9153161 |
| ENSG00000105677.11 | TMEM147 | 0.542196 | 0.666472 | 0.16509 | 0.6527736 | -0.376418 | 5.10E-05 | -0.30564 | 0.000847916 | -0.29546 | 0.01560716 | -1.408766 | 0.5344421 |
| ENSG00000105696.8 | TMEM59L | 0.537481 | 0.6214382 | -0.105686 | 0.6527736 | -1.264238 | 0.3947164 | 0.164897 | 0.8094417 | 1.988118 | 0.02757863 | -1.089013 | 0.6885065 |
| ENSG00000105698.15 | USF2 | -0.332475 | 0.8006286 | -0.601929 | 0.3977905 | -0.77947 | 0.07810551 | 0.803068 | 3.96E-05 | 1.616655 | 3.26E-05 | -0.035496 | 0.9377765 |
| ENSG00000105701.15 | FKBP8 | -0.881051 | 0.6214382 | 0.30008 | 0.3121527 | -0.641534 | 0.01020352 | 1.300945 | 0.04811661 | 1.497156 | 0.3321332 | 0.140587 | 0.6885065 |
| ENSG00000105722.9 | ERF | 0.865839 | 0.8220319 | 0.025906 | 0.9449715 | -0.64251 | 0.03037326 | 0.855735 | 0.000543727 | 0.962037 | 0.627383 | -0.402203 | 1 |
| ENSG00000105737.9 | GRIK5 | 1.878439 | 0.714717 | 1.180404 | 0.4001101 | 0.411764 | 0.9067415 | 1.997178 | 1.05E-06 | 2.742187 | 0.001005217 | 1.114364 | 0.7193722 |
| ENSG00000105767.2 | CADM4 | -0.490848 | 0.8587992 | -0.382963 | 0.3494625 | -0.718503 | 0.000443129 | 1.547264 | 1.06E-16 | 1.057522 | 0.05740199 | 0.348904 | 0.6885065 |
| ENSG00000105778.17 | AVL9 | -0.581521 | 0.6214382 | 0.059973 | 0.9364384 | 0.703854 | 0.4602617 | 1.132719 | 0.04242491 | -0.040089 | 0.8216963 | 0.784067 | 0.8479431 |
| ENSG00000105810.9 | CDK6 | 2.316719 | 0.6214382 | -0.592394 | 0.02491789 | 1.025623 | 0.001204 | 0.961021 | 6.34E-05 | -0.083361 | 0.5302919 | -0.795521 | 0.6885065 |
| ENSG00000105821.14 | DNAJC2 | -0.966158 | 0.6817555 | 0.297259 | 0.9281383 | 0.371901 | 0.05977368 | 0.328267 | 0.7854938 | 0.166422 | 0.0454681 | -0.582317 | 0.6829816 |
| ENSG00000105835.11 | NAMPT | 0.134214 | 0.884816 | 0.044164 | 0.9977474 | -0.290697 | 0.01982567 | 0.154252 | 0.02946863 | 0.752984 | 0.8507684 | 1.332906 | 0.6885065 |
| ENSG00000105849.5 | TWISTNB | -0.019424 | 0.6214382 | 0.137861 | 0.9452791 | 0.585765 | 0.004494928 | 0.221452 | 0.6625684 | 0.150484 | 0.4020737 | -0.436493 | 0.6885065 |
| ENSG00000105854.12 | PON2 | -2.119431 | 0.07017518 | -0.27616 | 0.1548653 | 0.256946 | 0.5018872 | 0.155486 | 0.0491633 | -0.307398 | 0.02704086 | 0.436851 | 0.8949489 |
| ENSG00000105855.9 | ITGB8 | 0.699377 | 0.5711449 | -0.41427 | 0.03619211 | 1.321709 | 4.55E-08 | 1.145241 | 2.12E-05 | 1.17135 | 0.7233645 | -0.566902 | 0.8722482 |
| ENSG00000105887.10 | MTPN | -0.837626 | 0.5711449 | -0.124524 | 0.6527736 | 0.421196 | 4.80E-07 | 0.36777 | 0.6996942 | -0.227086 | 0.05223546 | -0.112086 | 0.6885065 |
| ENSG00000105894.11 | PTN | -0.610454 | 0.6214382 | -0.214605 | 0.02139399 | 0.11037 | 0.5290845 | 0.072722 | 0.2010829 | -0.112059 | 0.001883415 | 0.465325 | 0.8419703 |
| ENSG00000105939.12 | ZC3HAV1 | 1.91127 | 0.7736255 | -0.054242 | 0.6527736 | 0.747082 | 0.2913878 | 1.689722 | 0.0410256 | -0.795674 | 0.4883149 | -1.912522 | 1 |
| ENSG00000105953.14 | OGDH | 1.363299 | 0.1480428 | -0.551826 | 0.5012872 | 0.464899 | 1 | 1.152708 | 0.01227431 | 1.193411 | 0.07956883 | 0.023404 | 0.8479431 |
| ENSG00000105968.18 | H2AFV | -0.890037 | 0.9667813 | -0.03465 | 0.4491636 | -0.246238 | 0.01339109 | 0.558091 | 0.1994706 | 0.16471 | 0.5196843 | 0.895508 | 0.9071717 |
| ENSG00000105976.14 | MET | 0.496574 | 1 | -0.948454 | 0.3381481 | 2.323105 | 0.934576 | 1.370378 | 4.33E-06 | -1.054371 | 4.97E-13 | -0.042615 | 0.5223622 |
| ENSG00000105983.19 | LMBR1 | 0.670354 | 0.6014343 | -0.21284 | 0.9002426 | 0.362023 | 0.8969395 | 1.422447 | 0.001660058 | 1.046946 | 0.3645298 | -0.091181 | 0.9399525 |
| ENSG00000105993.14 | DNAJB6 | 0.013333 | 1 | 0.116375 | 1 | 0.334088 | 0.04342208 | 1.112113 | 0.613586 | 1.121461 | 0.006705117 | 0.3862 | 0.4004082 |
| ENSG00000106028.10 | SSBP1 | -0.2742 | 0.6570323 | 0.01887 | 0.8953649 | -0.157731 | 7.92E-05 | 0.171801 | 0.3749933 | -0.050197 | 0.01523426 | -0.46545 | 0.3358843 |
| ENSG00000106052.13 | TAX1BP1 | 0.089647 | 0.6542137 | 0.235234 | 0.8944225 | 0.961251 | 8.46E-11 | 0.752539 | 0.9133426 | -0.280246 | 0.02559642 | 0.109378 | 0.6885065 |
| ENSG00000106105.13 | GARS | -0.642577 | 0.3705372 | 0.091835 | 0.6170592 | 0.458441 | 0.06504457 | 0.692321 | 0.000107693 | 0.509528 | 0.5759579 | 0.958491 | 0.8479431 |
| ENSG00000106144.19 | CASP2 | -1.382771 | 0.8006286 | -0.41698 | 0.5154617 | 0.161751 | 0.1333632 | 1.493799 | 0.003720848 | 1.405915 | 0.06956234 | 0.085654 | 0.6885065 |
| ENSG00000106153.12 | CHCHD2 | -1.050451 | 0.1575216 | 0.247534 | 9.28E-16 | -0.290247 | 1.19E-06 | 0.081738 | 0.03651792 | -0.219272 | 2.83E-07 | -0.031463 | 0.06476298 |
| ENSG00000106211.8 | HSPB1 | -1.731593 | 2.94E-05 | -0.190119 | 0.5334324 | -0.641385 | 5.47E-10 | 0.105192 | 0.008309395 | 0.5653 | 0.759959 | -0.368157 | 0.8427621 |
| ENSG00000106244.12 | PDAP1 | -0.543521 | 0.8240785 | -0.212867 | 0.1885534 | 0.278571 | 0.706876 | -0.022126 | 0.08529855 | -0.133822 | 0.01286342 | 0.283826 | 0.5066477 |
| ENSG00000106261.16 | ZKSCAN1 | -0.81232 | 0.7676825 | -0.290932 | 0.24796 | 0.27088 | 0.037209 | 0.588711 | 1.70E-06 | 1.096854 | 0.0120402 | 1.189592 | 0.6885065 |
| ENSG00000106263.17 | EIF3B | 0.165358 | 0.8587992 | 0.195797 | 0.6527736 | -0.104304 | 0.8613757 | 1.080734 | 1.36E-07 | 0.896714 | 0.04948871 | -0.46016 | 0.6885065 |
| ENSG00000106268.15 | NUDT1 | -2.083252 | 0.2674063 | -0.376239 | 0.5953732 | -0.739893 | 0.000521203 | -0.19103 | 0.9795096 | -0.053947 | 0.4408957 | 0.334045 | 0.6885065 |
| ENSG00000106278.11 | PTPRZ1 | -0.201695 | 0.4246706 | -0.31607 | 3.92E-05 | 0.868627 | 1.49E-16 | 0.61563 | 0.000419861 | 0.479034 | 0.7329173 | -0.000887 | 0.6885065 |
| ENSG00000106299.7 | WASL | -0.619028 | 0.7737941 | 0.218767 | 0.8500718 | 0.779019 | 1.02E-06 | 1.05068 | 2.76E-06 | 0.385781 | 0.8021966 | 0.651573 | 1 |
| ENSG00000106305.9 | AIMP2 | 0.0599 | 0.9410335 | -0.261283 | 0.5010139 | -0.455238 | 0.004950796 | 0.2001 | 0.1141629 | 0.161258 | 0.2364542 | 0.251163 | 0.6829816 |
| ENSG00000106344.8 | RBM28 | 0.235104 | 0.714717 | 0.434942 | 0.7765491 | 0.431356 | 0.4064599 | 1.516218 | 0.001484126 | -0.184135 | 0.1743517 | -1.498958 | 0.5344421 |
| ENSG00000106346.11 | USP42 | 0.213303 | 0.9438503 | -0.317103 | 1 | -0.495455 | 0.5457055 | 1.447459 | 0.01816429 | 1.703527 | 0.04367249 | -1.745485 | 0.6885065 |
| ENSG00000106355.9 | LSM5 | 0.151772 | 0.7204951 | -0.211858 | 0.2374402 | -0.110211 | 0.08636277 | -0.315316 | 0.005431497 | -0.018964 | 0.00044159 | 0.676942 | 0.985679 |
| ENSG00000106366.8 | SERPINE1 | -0.094543 | 0.6014343 | 0.343884 | 0.02260499 | -0.422224 | 0.01272831 | 0.829447 | 0.02533603 | 0.509947 | 0.3974318 | 0.621272 | 0.5733051 |
| ENSG00000106367.13 | AP1S1 | 0.342431 | 0.529676 | 0.359516 | 0.6045862 | -0.17573 | 4.82E-05 | 0.233102 | 0.835045 | -0.15659 | 0.04937944 | -0.297484 | 0.6206888 |
| ENSG00000106397.11 | PLOD3 | -0.562468 | 0.8639532 | -0.17203 | 0.7201934 | -0.430443 | 0.3872096 | 0.322688 | 0.03329343 | 1.541754 | 0.04819759 | 0.10829 | 0.8479431 |
| ENSG00000106399.11 | RPA3 | 0.81866 | 0.8291333 | 0.663307 | 0.3969148 | -0.136596 | 0.5516645 | -0.472144 | 0.01671469 | 0.249485 | 0.1680675 | -0.855771 | 0.9303247 |
| ENSG00000106400.11 | ZNHIT1 | -0.239901 | 0.6494684 | 0.096717 | 0.9856582 | 0.142801 | 0.813417 | 0.280827 | 0.1875735 | -0.558512 | 0.00018356 | 0.249091 | 0.9270789 |
| ENSG00000106415.12 | GLCCI1 | -1.36609 | 0.7736255 | 0.798642 | 1 | -0.221881 | 0.4064599 | 2.876766 | 0.03254613 | 0.012874 | 0.2518976 | 0.663331 | 1 |
| ENSG00000106460.18 | TMEM106B | -0.355605 | 1 | -0.143504 | 0.8064295 | 1.159874 | 0.002952376 | 0.367393 | 1 | 1.02326 | 0.9138243 | -0.767702 | 0.8614062 |
| ENSG00000106524.8 | ANKMY2 | 2.782238 | 0.6214382 | -0.265451 | 0.6527736 | 1.167236 | 0.1686141 | 0.824086 | 0.03363906 | 2.276792 | 0.000174021 | -0.380391 | 0.8334258 |
| ENSG00000106554.11 | CHCHD3 | 0.187288 | 0.6214382 | -0.546993 | 0.02797767 | 0.383323 | 0.2801232 | 0.187127 | 0.6861168 | 0.279345 | 0.8075275 | 0.079348 | 0.6267144 |
| ENSG00000106588.10 | PSMA2 | -0.84946 | 0.1743759 | 0.206294 | 0.468656 | -0.421687 | 2.06E-08 | -0.032327 | 0.2054284 | -0.474608 | 2.89E-10 | -0.24964 | 0.1508459 |
| ENSG00000106609.16 | TMEM248 | -2.027689 | 0.06849962 | 0.145482 | 1 | 0.277002 | 0.1951855 | 0.625281 | 0.01826913 | 0.840647 | 0.510841 | -0.539931 | 0.8749632 |
| ENSG00000106624.8 | AEBP1 | -1.255039 | 0.6214382 | -0.092803 | 0.8085222 | -1.76179 | 0.001017927 | 0.317229 | 0.1287006 | 1.388017 | 0.02779405 | -0.45709 | 1 |
| ENSG00000106635.7 | BCL7B | 0.45521 | 0.3379933 | -0.08958 | 0.937266 | -0.241125 | 0.008002535 | 0.730739 | 0.1232005 | 0.910088 | 0.5196843 | -0.261455 | 0.854875 |
| ENSG00000106636.7 | YKT6 | 0.235411 | 0.9268531 | 0.219559 | 0.1402928 | -0.037781 | 0.5290845 | 0.45098 | 0.002169083 | 0.403052 | 0.9234314 | 0.725917 | 0.7341273 |
| ENSG00000106665.15 | CLIP2 | -1.239396 | 0.6214382 | -0.039347 | 0.5465944 | -0.439969 | 0.5030582 | 0.931458 | 0.001345975 | 0.639159 | 0.2027773 | -1.13033 | 0.9576568 |
| ENSG00000106682.14 | EIF4H | -0.34075 | 0.8189164 | -0.056562 | 0.618047 | 0.304034 | 0.1372475 | 0.524354 | 0.02339529 | 0.440282 | 0.8493095 | -0.223873 | 0.6576742 |
| ENSG00000106701.11 | FSD1L | 1.759828 | 0.6214382 | -2.116463 | 0.8691081 | -1.387054 | 0.7985849 | 1.122818 | 0.9185245 | 1.742578 | 0.004919348 | -2.046338 | 1 |
| ENSG00000106723.16 | SPIN1 | 0.139044 | 0.8276028 | -0.177346 | 0.5010139 | 0.741135 | 0.3396353 | 0.966103 | 0.02823854 | 0.660681 | 0.2900175 | -1.207345 | 0.9171058 |
| ENSG00000106771.12 | TMEM245 | 0.283753 | 0.9686406 | 0.108347 | 0.6984883 | 0.708863 | 0.2348326 | 1.720311 | 0.000289796 | 1.86685 | 0.1806314 | 0.504667 | 0.8479431 |
| ENSG00000106772.17 | PRUNE2 | -0.268152 | 0.563505 | 0.45654 | 0.8545414 | 3.233227 | 0.1992941 | 1.570561 | 0.08014509 | 3.799435 | 0.007184823 | -1.012218 | 0.3769331 |
| ENSG00000106799.12 | TGFBR1 | 1.481318 | 0.714717 | -0.53994 | 0.5834943 | 0.272029 | 1 | -0.068653 | 0.01020107 | 0.771937 | 0.2637008 | 1.651428 | 0.9517616 |
| ENSG00000106803.9 | SEC61B | 0.207346 | 0.305762 | 0.156707 | 0.8953649 | -0.003105 | 0.5632235 | -0.291123 | 0.000135252 | -0.360771 | 0.000811934 | 0.244535 | 0.9181784 |
| ENSG00000106948.16 | AKNA | -0.301592 | 0.9198482 | 2.515003 | 0.6527736 | 1.397442 | 0.7796906 | 6.911043 | 0.000182145 | 3.708881 | 0.06809273 | 3.285537 | 0.6885065 |
| ENSG00000106976.19 | DNM1 | 0.920345 | 0.8587992 | 0.317234 | 0.8804703 | -0.489454 | 0.1048414 | 2.395595 | 2.80E-05 | 2.713544 | 9.78E-05 | 0.092549 | 0.9147227 |
| ENSG00000106993.11 | CDC37L1 | 0.278492 | 0.7000696 | -0.227212 | 0.7063369 | 0.857159 | 0.01393234 | 1.228874 | 0.7374141 | 0.699801 | 1 | 0.347344 | 1 |
| ENSG00000107020.9 | PLGRKT | -0.58043 | 0.6130622 | -0.214466 | 0.6527736 | 0.307112 | 0.5480968 | 0.111565 | 0.01207805 | -0.503101 | 0.03345076 | 0.642213 | 0.9875948 |
| ENSG00000107130.9 | NCS1 | -0.541601 | 0.6214382 | -0.401803 | 0.6527736 | 0.433539 | 0.5290845 | 1.511063 | 0.003228316 | 0.805006 | 0.4703084 | 1.62676 | 0.6829816 |
| ENSG00000107140.15 | TESK1 | 0.588288 | 0.6214382 | 0.300855 | 0.6527736 | -0.734492 | 0.5266754 | 1.717043 | 0.008473218 | -0.79976 | 1 | 0.634915 | 0.6885065 |
| ENSG00000107164.15 | FUBP3 | 0.678722 | 0.529676 | -0.030971 | 0.6905553 | -0.033322 | 0.6419675 | 2.100182 | 6.12E-06 | 1.942171 | 0.06271733 | -0.077935 | 0.701409 |
| ENSG00000107175.10 | CREB3 | 0.221433 | 0.8006286 | 0.198719 | 0.6527736 | -0.773933 | 0.01526418 | -0.67309 | 0.3377327 | 0.732165 | 0.1219119 | -0.737719 | 0.6778167 |
| ENSG00000107223.12 | EDF1 | -1.659603 | 0.4764974 | -0.051333 | 0.1356646 | -0.076507 | 0.007078775 | 0.022204 | 0.1917431 | -0.40089 | 1.29E-05 | 0.248574 | 0.6885065 |
| ENSG00000107281.9 | NPDC1 | 0.113923 | 0.9183025 | -0.103539 | 0.5919292 | -1.015294 | 6.72E-05 | 1.674939 | 0.00100326 | 1.5178 | 0.002918613 | -0.650615 | 0.9595173 |
| ENSG00000107290.13 | SETX | -0.321359 | 0.980174 | -0.600425 | 0.8800335 | 1.10695 | 3.75E-06 | 1.32685 | 0.01064038 | 1.542509 | 0.1194023 | 0.998733 | 0.6885065 |
| ENSG00000107317.11 | PTGDS | -1.364823 | 0.4469329 | -1.218995 | 1 | -1.492074 | 0.003635137 | 0.378843 | 0.2047258 | 0.519536 | 0.4314015 | -0.875669 | 0.795266 |
| ENSG00000107341.4 | UBE2R2 | 0.811675 | 0.7090887 | -0.340858 | 0.09142325 | 0.730325 | 0.09123437 | 1.420223 | 0.01663572 | 1.285664 | 0.01550204 | 0.285035 | 0.6885065 |
| ENSG00000107362.13 | ABHD17B | 3.965446 | 0.6214382 | -0.160485 | 0.859711 | -0.308173 | 0.6011545 | 2.197079 | 0.01909942 | 0.804011 | 0.1395844 | 1.619262 | 0.6885065 |
| ENSG00000107372.12 | ZFAND5 | 0.475231 | 0.8024091 | -0.020182 | 0.8690735 | 0.384447 | 0.2348326 | 0.980445 | 0.002813711 | 0.681326 | 0.2640175 | 0.261285 | 0.9161136 |
| ENSG00000107581.12 | EIF3A | 0.48261 | 0.00219167 | -0.172695 | 0.3776134 | 0.747257 | 6.18E-07 | 0.616523 | 0.04773307 | 0.645301 | 0.7894421 | 0.256193 | 0.8479431 |
| ENSG00000107625.12 | DDX50 | 0.807968 | 0.8353323 | 0.325646 | 0.8379413 | 1.188773 | 0.01393234 | 0.781789 | 0.746754 | -0.018298 | 0.778506 | -0.00384 | 0.6767422 |
| ENSG00000107643.15 | MAPK8 | 0.433913 | 0.8421394 | 0.302706 | 0.6527736 | -2.429497 | 0.4064599 | 1.367493 | 0.006264535 | 1.133233 | 0.6093143 | 0.858498 | 0.8479431 |
| ENSG00000107651.12 | SEC23IP | -0.31941 | 0.8006286 | 0.322985 | 0.5240506 | -0.194926 | 0.2283118 | 0.913711 | 0.01962794 | -0.067968 | 0.6185494 | -0.642084 | 0.6885065 |
| ENSG00000107672.14 | NSMCE4A | 2.330683 | 0.714717 | 0.868346 | 0.6817396 | -1.44158 | 0.003862177 | -1.40735 | 0.05012515 | 0.007045 | 0.7286448 | 0.007676 | 0.938661 |
| ENSG00000107731.12 | UNC5B | 2.64295 | 0.6014343 | 1.103651 | 0.7526929 | 0.797136 | 0.6827367 | 2.661024 | 0.009744773 | 1.902762 | 0.6302173 | 1.805231 | 0.6829816 |
| ENSG00000107815.7 | C10orf2 | -2.398715 | 0.03700106 | -0.743433 | 0.6031515 | 1.4112 | 0.1367134 | -1.116963 | 0.6914224 | 0.064399 | 0.6956242 | -0.724033 | 1 |
| ENSG00000107862.4 | GBF1 | -1.414489 | 0.8006286 | 0.306579 | 1 | 0.055877 | 0.8068696 | 0.894098 | 0.04663016 | 1.848397 | 0.1089968 | -0.320195 | 0.9631343 |
| ENSG00000107863.16 | ARHGAP21 | -0.071636 | 0.6744243 | -0.167859 | 0.6527736 | 0.986149 | 0.04634161 | 1.316839 | 0.000182652 | 0.906803 | 0.8069853 | -0.204343 | 0.786868 |
| ENSG00000107872.12 | FBXL15 | -0.059915 | 0.9678883 | 0.70482 | 0.7955884 | -0.932556 | 0.03192898 | -0.466544 | 0.2696559 | -0.168806 | 0.6258535 | -2.014348 | 0.9539924 |
| ENSG00000107890.16 | ANKRD26 | -0.414685 | 0.563505 | -0.378968 | 0.6527736 | 0.28827 | 0.8837048 | 1.571999 | 0.004010145 | 1.473812 | 0.4481846 | 1.38991 | 0.9062221 |
| ENSG00000107929.14 | LARP4B | -0.402194 | 0.6214382 | 0.337881 | 0.9452791 | -1.0725 | 0.9619523 | 0.750361 | 0.03020277 | 0.594723 | 0.7067087 | -2.524367 | 0.9147227 |
| ENSG00000107937.18 | GTPBP4 | -0.04484 | 0.5711449 | 0.154072 | 0.5747259 | 0.919669 | 0.04944273 | 0.610675 | 0.9309759 | 0.617359 | 0.7951357 | -0.03989 | 0.6885065 |
| ENSG00000107949.16 | BCCIP | -0.32397 | 0.8916027 | -0.163632 | 0.4487482 | -0.24226 | 0.4611303 | -0.065993 | 0.033497 | -0.277195 | 0.08657607 | -0.322341 | 0.6885065 |
| ENSG00000108021.19 | FAM208B | 0.086627 | 0.714717 | -0.299623 | 0.8263112 | 0.648446 | 0.2716766 | 1.954517 | 0.01983441 | 2.2665 | 0.03024717 | -0.47655 | 1 |
| ENSG00000108055.9 | SMC3 | -0.108057 | 0.7736255 | 0.428773 | 0.949642 | 1.076238 | 1.69E-07 | 1.094734 | 0.184613 | 0.173982 | 0.4478429 | -1.797811 | 0.01138072 |
| ENSG00000108061.11 | SHOC2 | 0.034048 | 0.6274396 | -0.304685 | 0.6737951 | 0.227114 | 0.0443262 | -1.044974 | 0.07150683 | -2.061328 | 0.3375835 | 0.062942 | 0.6885065 |
| ENSG00000108106.13 | UBE2S | -1.118955 | 0.4455568 | 0.31026 | 0.592336 | -2.12141 | 2.89E-23 | 0.930738 | 6.96E-06 | 1.424255 | 2.92E-06 | -0.846171 | 0.4061635 |
| ENSG00000108107.12 | RPL28 | -0.206747 | 0.8339661 | 0.151557 | 0.3791327 | -0.586536 | 4.74E-15 | 0.118332 | 0.7144938 | -0.193032 | 1.69E-05 | 0.398807 | 0.6231061 |
| ENSG00000108175.16 | ZMIZ1 | -1.214526 | 0.5858981 | -0.209543 | 0.95708 | -0.190651 | 0.9522762 | 1.482033 | 0.000559716 | 1.084933 | 0.000752491 | -0.797549 | 0.9363384 |
| ENSG00000108256.8 | NUFIP2 | -0.586407 | 0.8114707 | -0.112159 | 0.7257098 | 1.274816 | 1.74E-05 | 1.067938 | 3.72E-05 | 1.040363 | 0.01078126 | 0.431987 | 0.8479431 |
| ENSG00000108298.9 | RPL19 | -0.152374 | 0.2656681 | -0.029441 | 0.8264635 | 0.076156 | 0.5290845 | -0.216353 | 8.85E-05 | -0.029471 | 0.1580959 | -0.02145 | 0.3746505 |
| ENSG00000108312.14 | UBTF | -0.502608 | 0.5048401 | -0.143476 | 0.5222411 | 0.446421 | 0.8382626 | 0.864816 | 0.000366817 | 1.191256 | 0.6907982 | -0.929158 | 0.426223 |
| ENSG00000108344.14 | PSMD3 | -0.603824 | 0.9282145 | 0.05234 | 0.8736558 | -0.31096 | 0.004150655 | 0.291042 | 0.8622319 | 0.566234 | 0.6112598 | 0.928013 | 0.6885065 |
| ENSG00000108349.14 | CASC3 | 0.065252 | 0.8006286 | 0.554275 | 0.77388 | 0.275068 | 0.3564736 | 1.188811 | 0.01276329 | 0.723928 | 0.05208149 | -0.496504 | 0.6517664 |
| ENSG00000108389.9 | MTMR4 | -1.230409 | 0.9282145 | 0.41084 | 0.8217443 | -0.077961 | 0.7832847 | 1.874111 | 3.19E-06 | 1.0081 | 0.1172289 | 1.556747 | 0.8749632 |
| ENSG00000108395.13 | TRIM37 | 0.657054 | 1 | -0.268378 | 0.5953732 | 0.432549 | 0.04165667 | 0.014353 | 0.3116077 | 0.371596 | 0.1342636 | 0.91857 | 0.8183573 |
| ENSG00000108424.9 | KPNB1 | -1.442447 | 5.98E-07 | -0.340194 | 0.04907665 | 0.793117 | 2.24E-07 | 0.886965 | 0.001044563 | 0.280625 | 0.6236538 | -1.258884 | 0.000939089 |
| ENSG00000108443.13 | RPS6KB1 | 0.224978 | 0.956309 | 0.294602 | 0.704767 | 1.02563 | 9.61E-05 | 2.002969 | 0.005649521 | 1.199312 | 0.4671878 | 1.333305 | 0.6885065 |
| ENSG00000108448.20 | TRIM16L | 0.4562 | 0.9183025 | 0.379498 | 0.6527736 | -0.92793 | 0.003493635 | 1.560249 | 0.0425712 | 1.059985 | 0.7232762 | 2.461493 | 0.6885065 |
| ENSG00000108468.14 | CBX1 | -0.088212 | 0.5428806 | -0.349994 | 0.5012872 | 0.043548 | 0.6686219 | -0.022027 | 0.2335397 | -0.672768 | 0.001277837 | -0.530366 | 0.2936114 |
| ENSG00000108510.9 | MED13 | -0.033035 | 0.6014343 | -0.227173 | 0.7383998 | 0.865927 | 0.01022277 | 1.803919 | 1.27E-08 | 0.413434 | 0.08729226 | 0.196574 | 0.7767294 |
| ENSG00000108518.7 | PFN1 | 0.260945 | 8.02E-05 | 0.052672 | 0.9790409 | -0.931499 | 9.40E-45 | 0.103763 | 0.03519259 | -0.176996 | 0.000696277 | -0.270823 | 0.2068179 |
| ENSG00000108528.13 | SLC25A11 | -0.398731 | 0.6214382 | -0.132667 | 0.4375617 | -0.48724 | 0.04986294 | 1.064656 | 0.2802171 | -0.0582 | 0.5196843 | -1.358113 | 0.02166254 |
| ENSG00000108551.4 | RASD1 | 1.194574 | 7.45E-05 | 0.118257 | 0.6527736 | -0.67911 | 0.005073901 | -0.67126 | 0.05637094 | 2.609766 | 0.02356055 | 0.071839 | 1 |
| ENSG00000108557.17 | RAI1 | 0.111421 | 0.9047689 | 1.224135 | 0.5010139 | -0.021003 | 0.6698942 | 0.278614 | 0.4614976 | 1.622776 | 9.25E-06 | -3.319583 | 0.8949489 |
| ENSG00000108561.8 | C1QBP | 0.174887 | 0.6214382 | 0.203109 | 0.7949484 | -1.613112 | 2.24E-43 | -1.182428 | 4.81E-09 | 1.071658 | 1 | -0.42754 | 0.1394837 |
| ENSG00000108582.11 | CPD | -1.192199 | 0.9183025 | -0.287695 | 0.5953732 | 0.263992 | 0.2450435 | 0.183248 | 0.003633834 | 1.555143 | 0.3974318 | -0.23963 | 0.97072 |
| ENSG00000108587.14 | GOSR1 | -0.267411 | 0.9045883 | -0.034382 | 0.6527736 | 0.449425 | 0.04854632 | 0.108157 | 0.8434159 | 0.611989 | 0.7599374 | -0.742873 | 0.6885065 |
| ENSG00000108588.13 | CCDC47 | 0.436296 | 0.2656681 | 0.457131 | 0.6146853 | 0.083981 | 0.2536679 | 1.336843 | 0.000104536 | -0.25198 | 0.4703932 | 0.184108 | 0.6885065 |
| ENSG00000108590.10 | MED31 | -0.25286 | 0.7833919 | 0.319082 | 1 | -0.632794 | 0.03323509 | -0.312145 | 0.7994337 | 0.946169 | 0.3797324 | -0.075069 | 0.744192 |
| ENSG00000108599.14 | AKAP10 | 0.866676 | 1 | -0.251738 | 0.8298739 | -0.121582 | 0.895996 | 3.181596 | 0.02238098 | 1.456452 | 0.3063108 | 1.851768 | 0.9530808 |
| ENSG00000108654.11 | DDX5 | -0.519208 | 0.9972793 | 0.09918 | 0.9863789 | 0.438691 | 0.03964111 | 0.635236 | 0.05235768 | 0.372816 | 1 | 0.598319 | 0.6517664 |
| ENSG00000108669.16 | CYTH1 | 1.968201 | 0.3166974 | 0.220001 | 0.9134218 | -0.255764 | 0.2380533 | 0.96783 | 0.01859259 | -0.068415 | 0.7377262 | 1.360853 | 0.9335118 |
| ENSG00000108671.9 | PSMD11 | -0.729102 | 0.8456603 | 0.01686 | 0.7913203 | 0.83393 | 0.0479416 | 0.826189 | 0.3960072 | 0.276767 | 0.4289446 | 0.415505 | 0.9232737 |
| ENSG00000108679.12 | LGALS3BP | -1.358443 | 0.8787256 | 0.097454 | 0.8263112 | -0.46236 | 0.02591979 | 0.047903 | 0.7493034 | 0.890649 | 0.5196843 | 0.885249 | 0.9576568 |
| ENSG00000108691.9 | CCL2 | -2.423116 | 0.2025214 | -0.497819 | 0.971348 | -1.899521 | 0.000143487 | -0.406577 | 6.80E-06 | -1.346836 | 0.1206053 | -1.045016 | 0.9831628 |
| ENSG00000108773.10 | KAT2A | 1.295755 | 0.6214382 | 0.381678 | 0.8353216 | -0.529441 | 0.8061604 | 0.439368 | 0.3449579 | 1.036805 | 0.001415534 | 0.760458 | 0.8949489 |
| ENSG00000108774.14 | RAB5C | -0.034185 | 0.8867323 | 0.786019 | 0.1544733 | -0.376921 | 0.02290461 | 0.708206 | 0.4299976 | -0.079683 | 0.4090464 | 0.045067 | 0.6961634 |
| ENSG00000108819.10 | PPP1R9B | -0.435063 | 0.9537791 | -0.262636 | 0.7030131 | -0.195184 | 0.09968925 | 1.455722 | 0.000157153 | 2.70207 | 0.000794558 | 0.476718 | 0.6931304 |
| ENSG00000108848.15 | LUC7L3 | 0.094519 | 0.4406957 | 0.037037 | 0.5010139 | 0.712288 | 6.13E-10 | 1.127364 | 8.12E-06 | 0.222094 | 0.9226444 | -0.40096 | 0.5066477 |
| ENSG00000108946.14 | PRKAR1A | 0.324886 | 0.2221433 | 0.111984 | 0.9389787 | 0.139377 | 0.7164333 | 0.576027 | 0.02160238 | 0.83522 | 0.9002423 | 0.237501 | 0.9576568 |
| ENSG00000108953.16 | YWHAE | -0.043587 | 0.0318879 | -0.035447 | 0.7680378 | 0.183072 | 0.2756445 | 0.679643 | 1.19E-06 | 0.179089 | 0.5067421 | -0.077674 | 0.6195002 |
| ENSG00000109046.14 | WSB1 | -0.151212 | 0.6214382 | 0.438083 | 0.08356534 | 0.356418 | 0.2932027 | 1.417868 | 2.65E-07 | 1.676998 | 0.000250018 | 0.491747 | 0.9801657 |
| ENSG00000109062.9 | SLC9A3R1 | -1.045508 | 0.9183025 | -0.243448 | 0.6833417 | -1.435444 | 0.01851123 | 0.739503 | 0.4646256 | 1.468846 | 0.9266595 | -0.63696 | 0.9693119 |
| ENSG00000109079.9 | TNFAIP1 | 0.501341 | 0.8787256 | -0.184349 | 0.6077765 | 0.365789 | 0.9147533 | 2.069586 | 1.69E-06 | 0.344294 | 0.5196843 | 0.713991 | 0.7767294 |
| ENSG00000109099.13 | PMP22 | 0.127904 | 1 | -0.039016 | 0.7156572 | -0.22176 | 0.3191833 | 0.312661 | 0.00701136 | -0.875063 | 0.00456659 | -0.320077 | 1 |
| ENSG00000109118.13 | PHF12 | 1.930448 | 0.714717 | -0.046042 | 0.7594702 | 0.782435 | 0.6563895 | -0.050203 | 0.005840505 | 1.172549 | 0.5196843 | 4.646628 | 0.00077615 |
| ENSG00000109133.12 | TMEM33 | -0.074285 | 0.6014343 | -0.074222 | 0.6527736 | 0.410038 | 0.3564736 | 1.639333 | 0.00325396 | -0.551503 | 0.3683681 | -0.152595 | 1 |
| ENSG00000109171.14 | SLAIN2 | 0.685502 | 0.6214382 | -0.006594 | 0.6556516 | 1.277315 | 0.02821202 | -0.314093 | 0.8519978 | 1.798673 | 0.5826073 | 0.25515 | 0.803721 |
| ENSG00000109184.14 | DCUN1D4 | 1.252066 | 0.4764974 | 0.3807 | 0.6083172 | 1.052161 | 0.231686 | 0.920994 | 0.04393656 | 2.905836 | 0.09409813 | 0.812863 | 0.5733051 |
| ENSG00000109265.12 | KIAA1211 | -0.956776 | 0.8006286 | 0.45011 | 0.9222921 | -1.694304 | 0.04731355 | -0.880392 | 0.6578413 | 1.09621 | 0.8608026 | 2.011309 | 0.8479431 |
| ENSG00000109320.11 | NFKB1 | -3.082745 | 0.563505 | -0.31702 | 0.6527736 | 0.151063 | 0.7030899 | 0.63792 | 0.02429743 | 1.091815 | 0.777819 | 0.344145 | 0.9043938 |
| ENSG00000109339.18 | MAPK10 | -1.463194 | 0.8232133 | -0.683366 | 0.8465126 | 0.66158 | 0.1384986 | 1.411722 | 0.003915982 | 0.735108 | 0.4821192 | -1.457012 | 0.9646502 |
| ENSG00000109390.11 | NDUFC1 | 0.768448 | 0.4406957 | -0.123464 | 0.7765491 | -0.750226 | 1.21E-06 | -0.584038 | 1.65E-05 | -0.93364 | 0.003055278 | -0.96914 | 0.4406328 |
| ENSG00000109458.8 | GAB1 | -1.754263 | 0.8006286 | 0.018865 | 0.4921751 | 0.432522 | 0.2433478 | 2.567585 | 0.01099154 | 1.343901 | 0.1404709 | -0.912287 | 0.8949489 |
| ENSG00000109472.13 | CPE | -0.51478 | 0.1066198 | -0.475093 | 0.002183006 | -1.387264 | 0.002138815 | 1.44284 | 0.002324479 | 0.63978 | 0.260732 | 1.347857 | 1 |
| ENSG00000109475.16 | RPL34 | -0.562981 | 0.2025214 | -0.218728 | 0.1667081 | 0.033916 | 0.9070103 | 0.040043 | 0.1185248 | -0.302627 | 3.56E-05 | -0.08801 | 0.06111237 |
| ENSG00000109572.13 | CLCN3 | -0.546501 | 0.9883587 | -0.460057 | 0.2579583 | 1.024607 | 0.000413721 | 0.336286 | 0.4987772 | -0.172693 | 0.2828072 | 0.935619 | 1 |
| ENSG00000109586.11 | GALNT7 | -2.162208 | 0.5711449 | -0.26994 | 0.9597916 | 1.105249 | 0.03905198 | -0.318228 | 0.1633858 | -1.926304 | 1 | 0.538401 | 0.9617345 |
| ENSG00000109606.12 | DHX15 | 1.087668 | 0.5791794 | -0.111832 | 0.5394427 | 0.042059 | 0.001492356 | 1.156498 | 0.08271875 | 0.062583 | 0.2105971 | 0.727684 | 0.6185816 |
| ENSG00000109654.14 | TRIM2 | 1.282983 | 0.9671549 | 0.206673 | 0.8599148 | 1.440661 | 0.08238101 | 1.174524 | 0.007239704 | 1.029467 | 0.3616227 | 0.288608 | 1 |
| ENSG00000109670.13 | FBXW7 | 0.435593 | 0.6214382 | 0.079567 | 0.8263112 | 0.130102 | 0.4682361 | 1.864472 | 0.000967449 | 1.723687 | 0.621723 | -1.469765 | 0.9562356 |
| ENSG00000109685.17 | WHSC1 | -0.641465 | 0.6014343 | -0.228347 | 0.4768798 | 0.130298 | 0.9829733 | 0.295079 | 0.03225205 | 1.058188 | 0.01299974 | -0.246137 | 0.6885065 |
| ENSG00000109787.12 | KLF3 | 0.39001 | 0.7818196 | 0.225818 | 1 | 0.826519 | 0.5962642 | 1.191423 | 0.02750333 | 0.288482 | 0.9291558 | -1.799793 | 0.9147227 |
| ENSG00000109846.7 | CRYAB | -1.167884 | 3.02E-08 | -0.123241 | 0.04948531 | -0.021091 | 0.4064599 | -0.612413 | 0.000214893 | 0.072534 | 0.08374036 | -0.395382 | 0.7379523 |
| ENSG00000109919.9 | MTCH2 | -0.377043 | 0.9275335 | 0.341939 | 0.9985426 | -0.18875 | 0.03767703 | 1.057042 | 0.01506498 | -0.063259 | 0.2260152 | -0.183947 | 0.2856628 |
| ENSG00000109956.12 | B3GAT1 | 0.160152 | 0.529676 | -0.180239 | 0.681763 | 0.838398 | 0.4429676 | 1.168306 | 0.000360536 | 2.022716 | 0.4048755 | -1.33505 | 0.8949489 |
| ENSG00000109971.13 | HSPA8 | 0.03547 | 0.1487131 | 0.128725 | 0.4487482 | 0.648434 | 4.87E-11 | 0.317633 | 0.2784876 | 0.888012 | 5.71E-05 | -0.843141 | 0.000258077 |
| ENSG00000110076.18 | NRXN2 | -0.05138 | 0.9785048 | -0.213264 | 0.7436238 | -0.469193 | 0.4064599 | 0.92265 | 0.04593723 | 0.534731 | 0.3271351 | -0.025164 | 0.8479431 |
| ENSG00000110092.3 | CCND1 | -0.184863 | 1 | -0.432866 | 0.2799981 | 0.350725 | 0.01392126 | 1.066033 | 0.008125377 | -0.802456 | 0.005697906 | 0.030133 | 0.01821047 |
| ENSG00000110107.8 | PRPF19 | 0.092778 | 0.9174856 | -0.234116 | 0.3212092 | 0.163589 | 0.3365717 | 0.643642 | 0.02305301 | 0.792646 | 0.7377262 | 0.209384 | 0.8968136 |
| ENSG00000110171.18 | TRIM3 | -5.309499 | 0.4764974 | 0.682822 | 0.8953649 | 1.441464 | 0.3803088 | 1.402371 | 0.02693089 | 2.047677 | 0.3458308 | -0.854917 | 0.8749632 |
| ENSG00000110172.11 | CHORDC1 | -0.072382 | 1 | -0.000508 | 0.9828321 | -0.040578 | 0.000688818 | 0.442933 | 0.3192641 | 0.498395 | 0.4671878 | -0.842077 | 0.6232929 |
| ENSG00000110321.15 | EIF4G2 | -0.242041 | 0.529676 | -0.065119 | 0.3250176 | 0.4836 | 0.000429922 | 1.005724 | 3.59E-11 | 0.656262 | 0.2554415 | 0.282563 | 0.6517664 |
| ENSG00000110367.11 | DDX6 | 0.759151 | 0.3982827 | -0.39543 | 0.2919928 | 0.895577 | 0.004925626 | 1.31905 | 0.000159811 | 1.280487 | 0.06357357 | -0.44928 | 0.8479431 |
| ENSG00000110395.5 | CBL | -0.52133 | 0.8898405 | 0.045482 | 0.5953732 | 1.557314 | 0.00702513 | 0.822305 | 0.1606922 | 0.202509 | 0.93769 | 5.68735 | 9.82E-10 |
| ENSG00000110492.15 | MDK | -0.361093 | 0.5788332 | 0.333171 | 0.4409757 | -0.807923 | 2.17E-21 | 0.481625 | 0.9431066 | 0.486583 | 0.5084807 | -0.020586 | 0.2809502 |
| ENSG00000110514.19 | MADD | -3.026462 | 0.1364817 | 0.467745 | 1 | 0.720178 | 0.0361597 | 1.187621 | 0.03527959 | 0.318318 | 0.3974318 | 0.179408 | 0.7988798 |
| ENSG00000110536.13 | PTPMT1 | 0.390044 | 0.6214382 | -0.371968 | 0.6527736 | -1.278295 | 0.000380904 | 0.387692 | 0.1252108 | 0.108406 | 0.5377444 | 0.146089 | 0.8278576 |
| ENSG00000110583.12 | NAA40 | -1.190843 | 0.5930856 | -0.087213 | 0.987316 | 0.004383 | 0.0622543 | 1.681759 | 1.54E-07 | 0.566936 | 0.000194417 | -0.739387 | 0.6231061 |
| ENSG00000110619.16 | CARS | 1.050501 | 0.6214382 | 0.270369 | 0.9803143 | 0.691101 | 0.001412382 | 0.416961 | 0.7144938 | 0.810093 | 0.5196843 | -0.693695 | 0.922577 |
| ENSG00000110651.11 | CD81 | -0.907242 | 0.2351592 | -0.222039 | 0.1920223 | -0.101957 | 0.4064599 | 0.37545 | 0.1865665 | 1.544526 | 1.93E-05 | 0.074002 | 0.9663203 |
| ENSG00000110693.15 | SOX6 | -1.093472 | 0.59287 | -0.165224 | 0.9839225 | 0.872814 | 0.005437942 | 1.517819 | 0.05080366 | -0.739316 | 0.8786084 | -0.442982 | 0.8786493 |
| ENSG00000110696.9 | C11orf58 | -0.227576 | 0.59287 | 0.228671 | 0.6723689 | 0.706036 | 0.004266661 | 0.338133 | 0.04782381 | -0.182432 | 3.80E-05 | 0.323952 | 0.6885065 |
| ENSG00000110700.6 | RPS13 | -0.079488 | 0.01653545 | 0.030187 | 1 | 0.116304 | 0.3331553 | -0.444107 | 6.52E-12 | -0.153751 | 0.003344754 | 0.093987 | 0.3058061 |
| ENSG00000110711.9 | AIP | 0.578789 | 0.6214382 | -0.191219 | 0.9699228 | -0.878088 | 0.2728594 | -0.264519 | 0.02001972 | -0.185231 | 0.5196843 | -1.238915 | 0.4629479 |
| ENSG00000110713.15 | NUP98 | 0.137233 | 0.9360347 | 0.208857 | 0.6527736 | 0.895279 | 0.5290845 | 1.273079 | 0.2762756 | 2.296737 | 0.03760627 | 1.237021 | 0.964726 |
| ENSG00000110717.10 | NDUFS8 | 0.116734 | 0.529676 | -0.233944 | 0.6527736 | -0.019239 | 0.005898071 | -0.516845 | 0.003234961 | -0.153498 | 0.006268713 | -0.557076 | 0.6829816 |
| ENSG00000110756.17 | HPS5 | -0.637331 | 0.9401246 | 0.002328 | 0.8697479 | 0.230431 | 0.1496289 | 1.611312 | 0.009394191 | 0.651951 | 0.5715516 | 3.210293 | 0.9517616 |
| ENSG00000110768.11 | GTF2H1 | 0.122444 | 0.9045883 | 0.175404 | 0.9540226 | 0.59516 | 0.001863773 | -0.099795 | 1 | 0.522352 | 0.7850392 | -0.443857 | 0.9147227 |
| ENSG00000110801.13 | PSMD9 | -0.601142 | 0.7157845 | 0.387179 | 0.2945701 | -0.719046 | 0.000163622 | 0.90121 | 0.8444766 | -0.246217 | 0.8455691 | 0.204517 | 1 |
| ENSG00000110880.10 | CORO1C | 0.071507 | 0.9047689 | 0.085048 | 0.8703522 | 0.622713 | 0.9343369 | 1.125699 | 0.000354626 | 0.802826 | 0.02116689 | 0.032889 | 0.6885065 |
| ENSG00000110881.11 | ASIC1 | -0.307236 | 0.9183025 | -2.732122 | 0.05090264 | 1.610919 | 0.5610176 | 1.215505 | 0.009778502 | 2.589918 | 2.76E-05 | -4.865209 | 0.6517664 |
| ENSG00000110888.17 | CAPRIN2 | 0.253965 | 0.8867323 | -0.254128 | 0.6532996 | 0.357382 | 0.4064599 | 1.041615 | 0.02232997 | 1.935488 | 0.2905045 | -0.781055 | 0.6490419 |
| ENSG00000110925.6 | CSRNP2 | -2.285256 | 0.737591 | 0.196371 | 0.9931623 | -0.728143 | 0.5290845 | 1.115062 | 0.01804728 | 1.733282 | 0.01560716 | 0.568499 | 0.8250047 |
| ENSG00000110931.18 | CAMKK2 | 0.116409 | 0.6798783 | 0.15603 | 0.7581351 | 0.568087 | 0.9732741 | 0.982428 | 0.06383186 | 0.373252 | 0.03128833 | -0.612586 | 0.9813494 |
| ENSG00000110958.15 | PTGES3 | -0.344486 | 0.7035386 | 0.150806 | 0.6527736 | 0.443286 | 0.006469424 | 0.642435 | 0.08645516 | 0.61433 | 0.597842 | -0.20566 | 0.6885065 |
| ENSG00000111011.17 | RSRC2 | 0.328965 | 0.6214382 | -0.324202 | 0.5981187 | 0.493171 | 4.63E-07 | -0.017959 | 0.8472142 | 0.63046 | 0.8595963 | 0.662341 | 0.9368176 |
| ENSG00000111142.13 | METAP2 | -0.238588 | 0.4424206 | 0.107751 | 0.9973179 | 0.50653 | 0.002251313 | 0.302208 | 0.009887363 | -0.65302 | 6.14E-06 | -0.051119 | 0.7564421 |
| ENSG00000111196.9 | MAGOHB | -0.143656 | 0.6214382 | -0.015116 | 0.6833501 | -0.516178 | 0.01330515 | -0.324429 | 0.02691382 | 0.740226 | 0.2812673 | -0.619604 | 0.9147227 |
| ENSG00000111229.15 | ARPC3 | -0.746318 | 0.4764974 | 0.095448 | 0.7765491 | -0.204005 | 0.3504447 | 0.176664 | 0.5230877 | -0.252242 | 0.001005217 | 0.880468 | 0.9368176 |
| ENSG00000111231.8 | GPN3 | 0.322079 | 1 | -0.071072 | 0.6527736 | 0.360616 | 0.2147411 | -0.368817 | 0.03376927 | -0.922114 | 0.01082535 | -0.108829 | 0.6885065 |
| ENSG00000111266.8 | DUSP16 | 0.161537 | 0.9047689 | -0.426802 | 0.6527736 | 1.181905 | 0.02228582 | 1.141406 | 0.07654251 | 1.390058 | 0.007330751 | -0.364824 | 0.6231061 |
| ENSG00000111276.10 | CDKN1B | -0.060893 | 0.8006286 | 0.036758 | 0.8721583 | 0.505185 | 1.94E-06 | 0.231001 | 0.4919368 | 0.494407 | 0.5481917 | 1.900831 | 0.9677031 |
| ENSG00000111300.9 | NAA25 | -0.087609 | 0.737082 | 0.283143 | 0.8962992 | 0.80987 | 0.002841623 | 1.122006 | 0.1096608 | 2.444475 | 0.311393 | -2.505707 | 0.05841333 |
| ENSG00000111328.6 | CDK2AP1 | 0.175345 | 0.7308719 | 0.264097 | 0.6527736 | -1.085967 | 0.000918265 | 0.321449 | 0.3391173 | 3.221176 | 0.03099545 | -1.25265 | 0.3532606 |
| ENSG00000111364.15 | DDX55 | -0.498922 | 0.9794907 | 0.006479 | 0.6384858 | -0.589833 | 0.397212 | 1.532968 | 0.009724107 | 0.31584 | 0.4496768 | -2.206219 | 0.6885065 |
| ENSG00000111371.15 | SLC38A1 | -0.438536 | 0.6477491 | -0.027367 | 0.7945574 | 1.357538 | 0.002144312 | 1.082251 | 2.99E-07 | 0.34566 | 0.3976948 | -0.758047 | 0.7780985 |
| ENSG00000111530.12 | CAND1 | 0.598957 | 0.9183025 | 0.343161 | 0.7800195 | -0.309012 | 0.6149301 | 0.387853 | 0.000572729 | 0.312661 | 0.01053511 | 0.475867 | 0.9429078 |
| ENSG00000111605.16 | CPSF6 | 0.558918 | 0.9047689 | 0.687739 | 0.3803801 | 0.541405 | 0.2348326 | 1.526622 | 0.003210013 | -0.26509 | 0.7395971 | -0.661451 | 0.6885065 |
| ENSG00000111639.7 | MRPL51 | -0.606598 | 0.3266815 | -0.126677 | 0.6527736 | -0.12378 | 0.002892406 | -0.14656 | 0.01305491 | 0.082954 | 0.001955021 | -0.424088 | 0.09885076 |
| ENSG00000111640.14 | GAPDH | -0.611724 | 8.03E-06 | 0.33038 | 6.04E-22 | 0.159193 | 0.00436005 | -0.049495 | 0.8925294 | 0.206548 | 0.7777374 | 0.281817 | 0.6185816 |
| ENSG00000111641.10 | NOP2 | 0.624597 | 0.6214382 | -0.185745 | 0.6834954 | -0.07184 | 1 | 1.342646 | 0.03527869 | 1.408233 | 0.1045014 | 0.868976 | 0.5066477 |
| ENSG00000111642.14 | CHD4 | -2.078205 | 0.5119183 | 0.114959 | 0.6527736 | 0.571127 | 5.88E-05 | 0.385506 | 0.05689288 | 0.697762 | 0.5279687 | 0.122952 | 0.6885065 |
| ENSG00000111647.12 | UHRF1BP1L | 0.283028 | 0.563505 | -0.428352 | 0.293519 | 2.077084 | 0.000101611 | 0.326703 | 0.5874595 | 0.32831 | 0.8768974 | -0.430162 | 0.9618275 |
| ENSG00000111669.14 | TPI1 | 0.318195 | 0.01711078 | 0.242824 | 0.006625527 | 0.03371 | 0.3263442 | -0.139547 | 0.002317745 | -0.002822 | 0.03114012 | 0.206629 | 0.6829816 |
| ENSG00000111671.9 | SPSB2 | 0.750311 | 0.6214382 | 0.323737 | 0.4255557 | -1.029312 | 0.002947763 | -0.385348 | 0.2110701 | -0.73936 | 0.8605751 | -1.98744 | 0.8539279 |
| ENSG00000111674.8 | ENO2 | -1.25117 | 2.62E-05 | 0.550408 | 0.000181022 | 0.069669 | 0.3818497 | 0.327372 | 0.9051123 | -0.391399 | 0.000118883 | 0.868419 | 0.8950308 |
| ENSG00000111678.10 | C12orf57 | -0.0347 | 0.8774628 | -0.08907 | 0.5953732 | -0.085391 | 0.002391157 | -0.378432 | 0.02033164 | -0.309609 | 0.005873149 | 0.078022 | 0.9884778 |
| ENSG00000111707.11 | SUDS3 | 0.801885 | 0.9045883 | -0.455066 | 0.6527736 | -0.744983 | 0.5752098 | -0.740442 | 0.02516296 | 1.350362 | 0.3336184 | 1.261424 | 0.9169962 |
| ENSG00000111711.9 | GOLT1B | 0.324478 | 0.9016905 | 0.605469 | 0.2989839 | 0.336857 | 0.2771264 | 0.867601 | 0.4360536 | -0.269407 | 0.03812753 | 1.068173 | 0.6885065 |
| ENSG00000111716.12 | LDHB | -1.565086 | 0.1239011 | -0.216663 | 0.1054706 | 0.295018 | 0.000152727 | -0.391327 | 7.47E-06 | -0.471395 | 4.92E-05 | 0.069768 | 0.6185816 |
| ENSG00000111737.11 | RAB35 | -0.423661 | 0.9243498 | 0.318759 | 1 | -0.747402 | 0.01526418 | 0.78655 | 0.06928411 | 0.315643 | 0.4317284 | 0.379003 | 0.6885065 |
| ENSG00000111752.10 | PHC1 | -0.560176 | 0.8867323 | 0.399439 | 0.592336 | -0.047624 | 0.5290845 | 0.99193 | 0.000856657 | 1.17838 | 0.0127696 | -0.59827 | 0.4578379 |
| ENSG00000111775.2 | COX6A1 | 0.769586 | 5.68E-14 | -0.026701 | 1 | -0.525507 | 1.40E-14 | -0.054733 | 0.0038114 | 0.007652 | 0.000391835 | -0.084301 | 0.05841333 |
| ENSG00000111786.8 | SRSF9 | -0.076827 | 0.6214382 | -0.072637 | 0.8158046 | -0.3218 | 0.003574973 | 0.855229 | 0.06030703 | 1.569155 | 0.03616584 | -0.368096 | 0.6885065 |
| ENSG00000111817.16 | DSE | -0.392675 | 0.8006286 | 1.376134 | 0.5751014 | 0.902591 | 0.5290845 | 2.497685 | 0.04349377 | 1.71564 | 0.2880317 | 0.71859 | 0.8681734 |
| ENSG00000111843.13 | TMEM14C | -0.303942 | 0.8353323 | 0.520288 | 0.3121527 | -0.595846 | 1.49E-05 | 0.100857 | 0.5911715 | -0.437229 | 0.000328504 | 0.046486 | 0.6885065 |
| ENSG00000111845.4 | PAK1IP1 | 0.206726 | 0.529676 | -0.023987 | 0.8771169 | 0.187222 | 0.5290845 | -0.064586 | 0.01321152 | -0.371555 | 0.001363097 | -0.451669 | 0.7315846 |
| ENSG00000111860.13 | CEP85L | -0.926077 | 0.714717 | 0.124045 | 0.8703522 | 1.185671 | 0.001959195 | 0.311069 | 0.6445029 | 2.867312 | 0.154771 | 2.606017 | 0.8479431 |
| ENSG00000111897.6 | SERINC1 | 0.152858 | 1 | 0.028997 | 0.8703522 | 0.792075 | 1.85E-06 | 0.670073 | 0.5034517 | 0.538609 | 0.5475627 | 0.997642 | 0.6885065 |
| ENSG00000111906.17 | HDDC2 | -0.277189 | 0.6214382 | -0.063954 | 0.9683845 | 0.00698 | 0.2471551 | -0.244928 | 0.03549107 | -0.290277 | 0.1170868 | 0.129554 | 0.9391684 |
| ENSG00000111912.19 | NCOA7 | 2.478237 | 0.4764974 | 0.635535 | 0.8953649 | 1.243509 | 0.04360461 | -0.088452 | 0.2162673 | 0.187776 | 0.6240613 | 4.264448 | 0.8241214 |
| ENSG00000111962.7 | UST | -0.264793 | 0.8234926 | -1.271321 | 0.6527736 | 0.930743 | 0.5290845 | 1.157818 | 0.04811661 | 0.26561 | 1 | 0.813722 | 1 |
| ENSG00000112062.10 | MAPK14 | -1.972551 | 0.7381604 | 0.193103 | 0.6527736 | -1.183057 | 0.4893071 | 1.600355 | 0.004026498 | 1.961081 | 0.06448943 | -0.213873 | 0.8712617 |
| ENSG00000112078.13 | KCTD20 | -0.230223 | 0.9047689 | 0.193977 | 0.7383998 | 0.353485 | 0.3984193 | 1.885242 | 0.03303111 | 0.754401 | 0.03734702 | -0.855214 | 0.9382274 |
| ENSG00000112079.8 | STK38 | -0.828184 | 0.6014343 | -0.693987 | 0.2756663 | -0.792646 | 0.7967238 | 1.904207 | 0.01413636 | 0.566699 | 0.4384759 | -0.096419 | 1 |
| ENSG00000112096.16 | SOD2 | -1.106344 | 0.2449115 | -0.020519 | 0.378582 | -1.179548 | 2.13E-09 | 0.26484 | 0.1292522 | -0.196138 | 0.9128939 | 0.437183 | 0.6885065 |
| ENSG00000112118.17 | MCM3 | -0.134251 | 1 | -0.118405 | 0.6867931 | -1.072609 | 0.241585 | 0.807339 | 9.02E-05 | 0.004837 | 0.8387831 | -0.090093 | 0.8188271 |
| ENSG00000112146.16 | FBXO9 | -0.406018 | 1 | -0.27566 | 0.5010139 | 0.482046 | 0.2769014 | 0.858906 | 0.04638111 | 0.698368 | 0.9527758 | -0.775807 | 0.6885065 |
| ENSG00000112159.11 | MDN1 | 0.470297 | 0.7686507 | -0.567403 | 0.6107341 | -1.067412 | 0.8984213 | 2.677135 | 0.000263274 | 2.615935 | 0.001415534 | 1.633866 | 0.7623154 |
| ENSG00000112200.16 | ZNF451 | -1.23574 | 0.6214382 | -0.543702 | 1 | 0.752638 | 6.62E-05 | 0.643178 | 0.04775009 | 1.4191 | 0.04542333 | 0.740594 | 0.9789121 |
| ENSG00000112242.14 | E2F3 | -0.471467 | 0.563505 | -0.278297 | 0.6527736 | -0.578839 | 0.5290845 | 0.891529 | 0.001309599 | 1.588034 | 0.4020737 | -2.17631 | 0.6885065 |
| ENSG00000112249.13 | ASCC3 | -0.740669 | 0.714717 | -0.565432 | 0.09148334 | 0.930307 | 0.04702712 | 0.520266 | 0.1865665 | 1.594726 | 1 | 1.947061 | 0.7767294 |
| ENSG00000112282.17 | MED23 | -0.479966 | 0.7736255 | 0.390632 | 0.6527736 | -0.619624 | 0.8428641 | 0.857769 | 0.1248486 | 1.049907 | 0.03832539 | 3.04658 | 0.9147227 |
| ENSG00000112304.10 | ACOT13 | 0.694284 | 0.8052741 | -0.649277 | 0.09704869 | -0.096527 | 0.01143551 | 1.060884 | 0.05394541 | 0.066872 | 1 | 0.143633 | 0.6885065 |
| ENSG00000112305.14 | SMAP1 | 0.015323 | 1 | 0.15178 | 0.8953649 | 1.016949 | 0.1418079 | 0.312996 | 0.4919368 | -0.508017 | 0.006214714 | 2.753455 | 0.826754 |
| ENSG00000112306.7 | RPS12 | -0.571186 | 0.09810899 | 0.067696 | 0.6527736 | 0.014041 | 0.5840658 | -0.063527 | 0.2598457 | 0.024819 | 0.003786605 | 0.118906 | 0.5394145 |
| ENSG00000112308.12 | C6orf62 | 0.114736 | 0.8006286 | 0.173062 | 0.8314396 | 0.584183 | 0.2129549 | 1.605927 | 0.000123766 | 0.541052 | 0.6700639 | 0.807425 | 0.9153161 |
| ENSG00000112320.11 | SOBP | -1.252025 | 0.6014343 | -0.500365 | 0.4409757 | -0.169935 | 0.6387506 | 2.251596 | 1.82E-06 | 1.496897 | 0.2834684 | -0.200411 | 0.960853 |
| ENSG00000112333.11 | NR2E1 | -3.409582 | 0.6214382 | 0.321098 | 0.8863478 | -0.305259 | 0.7687948 | -1.311464 | 1 | -1.999443 | 1 |  |  |
| ENSG00000112335.14 | SNX3 | -0.656862 | 0.9282145 | -0.199613 | 0.5649416 | -0.779101 | 2.41E-05 | 0.608505 | 0.3467508 | 0.478378 | 0.9782718 | 0.334454 | 0.7767607 |
| ENSG00000112379.8 | ARFGEF3 | 0.524703 | 0.9047689 | -0.101182 | 0.9829438 | 1.036657 | 0.3395668 | 2.273628 | 8.89E-05 | -0.153494 | 0.01777855 | 2.771048 | 0.6885065 |
| ENSG00000112394.16 | SLC16A10 | 2.265258 | 1 | 1.086339 | 0.4571222 | 2.095492 | 0.5606143 | 2.705665 | 6.61E-05 | 0.55473 | 0.2357139 | 3.229351 | 0.6829816 |
| ENSG00000112406.4 | HECA | -0.408578 | 0.9047689 | -0.673102 | 0.4031681 | 2.659649 | 0.00089766 | 0.378847 | 0.2367097 | 0.570984 | 0.04945669 | 4.077185 | 0.6490419 |
| ENSG00000112511.17 | PHF1 | 0.247786 | 0.8006902 | 0.109992 | 0.8263112 | -0.434273 | 0.497211 | 0.747227 | 0.1557388 | 1.463921 | 0.03922063 | -1.233289 | 0.8930213 |
| ENSG00000112514.15 | CUTA | -0.216728 | 0.8867323 | -0.104567 | 0.6737951 | -0.066299 | 0.3205816 | 0.254976 | 0.9085299 | -0.208205 | 0.007522843 | 0.860356 | 0.7385207 |
| ENSG00000112531.16 | QKI | -0.188464 | 0.8963042 | -0.119507 | 0.4447592 | 0.676026 | 1.92E-13 | 1.028303 | 3.65E-07 | 0.457641 | 0.2523958 | 0.638778 | 0.9037465 |
| ENSG00000112640.14 | PPP2R5D | -0.50579 | 0.9183025 | -0.370183 | 0.9156685 | -0.4791 | 0.5290845 | 1.48157 | 0.2696559 | 1.961083 | 0.008247924 | -0.124354 | 0.6885065 |
| ENSG00000112651.11 | MRPL2 | -1.269419 | 0.7345905 | -0.142077 | 1 | -0.381947 | 0.000730534 | -0.487831 | 0.05306874 | -1.234255 | 0.007322463 | 0.552588 | 0.7215554 |
| ENSG00000112658.7 | SRF | -0.73244 | 0.3946735 | 0.655571 | 0.4375617 | 0.395782 | 0.7437308 | 1.393175 | 2.88E-05 | 0.692715 | 0.04422904 | 0.21952 | 0.8479431 |
| ENSG00000112685.13 | EXOC2 | 0.260539 | 1 | 0.299691 | 0.7383998 | -0.257408 | 0.7382221 | 1.426378 | 0.01970714 | 0.965204 | 0.930702 | -0.274064 | 1 |
| ENSG00000112695.11 | COX7A2 | 0.238906 | 0.007970027 | 0.168342 | 0.6527736 | -0.363956 | 1.11E-10 | -0.049102 | 0.02506126 | -0.141021 | 0.006549292 | -0.247737 | 0.2159565 |
| ENSG00000112697.15 | TMEM30A | 0.005967 | 0.8108384 | 0.031898 | 0.6077765 | -0.033197 | 0.8138435 | 0.767577 | 0.03655283 | 0.122185 | 0.18913 | -0.061203 | 0.8986153 |
| ENSG00000112701.17 | SENP6 | -0.421771 | 0.8203421 | -0.311424 | 0.2792759 | 0.892269 | 0.209738 | 0.947406 | 0.02279953 | 0.328216 | 0.5196843 | 0.237397 | 0.9824843 |
| ENSG00000112715.20 | VEGFA | -1.353076 | 0.3579569 | 0.770218 | 1.42E-23 | -0.465531 | 0.06427144 | 0.901795 | 0.001602625 | 1.657097 | 0.005343123 | 1.276205 | 0.4771551 |
| ENSG00000112739.16 | PRPF4B | -0.324712 | 0.6014343 | 0.001815 | 0.6527736 | 0.349368 | 0.000192559 | 0.265886 | 0.9298594 | 1.337402 | 0.1592962 | 0.358205 | 1 |
| ENSG00000112787.12 | FBRSL1 | -1.479915 | 0.6014343 | 0.413223 | 0.6414243 | -1.48196 | 0.5290845 | 0.818091 | 0.1133161 | 2.705157 | 0.003599031 | 1.852533 | 0.6829816 |
| ENSG00000112851.14 | ERBB2IP | -0.972142 | 0.329461 | -0.386791 | 0.01039093 | 0.638613 | 0.6976695 | 0.770258 | 0.1241339 | -0.793834 | 0.4780477 | 1.565505 | 0.868568 |
| ENSG00000112893.9 | MAN2A1 | -0.292123 | 0.8867323 | -0.964165 | 0.01902676 | -1.932026 | 0.1136698 | 2.972863 | 0.000746863 | 0.578946 | 0.5196843 | 0.44147 | 0.6517664 |
| ENSG00000112972.14 | HMGCS1 | -1.33106 | 0.6014343 | 0.453768 | 0.5306483 | -0.244948 | 0.7648735 | 1.251206 | 9.24E-05 | 0.27638 | 0.6834308 | -0.11117 | 0.6829816 |
| ENSG00000112983.17 | BRD8 | -5.267332 | 0.9036323 | 0.285023 | 0.8263112 | 0.193414 | 0.7824741 | 0.507515 | 0.01639512 | 1.048829 | 0.7377262 | -1.986354 | 0.7140075 |
| ENSG00000113140.10 | SPARC | -1.357109 | 0.000878274 | -0.135882 | 0.4886958 | -0.738569 | 3.46E-10 | -0.323829 | 5.01E-05 | -0.524625 | 7.89E-07 | 0.510554 | 0.6829816 |
| ENSG00000113141.15 | IK | -0.190803 | 0.8006286 | -0.149414 | 0.2801475 | 0.531883 | 0.000871313 | 0.252839 | 0.9924894 | 0.06107 | 0.009868435 | -0.731638 | 1 |
| ENSG00000113240.12 | CLK4 | -1.386026 | 1 | -0.550642 | 0.8953649 | 1.09407 | 0.0171631 | 0.956849 | 0.4071802 | 0.19814 | 0.8840084 | 0.955219 | 0.8654874 |
| ENSG00000113282.13 | CLINT1 | -2.151021 | 0.6214382 | -0.088904 | 0.69486 | 0.613979 | 0.000258451 | -0.322321 | 0.7273176 | 0.133977 | 0.3699044 | -1.748996 | 0.7815439 |
| ENSG00000113300.11 | CNOT6 | 1.300044 | 0.6214382 | -0.144848 | 0.8545414 | 0.575248 | 0.5042974 | 0.140637 | 0.0187186 | 1.15268 | 0.1446007 | 2.084138 | 0.6885065 |
| ENSG00000113312.10 | TTC1 | 0.318616 | 0.7736255 | 0.562557 | 0.570279 | 0.720416 | 0.0539913 | -0.490306 | 0.09060518 | -0.468767 | 0.000254175 | 0.269246 | 0.7523447 |
| ENSG00000113360.16 | DROSHA | -0.368309 | 1 | 0.270855 | 0.6527736 | 0.969408 | 0.2552655 | 0.983738 | 0.01292204 | 1.618868 | 0.3368344 | 0.368039 | 0.8949489 |
| ENSG00000113368.11 | LMNB1 | -2.013352 | 0.6168416 | -0.39488 | 0.5010139 | -0.026754 | 0.4577356 | 1.388766 | 0.006123422 | 1.996726 | 0.000289143 | 1.140714 | 0.6185816 |
| ENSG00000113369.8 | ARRDC3 | 1.373441 | 0.02815464 | 0.179153 | 0.6218841 | -0.027772 | 0.3017434 | 0.878436 | 0.03918882 | 1.515488 | 0.012842 | 0.213198 | 0.6829816 |
| ENSG00000113387.11 | SUB1 | 0.289401 | 0.4709027 | -0.030657 | 0.6012415 | -0.48001 | 0.000455376 | 0.200631 | 0.8896226 | -0.421844 | 0.01579695 | -0.218847 | 0.6885065 |
| ENSG00000113407.13 | TARS | -0.022106 | 0.714717 | -0.079086 | 0.6527736 | 0.591103 | 8.16E-06 | -0.145068 | 0.25709 | -0.463163 | 0.3974318 | -0.164923 | 0.8479431 |
| ENSG00000113441.15 | LNPEP | 0.945109 | 1 | -0.624246 | 0.2717173 | 0.826116 | 0.6767928 | 0.480643 | 0.02310609 | 1.263401 | 0.7347147 | -1.194068 | 0.6945946 |
| ENSG00000113448.16 | PDE4D | 0.058572 | 0.6214382 | 0.832134 | 0.6387692 | 1.617714 | 0.1126151 | -0.538364 | 0.02918812 | 1.510025 | 0.06320874 | 0.993788 | 0.6185816 |
| ENSG00000113522.13 | RAD50 | 0.336296 | 0.6214382 | 0.264129 | 0.6527736 | 0.931558 | 0.00643652 | 0.932171 | 0.06955268 | 0.245111 | 0.8589299 | 0.69398 | 0.8309897 |
| ENSG00000113558.18 | SKP1 | -0.139635 | 0.4399567 | 0.112259 | 0.8504478 | 0.155655 | 0.8123647 | -0.071258 | 1.62E-06 | -0.011357 | 6.77E-05 | -0.198379 | 0.4793294 |
| ENSG00000113580.14 | NR3C1 | -1.401797 | 0.1754689 | 0.20822 | 0.9104976 | 0.221674 | 0.2149145 | 0.604191 | 0.03575163 | 0.433224 | 0.6813171 | 3.070519 | 0.6829816 |
| ENSG00000113643.8 | RARS | 0.498701 | 1 | 0.20291 | 0.8545414 | 0.540196 | 0.3054882 | 0.256535 | 0.3725684 | -0.48111 | 0.03105236 | 0.822135 | 1 |
| ENSG00000113648.16 | H2AFY | -0.591694 | 0.06581042 | -0.021656 | 0.9618421 | -0.32454 | 0.3331553 | 0.534378 | 0.7398829 | -0.104275 | 0.03156166 | 0.626143 | 0.6885065 |
| ENSG00000113649.11 | TCERG1 | -0.248634 | 1 | -0.134304 | 0.8196487 | 0.619237 | 0.0133227 | 0.522343 | 0.572865 | 0.267638 | 0.8021966 | -0.673233 | 0.9705209 |
| ENSG00000113657.12 | DPYSL3 | -0.108401 | 1 | -0.295674 | 0.004197116 | -0.311338 | 0.09968925 | 1.076533 | 6.44E-07 | 0.488499 | 0.5929557 | -0.542378 | 0.566849 |
| ENSG00000113658.16 | SMAD5 | -0.287316 | 0.9223024 | -0.677728 | 0.06427071 | 0.133714 | 0.01834513 | 0.266569 | 0.06940845 | 0.661968 | 0.5381878 | -0.400907 | 0.826754 |
| ENSG00000113712.16 | CSNK1A1 | 0.129215 | 0.6014343 | 0.442324 | 0.5676698 | 0.773075 | 4.89E-06 | 0.797177 | 0.4185068 | -0.269826 | 0.2876398 | -2.020643 | 0.6829816 |
| ENSG00000113719.15 | ERGIC1 | 0.488417 | 0.8364538 | -0.613727 | 0.08356534 | -1.063583 | 0.2580724 | 1.84235 | 6.15E-05 | 1.143218 | 0.01165524 | -0.665379 | 0.7431136 |
| ENSG00000113732.8 | ATP6V0E1 | -0.078253 | 0.8006286 | 0.30484 | 0.5953732 | -0.416277 | 0.001744726 | 0.127466 | 2.40E-05 | 0.199703 | 0.1411968 | 0.623328 | 0.8213017 |
| ENSG00000113742.12 | CPEB4 | -0.624404 | 0.9411767 | -0.317635 | 0.5836731 | 0.461589 | 0.1415624 | 1.074467 | 0.06727595 | 1.653563 | 0.02825887 | 1.355647 | 0.9270789 |
| ENSG00000113758.13 | DBN1 | 0.159481 | 0.238424 | -0.049613 | 0.6527736 | 0.279256 | 0.7437308 | 1.07381 | 0.005310674 | 1.007723 | 0.103841 | 0.039681 | 0.348294 |
| ENSG00000113810.15 | SMC4 | -4.795561 | 0.6214382 | -0.370417 | 0.1146189 | 0.709154 | 3.51E-06 | -0.017479 | 0.4855817 | 0.295905 | 0.7030821 | -0.390362 | 0.6885065 |
| ENSG00000113811.10 | SELK | 0.310768 | 0.5711449 | 0.514079 | 0.002098572 | -0.036414 | 0.2839251 | -0.326471 | 5.94E-07 | -0.473626 | 0.000419524 | -0.630943 | 0.1520456 |
| ENSG00000113916.17 | BCL6 | -0.95942 | 0.8006286 | -0.980112 | 0.2709731 | -0.251158 | 0.7967238 | 0.860736 | 0.03997638 | 1.364362 | 0.04948871 | 1.541131 | 0.6885065 |
| ENSG00000113971.18 | NPHP3 | 0.211104 | 0.714717 | 0.39151 | 0.8089431 | -0.49812 | 0.8895055 | 4.261925 | 0.01477387 | 1.298326 | 0.8953039 | -3.180866 | 0.5553892 |
| ENSG00000114019.14 | AMOTL2 | 0.85409 | 0.9047689 | -0.005034 | 0.8263112 | 0.846379 | 0.8389806 | 2.256529 | 0.01608402 | 1.2693 | 0.2570214 | 2.235732 | 0.6185816 |
| ENSG00000114023.15 | FAM162A | -0.218595 | 0.8809848 | 0.583538 | 0.000379657 | -0.621757 | 0.0181549 | -0.254959 | 0.00162957 | -0.120496 | 0.01296756 | 0.042103 | 0.744192 |
| ENSG00000114062.17 | UBE3A | -0.201025 | 0.563505 | -0.140339 | 0.737997 | 0.612928 | 0.1356046 | 1.102295 | 0.001627599 | 0.306596 | 0.9316648 | -2.314791 | 1 |
| ENSG00000114120.11 | SLC25A36 | 0.118757 | 0.563505 | -0.009359 | 0.6414243 | 0.763517 | 0.01330515 | 1.424634 | 0.1288377 | 1.065621 | 0.5196843 | 0.317605 | 1 |
| ENSG00000114125.13 | RNF7 | 0.199376 | 0.69835 | 0.045112 | 0.9443543 | -0.498379 | 0.1217567 | 0.104147 | 0.8624565 | -1.136551 | 0.0127696 | -0.282556 | 0.8949489 |
| ENSG00000114127.10 | XRN1 | 1.193303 | 0.9047689 | -0.393237 | 0.8703522 | 1.633623 | 0.04089497 | 1.483701 | 0.4103395 | 1.582462 | 0.7687933 | 1.024918 | 0.9632895 |
| ENSG00000114200.9 | BCHE | 0.002553 | 0.9799552 | -0.83815 | 0.5010139 | 0.956562 | 0.009595218 | -0.609757 | 0.1578146 | 0.450842 | 0.5152872 | 0.548995 | 0.6885065 |
| ENSG00000114209.14 | PDCD10 | -1.13457 | 0.764956 | 0.240988 | 0.6833417 | 0.628538 | 0.5290845 | -0.434873 | 0.06916271 | -1.104187 | 0.01942541 | -0.90943 | 0.920468 |
| ENSG00000114270.15 | COL7A1 | -1.552016 | 1 | 0.034377 | 0.8703522 | -1.56883 | 0.8454005 | 2.438562 | 9.99E-07 | 0.981444 | 0.621723 | -0.952772 | 0.8949489 |
| ENSG00000114302.15 | PRKAR2A | 0.552463 | 0.6214382 | -0.407221 | 0.2217425 | 0.144229 | 0.339353 | 1.797865 | 0.001769389 | 0.947672 | 0.09561541 | 0.884545 | 0.8732786 |
| ENSG00000114315.3 | HES1 | -1.460065 | 0.4764974 | -0.227061 | 0.8263112 | -0.057572 | 0.8454005 | 0.126023 | 0.01269524 | 1.05817 | 0.8722349 | 0.617596 | 0.6885065 |
| ENSG00000114383.9 | TUSC2 | 1.341578 | 0.2588298 | 0.119961 | 0.8046865 | -0.7544 | 8.73E-06 | -0.544516 | 0.4569212 | 1.081725 | 0.9443862 | -1.100095 | 0.7419782 |
| ENSG00000114391.12 | RPL24 | -0.268671 | 0.8501558 | 0.029715 | 0.9848758 | 0.126233 | 0.5772092 | -0.167955 | 4.85E-06 | 0.007109 | 0.009755487 | 0.01811 | 0.4507434 |
| ENSG00000114395.10 | CYB561D2 | 0.427882 | 0.8264007 | 0.315476 | 0.9458404 | -1.066494 | 0.04517233 | 0.193297 | 0.5169969 | -1.097362 | 0.8138996 | 0.985192 | 0.8768631 |
| ENSG00000114416.17 | FXR1 | -0.472782 | 0.59287 | -0.135775 | 0.7985277 | 0.592836 | 6.74E-07 | 0.378121 | 0.266568 | -0.910415 | 0.001955021 | -0.700838 | 0.703227 |
| ENSG00000114439.18 | BBX | 0.531689 | 0.4511893 | 0.272013 | 0.6527736 | 1.529032 | 1.09E-06 | 1.477862 | 0.02072474 | 1.406802 | 0.1207057 | 0.641746 | 0.8919887 |
| ENSG00000114503.10 | NCBP2 | -0.71225 | 0.6214382 | -0.0359 | 0.5981187 | 0.748828 | 0.238938 | -0.640797 | 0.02140076 | 0.399541 | 0.2221977 | -0.565655 | 0.6829816 |
| ENSG00000114554.11 | PLXNA1 | 2.416611 | 0.6014343 | -0.190948 | 0.9951789 | -0.000988 | 0.9860195 | 0.965328 | 0.000132473 | 0.484076 | 0.1188982 | -0.899346 | 0.868568 |
| ENSG00000114573.9 | ATP6V1A | 0.657389 | 0.3781434 | -0.207018 | 0.9791082 | 1.110064 | 0.06670585 | 1.793747 | 0.02377077 | 0.886014 | 0.9274968 | 0.009956 | 0.6504862 |
| ENSG00000114631.10 | PODXL2 | -0.6736 | 0.3952859 | 0.190302 | 0.6527736 | -3.705573 | 6.92E-06 | 0.459268 | 0.9912905 | 1.176232 | 0.5475627 | -0.18685 | 0.6185816 |
| ENSG00000114742.13 | WDR48 | 0.941445 | 0.6214382 | 0.133257 | 0.8064295 | 0.400993 | 0.01012156 | 1.025855 | 0.04301863 | 1.22358 | 0.06175461 | 0.475226 | 0.8949489 |
| ENSG00000114784.3 | EIF1B | 0.181298 | 0.714717 | 0.58001 | 0.6527736 | 0.079547 | 0.8413311 | 0.225408 | 0.4850832 | -0.195757 | 0.04054872 | -0.472448 | 0.8479431 |
| ENSG00000114857.17 | NKTR | 0.085222 | 0.9511192 | 0.04487 | 0.7383998 | 0.689677 | 0.001963 | 0.352708 | 7.08E-07 | 1.447028 | 0.00134652 | -0.369635 | 1 |
| ENSG00000114867.19 | EIF4G1 | 0.312822 | 0.8612279 | 0.230083 | 0.9775912 | 0.052318 | 0.7614797 | 1.299314 | 3.53E-07 | 0.925455 | 0.2527216 | -0.686294 | 0.2202646 |
| ENSG00000114902.13 | SPCS1 | -0.232576 | 0.8658573 | 0.451194 | 0.2271903 | -0.155896 | 0.003248071 | 0.068049 | 0.003270086 | -0.158826 | 0.002398345 | -0.505759 | 0.6517664 |
| ENSG00000114933.15 | INO80D | -1.219507 | 0.8402193 | -0.044616 | 1 | 0.514342 | 0.01603959 | 1.26677 | 0.04997317 | 0.536914 | 0.2919943 | -1.721866 | 0.967026 |
| ENSG00000114942.13 | EEF1B2 | -1.735543 | 0.6214382 | 0.359923 | 0.4571222 | 0.261912 | 0.1027536 | 0.021161 | 0.4251194 | -0.397614 | 0.03738376 | 0.282591 | 0.186512 |
| ENSG00000114956.19 | DGUOK | -0.428283 | 0.8006286 | 0.298697 | 0.7383998 | -0.166905 | 0.4203906 | -0.247265 | 0.002397755 | 0.154291 | 0.2262118 | 0.769739 | 0.9289263 |
| ENSG00000114978.17 | MOB1A | 1.570307 | 0.714717 | 0.530628 | 0.8587199 | 1.270811 | 0.01265738 | 1.154624 | 0.176489 | 1.365981 | 0.3974318 | -1.491139 | 0.8168211 |
| ENSG00000114982.17 | KANSL3 | 4.98298 | 0.563505 | -0.953144 | 0.2866878 | -0.831373 | 0.4669981 | 0.746744 | 0.000126905 | 1.52396 | 0.183138 | 0.87781 | 1 |
| ENSG00000114999.7 | TTL | -0.331735 | 0.6562396 | -0.055104 | 0.6055625 | 0.033017 | 0.2405863 | 1.353028 | 3.28E-05 | 1.183511 | 0.1826169 | -0.462289 | 0.6885065 |
| ENSG00000115020.16 | PIKFYVE | -1.577674 | 0.6214382 | 0.512548 | 0.8504311 | 0.700522 | 0.09737406 | 0.731643 | 0.01812133 | 1.849164 | 0.1953258 | 0.59297 | 0.6647029 |
| ENSG00000115041.12 | KCNIP3 | -3.448881 | 0.6214382 | 1.995249 | 1 | -1.265071 | 0.7437308 | 2.255708 | 0.3467508 | 3.460009 | 0.4020737 |  |  |
| ENSG00000115053.15 | NCL | 0.3112 | 0.0505434 | -0.019776 | 0.628885 | 0.512343 | 1.79E-07 | 0.756206 | 0.005526995 | 0.20204 | 0.7377262 | 0.112875 | 0.2768456 |
| ENSG00000115073.7 | ACTR1B | -0.252613 | 0.6214382 | 0.452481 | 0.6200225 | 0.618368 | 0.8348198 | 0.214754 | 0.04939234 | 1.611154 | 0.2556593 | 1.595569 | 0.9278206 |
| ENSG00000115109.13 | EPB41L5 | -1.636195 | 0.714717 | -0.364175 | 0.7971908 | 0.817802 | 0.0334183 | 1.112783 | 0.5327414 | 1.471273 | 0.07182047 | -0.063676 | 0.6885065 |
| ENSG00000115128.6 | SF3B6 | 0.061889 | 1 | 0.070352 | 0.9848758 | -0.305838 | 0.00162376 | 0.216512 | 0.3870335 | -0.475212 | 4.36E-06 | -0.142619 | 0.3820098 |
| ENSG00000115170.13 | ACVR1 | 0.552606 | 0.6214382 | 0.372338 | 0.9102076 | 0.344831 | 0.9567706 | 1.02169 | 0.02448063 | 3.859854 | 0.2254389 | 2.04328 | 0.6044994 |
| ENSG00000115183.13 | TANC1 | -0.080296 | 0.825688 | -0.258066 | 0.5861593 | 0.209016 | 0.1183712 | 2.041802 | 0.005614985 | 1.388529 | 0.6626174 | 1.351884 | 0.9372892 |
| ENSG00000115204.14 | MPV17 | -1.302175 | 0.5428806 | 0.162089 | 0.5953732 | -0.795092 | 0.02908326 | -0.17517 | 0.3445634 | 0.103177 | 0.5279031 | -0.179218 | 0.9352001 |
| ENSG00000115207.13 | GTF3C2 | 1.706914 | 0.4764974 | 0.840686 | 0.8263112 | 0.898747 | 0.1781189 | 2.050672 | 0.006293281 | 1.247977 | 0.5770961 | -0.141919 | 1 |
| ENSG00000115211.15 | EIF2B4 | 0.24362 | 0.8787256 | -0.00709 | 0.6527736 | -0.412928 | 0.01753901 | 1.294428 | 2.62E-05 | 0.939739 | 0.08159642 | -0.044954 | 0.9724348 |
| ENSG00000115233.11 | PSMD14 | 0.315219 | 0.8006902 | 0.190836 | 0.7834199 | 0.069101 | 0.4064599 | -0.006589 | 0.0237393 | 0.318366 | 0.002470028 | 0.002191 | 0.6885065 |
| ENSG00000115234.10 | SNX17 | -2.694144 | 0.6792949 | 0.180226 | 0.9625534 | 0.564026 | 0.8357355 | 0.043444 | 0.8016097 | -1.353173 | 0.03327974 | 0.34116 | 0.9965354 |
| ENSG00000115241.10 | PPM1G | -0.473983 | 0.5657211 | -0.081034 | 0.2250307 | -1.208031 | 0.0107484 | 0.133047 | 0.3760083 | 0.434525 | 0.07029776 | 0.030266 | 0.359138 |
| ENSG00000115266.11 | APC2 | 0.760331 | 0.8006286 | -0.403974 | 0.8845769 | -1.162129 | 0.5654652 | 1.930645 | 0.002892901 | 1.597215 | 0.000383261 | -0.684827 | 0.6939365 |
| ENSG00000115268.9 | RPS15 | -0.523469 | 0.3946735 | -0.197027 | 0.159232 | -0.330393 | 2.13E-06 | 0.148884 | 0.7144938 | 0.380974 | 0.1852973 | -0.261158 | 0.08347771 |
| ENSG00000115275.11 | MOGS | 0.215998 | 0.9282145 | 0.173416 | 0.8953649 | -0.191455 | 0.008787921 | 0.39873 | 0.000316073 | 1.715452 | 0.09855349 | -0.465116 | 0.8494077 |
| ENSG00000115286.19 | NDUFS7 | 0.546755 | 0.6214382 | -0.097197 | 0.6200225 | 0.241822 | 0.7943041 | -0.032516 | 0.007728367 | 0.47068 | 0.4286052 | -0.233838 | 0.6829816 |
| ENSG00000115289.12 | PCGF1 | -0.859702 | 0.934454 | 0.919582 | 0.4929255 | -2.655528 | 0.002317644 | -0.234215 | 0.7211844 | -0.686448 | 0.4651854 | 0.808176 | 0.8309897 |
| ENSG00000115306.15 | SPTBN1 | -0.836512 | 0.6014343 | -0.798204 | 0.000145333 | 0.43811 | 0.1469899 | 0.522408 | 0.003544531 | 2.292962 | 0.004089181 | -0.592733 | 0.9169962 |
| ENSG00000115310.17 | RTN4 | 0.043183 | 0.529676 | -0.226253 | 0.2388066 | 0.461168 | 0.008642232 | 0.884855 | 1.36E-06 | 0.623043 | 0.7731787 | 0.306222 | 0.6517664 |
| ENSG00000115350.11 | POLE4 | -1.255472 | 0.6792949 | -0.335107 | 0.704767 | -0.479618 | 8.03E-05 | 0.380754 | 0.9039446 | 0.884444 | 0.8481737 | -0.26295 | 0.6579597 |
| ENSG00000115355.15 | CCDC88A | -1.060281 | 0.007696271 | -0.117898 | 0.6527736 | 1.546009 | 1.15E-23 | 0.654924 | 0.04221781 | 0.391937 | 0.715399 | 0.306806 | 0.8479431 |
| ENSG00000115380.19 | EFEMP1 | -5.171373 | 0.6214382 | 0.018965 | 0.7881421 | -1.170809 | 0.3266608 | 0.823394 | 0.02380383 | 1.010766 | 0.9917421 | 0.277019 | 0.6567003 |
| ENSG00000115415.18 | STAT1 | 0.27077 | 0.3009675 | -0.286947 | 0.5905303 | 0.182693 | 0.8425963 | 1.174137 | 0.02773246 | 1.679086 | 0.05688148 | 0.983138 | 0.6885065 |
| ENSG00000115419.12 | GLS | -0.64528 | 0.7736255 | 0.257513 | 0.8718989 | 0.339743 | 0.000217556 | 1.084638 | 0.133515 | 2.01543 | 0.005390303 | -0.841049 | 0.6885065 |
| ENSG00000115421.12 | PAPOLG | 1.880926 | 0.8867323 | -0.566713 | 0.8679705 | -0.774819 | 0.7974083 | 2.310421 | 0.03581853 | 0.07514 | 0.6568257 | 3.171749 | 0.6885065 |
| ENSG00000115457.9 | IGFBP2 | 0.023408 | 0.928247 | 0.928917 | 9.16E-16 | -1.44748 | 4.40E-19 | 1.136905 | 0.002778742 | 0.439198 | 0.6367485 | -0.270992 | 0.08805564 |
| ENSG00000115461.4 | IGFBP5 | -2.147215 | 0.529676 | 0.461212 | 5.94E-21 | -0.639197 | 0.002034972 | 1.022566 | 2.47E-08 | 0.406353 | 0.256083 | -0.126607 | 0.2159565 |
| ENSG00000115464.14 | USP34 | -0.618952 | 0.4764974 | 0.163402 | 0.7808545 | 0.584243 | 0.1449399 | 1.517353 | 1.78E-07 | 0.743288 | 0.5196843 | -0.180963 | 0.9377765 |
| ENSG00000115486.11 | GGCX | -2.601334 | 0.3613761 | -0.229143 | 0.7550531 | -1.266417 | 0.02661331 | 0.927004 | 0.03289865 | 0.222126 | 1 | 0.795705 | 0.9276719 |
| ENSG00000115504.14 | EHBP1 | -0.337158 | 0.3982827 | -0.599515 | 0.04105643 | 0.036098 | 0.9753354 | 0.403308 | 0.2137901 | 0.295725 | 0.933176 | -0.234895 | 0.6861021 |
| ENSG00000115524.15 | SF3B1 | -0.420162 | 0.6214382 | -0.087996 | 0.3121527 | 0.478246 | 3.60E-12 | 0.096169 | 0.5255202 | 0.393672 | 0.7991562 | 0.039159 | 0.6813272 |
| ENSG00000115525.16 | ST3GAL5 | 0.185343 | 0.8434435 | -0.684612 | 0.1777918 | -0.262927 | 0.607072 | 1.703938 | 6.26E-06 | 0.610603 | 0.2546941 | -0.251707 | 0.6829816 |
| ENSG00000115526.10 | CHST10 | -1.98922 | 0.6214382 | 0.924243 | 0.9017558 | 1.983592 | 0.953937 | 1.798262 | 0.000759227 | -0.696187 | 0.3114707 | 0.84902 | 0.8949489 |
| ENSG00000115539.13 | PDCL3 | -0.680695 | 0.6214382 | -0.138835 | 0.8149803 | 0.02485 | 0.5290845 | -0.856944 | 0.02076594 | 0.485217 | 0.06268192 | -2.439561 | 0.4650482 |
| ENSG00000115541.10 | HSPE1 | -0.527593 | 0.4692975 | -0.033118 | 0.8703522 | -0.485117 | 1.44E-08 | -0.027911 | 0.000104536 | 0.218712 | 0.000167471 | -0.551669 | 0.05025511 |
| ENSG00000115548.16 | KDM3A | 1.723475 | 0.4764974 | 0.508154 | 0.9329105 | 1.430141 | 0.02744378 | -0.569702 | 0.8185887 | 1.543262 | 0.03056663 | 0.989723 | 0.9687428 |
| ENSG00000115649.15 | CNPPD1 | -0.41336 | 0.8587992 | -0.222199 | 0.4375617 | 0.588825 | 0.7437308 | -1.081878 | 0.00253872 | 0.525876 | 0.8385533 | -0.271182 | 0.6829816 |
| ENSG00000115677.16 | HDLBP | 0.025772 | 1 | 0.221183 | 0.6527736 | 0.713911 | 0.000295921 | 0.803637 | 0.3647127 | -0.379484 | 0.4629348 | 0.109002 | 0.8168211 |
| ENSG00000115694.14 | STK25 | -0.918994 | 0.8006286 | 0.066713 | 0.7139087 | -0.33488 | 0.3079733 | 0.674951 | 0.04680542 | 0.278431 | 0.311879 | -1.262707 | 0.8949489 |
| ENSG00000115750.16 | TAF1B | 0.944983 | 0.6552695 | 0.302818 | 0.9918484 | 0.758809 | 0.03578271 | -0.300844 | 0.4919368 | -0.146586 | 0.1863088 | -2.873407 | 0.8406977 |
| ENSG00000115756.12 | HPCAL1 | 0.476534 | 0.6214382 | 0.235184 | 0.7958166 | -0.209275 | 0.5290845 | 0.992312 | 0.6480023 | 2.446648 | 0.01412561 | 0.715938 | 0.9270789 |
| ENSG00000115758.12 | ODC1 | -1.00738 | 0.1835496 | -0.084539 | 0.6250195 | -0.141831 | 0.8454005 | 1.751365 | 4.81E-09 | 0.995891 | 0.0155799 | -0.002082 | 0.2068179 |
| ENSG00000115760.13 | BIRC6 | -0.770447 | 0.7736255 | 0.293047 | 0.9931449 | 1.243192 | 0.06103368 | 2.091428 | 7.29E-05 | 0.875797 | 0.4398108 | -0.763874 | 0.9581586 |
| ENSG00000115761.15 | NOL10 | 0.031943 | 0.9185653 | -0.582244 | 0.2416476 | -0.219407 | 0.7479576 | 0.553897 | 0.04691967 | 1.763202 | 0.621723 | -0.903827 | 0.7419782 |
| ENSG00000115816.13 | CEBPZ | 0.226711 | 0.6014343 | -0.125327 | 0.8976809 | 0.253067 | 0.0479416 | 1.239225 | 0.000730442 | 0.938481 | 0.6617637 | 1.004782 | 0.9106535 |
| ENSG00000115825.9 | PRKD3 | -1.584937 | 0.6792949 | -0.599823 | 0.1834552 | 1.459659 | 0.00041897 | 0.404716 | 0.4919368 | 0.217302 | 0.5908104 | -0.072731 | 0.6885065 |
| ENSG00000115866.10 | DARS | -3.40963 | 0.8006286 | 0.217379 | 0.8244383 | 0.685733 | 0.6928639 | 0.085964 | 0.7620448 | -0.625644 | 0.01511994 | 0.00831 | 0.8949489 |
| ENSG00000115884.10 | SDC1 | 0.057011 | 1 | -1.343506 | 0.5949213 | 0.541713 | 0.768155 | 1.483571 | 0.03460858 | 2.382925 | 0.2840365 | -0.415285 | 0.6262252 |
| ENSG00000115904.12 | SOS1 | 1.70543 | 0.8006286 | -0.109981 | 0.8472233 | 1.235454 | 0.02207811 | 1.218457 | 0.04689817 | 2.035257 | 0.08729226 | 1.454692 | 0.6885065 |
| ENSG00000115946.7 | PNO1 | -0.370804 | 0.737082 | -0.28228 | 0.5953732 | 0.272209 | 0.7575356 | -0.447253 | 0.09720862 | 0.532471 | 0.0463146 | 1.459256 | 0.9915086 |
| ENSG00000115963.13 | RND3 | -0.649098 | 0.9183025 | -0.027506 | 0.9425616 | -0.920752 | 0.4792527 | 0.612679 | 0.01263687 | -0.191613 | 0.3974318 | 0.721818 | 0.6885065 |
| ENSG00000115970.18 | THADA | -1.575876 | 0.6792949 | -0.203667 | 0.7952779 | 0.560435 | 0.5071792 | 2.156967 | 0.02466897 | -0.511983 | 0.9463565 | -2.241699 | 0.8949489 |
| ENSG00000115993.11 | TRAK2 | -1.062142 | 0.8144419 | 0.121817 | 0.8818962 | 0.319338 | 0.0140601 | 0.938934 | 6.06E-06 | 1.038307 | 0.94203 | -0.326892 | 0.6885065 |
| ENSG00000116001.15 | TIA1 | -1.382102 | 0.8859625 | -0.203165 | 0.5939722 | 0.351612 | 0.07724106 | 1.228476 | 0.01427716 | 0.379175 | 0.9337825 | 0.433545 | 0.6885065 |
| ENSG00000116005.11 | PCYOX1 | -1.531652 | 0.5706649 | -0.207552 | 0.09999267 | 1.442174 | 0.08151947 | 0.645558 | 0.04798097 | 0.562351 | 0.5479355 | 1.447623 | 0.9912078 |
| ENSG00000116017.10 | ARID3A | -2.081959 | 0.563505 | 0.362296 | 0.7710216 | 1.316299 | 0.528724 | 1.567241 | 0.4919368 | 3.156285 | 0.003116612 | -1.095615 | 0.5733051 |
| ENSG00000116096.5 | SPR | 1.052254 | 0.6014343 | 0.001761 | 0.8472204 | -2.113477 | 0.03616489 | -0.81576 | 0.3760083 | 2.656315 | 0.7242087 | 0.213036 | 0.826754 |
| ENSG00000116133.11 | DHCR24 | 1.111534 | 0.185057 | -0.965904 | 0.2521066 | 0.274409 | 0.9063164 | 1.85535 | 0.03409007 | -0.906995 | 0.8173955 | -0.566257 | 0.9403369 |
| ENSG00000116138.12 | DNAJC16 | -1.218049 | 1 | 1.206705 | 0.6527736 | 2.023861 | 0.1651076 | 1.715673 | 0.01644999 | -0.056046 | 0.5475627 | 1.821231 | 0.9002633 |
| ENSG00000116147.16 | TNR | 0.10734 | 0.8587992 | -0.049137 | 0.2713328 | 1.271419 | 0.04560151 | -0.50138 | 0.954187 | 2.055001 | 0.2854946 | 4.840226 | 0.8949489 |
| ENSG00000116171.16 | SCP2 | 0.205498 | 1 | 0.249948 | 0.8263112 | 0.385235 | 0.996049 | 0.56855 | 0.429227 | -0.622625 | 0.000500995 | 0.881304 | 0.8479431 |
| ENSG00000116212.14 | LRRC42 | -1.69167 | 0.6214382 | 0.290739 | 0.9043399 | 1.295117 | 0.005189828 | 0.338769 | 1 | -0.603137 | 0.311879 | -0.223073 | 1 |
| ENSG00000116251.9 | RPL22 | -0.65417 | 0.529676 | -0.030444 | 0.6527736 | 0.107524 | 0.2279632 | -0.029321 | 0.03423802 | -0.368441 | 4.57E-06 | 0.015102 | 0.6885065 |
| ENSG00000116254.17 | CHD5 | 0.484538 | 1 | -2.53916 | 0.4487482 | -1.089218 | 0.5291749 | -0.416533 | 0.7858185 | 1.868675 | 0.875882 |  |  |
| ENSG00000116260.16 | QSOX1 | 0.949113 | 0.3132333 | 0.80406 | 0.5296018 | -0.192441 | 0.6928639 | 1.252248 | 0.04188543 | 0.579225 | 0.9387909 | 0.8401 | 0.8479431 |
| ENSG00000116288.12 | PARK7 | -0.235812 | 0.6792949 | -0.232439 | 0.02065288 | 0.137901 | 0.5430149 | 0.354904 | 0.9547491 | 0.052898 | 0.1172772 | -0.289344 | 0.6885065 |
| ENSG00000116337.15 | AMPD2 | -0.399473 | 1 | 0.185595 | 1 | 0.288479 | 1 | 2.105885 | 0.002167618 | 1.395987 | 0.06196625 | 0.827226 | 0.715468 |
| ENSG00000116459.10 | ATP5F1 | -0.004089 | 0.4755527 | 0.09056 | 0.7719688 | 0.133508 | 0.6852651 | -0.220297 | 0.007812163 | 0.03468 | 0.1318682 | -0.81134 | 0.4578379 |
| ENSG00000116514.16 | RNF19B | 1.304554 | 1 | 0.256674 | 1 | -2.807979 | 0.02155302 | 1.402798 | 0.08376554 | 2.201594 | 0.3974318 | -0.825885 | 0.6885065 |
| ENSG00000116521.10 | SCAMP3 | -0.01266 | 0.6014343 | 0.333053 | 0.8618881 | -0.413633 | 0.000646368 | 0.019123 | 0.4271336 | 0.140262 | 0.5196843 | -1.180575 | 1 |
| ENSG00000116539.10 | ASH1L | 0.421748 | 0.9662185 | -0.105552 | 0.2913444 | 0.830314 | 0.000291169 | 1.179698 | 0.009703093 | 0.868499 | 0.01657577 | 0.155611 | 0.8949489 |
| ENSG00000116560.10 | SFPQ | -0.354838 | 0.6214382 | -0.031475 | 0.7383998 | 0.215687 | 0.5290845 | 1.913858 | 1.50E-17 | 1.430423 | 7.87E-09 | -0.499611 | 0.2877472 |
| ENSG00000116586.11 | LAMTOR2 | -0.608713 | 0.9864501 | -0.033365 | 0.6527736 | -0.581095 | 0.000193258 | -0.137568 | 0.001931655 | -0.535826 | 0.5196843 | 0.574495 | 0.7780985 |
| ENSG00000116649.9 | SRM | 0.002237 | 0.8867323 | -0.166765 | 0.6444482 | -0.673827 | 1.24E-05 | 1.208586 | 2.53E-05 | 0.832306 | 0.3796044 | 1.173394 | 0.750045 |
| ENSG00000116670.14 | MAD2L2 | -0.281866 | 0.8092404 | -0.141731 | 0.7959668 | -0.582129 | 3.41E-05 | -0.284053 | 0.2131424 | 0.248894 | 0.4480469 | -0.010392 | 1 |
| ENSG00000116675.15 | DNAJC6 | 0.808269 | 0.714717 | -0.016093 | 0.9845079 | -0.82933 | 0.9139205 | 1.141282 | 0.0491633 | 0.50293 | 0.6340892 | -0.662254 | 0.9504164 |
| ENSG00000116688.16 | MFN2 | -1.158609 | 0.5711449 | 0.288132 | 0.9787499 | 0.178861 | 0.5232787 | 2.31253 | 0.000779335 | 0.336158 | 0.4281787 | 0.078091 | 0.7866071 |
| ENSG00000116698.20 | SMG7 | 1.007149 | 0.9569568 | 0.447762 | 0.9458404 | 1.1861 | 0.1595644 | 1.029474 | 0.01338072 | 1.491229 | 0.05411996 | 1.296692 | 0.8979257 |
| ENSG00000116729.13 | WLS | 0.252013 | 0.6180303 | -0.248729 | 0.2758583 | 0.152816 | 0.8650868 | 0.209699 | 0.02863136 | 0.335776 | 0.4073656 | 0.827468 | 0.7753492 |
| ENSG00000116731.22 | PRDM2 | -1.576898 | 0.7795461 | 0.415335 | 0.6636529 | 0.271308 | 0.2385939 | 0.960118 | 0.02974344 | 1.321298 | 0.1134992 | -1.258871 | 0.9581586 |
| ENSG00000116747.12 | TROVE2 | -2.943176 | 0.3431796 | 0.33664 | 0.2913444 | 0.559474 | 1.89E-06 | 0.509756 | 0.04032816 | 0.72459 | 0.5196843 | -1.23105 | 0.6829816 |
| ENSG00000116752.5 | BCAS2 | 0.730354 | 0.6214382 | 0.424467 | 0.7140253 | 0.125627 | 0.4430849 | 0.021816 | 0.05829131 | -0.042773 | 0.001459778 | 0.187689 | 0.8989893 |
| ENSG00000116754.13 | SRSF11 | 0.391997 | 0.8867323 | -0.219063 | 0.04321456 | 0.769356 | 2.24E-14 | 0.663415 | 0.5906907 | 0.34941 | 0.6445432 | -0.387864 | 0.9034098 |
| ENSG00000116852.14 | KIF21B | 0.945139 | 1 | -0.421402 | 0.9222921 | 1.658427 | 0.1157816 | 2.29614 | 0.008527215 | 0.313547 | 0.6500471 | -0.318791 | 0.9127491 |
| ENSG00000116857.16 | TMEM9 | 2.382238 | 0.007347962 | 0.275119 | 0.542173 | 0.370189 | 0.8333781 | 0.433207 | 0.7293477 | 0.954731 | 0.6097096 | 0.55214 | 0.6885065 |
| ENSG00000116871.15 | MAP7D1 | -0.158298 | 0.8587992 | -0.452614 | 0.6527736 | 0.283933 | 0.8386403 | 1.131706 | 0.009766286 | 1.025212 | 0.1427939 | -0.744352 | 0.7182399 |
| ENSG00000116898.11 | MRPS15 | 1.025946 | 0.6014343 | -0.13947 | 0.9848758 | -0.40229 | 0.004185562 | 0.884582 | 0.1505455 | -0.446861 | 0.7101365 | 0.12123 | 0.6885065 |
| ENSG00000116906.11 | GNPAT | 0.614301 | 1 | 0.20617 | 0.6542448 | -0.145452 | 0.7064607 | 1.207821 | 0.01800638 | 0.081654 | 0.7684919 | -2.663335 | 0.3394425 |
| ENSG00000116991.10 | SIPA1L2 | -3.253705 | 0.3111218 | -0.360838 | 0.3629265 | 0.765506 | 0.06987922 | 0.144486 | 0.1825859 | 1.470364 | 0.04511077 | -0.20521 | 1 |
| ENSG00000117000.8 | RLF | -0.033551 | 0.9047689 | -0.175651 | 0.8202471 | 0.691458 | 0.03056815 | 0.848409 | 0.2467163 | 1.434425 | 0.8092298 | 1.155963 | 0.6885065 |
| ENSG00000117020.16 | AKT3 | -0.480039 | 0.8265276 | 1.019717 | 0.86041 | 0.872498 | 0.02737868 | 1.453946 | 0.000851991 | 0.937349 | 0.04666956 | 2.102979 | 0.6885065 |
| ENSG00000117118.9 | SDHB | 0.848218 | 0.529676 | -0.011543 | 0.8637394 | 0.03482 | 0.630802 | -0.942478 | 0.01391062 | -0.28217 | 0.1134992 | 1.129919 | 0.8951582 |
| ENSG00000117139.16 | KDM5B | 0.058494 | 0.7644768 | -0.512072 | 0.78277 | 1.068434 | 0.000676574 | 1.152717 | 0.1476731 | 1.022055 | 0.5713168 | -1.351349 | 0.6885065 |
| ENSG00000117174.10 | ZNHIT6 | -0.646035 | 0.7780681 | -0.040065 | 0.8953649 | -0.012726 | 0.8454005 | 1.240172 | 0.01314281 | -0.499773 | 0.7377262 | 0.929493 | 0.7600927 |
| ENSG00000117318.8 | ID3 | -0.547206 | 0.9560231 | 0.364062 | 0.2779213 | -1.174248 | 1.85E-05 | -1.200986 | 1.23E-07 | -0.41156 | 0.01494005 | -1.044844 | 0.8102233 |
| ENSG00000117335.18 | CD46 | -1.088438 | 0.6214382 | 0.078464 | 0.9791082 | 0.187924 | 0.05977368 | 1.225682 | 0.01208972 | 1.097016 | 1 | 0.65878 | 0.8662714 |
| ENSG00000117360.12 | PRPF3 | 0.596182 | 1 | 0.008631 | 0.6527736 | 1.283282 | 0.009327078 | 0.790356 | 1 | 1.008838 | 0.6610687 | 0.001078 | 0.8343319 |
| ENSG00000117394.19 | SLC2A1 | 3.555009 | 0.6214382 | 0.958742 | 0.00432258 | -0.885727 | 0.04954543 | 0.566841 | 0.3388812 | 0.046839 | 0.5196843 | -0.430069 | 0.9188279 |
| ENSG00000117410.13 | ATP6V0B | 0.621185 | 0.4322665 | 0.217244 | 0.633346 | -0.107941 | 0.004852497 | 0.015876 | 0.05147036 | 0.186725 | 0.5196843 | 0.678845 | 0.9147227 |
| ENSG00000117450.13 | PRDX1 | 0.828221 | 0.6214382 | 0.10312 | 0.6813155 | 0.150859 | 0.5290845 | 0.216579 | 0.3719356 | 0.108122 | 0.04159686 | -0.128451 | 0.6879794 |
| ENSG00000117475.13 | BLZF1 | -1.135381 | 0.8006286 | -0.359592 | 0.6527736 | 1.403915 | 0.01812922 | 1.105463 | 0.9010661 | 0.574821 | 0.7377262 | 0.950455 | 0.826754 |
| ENSG00000117523.15 | PRRC2C | 0.061964 | 0.7383533 | -0.022499 | 0.5649416 | 0.90949 | 1.68E-14 | 1.308329 | 1.09E-09 | 0.844938 | 0.08708746 | 0.812891 | 0.9713975 |
| ENSG00000117592.8 | PRDX6 | -1.438121 | 0.1105572 | -0.05522 | 0.6527736 | -0.323183 | 0.00401426 | 0.143897 | 0.0690253 | 0.36291 | 0.08425073 | 0.081925 | 1 |
| ENSG00000117614.9 | SYF2 | 1.14025 | 0.9282145 | 0.198847 | 1 | 0.034543 | 0.2768914 | 0.497605 | 0.3196248 | -0.337585 | 0.006214297 | 0.542255 | 1 |
| ENSG00000117616.17 | RSRP1 | -1.281353 | 0.714717 | -0.137963 | 0.9104667 | 0.411702 | 0.1048414 | 1.357228 | 0.002447063 | 1.431632 | 0.06904104 | 1.017088 | 0.9506884 |
| ENSG00000117625.13 | RCOR3 | -0.388829 | 0.6214382 | 0.132917 | 0.6527736 | 0.498881 | 0.005011579 | 1.319063 | 0.1625797 | 1.388609 | 0.1924287 | -0.394214 | 0.6829816 |
| ENSG00000117632.20 | STMN1 | 0.00698 | 0.6603437 | -0.200215 | 0.5953732 | 0.157957 | 0.5501546 | 0.517392 | 0.4919368 | 0.093428 | 0.2358413 | -0.616395 | 0.01369193 |
| ENSG00000117707.15 | PROX1 | -1.098328 | 0.6014343 | -0.192991 | 0.85618 | 1.261386 | 4.09E-05 | 0.676264 | 0.03891837 | 0.885501 | 0.03247574 | -2.281365 | 0.9281839 |
| ENSG00000117713.17 | ARID1A | -0.42848 | 0.9478876 | -0.3338 | 0.4072962 | 0.857966 | 0.08165435 | 1.617343 | 1.33E-05 | 1.488022 | 0.01281183 | 0.7056 | 0.6885065 |
| ENSG00000117758.13 | STX12 | 1.499985 | 0.8006286 | -0.117279 | 0.7233606 | 0.694131 | 0.005304962 | 0.964443 | 0.9556238 | 0.6443 | 0.2444518 | -0.442513 | 0.7780985 |
| ENSG00000117868.15 | ESYT2 | -0.97776 | 0.6214382 | 0.182817 | 1 | 0.576568 | 0.7105729 | 0.415832 | 0.008110023 | 1.885722 | 0.1249186 | -1.094781 | 0.7961473 |
| ENSG00000117877.10 | CD3EAP | 0.081539 | 0.7031212 | -0.498367 | 0.4409757 | 0.244379 | 1 | 1.445685 | 0.02018338 | -0.562586 | 0.3380778 | -0.956261 | 0.5202568 |
| ENSG00000117984.12 | CTSD | 1.431207 | 0.5706649 | 0.186259 | 0.48118 | -0.510761 | 0.04729265 | 1.172505 | 0.2077593 | -0.482427 | 0.02095403 | 0.614323 | 0.5601856 |
| ENSG00000118046.14 | STK11 | 1.767017 | 0.529676 | -0.258184 | 0.7150943 | -0.816365 | 0.6198222 | 2.161236 | 0.000376492 | 2.407524 | 0.001791482 | 1.4918 | 0.9771994 |
| ENSG00000118058.20 | KMT2A | 0.193458 | 1 | -0.404906 | 0.05116306 | 0.645005 | 1.31E-08 | 0.627167 | 0.001116302 | 0.201148 | 0.08166633 | -2.34098 | 0.1508459 |
| ENSG00000118181.10 | RPS25 | -0.696223 | 0.01796694 | 0.180997 | 0.5818005 | 0.044904 | 0.7437308 | 0.214035 | 0.8224647 | 0.284693 | 0.3753397 | 0.066019 | 0.6185816 |
| ENSG00000118200.14 | CAMSAP2 | 0.191997 | 0.9737959 | -0.321553 | 0.4607773 | 0.371061 | 0.05838784 | 1.25729 | 1.35E-06 | 1.182534 | 0.1401461 | 0.859516 | 0.9147227 |
| ENSG00000118217.5 | ATF6 | -0.102356 | 0.8006286 | -0.285236 | 0.3121527 | 0.683214 | 0.007013421 | 0.407224 | 0.4919368 | -0.682534 | 0.1869176 | 0.814866 | 0.6829816 |
| ENSG00000118246.13 | FASTKD2 | 1.161112 | 0.5467762 | 0.096714 | 0.704767 | -0.795866 | 0.5885306 | 2.112695 | 0.02777876 | -0.607911 | 0.8346216 | -1.407593 | 0.7824117 |
| ENSG00000118263.14 | KLF7 | 0.594638 | 0.6214382 | -0.134705 | 0.6527736 | -0.65058 | 0.5290845 | 1.69017 | 0.005069884 | -0.205205 | 0.2880317 | 0.042304 | 0.8479431 |
| ENSG00000118363.11 | SPCS2 | -0.257351 | 0.7736255 | 0.303802 | 0.5953732 | 0.112709 | 0.401001 | -0.070967 | 0.05954403 | -0.498289 | 0.000828531 | 0.519534 | 0.6885065 |
| ENSG00000118402.5 | ELOVL4 | 0.01552 | 0.563505 | 0.229554 | 0.7894251 | 0.933214 | 0.03566389 | 0.985759 | 0.01735437 | -0.097133 | 0.4874815 | 0.641601 | 0.8949489 |
| ENSG00000118412.12 | CASP8AP2 | -0.202654 | 0.8103468 | -0.099118 | 0.5010139 | 0.022971 | 0.7437308 | 1.178044 | 0.02231367 | 0.872657 | 0.367828 | 2.147671 | 0.7356415 |
| ENSG00000118482.11 | PHF3 | -0.74244 | 0.8587992 | -0.146077 | 0.6527736 | 1.586009 | 2.06E-07 | 1.322946 | 0.03032508 | 0.331689 | 0.2372778 | -0.247452 | 0.6829816 |
| ENSG00000118496.4 | FBXO30 | -2.648344 | 0.6014343 | 0.949222 | 0.4011918 | 0.093906 | 0.1896632 | 2.194595 | 0.007661563 | 0.880808 | 0.8719628 | -0.828675 | 1 |
| ENSG00000118596.11 | SLC16A7 | 1.039925 | 1 | -0.435072 | 0.5633052 | 1.376141 | 0.5290845 | 3.557354 | 0.03968708 | -1.836877 | 0.9343644 | inf | 0.5344421 |
| ENSG00000118620.12 | ZNF430 | -2.296288 | 0.4692975 | -0.445832 | 0.5953732 | 0.231297 | 0.1958214 | 0.09522 | 0.1360198 | 2.024121 | 0.01787409 | -5.116019 | 0.6829816 |
| ENSG00000118689.14 | FOXO3 | -0.69927 | 0.9441665 | -0.220399 | 0.6054308 | 0.46645 | 0.01735633 | 1.018906 | 0.01144777 | 0.702542 | 0.152231 | -0.674875 | 0.9786489 |
| ENSG00000118762.7 | PKD2 | -0.337988 | 0.6214382 | -0.769081 | 0.6527736 | 0.48812 | 0.6713035 | 1.290108 | 0.004190918 | 1.966036 | 0.06067043 | 1.36479 | 0.9380942 |
| ENSG00000118785.13 | SPP1 | -0.473762 | 0.8946975 | -0.753967 | 1.11E-34 | -0.168484 | 0.1076488 | -0.001358 | 0.6985173 | -1.711768 | 0.001493672 | 0.850366 | 2.43E-05 |
| ENSG00000118816.9 | CCNI | -0.306153 | 0.5119304 | 0.154424 | 0.8953649 | 0.15939 | 0.001054705 | 0.237251 | 0.9524866 | 0.404294 | 0.9056171 | 0.144106 | 0.6517664 |
| ENSG00000118873.15 | RAB3GAP2 | -0.058566 | 0.8006286 | -0.276317 | 0.386448 | 0.345058 | 0.2518622 | 1.112374 | 0.002892901 | 0.361475 | 1 | -1.042192 | 0.9485566 |
| ENSG00000118894.14 | EEF2KMT | 0.027521 | 0.739831 | 0.255111 | 0.7383998 | -1.963907 | 0.000739632 | -0.787365 | 0.04856638 | -0.920686 | 0.4459269 | 1.433219 | 0.9238214 |
| ENSG00000118900.14 | UBN1 | 0.127596 | 1 | 0.160725 | 0.7304297 | 0.491291 | 0.5476072 | 1.899436 | 0.04089319 | 1.265079 | 0.2581597 | 1.512837 | 0.6885065 |
| ENSG00000118922.16 | KLF12 | 0.00834 | 0.8587992 | -0.356296 | 1 | 1.395538 | 0.2701559 | 0.655296 | 0.03329343 | 0.960411 | 0.1480744 | 0.561832 | 0.6885065 |
| ENSG00000118939.17 | UCHL3 | 0.543903 | 0.6014343 | -0.708978 | 0.6527736 | -0.862561 | 0.004787728 | 0.588948 | 0.4919368 | 0.820435 | 0.7175176 | 0.248501 | 0.7817802 |
| ENSG00000118946.11 | PCDH17 | 0.328911 | 0.7231646 | -0.736072 | 0.1689344 | 0.481702 | 1.60E-09 | 0.43368 | 0.05157198 | 1.87529 | 0.0148673 | 0.743717 | 0.9581586 |
| ENSG00000118971.7 | CCND2 | -0.211866 | 0.6615279 | -0.35018 | 0.03324584 | 1.155649 | 1.41E-19 | 0.805177 | 0.01595271 | 0.811632 | 1.33E-17 | 0.292205 | 0.9532204 |
| ENSG00000119004.14 | CYP20A1 | -0.804183 | 0.5468595 | 0.586903 | 0.4625157 | 0.016163 | 0.5259115 | 1.516801 | 0.005871422 | 0.138093 | 0.2664037 | -1.026241 | 0.8842299 |
| ENSG00000119013.8 | NDUFB3 | -1.54509 | 0.2934592 | 0.168785 | 0.918309 | -0.406494 | 6.09E-09 | -0.579383 | 3.62E-06 | -0.205948 | 0.02117621 | -0.365675 | 0.2835947 |
| ENSG00000119042.16 | SATB2 | 1.475278 | 0.65581 | 0.028233 | 0.7331946 | 0.476127 | 0.9447956 | 1.444702 | 0.00051747 | 2.668771 | 0.4702212 | -1.121917 | 0.6885065 |
| ENSG00000119185.12 | ITGB1BP1 | 0.493116 | 0.8569863 | -0.117085 | 0.6530513 | -0.373964 | 0.01419177 | -0.028487 | 0.1273493 | 0.085471 | 0.2121356 | -0.074331 | 0.8479431 |
| ENSG00000119242.8 | CCDC92 | -0.540531 | 0.7703009 | 0.447063 | 0.6527736 | -1.185325 | 0.4189965 | -0.23781 | 0.04239671 | 1.659361 | 0.08714613 | 1.821074 | 0.6885065 |
| ENSG00000119314.15 | PTBP3 | -0.6542 | 1 | -0.975228 | 0.1834552 | 1.06126 | 0.1936049 | 1.375881 | 0.02110829 | -0.053527 | 0.7662962 | -1.027233 | 0.9147227 |
| ENSG00000119318.12 | RAD23B | -0.031205 | 0.8006286 | -0.066611 | 0.6527736 | 0.025121 | 0.9447956 | 0.989591 | 1.99E-05 | 0.548545 | 0.4491712 | -0.644629 | 0.8183573 |
| ENSG00000119333.11 | WDR34 | -1.545247 | 0.8006286 | -0.145913 | 0.6331318 | -0.867505 | 0.002391157 | 0.104532 | 0.2732557 | 0.725159 | 0.6770529 | -2.505378 | 0.6885065 |
| ENSG00000119335.16 | SET | -0.476897 | 0.7486452 | 0.007776 | 0.6527736 | 0.522529 | 3.05E-08 | 0.656454 | 3.84E-05 | 0.697382 | 0.2339043 | 0.331089 | 0.2159565 |
| ENSG00000119392.14 | GLE1 | 0.617334 | 0.6214382 | -0.422043 | 0.6527736 | 1.022286 | 0.5645295 | 0.013824 | 0.02277414 | 1.326272 | 0.5196843 | -1.771552 | 0.8421466 |
| ENSG00000119396.10 | RAB14 | -0.086016 | 0.6014343 | 0.250714 | 0.9500565 | 0.464434 | 0.001345929 | 0.65235 | 0.8987922 | -0.045227 | 0.355045 | 1.033885 | 1 |
| ENSG00000119408.16 | NEK6 | 0.49391 | 0.7400552 | -0.377167 | 0.0168514 | -0.453895 | 0.000233683 | 0.886531 | 0.4360187 | 0.970908 | 0.7219788 | 0.225888 | 0.8995933 |
| ENSG00000119421.6 | NDUFA8 | 0.015764 | 0.8006286 | -0.186853 | 0.6527736 | -0.719945 | 0.002185914 | -0.353241 | 0.05080366 | -0.279596 | 0.002351616 | 0.235064 | 0.8479431 |
| ENSG00000119446.13 | RBM18 | 0.698916 | 0.3098608 | 0.208673 | 1 | 0.139141 | 1 | 1.004013 | 0.3732949 | 0.880043 | 0.04781087 | -0.206698 | 0.9222391 |
| ENSG00000119537.15 | KDSR | -1.019169 | 0.008112828 | -0.148218 | 0.9187044 | 0.467219 | 0.5220842 | -0.09825 | 0.1464185 | 0.002781 | 0.3204341 | 0.49221 | 1 |
| ENSG00000119574.12 | ZBTB45 | 1.289528 | 1 | 0.877402 | 0.5953732 | -2.801888 | 9.68E-05 | 2.345821 | 0.003119219 | 1.642824 | 0.7529496 | 1.150469 | 0.8479431 |
| ENSG00000119596.17 | YLPM1 | 0.226633 | 0.9666953 | -0.460976 | 0.2902651 | 1.162927 | 0.01793392 | 1.049069 | 0.3467508 | 0.018895 | 0.9201079 | 0.591828 | 1 |
| ENSG00000119650.12 | IFT43 | 0.865114 | 0.6014343 | -0.002555 | 0.8326284 | 0.452922 | 0.4176494 | 0.301677 | 0.02808548 | 0.860936 | 0.5196843 | 0.350594 | 0.7969605 |
| ENSG00000119655.8 | NPC2 | -0.491046 | 0.6214382 | 0.621757 | 0.4531355 | -1.323407 | 0.3994133 | 0.064091 | 0.04352887 | 0.505562 | 0.6188988 | 0.58282 | 0.6885065 |
| ENSG00000119669.4 | IRF2BPL | 0.051478 | 0.941967 | -0.462821 | 0.2215154 | 0.141045 | 0.5461593 | 1.253073 | 0.001643361 | 1.520691 | 0.01472873 | 0.030099 | 0.9733288 |
| ENSG00000119682.16 | AREL1 | -0.443898 | 0.6214382 | -0.040498 | 0.7881421 | -0.110884 | 0.8984213 | 2.984434 | 0.000100542 | 2.285802 | 0.597842 | -1.533576 | 0.6885065 |
| ENSG00000119705.9 | SLIRP | 0.298278 | 0.001633463 | -0.138562 | 0.4293459 | -0.461981 | 1.38E-06 | -0.034512 | 0.02509447 | -0.468743 | 0.00648229 | -0.101292 | 0.186512 |
| ENSG00000119707.13 | RBM25 | -0.126377 | 0.04094829 | -0.303641 | 0.024673 | 0.410688 | 0.00098187 | 1.185052 | 5.16E-11 | 0.563773 | 0.4148547 | 0.031514 | 0.8949489 |
| ENSG00000119729.10 | RHOQ | 0.919152 | 0.9183025 | -0.383461 | 0.2913444 | 0.278634 | 0.9289111 | 1.621305 | 0.03894757 | 0.004883 | 0.5757457 | -0.956177 | 0.6185816 |
| ENSG00000119772.16 | DNMT3A | 0.076517 | 0.9495183 | -0.397674 | 0.8208804 | 0.067463 | 0.2879524 | 2.053918 | 0.01841704 | 0.587731 | 0.5804605 | -0.149345 | 0.9745541 |
| ENSG00000119778.14 | ATAD2B | -1.800007 | 0.714717 | 0.605616 | 0.8217443 | 2.41712 | 0.1465489 | 4.014989 | 0.000945659 | 0.997138 | 0.6217996 | -1.82403 | 0.6185816 |
| ENSG00000119820.10 | YIPF4 | 0.067334 | 0.6214382 | -0.108205 | 0.6529973 | 0.126872 | 0.5020068 | 1.618848 | 2.15E-06 | 0.899582 | 0.5131507 | -0.904342 | 0.6413438 |
| ENSG00000119844.15 | AFTPH | 0.41595 | 0.8006286 | -0.525604 | 0.5010139 | -0.048066 | 0.6517876 | 2.076641 | 0.002926985 | 0.398276 | 0.1166035 | 0.735699 | 0.9830322 |
| ENSG00000119878.5 | CRIPT | 0.210229 | 0.6214382 | -0.051194 | 0.9593601 | -0.162659 | 0.5290845 | 0.192755 | 0.9613679 | -1.107333 | 0.03048683 | 0.187976 | 0.922577 |
| ENSG00000119906.11 | SLF2 | -1.639516 | 0.1956577 | -0.824312 | 0.2401656 | 0.340077 | 0.1616046 | 1.860322 | 0.03284013 | 0.110079 | 0.4278631 | -0.404356 | 0.4120408 |
| ENSG00000119950.20 | MXI1 | -0.953586 | 0.7709851 | 0.366517 | 0.6527736 | 1.64973 | 0.01366175 | 0.099412 | 0.7144938 | 1.128325 | 0.367138 | -1.421041 | 1 |
| ENSG00000119969.14 | HELLS | -1.215863 | 0.8006286 | -0.743117 | 0.6527736 | 0.984095 | 0.2490155 | 1.283376 | 0.04615682 | 0.978049 | 0.506562 | 0.765055 | 0.9727615 |
| ENSG00000120063.9 | GNA13 | 0.780558 | 0.8006286 | -0.169928 | 0.592336 | 2.036078 | 0.06376737 | 0.546762 | 0.005901793 | 1.334151 | 0.122358 | 0.150643 | 0.9899706 |
| ENSG00000120071.12 | KANSL1 | 0.694538 | 0.5711449 | 0.191559 | 0.9096651 | 0.19313 | 0.4727153 | 1.859229 | 0.000635083 | 1.109505 | 0.04057216 | -1.995793 | 0.6885065 |
| ENSG00000120129.5 | DUSP1 | 1.863518 | 0.191215 | 0.479228 | 0.3381481 | 0.45506 | 0.626664 | 0.677966 | 0.7087335 | 1.047651 | 0.000279287 | 1.031647 | 0.8564155 |
| ENSG00000120137.6 | PANK3 | 0.307026 | 0.9581872 | 0.223004 | 0.9638587 | 0.393861 | 0.1407419 | 0.703541 | 0.04595665 | 0.618348 | 0.687566 | -0.707294 | 0.8462399 |
| ENSG00000120251.18 | GRIA2 | -1.214813 | 0.6014343 | -0.160593 | 0.455084 | 0.996865 | 0.03314985 | 1.184748 | 0.002337486 | 1.095687 | 0.0250399 | -0.506203 | 0.9708573 |
| ENSG00000120265.16 | PCMT1 | 0.476128 | 0.6014343 | 0.347314 | 0.6527736 | 0.377493 | 0.6860841 | -0.314828 | 0.01983441 | -0.528697 | 0.01417919 | -0.027455 | 0.9617345 |
| ENSG00000120306.9 | CYSTM1 | -1.653035 | 0.08559909 | -0.029369 | 0.6527736 | -0.153303 | 0.001735405 | -0.702172 | 4.34E-05 | 0.210018 | 0.8794954 | -0.213749 | 0.9705209 |
| ENSG00000120438.11 | TCP1 | -0.557877 | 0.5119304 | 0.070001 | 0.9766239 | 0.605314 | 0.000213147 | -0.036955 | 0.7750325 | 0.580502 | 1 | -0.981238 | 0.6829816 |
| ENSG00000120451.10 | SNX19 | -1.076372 | 1 | -0.779077 | 0.6527736 | 0.467796 | 0.4761297 | 1.469672 | 0.006394826 | 0.565248 | 0.6829309 | -2.48246 | 0.7767294 |
| ENSG00000120509.10 | PDZD11 | -0.104485 | 0.8218433 | -0.128925 | 0.7755527 | -0.341471 | 0.06036305 | 0.35585 | 0.1100424 | -1.279089 | 0.00356938 | -0.400743 | 0.6185816 |
| ENSG00000120533.12 | ENY2 | 0.794378 | 0.3582986 | 0.433904 | 0.09467696 | -0.356662 | 0.4115773 | -0.528907 | 0.00233473 | -0.802332 | 2.10E-05 | 1.069577 | 1 |
| ENSG00000120616.15 | EPC1 | 0.475455 | 0.9791227 | 0.176751 | 1 | 0.872046 | 0.04027253 | 1.632749 | 0.09853275 | 0.660498 | 0.819072 | -0.455348 | 0.7804874 |
| ENSG00000120685.19 | PROSER1 | -0.134198 | 0.9047689 | -0.630105 | 0.7383998 | -1.030854 | 0.9147533 | 2.636986 | 0.001532897 | 1.33292 | 0.3053598 | 0.481312 | 0.8479431 |
| ENSG00000120694.19 | HSPH1 | 0.372231 | 0.8006286 | -0.615638 | 0.6527736 | 1.248008 | 1.32E-20 | 0.788525 | 0.03560962 | 0.738684 | 0.002886272 | -1.228515 | 0.5680347 |
| ENSG00000120696.8 | KBTBD7 | -1.932812 | 0.6214382 | 1.393408 | 1 | -1.047435 | 0.8108126 | 1.560364 | 0.4791525 | 2.634222 | 0.01467528 | inf | 0.6829816 |
| ENSG00000120697.8 | ALG5 | -0.353337 | 0.9853464 | 0.226255 | 0.8441696 | 0.544585 | 0.7693698 | 0.444338 | 0.2445427 | -0.634134 | 0.003235282 | 0.131018 | 0.7045294 |
| ENSG00000120705.12 | ETF1 | -0.45784 | 0.8763089 | 0.110207 | 0.8776804 | 0.693263 | 0.000169802 | 0.712312 | 0.2304771 | 0.170373 | 0.8815636 | -0.038009 | 0.6829816 |
| ENSG00000120733.13 | KDM3B | -0.118744 | 0.6792949 | -0.073969 | 0.9829438 | 0.566817 | 0.9441513 | 0.928527 | 0.001549484 | 1.058338 | 0.05340951 | 0.007229 | 0.6885065 |
| ENSG00000120738.7 | EGR1 | -0.691328 | 0.9184732 | -0.037133 | 0.867685 | -0.047705 | 0.3871766 | 0.832839 | 1.25E-05 | 0.502795 | 0.6962936 | 0.209017 | 0.8949398 |
| ENSG00000120742.10 | SERP1 | -0.023551 | 0.6214382 | 0.272657 | 0.8619483 | -1.239062 | 1.00E-30 | 0.899735 | 0.9583743 | 0.584144 | 0.621723 | 0.122786 | 0.8479431 |
| ENSG00000120800.4 | UTP20 | -0.383633 | 0.8913118 | -0.107156 | 0.6218841 | 0.223019 | 0.8312809 | 1.023367 | 0.03147009 | -0.005178 | 0.597842 | -0.516707 | 0.8787675 |
| ENSG00000120860.10 | CCDC53 | -0.802403 | 0.6214382 | -0.042147 | 0.8784109 | -0.13071 | 0.3276184 | 0.170631 | 0.01455697 | -0.196226 | 0.6382061 | -0.226507 | 0.6885065 |
| ENSG00000120885.19 | CLU | -0.594092 | 0.01740848 | 0.242992 | 1.00E-05 | -0.484414 | 1.17E-05 | -0.273045 | 5.02E-07 | -0.465843 | 2.77E-05 | 0.638116 | 0.6885065 |
| ENSG00000120896.13 | SORBS3 | -1.967132 | 0.2738943 | -1.341362 | 0.3142883 | -0.475109 | 0.009429051 | 1.11238 | 0.320553 | 0.730954 | 0.9316101 | -1.029896 | 0.8777121 |
| ENSG00000120963.11 | ZNF706 | 0.377033 | 0.6214382 | 0.336212 | 0.7881421 | -0.727331 | 0.003920665 | -0.543655 | 0.09450599 | 0.626825 | 0.3974318 | 0.599807 | 0.7969817 |
| ENSG00000121005.8 | CRISPLD1 | 0.538769 | 1 | -0.195071 | 0.3225827 | 2.576787 | 0.01531036 | -2.303174 | 0.4919368 | 1.809686 | 0.8387831 | 0.135407 | 0.744192 |
| ENSG00000121057.12 | AKAP1 | 0.318178 | 0.8218433 | 0.527072 | 0.6433679 | 0.772789 | 0.3488638 | 1.00339 | 0.009798458 | 1.000159 | 0.5617152 | -5.736494 | 0.5344421 |
| ENSG00000121060.14 | TRIM25 | -2.15878 | 0.3946735 | -0.263227 | 0.7057076 | -0.477288 | 0.1811887 | 2.108405 | 0.000347541 | -0.198611 | 0.9245352 | 0.495814 | 0.6829816 |
| ENSG00000121274.12 | PAPD5 | 0.634077 | 0.8006286 | -0.341758 | 0.8263112 | 0.115404 | 0.2646872 | 2.880049 | 0.001317823 | 0.972835 | 0.4387431 | -0.596847 | 0.6185816 |
| ENSG00000121390.18 | PSPC1 | -0.029161 | 0.9805839 | -0.016241 | 0.9686021 | -0.777562 | 0.0398183 | -0.135079 | 0.9675586 | 1.185725 | 0.8606292 | 0.649251 | 0.8634793 |
| ENSG00000121406.8 | ZNF549 | -0.281583 | 0.7736255 | 0.965601 | 0.6200225 | -0.63337 | 0.6260663 | 0.907763 | 0.2168885 | 1.053453 | 0.01318296 | -0.176709 | 0.9270338 |
| ENSG00000121579.12 | NAA50 | 0.420945 | 0.563505 | 0.340283 | 0.9102749 | 0.218716 | 0.4358165 | 1.131536 | 0.001392502 | -0.000166 | 0.4338963 | -1.137919 | 0.6885065 |
| ENSG00000121644.18 | DESI2 | -0.522054 | 0.9678883 | -0.093352 | 0.7596487 | 0.510935 | 0.9377685 | 1.956733 | 0.03362921 | 2.651327 | 0.4213485 | 2.127711 | 0.7767294 |
| ENSG00000121653.11 | MAPK8IP1 | -0.728865 | 0.6214382 | 0.186409 | 0.7383998 | -0.610542 | 0.9936722 | 0.604683 | 0.01974122 | 1.098839 | 0.1736796 | -0.089234 | 0.9147227 |
| ENSG00000121680.15 | PEX16 | -1.475864 | 0.529676 | 1.328903 | 0.4882009 | -0.720806 | 0.02566289 | -0.142161 | 0.6702963 | -1.263164 | 0.6447644 | 1.070649 | 0.7486771 |
| ENSG00000121741.16 | ZMYM2 | 0.248558 | 0.7648042 | 0.855802 | 0.8654277 | 1.243006 | 0.000869507 | 1.572052 | 0.0042943 | 1.669573 | 0.2884546 | 1.201156 | 1 |
| ENSG00000121749.15 | TBC1D15 | 0.574199 | 1 | -0.001094 | 0.8465126 | 0.162573 | 0.03961657 | 0.662718 | 0.6808574 | -1.168494 | 0.1998978 | -1.527335 | 0.7501684 |
| ENSG00000121753.12 | ADGRB2 | 2.860473 | 0.563505 | 0.252481 | 0.6527736 | -0.432409 | 0.1489368 | 1.467922 | 0.1208655 | 1.67882 | 0.001939793 | 0.850368 | 0.8167193 |
| ENSG00000121766.14 | ZCCHC17 | -0.983121 | 0.6214382 | 0.378809 | 0.8485366 | 0.491118 | 0.5290845 | 0.330473 | 0.3467508 | -0.344394 | 0.03854791 | -0.285344 | 0.9147227 |
| ENSG00000121774.17 | KHDRBS1 | -1.558721 | 0.000761938 | 0.441988 | 0.7944781 | 0.232825 | 0.3807776 | 0.02779 | 0.2047258 | 0.099138 | 0.6930133 | -0.052567 | 0.5733051 |
| ENSG00000121864.9 | ZNF639 | 0.049157 | 0.6214382 | -0.101573 | 0.9525045 | 0.080939 | 0.5480968 | 1.341025 | 2.12E-05 | 2.551703 | 0.06629523 | -0.855753 | 0.886525 |
| ENSG00000121879.3 | PIK3CA | -2.978453 | 0.3946735 | -0.579131 | 0.6305654 | 0.827168 | 0.003148427 | 1.850917 | 0.2099685 | 0.760646 | 0.8350514 | -2.394483 | 0.9130454 |
| ENSG00000121892.14 | PDS5A | -0.647512 | 0.7762699 | -0.419479 | 0.4727414 | 1.1175 | 0.02799093 | 2.156377 | 1.42E-05 | 1.065285 | 0.2973323 | -0.118279 | 0.6885065 |
| ENSG00000121904.17 | CSMD2 | 0.452669 | 0.9183025 | -0.972571 | 0.5953732 | 1.534442 | 0.01791645 | 1.148905 | 0.000165864 | 2.794477 | 0.007470624 | -1.425895 | 0.8479431 |
| ENSG00000121931.15 | LRIF1 | 0.19799 | 0.6214382 | 0.796788 | 0.5953732 | 0.338983 | 0.04392145 | 0.686146 | 0.4919368 | 0.237964 | 0.5196843 | -1.206074 | 0.5733051 |
| ENSG00000121964.14 | GTDC1 | -0.135959 | 0.7736255 | 0.362807 | 1 | 1.158564 | 0.8610253 | 1.502768 | 0.08788191 | 1.493657 | 0.021599 | 2.49661 | 0.8255112 |
| ENSG00000121989.14 | ACVR2A | -1.251661 | 0.6214382 | -0.822017 | 0.109154 | -0.273053 | 0.6505701 | 1.248992 | 0.008133114 | 1.46322 | 0.3516806 | 0.590507 | 0.9699068 |
| ENSG00000122026.10 | RPL21 | -2.731087 | 2.90E-47 | -0.145701 | 0.9994369 | 3.500549 | 0 | -0.217261 | 2.33E-05 | -0.116056 | 0.001158874 | -0.067227 | 0.2276299 |
| ENSG00000122033.14 | MTIF3 | -0.207969 | 0.934454 | 0.298888 | 0.6124714 | 0.409928 | 0.589407 | 0.189349 | 0.7179845 | -0.786885 | 0.004324004 | -0.80175 | 0.6829816 |
| ENSG00000122034.12 | GTF3A | 0.179176 | 0.8648814 | -0.062103 | 0.6679548 | -0.784701 | 0.000154657 | 0.487561 | 0.7796834 | 1.007695 | 0.5196843 | 1.005574 | 0.6885065 |
| ENSG00000122042.9 | UBL3 | -0.03577 | 0.7405237 | -0.138569 | 0.6129073 | 0.958901 | 0.03174585 | -0.281533 | 0.4919368 | 2.137356 | 0.1278327 | 0.048937 | 1 |
| ENSG00000122068.12 | FYTTD1 | 0.159201 | 0.8711787 | 0.290727 | 0.9256141 | 0.664719 | 0.006271249 | 1.248412 | 0.4284626 | -0.482061 | 0.5387167 | -0.884998 | 0.8879291 |
| ENSG00000122218.14 | COPA | 0.507129 | 0.9785048 | 0.299037 | 0.5671096 | 0.636048 | 0.8454005 | 1.260069 | 2.56E-07 | 1.377214 | 0.007826347 | 0.953179 | 0.7767294 |
| ENSG00000122257.18 | RBBP6 | -0.197281 | 0.9462203 | 0.491716 | 0.7449708 | 0.7988 | 0.00141721 | 1.050106 | 5.72E-06 | 0.835161 | 0.2090202 | -2.920668 | 0.6829816 |
| ENSG00000122299.11 | ZC3H7A | -0.183153 | 0.6214382 | 0.400619 | 0.8653838 | 1.239705 | 0.09429007 | 0.256342 | 0.002084874 | 1.082473 | 0.506562 | -0.069277 | 0.6885065 |
| ENSG00000122386.10 | ZNF205 | -5.026302 | 0.04178429 | -0.425414 | 0.6527736 | -0.009078 | 0.03757911 | 2.328199 | 4.41E-07 | 0.345253 | 0.3371898 | -1.809852 | 0.6185816 |
| ENSG00000122406.12 | RPL5 | -0.583275 | 0.6214382 | 0.152833 | 0.8504478 | 0.298221 | 0.001421368 | -0.050781 | 0.1707261 | -0.095526 | 0.000364991 | -0.055727 | 0.8479431 |
| ENSG00000122435.9 | TRMT13 | -1.133953 | 0.8006286 | -0.297006 | 0.5953732 | 1.298749 | 0.01465168 | 0.215062 | 0.8859784 | 0.950074 | 1 | -0.86905 | 0.8056071 |
| ENSG00000122482.20 | ZNF644 | 0.528071 | 0.6214382 | 0.231039 | 0.8837097 | 0.958679 | 6.28E-05 | 0.069314 | 0.7525043 | 0.389594 | 0.8936373 | 1.413575 | 0.8479431 |
| ENSG00000122515.14 | ZMIZ2 | 1.95245 | 0.714717 | 0.272249 | 0.7398038 | 0.061511 | 0.8322087 | 1.338237 | 0.004962482 | 1.702832 | 4.45E-05 | 0.445909 | 0.9083382 |
| ENSG00000122545.18 | 7-Sep | -0.308622 | 0.6214382 | 0.008534 | 0.7905529 | 0.808159 | 2.28E-16 | -0.068551 | 0.04364143 | -0.212129 | 0.000150248 | -0.253401 | 0.6885065 |
| ENSG00000122565.18 | CBX3 | 0.088949 | 0.07604642 | 0.071092 | 0.9520488 | 0.432455 | 0.000714677 | 0.098762 | 0.6458762 | -0.145717 | 0.03685235 | -0.688329 | 0.0620743 |
| ENSG00000122566.20 | HNRNPA2B1 | -2.033593 | 3.81E-08 | -0.090322 | 0.6126945 | -0.211019 | 1 | 0.748079 | 3.41E-09 | 1.603421 | 1.43E-38 | -1.072648 | 3.68E-06 |
| ENSG00000122584.12 | NXPH1 | -1.108921 | 0.714717 | -1.068566 | 0.01398133 | 1.085455 | 0.5681154 | 0.270059 | 0.9092206 | 1.611634 | 1 | 0.225663 | 0.863256 |
| ENSG00000122591.11 | FAM126A | -1.260336 | 0.529676 | -0.443321 | 0.02049533 | 0.276898 | 0.2011368 | 0.528947 | 0.5249905 | 0.510993 | 0.966632 | 0.238427 | 0.9077431 |
| ENSG00000122644.12 | ARL4A | 0.089697 | 1 | -0.14655 | 0.9102749 | 0.85161 | 0.000319989 | -0.121505 | 0.8549654 | 0.429528 | 0.9583141 | -0.372415 | 0.9147227 |
| ENSG00000122705.16 | CLTA | -0.514353 | 0.6014343 | 0.131102 | 0.9671539 | -0.716593 | 2.10E-09 | -0.193768 | 0.006900343 | 0.172981 | 0.01343337 | 0.387796 | 0.6829816 |
| ENSG00000122733.12 | PHF24 | -1.893696 | 0.9047689 | 0.757035 | 0.6527736 | 0.106112 | 1 | -1.114114 | 0.6468205 | 4.39557 | 0.04279185 | -1.650931 | 1 |
| ENSG00000122756.14 | CNTFR | -1.57552 | 0.6214382 | 0.713479 | 0.6527736 | -0.602343 | 0.02875454 | 0.658689 | 0.3208782 | 1.869544 | 0.000152282 | -0.5313 | 0.6885065 |
| ENSG00000122778.9 | KIAA1549 | 0.622688 | 0.8867323 | -1.840952 | 0.7384164 | 0.078981 | 0.1636297 | 1.059426 | 0.00029798 | 0.463311 | 0.4490861 | -0.346078 | 0.6885065 |
| ENSG00000122779.16 | TRIM24 | -0.336645 | 0.6562396 | -0.718031 | 0.008951202 | 0.286655 | 1.77E-05 | 1.03535 | 0.00266341 | 1.055449 | 0.8865876 | -0.621008 | 0.5733051 |
| ENSG00000122873.11 | CISD1 | 0.449189 | 0.529676 | -0.182619 | 0.1814896 | 0.005043 | 0.7437308 | 0.265236 | 0.5448609 | -1.116919 | 0.01491143 | -0.832872 | 0.4578379 |
| ENSG00000122966.13 | CIT | 0.692094 | 0.6559753 | 0.136449 | 0.8670464 | 1.591024 | 0.5290845 | 1.654668 | 0.006958854 | 3.294662 | 0.000278079 | -2.386709 | 0.815904 |
| ENSG00000123066.7 | MED13L | -2.067968 | 0.2531125 | -0.751652 | 0.3530125 | 1.147781 | 0.000155505 | 1.5375 | 2.33E-05 | 0.380002 | 0.913409 | 1.39002 | 1 |
| ENSG00000123091.4 | RNF11 | 0.205511 | 0.8006286 | -0.22453 | 0.09997869 | 0.016341 | 0.5832822 | 2.005934 | 0.03130918 | 1.106233 | 0.2364542 | -0.674198 | 0.6042097 |
| ENSG00000123094.15 | RASSF8 | 0.863534 | 0.3982827 | -0.490272 | 0.9081785 | 0.623564 | 0.6876227 | 0.933727 | 0.003680048 | 1.021682 | 0.9733105 | -1.669534 | 0.6829816 |
| ENSG00000123131.12 | PRDX4 | 0.482625 | 0.2351592 | 0.529297 | 0.03905952 | -0.248981 | 0.176831 | -0.134687 | 0.002136901 | 0.945069 | 0.8387831 | -1.044115 | 0.5653062 |
| ENSG00000123136.14 | DDX39A | 0.167301 | 0.9047689 | 0.385169 | 0.3610905 | 0.190762 | 0.9162151 | 0.529764 | 0.0743417 | -0.146416 | 0.03636076 | 0.379337 | 0.703227 |
| ENSG00000123143.12 | PKN1 | -0.68066 | 0.9473277 | 0.561118 | 0.4015099 | 0.319502 | 0.7523084 | 0.376006 | 0.04540425 | 0.377429 | 0.03064117 | -1.077973 | 1 |
| ENSG00000123144.10 | C19orf43 | -0.787216 | 0.001420982 | 0.41767 | 0.4914864 | 0.206423 | 0.4011225 | 0.234871 | 1 | 0.604195 | 0.9317563 | 0.046547 | 0.8479431 |
| ENSG00000123159.15 | GIPC1 | -0.515836 | 0.7833919 | 0.12793 | 0.6364102 | -0.939514 | 0.002893001 | 0.600479 | 1 | 0.386585 | 0.6950751 | 0.2588 | 0.9352001 |
| ENSG00000123178.14 | SPRYD7 | -1.21907 | 0.866076 | -0.25834 | 0.9449715 | -1.044061 | 0.003520218 | 0.740391 | 0.6296888 | 1.019296 | 0.7377262 | -2.355364 | 0.3406188 |
| ENSG00000123179.13 | EBPL | -0.950115 | 0.2600418 | 0.660319 | 0.7276361 | -2.599217 | 1.43E-06 | -0.241888 | 0.1453248 | 1.32049 | 0.3045911 | 0.823418 | 0.9011704 |
| ENSG00000123200.16 | ZC3H13 | -0.079978 | 0.934454 | 0.025218 | 0.1982022 | 0.878743 | 7.41E-08 | 1.637748 | 0.01826913 | -0.181086 | 0.08966305 | 0.679855 | 0.8868882 |
| ENSG00000123213.22 | NLN | -0.573406 | 0.8375411 | -0.320518 | 0.1014605 | -0.339656 | 0.1313518 | 1.148071 | 0.2391379 | 1.608815 | 0.01966133 | 0.926085 | 0.6885065 |
| ENSG00000123297.17 | TSFM | 0.529427 | 0.7749577 | 0.56408 | 0.3907323 | -0.290691 | 0.4064599 | 1.112316 | 0.1860003 | -0.05182 | 0.000229075 | 1.089171 | 0.9510409 |
| ENSG00000123349.13 | PFDN5 | -1.429011 | 0.000882101 | 0.144214 | 0.8866333 | -0.271764 | 0.000176781 | 0.281714 | 0.6996942 | -0.148251 | 0.1447676 | 0.221803 | 0.5177503 |
| ENSG00000123352.17 | SPATS2 | -0.331484 | 0.928247 | -0.171156 | 0.6527736 | 0.430217 | 0.003663876 | 0.654825 | 0.4252723 | 0.228303 | 0.4167039 | -0.977977 | 0.8479431 |
| ENSG00000123358.19 | NR4A1 | -1.138167 | 0.6214382 | 0.313141 | 0.6527736 | 0.375865 | 0.6717367 | 1.302497 | 0.14419 | 2.965307 | 0.003209786 | 1.381143 | 0.3728408 |
| ENSG00000123416.15 | TUBA1B | 0.200386 | 0.02212045 | -0.137416 | 0.6617347 | -0.051376 | 0.4064599 | -0.16074 | 0.02084521 | 0.172948 | 0.8044619 | -0.158871 | 0.539366 |
| ENSG00000123472.12 | ATPAF1 | 0.428069 | 0.8587992 | 0.324686 | 0.6527736 | 0.035772 | 0.4260085 | 0.546456 | 0.01873068 | 0.874021 | 0.9953752 | -0.11167 | 0.6829816 |
| ENSG00000123505.14 | AMD1 | -0.720453 | 0.3253495 | 0.108147 | 0.7881421 | 0.41975 | 0.001425147 | 0.522179 | 0.9369315 | -0.104703 | 0.4773416 | 0.245258 | 0.9147227 |
| ENSG00000123562.16 | MORF4L2 | 0.218628 | 0.6014343 | 0.333476 | 0.6527736 | 0.463259 | 0.006444507 | 0.144348 | 0.8978839 | 0.95502 | 0.305246 | 0.717726 | 1 |
| ENSG00000123607.14 | TTC21B | 2.457796 | 1 | -0.876468 | 0.8598627 | 0.906246 | 0.006780231 | 3.544793 | 0.1381463 | 1.952009 | 0.3974318 | 4.690107 | 0.278077 |
| ENSG00000123636.17 | BAZ2B | 0.199609 | 0.9663927 | 0.074687 | 0.8749658 | 0.123071 | 0.09417047 | 1.495104 | 0.01841683 | 1.632435 | 0.09488473 | 0.198374 | 0.9147227 |
| ENSG00000123684.12 | LPGAT1 | -0.330666 | 0.8006286 | 0.247988 | 0.7917816 | -0.272381 | 0.2879524 | 1.06838 | 0.04643737 | 1.286049 | 0.08810529 | 0.58053 | 0.7767294 |
| ENSG00000123810.7 | B9D2 | -1.478033 | 1 | 0.636552 | 0.67548 | -1.708612 | 0.009865723 | 0.784332 | 0.6716329 | 0.676589 | 0.5668961 | -3.201368 | 0.2768456 |
| ENSG00000123908.11 | AGO2 | -1.23243 | 0.6214382 | 0.298295 | 0.9797314 | 1.227243 | 0.000184109 | 0.249928 | 0.005400628 | 1.100759 | 0.2581597 | 0.076955 | 1 |
| ENSG00000123933.14 | MXD4 | -0.158149 | 1 | 0.172669 | 0.979665 | 0.399119 | 0.6462166 | 0.768204 | 0.05469418 | 2.269402 | 0.0130078 | -2.144206 | 0.7136557 |
| ENSG00000123983.13 | ACSL3 | -0.548856 | 0.6792949 | 0.301017 | 0.7869577 | 0.712028 | 0.06624586 | 0.575683 | 0.1385281 | 2.818915 | 0.03056663 | -0.348222 | 0.7863554 |
| ENSG00000123989.13 | CHPF | -0.095859 | 0.8006286 | 0.110601 | 0.455084 | -0.705853 | 5.45E-07 | 0.288629 | 0.4919368 | -0.152644 | 0.7395277 | 0.58857 | 0.6885065 |
| ENSG00000124006.14 | OBSL1 | -0.606107 | 0.7820597 | 0.511015 | 0.5676172 | -0.974227 | 0.7437308 | 1.190684 | 0.1504201 | 2.072798 | 0.006569714 | 0.30681 | 1 |
| ENSG00000124067.16 | SLC12A4 | -1.938501 | 0.6214382 | -0.348645 | 0.6274147 | 1.074947 | 0.8656249 | 1.557849 | 0.02882544 | -0.589891 | 0.7377262 | -0.275438 | 0.8207814 |
| ENSG00000124098.9 | FAM210B | -0.619618 | 0.8006286 | -0.613914 | 0.2570642 | 0.818397 | 0.9733888 | 2.589368 | 0.000326842 | 0.866705 | 0.3993666 | 0.982666 | 0.8260537 |
| ENSG00000124120.10 | TTPAL | 0.319753 | 0.6214382 | 0.755712 | 0.7342083 | 0.180629 | 0.7613083 | -0.020515 | 0.002832117 | -0.729298 | 0.8387831 | 0.183891 | 1 |
| ENSG00000124151.18 | NCOA3 | -0.649398 | 1 | -1.074864 | 0.688619 | -0.228124 | 0.1275383 | 3.870137 | 0.000302031 | 1.844792 | 0.2951202 | 0.528282 | 0.958512 |
| ENSG00000124155.16 | PIGT | -0.116859 | 0.9195601 | -0.507525 | 0.6527736 | -0.51583 | 2.13E-06 | 0.598393 | 0.002513517 | 0.415441 | 0.4631212 | -0.081177 | 0.6200022 |
| ENSG00000124160.11 | NCOA5 | 0.91806 | 0.9384913 | -0.130493 | 0.9314977 | -0.050362 | 0.5290845 | 2.353047 | 0.005177972 | 0.248839 | 0.5196843 | 1.953053 | 0.9147227 |
| ENSG00000124164.15 | VAPB | 1.132602 | 0.6692173 | 0.065663 | 0.3028222 | -0.645268 | 0.001005672 | 0.27796 | 0.5930297 | 0.312051 | 0.5743138 | -1.908082 | 0.6829816 |
| ENSG00000124172.9 | ATP5E | 0.605639 | 1.16E-10 | -0.361741 | 1.18E-05 | -0.455175 | 1.05E-15 | 0.42524 | 5.39E-08 | 0.223151 | 0.6854585 | -0.242809 | 0.008724428 |
| ENSG00000124177.14 | CHD6 | 0.092808 | 0.8006286 | -0.357431 | 0.9583742 | 1.170456 | 0.00277844 | 0.473055 | 0.5099779 | 1.44048 | 0.05313626 | 0.438427 | 0.8194787 |
| ENSG00000124193.14 | SRSF6 | -0.627241 | 0.6836146 | 0.408447 | 0.8632241 | 0.595066 | 0.8130993 | 0.742593 | 0.000259537 | 0.962166 | 0.5196843 | -1.268309 | 0.6829816 |
| ENSG00000124207.16 | CSE1L | 1.007083 | 0.5815318 | -0.08387 | 0.7647276 | 0.551178 | 0.7380232 | 1.141055 | 0.02752294 | 1.287202 | 0.183138 | 0.387116 | 0.8479431 |
| ENSG00000124209.3 | RAB22A | -0.086035 | 0.3610267 | -0.555082 | 0.5117578 | 0.290817 | 0.6057253 | 1.332901 | 0.03345424 | 0.468201 | 0.2880317 | 1.784629 | 0.5137734 |
| ENSG00000124214.19 | STAU1 | 0.16225 | 0.563505 | 0.016369 | 0.6200225 | 0.321182 | 0.000455376 | -0.311626 | 0.4919368 | 0.196546 | 0.2019818 | -1.481986 | 1 |
| ENSG00000124217.4 | MOCS3 | 0.264698 | 1 | 0.879283 | 0.8903384 | 0.666684 | 0.8862589 | 0.0356 | 0.01052272 | 1.037966 | 0.119277 | -1.27361 | 0.9565287 |
| ENSG00000124222.21 | STX16 | -1.032383 | 0.6214382 | -0.691012 | 0.4011918 | 0.213918 | 0.138687 | 1.70639 | 0.001082576 | 0.483681 | 0.1399344 | -1.613259 | 0.8318522 |
| ENSG00000124225.15 | PMEPA1 | -1.591927 | 0.06126222 | -0.197538 | 0.6527736 | -0.819395 | 0.1821132 | 1.184421 | 5.38E-10 | 1.129694 | 0.2232935 | 0.197597 | 0.6885065 |
| ENSG00000124383.8 | MPHOSPH10 | -1.115489 | 0.1367467 | 0.027759 | 0.9335892 | 0.628252 | 0.01010057 | 0.298122 | 0.2559244 | 0.318593 | 0.4218601 | -1.305971 | 0.6885065 |
| ENSG00000124422.11 | USP22 | 0.571911 | 0.4406957 | -0.531754 | 0.4571222 | -0.489948 | 0.2707921 | 1.691512 | 1.55E-06 | 1.149426 | 0.000459067 | -1.865681 | 0.05841333 |
| ENSG00000124486.12 | USP9X | -0.212248 | 0.8212257 | -0.851639 | 0.1378664 | 0.865107 | 0.001137674 | 1.098372 | 0.07585453 | 1.304777 | 0.02611546 | 0.976337 | 1 |
| ENSG00000124496.12 | TRERF1 | 0.45565 | 0.6214382 | 1.919158 | 0.7362582 | 0.707611 | 0.8547576 | 4.424519 | 0.04310756 | 3.300408 | 0.08266705 | 0.791848 | 0.6829816 |
| ENSG00000124535.15 | WRNIP1 | 1.029016 | 0.8006286 | -0.489104 | 0.5785445 | 0.046102 | 0.9822579 | -0.130272 | 0.1971461 | 2.201805 | 0.001867602 | -0.441121 | 1 |
| ENSG00000124562.9 | SNRPC | -0.970365 | 0.563505 | -0.107333 | 0.3736565 | 0.025878 | 0.3314981 | 0.52498 | 0.2838987 | -0.046506 | 6.72E-07 | -0.456342 | 0.4304069 |
| ENSG00000124588.19 | NQO2 | -2.994831 | 0.563505 | 0.738584 | 0.6527736 | -0.558603 | 0.000212496 | 1.260611 | 0.9414643 | -0.517406 | 0.7579722 | 1.999691 | 0.6963763 |
| ENSG00000124596.16 | OARD1 | 0.707105 | 0.7736255 | -0.226008 | 0.6527736 | 0.213134 | 0.8100781 | 0.924443 | 0.01960715 | 0.399586 | 0.891217 | -0.164614 | 0.9174022 |
| ENSG00000124614.13 | RPS10 | -1.168669 | 4.63E-09 | 0.549156 | 0.003152636 | 0.189162 | 0.8042144 | 0.328964 | 0.3525754 | -0.022862 | 0.001063327 | 0.094618 | 0.3561917 |
| ENSG00000124641.14 | MED20 | -1.671641 | 0.9047689 | 0.245534 | 0.8953649 | 1.009603 | 0.5312449 | 1.760611 | 0.001302451 | 0.719413 | 1 | -0.425079 | 0.9532204 |
| ENSG00000124733.3 | MEA1 | -0.62291 | 0.4406957 | 0.263834 | 0.8321919 | -0.423799 | 1.07E-05 | 0.164752 | 1 | 0.25821 | 0.6668636 | -0.14153 | 0.8479431 |
| ENSG00000124766.5 | SOX4 | -1.087704 | 0.8006902 | -0.472242 | 0.2895653 | -0.143378 | 0.5290845 | 1.746564 | 6.57E-09 | 0.999993 | 3.02E-07 | -0.776316 | 0.4149735 |
| ENSG00000124772.11 | CPNE5 | -1.049904 | 0.6214382 | -2.087406 | 0.2174163 | 0.77323 | 0.7236298 | -0.167945 | 0.4919368 | 4.314142 | 0.04279185 | 0.01713 | 1 |
| ENSG00000124783.12 | SSR1 | 0.447309 | 0.563505 | 0.026843 | 0.8208804 | 0.050544 | 0.04750321 | 1.199933 | 0.04148071 | 0.416275 | 1 | 0.249731 | 0.868568 |
| ENSG00000124784.8 | RIOK1 | -0.452971 | 1 | -0.298322 | 0.6527736 | 0.508403 | 0.02388094 | 0.021155 | 0.9732039 | -0.654025 | 0.04652304 | 0.350223 | 0.8639532 |
| ENSG00000124788.17 | ATXN1 | -0.447918 | 0.529676 | -0.764596 | 0.03321267 | 1.293868 | 0.0173006 | 0.955787 | 0.03950552 | -0.237908 | 0.5520748 | 0.539184 | 0.6885065 |
| ENSG00000124789.11 | NUP153 | 0.815378 | 0.9543887 | -0.118481 | 0.8710975 | 0.333739 | 0.03729056 | -0.549307 | 0.004807921 | 0.657643 | 0.8889533 | 0.167438 | 0.9786489 |
| ENSG00000124802.11 | EEF1E1 | -1.308472 | 0.00598845 | -0.057821 | 0.6628721 | 0.126068 | 0.5290845 | 0.228268 | 0.5378596 | -0.412301 | 0.2433356 | 0.09539 | 0.8016607 |
| ENSG00000125107.16 | CNOT1 | -0.112361 | 0.9542165 | 0.037263 | 0.8566182 | 0.669605 | 0.07847327 | 0.870209 | 0.02111672 | 0.230873 | 0.8389957 | -0.173929 | 0.9532204 |
| ENSG00000125144.13 | MT1G | 1.331189 | 0.4692975 | 0.927432 | 0.4905237 | -1.033815 | 0.001961848 | -1.176804 | 0.001510571 | 0.455861 | 0.3647106 | 1.72136 | 0.8637809 |
| ENSG00000125148.6 | MT2A | 0.375881 | 0.6185251 | -0.222078 | 0.7906186 | -0.293188 | 0.03794911 | -0.721226 | 3.80E-07 | 0.342074 | 0.09575711 | 0.644178 | 0.530077 |
| ENSG00000125166.12 | GOT2 | 2.298106 | 0.04178429 | -0.126691 | 0.95708 | 0.67228 | 0.3228353 | 0.945317 | 0.7875686 | 0.269615 | 0.8785838 | -3.22197 | 0.5733051 |
| ENSG00000125249.6 | RAP2A | -0.912851 | 0.563505 | 0.391983 | 0.592336 | 1.393432 | 0.000361019 | 1.566419 | 0.1179233 | 1.467453 | 0.521264 | 0.000972 | 1 |
| ENSG00000125351.10 | UPF3B | -0.651866 | 0.563505 | -0.04034 | 0.7811664 | 0.453448 | 0.3871766 | 0.441454 | 0.4055117 | -0.39897 | 0.000746459 | 1.626205 | 0.9425307 |
| ENSG00000125356.6 | NDUFA1 | 0.634255 | 0.001073927 | 0.044315 | 0.7955884 | -0.629689 | 3.10E-09 | -0.160356 | 0.000776972 | -0.276138 | 0.000106562 | 0.242251 | 0.9264343 |
| ENSG00000125386.14 | FAM193A | -1.191155 | 0.7766974 | -0.07577 | 0.5394427 | 0.574274 | 0.3714078 | 1.36338 | 0.006353792 | -1.193018 | 0.800077 | 1.609909 | 0.7955389 |
| ENSG00000125388.19 | GRK4 | -0.271578 | 1 | -0.667438 | 0.9203962 | 0.197002 | 0.8473149 | 0.707143 | 0.125732 | 3.045443 | 0.02673647 | 0.845685 | 0.6829816 |
| ENSG00000125398.5 | SOX9 | -0.755037 | 0.6214382 | -0.186528 | 0.4872949 | 0.6404 | 6.49E-05 | 0.515174 | 0.2115588 | 0.370551 | 0.325647 | 0.839989 | 0.6185816 |
| ENSG00000125445.10 | MRPS7 | -0.189017 | 0.9181195 | 0.25213 | 0.7112271 | -0.514264 | 0.000254115 | 0.065533 | 0.381621 | 0.256192 | 0.2880317 | -0.961721 | 0.5344421 |
| ENSG00000125447.16 | GGA3 | 1.194929 | 0.6214382 | -1.207968 | 0.2521066 | -0.238906 | 0.5875392 | 3.19876 | 0.01033382 | 0.96925 | 0.2053281 | -2.473518 | 0.8573709 |
| ENSG00000125458.6 | NT5C | 0.080127 | 0.6214382 | -0.094048 | 0.7608509 | -0.360171 | 1.44E-06 | -0.357553 | 0.09720862 | -0.202742 | 0.2578171 | 1.095978 | 0.6885065 |
| ENSG00000125459.14 | MSTO1 | -0.399591 | 0.8006286 | 0.591204 | 0.4403562 | -1.10206 | 0.001821162 | 1.566402 | 0.2287146 | -1.68897 | 0.4538479 | -2.408016 | 0.6885065 |
| ENSG00000125462.16 | C1orf61 | -1.258043 | 0.5468595 | -0.287966 | 0.5394427 | -0.389941 | 5.98E-06 | 0.098284 | 0.007660097 | -0.103831 | 0.01472525 | -0.423868 | 0.05234973 |
| ENSG00000125482.12 | TTF1 | 0.268151 | 0.714717 | 0.54735 | 0.7750859 | 1.066864 | 0.01345503 | 1.10151 | 0.6288398 | 1.080606 | 0.5608493 | 1.344046 | 0.9270789 |
| ENSG00000125484.11 | GTF3C4 | 0.621907 | 0.8218433 | -0.160317 | 0.7236203 | 0.751597 | 0.01152691 | 0.971987 | 0.1292751 | 0.621903 | 0.01871809 | -0.694615 | 1 |
| ENSG00000125485.17 | DDX31 | 0.185291 | 0.9367821 | 0.010107 | 0.8263112 | 0.656508 | 0.3201582 | 0.701957 | 0.005738205 | 1.895668 | 0.0212476 | -2.596762 | 0.5733051 |
| ENSG00000125503.12 | PPP1R12C | 0.600668 | 1 | 0.303955 | 0.8829419 | -0.636216 | 0.3332424 | 1.236771 | 0.002136901 | 1.247706 | 0.587307 | -0.519677 | 0.6885065 |
| ENSG00000125534.9 | PPDPF | -0.449984 | 0.1824223 | -0.320345 | 0.1772534 | -0.515239 | 8.32E-14 | 0.011865 | 0.7873881 | 0.396496 | 0.3974318 | -0.371413 | 0.2321124 |
| ENSG00000125629.14 | INSIG2 | -0.103035 | 0.9348915 | 0.561233 | 0.3961673 | 1.297341 | 0.01935509 | 0.684051 | 0.7227707 | 0.916171 | 0.7938524 | -0.268131 | 0.6829816 |
| ENSG00000125630.15 | POLR1B | -0.173422 | 1 | -0.221295 | 0.8217443 | 0.954482 | 0.2969884 | 3.404869 | 0.02595271 | 0.773392 | 0.4120285 | -1.244006 | 0.826754 |
| ENSG00000125633.10 | CCDC93 | -0.76539 | 0.6214382 | 0.339004 | 0.9622004 | 0.366159 | 0.6574966 | -0.205187 | 0.05545657 | 0.982785 | 0.02709242 | 0.440221 | 1 |
| ENSG00000125651.13 | GTF2F1 | -0.092897 | 0.6214382 | -0.079386 | 0.6286991 | 0.112822 | 0.0447881 | -0.484972 | 0.7858185 | 0.521304 | 0.8019395 | -0.003409 | 1 |
| ENSG00000125656.8 | CLPP | -0.592087 | 1 | -0.427152 | 0.6200225 | -0.343853 | 0.02646748 | -0.148792 | 0.1170114 | -0.419464 | 0.4684593 | -0.13068 | 0.6885065 |
| ENSG00000125675.17 | GRIA3 | 0.706286 | 0.8006286 | 0.157642 | 0.842947 | 1.090687 | 0.4064599 | 0.699615 | 0.006286247 | 0.644049 | 0.5196843 | -2.169832 | 0.9576568 |
| ENSG00000125676.19 | THOC2 | -0.678037 | 0.6214382 | 0.351919 | 0.9680711 | 0.720085 | 0.000299694 | 1.274571 | 0.05964933 | 0.018852 | 0.07066598 | 1.02209 | 0.8816797 |
| ENSG00000125734.13 | GPR108 | -0.823075 | 0.8256267 | 0.937335 | 0.5517607 | -0.489667 | 0.2042955 | -1.043763 | 0.01460268 | 1.079449 | 0.8390546 | 1.174644 | 1 |
| ENSG00000125740.13 | FOSB | -1.472526 | 0.714717 | 0.003207 | 0.7383998 | -0.614842 | 0.9143308 | 0.687147 | 0.07141399 | 5.716471 | 0.002918613 | 1.056143 | 0.5344421 |
| ENSG00000125743.10 | SNRPD2 | -0.350829 | 0.8716459 | 0.069099 | 0.7479736 | -0.333014 | 1.50E-08 | -0.077042 | 0.02806712 | 0.012641 | 0.00108844 | 0.262408 | 0.6108304 |
| ENSG00000125755.18 | SYMPK | -0.035597 | 0.7736255 | 0.68463 | 0.5953732 | -0.437922 | 0.0757302 | 0.278948 | 0.1586932 | 2.01517 | 0.03056663 | -1.075114 | 0.9335118 |
| ENSG00000125772.12 | GPCPD1 | -1.246828 | 0.6792949 | -0.285164 | 0.7427243 | 0.31247 | 1 | 1.115289 | 0.01034389 | 2.251219 | 0.1020081 | -0.65317 | 0.8949489 |
| ENSG00000125779.21 | PANK2 | -1.005228 | 0.6214382 | 0.199906 | 0.592336 | 0.237594 | 0.5290845 | 2.008606 | 0.002259593 | 1.21589 | 0.1477952 | 0.139464 | 0.9569765 |
| ENSG00000125817.7 | CENPB | 0.465204 | 0.6214382 | -0.263506 | 0.9561974 | -0.040726 | 1 | 0.255681 | 0.001467926 | 1.543364 | 0.06557358 | -0.099846 | 0.8599171 |
| ENSG00000125821.11 | DTD1 | -0.154936 | 0.6214382 | -0.098263 | 0.937266 | 0.011232 | 0.2456481 | 0.086294 | 0.346924 | 0.038911 | 0.006379689 | -0.014488 | 0.359138 |
| ENSG00000125835.17 | SNRPB | 0.224995 | 0.6014343 | -0.013695 | 0.9028136 | -0.205297 | 1.91E-05 | 0.878426 | 0.6575769 | 0.84594 | 0.3974318 | -0.227047 | 0.2281002 |
| ENSG00000125841.12 | NRSN2 | 0.857833 | 0.9601425 | -0.131514 | 0.7806598 | -0.256522 | 0.0479416 | 0.558501 | 0.1850748 | 0.629556 | 0.9958488 | -1.248245 | 0.7302103 |
| ENSG00000125868.15 | DSTN | -0.023573 | 0.9047689 | -0.078644 | 0.8200822 | 0.165508 | 0.5693693 | -0.068769 | 0.05914301 | -0.373061 | 3.81E-06 | 0.043092 | 0.5344421 |
| ENSG00000125875.13 | TBC1D20 | 1.541968 | 1 | -0.119535 | 0.9222921 | -1.00483 | 0.5442731 | 0.230425 | 0.05212385 | 2.104199 | 0.03064117 | 0.719493 | 0.5733051 |
| ENSG00000125877.12 | ITPA | -0.065257 | 0.9470141 | 0.398642 | 0.9337922 | -0.106113 | 0.005118679 | -0.251767 | 0.1474483 | -0.458561 | 0.001390045 | -0.146067 | 0.6525228 |
| ENSG00000125885.13 | MCM8 | 0.395672 | 1 | 1.295481 | 1 | -0.253202 | 0.7781174 | 2.631163 | 0.000156107 | 0.820339 | 0.5196843 | -2.193011 | 0.7728704 |
| ENSG00000125901.5 | MRPS26 | 0.242897 | 0.6214382 | 0.117299 | 0.4158789 | -0.812927 | 1.33E-13 | 0.531658 | 0.03370854 | 1.3097 | 0.5327129 | -0.582236 | 0.6749528 |
| ENSG00000125931.10 | CITED1 | 0.253626 | 0.4605085 | -0.707412 | 0.007219487 | -0.288076 | 0.2742937 | -0.324817 | 0.01035501 | -0.535888 | 0.5674589 | 0.149564 | 0.6885065 |
| ENSG00000125966.9 | MMP24 | 2.953135 | 0.6214382 | 2.921222 | 0.8263112 | 1.112066 | 0.7354314 | 0.906965 | 0.01223151 | 0.849272 | 0.07671111 | 1.851182 | 0.9147227 |
| ENSG00000125968.8 | ID1 | 0.150921 | 0.2299131 | -0.202893 | 0.3386108 | -2.154439 | 2.11E-28 | -0.546212 | 0.000153284 | 0.587408 | 0.6217996 | 0.415062 | 1 |
| ENSG00000125971.16 | DYNLRB1 | -0.324093 | 0.6214382 | -0.12775 | 0.8146081 | 0.0983 | 0.846701 | 0.083965 | 0.07611178 | -0.445798 | 0.01116125 | 0.909981 | 0.9104104 |
| ENSG00000125977.6 | EIF2S2 | -2.426614 | 0.6214382 | 0.19281 | 0.6527736 | 0.073038 | 0.700861 | 0.187982 | 0.22773 | -0.637234 | 5.82E-08 | -0.221245 | 0.5653062 |
| ENSG00000125991.18 | ERGIC3 | 0.828683 | 0.4631167 | 0.452417 | 0.6527736 | -0.050491 | 0.2879524 | 0.500374 | 0.1220768 | -0.852426 | 0.003061704 | -0.154137 | 0.3746505 |
| ENSG00000125995.15 | ROMO1 | 0.359139 | 0.4406957 | -0.00183 | 0.9564082 | -0.791675 | 1.16E-24 | -0.210467 | 0.002666913 | -0.237571 | 9.74E-05 | -0.257108 | 0.3542189 |
| ENSG00000126003.6 | PLAGL2 | -0.104732 | 1 | 0.104107 | 0.8263112 | 0.383189 | 0.7039521 | 3.780644 | 0.000414622 | 1.637834 | 0.2707324 | -0.326276 | 0.7833507 |
| ENSG00000126005.16 | MMP24-AS1 | -2.860159 | 0.6214382 | 0.06853 | 0.6597022 | -1.495578 | 0.002678974 | 0.297003 | 0.2335397 | 2.139348 | 0.5168571 | 1.435798 | 0.9461816 |
| ENSG00000126012.11 | KDM5C | -1.492142 | 0.6214382 | 0.322453 | 0.6200225 | 0.620339 | 0.761562 | 1.135232 | 0.01814921 | 0.514289 | 0.1806765 | -1.058521 | 0.7501243 |
| ENSG00000126067.11 | PSMB2 | 0.569667 | 0.1835496 | -0.317725 | 0.3923745 | -0.097638 | 0.05182658 | 0.180102 | 0.01888407 | 0.09204 | 0.2883445 | -0.386478 | 0.6829816 |
| ENSG00000126070.19 | AGO3 | -1.793026 | 0.3009675 | 0.976082 | 0.9720618 | 0.782769 | 1.52E-05 | 1.619919 | 5.39E-05 | 1.590983 | 0.003057286 | -2.69638 | 0.7963477 |
| ENSG00000126088.12 | UROD | 1.333073 | 0.4704049 | 0.527455 | 0.5445142 | -0.122055 | 0.4958973 | -0.398399 | 0.00542013 | -1.025273 | 0.05231538 | -0.647014 | 0.5653062 |
| ENSG00000126214.21 | KLC1 | -0.299202 | 0.6014343 | 0.05047 | 0.9696336 | 0.296671 | 0.5290845 | 0.615501 | 0.008116431 | 1.089398 | 0.01373222 | 0.407424 | 0.6885065 |
| ENSG00000126216.12 | TUBGCP3 | -3.566518 | 0.02220694 | -0.00606 | 1 | -0.326105 | 0.2983168 | 1.32416 | 0.2358939 | 0.349728 | 0.2019818 | -0.182472 | 0.6885065 |
| ENSG00000126217.20 | MCF2L | 2.438532 | 0.8006286 | -1.172658 | 0.1547783 | 0.731876 | 0.578568 | 2.440777 | 0.02457347 | 0.060782 | 0.2545482 | 0.693798 | 0.9532204 |
| ENSG00000126247.10 | CAPNS1 | -0.339864 | 0.8006286 | -0.232329 | 0.2296726 | -0.407022 | 0.01012156 | -0.257054 | 0.6821984 | -0.239754 | 0.3663286 | -0.044152 | 0.8171275 |
| ENSG00000126249.7 | PDCD2L | 0.269387 | 0.7840525 | 0.274973 | 0.7952779 | -1.05852 | 0.0445227 | 0.487778 | 0.5396604 | 1.159484 | 0.8608026 | 4.895208 | 0.6885065 |
| ENSG00000126254.11 | RBM42 | -0.511171 | 0.5711449 | 0.459301 | 0.1812803 | -0.635116 | 8.18E-11 | 0.397517 | 0.6628765 | 0.087644 | 0.1213481 | -0.513465 | 0.4304069 |
| ENSG00000126261.12 | UBA2 | -1.019465 | 0.9678883 | -0.09721 | 0.3803801 | 0.182558 | 0.4970176 | 1.020588 | 0.03242844 | 0.818325 | 0.2714646 | -1.43277 | 0.6829816 |
| ENSG00000126267.8 | COX6B1 | -0.326727 | 0.6014343 | -0.047808 | 0.6527736 | -0.204756 | 6.72E-08 | 0.311331 | 0.4109963 | -0.115641 | 0.000585203 | -0.060877 | 0.4498785 |
| ENSG00000126391.13 | FRMD8 | 0.664532 | 0.7721746 | 0.056912 | 0.8071936 | -1.74323 | 0.002110125 | 0.258813 | 0.6945351 | 0.217388 | 0.9510798 | 0.876221 | 0.8532529 |
| ENSG00000126432.13 | PRDX5 | -0.166518 | 0.6214382 | 0.039444 | 1 | -0.190467 | 0.04187837 | -0.529388 | 0.000323618 | -1.340719 | 2.61E-07 | 0.160954 | 0.6885065 |
| ENSG00000126457.21 | PRMT1 | -0.323605 | 0.8774628 | 0.503508 | 0.3610905 | -0.07885 | 0.000460475 | 0.423395 | 0.8224647 | 0.404249 | 0.5196843 | 0.262068 | 0.6829816 |
| ENSG00000126461.14 | SCAF1 | 1.581841 | 0.6214382 | -0.366057 | 0.9225117 | -0.1333 | 0.3393299 | 1.64281 | 0.2307812 | 2.502418 | 0.00129205 | -0.025063 | 0.7354932 |
| ENSG00000126464.13 | PRR12 | -1.903537 | 0.563505 | 0.698194 | 1 | 0.935311 | 0.7447977 | 1.237756 | 0.1862039 | 1.292626 | 0.009383589 | -2.85746 | 0.5733051 |
| ENSG00000126602.10 | TRAP1 | 0.048562 | 0.6991852 | -0.129078 | 0.4559872 | -0.070156 | 0.02654338 | -0.024372 | 0.5156115 | 0.420328 | 0.6314075 | -0.647075 | 0.3532606 |
| ENSG00000126653.15 | NSRP1 | -1.027943 | 0.08293861 | -0.245186 | 0.2945701 | 0.429397 | 0.009242744 | 0.793281 | 1 | 0.497618 | 0.01897976 | 0.554653 | 0.8278576 |
| ENSG00000126698.10 | DNAJC8 | -0.341699 | 0.9047689 | 0.144978 | 0.9502423 | 0.169462 | 0.5726085 | -0.635141 | 0.004987918 | -0.516063 | 4.12E-06 | 0.063577 | 0.6829816 |
| ENSG00000126705.13 | AHDC1 | -1.542329 | 0.8240785 | -0.401631 | 0.6527736 | -1.271933 | 0.5363837 | 0.984494 | 0.01043677 | 2.749894 | 0.000665363 | 0.603942 | 0.8479431 |
| ENSG00000126709.14 | IFI6 | -0.319453 | 0.9548281 | -0.459058 | 0.1054706 | -1.541174 | 2.77E-18 | 0.71812 | 0.2969997 | -0.048573 | 0.03056663 | 0.064308 | 0.7821477 |
| ENSG00000126746.17 | ZNF384 | -0.470981 | 1 | 0.147805 | 0.8468859 | 0.621869 | 0.1120987 | 1.109328 | 0.03335045 | 1.970462 | 0.1812232 | 0.038074 | 0.9147227 |
| ENSG00000126756.11 | UXT | 0.019912 | 0.9328506 | -0.053904 | 0.3855014 | -0.160725 | 0.001551112 | 0.465887 | 0.7144938 | -0.346709 | 0.04196275 | -0.79138 | 0.4304069 |
| ENSG00000126767.17 | ELK1 | -1.412521 | 0.6169802 | -0.330163 | 0.9821504 | 0.324042 | 0.7935863 | 0.953889 | 0.04996903 | 2.830615 | 0.008476363 | -1.090315 | 1 |
| ENSG00000126768.12 | TIMM17B | 0.083484 | 0.9356481 | 0.312801 | 0.4487482 | -0.398091 | 0.06092656 | -0.502914 | 0.03625938 | -1.033414 | 0.003224625 | 0.513942 | 0.9033538 |
| ENSG00000126773.12 | PCNXL4 | -0.373729 | 0.8587992 | -0.221065 | 0.8608512 | 0.79418 | 0.5290845 | 0.883973 | 0.001221598 | 0.786476 | 0.09247983 | -1.65188 | 0.8430553 |
| ENSG00000126777.17 | KTN1 | 0.129572 | 0.2656681 | 0.061093 | 0.6527736 | 0.079422 | 0.01110157 | 0.757935 | 0.2602146 | -0.129475 | 0.5196843 | -0.087278 | 0.7479903 |
| ENSG00000126804.13 | ZBTB1 | 0.094495 | 0.9183304 | -0.368615 | 0.8689748 | 1.107105 | 0.2953579 | -0.127582 | 0.9426777 | 2.149321 | 0.03757692 | 3.527686 | 0.6885065 |
| ENSG00000126814.6 | TRMT5 | -0.190014 | 0.8218433 | 0.091093 | 0.9624823 | -0.348348 | 0.01612867 | -0.376434 | 0.9069344 | 1.752553 | 0.6630768 | -0.169841 | 0.6885065 |
| ENSG00000126858.16 | RHOT1 | 1.023183 | 0.7035386 | 0.574772 | 1 | 0.567881 | 0.7230879 | 1.314946 | 0.1266405 | 2.096376 | 0.02795992 | 0.497628 | 0.9726757 |
| ENSG00000126861.4 | OMG | -5.809538 | 0.8867323 | 1.666856 | 1 | 0.840009 | 0.8832504 | 1.914583 | 0.3467508 | 0.577581 | 0.7377262 |  |  |
| ENSG00000126883.16 | NUP214 | 1.593786 | 1 | 0.243925 | 0.7079982 | 0.976629 | 0.03334989 | -0.503565 | 0.8764761 | 1.67999 | 0.01162002 | -1.588934 | 0.6885065 |
| ENSG00000126934.13 | MAP2K2 | -1.716212 | 0.08293861 | -0.068553 | 0.8263112 | -0.190555 | 0.8221252 | 1.372362 | 0.01492078 | 2.533466 | 0.000105236 | 0.370558 | 1 |
| ENSG00000127022.14 | CANX | 0.342316 | 0.6214382 | 0.216413 | 0.6131773 | 0.660599 | 3.70E-10 | 0.998124 | 2.01E-07 | 0.233424 | 0.2477719 | -0.037091 | 0.5733051 |
| ENSG00000127080.9 | IPPK | -0.839149 | 0.8006286 | 0.477901 | 0.5010139 | 0.250799 | 0.8767234 | 2.444625 | 0.004392971 | -0.131782 | 0.8323733 | 2.354715 | 0.826754 |
| ENSG00000127081.13 | ZNF484 | -0.935372 | 1 | -1.080333 | 0.8128893 | 1.619727 | 0.04727701 | 0.070604 | 0.8638749 | 0.544239 | 0.5488317 | -3.740699 | 0.8479431 |
| ENSG00000127124.13 | HIVEP3 | 0.360409 | 0.8006286 | 0.01836 | 0.3324257 | 0.437218 | 0.2611212 | 0.656423 | 3.19E-05 | -0.66808 | 0.2850314 | -0.153791 | 0.8479431 |
| ENSG00000127184.10 | COX7C | -0.26482 | 1 | -0.026794 | 0.6527736 | -0.43218 | 2.02E-13 | 0.177717 | 0.3830392 | -0.184827 | 4.36E-06 | -0.157828 | 0.1683022 |
| ENSG00000127325.18 | BEST3 | -0.482426 | 0.8052741 | -0.236192 | 0.6331318 | 1.70264 | 0.5290845 | 1.55724 | 0.02290567 | -1.105985 | 0.2880317 | 0.264789 | 0.9148375 |
| ENSG00000127328.21 | RAB3IP | -1.444134 | 0.000737572 | -0.112405 | 0.5394427 | 0.403932 | 0.4064599 | 0.431513 | 0.0202141 | 0.713229 | 0.6738424 | 0.238028 | 0.9294466 |
| ENSG00000127445.13 | PIN1 | 1.354774 | 0.5540319 | -0.069181 | 0.5244682 | -0.527232 | 2.81E-05 | 0.743738 | 0.9352041 | 0.295909 | 0.4064677 | -0.967303 | 0.2947018 |
| ENSG00000127481.14 | UBR4 | 2.413106 | 0.8006286 | -0.629467 | 0.4264625 | 1.096705 | 5.98E-05 | 1.337355 | 0.1740326 | 3.1859 | 0.06237965 | 3.134189 | 0.6990228 |
| ENSG00000127483.17 | HP1BP3 | -0.104466 | 0.9840028 | 0.098029 | 0.6527736 | 0.578344 | 0.00037947 | 0.622948 | 0.08491499 | 1.029953 | 0.1092614 | 0.082767 | 0.6829816 |
| ENSG00000127511.9 | SIN3B | -0.988262 | 0.6214382 | -0.178449 | 1 | -1.086877 | 0.000681624 | 1.563089 | 0.008159015 | 0.780179 | 0.8102157 | 0.258688 | 0.8888976 |
| ENSG00000127526.13 | SLC35E1 | -0.355646 | 0.9531264 | 0.710841 | 0.4358838 | -0.089165 | 0.896092 | 1.80432 | 0.002983903 | 1.020743 | 0.5464122 | 1.714208 | 0.7901525 |
| ENSG00000127540.11 | UQCR11 | 0.370154 | 0.01177121 | -0.26844 | 0.004201507 | -0.652369 | 5.44E-17 | -0.177935 | 0.001059345 | -0.360121 | 3.32E-06 | 0.059759 | 0.6490419 |
| ENSG00000127586.16 | CHTF18 | 1.195773 | 0.8587992 | -0.663876 | 0.5953732 | -1.536427 | 0.03328468 | 1.888279 | 0.00460644 | 1.825148 | 0.01532942 | -0.576413 | 0.6990228 |
| ENSG00000127603.23 | MACF1 | -0.653408 | 0.6214382 | -0.496052 | 0.0108578 | 0.76375 | 0.03056815 | 1.375104 | 2.32E-07 | 1.486871 | 0.2206396 | -0.093594 | 0.9657663 |
| ENSG00000127616.17 | SMARCA4 | -0.35175 | 0.6214382 | -0.099039 | 0.1395298 | -0.181004 | 0.6558625 | 0.436308 | 0.02239099 | 0.798844 | 0.2601554 | -0.321592 | 0.6885065 |
| ENSG00000127663.14 | KDM4B | -0.328872 | 0.9562681 | 0.072289 | 0.6527736 | -0.985291 | 0.4540623 | 1.025986 | 0.001747724 | 2.011932 | 3.28E-05 | -0.153383 | 0.8479431 |
| ENSG00000127774.6 | EMC6 | -0.116903 | 0.7247082 | 0.353052 | 0.543247 | -0.476998 | 0.000111601 | -0.244712 | 0.05000235 | -0.110361 | 0.01238434 | 0.149464 | 0.9147227 |
| ENSG00000127804.12 | METTL16 | 0.101192 | 0.9847089 | 0.066248 | 0.9192533 | 0.011027 | 0.03524809 | 1.446954 | 0.5668539 | -0.080169 | 0.6732466 | -3.601136 | 0.2850719 |
| ENSG00000127837.9 | AAMP | -3.591607 | 0.8006286 | 0.404216 | 0.3947664 | -0.469709 | 0.009690441 | 1.023632 | 0.655488 | 0.638189 | 0.3974318 | 0.675245 | 0.6885065 |
| ENSG00000127838.13 | PNKD | 0.860364 | 0.6214382 | -0.234827 | 1 | 0.049749 | 0.02474132 | -0.044537 | 0.4919368 | 0.046544 | 0.2825999 | -0.518595 | 0.8818722 |
| ENSG00000127884.4 | ECHS1 | 0.516895 | 0.1239011 | 0.703022 | 0.2538827 | -0.020169 | 0.2376942 | 1.02717 | 0.000184862 | 0.785338 | 0.5079513 | -0.291971 | 0.3560513 |
| ENSG00000127914.16 | AKAP9 | 1.791374 | 0.2515428 | -0.3093 | 0.5561493 | 1.326644 | 9.77E-12 | 0.764466 | 7.52E-05 | 0.960298 | 0.8387831 | -0.834962 | 0.6885065 |
| ENSG00000127922.9 | SHFM1 | 0.392297 | 0.4218113 | 0.171829 | 0.8263112 | -0.43539 | 3.19E-12 | -0.099379 | 0.0463235 | -0.828142 | 5.33E-10 | -0.565043 | 0.02886423 |
| ENSG00000127946.16 | HIP1 | 0.494899 | 0.9870834 | -0.553864 | 0.1722198 | 0.532171 | 0.01192579 | 1.429765 | 0.001051713 | 1.428867 | 0.03792279 | 0.745383 | 0.7465357 |
| ENSG00000127952.16 | STYXL1 | -3.128231 | 0.4692975 | 0.294282 | 0.9547269 | -0.409957 | 0.03404331 | 0.099143 | 0.5184642 | 1.638474 | 0.9372936 | 0.962067 | 0.8375052 |
| ENSG00000127995.16 | CASD1 | -1.609234 | 0.4234646 | -0.316785 | 0.6861706 | 1.355995 | 0.1131161 | 2.588737 | 0.01811201 | 1.146566 | 0.294428 | -1.251506 | 0.6885065 |
| ENSG00000128016.5 | ZFP36 | -2.084815 | 0.8006286 | -0.047054 | 0.8500718 | -0.989285 | 0.001952285 | 0.481629 | 0.8114104 | -0.052468 | 0.7076897 | 0.59485 | 0.5553892 |
| ENSG00000128039.10 | SRD5A3 | -0.652207 | 1 | -1.901396 | 0.3126509 | -3.920146 | 0.0259659 | -0.099129 | 0.3912177 | 2.481515 | 0.5196843 | 2.40875 | 0.7915993 |
| ENSG00000128050.8 | PAICS | 0.677924 | 0.6171259 | -0.93755 | 9.61E-06 | 0.887909 | 0.09046443 | 0.180848 | 0.7022911 | 0.189853 | 0.4927467 | -1.062728 | 0.5344421 |
| ENSG00000128059.8 | PPAT | 0.293685 | 0.9666561 | -0.124288 | 0.9931449 | 1.149237 | 0.009319514 | 1.874611 | 0.09856702 | 0.325719 | 0.8881192 | 0.124173 | 0.6885065 |
| ENSG00000128159.11 | TUBGCP6 | -2.041687 | 0.6214382 | -0.965088 | 0.8528295 | 0.976363 | 1 | 3.693872 | 0.08446449 | 2.227625 | 0.0269531 | 3.128436 | 0.7501684 |
| ENSG00000128191.13 | DGCR8 | -1.010329 | 0.6214382 | -0.689491 | 0.5953732 | -0.167103 | 0.7214653 | 3.108987 | 2.62E-05 | 1.815098 | 0.2024552 | -1.998444 | 0.5653062 |
| ENSG00000128228.4 | SDF2L1 | 0.198886 | 0.9282145 | 0.708208 | 0.1950725 | -1.487428 | 7.99E-14 | -0.29028 | 0.06702551 | 0.294526 | 0.621723 | -0.214409 | 0.6885065 |
| ENSG00000128242.12 | GAL3ST1 | -0.069082 | 0.6214382 | 2.172842 | 0.5953732 | 0.116293 | 0.8100781 | 0.381891 | 0.8472142 | -1.052271 | 0.5196843 |  |  |
| ENSG00000128245.14 | YWHAH | -0.02992 | 0.8587992 | 0.106214 | 0.6527736 | 0.392528 | 1 | 0.99099 | 6.12E-05 | 1.360409 | 0.005074025 | 0.282846 | 0.4981184 |
| ENSG00000128266.8 | GNAZ | -0.679071 | 0.3946735 | -0.53378 | 0.635001 | 0.077927 | 0.9539224 | 0.674188 | 0.000759227 | 0.792309 | 0.5614302 | -0.969686 | 0.9174022 |
| ENSG00000128272.14 | ATF4 | -1.08259 | 0.007594835 | 0.147383 | 0.7417582 | -0.014459 | 0.8698024 | 0.595261 | 2.06E-05 | 0.859811 | 4.17E-05 | 0.365148 | 0.9971132 |
| ENSG00000128283.6 | CDC42EP1 | -0.663893 | 0.714717 | -0.143093 | 0.7276361 | -1.091212 | 0.000126412 | 0.056465 | 0.01391062 | 1.277424 | 0.65378 | 2.941179 | 0.7733704 |
| ENSG00000128309.16 | MPST | -0.621713 | 0.9522143 | -0.837687 | 0.2758186 | -0.442272 | 0.04887835 | -0.472359 | 0.03820042 | -0.010904 | 0.9451467 | -1.039677 | 0.8949489 |
| ENSG00000128335.13 | APOL2 | -1.43752 | 0.939666 | 0.001464 | 0.8789521 | -0.631324 | 0.003472498 | 0.600632 | 1 | 1.295707 | 0.9983594 | -1.006363 | 0.797124 |
| ENSG00000128463.12 | EMC4 | 0.172928 | 0.8006286 | 0.247553 | 0.3996669 | -0.006466 | 0.055326 | -0.337656 | 4.85E-06 | -0.161541 | 0.0177734 | 0.474929 | 0.7081915 |
| ENSG00000128482.15 | RNF112 | -0.657257 | 0.6214382 | -0.39966 | 0.5953732 | 0.359431 | 0.2814985 | 0.238437 | 0.7872531 | 2.27686 | 0.04281867 | -0.210999 | 0.8947452 |
| ENSG00000128487.16 | SPECC1 | -0.604124 | 0.9294053 | -0.308767 | 0.1011415 | 0.77174 | 9.91E-05 | 1.655708 | 0.01778999 | 0.956029 | 0.2357785 | -2.154801 | 0.9062221 |
| ENSG00000128512.19 | DOCK4 | 0.419211 | 0.6214382 | 0.065764 | 0.704767 | 0.124239 | 0.8100781 | 1.824248 | 0.000687068 | 0.702501 | 0.3698471 | 2.645777 | 0.6885065 |
| ENSG00000128524.4 | ATP6V1F | 0.482283 | 0.6815849 | -0.315909 | 0.6527736 | -0.595755 | 4.94E-10 | -0.405191 | 0.000795168 | 0.112394 | 0.2880317 | -0.004942 | 0.4498785 |
| ENSG00000128564.6 | VGF | 0.294073 | 0.000294684 | -0.743853 | 0.000916482 | -0.887493 | 5.92E-15 | 0.700749 | 7.65E-12 | 0.945024 | 0.3974318 | -0.306009 | 0.5223622 |
| ENSG00000128578.9 | STRIP2 | 1.74085 | 0.8006286 | -1.31847 | 0.2181475 | 0.515397 | 0.5704395 | 3.249301 | 0.01020509 | 1.035971 | 0.4511061 | -0.547214 | 0.7998818 |
| ENSG00000128581.15 | IFT22 | 0.431224 | 1 | -0.063719 | 0.6645701 | -0.630321 | 0.03835208 | 0.385674 | 0.9042415 | -1.89644 | 0.01099018 | 0.432816 | 0.8868882 |
| ENSG00000128585.17 | MKLN1 | 0.338762 | 0.9790432 | 0.140352 | 0.7487049 | 1.015224 | 1.96E-05 | 0.435462 | 0.01960715 | 1.276448 | 0.005248249 | 0.073036 | 0.715481 |
| ENSG00000128590.4 | DNAJB9 | 0.624631 | 0.9438503 | 0.65828 | 0.04363252 | -0.189022 | 0.7926387 | -0.361385 | 0.8509531 | -0.609461 | 0.1351765 | 0.261126 | 0.7767294 |
| ENSG00000128591.15 | FLNC | 1.450702 | 0.714717 | 0.425835 | 0.09946479 | -1.231677 | 0.05955588 | 1.054693 | 0.01980759 | 2.4261 | 0.02072565 | 0.314192 | 0.782216 |
| ENSG00000128594.7 | LRRC4 | 0.588927 | 0.7693197 | -0.784532 | 0.8703522 | 0.023734 | 0.7437308 | 2.913679 | 0.001566916 | 1.801411 | 0.432453 | -1.088864 | 0.897797 |
| ENSG00000128595.16 | CALU | 0.570899 | 0.6214382 | 0.174652 | 0.5757493 | 0.443194 | 0.03759805 | 0.859846 | 0.001489124 | 0.300828 | 0.2682211 | -0.061834 | 0.8949489 |
| ENSG00000128607.13 | KLHDC10 | 0.758087 | 0.9375802 | -0.282684 | 0.6401456 | 0.434803 | 0.0458126 | 1.192715 | 0.001012319 | 1.640873 | 0.00489708 | 0.425202 | 0.9174022 |
| ENSG00000128626.11 | MRPS12 | -0.909264 | 1 | -0.23497 | 0.1794587 | -0.43855 | 0.02821202 | -0.21168 | 0.004136774 | 0.73307 | 1 | -0.218336 | 0.3446658 |
| ENSG00000128654.13 | MTX2 | -5.288148 | 0.3772759 | 0.527423 | 1 | 0.20528 | 0.9189902 | 0.076551 | 0.8914089 | -0.727509 | 0.01243284 | -1.536496 | 0.6185816 |
| ENSG00000128683.13 | GAD1 | -0.784386 | 0.6214382 | 0.866248 | 0.7944781 | 0.708366 | 0.009192678 | 0.544231 | 0.5056817 | 1.927736 | 0.9670623 | 1.264227 | 0.9147227 |
| ENSG00000128731.15 | HERC2 | -0.042048 | 0.714717 | 0.668752 | 0.8109973 | 0.441723 | 0.5290845 | 0.341215 | 0.03996698 | 1.352076 | 0.06196625 | -1.674297 | 0.9543956 |
| ENSG00000128739.21 | SNRPN | -2.091252 | 0.563505 | -0.374233 | 0.8953649 | -0.109249 | 0.4637903 | 0.086638 | 0.7144938 | -0.474654 | 0.04088924 | 0.064884 | 0.9562147 |
| ENSG00000128881.16 | TTBK2 | -1.19896 | 0.529676 | 0.595578 | 0.4671295 | 0.71517 | 0.2348326 | 0.344705 | 0.1731044 | 1.458218 | 0.03863239 | 2.950708 | 0.9264343 |
| ENSG00000128973.11 | CLN6 | 0.465009 | 0.7736255 | 1.956294 | 0.1550729 | -2.523129 | 0.02588165 | 0.102828 | 0.4919368 | 3.309158 | 0.8021966 | 0.69756 | 0.6829816 |
| ENSG00000129003.15 | VPS13C | -1.441361 | 0.6014343 | -0.423788 | 0.1306777 | 0.53991 | 0.01148268 | 2.311595 | 0.000635083 | 0.773796 | 0.4972851 | 0.802187 | 0.8202624 |
| ENSG00000129083.12 | COPB1 | -0.281788 | 0.714717 | 0.364756 | 1 | 0.762825 | 4.56E-06 | -0.469291 | 0.3644423 | 0.622947 | 0.2709634 | 1.363385 | 0.9147227 |
| ENSG00000129084.17 | PSMA1 | -0.281251 | 0.7736255 | 0.316539 | 0.5277265 | 0.059222 | 0.3652139 | 0.175895 | 0.5160642 | -0.043299 | 0.001818441 | 0.413481 | 0.5344421 |
| ENSG00000129103.17 | SUMF2 | -0.148575 | 0.8006286 | -0.058823 | 0.9993161 | 0.43173 | 0.5290845 | 0.472543 | 0.03640452 | 1.667913 | 0.4218601 | 0.96904 | 0.8539279 |
| ENSG00000129116.17 | PALLD | -1.334496 | 0.4764974 | -0.049372 | 0.7235573 | 0.778071 | 0.002892406 | 0.261237 | 0.9546857 | 0.794279 | 0.05520941 | -0.136019 | 1 |
| ENSG00000129159.6 | KCNC1 | 0.245574 | 0.8006286 | -0.60551 | 0.6527736 | 1.155708 | 0.5290845 | 2.537437 | 0.0215374 | 2.10815 | 0.03964182 | -5.494682 | 0.8260537 |
| ENSG00000129197.14 | RPAIN | -0.165885 | 0.9638653 | 0.135539 | 0.9918484 | 0.101818 | 0.2810805 | 0.770336 | 0.02262926 | 0.315905 | 0.8530275 | -1.035497 | 0.008724428 |
| ENSG00000129235.10 | TXNDC17 | 0.161323 | 1 | 0.169277 | 0.9676809 | -0.380393 | 0.001189836 | -0.383338 | 4.83E-05 | -0.288824 | 4.66E-05 | -0.427838 | 0.06476298 |
| ENSG00000129245.11 | FXR2 | 1.188066 | 0.6368609 | 0.686609 | 0.6527736 | -0.143575 | 0.5290845 | 2.171584 | 0.001432094 | 1.419601 | 0.000665864 | 0.022986 | 0.9326505 |
| ENSG00000129250.11 | KIF1C | -0.106617 | 0.9616839 | 0.108036 | 0.5837659 | 2.35246 | 0.5182859 | 0.978377 | 0.008154536 | 0.904302 | 0.5196843 | -1.607271 | 0.8746228 |
| ENSG00000129255.14 | MPDU1 | -0.66236 | 0.6214382 | 0.091262 | 0.5794755 | -1.002959 | 0.1958328 | 1.078882 | 0.484078 | -0.052962 | 0.09748147 | 0.03743 | 0.04261865 |
| ENSG00000129292.20 | PHF20L1 | 2.143051 | 0.1953452 | -0.022733 | 0.6527736 | 1.146331 | 0.001098914 | 1.73264 | 0.000483583 | 0.629918 | 0.7477593 | 0.755616 | 0.9690603 |
| ENSG00000129315.9 | CCNT1 | 0.431687 | 0.8430867 | 0.481181 | 0.6527736 | 0.600204 | 0.4064599 | 2.846656 | 0.003358301 | 2.432623 | 0.0100413 | -0.566372 | 0.6939365 |
| ENSG00000129351.17 | ILF3 | -1.138991 | 0.0379753 | -0.006968 | 0.5010139 | 0.143808 | 0.2287399 | 0.740861 | 0.001320529 | 1.106417 | 0.001280028 | -1.159243 | 0.3846916 |
| ENSG00000129473.9 | BCL2L2 | 0.712832 | 0.6014343 | -0.56187 | 0.5975609 | -0.94134 | 0.5476072 | -1.129412 | 0.7144938 | 3.72246 | 0.04272514 | -0.174238 | 0.8278576 |
| ENSG00000129515.18 | SNX6 | -1.315307 | 0.6214382 | 0.117427 | 0.6527736 | 0.724754 | 0.004611001 | -1.550896 | 0.02688725 | -0.571832 | 0.1948061 | 1.089989 | 0.9019021 |
| ENSG00000129518.8 | EAPP | 0.18432 | 0.714717 | 0.799727 | 0.6875693 | 1.059276 | 1 | 1.121213 | 0.3281061 | -0.928724 | 0.03223452 | 0.35838 | 0.9147227 |
| ENSG00000129521.13 | EGLN3 | 0.592244 | 0.8006286 | 0.777865 | 0.01853084 | 1.596403 | 0.2101108 | -0.301108 | 0.2232632 | 0.591955 | 0.7545966 | 0.337698 | 0.6885065 |
| ENSG00000129559.12 | NEDD8 | 0.247347 | 0.9850641 | 0.0158 | 0.9111975 | -0.185208 | 8.99E-07 | 0.168904 | 0.2696559 | 0.190418 | 0.2725295 | -0.616448 | 0.8350672 |
| ENSG00000129562.10 | DAD1 | -1.111153 | 0.2600418 | -0.004418 | 0.7383998 | -0.10761 | 0.4455303 | -0.086572 | 0.001362045 | -0.390458 | 2.95E-05 | 1.459595 | 0.8278576 |
| ENSG00000129595.12 | EPB41L4A | 0.777582 | 1 | 0.419373 | 0.6527736 | -1.512957 | 0.239172 | 1.109133 | 0.01791766 | -0.437972 | 0.9072651 | -3.721731 | 0.6517664 |
| ENSG00000129657.14 | SEC14L1 | 0.123416 | 0.9690262 | -0.346128 | 0.3959708 | 0.728612 | 1.85E-05 | 0.959317 | 0.8980137 | 1.452607 | 0.8927617 | 0.375862 | 0.6517664 |
| ENSG00000129911.8 | KLF16 | 1.236407 | 1 | -0.493869 | 0.9104869 | -0.950298 | 0.9133384 | 1.207818 | 0.006323922 | 2.085643 | 0.000475026 | 1.851031 | 0.6185816 |
| ENSG00000129932.7 | DOHH | -0.33037 | 0.6214382 | 0.812244 | 0.3610905 | 1.197676 | 0.000142779 | 0.353972 | 0.1266762 | 0.484799 | 0.2622563 | 0.960846 | 0.9276623 |
| ENSG00000129933.20 | MAU2 | 1.59619 | 0.4764974 | -0.471086 | 0.3795682 | 0.500792 | 0.9325352 | 2.692286 | 0.0047097 | 0.286519 | 0.002884113 | -0.902871 | 0.6885065 |
| ENSG00000129946.10 | SHC2 | 1.230196 | 0.714717 | -1.268331 | 0.7789922 | -1.631754 | 0.8505612 | 1.035454 | 0.2335397 | 0.081986 | 0.04834983 | 1.040641 | 0.9147227 |
| ENSG00000129968.15 | ABHD17A | -1.478652 | 0.6014343 | 0.477917 | 0.6511425 | -0.798356 | 0.000186174 | 0.258611 | 0.04042573 | 1.35454 | 0.4209851 | -1.294569 | 0.9147227 |
| ENSG00000130024.14 | PHF10 | -0.650855 | 0.6214382 | 0.231198 | 0.8263112 | 1.495382 | 0.03056815 | 0.993114 | 0.001673552 | 0.950275 | 0.1806765 | -0.080715 | 0.6885065 |
| ENSG00000130066.16 | SAT1 | -0.970243 | 0.4406957 | -0.007773 | 1 | 0.07711 | 0.4064599 | -0.772952 | 0.0011483 | -0.009524 | 0.5196843 | 0.999589 | 0.01991945 |
| ENSG00000130147.15 | SH3BP4 | 1.245232 | 0.529676 | -0.191691 | 0.877419 | -0.006068 | 0.4064599 | -0.615908 | 0.7144938 | 2.257508 | 0.003203399 | 0.437765 | 1 |
| ENSG00000130175.9 | PRKCSH | 0.161933 | 0.9626331 | 0.026923 | 0.5953732 | 0.381986 | 0.3308096 | 2.223471 | 0.000888791 | 1.65817 | 0.1555238 | -0.095723 | 0.7087388 |
| ENSG00000130193.7 | THEM6 | -1.975698 | 0.9183025 | 0.228899 | 0.8814631 | -1.304789 | 0.4657921 | 1.059827 | 0.01413636 | -0.299273 | 1 | 2.321388 | 0.9346461 |
| ENSG00000130203.9 | APOE | -2.366181 | 1.37E-11 | 0.63191 | 2.54E-23 | -4.586508 | 1.18E-20 | -0.585186 | 0.02352122 | 2.397665 | 0.3835184 | 1.461883 | 0.6185816 |
| ENSG00000130204.12 | TOMM40 | -0.131185 | 0.8006286 | 0.209912 | 0.838909 | -0.538383 | 0.2078638 | 0.516485 | 0.2651362 | 2.090383 | 0.03223591 | 0.615171 | 0.7072982 |
| ENSG00000130222.10 | GADD45G | -2.547759 | 0.6214382 | -0.417339 | 0.289717 | -0.042346 | 0.4653674 | 0.112598 | 0.6614105 | 1.37234 | 0.000447908 | -1.401065 | 0.6885065 |
| ENSG00000130226.16 | DPP6 | -0.595592 | 0.9789948 | -0.399817 | 0.1623079 | 1.114009 | 9.40E-05 | 1.028779 | 0.0128045 | 1.605756 | 0.005969109 | -0.958258 | 0.956933 |
| ENSG00000130227.16 | XPO7 | 0.32654 | 0.6792949 | -0.616625 | 0.6527736 | 1.145733 | 0.01834513 | 0.250028 | 0.0202141 | 0.990421 | 0.3798365 | -1.455606 | 0.8009135 |
| ENSG00000130244.12 | FAM98C | 3.075992 | 0.3098608 | -0.438043 | 0.6527736 | 0.354897 | 0.9750611 | 1.416952 | 0.01836355 | 1.327994 | 0.3283682 | -3.086558 | 0.9062221 |
| ENSG00000130254.11 | SAFB2 | 0.234805 | 0.6493358 | 0.033807 | 0.682287 | 0.687706 | 0.1506649 | 1.435662 | 0.000105024 | 0.390587 | 0.3974318 | -0.447923 | 0.7408601 |
| ENSG00000130255.12 | RPL36 | 0.232718 | 0.191343 | 0.046105 | 0.8217443 | -0.386046 | 8.83E-10 | 0.138696 | 0.9035871 | -0.180569 | 7.65E-05 | -0.108875 | 0.6345741 |
| ENSG00000130287.13 | NCAN | 1.079225 | 0.6487354 | -0.374481 | 0.5656503 | -0.74622 | 0.7321183 | 1.825586 | 3.45E-07 | 1.762497 | 0.001807581 | -1.125153 | 0.2955728 |
| ENSG00000130312.6 | MRPL34 | -0.554981 | 0.6559753 | -0.629333 | 0.7078432 | -0.047531 | 0.7547061 | -1.086231 | 0.007249297 | 0.081789 | 0.1305014 | 1.695903 | 0.8949489 |
| ENSG00000130313.6 | PGLS | -0.023864 | 1 | -0.294278 | 0.5224817 | -1.076759 | 0.000265497 | -0.39251 | 0.03043518 | -0.596941 | 0.1213641 | -0.195762 | 0.6185816 |
| ENSG00000130332.14 | LSM7 | 0.26616 | 0.4392896 | -0.175943 | 0.3121527 | -0.164019 | 0.006271249 | 0.151912 | 0.8198588 | 0.191338 | 0.1610796 | -0.306742 | 0.05044736 |
| ENSG00000130349.9 | C6orf203 | -0.550006 | 0.563505 | 0.206036 | 0.9755916 | -0.808564 | 0.01370691 | 0.858029 | 1 | -0.038993 | 0.2228691 | 2.46856 | 0.7767294 |
| ENSG00000130382.8 | MLLT1 | -0.449088 | 0.9183025 | -0.254398 | 0.9041213 | -1.1607 | 0.6582609 | 1.226403 | 0.09722202 | 1.954353 | 0.04590921 | 1.596648 | 0.6185816 |
| ENSG00000130396.20 | MLLT4 | 0.381685 | 0.8006286 | -0.175789 | 0.5953732 | -0.18351 | 0.3125876 | 0.924855 | 4.14E-06 | 1.986866 | 2.14E-05 | 2.031644 | 1 |
| ENSG00000130402.11 | ACTN4 | -5.889644 | 0.9483118 | 0.199901 | 1 | -0.579201 | 0.5290845 | 1.157042 | 2.31E-05 | 1.03547 | 0.2422209 | -0.659361 | 0.8629701 |
| ENSG00000130429.12 | ARPC1B | -2.059956 | 1 | 0.001225 | 0.9178537 | -0.100576 | 0.01833498 | 0.002386 | 0.3467508 | 0.466931 | 0.3029081 | 0.1194 | 0.9684244 |
| ENSG00000130477.14 | UNC13A | -0.530479 | 0.9501272 | -0.34223 | 0.4487482 | -0.194858 | 0.182889 | 0.992938 | 0.001953302 | 0.445795 | 0.04497572 | -1.196809 | 0.9565287 |
| ENSG00000130479.10 | MAP1S | 0.387834 | 0.6214382 | -0.50168 | 1 | -1.384898 | 0.0546432 | -0.816084 | 0.8042668 | 1.809548 | 0.03056663 | -0.827634 | 0.7111224 |
| ENSG00000130508.10 | PXDN | -1.039674 | 0.8006286 | 0.193809 | 0.9452791 | 1.007164 | 0.141837 | 0.082266 | 0.4919368 | 1.992414 | 0.001624447 | -0.062494 | 1 |
| ENSG00000130511.15 | SSBP4 | -0.234268 | 0.8006286 | -0.156699 | 0.6527736 | -0.304757 | 0.3147505 | 0.854897 | 0.3745792 | 0.554592 | 0.03308831 | -0.610772 | 0.826754 |
| ENSG00000130517.13 | PGPEP1 | 0.092943 | 0.6798783 | -0.24481 | 0.6427295 | 1.184651 | 0.03234075 | 1.046799 | 0.01576557 | 1.434285 | 0.5196843 | 0.219548 | 0.7973567 |
| ENSG00000130520.10 | LSM4 | 0.594708 | 0.4095545 | -0.037475 | 0.797408 | -0.551884 | 5.03E-06 | -0.307171 | 0.01820927 | 0.331015 | 0.7805377 | -0.337696 | 0.8893816 |
| ENSG00000130522.5 | JUND | -1.344585 | 0.6014343 | -0.204094 | 0.7322691 | -0.844013 | 0.5290845 | 0.890872 | 0.01608867 | 1.544213 | 1.15E-06 | 0.449754 | 0.8384951 |
| ENSG00000130529.15 | TRPM4 | -1.362968 | 0.6214382 | -0.036654 | 0.8703522 | 1.997417 | 0.7644337 | 3.763527 | 0.03003616 | 1.490813 | 0.0641349 | -0.694138 | 0.6885065 |
| ENSG00000130544.11 | ZNF557 | -0.101925 | 0.9282145 | 0.474436 | 0.5012872 | -0.872309 | 0.2707921 | 1.306559 | 0.03284013 | 0.942731 | 0.891217 | 1.149501 | 0.9606764 |
| ENSG00000130560.8 | UBAC1 | -0.607813 | 0.6457414 | 0.024919 | 0.8263112 | 0.498558 | 0.3408538 | 1.031619 | 0.2454821 | 1.65891 | 0.04567032 | -0.374222 | 0.8876567 |
| ENSG00000130684.13 | ZNF337 | -3.492329 | 0.007597665 | 0.599597 | 0.9848758 | 0.395059 | 0.8874786 | 0.546243 | 0.1623848 | 2.626748 | 0.2956699 | -0.154954 | 0.9554184 |
| ENSG00000130706.12 | ADRM1 | 0.245573 | 0.3610267 | 0.22575 | 0.8748587 | -0.368138 | 0.01332102 | 0.06383 | 0.1557388 | 0.127779 | 0.3974318 | 1.130002 | 0.9147227 |
| ENSG00000130723.17 | PRRC2B | -1.324619 | 0.563505 | -0.265637 | 0.2810188 | 0.119836 | 0.5290845 | 1.194845 | 3.81E-07 | 1.301822 | 3.26E-05 | 0.078826 | 0.6885065 |
| ENSG00000130724.8 | CHMP2A | -0.011586 | 0.6214382 | 0.091064 | 0.8923091 | -0.111214 | 0.4064599 | -0.236202 | 0.001846366 | -0.251273 | 0.005044997 | 1.39684 | 0.9389125 |
| ENSG00000130725.7 | UBE2M | 0.333993 | 0.9686406 | 0.000147 | 0.8814279 | -0.361224 | 0.000948897 | 0.803163 | 2.28E-06 | 1.762862 | 3.71E-08 | 0.645509 | 0.9576568 |
| ENSG00000130726.11 | TRIM28 | -0.241014 | 0.6214382 | -0.258905 | 0.9676809 | -0.356894 | 0.09431921 | 1.913771 | 3.66E-08 | 0.842035 | 6.76E-07 | 0.366788 | 0.7525885 |
| ENSG00000130731.15 | C16orf13 | 0.696822 | 0.4406957 | 0.047411 | 0.8863806 | -0.666767 | 8.71E-05 | -0.113977 | 0.1709254 | 0.394448 | 0.2349698 | 0.037723 | 0.8949489 |
| ENSG00000130734.9 | ATG4D | -0.653382 | 0.6214382 | 0.148137 | 0.9452791 | -0.768814 | 0.01071855 | 1.367485 | 0.1571591 | 0.229306 | 0.3796044 | 1.873536 | 0.9524428 |
| ENSG00000130748.6 | TMEM160 | 0.330391 | 0.6214382 | -0.517276 | 0.01738075 | -3.226303 | 7.53E-28 | -0.593713 | 0.02113961 | 0.144233 | 0.5196843 | 1.09445 | 0.9554184 |
| ENSG00000130758.7 | MAP3K10 | -2.556925 | 0.8006286 | 0.467682 | 0.6636529 | 0.158481 | 0.5290845 | 1.080504 | 0.01002573 | 1.569652 | 0.3762104 | -1.482207 | 0.8479431 |
| ENSG00000130766.4 | SESN2 | 0.267789 | 0.9348915 | 0.824287 | 0.5042975 | -0.838394 | 0.4791605 | 1.482878 | 0.02455892 | 1.063205 | 0.04017399 | -1.16796 | 0.8479431 |
| ENSG00000130770.17 | ATPIF1 | 0.256242 | 1 | -0.178365 | 0.6527736 | -0.084134 | 0.02389585 | 0.084615 | 0.03997708 | 0.125338 | 0.09510483 | 0.390388 | 1 |
| ENSG00000130779.19 | CLIP1 | -0.314449 | 0.7107241 | 0.079726 | 0.9854207 | 0.337544 | 0.0629293 | 1.710885 | 2.76E-05 | 0.908537 | 0.380721 | 0.364715 | 0.8479431 |
| ENSG00000130810.19 | PPAN | 0.281355 | 0.7795461 | -0.458686 | 0.5170005 | -1.1063 | 0.03404048 | -0.276708 | 0.8949329 | 2.078746 | 0.5196843 | -3.059192 | 0.4304069 |
| ENSG00000130816.14 | DNMT1 | 0.275154 | 0.8006286 | 0.06177 | 0.9810075 | 0.829526 | 0.002285028 | 0.221425 | 0.7045986 | 0.367219 | 0.7016423 | 1.131453 | 0.7832887 |
| ENSG00000130821.15 | SLC6A8 | -0.360533 | 1 | 0.10732 | 0.6015027 | -0.856861 | 0.7064607 | 1.180155 | 0.06835461 | 2.42879 | 0.004512036 | 0.146966 | 1 |
| ENSG00000130827.6 | PLXNA3 | -0.807453 | 0.8871837 | 0.167601 | 0.5818005 | 0.314873 | 0.7437308 | 1.271457 | 0.005698923 | 1.652933 | 0.004607062 | -1.831011 | 0.2281002 |
| ENSG00000130956.13 | HABP4 | -1.266961 | 0.1366571 | -0.561124 | 0.01738344 | 0.143854 | 0.7273239 | 0.374016 | 0.07128885 | 1.097705 | 0.4274216 | -1.117996 | 0.7408828 |
| ENSG00000130958.11 | SLC35D2 | -0.314595 | 0.824666 | 0.090932 | 0.7383618 | -0.395985 | 0.01109271 | -0.635843 | 0.5409218 | 0.586243 | 0.6460384 | -1.936849 | 0.8489889 |
| ENSG00000130985.16 | UBA1 | -0.180853 | 0.8311338 | -0.029329 | 0.875206 | -0.31026 | 0.3395668 | 1.207439 | 0.007159055 | 1.201964 | 0.01178442 | -1.129136 | 0.6885065 |
| ENSG00000131016.16 | AKAP12 | 0.168586 | 0.835624 | 0.477783 | 1.15E-05 | 1.395393 | 0.003137943 | 0.881542 | 0.1471756 | 0.224329 | 0.2286099 | 0.226413 | 0.8479431 |
| ENSG00000131018.22 | SYNE1 | 0.273012 | 1 | 0.485835 | 0.8263112 | 0.795447 | 0.06380182 | -0.808761 | 0.000783168 | 2.370539 | 0.00011686 | -0.586338 | 0.9518271 |
| ENSG00000131051.21 | RBM39 | -1.034436 | 0.04610452 | 0.085793 | 0.796879 | 0.253054 | 3.75E-06 | -0.07366 | 0.5860824 | 0.59633 | 0.9228722 | 0.691359 | 0.9375673 |
| ENSG00000131067.16 | GGT7 | 0.147668 | 0.7402994 | 1.429179 | 0.5311184 | -0.365954 | 0.5290845 | 1.800787 | 4.04E-07 | 0.174104 | 0.3036845 | -1.445811 | 0.6576742 |
| ENSG00000131094.3 | C1QL1 | -0.413621 | 0.4511893 | -0.247536 | 1 | -2.235272 | 0.001922036 | 0.881684 | 0.006455731 | -0.087749 | 0.8597357 | -1.471322 | 0.3820098 |
| ENSG00000131095.11 | GFAP | -0.99515 | 6.88E-05 | -0.131689 | 1 | -0.959946 | 7.36E-05 | -0.053463 | 0.01551148 | 1.377395 | 0.8608026 | -0.122977 | 0.6516659 |
| ENSG00000131100.12 | ATP6V1E1 | -0.578454 | 0.6214382 | 0.304285 | 0.6527736 | 0.025086 | 0.7321183 | -0.357871 | 0.006693037 | 0.454907 | 0.3037489 | 0.854217 | 0.9270789 |
| ENSG00000131116.11 | ZNF428 | -1.267911 | 0.3589335 | -0.209097 | 0.3121527 | -1.005207 | 5.16E-16 | 1.020973 | 9.04E-09 | 0.503696 | 0.2665595 | -0.395176 | 0.2420703 |
| ENSG00000131143.8 | COX4I1 | 0.107557 | 0.9305688 | -0.074293 | 0.971348 | -0.173515 | 0.008705651 | -0.076061 | 0.1126732 | -0.125562 | 0.000298606 | -0.23596 | 0.02166254 |
| ENSG00000131148.8 | EMC8 | -0.599915 | 1 | 0.178167 | 0.6527736 | -0.493531 | 0.000576834 | 0.399941 | 0.8292778 | 0.216513 | 1 | 1.003714 | 0.9518271 |
| ENSG00000131149.17 | GSE1 | 1.024099 | 1 | -0.801453 | 0.5228465 | -0.296839 | 0.9213665 | 1.338372 | 0.0001486 | 0.91285 | 0.006475539 | 1.100666 | 0.9037836 |
| ENSG00000131171.12 | SH3BGRL | -0.671252 | 0.6014343 | 0.037676 | 0.6527736 | 0.603759 | 0.6099207 | 0.739427 | 0.3586405 | -0.263835 | 0.03553583 | 0.509837 | 0.9264343 |
| ENSG00000131174.4 | COX7B | 0.414375 | 0.09715794 | 0.116415 | 0.7383998 | -0.161652 | 5.26E-08 | 0.011626 | 0.1273493 | 0.275349 | 0.2852543 | -0.063713 | 0.2562169 |
| ENSG00000131187.9 | F12 | -1.318436 | 0.8867323 | -0.162872 | 0.5162778 | -1.020291 | 0.03968583 | 0.667851 | 0.08649112 | 0.108741 | 0.6525526 | -1.467106 | 0.2768456 |
| ENSG00000131188.11 | PRR7 | -0.39706 | 0.4995708 | 0.118906 | 0.86041 | -0.881155 | 0.1078054 | 1.466267 | 0.006293281 | 2.494063 | 0.001335033 | -0.420473 | 0.9733982 |
| ENSG00000131242.15 | RAB11FIP4 | -0.903338 | 0.6172955 | 0.554436 | 0.7846957 | -0.14009 | 0.8591179 | 1.123134 | 0.6308467 | 1.8863 | 0.04430596 | 2.700352 | 0.8479431 |
| ENSG00000131263.12 | RLIM | -0.825864 | 0.6214382 | -0.043293 | 0.8629577 | 0.235535 | 0.01694832 | 1.810705 | 0.02308634 | 0.653801 | 0.2880317 | 0.135971 | 0.6885065 |
| ENSG00000131375.9 | CAPN7 | -1.297429 | 0.8437756 | -0.081018 | 0.9103295 | 0.393908 | 0.1992941 | 0.322455 | 0.3017176 | 2.535022 | 0.02205163 | -0.667452 | 0.6885065 |
| ENSG00000131378.13 | RFTN1 | -0.546047 | 1 | -0.249079 | 0.6527736 | -0.544734 | 0.01158143 | -2.097999 | 1 | -0.329466 | 1 | -0.719509 | 0.7308361 |
| ENSG00000131408.13 | NR1H2 | 0.189605 | 0.6214382 | 0.27188 | 1 | -2.066752 | 0.00381867 | -0.373375 | 0.5205536 | 0.954227 | 0.6733743 | -0.408531 | 0.6090735 |
| ENSG00000131409.12 | LRRC4B | -0.901167 | 0.7401773 | -0.466387 | 0.4001101 | -1.231773 | 0.03852063 | 1.535867 | 0.04638111 | 1.243974 | 0.4620283 | -1.756986 | 0.5517992 |
| ENSG00000131437.15 | KIF3A | -1.312756 | 0.8006286 | 0.608293 | 0.7881421 | 1.229967 | 0.03018847 | 1.043403 | 0.007145323 | 0.117589 | 0.7377262 | -0.931559 | 0.5617792 |
| ENSG00000131467.10 | PSME3 | -0.525651 | 0.3132333 | 0.451913 | 0.6527736 | 0.414623 | 0.9139205 | 0.663755 | 0.02131841 | 0.86192 | 0.7131321 | 0.356124 | 0.4301971 |
| ENSG00000131469.12 | RPL27 | 0.423035 | 1.09E-05 | -0.130945 | 0.1143434 | -0.037963 | 0.1433827 | -0.121828 | 0.01003073 | 0.026288 | 0.1381172 | -0.108903 | 0.09368354 |
| ENSG00000131470.14 | PSMC3IP | -0.098117 | 0.714717 | -0.092771 | 0.7786995 | -0.743689 | 0.07756993 | 1.072113 | 0.04341242 | 1.107536 | 0.7052211 | -1.525736 | 0.6885065 |
| ENSG00000131473.16 | ACLY | -3.347518 | 0.6214382 | -0.075718 | 0.873339 | -0.102499 | 0.6615602 | 0.911003 | 0.000162176 | 0.135877 | 0.7775367 | 0.43204 | 0.6675976 |
| ENSG00000131495.8 | NDUFA2 | -0.037312 | 1 | 0.41456 | 0.6527736 | -0.235823 | 0.000948578 | 0.280597 | 0.02086458 | -0.281552 | 0.08966305 | -0.316474 | 0.5344421 |
| ENSG00000131504.15 | DIAPH1 | -0.098411 | 0.9075912 | -0.020815 | 0.9028136 | 0.129708 | 0.1765044 | 1.060876 | 0.209559 | 2.195599 | 0.01555347 | 2.184997 | 0.8384921 |
| ENSG00000131507.10 | NDFIP1 | -0.313113 | 0.8353323 | -0.040927 | 0.5988524 | -0.454963 | 0.000974653 | 0.016697 | 0.2335397 | 0.591891 | 0.5196843 | 0.734375 | 0.9699068 |
| ENSG00000131508.15 | UBE2D2 | -0.132266 | 0.6214382 | -0.251246 | 0.2416476 | -0.194709 | 0.003399897 | 0.16836 | 0.4850832 | 0.22602 | 0.8390546 | 0.285575 | 0.6829816 |
| ENSG00000131558.14 | EXOC4 | 1.857268 | 0.6214382 | -0.168324 | 0.8703522 | 1.481234 | 0.1336022 | 1.7257 | 0.004807921 | 0.911913 | 0.07373239 | 0.939429 | 0.9365938 |
| ENSG00000131653.12 | TRAF7 | -2.995382 | 0.7381604 | -0.323984 | 0.9671539 | -0.7659 | 0.004077784 | 1.346772 | 0.3760083 | 0.786746 | 0.8328283 | 1.099121 | 0.7866135 |
| ENSG00000131711.14 | MAP1B | 0.070392 | 0.08197923 | -0.456305 | 0.000176221 | 0.662912 | 1.03E-12 | 0.536327 | 1.23E-05 | 0.369822 | 0.3974318 | -0.228287 | 0.5524003 |
| ENSG00000131773.13 | KHDRBS3 | -0.90351 | 0.9741156 | -0.176692 | 0.5010139 | 0.526525 | 0.7236298 | 1.134992 | 0.01087618 | 0.604858 | 0.2089173 | 1.212453 | 0.8749632 |
| ENSG00000131791.7 | PRKAB2 | -1.649319 | 0.7736255 | -0.238046 | 0.9113581 | 1.402253 | 0.1198737 | 1.148984 | 0.01291313 | 1.421592 | 0.1686314 | -0.796911 | 0.9750536 |
| ENSG00000131828.13 | PDHA1 | -0.487561 | 0.622607 | -0.05352 | 0.8826101 | 0.035493 | 0.5290845 | -0.051307 | 0.1890064 | -0.281421 | 0.01619679 | 0.873831 | 0.6885065 |
| ENSG00000131871.14 | VIMP | 0.133582 | 0.6014343 | 0.525371 | 0.03324584 | -0.06453 | 0.03892382 | 0.086124 | 0.2752288 | 0.352847 | 0.2363864 | 0.384426 | 0.9375673 |
| ENSG00000131899.10 | LLGL1 | 2.670217 | 0.6014343 | 0.680807 | 0.9622004 | 0.759925 | 0.8450094 | -0.196379 | 0.9645265 | 2.018275 | 0.000597132 | -3.000353 | 0.4578379 |
| ENSG00000131981.15 | LGALS3 | -1.656277 | 0.6214382 | 0.274383 | 0.08172856 | -0.283318 | 3.36E-05 | -0.287037 | 7.32E-05 | -0.417782 | 0.06557358 | 0.67172 | 0.9175671 |
| ENSG00000132002.7 | DNAJB1 | 0.478941 | 0.6014343 | 0.035664 | 0.2350177 | 1.085854 | 1.68E-08 | 1.234665 | 0.2351957 | 1.137178 | 8.61E-05 | 0.213667 | 0.7302807 |
| ENSG00000132155.11 | RAF1 | -0.436124 | 0.8587992 | -0.039853 | 0.6527736 | 0.306159 | 0.8721477 | 1.61248 | 0.09250538 | 1.986807 | 0.000723013 | 1.771983 | 0.5733051 |
| ENSG00000132182.11 | NUP210 | -0.617089 | 0.6214382 | -1.783921 | 0.2756432 | 0.835482 | 0.7437308 | 0.548053 | 0.07163353 | 1.838063 | 0.02912021 | -1.257775 | 0.9147227 |
| ENSG00000132254.12 | ARFIP2 | 0.33738 | 0.9930335 | 0.67311 | 0.5953732 | -0.205125 | 0.007587251 | 0.362778 | 0.4996012 | 0.868839 | 0.6016665 | 0.297868 | 0.9663203 |
| ENSG00000132286.11 | TIMM10B | 1.061097 | 0.4406957 | 0.219727 | 0.9008453 | 0.371774 | 0.9733888 | 1.992124 | 0.002228985 | -1.038561 | 0.4931295 | -0.193025 | 0.7901525 |
| ENSG00000132341.11 | RAN | -0.412359 | 0.6014343 | 0.108056 | 0.880593 | -0.151082 | 0.000788252 | 0.215676 | 1 | 0.16498 | 0.07284488 | -0.521997 | 0.003689052 |
| ENSG00000132388.12 | UBE2G1 | 0.129203 | 0.8413376 | -0.513171 | 0.3651168 | -0.330963 | 0.1316091 | 1.401825 | 0.01438365 | 0.571802 | 0.5389054 | 0.120178 | 0.8391483 |
| ENSG00000132405.18 | TBC1D14 | 0.029693 | 0.6805898 | 1.092445 | 0.698316 | 0.06762 | 0.468897 | 0.011578 | 0.5998925 | 2.448595 | 0.04558844 | 1.289316 | 0.960853 |
| ENSG00000132424.14 | PNISR | -0.735502 | 0.1575101 | -0.11981 | 0.6396549 | 0.681535 | 0.000531237 | 2.192354 | 1.58E-08 | 1.33349 | 2.12E-06 | -0.248336 | 0.7300762 |
| ENSG00000132432.13 | SEC61G | -0.418055 | 0.8006286 | 0.476888 | 5.88E-45 | -0.506476 | 1.33E-12 | -0.096427 | 0.000480996 | -0.255001 | 4.14E-05 | -0.584852 | 0.3946031 |
| ENSG00000132434.9 | LANCL2 | -0.380595 | 0.954995 | -0.069322 | 0.8946072 | 0.376581 | 0.000103573 | 2.350624 | 0.005422346 | 0.77134 | 0.750425 | 0.724956 | 0.9618793 |
| ENSG00000132463.13 | GRSF1 | -0.572146 | 0.8006286 | 0.126668 | 0.6527736 | 0.76832 | 0.2220261 | 1.164407 | 0.03440956 | 0.997524 | 0.4608523 | -1.187619 | 0.6829816 |
| ENSG00000132466.17 | ANKRD17 | -0.534263 | 0.8293347 | -0.135906 | 0.9365571 | 0.734872 | 2.09E-09 | 0.363193 | 0.005790493 | 1.091291 | 0.2011388 | 0.63705 | 0.9556969 |
| ENSG00000132475.8 | H3F3B | 0.197676 | 0.00010072 | -0.012767 | 0.8902958 | 0.13671 | 0.3562859 | 0.205978 | 0.213019 | 0.121607 | 0.8147074 | 0.13392 | 0.6885065 |
| ENSG00000132481.6 | TRIM47 | -2.296215 | 0.1582834 | -0.16226 | 0.4951754 | -0.291399 | 0.150887 | 1.31421 | 0.04459352 | 1.423471 | 0.2322731 | 0.452664 | 0.9617345 |
| ENSG00000132485.12 | ZRANB2 | -1.148643 | 0.6014343 | 0.240732 | 0.890901 | 0.50579 | 0.001229093 | 0.488039 | 0.8379111 | 0.185648 | 0.7269042 | -0.029468 | 0.5344421 |
| ENSG00000132507.17 | EIF5A | 0.562818 | 4.85E-06 | 0.211902 | 0.6527736 | -0.848289 | 1.29E-12 | 0.46288 | 0.1406237 | 0.123431 | 0.819072 | 0.26228 | 0.316809 |
| ENSG00000132510.10 | KDM6B | -3.598916 | 0.07027023 | 0.061485 | 0.6527736 | -0.279968 | 0.7482391 | 1.276997 | 0.008845667 | 1.535335 | 0.05609994 | -2.266225 | 0.6885065 |
| ENSG00000132535.18 | DLG4 | -0.761695 | 0.6214382 | -0.205097 | 0.923953 | -0.650555 | 0.1186365 | 1.052634 | 6.68E-06 | 1.772091 | 0.2404108 | -0.570352 | 0.6829816 |
| ENSG00000132541.10 | HRSP12 | -0.250818 | 0.6214382 | -0.385494 | 0.2713328 | 0.024145 | 0.5999904 | -0.819508 | 0.01641403 | 0.286447 | 0.3974318 | -1.243228 | 0.807533 |
| ENSG00000132563.15 | REEP2 | 0.183661 | 0.8082374 | -0.597557 | 0.2580593 | 0.289181 | 0.09046443 | 0.291301 | 0.4278854 | 2.13419 | 0.04937944 | -1.673449 | 0.6829816 |
| ENSG00000132613.14 | MTSS1L | 0.286219 | 0.6708674 | -0.308573 | 0.5350015 | -0.105716 | 0.7237646 | 1.362987 | 0.01043677 | 1.135024 | 0.001365086 | -0.756815 | 0.7918118 |
| ENSG00000132640.14 | BTBD3 | 0.443418 | 1 | -2.099943 | 0.1938583 | -0.302003 | 0.8100781 | 1.863696 | 0.01291313 | 1.029747 | 0.2975229 | -2.543265 | 0.6885065 |
| ENSG00000132661.3 | NXT1 | -0.684863 | 0.9309704 | 0.166866 | 0.8955188 | -0.328239 | 0.004226592 | 0.497792 | 0.5934221 | 0.211041 | 0.2554619 | -0.588526 | 0.6829816 |
| ENSG00000132664.11 | POLR3F | -4.722126 | 0.03957161 | 0.170178 | 0.3250176 | -0.309195 | 0.7899263 | 0.93874 | 0.01381864 | 1.004583 | 0.5281554 | 0.59767 | 0.7901525 |
| ENSG00000132670.20 | PTPRA | -0.038668 | 0.8787256 | -0.327446 | 0.2416476 | 0.126128 | 0.4722012 | 1.125268 | 0.01002573 | 1.457012 | 0.02663358 | 0.283371 | 0.8375052 |
| ENSG00000132688.10 | NES | -0.893541 | 6.45E-06 | -0.249431 | 0.1943572 | -0.164177 | 0.6295125 | 0.718611 | 7.64E-06 | 0.556491 | 0.1082296 | 0.091779 | 0.4730982 |
| ENSG00000132692.18 | BCAN | -0.123909 | 0.01711078 | -0.14785 | 0.2134316 | 0.225677 | 0.001147586 | 0.41657 | 0.8537667 | 0.944254 | 0.211283 | -0.518474 | 0.03362812 |
| ENSG00000132716.18 | DCAF8 | -0.297555 | 1 | -0.054574 | 0.6487807 | -0.363825 | 0.3683009 | 1.131741 | 0.02485579 | 1.944654 | 0.3974318 | -0.165319 | 0.7827587 |
| ENSG00000132718.8 | SYT11 | 0.122374 | 0.844083 | -0.032343 | 0.6527736 | 0.816782 | 3.90E-06 | -0.003093 | 0.1052874 | 0.834439 | 0.3974318 | 0.267679 | 0.8214575 |
| ENSG00000132763.14 | MMACHC | -1.533851 | 0.6214382 | -0.299512 | 0.9696336 | 0.870774 | 0.7697435 | 2.7471 | 0.01282542 | -0.399606 | 1 | 0.188298 | 0.9684849 |
| ENSG00000132792.18 | CTNNBL1 | -0.585867 | 0.7218619 | 0.052135 | 0.8845769 | 0.245138 | 0.7437308 | 0.666389 | 0.6141936 | -0.162452 | 0.0167936 | 0.449488 | 0.5733051 |
| ENSG00000132819.16 | RBM38 | -1.632404 | 0.6214382 | 0.206929 | 0.9430408 | -1.233201 | 0.04702712 | 0.438092 | 0.02251378 | 2.178031 | 0.006978076 | 0.223784 | 0.6885065 |
| ENSG00000132912.12 | DCTN4 | 0.257962 | 0.6014343 | -0.000279 | 0.5646148 | 0.722306 | 0.01132726 | 0.961142 | 0.03242844 | 0.260658 | 0.632794 | 1.343318 | 0.6885065 |
| ENSG00000132950.18 | ZMYM5 | 1.266688 | 0.9263323 | 0.211171 | 0.8954254 | 1.248515 | 0.000715402 | 2.052504 | 0.4919368 | 0.461964 | 0.7139976 | 3.346004 | 0.6185816 |
| ENSG00000132952.11 | USPL1 | -0.670721 | 0.8075018 | 0.438001 | 0.7595412 | 1.259789 | 0.01151007 | 0.447046 | 0.6655012 | 0.983383 | 0.2353817 | -1.21376 | 0.6885065 |
| ENSG00000132963.7 | POMP | -0.001798 | 0.529676 | 0.337892 | 0.577543 | -0.271753 | 0.000148895 | -0.124797 | 0.001688208 | -0.71375 | 2.07E-09 | 0.299658 | 0.3532606 |
| ENSG00000133026.12 | MYH10 | 0.895622 | 0.7558063 | -0.083044 | 0.5309968 | 0.819133 | 0.00210624 | 1.478236 | 0.001407826 | 0.379479 | 1 | -1.376298 | 0.1834181 |
| ENSG00000133030.19 | MPRIP | -0.732315 | 0.81271 | -0.088697 | 0.5872292 | 0.157828 | 0.05669581 | 0.589066 | 0.000170762 | 1.247804 | 0.03157649 | 0.823175 | 0.5191309 |
| ENSG00000133112.16 | TPT1 | -0.394851 | 0.2683603 | 0.215022 | 0.3835936 | 0.178884 | 0.002630627 | 0.516382 | 0.1862039 | 0.028997 | 0.2826025 | 0.464075 | 0.8278576 |
| ENSG00000133142.17 | TCEAL4 | -0.136136 | 0.7820597 | 0.353426 | 0.8263112 | 0.028264 | 0.2808121 | 0.172084 | 0.5041467 | 0.415039 | 0.04102581 | 0.222687 | 0.9154868 |
| ENSG00000133169.5 | BEX1 | -0.068097 | 0.7349481 | -0.067723 | 0.9768713 | -0.222286 | 0.1725688 | 0.491857 | 0.042941 | -0.277461 | 0.01275075 | -0.566751 | 0.05841333 |
| ENSG00000133226.16 | SRRM1 | -0.283381 | 0.8133722 | 0.115593 | 0.9507102 | 1.109738 | 8.13E-12 | 0.056932 | 0.02384542 | 0.310172 | 0.5196843 | 0.469005 | 0.8479431 |
| ENSG00000133243.8 | BTBD2 | 1.054545 | 0.563505 | -0.190604 | 0.873339 | -0.166462 | 0.4382561 | 0.65661 | 0.002812957 | 1.13058 | 0.01207519 | 0.866394 | 0.6138783 |
| ENSG00000133265.10 | HSPBP1 | -0.326655 | 1 | 0.553091 | 0.5953732 | -0.382254 | 0.000320231 | 0.583726 | 0.5204107 | 0.578579 | 0.8646421 | -0.764159 | 0.3358843 |
| ENSG00000133275.15 | CSNK1G2 | -0.51177 | 1 | 0.206134 | 0.9324033 | -0.904871 | 0.9101956 | 0.834395 | 0.05392813 | 1.320962 | 0.005195662 | 0.055032 | 0.7733704 |
| ENSG00000133318.13 | RTN3 | -0.788793 | 0.8213466 | -0.146704 | 0.7613037 | 0.786294 | 2.65E-09 | 0.575456 | 0.7073943 | 0.775352 | 0.3798869 | -0.676484 | 0.5733051 |
| ENSG00000133398.3 | MED10 | 0.393182 | 0.6633925 | 0.67469 | 0.03499356 | 0.106731 | 0.06507405 | 0.925747 | 0.2419222 | 0.225586 | 0.5196843 | 0.612376 | 1 |
| ENSG00000133597.9 | ADCK2 | -3.356598 | 0.4764974 | -0.153696 | 1 | 0.914024 | 0.6175829 | 2.148636 | 0.007052959 | 2.247093 | 0.1303523 | 1.397012 | 0.7419782 |
| ENSG00000133612.18 | AGAP3 | -0.754354 | 0.6014343 | -0.134962 | 0.4487482 | 0.686122 | 0.5736278 | 0.850392 | 0.00249849 | 1.0014 | 0.01759495 | 0.406194 | 0.9704304 |
| ENSG00000133627.17 | ACTR3B | 0.624813 | 0.8006286 | 0.500118 | 0.9834088 | -0.33281 | 0.589407 | 3.180504 | 0.002427199 | 0.835389 | 0.280178 | -1.769379 | 0.6885065 |
| ENSG00000133657.14 | ATP13A3 | -3.509698 | 0.6214382 | -0.232026 | 0.6376243 | 1.081813 | 0.3669917 | 1.002387 | 0.003636421 | 0.386379 | 0.5196843 | -2.020906 | 0.9671638 |
| ENSG00000133706.17 | LARS | -0.4248 | 0.824966 | -0.221317 | 0.03414591 | 0.586751 | 9.12E-07 | 0.220507 | 0.4853154 | 0.174341 | 0.3255801 | 0.977062 | 0.9860669 |
| ENSG00000133812.14 | SBF2 | 1.138003 | 0.8006286 | -0.296379 | 0.6527736 | -0.328479 | 0.9083351 | 1.209373 | 0.0171066 | 0.448771 | 0.1710877 | -2.728665 | 0.8479431 |
| ENSG00000133858.15 | ZFC3H1 | -0.269431 | 0.6214382 | -0.875895 | 0.7785607 | 1.096507 | 0.001412382 | 0.768169 | 0.3010358 | 2.45643 | 0.000566666 | 0.578213 | 0.8479431 |
| ENSG00000133874.1 | RNF122 | -0.874789 | 0.9047689 | -0.604264 | 0.6527736 | -3.305456 | 0.03847234 | 2.180344 | 0.01389293 | 1.704305 | 0.006262912 | 2.106536 | 1 |
| ENSG00000133878.8 | DUSP26 | 1.132896 | 0.6014343 | -1.160266 | 0.1809109 | -1.3514 | 0.528724 | 0.929894 | 0.006524058 | 0.370666 | 0.8993413 | -0.422428 | 0.7419782 |
| ENSG00000133935.6 | C14orf1 | -1.141613 | 0.3946735 | -0.346842 | 0.6527736 | 1.51923 | 0.5290845 | 0.57525 | 0.5681026 | -0.727692 | 0.02431778 | -2.558122 | 0.6885065 |
| ENSG00000133983.14 | COX16 | 0.314543 | 0.5454431 | 1.066057 | 0.8115448 | -2.872503 | 3.28E-05 | 0.372448 | 0.6166893 | 1.432255 | 0.8752597 | 2.932664 | 0.6885065 |
| ENSG00000134030.13 | CTIF | 0.263063 | 1 | -0.546979 | 0.6107341 | 0.60967 | 1 | 1.526099 | 0.03266133 | 1.535632 | 0.5196843 | -0.26445 | 0.9552941 |
| ENSG00000134056.11 | MRPS36 | 0.156352 | 0.8901384 | 0.306064 | 0.6527736 | -0.794823 | 0.001289673 | 0.069338 | 0.0910202 | 0.233818 | 0.1939847 | -1.934918 | 0.6885065 |
| ENSG00000134086.7 | VHL | 0.612108 | 0.8867323 | 0.148625 | 0.6527736 | 1.021129 | 0.000331022 | 0.689832 | 0.06316214 | 1.346301 | 0.2016786 | -2.098377 | 1 |
| ENSG00000134107.4 | BHLHE40 | 0.766013 | 0.8006286 | -0.215349 | 0.6527736 | -0.611176 | 0.005437942 | -0.194446 | 0.3274847 | 0.323954 | 0.4954444 | 2.876049 | 0.1508168 |
| ENSG00000134108.12 | ARL8B | -1.139001 | 0.3946735 | 0.076461 | 0.6540268 | 0.922406 | 0.3943115 | 0.793264 | 0.7981169 | 1.750201 | 0.02900841 | -0.599606 | 0.794948 |
| ENSG00000134152.10 | KATNBL1 | 0.172475 | 0.6337971 | -0.329232 | 0.6514723 | 0.964263 | 0.002089198 | -0.239224 | 0.8042668 | 0.163133 | 0.9463565 | 1.340858 | 1 |
| ENSG00000134153.9 | EMC7 | 0.108808 | 0.6214382 | 0.272437 | 0.8934507 | -0.001434 | 0.4258949 | -0.08042 | 0.07227293 | -0.756045 | 0.00360099 | 0.029027 | 0.9517616 |
| ENSG00000134186.11 | PRPF38B | -0.400063 | 0.4673415 | -0.245816 | 0.5394427 | 0.813253 | 2.48E-07 | 0.704182 | 0.3224734 | 0.286843 | 0.2773103 | 0.007912 | 0.9532204 |
| ENSG00000134243.11 | SORT1 | -0.301058 | 0.6214382 | -0.098772 | 0.5733119 | 0.734167 | 0.6651783 | 0.764127 | 0.01428451 | -0.126824 | 0.742061 | 0.43836 | 0.9536927 |
| ENSG00000134248.13 | LAMTOR5 | -0.137721 | 0.8744211 | 0.147775 | 0.7725713 | -0.121603 | 0.01282359 | 0.213733 | 0.003173628 | 0.248387 | 0.03484403 | 0.540981 | 0.7870938 |
| ENSG00000134294.13 | SLC38A2 | 0.220759 | 0.6214382 | 0.090417 | 0.8703522 | 0.401377 | 6.48E-05 | 0.433753 | 0.001646633 | 0.406329 | 0.02556428 | 0.980406 | 0.8278546 |
| ENSG00000134308.13 | YWHAQ | 0.349453 | 0.2670389 | -0.031572 | 0.6527736 | 0.167136 | 0.7354314 | 0.55859 | 0.1727122 | 0.56013 | 0.9622018 | -1.777103 | 0.005067357 |
| ENSG00000134313.14 | KIDINS220 | 0.350486 | 0.6956464 | 0.453077 | 0.846862 | 0.274425 | 0.8966914 | 1.06314 | 4.30E-05 | 0.32516 | 0.8458122 | -1.304822 | 0.577079 |
| ENSG00000134318.13 | ROCK2 | 0.355705 | 0.7736255 | -0.254111 | 0.4562492 | 1.419612 | 0.01692749 | 2.561414 | 8.57E-07 | 1.120178 | 0.8385425 | -0.504363 | 1 |
| ENSG00000134324.11 | LPIN1 | -1.355863 | 0.01020723 | 0.191401 | 0.7331946 | 0.912348 | 0.1255046 | 1.041204 | 0.03096313 | 1.793845 | 0.04382212 | -0.696026 | 0.9147227 |
| ENSG00000134352.19 | IL6ST | -0.569923 | 0.7669132 | -0.276267 | 0.04799695 | 0.598092 | 0.4664295 | 1.29992 | 6.26E-06 | 1.170026 | 0.6895464 | 1.663696 | 0.8479431 |
| ENSG00000134369.15 | NAV1 | -0.042061 | 0.9371596 | -0.314609 | 0.2837273 | 0.519517 | 0.02271339 | 0.894822 | 0.002892901 | 0.280792 | 0.1697005 | -1.195254 | 0.6829816 |
| ENSG00000134376.14 | CRB1 | -1.460867 | 0.8587992 | 0.792885 | 0.6481154 | 0.355643 | 0.6145827 | 3.749215 | 0.004575151 | 0.06169 | 0.6357065 | -2.168533 | 0.7767294 |
| ENSG00000134419.15 | RPS15A | -0.652052 | 0.03176252 | -0.039106 | 0.7841336 | 0.080814 | 0.5290845 | -0.23518 | 0.002321358 | 0.203495 | 0.3259578 | 0.035076 | 0.5344421 |
| ENSG00000134440.11 | NARS | 0.376769 | 0.6214382 | 0.16439 | 1 | 0.471031 | 0.007067549 | 0.326309 | 0.7144938 | -0.335941 | 0.002717053 | 0.204161 | 0.9581586 |
| ENSG00000134531.9 | EMP1 | 1.361301 | 0.003228446 | -0.497862 | 0.1054706 | 0.285093 | 0.1182794 | 0.093928 | 0.4047786 | 1.108259 | 0.5301735 | 0.082484 | 0.8214575 |
| ENSG00000134590.13 | FAM127A | -0.165648 | 0.7640133 | 0.172553 | 0.9222921 | -0.594056 | 8.77E-07 | -0.280921 | 0.1331699 | 0.437384 | 0.5196843 | -0.24438 | 1 |
| ENSG00000134594.4 | RAB33A | -0.239637 | 0.8587992 | -1.259395 | 0.9192533 | 0.352493 | 0.2879524 | -0.96122 | 0.01014885 | -1.777244 | 0.02681478 | -3.182599 | 0.1427307 |
| ENSG00000134644.15 | PUM1 | 1.500883 | 0.1575216 | 0.16146 | 0.6527736 | 0.311146 | 0.06805893 | 1.646278 | 0.001768971 | 1.607513 | 5.77E-05 | -0.860502 | 0.8560727 |
| ENSG00000134684.10 | YARS | 0.021973 | 0.7259912 | 0.220239 | 0.9859883 | -0.232412 | 0.896092 | 0.708294 | 2.54E-05 | 0.676492 | 0.2791546 | 0.308814 | 0.9562356 |
| ENSG00000134686.16 | PHC2 | -1.003322 | 0.6214382 | -0.061617 | 0.8320884 | -0.095069 | 1 | 0.928505 | 3.33E-06 | 1.0104 | 0.000101933 | 0.884735 | 0.9161136 |
| ENSG00000134744.13 | ZCCHC11 | -1.024831 | 1 | 0.261047 | 0.6934038 | 1.227425 | 0.001191301 | 0.320164 | 0.6602753 | 1.123328 | 0.7528548 | 0.438319 | 1 |
| ENSG00000134748.12 | PRPF38A | -1.199777 | 0.9762384 | 0.028414 | 0.8106695 | -0.233505 | 0.4388213 | 1.21703 | 0.02549987 | 0.408528 | 0.6913914 | 0.575675 | 0.9428037 |
| ENSG00000134759.13 | ELP2 | 0.406268 | 0.9794907 | 0.145775 | 0.9008453 | 0.476646 | 0.02505569 | 0.586326 | 0.007043323 | 0.54272 | 0.427678 | -1.299329 | 0.9562356 |
| ENSG00000134809.8 | TIMM10 | -1.292103 | 0.6014343 | -0.412159 | 0.7852785 | -1.069893 | 2.36E-05 | -0.126037 | 0.132669 | -0.634236 | 0.1119542 | 0.188296 | 0.4629017 |
| ENSG00000134825.13 | TMEM258 | 0.423327 | 0.4406957 | 0.070614 | 0.9893015 | -0.134151 | 0.000130234 | -0.34853 | 6.30E-05 | -0.337222 | 3.39E-06 | -0.171383 | 0.5344421 |
| ENSG00000134851.12 | TMEM165 | -1.781524 | 1 | -0.387989 | 0.6239827 | -0.623994 | 0.01098563 | 0.65857 | 0.02592012 | 1.88769 | 0.04810533 | 0.62238 | 0.5344421 |
| ENSG00000134852.14 | CLOCK | 2.305222 | 0.1832812 | -0.793439 | 0.2578431 | 0.920337 | 0.03337556 | -0.040902 | 0.1565606 | 0.341252 | 0.2880317 | 0.564887 | 0.8663281 |
| ENSG00000134874.17 | DZIP1 | -0.279177 | 0.8587992 | -0.681279 | 0.5444891 | 1.030103 | 0.002124477 | 1.595666 | 0.1213293 | 0.963064 | 0.2019818 | 1.320956 | 0.9634357 |
| ENSG00000134884.13 | ARGLU1 | -0.054444 | 0.9241468 | -0.316205 | 0.5010139 | 0.221112 | 0.3519576 | 0.759192 | 0.02533603 | 0.8346 | 0.4315259 | 0.208378 | 0.7153093 |
| ENSG00000134900.11 | TPP2 | -0.50426 | 0.6214382 | -0.059461 | 0.8639685 | -0.177975 | 0.5513837 | 1.563642 | 0.0438069 | 1.293131 | 0.6296645 | 1.05402 | 0.6885065 |
| ENSG00000134910.12 | STT3A | 0.625911 | 0.2979597 | 0.066717 | 0.5907462 | 0.369507 | 1 | 0.908137 | 0.01757389 | 0.665442 | 0.9794739 | -0.588814 | 0.6885065 |
| ENSG00000134970.13 | TMED7 | 1.688118 | 0.292386 | 0.169345 | 0.6427295 | -0.927087 | 0.000652166 | 0.920637 | 0.03242844 | 1.680588 | 0.1541634 | -0.252685 | 1 |
| ENSG00000134982.16 | APC | 1.277387 | 0.1551568 | -0.503654 | 0.005446727 | 0.74608 | 5.95E-07 | 1.162988 | 0.0001957 | 0.683522 | 0.5196843 | -0.067383 | 0.6829816 |
| ENSG00000134986.13 | NREP | 0.545964 | 0.2177303 | -0.505001 | 0.1214069 | 0.138951 | 0.8318275 | 1.124126 | 0.01964373 | 0.143888 | 0.584557 | -0.722657 | 0.4773627 |
| ENSG00000134987.11 | WDR36 | -0.817218 | 0.5242565 | 1.05603 | 0.5681739 | 0.204482 | 0.4382561 | 1.642311 | 0.009399882 | 1.239502 | 0.2455619 | 2.12467 | 0.6885065 |
| ENSG00000135018.13 | UBQLN1 | -0.473236 | 0.6214382 | 0.109776 | 0.9596929 | 0.433482 | 0.2227885 | 0.476758 | 0.01757389 | 0.669463 | 0.1486318 | -0.004764 | 0.8479431 |
| ENSG00000135047.14 | CTSL | -0.423982 | 0.7472097 | 0.666893 | 1.51E-05 | -0.215932 | 0.005898071 | 0.573875 | 0.7144938 | 0.360406 | 0.5196843 | 1.034874 | 0.6885065 |
| ENSG00000135070.13 | ISCA1 | -0.112556 | 1 | -0.187182 | 0.9440625 | -1.217445 | 0.2348326 | 0.847644 | 0.0476234 | -0.187418 | 0.6860878 | 1.31448 | 0.6829816 |
| ENSG00000135090.13 | TAOK3 | -0.132361 | 1 | -0.074141 | 0.8953649 | 0.264815 | 5.24E-05 | 1.582038 | 0.2137901 | 0.796069 | 0.9247793 | -1.211577 | 0.7530716 |
| ENSG00000135093.12 | USP30 | -0.245761 | 0.8006286 | 0.587101 | 0.6527736 | 0.537188 | 0.8935082 | 2.627447 | 0.02116525 | 1.350184 | 0.7775367 | -1.199037 | 0.6885065 |
| ENSG00000135097.6 | MSI1 | 0.852707 | 0.6214382 | 0.315683 | 0.8944225 | 0.52746 | 0.6295125 | 0.76151 | 0.007052959 | 0.828835 | 0.05112503 | -0.691931 | 0.6885065 |
| ENSG00000135108.14 | FBXO21 | -0.128626 | 0.6214382 | 0.489467 | 0.9478771 | 0.817139 | 0.0002759 | 0.003554 | 0.5333586 | -0.035348 | 0.3809978 | 0.296903 | 0.6206888 |
| ENSG00000135144.7 | DTX1 | 0.451123 | 0.714717 | -0.242033 | 0.6527736 | -0.324017 | 0.5890545 | 2.173461 | 0.1571142 | 1.070518 | 0.03056663 | 1.163692 | 0.7901525 |
| ENSG00000135164.18 | DMTF1 | 0.042696 | 0.6214382 | 0.517271 | 0.9461836 | 1.198541 | 7.35E-05 | 1.290129 | 0.2389024 | 1.123543 | 0.6217705 | 0.037213 | 0.7942034 |
| ENSG00000135205.14 | CCDC146 | -3.039229 | 0.8587992 | 0.453742 | 0.7680226 | 1.538457 | 0.8656249 | 3.394457 | 0.01413636 | 1.410977 | 1 | 2.738221 | 0.6885065 |
| ENSG00000135241.16 | PNPLA8 | -0.179997 | 1 | 0.01559 | 0.8263112 | 0.839039 | 4.89E-06 | 0.122459 | 0.4919368 | 0.747945 | 0.95261 | 0.306589 | 1 |
| ENSG00000135245.9 | HILPDA | -0.107347 | 1 | 0.547985 | 0.000189974 | 0.103292 | 0.5666039 | 0.739118 | 0.4956881 | -0.45475 | 0.8472945 | 0.025859 | 0.7176063 |
| ENSG00000135250.16 | SRPK2 | 0.597575 | 0.5711449 | 0.079336 | 0.6527736 | 0.833609 | 2.23E-07 | 0.543023 | 0.2444151 | 0.381985 | 0.8608026 | -1.634958 | 0.2768456 |
| ENSG00000135298.13 | ADGRB3 | -2.05539 | 0.529676 | -0.338756 | 0.9707115 | 1.242578 | 0.3399293 | 4.779392 | 0.1452614 | 1.578351 | 0.03666503 | -1.240212 | 0.9147227 |
| ENSG00000135315.11 | CEP162 | 1.553613 | 0.8867323 | 0.88799 | 0.7647276 | 0.239967 | 0.35286 | 0.870901 | 0.1219909 | -1.152163 | 0.04034828 | -1.523298 | 0.7298895 |
| ENSG00000135316.17 | SYNCRIP | -0.892712 | 0.4764974 | -0.202036 | 0.5010139 | 1.227821 | 5.16E-07 | 0.701452 | 0.008366469 | 0.285621 | 0.8574115 | 0.58103 | 1 |
| ENSG00000135318.11 | NT5E | 0.341835 | 0.9624741 | -0.062427 | 0.8504311 | 0.486396 | 0.3393299 | 1.334889 | 8.92E-05 | 1.463886 | 0.4493174 | 0.647812 | 0.8949489 |
| ENSG00000135338.13 | LCA5 | 0.257366 | 1 | 0.969332 | 0.5967879 | 1.097339 | 0.002317644 | 2.294459 | 0.5194681 | 0.503955 | 0.937993 | inf | 0.6185816 |
| ENSG00000135363.11 | LMO2 | -0.034232 | 0.8006286 | -0.229771 | 0.4284444 | 0.001319 | 0.5290845 | -1.149756 | 0.01797834 | -1.27437 | 0.02813969 | -4.140034 | 0.5344421 |
| ENSG00000135365.15 | PHF21A | -1.076732 | 0.8587992 | -0.656374 | 0.5010139 | 0.758364 | 0.01709712 | 1.115505 | 0.683125 | 0.223089 | 0.7134836 | -0.480656 | 0.6885065 |
| ENSG00000135372.8 | NAT10 | 1.379075 | 0.6214382 | -0.582868 | 0.4442782 | 1.100929 | 0.3843036 | 2.145362 | 4.07E-06 | 0.782387 | 0.05413046 | -0.977312 | 0.4998763 |
| ENSG00000135387.19 | CAPRIN1 | 0.357899 | 0.8006286 | -0.078107 | 0.5296245 | -0.240508 | 1 | 1.06503 | 1.54E-08 | 0.4354 | 0.5196843 | 0.900403 | 0.9320234 |
| ENSG00000135390.17 | ATP5G2 | -0.099172 | 0.6214382 | -0.172983 | 0.1643663 | -0.295202 | 0.003675986 | 0.314392 | 1 | -0.142618 | 0.00339458 | -0.788051 | 0.04864488 |
| ENSG00000135404.11 | CD63 | -0.567098 | 0.6603437 | -0.153011 | 0.03594505 | -0.009991 | 0.04009247 | 0.076303 | 0.9788705 | -0.251522 | 1.70E-06 | 0.171055 | 1 |
| ENSG00000135414.9 | GDF11 | -0.159485 | 0.7646067 | -0.439631 | 0.6909081 | -0.366539 | 0.5607166 | 1.635674 | 6.42E-05 | 1.248249 | 0.167884 | 0.239772 | 0.8949489 |
| ENSG00000135424.15 | ITGA7 | 2.209599 | 0.5242565 | 0.000379 | 0.5953732 | -0.751469 | 0.9642892 | 1.449858 | 0.0001486 | 0.83259 | 0.2686155 | 0.530608 | 0.7767294 |
| ENSG00000135441.7 | BLOC1S1 | 0.233553 | 0.342661 | 0.075261 | 0.9876393 | -0.870495 | 1.88E-09 | 0.312206 | 0.384552 | -0.229935 | 0.0404569 | 0.535081 | 0.4553016 |
| ENSG00000135452.9 | TSPAN31 | -0.0501 | 0.9047689 | 0.1383 | 0.9111975 | -0.695267 | 0.1607212 | 0.873984 | 0.5288997 | 0.165464 | 0.03788656 | 0.925883 | 0.744192 |
| ENSG00000135454.13 | B4GALNT1 | -0.369298 | 0.714717 | -0.239672 | 0.6527736 | 0.498181 | 0.9953601 | 0.056558 | 0.6894417 | -0.317549 | 0.000142653 | -0.607935 | 0.6829816 |
| ENSG00000135486.17 | HNRNPA1 | -0.943686 | 0.000286034 | -0.032804 | 0.5953732 | 0.455049 | 0.000121848 | 0.091637 | 0.900061 | 0.435515 | 0.5196843 | -0.332751 | 0.05824041 |
| ENSG00000135503.12 | ACVR1B | -0.080744 | 0.9464322 | -1.222014 | 0.1573875 | -1.055523 | 0.9683466 | 1.289131 | 0.03061816 | 1.74012 | 0.1144781 | -0.579875 | 0.8309897 |
| ENSG00000135506.15 | OS9 | 0.366958 | 0.6214382 | -0.059285 | 0.8953649 | 0.684025 | 0.5290845 | 0.854558 | 0.0203976 | 0.432545 | 0.5716684 | -0.014205 | 0.6885065 |
| ENSG00000135540.11 | NHSL1 | 1.89669 | 0.7736255 | -1.256465 | 0.5840314 | 0.198831 | 1 | 1.879421 | 0.03308457 | 1.812748 | 0.05118678 | -3.078282 | 0.6556863 |
| ENSG00000135541.20 | AHI1 | 1.359338 | 0.9305688 | -0.287142 | 0.6527736 | 0.140179 | 0.08694649 | 1.20929 | 0.004267356 | 0.534342 | 0.3797324 | -3.152964 | 0.9147227 |
| ENSG00000135549.14 | PKIB | 1.215208 | 0.6014343 | 0.267431 | 1 | 0.040195 | 0.01921588 | 0.446177 | 0.8224647 | -0.479635 | 0.007549975 | -0.529692 | 0.9134607 |
| ENSG00000135596.17 | MICAL1 | 1.354013 | 0.714717 | 0.627443 | 0.6527736 | 0.246704 | 0.8832504 | 1.349761 | 0.6493189 | 1.169693 | 0.03825809 | 1.746926 | 0.8770536 |
| ENSG00000135597.18 | REPS1 | 0.92461 | 0.8006286 | -0.336977 | 0.8703522 | 0.01999 | 0.9299562 | 0.52187 | 0.0215755 | 1.602309 | 0.3424233 | 1.275074 | 0.9067556 |
| ENSG00000135617.3 | PRADC1 | -0.327142 | 0.8006286 | 0.062419 | 0.3946411 | -0.162814 | 0.03616489 | -0.141473 | 0.7910526 | 1.294431 | 0.5196843 | -0.014775 | 0.9379128 |
| ENSG00000135643.4 | KCNMB4 | -2.184113 | 0.5084093 | -0.206501 | 0.01857467 | -1.818219 | 0.1431991 | 1.379433 | 0.09126843 | 1.861506 | 0.01759495 | -2.884229 | 0.7333942 |
| ENSG00000135655.13 | USP15 | -0.866379 | 0.6014343 | 0.433401 | 0.4011918 | 0.323828 | 0.02196322 | 0.570038 | 0.9074057 | -0.716049 | 0.5196843 | -0.696401 | 0.6885065 |
| ENSG00000135679.21 | MDM2 | 0.268327 | 0.8006286 | -0.219825 | 0.5953732 | 0.264467 | 0.342534 | 1.843989 | 1.25E-05 | 1.019098 | 0.09943621 | 0.460815 | 0.9565287 |
| ENSG00000135709.12 | KIAA0513 | -0.759482 | 0.9047689 | -0.191955 | 0.8703522 | 1.118404 | 0.2269434 | -0.286815 | 0.2047494 | 2.120024 | 0.01299526 | -0.588297 | 0.9581926 |
| ENSG00000135720.12 | DYNC1LI2 | -3.291066 | 0.6214382 | -0.036222 | 0.6527736 | 0.674374 | 0.02484612 | 1.189291 | 0.000218648 | 1.172279 | 0.1065267 | -0.995118 | 0.9147227 |
| ENSG00000135749.18 | PCNXL2 | -0.112388 | 0.6504346 | -0.784268 | 0.3045812 | 0.823323 | 0.003019379 | 0.906374 | 0.07318798 | 2.031601 | 0.225578 | -0.844629 | 0.9713975 |
| ENSG00000135821.16 | GLUL | 0.594608 | 0.05468731 | 0.318162 | 0.003161586 | -0.1933 | 0.02344843 | 0.434339 | 0.1085878 | 0.766865 | 0.3289736 | 1.758714 | 0.04073667 |
| ENSG00000135823.13 | STX6 | -0.315958 | 0.759776 | -0.508717 | 0.4882009 | 0.819469 | 0.4064599 | 1.095346 | 0.01479058 | 0.189184 | 0.9245349 | 1.571197 | 0.8746228 |
| ENSG00000135829.16 | DHX9 | 0.52967 | 0.8485222 | -0.225707 | 0.6527736 | 0.168155 | 0.2251962 | 0.836755 | 0.03258538 | 0.648658 | 0.5196843 | -0.704059 | 0.6885065 |
| ENSG00000135837.15 | CEP350 | 1.06716 | 0.6014343 | -0.06362 | 0.3298988 | 0.256511 | 0.02391163 | 1.287542 | 0.001680614 | 2.279883 | 0.01557103 | 2.116794 | 0.5137734 |
| ENSG00000135870.11 | RC3H1 | -1.149116 | 0.5901699 | 0.107279 | 0.8306148 | 0.405347 | 0.9063159 | 0.246437 | 0.03328858 | 1.555096 | 0.09889004 | 0.437712 | 0.8383091 |
| ENSG00000135905.18 | DOCK10 | 1.323867 | 0.6214382 | 0.531204 | 0.6527736 | 1.848374 | 0.5018872 | 1.806607 | 0.02534851 | 1.466548 | 0.1407039 | 1.253502 | 0.9562356 |
| ENSG00000135913.10 | USP37 | -0.640226 | 0.7736255 | -0.079812 | 0.9822141 | 1.754206 | 0.1172087 | 2.641034 | 0.01134548 | 0.728153 | 0.9349828 | -0.207437 | 0.8949489 |
| ENSG00000135916.15 | ITM2C | -1.547167 | 0.002148045 | 0.037061 | 0.6132238 | 0.332787 | 0.5374158 | 0.045961 | 1.75E-05 | 0.257981 | 0.2141599 | 0.149072 | 1 |
| ENSG00000135919.12 | SERPINE2 | -0.151756 | 1 | -0.247414 | 0.1905717 | -0.04531 | 0.1765415 | 0.281699 | 0.04775009 | -0.057177 | 0.9975733 | -0.198977 | 0.6885065 |
| ENSG00000135932.10 | CAB39 | -0.820876 | 0.4417256 | 0.026603 | 0.9164212 | 1.253153 | 0.000165877 | 2.118337 | 0.06285065 | 0.105756 | 0.6574572 | -0.032285 | 0.6829816 |
| ENSG00000135940.6 | COX5B | -0.135581 | 0.4764974 | 0.005366 | 0.971348 | -0.216392 | 6.08E-07 | -0.154872 | 0.01223151 | -0.61336 | 1.07E-06 | -0.038224 | 0.1177824 |
| ENSG00000135966.12 | TGFBRAP1 | -0.09363 | 0.7736255 | 0.351278 | 0.9977627 | 0.543446 | 0.9755686 | 0.874746 | 0.01537076 | -0.247795 | 1 | -2.087289 | 0.7561621 |
| ENSG00000135968.19 | GCC2 | -1.546634 | 0.1367467 | 0.226963 | 0.8790112 | 1.432048 | 1.49E-06 | 0.932271 | 0.4919368 | 0.25933 | 1 | 1.997367 | 0.9565287 |
| ENSG00000135976.16 | ANKRD36 | 0.781791 | 0.6014343 | -0.339632 | 0.6527736 | 0.192819 | 0.06336058 | 2.403564 | 2.91E-05 | 0.903647 | 0.000140484 | 2.286835 | 0.6885065 |
| ENSG00000136021.18 | SCYL2 | -4.850489 | 0.6214382 | -0.395309 | 0.8070656 | 0.565742 | 0.002799527 | 0.441014 | 0.1562241 | 1.047189 | 0.5196843 | 1.079538 | 0.8579425 |
| ENSG00000136026.13 | CKAP4 | 0.566971 | 1 | -0.634345 | 0.01825744 | -0.075783 | 0.4435829 | 0.747866 | 0.3314174 | 0.646835 | 0.9032611 | 0.820124 | 0.7123894 |
| ENSG00000136045.11 | PWP1 | -0.545201 | 0.4764974 | -0.156909 | 0.900087 | 0.42303 | 0.04212583 | -0.126249 | 0.9716021 | -0.143282 | 0.007301051 | -1.657041 | 0.5956045 |
| ENSG00000136051.13 | KIAA1033 | -0.44062 | 0.7811112 | -0.126819 | 0.9502109 | 0.554726 | 0.009884704 | 1.01362 | 0.005871422 | 0.702731 | 1 | -0.144501 | 0.6885065 |
| ENSG00000136099.13 | PCDH8 | 4.234017 | 0.2335514 | -0.136138 | 0.7828571 | -1.291264 | 0.02505569 | 0.089487 | 0.8224647 | -0.790504 | 0.5196843 | 1.241559 | 0.8249021 |
| ENSG00000136108.14 | CKAP2 | -1.257711 | 0.4692975 | -0.198304 | 0.5010139 | 1.363063 | 9.23E-05 | -0.454568 | 0.5712232 | 0.090253 | 0.8880448 | 0.387191 | 0.5026528 |
| ENSG00000136141.14 | LRCH1 | 2.186652 | 0.8006286 | 0.330393 | 0.8872744 | -0.864603 | 0.5290845 | -0.593386 | 1 | 3.738514 | 0.02713558 | 1.229692 | 0.9147227 |
| ENSG00000136156.12 | ITM2B | 0.023222 | 0.6014343 | -0.164834 | 0.7057076 | -0.485029 | 1.88E-14 | -0.002654 | 0.002036528 | 0.428051 | 0.5196843 | 0.283594 | 0.9536927 |
| ENSG00000136158.10 | SPRY2 | 0.319607 | 0.6214382 | 0.130565 | 1 | 0.404882 | 0.2423039 | 1.668203 | 0.002854551 | 1.815542 | 0.3796044 | 0.829861 | 0.7767294 |
| ENSG00000136193.16 | SCRN1 | -1.515429 | 0.6214382 | -0.389677 | 0.01886794 | 0.399752 | 0.001653331 | 0.105432 | 0.1020163 | 1.178268 | 0.1446007 | 0.225874 | 0.6885065 |
| ENSG00000136238.17 | RAC1 | -0.605979 | 0.015117 | -0.01141 | 0.3947664 | 0.711527 | 0.03156933 | 0.072516 | 0.2322369 | 0.354868 | 0.3270819 | -0.171815 | 0.2936114 |
| ENSG00000136240.9 | KDELR2 | -0.289987 | 0.6014343 | 0.08695 | 0.7985277 | -0.204306 | 7.55E-06 | 0.013713 | 1 | 0.667724 | 0.4055432 | 0.061742 | 0.9946449 |
| ENSG00000136247.14 | ZDHHC4 | -1.32447 | 0.7338281 | 0.438728 | 0.9783445 | 0.018747 | 0.4217249 | 0.65045 | 0.6929659 | -1.17327 | 0.002015865 | -0.568278 | 0.6885065 |
| ENSG00000136261.14 | BZW2 | -4.90083 | 0.02230429 | -0.509777 | 0.01245541 | 0.243085 | 0.1089188 | 0.525655 | 0.7479718 | -0.204316 | 0.001005217 | 0.00703 | 0.7235618 |
| ENSG00000136274.8 | NACAD | 1.000827 | 0.8212257 | 0.827885 | 0.708003 | 1.579526 | 0.4157337 | 1.18379 | 0.03905495 | 0.886381 | 0.5196843 | 0.08176 | 0.6829816 |
| ENSG00000136279.18 | DBNL | 1.242242 | 0.3544337 | -0.159808 | 0.9618421 | -0.915331 | 0.5290845 | 1.502633 | 0.0410256 | -0.302897 | 0.235777 | 0.435651 | 0.7737167 |
| ENSG00000136295.14 | TTYH3 | -0.658337 | 0.9449394 | -0.022676 | 0.7952779 | -1.054256 | 0.000320027 | 0.920385 | 0.001176646 | 1.869288 | 4.49E-05 | -0.358077 | 0.6885065 |
| ENSG00000136381.12 | IREB2 | 0.094703 | 0.9708532 | -0.434256 | 0.6527736 | -0.033653 | 0.5290845 | 1.503301 | 0.01934312 | 2.040587 | 0.01667537 | 2.160893 | 0.6517664 |
| ENSG00000136425.12 | CIB2 | 1.979263 | 0.6014343 | -0.352064 | 0.8263112 | -0.334839 | 0.5705225 | 2.115101 | 0.01774739 | 0.557667 | 0.1300713 | 1.325784 | 0.6885065 |
| ENSG00000136450.12 | SRSF1 | 0.746007 | 0.714717 | 0.388699 | 0.9283624 | 0.008501 | 0.3228353 | 0.846971 | 5.35E-05 | 0.827438 | 0.2440725 | -0.158438 | 0.6517664 |
| ENSG00000136451.8 | VEZF1 | -1.236009 | 0.8894962 | -0.352666 | 0.3651168 | 0.602239 | 0.009993063 | 1.324999 | 0.05310272 | 0.835319 | 0.1585606 | -0.753892 | 0.6243454 |
| ENSG00000136485.14 | DCAF7 | -1.088665 | 0.8867323 | -0.187325 | 0.2416476 | 0.847036 | 6.22E-05 | 1.067132 | 0.004603223 | 0.126895 | 0.9469705 | -0.294161 | 0.6885065 |
| ENSG00000136521.12 | NDUFB5 | 0.954012 | 0.1863043 | 0.067408 | 0.9047759 | 0.188808 | 0.8823128 | 0.204179 | 0.06228248 | -0.504447 | 0.01618838 | 0.578564 | 0.8088001 |
| ENSG00000136522.13 | MRPL47 | 0.329788 | 0.9887371 | 0.329726 | 0.9452791 | -0.180769 | 0.3087922 | -0.104219 | 0.01430575 | -0.154612 | 0.2880317 | 0.634068 | 0.5733051 |
| ENSG00000136527.17 | TRA2B | 0.684303 | 0.6214382 | -0.046053 | 0.6527736 | 0.556682 | 0.000460475 | 0.216455 | 0.1534232 | 0.546614 | 0.2880317 | -0.135481 | 0.6898338 |
| ENSG00000136536.14 | 7-Mar | 0.277206 | 0.3098608 | -0.076302 | 0.6527736 | 1.159007 | 0.01631626 | 1.427216 | 0.04798097 | 1.278562 | 0.06416308 | 1.176093 | 0.6885065 |
| ENSG00000136603.13 | SKIL | -0.921497 | 0.6214382 | 0.399282 | 0.6527736 | 0.695873 | 0.000688818 | 1.679497 | 0.000122804 | 0.746755 | 0.608741 | -2.030944 | 0.5344421 |
| ENSG00000136628.17 | EPRS | -0.075583 | 0.2990151 | 0.245763 | 1 | 0.705805 | 0.000744933 | 1.04689 | 0.2176454 | -0.009855 | 0.4440658 | 1.128254 | 0.8587735 |
| ENSG00000136636.12 | KCTD3 | -0.54157 | 0.714717 | 0.326287 | 0.7594702 | 1.039261 | 0.2818457 | 0.853214 | 0.01561965 | 2.324243 | 0.000305483 | -2.111138 | 0.9147227 |
| ENSG00000136682.14 | CBWD2 | -0.428289 | 0.8867323 | 1.07593 | 0.6527736 | 0.819625 | 0.01143805 | -0.876139 | 0.04310756 | -0.22787 | 0.7728137 | 1.368236 | 0.9258627 |
| ENSG00000136715.17 | SAP130 | 0.398301 | 0.7736255 | 0.016164 | 0.7420724 | 0.250799 | 0.1464093 | -0.150577 | 0.4936416 | 1.861187 | 0.006387211 | -0.576912 | 0.9147227 |
| ENSG00000136717.14 | BIN1 | -1.096415 | 0.6214382 | 0.410767 | 0.6527736 | 0.167745 | 0.9769181 | 0.164462 | 0.6864771 | 1.360361 | 0.03214838 | -2.309924 | 0.6576742 |
| ENSG00000136731.12 | UGGT1 | -0.467337 | 0.7083453 | 0.542118 | 0.8719537 | 0.466607 | 0.8360224 | 2.295708 | 0.000212025 | 0.009696 | 0.652246 | -1.168603 | 0.7904104 |
| ENSG00000136810.12 | TXN | 0.222203 | 0.8867323 | 0.005154 | 0.7869577 | -0.18611 | 6.02E-07 | -0.157955 | 0.003961012 | -0.481942 | 1.48E-06 | -0.2928 | 0.2440556 |
| ENSG00000136811.16 | ODF2 | 0.922395 | 0.6214382 | 0.746424 | 1 | 0.586539 | 0.005893275 | 0.514102 | 0.7858185 | -0.141682 | 0.9669702 | 2.467622 | 0.6885065 |
| ENSG00000136813.14 | KIAA0368 | 0.424991 | 1 | 0.083966 | 0.9766239 | 0.170941 | 0.7263718 | 0.689222 | 0.001350605 | 2.51964 | 0.225578 | -0.163069 | 0.9025407 |
| ENSG00000136819.14 | C9orf78 | -0.088433 | 0.6991852 | 0.108628 | 0.8338318 | 0.082323 | 1 | 0.735517 | 0.5627418 | -0.361342 | 0.03593768 | 0.6095 | 0.6829816 |
| ENSG00000136842.13 | TMOD1 | 0.237283 | 0.6214382 | -0.139741 | 0.6974771 | -3.522448 | 0.01760987 | 0.890262 | 0.6901888 | 1.378956 | 0.5196843 | 1.188354 | 0.8479431 |
| ENSG00000136848.16 | DAB2IP | -1.258521 | 1 | -1.323622 | 0.5953732 | -0.653773 | 0.8042144 | 1.452672 | 0.02217595 | 0.655222 | 0.08170657 | -0.370296 | 0.744192 |
| ENSG00000136866.13 | ZFP37 | -5.90468 | 0.4764974 | -0.391139 | 0.7383998 | 2.337441 | 0.00759642 | 2.415376 | 0.07244044 | 0.598319 | 0.6715871 | 2.747656 | 0.9276623 |
| ENSG00000136875.12 | PRPF4 | 0.868386 | 0.2520456 | 0.14858 | 0.6200225 | 0.173137 | 0.04184862 | 0.652814 | 0.4919368 | 1.015406 | 0.636096 | -0.997219 | 0.9147227 |
| ENSG00000136888.6 | ATP6V1G1 | -0.406565 | 0.6214382 | 0.061548 | 0.6527736 | -0.036103 | 0.7102786 | 0.0006 | 0.9741026 | -0.278887 | 0.004324077 | 0.052158 | 0.9296174 |
| ENSG00000136908.17 | DPM2 | 0.127289 | 0.6214382 | 0.201242 | 0.8819808 | 0.520863 | 0.9395155 | 0.900939 | 0.7680758 | -0.303048 | 0.003188443 | -1.24819 | 0.7385207 |
| ENSG00000136937.12 | NCBP1 | -2.381853 | 0.6214382 | -0.294652 | 0.6527736 | 1.021539 | 0.02382767 | -0.422969 | 0.1094014 | 0.945892 | 0.5674589 | 0.775336 | 0.6885065 |
| ENSG00000136938.8 | ANP32B | -1.254467 | 0.6214382 | -0.195857 | 0.5676172 | -0.137757 | 0.02173691 | -0.231196 | 0.1842832 | -0.120283 | 0.1182681 | -0.467316 | 0.3719182 |
| ENSG00000136942.14 | RPL35 | 0.094044 | 0.005603822 | -0.167437 | 0.3027683 | -0.101189 | 0.1296631 | -0.008575 | 0.4421404 | -0.187379 | 0.004497962 | -0.09503 | 0.4784299 |
| ENSG00000136950.13 | ARPC5L | -0.121318 | 0.6214382 | 0.541959 | 0.2374402 | -0.418139 | 0.000117345 | 0.294316 | 0.7939991 | 0.698853 | 0.5529159 | 0.754695 | 0.8265897 |
| ENSG00000137038.7 | TMEM261 | -0.516118 | 0.934454 | -0.048653 | 0.9534593 | -0.743626 | 0.000261718 | -0.178991 | 0.0595112 | 0.01156 | 0.02041193 | 0.196277 | 0.9264343 |
| ENSG00000137040.9 | RANBP6 | 0.399142 | 0.6214382 | -0.056962 | 0.9930412 | 0.760594 | 0.01352956 | 0.51957 | 0.2545636 | 2.06871 | 0.036961 | 0.302428 | 0.6885065 |
| ENSG00000137100.15 | DCTN3 | 0.520117 | 0.6214382 | 0.094135 | 0.8263112 | -0.445404 | 0.002254981 | 0.429164 | 0.7286701 | -0.035256 | 0.04251491 | 1.102651 | 0.6829816 |
| ENSG00000137133.10 | HINT2 | -0.182895 | 0.9803525 | -0.03735 | 0.6527736 | -0.949978 | 2.38E-06 | -0.963915 | 0.06977289 | -0.937155 | 0.1219535 | -0.708899 | 0.6885065 |
| ENSG00000137145.20 | DENND4C | -1.711618 | 0.8006286 | -0.923412 | 0.1191613 | 0.29642 | 0.5118662 | 2.11409 | 0.2696559 | 3.952664 | 0.000173452 | -4.545684 | 0.9147227 |
| ENSG00000137154.12 | RPS6 | -0.275886 | 0.928247 | 0.157442 | 0.5394427 | -0.014192 | 0.7437308 | -0.236857 | 9.73E-05 | -0.485281 | 5.09E-10 | 0.07153 | 0.7486771 |
| ENSG00000137161.16 | CNPY3 | -1.254494 | 0.6214382 | 0.074805 | 0.9790409 | 0.233375 | 0.5809753 | 0.493576 | 0.4124184 | -0.679394 | 0.04927588 | 1.635915 | 0.9875948 |
| ENSG00000137210.13 | TMEM14B | -0.200866 | 0.714717 | 0.319255 | 0.4333971 | -0.075372 | 0.02397417 | 0.344477 | 0.01576557 | -0.521744 | 0.00901316 | -0.746855 | 0.6885065 |
| ENSG00000137266.14 | SLC22A23 | 0.479243 | 0.6214382 | -0.162618 | 0.9104976 | 0.129149 | 0.5290845 | 0.615827 | 0.05738778 | 3.060124 | 0.04407967 | 0.377605 | 0.8553819 |
| ENSG00000137267.5 | TUBB2A | 0.062544 | 0.563505 | -0.265749 | 0.1015619 | 0.13851 | 0.5290845 | 0.027423 | 0.03737114 | -0.350598 | 0.000562657 | 0.015765 | 0.7020544 |
| ENSG00000137269.14 | LRRC1 | 0.226295 | 1 | 0.495026 | 0.6757195 | 0.433399 | 0.06474051 | 2.304148 | 0.03360361 | 1.081761 | 1 | -2.809226 | 0.6885065 |
| ENSG00000137285.9 | TUBB2B | -0.737298 | 0.03648071 | -0.237958 | 0.04760656 | -0.039001 | 0.8473149 | 0.163075 | 0.01153806 | -0.006875 | 0.3974318 | -0.07182 | 0.6829816 |
| ENSG00000137288.9 | UQCC2 | 0.036816 | 0.6214382 | -0.557578 | 0.1042612 | -1.133547 | 9.51E-08 | 0.154516 | 0.4744283 | -0.600771 | 0.04272514 | 0.977622 | 0.7499106 |
| ENSG00000137309.19 | HMGA1 | -0.36589 | 0.6776314 | -0.682078 | 0.009011566 | -0.590884 | 3.15E-05 | 0.00556 | 0.4919368 | -0.178214 | 0.4931295 | 0.289754 | 0.6185816 |
| ENSG00000137312.14 | FLOT1 | -0.880829 | 0.001974556 | 0.088745 | 1 | 0.0316 | 0.08371683 | -0.067604 | 0.3724881 | -0.224558 | 0.07818107 | 0.060774 | 0.6885065 |
| ENSG00000137337.14 | MDC1 | -0.986624 | 0.8867323 | -0.43382 | 0.9315666 | 0.971843 | 0.4229542 | 2.738634 | 0.0293315 | 1.81136 | 0.2589228 | -1.49112 | 0.9791458 |
| ENSG00000137409.18 | MTCH1 | 0.816119 | 0.007074581 | 0.200622 | 0.6527736 | -0.662589 | 0.08743214 | 0.673328 | 0.320553 | 0.134708 | 0.8163214 | 0.41007 | 0.9576568 |
| ENSG00000137449.15 | CPEB2 | 4.516897 | 0.6214382 | -0.067714 | 0.7269428 | -0.226361 | 0.9755686 | -0.67504 | 0.001085735 | 0.363749 | 0.1346549 | 0.491739 | 0.8965366 |
| ENSG00000137478.14 | FCHSD2 | -5.389605 | 0.8867323 | 0.334052 | 0.8472233 | 0.499418 | 0.4261006 | 0.772166 | 0.2326291 | 2.408929 | 0.000447908 | -0.749333 | 0.9576568 |
| ENSG00000137497.17 | NUMA1 | -1.839103 | 0.2299131 | 0.508821 | 0.4257977 | -0.401683 | 0.005966985 | 1.327825 | 2.60E-14 | 1.037601 | 0.001075709 | 0.168336 | 0.6829816 |
| ENSG00000137500.9 | CCDC90B | -0.173123 | 0.8253544 | -0.187674 | 0.7550531 | 0.436996 | 0.6140882 | -0.07735 | 0.04122336 | -0.940133 | 0.006378399 | -1.284951 | 0.7767294 |
| ENSG00000137574.10 | TGS1 | -1.684232 | 0.4764974 | 0.196488 | 0.6527736 | 0.446179 | 0.1674476 | 0.068075 | 0.9699324 | -0.712673 | 0.02226811 | 0.042041 | 0.8479431 |
| ENSG00000137714.2 | FDX1 | -1.891278 | 0.5048834 | 0.008381 | 0.9337922 | -0.397176 | 0.01787713 | 0.505052 | 0.4919368 | 0.912533 | 0.3762104 | 0.982249 | 0.6885065 |
| ENSG00000137726.16 | FXYD6 | 0.342023 | 0.6014343 | -0.572002 | 0.002588938 | 0.360254 | 0.1105011 | 0.543544 | 0.042941 | 0.677035 | 0.8785838 | -0.569552 | 0.186512 |
| ENSG00000137770.13 | CTDSPL2 | -0.871529 | 0.8371577 | 0.252927 | 1 | 0.784721 | 0.01975766 | 0.886517 | 0.07174536 | 0.970864 | 0.2630496 | 0.730476 | 0.9618424 |
| ENSG00000137776.16 | SLTM | -0.54688 | 0.4463293 | -0.264917 | 0.01857467 | 0.498464 | 0.01436828 | 0.553669 | 0.5833826 | 0.339957 | 0.8997439 | -0.125563 | 0.6885065 |
| ENSG00000137802.13 | MAPKBP1 | 0.076733 | 0.8732635 | -1.151833 | 0.4797514 | 0.277959 | 0.6287629 | 0.856768 | 0.004904019 | 2.052239 | 0.01257237 | 2.661581 | 0.5653062 |
| ENSG00000137815.14 | RTF1 | -0.253786 | 0.7217504 | 0.021954 | 0.979665 | 0.380574 | 0.0142718 | 1.159154 | 0.02071081 | 0.125212 | 0.296405 | -1.162584 | 0.8479431 |
| ENSG00000137817.16 | PARP6 | 0.466355 | 0.9585311 | 0.397417 | 0.6527736 | 1.224679 | 0.4602617 | 0.81337 | 0.01064038 | 1.353948 | 0.006124168 | 2.423473 | 0.4081608 |
| ENSG00000137818.11 | RPLP1 | -0.072215 | 0.2949764 | -0.4338 | 1.97E-13 | 0.113414 | 0.04447082 | 0.027282 | 0.7858185 | -0.236219 | 1.29E-05 | 0.210861 | 0.741293 |
| ENSG00000137857.17 | DUOX1 | -3.893566 | 0.8006286 | 1.1055 | 0.8263112 | 0.221624 | 0.7354314 | -2.298137 | 0.4919368 | -0.546036 | 1 |  |  |
| ENSG00000137872.15 | SEMA6D | 2.080193 | 0.4764974 | -0.307519 | 0.9719177 | 0.949347 | 0.5290845 | 0.218298 | 0.3534472 | 1.71895 | 0.001932804 | 2.218452 | 0.8949489 |
| ENSG00000137876.9 | RSL24D1 | 0.49672 | 0.6014343 | 0.231702 | 0.5394427 | 0.39518 | 0.001006486 | -0.069794 | 0.01555457 | 0.386845 | 0.6449337 | -1.123849 | 1 |
| ENSG00000137941.16 | TTLL7 | 0.714336 | 0.8867323 | -0.50572 | 0.4961426 | 0.813366 | 0.2273012 | -0.82318 | 0.02485579 | 1.673232 | 0.17142 | 0.507826 | 0.8479431 |
| ENSG00000137942.16 | FNBP1L | 0.918076 | 0.8587992 | 0.622355 | 0.7886006 | 0.865909 | 0.002954378 | 0.6696 | 0.3760083 | 0.250354 | 0.6351398 | -1.024738 | 0.6885065 |
| ENSG00000137965.10 | IFI44 | -1.412384 | 0.8006286 | 0.174068 | 0.9433711 | -0.175882 | 0.0332592 | 0.301649 | 0.4919368 | 0.031431 | 0.7040169 | 2.356912 | 0.6885065 |
| ENSG00000137992.14 | DBT | 2.502422 | 0.8006286 | -0.00852 | 0.9832381 | 1.172471 | 0.4523625 | 1.336519 | 0.02513874 | -0.172688 | 0.7771194 | -1.416631 | 0.8675096 |
| ENSG00000138018.17 | EPT1 | -0.794363 | 0.737082 | 0.183232 | 0.9315666 | 0.824157 | 0.8486387 | 0.655644 | 0.02914838 | 1.149495 | 0.573449 | 0.367076 | 0.7119977 |
| ENSG00000138032.20 | PPM1B | -1.905136 | 0.6214382 | 0.224433 | 0.7337542 | 0.868196 | 0.1182794 | 0.883845 | 0.166888 | 1.928419 | 0.04554435 | 0.783005 | 0.762714 |
| ENSG00000138050.14 | THUMPD2 | 0.44295 | 0.7591961 | -0.806331 | 0.186686 | 1.209273 | 0.002661796 | 0.646176 | 0.8284321 | 1.170379 | 1 | 0.494749 | 0.9526606 |
| ENSG00000138071.13 | ACTR2 | -0.179181 | 0.714717 | -0.039638 | 0.5940759 | 1.365573 | 5.94E-07 | 1.141382 | 0.1086422 | 0.928526 | 0.2019818 | 0.794699 | 0.6885065 |
| ENSG00000138073.13 | PREB | -0.15208 | 0.8014532 | 0.15871 | 0.9909831 | -1.576674 | 0.009261863 | 0.969294 | 0.03096616 | 1.611478 | 0.2055716 | -0.992672 | 0.5331811 |
| ENSG00000138081.19 | FBXO11 | 0.129925 | 0.5711449 | -0.170432 | 0.8217443 | 0.342151 | 0.04447082 | 2.637074 | 8.57E-05 | 1.389938 | 0.03056663 | 0.676243 | 0.6829816 |
| ENSG00000138095.18 | LRPPRC | -0.349347 | 0.9812137 | 0.045102 | 0.725047 | 0.563316 | 0.4035408 | 1.561578 | 0.01937453 | 0.818073 | 0.6185494 | -1.097058 | 0.6167027 |
| ENSG00000138162.18 | TACC2 | -0.379416 | 0.8867323 | -0.477819 | 0.4440278 | 0.106518 | 0.5290845 | 4.091455 | 0.000474841 | 2.290831 | 0.08578011 | -1.189132 | 0.4306015 |
| ENSG00000138166.5 | DUSP5 | -0.141556 | 1 | 0.136825 | 1 | 0.339059 | 0.5991171 | 1.39436 | 1.77E-05 | 3.788016 | 0.001037329 | 1.223925 | 0.985679 |
| ENSG00000138175.8 | ARL3 | -0.720578 | 0.6214382 | 0.148624 | 0.9030766 | 0.383877 | 0.5873317 | -0.305334 | 0.003130059 | -0.482858 | 0.195584 | 0.392736 | 0.6829816 |
| ENSG00000138190.16 | EXOC6 | 1.289733 | 0.6214382 | -0.443065 | 0.6527736 | 1.810889 | 0.0032438 | 0.235596 | 0.3434373 | 1.164723 | 1 | -1.550121 | 0.8597926 |
| ENSG00000138246.15 | DNAJC13 | -1.891212 | 0.6214382 | -0.844343 | 0.5953732 | 0.365423 | 0.03420804 | 1.188792 | 0.5523657 | 1.190282 | 0.7963151 | 0.738761 | 0.6885065 |
| ENSG00000138380.17 | CARF | 1.540508 | 1 | 0.127412 | 0.9222921 | 2.060812 | 0.7670335 | 3.72958 | 0.04186523 | 2.279907 | 0.8757379 | -2.157801 | 0.8949489 |
| ENSG00000138382.13 | METTL5 | 1.767277 | 0.4764974 | 0.044269 | 0.6427461 | 0.0711 | 0.5020747 | -0.075045 | 0.006878435 | 0.149044 | 0.4357714 | 0.197493 | 0.557022 |
| ENSG00000138385.15 | SSB | -0.498169 | 0.4764974 | -0.068504 | 0.5634345 | 0.40323 | 0.0396086 | 0.285453 | 0.5033921 | 0.202838 | 0.0013053 | 0.178906 | 0.4578379 |
| ENSG00000138398.15 | PPIG | -0.318838 | 0.6214382 | 0.190069 | 1 | 0.93217 | 7.21E-09 | 0.477146 | 0.3631319 | -0.859916 | 0.002564532 | 0.316463 | 1 |
| ENSG00000138399.17 | FASTKD1 | -0.520321 | 0.8006286 | -0.017507 | 0.6527736 | 0.571246 | 0.4157337 | 1.687509 | 0.001297763 | 0.697444 | 1 | -0.445482 | 1 |
| ENSG00000138434.16 | SSFA2 | -2.060915 | 0.6214382 | 0.026834 | 0.8779254 | 0.690051 | 0.02506782 | 0.749819 | 0.06044308 | 1.823402 | 0.0630294 | -0.216845 | 0.9532204 |
| ENSG00000138443.15 | ABI2 | -0.35329 | 0.7461932 | -0.064469 | 0.704767 | 0.778023 | 0.009910793 | 0.774196 | 7.64E-06 | 1.057336 | 0.0260978 | 0.205509 | 0.530077 |
| ENSG00000138448.11 | ITGAV | 0.268919 | 0.8216577 | -0.416475 | 0.4016878 | 0.227483 | 0.7021582 | 1.156986 | 0.000141101 | 1.1376 | 0.2601554 | 0.488241 | 0.6829816 |
| ENSG00000138495.6 | COX17 | 0.071148 | 0.7677129 | 0.218277 | 0.6755195 | -0.668236 | 1.38E-08 | -0.171003 | 0.001280431 | -0.021037 | 0.030568 | -0.297104 | 0.6885065 |
| ENSG00000138592.13 | USP8 | -0.598487 | 0.6014343 | 0.161719 | 0.9403597 | 1.015095 | 3.78E-07 | 0.284856 | 0.9012568 | 0.391851 | 0.5757394 | 1.426334 | 0.8479431 |
| ENSG00000138593.8 | SECISBP2L | -0.809424 | 0.9794907 | 0.019152 | 0.8237948 | 0.134154 | 0.08414334 | 0.934376 | 0.04645593 | 1.079519 | 0.4050482 | 0.643988 | 0.6556863 |
| ENSG00000138594.12 | TMOD3 | 0.08225 | 0.9591936 | -0.187582 | 0.6527736 | 0.985605 | 0.01026377 | 1.240585 | 0.6996942 | 0.005992 | 0.2751005 | -1.441298 | 0.6829816 |
| ENSG00000138606.19 | SHF | -0.799769 | 0.8006286 | -0.906058 | 0.6527736 | -0.392112 | 0.9140124 | 1.64015 | 0.162841 | 3.043737 | 0.002401816 | 3.224772 | 0.6885065 |
| ENSG00000138623.9 | SEMA7A | -1.561849 | 0.714717 | -0.680181 | 0.7289854 | 1.103365 | 0.9215126 | 0.14881 | 0.02076542 | 1.489472 | 0.5724789 | -1.023066 | 0.6885065 |
| ENSG00000138640.14 | FAM13A | 1.205064 | 0.6214382 | -0.162928 | 0.8263112 | 2.236735 | 0.04735417 | 0.516378 | 1 | 0.474868 | 0.2354226 | -0.965322 | 0.9270338 |
| ENSG00000138650.8 | PCDH10 | 2.721298 | 0.6014343 | -0.282451 | 0.7079982 | 0.372189 | 0.04447082 | 0.766211 | 0.6337475 | 1.030265 | 0.9443862 | -1.533123 | 0.9398376 |
| ENSG00000138660.11 | AP1AR | 0.37476 | 0.9282145 | 0.311128 | 0.6527736 | -0.174006 | 0.4870146 | 2.170801 | 0.003704982 | 1.167532 | 0.6355147 | 1.523986 | 1 |
| ENSG00000138668.18 | HNRNPD | -1.311517 | 2.91E-06 | 0.05546 | 0.704767 | -0.071912 | 0.5290845 | 0.178146 | 0.8420774 | 0.020055 | 0.2167378 | -0.028144 | 0.2967538 |
| ENSG00000138674.16 | SEC31A | -0.882435 | 0.6014343 | -0.110269 | 0.6527736 | 0.849714 | 1.99E-10 | 0.499946 | 0.9172514 | 0.559289 | 0.5436865 | -1.013758 | 0.9147227 |
| ENSG00000138688.15 | KIAA1109 | 1.913788 | 0.5711449 | 0.078444 | 0.6527736 | 0.787182 | 0.01702556 | 1.416558 | 0.002661105 | 1.516657 | 1 | -2.416152 | 0.359138 |
| ENSG00000138709.17 | LARP1B | -0.612533 | 0.6214382 | -0.172256 | 0.8688631 | 0.820811 | 0.03274039 | -0.864114 | 0.9083039 | 0.852302 | 0.6894088 | 2.4711 | 0.8528312 |
| ENSG00000138757.14 | G3BP2 | 0.789499 | 0.5135558 | 0.261942 | 0.9104976 | 1.109121 | 0.004240838 | 1.601217 | 0.002878546 | 0.681856 | 0.8171444 | 0.420023 | 0.8954041 |
| ENSG00000138758.11 | 11-Sep | 0.074157 | 0.8567515 | -0.061505 | 0.5953732 | 0.824886 | 3.50E-05 | 1.597992 | 0.1004331 | 0.095806 | 0.4840581 | -0.327395 | 0.9091709 |
| ENSG00000138760.8 | SCARB2 | 0.593937 | 0.563505 | 0.24919 | 0.6527736 | -0.301445 | 0.5784791 | 0.864428 | 0.02803104 | 1.17729 | 0.189278 | 0.142752 | 0.792724 |
| ENSG00000138764.13 | CCNG2 | -1.110174 | 1 | 0.201961 | 0.9329105 | -0.59158 | 0.9295516 | -0.130465 | 0.0298972 | 0.393189 | 1 | -0.309172 | 0.9326505 |
| ENSG00000138767.12 | CNOT6L | -0.261419 | 0.9332642 | -0.170759 | 0.6779949 | 2.310728 | 0.000594743 | 1.502054 | 0.008499454 | 1.310374 | 0.5196843 | 2.529993 | 0.6829816 |
| ENSG00000138777.19 | PPA2 | 0.097508 | 1 | 0.111042 | 0.9452791 | 0.205631 | 0.8473149 | -0.563419 | 0.001010299 | 0.435019 | 0.9865424 | -0.476506 | 0.9562356 |
| ENSG00000138814.16 | PPP3CA | 0.778533 | 0.3758557 | 0.034387 | 0.8101901 | 1.080359 | 0.004895182 | 0.181795 | 0.5271934 | 0.524955 | 0.3153504 | -0.269575 | 0.6885065 |
| ENSG00000138834.12 | MAPK8IP3 | -1.501518 | 0.2491762 | 0.436032 | 0.6527736 | 0.157513 | 0.905461 | 1.510896 | 0.01231478 | 1.619878 | 0.06513754 | -0.435392 | 1 |
| ENSG00000138867.16 | GUCD1 | -0.585063 | 1 | -0.225777 | 0.6527736 | 0.128242 | 0.8473149 | 1.781712 | 0.01089687 | 2.652626 | 0.2065601 | 0.52901 | 0.8479431 |
| ENSG00000139112.10 | GABARAPL1 | 0.060153 | 0.8597396 | 0.110199 | 1 | 1.260879 | 0.003313359 | -0.288604 | 0.5179377 | 0.62966 | 0.8021966 | 0.702179 | 0.6829816 |
| ENSG00000139116.17 | KIF21A | 0.77675 | 0.8969082 | 0.136877 | 0.5010139 | 0.565719 | 0.01090776 | 0.959409 | 2.93E-06 | 0.607013 | 0.5196843 | -1.706631 | 0.7419782 |
| ENSG00000139163.15 | ETNK1 | -1.958555 | 0.714717 | 0.130517 | 0.5010139 | 1.186342 | 0.007979364 | -0.163587 | 0.06675455 | 0.636915 | 1 | 1.986601 | 0.6885065 |
| ENSG00000139168.7 | ZCRB1 | -1.315176 | 0.002561164 | 0.071921 | 1 | 0.252773 | 0.8732897 | -0.478509 | 0.002048344 | -0.179581 | 0.02847549 | 0.776827 | 0.7352073 |
| ENSG00000139182.13 | CLSTN3 | 2.310883 | 0.8006286 | 0.467787 | 0.5953732 | -1.579424 | 0.4780312 | 2.153852 | 0.007052959 | 2.640051 | 0.01075306 | 1.492417 | 0.8391483 |
| ENSG00000139218.17 | SCAF11 | 0.698065 | 0.08143052 | 0.189687 | 0.7062105 | 1.260493 | 9.51E-06 | 0.701374 | 0.06932471 | 0.755766 | 0.5537753 | 0.347978 | 0.9994343 |
| ENSG00000139266.5 | 9-Mar | -0.068502 | 0.714717 | -0.030707 | 0.9918484 | 0.04577 | 0.6635072 | 0.551652 | 0.000270298 | 1.403042 | 1.53E-06 | -0.723633 | 0.947307 |
| ENSG00000139278.9 | GLIPR1 | -0.408378 | 1 | 0.896219 | 0.2011142 | -1.002749 | 0.1212319 | -0.313912 | 0.284075 | -1.246448 | 0.04490897 | 1.086152 | 0.936657 |
| ENSG00000139324.11 | TMTC3 | -0.673261 | 1 | -0.174341 | 0.8653838 | -1.115726 | 0.6401781 | 2.059836 | 0.01837465 | 1.111382 | 0.5070917 | -0.133363 | 0.661816 |
| ENSG00000139343.10 | SNRPF | -0.383995 | 1 | 0.005332 | 0.797408 | -0.192916 | 0.001032485 | -0.010857 | 0.14454 | -0.329805 | 0.003550296 | -0.922668 | 0.0947638 |
| ENSG00000139364.10 | TMEM132B | 0.873723 | 0.5873352 | -0.564788 | 0.2388066 | 0.602451 | 1 | 1.934261 | 1.08E-07 | 0.690069 | 0.0129887 | -0.993063 | 0.6885065 |
| ENSG00000139438.5 | FAM222A | -0.65181 | 0.714717 | 2.092018 | 0.8263112 | 0.926055 | 0.4712381 | 2.806 | 0.0149074 | 1.462153 | 0.05970807 | -1.923472 | 0.6885065 |
| ENSG00000139496.15 | NUP58 | -0.715329 | 0.7364985 | -0.621619 | 0.9031929 | 1.060076 | 0.000171547 | 1.149193 | 0.01087571 | 0.494091 | 0.9126976 | -0.429165 | 0.8708371 |
| ENSG00000139546.10 | TARBP2 | 1.753529 | 0.9183025 | 0.220965 | 0.9800936 | -1.079055 | 0.04616149 | -0.086511 | 0.6261611 | 0.867636 | 0.8526446 | -0.116485 | 0.8479431 |
| ENSG00000139613.11 | SMARCC2 | 0.165753 | 0.6039409 | 0.021818 | 0.6527736 | 0.85217 | 0.001640426 | 0.552536 | 0.5509566 | 0.817044 | 0.5196843 | -1.282967 | 0.6885065 |
| ENSG00000139620.12 | KANSL2 | -1.545265 | 0.563505 | -0.090924 | 0.4011918 | 0.085381 | 0.8523164 | -0.391375 | 0.7762366 | 0.76454 | 0.01065979 | 0.014468 | 0.6885065 |
| ENSG00000139624.12 | CERS5 | -1.321241 | 0.9678883 | 0.383381 | 0.6527736 | 0.038418 | 1 | 2.18546 | 0.02824263 | 1.693319 | 0.1278327 | -0.507809 | 0.9875948 |
| ENSG00000139625.12 | MAP3K12 | 0.484851 | 0.7777441 | 0.466601 | 0.5656503 | 0.798991 | 0.8071601 | 2.062088 | 3.69E-12 | 0.939491 | 0.8993413 | -0.851794 | 0.5653062 |
| ENSG00000139636.15 | LMBR1L | 2.775249 | 0.3098608 | 0.820663 | 0.6200225 | 0.184894 | 0.5053762 | 0.967135 | 0.2560809 | 1.136495 | 0.03368782 | -1.790278 | 0.821948 |
| ENSG00000139644.12 | TMBIM6 | 0.948642 | 0.03991323 | 0.156454 | 0.7968942 | 0.856444 | 5.13E-08 | 0.927385 | 0.3235822 | 0.659498 | 0.6842944 | 0.52241 | 0.6055076 |
| ENSG00000139645.9 | ANKRD52 | 0.149164 | 0.6214382 | 0.013584 | 0.6527736 | 0.008858 | 0.5290845 | 1.470569 | 0.000776635 | 1.876614 | 0.0963637 | 0.963121 | 0.8757616 |
| ENSG00000139668.8 | WDFY2 | -0.000514 | 0.8006286 | -0.267297 | 0.6107341 | 0.740841 | 0.1951987 | 3.240788 | 0.002169083 | 2.386539 | 0.0166049 | -2.088556 | 0.9425055 |
| ENSG00000139675.11 | HNRNPA1L2 | -1.906462 | 4.02E-06 | -0.079968 | 0.8513833 | 2.697079 | 5.84E-35 | -0.506207 | 1.82E-07 | 0.575555 | 0.8562558 | -0.440684 | 0.01006513 |
| ENSG00000139697.11 | SBNO1 | -0.078396 | 1 | 0.377496 | 0.8653057 | 1.518863 | 6.76E-07 | 2.232257 | 0.000104536 | 1.07925 | 0.5196843 | -1.012449 | 0.6885065 |
| ENSG00000139746.15 | RBM26 | -1.558959 | 0.3946735 | -1.10666 | 0.000277555 | 0.545262 | 0.000522619 | 1.549412 | 0.03820042 | 0.874567 | 0.2660195 | -1.635831 | 0.4499227 |
| ENSG00000139842.14 | CUL4A | 0.770951 | 0.6562396 | 0.144647 | 0.6527736 | -1.011753 | 0.8776881 | 1.263226 | 0.7120628 | 1.636014 | 0.03030386 | 1.659128 | 0.7767294 |
| ENSG00000139910.19 | NOVA1 | -1.813381 | 0.000114999 | -0.286572 | 0.3121527 | 0.533829 | 1.09E-07 | 0.386562 | 0.009224418 | 0.975663 | 0.1118369 | -0.301327 | 0.6490419 |
| ENSG00000139946.9 | PELI2 | 1.449104 | 1 | -2.599946 | 0.3171575 | 2.013427 | 0.7437308 | 1.241849 | 0.0215374 | 0.610007 | 0.5196843 | -0.636362 | 1 |
| ENSG00000139973.15 | SYT16 | -2.049496 | 0.7736255 | -3.40435 | 0.8263112 | 2.414943 | 0.9126812 | 1.021653 | 0.8622871 | 0.521791 | 0.04088424 | -2.786747 | 0.6885065 |
| ENSG00000139990.17 | DCAF5 | 0.356435 | 0.4692975 | 0.001135 | 0.8263112 | 0.931949 | 0.7437308 | 1.154565 | 0.02090518 | 1.241095 | 0.3380778 | -1.411402 | 0.6885065 |
| ENSG00000140262.17 | TCF12 | 0.350822 | 0.1057507 | -0.756412 | 0.02491789 | 0.488126 | 2.35E-12 | 0.670922 | 0.273789 | 0.252542 | 0.7242087 | -0.106109 | 1 |
| ENSG00000140263.13 | SORD | -0.578291 | 0.9589706 | -1.014058 | 0.5953732 | 0.249689 | 0.8774944 | -0.765332 | 0.00203379 | -0.391254 | 0.2404349 | -4.399585 | 0.7767294 |
| ENSG00000140264.19 | SERF2 | 0.806599 | 1.61E-08 | -0.085209 | 0.6527736 | -0.587764 | 3.75E-15 | 0.141026 | 0.1093889 | 0.087503 | 0.1118369 | 0.68094 | 0.9126404 |
| ENSG00000140265.12 | ZSCAN29 | -0.161218 | 0.9464322 | -0.22735 | 0.6809524 | 0.816872 | 0.7437308 | 1.137452 | 0.02980702 | 1.311062 | 0.04783049 | 1.630544 | 0.9618424 |
| ENSG00000140307.10 | GTF2A2 | -0.75548 | 0.9141126 | 0.164301 | 0.5010139 | -0.116421 | 0.01133138 | -0.169254 | 0.2362617 | 0.151779 | 0.01816971 | -0.159911 | 0.6829816 |
| ENSG00000140319.10 | SRP14 | -0.369626 | 0.9611425 | 0.043368 | 0.8263112 | -0.079485 | 0.001329646 | 0.369654 | 0.4919368 | 0.000346 | 0.00581755 | 0.587807 | 0.8207814 |
| ENSG00000140320.11 | BAHD1 | 0.226996 | 0.8435833 | -1.101828 | 0.0764189 | 0.119497 | 0.5990619 | 1.090177 | 0.04081668 | -0.137021 | 0.5196843 | 1.601526 | 0.8949489 |
| ENSG00000140332.15 | TLE3 | -0.095279 | 0.9282145 | 0.11604 | 0.7739688 | -0.058461 | 0.5674315 | 1.365333 | 0.001639525 | 0.827509 | 0.001883415 | 0.24749 | 0.8925226 |
| ENSG00000140391.14 | TSPAN3 | -0.142704 | 0.6214382 | 0.044446 | 0.7013839 | -0.657715 | 7.50E-07 | 0.675593 | 0.2676019 | 0.340803 | 0.812506 | -0.23539 | 0.8589196 |
| ENSG00000140406.3 | MESDC1 | 2.437703 | 0.8006286 | -0.109327 | 0.490383 | -0.346887 | 0.3512525 | 2.217222 | 0.04864249 | 1.453152 | 0.04464991 | 3.643205 | 0.6885065 |
| ENSG00000140416.19 | TPM1 | -0.008773 | 0.8587992 | 0.389428 | 0.4851871 | -0.884168 | 0.1211723 | 0.782539 | 0.04484561 | 0.678733 | 0.5196843 | -0.113626 | 0.6885065 |
| ENSG00000140443.13 | IGF1R | 0.121034 | 0.8689227 | 1.22214 | 0.3154845 | 0.117135 | 0.8100781 | 1.455825 | 6.83E-05 | -0.22931 | 0.4402432 | -2.807228 | 0.6885065 |
| ENSG00000140455.16 | USP3 | -0.286494 | 0.9768625 | 0.375629 | 0.8605299 | 1.423979 | 0.007413158 | 0.31565 | 0.6197784 | 0.672282 | 0.8718069 | -0.798373 | 0.8479431 |
| ENSG00000140521.11 | POLG | 0.910131 | 0.8408841 | -0.435416 | 0.7762225 | -0.917736 | 0.2339546 | 2.337434 | 0.001757507 | 1.250593 | 0.05602322 | -2.106888 | 0.8019827 |
| ENSG00000140564.10 | FURIN | 0.42908 | 0.8867323 | -0.630639 | 0.787908 | -1.63284 | 0.1455233 | 0.764113 | 0.1902046 | 3.844714 | 0.002032073 | 1.076499 | 1 |
| ENSG00000140575.12 | IQGAP1 | -1.099745 | 0.6214382 | -0.234839 | 0.8263112 | 0.742112 | 9.39E-06 | 1.294586 | 1 | -0.132572 | 0.2880317 | 1.57746 | 0.3406188 |
| ENSG00000140688.16 | C16orf58 | -0.061104 | 0.8240785 | 0.410483 | 0.9033552 | -0.18459 | 0.2558855 | -0.213191 | 0.7144938 | 1.257097 | 0.01558003 | -0.593811 | 0.9733982 |
| ENSG00000140750.16 | ARHGAP17 | -1.096145 | 0.6014343 | 1.233166 | 0.422236 | 1.617295 | 0.01142469 | 2.131526 | 0.228926 | -1.262059 | 0.9469636 | 1.247307 | 0.6829816 |
| ENSG00000140859.15 | KIFC3 | -2.106476 | 0.007523103 | -0.38254 | 0.6527736 | 0.7273 | 0.1903029 | 0.075101 | 0.4021022 | -0.341351 | 0.3974318 | 0.370254 | 0.7604024 |
| ENSG00000140905.9 | GCSH | -0.199607 | 0.6214382 | 0.365477 | 0.9104976 | -0.377202 | 0.002454097 | 0.421832 | 0.192668 | -0.385764 | 0.1213641 | -1.305623 | 0.6267144 |
| ENSG00000140931.19 | CMTM3 | -0.994563 | 0.006476816 | 0.288416 | 0.8937196 | -0.221064 | 0.8454005 | 0.941141 | 0.1526417 | 0.63966 | 0.09069599 | 0.207094 | 0.8479431 |
| ENSG00000140945.15 | CDH13 | -0.369391 | 1 | -0.538859 | 0.7841336 | 1.209246 | 0.5841679 | 0.088826 | 0.7834204 | 1.500814 | 1 |  |  |
| ENSG00000140983.13 | RHOT2 | -0.250698 | 0.529676 | -0.215538 | 1 | 0.139465 | 0.1538092 | 1.478747 | 3.70E-06 | 0.875768 | 0.009755487 | -0.143285 | 0.6885065 |
| ENSG00000140988.15 | RPS2 | -1.36382 | 6.06E-34 | -0.170376 | 0.01738075 | 0.073422 | 0.7652852 | -0.175044 | 0.00145517 | -0.123868 | 0.000495044 | -0.022975 | 0.3565625 |
| ENSG00000140990.14 | NDUFB10 | -0.173803 | 0.9444548 | 0.080581 | 0.6859453 | -0.306204 | 0.001057813 | 0.507367 | 0.3718594 | 0.477742 | 0.1535504 | 0.485199 | 0.6885065 |
| ENSG00000140995.16 | DEF8 | 0.56883 | 0.4004497 | 0.336894 | 1 | -0.412048 | 0.02101399 | 0.652017 | 0.7146594 | 0.305031 | 0.4492321 | -2.07689 | 0.6885065 |
| ENSG00000141027.20 | NCOR1 | 0.127395 | 0.714717 | -0.286005 | 0.1581444 | 0.783396 | 4.45E-06 | 1.118638 | 0.000155312 | 1.332288 | 0.06634352 | 1.179561 | 0.6885065 |
| ENSG00000141098.12 | GFOD2 | -1.463224 | 0.529676 | 0.844059 | 0.8703522 | -0.324447 | 0.9417779 | 2.150104 | 0.01539252 | 0.90245 | 0.1291723 | -0.981114 | 0.6885065 |
| ENSG00000141219.15 | C17orf80 | -1.034312 | 0.6214382 | -0.310809 | 0.8116276 | 1.130178 | 0.03804819 | -0.197997 | 0.4919368 | 1.138394 | 0.7146549 | -0.08849 | 0.6885065 |
| ENSG00000141232.4 | TOB1 | 0.778094 | 0.714717 | 0.50031 | 0.8263112 | -0.711129 | 0.9648047 | 0.685946 | 0.8095051 | 1.941016 | 0.004636219 | -2.659825 | 0.9290896 |
| ENSG00000141252.19 | VPS53 | -0.7534 | 0.3166974 | -0.044509 | 0.8939284 | 0.210712 | 0.3781922 | 1.512134 | 0.03256381 | 2.216542 | 0.000696277 | 1.171132 | 0.7532781 |
| ENSG00000141258.12 | SGSM2 | -1.532284 | 0.6014343 | 0.083133 | 0.9618682 | -0.542564 | 0.5108004 | 0.547304 | 0.2143686 | 1.601241 | 0.008567685 | 1.480957 | 0.7243667 |
| ENSG00000141279.15 | NPEPPS | -3.971255 | 0.7750119 | 0.291177 | 0.6527736 | 0.903769 | 0.02747485 | 1.537671 | 0.1648328 | 0.166856 | 0.2880317 | 2.209893 | 0.6885065 |
| ENSG00000141349.8 | G6PC3 | -0.107181 | 0.7152438 | -0.005868 | 0.5671096 | -0.872062 | 0.004806327 | -0.040084 | 0.6575769 | 0.25304 | 0.8205332 | 1.011042 | 0.8479431 |
| ENSG00000141367.11 | CLTC | 0.090953 | 0.7648042 | -0.053022 | 0.7449708 | 0.20317 | 0.002920147 | 1.235594 | 0.00045649 | 0.920776 | 0.02635043 | -0.092769 | 0.6885065 |
| ENSG00000141424.12 | SLC39A6 | -0.373758 | 1 | -0.266045 | 0.478664 | 0.647534 | 0.04230269 | 1.598621 | 0.01305884 | 0.74441 | 0.8915767 | -0.783127 | 0.5066477 |
| ENSG00000141446.10 | ESCO1 | -1.195515 | 0.6528194 | -0.161031 | 0.8238638 | 1.618897 | 4.63E-05 | -0.627704 | 0.7226722 | 1.94309 | 0.4310728 | -2.480811 | 0.9526606 |
| ENSG00000141447.16 | OSBPL1A | 0.484663 | 0.8006286 | -0.2068 | 0.7881421 | 0.758218 | 0.01061134 | -0.668928 | 1 | 1.371365 | 0.8049185 | 0.867093 | 0.9577321 |
| ENSG00000141503.15 | MINK1 | 0.160483 | 0.8006286 | 1.608356 | 0.0205677 | -1.109061 | 1 | 4.140031 | 0.162841 | 2.380488 | 0.0427337 | -2.537379 | 0.8883513 |
| ENSG00000141510.16 | TP53 | -0.303009 | 0.8006286 | -0.32807 | 0.3255429 | -0.492856 | 0.02030933 | 0.389871 | 1 | 0.793612 | 0.684036 | 2.156712 | 0.6200022 |
| ENSG00000141543.9 | EIF4A3 | -0.321508 | 0.1487841 | 0.154446 | 0.7647276 | -0.763665 | 1.46E-09 | 0.086356 | 0.1758005 | -0.138891 | 0.06767349 | 0.143506 | 0.7753492 |
| ENSG00000141552.17 | ANAPC11 | -0.039828 | 0.6214382 | -0.179628 | 0.2801475 | -0.118533 | 0.000844891 | -0.119737 | 0.000315064 | -0.424493 | 7.81E-06 | -0.519306 | 0.1846822 |
| ENSG00000141556.20 | TBCD | 2.083177 | 0.714717 | -0.287033 | 0.9563625 | -0.250056 | 0.3301939 | 1.361583 | 0.013996 | 1.218822 | 0.7762041 | 0.959846 | 0.6462656 |
| ENSG00000141568.19 | FOXK2 | -0.335115 | 0.8263393 | -1.201274 | 0.01245541 | 0.598887 | 0.7235224 | 0.91202 | 0.01114426 | -0.017403 | 0.07842487 | -0.209271 | 1 |
| ENSG00000141576.14 | RNF157 | -1.562521 | 0.005603822 | -0.5871 | 0.02412696 | -0.697593 | 1 | 0.76673 | 0.02699969 | 0.155507 | 0.8021966 | 1.112791 | 0.809678 |
| ENSG00000141580.15 | WDR45B | 0.107781 | 0.6214382 | 0.099568 | 0.6527736 | 0.350893 | 0.9836024 | -0.345593 | 0.2135278 | -0.477151 | 0.000208777 | -0.521527 | 0.9406771 |
| ENSG00000141582.14 | CBX4 | -2.78584 | 0.07216456 | 0.580725 | 0.920955 | -0.628854 | 0.7133511 | 2.050881 | 0.000313019 | 1.963568 | 2.11E-05 | -0.438628 | 0.6829816 |
| ENSG00000141639.11 | MAPK4 | -1.040107 | 1 | 0.437406 | 0.8600039 | -3.437313 | 0.02997543 | 0.417352 | 0.8984702 | -0.128514 | 0.6188988 | -1.335539 | 0.8479431 |
| ENSG00000141644.17 | MBD1 | 0.430648 | 0.6214382 | -0.244054 | 0.9616396 | 0.440464 | 0.6345169 | 1.176009 | 0.007378328 | 1.604614 | 0.3803925 | 0.148298 | 0.8663014 |
| ENSG00000141646.13 | SMAD4 | 0.134844 | 0.529676 | -0.076947 | 0.9069847 | 0.580965 | 0.09914996 | 1.223561 | 0.008024982 | 1.314126 | 0.000826124 | -0.953545 | 0.9144171 |
| ENSG00000141664.9 | ZCCHC2 | -0.441983 | 0.739831 | -0.517263 | 0.5362867 | 0.57559 | 0.6669178 | 1.740904 | 0.02785789 | 1.14406 | 0.5196843 | -0.59846 | 0.7767294 |
| ENSG00000141696.12 | P3H4 | -1.247289 | 0.6214382 | 0.068395 | 1 | -0.133116 | 0.04027253 | 1.38585 | 0.3206397 | 1.447306 | 0.7110301 | -1.816434 | 0.6885065 |
| ENSG00000141699.10 | FAM134C | 1.228822 | 0.8006286 | -0.080128 | 0.6527736 | 0.245374 | 0.9676705 | 1.489724 | 0.006455731 | 0.270985 | 0.3251443 | 1.165913 | 0.8257547 |
| ENSG00000141741.11 | MIEN1 | -2.87911 | 0.8587992 | 0.062073 | 0.804493 | -0.025067 | 0.4176494 | -0.231108 | 0.003144767 | 0.165397 | 0.0564468 | 0.026258 | 0.4578379 |
| ENSG00000141759.14 | TXNL4A | -0.844072 | 0.422882 | 0.133028 | 0.9856559 | -0.467123 | 2.50E-09 | 0.458334 | 0.1833938 | 0.376174 | 0.7377262 | 0.761662 | 0.8479431 |
| ENSG00000141837.18 | CACNA1A | 0.141357 | 0.9747122 | -1.009506 | 0.5761862 | 1.725849 | 0.9381042 | 1.338168 | 0.001070539 | 1.976273 | 7.12E-05 | 2.00757 | 0.9004786 |
| ENSG00000141858.11 | SAMD1 | -0.920722 | 0.8006286 | 0.416909 | 0.8962595 | -0.336041 | 0.3144442 | 0.121654 | 0.1343519 | 1.783397 | 4.27E-05 | -1.278088 | 0.84525 |
| ENSG00000141867.17 | BRD4 | -0.460449 | 1 | -0.18925 | 0.5732186 | 0.985345 | 1.91E-06 | 1.696767 | 0.000251844 | 1.151508 | 0.04407967 | 2.268069 | 0.6185816 |
| ENSG00000141873.10 | SLC39A3 | -3.550455 | 0.3166974 | -0.099507 | 0.86041 | -1.586723 | 5.89E-07 | -0.686797 | 0.02462981 | -0.563248 | 0.213644 | 0.848624 | 0.6885065 |
| ENSG00000141905.17 | NFIC | -0.025035 | 0.06849962 | -0.526374 | 0.002621603 | 0.425342 | 0.1432858 | 0.984242 | 0.006297858 | 1.285645 | 0.2880317 | -0.470696 | 0.9270789 |
| ENSG00000141956.13 | PRDM15 | 1.667673 | 0.8867323 | 1.422877 | 0.8465126 | -1.802693 | 0.7437308 | 1.337815 | 0.1293551 | 2.525067 | 0.03043237 | -2.808101 | 0.7767294 |
| ENSG00000141965.4 | FEM1A | -0.195051 | 0.5858981 | -0.419383 | 0.4492227 | -0.25989 | 0.7437308 | 0.561937 | 0.027258 | 1.60784 | 0.07956883 | -1.464566 | 0.6829816 |
| ENSG00000141971.12 | MVB12A | -0.684512 | 1 | -0.536753 | 0.64507 | -0.476374 | 0.001809352 | -0.3275 | 0.5860824 | 0.136933 | 0.5453018 | -0.967747 | 0.9208967 |
| ENSG00000141985.9 | SH3GL1 | 0.672401 | 0.5706649 | 0.124478 | 0.87788 | -0.763498 | 0.02097515 | 0.293552 | 0.4919368 | 1.315392 | 0.119277 | -0.030109 | 0.8479431 |
| ENSG00000142039.3 | CCDC97 | -0.079004 | 0.8006286 | -0.555055 | 0.703649 | -2.275995 | 0.1371748 | 1.483716 | 0.04072971 | 0.558184 | 0.584557 | 1.499284 | 0.7285797 |
| ENSG00000142089.15 | IFITM3 | -3.245441 | 0.563505 | -0.044467 | 0.8519642 | -1.348225 | 1.97E-22 | -0.852602 | 4.97E-08 | -1.643137 | 0.002668025 | -0.123169 | 0.9368176 |
| ENSG00000142156.14 | COL6A1 | 1.865401 | 0.6214382 | -0.423234 | 0.6433679 | -0.149043 | 0.5194179 | 0.181937 | 0.04615682 | 0.220865 | 1 | -0.08593 | 0.8768029 |
| ENSG00000142166.12 | IFNAR1 | -0.346455 | 0.6347687 | 0.214826 | 1 | -0.200295 | 0.02852264 | 1.375179 | 0.4043669 | 0.689358 | 0.9705634 | 1.130358 | 1 |
| ENSG00000142168.14 | SOD1 | -0.773375 | 0.3787871 | -0.260421 | 0.2238857 | -0.20525 | 0.2358121 | 0.06095 | 0.01034389 | 0.009802 | 0.2363864 | 0.247418 | 0.5653062 |
| ENSG00000142192.20 | APP | -0.22388 | 0.6214382 | 0.179479 | 0.8703522 | 0.137776 | 0.4064599 | 1.254353 | 6.44E-07 | 1.047961 | 0.4118087 | -0.444581 | 0.7672365 |
| ENSG00000142197.12 | DOPEY2 | -0.677433 | 1 | 0.290709 | 0.9772169 | 0.320683 | 0.5884821 | 0.241473 | 0.09071365 | 2.481912 | 0.03941972 | 0.314413 | 0.8479431 |
| ENSG00000142207.6 | URB1 | -0.930344 | 0.6214382 | -0.126028 | 0.5010139 | -0.900099 | 0.3871766 | 1.828862 | 0.01514072 | 1.799273 | 0.9275881 | 1.114945 | 0.8479431 |
| ENSG00000142235.8 | LMTK3 | -0.72911 | 0.6214382 | 0.299773 | 0.80335 | -1.380024 | 0.3901219 | 2.897706 | 0.000645918 | -0.067115 | 0.09409698 | 1.991977 | 0.5733051 |
| ENSG00000142303.13 | ADAMTS10 | 1.094323 | 0.8587992 | 0.557915 | 0.9104976 | 5.38407 | 0.3276184 | 1.798742 | 0.3570293 | 3.626877 | 0.02652126 | -3.622665 | 0.6517664 |
| ENSG00000142409.5 | ZNF787 | 2.553567 | 0.4764974 | -1.280229 | 0.05116306 | 0.681283 | 0.5290845 | 1.337955 | 0.009082928 | 2.856698 | 8.95E-05 | 1.419386 | 0.8956025 |
| ENSG00000142444.6 | C19orf52 | 0.739301 | 0.6214382 | 0.114266 | 0.913312 | -1.528884 | 0.001904645 | 1.545319 | 0.1436444 | 2.239305 | 0.2105971 | 1.681143 | 0.9704304 |
| ENSG00000142453.11 | CARM1 | 0.659589 | 0.8432999 | -0.072139 | 0.6527736 | 0.373862 | 0.9143308 | 1.525538 | 0.02970607 | 0.523703 | 0.4225841 | -1.185318 | 0.9147227 |
| ENSG00000142507.9 | PSMB6 | -0.116252 | 0.6214382 | 0.063644 | 1 | -0.107982 | 0.06659945 | -0.197078 | 6.71E-05 | -0.571319 | 3.04E-08 | 0.010018 | 0.05044736 |
| ENSG00000142541.16 | RPL13A | -0.939849 | 0.000601649 | 0.355412 | 0.000150637 | 0.095637 | 0.07970508 | 0.099737 | 0.7144938 | 0.166022 | 0.376032 | 0.278535 | 0.8384921 |
| ENSG00000142599.17 | RERE | -0.701246 | 1 | -0.04168 | 0.6527736 | 0.438633 | 1 | 2.06913 | 0.001070539 | 1.269543 | 0.00018959 | 4.410277 | 0.6885065 |
| ENSG00000142669.13 | SH3BGRL3 | -0.03654 | 0.7736255 | -0.252087 | 0.4011918 | -0.298706 | 3.90E-06 | -0.200347 | 0.00077914 | -0.113377 | 0.000366611 | 0.108948 | 0.5008296 |
| ENSG00000142676.12 | RPL11 | -0.756893 | 0.04094829 | -0.15083 | 0.2784584 | 0.261321 | 6.68E-05 | -0.041134 | 0.01572454 | -0.235643 | 4.70E-06 | 0.095822 | 0.9146615 |
| ENSG00000142684.7 | ZNF593 | -0.131893 | 0.8562417 | -0.236819 | 0.8818962 | -0.048923 | 0.000535742 | -0.006141 | 0.04803716 | -0.552278 | 0.07584831 | -0.514906 | 0.8756022 |
| ENSG00000142687.17 | KIAA0319L | -0.371256 | 0.824666 | -0.400176 | 0.6527736 | 0.001544 | 0.3312075 | 0.940096 | 0.04823745 | 1.292834 | 0.01157352 | 1.147621 | 0.8479431 |
| ENSG00000142751.14 | GPN2 | -0.370125 | 1 | -0.21144 | 0.7059293 | -0.098423 | 0.7638631 | 1.032747 | 0.04796113 | 0.578624 | 0.1407934 | 1.673994 | 0.2321124 |
| ENSG00000142864.14 | SERBP1 | -0.466712 | 0.6214382 | -0.3392 | 0.007131869 | 0.559131 | 2.91E-08 | 0.247191 | 0.3874059 | 0.174059 | 0.01099018 | -0.405167 | 0.3394425 |
| ENSG00000142871.15 | CYR61 | -1.672459 | 0.4506214 | 0.366464 | 6.04E-06 | -0.224893 | 0.04944273 | 0.435024 | 0.9230023 | -0.7229 | 0.8795727 | -1.534123 | 0.6829816 |
| ENSG00000142875.19 | PRKACB | -0.291825 | 1 | -0.204339 | 0.7150943 | 1.379723 | 0.03020129 | -0.075005 | 0.1248486 | -0.206989 | 1 | -0.40316 | 0.9441995 |
| ENSG00000142937.11 | RPS8 | -0.370294 | 0.6014343 | -0.072928 | 0.6200225 | 0.298555 | 1.25E-06 | -0.181571 | 0.00045875 | 0.17048 | 0.5196843 | 0.115187 | 0.6885065 |
| ENSG00000142945.12 | KIF2C | -1.241878 | 0.6214382 | -0.200439 | 0.4011918 | -0.066354 | 0.1778406 | -0.005935 | 0.6936897 | 1.837691 | 1.84E-06 | -0.642663 | 0.6004498 |
| ENSG00000143033.17 | MTF2 | 0.626672 | 0.728867 | 0.351895 | 0.6527736 | 1.043509 | 1.16E-06 | 0.372524 | 0.1704381 | 0.57795 | 0.7344498 | -0.433698 | 0.6829816 |
| ENSG00000143079.14 | CTTNBP2NL | -3.394312 | 0.1835496 | 0.28719 | 0.8640615 | 0.675522 | 0.04925267 | 1.22234 | 0.2113251 | 0.951505 | 0.5863663 | -0.101471 | 0.9576568 |
| ENSG00000143093.14 | STRIP1 | 2.245106 | 0.6214382 | -0.461539 | 0.6527736 | 0.229182 | 0.5290845 | 0.2465 | 0.7794319 | 3.587368 | 1.62E-05 | 0.341041 | 0.8309897 |
| ENSG00000143106.12 | PSMA5 | 0.105037 | 0.8731872 | 0.051371 | 0.9433181 | 0.00461 | 0.3757247 | -0.095863 | 0.00608761 | -0.103161 | 0.1845956 | 0.372457 | 0.8309897 |
| ENSG00000143126.7 | CELSR2 | -0.476003 | 0.8587992 | -0.022917 | 0.1256222 | 0.305085 | 0.6163624 | 0.095855 | 0.01911511 | 1.531481 | 0.001280028 | 1.475031 | 0.7553807 |
| ENSG00000143157.11 | POGK | -0.039538 | 0.8006286 | 0.130427 | 0.8101097 | 0.084365 | 0.1911841 | 2.092412 | 2.01E-07 | 0.664185 | 0.1918853 | -1.195622 | 0.6885065 |
| ENSG00000143158.10 | MPC2 | -0.309533 | 0.7090887 | -0.057328 | 0.6527736 | -0.215954 | 0.1253014 | -0.222871 | 0.1036641 | -0.232607 | 0.01491851 | 0.321502 | 0.8949489 |
| ENSG00000143162.7 | CREG1 | -1.644787 | 0.6214382 | 1.081829 | 0.00078029 | -1.195029 | 0.2348326 | 1.72391 | 0.05560288 | 1.535091 | 0.3974318 | 1.238938 | 0.773978 |
| ENSG00000143183.16 | TMCO1 | -2.586555 | 0.9975232 | 0.206084 | 0.8101901 | 0.092943 | 0.5290845 | 0.312121 | 0.2851303 | -0.067336 | 0.008295065 | 0.362339 | 0.8315889 |
| ENSG00000143190.21 | POU2F1 | -1.331759 | 0.6014343 | -0.53843 | 0.5394427 | 0.37315 | 0.08493438 | 0.291789 | 0.02615249 | 2.074582 | 0.002340489 | -0.056194 | 0.8949489 |
| ENSG00000143198.12 | MGST3 | -0.235095 | 1 | -0.002597 | 0.6795004 | -0.42789 | 0.04579444 | -0.463238 | 3.89E-06 | 0.179681 | 0.3928351 | -0.542825 | 0.4650016 |
| ENSG00000143207.19 | RFWD2 | -0.979388 | 0.6434318 | 0.144986 | 0.6527736 | 0.980461 | 0.6288509 | 1.52755 | 0.02262926 | 1.468625 | 0.6214624 | 0.398389 | 0.960853 |
| ENSG00000143256.4 | PFDN2 | -0.062767 | 0.4406957 | -0.143273 | 0.2876783 | -0.285287 | 7.61E-07 | 0.179105 | 0.1122665 | -0.26105 | 0.0166049 | 0.676058 | 0.7767294 |
| ENSG00000143258.15 | USP21 | 0.864773 | 0.6214382 | -0.178226 | 0.8657239 | -0.859437 | 0.7145461 | 1.761342 | 0.1766588 | 2.392384 | 0.0389724 | -1.991291 | 0.8479431 |
| ENSG00000143314.12 | MRPL24 | 0.12604 | 0.6214382 | 0.139742 | 0.6945593 | -0.52846 | 0.000195417 | -0.040341 | 0.1134309 | -1.250859 | 0.005440811 | -1.534926 | 0.530077 |
| ENSG00000143315.6 | PIGM | -1.967071 | 0.6214382 | -0.146591 | 0.6936532 | 1.694247 | 0.1576355 | 1.719989 | 0.002674322 | 1.955053 | 0.1476257 | 2.512305 | 0.8877351 |
| ENSG00000143321.18 | HDGF | -0.50552 | 0.9907591 | -0.062221 | 0.5108497 | 0.789528 | 0.04874163 | 0.301443 | 0.4919368 | 0.992099 | 0.00533696 | -0.376194 | 0.834639 |
| ENSG00000143322.19 | ABL2 | 0.109526 | 0.845546 | -0.377526 | 0.5394427 | 0.474188 | 0.6453005 | 1.957088 | 0.001018841 | 0.175582 | 0.5196843 | 1.897163 | 0.9228374 |
| ENSG00000143368.9 | SF3B4 | -1.082295 | 0.08143052 | -0.276841 | 0.6527736 | 0.081191 | 0.238948 | -0.975762 | 0.000159811 | 0.48817 | 0.5407965 | -0.415242 | 0.8353355 |
| ENSG00000143384.12 | MCL1 | -0.827369 | 0.6214382 | -0.085709 | 0.2801475 | 0.008563 | 0.5505943 | 1.253685 | 2.01E-07 | 1.79136 | 0.002269503 | -0.421073 | 0.8479431 |
| ENSG00000143398.19 | PIP5K1A | 0.33223 | 0.9821531 | 0.367429 | 0.5608611 | -0.057272 | 0.1991342 | -0.034612 | 3.54E-05 | 0.076245 | 0.1169767 | 0.452258 | 1 |
| ENSG00000143420.17 | ENSA | 0.518741 | 0.4584886 | -0.347275 | 0.4987653 | 0.24593 | 0.5115748 | 0.756872 | 0.01318206 | 0.574655 | 0.7377262 | -0.085559 | 0.6910517 |
| ENSG00000143434.15 | SEMA6C | 0.128129 | 0.8587992 | -0.345138 | 0.6527736 | 1.033132 | 1 | 0.702683 | 0.2696559 | 3.09499 | 0.003393002 | 0.232776 | 0.7501684 |
| ENSG00000143442.21 | POGZ | 0.212934 | 0.8867323 | 0.041181 | 0.8703522 | 0.84009 | 0.01370691 | 0.886941 | 6.99E-05 | 0.48473 | 0.07922936 | 0.910344 | 1 |
| ENSG00000143443.9 | C1orf56 | 0.611506 | 0.4764192 | 0.203488 | 0.6527736 | 0.562977 | 0.9265011 | 0.864689 | 0.001300776 | 1.122749 | 0.8649429 | -0.743547 | 0.9918431 |
| ENSG00000143458.11 | GABPB2 | -1.545283 | 0.1835496 | 0.455687 | 1 | 0.225779 | 0.5645295 | 1.312593 | 0.007198196 | 1.638885 | 0.6427258 | 2.96118 | 1 |
| ENSG00000143514.16 | TP53BP2 | -1.782294 | 0.1835496 | -0.432135 | 0.04469129 | 0.599481 | 0.08140955 | 1.124534 | 0.3547738 | -1.108122 | 0.5196843 | 0.954636 | 0.8949489 |
| ENSG00000143515.16 | ATP8B2 | -0.250993 | 0.6796903 | 0.185493 | 0.6527736 | 0.225174 | 0.9757648 | 1.28296 | 0.03756797 | 0.98855 | 0.5196843 | 0.7505 | 0.8876567 |
| ENSG00000143543.14 | JTB | 0.006082 | 0.6169321 | 0.086225 | 0.9198439 | -0.708999 | 1.32E-07 | 0.047011 | 0.6855744 | 0.276029 | 0.6112598 | -0.212007 | 0.3820098 |
| ENSG00000143549.19 | TPM3 | 0.696663 | 0.02641092 | 0.234132 | 1 | 0.446794 | 0.000646362 | 0.541302 | 0.8337841 | -0.06077 | 0.19459 | 0.135542 | 0.6206888 |
| ENSG00000143553.10 | SNAPIN | 1.316539 | 0.4392896 | 0.410955 | 0.8549935 | -0.381033 | 0.012381 | -0.321987 | 0.6357772 | 0.11952 | 0.5196843 | 0.635185 | 1 |
| ENSG00000143612.18 | C1orf43 | -0.910395 | 0.06849962 | 0.244976 | 0.6527736 | -0.212752 | 0.2723898 | -0.19714 | 0.1557388 | -0.401704 | 3.65E-05 | -0.576631 | 0.5066477 |
| ENSG00000143614.8 | GATAD2B | 0.337878 | 0.9047689 | -0.06359 | 0.5971742 | 0.818482 | 0.004168428 | 1.011673 | 4.13E-06 | 1.50369 | 2.07E-05 | 0.436549 | 0.8467036 |
| ENSG00000143630.9 | HCN3 | -1.943681 | 0.529676 | -2.272095 | 0.5953732 | 0.842689 | 0.907132 | 1.010102 | 0.4109963 | 2.033748 | 0.04969669 | 6.296267 | 0.1224713 |
| ENSG00000143669.13 | LYST | -0.059217 | 0.8006286 | -0.098821 | 0.8815325 | 0.461166 | 0.008254603 | 0.968781 | 0.07107542 | 1.008683 | 0.8485911 | -0.148551 | 0.9147227 |
| ENSG00000143702.15 | CEP170 | 0.493983 | 0.624814 | -0.609317 | 0.4987653 | 0.550316 | 0.000950538 | 0.933146 | 0.0410256 | 0.866861 | 0.8323733 | 0.223416 | 0.6885065 |
| ENSG00000143727.15 | ACP1 | -0.841722 | 0.8673742 | 0.139768 | 0.9342002 | -0.261197 | 0.3695136 | 0.556739 | 0.3033815 | -0.590052 | 0.03730835 | 0.792974 | 0.8105922 |
| ENSG00000143742.12 | SRP9 | 0.275673 | 0.2325976 | 0.306662 | 0.2374402 | 0.064839 | 0.4064599 | -0.143083 | 0.02984614 | -1.064713 | 8.86E-11 | -0.857121 | 0.4629479 |
| ENSG00000143751.9 | SDE2 | -0.014668 | 0.6214382 | 0.620326 | 0.8263112 | 0.623865 | 0.2150698 | -0.073992 | 0.7144938 | 1.360268 | 0.04137976 | 0.414535 | 0.8145966 |
| ENSG00000143756.11 | FBXO28 | -1.301016 | 0.934454 | 0.171738 | 1 | -0.946491 | 0.776456 | 0.681712 | 0.01087494 | 0.854347 | 0.185325 | -0.773668 | 0.9576568 |
| ENSG00000143761.13 | ARF1 | 0.577206 | 0.6014343 | -0.057741 | 0.5394427 | 0.362226 | 7.69E-08 | 0.328049 | 0.1016405 | 0.878814 | 0.3690304 | 0.51783 | 0.9147227 |
| ENSG00000143771.11 | CNIH4 | -0.472162 | 0.6214382 | 0.0283 | 0.8879683 | -0.480582 | 0.005903382 | -0.277852 | 0.002110492 | 0.085253 | 0.2744651 | 0.184735 | 0.9576568 |
| ENSG00000143772.9 | ITPKB | -3.830267 | 0.04197057 | 0.103465 | 0.5350015 | -1.640671 | 0.4170444 | 0.334191 | 1 | 3.60864 | 0.5196843 | -0.309128 | 0.8479431 |
| ENSG00000143774.16 | GUK1 | -0.307639 | 0.8055273 | 0.228081 | 0.5814384 | -0.551956 | 5.86E-12 | -0.193327 | 5.45E-08 | 0.076139 | 0.1099632 | 0.392238 | 0.6829816 |
| ENSG00000143776.18 | CDC42BPA | -0.308941 | 0.9059623 | 0.316293 | 1 | 0.814574 | 0.001733944 | 1.613995 | 0.06618271 | 0.206112 | 0.1858395 | -0.276805 | 0.9539924 |
| ENSG00000143793.12 | C1orf35 | 0.007748 | 1 | -0.237616 | 0.1834552 | -0.248179 | 0.02619358 | -0.841027 | 0.1000079 | -0.118199 | 0.6245743 | -0.0959 | 0.6829816 |
| ENSG00000143797.11 | MBOAT2 | 0.281115 | 0.4541625 | 0.229998 | 1 | 0.72013 | 0.01106777 | 1.762519 | 0.006155218 | 0.67559 | 0.3974318 | -0.269857 | 0.6885065 |
| ENSG00000143799.12 | PARP1 | -0.008097 | 0.8470618 | -0.232981 | 0.02497202 | 0.333094 | 0.2348326 | 0.394552 | 0.613025 | -0.212682 | 0.08223235 | -0.124488 | 0.6885065 |
| ENSG00000143811.16 | PYCR2 | -0.415103 | 0.714717 | 0.102084 | 0.681763 | -0.624109 | 0.009692558 | 0.493833 | 0.8645693 | 0.20504 | 0.8496491 | -1.509614 | 0.961839 |
| ENSG00000143815.14 | LBR | 0.605553 | 0.9660565 | 0.137133 | 0.7079982 | 1.211773 | 0.000685504 | 1.463868 | 0.001953302 | 0.987289 | 0.9588216 | 0.781452 | 0.9037465 |
| ENSG00000143870.12 | PDIA6 | 0.675813 | 0.09713735 | 0.10832 | 0.7079982 | 0.346452 | 0.09093274 | 0.061489 | 0.2696559 | -0.795772 | 4.37E-07 | -0.669833 | 0.4815789 |
| ENSG00000143878.9 | RHOB | 0.673712 | 0.06126222 | 0.462682 | 0.4281695 | 0.664282 | 0.7467693 | 2.062207 | 3.13E-05 | 1.534281 | 0.01390676 | -0.313218 | 0.9726757 |
| ENSG00000143924.18 | EML4 | 0.257453 | 0.824666 | -0.029907 | 0.5499727 | -0.169399 | 0.4103761 | 1.830725 | 0.01341412 | 0.781739 | 0.565496 | -0.511827 | 0.9705209 |
| ENSG00000143933.16 | CALM2 | 0.132769 | 0.03176252 | 0.18177 | 0.4487482 | -0.189933 | 0.05094688 | 0.159945 | 0.2869336 | -0.311801 | 3.75E-06 | 0.022166 | 0.5203855 |
| ENSG00000143942.4 | CHAC2 | -1.886949 | 0.6214382 | 0.198241 | 1 | -0.502004 | 0.2817378 | -1.497911 | 0.04997317 | -0.770057 | 0.1401461 | 0.794316 | 0.8949489 |
| ENSG00000143947.12 | RPS27A | -0.115624 | 0.6214382 | 0.056159 | 0.6645701 | 0.122259 | 0.1982205 | -0.114713 | 0.04737357 | -0.304584 | 9.54E-07 | -0.189167 | 0.3358843 |
| ENSG00000143977.13 | SNRPG | -0.078546 | 0.4318827 | 0.13545 | 0.8146081 | -0.444036 | 1.45E-06 | -0.005003 | 0.08103448 | -0.190722 | 0.005714101 | -0.269282 | 0.1570836 |
| ENSG00000144028.14 | SNRNP200 | 0.171954 | 0.8576226 | -0.082749 | 0.5321133 | 0.015979 | 1 | 1.143813 | 4.01E-07 | 1.722102 | 5.23E-06 | 0.950737 | 0.9783488 |
| ENSG00000144034.14 | TPRKB | -0.488475 | 0.858385 | 0.546164 | 0.5953732 | -0.110096 | 0.3095119 | 0.449944 | 0.335352 | -1.089465 | 0.002998326 | -0.546339 | 0.6885065 |
| ENSG00000144036.15 | EXOC6B | 0.14867 | 1 | -0.674656 | 0.6218841 | 0.66408 | 0.5290845 | 0.993077 | 0.008388443 | 0.752261 | 0.5196843 | -1.862184 | 0.9801417 |
| ENSG00000144043.11 | TEX261 | 1.28007 | 0.6214382 | -0.332102 | 0.8703522 | -0.061611 | 0.7437308 | 1.757002 | 0.000284065 | -0.796344 | 0.8795727 | -1.40464 | 0.7767294 |
| ENSG00000144136.10 | SLC20A1 | 0.542136 | 0.6214382 | -0.818678 | 0.3191072 | 0.53705 | 0.6273532 | 1.402948 | 0.006838718 | 1.904183 | 0.004259492 | -0.359678 | 0.8469357 |
| ENSG00000144218.18 | AFF3 | 0.015703 | 0.8867323 | 0.11526 | 0.9798987 | 0.441973 | 0.1827981 | 1.859262 | 0.009064554 | 0.990408 | 0.4081362 | 0.746195 | 0.8949489 |
| ENSG00000144224.16 | UBXN4 | -1.359632 | 0.07604642 | 0.385559 | 0.5835893 | 1.071046 | 1.63E-12 | 0.116102 | 0.7396717 | 0.30177 | 0.02806531 | -1.468737 | 0.1928225 |
| ENSG00000144231.10 | POLR2D | -0.876878 | 0.6214382 | -0.195109 | 0.9078096 | -0.117437 | 0.7455873 | 1.350823 | 9.70E-08 | 1.442644 | 0.003137382 | -0.493773 | 0.6885065 |
| ENSG00000144283.21 | PKP4 | 1.331902 | 0.6214382 | -0.127144 | 0.9385836 | 0.599959 | 0.5872968 | 0.952152 | 0.007368882 | 2.693777 | 0.00707925 | 0.15846 | 0.6185816 |
| ENSG00000144369.12 | FAM171B | 0.325969 | 0.71468 | -0.279838 | 0.6685387 | 0.441287 | 0.006091879 | 0.129185 | 0.9595908 | 0.387937 | 0.9636876 | -0.430744 | 0.8020906 |
| ENSG00000144381.16 | HSPD1 | 0.083255 | 0.6214382 | -0.098009 | 0.7039714 | 0.591216 | 1.65E-16 | 0.501329 | 0.550044 | 0.124055 | 0.01083921 | -0.459754 | 0.02030807 |
| ENSG00000144401.14 | METTL21A | 0.375257 | 0.6473062 | -1.789495 | 0.04771739 | -0.007039 | 0.4876164 | -0.348384 | 0.3428367 | 1.007942 | 1 | 2.576536 | 0.8578487 |
| ENSG00000144406.18 | UNC80 | -1.293181 | 0.714717 | -1.455195 | 0.4943499 | 0.218305 | 0.4928603 | 3.218602 | 0.00101636 | 2.716644 | 0.2880317 | 2.384901 | 0.8949489 |
| ENSG00000144485.10 | HES6 | -0.613513 | 0.529676 | -0.129935 | 1 | 0.480036 | 0.6400305 | 0.561063 | 0.9327635 | 2.118487 | 4.59E-08 | -1.003038 | 0.6042097 |
| ENSG00000144566.10 | RAB5A | 0.090904 | 0.8698122 | -0.024455 | 0.9330607 | 0.708209 | 0.01904784 | 1.141202 | 0.04837687 | 0.812266 | 0.5196843 | 0.942031 | 0.96412 |
| ENSG00000144674.16 | GOLGA4 | -1.427861 | 0.2423804 | 0.024375 | 0.5953732 | 1.126971 | 1.73E-12 | 1.689869 | 0.00144579 | 0.774048 | 0.4111602 | 0.534804 | 0.7979751 |
| ENSG00000144711.13 | IQSEC1 | -1.196246 | 0.7736255 | 0.076936 | 0.6527736 | -1.502323 | 0.7026259 | 1.234361 | 0.0326712 | 1.427214 | 0.4407481 | 3.161382 | 1 |
| ENSG00000144713.12 | RPL32 | -0.045699 | 0.4562592 | -0.020834 | 0.6527736 | 0.032103 | 1 | -0.141863 | 0.01260208 | -0.241075 | 1.38E-05 | 0.232716 | 0.5137734 |
| ENSG00000144724.18 | PTPRG | -1.70322 | 0.6214382 | -0.213553 | 0.5953732 | 0.617677 | 0.5619923 | 2.170495 | 8.92E-05 | 0.552686 | 0.7253092 | -1.892808 | 0.5733051 |
| ENSG00000144746.6 | ARL6IP5 | -0.125138 | 0.7134849 | 0.206768 | 0.9296311 | 0.039845 | 0.282157 | -0.231916 | 0.001790909 | -0.134692 | 0.05194075 | -0.329181 | 0.6185816 |
| ENSG00000144747.14 | TMF1 | 0.627227 | 0.9067124 | 0.644382 | 0.5961176 | 0.981351 | 0.01828758 | 0.692179 | 0.184576 | -0.934881 | 0.4557081 | 0.636366 | 0.6885065 |
| ENSG00000144749.13 | LRIG1 | 0.944674 | 0.6314957 | -0.251793 | 0.6200225 | -0.443378 | 0.3393299 | 0.637214 | 0.01766606 | 0.935417 | 0.130985 | 1.908871 | 0.4149735 |
| ENSG00000144791.9 | LIMD1 | -0.53971 | 1 | -0.030017 | 0.6527736 | -1.109501 | 0.8454005 | 2.013465 | 0.03793568 | 0.798669 | 0.4201444 | -2.005476 | 0.6267144 |
| ENSG00000144815.14 | NXPE3 | 2.759151 | 0.3946735 | 0.939514 | 0.6200225 | 0.436337 | 0.9148391 | 1.04601 | 0.02029956 | 3.686468 | 0.03205693 | 0.465444 | 0.8354109 |
| ENSG00000144909.7 | OSBPL11 | -1.641821 | 0.6014343 | -0.233989 | 0.8192174 | 1.265467 | 0.002285028 | 0.563663 | 0.9693526 | 1.288475 | 0.696089 | -0.008384 | 0.7417751 |
| ENSG00000144959.9 | NCEH1 | 2.570461 | 0.8006286 | -0.017315 | 0.9815352 | -2.073213 | 0.5508895 | 3.209951 | 0.03153022 | -2.286485 | 0.8387831 | 0.828541 | 0.8949489 |
| ENSG00000145012.12 | LPP | -3.816287 | 0.6014343 | 0.485468 | 0.9682923 | 1.79584 | 0.000838873 | 1.271798 | 0.02074892 | 0.534739 | 0.6170361 | -0.608589 | 0.9708573 |
| ENSG00000145014.17 | TMEM44 | -1.095604 | 0.6214382 | -1.407913 | 0.6527736 | -0.610551 | 0.07205032 | 0.862922 | 0.00013631 | 2.161078 | 0.003418989 | -0.09385 | 0.6885065 |
| ENSG00000145050.15 | MANF | -1.349125 | 0.1710381 | 0.138982 | 0.3583128 | -0.314424 | 0.002474334 | -0.146389 | 0.04090345 | -0.364097 | 0.1757142 | 0.825963 | 0.6885065 |
| ENSG00000145284.11 | SCD5 | -0.180816 | 0.714717 | -0.700908 | 0.000125296 | 1.107606 | 0.08425552 | 0.429175 | 0.7858185 | 0.389123 | 0.940485 | 0.389172 | 0.8507262 |
| ENSG00000145337.4 | PYURF | 0.507537 | 0.3277506 | 0.13549 | 0.7299408 | -0.296271 | 0.04461904 | -0.206488 | 0.004545648 | -0.118563 | 0.0963637 | -0.14805 | 0.88512 |
| ENSG00000145349.16 | CAMK2D | 0.52166 | 0.6214382 | 0.022254 | 0.7793794 | 1.097571 | 4.53E-05 | 1.001405 | 0.01145825 | -0.065351 | 0.4646869 | 0.026806 | 0.9388279 |
| ENSG00000145354.9 | CISD2 | 0.838584 | 0.01509627 | -0.057096 | 0.8948108 | 0.297543 | 1 | 0.084219 | 0.6575769 | -0.392569 | 0.009087815 | 1.613082 | 1 |
| ENSG00000145362.17 | ANK2 | 1.489573 | 0.563505 | -0.575556 | 0.01536795 | 0.490798 | 0.5838725 | 1.161546 | 0.06671646 | 2.10916 | 0.05110354 | 2.87175 | 0.2068179 |
| ENSG00000145390.11 | USP53 | -0.722941 | 0.8867323 | -0.265044 | 0.704767 | 1.352876 | 0.005654245 | 0.598226 | 0.8224647 | 2.865107 | 0.7377262 | 0.620957 | 0.8949489 |
| ENSG00000145425.9 | RPS3A | -0.100116 | 0.6014343 | 0.807938 | 1.90E-18 | 0.637367 | 1.12E-31 | 0.112531 | 0.8905977 | -0.101193 | 0.003323382 | -0.181967 | 0.05728824 |
| ENSG00000145431.10 | PDGFC | 1.638776 | 0.538507 | 0.094998 | 0.8228029 | -0.300603 | 0.7371191 | 1.865409 | 1.55E-06 | 1.871544 | 0.6141002 | -0.40906 | 0.6885065 |
| ENSG00000145494.11 | NDUFS6 | -3.350747 | 0.4579069 | -0.13101 | 0.3045812 | -0.419189 | 4.27E-08 | 0.949939 | 0.4919368 | -0.03967 | 0.3245161 | 0.500164 | 0.4773627 |
| ENSG00000145495.14 | 6-Mar | -0.596787 | 0.4826978 | 0.337187 | 0.8461262 | 0.698024 | 0.000950538 | 1.807108 | 7.72E-05 | 0.673423 | 0.03102113 | 0.392708 | 0.7497966 |
| ENSG00000145555.14 | MYO10 | -0.339431 | 0.6014343 | 0.045915 | 0.6453628 | 0.156226 | 0.000682191 | 0.387191 | 0.9439697 | -0.172483 | 0.05392398 | -2.411483 | 0.6885065 |
| ENSG00000145592.13 | RPL37 | -0.357117 | 0.4023605 | -0.099654 | 0.4002287 | -0.063992 | 0.6272378 | -0.033829 | 0.2322369 | 0.084684 | 0.0196853 | 0.031011 | 0.6885065 |
| ENSG00000145675.14 | PIK3R1 | -0.878962 | 0.7939851 | -0.837238 | 0.07408047 | 0.363125 | 0.01128174 | 0.193713 | 0.1881597 | -1.214146 | 0.8021966 | 1.379656 | 0.9147227 |
| ENSG00000145685.13 | LHFPL2 | 0.483033 | 0.8867323 | -0.193259 | 0.9222921 | -0.797895 | 0.5290845 | 1.161378 | 0.03463891 | 2.879428 | 0.2016786 | 2.04453 | 0.7234069 |
| ENSG00000145725.19 | PPIP5K2 | -1.43894 | 1 | -0.215284 | 0.8953649 | -0.26408 | 0.5290845 | 1.506604 | 0.01035501 | 0.441504 | 0.5196843 | 1.533737 | 0.8949489 |
| ENSG00000145734.18 | BDP1 | 1.019376 | 0.6214382 | -0.105148 | 0.6527736 | 0.923717 | 4.10E-07 | 1.053839 | 7.02E-05 | 1.28225 | 0.002862708 | 1.629623 | 0.6517664 |
| ENSG00000145740.18 | SLC30A5 | -0.490706 | 0.7736255 | -0.641029 | 0.4324806 | -0.309533 | 0.3220802 | 1.481056 | 5.94E-07 | 1.29381 | 0.3079802 | -0.163474 | 0.6885065 |
| ENSG00000145741.15 | BTF3 | -0.324604 | 0.8587992 | 0.064538 | 0.9234066 | -0.043825 | 0.891888 | 0.141921 | 0.03471188 | 0.171622 | 0.3899638 | 0.147563 | 0.6185816 |
| ENSG00000145743.15 | FBXL17 | -0.090775 | 0.9047689 | -0.21963 | 0.9159517 | -1.673679 | 0.1379036 | 2.359129 | 0.01754162 | 1.300124 | 0.6668542 | 0.771543 | 0.826754 |
| ENSG00000145833.15 | DDX46 | -0.046535 | 0.7703009 | 0.355585 | 0.7269428 | 0.831327 | 0.000196135 | 0.271106 | 0.4919368 | 0.083929 | 0.1946233 | -0.709122 | 0.6490419 |
| ENSG00000145860.11 | RNF145 | -0.551993 | 0.2643306 | 0.299513 | 0.824352 | 0.171718 | 0.7030899 | 0.492913 | 0.001607334 | 1.023805 | 0.1589272 | 0.16456 | 0.7952164 |
| ENSG00000145868.16 | FBXO38 | 2.163743 | 0.4764974 | 0.499487 | 0.9768713 | 0.076403 | 0.8100781 | 2.023337 | 0.03360361 | 0.540425 | 0.5497593 | -1.479019 | 0.9359415 |
| ENSG00000145901.14 | TNIP1 | 0.524843 | 0.9183025 | -0.25647 | 0.7211571 | -0.4725 | 0.4176494 | 0.059929 | 0.004203678 | -0.184251 | 0.4973025 | 0.646678 | 0.8949489 |
| ENSG00000145907.14 | G3BP1 | -0.22661 | 1 | -0.001831 | 0.6527736 | 0.496725 | 0.005063536 | 0.418176 | 0.4791525 | 0.109035 | 1 | 0.951307 | 0.6829816 |
| ENSG00000145908.12 | ZNF300 | 3.437405 | 0.3098608 | -0.143006 | 1 | 1.514075 | 0.1163266 | 1.557465 | 0.02962288 | 0.698824 | 0.6177238 | 0.343383 | 0.6885065 |
| ENSG00000145912.8 | NHP2 | 0.318056 | 0.1788327 | 0.123677 | 0.8217443 | 0.002048 | 0.1457035 | -0.166037 | 0.04645593 | -0.352354 | 0.000300526 | 0.492143 | 0.6885065 |
| ENSG00000145990.10 | GFOD1 | -0.313732 | 0.8867323 | -0.96176 | 0.4531355 | 0.364676 | 0.765363 | 2.331644 | 0.001756941 | 1.485381 | 0.4096664 | 1.105561 | 0.6885065 |
| ENSG00000145996.11 | CDKAL1 | -0.141083 | 1 | 0.234332 | 0.979665 | -0.539456 | 0.6879111 | 3.019772 | 0.02076542 | 0.06448 | 0.42196 | -0.389212 | 0.8903882 |
| ENSG00000146006.7 | LRRTM2 | 1.684103 | 0.6014343 | -1.114548 | 0.01272782 | -0.135863 | 0.7354314 | 1.302315 | 0.000220402 | 1.893408 | 0.000364991 | -0.270793 | 0.8479431 |
| ENSG00000146007.10 | ZMAT2 | -0.41008 | 0.6798783 | 0.288607 | 0.575271 | -0.012019 | 0.9533414 | 0.339067 | 0.2869336 | -0.493178 | 0.01316432 | 0.861846 | 0.9147227 |
| ENSG00000146063.18 | TRIM41 | 0.890287 | 0.8587992 | 0.152841 | 0.8703522 | -1.349038 | 0.3118792 | 1.487581 | 0.04825715 | 4.049504 | 0.000204097 | 1.375125 | 0.6885065 |
| ENSG00000146066.2 | HIGD2A | 0.055699 | 0.85228 | 0.037007 | 0.5651089 | -0.133406 | 0.2594975 | 0.176593 | 0.04341242 | -0.313859 | 0.06053149 | -1.700976 | 0.1000281 |
| ENSG00000146067.15 | FAM193B | 0.345505 | 1 | 0.343743 | 0.8485436 | 0.168976 | 0.8267131 | 0.558053 | 0.004641792 | 1.121345 | 0.007120097 | 0.118227 | 1 |
| ENSG00000146083.11 | RNF44 | -2.074771 | 0.4322214 | 0.184855 | 0.7383998 | 0.686152 | 0.04259207 | 0.821306 | 0.5968653 | 1.200323 | 0.0963637 | 0.363714 | 0.6885065 |
| ENSG00000146109.4 | ABT1 | -0.515282 | 0.6508875 | 0.249431 | 0.9988717 | -1.477221 | 0.009655101 | 1.39696 | 1.94E-06 | 1.576321 | 0.005714101 | -1.742373 | 0.6885065 |
| ENSG00000146216.11 | TTBK1 | -1.036453 | 0.6214382 | -2.835821 | 0.5394427 | -1.342769 | 0.5290845 | 0.309213 | 0.04239959 | 0.483941 | 0.5551535 | -2.32764 | 0.7501684 |
| ENSG00000146223.14 | RPL7L1 | -0.123943 | 0.7736255 | 0.463135 | 0.3255429 | 0.2649 | 0.1851823 | 1.790033 | 1.57E-07 | 0.357974 | 0.4055596 | -0.15585 | 0.553308 |
| ENSG00000146232.14 | NFKBIE | 0.372588 | 0.8264007 | -0.980819 | 0.592336 | 0.10307 | 0.04895234 | 0.727344 | 0.9307757 | -0.106487 | 0.3797324 | 1.597614 | 0.8643622 |
| ENSG00000146247.13 | PHIP | 0.187045 | 0.6014343 | 0.126109 | 0.8263112 | 1.569415 | 1.65E-07 | 1.802242 | 5.18E-10 | 1.485968 | 0.01083921 | 1.386352 | 0.7901525 |
| ENSG00000146267.11 | FAXC | -1.883543 | 0.6214382 | -0.16814 | 0.7881421 | 1.153423 | 0.6472573 | 4.189515 | 0.01870689 | -0.141605 | 0.9735552 | -3.924854 | 0.7767294 |
| ENSG00000146278.10 | PNRC1 | -1.463706 | 0.6214382 | 0.225515 | 0.6088437 | 0.578678 | 0.7030715 | 0.360584 | 0.5930297 | 1.621728 | 0.01796938 | 1.727729 | 0.7265625 |
| ENSG00000146282.17 | RARS2 | -0.939977 | 0.4464112 | 0.02687 | 0.6828727 | 0.513056 | 0.03798493 | -0.910817 | 0.4379912 | -0.175488 | 0.1029519 | 0.876756 | 0.6885065 |
| ENSG00000146373.16 | RNF217 | 2.967871 | 0.8006286 | -5.095656 | 0.6527736 | 1.608745 | 0.4064599 | 0.600933 | 0.3404336 | 3.245759 | 0.00513524 | #NAME? | 0.6885065 |
| ENSG00000146386.7 | ABRACL | -0.096407 | 0.714717 | 0.229832 | 0.6731614 | -0.534332 | 0.6141868 | 0.008315 | 0.1061598 | -0.741952 | 0.04233873 | 1.24122 | 0.4403284 |
| ENSG00000146414.15 | SHPRH | 0.174664 | 0.8867323 | 0.455004 | 0.8703522 | 0.724813 | 0.01722079 | 2.775626 | 0.07312586 | 1.997852 | 0.2167378 | 2.233922 | 0.8479431 |
| ENSG00000146425.10 | DYNLT1 | -0.191703 | 0.9483118 | 0.064178 | 0.5953732 | -0.192154 | 0.07205032 | -0.155806 | 0.000822785 | -0.07596 | 0.00685221 | -0.149718 | 0.8056213 |
| ENSG00000146433.8 | TMEM181 | 0.784168 | 0.6214382 | 0.339 | 0.6527736 | 0.023574 | 0.7788986 | 2.148089 | 1.92E-10 | 2.022373 | 0.007819554 | 0.088793 | 0.9849258 |
| ENSG00000146457.14 | WTAP | -0.484079 | 0.392544 | -0.176317 | 0.6109044 | -0.863443 | 0.002199471 | 0.51135 | 0.006286247 | 0.976219 | 0.1427939 | 0.445999 | 0.6262252 |
| ENSG00000146535.13 | GNA12 | -5.76723 | 0.8006286 | 0.289957 | 0.6527736 | -0.388121 | 0.3573028 | 0.187417 | 0.03400141 | 1.51849 | 0.3027771 | 1.446628 | 0.6885065 |
| ENSG00000146540.14 | C7orf50 | -0.378509 | 0.9548281 | 0.251395 | 0.6527736 | -0.595426 | 2.65E-09 | -0.006709 | 0.501923 | 0.56714 | 0.4356309 | -0.177355 | 0.5828283 |
| ENSG00000146576.12 | C7orf26 | -1.068539 | 0.6214382 | -0.191144 | 0.6527736 | -1.107058 | 0.06328476 | -0.275868 | 0.277184 | 1.984357 | 0.03711912 | -3.846193 | 0.8479431 |
| ENSG00000146587.17 | RBAK | 0.842155 | 1 | -0.133421 | 0.5732186 | 1.066292 | 1.27E-26 | 0.86017 | 0.4562461 | 2.020943 | 0.03778507 | -1.669326 | 0.7020544 |
| ENSG00000146592.16 | CREB5 | 0.477837 | 1 | -0.217311 | 0.6527736 | 1.863393 | 7.03E-16 | 0.572814 | 0.4583363 | 0.997906 | 0.01233229 | -1.85671 | 0.954498 |
| ENSG00000146648.15 | EGFR | 0.168264 | 0.9453571 | -0.317584 | 0.000501508 | 0.304612 | 0.004064813 | 0.972795 | 0.04461712 | 0.892977 | 0.2387526 | -2.322175 | 0.8949489 |
| ENSG00000146676.6 | PURB | -1.922111 | 0.6522981 | 0.291223 | 0.628885 | 1.093467 | 2.91E-08 | 1.063733 | 0.0427187 | 1.122753 | 0.04754247 | 0.91335 | 0.9606764 |
| ENSG00000146731.10 | CCT6A | -0.901638 | 0.5139066 | 0.181828 | 7.56E-06 | 0.553776 | 2.13E-06 | 0.564131 | 0.05976993 | 0.150915 | 0.2204656 | 0.576161 | 0.5647157 |
| ENSG00000146757.13 | ZNF92 | -0.610157 | 0.9183025 | 0.10962 | 0.7702247 | 0.690053 | 0.4064599 | 0.951609 | 0.01813701 | 1.166024 | 0.09168286 | 0.83374 | 0.6885065 |
| ENSG00000146776.14 | ATXN7L1 | -0.754981 | 1 | -0.006067 | 1 | 4.15725 | 0.5290845 | 3.433854 | 0.04008401 | 1.33672 | 1 | 1.34314 | 0.6885065 |
| ENSG00000146830.9 | GIGYF1 | -2.047784 | 0.6214382 | 0.369098 | 0.6527736 | -0.410828 | 0.3009727 | 1.172686 | 0.06404674 | 1.908606 | 0.000281589 | -2.497729 | 0.6829816 |
| ENSG00000146833.15 | TRIM4 | -1.131658 | 0.6792949 | -0.439563 | 0.6527736 | 0.745031 | 0.3278254 | 1.658516 | 0.008552578 | 0.866746 | 0.5414081 | 0.691473 | 0.9147227 |
| ENSG00000146834.13 | MEPCE | 1.193907 | 0.6014343 | 0.29744 | 0.9839225 | 0.074299 | 0.4064599 | 1.213715 | 0.000361214 | 0.803448 | 0.3657673 | -0.89818 | 0.826754 |
| ENSG00000146842.16 | TMEM209 | 1.677991 | 0.529676 | 0.150491 | 0.6527736 | 1.549684 | 0.01347034 | 0.333706 | 0.7144938 | 0.969247 | 0.8589299 | -0.275111 | 0.9617345 |
| ENSG00000146938.14 | NLGN4X | -0.032751 | 0.810319 | -0.285244 | 0.5260916 | 2.152497 | 0.009884704 | 1.471924 | 8.17E-06 | 0.604849 | 0.4458415 | -0.463494 | 0.9264343 |
| ENSG00000147010.17 | SH3KBP1 | -0.053942 | 0.6214382 | -0.72943 | 0.01296324 | 1.255926 | 0.1126036 | -0.968666 | 0.001555312 | -0.959313 | 0.1907698 | 2.07735 | 0.6885065 |
| ENSG00000147027.3 | TMEM47 | 2.465693 | 0.6214382 | 0.136381 | 1 | 0.88733 | 0.1357261 | -1.614572 | 1 | 1.445272 | 1 |  |  |
| ENSG00000147044.20 | CASK | -0.818997 | 0.5768305 | -0.513615 | 0.3583128 | 0.98288 | 0.1923336 | 0.58292 | 0.002854442 | 0.834766 | 0.06231981 | -0.522453 | 0.5191309 |
| ENSG00000147123.10 | NDUFB11 | 0.250618 | 0.5428806 | 0.054891 | 0.9794341 | -0.400686 | 0.002498065 | 0.010966 | 0.04943624 | -0.495335 | 0.000323417 | 0.301093 | 0.5344421 |
| ENSG00000147133.15 | TAF1 | -3.685222 | 0.2717194 | 0.18939 | 0.846655 | 0.413263 | 0.9836024 | 1.306836 | 5.01E-05 | 1.660217 | 0.04281867 | -1.950771 | 0.9147227 |
| ENSG00000147140.15 | NONO | 0.990133 | 0.01177678 | -0.057426 | 0.7662689 | 0.421329 | 0.3999877 | 1.447304 | 1.19E-05 | 0.949174 | 0.02368927 | -0.295766 | 0.4727232 |
| ENSG00000147162.13 | OGT | 0.499656 | 0.563505 | -0.937313 | 0.5700388 | 1.092121 | 0.4382561 | 0.832346 | 0.0873308 | 2.26098 | 3.20E-05 | -1.480702 | 1 |
| ENSG00000147180.16 | ZNF711 | -0.11887 | 0.8324763 | -1.182093 | 0.08794836 | 0.994323 | 0.0926263 | 1.06918 | 0.01164354 | 0.489574 | 0.08874741 | 1.242236 | 1 |
| ENSG00000147274.14 | RBMX | -0.353135 | 1 | -0.067245 | 0.3160414 | 0.225019 | 0.02083316 | 0.273742 | 0.2813112 | 0.006965 | 0.7104629 | -0.676439 | 0.01138072 |
| ENSG00000147316.12 | MCPH1 | 0.222964 | 0.563505 | -0.838577 | 0.03324584 | 0.193108 | 0.01918694 | 0.882982 | 0.7953528 | 0.100984 | 0.216249 | -3.452608 | 0.7827587 |
| ENSG00000147400.8 | CETN2 | 0.576823 | 0.529676 | 0.420406 | 0.6996212 | -0.012803 | 0.4064599 | -0.612308 | 4.58E-05 | -0.924414 | 0.01759495 | 0.185574 | 0.6885065 |
| ENSG00000147403.16 | RPL10 | -0.420827 | 0.000670282 | 0.116696 | 0.5953732 | 0.248242 | 3.25E-07 | -0.022899 | 0.1681854 | 0.098406 | 0.1196747 | 0.151747 | 1 |
| ENSG00000147416.10 | ATP6V1B2 | 0.859568 | 0.02153352 | -0.031242 | 0.8760329 | 0.401994 | 0.8811167 | 1.776961 | 0.000687068 | -0.105967 | 0.8425408 | 0.551714 | 0.6829816 |
| ENSG00000147421.17 | HMBOX1 | -1.353835 | 0.1448298 | 0.314627 | 0.6527736 | 0.861955 | 0.000873782 | 1.088171 | 0.2009221 | 0.978198 | 1 | 0.84836 | 0.9147227 |
| ENSG00000147454.13 | SLC25A37 | 1.379533 | 0.6530318 | -0.755827 | 0.121014 | 0.9058 | 0.002654725 | 1.248134 | 0.01859259 | 0.645594 | 0.9897941 | 0.390093 | 0.8360203 |
| ENSG00000147475.14 | ERLIN2 | 0.459549 | 0.8587992 | -0.287168 | 0.5517607 | -0.149037 | 0.2879524 | 0.213595 | 0.02131309 | -0.456639 | 0.6636264 | 1.2628 | 1 |
| ENSG00000147533.16 | GOLGA7 | 0.703789 | 0.000810564 | 0.059358 | 0.8953649 | 0.785002 | 0.2980055 | 0.132817 | 0.4529832 | 0.417578 | 1 | 0.265868 | 0.9526606 |
| ENSG00000147548.16 | WHSC1L1 | -1.186321 | 0.6014343 | -0.293398 | 0.32473 | 1.147924 | 3.18E-06 | 1.538201 | 0.001827364 | 0.647525 | 0.06557358 | -0.215983 | 0.8309897 |
| ENSG00000147586.9 | MRPS28 | -1.316644 | 0.8006286 | -0.176561 | 0.7801026 | -0.008066 | 0.4018207 | -0.714754 | 0.01657862 | 0.319475 | 0.8266278 | -0.993315 | 0.9368176 |
| ENSG00000147588.6 | PMP2 | -0.613668 | 0.1492051 | -0.138276 | 0.3242827 | 1.010526 | 7.06E-15 | 0.205917 | 0.1926368 | 0.126476 | 0.001570875 | -1.102898 | 0.009027293 |
| ENSG00000147604.13 | RPL7 | -0.583667 | 0.02247068 | 0.841686 | 2.42E-08 | 0.668227 | 3.08E-22 | -0.067322 | 0.01401338 | 0.209097 | 0.1095202 | -0.385603 | 0.02166254 |
| ENSG00000147649.9 | MTDH | -0.335282 | 1 | 0.255507 | 0.542312 | 0.306534 | 0.1625971 | 0.758892 | 0.02455892 | 0.659305 | 0.1688763 | -1.160783 | 0.4301971 |
| ENSG00000147669.10 | POLR2K | -1.202019 | 0.529676 | 0.653004 | 0.5395105 | 0.018643 | 0.00794631 | 0.151111 | 0.5543343 | -0.470129 | 0.000896152 | -0.560367 | 0.4578379 |
| ENSG00000147677.10 | EIF3H | -0.386975 | 0.6214382 | 0.092081 | 0.8484484 | -0.054113 | 0.9381042 | -0.13235 | 0.1241064 | -0.405177 | 0.01083921 | 0.267729 | 0.6885065 |
| ENSG00000147684.7 | NDUFB9 | -1.422855 | 0.04459519 | -0.242575 | 0.1232781 | -0.285317 | 0.00162993 | 0.177009 | 0.1003607 | -0.153823 | 0.000696277 | -0.49565 | 0.04261865 |
| ENSG00000147854.16 | UHRF2 | -0.6702 | 0.8006286 | 0.127547 | 0.9378151 | -0.596854 | 0.1394359 | 0.743714 | 0.002367512 | 1.623875 | 2.04E-05 | 0.013256 | 0.9147227 |
| ENSG00000147862.14 | NFIB | -0.47731 | 0.9678883 | -0.498842 | 0.01825744 | 0.523067 | 0.000425536 | 0.981044 | 1.36E-06 | 1.524744 | 1.29E-05 | 0.7531 | 0.6829816 |
| ENSG00000147872.9 | PLIN2 | -0.166826 | 0.8014507 | 0.873746 | 0.000338069 | -0.995283 | 0.03124823 | 1.193756 | 0.5501287 | -3.344105 | 0.3255801 | 0.686142 | 0.9216612 |
| ENSG00000148019.12 | CEP78 | 0.116128 | 0.9282145 | -0.674429 | 0.5943878 | -0.297328 | 0.8587255 | 1.292575 | 0.01711435 | 0.561254 | 0.4884683 | -0.809727 | 0.951024 |
| ENSG00000148082.9 | SHC3 | 0.353505 | 0.3152417 | -0.391598 | 0.57799 | 0.594258 | 0.3378366 | 1.533757 | 6.44E-07 | 1.769673 | 0.5196843 | -1.016625 | 0.9174022 |
| ENSG00000148110.15 | HIATL1 | -3.751193 | 0.06285358 | -0.274411 | 0.6853354 | 0.225702 | 0.6140882 | 1.832553 | 0.003059717 | -1.2674 | 0.8617983 | 2.217006 | 0.4322251 |
| ENSG00000148143.12 | ZNF462 | -1.260327 | 0.3946735 | -0.084943 | 1 | 0.333279 | 0.08944676 | 1.621454 | 9.05E-06 | 0.593007 | 0.4780862 | -1.393584 | 0.8479431 |
| ENSG00000148154.9 | UGCG | -0.620786 | 0.6214382 | -0.314931 | 0.6532996 | 1.11106 | 0.002081575 | 0.625872 | 0.4486565 | 0.551989 | 0.9243642 | 1.041488 | 0.5733051 |
| ENSG00000148158.16 | SNX30 | 2.507316 | 0.6214382 | 0.266528 | 0.8549935 | 0.339399 | 0.6821321 | 2.16215 | 0.08298408 | 1.966145 | 0.03562571 | -0.669232 | 0.6829816 |
| ENSG00000148248.13 | SURF4 | 1.147324 | 0.01172921 | 0.236964 | 0.577543 | 0.235565 | 0.9449308 | 0.134946 | 0.791347 | 1.144072 | 0.4249407 | 1.677269 | 0.8233128 |
| ENSG00000148303.16 | RPL7A | -0.663286 | 0.001177428 | 0.007365 | 0.9452791 | 0.009859 | 0.6827367 | 0.242836 | 0.2480199 | 0.161426 | 0.1739619 | 0.093855 | 0.426223 |
| ENSG00000148341.17 | SH3GLB2 | -0.084096 | 1 | -1.289644 | 0.4011918 | -0.182297 | 0.6933226 | 2.01757 | 0.03774232 | 0.92492 | 0.07222869 | 1.407097 | 0.7429541 |
| ENSG00000148358.19 | GPR107 | -1.973521 | 0.3534588 | -0.239699 | 0.5553512 | 0.470039 | 0.01935118 | 0.306598 | 0.02021101 | 0.815561 | 0.4967944 | -0.398592 | 0.7494298 |
| ENSG00000148362.10 | C9orf142 | 1.110066 | 0.4406957 | -0.195339 | 0.5953732 | -0.478122 | 0.01670394 | 0.8872 | 0.7503704 | -0.329506 | 0.4657463 | -1.004043 | 0.8089706 |
| ENSG00000148384.11 | INPP5E | 2.024364 | 0.6787488 | -0.798844 | 0.8953649 | -0.709274 | 0.8100781 | 3.472612 | 0.003001948 | 2.01551 | 0.01353696 | 0.097192 | 0.6885065 |
| ENSG00000148396.18 | SEC16A | -1.947939 | 0.3946735 | -0.02455 | 0.9222921 | 0.131456 | 0.2082075 | 0.877333 | 0.001311392 | 0.672527 | 0.1778915 | -0.6103 | 0.8949489 |
| ENSG00000148399.11 | DPH7 | -0.572285 | 0.9838363 | -0.191698 | 0.8731418 | -0.364352 | 0.01112682 | 0.921065 | 0.127053 | 0.381573 | 0.8672172 | -0.252014 | 0.958512 |
| ENSG00000148400.9 | NOTCH1 | -0.140298 | 0.9183025 | -0.153344 | 0.592336 | -0.375497 | 0.7786988 | -0.316495 | 0.1048458 | 1.676007 | 0.001983529 | 0.436096 | 0.6829816 |
| ENSG00000148411.7 | NACC2 | 0.820183 | 0.7736255 | -0.121161 | 0.8238603 | -0.179399 | 0.6848226 | 1.288831 | 0.04503337 | 0.968142 | 0.5196843 | 1.75959 | 0.6885065 |
| ENSG00000148468.16 | FAM171A1 | 0.599731 | 0.6214382 | -1.435037 | 0.6527736 | -0.528829 | 1 | 0.963582 | 0.008649388 | 1.09268 | 0.3920048 | 1.668516 | 0.6885065 |
| ENSG00000148484.17 | RSU1 | 0.180237 | 0.7736255 | 0.223673 | 0.8272197 | -0.647825 | 0.4804635 | -0.734759 | 0.03682396 | 0.09967 | 0.5196843 | -1.526897 | 0.7232476 |
| ENSG00000148516.21 | ZEB1 | -0.048607 | 0.1823038 | -0.297566 | 0.1141252 | 0.837025 | 0.003647393 | 1.201723 | 0.005440612 | 0.104284 | 0.3244978 | 1.253812 | 0.8479431 |
| ENSG00000148672.8 | GLUD1 | -0.434533 | 0.7833919 | 0.263623 | 0.8938811 | -0.088421 | 0.7354314 | 1.130123 | 0.02840309 | 1.631554 | 0.2019818 | -0.689242 | 0.6829816 |
| ENSG00000148730.6 | EIF4EBP2 | 0.919263 | 0.6214382 | -0.049597 | 0.4017139 | -0.540611 | 0.9914608 | 1.371405 | 0.002023283 | 1.875492 | 0.004806412 | -0.519914 | 0.5344421 |
| ENSG00000148834.12 | GSTO1 | 0.686952 | 0.529676 | 0.14855 | 0.4011918 | -0.88614 | 0.005259102 | 0.258955 | 0.3669262 | -0.052705 | 0.6217705 | -0.20193 | 0.6885065 |
| ENSG00000148840.10 | PPRC1 | 0.071744 | 0.8867323 | -0.268907 | 0.8703522 | 0.691078 | 0.08771684 | 1.385213 | 0.003665166 | 1.807674 | 1 | -0.512878 | 0.8464135 |
| ENSG00000148841.15 | ITPRIP | -0.167997 | 0.7422989 | -0.42025 | 0.6782932 | -1.166283 | 0.4189965 | 0.539907 | 0.02685789 | -0.345737 | 0.7268459 | -1.45555 | 0.6829816 |
| ENSG00000148843.13 | PDCD11 | -0.193201 | 0.7337791 | 0.307146 | 0.9638587 | 0.78609 | 0.00489178 | 1.546491 | 0.2467163 | 0.230119 | 0.5196843 | -0.48153 | 0.790982 |
| ENSG00000148950.9 | IMMP1L | -0.090143 | 0.9474844 | -0.022032 | 0.9653874 | -0.103226 | 0.006870424 | -0.412521 | 0.09170208 | -0.590153 | 0.2364542 | 0.950244 | 0.9380942 |
| ENSG00000149084.11 | HSD17B12 | 0.607011 | 0.1508129 | -0.084204 | 0.5394427 | 1.171399 | 9.68E-06 | 0.7152 | 0.3340419 | 0.314183 | 0.7377262 | 0.198862 | 0.8479431 |
| ENSG00000149091.15 | DGKZ | -0.08323 | 0.7736255 | 1.304025 | 0.002750998 | -0.046205 | 0.4382561 | 1.540955 | 0.2362617 | 0.700278 | 0.3856748 | -1.892674 | 0.8316849 |
| ENSG00000149115.13 | TNKS1BP1 | -1.38785 | 1 | -0.89511 | 0.5953732 | -0.049944 | 0.7207737 | 2.012236 | 0.000882846 | 3.118906 | 0.2913839 | -0.958649 | 0.6829816 |
| ENSG00000149136.8 | SSRP1 | 0.471201 | 0.3254792 | 0.015664 | 0.5953732 | 0.559323 | 0.000173496 | 0.527508 | 0.7760884 | -0.003177 | 0.09023571 | -0.192402 | 0.6885065 |
| ENSG00000149187.17 | CELF1 | -0.202697 | 1 | -0.295619 | 0.8225043 | 0.617641 | 0.00036892 | 1.452969 | 1.33E-09 | 0.589445 | 0.05375272 | 0.169025 | 0.9147227 |
| ENSG00000149196.15 | C11orf73 | -1.031215 | 0.4406957 | 0.275306 | 0.5394427 | -0.005251 | 0.007979364 | 0.53663 | 0.4919368 | -0.321993 | 0.000260231 | -2.207244 | 0.5066477 |
| ENSG00000149212.11 | SESN3 | -1.096315 | 0.6214382 | 0.167981 | 0.9525045 | 1.456425 | 0.005094059 | 1.702559 | 0.1589128 | 0.196702 | 0.4011029 | 0.675652 | 0.9147227 |
| ENSG00000149218.4 | ENDOD1 | 0.499275 | 0.7725499 | -0.073987 | 0.3126509 | 0.37421 | 0.9354988 | 1.258103 | 0.02521105 | 1.90405 | 0.7216224 | 0.985021 | 0.9589332 |
| ENSG00000149231.11 | CCDC82 | -0.455657 | 0.8641853 | 0.258266 | 0.8849675 | 0.715959 | 0.009242744 | -0.262055 | 0.9805636 | -0.164141 | 0.07255698 | 0.942944 | 0.8202624 |
| ENSG00000149256.14 | TENM4 | -2.735566 | 0.6214382 | -0.609176 | 0.7949484 | -0.059442 | 0.7898157 | 1.700892 | 0.0206566 | -3.156784 | 0.4671878 | 1.839534 | 0.8949489 |
| ENSG00000149273.14 | RPS3 | -0.607828 | 0.01068364 | -0.006364 | 1 | 0.127712 | 0.06860375 | 0.10254 | 0.1469461 | -0.376295 | 5.55E-08 | -0.322906 | 0.04613118 |
| ENSG00000149294.16 | NCAM1 | -0.131761 | 0.714717 | -0.495071 | 0.00031583 | 0.712642 | 0.000478237 | 1.045494 | 1.73E-06 | 0.985299 | 0.001842495 | 0.286004 | 0.6885065 |
| ENSG00000149308.16 | NPAT | -0.901528 | 0.6214382 | -0.209175 | 1 | 1.128456 | 0.01624726 | 0.053979 | 0.841123 | 0.903492 | 1 | -0.631697 | 0.9428037 |
| ENSG00000149311.17 | ATM | -0.853807 | 0.7683685 | -0.38866 | 0.5953732 | 0.683732 | 0.4189965 | 1.194474 | 0.008845667 | 1.23274 | 0.1764893 | 0.857594 | 0.9526606 |
| ENSG00000149313.10 | AASDHPPT | 0.562616 | 0.3987745 | -0.137583 | 0.8721183 | 0.808842 | 0.1778406 | 0.509703 | 0.1894794 | -1.111268 | 0.02294687 | -0.524827 | 0.6885065 |
| ENSG00000149357.9 | LAMTOR1 | 0.636631 | 0.06126222 | 0.059997 | 0.906244 | -0.407065 | 0.1084149 | 0.115462 | 0.03998217 | -0.900823 | 0.001233426 | 0.310747 | 0.9801657 |
| ENSG00000149480.6 | MTA2 | 0.478185 | 0.529676 | -0.04485 | 0.9563746 | -0.78897 | 0.5290845 | 1.229501 | 0.000148363 | 1.082442 | 0.1034049 | 0.644095 | 0.7767294 |
| ENSG00000149503.12 | INCENP | -0.601833 | 0.8006286 | -2.379158 | 0.07233965 | 0.332337 | 0.9760024 | -1.417578 | 0.8877935 | 3.045183 | 0.02062054 | 1.353775 | 0.9476473 |
| ENSG00000149532.15 | CPSF7 | 1.196533 | 0.5139066 | 0.493509 | 0.5010139 | 0.235747 | 0.6462166 | 0.317081 | 0.08788191 | 0.362294 | 0.0117725 | 1.722447 | 1 |
| ENSG00000149557.12 | FEZ1 | 0.107357 | 0.838572 | 0.04233 | 0.6527736 | 0.238498 | 0.2348326 | 0.212636 | 0.08414325 | -0.271966 | 0.002003795 | 0.581509 | 0.9456372 |
| ENSG00000149577.15 | SIDT2 | 1.10897 | 0.8006286 | -1.089529 | 0.3959708 | 0.299344 | 0.5290845 | 1.194834 | 0.9105922 | 3.789702 | 0.04281867 | 2.833688 | 0.5733051 |
| ENSG00000149639.14 | SOGA1 | -0.434968 | 0.9047689 | -0.277482 | 0.7720357 | 0.404123 | 0.5290845 | 1.294002 | 0.00012979 | 1.805765 | 0.002407718 | 0.829071 | 0.782216 |
| ENSG00000149658.17 | YTHDF1 | 1.171155 | 0.8867323 | 0.174791 | 0.9967095 | 0.254891 | 0.9249654 | 2.440171 | 1.82E-06 | 0.244916 | 0.00746047 | -0.062356 | 0.8309897 |
| ENSG00000149806.10 | FAU | 0.202577 | 0.003228446 | 0.141749 | 0.3673142 | -0.201601 | 0.001739388 | -0.129514 | 0.005792373 | -0.104219 | 0.01542573 | -0.15469 | 0.1830592 |
| ENSG00000149809.14 | TM7SF2 | -4.207598 | 0.02378468 | -1.141602 | 0.3583128 | 2.31928 | 0.7669328 | 0.908532 | 0.7914774 | -0.519598 | 0.5196843 | -2.623529 | 1 |
| ENSG00000149925.16 | ALDOA | -0.452135 | 0.3946735 | 0.241826 | 0.005047786 | 0.06253 | 0.4064599 | 0.020054 | 0.1083296 | 0.008406 | 0.001502788 | 0.214486 | 0.9576568 |
| ENSG00000149926.13 | FAM57B | -1.161475 | 0.1675981 | 1.155155 | 0.6200225 | 0.417834 | 0.7437308 | 1.230823 | 0.1099928 | 1.5141 | 0.002514978 | -0.020146 | 0.9002633 |
| ENSG00000149930.17 | TAOK2 | 0.448249 | 0.9282145 | 0.7623 | 0.5953732 | -0.379821 | 0.6997526 | 1.844813 | 8.90E-05 | 1.05906 | 0.06557358 | -0.191865 | 0.6829816 |
| ENSG00000149932.16 | TMEM219 | 0.227987 | 0.9257047 | 0.100073 | 1 | -0.556798 | 0.01732351 | 0.339311 | 0.169916 | 0.180094 | 0.9466051 | 0.927112 | 1 |
| ENSG00000149970.14 | CNKSR2 | -0.12373 | 0.9047689 | -2.278739 | 0.01326762 | 1.729356 | 0.004736971 | 2.042101 | 0.002259593 | 1.490098 | 0.3100093 | 2.637342 | 0.9506884 |
| ENSG00000150347.14 | ARID5B | -0.80786 | 0.9695108 | -0.324016 | 0.0619414 | 0.71263 | 0.001932565 | -0.120922 | 1 | 0.62087 | 0.9675035 | 0.268093 | 0.8490512 |
| ENSG00000150459.12 | SAP18 | 0.002864 | 0.3582986 | -0.030338 | 0.6527736 | -0.024754 | 0.1674476 | 0.573918 | 0.7873955 | 0.167998 | 0.02276258 | 0.562692 | 0.553308 |
| ENSG00000150471.15 | ADGRL3 | 0.733211 | 0.6214382 | -0.905848 | 0.0125189 | 0.618752 | 0.001708461 | 1.039273 | 0.01696554 | 0.269678 | 0.7346833 | -0.124766 | 0.6185816 |
| ENSG00000150551.10 | LYPD1 | -0.444773 | 0.9553313 | -0.436541 | 0.01738075 | -0.557279 | 3.57E-06 | 0.468866 | 0.3292457 | 0.452259 | 0.1225306 | -1.253359 | 0.02166254 |
| ENSG00000150625.16 | GPM6A | 0.381799 | 0.4406957 | -0.327413 | 0.06515464 | 0.62514 | 6.45E-08 | 0.850027 | 0.000132579 | 0.489102 | 0.1346421 | 0.098493 | 0.6572299 |
| ENSG00000150687.11 | PRSS23 | -1.31435 | 0.8006286 | -0.000382 | 0.7881421 | -0.285632 | 0.01606964 | -0.516119 | 0.02889839 | -0.449855 | 0.3974318 | -0.506939 | 1 |
| ENSG00000150779.11 | TIMM8B | 0.026541 | 0.7736255 | 0.313692 | 0.6527736 | -0.304909 | 0.000860168 | -0.046 | 0.2324655 | -0.502441 | 0.01243284 | 0.206784 | 0.5223622 |
| ENSG00000150787.7 | PTS | 0.660885 | 0.1835496 | -0.107426 | 0.9618682 | -0.408569 | 0.1167096 | -0.010553 | 0.8138963 | -0.803684 | 0.000197614 | -0.195467 | 0.4304069 |
| ENSG00000150938.9 | CRIM1 | 2.79217 | 0.6014343 | -0.071168 | 0.9120034 | 0.897485 | 0.6575661 | 2.749836 | 0.00508131 | -0.190202 | 0.07956883 | 0.628328 | 0.6885065 |
| ENSG00000150991.14 | UBC | 0.438627 | 0.001815581 | 0.122229 | 0.368162 | 0.213225 | 0.000472418 | 0.04465 | 0.00517312 | 0.707248 | 0.1449763 | -0.042145 | 0.4304069 |
| ENSG00000151023.16 | ENKUR | -0.488784 | 0.8006286 | -0.385672 | 0.7834199 | 0.425812 | 0.8100781 | -1.190637 | 0.03511444 | -0.571263 | 0.3974318 | -2.431353 | 0.071337 |
| ENSG00000151150.20 | ANK3 | -0.995145 | 0.4764974 | -1.501182 | 0.2293481 | 0.153104 | 0.339353 | 0.32987 | 0.03897443 | -0.354041 | 0.1449215 | 0.864311 | 0.8479431 |
| ENSG00000151229.12 | SLC2A13 | -0.406805 | 0.826609 | -0.74009 | 0.1012381 | 1.138518 | 0.2295335 | 1.206318 | 0.02261337 | 0.25465 | 0.7680282 | 2.694987 | 0.9352766 |
| ENSG00000151233.10 | GXYLT1 | -1.425685 | 1 | -0.147476 | 0.6527736 | -0.097465 | 0.8100781 | 4.258018 | 0.04333874 | 0.825636 | 0.2653803 | 1.062259 | 0.6885065 |
| ENSG00000151276.23 | MAGI1 | -2.78751 | 0.705434 | -1.080212 | 0.2863181 | 1.025797 | 0.05717048 | 2.024994 | 3.58E-05 | 1.936393 | 0.03012886 | 0.538832 | 0.4773627 |
| ENSG00000151304.5 | SRFBP1 | -0.623964 | 0.845546 | -0.199808 | 0.9984248 | 1.195345 | 0.01722513 | -0.092187 | 0.841123 | -0.564843 | 0.03299258 | -0.549795 | 0.9531804 |
| ENSG00000151320.10 | AKAP6 | -0.619524 | 0.8205765 | -0.280306 | 0.3251036 | 1.537421 | 0.01267166 | -0.474161 | 0.2206253 | 1.978301 | 0.6668693 | 0.545115 | 0.8479431 |
| ENSG00000151322.18 | NPAS3 | -2.993637 | 0.01158252 | -0.175128 | 0.4493532 | 0.37204 | 0.5290845 | 1.332197 | 0.01773501 | 2.391853 | 0.15985 | 0.276914 | 0.752198 |
| ENSG00000151327.12 | FAM177A1 | 0.119286 | 0.2646568 | 0.093602 | 0.9187064 | -0.236203 | 0.02449485 | 0.032184 | 0.5911774 | 0.270742 | 0.5196843 | 0.580026 | 0.6710264 |
| ENSG00000151366.12 | NDUFC2 | 0.122519 | 0.1004496 | -0.09294 | 0.6527736 | -0.352163 | 3.16E-09 | -0.216523 | 0.000315064 | 0.043443 | 0.04279185 | -0.04776 | 0.5344421 |
| ENSG00000151422.12 | FER | -0.85152 | 0.8867323 | -0.454559 | 0.5110299 | 1.789169 | 0.001420521 | 0.115539 | 0.5576214 | 0.67339 | 0.6155434 | -0.210014 | 0.9147227 |
| ENSG00000151458.11 | ANKRD50 | -1.022593 | 0.6214382 | -0.335273 | 0.7456226 | -0.948141 | 0.0550398 | 0.614733 | 0.03582212 | 0.362601 | 0.2040744 | 1.504069 | 0.8479431 |
| ENSG00000151461.19 | UPF2 | 0.831547 | 0.9624741 | 0.940517 | 0.7212499 | 0.385678 | 0.01050817 | -0.055153 | 1 | 0.093262 | 0.7902863 | 3.340004 | 0.7554837 |
| ENSG00000151474.19 | FRMD4A | -0.280573 | 0.7736255 | 0.498438 | 0.906244 | 0.433875 | 0.6378982 | 1.646787 | 8.97E-06 | 0.511714 | 0.405494 | -2.805503 | 0.3956268 |
| ENSG00000151500.14 | THYN1 | 0.435716 | 0.8648814 | -0.184657 | 0.3423324 | 0.340284 | 0.8391286 | 0.05683 | 0.000407728 | -0.178114 | 0.06142639 | -0.745473 | 0.2768456 |
| ENSG00000151503.12 | NCAPD3 | 1.856809 | 0.9183025 | 0.045273 | 0.9080086 | 1.595305 | 0.01082418 | 1.402493 | 0.3467508 | 0.6202 | 0.8724107 | -1.696479 | 0.8949489 |
| ENSG00000151575.14 | TEX9 | 3.158819 | 1 | 0.390003 | 0.4892267 | 1.814166 | 0.01317874 | -0.15162 | 0.9271763 | -0.736007 | 0.5196843 | -2.34325 | 0.6885065 |
| ENSG00000151612.15 | ZNF827 | -1.20742 | 0.9591936 | 0.113293 | 0.6527736 | 0.336212 | 0.3288765 | 0.142962 | 0.001195669 | 1.963831 | 0.1687325 | -1.309931 | 0.8168211 |
| ENSG00000151640.12 | DPYSL4 | 0.154961 | 0.8006286 | -0.572723 | 0.8953649 | 0.76652 | 0.1394359 | 1.605093 | 0.2017005 | 1.772296 | 0.03224233 | -0.491819 | 0.7688175 |
| ENSG00000151692.14 | RNF144A | 1.124011 | 1 | -0.616503 | 0.7383998 | 1.170682 | 0.6341354 | 1.05664 | 0.01959123 | 0.606692 | 0.8686756 | -0.31517 | 1 |
| ENSG00000151746.13 | BICD1 | 0.876126 | 0.9438474 | 0.023933 | 0.6527736 | 0.69979 | 0.001289673 | 0.938222 | 0.02445851 | 0.307894 | 0.3974318 | -0.035612 | 0.6885065 |
| ENSG00000151806.13 | GUF1 | 0.741759 | 0.6014343 | -0.16574 | 1 | 1.412675 | 0.182889 | 2.450981 | 0.01317636 | -0.620923 | 0.5240903 | 0.503578 | 0.7554837 |
| ENSG00000151835.13 | SACS | -1.016744 | 0.6956464 | -0.163827 | 0.5171407 | 0.361647 | 0.208236 | 1.23174 | 0.04772157 | 1.395805 | 0.02228207 | 0.182608 | 0.9602719 |
| ENSG00000151846.8 | PABPC3 | 0.630596 | 0.6570323 | 1.556771 | 0.1940102 | -2.023018 | 3.58E-06 | 0.799299 | 0.945034 | 0.575968 | 0.6913898 | -1.179569 | 0.6885065 |
| ENSG00000151892.14 | GFRA1 | 1.765886 | 0.4547783 | -1.300004 | 0.8703522 | -0.484881 | 0.8100781 | 1.696961 | 0.003936151 | 1.015501 | 0.2138958 | 1.074828 | 1 |
| ENSG00000151914.18 | DST | 0.37379 | 0.5568401 | -0.205572 | 0.03901314 | 0.460527 | 0.000101817 | 1.539226 | 3.63E-12 | 0.69668 | 0.0623771 | 0.501005 | 0.8167193 |
| ENSG00000151929.9 | BAG3 | -1.739612 | 0.8587992 | 0.101513 | 1 | 0.452996 | 0.7305312 | 0.179562 | 0.6650041 | 3.115009 | 0.02724457 | -0.742481 | 0.9777201 |
| ENSG00000152061.22 | RABGAP1L | -1.683517 | 0.563505 | -0.624025 | 0.7335559 | 1.150211 | 0.2113831 | 2.337114 | 0.000198834 | -0.117022 | 0.9659049 | -2.204765 | 0.6885065 |
| ENSG00000152082.13 | MZT2B | -1.040737 | 0.08197923 | -0.288091 | 0.02922882 | -1.460025 | 1.01E-31 | -0.117905 | 0.007056223 | 0.664112 | 0.2179726 | -0.155145 | 0.5215899 |
| ENSG00000152102.17 | FAM168B | -1.168622 | 0.5428806 | -0.071562 | 0.9767342 | 0.150765 | 0.08654144 | 0.981058 | 0.003530003 | 0.612738 | 0.3081469 | -0.711371 | 0.5339044 |
| ENSG00000152127.8 | MGAT5 | 0.719341 | 0.7736255 | 0.285856 | 0.7907123 | 0.516705 | 0.7133122 | 0.182488 | 0.2122728 | 1.900661 | 0.004043844 | 0.495948 | 0.9638718 |
| ENSG00000152208.11 | GRID2 | -0.829497 | 0.6792949 | 0.499952 | 1 | 3.133253 | 0.000140916 | 0.600998 | 0.9311726 | 2.202856 | 0.07529074 | -2.467991 | 0.6885065 |
| ENSG00000152217.16 | SETBP1 | -0.541088 | 0.7795461 | 0.686636 | 0.6527736 | 1.707647 | 0.05667349 | 2.071237 | 0.01153566 | 1.901012 | 0.09276866 | -0.084073 | 0.7392448 |
| ENSG00000152234.15 | ATP5A1 | -0.652688 | 0.1268124 | -0.029482 | 0.6414243 | 0.086636 | 0.672594 | 0.279563 | 0.2335397 | -0.068202 | 0.001779951 | 0.038739 | 0.4192265 |
| ENSG00000152291.13 | TGOLN2 | -0.989035 | 0.6075174 | 0.206472 | 0.6527736 | 0.424144 | 0.005984437 | 0.813424 | 0.5288736 | 1.132584 | 0.2226329 | 1.301592 | 0.8949489 |
| ENSG00000152332.15 | UHMK1 | -0.900417 | 0.7736255 | -0.182799 | 0.6527736 | 0.442334 | 0.000773228 | 2.777885 | 0.001720137 | 0.287887 | 0.8385425 | -1.385349 | 0.6885065 |
| ENSG00000152377.13 | SPOCK1 | 0.064409 | 0.6214382 | -1.486769 | 0.2832272 | 0.515216 | 0.8061885 | 0.720296 | 0.000243547 | 1.094913 | 0.898446 | -0.616269 | 0.02166254 |
| ENSG00000152402.10 | GUCY1A2 | 1.150482 | 0.7679975 | -0.092941 | 0.8037648 | 1.33327 | 0.07264285 | 0.878638 | 0.01150633 | 2.478362 | 0.3911206 | -1.566041 | 0.8932168 |
| ENSG00000152404.15 | CWF19L2 | 1.379915 | 0.6214382 | -0.076541 | 0.791793 | 1.236155 | 0.02251206 | 0.215033 | 0.1557388 | 1.25013 | 0.9342997 | 0.715292 | 0.9147227 |
| ENSG00000152413.14 | HOMER1 | -0.434852 | 0.8434435 | -0.297638 | 0.5531715 | 0.716766 | 0.5290845 | 1.215258 | 0.001509717 | 0.799436 | 0.9376555 | -0.790011 | 0.8883513 |
| ENSG00000152492.13 | CCDC50 | 0.681231 | 0.6214382 | -0.3469 | 0.6527736 | -0.040868 | 0.353211 | 1.13922 | 0.04148071 | 0.897026 | 0.3974318 | -1.497929 | 1 |
| ENSG00000152495.10 | CAMK4 | -2.618813 | 0.4764974 | 0.967123 | 0.6527736 | -0.473028 | 0.896092 | 3.823913 | 0.01472886 | 1.375317 | 0.3060685 | -1.461051 | 1 |
| ENSG00000152578.12 | GRIA4 | 0.690263 | 0.4267377 | -0.317707 | 0.5953732 | 0.591459 | 0.5290845 | 1.405213 | 0.000407235 | 0.229735 | 0.2947305 | -2.121366 | 0.8856541 |
| ENSG00000152583.12 | SPARCL1 | -0.423029 | 9.65E-05 | 0.001833 | 0.9153026 | 0.23714 | 0.8922159 | -0.097261 | 2.12E-05 | -1.476065 | 0.01663105 | -0.669854 | 0.6885065 |
| ENSG00000152620.12 | NADK2 | -0.7812 | 1 | 0.00712 | 0.217888 | -0.178956 | 0.7605316 | 0.790691 | 0.1104339 | 2.820219 | 0.02878878 | 1.725503 | 0.6885065 |
| ENSG00000152795.17 | HNRNPDL | -0.956027 | 0.06050977 | -0.11869 | 0.1573875 | 0.680065 | 2.28E-08 | 0.03925 | 0.6590945 | 0.279137 | 0.08892368 | 0.33575 | 0.5344421 |
| ENSG00000152926.14 | ZNF117 | 1.742954 | 0.714717 | -0.921233 | 0.2166933 | 1.518177 | 0.0323305 | 1.683531 | 0.00025815 | 1.456284 | 0.00105812 | 1.192547 | 0.6885065 |
| ENSG00000152952.11 | PLOD2 | 1.013456 | 0.6214382 | 0.493545 | 0.002021575 | 0.460669 | 0.3679569 | 1.214134 | 0.8108755 | 0.357619 | 0.5183105 | -0.885439 | 0.8489889 |
| ENSG00000152953.12 | STK32B | 0.340336 | 0.8006286 | -0.356641 | 0.5953732 | 0.864566 | 0.5262128 | 1.935339 | 0.001623033 | 0.929726 | 0.5196843 | 0.648376 | 0.8349295 |
| ENSG00000152990.13 | ADGRA3 | -0.069515 | 0.8006286 | 0.037248 | 0.6239827 | 0.909595 | 0.7807636 | 0.382521 | 0.01305801 | 1.10561 | 0.1401461 | -1.47677 | 0.9391913 |
| ENSG00000153006.14 | SREK1IP1 | -2.740466 | 0.1985849 | 0.232293 | 1 | 0.418168 | 0.000906455 | 1.051071 | 0.1783688 | 0.921787 | 0.8159852 | -0.615766 | 0.6829816 |
| ENSG00000153012.11 | LGI2 | -0.065015 | 0.9047689 | -2.582031 | 0.09707294 | -2.672906 | 0.5290845 | 2.110435 | 0.006293281 | 0.875948 | 0.5196843 | -2.452678 | 0.6517664 |
| ENSG00000153037.13 | SRP19 | -1.003694 | 0.9738101 | -0.19455 | 0.8200822 | 0.209907 | 0.640899 | 0.588879 | 0.3468995 | -0.150253 | 0.03653915 | 0.915785 | 0.8479431 |
| ENSG00000153140.8 | CETN3 | -0.05243 | 1 | 0.560928 | 0.9177385 | 0.106814 | 0.5290845 | -0.028747 | 0.4955519 | 0.011318 | 0.02275793 | -1.809204 | 0.5915777 |
| ENSG00000153147.5 | SMARCA5 | -0.659603 | 0.6014343 | 0.2025 | 0.6527736 | 0.551687 | 4.04E-10 | 0.575852 | 0.3157928 | 0.205809 | 0.2495094 | -0.450613 | 0.2068179 |
| ENSG00000153187.16 | HNRNPU | -0.483876 | 0.8739908 | -0.082282 | 0.501917 | 0.402869 | 0.01649772 | 1.220997 | 1.26E-08 | 0.586413 | 0.5757457 | -0.003503 | 0.6885065 |
| ENSG00000153201.15 | RANBP2 | -0.967728 | 0.6897197 | 0.255341 | 0.8902958 | 0.958829 | 0.03418657 | 0.750587 | 1.36E-05 | -0.39182 | 0.8001666 | -0.035313 | 0.6829816 |
| ENSG00000153207.14 | AHCTF1 | -0.314678 | 0.6214382 | -0.547184 | 0.8263112 | 0.17033 | 0.4055474 | 0.531337 | 0.01120092 | 1.635515 | 0.1162536 | 0.605799 | 0.8167193 |
| ENSG00000153253.15 | SCN3A | 0.766795 | 0.5533102 | -0.071179 | 0.9330407 | 0.11457 | 0.2807066 | 1.542379 | 0.000100542 | 1.975556 | 0.02278366 | -0.486264 | 0.9037465 |
| ENSG00000153317.14 | ASAP1 | 0.928876 | 0.6214382 | 0.103263 | 0.6045862 | 0.998369 | 0.003433748 | 1.317724 | 0.02451119 | 0.526558 | 0.9386152 | -0.629282 | 0.5344421 |
| ENSG00000153395.9 | LPCAT1 | -1.282367 | 0.6214382 | 0.336566 | 0.5953732 | 0.060446 | 0.5290845 | 1.121539 | 2.73E-07 | 1.040644 | 0.003137382 | 0.071001 | 0.8479431 |
| ENSG00000153443.12 | UBALD1 | 0.842118 | 0.6214382 | -0.156857 | 0.7881421 | -0.025076 | 0.9790072 | 2.810493 | 0.267865 | 3.435286 | 0.01026642 | -0.355728 | 0.8949489 |
| ENSG00000153560.11 | UBP1 | -0.69646 | 0.6798783 | 0.039044 | 0.8217443 | 0.170956 | 0.9817784 | 0.75928 | 0.02455892 | 2.50336 | 4.27E-05 | 0.11196 | 0.8479431 |
| ENSG00000153561.12 | RMND5A | -1.469465 | 0.6214382 | -0.505622 | 0.8115344 | 0.338196 | 0.004787728 | 0.201173 | 0.3467508 | 0.539308 | 0.7377262 | -0.14306 | 0.9326505 |
| ENSG00000153707.15 | PTPRD | -1.418719 | 0.6214382 | -0.119718 | 0.5134635 | 1.386729 | 0.3118792 | 3.284114 | 0.3544716 | 1.831688 | 0.02023917 | -3.362433 | 0.6885065 |
| ENSG00000153774.8 | CFDP1 | 0.089939 | 0.8587992 | -0.304506 | 0.5010139 | -0.015069 | 0.5290845 | 0.01664 | 0.02321037 | -0.543223 | 0.000152282 | -0.532406 | 0.6885065 |
| ENSG00000153786.12 | ZDHHC7 | 3.591911 | 0.3946735 | -0.048459 | 0.8263112 | 0.828771 | 0.1915201 | 2.456465 | 0.005525534 | 1.246631 | 0.278178 | 0.014108 | 0.9161136 |
| ENSG00000153815.16 | CMIP | -0.624527 | 0.6927538 | -1.104182 | 0.2626761 | 0.051526 | 0.2806789 | 1.280851 | 0.02018338 | 0.93637 | 0.1196747 | 1.321352 | 0.6829816 |
| ENSG00000153827.13 | TRIP12 | 1.111349 | 0.8587992 | -0.370734 | 0.8392698 | 0.910335 | 0.009585109 | 0.7972 | 0.0805996 | 0.687426 | 0.4157048 | 0.151823 | 0.9612903 |
| ENSG00000153904.18 | DDAH1 | -0.061949 | 0.8006286 | -0.420946 | 0.7480928 | 0.383643 | 0.5194179 | 0.709111 | 0.002129222 | 1.443818 | 0.3183997 | 1.716144 | 0.6517664 |
| ENSG00000153914.15 | SREK1 | 0.471584 | 0.9047689 | -0.211681 | 0.3240152 | 0.415659 | 0.007815536 | 1.360311 | 3.53E-05 | 0.164809 | 1 | 0.610006 | 0.8949489 |
| ENSG00000153922.10 | CHD1 | 0.592816 | 0.928247 | -0.032702 | 0.7987376 | 0.615348 | 1.35E-07 | 0.647233 | 0.007052959 | 0.533286 | 0.6217705 | 0.304387 | 0.7787921 |
| ENSG00000153944.10 | MSI2 | 1.773365 | 0.6347536 | 0.398488 | 0.6527736 | 0.532112 | 4.47E-06 | 0.121002 | 0.09118013 | 0.30772 | 0.5722804 | 0.407337 | 0.8479431 |
| ENSG00000153956.15 | CACNA2D1 | 1.832071 | 0.9047689 | -0.271139 | 1 | 0.344747 | 0.7238584 | 2.062266 | 0.06727595 | 0.730384 | 0.01281183 | -0.653605 | 0.9531804 |
| ENSG00000153993.13 | SEMA3D | 0.042325 | 1 | -0.692557 | 1 | -1.42579 | 0.8100781 | -2.87189 | 0.3760083 | 2.361347 | 0.7377262 |  |  |
| ENSG00000154001.13 | PPP2R5E | -0.160621 | 0.8783629 | -0.210148 | 0.2851802 | 0.058904 | 0.313385 | 0.668653 | 0.01980759 | 0.310694 | 0.9275881 | -0.681378 | 0.6885065 |
| ENSG00000154122.12 | ANKH | 1.009322 | 0.06285358 | 0.193859 | 0.6580135 | 0.35911 | 0.965327 | 0.856517 | 0.02424875 | 3.02619 | 0.001848206 | -0.739979 | 0.6517664 |
| ENSG00000154174.7 | TOMM70A | -1.764788 | 0.6214382 | 0.077082 | 0.9577957 | 0.470913 | 0.2134736 | 0.670286 | 0.01223151 | 1.106852 | 0.322123 | -1.098303 | 0.9169962 |
| ENSG00000154277.12 | UCHL1 | 0.108903 | 0.9310946 | 0.009939 | 0.8263112 | 0.145685 | 0.1089077 | 0.027124 | 0.01846326 | -0.265943 | 0.002398345 | -0.861753 | 0.000373203 |
| ENSG00000154305.16 | MIA3 | 1.099969 | 0.3326528 | -0.188765 | 0.6527736 | 0.864135 | 0.000960436 | 0.401152 | 0.08651083 | -0.179732 | 0.840404 | -0.139471 | 0.6885065 |
| ENSG00000154310.16 | TNIK | 0.976452 | 0.6014343 | -0.094219 | 0.6527736 | 1.517668 | 0.06323885 | 1.412597 | 0.04611506 | 0.33131 | 0.2581597 | -0.834 | 0.9147227 |
| ENSG00000154359.12 | LONRF1 | 0.384907 | 0.8747535 | -0.250854 | 0.5821728 | -0.33428 | 0.8221252 | 1.002679 | 0.03761421 | 1.102136 | 0.1831888 | -2.666208 | 0.7901525 |
| ENSG00000154370.13 | TRIM11 | -1.232368 | 0.1364817 | -0.985859 | 0.6384858 | 0.309512 | 0.01483275 | 0.492492 | 0.0909425 | -0.011309 | 0.4826781 | 0.224449 | 0.960853 |
| ENSG00000154380.17 | ENAH | -0.521717 | 0.8297654 | -0.274585 | 0.1053664 | 1.333616 | 3.26E-07 | 1.026224 | 0.01181088 | 0.809353 | 0.4247809 | 0.64002 | 0.8479431 |
| ENSG00000154511.11 | FAM69A | -0.876221 | 0.8587992 | 1.111142 | 0.6514723 | 2.498223 | 0.3394407 | -2.006001 | 0.03028101 | 1.911141 | 0.7232762 | -0.365707 | 0.8479431 |
| ENSG00000154518.9 | ATP5G3 | -0.271423 | 0.7076572 | -1.030554 | 0.24796 | -0.016129 | 0.3095404 | -0.261427 | 0.007662836 | -0.529376 | 2.12E-05 | -0.125136 | 0.1092491 |
| ENSG00000154582.16 | TCEB1 | 0.440941 | 0.714717 | 0.040192 | 0.9104976 | -0.264919 | 6.92E-06 | -0.095436 | 0.0871018 | -0.033622 | 0.01073185 | -0.390435 | 0.3394425 |
| ENSG00000154640.14 | BTG3 | -0.873439 | 0.9146427 | -0.200955 | 0.3395022 | -0.588613 | 0.00095425 | -0.268234 | 0.001646819 | 0.158626 | 0.3974318 | 1.288067 | 0.9741465 |
| ENSG00000154654.14 | NCAM2 | 0.439234 | 0.8408841 | 0.35403 | 0.6527736 | 0.924233 | 0.50389 | 1.944538 | 0.03325559 | 1.363991 | 0.3013132 | -2.967628 | 1 |
| ENSG00000154723.12 | ATP5J | 0.013324 | 0.563505 | 0.063262 | 0.7257098 | -0.57209 | 1.18E-11 | -0.146014 | 0.003758342 | -0.265173 | 3.17E-06 | -0.494504 | 0.03227675 |
| ENSG00000154781.15 | CCDC174 | -4.671711 | 0.8587992 | 0.081136 | 0.864346 | 0.758879 | 0.001779562 | 0.335676 | 0.7144938 | 0.105238 | 0.2169837 | -1.609046 | 0.9614796 |
| ENSG00000154803.12 | FLCN | -0.451981 | 1 | -0.479756 | 0.6527736 | -0.461148 | 0.7090807 | 0.423339 | 0.03847618 | 0.554073 | 0.5491251 | 0.017753 | 0.9077431 |
| ENSG00000154845.15 | PPP4R1 | 0.255837 | 0.6214382 | 0.340255 | 0.5953732 | 0.984512 | 0.6608038 | 0.649013 | 0.2520989 | 1.112468 | 0.007605515 | 0.340165 | 0.6885065 |
| ENSG00000154856.12 | APCDD1 | -1.924049 | 0.714717 | -0.712256 | 0.5519081 | -1.232197 | 0.02213292 | 1.150749 | 0.01064005 | 0.410925 | 0.3987518 | -2.841157 | 0.8385963 |
| ENSG00000154978.12 | VOPP1 | -0.13438 | 0.9223024 | -0.009924 | 0.1953788 | -0.388323 | 0.3569457 | 0.349351 | 0.01279765 | 0.8394 | 0.1053151 | 0.357649 | 0.8208081 |
| ENSG00000155090.14 | KLF10 | 1.414972 | 0.3903167 | -0.181314 | 0.5569064 | -0.326223 | 0.8247492 | 0.284693 | 0.008439201 | 1.180572 | 0.05504727 | -1.419576 | 0.6667259 |
| ENSG00000155100.10 | OTUD6B | -0.428228 | 0.959823 | -0.543381 | 0.6527736 | 0.935744 | 0.1011529 | 1.430526 | 0.01816429 | 1.290897 | 0.547662 | -1.20546 | 0.8479431 |
| ENSG00000155111.14 | CDK19 | -2.123574 | 0.3946735 | -0.676153 | 0.6527736 | 0.49924 | 0.3452689 | 1.983277 | 0.1044159 | 1.386314 | 0.01236688 | -1.521673 | 0.6885065 |
| ENSG00000155115.6 | GTF3C6 | -0.457922 | 0.6760326 | 0.008093 | 0.5953732 | 0.3902 | 0.04122184 | 0.152501 | 0.4401092 | -1.023142 | 0.01038567 | 1.364256 | 0.96412 |
| ENSG00000155304.5 | HSPA13 | 0.01436 | 1 | 0.175023 | 0.501917 | 1.076719 | 0.02168833 | 1.292257 | 0.01989019 | 0.152101 | 0.1558437 | -0.455236 | 0.9264343 |
| ENSG00000155330.9 | C16orf87 | -1.178086 | 0.3582986 | -0.460922 | 0.168108 | 0.840417 | 0.009692558 | -0.075661 | 0.01118812 | 0.572464 | 0.2726807 | 1.832832 | 0.9147227 |
| ENSG00000155366.16 | RHOC | -0.048157 | 0.714717 | 0.189349 | 0.7383998 | 0.121371 | 0.8042144 | -0.081835 | 0.001255306 | 0.140552 | 0.02212756 | -0.066169 | 0.6885065 |
| ENSG00000155368.16 | DBI | -0.973907 | 1 | -0.431715 | 1.25E-09 | -0.166427 | 0.05282253 | -0.1869 | 3.30E-06 | -0.087358 | 0.01434427 | -0.130256 | 0.2846541 |
| ENSG00000155380.11 | SLC16A1 | -0.187906 | 0.8964996 | 0.290351 | 0.08794836 | 0.367406 | 0.5729508 | 1.256139 | 0.03560962 | 1.510861 | 0.6145561 | -0.260212 | 0.6901786 |
| ENSG00000155393.12 | HEATR3 | -1.758435 | 0.714717 | 0.816351 | 0.8819808 | 1.086259 | 0.6928639 | 3.227923 | 0.001791764 | 2.462109 | 0.00513524 | -1.346992 | 1 |
| ENSG00000155506.16 | LARP1 | -1.04414 | 0.529676 | -0.367089 | 0.2709731 | 0.331917 | 0.5290845 | 0.789493 | 0.001265981 | 1.064793 | 0.03711912 | 0.121364 | 0.6885065 |
| ENSG00000155545.19 | MIER3 | 1.132294 | 0.8867323 | 0.780047 | 0.8895498 | 0.855561 | 0.1016264 | 0.264787 | 0.6558628 | 2.151351 | 0.01177698 | -0.693163 | 0.868568 |
| ENSG00000155561.14 | NUP205 | 1.698078 | 0.529676 | -0.013921 | 0.8519642 | 0.485324 | 0.8100781 | 1.666923 | 0.000125549 | 1.371424 | 0.1384387 | -0.282145 | 0.6885065 |
| ENSG00000155755.18 | TMEM237 | 0.179369 | 0.8587992 | 0.02469 | 1 | -0.449569 | 0.5864994 | 0.555182 | 0.008737569 | 1.36035 | 0.3628287 | 0.380767 | 0.6885065 |
| ENSG00000155876.5 | RRAGA | -0.405699 | 0.7259912 | 0.546944 | 0.8304896 | -1.119776 | 1.96E-06 | -0.075037 | 0.09950505 | 1.555047 | 0.5196843 | -1.713774 | 0.6885065 |
| ENSG00000155957.17 | TMBIM4 | -0.214055 | 0.7135744 | 0.272938 | 0.8263112 | -0.895017 | 2.24E-05 | 1.022569 | 0.549329 | -0.127796 | 0.116144 | -0.284467 | 0.9726757 |
| ENSG00000155959.10 | VBP1 | -0.461717 | 0.9610968 | -0.144738 | 0.7020147 | 0.766477 | 0.06202009 | -0.308619 | 0.03519259 | -0.188935 | 0.03621229 | -0.818849 | 0.6675976 |
| ENSG00000155961.4 | RAB39B | -1.715052 | 0.4406957 | 0.112035 | 0.7688491 | 0.716449 | 0.0104496 | 1.085169 | 0.8241581 | 0.442283 | 0.97225 | -0.47455 | 0.6885065 |
| ENSG00000155970.11 | MICU3 | -2.656453 | 0.6562038 | 0.930648 | 0.9597479 | 1.168601 | 0.5290845 | 1.244109 | 0.007145323 | 1.393468 | 0.665227 | -1.125634 | 0.7780985 |
| ENSG00000155980.11 | KIF5A | -1.591844 | 0.4375066 | -0.318174 | 0.918309 | -0.700141 | 0.3695136 | 1.729272 | 8.86E-07 | 0.641573 | 0.104272 | -0.390558 | 0.6490419 |
| ENSG00000156030.12 | ELMSAN1 | -1.758164 | 0.6014343 | -0.149925 | 0.5010139 | 0.396849 | 0.5374158 | 2.024855 | 0.01116345 | 1.666446 | 0.1146485 | 0.034741 | 0.9100805 |
| ENSG00000156052.10 | GNAQ | -0.852628 | 0.9183025 | -0.055768 | 0.1290746 | 0.236623 | 0.1922832 | 0.719664 | 0.01124605 | 0.926352 | 0.4760227 | 1.214012 | 1 |
| ENSG00000156103.15 | MMP16 | -0.090018 | 0.6214382 | -0.247895 | 0.5670497 | 0.399026 | 0.001010625 | 1.902217 | 9.59E-07 | 0.854139 | 0.2532208 | -0.274602 | 0.6885065 |
| ENSG00000156113.20 | KCNMA1 | 0.213428 | 1 | -0.25394 | 0.5394427 | -0.51676 | 0.7353381 | 1.513649 | 0.002613865 | 1.676046 | 0.5196843 | 2.024452 | 0.6885065 |
| ENSG00000156136.9 | DCK | -1.259789 | 0.7795461 | 0.644368 | 0.6527736 | 0.546308 | 0.8544426 | 0.246078 | 0.00072658 | 0.023856 | 0.677167 | -0.56275 | 0.7880595 |
| ENSG00000156298.12 | TSPAN7 | 0.093779 | 0.04098339 | -0.591869 | 0.007219487 | 0.363953 | 0.5258213 | 0.249974 | 0.9204585 | 0.224868 | 0.6344744 | -0.045039 | 0.4578379 |
| ENSG00000156313.12 | RPGR | 0.683682 | 0.9289591 | 0.993589 | 0.8920257 | 0.922678 | 0.03873586 | 0.875184 | 0.09711314 | 0.823726 | 0.8021966 | -0.487665 | 0.8708442 |
| ENSG00000156411.9 | C14orf2 | 1.13676 | 0.007347962 | 0.009492 | 0.5953732 | -0.229833 | 0.0104496 | 0.441557 | 0.7047221 | -0.524637 | 0.001848206 | 0.123183 | 0.6885065 |
| ENSG00000156453.13 | PCDH1 | 0.059115 | 0.739831 | 0.973136 | 0.03241867 | 0.949899 | 0.6287629 | 1.084183 | 0.3113668 | 0.383916 | 0.9208937 | -0.089087 | 0.5066477 |
| ENSG00000156467.9 | UQCRB | -0.535108 | 0.5084093 | -0.430182 | 0.001905827 | 0.123852 | 0.9342469 | -0.03749 | 0.1040467 | -0.129762 | 0.02800883 | -0.185318 | 0.04261865 |
| ENSG00000156504.16 | FAM122B | 2.153507 | 0.4764974 | 0.484917 | 0.9790409 | 0.537702 | 0.1223306 | -0.434419 | 0.3530282 | 2.513546 | 0.008813299 | 0.373717 | 0.8378323 |
| ENSG00000156508.17 | EEF1A1 | -0.672054 | 6.93E-18 | 0.120372 | 0.004581147 | 0.593696 | 6.58E-43 | 0.273636 | 0.02516068 | -0.058389 | 0.07002445 | 0.015178 | 0.8472204 |
| ENSG00000156587.15 | UBE2L6 | -0.206812 | 0.9734775 | -0.146038 | 0.6899483 | -0.382824 | 1.16E-05 | -0.627788 | 0.08188372 | -0.250117 | 0.02444352 | 0.278118 | 1 |
| ENSG00000156599.10 | ZDHHC5 | 2.726827 | 0.5694264 | -0.22212 | 0.4886958 | 0.486207 | 0.424857 | -0.229568 | 0.6174179 | 3.497882 | 0.02733575 | 0.396308 | 0.7767294 |
| ENSG00000156671.12 | SAMD8 | -0.795094 | 0.4692975 | 0.161433 | 0.9754015 | 1.35882 | 0.00058661 | 0.935662 | 0.655488 | 1.147246 | 0.9056171 | 1.109738 | 0.6885065 |
| ENSG00000156928.4 | MALSU1 | 0.327665 | 1 | 0.130624 | 0.9848758 | 0.026861 | 0.02701106 | 0.061675 | 0.70099 | 0.762329 | 0.8901278 | -0.963966 | 0.6185816 |
| ENSG00000156931.15 | VPS8 | -1.381744 | 0.822435 | -0.279814 | 0.9981323 | 0.84228 | 0.8264533 | 1.195233 | 0.03114126 | 0.80067 | 0.3796044 | 1.045647 | 0.8479431 |
| ENSG00000156976.14 | EIF4A2 | -0.734994 | 0.1004496 | 0.425117 | 0.3160414 | 0.416946 | 0.01796672 | 0.033098 | 0.8476371 | 0.45522 | 1 | 0.505197 | 0.6885065 |
| ENSG00000157020.17 | SEC13 | -0.437228 | 0.529676 | 0.29776 | 0.7361049 | 0.130396 | 0.8459315 | -0.072913 | 3.89E-06 | -0.940666 | 0.009870185 | 0.875765 | 0.936179 |
| ENSG00000157064.10 | NMNAT2 | 0.176111 | 1 | -0.322556 | 0.8928139 | 1.803196 | 0.3871766 | 1.552534 | 0.03364031 | -0.889173 | 0.6277673 | -2.254953 | 0.8949489 |
| ENSG00000157087.16 | ATP2B2 | 0.757422 | 0.8587992 | -1.896082 | 0.3795682 | 1.728326 | 0.04249714 | 1.529168 | 0.03727306 | 0.315996 | 0.9570975 |  |  |
| ENSG00000157106.16 | SMG1 | -0.853129 | 0.8862706 | -0.180771 | 0.6580135 | 0.09365 | 0.02831409 | 0.543946 | 0.001921065 | 0.677169 | 0.6496023 | 0.031466 | 0.9562147 |
| ENSG00000157193.14 | LRP8 | 0.105423 | 0.9531264 | 0.621125 | 0.7991916 | -0.812591 | 0.5290845 | 0.898537 | 1.36E-06 | 1.241369 | 0.1421028 | -0.851677 | 0.8754943 |
| ENSG00000157212.18 | PAXIP1 | -0.693786 | 0.6793885 | 0.282973 | 0.8281295 | 0.820174 | 0.2044962 | 2.119616 | 0.04843889 | 0.687354 | 0.4749013 | 1.358079 | 0.9524428 |
| ENSG00000157216.15 | SSBP3 | -0.284351 | 0.6214382 | 0.094757 | 0.77388 | 0.047853 | 0.7882906 | 1.343942 | 0.03928446 | 1.301972 | 0.008596796 | 0.873413 | 0.8856541 |
| ENSG00000157224.15 | CLDN12 | 2.293011 | 0.6214382 | 0.11364 | 0.9120034 | 0.55324 | 0.1696305 | 1.15333 | 0.03897443 | 0.849299 | 0.1923385 | 1.035672 | 0.9007097 |
| ENSG00000157259.6 | GATAD1 | -1.760931 | 0.004432297 | 0.764657 | 0.5821728 | -0.116448 | 0.2349576 | 1.553951 | 0.000548681 | 0.602102 | 0.7846335 | 1.861726 | 0.7370883 |
| ENSG00000157326.18 | DHRS4 | -0.189959 | 0.6214382 | 0.287603 | 0.6364102 | 0.811866 | 0.8102513 | -0.651031 | 0.0200881 | 0.000742 | 0.8788811 | -1.348672 | 0.8479431 |
| ENSG00000157350.12 | ST3GAL2 | 1.049406 | 0.6214382 | 0.749519 | 0.592336 | 0.982306 | 0.6295125 | 2.18168 | 0.01430877 | 0.556989 | 0.05408505 | 1.465415 | 1 |
| ENSG00000157450.15 | RNF111 | 1.681199 | 0.6214382 | 0.184176 | 0.8703522 | 0.639059 | 0.3522448 | 0.782204 | 0.03857076 | 1.676977 | 0.4761828 | 2.8629 | 0.3565625 |
| ENSG00000157500.10 | APPL1 | -0.771021 | 0.714717 | -0.576398 | 0.06013553 | 0.896674 | 0.00642867 | 1.289787 | 0.7004598 | 0.288377 | 0.566356 | -0.577891 | 0.6829816 |
| ENSG00000157540.19 | DYRK1A | -0.768238 | 0.9183025 | -0.335895 | 0.5150959 | 0.242164 | 0.7437308 | 0.809441 | 0.00692364 | 1.267697 | 0.02246628 | 1.076985 | 1 |
| ENSG00000157557.11 | ETS2 | 0.870988 | 0.6214382 | 0.082135 | 0.7984762 | 0.592209 | 0.7354314 | -0.184484 | 0.06674489 | 3.315007 | 0.000301829 | 3.484909 | 0.9532204 |
| ENSG00000157625.15 | TAB3 | 1.529405 | 0.8006286 | -1.024047 | 0.6200225 | 0.954201 | 0.4405609 | 0.112597 | 0.001546506 | 0.661026 | 0.1602882 | -3.684985 | 1 |
| ENSG00000157637.12 | SLC38A10 | 0.258614 | 1 | 0.475345 | 0.7855215 | -1.520467 | 0.04991908 | 0.059961 | 0.7806983 | 1.8382 | 0.195661 | 1.528136 | 0.8479431 |
| ENSG00000157741.14 | UBN2 | -1.481058 | 0.6214382 | 0.163203 | 1 | 0.607163 | 0.01230171 | 2.239826 | 0.000956777 | 1.079547 | 0.0124237 | -0.12149 | 0.6939365 |
| ENSG00000157766.15 | ACAN | 1.25912 | 0.6214382 | -2.705075 | 0.6527736 | -2.31137 | 0.8454005 | 0.881188 | 0.04242491 | 2.636852 | 0.2187443 | 2.582494 | 0.9147227 |
| ENSG00000157827.19 | FMNL2 | -0.438572 | 0.8587992 | 0.200504 | 0.2388066 | -0.002082 | 0.8026663 | 1.158766 | 0.000368412 | 2.298346 | 0.00052597 | -0.752337 | 0.7694423 |
| ENSG00000157837.15 | SPPL3 | 0.299823 | 0.9489851 | -0.461008 | 0.1712543 | -0.434293 | 0.9968942 | 0.575934 | 0.01020509 | 0.691463 | 0.2047158 | -0.560077 | 0.6885065 |
| ENSG00000157851.16 | DPYSL5 | -0.649892 | 0.6014343 | -0.062952 | 0.7105285 | 0.0911 | 0.8220688 | 1.678022 | 3.78E-05 | 0.559175 | 0.6171067 | -0.27744 | 0.960853 |
| ENSG00000157916.18 | RER1 | 0.422548 | 0.529676 | 0.030092 | 0.7382578 | -0.297563 | 0.000156275 | 1.162189 | 0.9788705 | 1.468183 | 0.3013132 | 1.024951 | 0.6490419 |
| ENSG00000157933.9 | SKI | -1.132824 | 0.6014343 | -0.31974 | 0.2588352 | -0.521623 | 0.02213292 | 0.810484 | 0.0371867 | 1.372423 | 0.09409813 | -0.649056 | 0.8479431 |
| ENSG00000157985.18 | AGAP1 | -0.1691 | 1 | 0.138073 | 0.9439537 | -0.139199 | 0.8616087 | 1.635136 | 0.00035884 | 1.301877 | 0.3974318 | -0.232743 | 0.8181167 |
| ENSG00000158042.8 | MRPL17 | -0.892893 | 0.563505 | -0.024929 | 0.8263112 | -0.116444 | 0.02521457 | -0.022846 | 0.1713484 | -0.399583 | 0.1169767 | 1.583835 | 0.9161136 |
| ENSG00000158109.14 | TPRG1L | 2.424663 | 1 | 0.260607 | 0.6527736 | -1.332828 | 0.09058387 | 3.399144 | 0.002572669 | 0.403429 | 0.9549135 | 2.975868 | 0.9381693 |
| ENSG00000158195.10 | WASF2 | -0.516572 | 1 | 0.036852 | 0.6834954 | 0.714569 | 0.01800539 | -0.103441 | 0.5259364 | 1.637048 | 0.621723 | 1.617685 | 0.6829816 |
| ENSG00000158290.16 | CUL4B | 0.777583 | 0.9183025 | -0.95238 | 0.04907665 | 0.060941 | 0.03414511 | 0.150417 | 0.1503671 | 0.69148 | 0.5976635 | -0.050387 | 0.8573709 |
| ENSG00000158301.18 | GPRASP2 | -2.683742 | 0.5711449 | 0.420734 | 0.8263112 | -0.595884 | 0.4064599 | 2.828003 | 0.0047097 | -0.122383 | 0.7494748 | 2.005889 | 0.9270338 |
| ENSG00000158321.15 | AUTS2 | -1.193784 | 0.6214382 | -0.011374 | 0.8703522 | 0.39727 | 0.4377153 | 2.035407 | 1.39E-06 | 0.595937 | 0.1585606 | -1.221371 | 0.4578379 |
| ENSG00000158417.10 | EIF5B | -1.460643 | 1.55E-08 | -0.131616 | 0.05678909 | 1.06764 | 2.01E-18 | 0.794853 | 0.3405933 | -0.155213 | 0.009713756 | -0.300174 | 0.3746505 |
| ENSG00000158445.8 | KCNB1 | -0.743087 | 1 | -2.185932 | 0.7383998 | 0.837151 | 0.6827367 | 3.201114 | 0.000315064 | 3.660907 | 3.02E-09 | 2.503094 | 0.6829816 |
| ENSG00000158470.5 | B4GALT5 | 0.929962 | 1 | 0.028311 | 0.9068973 | 0.482537 | 0.02837269 | 0.933531 | 0.03164253 | 0.71676 | 0.5196843 | 0.500104 | 0.8963493 |
| ENSG00000158526.7 | TSR2 | -0.752704 | 0.9085559 | 0.897436 | 0.3121527 | -0.028047 | 0.5290845 | 0.980637 | 0.4325994 | -0.520422 | 0.01972457 | 0.434329 | 0.930152 |
| ENSG00000158528.11 | PPP1R9A | -1.252205 | 0.3946735 | 0.638969 | 0.978688 | 2.234789 | 4.68E-06 | 0.542081 | 0.269482 | -1.040867 | 0.2165533 | 2.231224 | 0.8479431 |
| ENSG00000158545.15 | ZC3H18 | -0.223007 | 0.8867323 | 0.363918 | 0.8717486 | 0.089007 | 0.139989 | 1.190476 | 0.04333874 | 1.050493 | 0.05598594 | -0.457387 | 0.9352001 |
| ENSG00000158615.8 | PPP1R15B | -1.143094 | 0.6792949 | 0.362263 | 0.6527736 | 0.252189 | 0.4008897 | 1.132787 | 0.04785592 | 0.813796 | 0.007103428 | 0.565624 | 0.6490419 |
| ENSG00000158636.16 | C11orf30 | -0.200742 | 0.8006286 | 0.045262 | 0.6660806 | 0.67723 | 0.1388313 | 1.644879 | 0.02685963 | 0.416508 | 0.6007569 | -2.301954 | 0.6885065 |
| ENSG00000158669.11 | GPAT4 | 0.017316 | 0.6642509 | -0.067907 | 0.7276361 | 0.974916 | 0.4109319 | 0.859087 | 0.000253987 | 0.800828 | 0.5845416 | -0.464039 | 0.7979751 |
| ENSG00000158711.13 | ELK4 | -0.289298 | 0.9421205 | -0.338935 | 0.6527736 | 0.364441 | 0.5290845 | 1.944835 | 0.02029956 | 1.924807 | 0.06437961 | -1.136732 | 0.9969118 |
| ENSG00000158773.14 | USF1 | 0.202127 | 0.7736255 | -0.431401 | 0.6200225 | 0.181595 | 0.4775324 | -0.466182 | 0.3467508 | 2.604813 | 0.000280026 | -0.822415 | 0.9526606 |
| ENSG00000158805.11 | ZNF276 | 0.199782 | 0.8454066 | -0.31269 | 0.8217443 | -0.052591 | 0.003743584 | 1.045157 | 0.09671702 | 1.492904 | 0.01562084 | 0.980495 | 0.6885065 |
| ENSG00000158863.21 | FAM160B2 | -0.953363 | 0.6214382 | 0.641336 | 0.6208241 | -0.560583 | 0.7437308 | 1.421062 | 0.2456134 | 2.443814 | 0.04558844 | 0.955709 | 0.9147227 |
| ENSG00000158941.16 | CCAR2 | -0.634237 | 0.9485354 | 0.576771 | 0.6527736 | 0.854844 | 0.7354314 | 1.459142 | 0.000446152 | 1.541837 | 0.2795958 | 2.665204 | 0.7958826 |
| ENSG00000159079.18 | C21orf59 | -1.763503 | 0.07604642 | -0.140448 | 0.3583128 | -0.112163 | 0.04753476 | -0.291302 | 0.04615682 | -0.345256 | 0.01055545 | 0.794026 | 0.6855855 |
| ENSG00000159086.14 | PAXBP1 | -0.390343 | 0.8867323 | 0.203796 | 0.3045812 | -0.457516 | 0.5290845 | 0.356738 | 0.2655956 | 1.189469 | 0.003778489 | 0.24253 | 0.8949489 |
| ENSG00000159128.14 | IFNGR2 | 0.595566 | 0.8006286 | -0.58223 | 0.5010139 | -0.622569 | 0.3118792 | 0.709662 | 0.03479536 | 1.631279 | 0.1823192 | 0.684957 | 0.8479431 |
| ENSG00000159140.17 | SON | -0.64291 | 0.7821041 | 0.115683 | 1 | 0.061444 | 0.000927701 | 0.793331 | 0.000555177 | 0.83315 | 0.2812231 | 0.370206 | 0.859732 |
| ENSG00000159147.17 | DONSON | -2.97913 | 0.8006286 | -0.313536 | 0.6527736 | -1.425386 | 0.5337564 | 0.049616 | 0.4919368 | 2.271799 | 0.0292479 | 0.37088 | 0.6885065 |
| ENSG00000159164.9 | SV2A | 0.649187 | 0.01796694 | -0.496941 | 0.3212364 | 0.186724 | 0.7437308 | 0.734191 | 0.1620965 | 0.234327 | 0.4123379 | -0.517579 | 0.9127491 |
| ENSG00000159199.13 | ATP5G1 | 0.05308 | 0.984876 | -0.170202 | 0.4011918 | -0.748788 | 1.50E-08 | 0.144126 | 0.1093889 | 0.522355 | 0.7594537 | -0.2348 | 0.01991945 |
| ENSG00000159202.17 | UBE2Z | 1.72065 | 0.4215509 | 0.262092 | 0.6459834 | 0.949182 | 0.1778406 | 1.257368 | 0.00964678 | 0.510864 | 0.4853775 | -1.649457 | 0.9270789 |
| ENSG00000159216.18 | RUNX1 | -2.373877 | 0.3946735 | 0.245154 | 0.6527736 | -0.652192 | 1 | 1.603508 | 2.57E-06 | 0.634688 | 0.6830899 | 0.682858 | 0.5733051 |
| ENSG00000159228.12 | CBR1 | -0.38517 | 0.8434435 | 0.139808 | 0.8785221 | 0.096771 | 0.5290845 | -0.431091 | 0.001345975 | -0.641464 | 0.008945948 | 0.106987 | 1 |
| ENSG00000159335.15 | PTMS | -1.339562 | 0.563505 | 0.275299 | 0.6628666 | -1.665594 | 1.45E-12 | 1.418188 | 3.11E-19 | 1.362758 | 1.48E-11 | -0.189541 | 0.1508459 |
| ENSG00000159377.10 | PSMB4 | -0.080776 | 0.8006286 | 0.144446 | 0.7383998 | -0.498274 | 0.00254375 | -0.131368 | 0.01124156 | -0.605325 | 5.99E-05 | 0.273406 | 0.3565625 |
| ENSG00000159461.14 | AMFR | 0.301789 | 0.6014343 | -0.314844 | 0.6532996 | 0.173144 | 0.8994235 | 1.041539 | 0.03434133 | 1.577395 | 0.1993336 | 2.116731 | 0.6767422 |
| ENSG00000159579.13 | RSPRY1 | 0.32128 | 0.6014343 | 0.238219 | 0.8101094 | 1.406913 | 0.000140916 | 1.157573 | 0.1698448 | 0.411604 | 0.851927 | 1.58428 | 0.961427 |
| ENSG00000159592.10 | GPBP1L1 | 0.28601 | 0.8006286 | -0.138942 | 0.8895089 | -0.313215 | 0.2013354 | 0.993534 | 0.00015275 | 0.705662 | 0.2534005 | -1.050533 | 0.9531804 |
| ENSG00000159596.6 | TMEM69 | -0.922555 | 0.7736255 | -0.601889 | 0.6109044 | 0.33726 | 0.7623935 | -1.947855 | 0.0001486 | 0.920514 | 0.7377262 | 1.426696 | 0.8479431 |
| ENSG00000159692.15 | CTBP1 | 0.472843 | 0.03176252 | -0.220912 | 0.6527736 | -0.141049 | 0.4064599 | 1.232505 | 4.91E-05 | 0.488622 | 0.09515997 | -0.428629 | 0.6885065 |
| ENSG00000159720.11 | ATP6V0D1 | 0.478063 | 0.6014343 | 0.361015 | 0.5296018 | -0.055616 | 0.7221803 | 0.676238 | 1 | -0.96988 | 0.000556679 | 1.353312 | 0.826754 |
| ENSG00000159784.17 | FAM131B | -1.362584 | 0.7736255 | -0.542688 | 0.3651168 | 0.599063 | 0.6174506 | 0.530167 | 0.03997638 | 1.059136 | 0.8021966 | 0.380085 | 0.9581114 |
| ENSG00000159840.15 | ZYX | -0.368483 | 0.6214382 | -0.127645 | 0.8649393 | 0.23323 | 0.004221161 | 0.831313 | 0.001044563 | 0.010861 | 0.682928 | -0.031067 | 0.8432645 |
| ENSG00000160007.17 | ARHGAP35 | 1.798211 | 0.6214382 | -0.076937 | 0.8953649 | -0.07935 | 0.8218004 | 0.88772 | 0.000328298 | 1.942233 | 0.000260653 | -1.549968 | 0.6829816 |
| ENSG00000160049.11 | DFFA | -0.384298 | 0.59287 | -0.107656 | 0.9035383 | 0.537348 | 0.02837269 | 0.575557 | 0.234675 | 0.208956 | 0.8982465 | -0.087217 | 0.8479431 |
| ENSG00000160087.20 | UBE2J2 | -0.077077 | 1 | 0.03831 | 0.6974771 | -1.628713 | 2.36E-12 | 0.295208 | 0.9317757 | 0.623453 | 0.1743517 | -0.812713 | 0.9264343 |
| ENSG00000160113.5 | NR2F6 | 0.05848 | 0.7563254 | 0.103019 | 0.9626414 | -0.891389 | 0.2022843 | 0.84497 | 0.1557388 | 1.495771 | 0.01102216 | -0.917183 | 0.926878 |
| ENSG00000160194.17 | NDUFV3 | -0.590201 | 0.529676 | -0.465623 | 0.03144751 | -0.445061 | 0.000311976 | 0.972721 | 0.7313084 | -0.324971 | 0.007372379 | 0.640277 | 0.8145966 |
| ENSG00000160201.11 | U2AF1 | -0.065714 | 0.9047689 | 0.360568 | 0.9618421 | 0.434848 | 0.5290845 | 3.743399 | 7.97E-06 | 2.134814 | 0.01038567 | -1.071612 | 0.6656984 |
| ENSG00000160213.5 | CSTB | -0.870744 | 0.529676 | 0.014682 | 0.8263112 | -0.653392 | 1.74E-06 | 0.306086 | 0.005117581 | -0.264394 | 0.04558844 | 1.2427 | 1 |
| ENSG00000160216.18 | AGPAT3 | -0.029614 | 0.9047689 | -0.253707 | 0.5296018 | -0.350412 | 0.85126 | 1.706054 | 0.1374552 | 2.490469 | 0.006167715 | 0.473382 | 0.6885065 |
| ENSG00000160229.11 | ZNF66 | 1.192286 | 0.8006286 | 0.085109 | 0.8217443 | 4.831336 | 0.02458539 | -0.704392 | 0.8472142 | 1.543747 | 0.3341786 | 0.249459 | 0.6885065 |
| ENSG00000160271.14 | RALGDS | 1.326595 | 0.8556787 | -0.677575 | 0.3565973 | 0.181165 | 0.6295125 | 2.019812 | 6.88E-06 | 1.035867 | 0.04058627 | -0.456425 | 0.9264343 |
| ENSG00000160293.16 | VAV2 | 0.633392 | 0.7736255 | -0.778642 | 0.592336 | -0.009667 | 0.5290845 | 0.7115 | 0.007284783 | 1.247369 | 0.2660195 | 0.153099 | 0.6885065 |
| ENSG00000160294.10 | MCM3AP | 0.375357 | 0.9543822 | 0.055943 | 0.7733738 | -1.202634 | 0.9799645 | 1.272016 | 0.02400653 | 1.710808 | 0.0171907 | -0.286307 | 0.8457101 |
| ENSG00000160305.17 | DIP2A | -0.534169 | 0.8006286 | -0.18766 | 0.8217443 | -0.978374 | 1 | 1.684085 | 0.000450429 | 0.93451 | 0.0155799 | 1.220336 | 0.8910687 |
| ENSG00000160307.9 | S100B | -0.092735 | 0.9569992 | -0.478723 | 0.06013553 | -0.230985 | 0.6406246 | -0.055367 | 0.0213259 | -0.260027 | 0.002297051 | -0.27128 | 0.8479431 |
| ENSG00000160352.15 | ZNF714 | -0.533131 | 0.6214382 | -0.186009 | 0.1997538 | 0.300825 | 0.3644618 | -1.16066 | 0.04053302 | 1.067032 | 0.06448943 | 1.986433 | 0.8479431 |
| ENSG00000160360.11 | GPSM1 | 1.445493 | 0.3567736 | -0.373078 | 0.6527736 | -0.510112 | 0.08298866 | 0.825844 | 0.002078872 | 1.465672 | 0.00053595 | 0.104837 | 0.9809768 |
| ENSG00000160439.15 | RDH13 | 1.178159 | 0.563505 | -0.555246 | 0.8124878 | 0.304691 | 0.8106078 | 3.165406 | 0.0212482 | 1.146409 | 0.808821 | -3.346946 | 0.5733051 |
| ENSG00000160445.10 | ZER1 | -0.332492 | 0.8587992 | -0.126284 | 1 | 1.872328 | 0.4382561 | 2.736282 | 0.3129763 | 2.798068 | 0.005112807 | 0.188478 | 0.5855893 |
| ENSG00000160469.16 | BRSK1 | -0.043752 | 0.8006286 | 0.773931 | 0.5468242 | -0.387254 | 0.1232614 | 0.893248 | 0.0109765 | 2.217696 | 0.000462566 | 2.193629 | 0.736021 |
| ENSG00000160551.9 | TAOK1 | -0.249631 | 0.6214382 | 0.080106 | 0.9906117 | 1.382271 | 2.10E-10 | 1.767802 | 2.40E-05 | 1.88292 | 0.000531075 | 0.854061 | 1 |
| ENSG00000160584.15 | SIK3 | -0.750857 | 1 | 0.138958 | 0.8263112 | 0.896437 | 0.005903382 | 1.131572 | 0.1648328 | 1.959848 | 0.002216678 | 1.154602 | 0.9562147 |
| ENSG00000160613.12 | PCSK7 | 1.564306 | 0.9574742 | 0.246736 | 0.9836728 | 0.260453 | 0.8454005 | 2.207747 | 0.000699288 | 0.929137 | 0.3917336 | 0.316148 | 0.6885065 |
| ENSG00000160633.12 | SAFB | -1.147574 | 0.6214382 | -0.479563 | 0.6184438 | 0.739955 | 0.002915961 | 0.012939 | 0.5117776 | 0.709634 | 0.5497035 | 0.566798 | 0.9435254 |
| ENSG00000160710.15 | ADAR | 0.396368 | 0.8006286 | -0.342102 | 0.4900459 | 0.512731 | 0.9817751 | 0.744412 | 0.001098587 | 0.951331 | 0.03043237 | -0.437445 | 0.8479431 |
| ENSG00000160714.9 | UBE2Q1 | -2.060685 | 0.1004496 | 0.182928 | 0.6940633 | -1.028758 | 0.1892891 | 1.414382 | 0.003235598 | 2.063196 | 0.001555174 | 0.786423 | 0.8479431 |
| ENSG00000160753.15 | RUSC1 | -2.000287 | 0.02822298 | -0.263901 | 0.6527736 | 0.691264 | 0.7999551 | -0.644438 | 0.2187288 | -0.216029 | 0.5430328 | 0.521704 | 0.6885065 |
| ENSG00000160785.13 | SLC25A44 | -0.836028 | 0.6214382 | 0.068178 | 0.7881421 | 0.67347 | 0.7190639 | 1.773752 | 0.02984841 | 1.106506 | 0.4127764 | -1.185707 | 0.6576742 |
| ENSG00000160789.19 | LMNA | -0.254245 | 0.6214382 | -0.067699 | 0.6573885 | -0.2837 | 0.03072077 | 0.516481 | 0.6865487 | 0.957335 | 0.6700181 | -0.04391 | 0.9510409 |
| ENSG00000160813.6 | PPP1R35 | -0.73586 | 0.8587992 | -0.108103 | 0.5515692 | -0.678215 | 4.66E-05 | 0.364566 | 0.4919368 | 1.666501 | 0.02125501 | 0.241067 | 0.936657 |
| ENSG00000160818.16 | GPATCH4 | -0.396544 | 0.6014343 | -0.052659 | 0.5818005 | 0.326145 | 0.912181 | 1.214499 | 1 | -0.534945 | 0.002514978 | -0.541297 | 0.426223 |
| ENSG00000160877.5 | NACC1 | 0.253919 | 0.6362998 | 0.335022 | 0.7383998 | -0.245923 | 0.2100026 | 1.242944 | 0.1088665 | 0.974352 | 0.01586635 | 0.21297 | 0.7787921 |
| ENSG00000160888.6 | IER2 | -1.10611 | 1 | 0.113203 | 0.7674398 | -0.741899 | 4.24E-18 | 0.055832 | 0.4919368 | 0.884524 | 0.09283086 | 1.264403 | 0.5066477 |
| ENSG00000160917.14 | CPSF4 | -0.993519 | 0.3026228 | -0.050079 | 0.7118133 | -0.265762 | 0.935949 | 1.871183 | 0.01556729 | 0.637111 | 0.0619892 | -1.984093 | 1 |
| ENSG00000160932.10 | LY6E | -0.294144 | 0.8006286 | 0.421966 | 0.6527736 | -0.954009 | 6.28E-05 | 0.446821 | 0.7325725 | -0.536683 | 0.3284643 | 0.487197 | 0.8369896 |
| ENSG00000160961.11 | ZNF333 | -1.609794 | 0.6014343 | 0.016081 | 0.9002426 | -0.690212 | 0.8656249 | 1.902176 | 0.02471293 | 0.168278 | 0.2922943 | 1.250547 | 0.6167027 |
| ENSG00000160991.15 | ORAI2 | -0.747446 | 0.6214382 | -0.529533 | 0.01857467 | -0.269931 | 0.9737329 | 0.576082 | 0.001371073 | 1.020268 | 0.2019818 | -4.80E-05 | 0.8361696 |
| ENSG00000160993.3 | ALKBH4 | 0.239071 | 0.8263393 | -0.377299 | 0.69914 | -0.363268 | 0.5257946 | 1.647367 | 0.4919368 | -0.861054 | 0.04945669 | 1.232052 | 0.6885065 |
| ENSG00000161011.19 | SQSTM1 | 0.135147 | 0.4764974 | 0.214075 | 0.5395105 | -0.249832 | 0.000556776 | -0.19188 | 0.004493672 | 0.816765 | 0.5669432 | 0.287808 | 1 |
| ENSG00000161013.16 | MGAT4B | 2.770993 | 0.6214382 | -0.107405 | 0.8953649 | -1.223805 | 0.1602815 | -0.192246 | 0.004779477 | 1.617799 | 0.2880317 | 0.155752 | 0.9391913 |
| ENSG00000161016.15 | RPL8 | -0.156466 | 0.6014343 | -0.04084 | 0.8100352 | 0.120398 | 0.01877144 | 0.20472 | 0.4919368 | -0.048533 | 0.06067043 | -0.016666 | 0.5733051 |
| ENSG00000161036.10 | LRWD1 | -0.364874 | 0.8867323 | 0.820921 | 0.6258673 | -0.47831 | 0.0153109 | 0.471507 | 0.7144938 | 0.329781 | 0.621723 | 0.921265 | 0.6829816 |
| ENSG00000161048.11 | NAPEPLD | -3.169701 | 0.563505 | -1.238247 | 0.3250176 | -0.810018 | 0.8616665 | 3.157268 | 0.01017765 | 1.516578 | 0.3651006 | 0.442774 | 0.7942023 |
| ENSG00000161057.10 | PSMC2 | 0.997459 | 0.03720286 | 0.262461 | 0.7714112 | 0.104654 | 0.897684 | 0.049354 | 0.07777871 | 0.534567 | 0.4089712 | -1.700858 | 0.2159896 |
| ENSG00000161082.12 | CELF5 | -0.272837 | 0.8006286 | -1.758738 | 0.4487482 | -0.06032 | 0.5290845 | 1.727759 | 0.000174739 | 1.066727 | 0.000172682 | 0.870587 | 0.6885065 |
| ENSG00000161091.12 | MFSD12 | 0.638382 | 0.6014343 | -0.281257 | 0.9878938 | 0.048959 | 0.5290845 | 0.688603 | 0.04148071 | 0.590783 | 0.1205422 | 1.419555 | 0.6885065 |
| ENSG00000161179.13 | YDJC | -0.315712 | 1 | 0.253647 | 0.7450389 | -2.517733 | 0.008038484 | 0.233775 | 0.2821571 | 1.37117 | 0.8482446 | 1.781981 | 0.6885065 |
| ENSG00000161202.17 | DVL3 | -1.285199 | 0.563505 | 0.304664 | 0.7049761 | -0.48176 | 0.9870138 | 0.906419 | 0.002014056 | 1.553247 | 0.03164928 | -1.721095 | 0.9936577 |
| ENSG00000161298.16 | ZNF382 | 4.427891 | 0.6214382 | 0.312879 | 0.9222921 | -0.492262 | 0.4587626 | 2.752618 | 9.18E-07 | -0.011398 | 0.08857983 | 0.912634 | 0.9483075 |
| ENSG00000161533.11 | ACOX1 | 0.523442 | 0.8867323 | -0.112313 | 0.5953732 | 0.289909 | 0.1620803 | 1.439479 | 0.01576557 | 0.559316 | 0.6759491 | -0.162723 | 0.7896829 |
| ENSG00000161547.14 | SRSF2 | -1.105317 | 0.04134193 | 0.396527 | 0.1573875 | 0.241903 | 0.8100781 | 0.409513 | 0.7144938 | 0.292544 | 0.5196843 | -0.021202 | 0.4304069 |
| ENSG00000161638.10 | ITGA5 | 2.132842 | 0.6214382 | -0.029516 | 1 | 0.565565 | 0.5409989 | 0.351342 | 0.04690412 | -2.535546 | 1 | 1.174986 | 0.6140767 |
| ENSG00000161654.9 | LSM12 | -0.318912 | 1 | 0.084 | 0.6222236 | 0.502574 | 0.4721352 | -0.024109 | 0.01324589 | 1.04466 | 0.7346969 | -0.43315 | 0.4049998 |
| ENSG00000161671.16 | EMC10 | 0.530786 | 0.4680273 | -0.177057 | 0.5012872 | 0.547136 | 0.168473 | 0.951003 | 0.01305884 | 1.265104 | 0.3450415 | -0.307164 | 0.7213346 |
| ENSG00000161677.11 | JOSD2 | -0.518244 | 1 | 0.139406 | 0.6527736 | -1.349227 | 0.000950538 | 0.34788 | 0.6996942 | -1.116599 | 0.1148494 | -1.430684 | 0.8620547 |
| ENSG00000161692.17 | DBF4B | 1.185791 | 1 | 2.888888 | 0.6527736 | 0.039293 | 0.6752066 | 3.645627 | 0.004296663 | 3.160258 | 0.6841782 | 6.000925 | 0.6517664 |
| ENSG00000161791.13 | FMNL3 | -2.048063 | 0.9057626 | 0.447317 | 0.7615373 | -1.194957 | 0.09304354 | 1.340362 | 0.002623356 | 2.648425 | 0.006445486 | -0.447304 | 0.6885065 |
| ENSG00000161813.20 | LARP4 | -0.272141 | 0.9282145 | 0.551987 | 0.2335602 | 0.561378 | 0.01574431 | 1.103023 | 0.157809 | 1.496065 | 0.3907759 | 0.987214 | 0.7557776 |
| ENSG00000161904.11 | LEMD2 | 0.343138 | 0.9183025 | 0.946113 | 0.6527736 | 0.480591 | 0.492686 | -0.86952 | 0.006286247 | 2.707031 | 0.03387033 | -0.588189 | 0.9368176 |
| ENSG00000161960.14 | EIF4A1 | 0.042538 | 0.8006286 | -0.711552 | 0.7383998 | 1.591424 | 0.9304971 | -0.791448 | 0.7858185 | 1.761009 | 0.01498928 | 0.939868 | 0.6885065 |
| ENSG00000161970.12 | RPL26 | -1.0152 | 0.6014343 | 0.117751 | 0.704767 | -3.70E-05 | 0.8832504 | 0.012795 | 0.1331699 | 0.004635 | 0.01677568 | -0.101768 | 0.3245041 |
| ENSG00000161980.5 | POLR3K | 0.179903 | 0.6214382 | 0.310166 | 0.7149452 | -0.322616 | 0.04815049 | -0.196634 | 0.2676368 | -1.232009 | 0.002485204 | -0.396983 | 0.6490419 |
| ENSG00000161981.10 | SNRNP25 | 0.385799 | 0.8006286 | -0.06814 | 1 | -0.411247 | 0.000664824 | 0.231568 | 0.1524893 | 0.021177 | 0.1576769 | -0.946735 | 0.6885065 |
| ENSG00000162065.11 | TBC1D24 | 0.606729 | 0.6214382 | -0.030505 | 1 | -0.570771 | 0.5290845 | 1.291032 | 0.002422767 | 0.65803 | 0.4499166 | -0.455385 | 0.960853 |
| ENSG00000162066.14 | AMDHD2 | -1.048799 | 0.6214382 | 0.418667 | 0.5953732 | -1.640904 | 0.01534135 | 0.775416 | 0.8472142 | 1.925957 | 0.4049597 | 0.609232 | 0.5193439 |
| ENSG00000162073.13 | PAQR4 | 0.267574 | 0.8006286 | 0.238803 | 0.7881421 | 0.026838 | 0.5782905 | 1.51864 | 0.04638111 | 0.950373 | 0.4879833 | 0.867896 | 1 |
| ENSG00000162086.14 | ZNF75A | -1.694961 | 0.529676 | -0.295173 | 0.8703522 | 1.148857 | 0.5290845 | 1.409866 | 0.004981271 | 1.163986 | 0.5196843 | 0.13287 | 0.7169928 |
| ENSG00000162174.12 | ASRGL1 | -0.201516 | 0.6214382 | -0.429461 | 0.51543 | 0.628872 | 0.03643028 | -0.292948 | 0.4919368 | 0.493154 | 1 | -0.344904 | 0.789721 |
| ENSG00000162191.13 | UBXN1 | 0.083555 | 0.9282145 | 0.332081 | 0.6527736 | 0.481036 | 0.5705018 | 0.731724 | 0.2048887 | -0.897891 | 0.005218747 | 0.429382 | 0.6885065 |
| ENSG00000162244.10 | RPL29 | -0.110003 | 0.896426 | 0.107889 | 1 | -0.041929 | 0.2707921 | -0.056406 | 0.03776658 | 0.028809 | 0.0658242 | 0.005331 | 0.3746505 |
| ENSG00000162298.16 | SYVN1 | 0.188243 | 0.9363575 | -0.094982 | 0.7133822 | 0.657862 | 0.6209689 | 3.084836 | 0.01368152 | 1.00079 | 1 | 1.690854 | 0.7314808 |
| ENSG00000162368.13 | CMPK1 | -0.810702 | 0.8814146 | 0.024993 | 0.5398987 | 0.574271 | 0.01060758 | 0.12463 | 1 | -0.195025 | 0.1198138 | 1.829082 | 0.6829816 |
| ENSG00000162374.16 | ELAVL4 | 0.585731 | 0.9015252 | -0.639886 | 1 | 0.619817 | 0.002400545 | 1.111344 | 0.04612389 | -0.146107 | 0.2544437 | -0.182735 | 0.4684661 |
| ENSG00000162378.12 | ZYG11B | 2.037421 | 0.4511893 | -0.41807 | 0.3530125 | 0.021189 | 0.6057253 | 1.113465 | 5.17E-05 | 1.192415 | 0.03105236 | -0.908786 | 0.4258987 |
| ENSG00000162385.10 | MAGOH | -1.188811 | 0.6616343 | 0.213628 | 0.8657239 | -0.08535 | 0.000685504 | -0.566396 | 0.000155077 | -0.167958 | 0.02249455 | -0.153938 | 0.6885065 |
| ENSG00000162408.10 | NOL9 | 0.448351 | 0.9444548 | 0.013872 | 0.6527736 | 0.671792 | 0.6295125 | 1.579108 | 0.003473137 | 1.62621 | 0.02344635 | -2.408698 | 0.6517664 |
| ENSG00000162419.12 | GMEB1 | 0.634293 | 0.7090887 | -0.46966 | 0.5953732 | -0.747913 | 1 | 1.5248 | 0.000494198 | 1.59003 | 0.1158034 | 1.03098 | 0.6829816 |
| ENSG00000162426.14 | SLC45A1 | 0.292152 | 1 | 0.225238 | 0.704767 | -0.161798 | 0.6275552 | 2.216477 | 0.2888859 | -0.257288 | 0.6171067 |  |  |
| ENSG00000162430.16 | SEPN1 | 1.858794 | 0.6214382 | -0.356518 | 0.8263112 | -0.887174 | 0.6928639 | 1.011872 | 0.01797834 | 1.241581 | 0.1247593 | -0.219355 | 0.5733051 |
| ENSG00000162441.11 | LZIC | 1.786282 | 0.8006286 | 0.106936 | 0.9056349 | -0.14576 | 0.1138222 | -0.310992 | 0.9720008 | -1.316276 | 0.0124237 | 1.598279 | 0.9215032 |
| ENSG00000162490.6 | DRAXIN | 1.935129 | 0.6214382 | -0.324605 | 0.8953649 | 0.672646 | 1 | 0.941341 | 4.34E-05 | 0.937124 | 0.2063571 | 0.081449 | 0.8255112 |
| ENSG00000162512.15 | SDC3 | 0.10232 | 1 | -0.43786 | 0.6527736 | 0.065517 | 0.5322465 | 0.976754 | 0.000675459 | 0.539715 | 0.1170868 | 0.826981 | 0.6885065 |
| ENSG00000162520.14 | SYNC | -0.7253 | 1 | 1.128268 | 0.9674023 | 0.490425 | 0.8868899 | 2.040647 | 0.000766333 | 1.222779 | 0.95261 | 0.409441 | 0.773978 |
| ENSG00000162521.18 | RBBP4 | -0.316642 | 0.529676 | -0.380017 | 0.6746338 | 0.717545 | 0.000418292 | 1.04825 | 0.000105024 | 0.987541 | 0.09188663 | -0.974069 | 0.570375 |
| ENSG00000162545.5 | CAMK2N1 | -0.101334 | 0.9986678 | 0.11853 | 0.6527736 | 0.15732 | 0.8583913 | 1.116674 | 0.000353717 | 0.615717 | 0.1191954 | -0.083429 | 0.6885065 |
| ENSG00000162585.16 | FAAP20 | -0.120668 | 0.9506557 | 0.542623 | 0.7813551 | -2.5954 | 0.000731729 | 1.023843 | 0.4446428 | 1.286748 | 0.5196843 | 1.085438 | 0.6885065 |
| ENSG00000162599.15 | NFIA | -1.81872 | 0.6214382 | -0.21429 | 0.005018379 | 0.960676 | 6.93E-06 | 1.247251 | 0.005905424 | 0.498845 | 0.07401196 | 2.447478 | 0.5553892 |
| ENSG00000162601.9 | MYSM1 | -1.610645 | 0.8587992 | 0.121952 | 0.6527736 | 0.174877 | 0.583618 | 1.546731 | 0.04303665 | 1.000895 | 0.5196843 | 0.100893 | 0.6167027 |
| ENSG00000162607.12 | USP1 | -2.160584 | 0.2998756 | 0.127571 | 0.8379413 | 1.101335 | 0.002952376 | 0.835818 | 0.0229571 | 0.227113 | 0.3630456 | -1.008677 | 0.3565625 |
| ENSG00000162613.16 | FUBP1 | -0.513939 | 0.7996471 | -0.095726 | 0.9002351 | 0.671725 | 0.00546411 | 1.536659 | 7.52E-07 | 0.686938 | 0.5757457 | 0.192541 | 1 |
| ENSG00000162623.15 | TYW3 | 0.633093 | 1 | 0.754659 | 0.9441754 | 0.275168 | 0.6646119 | 0.183955 | 0.5584647 | 4.222843 | 0.04179319 | -0.307064 | 0.9147227 |
| ENSG00000162645.12 | GBP2 | -0.902361 | 0.8587992 | 0.116051 | 0.6527736 | -1.015201 | 0.000160681 | 0.116917 | 0.02464196 | 0.23054 | 0.3027771 | 0.993006 | 0.8167193 |
| ENSG00000162694.13 | EXTL2 | 2.217472 | 0.6214382 | -0.046317 | 0.8703522 | 1.62523 | 0.004463462 | 1.469585 | 0.01000591 | -0.738069 | 0.130167 | 0.204107 | 0.6885065 |
| ENSG00000162704.15 | ARPC5 | 0.173935 | 0.8222779 | 0.449402 | 0.4072962 | 0.382888 | 0.05005774 | -0.074705 | 0.008110694 | -0.018159 | 0.000159217 | 0.193594 | 0.6647029 |
| ENSG00000162706.12 | CADM3 | -3.029291 | 0.4764974 | 0.580202 | 1 | 1.117774 | 0.2814014 | 0.805474 | 0.04788096 | 0.425836 | 0.5659006 | 0.65707 | 0.9134607 |
| ENSG00000162734.12 | PEA15 | 0.323765 | 0.00438331 | -0.357435 | 0.137375 | 0.292663 | 0.4341104 | 0.147297 | 0.4690701 | 0.61709 | 0.506562 | 0.333774 | 1 |
| ENSG00000162735.18 | PEX19 | 0.94578 | 0.6792949 | 0.229392 | 0.8845769 | 1.218936 | 0.1787542 | 2.076047 | 0.03605383 | 0.252308 | 0.7377262 | -3.921308 | 0.6885065 |
| ENSG00000162738.5 | VANGL2 | 0.131186 | 0.8006286 | -0.499402 | 0.8703522 | 0.599522 | 0.3336662 | 2.093611 | 0.0241994 | 2.0291 | 0.002269503 | 2.34397 | 0.6829816 |
| ENSG00000162772.16 | ATF3 | -2.044516 | 0.714717 | 0.118325 | 0.6284076 | 0.048017 | 0.5290845 | -0.700841 | 0.02497053 | 0.062618 | 0.5196843 | 0.779103 | 0.6185816 |
| ENSG00000162775.14 | RBM15 | -1.638794 | 0.6214382 | -0.142941 | 0.8017884 | 0.356299 | 0.4064599 | 2.383941 | 0.0495995 | 1.975428 | 0.2523958 | -1.640602 | 0.8260537 |
| ENSG00000162783.10 | IER5 | -1.106866 | 0.529676 | 0.032566 | 0.8263112 | 0.122302 | 0.8089907 | -0.81468 | 0.8224647 | 1.150015 | 0.03734702 | 0.795569 | 0.6885065 |
| ENSG00000162849.15 | KIF26B | -2.379531 | 0.6214382 | 1.00113 | 0.5953732 | 0.923426 | 0.4136152 | 0.905074 | 0.043823 | -1.631694 | 0.5196843 | 1.599056 | 0.8949489 |
| ENSG00000162889.10 | MAPKAPK2 | -4.37344 | 0.8122564 | -0.037197 | 1 | -0.458745 | 0.5290845 | 0.853552 | 0.02160238 | 0.159022 | 0.8954063 | 1.79574 | 0.7767294 |
| ENSG00000162910.18 | MRPL55 | 0.33061 | 0.4923792 | 0.003377 | 0.9117805 | -0.605422 | 6.13E-10 | 0.170064 | 0.3059632 | 0.124606 | 0.01165836 | 0.631773 | 0.6885065 |
| ENSG00000162923.14 | WDR26 | -0.228733 | 0.8374767 | 0.190002 | 0.906244 | 1.185374 | 0.001345929 | 1.068856 | 0.004466975 | 1.078332 | 0.01640239 | -2.388785 | 0.8235317 |
| ENSG00000162961.13 | DPY30 | 0.068364 | 0.9591207 | -0.131027 | 0.9222921 | 0.264014 | 0.6930033 | -0.299872 | 0.01062276 | 0.128624 | 0.02427023 | -0.277381 | 0.7767294 |
| ENSG00000162975.4 | KCNF1 | -0.466829 | 0.781685 | -0.222446 | 0.5953732 | -0.78756 | 1 | 1.953558 | 4.94E-07 | -0.029354 | 0.5196843 | 2.725321 | 0.6829816 |
| ENSG00000162992.3 | NEUROD1 | -0.94782 | 0.779522 | -1.087812 | 0.06905985 | -1.927448 | 0.3389032 | 1.651479 | 0.007199007 | -0.12902 | 0.8769722 |  |  |
| ENSG00000163001.11 | CFAP36 | -0.784271 | 0.714717 | 0.443915 | 0.5103145 | 0.273938 | 0.2049447 | -0.686772 | 0.00062518 | -0.823449 | 0.009058141 | -0.475477 | 0.7191061 |
| ENSG00000163029.15 | SMC6 | -0.430907 | 0.6817555 | 0.461658 | 0.6527736 | 1.210683 | 0.1991342 | -0.387256 | 0.03932487 | 0.917198 | 0.2130361 | -1.746931 | 0.8699042 |
| ENSG00000163041.9 | H3F3A | -0.878895 | 0.04270195 | -0.253086 | 0.06746408 | 0.204867 | 0.006792414 | 0.165955 | 0.4719248 | -0.076441 | 0.2910737 | -0.394203 | 0.03362812 |
| ENSG00000163069.12 | SGCB | 0.015757 | 0.8240785 | -0.925751 | 0.005330032 | 0.777791 | 0.2716525 | 0.971016 | 0.7657271 | 0.184692 | 0.9062893 | 1.115612 | 0.8479431 |
| ENSG00000163104.17 | SMARCAD1 | -0.49041 | 0.6014343 | 0.562215 | 0.9622004 | 0.96491 | 0.2879524 | 0.60253 | 0.5485418 | -0.974691 | 0.01396457 | -0.422531 | 0.6829816 |
| ENSG00000163125.15 | RPRD2 | 1.149188 | 0.529676 | -0.032327 | 0.9008453 | 1.255484 | 0.02035361 | -0.486862 | 0.854549 | 1.238348 | 0.2206396 | -1.0662 | 0.8949489 |
| ENSG00000163155.11 | LYSMD1 | 0.238645 | 0.7736255 | 0.632356 | 0.5953732 | -1.888275 | 0.8502175 | 1.04064 | 0.4919368 | 5.090599 | 0.02707314 | 0.447315 | 1 |
| ENSG00000163156.11 | SCNM1 | -0.174527 | 0.7520517 | 0.159055 | 0.6890183 | -0.708842 | 0.000288113 | 0.681072 | 0.7376256 | -0.440378 | 0.2606029 | 0.156032 | 0.6885065 |
| ENSG00000163162.8 | RNF149 | 0.452105 | 0.8006286 | 0.360269 | 0.3531232 | -0.670618 | 0.4064599 | 2.936608 | 0.3760083 | 2.124325 | 0.006961238 | 0.890164 | 0.5344421 |
| ENSG00000163170.11 | BOLA3 | 0.062378 | 0.6014343 | -0.068224 | 0.9550491 | -0.764534 | 1.60E-09 | 0.375476 | 0.3946459 | -0.332453 | 0.1293467 | -0.006221 | 0.8273997 |
| ENSG00000163191.5 | S100A11 | 0.661108 | 0.8006286 | 0.167507 | 0.7206959 | -0.976301 | 5.69E-05 | 0.09443 | 0.3377101 | 0.700484 | 0.3820915 | 0.106749 | 0.6885065 |
| ENSG00000163214.20 | DHX57 | -0.565613 | 0.8006286 | 0.405857 | 0.3269793 | 0.938019 | 0.1951855 | 2.351549 | 0.00028898 | 3.33577 | 0.005218747 | -1.37045 | 0.6108304 |
| ENSG00000163291.14 | PAQR3 | 1.51627 | 0.6214382 | -0.273015 | 1 | -0.606963 | 0.2273924 | 2.199303 | 0.04977764 | 1.532708 | 0.6573569 | -0.650189 | 0.7767294 |
| ENSG00000163319.10 | MRPS18C | -0.327116 | 0.8530607 | -0.064574 | 0.9793419 | -0.330339 | 0.001726757 | -0.423238 | 0.01868866 | -0.730996 | 0.01777855 | -0.122612 | 0.6885065 |
| ENSG00000163328.13 | GPR155 | 1.345982 | 0.6214382 | 0.027034 | 0.7806263 | 0.641409 | 0.7493088 | 1.351257 | 0.04382913 | 1.353154 | 0.811017 | 1.994122 | 0.6885065 |
| ENSG00000163344.5 | PMVK | 0.691579 | 1 | -0.153258 | 0.87788 | -0.6299 | 0.000803259 | 0.094753 | 0.616161 | 0.92444 | 0.7339134 | 0.378853 | 0.9391913 |
| ENSG00000163348.3 | PYGO2 | 0.161566 | 0.8006286 | -0.918389 | 0.3442044 | -0.740347 | 0.5424025 | 1.99789 | 6.59E-06 | 1.495462 | 0.2089173 | -2.414081 | 0.9532204 |
| ENSG00000163349.21 | HIPK1 | -0.660709 | 0.6214382 | -0.046452 | 1 | 0.815752 | 0.05529066 | 1.207562 | 0.006354984 | 0.813173 | 0.06141896 | -0.501071 | 0.6885065 |
| ENSG00000163374.19 | YY1AP1 | -1.854213 | 0.563505 | -0.044936 | 0.8148374 | 0.056963 | 0.8800236 | 0.409706 | 0.1722277 | 1.629381 | 0.03043237 | -0.233249 | 0.9770956 |
| ENSG00000163428.3 | LRRC58 | -0.243913 | 0.9281698 | 0.026646 | 0.8513833 | 1.187037 | 0.02831164 | 0.362245 | 0.06765408 | 0.884465 | 0.5196843 | -0.226211 | 0.9147227 |
| ENSG00000163453.11 | IGFBP7 | -3.859955 | 0.000100952 | 0.039034 | 0.7662689 | -3.068637 | 0.02505569 | -0.892235 | 0.007816005 | 1.158693 | 0.9013748 | 0.384168 | 0.6829816 |
| ENSG00000163463.11 | KRTCAP2 | -0.27912 | 0.7865587 | -0.919424 | 0.6527736 | -0.640323 | 0.01138584 | 0.569418 | 0.7383668 | 0.745396 | 1 | -2.920135 | 0.6885065 |
| ENSG00000163466.15 | ARPC2 | -0.402386 | 0.3585928 | 0.309481 | 0.3903174 | 0.021324 | 1 | -0.196014 | 0.001255635 | -0.002645 | 0.001005217 | 0.13354 | 0.4149735 |
| ENSG00000163482.11 | STK36 | -0.802325 | 0.6214382 | -0.062031 | 1 | -0.4379 | 0.3403162 | 1.972801 | 0.02179676 | 2.842824 | 0.7256106 | 1.258452 | 0.6490419 |
| ENSG00000163510.13 | CWC22 | -1.302766 | 0.9882924 | 0.301321 | 0.8872169 | 0.644895 | 0.0349371 | -0.183907 | 0.9294852 | 1.250502 | 0.5284694 | -0.939673 | 0.6829816 |
| ENSG00000163512.13 | AZI2 | -0.507939 | 0.9473277 | -0.314659 | 0.7740003 | 1.090496 | 0.01090776 | 0.71643 | 0.6122347 | -0.460442 | 0.09569036 | 0.405495 | 0.9147227 |
| ENSG00000163527.9 | STT3B | 1.330016 | 0.5104673 | -0.042296 | 0.2482908 | 1.12193 | 0.5411385 | 1.325328 | 1.59E-06 | 1.29055 | 0.133248 | -0.6196 | 1 |
| ENSG00000163531.15 | NFASC | -0.390108 | 0.6214382 | 0.439063 | 0.2217175 | -0.650728 | 0.5290845 | 0.820792 | 0.001660058 | 2.34831 | 0.02269293 | -0.449315 | 0.8460322 |
| ENSG00000163539.16 | CLASP2 | -0.791332 | 0.961441 | -0.949081 | 0.6369839 | 1.118426 | 0.02006313 | 1.226648 | 9.33E-06 | 1.431857 | 0.000197614 | -2.066868 | 0.8739261 |
| ENSG00000163541.11 | SUCLG1 | -0.251748 | 0.6956464 | 0.116627 | 0.7765491 | -0.049918 | 0.5772181 | -0.090637 | 0.005179108 | -0.420409 | 6.14E-06 | 0.117364 | 0.7385207 |
| ENSG00000163558.12 | PRKCI | 0.658219 | 0.9437365 | 0.197361 | 0.957632 | 1.603372 | 0.02862298 | 0.256315 | 0.3825322 | -0.772227 | 0.4099199 | 0.627315 | 0.8841423 |
| ENSG00000163565.18 | IFI16 | 0.167226 | 0.7706531 | -0.154719 | 0.3250176 | 0.307939 | 0.3228618 | 0.026149 | 0.001646819 | 0.267821 | 0.280488 | 0.64173 | 0.8316849 |
| ENSG00000163584.17 | RPL22L1 | -0.172033 | 0.6214382 | -0.042317 | 0.8082353 | -0.608126 | 0.001804989 | -0.477637 | 0.06040875 | -0.638839 | 0.2880317 | 0.297219 | 0.1652728 |
| ENSG00000163590.13 | PPM1L | 1.447339 | 0.714717 | 1.354965 | 0.9104976 | 0.998505 | 0.2735701 | 2.074995 | 0.01576557 | 1.295383 | 0.3588972 | 2.796487 | 0.6829816 |
| ENSG00000163602.9 | RYBP | -1.980061 | 0.529676 | 0.053924 | 0.8653838 | 0.207618 | 0.2026939 | 0.635551 | 0.1886176 | 2.069228 | 0.009884707 | -0.639842 | 0.9467563 |
| ENSG00000163605.14 | PPP4R2 | 0.025834 | 0.714717 | 0.261212 | 0.6527736 | 0.795189 | 0.0227862 | 0.829973 | 0.04738219 | 1.013991 | 0.6454194 | -0.918921 | 0.8479431 |
| ENSG00000163618.17 | CADPS | 0.544794 | 1 | -1.748531 | 0.5010139 | 0.113583 | 0.8656249 | 1.646417 | 2.84E-05 | 1.666127 | 0.621723 | -1.187028 | 0.834639 |
| ENSG00000163625.15 | WDFY3 | -0.272299 | 0.9282145 | -0.234641 | 0.8832909 | 2.335833 | 0.05147203 | 3.337518 | 0.000688376 | 1.194478 | 0.2433502 | 0.153632 | 0.8818722 |
| ENSG00000163629.12 | PTPN13 | 1.061823 | 0.9500536 | -0.413039 | 0.04079597 | 1.352554 | 0.002570252 | -0.501443 | 0.8216874 | 0.550754 | 0.3467589 | 1.249127 | 1 |
| ENSG00000163634.11 | THOC7 | -0.654683 | 0.6214382 | 0.373196 | 0.8703522 | -0.490101 | 0.3683009 | 0.320117 | 0.01258996 | 0.225731 | 0.5101933 | 0.968111 | 0.897483 |
| ENSG00000163637.11 | PRICKLE2 | -1.292132 | 0.7736255 | 0.185703 | 0.6489085 | 0.942269 | 0.776456 | 2.624513 | 0.01446329 | 3.134584 | 0.6638875 | -1.321619 | 0.7333942 |
| ENSG00000163638.13 | ADAMTS9 | 0.344182 | 0.6014343 | -0.572985 | 0.6107341 | 0.995256 | 0.00482539 | 0.707844 | 0.1052874 | -1.154415 | 0.3815761 | -1.488465 | 0.5653062 |
| ENSG00000163644.14 | PPM1K | 0.035127 | 0.9991686 | -0.251508 | 0.6615629 | 0.997667 | 1.80E-05 | 1.257885 | 0.1524893 | 0.378777 | 0.5196843 | -2.80352 | 0.6463174 |
| ENSG00000163655.15 | GMPS | 0.400982 | 1 | -0.000701 | 0.971348 | 1.360785 | 0.006171149 | 2.348231 | 0.6181175 | 0.773847 | 0.7767786 | 0.532461 | 1 |
| ENSG00000163660.11 | CCNL1 | -0.930117 | 0.1187072 | 0.092789 | 0.9794341 | 0.4315 | 0.00208144 | 0.96989 | 0.5060864 | 0.709698 | 0.5196843 | 0.577381 | 0.4081608 |
| ENSG00000163682.15 | RPL9 | -2.147694 | 2.84E-26 | -0.001255 | 0.5953732 | 1.240386 | 2.25E-52 | -0.266424 | 1.55E-06 | -0.306649 | 4.37E-07 | -0.17731 | 0.005067357 |
| ENSG00000163684.11 | RPP14 | -5.124413 | 0.8867323 | 0.216541 | 0.979665 | 0.48624 | 0.4165649 | 2.020706 | 0.03166341 | -0.171269 | 0.8733325 | -2.776197 | 0.8167193 |
| ENSG00000163697.16 | APBB2 | -1.394659 | 0.4764974 | -0.968231 | 0.1276613 | 0.506052 | 0.04717406 | 1.940536 | 0.2683327 | 0.317848 | 0.8487036 | -2.568153 | 0.9147227 |
| ENSG00000163701.18 | IL17RE | 0.5629 | 0.8006286 | 1.357938 | 0.6527736 | -1.458854 | 0.3897481 | 2.580785 | 0.04369391 | 3.16082 | 0.1691344 | -3.916602 | 0.8479431 |
| ENSG00000163714.17 | U2SURP | -0.470385 | 0.6214382 | 0.120367 | 0.6527736 | 0.101363 | 9.57E-07 | 0.569528 | 0.08786815 | 0.80847 | 0.812769 | -0.391847 | 0.7759607 |
| ENSG00000163719.18 | MTMR14 | -1.371458 | 0.6214382 | 0.353787 | 0.6527736 | -0.859832 | 0.01553581 | 1.372472 | 9.93E-06 | 0.888979 | 0.3369949 | 1.466999 | 0.6576742 |
| ENSG00000163734.4 | CXCL3 | -1.637923 | 0.6168416 | 1.304815 | 0.8217443 | -4.695268 | 0.000951691 | 1.813935 | 0.9218588 | -0.229726 | 0.7377262 | 0.298956 | 0.9278206 |
| ENSG00000163754.17 | GYG1 | -0.007856 | 0.7548823 | 0.335612 | 0.5329817 | 0.44444 | 0.8888327 | -0.562806 | 0.03640873 | -0.614194 | 0.621723 | 2.095237 | 0.6837711 |
| ENSG00000163781.12 | TOPBP1 | -0.897678 | 0.4620794 | 0.485495 | 0.7789922 | 0.887762 | 0.2159645 | 2.011712 | 0.003867006 | 0.419539 | 0.9961553 | -1.725581 | 0.525454 |
| ENSG00000163785.12 | RYK | -1.56852 | 0.5464743 | -0.116189 | 0.8953649 | 0.55618 | 0.4064599 | 0.685253 | 0.04071611 | 1.707848 | 0.03387033 | 0.097347 | 0.7698133 |
| ENSG00000163795.13 | ZNF513 | -0.352498 | 0.7854407 | 0.406607 | 0.6895019 | -1.676969 | 0.01732351 | 0.528053 | 0.1311915 | 2.019339 | 0.09385903 | -1.448549 | 0.9614796 |
| ENSG00000163848.18 | ZNF148 | 0.631175 | 0.6014343 | 0.2648 | 0.7881421 | 1.339403 | 1.16E-05 | 1.799095 | 0.01980018 | 0.411824 | 0.3114707 | -1.477793 | 0.8479431 |
| ENSG00000163870.14 | TPRA1 | -0.528425 | 0.6214382 | 0.633273 | 0.8740162 | -0.931552 | 0.01216325 | 1.472439 | 1 | 1.20398 | 0.6179592 | 0.803645 | 0.6829816 |
| ENSG00000163872.15 | YEATS2 | -0.659473 | 0.7736255 | -0.39254 | 0.6077765 | -0.630568 | 0.4382561 | 0.557862 | 0.1557388 | 1.872137 | 0.004705683 | -1.735696 | 0.7197883 |
| ENSG00000163900.10 | TMEM41A | -1.57876 | 0.1835496 | -1.228057 | 0.2873629 | -1.748906 | 0.0745984 | 1.783838 | 0.04462914 | 2.275022 | 0.3371898 | 0.267621 | 0.6829816 |
| ENSG00000163902.11 | RPN1 | 0.930329 | 0.1784527 | 0.228515 | 0.8263112 | 0.393593 | 0.01897636 | 0.005294 | 0.479858 | -0.166757 | 0.1993336 | 0.992314 | 0.9428375 |
| ENSG00000163930.9 | BAP1 | -1.939343 | 0.6214382 | 0.171209 | 0.8040003 | -0.374239 | 0.7437308 | 0.35078 | 0.01064038 | 0.059126 | 0.393677 | 0.785925 | 0.868568 |
| ENSG00000163938.16 | GNL3 | 0.016866 | 0.6014343 | 0.15012 | 0.8208804 | 0.37507 | 0.0332374 | -0.318921 | 0.1729313 | 0.284117 | 0.8021966 | -0.880852 | 0.6556863 |
| ENSG00000163939.18 | PBRM1 | -0.686486 | 0.7727139 | -0.287108 | 0.4901086 | 0.732686 | 0.6411711 | 0.201039 | 0.02574097 | 0.435919 | 0.5013099 | -0.765194 | 0.6885065 |
| ENSG00000163947.11 | ARHGEF3 | 2.653142 | 0.6014343 | 0.156576 | 0.8113877 | -0.981483 | 0.4064599 | 1.24717 | 0.00972096 | 1.759966 | 0.2739281 | 0.614702 | 0.7473996 |
| ENSG00000163950.12 | SLBP | 0.322441 | 0.9282145 | 0.178379 | 0.8931774 | -0.342202 | 0.01722079 | 1.854197 | 0.05885179 | 0.638774 | 0.3974318 | 0.042804 | 0.6885065 |
| ENSG00000163956.10 | LRPAP1 | 0.499221 | 0.6214382 | 0.427064 | 0.5953732 | 0.183163 | 0.5103391 | -0.125276 | 0.03666334 | 0.281437 | 0.367138 | 0.541636 | 0.7595705 |
| ENSG00000163960.11 | UBXN7 | -5.326788 | 0.1745642 | 0.325521 | 0.6527736 | 0.861702 | 0.000106566 | 1.165201 | 0.01058931 | 0.744401 | 0.004418782 | -0.72627 | 0.9589332 |
| ENSG00000163961.4 | RNF168 | 1.432129 | 0.1065073 | -0.23365 | 0.7283542 | 0.592935 | 0.001894492 | 0.666012 | 0.04638111 | 0.164818 | 0.5196843 | -2.924721 | 0.5733051 |
| ENSG00000164008.13 | C1orf50 | 1.356315 | 0.529676 | 0.064154 | 1 | -0.889759 | 0.04023886 | -0.745794 | 0.2199825 | 0.609908 | 0.97225 | -2.26491 | 0.9460594 |
| ENSG00000164031.16 | DNAJB14 | 0.00433 | 0.6214382 | -0.194635 | 0.4886958 | 0.988692 | 0.08102887 | 0.212177 | 0.1372769 | 1.307192 | 0.03937989 | 0.057908 | 0.6885065 |
| ENSG00000164032.11 | H2AFZ | -0.211657 | 0.9545168 | 0.350602 | 0.1275359 | 0.144753 | 0.9704422 | -0.227749 | 0.02029956 | 0.14805 | 0.3201039 | -0.295645 | 0.1729051 |
| ENSG00000164040.16 | PGRMC2 | 0.061879 | 0.8006286 | 0.003643 | 0.9779936 | -0.374119 | 0.8652199 | 0.42306 | 0.3961377 | 1.078362 | 0.03306085 | 0.360559 | 0.969706 |
| ENSG00000164050.12 | PLXNB1 | -1.704509 | 0.6214382 | 0.871125 | 0.7593936 | 0.277569 | 0.7887553 | 1.259335 | 0.003880245 | 1.866737 | 0.000408876 | -0.589929 | 0.6767422 |
| ENSG00000164056.10 | SPRY1 | -0.319231 | 0.8006286 | -0.327686 | 0.6527736 | 1.097907 | 0.05391221 | 0.459315 | 0.4078268 | 0.010284 | 0.02647761 | 2.922187 | 0.6885065 |
| ENSG00000164062.12 | APEH | 1.442754 | 0.6980706 | -0.377929 | 0.4487482 | 0.271506 | 0.4538695 | 1.824617 | 4.47E-05 | 2.197756 | 0.02150138 | -1.138425 | 0.2959482 |
| ENSG00000164066.12 | INTU | -1.297801 | 0.714717 | -2.39059 | 0.6048501 | 1.217331 | 9.23E-05 | -0.035358 | 0.6296888 | 0.597445 | 0.5963743 | -0.800601 | 1 |
| ENSG00000164077.13 | MON1A | 0.362633 | 0.9367821 | -0.30264 | 0.9222921 | -0.375647 | 0.4064599 | 0.757005 | 0.4919368 | 2.613495 | 0.007549975 | -0.686883 | 0.8949489 |
| ENSG00000164080.13 | RAD54L2 | -0.802988 | 0.529676 | -0.249137 | 0.6527736 | 0.583621 | 0.03615249 | 1.020761 | 6.66E-05 | 0.452346 | 0.898446 | 1.206766 | 0.9029338 |
| ENSG00000164081.12 | TEX264 | -0.600142 | 0.7736255 | 0.158238 | 0.860006 | 0.394677 | 0.1394359 | -1.367586 | 0.04116586 | -0.217884 | 0.5196843 | 1.244964 | 0.9955797 |
| ENSG00000164091.11 | WDR82 | -1.550489 | 0.2166167 | -0.192639 | 0.3468329 | 0.448202 | 0.1444425 | 0.684442 | 0.002860427 | 0.500604 | 0.6105008 | -1.395745 | 0.1966294 |
| ENSG00000164096.12 | C4orf3 | -0.331693 | 0.6424217 | 0.34684 | 0.2416476 | -0.064495 | 0.1896442 | 0.057799 | 0.01980018 | 1.486646 | 0.7594424 | -0.363317 | 0.8949489 |
| ENSG00000164105.3 | SAP30 | -0.830754 | 0.6171259 | -0.046171 | 0.8661807 | -1.433899 | 0.8656249 | 0.477586 | 0.02921025 | 2.340786 | 0.005058518 | 0.758162 | 0.9895571 |
| ENSG00000164106.7 | SCRG1 | -0.073401 | 0.9195607 | 0.049869 | 0.8472204 | -0.030682 | 0.640899 | -0.253881 | 4.76E-06 | -0.146962 | 0.1084796 | -0.243109 | 0.5256722 |
| ENSG00000164111.14 | ANXA5 | -0.609302 | 0.6014343 | -0.113847 | 0.3417686 | 0.270717 | 0.02727104 | -0.137225 | 0.000911272 | -0.006316 | 0.000174371 | -0.041047 | 0.6885065 |
| ENSG00000164114.18 | MAP9 | -0.719486 | 0.6214382 | -0.532498 | 0.5953732 | 1.072952 | 1.27E-06 | 0.263807 | 1 | -0.179024 | 0.1912543 | 1.396755 | 0.7698133 |
| ENSG00000164118.12 | CEP44 | -0.312999 | 0.6214382 | 0.471353 | 0.6527736 | 1.959126 | 0.01817763 | 0.267316 | 0.9491102 | -0.942162 | 0.5196843 | -1.083812 | 0.6576742 |
| ENSG00000164134.12 | NAA15 | 0.481456 | 0.529676 | -0.077523 | 0.6200225 | 0.937246 | 0.000361376 | 1.220139 | 1.82E-06 | 0.303464 | 0.8159852 | -2.086037 | 0.3237894 |
| ENSG00000164151.11 | ICE1 | -1.505121 | 0.9264299 | 0.222459 | 0.8703522 | 0.574092 | 0.04435432 | 1.181412 | 0.06727595 | 0.581636 | 0.2130361 | 3.336085 | 0.3746505 |
| ENSG00000164163.10 | ABCE1 | -0.046891 | 0.7072782 | 0.144747 | 0.8688631 | 0.565009 | 0.01410444 | 1.061803 | 0.7051646 | 1.142417 | 0.1845956 | 0.161678 | 0.8207291 |
| ENSG00000164167.9 | LSM6 | -1.518137 | 0.6214382 | -0.12758 | 0.4519375 | 0.030539 | 1 | -0.835938 | 0.008473218 | -0.524537 | 0.1559212 | 0.878058 | 0.841154 |
| ENSG00000164176.12 | EDIL3 | 0.600762 | 0.8587992 | 1.030892 | 0.8263112 | 0.600602 | 1 | 1.288562 | 0.04969589 | 0.339644 | 0.5196843 | -2.039369 | 0.8535547 |
| ENSG00000164187.6 | LMBRD2 | -0.210096 | 0.6214382 | -0.922136 | 0.4148108 | 1.212551 | 0.04511416 | 1.17649 | 0.320553 | 1.918477 | 0.3254721 | -2.003094 | 0.6885065 |
| ENSG00000164190.16 | NIPBL | 0.648056 | 0.6014343 | -0.151216 | 0.3832761 | 1.125922 | 1.29E-05 | 1.626089 | 0.001745564 | 1.355264 | 0.007103428 | 1.237298 | 1 |
| ENSG00000164219.9 | PGGT1B | 1.028219 | 0.9828209 | -0.051192 | 0.5814954 | 1.008517 | 0.01383124 | 0.957746 | 0.2727697 | 0.352517 | 0.7447555 | -0.078159 | 0.9620927 |
| ENSG00000164241.13 | C5orf63 | -0.508664 | 0.6214382 | -0.739861 | 0.7881421 | -1.030859 | 0.1281741 | 1.386654 | 0.02741305 | -2.335836 | 0.7377262 | -2.059072 | 0.9570429 |
| ENSG00000164258.11 | NDUFS4 | 0.225073 | 0.6214382 | 0.129836 | 0.8379413 | 0.068474 | 0.5290845 | -0.283031 | 0.005840505 | -0.285877 | 0.09907856 | 0.457485 | 0.6885065 |
| ENSG00000164284.14 | GRPEL2 | -0.489418 | 0.9529915 | 0.289073 | 0.9836728 | 0.158167 | 0.6272378 | 1.934695 | 0.03914854 | 1.931954 | 0.3974318 | 1.088128 | 1 |
| ENSG00000164292.12 | RHOBTB3 | -0.354386 | 0.7002524 | -0.144078 | 0.1708551 | 1.222931 | 0.006795695 | 0.786496 | 0.105236 | 0.492324 | 0.6561467 | -0.440231 | 0.6885065 |
| ENSG00000164305.17 | CASP3 | -0.096798 | 0.8006286 | -0.226689 | 0.7336348 | -0.041898 | 0.004296811 | 0.291159 | 0.6398564 | 0.615807 | 0.06075365 | 0.802924 | 0.9087282 |
| ENSG00000164323.12 | CFAP97 | -0.616895 | 0.5893271 | -0.199221 | 0.08410382 | 0.594805 | 0.007799552 | 0.602243 | 0.8224647 | 0.148642 | 0.3688237 | -0.041721 | 0.6829816 |
| ENSG00000164327.12 | RICTOR | -1.483519 | 0.6882604 | -0.105311 | 0.652973 | 0.359195 | 0.7437308 | 3.635583 | 0.000257207 | 2.622702 | 2.91E-05 | 1.665131 | 0.951232 |
| ENSG00000164338.9 | UTP15 | -0.632046 | 0.563505 | 0.109971 | 0.9069847 | 0.424204 | 0.8300085 | 1.639623 | 0.02232997 | 1.264118 | 0.9368147 | 0.480847 | 0.5344421 |
| ENSG00000164346.9 | NSA2 | -1.508698 | 0.4764974 | 0.282179 | 1 | 0.388698 | 0.3549672 | 0.233389 | 0.2141464 | -0.452121 | 0.01988356 | 0.414185 | 0.6829816 |
| ENSG00000164347.17 | GFM2 | 1.658442 | 0.3439898 | -0.237196 | 0.5883005 | 0.93836 | 0.5290845 | 1.211154 | 0.02522358 | 0.982395 | 0.5144657 | 2.083697 | 0.9147227 |
| ENSG00000164398.12 | ACSL6 | -0.123971 | 0.6214382 | -0.974992 | 0.9559854 | -3.020121 | 0.02590966 | 1.125595 | 0.03048442 | 1.633908 | 0.4173086 | -5.874044 | 0.8479431 |
| ENSG00000164405.10 | UQCRQ | -0.103573 | 0.01706039 | 0.013514 | 0.2884286 | -0.590332 | 1.44E-21 | -0.207165 | 0.03560962 | -0.349237 | 3.06E-07 | -0.059197 | 0.4004082 |
| ENSG00000164414.16 | SLC35A1 | 1.016086 | 0.8587992 | -0.192427 | 0.9787499 | 0.368744 | 0.7437308 | 1.705028 | 0.001549484 | 1.091051 | 0.7307303 | -3.462701 | 1 |
| ENSG00000164418.19 | GRIK2 | -0.454596 | 0.6214382 | -0.520037 | 0.5480151 | 2.080448 | 0.1618415 | 1.498298 | 0.001602625 | 1.185778 | 0.6669739 | 2.976264 | 0.7767294 |
| ENSG00000164434.11 | FABP7 | -4.42586 | 0.08029934 | -0.171874 | 0.4002695 | 0.106398 | 0.8996595 | -1.11163 | 0.00133523 | -1.045234 | 1.84E-05 | -0.407081 | 0.5606863 |
| ENSG00000164442.9 | CITED2 | -0.088373 | 0.8867323 | 0.137427 | 0.9329105 | 0.515736 | 0.8382521 | 0.706449 | 0.5930297 | 2.233011 | 0.03056663 | 5.592981 | 0.06111237 |
| ENSG00000164463.12 | CREBRF | -0.711733 | 0.8006286 | 0.033089 | 0.5953732 | 1.005238 | 1.90E-05 | 0.90083 | 0.4608889 | 1.266497 | 0.5196843 | 0.327823 | 0.186512 |
| ENSG00000164466.12 | SFXN1 | 0.267016 | 0.6792949 | -0.053584 | 0.6605989 | 0.978966 | 0.0627237 | -0.070175 | 0.003677139 | 0.844234 | 0.3974318 | -1.009213 | 0.6829816 |
| ENSG00000164506.14 | STXBP5 | -2.045925 | 0.8006286 | -0.889262 | 0.8703522 | 0.092283 | 0.8635439 | 0.801535 | 0.01996295 | 1.859244 | 0.08770913 | 2.884626 | 0.8479431 |
| ENSG00000164543.5 | STK17A | 0.095526 | 0.6014343 | -0.01314 | 1 | -0.027732 | 0.04815049 | 0.35241 | 0.312938 | 1.414987 | 0.4144335 | -0.563412 | 0.8479431 |
| ENSG00000164548.10 | TRA2A | 0.191964 | 0.1607609 | -0.212752 | 0.8887798 | 0.02493 | 0.7046867 | 0.946765 | 1.14E-08 | 0.549237 | 0.5196843 | -0.035343 | 0.9052108 |
| ENSG00000164587.11 | RPS14 | -0.103108 | 0.8867323 | 0.053062 | 0.9222921 | -0.001255 | 0.7720073 | 0.009624 | 0.3467508 | -0.02032 | 0.005296486 | -0.161307 | 0.3762381 |
| ENSG00000164609.9 | SLU7 | 0.859961 | 0.6214382 | 0.411792 | 0.6527736 | 0.969649 | 9.92E-06 | 0.014321 | 0.8638749 | 0.183772 | 0.6598924 | 1.07507 | 0.9545867 |
| ENSG00000164610.8 | RP9 | -2.799336 | 0.4406957 | -0.366464 | 0.5953732 | -0.394551 | 0.04230269 | -0.690223 | 0.1219241 | 0.258826 | 1 | -0.017169 | 0.6885065 |
| ENSG00000164611.12 | PTTG1 | 1.239464 | 0.2681166 | -0.429091 | 0.2670385 | 0.077407 | 0.7437308 | -0.279136 | 0.001221598 | 0.530275 | 0.9259304 | -0.23907 | 0.8285762 |
| ENSG00000164615.4 | CAMLG | -0.154063 | 0.9067124 | 0.271577 | 0.9577957 | 0.734672 | 0.01121461 | -0.159066 | 0.2960427 | -0.246087 | 0.00042409 | -0.621035 | 0.6885065 |
| ENSG00000164631.18 | ZNF12 | 0.104767 | 0.6016874 | 1.139031 | 0.4148108 | -0.044246 | 0.6998472 | 2.406136 | 0.09729777 | 2.051699 | 0.04581517 | 0.112291 | 0.8949489 |
| ENSG00000164663.13 | USP49 | -0.550038 | 0.9047689 | 0.97359 | 1 | 1.265573 | 0.2227885 | 4.13562 | 1.38E-05 | 0.907408 | 0.560774 | -2.460818 | 0.9427738 |
| ENSG00000164684.13 | ZNF704 | -1.879378 | 0.8006286 | -1.076523 | 0.4375617 | 0.583145 | 0.02583934 | 2.390244 | 0.001936852 | 0.874566 | 0.1355518 | 1.236073 | 0.7767294 |
| ENSG00000164687.10 | FABP5 | 2.067709 | 0.09749406 | -0.283722 | 0.04797328 | 0.207298 | 0.5290845 | 0.126729 | 0.2699721 | 0.262609 | 0.3257176 | -0.354635 | 0.05841333 |
| ENSG00000164741.14 | DLC1 | 0.510918 | 0.9057626 | -0.978794 | 0.02412696 | -1.482479 | 0.4382561 | 3.212264 | 0.04346603 | 2.618248 | 0.8888294 | -1.382762 | 1 |
| ENSG00000164742.14 | ADCY1 | 0.116645 | 0.8587992 | 0.039824 | 0.883333 | -2.201257 | 0.2728726 | 0.802653 | 0.04856638 | 0.605301 | 0.8387831 | -1.422359 | 0.6885065 |
| ENSG00000164754.12 | RAD21 | 0.028142 | 0.6214382 | 0.227231 | 0.6540268 | 0.655439 | 3.57E-09 | 0.259467 | 0.7345366 | 0.137989 | 0.6539066 | -0.387606 | 0.6829816 |
| ENSG00000164823.9 | OSGIN2 | 0.672661 | 0.739749 | -0.351955 | 0.8953649 | 1.513931 | 0.7317752 | 2.322059 | 0.02462981 | 0.534795 | 0.4481846 | -0.308288 | 0.8949489 |
| ENSG00000164828.17 | SUN1 | 0.123001 | 0.4246534 | 0.247705 | 0.9108153 | 0.664513 | 0.5290845 | 1.259412 | 0.000188115 | 1.291705 | 0.1100045 | -0.127993 | 0.7354313 |
| ENSG00000164896.19 | FASTK | 0.125903 | 0.7693197 | 0.485854 | 0.6745307 | -0.586317 | 0.02449485 | 0.672302 | 0.7308295 | -0.185974 | 0.1895901 | 1.009078 | 0.6829816 |
| ENSG00000164897.12 | TMUB1 | -2.090469 | 0.3004379 | 0.193766 | 0.846655 | -1.917431 | 2.56E-11 | 2.247946 | 0.005035035 | 2.139794 | 0.02294687 | -1.257159 | 0.9617345 |
| ENSG00000164916.10 | FOXK1 | -0.097406 | 0.8006902 | -0.171302 | 0.6527736 | 0.294505 | 0.04102475 | 2.069138 | 1.35E-15 | 1.554018 | 0.005245519 | -1.00548 | 0.7767294 |
| ENSG00000164919.10 | COX6C | -0.012201 | 0.3946735 | -0.353507 | 8.19E-06 | -0.476886 | 1.59E-08 | -0.222685 | 2.65E-05 | -0.275345 | 0.000159722 | -0.220707 | 0.03646438 |
| ENSG00000164924.17 | YWHAZ | -0.178935 | 0.8006286 | 0.203166 | 0.652973 | 0.269455 | 2.45E-05 | 0.176532 | 0.7061815 | 0.482005 | 0.9175623 | 0.214208 | 0.6185816 |
| ENSG00000164934.13 | DCAF13 | -0.344679 | 0.8937801 | 0.398026 | 0.4130975 | -0.162025 | 0.5044352 | 0.008942 | 0.8502343 | 0.108767 | 0.01677568 | 0.094752 | 0.6885065 |
| ENSG00000164941.13 | INTS8 | -0.041573 | 0.9183025 | 0.187956 | 0.8528295 | 1.332938 | 0.02397417 | 1.639974 | 0.000187612 | 1.335635 | 0.08601907 | -0.566015 | 0.6829816 |
| ENSG00000164944.11 | KIAA1429 | -1.367871 | 0.6214382 | 0.133249 | 0.7548431 | 0.200619 | 0.057908 | 2.322168 | 0.006958854 | 1.548938 | 0.148644 | 0.085661 | 0.6885065 |
| ENSG00000164949.7 | GEM | 1.034266 | 0.6014343 | -0.342068 | 0.9262666 | -0.491745 | 0.9425542 | 0.666742 | 0.01045382 | 1.097887 | 0.6739559 | -1.894199 | 0.8881296 |
| ENSG00000164951.15 | PDP1 | -0.252027 | 0.6214382 | 0.748512 | 0.7364864 | 0.420566 | 0.7621311 | 1.05196 | 0.04148071 | 1.140795 | 0.1191155 | -1.324897 | 0.1987453 |
| ENSG00000164985.14 | PSIP1 | -0.463554 | 0.9345595 | -0.066379 | 0.3002505 | -0.69104 | 0.5290845 | 1.038659 | 0.008380351 | 0.437223 | 0.2875639 | 0.187991 | 0.6885065 |
| ENSG00000165029.15 | ABCA1 | 0.860991 | 0.6574918 | 0.479307 | 0.1919448 | -1.330428 | 0.05489036 | 1.063979 | 0.002878396 | 1.340067 | 0.4879833 | 0.41853 | 0.8638276 |
| ENSG00000165055.15 | METTL2B | 0.083084 | 1 | 0.302802 | 1 | 0.109783 | 0.5588773 | 0.904968 | 0.02741529 | 1.161655 | 1 | -0.580461 | 0.8489889 |
| ENSG00000165119.18 | HNRNPK | 0.07625 | 0.4764974 | -0.125408 | 0.5876088 | 0.25121 | 0.1992941 | 0.265853 | 0.9513599 | -0.122657 | 0.004991493 | 0.037338 | 0.4304069 |
| ENSG00000165156.14 | ZHX1 | -0.503961 | 0.9695288 | -0.192899 | 0.09893397 | 0.078771 | 0.0133227 | 1.695963 | 0.1496981 | 0.802544 | 0.3974318 | -2.214736 | 0.8247617 |
| ENSG00000165185.14 | KIAA1958 | -1.474864 | 0.6014343 | -0.751188 | 0.5395105 | 0.429143 | 0.3734283 | 0.82393 | 0.1125707 | 2.394753 | 0.04771425 | 2.545003 | 0.6196823 |
| ENSG00000165194.14 | PCDH19 | 0.946524 | 0.9047689 | 0.01541 | 0.8703522 | 1.145905 | 0.00040418 | 1.489166 | 0.3740072 | -1.328367 | 0.4071955 | 0.373957 | 1 |
| ENSG00000165209.18 | STRBP | -0.028202 | 0.8006286 | -0.285917 | 0.6200225 | -0.52238 | 0.7081668 | 1.68383 | 0.01033382 | 0.956712 | 1 | 0.385803 | 0.9077431 |
| ENSG00000165219.21 | GAPVD1 | 0.496 | 0.7420743 | 0.44376 | 0.7383998 | 0.544211 | 0.001482617 | -0.343557 | 0.1860894 | 1.461785 | 0.201852 | 0.646732 | 0.8931285 |
| ENSG00000165244.6 | ZNF367 | 1.111728 | 0.8587992 | -0.672336 | 0.5012872 | -0.1457 | 0.4382561 | 2.411347 | 0.04187258 | 0.999415 | 0.4278631 | 2.321683 | 0.8949489 |
| ENSG00000165264.10 | NDUFB6 | 0.041046 | 0.6214382 | -0.059375 | 0.8263112 | -0.137272 | 0.01439134 | -0.196289 | 0.005283804 | -0.241784 | 7.81E-06 | -0.275611 | 0.2949856 |
| ENSG00000165271.16 | NOL6 | -0.296315 | 0.6014343 | -0.832529 | 0.5905303 | -0.834798 | 0.5363627 | 2.126723 | 7.52E-05 | -0.797458 | 0.643255 | 0.482828 | 0.9661659 |
| ENSG00000165283.15 | STOML2 | -0.67401 | 0.2614863 | 0.110165 | 0.5953732 | 0.177454 | 0.108041 | 0.131802 | 0.02530799 | 0.251232 | 0.003546663 | -0.345724 | 0.6185816 |
| ENSG00000165338.16 | HECTD2 | -0.696173 | 0.6214382 | -0.956845 | 0.704767 | 3.278488 | 0.02483941 | 0.436305 | 0.9073329 | 1.669987 | 0.04430596 | 2.302705 | 0.9532204 |
| ENSG00000165355.7 | FBXO33 | 0.911665 | 0.7736255 | 0.234835 | 0.9376227 | -0.273805 | 0.8581499 | 3.077779 | 0.002228985 | 2.59822 | 0.2087791 | 1.551274 | 0.6885065 |
| ENSG00000165392.9 | WRN | -1.367028 | 0.714717 | 0.956184 | 0.7017709 | 0.823624 | 0.001014832 | 0.618832 | 0.1640101 | 1.136836 | 0.5196843 | 1.900002 | 0.6185816 |
| ENSG00000165434.7 | PGM2L1 | 0.100678 | 0.8006286 | -0.33475 | 0.592336 | 0.403085 | 0.000169802 | 0.73903 | 0.002188595 | 1.076432 | 0.5196843 | -1.151039 | 0.9506884 |
| ENSG00000165475.13 | CRYL1 | -0.682427 | 0.8016328 | -0.486522 | 0.4011918 | -0.272484 | 0.5731622 | -0.508118 | 0.001448862 | -2.57137 | 0.009939173 | 1.479078 | 0.5628123 |
| ENSG00000165494.10 | PCF11 | 0.8422 | 1 | 0.54715 | 0.6527736 | 0.65512 | 0.001612906 | 0.264318 | 0.4679723 | 1.042848 | 0.0453052 | 1.440254 | 0.3848319 |
| ENSG00000165502.6 | RPL36AL | 0.107239 | 0.6214382 | -0.005526 | 0.6263369 | -0.258148 | 0.0030243 | 0.243873 | 0.1406301 | -0.72881 | 0.004839936 | 0.710865 | 0.995591 |
| ENSG00000165516.10 | KLHDC2 | 1.210718 | 0.563505 | -0.082099 | 0.5295559 | -1.108649 | 0.000429922 | 1.173403 | 0.6476974 | 1.011601 | 0.2011388 | -0.801724 | 0.7775901 |
| ENSG00000165525.17 | NEMF | 0.072725 | 0.6918831 | 0.082444 | 0.8653057 | 0.840213 | 0.005032306 | 1.144206 | 0.09114691 | 0.288636 | 0.5891826 | 0.028052 | 0.9725802 |
| ENSG00000165566.12 | AMER2 | 0.300794 | 1 | 0.043712 | 1 | 0.51684 | 0.4410989 | 1.049025 | 0.7144938 | 1.287237 | 0.01432977 | -1.059123 | 1 |
| ENSG00000165629.19 | ATP5C1 | 0.324276 | 0.4764974 | 0.209041 | 1 | 0.079099 | 0.4203906 | -0.105435 | 0.002338666 | -0.060959 | 0.02270501 | 0.979787 | 0.7953308 |
| ENSG00000165632.7 | TAF3 | -1.125101 | 0.6214382 | 0.062498 | 0.4886958 | 1.207114 | 0.02688048 | 0.621645 | 0.09487473 | 1.101433 | 0.8785196 | -2.279553 | 0.6910517 |
| ENSG00000165637.13 | VDAC2 | -0.43679 | 0.6214382 | 0.183563 | 0.6527736 | 0.116356 | 0.7389126 | 0.171944 | 0.03567679 | -0.432412 | 0.003187902 | 0.017227 | 0.6829816 |
| ENSG00000165650.11 | PDZD8 | -0.327692 | 1 | -0.26591 | 0.7615373 | 0.911857 | 0.1538296 | 2.061973 | 0.01330514 | 2.465513 | 0.01475557 | 1.044373 | 0.8949489 |
| ENSG00000165669.13 | FAM204A | -0.07667 | 0.8006902 | -0.092991 | 0.9574026 | 0.363317 | 0.3994594 | -0.076378 | 0.7917638 | -0.483497 | 0.04619379 | -0.755768 | 0.4301971 |
| ENSG00000165671.18 | NSD1 | -0.354124 | 0.7135744 | -0.497839 | 0.1054706 | 0.934719 | 0.06378607 | 1.414806 | 0.001653081 | 0.912382 | 0.1399344 | 0.807481 | 0.961427 |
| ENSG00000165689.16 | SDCCAG3 | -0.284165 | 0.970075 | -0.48006 | 0.5953732 | 0.02857 | 0.7149145 | 1.223409 | 0.1617509 | 2.58065 | 0.000459067 | -1.305887 | 0.8167193 |
| ENSG00000165699.13 | TSC1 | 0.350377 | 0.9426638 | 0.239183 | 0.797408 | 0.446328 | 0.5290845 | 1.16983 | 8.19E-07 | 1.215365 | 0.004758115 | -0.042936 | 0.7567976 |
| ENSG00000165724.5 | ZMYND19 | -0.41695 | 0.8587992 | -0.283562 | 0.949642 | 0.009264 | 0.07651684 | 0.853184 | 0.008823624 | 1.140637 | 0.1505337 | -0.076075 | 1 |
| ENSG00000165732.12 | DDX21 | 0.205978 | 0.6214382 | 0.456136 | 0.9979553 | 0.521868 | 5.90E-05 | 0.69732 | 0.8472142 | 0.227361 | 0.2363864 | -0.345308 | 0.7767294 |
| ENSG00000165733.7 | BMS1 | -0.514235 | 0.3254792 | 0.258525 | 0.9243892 | 1.08677 | 0.02566289 | -0.807767 | 0.8093083 | 0.593766 | 0.7986287 | 0.949734 | 0.806946 |
| ENSG00000165782.10 | TMEM55B | -0.647623 | 0.6214382 | 0.82921 | 0.09100182 | 0.545025 | 0.8998163 | 3.435876 | 0.001068983 | 3.291442 | 0.4248747 | 1.895161 | 0.868568 |
| ENSG00000165795.21 | NDRG2 | -2.05985 | 0.01985512 | -0.64524 | 0.1809109 | -0.424898 | 0.5290845 | 2.575802 | 0.7068176 | 0.683019 | 0.5196843 | 0.790288 | 0.9597711 |
| ENSG00000165801.9 | ARHGEF40 | 0.468052 | 0.4764974 | -0.157367 | 0.9669919 | -1.027557 | 0.3276184 | 2.43176 | 0.002283922 | 1.004207 | 0.1691344 | 1.893752 | 0.348294 |
| ENSG00000165802.19 | NSMF | 0.149305 | 0.563505 | 0.01794 | 1 | 0.001695 | 0.4255187 | 1.265964 | 0.001456391 | 1.774916 | 9.12E-05 | 0.296053 | 0.6829816 |
| ENSG00000165813.16 | CCDC186 | 1.165868 | 0.8006286 | 0.587091 | 0.6527736 | 0.510165 | 0.7437308 | -1.599671 | 0.4919368 | 2.264859 | 0.03847182 | 0.124933 | 0.7767294 |
| ENSG00000165821.11 | SALL2 | -3.532522 | 0.3946735 | 0.450034 | 1 | -0.032917 | 0.5290845 | 1.527816 | 0.02995742 | 2.81622 | 0.05546865 | 2.618355 | 0.6885065 |
| ENSG00000165868.12 | HSPA12A | 1.604285 | 1 | -0.165572 | 0.9648958 | -0.058136 | 1 | 2.845809 | 0.002260604 | 2.621478 | 0.3871425 | -3.956075 | 0.8949489 |
| ENSG00000165916.8 | PSMC3 | -0.221866 | 0.9816863 | 0.173118 | 0.9683094 | -0.083005 | 0.2655881 | 0.040129 | 0.5271934 | -0.436462 | 0.002514978 | 0.220154 | 0.5394145 |
| ENSG00000165997.4 | ARL5B | -1.181654 | 0.6214382 | 0.043384 | 0.9821162 | 1.324545 | 0.001345929 | 1.133161 | 0.07646099 | 0.834938 | 0.3559576 | -0.109039 | 0.6517664 |
| ENSG00000166012.14 | TAF1D | -0.178883 | 0.6214382 | -0.02324 | 0.7594721 | 0.630718 | 4.42E-07 | 1.073476 | 0.1016405 | 0.787453 | 0.9340732 | 0.511581 | 0.6885065 |
| ENSG00000166037.10 | CEP57 | -0.56228 | 0.6214382 | 0.093573 | 0.6200225 | 1.127467 | 3.28E-07 | 0.085554 | 0.3372721 | 0.129024 | 0.5196843 | 0.591707 | 0.9699068 |
| ENSG00000166068.12 | SPRED1 | -0.234661 | 1 | -0.196891 | 0.5306483 | 0.490178 | 0.03061788 | 0.627922 | 0.2048887 | 0.690646 | 0.5750522 | 1.285285 | 0.6191255 |
| ENSG00000166086.12 | JAM3 | 0.558109 | 0.563505 | 0.583982 | 0.8519557 | 0.337716 | 0.2569818 | 0.802923 | 0.02482187 | 0.606255 | 1 | -0.06852 | 0.9726757 |
| ENSG00000166128.12 | RAB8B | 1.020731 | 0.4845308 | 0.034482 | 1 | 1.164283 | 0.04802112 | 1.088128 | 0.02984614 | 1.138362 | 0.2376995 | 2.651491 | 0.5736188 |
| ENSG00000166136.15 | NDUFB8 | -0.334727 | 0.4923792 | -0.356282 | 0.7383998 | -0.739332 | 5.25E-08 | 0.309111 | 0.8759049 | 0.177071 | 0.1519109 | -0.671887 | 0.01006513 |
| ENSG00000166164.15 | BRD7 | -0.537763 | 0.8006286 | -0.234736 | 0.2626761 | 0.37762 | 0.5233268 | 0.765129 | 6.43E-05 | 1.790273 | 0.01074803 | 0.403695 | 0.7817802 |
| ENSG00000166165.12 | CKB | -0.737888 | 0.1607609 | -0.274754 | 0.02404456 | -0.771398 | 3.97E-09 | 0.474507 | 0.1231427 | 0.550331 | 0.4667651 | -0.748855 | 0.04529194 |
| ENSG00000166171.12 | DPCD | 0.449549 | 0.9367821 | -0.102065 | 0.7601699 | 0.34448 | 0.4077154 | -0.081817 | 0.239284 | -1.438935 | 0.02715552 | -0.356974 | 0.4629017 |
| ENSG00000166181.12 | API5 | -1.734008 | 0.6201989 | 0.361397 | 1 | 0.544056 | 0.01705104 | 0.16253 | 0.9137041 | 0.571972 | 0.9738386 | -1.41965 | 0.7935528 |
| ENSG00000166197.16 | NOLC1 | -0.233259 | 0.8006286 | -0.196336 | 0.24394 | 0.877169 | 0.000554427 | 0.655728 | 0.9133698 | 0.060207 | 0.7503278 | -0.074309 | 0.6167027 |
| ENSG00000166200.14 | COPS2 | -0.350402 | 0.845546 | 0.423331 | 0.6126945 | 0.843515 | 0.0328671 | -0.179517 | 0.05637094 | 0.935285 | 0.6180321 | 1.242564 | 0.7870028 |
| ENSG00000166224.16 | SGPL1 | 0.70962 | 0.7811112 | -0.10183 | 0.8429144 | 0.032407 | 1 | -0.157134 | 0.03470727 | 1.378844 | 0.320077 | -0.166525 | 0.8813817 |
| ENSG00000166226.12 | CCT2 | 0.563001 | 0.6214382 | -0.215014 | 0.2731087 | 0.374165 | 0.005457999 | 0.433312 | 0.7505319 | 0.427157 | 0.621723 | 0.45472 | 0.7973567 |
| ENSG00000166233.12 | ARIH1 | 0.309497 | 0.8867323 | 0.051042 | 0.570279 | 1.184301 | 3.76E-05 | 1.087456 | 0.02891696 | 1.624309 | 0.003737508 | -2.191934 | 0.9526606 |
| ENSG00000166266.13 | CUL5 | 1.815065 | 0.4665035 | -0.084967 | 0.7044612 | 0.642045 | 0.005631247 | 1.173039 | 0.001639525 | 0.485207 | 0.8844564 | 0.38012 | 0.7901525 |
| ENSG00000166326.6 | TRIM44 | -0.936991 | 0.529676 | -0.184138 | 0.5953732 | 0.507173 | 0.1278455 | 0.812898 | 0.001964851 | 0.508434 | 0.7360465 | 0.711811 | 0.9147227 |
| ENSG00000166337.9 | TAF10 | 0.796763 | 0.9054592 | -0.222942 | 0.6527736 | -0.738767 | 0.04028631 | 0.837673 | 0.1487277 | 2.100195 | 0.0619892 | 0.535313 | 0.7868517 |
| ENSG00000166341.7 | DCHS1 | 1.541197 | 0.8587992 | -0.230916 | 0.8591657 | -1.282436 | 0.1529176 | 1.563566 | 0.003228635 | 1.405964 | 0.02779405 | -3.510188 | 0.2525489 |
| ENSG00000166411.13 | IDH3A | -0.189245 | 0.6014343 | 0.031724 | 0.7976384 | 0.611746 | 0.574715 | 0.889073 | 0.0381379 | 0.31728 | 0.5196843 | -1.825495 | 0.8479431 |
| ENSG00000166441.12 | RPL27A | -0.758772 | 3.00E-09 | -0.128301 | 0.4403562 | -0.105445 | 0.2601653 | 0.660685 | 1.33E-20 | 0.43313 | 0.08374036 | -0.227042 | 0.2045823 |
| ENSG00000166450.12 | PRTG | -3.642643 | 0.4764974 | -0.321978 | 0.5517607 | 1.380332 | 0.01424104 | 0.463335 | 1 | 0.399766 | 0.4074192 | inf | 0.6829816 |
| ENSG00000166454.9 | ATMIN | -1.500263 | 0.5852558 | 0.085302 | 0.8263112 | 0.208508 | 0.9253493 | 3.081061 | 1.78E-05 | 0.128559 | 0.4430726 | 0.837232 | 0.7520694 |
| ENSG00000166471.10 | TMEM41B | 0.539228 | 0.4406957 | 0.662953 | 0.6527736 | 0.212714 | 0.01287997 | 0.315433 | 0.8095051 | 1.588528 | 0.2518976 | 1.964368 | 0.8949489 |
| ENSG00000166477.12 | LEO1 | 0.35982 | 0.5151741 | -0.06592 | 0.6077765 | 0.056587 | 0.4401212 | 0.032253 | 0.4919368 | -0.401802 | 0.01732668 | 0.289162 | 0.7812339 |
| ENSG00000166479.9 | TMX3 | -0.092968 | 0.6214382 | 0.182729 | 0.9473701 | 0.779089 | 0.3748806 | 1.010674 | 0.008359651 | -0.257051 | 0.3974318 | 0.108388 | 0.7175744 |
| ENSG00000166483.10 | WEE1 | 0.779572 | 0.6214382 | 0.208067 | 0.8703522 | -0.215474 | 0.5290845 | 1.443369 | 0.000126658 | 0.567245 | 0.01565028 | -0.424777 | 1 |
| ENSG00000166503.8 | RP11-382A20.3 | -0.397434 | 0.6991852 | 0.134002 | 0.9862305 | 0.666318 | 0.4148722 | 0.784349 | 0.008309395 | 0.383661 | 1 | -0.130389 | 0.9517616 |
| ENSG00000166508.17 | MCM7 | 2.089414 | 0.9183025 | 0.050856 | 0.8703522 | 0.304901 | 0.5290845 | 0.887606 | 0.03452489 | -0.072179 | 0.1772087 | 0.968937 | 0.5344421 |
| ENSG00000166532.15 | RIMKLB | 0.249617 | 0.8867323 | 0.298822 | 0.8263112 | 0.361647 | 0.06314447 | 0.336147 | 0.01455163 | 0.041603 | 0.8465018 | 1.774907 | 0.9589332 |
| ENSG00000166562.8 | SEC11C | 0.305309 | 0.6214382 | -0.037669 | 0.8450768 | -0.06611 | 0.2879524 | 0.08591 | 0.05619092 | 0.098939 | 0.01316432 | -0.440325 | 0.2100537 |
| ENSG00000166582.9 | CENPV | 0.952641 | 1 | -0.248245 | 0.1063235 | -0.844222 | 0.7354314 | 0.66012 | 0.04032816 | 1.222601 | 0.1102536 | -0.19184 | 0.6185816 |
| ENSG00000166595.11 | FAM96B | -0.470596 | 0.6214382 | 0.361013 | 0.3251036 | -0.215903 | 0.000596594 | -0.503005 | 0.001354845 | 0.364678 | 0.01472525 | -0.379626 | 0.4553016 |
| ENSG00000166598.12 | HSP90B1 | -0.0989 | 0.3946735 | 0.324104 | 1.18E-05 | 0.509217 | 2.63E-10 | 0.460269 | 0.4254824 | 0.260061 | 0.3289868 | 0.3992 | 1 |
| ENSG00000166681.13 | NGFRAP1 | -0.331957 | 0.8689227 | -0.332623 | 0.05116306 | 0.012786 | 0.008388637 | -0.13766 | 0.008427881 | 0.07927 | 0.1396126 | 0.064637 | 0.4553016 |
| ENSG00000166685.11 | COG1 | -0.994145 | 0.6214382 | 0.217282 | 1 | 1.284278 | 0.4737144 | 1.380929 | 0.007498335 | 1.018908 | 0.2153128 | 0.344203 | 0.9581586 |
| ENSG00000166710.17 | B2M | 0.156969 | 0.06605898 | 0.188616 | 8.19E-08 | -0.440162 | 3.95E-10 | -0.328566 | 3.42E-09 | 0.203821 | 0.007184823 | 0.766724 | 0.01991945 |
| ENSG00000166734.18 | CASC4 | -0.120799 | 0.7146291 | -0.443549 | 0.3150467 | 0.282762 | 0.4008897 | 1.145942 | 0.02232231 | 0.653069 | 0.725385 | 1.115782 | 0.7753492 |
| ENSG00000166747.12 | AP1G1 | -0.261657 | 1 | -0.533468 | 0.5953732 | 1.106416 | 0.01304157 | 1.714686 | 1.85E-05 | 0.638212 | 0.3553472 | -0.833008 | 0.8723012 |
| ENSG00000166780.10 | C16orf45 | -0.465737 | 0.6014343 | -0.645604 | 0.09470648 | 0.000399 | 0.7887553 | 0.467171 | 0.4846975 | -0.504795 | 0.000168042 | -0.06371 | 0.6517664 |
| ENSG00000166783.19 | KIAA0430 | 0.368772 | 0.7736255 | 0.309098 | 0.9495034 | 0.939942 | 0.1352928 | 1.966306 | 0.006916264 | 0.53782 | 0.03923004 | -0.448248 | 0.9724348 |
| ENSG00000166794.4 | PPIB | -0.203467 | 0.934454 | 0.011689 | 0.6527736 | -0.013628 | 0.1929317 | -0.263547 | 2.53E-05 | -0.522138 | 3.15E-05 | 0.464837 | 1 |
| ENSG00000166823.5 | MESP1 | 0.227956 | 0.9589706 | -0.128664 | 0.853835 | -3.002907 | 0.003850736 | 0.810671 | 0.8217677 | 0.393248 | 0.9147336 | 1.060624 | 0.9602719 |
| ENSG00000166833.19 | NAV2 | -2.972581 | 0.6014343 | -0.214428 | 0.8678846 | 0.060948 | 0.8100781 | 1.204587 | 8.69E-05 | 1.837573 | 0.3183218 | -0.793423 | 0.569325 |
| ENSG00000166847.9 | DCTN5 | -0.03261 | 0.980174 | 0.708973 | 0.6799811 | 0.383864 | 0.9151684 | 0.748652 | 0.09170208 | 1.992124 | 0.000405058 | -2.179719 | 0.9147227 |
| ENSG00000166848.5 | TERF2IP | 0.140422 | 0.8867323 | -0.00227 | 0.6527736 | 0.156673 | 0.6031248 | 0.266442 | 0.8589194 | -0.416972 | 1.72E-06 | -0.201252 | 0.8479431 |
| ENSG00000166912.16 | MTMR10 | -1.090329 | 0.6214382 | -0.420055 | 0.979665 | 0.725828 | 0.6876625 | 1.433177 | 0.009822711 | 2.072672 | 0.3339985 | -0.561354 | 0.8479431 |
| ENSG00000166922.8 | SCG5 | 0.021856 | 0.7063838 | 0.142247 | 0.8703522 | 0.960113 | 0.1683791 | 0.279808 | 0.7427511 | -0.795422 | 0.02384206 | 1.55079 | 0.6885065 |
| ENSG00000166925.8 | TSC22D4 | -1.556445 | 0.01068364 | -0.233507 | 0.01857467 | -0.013183 | 0.02630396 | 0.388004 | 0.3849974 | -0.179006 | 0.1912543 | -1.297628 | 0.05044736 |
| ENSG00000166938.12 | DIS3L | -1.617448 | 0.6792949 | -0.280928 | 0.5394427 | 0.410468 | 1 | 2.770651 | 0.007107074 | 0.121809 | 0.9171552 | -1.128909 | 0.6910517 |
| ENSG00000166963.12 | MAP1A | 0.925089 | 0.3610267 | 0.186676 | 0.9439293 | -0.799747 | 0.9516878 | 0.966813 | 0.004146686 | 1.337467 | 0.1202747 | -0.581426 | 0.9581586 |
| ENSG00000166987.14 | MBD6 | 0.242916 | 0.714717 | -0.203665 | 0.7811664 | -1.057401 | 0.5290845 | 2.278201 | 0.02352122 | -0.730976 | 0.05660876 | -1.294842 | 0.9192932 |
| ENSG00000167004.12 | PDIA3 | 0.506181 | 0.005239251 | 0.56856 | 0.000902393 | -0.101535 | 0.3393299 | 0.477784 | 0.5932107 | -0.487413 | 0.05169547 | 1.09376 | 0.6885065 |
| ENSG00000167005.13 | NUDT21 | -0.871681 | 0.6387595 | 0.302105 | 0.936943 | 0.771104 | 0.1819643 | 1.247975 | 0.009888774 | -0.026284 | 0.8274257 | -2.135747 | 0.8168211 |
| ENSG00000167011.8 | NAT16 | 1.315783 | 0.6214382 | -2.792301 | 0.2251635 | 0.652512 | 0.9423128 | 1.243701 | 0.7786328 | 2.494473 | 0.01432977 | 0.365209 | 0.9476473 |
| ENSG00000167081.16 | PBX3 | 1.476758 | 0.6214382 | 0.237265 | 0.8617702 | 0.301226 | 0.7893626 | 0.942481 | 0.009778502 | -0.487528 | 0.7397084 | 0.649582 | 0.9125446 |
| ENSG00000167088.10 | SNRPD1 | -1.812236 | 0.9268352 | -0.14755 | 0.2042514 | 0.040868 | 0.1135741 | 0.111197 | 0.6483393 | -0.100051 | 0.005747048 | -0.130082 | 0.01381583 |
| ENSG00000167100.14 | SAMD14 | -0.674311 | 0.7683685 | 0.081663 | 0.7279842 | -1.046351 | 0.4791605 | 3.598163 | 0.000214893 | -0.456799 | 0.7356185 | -0.426862 | 0.7969605 |
| ENSG00000167106.11 | FAM102A | 0.014049 | 1 | 0.423736 | 0.6527736 | -0.329918 | 0.3357291 | 0.236872 | 0.4791525 | 3.101906 | 0.0292479 | 1.420336 | 0.2597332 |
| ENSG00000167110.16 | GOLGA2 | -0.702311 | 0.1956577 | -0.442596 | 0.2588352 | 0.131303 | 0.5772029 | 1.018707 | 0.001446231 | 1.120543 | 0.2382141 | -0.075017 | 0.7753492 |
| ENSG00000167191.11 | GPRC5B | 1.007816 | 0.1835496 | 0.03123 | 1 | -0.003382 | 0.8100781 | 1.73788 | 2.12E-05 | 0.537893 | 1 | 0.479139 | 0.9377765 |
| ENSG00000167193.7 | CRK | -0.446323 | 0.4764974 | -0.048943 | 0.6527736 | 0.345647 | 0.03466643 | 0.671848 | 0.05182835 | 0.11256 | 1 | 0.365735 | 0.6885065 |
| ENSG00000167196.13 | FBXO22 | -2.127853 | 0.563505 | 0.463721 | 0.8647057 | 0.398813 | 0.3119492 | 1.591154 | 0.009766286 | 0.863248 | 0.3796044 | -0.026395 | 0.9581586 |
| ENSG00000167220.11 | HDHD2 | -1.00907 | 0.5434436 | -0.40784 | 0.5670497 | -0.024558 | 0.4264835 | 2.045438 | 3.14E-09 | 1.105501 | 0.07002445 | -0.249583 | 0.6609803 |
| ENSG00000167232.13 | ZNF91 | -0.307939 | 0.8523587 | 0.067945 | 0.9222921 | 1.138355 | 0.000418452 | 1.514513 | 0.01250537 | 0.806683 | 0.5290784 | 2.678847 | 0.7408601 |
| ENSG00000167272.10 | POP5 | -0.188632 | 0.8006902 | 0.380402 | 0.9289724 | -0.970599 | 0.002323402 | -1.334982 | 0.08891507 | 1.135861 | 0.82104 | -0.125074 | 0.7447257 |
| ENSG00000167283.7 | ATP5L | -0.090375 | 0.5441969 | 0.004588 | 0.4403562 | -0.239011 | 0.000283221 | 0.019933 | 0.009420049 | -0.183181 | 0.000361072 | 0.145298 | 0.186512 |
| ENSG00000167302.9 | ENTHD2 | -0.624172 | 1 | -0.146397 | 0.8785769 | 1.619992 | 0.8454352 | 1.312742 | 0.5860824 | 4.277808 | 0.004978746 | 0.761921 | 0.6885065 |
| ENSG00000167371.16 | PRRT2 | 2.764298 | 1 | -3.787108 | 0.7383998 | -0.68038 | 0.4382561 | -0.742276 | 0.1827153 | 1.464456 | 0.003057286 | 1.522748 | 0.9318796 |
| ENSG00000167378.8 | IRGQ | -1.865102 | 0.6170632 | -0.616979 | 0.7881421 | 0.278539 | 0.01287997 | 2.184281 | 1.58E-07 | 1.422666 | 0.0205332 | 0.104015 | 0.7110899 |
| ENSG00000167397.14 | VKORC1 | 0.588541 | 0.6214382 | 0.333853 | 0.5868806 | -0.048326 | 0.01037681 | 0.327825 | 0.184613 | -0.061568 | 0.04854239 | 0.154363 | 0.9627244 |
| ENSG00000167460.14 | TPM4 | -0.29124 | 0.9789948 | 0.007305 | 0.8483182 | 0.358216 | 0.1900111 | -0.052335 | 0.1257066 | -0.108468 | 0.02704086 | -0.184595 | 0.6885065 |
| ENSG00000167468.16 | GPX4 | -0.250974 | 0.8006286 | -0.136608 | 0.5736784 | 0.082282 | 0.8592855 | -0.190307 | 0.04747859 | -0.619792 | 3.40E-09 | 0.789443 | 0.6267144 |
| ENSG00000167470.12 | MIDN | -1.755464 | 0.008844659 | -0.266347 | 0.2330536 | -0.267884 | 0.2168297 | 1.365238 | 1.57E-11 | 1.361577 | 8.86E-11 | 0.011838 | 0.9382274 |
| ENSG00000167491.17 | GATAD2A | -0.426905 | 0.6214382 | 0.047385 | 0.8125086 | 0.523329 | 0.7804142 | 0.979381 | 5.87E-07 | 1.24542 | 0.01297638 | -0.381216 | 0.6829816 |
| ENSG00000167515.10 | TRAPPC2L | -0.144991 | 0.9922212 | -0.223777 | 0.9618421 | 0.211412 | 0.5736278 | 0.20694 | 0.2602205 | -1.159991 | 0.001731107 | -0.580185 | 0.5733051 |
| ENSG00000167522.14 | ANKRD11 | -1.537109 | 0.3207272 | 0.397162 | 0.5818005 | 0.135885 | 0.2273012 | 0.832001 | 4.23E-05 | 1.073372 | 4.76E-07 | -0.900627 | 0.6885065 |
| ENSG00000167524.14 | SGK494 | 2.288643 | 0.6214382 | 1.465685 | 0.5296018 | -0.066615 | 0.4400377 | 4.720444 | 0.004269593 | 1.600184 | 0.404173 | -4.151057 | 0.5066477 |
| ENSG00000167526.13 | RPL13 | -0.216177 | 0.6214382 | -0.020501 | 0.936943 | 0.119247 | 0.2490947 | -0.230668 | 9.66E-07 | -0.090895 | 0.003418989 | 0.246814 | 0.6885065 |
| ENSG00000167528.12 | ZNF641 | 0.67902 | 0.6214382 | 1.374028 | 0.8910308 | -1.488394 | 0.5611694 | 3.07472 | 0.04392434 | 1.495655 | 0.8152449 | 2.511908 | 1 |
| ENSG00000167543.15 | TP53I13 | -1.060618 | 0.9047689 | 0.524645 | 0.6384858 | -3.266044 | 1.15E-05 | -0.981693 | 0.01820927 | 0.091886 | 0.902884 | 1.532346 | 0.9733288 |
| ENSG00000167548.14 | KMT2D | -3.642396 | 0.6214382 | -0.031807 | 0.7156572 | 0.082505 | 0.9135623 | -0.038321 | 0.7936091 | -0.464006 | 0.01324469 | 3.109046 | 0.6587745 |
| ENSG00000167552.13 | TUBA1A | 0.069189 | 0.3907286 | -0.151617 | 0.5445828 | 0.099978 | 0.01986339 | 0.011525 | 0.7875686 | 0.042155 | 1 | -0.173592 | 0.2321124 |
| ENSG00000167553.14 | TUBA1C | -1.152866 | 0.8006286 | -0.049392 | 0.7834199 | -0.440651 | 2.01E-08 | 0.153527 | 0.9837599 | 0.486603 | 0.5912862 | 0.136434 | 0.6556863 |
| ENSG00000167614.13 | TTYH1 | -7.360238 | 0.1341967 | -0.304625 | 0.09704869 | -0.473239 | 0.02167033 | 0.537372 | 1 | 0.835238 | 0.3974318 | -2.673716 | 0.9225091 |
| ENSG00000167615.16 | LENG8 | 2.28564 | 0.6214382 | 1.099648 | 0.2858022 | -1.651396 | 0.3495868 | 1.267279 | 3.89E-07 | 2.085983 | 0.000203276 | 0.143887 | 0.6885065 |
| ENSG00000167635.11 | ZNF146 | -0.512444 | 0.4407366 | -0.150522 | 0.7828571 | -0.269987 | 0.8100781 | 1.453545 | 5.39E-11 | 0.984581 | 0.05158486 | -1.010095 | 0.6885065 |
| ENSG00000167645.16 | YIF1B | -4.596458 | 0.563505 | 0.206108 | 0.5010139 | -0.32364 | 0.01705 | 0.296032 | 0.8224647 | -0.327451 | 0.2069824 | 0.742679 | 0.8469357 |
| ENSG00000167654.17 | ATCAY | -0.215266 | 0.9798559 | -0.69718 | 0.8703522 | 0.857654 | 0.3276184 | 1.560942 | 4.17E-05 | 1.004063 | 0.010784 | -0.206892 | 0.4217518 |
| ENSG00000167657.11 | DAPK3 | -0.201194 | 0.6014343 | -0.256324 | 1 | -0.558355 | 0.02590966 | 0.054473 | 0.7394347 | 1.97247 | 0.1134992 | 1.497241 | 0.9576568 |
| ENSG00000167658.15 | EEF2 | -0.373206 | 0.4540036 | 0.0679 | 0.8545414 | 0.474712 | 2.43E-16 | 0.504937 | 0.109567 | 0.706031 | 0.2550605 | -0.228527 | 0.1629698 |
| ENSG00000167680.15 | SEMA6B | -2.505025 | 0.5453725 | 1.17314 | 0.4487482 | 1.968259 | 0.5010887 | 2.014621 | 0.04823745 | 2.217069 | 0.001086911 | 2.398284 | 0.9510409 |
| ENSG00000167685.14 | ZNF444 | -2.009151 | 0.6214382 | -0.82984 | 0.842947 | -1.170817 | 0.03228664 | 0.883997 | 0.4839588 | 1.538118 | 0.1236708 | 0.04408 | 0.9467893 |
| ENSG00000167693.16 | NXN | -2.664488 | 0.6214382 | -0.684642 | 0.5923406 | -0.176495 | 0.6641614 | 2.895192 | 0.0100786 | 1.286262 | 0.1778915 | -0.015758 | 1 |
| ENSG00000167703.14 | SLC43A2 | 0.359279 | 0.8867323 | 0.584012 | 0.9222921 | -0.102718 | 0.3590797 | -0.912314 | 0.7287133 | 1.55088 | 0.03465387 | 0.075065 | 0.8247617 |
| ENSG00000167721.10 | TSR1 | -0.126248 | 0.6214382 | 0.234136 | 0.8657239 | 0.614466 | 0.02654938 | 1.52572 | 0.05976993 | 0.856752 | 0.6372012 | 0.21742 | 0.752198 |
| ENSG00000167771.5 | RCOR2 | -3.009896 | 0.6214382 | 1.122679 | 0.6527736 | -0.307294 | 0.6624105 | 1.469087 | 4.05E-15 | 1.481076 | 1.01E-07 | -2.131634 | 0.6885065 |
| ENSG00000167772.11 | ANGPTL4 | 2.076886 | 0.6214382 | 0.590108 | 4.49E-05 | -1.16878 | 0.01143551 | 1.336153 | 0.8600661 | 1.574863 | 0.2206737 | 0.779603 | 0.501643 |
| ENSG00000167775.10 | CD320 | -0.254832 | 0.8867323 | -0.276465 | 0.6170592 | -0.500917 | 0.03837948 | 0.279594 | 0.9450585 | -0.287984 | 0.2501933 | -0.380163 | 0.5585965 |
| ENSG00000167778.8 | SPRYD3 | 1.446213 | 0.04901151 | 0.293909 | 1 | -0.454737 | 0.5105902 | 0.524392 | 0.9903542 | -2.253497 | 0.6670798 | -1.503571 | 0.5841302 |
| ENSG00000167785.8 | ZNF558 | -0.479718 | 0.9183025 | 0.275637 | 0.8703522 | 1.161591 | 0.4064599 | 1.214858 | 0.0978339 | 2.570259 | 0.01238434 | 2.869674 | 0.6185816 |
| ENSG00000167792.11 | NDUFV1 | -0.216304 | 1 | 0.190815 | 0.9596929 | -0.139727 | 0.06905414 | 0.062323 | 0.3122989 | -0.069136 | 0.03407544 | -1.14678 | 0.6829816 |
| ENSG00000167797.7 | CDK2AP2 | -2.86891 | 0.09749406 | 0.634288 | 0.6527736 | 0.063261 | 0.7887553 | -0.344542 | 0.02749053 | -0.564305 | 0.002998326 | 0.816099 | 0.6885065 |
| ENSG00000167815.11 | PRDX2 | 0.304873 | 0.4406957 | 0.359543 | 0.08794836 | -0.458293 | 1.90E-05 | 0.334947 | 0.3467508 | -0.02153 | 0.005775222 | -0.037643 | 0.3820098 |
| ENSG00000167862.9 | ICT1 | -0.793953 | 0.6214382 | -0.679279 | 0.5649416 | -0.371683 | 0.01878676 | 0.025071 | 0.6788669 | 1.222369 | 0.9461086 | -2.754802 | 0.5653062 |
| ENSG00000167863.11 | ATP5H | -0.986893 | 0.2490442 | -0.064395 | 0.3285933 | -0.251308 | 5.22E-05 | 0.220268 | 0.3762574 | -0.50897 | 0.00042294 | -0.034921 | 0.6885065 |
| ENSG00000167962.12 | ZNF598 | 0.205026 | 0.8006286 | -0.096143 | 1 | 1.033292 | 0.6054545 | 0.080459 | 0.665837 | 1.368574 | 0.0141271 | -1.041312 | 0.8479431 |
| ENSG00000167969.12 | ECI1 | -0.623453 | 0.6014343 | 0.418629 | 0.8653838 | -1.687914 | 4.79E-12 | 0.503237 | 0.3023295 | -0.092803 | 0.6517097 | -0.739708 | 0.7973567 |
| ENSG00000167971.15 | CASKIN1 | 0.451139 | 0.9047689 | -0.374733 | 0.9689271 | -0.556368 | 0.5290845 | 1.998397 | 0.04210134 | 3.563206 | 0.005660647 | 3.271202 | 0.5733051 |
| ENSG00000167977.8 | KCTD5 | -0.904671 | 0.6214382 | -0.432593 | 0.6327597 | 0.434074 | 0.402077 | -0.137447 | 0.4354743 | 1.572794 | 0.02209267 | -1.199502 | 1 |
| ENSG00000167978.16 | SRRM2 | -1.970921 | 5.33E-07 | -0.297687 | 0.4334445 | 0.321283 | 0.3695136 | 0.52455 | 0.140091 | 1.234921 | 0.000173452 | 0.309228 | 0.9270789 |
| ENSG00000167995.15 | BEST1 | -1.187099 | 0.6214382 | 1.721706 | 1.87E-06 | 0.432076 | 0.7391321 | 2.194395 | 0.000316394 | 0.344221 | 0.2364543 | 0.598193 | 0.09230697 |
| ENSG00000167996.15 | FTH1 | -0.132661 | 0.8621764 | 0.51823 | 1.07E-41 | -0.515677 | 2.11E-28 | -0.600869 | 4.16E-26 | -0.420773 | 4.69E-05 | 0.851502 | 1.99E-09 |
| ENSG00000168002.11 | POLR2G | -0.35307 | 0.644046 | 0.115835 | 0.8089431 | -0.096626 | 0.1266825 | -0.069635 | 0.02166535 | 0.111759 | 0.1145696 | -0.360395 | 0.8915926 |
| ENSG00000168003.16 | SLC3A2 | 0.158901 | 0.4406957 | 0.515521 | 3.02E-05 | 0.073105 | 0.5290845 | 0.053549 | 0.1588082 | 0.306046 | 0.7491724 | 0.262389 | 0.4306015 |
| ENSG00000168005.8 | C11orf84 | 1.249727 | 1 | 0.020537 | 0.604359 | 0.929787 | 0.7898157 | 2.334316 | 0.07128885 | 3.034356 | 0.002621255 | -1.040003 | 0.9147227 |
| ENSG00000168028.13 | RPSA | -1.612876 | 5.42E-05 | 0.165406 | 0.4691942 | 0.359671 | 0.000313507 | -0.198381 | 0.02591905 | -0.041423 | 0.05157802 | -0.072069 | 0.05824041 |
| ENSG00000168036.16 | CTNNB1 | 0.850872 | 0.01638188 | -0.217185 | 0.3013217 | 0.137765 | 0.4064599 | 0.384536 | 0.2316384 | 0.754778 | 0.2714492 | 0.130232 | 0.594353 |
| ENSG00000168040.4 | FADD | 1.10924 | 0.6014343 | 0.117584 | 0.9653874 | -1.068111 | 0.005172759 | 0.738972 | 0.8622871 | 0.916716 | 0.7406781 | -1.99764 | 0.6185816 |
| ENSG00000168056.14 | LTBP3 | -2.144281 | 0.6214382 | 0.029564 | 0.8471755 | -2.838879 | 0.003962782 | 0.357895 | 0.730292 | 0.235757 | 0.3974318 | -0.361297 | 1 |
| ENSG00000168066.20 | SF1 | 0.069511 | 0.1233388 | -0.249893 | 0.2388066 | 0.195547 | 0.8174109 | 0.826506 | 0.005887308 | 0.703836 | 0.4718093 | 0.493971 | 0.6885065 |
| ENSG00000168092.13 | PAFAH1B2 | 0.915804 | 0.3098608 | -0.537043 | 0.5953732 | 0.619016 | 0.000766168 | 1.212906 | 0.01381081 | 1.414998 | 0.2789974 | -0.957273 | 0.6185816 |
| ENSG00000168096.14 | ANKS3 | 0.949964 | 0.8444846 | -0.192487 | 0.9403621 | -1.185761 | 0.03089392 | 0.964969 | 0.1880603 | 0.915776 | 0.8489112 | -0.793506 | 0.9147227 |
| ENSG00000168116.13 | KIAA1586 | -0.089727 | 0.9729966 | -0.655877 | 0.6527736 | 0.53626 | 0.01162846 | 2.912408 | 0.2242006 | -0.151762 | 0.8387831 | -1.019446 | 0.9264343 |
| ENSG00000168137.15 | SETD5 | -0.320294 | 0.3946735 | 0.013659 | 0.8872857 | 0.502539 | 0.01116236 | 1.646524 | 1.73E-07 | 0.814318 | 0.01124704 | 0.335715 | 0.6885065 |
| ENSG00000168172.8 | HOOK3 | -0.863128 | 0.529676 | -0.303891 | 0.6433605 | 0.92096 | 1.84E-05 | 1.718794 | 0.008823624 | 0.888316 | 0.3977519 | 2.034088 | 0.5344421 |
| ENSG00000168214.20 | RBPJ | 0.063664 | 0.6980258 | 0.230607 | 0.9138985 | 1.283268 | 1.44E-08 | 0.462612 | 0.8837017 | 0.154881 | 0.7087352 | -0.25484 | 0.9704304 |
| ENSG00000168243.10 | GNG4 | 0.048121 | 0.59287 | 0.281796 | 1 | -0.229103 | 0.1663608 | 0.852092 | 0.02741529 | 0.830828 | 0.3974318 | -1.348715 | 0.4578379 |
| ENSG00000168246.5 | UBTD2 | -2.586856 | 0.4692975 | -0.696663 | 0.9398726 | 0.948742 | 0.8240808 | 2.628281 | 8.48E-05 | 1.963801 | 0.001346348 | -2.471805 | 0.7817802 |
| ENSG00000168255.19 | POLR2J3 | -0.591777 | 0.3526668 | 1.147946 | 0.2779213 | 0.213899 | 0.4192542 | 1.330369 | 0.000582515 | 1.433263 | 0.000153027 | -1.271092 | 0.8070118 |
| ENSG00000168259.14 | DNAJC7 | -0.023551 | 0.8765042 | 0.01507 | 0.7711097 | -0.076458 | 0.7119994 | 1.097147 | 0.000870034 | 0.84414 | 0.9178187 | 0.242444 | 0.6185816 |
| ENSG00000168264.10 | IRF2BP2 | 0.380924 | 0.6523873 | -0.049721 | 0.9036408 | -0.039293 | 0.8610371 | 1.434028 | 0.000851278 | 0.849623 | 0.1197221 | -0.227419 | 1 |
| ENSG00000168268.10 | NT5DC2 | 0.566507 | 0.824966 | -0.00592 | 0.5237469 | -0.219174 | 0.4664068 | 0.523249 | 1.92E-06 | 1.339196 | 0.000163167 | -0.277487 | 0.4168869 |
| ENSG00000168273.7 | SMIM4 | -0.134159 | 0.6708674 | 0.140021 | 0.4440278 | -0.938819 | 5.20E-05 | 0.049974 | 0.1052874 | 0.776755 | 0.5196843 | 0.540668 | 0.6185816 |
| ENSG00000168275.14 | COA6 | 0.327728 | 0.8140543 | -0.010091 | 0.8263112 | -0.223416 | 0.07028776 | 0.73582 | 0.3112291 | 0.002937 | 0.02599105 | 0.356187 | 0.6185816 |
| ENSG00000168280.16 | KIF5C | 1.024041 | 0.06126222 | -0.182698 | 0.7560798 | 0.392691 | 0.3801085 | 1.486682 | 4.23E-15 | 0.762445 | 0.03056663 | 0.275988 | 0.8949489 |
| ENSG00000168283.13 | BMI1 | 1.528062 | 0.4536837 | -0.215362 | 0.8263112 | 1.012886 | 0.1352928 | 2.46905 | 3.51E-05 | 2.050174 | 0.003872935 | 0.439708 | 0.6885065 |
| ENSG00000168298.6 | HIST1H1E | -0.868225 | 0.714717 | -1.350462 | 0.7819293 | 1.955354 | 0.000604182 | 0.141229 | 0.5449772 | 3.059099 | 0.5734675 | inf | 0.6185816 |
| ENSG00000168310.10 | IRF2 | 0.835339 | 1 | -0.593996 | 0.6527736 | 0.654576 | 0.4003628 | -0.007846 | 0.03626243 | 0.077168 | 0.544686 | 2.465995 | 0.4578379 |
| ENSG00000168374.10 | ARF4 | 0.275491 | 0.8456603 | -0.040699 | 0.9752243 | 0.362798 | 0.01366175 | -0.188224 | 0.000366314 | 0.230108 | 0.404173 | 0.785414 | 0.6885065 |
| ENSG00000168385.17 | 2-Sep | 0.35098 | 0.3982827 | -0.277838 | 0.04079597 | 0.378807 | 0.009631666 | 0.554898 | 0.148124 | 0.621049 | 0.634191 | -0.075004 | 0.6885065 |
| ENSG00000168393.12 | DTYMK | 0.294507 | 0.8639532 | -0.262942 | 0.6053838 | -0.523971 | 0.000163622 | -0.220104 | 0.1317969 | 0.060496 | 0.2215715 | 1.189383 | 0.6885065 |
| ENSG00000168394.10 | TAP1 | 0.977713 | 0.9510349 | -0.051872 | 0.8887513 | -0.778498 | 0.004077784 | 1.011674 | 0.861719 | 0.292705 | 0.46228 | 0.328426 | 1 |
| ENSG00000168411.13 | RFWD3 | 1.289896 | 0.714717 | -0.275049 | 0.2913444 | -0.07444 | 0.809226 | 0.78207 | 0.002892901 | 1.155989 | 0.3974318 | 1.0995 | 0.8614062 |
| ENSG00000168438.14 | CDC40 | -0.622407 | 0.6573292 | 0.380908 | 0.8346083 | 0.245487 | 0.0136474 | -0.267994 | 0.8987922 | 0.167757 | 0.6427258 | 0.934123 | 0.6885065 |
| ENSG00000168461.12 | RAB31 | 0.020834 | 0.9047689 | 0.065462 | 0.85618 | 0.720565 | 0.002954378 | 0.546148 | 0.833314 | 0.517816 | 0.7528548 | 0.072645 | 1 |
| ENSG00000168488.18 | ATXN2L | 0.237309 | 0.6214382 | 0.341634 | 0.8263112 | 0.453027 | 0.6527122 | 1.09723 | 0.02639972 | 1.095375 | 0.3974318 | 0.056866 | 1 |
| ENSG00000168528.11 | SERINC2 | -1.224554 | 0.8587992 | 0.072253 | 0.7763773 | -1.703515 | 0.01196514 | 0.617044 | 0.4919368 | -0.207402 | 0.7404912 | 1.197195 | 1 |
| ENSG00000168610.14 | STAT3 | -0.044167 | 0.7598375 | -0.138179 | 0.6527736 | 0.698702 | 0.8100781 | 0.527529 | 0.009064554 | -0.185996 | 0.5196843 | 0.212303 | 0.8009135 |
| ENSG00000168615.11 | ADAM9 | -0.811843 | 0.8242019 | -0.314232 | 0.4219211 | -0.098449 | 0.9368403 | 0.384617 | 0.03998217 | 2.389211 | 0.6684212 | 0.767234 | 1 |
| ENSG00000168646.12 | AXIN2 | -2.038798 | 0.6214382 | -0.458748 | 0.6527736 | 0.165237 | 0.3144442 | 1.647804 | 0.1242662 | 4.296591 | 0.0426606 | -1.847173 | 0.8479431 |
| ENSG00000168653.10 | NDUFS5 | -0.935626 | 0.6214382 | -0.125206 | 0.8649079 | -0.381506 | 3.44E-05 | 0.141074 | 0.01132536 | -0.565664 | 3.81E-06 | 0.445826 | 0.6829816 |
| ENSG00000168675.18 | LDLRAD4 | -2.117903 | 0.6014343 | -0.056125 | 1 | 2.127737 | 0.6894429 | 2.249824 | 0.4919368 | 1.699735 | 0.01998467 | 0.715735 | 0.5733051 |
| ENSG00000168701.18 | TMEM208 | 0.29947 | 0.9655291 | 0.297332 | 0.6527736 | -0.137921 | 0.03187308 | -0.445143 | 0.01011425 | -0.719682 | 0.01238434 | 0.534187 | 0.9175671 |
| ENSG00000168702.16 | LRP1B | -1.467401 | 0.8006286 | -0.719052 | 0.6527736 | 2.366243 | 0.03176686 | 1.382935 | 0.4051678 | -2.934174 | 0.5196843 | 2.640469 | 0.6885065 |
| ENSG00000168710.17 | AHCYL1 | -0.886041 | 0.6214382 | -0.487091 | 0.2713328 | 0.579446 | 0.19147 | 1.482566 | 0.1208655 | 1.983302 | 0.007823683 | 0.456062 | 0.9532204 |
| ENSG00000168734.13 | PKIG | 0.149928 | 1 | 0.086694 | 0.9353253 | -1.103108 | 0.03228664 | -0.377043 | 0.02029956 | -0.341519 | 0.4440658 | -2.172594 | 0.3565625 |
| ENSG00000168763.15 | CNNM3 | 0.972899 | 1 | 0.061898 | 0.863029 | -0.143522 | 0.6278045 | 3.319685 | 0.02710839 | 2.622972 | 0.2363864 | -2.613912 | 1 |
| ENSG00000168769.12 | TET2 | -0.358603 | 0.8006286 | -0.522947 | 0.6527736 | 0.976535 | 0.05530681 | 1.706728 | 0.023702 | 1.874865 | 0.01782745 | -0.658507 | 0.8479431 |
| ENSG00000168772.10 | CXXC4 | -1.123932 | 0.646639 | 2.813926 | 0.4487482 | 1.500062 | 0.1992941 | 1.669667 | 0.007453854 | 1.318923 | 0.006147839 | -1.165751 | 0.9452148 |
| ENSG00000168802.12 | CHTF8 | 0.694053 | 0.9047689 | 0.063568 | 0.5953732 | -0.889678 | 0.003164196 | 1.984847 | 0.01718177 | 0.475553 | 0.7377262 | 0.966417 | 0.6490419 |
| ENSG00000168806.7 | LCMT2 | -0.536715 | 1 | -0.10071 | 0.8478107 | 0.89782 | 1 | 3.022841 | 0.03932487 | -0.068181 | 0.9263124 | 2.397102 | 0.8479431 |
| ENSG00000168807.16 | SNTB2 | -3.777426 | 0.3098608 | 0.016532 | 0.8217443 | -0.206084 | 0.7437308 | 1.415128 | 0.001085659 | 0.002647 | 0.1401461 | 2.138437 | 0.8876753 |
| ENSG00000168818.9 | STX18 | -4.132093 | 0.04197057 | -0.77454 | 0.3919863 | 0.779876 | 0.5396515 | 0.664471 | 0.3361528 | -0.602421 | 0.5597051 | -0.4343 | 0.6885065 |
| ENSG00000168874.12 | ATOH8 | 0.710234 | 0.9047689 | -1.791753 | 0.3903174 | -1.483469 | 0.1824711 | 1.985848 | 0.007660097 | 2.052209 | 0.8021966 | 3.640734 | 0.6504862 |
| ENSG00000168876.8 | ANKRD49 | 1.18206 | 0.6792949 | 1.041891 | 1 | 0.156263 | 0.9409591 | 2.121476 | 0.01008289 | 1.611792 | 0.3474398 | -0.63179 | 0.6885065 |
| ENSG00000168887.10 | C2orf68 | 1.314136 | 0.9282145 | -0.096079 | 0.9278825 | 0.960899 | 0.5290845 | 2.148293 | 0.04547192 | 1.074127 | 0.4042092 | -4.123495 | 0.3800088 |
| ENSG00000168894.9 | RNF181 | 0.127984 | 0.954995 | 0.132246 | 0.9775912 | -0.801852 | 0.000119097 | -0.325404 | 0.001539144 | 0.230574 | 0.5196843 | -0.038693 | 0.6185816 |
| ENSG00000168899.4 | VAMP5 | 1.311596 | 0.7953772 | -0.113382 | 0.6527736 | -0.887659 | 0.000341591 | -0.442924 | 0.005079915 | 0.302709 | 0.2664633 | -0.251267 | 0.6829816 |
| ENSG00000168906.12 | MAT2A | -1.000851 | 0.4348699 | 0.062479 | 0.9330607 | 0.48304 | 0.9051127 | 0.842163 | 0.003665272 | 1.795342 | 6.14E-06 | 0.726924 | 0.9278206 |
| ENSG00000168916.15 | ZNF608 | -0.136089 | 0.9351938 | -0.563302 | 0.5722619 | 0.097274 | 0.05710177 | 1.867329 | 0.001300776 | 0.352748 | 0.8383298 | -1.12405 | 0.6185816 |
| ENSG00000168936.10 | TMEM129 | 0.06448 | 1 | 0.384286 | 0.9878938 | -0.777574 | 0.5287801 | 0.284506 | 0.3760083 | 1.179043 | 0.04698277 | -0.607041 | 0.6885065 |
| ENSG00000168993.14 | CPLX1 | 0.615653 | 0.6214382 | -0.787831 | 0.897729 | 2.892274 | 0.4382561 | 2.78669 | 0.01958024 | 2.047153 | 0.3390021 | -1.984809 | 0.9147227 |
| ENSG00000169018.5 | FEM1B | -0.611185 | 0.9183025 | 0.191698 | 0.5394427 | 0.502082 | 0.05928462 | 1.057753 | 0.001579715 | 0.949489 | 0.3974318 | -1.070438 | 0.6885065 |
| ENSG00000169020.9 | ATP5I | 0.648877 | 0.000710361 | 0.155829 | 0.7139087 | -0.575154 | 4.58E-17 | 0.077956 | 0.120765 | -0.31494 | 1.29E-05 | -0.187624 | 0.05261484 |
| ENSG00000169021.5 | UQCRFS1 | -0.12169 | 0.764956 | -0.033382 | 0.7015905 | -0.592045 | 1.41E-10 | -0.43857 | 0.01500599 | 0.252464 | 0.2364542 | -0.618552 | 0.9147227 |
| ENSG00000169045.17 | HNRNPH1 | 0.000501 | 0.6214382 | -0.021244 | 0.8381834 | 0.051253 | 0.331527 | 0.391151 | 4.96E-05 | 0.481819 | 0.02116972 | -0.359248 | 0.215927 |
| ENSG00000169062.14 | UPF3A | -0.470285 | 0.1524934 | 0.182981 | 0.9214744 | 0.521911 | 0.1138222 | 0.135318 | 0.7821218 | -0.697918 | 0.00951511 | 2.334533 | 0.9687888 |
| ENSG00000169085.11 | C8orf46 | -0.046997 | 0.1065073 | -0.460036 | 0.8483182 | 1.476374 | 0.04093178 | 0.175048 | 0.9029261 | 1.128282 | 0.5196843 | -1.617326 | 1 |
| ENSG00000169118.16 | CSNK1G1 | -1.642662 | 0.529676 | -0.545435 | 0.6045862 | 0.687825 | 0.5016363 | 1.788943 | 0.006512046 | 1.370375 | 0.8056086 | -0.524245 | 0.6885065 |
| ENSG00000169131.10 | ZNF354A | -0.575964 | 1 | -0.480056 | 0.8627375 | 0.118184 | 0.5567417 | 0.898275 | 0.008322128 | 0.303547 | 0.5196843 | -1.92314 | 0.9483075 |
| ENSG00000169139.11 | UBE2V2 | -0.309693 | 0.6562396 | 0.12631 | 0.9104976 | 0.143698 | 0.9368403 | 0.005629 | 0.3467508 | -0.199408 | 0.008448242 | 0.033482 | 0.6463174 |
| ENSG00000169155.9 | ZBTB43 | 0.682043 | 0.8006286 | -0.196394 | 0.7401221 | 1.544819 | 2.19E-05 | 1.815362 | 0.000595202 | 0.05466 | 1 | -1.616885 | 0.6885065 |
| ENSG00000169169.14 | CPT1C | 0.587449 | 0.7736255 | -0.361005 | 0.3121527 | 0.159089 | 0.7437308 | 1.210693 | 0.01413636 | 1.252244 | 0.5196843 | -0.686537 | 0.6885065 |
| ENSG00000169189.16 | NSMCE1 | -0.585478 | 0.8006286 | -0.297624 | 0.4330346 | -0.155528 | 0.8221252 | -0.808117 | 0.002972112 | -0.133795 | 0.02955035 | -3.537616 | 0.6153291 |
| ENSG00000169223.14 | LMAN2 | 0.916325 | 0.8587992 | -0.037266 | 0.6792212 | -0.188688 | 0.004020273 | 0.497399 | 0.3380727 | 0.594015 | 0.544002 | -0.555564 | 0.6885065 |
| ENSG00000169228.13 | RAB24 | 0.927583 | 0.8006286 | -0.16524 | 1 | -1.501291 | 0.02501833 | -1.180293 | 0.5243294 | 0.172515 | 0.7672587 | -1.517913 | 0.8479431 |
| ENSG00000169230.9 | PRELID1 | 0.107483 | 0.2644795 | 0.225227 | 0.872021 | -1.831776 | 1.01E-50 | 0.081057 | 0.6745348 | -0.178155 | 0.7574155 | -0.131742 | 0.3358843 |
| ENSG00000169255.13 | B3GALNT1 | -0.208758 | 0.8474876 | -0.194733 | 0.3725835 | 1.134453 | 0.5383177 | 1.36384 | 0.05458417 | 1.692702 | 0.01845345 | -0.23169 | 0.6185816 |
| ENSG00000169258.6 | GPRIN1 | -0.20652 | 0.8006286 | 0.908524 | 0.7949423 | -1.669213 | 0.02505728 | 2.224585 | 0.04666571 | 1.276009 | 0.01185816 | 0.994495 | 0.6885065 |
| ENSG00000169302.14 | STK32A | -0.885783 | 0.3439898 | -0.317765 | 0.26262 | 0.106181 | 0.2744947 | 1.572487 | 6.78E-05 | 1.502864 | 0.06986248 | 1.408785 | 0.7019165 |
| ENSG00000169330.8 | KIAA1024 | 2.172349 | 0.1366571 | -0.804047 | 0.8953649 | -0.82509 | 0.4132012 | -1.98369 | 0.6153993 | 1.679499 | 0.5746463 |  |  |
| ENSG00000169359.13 | SLC33A1 | -0.374834 | 0.6214382 | 0.150838 | 0.8263112 | 0.188397 | 0.5290845 | 1.208168 | 0.04545716 | -0.325778 | 0.5196843 | -1.543186 | 0.5733051 |
| ENSG00000169379.15 | ARL13B | -0.652855 | 0.7365772 | 0.472778 | 0.9795629 | 0.873788 | 0.003456105 | -0.13638 | 0.7328179 | -0.961005 | 0.4246135 | -3.460943 | 0.6885065 |
| ENSG00000169429.10 | CXCL8 | -2.04717 | 0.6131246 | 1.602683 | 4.92E-13 | 2.222084 | 0.7354314 | 0.821099 | 0.6996942 | -0.870959 | 0.7377262 | 1.687441 | 0.8949489 |
| ENSG00000169432.14 | SCN9A | 3.030422 | 0.6214382 | -0.732431 | 0.8703522 | 0.268464 | 0.2879524 | 2.378817 | 0.01173764 | 3.88529 | 0.2947291 | -0.108211 | 0.9270789 |
| ENSG00000169504.14 | CLIC4 | -0.343546 | 0.714717 | -0.018427 | 0.7568131 | 0.980035 | 0.006241239 | 0.59683 | 0.6227788 | 1.005209 | 0.6357065 | 0.803357 | 0.5716672 |
| ENSG00000169554.16 | ZEB2 | 0.186345 | 0.9734607 | 0.115331 | 0.7497389 | 0.560822 | 7.75E-05 | 1.289436 | 2.92E-05 | 0.103641 | 0.5196843 | 1.339341 | 0.8999026 |
| ENSG00000169567.11 | HINT1 | 0.23471 | 0.01541151 | 0.106043 | 0.9247252 | -0.081951 | 0.00241711 | -0.046572 | 0.005925879 | -0.157834 | 0.003684308 | -0.172339 | 0.3476515 |
| ENSG00000169592.14 | INO80E | -0.813413 | 0.8272675 | -0.153017 | 0.9081785 | -0.792955 | 0.001804798 | -0.289106 | 0.6180045 | 1.083016 | 0.6217705 | -0.609157 | 0.6829816 |
| ENSG00000169599.12 | NFU1 | -0.396341 | 0.928247 | -0.205368 | 0.5851668 | -0.062883 | 0.02821202 | -0.681695 | 0.4919368 | -0.255752 | 0.0933406 | -1.194814 | 0.8723509 |
| ENSG00000169612.3 | FAM103A1 | -1.663185 | 0.3975154 | 0.386445 | 0.8703522 | 0.259263 | 0.5290845 | 0.215886 | 0.448147 | -0.272616 | 0.03283007 | 1.685279 | 0.8479431 |
| ENSG00000169627.7 | BOLA2B | 0.095283 | 0.6214382 | -0.283928 | 0.6200225 | -0.364167 | 2.59E-08 | 0.075211 | 0.5899001 | -0.123031 | 0.007470624 | -0.650569 | 0.05357521 |
| ENSG00000169635.9 | HIC2 | 2.681989 | 0.6214382 | -0.117287 | 0.9452791 | 0.644255 | 0.1336022 | 1.258967 | 0.1292522 | 2.183793 | 0.04222895 | -1.168115 | 0.8479431 |
| ENSG00000169641.13 | LUZP1 | 0.369294 | 0.5468595 | -0.230277 | 0.2521066 | 0.690238 | 0.02142615 | 1.41705 | 0.1420578 | 1.143277 | 0.6014201 | 0.784692 | 0.5344421 |
| ENSG00000169689.14 | STRA13 | -0.700172 | 0.4406957 | 0.198854 | 0.8263112 | -0.096253 | 0.5290845 | -0.620319 | 0.002688417 | -0.453411 | 0.03125981 | 0.528082 | 0.6885065 |
| ENSG00000169710.7 | FASN | 0.809159 | 0.05018941 | -0.232593 | 0.7398503 | -0.902793 | 0.06330527 | 0.533201 | 0.02029956 | 0.769548 | 0.08617161 | -0.237063 | 0.6885065 |
| ENSG00000169714.16 | CNBP | -0.158523 | 0.9014318 | -0.032328 | 0.7585627 | 0.287323 | 0.006296326 | 0.346141 | 0.5193214 | 0.264621 | 0.3450415 | 0.462975 | 0.5733051 |
| ENSG00000169718.17 | DUS1L | 0.51623 | 0.8587992 | 0.411067 | 0.9005514 | -0.669027 | 0.06341897 | 2.306133 | 0.003447053 | 1.325513 | 0.2721917 | 0.269911 | 0.8579425 |
| ENSG00000169727.12 | GPS1 | 0.96303 | 0.5711449 | 0.489325 | 0.8263112 | 0.363819 | 0.8171765 | 2.07041 | 0.03632936 | 0.577719 | 0.9818344 | 0.412347 | 0.6885065 |
| ENSG00000169738.7 | DCXR | 0.047862 | 0.3529259 | 0.071781 | 0.7789922 | -0.496562 | 0.01296452 | 0.577261 | 0.2335648 | -0.36491 | 0.5213414 | 1.597768 | 0.8479431 |
| ENSG00000169750.8 | RAC3 | 0.727378 | 0.4974608 | 0.076204 | 0.8703522 | -1.209455 | 5.20E-05 | 0.898996 | 0.099043 | 0.797262 | 0.2880317 | -0.285897 | 0.06861843 |
| ENSG00000169760.17 | NLGN1 | -0.591044 | 0.7722327 | 0.749005 | 0.6895794 | 1.74841 | 0.006188158 | 0.848218 | 0.000124633 | 1.241014 | 0.02525373 | 1.539886 | 0.9264343 |
| ENSG00000169783.12 | LINGO1 | -1.387184 | 0.4605085 | -0.896214 | 0.3121527 | 0.165505 | 0.9807875 | 0.470049 | 0.1729147 | 1.185292 | 0.04652304 | 0.969533 | 0.7733704 |
| ENSG00000169855.19 | ROBO1 | -0.759359 | 0.6014343 | -0.241412 | 0.8263112 | 0.752646 | 0.8592813 | 1.197272 | 5.25E-08 | 0.274143 | 0.9216125 | -1.075324 | 1 |
| ENSG00000169862.18 | CTNND2 | -0.546953 | 0.3286597 | -0.326526 | 0.0410984 | 0.978558 | 0.000190296 | 0.419689 | 0.03776643 | 0.380106 | 1 | -0.710436 | 0.6885065 |
| ENSG00000169871.12 | TRIM56 | -3.438526 | 0.01706913 | -0.256841 | 0.9574619 | -0.030966 | 0.8457938 | 1.841555 | 0.00033715 | 0.94759 | 0.8667133 | 1.191426 | 0.6767422 |
| ENSG00000169895.5 | SYAP1 | -0.133152 | 0.3724613 | 0.010823 | 0.9403483 | 0.422479 | 0.1625971 | 0.182006 | 0.7238932 | -0.699821 | 0.03470409 | 2.069621 | 0.6885065 |
| ENSG00000169905.12 | TOR1AIP2 | 1.203031 | 0.529676 | 0.276782 | 0.6527736 | 0.716225 | 0.04280891 | 0.67224 | 0.4919368 | 0.3408 | 0.3280237 | 0.10135 | 0.8479431 |
| ENSG00000169908.10 | TM4SF1 | 0.079424 | 0.7727139 | -0.077165 | 0.3027683 | -0.297976 | 0.8148288 | 0.568437 | 0.6328348 | 0.06451 | 1.89E-05 | 0.081954 | 0.6185816 |
| ENSG00000169925.16 | BRD3 | -0.151989 | 0.6214382 | -0.222339 | 0.7765491 | 0.143563 | 0.2808088 | 1.063639 | 0.02577896 | 0.492253 | 1 | 0.453224 | 1 |
| ENSG00000169957.9 | ZNF768 | -1.425561 | 0.6214382 | -0.917151 | 0.5394427 | -0.673603 | 0.02506782 | -0.088358 | 0.2436291 | 1.040743 | 0.1169767 | -0.303246 | 0.6829816 |
| ENSG00000169967.16 | MAP3K2 | 0.953571 | 0.5711449 | 0.316227 | 0.7477293 | 0.768781 | 0.0458126 | 0.996959 | 6.26E-06 | 1.638943 | 0.003778489 | 0.079494 | 0.6885065 |
| ENSG00000169976.6 | SF3B5 | 0.063504 | 0.4406957 | 0.167466 | 0.8657239 | -0.387199 | 9.12E-07 | 0.513126 | 0.06674489 | 0.266513 | 0.3886833 | 0.010765 | 0.6885065 |
| ENSG00000169992.9 | NLGN2 | 0.340708 | 1 | 0.007955 | 0.9574152 | -0.035398 | 0.8925008 | 0.96621 | 0.4225718 | 2.337972 | 0.002918613 | -1.145552 | 0.7901525 |
| ENSG00000170004.16 | CHD3 | -1.048995 | 0.3009675 | 0.144602 | 0.6527736 | 1.013772 | 0.7245422 | 1.083981 | 0.004181679 | 1.369877 | 0.05634492 | -0.376247 | 0.6185816 |
| ENSG00000170017.12 | ALCAM | 0.863815 | 0.2387521 | 0.234849 | 0.7057076 | 1.026459 | 0.01852609 | 0.242719 | 0.1685514 | 0.758382 | 0.07671263 | 1.331279 | 0.9517616 |
| ENSG00000170027.6 | YWHAG | -0.113975 | 1 | -0.121265 | 0.6227004 | 0.23039 | 0.5740276 | 1.01919 | 1.43E-12 | 0.920664 | 0.000159403 | 0.050226 | 0.6885065 |
| ENSG00000170043.11 | TRAPPC1 | -0.870594 | 0.6014343 | 0.007589 | 0.7321214 | -0.110132 | 0.002422785 | 0.231496 | 0.1860894 | -0.818718 | 0.0141271 | 0.061719 | 0.5344421 |
| ENSG00000170100.13 | ZNF778 | 0.444947 | 0.8867323 | -0.460063 | 0.6435681 | -1.53641 | 0.9648047 | 1.28545 | 0.03359915 | 0.492168 | 0.6770529 | -1.377448 | 1 |
| ENSG00000170113.15 | NIPA1 | -2.854748 | 0.6168416 | 0.761088 | 0.9832381 | 0.975818 | 0.5507355 | 1.718932 | 0.01580491 | 1.437644 | 0.1228802 | -3.258757 | 0.8479431 |
| ENSG00000170144.18 | HNRNPA3 | -0.654471 | 0.5519048 | -0.418604 | 3.01E-05 | 0.27305 | 0.08139729 | 0.493891 | 0.03914854 | 0.237694 | 0.3974318 | -0.234201 | 0.2669075 |
| ENSG00000170145.4 | SIK2 | -2.457158 | 0.6214382 | -0.737877 | 0.5953732 | -0.028423 | 0.8616087 | 1.095176 | 0.03982882 | -0.635047 | 0.6196247 | -0.999505 | 0.9380942 |
| ENSG00000170153.10 | RNF150 | 1.66524 | 0.6214382 | 1.138512 | 0.2553839 | 3.320282 | 0.00047823 | 2.242312 | 0.03626243 | 0.510161 | 0.6009809 | 0.436258 | 1 |
| ENSG00000170234.12 | PWWP2A | -1.093223 | 0.7138283 | -0.36604 | 0.1928922 | 0.410088 | 0.7437308 | 2.035127 | 0.005135153 | 0.968211 | 0.2105971 | 1.733118 | 0.9228374 |
| ENSG00000170242.17 | USP47 | 0.39793 | 0.6214382 | -0.170906 | 0.04318493 | 0.816673 | 0.003399897 | 0.655889 | 0.000907911 | 0.993566 | 0.2985977 | 1.284116 | 0.6885065 |
| ENSG00000170265.11 | ZNF282 | 3.253339 | 0.8006286 | -0.800216 | 0.9193039 | 0.458003 | 0.8421791 | 2.789784 | 0.1385281 | 2.808767 | 0.02951102 | 4.438194 | 0.6885065 |
| ENSG00000170296.9 | GABARAP | 0.38038 | 0.7736255 | -0.930474 | 0.8599148 | 1.197819 | 0.5290845 | 0.333829 | 0.3926282 | 2.811982 | 0.01073185 | -3.623772 | 0.7670687 |
| ENSG00000170315.13 | UBB | 0.145987 | 0.1375427 | -0.144328 | 0.7262755 | 0.300848 | 0.003121318 | -0.279498 | 1.26E-08 | 0.201098 | 0.2538929 | -0.290444 | 0.06348693 |
| ENSG00000170345.9 | FOS | -0.274226 | 0.6924582 | 0.216064 | 2.43E-08 | 0.14286 | 0.151938 | -0.21321 | 0.1539482 | 0.592081 | 0.6145561 | 0.529677 | 0.3746505 |
| ENSG00000170385.9 | SLC30A1 | 0.252626 | 0.6014343 | 0.307041 | 0.6527736 | 2.35472 | 0.1100886 | 0.182088 | 0.701209 | 2.73708 | 0.03383784 | 0.312464 | 0.9127491 |
| ENSG00000170390.15 | DCLK2 | -1.136257 | 0.4579871 | -0.554728 | 0.007393953 | 0.076557 | 0.9481144 | 1.025587 | 0.08481253 | 1.02828 | 0.07366152 | 0.778422 | 0.8949489 |
| ENSG00000170439.6 | METTL7B | -1.042958 | 0.714717 | -0.771538 | 0.007452629 | -0.247779 | 0.4644258 | -0.801187 | 0.7803491 | -2.509274 | 0.5196843 | -0.395508 | 0.8663281 |
| ENSG00000170448.11 | NFXL1 | 0.680874 | 0.6214382 | -0.537998 | 0.8221408 | 0.55849 | 0.04996501 | 0.145925 | 0.7524047 | 1.52297 | 0.5196843 | 0.474609 | 0.9147227 |
| ENSG00000170456.14 | DENND5B | 0.360207 | 0.9529915 | -1.052549 | 0.485736 | 1.235492 | 0.001851701 | 2.296795 | 0.01671469 | 0.951734 | 0.034258 | -0.148946 | 0.6885065 |
| ENSG00000170464.9 | DNAJC18 | 0.131194 | 0.7896588 | -0.012173 | 0.6527736 | -0.053751 | 0.5290845 | -0.109225 | 1 | 1.586776 | 0.01013219 | 1.013164 | 0.6885065 |
| ENSG00000170471.14 | RALGAPB | 1.526523 | 0.6168416 | -0.88172 | 0.4106687 | -0.09027 | 0.5290845 | 1.642182 | 0.03168751 | 0.989532 | 0.05948264 | 2.541022 | 0.6885065 |
| ENSG00000170500.12 | LONRF2 | -3.81129 | 0.2335514 | -0.196013 | 0.6056121 | 1.482867 | 0.03054897 | 1.418028 | 0.04854501 | 1.022302 | 0.8726186 | -1.158651 | 1 |
| ENSG00000170515.13 | PA2G4 | -0.282367 | 0.9195601 | 0.202756 | 0.9831379 | 0.332316 | 0.4860816 | 0.214364 | 0.3517926 | 0.37431 | 0.01970273 | 0.195169 | 0.5653062 |
| ENSG00000170522.9 | ELOVL6 | -0.503094 | 0.7987522 | -0.699006 | 0.2021027 | 1.843716 | 2.01E-07 | 1.714888 | 0.000403778 | 0.379903 | 0.8387831 | -1.311382 | 0.9645832 |
| ENSG00000170540.14 | ARL6IP1 | -0.530068 | 0.563505 | -0.03492 | 0.7820217 | 0.458788 | 0.001762 | 0.771573 | 0.2335397 | 0.267468 | 0.7853444 | 0.634688 | 0.8868882 |
| ENSG00000170558.8 | CDH2 | -0.440944 | 0.7736255 | -0.230172 | 0.3045812 | 0.429754 | 0.04532125 | 0.726101 | 6.83E-05 | 0.712428 | 0.2011388 | -1.366313 | 0.2656042 |
| ENSG00000170606.13 | HSPA4 | 0.239663 | 0.8867323 | 0.1206 | 0.5943878 | 0.435302 | 0.0479416 | 0.532416 | 8.19E-05 | 0.861006 | 0.01472525 | -0.004616 | 0.5193439 |
| ENSG00000170632.13 | ARMC10 | -1.323058 | 0.2998756 | -0.237264 | 0.8518755 | 0.02978 | 0.5840658 | 1.113088 | 0.007539285 | 2.06656 | 0.04652304 | -0.603588 | 0.9417757 |
| ENSG00000170634.12 | ACYP2 | -1.013819 | 0.8408841 | 0.629262 | 0.5953732 | -0.238526 | 0.1562889 | -0.715526 | 0.001938814 | -1.096666 | 0.0928065 | 0.319842 | 0.7973567 |
| ENSG00000170667.14 | RASA4B | 0.609423 | 0.9360557 | 0.022087 | 0.7955884 | -0.935649 | 0.2983833 | 1.544648 | 0.01101017 | 1.204171 | 0.5578349 | 0.279119 | 0.7767294 |
| ENSG00000170677.5 | SOCS6 | -1.496478 | 0.6798783 | 0.114333 | 0.688619 | -0.08557 | 0.578623 | 3.169941 | 0.02741529 | 0.390882 | 0.3232893 | 1.4624 | 1 |
| ENSG00000170759.10 | KIF5B | 0.278189 | 0.08143052 | 0.224148 | 0.9668554 | 0.569656 | 1.88E-06 | 1.20658 | 3.24E-06 | -0.455711 | 0.3536738 | -0.150288 | 0.1117128 |
| ENSG00000170802.15 | FOXN2 | -1.503026 | 0.4579871 | -0.682446 | 0.6527736 | 0.761258 | 0.003033089 | 1.278621 | 0.0412511 | 2.418439 | 0.2880317 | -2.040797 | 1 |
| ENSG00000170836.11 | PPM1D | 2.194332 | 0.529676 | -0.334699 | 0.6527736 | 0.971566 | 0.5068969 | 1.980353 | 0.008024982 | 1.6053 | 0.3625413 | -0.467351 | 0.6885065 |
| ENSG00000170852.10 | KBTBD2 | -0.375614 | 0.6214382 | -0.402812 | 0.9452791 | 0.722606 | 0.2555203 | 2.099562 | 0.08574809 | 1.008327 | 0.002514978 | -1.297395 | 0.4998763 |
| ENSG00000170860.3 | LSM3 | 0.485671 | 0.5441969 | -0.107386 | 0.8263112 | -0.415947 | 0.000585923 | -0.51213 | 0.000139095 | 0.028884 | 0.02199816 | 0.209714 | 0.5344421 |
| ENSG00000170873.18 | MTSS1 | 1.314232 | 0.8587992 | -0.993373 | 0.6527736 | 1.345943 | 7.74E-05 | 1.379035 | 0.008905317 | 1.048555 | 0.02288874 | 0.660876 | 0.9733288 |
| ENSG00000170881.4 | RNF139 | 0.372817 | 0.8467545 | -0.560768 | 0.9439537 | 0.811561 | 0.5440704 | -0.834172 | 0.3620324 | 3.371754 | 0.002514978 | -4.137058 | 0.008590575 |
| ENSG00000170889.13 | RPS9 | -0.43742 | 0.4464112 | 0.016808 | 0.9329105 | -0.043521 | 0.1282668 | -0.163531 | 0.03432917 | 0.102445 | 0.2774095 | 0.031942 | 0.1268618 |
| ENSG00000170892.10 | TSEN34 | -1.097727 | 0.6201989 | 0.123997 | 0.8362229 | -0.565218 | 0.000399694 | 0.514178 | 0.3153792 | 2.167327 | 0.01642174 | -0.713522 | 0.985679 |
| ENSG00000170899.10 | GSTA4 | 1.945371 | 0.653693 | -0.313261 | 0.5010139 | 1.06485 | 0.02275597 | 1.05261 | 0.4919368 | 0.46173 | 0.6492082 | 0.448514 | 0.7975504 |
| ENSG00000170903.10 | MSANTD4 | -0.076426 | 0.682906 | 0.612848 | 0.7453671 | 0.780884 | 0.03732293 | -0.536888 | 0.2335397 | 1.546511 | 0.6419765 | 1.778493 | 0.6885065 |
| ENSG00000170906.15 | NDUFA3 | 0.589412 | 0.5671641 | -0.334442 | 0.1982022 | -1.195438 | 5.77E-13 | -0.025149 | 0.02566394 | -0.373678 | 0.008567685 | -0.259164 | 0.6042097 |
| ENSG00000170921.14 | TANC2 | -1.570502 | 0.8587992 | 0.279191 | 0.8703522 | 0.005932 | 0.000117345 | 1.129806 | 0.001566916 | 2.257569 | 0.00094153 | -0.920974 | 0.8278576 |
| ENSG00000171033.12 | PKIA | -0.029836 | 0.9870834 | 0.647924 | 0.6527736 | 0.431727 | 0.370503 | 1.895425 | 8.57E-07 | 0.041194 | 1 | 1.561115 | 0.6829816 |
| ENSG00000171055.14 | FEZ2 | 0.152621 | 0.6214382 | -0.118301 | 0.3651168 | 0.384659 | 0.3538945 | 0.953711 | 0.7111474 | -0.489707 | 0.000980785 | -0.574813 | 0.6261342 |
| ENSG00000171067.10 | C11orf24 | -0.404819 | 0.8203421 | -0.462538 | 0.6335433 | -0.09585 | 0.2733328 | 0.519779 | 0.04501672 | 0.784495 | 0.4174983 | 1.204909 | 0.9134166 |
| ENSG00000171097.13 | CCBL1 | 2.419037 | 0.714717 | 0.434431 | 1 | 0.7326 | 0.64645 | -0.496835 | 0.9119095 | 2.258975 | 0.03378388 | -1.650975 | 0.8949489 |
| ENSG00000171105.13 | INSR | -1.671231 | 0.563505 | 0.55049 | 0.9028136 | 0.269747 | 0.5608325 | 1.000956 | 0.01035831 | 1.907529 | 0.2487425 | -1.611284 | 0.8770536 |
| ENSG00000171132.13 | PRKCE | 0.464839 | 0.9289591 | -0.035871 | 0.9798987 | -1.756362 | 0.7878575 | 2.844436 | 0.01230964 | 2.335063 | 0.2922672 | 2.136271 | 0.7175744 |
| ENSG00000171159.4 | C9orf16 | 1.019645 | 0.3253495 | 0.528373 | 0.7156572 | -0.459126 | 0.08151947 | 0.451775 | 0.7361614 | -0.221515 | 0.01703808 | -0.009292 | 0.8318522 |
| ENSG00000171202.6 | TMEM126A | 0.436123 | 0.8006286 | 0.268344 | 0.8785714 | -0.660006 | 0.006577094 | 0.161837 | 0.0709208 | -0.10444 | 0.08724925 | -0.725453 | 0.9241789 |
| ENSG00000171204.12 | TMEM126B | -0.04539 | 0.8006286 | -0.267179 | 0.6527736 | 0.291138 | 0.8688128 | 0.127988 | 0.02261337 | 1.138746 | 0.6433334 | 0.363277 | 1 |
| ENSG00000171222.10 | SCAND1 | 0.112035 | 0.6214382 | 0.021856 | 0.6527736 | -0.735594 | 7.03E-11 | -0.037207 | 0.2174842 | -0.304723 | 0.03848503 | 1.290006 | 1 |
| ENSG00000171223.5 | JUNB | 0.505731 | 0.007696271 | 0.38022 | 0.000806275 | -1.593147 | 3.93E-28 | 0.699415 | 0.9069344 | 1.077133 | 0.5606837 | 0.227984 | 1 |
| ENSG00000171291.8 | ZNF439 | 2.166682 | 0.4764974 | -0.530487 | 0.8953649 | 0.834806 | 0.04759723 | 0.024932 | 1 | 1.925033 | 0.01596134 | -1.225374 | 0.7494298 |
| ENSG00000171310.10 | CHST11 | -2.872935 | 0.4406957 | 0.325105 | 0.8953649 | 1.412393 | 0.02506782 | 3.427631 | 0.3373026 | 0.608288 | 0.7541814 | 1.532939 | 0.6885065 |
| ENSG00000171314.8 | PGAM1 | 0.055073 | 0.5538099 | 0.391107 | 0.08640355 | 0.339172 | 0.2495461 | 0.210221 | 0.0438475 | 0.573073 | 0.8053132 | 0.509016 | 0.6721427 |
| ENSG00000171316.11 | CHD7 | 0.168934 | 0.8489642 | -0.637774 | 0.6483776 | 0.807138 | 0.000551725 | 2.01689 | 5.60E-08 | 1.744832 | 2.43E-08 | 0.545219 | 0.5700377 |
| ENSG00000171385.9 | KCND3 | -1.64713 | 1 | -1.221413 | 0.4011918 | 1.629823 | 0.03615611 | 2.051433 | 9.26E-07 | 0.898842 | 1 | -0.785706 | 0.9147227 |
| ENSG00000171421.12 | MRPL36 | 0.237885 | 0.6893253 | -0.011892 | 0.9103295 | -0.13101 | 0.08450958 | -0.458878 | 0.04115065 | -0.276668 | 0.5196843 | -0.631385 | 0.7870028 |
| ENSG00000171451.13 | DSEL | -0.003244 | 0.7411192 | 0.101489 | 0.8465126 | 0.595031 | 0.01652201 | 1.045494 | 9.93E-06 | 0.071274 | 1 | -0.862817 | 0.6885065 |
| ENSG00000171456.16 | ASXL1 | -0.47714 | 0.6214382 | -0.091774 | 0.5395105 | -0.644244 | 0.7720073 | 1.311546 | 2.58E-05 | 0.792136 | 0.06448943 | 2.292577 | 0.9726757 |
| ENSG00000171466.9 | ZNF562 | -0.97405 | 0.527339 | -0.163649 | 0.3206004 | 1.023171 | 0.06202009 | 0.941175 | 0.002041384 | 0.382061 | 0.3974318 | 0.474992 | 0.9147227 |
| ENSG00000171467.15 | ZNF318 | 0.364624 | 0.9183025 | -0.293947 | 0.9442632 | 0.4304 | 0.233483 | 0.534483 | 0.1880764 | 1.963893 | 0.02289798 | 0.677848 | 0.6885065 |
| ENSG00000171475.13 | WIPF2 | -0.069255 | 0.7703009 | 0.386563 | 0.944317 | -0.869049 | 1 | 1.203253 | 0.005559663 | 0.179514 | 0.5382216 | 0.574782 | 0.6885065 |
| ENSG00000171490.12 | RSL1D1 | -0.281608 | 0.8867323 | 0.154143 | 0.8263112 | 0.820783 | 4.65E-09 | 0.210016 | 0.1860003 | -0.396562 | 0.008692014 | -0.554022 | 0.5733051 |
| ENSG00000171530.13 | TBCA | -0.214365 | 0.9867064 | -0.200329 | 0.4011918 | -0.192494 | 0.001696806 | -0.09934 | 0.02991167 | -0.21252 | 0.002050046 | 0.16503 | 0.6885065 |
| ENSG00000171533.11 | MAP6 | -1.073248 | 0.6214382 | -1.024998 | 0.5953732 | 0.711103 | 0.5290845 | 2.106184 | 0.04009018 | 0.31341 | 0.00557683 | -1.61913 | 0.6885065 |
| ENSG00000171566.11 | PLRG1 | -0.840383 | 0.9198482 | 0.328533 | 0.5736784 | -0.22099 | 0.9941063 | 1.12125 | 0.0496401 | 1.080118 | 0.6294954 | 0.166617 | 0.7767294 |
| ENSG00000171587.14 | DSCAM | -1.036947 | 1 | -1.983305 | 0.5517607 | -3.02005 | 0.7437308 | -0.144313 | 0.006286247 | 0.75811 | 1 | -2.472859 | 0.9850898 |
| ENSG00000171603.16 | CLSTN1 | 0.969311 | 0.9183025 | -0.792117 | 0.004197116 | 0.491677 | 0.6292043 | 1.360515 | 1.36E-06 | 1.2741 | 3.49E-05 | 1.038028 | 0.7571519 |
| ENSG00000171606.17 | ZNF274 | -0.766401 | 0.6214382 | 0.087642 | 0.7778236 | -0.21971 | 0.6651783 | 1.661238 | 0.08656066 | 2.166599 | 0.002548226 | 1.56753 | 0.6517664 |
| ENSG00000171617.13 | ENC1 | -0.711398 | 0.954995 | 0.487469 | 0.9285327 | -0.327373 | 0.7284574 | 1.842877 | 6.44E-07 | 0.123795 | 0.3638582 | 1.309071 | 0.9270789 |
| ENSG00000171634.16 | BPTF | 0.339035 | 0.6214382 | 0.028135 | 0.5010139 | 0.784443 | 0.001437818 | 1.7058 | 2.98E-05 | 1.289344 | 0.03923426 | -2.391239 | 0.5812196 |
| ENSG00000171681.12 | ATF7IP | -0.578009 | 0.9647416 | -0.303978 | 0.6527736 | 0.9218 | 1.33E-06 | 0.695737 | 0.002470523 | 1.847659 | 0.000749346 | 2.129834 | 0.7904104 |
| ENSG00000171703.16 | TCEA2 | -0.402247 | 0.9802976 | 0.41838 | 0.4841414 | -1.425108 | 0.004032107 | 1.335793 | 0.630264 | 1.099184 | 0.3387021 | -0.597875 | 0.8208081 |
| ENSG00000171735.18 | CAMTA1 | -0.10775 | 0.8415359 | -0.341566 | 0.1834552 | -0.325886 | 0.04944273 | 0.011052 | 0.06377668 | -0.088984 | 0.3697675 | 0.623442 | 0.6552711 |
| ENSG00000171792.10 | RHNO1 | 0.24266 | 0.9047689 | -1.240213 | 0.02338178 | 0.413899 | 0.5290845 | 0.216798 | 0.9471387 | 0.581767 | 0.7377262 | -1.385519 | 0.6885065 |
| ENSG00000171793.13 | CTPS1 | -1.660739 | 0.6792949 | -0.324487 | 0.4011918 | 0.137813 | 0.01540585 | 1.121378 | 0.2930784 | 0.881616 | 0.4432323 | -0.385831 | 0.6185816 |
| ENSG00000171853.15 | TRAPPC12 | -1.164355 | 0.6214382 | -0.560883 | 0.5993025 | -1.398085 | 0.01347147 | 0.332446 | 0.2335397 | 1.805369 | 0.03157649 | -0.940647 | 0.8479431 |
| ENSG00000171858.17 | RPS21 | -0.46314 | 0.9660232 | 0.079542 | 0.7820217 | -0.543469 | 1.25E-24 | 0.232915 | 0.2526197 | -0.282464 | 4.35E-05 | 0.257129 | 0.5841302 |
| ENSG00000171862.9 | PTEN | 0.171772 | 1 | -0.238055 | 0.5010139 | 1.0245 | 0.192418 | 2.046097 | 0.06406476 | 2.250344 | 0.04590921 | 1.807881 | 0.6829816 |
| ENSG00000171863.12 | RPS7 | -0.445514 | 0.1413791 | 0.253596 | 0.4072962 | 0.225404 | 0.000256547 | -0.360214 | 2.92E-10 | -0.121129 | 0.003357137 | -0.031126 | 0.2281002 |
| ENSG00000171943.11 | SRGAP2C | 0.276225 | 0.8006286 | -0.136445 | 0.8703522 | 1.157489 | 1.94E-07 | 1.746776 | 0.384335 | 2.063134 | 0.1206097 | 0.289336 | 0.5524003 |
| ENSG00000171951.4 | SCG2 | 0.154363 | 3.45E-06 | 0.088579 | 0.6527736 | 0.339665 | 0.8380672 | 0.60444 | 0.04072971 | 0.156556 | 0.2880317 | 0.528274 | 0.6185816 |
| ENSG00000171960.10 | PPIH | -0.91836 | 0.6214382 | -0.107808 | 0.9100694 | -0.365309 | 0.04841948 | 0.441255 | 1 | -0.437678 | 0.2990962 | -0.068351 | 0.6885065 |
| ENSG00000171988.17 | JMJD1C | -0.455923 | 0.7225091 | -0.029147 | 0.8894405 | 0.60292 | 7.70E-05 | 1.629326 | 0.000466708 | 1.234502 | 0.7397084 | -1.094483 | 0.9589318 |
| ENSG00000172007.5 | RAB33B | -0.637919 | 1 | -0.140299 | 0.8149803 | 1.528763 | 0.4382561 | 2.80544 | 0.2481975 | 2.219933 | 0.0250399 | 1.679727 | 0.6885065 |
| ENSG00000172009.14 | THOP1 | 1.174772 | 0.6214382 | 0.072214 | 0.9486562 | 0.216557 | 0.606319 | 1.091988 | 0.02252143 | 1.41221 | 0.1327475 | -0.778778 | 1 |
| ENSG00000172020.12 | GAP43 | 0.350926 | 8.98E-12 | -0.279164 | 0.00031583 | 0.246784 | 0.3119492 | 0.14184 | 0.4919368 | -0.515634 | 3.86E-10 | -0.19176 | 0.4553016 |
| ENSG00000172057.9 | ORMDL3 | -1.009304 | 0.8744474 | -0.323617 | 0.930564 | 0.441343 | 0.9781587 | 1.103719 | 0.02806712 | 0.375581 | 0.5196843 | -0.096826 | 0.9037836 |
| ENSG00000172071.11 | EIF2AK3 | -0.832302 | 0.8006286 | 0.140898 | 0.5299416 | 0.293394 | 0.8644385 | 2.050085 | 0.03053125 | 1.667233 | 0.2105971 | -0.928517 | 1 |
| ENSG00000172115.8 | CYCS | -1.88197 | 0.2398916 | 0.09368 | 0.6527736 | -0.133529 | 0.07009909 | 0.407614 | 0.8502343 | -0.070979 | 0.002550804 | 0.160693 | 0.3414178 |
| ENSG00000172172.7 | MRPL13 | -0.277923 | 1 | 0.023587 | 0.9573504 | 0.45393 | 1 | -0.518505 | 0.000100715 | -0.081394 | 0.1312457 | 0.312131 | 0.6885065 |
| ENSG00000172175.12 | MALT1 | -2.252347 | 0.6214382 | 0.683407 | 0.8263112 | 0.545451 | 0.4655358 | -0.273128 | 0.4919368 | 1.23098 | 0.04222895 | 1.201204 | 0.8532529 |
| ENSG00000172183.14 | ISG20 | -0.777564 | 0.6214382 | -0.097434 | 0.4961426 | -0.475108 | 0.238948 | -0.127582 | 0.001350605 | -0.759335 | 0.643255 | 2.374794 | 0.9799292 |
| ENSG00000172201.10 | ID4 | -0.752777 | 0.4908084 | 0.071313 | 0.906244 | -0.165047 | 0.5290845 | 1.329804 | 0.002584584 | 0.871002 | 0.7964713 | -0.441529 | 0.7136264 |
| ENSG00000172262.11 | ZNF131 | -1.310997 | 0.980017 | 0.130962 | 1 | 0.520885 | 0.4344023 | 0.297723 | 0.9041574 | 1.929421 | 0.006360815 | 1.25367 | 0.8085394 |
| ENSG00000172270.18 | BSG | 0.361519 | 0.8996471 | 0.342822 | 0.02922882 | -0.768963 | 1.27E-08 | 0.892487 | 0.02343577 | 0.952936 | 0.6821496 | 0.124173 | 0.6885065 |
| ENSG00000172301.10 | COPRS | -0.378961 | 0.8587992 | -0.191915 | 0.6200786 | -1.342439 | 0.000162288 | 0.78496 | 0.06811579 | -0.521033 | 0.7701253 | 0.351645 | 0.8489889 |
| ENSG00000172315.5 | TP53RK | 0.343762 | 0.4764974 | 0.273993 | 0.6527736 | -1.59925 | 3.81E-06 | 0.854452 | 0.338872 | -0.143822 | 0.9438109 | 2.443567 | 0.881999 |
| ENSG00000172336.4 | POP7 | -0.445396 | 0.3787871 | 0.097495 | 1 | 0.294451 | 0.7437308 | 0.091098 | 0.6902435 | 0.128097 | 0.01388672 | 0.86879 | 0.9077431 |
| ENSG00000172354.9 | GNB2 | -0.017633 | 0.4406957 | -0.211522 | 0.1583423 | -0.330842 | 9.49E-06 | 0.562793 | 0.005383314 | 0.952121 | 0.002229761 | -0.221382 | 0.5026529 |
| ENSG00000172366.19 | FAM195A | -0.900377 | 0.7648042 | -0.190336 | 1 | -0.351236 | 0.003030989 | 0.896325 | 0.8224647 | -0.686752 | 0.5788462 | -0.205997 | 0.6490419 |
| ENSG00000172379.18 | ARNT2 | -1.401588 | 0.6014343 | -0.055848 | 0.4281695 | -0.043072 | 0.9940965 | 1.601371 | 0.000113249 | 0.256529 | 0.464861 | 1.832298 | 1 |
| ENSG00000172380.5 | GNG12 | -2.349424 | 1 | -0.141349 | 0.6527736 | 1.342819 | 0.01192579 | 0.239652 | 0.8059372 | 1.177395 | 1 | 1.058404 | 0.530077 |
| ENSG00000172428.10 | MYEOV2 | 0.617248 | 0.004460494 | 0.1109 | 1 | -0.781299 | 2.72E-14 | -0.084263 | 0.006264535 | -0.301562 | 9.04E-06 | -0.336913 | 0.2093204 |
| ENSG00000172466.15 | ZNF24 | -0.412002 | 0.6014343 | -0.28025 | 0.2079972 | 0.722884 | 2.73E-07 | 0.592342 | 0.1796423 | 0.734538 | 0.9804192 | 0.66025 | 0.6885065 |
| ENSG00000172469.14 | MANEA | 2.820561 | 0.6214382 | 0.670151 | 0.6527736 | 1.705793 | 0.1157816 | 2.43082 | 0.03741247 | 1.380677 | 0.8293855 | -0.961954 | 0.9438593 |
| ENSG00000172534.13 | HCFC1 | 1.474488 | 0.651389 | -1.043524 | 0.5010139 | -0.443549 | 0.5451095 | 0.836451 | 0.06070451 | 2.017043 | 3.16E-05 | -0.447178 | 0.6185816 |
| ENSG00000172590.18 | MRPL52 | 0.174559 | 0.8587992 | -0.392711 | 0.03342537 | -0.180696 | 1.91E-06 | 0.136931 | 0.7144938 | 0.055493 | 0.1688763 | -0.092685 | 0.468071 |
| ENSG00000172667.10 | ZMAT3 | -0.083041 | 0.8006286 | -0.74431 | 0.06357043 | 1.014769 | 0.5071415 | 0.24186 | 0.0381379 | -0.084197 | 0.7544768 | 0.248132 | 0.6267144 |
| ENSG00000172717.15 | FAM71D | 1.235828 | 0.8587992 | 1.771734 | 0.5010139 | -2.3947 | 1.52E-14 | 0.3091 | 0.8224647 | 0.790164 | 0.8013652 | 0.554593 | 0.6829816 |
| ENSG00000172757.12 | CFL1 | -0.440714 | 0.928247 | -0.086263 | 0.5967879 | -0.028805 | 0.06249689 | 0.274289 | 0.4919368 | -0.029721 | 0.03593768 | -0.288582 | 0.05186339 |
| ENSG00000172766.18 | NAA16 | -2.706628 | 0.6214382 | -0.500241 | 0.3530125 | 0.833761 | 4.49E-06 | 1.551235 | 0.01659537 | 1.457963 | 0.3300883 | 1.081466 | 0.958512 |
| ENSG00000172775.16 | FAM192A | -0.407841 | 0.8006286 | -0.318339 | 0.5227713 | 0.061638 | 0.1142985 | 0.27912 | 0.7339795 | 0.019149 | 0.003493004 | -0.357843 | 0.6885065 |
| ENSG00000172785.18 | CBWD1 | 0.222714 | 1 | 1.164568 | 0.4530135 | 0.383394 | 0.8535417 | -0.767818 | 0.02631783 | -0.164639 | 0.7285653 | -0.111074 | 1 |
| ENSG00000172795.15 | DCP2 | -1.536446 | 0.9047689 | 0.29401 | 0.8177316 | 0.522116 | 0.01090776 | 0.425949 | 0.1525415 | 1.564994 | 0.06141896 | 2.567233 | 0.6885065 |
| ENSG00000172809.12 | RPL38 | 0.046216 | 0.714717 | -0.033088 | 0.780648 | -0.4561 | 8.13E-14 | 0.31948 | 0.3186265 | 0.087172 | 0.120105 | -0.335478 | 0.02166254 |
| ENSG00000172845.13 | SP3 | 0.079066 | 0.9575547 | 0.342561 | 0.5953732 | 0.060602 | 0.5881008 | 1.22864 | 0.006807311 | 1.857006 | 0.03257419 | 1.224922 | 0.6517664 |
| ENSG00000172890.11 | NADSYN1 | -0.212787 | 0.6214382 | 1.235011 | 0.2758583 | 0.645895 | 0.03256043 | 1.124092 | 0.1633858 | 2.977852 | 0.6252573 | 2.671456 | 0.6885065 |
| ENSG00000172915.18 | NBEA | -0.660755 | 0.7427276 | -0.993396 | 0.6515605 | 1.06443 | 0.0993183 | 1.163795 | 0.01773501 | 0.514048 | 0.5196843 | 0.021447 | 0.7864577 |
| ENSG00000172943.18 | PHF8 | 0.976792 | 0.822435 | -0.418587 | 0.6527736 | 0.462201 | 0.5290845 | 1.820713 | 0.007615897 | 0.376954 | 0.8021966 | 1.694471 | 0.6885065 |
| ENSG00000172985.10 | SH3RF3 | -1.212478 | 1 | -0.02311 | 0.6527736 | -1.304473 | 0.5430378 | 2.008664 | 0.02248256 | 1.972006 | 0.5196843 | 3.085115 | 0.9264343 |
| ENSG00000173064.10 | HECTD4 | -1.618736 | 0.714717 | 0.819057 | 0.8188012 | 0.475628 | 0.8607697 | 1.421874 | 0.000214251 | 2.872262 | 7.78E-05 | -1.277462 | 0.6829816 |
| ENSG00000173085.13 | COQ2 | -0.618235 | 0.6014343 | 0.308737 | 0.8113877 | -1.294751 | 0.03832258 | 0.957049 | 0.6663027 | 0.240121 | 0.905839 | -0.560158 | 0.9403369 |
| ENSG00000173110.7 | HSPA6 | -1.679588 | 0.714717 | -0.253467 | 0.8893354 | 1.380505 | 1.05E-08 | -0.093928 | 0.9733822 | 2.266824 | 0.01193561 | 1.182134 | 1 |
| ENSG00000173113.6 | TRMT112 | -0.048375 | 0.7855557 | 0.035783 | 0.9217053 | -0.354761 | 2.18E-06 | -0.173096 | 0.000190613 | -0.006054 | 0.009236574 | -0.080906 | 0.7417751 |
| ENSG00000173114.12 | LRRN3 | -1.156744 | 0.7736255 | -0.407684 | 0.2296726 | -1.097729 | 0.290478 | 0.491168 | 0.5129328 | 1.735059 | 0.004512036 | -4.872614 | 0.6885065 |
| ENSG00000173120.14 | KDM2A | -0.227522 | 0.9678883 | -0.135883 | 0.7972162 | 0.223076 | 0.9526702 | 1.255764 | 8.62E-05 | 0.943528 | 0.01618785 | -1.714727 | 0.6200022 |
| ENSG00000173141.4 | MRPL57 | -0.299676 | 0.9729215 | 0.150411 | 0.7881421 | -0.505789 | 0.01845196 | -0.703454 | 0.005933699 | -0.009139 | 0.1665916 | 0.27824 | 0.348294 |
| ENSG00000173153.13 | ESRRA | -0.685649 | 0.6014343 | 0.296687 | 0.9403621 | 0.341435 | 0.7437308 | 1.246045 | 0.01714385 | 0.376857 | 0.9362282 | 0.208827 | 0.5023959 |
| ENSG00000173171.14 | MTX1 | 0.112 | 0.6214382 | 0.088853 | 0.7483725 | -1.040133 | 0.002629245 | 0.138457 | 1 | -1.286347 | 0.1633784 | -3.05511 | 0.5552649 |
| ENSG00000173209.22 | AHSA2 | -2.679386 | 0.3946735 | 0.20977 | 0.9856559 | 0.288496 | 0.4064599 | 0.621034 | 0.5934221 | 1.373747 | 0.0229376 | 2.508154 | 0.7728704 |
| ENSG00000173230.15 | GOLGB1 | 1.609104 | 0.6214382 | 0.484928 | 0.9834088 | 0.867524 | 1.87E-06 | 0.391907 | 0.05894564 | 0.428519 | 0.8965775 | 0.819214 | 0.9002633 |
| ENSG00000173264.13 | GPR137 | 1.150448 | 0.6214382 | -0.615009 | 0.9899626 | 0.589683 | 0.5290845 | 2.726579 | 0.0362763 | -1.686116 | 0.9383375 | 1.181125 | 0.8479431 |
| ENSG00000173272.13 | MZT2A | -0.433887 | 0.6799027 | -0.448016 | 0.07080539 | -1.685796 | 5.98E-12 | 0.332698 | 0.4919368 | 0.440331 | 0.6006688 | 0.605028 | 0.5344421 |
| ENSG00000173273.15 | TNKS | -0.121134 | 0.6409583 | 0.12742 | 0.9873764 | 0.422589 | 0.000521203 | 1.421163 | 0.1292522 | 0.945594 | 0.1976806 | -1.815495 | 0.7120122 |
| ENSG00000173276.13 | ZBTB21 | -1.058819 | 0.6214382 | -0.026906 | 0.9798987 | 0.274814 | 0.02583133 | 0.477208 | 0.4716463 | 0.517593 | 0.6019412 | 1.221626 | 0.6885065 |
| ENSG00000173281.4 | PPP1R3B | -2.295787 | 0.08764673 | 0.017092 | 1 | -0.974083 | 0.5290845 | 0.937956 | 0.03043518 | 1.384903 | 1 | 1.297612 | 0.7095467 |
| ENSG00000173320.9 | STOX2 | 2.027438 | 0.3946735 | -0.733581 | 0.9339858 | -0.488319 | 0.5056489 | 0.842392 | 0.01755988 | 1.214717 | 0.000923521 | 1.514504 | 0.9526606 |
| ENSG00000173404.4 | INSM1 | 3.105343 | 0.2404198 | 0.211691 | 0.9821542 | -2.682376 | 0.01267166 | 1.514067 | 3.33E-06 | 1.845692 | 3.89E-06 | 1.068182 | 0.6829816 |
| ENSG00000173418.11 | NAA20 | -0.87956 | 0.1118348 | 0.96122 | 0.0172833 | -0.187323 | 0.5290845 | -0.163156 | 0.000301883 | -0.299367 | 0.02269293 | -0.407983 | 0.9931654 |
| ENSG00000173436.13 | MINOS1 | 0.814861 | 0.5918523 | -0.058846 | 0.7457394 | -0.580299 | 8.09E-09 | -0.308676 | 0.0765766 | -0.58106 | 0.000783481 | -0.74183 | 0.06501308 |
| ENSG00000173451.6 | THAP2 | -2.602254 | 0.6014343 | 0.032834 | 0.8212472 | 0.069717 | 0.001625198 | 2.537024 | 0.08534343 | 0.326498 | 1 | -0.817001 | 0.9460594 |
| ENSG00000173457.10 | PPP1R14B | -4.176428 | 0.001736947 | -0.087307 | 0.4580322 | -0.890052 | 1.69E-11 | 0.53849 | 0.003240262 | 2.046193 | 2.79E-13 | 0.085386 | 0.5256722 |
| ENSG00000173465.7 | SSSCA1 | -0.622302 | 0.714717 | -0.627919 | 0.4208458 | -0.823143 | 0.009112063 | -0.546626 | 0.01886265 | -0.506397 | 0.8438512 | -1.045347 | 0.6885065 |
| ENSG00000173473.10 | SMARCC1 | -0.652811 | 0.6214382 | -0.1655 | 0.4420234 | 0.30399 | 0.000130234 | 1.358555 | 0.001089214 | 1.403669 | 0.04581517 | 0.191102 | 0.6885065 |
| ENSG00000173486.12 | FKBP2 | -0.346985 | 0.8006286 | -0.440392 | 0.1290746 | -0.307828 | 0.000171674 | 0.297457 | 0.4919368 | -0.000963 | 0.1310859 | -0.11645 | 0.6489815 |
| ENSG00000173517.10 | PEAK1 | -3.963222 | 0.06285358 | 0.034256 | 0.6527736 | 1.875723 | 0.03475121 | 0.290394 | 0.5888232 | 2.539991 | 0.00095954 | -0.690802 | 0.9733982 |
| ENSG00000173540.12 | GMPPB | -2.447292 | 0.2430963 | -0.620879 | 0.6527736 | -0.779286 | 0.01893482 | 1.114372 | 0.7002918 | 0.827922 | 0.8387831 | 0.380718 | 0.926878 |
| ENSG00000173542.8 | MOB1B | 0.060555 | 0.9355297 | -1.041425 | 0.06481976 | 0.604658 | 0.000483747 | 1.543093 | 0.0240678 | 1.846725 | 0.009755487 | 0.213905 | 0.4306015 |
| ENSG00000173545.4 | ZNF622 | 1.076668 | 0.6214382 | -0.067098 | 0.8954254 | -0.025561 | 1 | 0.230127 | 0.009221742 | -0.556718 | 0.1436435 | -0.775053 | 0.6885065 |
| ENSG00000173575.18 | CHD2 | -0.874829 | 0.3619888 | -0.154252 | 0.1335004 | 0.616702 | 8.17E-06 | 1.075982 | 0.02377077 | 0.942877 | 0.04521787 | 0.868088 | 0.6885065 |
| ENSG00000173598.13 | NUDT4 | -0.98527 | 0.9645544 | -0.303742 | 0.1606176 | 0.171478 | 0.5856773 | 0.848918 | 0.000269574 | 0.372423 | 0.5196843 | -0.373878 | 0.5344421 |
| ENSG00000173621.8 | LRFN4 | -0.433389 | 0.9401246 | 0.088138 | 1 | -1.364715 | 0.000521203 | -0.02333 | 0.4986906 | 1.194528 | 0.04969669 | -0.053446 | 0.8479431 |
| ENSG00000173638.18 | SLC19A1 | 1.736469 | 0.6214382 | -0.146251 | 0.4053594 | -2.655633 | 0.4644258 | 2.254674 | 0.04811661 | 0.892693 | 0.2010318 | 0.913415 | 0.8949489 |
| ENSG00000173653.7 | RCE1 | -2.801822 | 0.5711449 | 0.269345 | 0.8153304 | -0.72403 | 0.01250308 | -0.601975 | 0.1847644 | 0.77003 | 0.963643 | 1.972149 | 0.6885065 |
| ENSG00000173660.11 | UQCRH | -0.002619 | 0.9700233 | -0.188484 | 0.1042612 | -0.072118 | 0.4064599 | 0.070681 | 0.1436444 | -0.256539 | 0.000710986 | -0.658142 | 0.04053673 |
| ENSG00000173674.10 | EIF1AX | 0.173723 | 0.563505 | 0.151493 | 0.9315666 | 0.707762 | 0.000207276 | 0.317491 | 0.7356286 | 0.186873 | 0.6868144 | 0.534617 | 0.9618424 |
| ENSG00000173681.16 | CXorf23 | -1.438573 | 0.7363511 | -0.345006 | 0.8215564 | 0.954243 | 0.8959394 | 2.265509 | 0.05672417 | 2.567671 | 0.03327835 | 0.664074 | 0.7126184 |
| ENSG00000173692.12 | PSMD1 | -0.383313 | 0.5242565 | -0.219718 | 0.4001101 | 0.644533 | 2.07E-05 | 0.724248 | 0.8319427 | 0.277268 | 0.06028616 | 1.104438 | 0.6885065 |
| ENSG00000173744.17 | AGFG1 | -0.257046 | 0.6226158 | -0.067944 | 0.9403621 | 0.683981 | 0.1889106 | 0.639438 | 0.03351837 | 0.706844 | 0.0839525 | 1.268412 | 0.8949489 |
| ENSG00000173757.9 | STAT5B | -0.153443 | 0.8587992 | -0.76501 | 0.5813882 | 1.139002 | 0.2879524 | 1.851327 | 0.02496282 | 1.444598 | 0.01024338 | -0.796058 | 0.6829816 |
| ENSG00000173786.16 | CNP | -0.992607 | 0.1119893 | -0.53543 | 0.002819794 | -0.092872 | 0.445851 | 0.140276 | 0.8253115 | 0.275289 | 0.9963991 | 0.884033 | 0.8218739 |
| ENSG00000173846.12 | PLK3 | 1.145649 | 0.529676 | 0.22366 | 0.6527736 | 0.866789 | 0.8100781 | 1.584615 | 0.07602296 | 2.116428 | 0.01618785 | 2.812721 | 0.6185816 |
| ENSG00000173875.13 | ZNF791 | -0.051091 | 0.6214382 | -0.114812 | 0.8263112 | 0.434189 | 0.0686743 | 2.339335 | 0.02238098 | 1.330211 | 0.02209058 | 0.039392 | 0.9618424 |
| ENSG00000173889.15 | PHC3 | -0.659349 | 0.6214382 | -0.175049 | 0.8263112 | 1.380696 | 2.08E-05 | 1.667884 | 0.007421547 | -0.818564 | 0.1265615 | 0.410173 | 0.9646502 |
| ENSG00000173898.11 | SPTBN2 | 0.513567 | 0.714717 | -0.85639 | 0.125909 | -0.777165 | 0.3144442 | 1.18882 | 0.000865519 | 3.238796 | 0.000279287 | 1.9953 | 0.9661868 |
| ENSG00000173915.12 | USMG5 | 0.10975 | 0.2506697 | 0.571211 | 0.1003599 | -0.550022 | 3.26E-08 | -0.077998 | 0.01114426 | -0.564769 | 0.000432014 | -0.240155 | 0.1508459 |
| ENSG00000173960.12 | UBXN2A | 0.640913 | 0.8756719 | 0.2619 | 0.9766519 | 0.787753 | 0.009881126 | 0.572862 | 0.1751371 | 0.960366 | 0.4499166 | 0.123758 | 0.9148375 |
| ENSG00000173991.5 | TCAP | -3.608755 | 0.6214382 | -1.920583 | 0.2758186 | -1.637861 | 0.4655358 | 2.32218 | 0.03985619 | 3.20724 | 0.03399884 | -3.321384 | 0.7767294 |
| ENSG00000174013.7 | FBXO45 | -0.369597 | 0.7898271 | 0.443677 | 0.6527736 | 1.061969 | 0.02290791 | 0.726618 | 0.01289896 | 1.534877 | 0.0641349 | -1.600002 | 0.9506884 |
| ENSG00000174021.10 | GNG5 | -0.544692 | 0.9667286 | -0.031102 | 0.8263112 | -0.169161 | 0.1134172 | -0.445482 | 2.60E-08 | -0.243878 | 0.1197239 | 0.040078 | 0.468071 |
| ENSG00000174106.2 | LEMD3 | -1.780688 | 0.8006286 | -0.095875 | 0.8703522 | 0.719734 | 0.6427115 | 2.889448 | 0.000688376 | 3.279698 | 0.04249027 | 0.645234 | 0.9094844 |
| ENSG00000174136.11 | RGMB | 0.569804 | 0.5925284 | -0.126169 | 0.9009117 | -0.042416 | 0.9682148 | 1.075517 | 2.49E-11 | 1.178867 | 0.001255969 | -0.457316 | 0.4974178 |
| ENSG00000174151.14 | CYB561D1 | 1.02471 | 0.8006286 | 1.610789 | 0.7049761 | -0.284205 | 0.7720073 | 2.754991 | 0.01754162 | 1.566086 | 0.5052437 | -2.785659 | 0.8606444 |
| ENSG00000174197.16 | MGA | 0.07826 | 0.9426668 | 0.494618 | 0.704767 | 0.426427 | 0.01466097 | 0.714482 | 0.042808 | 1.053808 | 0.08559807 | 1.858116 | 0.6885065 |
| ENSG00000174227.15 | PIGG | 1.373513 | 0.6169802 | -0.364686 | 0.6527736 | 0.519816 | 0.609629 | 1.745233 | 0.03641313 | 1.673478 | 0.7117629 | 0.778238 | 0.9147227 |
| ENSG00000174231.16 | PRPF8 | 0.470609 | 0.8006286 | 0.139932 | 0.6527736 | 0.509054 | 0.009867582 | 1.307665 | 0.01104565 | 0.930193 | 0.7620165 | 0.158814 | 0.9204395 |
| ENSG00000174233.11 | ADCY6 | -0.583678 | 0.6214382 | 0.334067 | 0.9766239 | -1.099839 | 0.5752162 | 0.688237 | 0.06019179 | 1.808764 | 0.0325649 | 1.340439 | 0.5344421 |
| ENSG00000174306.21 | ZHX3 | 0.099064 | 0.6214382 | -0.307086 | 0.6527736 | -0.980215 | 1 | 2.073939 | 0.001320651 | 0.717273 | 0.6008953 | -0.293054 | 0.6885065 |
| ENSG00000174307.6 | PHLDA3 | 1.068066 | 0.8716459 | -0.408496 | 0.7232541 | -2.73988 | 0.02506782 | 1.222553 | 0.5837245 | 3.01462 | 0.5196843 | -2.527644 | 0.9617345 |
| ENSG00000174437.16 | ATP2A2 | 0.443837 | 0.8006286 | 0.05754 | 0.9829027 | -0.101326 | 0.772371 | 1.867278 | 7.15E-10 | 1.409238 | 0.01100915 | 1.294398 | 0.6885065 |
| ENSG00000174444.14 | RPL4 | -1.193321 | 4.95E-10 | -0.062075 | 0.8703522 | 0.592651 | 4.58E-14 | 0.295525 | 0.7858185 | -0.081374 | 0.01078935 | -0.537674 | 0.5733051 |
| ENSG00000174446.12 | SNAPC5 | -0.67056 | 0.6214382 | -0.306761 | 0.5953732 | -0.814562 | 0.03372233 | 0.210464 | 0.7144938 | -0.395597 | 0.5196843 | -3.726893 | 0.5344421 |
| ENSG00000174501.14 | ANKRD36C | 2.831332 | 0.2381383 | -0.21336 | 0.5484483 | 0.439944 | 1 | 1.320414 | 1.36E-05 | 2.153954 | 0.01138348 | -0.16047 | 0.6829816 |
| ENSG00000174521.7 | TTC9B | 0.199974 | 1 | -1.093962 | 0.193606 | 0.093241 | 0.7170523 | 0.678145 | 0.2940218 | 0.783427 | 0.01055545 | 0.282563 | 0.6517664 |
| ENSG00000174547.13 | MRPL11 | 0.333717 | 0.9051519 | -0.242524 | 0.7317746 | -1.014999 | 2.35E-05 | -0.262364 | 0.09844449 | -1.757887 | 0.02749219 | 1.05189 | 0.6885065 |
| ENSG00000174652.17 | ZNF266 | -2.131191 | 0.6214382 | 0.668243 | 0.8647637 | 0.434274 | 0.4960197 | 3.143777 | 0.1231799 | 0.398369 | 0.04654734 | -0.907468 | 0.7764933 |
| ENSG00000174672.15 | BRSK2 | 1.362174 | 0.714717 | -0.033171 | 1 | -0.242599 | 0.742077 | 0.886697 | 6.79E-05 | 1.879646 | 9.07E-06 | -0.999151 | 0.8479431 |
| ENSG00000174695.9 | TMEM167A | -0.370787 | 0.9047689 | 0.41214 | 0.4487482 | 0.05836 | 0.4575892 | 0.179558 | 0.4919368 | -0.474632 | 0.000219032 | 0.154138 | 0.6885065 |
| ENSG00000174720.15 | LARP7 | -0.81937 | 0.1236965 | 0.220349 | 0.7383998 | 0.303498 | 0.003251482 | 0.148401 | 0.1654413 | 0.107231 | 0.08811391 | -0.416545 | 0.4304069 |
| ENSG00000174748.18 | RPL15 | -0.035369 | 0.3787871 | 0.139084 | 0.5953732 | 0.003547 | 0.6675523 | 0.064843 | 0.02389999 | 0.228576 | 0.5948759 | -0.00746 | 0.3565625 |
| ENSG00000174780.15 | SRP72 | -0.006526 | 0.4406957 | -0.075846 | 0.7955905 | 0.960751 | 8.77E-07 | 0.853153 | 0.8126451 | -1.191948 | 2.03E-06 | -1.211822 | 0.5344421 |
| ENSG00000174840.8 | PDE12 | -0.241904 | 0.8867323 | -0.845297 | 0.5544987 | 0.859961 | 0.03010539 | 1.790684 | 0.2092092 | -0.72492 | 0.5196843 | 1.931543 | 0.8763843 |
| ENSG00000174851.14 | YIF1A | -0.411524 | 0.9779807 | 0.607983 | 0.2670385 | -0.580239 | 0.000320495 | -0.075071 | 0.05147036 | -0.311177 | 0.00564434 | 0.241453 | 0.7880595 |
| ENSG00000174871.10 | CNIH2 | -0.378072 | 0.8867323 | 0.883003 | 0.4901086 | -1.129898 | 0.6617607 | 1.035028 | 0.00581152 | 2.469892 | 0.000632925 | -0.759974 | 0.6490419 |
| ENSG00000174886.12 | NDUFA11 | -0.443654 | 0.714717 | 0.161036 | 0.7711547 | -0.324415 | 9.53E-07 | -0.217933 | 0.09206469 | -0.387447 | 0.000279287 | -0.060872 | 0.6829816 |
| ENSG00000174891.12 | RSRC1 | -1.365535 | 0.2506697 | 0.49062 | 0.949642 | 0.450242 | 0.001612394 | -0.654338 | 0.006652577 | -0.694593 | 0.001277525 | -0.576046 | 0.6829816 |
| ENSG00000174917.8 | C19orf70 | -0.293862 | 0.6777991 | -0.417758 | 0.01322982 | -0.936451 | 2.73E-06 | 0.139668 | 0.0028842 | -1.31728 | 0.01955383 | 0.438906 | 0.7596076 |
| ENSG00000174939.10 | ASPHD1 | -0.571724 | 0.4874319 | -0.060968 | 0.9403381 | -0.502708 | 0.006698259 | 1.170921 | 0.04385686 | 1.072824 | 0.9854824 | 1.926494 | 0.6885065 |
| ENSG00000174943.9 | KCTD13 | 0.700953 | 0.59287 | 1.257683 | 0.6527736 | -0.271127 | 0.04854632 | 0.424082 | 0.1782976 | 0.907312 | 0.7573677 | -0.284691 | 0.9581586 |
| ENSG00000174953.13 | DHX36 | -0.409484 | 0.6214382 | 0.145551 | 0.937266 | 0.688997 | 2.44E-10 | 0.223326 | 0.7407444 | 0.079127 | 0.00858852 | -1.53118 | 0.6885065 |
| ENSG00000174996.11 | KLC2 | 0.715929 | 0.6214382 | -0.117545 | 0.9709313 | -0.7515 | 0.01586198 | -0.804672 | 0.6079649 | 0.225067 | 0.9690603 | -1.117653 | 0.9562356 |
| ENSG00000175040.5 | CHST2 | 0.84304 | 0.8228711 | -0.262026 | 0.5296018 | -0.51128 | 0.8220688 | 1.861134 | 0.01184951 | 1.84284 | 0.5452636 | -0.148946 | 0.6906533 |
| ENSG00000175073.7 | VCPIP1 | -4.550667 | 0.02807604 | -0.501802 | 0.5205034 | 0.041413 | 0.2684183 | 1.639774 | 0.04207541 | -0.504325 | 0.8888294 | -0.009343 | 0.957527 |
| ENSG00000175104.14 | TRAF6 | -0.400355 | 0.8006286 | 1.26333 | 1 | 0.894838 | 0.02825523 | 2.137977 | 0.3467508 | 0.645522 | 0.7934534 | 8.606435 | 0.2768456 |
| ENSG00000175105.5 | ZNF654 | 0.209621 | 0.7736255 | -0.581592 | 0.2038521 | 2.094285 | 0.001184957 | 2.504369 | 0.1266311 | 1.471415 | 0.5196843 | -2.734164 | 0.7950024 |
| ENSG00000175115.11 | PACS1 | 0.25524 | 0.7401773 | -0.34712 | 0.7321384 | -0.410793 | 0.5290845 | 0.312543 | 0.020826 | 0.673594 | 0.1821909 | -0.050588 | 0.6835249 |
| ENSG00000175130.6 | MARCKSL1 | -0.211572 | 0.8006286 | -0.258884 | 0.2758583 | -0.374878 | 8.73E-12 | 0.828664 | 0.000500492 | 0.91883 | 1.36E-05 | -0.99923 | 0.007627081 |
| ENSG00000175166.16 | PSMD2 | 0.334562 | 0.6214382 | -0.207991 | 0.6527736 | 0.134318 | 0.5814611 | 0.750535 | 1.54E-06 | 1.164381 | 0.008242888 | 0.190305 | 0.5653062 |
| ENSG00000175182.13 | FAM131A | -1.229188 | 0.4383535 | 0.984497 | 0.5953732 | 1.24405 | 1 | 0.242017 | 0.8224647 | 2.135957 | 0.01177698 | -1.580578 | 0.6885065 |
| ENSG00000175183.9 | CSRP2 | -1.446273 | 0.6214382 | 0.3543 | 0.5239144 | 0.265648 | 0.6735793 | -1.313956 | 0.000312917 | 1.619007 | 0.5668236 | 0.54289 | 0.9016395 |
| ENSG00000175197.10 | DDIT3 | -0.281037 | 0.6014343 | 0.570523 | 0.01593197 | -0.002997 | 1 | 0.278437 | 0.7967812 | -0.160311 | 0.01717712 | 2.600962 | 0.5202568 |
| ENSG00000175216.14 | CKAP5 | -1.052106 | 0.4634314 | -0.124558 | 0.5953732 | 0.471366 | 1.07E-05 | 0.120962 | 0.886985 | 1.47575 | 0.02444442 | -0.585807 | 0.9153161 |
| ENSG00000175221.14 | MED16 | -0.170282 | 0.8006286 | 0.040063 | 0.9377864 | -0.156663 | 0.5290845 | 2.006098 | 0.03175449 | -0.576591 | 0.8963489 | 0.609886 | 0.9264343 |
| ENSG00000175224.16 | ATG13 | 1.954327 | 0.714717 | 0.158344 | 0.8703522 | 1.676185 | 0.04693636 | 1.234982 | 0.3748688 | 0.626324 | 0.3890863 | 0.498287 | 0.6885065 |
| ENSG00000175265.17 | GOLGA8A | -1.866832 | 0.4764974 | 0.605324 | 0.6107341 | 0.10109 | 0.7394795 | 0.989968 | 0.0028842 | 0.712518 | 0.356568 | -0.251598 | 0.9094056 |
| ENSG00000175274.18 | TP53I11 | -2.134731 | 0.01118076 | 0.297686 | 0.9790409 | -0.56511 | 0.5497912 | 0.424279 | 0.7144938 | -0.478506 | 0.01828815 | -0.820746 | 0.8479431 |
| ENSG00000175309.14 | PHYKPL | -1.507892 | 0.7800288 | 0.697003 | 0.4886958 | 0.777662 | 0.5905862 | 1.434969 | 0.000459847 | 0.719756 | 0.5196843 | 0.938125 | 0.7901525 |
| ENSG00000175324.9 | LSM1 | -0.369705 | 1 | 0.307045 | 0.9028136 | -0.266081 | 0.02030933 | -0.236134 | 0.3467508 | 0.175802 | 0.4146677 | -0.663697 | 0.5137734 |
| ENSG00000175334.7 | BANF1 | -0.203349 | 1 | 0.086388 | 1 | -0.379574 | 4.80E-07 | -0.263004 | 0.01408544 | -0.318467 | 0.002293693 | -0.743694 | 0.6517664 |
| ENSG00000175387.15 | SMAD2 | 0.701816 | 0.529676 | -0.328596 | 0.5010139 | 0.720668 | 0.3528065 | 0.01066 | 0.3046458 | 1.185122 | 0.03477973 | -0.116123 | 1 |
| ENSG00000175395.15 | ZNF25 | 1.833676 | 0.3254792 | 0.116793 | 0.9213306 | 0.321875 | 0.7437308 | 1.975372 | 0.02882544 | -1.223127 | 0.705576 | -0.480559 | 0.8469357 |
| ENSG00000175414.6 | ARL10 | -1.896347 | 0.4406957 | 0.406816 | 0.5553512 | -0.240129 | 0.5290845 | 0.030521 | 0.0427187 | 0.301287 | 0.165394 | -2.464374 | 0.9500331 |
| ENSG00000175416.12 | CLTB | -0.592076 | 0.5084093 | -0.104723 | 0.6433679 | -0.736229 | 0.000950517 | 0.444561 | 0.6282756 | -0.196987 | 1 | 1.275083 | 1 |
| ENSG00000175445.14 | LPL | 0.976422 | 0.8006286 | 0.01847 | 0.8263112 | 0.576879 | 0.8914851 | -0.136479 | 0.3518093 | -2.852348 | 2.83E-07 | -0.545227 | 0.9581586 |
| ENSG00000175470.18 | PPP2R2D | -0.180147 | 0.8867323 | 0.653084 | 0.6427295 | 0.340451 | 0.3139657 | 3.502504 | 0.00033715 | 0.806341 | 0.3374377 | 0.482929 | 0.8949489 |
| ENSG00000175550.7 | DRAP1 | -0.063204 | 0.6214382 | 0.151143 | 0.9134218 | -0.552648 | 1.19E-14 | 0.006495 | 0.1329452 | 0.105777 | 0.1592962 | 0.406472 | 0.6829816 |
| ENSG00000175573.6 | C11orf68 | -2.512219 | 1 | 0.11291 | 0.8870041 | -1.193299 | 0.02690462 | 0.427765 | 0.2887261 | 2.187803 | 0.5196843 | 1.751749 | 0.8230591 |
| ENSG00000175582.19 | RAB6A | -0.155005 | 0.8867323 | -0.187607 | 0.9641349 | 0.184306 | 0.8436887 | 1.231294 | 0.001996903 | 1.086509 | 0.2187443 | 2.172746 | 0.6185816 |
| ENSG00000175595.14 | ERCC4 | 1.376911 | 0.6214382 | -0.33119 | 0.6527736 | 0.076265 | 0.5290845 | 2.769032 | 5.41E-05 | 1.003471 | 0.6217705 | 0.149211 | 0.8724921 |
| ENSG00000175602.3 | CCDC85B | -0.395284 | 0.2182308 | 0.165627 | 0.706677 | -2.141219 | 3.66E-14 | 0.50179 | 0.8663822 | 1.76986 | 0.9045635 | 0.485514 | 1 |
| ENSG00000175606.10 | TMEM70 | -0.53437 | 0.6214382 | 0.070584 | 0.8953649 | -0.318723 | 0.9304971 | -0.284915 | 0.2043429 | 0.354665 | 0.001346348 | -0.629274 | 0.4306015 |
| ENSG00000175701.10 | LINC00116 | -0.225391 | 0.563505 | 0.169291 | 0.9989119 | -1.090711 | 7.23E-08 | -0.692412 | 5.41E-05 | -0.409939 | 0.4374622 | 0.834529 | 1 |
| ENSG00000175727.13 | MLXIP | -0.073298 | 1 | -0.223495 | 0.7553826 | 0.899431 | 0.999519 | 1.608432 | 0.000107574 | 2.74407 | 0.009713756 | -0.703842 | 0.6885065 |
| ENSG00000175745.11 | NR2F1 | -2.172835 | 0.6172955 | -0.08772 | 0.3216261 | -0.138605 | 0.9088563 | 0.12683 | 0.004953157 | 0.735885 | 0.2427185 | 1.802238 | 0.9011704 |
| ENSG00000175756.13 | AURKAIP1 | -0.338311 | 0.7686507 | 0.402743 | 0.1129216 | -0.624028 | 1.99E-10 | 0.262794 | 0.4134284 | -0.130472 | 1 | -0.938141 | 0.8479431 |
| ENSG00000175768.12 | TOMM5 | 1.968734 | 0.06475284 | 0.79451 | 0.8263112 | -1.185367 | 2.94E-10 | 0.577992 | 0.5325892 | 1.209025 | 0.5196843 | -1.362416 | 0.8479431 |
| ENSG00000175782.9 | SLC35E3 | -0.962552 | 0.7098524 | 0.993157 | 0.377938 | 0.112528 | 0.5518142 | 0.844222 | 0.1653601 | 3.278447 | 0.000278079 | 2.834789 | 0.9589318 |
| ENSG00000175792.11 | RUVBL1 | -1.11043 | 0.6014343 | 0.068736 | 0.2779213 | 0.048722 | 0.4164245 | -0.299209 | 0.006162612 | -0.512792 | 0.008579021 | 0.15516 | 0.278077 |
| ENSG00000175826.11 | CTDNEP1 | 0.231387 | 0.6432291 | -0.486294 | 0.4011918 | -0.663351 | 0.001395263 | 0.802164 | 0.1123449 | 0.616935 | 0.08838444 | -0.304184 | 0.6369988 |
| ENSG00000175854.11 | SWI5 | -0.368271 | 0.7709851 | 0.25932 | 0.9478771 | -0.669061 | 0.003892601 | -0.569022 | 0.04081856 | -0.591248 | 0.1920753 | -1.988723 | 0.863256 |
| ENSG00000175895.3 | PLEKHF2 | 1.658469 | 0.563505 | -1.225416 | 0.190163 | 1.68181 | 0.4580147 | 1.873654 | 0.1397302 | 2.782306 | 0.02728895 | -4.001731 | 0.6885065 |
| ENSG00000175931.12 | UBE2O | -0.9158 | 0.8006286 | -0.71822 | 0.453951 | -0.174669 | 0.8454005 | 0.923851 | 0.001827364 | 1.203041 | 0.004246986 | -1.189811 | 0.6885065 |
| ENSG00000176014.12 | TUBB6 | -1.988008 | 0.8006286 | 0.357348 | 0.749091 | -0.404457 | 0.005012622 | 0.575802 | 0.2869867 | 1.307236 | 0.6226215 | 0.085651 | 0.9944282 |
| ENSG00000176022.4 | B3GALT6 | -1.093138 | 0.529676 | -0.606864 | 0.468656 | -0.553157 | 0.9681982 | 0.590337 | 0.03495703 | -0.632778 | 0.7961103 | 2.141791 | 0.6885065 |
| ENSG00000176046.8 | NUPR1 | -1.557079 | 0.6014343 | 0.666108 | 0.1210906 | -0.866414 | 2.92E-05 | 0.437894 | 0.8009899 | 0.166326 | 0.6566095 | 1.75371 | 0.501643 |
| ENSG00000176058.11 | TPRN | -0.798578 | 0.714717 | 0.613865 | 0.7057076 | -1.605489 | 0.1708509 | 2.445693 | 0.000326842 | 3.833813 | 0.009646741 | 1.827638 | 0.6829816 |
| ENSG00000176095.11 | IP6K1 | -0.781352 | 0.6214382 | 0.097707 | 0.8238638 | -0.058718 | 0.2128224 | 0.801792 | 0.007639344 | 0.266738 | 0.864108 | 0.234751 | 0.7520257 |
| ENSG00000176101.11 | SSNA1 | 1.096136 | 0.7545846 | 0.273677 | 0.8243157 | -0.401796 | 0.001723117 | 0.380971 | 0.7445881 | 0.061844 | 0.5145436 | 1.153716 | 0.8479431 |
| ENSG00000176102.11 | CSTF3 | 0.395951 | 0.6214382 | -0.207135 | 0.6015027 | 0.670739 | 0.01909842 | 0.103312 | 0.4919368 | 0.324582 | 0.7958526 | 0.651826 | 0.5733051 |
| ENSG00000176105.13 | YES1 | 0.410744 | 0.6014343 | 0.208109 | 0.9449715 | 2.011113 | 0.000436208 | -0.324153 | 0.4647402 | 0.876798 | 1 | 0.58742 | 0.7973567 |
| ENSG00000176170.13 | SPHK1 | -1.041177 | 0.529676 | -0.208699 | 0.8360642 | -1.073923 | 0.02228582 | -0.517279 | 0.05603949 | 3.555232 | 0.8728425 | 2.962796 | 0.6829816 |
| ENSG00000176171.11 | BNIP3 | 0.242582 | 0.529676 | 0.863998 | 3.02E-05 | -0.271421 | 0.4064599 | -0.278957 | 0.000195994 | -0.712924 | 0.000813403 | -0.084839 | 1 |
| ENSG00000176209.11 | SMIM19 | 0.207883 | 0.9005211 | 0.161893 | 0.5338232 | -0.732237 | 0.003437671 | -0.237392 | 0.1216426 | 0.265189 | 0.1641325 | -1.023773 | 0.8676378 |
| ENSG00000176248.8 | ANAPC2 | -0.254808 | 0.9289591 | 1.611168 | 0.5233192 | -1.717956 | 0.04944273 | 2.61882 | 0.1093889 | 1.106319 | 0.2682211 | -2.119229 | 0.8009135 |
| ENSG00000176261.15 | ZBTB8OS | -5.323241 | 0.6214382 | 0.880244 | 0.5774973 | -0.480582 | 0.000125894 | -0.278413 | 0.01868866 | 0.182728 | 0.5196843 | 0.640076 | 0.6829816 |
| ENSG00000176293.19 | ZNF135 | 0.704007 | 1 | 0.169942 | 1 | 0.136402 | 1 | 2.084716 | 0.3760083 | 5.371757 | 0.0203964 | -1.11387 | 0.9147227 |
| ENSG00000176340.3 | COX8A | -0.027949 | 1 | -0.266718 | 0.1504499 | -0.418653 | 5.74E-08 | -0.247792 | 1.55E-05 | -0.116747 | 0.004919348 | -0.146751 | 0.2263475 |
| ENSG00000176386.8 | CDC26 | 0.077967 | 1 | 0.317911 | 0.6527736 | 0.2217 | 0.313385 | -0.40019 | 0.02485579 | -1.288755 | 0.113404 | 0.110168 | 0.7670687 |
| ENSG00000176444.18 | CLK2 | 2.04603 | 0.8006286 | 0.440787 | 0.8217443 | -0.841051 | 0.3276184 | 1.726144 | 0.02859072 | 1.259875 | 0.2632425 | 1.788203 | 0.9713109 |
| ENSG00000176476.8 | SGF29 | 0.79364 | 0.714717 | 0.450039 | 0.8263112 | 0.501099 | 1 | -0.486978 | 0.04408235 | 0.287894 | 0.3056603 | 6.052603 | 0.8479431 |
| ENSG00000176619.10 | LMNB2 | 0.318039 | 0.9575547 | -0.644898 | 0.1015619 | 0.015034 | 1 | 0.771082 | 0.2930837 | 1.763479 | 0.02662532 | 0.19434 | 0.9029282 |
| ENSG00000176624.10 | MEX3C | -0.296954 | 0.8006286 | -0.054941 | 0.8703522 | 0.224728 | 0.827962 | 0.320544 | 0.4582864 | 2.114507 | 0.001112557 | 1.585632 | 0.8637809 |
| ENSG00000176731.11 | C8orf59 | -0.006412 | 1 | 0.521019 | 0.1943572 | -0.134888 | 0.000680459 | -0.331376 | 0.000628815 | -0.358885 | 0.5196843 | 1.1811 | 0.6836609 |
| ENSG00000176749.8 | CDK5R1 | -0.643388 | 0.8006286 | -1.78779 | 0.1907035 | 0.275969 | 0.6608587 | 2.116023 | 8.62E-05 | 0.80289 | 0.08266705 | -0.357686 | 0.9653202 |
| ENSG00000176788.8 | BASP1 | -0.362702 | 0.8587992 | 0.254921 | 0.9217775 | -0.362089 | 0.1108855 | 0.863431 | 0.03328858 | 1.443178 | 4.60E-10 | 0.081256 | 0.8171275 |
| ENSG00000176871.8 | WSB2 | 0.585548 | 0.5711449 | -0.091555 | 0.2945701 | -0.555974 | 0.6720169 | 0.590651 | 0.03886295 | 0.131797 | 0.897758 | 0.646962 | 0.8949489 |
| ENSG00000176887.6 | SOX11 | 0.429371 | 0.3439898 | -0.393878 | 0.2707674 | 0.695724 | 0.001193444 | 1.858772 | 4.18E-27 | 0.729547 | 1.96E-05 | -0.547775 | 0.01138072 |
| ENSG00000176890.15 | TYMS | 0.354427 | 0.9478876 | -0.657247 | 0.2284406 | -1.403241 | 0.001988877 | -0.178178 | 0.765156 | 1.698709 | 0.006997413 | 0.259049 | 0.3960489 |
| ENSG00000176894.9 | PXMP2 | 0.470521 | 0.714717 | 0.351026 | 0.9859883 | -2.031642 | 0.000125282 | -1.081246 | 0.3373026 | 1.639522 | 0.8280888 | -2.489679 | 1 |
| ENSG00000176907.4 | C8orf4 | -0.311217 | 0.529676 | 0.088619 | 0.8703522 | -0.315378 | 0.6383858 | -1.145149 | 4.93E-05 | 1.357635 | 0.5810521 | -2.953936 | 0.6829816 |
| ENSG00000176915.14 | ANKLE2 | -0.766267 | 0.6214382 | 0.378035 | 0.8653057 | -0.191547 | 0.5194764 | 1.162756 | 0.002637549 | 1.581288 | 0.02746989 | 0.792296 | 0.8421466 |
| ENSG00000176973.7 | FAM89B | -1.05001 | 0.8006286 | -0.163551 | 0.9140014 | -0.91354 | 0.5697005 | 1.475235 | 0.01151071 | 2.415374 | 1.79E-05 | 0.094814 | 0.6885065 |
| ENSG00000176978.13 | DPP7 | -0.556844 | 0.6362998 | 0.277973 | 0.9839443 | -0.868582 | 8.43E-05 | 1.571958 | 4.81E-09 | 1.583625 | 0.4232345 | -1.082303 | 0.3565625 |
| ENSG00000176994.10 | SMCR8 | -2.835327 | 0.4764974 | 0.054463 | 0.8810704 | 1.63025 | 0.02000886 | -0.698134 | 0.931053 | 0.988521 | 0.9202782 | 2.993124 | 0.8910334 |
| ENSG00000177034.14 | MTX3 | -2.080393 | 0.529676 | 0.321031 | 0.9846935 | 1.091126 | 0.00210624 | 2.027972 | 0.002760806 | 0.771011 | 0.5849259 | -0.304819 | 0.9627406 |
| ENSG00000177051.5 | FBXO46 | 2.138353 | 0.4764974 | 1.12933 | 0.6527736 | -1.571892 | 0.4705492 | 2.478314 | 0.02076542 | 0.783742 | 0.3831662 | -2.394412 | 1 |
| ENSG00000177058.11 | SLC38A9 | 2.674931 | 0.8006286 | 1.949991 | 0.6527736 | -0.696576 | 0.8519804 | 2.543678 | 0.04363231 | 0.672582 | 0.7080077 | 3.015341 | 0.7767294 |
| ENSG00000177108.5 | ZDHHC22 | -0.427919 | 0.6214382 | -2.451959 | 0.911101 | 1.152951 | 0.7643073 | 2.001242 | 0.04043331 | 0.338176 | 0.8374019 | -2.984445 | 0.8777121 |
| ENSG00000177156.10 | TALDO1 | 0.18513 | 0.5791344 | 0.137496 | 0.7044791 | -0.311549 | 0.000297804 | 0.230605 | 0.8230242 | -0.030766 | 0.1834377 | -0.353271 | 0.6185816 |
| ENSG00000177169.9 | ULK1 | -0.030645 | 0.8587992 | 1.251967 | 0.5953732 | 0.609333 | 0.2902538 | 2.587 | 0.06852575 | 0.955924 | 0.03827613 | 2.418617 | 0.4771551 |
| ENSG00000177189.12 | RPS6KA3 | -0.830916 | 0.901215 | -0.094424 | 0.8217443 | 0.862044 | 0.233483 | 1.04292 | 1.44E-05 | 1.982662 | 0.1130762 | 1.678013 | 0.6885065 |
| ENSG00000177192.13 | PUS1 | -2.486934 | 0.09749406 | -0.928009 | 0.3606957 | -1.124571 | 7.39E-05 | 0.091496 | 0.4919368 | 0.377197 | 0.5196843 | -0.590806 | 0.9161136 |
| ENSG00000177200.15 | CHD9 | -2.903372 | 0.2765744 | 0.269163 | 0.4242684 | 1.110461 | 1.27E-09 | 1.265963 | 0.000270591 | 0.864422 | 0.09288315 | -0.385147 | 0.6885065 |
| ENSG00000177302.14 | TOP3A | -1.969214 | 0.6214382 | 0.408131 | 0.7801026 | -0.164347 | 0.7974574 | 1.95143 | 0.000376119 | -0.913069 | 0.5196843 | -0.015828 | 0.6885065 |
| ENSG00000177352.9 | CCDC71 | 0.133669 | 1 | 0.338852 | 0.8873752 | -1.46864 | 0.007919787 | 2.31768 | 0.2335397 | 0.449868 | 0.9104863 | -1.835593 | 0.6185816 |
| ENSG00000177380.13 | PPFIA3 | 3.169496 | 0.6214382 | 1.77315 | 0.2570642 | 0.048293 | 0.8454242 | 0.981702 | 0.003110801 | 2.326612 | 0.02779405 | -0.553282 | 0.9476473 |
| ENSG00000177425.10 | PAWR | 1.613442 | 0.7736255 | 0.234145 | 0.704767 | -4.368284 | 0.03873586 | 2.019323 | 0.08268254 | 0.533558 | 0.9201079 | -5.02449 | 0.7767294 |
| ENSG00000177479.19 | ARIH2 | -0.71911 | 0.2019999 | 0.141488 | 0.7432204 | 0.730143 | 4.45E-08 | 0.01855 | 0.5209719 | 0.833107 | 0.9345111 | -1.511365 | 0.3469679 |
| ENSG00000177485.6 | ZBTB33 | 0.134684 | 1 | 0.427124 | 0.7181571 | 0.989438 | 0.008092781 | 2.204811 | 0.0522366 | 2.691838 | 0.01565028 | -0.741511 | 0.7040119 |
| ENSG00000177556.11 | ATOX1 | -0.663445 | 0.7574394 | -0.514749 | 0.2666086 | -0.516586 | 1.47E-06 | -0.175574 | 0.02205499 | -0.89551 | 0.00139047 | 0.358035 | 1 |
| ENSG00000177565.15 | TBL1XR1 | -0.970549 | 0.8665937 | -0.258945 | 1 | 0.83782 | 0.04792947 | 1.256799 | 0.04680542 | 0.878205 | 0.3602046 | -0.318715 | 0.9687984 |
| ENSG00000177600.8 | RPLP2 | -0.304213 | 0.6014343 | -0.091612 | 0.2731087 | -0.160604 | 0.000654521 | 0.214155 | 0.6530221 | -0.075529 | 0.008782219 | 0.285659 | 0.9526606 |
| ENSG00000177606.6 | JUN | -0.14553 | 0.3772759 | 0.17865 | 0.04122189 | -0.312576 | 6.54E-09 | 0.36833 | 0.6338904 | 0.918218 | 0.03383784 | -0.384302 | 0.6885065 |
| ENSG00000177614.9 | PGBD5 | -1.421373 | 0.3166974 | 1.79584 | 0.9222921 | -0.18079 | 0.4064599 | 1.255369 | 0.04055961 | 1.0668 | 0.01149382 | -0.786083 | 0.6885065 |
| ENSG00000177666.15 | PNPLA2 | -0.168007 | 0.8006286 | 0.559121 | 0.8263112 | -2.052896 | 0.000133971 | 0.350777 | 0.791347 | 0.243807 | 0.5886882 | 2.006594 | 0.6530179 |
| ENSG00000177683.13 | THAP5 | 0.619026 | 0.6014343 | -0.039363 | 1 | 0.622883 | 0.001268929 | 1.101222 | 0.4919368 | -1.039171 | 0.7377262 | 2.15013 | 0.8479431 |
| ENSG00000177697.17 | CD151 | -0.31171 | 0.8474876 | 0.308711 | 0.6527736 | -0.15847 | 0.5290845 | 0.366481 | 0.1202644 | -0.124496 | 0.0463146 | 0.131803 | 0.8479431 |
| ENSG00000177700.5 | POLR2L | 0.471766 | 0.3966394 | -0.024866 | 0.5553512 | -0.841759 | 4.85E-26 | 0.275214 | 0.7349809 | -0.081155 | 0.000178438 | -0.261412 | 0.0739526 |
| ENSG00000177706.8 | FAM20C | -0.759587 | 0.7996471 | 0.382362 | 0.6527736 | -2.04345 | 7.35E-05 | 0.478733 | 0.01230964 | 0.670384 | 0.3817941 | -0.025698 | 0.9270789 |
| ENSG00000177732.7 | SOX12 | -1.459692 | 0.6799027 | -0.262165 | 0.5010139 | -0.383279 | 0.5290845 | 1.902137 | 0.00015851 | 1.896205 | 2.37E-05 | 0.792506 | 0.9851284 |
| ENSG00000177733.6 | HNRNPA0 | 0.114952 | 0.8006286 | 0.142904 | 0.8263112 | -0.245415 | 0.02389585 | 0.760377 | 0.000965756 | 1.246633 | 0.0127696 | -0.174327 | 0.6829816 |
| ENSG00000177853.14 | ZNF518A | -1.968305 | 0.8867323 | 0.3459 | 0.6807583 | -0.871747 | 0.2348326 | 2.457367 | 0.007146934 | -1.322943 | 0.5196843 | -1.120636 | 0.8479431 |
| ENSG00000177885.13 | GRB2 | -0.657771 | 1 | 0.019788 | 0.9994369 | -0.191469 | 0.5290845 | -0.126889 | 0.1783688 | 0.098763 | 0.0426606 | -0.365817 | 0.6885065 |
| ENSG00000177954.11 | RPS27 | -2.42287 | 3.17E-20 | 0.275086 | 0.5949213 | 0.611005 | 2.33E-20 | -0.165612 | 0.00118511 | -0.186526 | 0.000114109 | -0.021187 | 0.5344421 |
| ENSG00000177971.8 | IMP3 | -0.71641 | 0.260849 | -0.173512 | 0.7211571 | -0.57873 | 0.000587267 | 0.310395 | 0.9480626 | 0.523268 | 0.6107287 | -0.402754 | 0.88512 |
| ENSG00000178035.11 | IMPDH2 | -0.139684 | 0.4392896 | -0.287927 | 0.03414591 | 0.295101 | 0.7419117 | 0.125308 | 0.1482986 | 0.736767 | 0.5381878 | -1.094232 | 0.6885065 |
| ENSG00000178053.17 | MLF1 | -0.72908 | 0.8006286 | 0.46177 | 0.7248185 | -0.432764 | 0.8357355 | -0.619406 | 0.04351982 | 0.292705 | 0.819204 | -0.250214 | 0.9100805 |
| ENSG00000178057.14 | NDUFAF3 | -0.521792 | 0.7856294 | -0.72442 | 0.05593321 | -1.272965 | 1.43E-05 | -0.340112 | 0.000186489 | 0.273609 | 0.1406207 | 0.527124 | 0.9147227 |
| ENSG00000178096.8 | BOLA1 | -0.394819 | 1 | 0.463069 | 0.7431188 | -0.972288 | 0.003738609 | -0.790105 | 0.1397979 | 2.06715 | 0.5196843 | 2.839337 | 0.9593131 |
| ENSG00000178104.19 | PDE4DIP | -0.267816 | 0.8006286 | -0.333704 | 0.6527736 | -0.452066 | 0.5290845 | 1.021011 | 0.05689288 | 0.509991 | 0.00056255 | 0.018967 | 0.8033485 |
| ENSG00000178127.12 | NDUFV2 | 0.001016 | 0.6860016 | 1.075415 | 0.1021383 | -0.917811 | 5.58E-09 | 0.556328 | 0.2797675 | -0.433085 | 0.2439714 | 1.521576 | 0.9777201 |
| ENSG00000178150.8 | ZNF114 | 2.292769 | 0.6214382 | -1.553099 | 0.4956727 | -0.841049 | 1 | -0.864257 | 0.6866205 | 2.455623 | 0.003375492 | 0.698151 | 0.826754 |
| ENSG00000178177.14 | LCORL | -1.338572 | 0.7991968 | 0.522694 | 1 | 1.454089 | 0.01290545 | 0.341991 | 0.8180347 | 0.986861 | 0.7728537 | -0.147684 | 0.7372094 |
| ENSG00000178188.14 | SH2B1 | 0.062442 | 0.714717 | 0.00122 | 1 | -0.139468 | 0.877922 | 1.441637 | 0.4410502 | 2.851747 | 0.01472525 | -1.387861 | 0.6885065 |
| ENSG00000178229.7 | ZNF543 | 1.625403 | 0.6214382 | 1.227053 | 0.6527736 | 3.120085 | 0.003012856 | 2.789335 | 0.3467508 | 1.382029 | 1 | -1.080779 | 0.9147227 |
| ENSG00000178252.17 | WDR6 | -0.123268 | 0.6214382 | -0.091119 | 1 | -0.544257 | 0.4264136 | 0.725364 | 0.03646961 | -0.058 | 0.8021966 | 0.109673 | 0.7643634 |
| ENSG00000178295.14 | GEN1 | -0.013177 | 0.8587992 | 1.660826 | 0.8263112 | -0.15952 | 0.09717325 | 1.810622 | 0.000107574 | 0.861234 | 0.2645742 | 1.794633 | 0.7767294 |
| ENSG00000178381.11 | ZFAND2A | 0.347119 | 0.8748952 | -0.097515 | 1 | -0.116477 | 0.04364931 | 0.558523 | 0.4919368 | 0.500709 | 0.2825999 | -1.636291 | 0.7767294 |
| ENSG00000178397.12 | FAM220A | 1.067189 | 0.822435 | 0.370931 | 0.9906117 | 0.719667 | 0.2289298 | 3.328846 | 0.01841683 | 0.758321 | 0.3516806 | -0.206318 | 0.8144829 |
| ENSG00000178449.8 | COX14 | 0.305132 | 0.906197 | -0.381742 | 0.3682639 | -0.329157 | 0.007313234 | 0.309807 | 0.2209365 | 0.230353 | 0.08151074 | 0.998963 | 0.9131264 |
| ENSG00000178531.5 | CTXN1 | -0.928104 | 0.6214382 | 0.50068 | 0.6527736 | -0.847345 | 0.1287908 | 0.96799 | 0.04778811 | 0.608866 | 0.02827891 | -0.977082 | 0.6140824 |
| ENSG00000178538.9 | CA8 | -0.172458 | 0.6214382 | 1.145654 | 0.9473701 | 1.210387 | 0.4382561 | 3.060583 | 0.861719 | 0.828538 | 0.3379763 |  |  |
| ENSG00000178567.7 | EPM2AIP1 | 0.146672 | 0.9282145 | -0.701347 | 0.4402768 | -0.110841 | 0.6876894 | 1.784732 | 0.04809897 | 2.019656 | 7.60E-05 | -0.354998 | 0.6517664 |
| ENSG00000178585.14 | CTNNBIP1 | 1.507393 | 0.8867323 | -0.414755 | 0.3033464 | 0.025489 | 0.9391584 | 2.383623 | 0.02141464 | 1.242625 | 0.3440157 | -0.620347 | 0.8595618 |
| ENSG00000178695.5 | KCTD12 | -2.337229 | 0.6014343 | -0.43379 | 0.03439805 | 0.133817 | 0.7346746 | 1.396926 | 0.1637266 | 1.312266 | 1 | 1.042814 | 0.9147227 |
| ENSG00000178719.16 | GRINA | -0.501851 | 0.8587992 | 0.473411 | 0.7071958 | -0.593086 | 9.47E-06 | 0.718959 | 0.1097499 | 1.012646 | 0.2063571 | -0.644054 | 0.5331811 |
| ENSG00000178741.11 | COX5A | -0.081547 | 0.9047689 | -0.020875 | 0.5228465 | -0.588799 | 1.23E-08 | 0.318435 | 0.4608889 | -0.303077 | 0.02891048 | -0.137293 | 0.6885065 |
| ENSG00000178896.6 | EXOSC4 | 0.288123 | 0.7207113 | -0.479558 | 0.5737754 | -0.817499 | 7.12E-05 | -0.229514 | 0.6806715 | -0.869346 | 0.5423956 | -1.171731 | 0.6683212 |
| ENSG00000178904.18 | DPY19L3 | -0.456032 | 0.8006286 | -0.669882 | 0.2731087 | 0.824337 | 0.05245541 | 1.154694 | 0.000459847 | 1.751588 | 0.3974318 | 0.391368 | 1 |
| ENSG00000178913.7 | TAF7 | -0.360364 | 0.9316827 | -0.082515 | 0.6527736 | 0.195415 | 0.0106224 | 0.165587 | 0.8079576 | 0.385449 | 0.6378269 | -0.69261 | 1 |
| ENSG00000178921.13 | PFAS | 0.134512 | 0.6214382 | -0.088523 | 0.8484992 | -0.985711 | 0.5290845 | 0.782802 | 0.03820042 | 2.493305 | 0.7371141 | -1.525788 | 0.6606272 |
| ENSG00000178922.16 | HYI | 0.673256 | 1 | -0.369606 | 0.983232 | -2.880053 | 0.000544099 | 0.559112 | 0.9876125 | 1.259693 | 0.4391732 | -0.808954 | 0.9147227 |
| ENSG00000178951.8 | ZBTB7A | -2.368285 | 0.07216456 | -0.446146 | 0.5953732 | -0.478911 | 0.5130307 | 1.858083 | 9.68E-06 | 1.942205 | 0.01299526 | 1.302354 | 0.9803827 |
| ENSG00000178980.14 | SEPW1 | 0.752944 | 0.08969904 | -0.167409 | 0.2553839 | -0.326821 | 0.0047011 | -0.004587 | 0.0725107 | 0.425528 | 0.681502 | 0.237018 | 0.6885065 |
| ENSG00000178982.9 | EIF3K | 0.494478 | 0.4923792 | 0.019332 | 0.6527736 | -0.114298 | 0.002245673 | 0.20325 | 0.287279 | -0.437227 | 0.06026757 | 0.633774 | 0.9215399 |
| ENSG00000178988.10 | MRFAP1L1 | -1.375161 | 0.02363633 | -0.138781 | 0.6527736 | 0.008913 | 1 | 0.630005 | 0.9221823 | 0.238347 | 0.2228691 | -0.841536 | 0.4650016 |
| ENSG00000179010.14 | MRFAP1 | -0.884392 | 0.000248137 | -0.123833 | 0.1846751 | 0.473293 | 0.5290845 | -0.364692 | 8.88E-05 | 0.693058 | 0.7078772 | 0.155185 | 0.6829816 |
| ENSG00000179021.9 | C3orf38 | 1.828345 | 0.3166974 | 0.274332 | 0.7247706 | 0.879029 | 0.04494191 | 0.920993 | 0.4312634 | 1.463051 | 0.6214624 | -1.30384 | 0.7833507 |
| ENSG00000179041.3 | RRS1 | -1.095391 | 0.1082254 | -0.188001 | 0.6527736 | -0.708481 | 0.01776045 | 0.568517 | 0.01244221 | 0.717248 | 0.5196843 | -0.091712 | 0.2971265 |
| ENSG00000179051.13 | RCC2 | -1.756847 | 0.04993189 | 0.020844 | 0.6562527 | 0.113421 | 0.6538987 | 0.762259 | 2.93E-05 | 1.125728 | 0.008341119 | 0.081153 | 0.6885065 |
| ENSG00000179085.7 | DPM3 | -0.170296 | 0.921979 | 0.070358 | 0.8544073 | -0.551176 | 0.003206137 | 0.602553 | 0.5627564 | 0.165566 | 0.5196843 | -0.137685 | 0.9576568 |
| ENSG00000179091.4 | CYC1 | 0.5671 | 0.03631088 | 0.037606 | 0.6200225 | -1.576219 | 7.64E-30 | -0.382715 | 0.001789677 | -0.368315 | 0.4970703 | -0.350186 | 0.6829816 |
| ENSG00000179104.8 | TMTC2 | 1.387011 | 1 | -0.495809 | 0.5949213 | -0.861916 | 0.9368403 | 1.552085 | 0.03675516 | 2.56185 | 0.5196843 | 0.247389 | 0.6891436 |
| ENSG00000179119.14 | SPTY2D1 | -0.390993 | 0.529676 | -0.442938 | 0.8637961 | 0.863253 | 0.01563975 | 1.592752 | 0.7638599 | 1.144848 | 0.750425 | -1.516947 | 0.920417 |
| ENSG00000179134.14 | SAMD4B | -1.765468 | 0.3166974 | 0.665816 | 0.5520739 | 0.425812 | 0.7643073 | 0.574326 | 0.4701433 | 1.369058 | 0.01238434 | -1.023763 | 0.583558 |
| ENSG00000179195.15 | ZNF664 | 0.667653 | 0.9571464 | -0.07702 | 0.5681739 | 0.689695 | 0.4382561 | 1.769693 | 0.00061113 | 0.423554 | 0.1883232 | -0.322444 | 0.9791298 |
| ENSG00000179222.17 | MAGED1 | -0.263369 | 0.6214382 | -0.458504 | 0.06545638 | 0.413965 | 0.02880594 | 0.315888 | 0.9547789 | -0.305189 | 0.01789147 | -0.743243 | 0.3574139 |
| ENSG00000179241.12 | LDLRAD3 | 0.54301 | 0.6214382 | -0.845805 | 0.04558655 | 0.653537 | 0.4064599 | 0.324922 | 0.6996942 | 0.784824 | 0.1875029 | -0.524049 | 0.7095543 |
| ENSG00000179262.9 | RAD23A | 0.188403 | 0.2335514 | 0.058993 | 0.9486562 | 0.37003 | 0.05678161 | -0.507741 | 0.005138598 | 0.128067 | 0.621723 | 0.679836 | 0.7866135 |
| ENSG00000179271.2 | GADD45GIP1 | -0.029196 | 0.9444548 | -0.053804 | 0.8703522 | -0.642805 | 1.88E-08 | -0.174957 | 0.003104085 | -0.238944 | 0.04233873 | -0.2237 | 0.6185816 |
| ENSG00000179295.16 | PTPN11 | -0.660975 | 0.1487841 | -0.161069 | 0.3805897 | 0.435525 | 0.2533595 | 0.874143 | 0.01033382 | -0.114248 | 0.1945729 | 0.796039 | 0.8479431 |
| ENSG00000179335.18 | CLK3 | 0.781121 | 0.6214382 | -0.404753 | 0.8263112 | -0.739049 | 0.7623935 | -0.1453 | 0.8903193 | 1.903109 | 0.03043237 | 1.759396 | 0.8479431 |
| ENSG00000179388.8 | EGR3 | 1.431014 | 0.3951489 | -0.482184 | 0.5394427 | -0.095448 | 0.9790188 | 2.449533 | 0.03303111 | 1.484103 | 0.8021966 | -1.931465 | 0.8479431 |
| ENSG00000179431.6 | FJX1 | -1.614 | 0.6279301 | -0.669602 | 0.02540926 | -0.108927 | 0.7173462 | 0.550579 | 0.2174487 | -0.078504 | 0.009938442 | 1.633158 | 0.7764132 |
| ENSG00000179456.10 | ZBTB18 | 0.561638 | 0.714717 | -0.408793 | 0.4072962 | 0.827012 | 0.393184 | 1.469537 | 0.0014943 | 0.34054 | 0.04090365 | -0.483244 | 0.8070118 |
| ENSG00000179526.16 | SHARPIN | -0.324323 | 0.8493201 | 0.429585 | 1 | -3.09181 | 0.000455376 | -0.075556 | 0.6575769 | 0.950998 | 0.4603648 | -2.955665 | 0.6829816 |
| ENSG00000179562.2 | GCC1 | 0.671345 | 0.9183025 | -0.045409 | 0.83982 | -1.697738 | 0.2437457 | 2.016711 | 2.01E-07 | 0.665329 | 0.02906571 | -0.275309 | 0.9662158 |
| ENSG00000179598.5 | PLD6 | -0.161909 | 0.8006286 | 0.087935 | 0.9590571 | -2.206554 | 0.03807631 | -0.228262 | 0.4919368 | 0.444805 | 0.7243765 | 1.944938 | 0.9382274 |
| ENSG00000179632.9 | MAF1 | 0.950199 | 0.4764974 | -0.2832 | 0.9084242 | -1.192277 | 4.62E-05 | -1.429917 | 0.4919368 | 0.845337 | 0.1401461 | 1.948391 | 0.8886024 |
| ENSG00000179832.17 | MROH1 | 1.05493 | 1 | 1.320093 | 0.4093377 | -1.7961 | 0.3013477 | 1.605708 | 0.02029956 | 1.111034 | 1 | -0.978987 | 0.7753492 |
| ENSG00000179950.13 | PUF60 | 0.348524 | 0.6014343 | -0.147424 | 0.9104976 | -0.216239 | 0.03540251 | 0.269465 | 0.701149 | 0.153999 | 0.7219826 | -0.184354 | 0.6675976 |
| ENSG00000179979.8 | CRIPAK | 4.248889 | 0.6214382 | 0.471136 | 0.8263112 | 1.096076 | 0.6821321 | 4.475774 | 0.04333874 | 2.729911 | 1 | inf | 0.5344421 |
| ENSG00000180011.6 | ZADH2 | 0.2906 | 0.8006286 | -0.459141 | 0.6527736 | 0.657986 | 0.3266608 | 1.537394 | 0.04684857 | 1.99373 | 0.0003226 | -0.297983 | 0.97804 |
| ENSG00000180104.15 | EXOC3 | -0.189758 | 0.8006902 | -0.113314 | 0.8764311 | -0.591585 | 0.06843866 | 0.467269 | 0.01989019 | 0.569818 | 0.3513722 | 0.872204 | 1 |
| ENSG00000180198.15 | RCC1 | 0.496534 | 0.8867323 | -0.05323 | 0.829342 | -0.214028 | 0.5290845 | 0.870256 | 0.02905006 | 0.6544 | 0.5357356 | 0.521972 | 0.9626957 |
| ENSG00000180354.15 | MTURN | -1.235457 | 0.824666 | -0.161862 | 0.6527736 | 1.772516 | 0.5290845 | 2.216528 | 0.000293083 | 0.659562 | 0.08177415 | -0.001613 | 0.4304069 |
| ENSG00000180357.9 | ZNF609 | -2.158086 | 0.529676 | 0.142682 | 0.6527736 | 0.864344 | 0.005574707 | 1.519615 | 0.07581879 | 2.405646 | 0.0128419 | 1.760415 | 0.8105922 |
| ENSG00000180370.10 | PAK2 | -0.432407 | 0.8041679 | 0.003274 | 0.507133 | 0.467608 | 0.000267405 | 0.84678 | 0.4919368 | -0.114348 | 0.00531443 | -0.077902 | 0.4578379 |
| ENSG00000180389.6 | ATP5EP2 | 1.487615 | 0.2215131 | 0.022991 | 0.8304896 | -2.506657 | 7.48E-42 | 0.929259 | 0.0412511 | -0.507684 | 0.01760693 | -0.8081 | 0.8479431 |
| ENSG00000180398.11 | MCFD2 | 0.172922 | 0.5467816 | 0.685225 | 0.04965752 | 0.100152 | 0.7540344 | 0.445023 | 0.541717 | -0.15411 | 0.0220309 | 0.232479 | 0.9770956 |
| ENSG00000180488.14 | FAM73A | 0.030813 | 0.6792949 | 0.692377 | 0.3225624 | 0.135838 | 0.01887916 | 0.592328 | 0.7439741 | 2.02058 | 0.7468048 | 0.667606 | 0.5689187 |
| ENSG00000180530.9 | NRIP1 | 0.782809 | 0.563505 | 0.173986 | 0.8089431 | 1.249281 | 0.02182793 | 1.094137 | 0.004603223 | 1.045919 | 0.5196843 | 0.520472 | 1 |
| ENSG00000180817.11 | PPA1 | -2.612778 | 0.6214382 | -0.094272 | 0.6527736 | 0.201676 | 0.9000186 | 0.244131 | 0.03055383 | 0.115997 | 0.5261 | -0.707261 | 0.6885065 |
| ENSG00000180822.11 | PSMG4 | -0.68434 | 0.714717 | 0.765191 | 0.6527736 | -2.035058 | 5.63E-05 | -0.628101 | 0.960229 | 0.254205 | 0.8951154 | -0.190697 | 0.8469357 |
| ENSG00000180879.13 | SSR4 | 0.625015 | 0.8006286 | 0.303271 | 0.6527736 | -0.336101 | 2.66E-06 | -0.258407 | 0.1181176 | 0.117145 | 0.1993336 | 0.23524 | 0.6885065 |
| ENSG00000180891.12 | CUEDC1 | 2.663265 | 0.6014343 | -0.569737 | 0.109154 | -1.287382 | 0.1002947 | 0.775422 | 0.1000303 | 1.755146 | 0.00360099 | 1.569921 | 0.7963477 |
| ENSG00000180957.17 | PITPNB | 1.020007 | 0.3946735 | 0.217408 | 1 | 0.835974 | 0.009914828 | 0.476931 | 0.3816997 | 0.810421 | 1 | -0.353929 | 0.8009135 |
| ENSG00000180992.6 | MRPL14 | -0.01097 | 0.8734895 | -0.379522 | 0.1232781 | -0.343399 | 0.000556776 | 0.17829 | 0.7624241 | 0.468036 | 0.4911506 | -0.831312 | 0.06085273 |
| ENSG00000180998.11 | GPR137C | 0.267417 | 1 | 2.181087 | 0.8217443 | 3.907954 | 0.8454005 | 4.869868 | 0.01228639 | 1.455577 | 0.06405667 | inf | 0.5344421 |
| ENSG00000181007.8 | ZFP82 | -1.13646 | 0.6214382 | 0.161961 | 0.8626845 | 0.692676 | 0.8597061 | 1.788989 | 0.002447063 | 1.244803 | 0.1300713 | 1.016548 | 0.6885065 |
| ENSG00000181026.14 | AEN | 0.131413 | 0.9946988 | -0.149062 | 0.8873752 | -0.019844 | 1 | 1.505724 | 0.01146035 | 1.048914 | 0.5196843 | -1.623986 | 0.9370888 |
| ENSG00000181029.8 | TRAPPC5 | -0.466518 | 0.6214382 | 1.638389 | 0.4375617 | 0.244845 | 0.5290845 | -1.656198 | 2.54E-06 | 0.644146 | 0.7771194 | 0.239485 | 0.9181784 |
| ENSG00000181061.13 | HIGD1A | 0.799889 | 0.09713735 | 0.181659 | 0.638704 | -0.065982 | 0.1663608 | -0.032352 | 0.01278236 | -0.725094 | 0.07302247 | -0.376874 | 0.6885065 |
| ENSG00000181104.6 | F2R | -1.02152 | 0.9500536 | -1.104793 | 0.02260499 | -0.131529 | 0.9594981 | -0.058569 | 0.07723471 | 2.114641 | 0.7377262 | -2.443778 | 0.4018772 |
| ENSG00000181163.13 | NPM1 | 0.088697 | 0.6014343 | -0.043829 | 0.5632694 | 0.087112 | 0.1450122 | 1.115304 | 9.64E-28 | 0.783514 | 0.000477112 | -0.374421 | 0.001751751 |
| ENSG00000181220.15 | ZNF746 | -0.118039 | 0.8006286 | 0.203757 | 0.8880028 | -0.217822 | 0.9739697 | 3.419124 | 0.02848393 | 2.483487 | 0.5087334 | -1.12831 | 0.9565287 |
| ENSG00000181315.10 | ZNF322 | 0.513421 | 1 | 0.255938 | 0.9385836 | 0.921702 | 0.005089577 | 2.427985 | 0.005277871 | 0.520076 | 0.1807246 | 1.230061 | 0.6997204 |
| ENSG00000181350.11 | LRRC75A | 0.714545 | 0.529676 | 0.030928 | 1 | 0.14615 | 0.8247492 | 0.850377 | 0.1016278 | -1.215277 | 5.84E-05 | -0.221947 | 0.006637347 |
| ENSG00000181449.3 | SOX2 | 0.760314 | 1.30E-05 | -0.043905 | 0.6413306 | 0.520152 | 4.41E-07 | 0.837221 | 1.03E-05 | 0.813965 | 0.0656862 | 0.782647 | 0.9741465 |
| ENSG00000181555.19 | SETD2 | -0.484807 | 0.6014343 | -0.040136 | 0.9345859 | 1.093024 | 3.81E-06 | 1.625652 | 0.4919368 | 0.736687 | 0.7437196 | 0.915405 | 1 |
| ENSG00000181588.16 | MEX3D | 0.97018 | 0.8006286 | 0.678235 | 0.9854207 | -1.406495 | 0.1962082 | 1.605122 | 2.21E-06 | 0.438464 | 0.001365086 | 1.288928 | 1 |
| ENSG00000181619.11 | GPR135 | 1.034398 | 1 | -0.651977 | 0.8703522 | 0.582353 | 0.7354314 | 0.590266 | 0.3760083 | 0.803693 | 0.7377262 |  |  |
| ENSG00000181649.5 | PHLDA2 | 0.676238 | 0.928247 | 1.577829 | 0.6527736 | -1.479159 | 2.67E-21 | 0.299766 | 0.7554296 | 1.739417 | 0.1845956 | 0.42019 | 0.6885065 |
| ENSG00000181666.17 | HKR1 | -0.274173 | 0.7736255 | -0.220652 | 0.8217443 | 0.639112 | 0.06336058 | 2.249928 | 0.4355714 | 2.975812 | 0.000912166 | 0.599154 | 0.7901525 |
| ENSG00000181722.15 | ZBTB20 | -1.326742 | 0.4947716 | 0.082209 | 0.1583423 | 1.941029 | 1.35E-07 | 0.521366 | 0.006565183 | 0.751586 | 0.1119361 | -1.931812 | 0.6185816 |
| ENSG00000181817.5 | LSM10 | 0.086213 | 0.6962116 | -0.135703 | 1 | 0.036527 | 0.01261436 | -0.327912 | 0.01554383 | 0.025097 | 0.3254391 | -0.456688 | 0.7109418 |
| ENSG00000181827.14 | RFX7 | 0.492691 | 1 | -0.886776 | 0.2770455 | 1.78898 | 0.000129607 | 6.468653 | 0.009420281 | 2.936049 | 0.000890215 | -3.360542 | 0.8249021 |
| ENSG00000181904.8 | C5orf24 | -1.166139 | 0.1956577 | 0.025012 | 0.7972162 | 0.066359 | 0.5290845 | 1.12799 | 1.94E-08 | 1.839007 | 1.21E-05 | -0.529997 | 0.7333942 |
| ENSG00000181915.4 | ADO | -0.203385 | 0.6214382 | 0.713981 | 0.8953649 | 0.419516 | 0.7437308 | 1.70882 | 0.01974122 | 0.563438 | 0.5572347 | 0.149112 | 0.6885065 |
| ENSG00000181991.15 | MRPS11 | -1.110868 | 0.1835496 | -0.285374 | 0.4011918 | 0.154415 | 0.004645586 | 0.3617 | 0.6864296 | 0.494292 | 0.9893414 | 1.841488 | 0.868568 |
| ENSG00000182004.12 | SNRPE | -0.628993 | 0.8006286 | -0.030571 | 0.8689748 | -0.261163 | 2.66E-06 | 0.0202 | 0.3202165 | -0.49776 | 1.92E-05 | -0.543818 | 0.07512959 |
| ENSG00000182013.17 | PNMAL1 | 1.141531 | 0.8006286 | -0.267597 | 0.3059816 | -0.697837 | 0.6827367 | 2.011708 | 4.06E-05 | -0.786787 | 0.4999956 | 2.125759 | 0.6517664 |
| ENSG00000182054.9 | IDH2 | 0.276974 | 0.6274494 | -0.0607 | 1 | -0.446791 | 0.004494928 | -0.057926 | 0.8230242 | -0.522609 | 0.5509506 | 0.76731 | 0.7565328 |
| ENSG00000182087.12 | TMEM259 | -0.165396 | 0.6685525 | 0.173878 | 0.7166727 | -0.303351 | 0.002917154 | 0.531031 | 0.3978915 | 1.143495 | 1.47E-09 | 1.350347 | 0.6885065 |
| ENSG00000182093.14 | WRB | 0.495132 | 0.6214382 | 0.480141 | 0.8593459 | 0.647145 | 0.7250004 | -0.569187 | 0.04053302 | -0.609037 | 0.002548226 | -0.881161 | 0.1570836 |
| ENSG00000182108.9 | DEXI | -0.771765 | 0.9538897 | 0.343631 | 1 | -0.687366 | 0.000495317 | 0.48499 | 0.8728066 | 1.146814 | 0.05238498 | 1.308134 | 0.8479431 |
| ENSG00000182117.5 | NOP10 | 0.894092 | 0.04901151 | 0.014286 | 0.7281537 | -0.111212 | 6.83E-05 | -0.457761 | 0.000342251 | -0.270272 | 5.62E-05 | 0.163348 | 0.8949489 |
| ENSG00000182141.9 | ZNF708 | -0.2889 | 0.8448862 | -0.017915 | 0.9463185 | 1.965072 | 0.000251539 | 0.614005 | 0.5605366 | -0.092521 | 1 | 3.86272 | 1 |
| ENSG00000182149.20 | IST1 | 0.312527 | 0.6214382 | 0.210523 | 0.9973179 | 0.614872 | 0.002635969 | 1.447437 | 0.02018338 | 0.47191 | 0.6332738 | -1.522245 | 0.6829816 |
| ENSG00000182150.15 | ERCC6L2 | -0.405421 | 0.714717 | -0.320418 | 0.2588352 | 1.398996 | 0.001831984 | -0.140926 | 0.6599507 | 0.450758 | 0.7134836 | -4.106911 | 0.9547642 |
| ENSG00000182154.7 | MRPL41 | 1.016167 | 0.000365532 | 0.174111 | 1 | -1.203396 | 1.47E-12 | -0.607982 | 0.1507125 | 0.528294 | 0.4022846 | 0.668 | 0.9565287 |
| ENSG00000182173.12 | TSEN54 | -0.710974 | 0.8867323 | -0.50567 | 0.6527736 | 0.199014 | 0.5018872 | 0.819188 | 0.04434957 | 1.124271 | 0.650378 | -1.02043 | 0.8949489 |
| ENSG00000182247.9 | UBE2E2 | 0.042529 | 0.8587992 | -0.637847 | 0.6274147 | -0.685361 | 0.4674196 | 0.399506 | 0.01751854 | 0.024772 | 0.7377262 | -0.505191 | 1 |
| ENSG00000182263.13 | FIGN | 0.006941 | 1 | -0.052376 | 0.2792759 | 1.023295 | 0.5205494 | 1.029021 | 0.04963161 | 1.090703 | 0.9695178 | 1.375224 | 0.7915993 |
| ENSG00000182287.13 | AP1S2 | 0.296305 | 0.6214382 | -0.106259 | 0.6527736 | -0.057688 | 0.02645855 | -0.061231 | 0.4919368 | 0.375223 | 0.8350514 | -0.02615 | 0.6517664 |
| ENSG00000182307.12 | C8orf33 | -2.160977 | 0.8006286 | 0.079611 | 0.9104976 | 0.176905 | 0.8432413 | 0.176772 | 0.8024673 | -0.813692 | 7.40E-05 | -0.789739 | 0.1570836 |
| ENSG00000182359.14 | KBTBD3 | -0.19849 | 1 | 0.574808 | 0.9420234 | 1.279398 | 0.394998 | -1.373775 | 0.02029956 | -0.601874 | 1 | -1.842048 | 0.9427738 |
| ENSG00000182446.13 | NPLOC4 | 0.196133 | 0.8862706 | 0.021495 | 0.9696336 | 0.023251 | 0.539885 | 0.299975 | 8.35E-05 | 1.75161 | 0.09250305 | 0.524379 | 0.737834 |
| ENSG00000182473.21 | EXOC7 | 0.002445 | 0.8807947 | -0.220779 | 0.5953732 | 0.152622 | 0.2507965 | 0.731328 | 0.02918812 | 0.439178 | 0.7366496 | 0.008333 | 0.9352766 |
| ENSG00000182504.10 | CEP97 | 1.181543 | 0.6014343 | -0.072021 | 0.7566832 | 1.22883 | 0.001932134 | 0.746079 | 0.2595185 | 0.721526 | 0.1591258 | 0.232688 | 0.6885065 |
| ENSG00000182512.4 | GLRX5 | -0.714698 | 0.6501391 | -0.720384 | 0.07401424 | 0.084413 | 0.8330392 | 1.325273 | 9.69E-07 | -0.206001 | 0.1711844 | 0.985683 | 0.7427169 |
| ENSG00000182534.13 | MXRA7 | -0.360677 | 0.9362121 | 0.291087 | 0.7383998 | 0.016458 | 0.5864994 | 0.682937 | 0.01009829 | 1.078894 | 0.7377262 | -0.338607 | 0.6885065 |
| ENSG00000182541.17 | LIMK2 | -2.969456 | 0.6214382 | 0.387565 | 0.592336 | -2.074135 | 0.01540444 | 0.600629 | 0.3609648 | 0.17182 | 0.768601 | -2.705853 | 0.8309897 |
| ENSG00000182544.8 | MFSD5 | 0.846974 | 0.9497414 | 0.672034 | 0.7383998 | -0.690172 | 0.342534 | 0.409099 | 0.04291 | 1.665266 | 1 | -1.577713 | 0.7213346 |
| ENSG00000182568.16 | SATB1 | 0.645302 | 0.9025813 | 0.886795 | 0.4211636 | 1.062944 | 0.000414656 | -0.263234 | 0.1174432 | 0.971144 | 0.01964411 | -2.094065 | 0.6490419 |
| ENSG00000182621.16 | PLCB1 | -0.296132 | 0.6214382 | 0.030215 | 0.592336 | 1.717339 | 7.09E-06 | 0.161103 | 0.2603229 | 0.628966 | 0.9527758 | 0.719991 | 0.7080983 |
| ENSG00000182636.5 | NDN | -0.430631 | 0.6014343 | -4.926239 | 1 | -2.546384 | 0.03807631 | 1.493368 | 0.006469687 | 1.109786 | 0.1374127 | 2.058118 | 0.6885065 |
| ENSG00000182667.14 | NTM | -0.267829 | 1 | -0.308568 | 0.6413306 | 0.157255 | 0.2643875 | 1.352056 | 0.001267967 | -0.534261 | 0.03792058 | -0.431056 | 0.9575366 |
| ENSG00000182670.13 | TTC3 | -0.162426 | 0.8972807 | -0.103023 | 0.3218572 | 0.329422 | 0.001404305 | 1.589037 | 9.51E-20 | 0.359443 | 0.7136911 | -0.017265 | 0.6885065 |
| ENSG00000182700.4 | IGIP | -1.499863 | 0.6214382 | -0.942306 | 0.6527736 | 0.721997 | 0.6406246 | 1.473693 | 0.02326622 | 1.597434 | 0.6206166 | 3.299565 | 0.782216 |
| ENSG00000182718.16 | ANXA2 | -1.172504 | 0.714717 | -0.017875 | 0.8243441 | 0.0437 | 0.06837429 | -0.36383 | 2.76E-06 | -0.293449 | 9.12E-05 | 0.603426 | 0.6885065 |
| ENSG00000182749.5 | PAQR7 | -0.985676 | 0.9289591 | 0.431245 | 1 | -0.444366 | 0.1625971 | 1.788205 | 0.03163649 | -0.733686 | 0.9268123 | -0.154815 | 0.6885065 |
| ENSG00000182774.10 | RPS17 | -0.531452 | 0.0180643 | -0.117604 | 0.5395105 | -0.009786 | 0.814639 | -0.052455 | 0.1219241 | -0.159964 | 0.01007426 | 0.56501 | 0.9539924 |
| ENSG00000182809.10 | CRIP2 | 1.0739 | 0.01796694 | -0.328683 | 0.3160414 | -0.300043 | 0.4923004 | -0.583247 | 0.05157198 | 0.02515 | 0.2415558 | -0.448297 | 0.9264343 |
| ENSG00000182831.11 | C16orf72 | 0.887408 | 0.937674 | -0.032382 | 0.5953732 | -0.876815 | 0.5290845 | 1.300082 | 0.000953523 | 1.454883 | 0.04847197 | 0.385759 | 0.7870938 |
| ENSG00000182871.14 | COL18A1 | 1.055294 | 1 | -1.283069 | 0.0511244 | -3.517834 | 0.8656249 | 0.288182 | 0.004807921 | 3.729194 | 0.08640932 | 0.017778 | 0.6885065 |
| ENSG00000182899.14 | RPL35A | -0.450516 | 0.1634942 | -0.106118 | 0.1940102 | -0.049622 | 0.1929588 | -0.099062 | 0.02205499 | -0.063476 | 0.007331181 | 0.130998 | 0.3887828 |
| ENSG00000182901.15 | RGS7 | -0.183319 | 1 | -1.069009 | 0.6200225 | -1.166794 | 0.6295125 | 4.470337 | 0.3760083 | -1.576231 | 0.7253092 |  |  |
| ENSG00000182944.17 | EWSR1 | -0.405887 | 0.7552987 | 0.187828 | 0.9104976 | 0.257175 | 0.7368441 | 0.483049 | 0.002422189 | 1.125445 | 0.04652304 | -0.197693 | 0.1110081 |
| ENSG00000182979.17 | MTA1 | -0.606801 | 0.4406957 | 0.243005 | 0.8953649 | -1.492185 | 0.00402959 | 1.016842 | 0.02635912 | 1.626921 | 0.2063571 | -2.269935 | 0.9130454 |
| ENSG00000182986.12 | ZNF320 | 2.023375 | 0.8006286 | 0.141872 | 0.9061805 | 0.434178 | 0.5093362 | 1.816386 | 0.04942714 | 1.236404 | 0.131029 | -0.21345 | 0.7901525 |
| ENSG00000183011.13 | NAA38 | -0.494875 | 0.6214382 | -0.523238 | 0.2731087 | -0.884149 | 8.28E-10 | -0.025265 | 0.06999546 | 0.085396 | 0.06763574 | -0.743513 | 0.9354405 |
| ENSG00000183044.11 | ABAT | -1.958198 | 0.8249143 | -0.106074 | 0.8953649 | 0.523201 | 0.5900918 | 1.382562 | 0.1201174 | 1.659037 | 0.005257664 | -1.652381 | 0.9147227 |
| ENSG00000183117.18 | CSMD1 | -1.702386 | 0.6014343 | -0.323343 | 0.8263112 | -1.330898 | 1 | -0.275467 | 0.4722399 | 3.66292 | 0.2342999 |  |  |
| ENSG00000183155.4 | RABIF | 0.199492 | 0.5740533 | 0.387095 | 0.7881421 | -0.810869 | 0.5290845 | 0.812596 | 0.4281238 | -1.340332 | 0.03068444 | 0.772141 | 0.8949489 |
| ENSG00000183172.8 | SMDT1 | -0.84075 | 0.7693197 | -0.152088 | 0.6527736 | -0.764289 | 0.002366518 | -0.097194 | 0.2453102 | 0.026091 | 0.5196843 | -0.819844 | 0.6037129 |
| ENSG00000183207.12 | RUVBL2 | -0.36596 | 1 | 0.37807 | 0.6527736 | -0.551094 | 0.03767129 | 0.176966 | 0.1685514 | -0.369569 | 0.2906522 | 0.130543 | 0.6885065 |
| ENSG00000183248.11 | PRR36 | -2.283307 | 0.563505 | -0.034789 | 0.8703522 | 0.833197 | 0.6295125 | 3.035694 | 0.4127967 | 2.514663 | 0.005747048 | 1.608319 | 0.6829816 |
| ENSG00000183291.15 | 15-Sep | 0.092343 | 0.8859625 | 0.458527 | 0.008676432 | 0.244351 | 1 | -0.184524 | 6.83E-05 | -0.338789 | 0.001452355 | -1.263146 | 0.4038239 |
| ENSG00000183323.12 | CCDC125 | -2.644984 | 0.7736255 | 0.88732 | 0.6527736 | 2.378099 | 0.001932565 | -2.34107 | 0.2174461 | 1.244875 | 0.844493 | 1.56398 | 0.5066477 |
| ENSG00000183426.15 | NPIPA1 | 0.505081 | 0.1754689 | 0.0984 | 0.6527736 | -0.174237 | 0.8889153 | 0.862604 | 6.15E-05 | 1.89041 | 0.0963637 | -1.087557 | 0.8818722 |
| ENSG00000183431.11 | SF3A3 | 1.045409 | 0.6214382 | 0.013494 | 0.6529973 | 0.194973 | 0.8153577 | 1.108017 | 0.00597044 | 0.208508 | 0.5923787 | 1.802059 | 0.9045641 |
| ENSG00000183495.13 | EP400 | -0.106527 | 0.9501272 | -0.043461 | 0.5394427 | -0.231483 | 0.5290845 | 1.651705 | 0.000284065 | 1.850072 | 0.01352068 | 1.152801 | 0.9330765 |
| ENSG00000183520.11 | UTP11L | 0.496001 | 0.8006286 | 0.11327 | 0.9028136 | 0.121418 | 0.4257919 | 0.382108 | 0.4919368 | -0.078964 | 0.00574202 | 0.772985 | 0.7523447 |
| ENSG00000183530.13 | PRR14L | -1.154778 | 0.6214382 | 0.20414 | 0.8953649 | -0.065035 | 0.003949334 | 0.5709 | 0.1484173 | 0.127466 | 0.9606942 | 1.359559 | 0.9589318 |
| ENSG00000183617.4 | MRPL54 | 0.160115 | 0.7199598 | -0.228727 | 0.7321384 | -0.62607 | 0.001201424 | -1.046319 | 0.384552 | 0.548117 | 0.5196843 | 0.368527 | 1 |
| ENSG00000183624.13 | HMCES | -0.019847 | 0.8599919 | 0.286423 | 0.8263112 | -0.421913 | 0.110985 | 1.256197 | 0.04077943 | 0.334662 | 0.3974318 | -0.594368 | 0.9147227 |
| ENSG00000183648.9 | NDUFB1 | -0.276432 | 0.9444548 | -0.028374 | 0.5445556 | -0.683505 | 3.31E-09 | -0.322104 | 0.01404551 | -0.736168 | 1.37E-06 | 0.58466 | 0.8270227 |
| ENSG00000183684.7 | ALYREF | 0.698294 | 0.9591207 | -0.070129 | 0.8760574 | 0.259923 | 0.8400653 | 1.549498 | 0.007135165 | 1.30788 | 0.1594797 | -0.890072 | 0.4553016 |
| ENSG00000183696.13 | UPP1 | 0.333234 | 0.9198482 | -0.05466 | 0.9104976 | -0.078459 | 0.5290845 | 0.22181 | 0.4097742 | -1.575923 | 0.04756918 | -0.054775 | 0.807957 |
| ENSG00000183715.13 | OPCML | 3.392012 | 0.563505 | -0.540854 | 0.9222921 | 2.998574 | 0.1484182 | 0.996831 | 0.01663572 | 1.470239 | 0.02508037 | -1.962203 | 0.8404592 |
| ENSG00000183722.7 | LHFP | -0.79527 | 0.8587992 | 0.032686 | 0.5010139 | 0.475944 | 0.5290845 | 1.139028 | 0.754102 | -1.237771 | 1.14E-05 | 0.290737 | 1 |
| ENSG00000183723.12 | CMTM4 | -0.226693 | 1 | -1.100502 | 0.5010139 | -0.97374 | 0.7231855 | 0.756866 | 0.2049335 | 2.577149 | 0.01495364 | -2.676378 | 0.9264343 |
| ENSG00000183726.10 | TMEM50A | -0.163475 | 0.9962523 | -0.165983 | 0.6527736 | 0.248507 | 0.9765029 | -0.358414 | 0.01221979 | -0.22817 | 0.009329853 | -0.022078 | 0.8681734 |
| ENSG00000183735.9 | TBK1 | 0.665551 | 0.6792949 | -0.360499 | 0.4883536 | 0.099061 | 0.04299809 | 0.843874 | 0.1052874 | 0.711103 | 0.7257217 | 2.473295 | 0.359138 |
| ENSG00000183741.11 | CBX6 | -0.115156 | 0.9437365 | -0.146534 | 0.6615629 | -0.954094 | 0.8059492 | 1.272225 | 9.93E-06 | 1.344165 | 0.4254525 | -0.89328 | 0.826754 |
| ENSG00000183793.13 | NPIPA5 | 1.436809 | 0.714717 | 0.531431 | 0.9697628 | -0.228194 | 0.4198037 | 1.278732 | 0.007565047 | 2.933335 | 0.001312023 | -1.502168 | 1 |
| ENSG00000183808.11 | RBM12B | 1.328754 | 0.8006286 | 0.314512 | 0.832961 | 1.856912 | 0.000399937 | 1.176915 | 0.3359984 | 3.030847 | 0.000942426 | -1.703941 | 0.9398376 |
| ENSG00000183943.5 | PRKX | 0.091129 | 0.6214382 | 0.130875 | 0.7246867 | 0.359583 | 0.2091705 | 2.016873 | 0.01223151 | 0.627657 | 0.01500431 | 0.9186 | 0.8949489 |
| ENSG00000183955.12 | SETD8 | -0.232568 | 0.8024966 | 0.08033 | 0.9104976 | -0.183382 | 0.08445234 | 0.929836 | 0.01026578 | 0.970302 | 0.1081438 | -0.0363 | 0.4403284 |
| ENSG00000183978.7 | COA3 | 0.403527 | 0.2223864 | -0.279516 | 0.6580135 | -0.946685 | 0.00715996 | 0.096588 | 0.5162824 | 0.140879 | 0.2680464 | 0.536343 | 0.8719127 |
| ENSG00000184007.17 | PTP4A2 | 0.646953 | 0.6214382 | 0.138691 | 0.797408 | 0.492464 | 0.06008581 | 0.294617 | 0.000334033 | 0.691759 | 0.6517775 | -0.227189 | 0.744192 |
| ENSG00000184009.9 | ACTG1 | -0.12357 | 0.000286034 | -0.141683 | 0.2518636 | 0.167527 | 0.1656241 | 0.250587 | 0.03048442 | 0.338463 | 0.2511196 | 0.069442 | 0.7573485 |
| ENSG00000184014.7 | DENND5A | 0.040256 | 0.6214382 | 0.094426 | 0.780119 | 0.929922 | 0.2839251 | 1.569876 | 0.01566609 | 1.034171 | 0.05359226 | 3.036524 | 0.6885065 |
| ENSG00000184076.12 | UQCR10 | 0.567803 | 0.0886827 | -0.116295 | 0.3530125 | -0.552327 | 5.97E-13 | 0.12814 | 0.02255629 | -0.311189 | 1.36E-05 | -0.599443 | 0.7663039 |
| ENSG00000184163.3 | FAM132A | 0.051184 | 0.714717 | 0.648404 | 0.6527736 | -2.864373 | 1 | 0.883988 | 0.4919368 | 1.195726 | 0.8021966 |  |  |
| ENSG00000184164.14 | CRELD2 | 0.558732 | 1 | 0.143309 | 0.5953732 | -1.622809 | 1.36E-10 | 0.40219 | 0.1696878 | 1.315614 | 0.8161922 | -2.02299 | 0.6885065 |
| ENSG00000184194.5 | GPR173 | 2.320962 | 1 | 1.612069 | 0.8217443 | -0.936161 | 0.1669069 | 1.403063 | 0.008552578 | 1.481964 | 0.000327706 | -1.592036 | 1 |
| ENSG00000184209.14 | SNRNP35 | -0.296604 | 0.9513755 | 0.613044 | 0.8703522 | 0.221208 | 0.7333189 | 1.832437 | 0.009864083 | 0.812558 | 0.3974318 | -0.038212 | 0.8479431 |
| ENSG00000184216.11 | IRAK1 | -1.009065 | 0.9789948 | 0.187143 | 0.8217587 | 0.192152 | 0.5473521 | 0.287073 | 0.01453163 | 0.844641 | 0.7253092 | 0.065711 | 0.9264343 |
| ENSG00000184221.12 | OLIG1 | -2.405332 | 0.000286034 | -0.883424 | 0.4069075 | 0.363059 | 0.02588737 | 1.741336 | 0.02314265 | 1.185625 | 0.1310058 | -0.400607 | 0.7920929 |
| ENSG00000184226.14 | PCDH9 | 0.076332 | 0.714717 | -0.647963 | 0.04980181 | 0.639287 | 9.69E-10 | 0.083312 | 0.3881084 | 0.399691 | 1 | 3.362074 | 0.6185816 |
| ENSG00000184271.15 | POU6F1 | -3.438064 | 0.6792949 | 0.232414 | 0.8641479 | -0.225283 | 0.9665355 | 2.108477 | 9.15E-06 | 1.655499 | 0.0003051 | 1.271995 | 0.6885065 |
| ENSG00000184277.12 | TM2D3 | -2.148159 | 0.563505 | -0.13399 | 0.9490364 | -0.132463 | 0.186269 | 0.170226 | 0.9953945 | 0.11522 | 0.03179738 | 0.956532 | 0.6885065 |
| ENSG00000184371.13 | CSF1 | -3.091291 | 0.563505 | -0.431631 | 0.1957576 | -1.142868 | 0.1958328 | 1.700547 | 0.04287183 | -0.283217 | 1 | -0.409229 | 0.7767294 |
| ENSG00000184402.14 | SS18L1 | -0.64945 | 0.6574918 | 0.540902 | 0.8263112 | 0.086678 | 0.1130704 | 1.71455 | 0.002427199 | 0.665465 | 0.9519348 | 0.249006 | 0.8489643 |
| ENSG00000184432.9 | COPB2 | -0.181269 | 0.9047689 | 0.092398 | 0.8953649 | 0.707085 | 0.001855825 | 0.051463 | 0.613586 | 0.041812 | 0.5196843 | -0.90525 | 1 |
| ENSG00000184486.8 | POU3F2 | -1.532926 | 0.5687266 | -0.148902 | 0.3465098 | -0.022302 | 0.1674763 | 1.59158 | 0.004555736 | 1.283796 | 0.009828167 | 1.116085 | 0.3058061 |
| ENSG00000184515.10 | BEX5 | -3.11316 | 0.8814146 | 0.125469 | 1 | -0.087862 | 0.1697893 | -0.201698 | 0.04089425 | -0.192514 | 0.2187443 | -0.552816 | 0.9646502 |
| ENSG00000184545.10 | DUSP8 | 0.19946 | 0.8006286 | -1.183146 | 0.7269578 | -0.466466 | 0.736148 | 1.903384 | 0.01262694 | 0.400883 | 3.20E-05 | -0.837153 | 0.6885065 |
| ENSG00000184557.4 | SOCS3 | -0.118258 | 0.3946735 | 0.453667 | 0.5953732 | -1.086866 | 0.002328365 | 0.852252 | 0.7945946 | 0.384423 | 1 | -0.647467 | 0.9228374 |
| ENSG00000184575.11 | XPOT | 0.164204 | 0.8474876 | 0.039699 | 0.8056038 | 0.046645 | 0.7356546 | 1.181146 | 0.00106462 | 0.254651 | 0.699868 | -0.43998 | 0.6885065 |
| ENSG00000184588.17 | PDE4B | -1.05625 | 0.4764974 | -0.555637 | 0.6527736 | 1.329602 | 0.00322036 | 1.699836 | 0.6109338 | 0.386924 | 0.7377262 | 0.560833 | 0.7920121 |
| ENSG00000184602.5 | SNN | 1.164152 | 0.714717 | 0.684392 | 0.3465098 | 0.936326 | 0.7437308 | 1.280599 | 0.005022021 | 1.069044 | 0.162379 | -1.366517 | 0.2658789 |
| ENSG00000184634.15 | MED12 | 2.511173 | 0.714717 | -0.745548 | 0.6527736 | -1.721595 | 0.5835895 | 2.934154 | 0.03168293 | 1.539791 | 0.2603432 | 1.764929 | 0.6885065 |
| ENSG00000184752.12 | NDUFA12 | -0.112899 | 0.6214382 | -0.164237 | 0.401251 | -0.523524 | 0.009908236 | -0.619744 | 2.82E-06 | -0.306584 | 0.00854123 | 0.797475 | 0.5599477 |
| ENSG00000184787.18 | UBE2G2 | -1.935171 | 0.007597665 | 0.182773 | 0.9222921 | 0.072053 | 0.9977776 | 1.251925 | 2.00E-06 | 0.779472 | 0.5196843 | -0.18716 | 0.9131264 |
| ENSG00000184792.15 | OSBP2 | -1.916753 | 0.9667286 | -1.173607 | 0.3381481 | 0.622016 | 0.3931442 | 0.836663 | 0.01062276 | 2.03055 | 0.2363864 | -1.09945 | 0.8479431 |
| ENSG00000184840.11 | TMED9 | 0.241479 | 0.2065712 | 0.191849 | 0.5953732 | -0.448535 | 0.000544144 | -0.196614 | 0.006382598 | 0.096824 | 0.00858852 | -0.363168 | 0.9581926 |
| ENSG00000184863.10 | RBM33 | -0.173504 | 1 | 0.28267 | 0.6816051 | 0.749493 | 0.02449485 | 0.840988 | 0.05983567 | 0.340233 | 0.3812741 | 2.213858 | 0.6885065 |
| ENSG00000184897.5 | H1FX | -1.425916 | 0.6014343 | 0.053329 | 0.9944928 | -1.939421 | 1.99E-11 | 0.392664 | 0.9085299 | 1.417475 | 1.02E-10 | 1.043573 | 0.8479431 |
| ENSG00000184979.9 | USP18 | -0.496419 | 1 | 0.770701 | 0.9764368 | -0.999452 | 0.003775625 | 1.67498 | 0.8270451 | 0.765651 | 1 | 2.694961 | 0.5976937 |
| ENSG00000184983.9 | NDUFA6 | -1.433291 | 0.2398916 | 0.03401 | 0.6954394 | -0.246639 | 4.37E-05 | -0.319612 | 4.27E-07 | -0.549748 | 0.000399825 | 0.181701 | 0.6829816 |
| ENSG00000184990.12 | SIVA1 | 0.218714 | 0.8006286 | 0.489161 | 0.01902676 | -0.434881 | 0.02737567 | -0.235748 | 0.1985251 | -0.708495 | 0.0707843 | 0.627899 | 1 |
| ENSG00000185024.16 | BRF1 | -1.074416 | 0.6214382 | -0.445593 | 0.7336348 | 0.23281 | 0.9481144 | 1.100385 | 0.02516296 | 1.205167 | 0.6367485 | -1.432571 | 1 |
| ENSG00000185033.14 | SEMA4B | -0.220445 | 1 | -2.061798 | 0.3530125 | 2.026346 | 0.6113027 | 2.08974 | 0.4701537 | 1.937294 | 0.01236688 | 0.766268 | 0.6945946 |
| ENSG00000185043.10 | CIB1 | 1.437251 | 0.1004496 | 0.340021 | 0.5613062 | -0.129556 | 0.000921272 | -0.085712 | 0.01521201 | -0.156757 | 0.2880317 | -0.385142 | 0.9257687 |
| ENSG00000185052.11 | SLC24A3 | -1.945584 | 0.781685 | -0.253323 | 1 | 1.797667 | 0.5290845 | 1.664477 | 0.03205589 | -0.58628 | 0.8390546 | -3.985341 | 0.6885065 |
| ENSG00000185088.13 | RPS27L | -0.055371 | 0.9371596 | -0.064945 | 0.6527736 | -0.108493 | 0.01099163 | -0.113041 | 0.01645598 | 0.308825 | 0.3974318 | -0.028346 | 0.6885065 |
| ENSG00000185122.10 | HSF1 | -0.28606 | 0.8006286 | 0.105671 | 0.7062105 | -0.935288 | 0.000418292 | 1.185412 | 3.17E-05 | 1.661813 | 4.36E-06 | 1.411852 | 1 |
| ENSG00000185129.5 | PURA | -2.155183 | 0.01644604 | -0.06393 | 0.8445455 | 0.951549 | 0.1527554 | 2.056426 | 0.01691853 | 0.694611 | 0.1461268 | -0.373527 | 0.6978132 |
| ENSG00000185163.9 | DDX51 | -0.502114 | 0.9345595 | 0.510061 | 0.8202471 | 0.403993 | 0.9407871 | 1.331654 | 0.01970922 | 1.437351 | 0.03997091 | 1.213545 | 0.6885065 |
| ENSG00000185219.16 | ZNF445 | -0.761269 | 0.8234926 | 0.111342 | 0.9439293 | -1.125408 | 0.4159143 | 1.165999 | 0.008093217 | 1.206519 | 0.02023917 | -0.784141 | 0.8497806 |
| ENSG00000185222.7 | WBP5 | 0.310757 | 0.8006286 | 0.251702 | 0.6136843 | 0.223566 | 0.5290845 | 0.042314 | 0.03423155 | 0.355027 | 0.0554405 | 0.622762 | 0.9565287 |
| ENSG00000185236.11 | RAB11B | 0.404277 | 0.4195916 | 0.451506 | 0.5649416 | -0.692729 | 0.001002012 | 0.130029 | 0.7145467 | -0.075413 | 0.8202133 | 0.001668 | 0.9906609 |
| ENSG00000185246.17 | PRPF39 | 0.448845 | 0.4634325 | 0.24494 | 0.8465126 | 0.398582 | 0.7044335 | 2.35883 | 0.04352887 | 0.90468 | 0.9629537 | -1.220612 | 0.6885065 |
| ENSG00000185252.17 | ZNF74 | 0.689017 | 0.7678046 | 0.25784 | 0.9628321 | -1.36753 | 0.5519203 | -0.271987 | 0.5747952 | 2.526672 | 0.01427594 | -3.570376 | 0.1783569 |
| ENSG00000185262.8 | UBALD2 | -1.70591 | 0.6014343 | -0.447683 | 0.007219487 | 1.088597 | 0.001006486 | 0.945715 | 0.9948759 | 0.704243 | 0.6351398 | -0.551736 | 0.6885065 |
| ENSG00000185298.12 | CCDC137 | -0.445019 | 0.4764974 | -0.572542 | 0.03324584 | -0.004658 | 0.5290845 | -0.059621 | 0.4919368 | -0.472299 | 0.195584 | -0.712214 | 0.435771 |
| ENSG00000185344.13 | ATP6V0A2 | -2.406494 | 0.1835496 | -0.266644 | 0.4145756 | 0.929706 | 0.8357355 | 1.705364 | 9.18E-05 | 0.518388 | 0.002918613 | 0.172016 | 0.7046514 |
| ENSG00000185418.15 | TARSL2 | -0.105358 | 0.7648042 | 0.051254 | 0.8154382 | 0.181542 | 0.603123 | 0.759386 | 0.02455892 | 0.061601 | 0.573869 | 2.393751 | 0.9600631 |
| ENSG00000185420.18 | SMYD3 | 1.613907 | 0.3946735 | 0.242222 | 0.6527736 | -0.626866 | 0.03232856 | -0.300866 | 0.04066691 | -0.332594 | 0.2906522 | -0.250606 | 0.9539154 |
| ENSG00000185475.8 | TMEM179B | -0.207433 | 0.8006286 | -0.136095 | 0.6710279 | -0.659441 | 9.91E-05 | -0.235039 | 0.2206253 | -0.066332 | 0.3932047 | 0.386816 | 0.97072 |
| ENSG00000185507.19 | IRF7 | -0.59834 | 0.6214382 | 0.109709 | 0.8263112 | -1.844116 | 4.90E-05 | 1.071944 | 0.4919368 | -0.300308 | 0.620182 | 2.606942 | 0.6861518 |
| ENSG00000185565.11 | LSAMP | 0.947243 | 0.6214382 | -0.128032 | 0.5956071 | 1.928119 | 9.29E-06 | 0.886275 | 0.006698399 | 0.448512 | 0.2206737 | -1.25159 | 0.8560031 |
| ENSG00000185608.8 | MRPL40 | -0.389219 | 0.9183025 | 0.309478 | 0.7785607 | -0.482958 | 0.007667511 | 0.53802 | 0.689137 | -0.065423 | 0.1389793 | -0.214829 | 0.6829816 |
| ENSG00000185624.14 | P4HB | 1.173772 | 0.001302176 | 0.219034 | 0.6527736 | -0.147904 | 0.801943 | 0.56484 | 0.4782556 | -0.028304 | 0.2890193 | -0.21516 | 0.6517664 |
| ENSG00000185630.18 | PBX1 | -0.371345 | 0.7795461 | 0.788543 | 0.6527736 | 1.745408 | 0.02483941 | 1.090503 | 0.2955819 | 0.85139 | 0.4987807 | 1.136045 | 0.9154868 |
| ENSG00000185650.9 | ZFP36L1 | -2.474715 | 0.008115756 | -0.259792 | 0.354537 | 0.135811 | 0.5290845 | 0.117223 | 0.7144938 | 0.514693 | 0.8589299 | 0.349016 | 0.5344421 |
| ENSG00000185651.14 | UBE2L3 | -0.535091 | 0.6809954 | -0.130389 | 0.2801475 | 0.159189 | 0.5290845 | 0.060277 | 0.2602146 | -0.466443 | 3.20E-05 | -0.807235 | 0.4018772 |
| ENSG00000185658.13 | BRWD1 | 0.229516 | 0.8867323 | 0.173083 | 0.8196487 | 1.135016 | 0.000695965 | 1.103475 | 0.002986579 | 2.020251 | 0.00139047 | -0.28357 | 0.7973567 |
| ENSG00000185721.11 | DRG1 | -1.076208 | 0.8006286 | 0.160105 | 0.9104976 | 0.278079 | 0.8026512 | 1.125165 | 0.04310756 | -0.224449 | 0.2719076 | 0.102594 | 0.8479431 |
| ENSG00000185722.16 | ANKFY1 | -4.702454 | 0.7736255 | 0.127936 | 1 | 0.122838 | 1 | 0.623761 | 0.0484318 | 1.50082 | 0.04948871 | 0.574081 | 0.8829971 |
| ENSG00000185728.16 | YTHDF3 | -1.050867 | 0.5356155 | -0.093981 | 0.9217053 | 0.448187 | 0.5507355 | 1.511789 | 9.93E-06 | 0.587911 | 0.09570889 | -1.397894 | 0.8935461 |
| ENSG00000185760.15 | KCNQ5 | -0.961083 | 0.563505 | 0.444415 | 0.6858761 | -1.101285 | 0.5290845 | -0.094244 | 0.02497894 | -1.398792 | 0.7377262 | 2.897686 | 0.7767294 |
| ENSG00000185787.14 | MORF4L1 | 0.383682 | 0.6014343 | 0.217688 | 0.826013 | 0.287204 | 0.01739558 | 0.222394 | 0.1278163 | -0.14708 | 0.000118883 | -0.749098 | 0.6829816 |
| ENSG00000185818.7 | NAT8L | -0.905827 | 0.8220319 | -0.045385 | 0.6630531 | -0.231354 | 0.5966424 | 2.034291 | 0.000760156 | 0.025806 | 0.9566575 | 1.798051 | 0.6885065 |
| ENSG00000185825.15 | BCAP31 | -0.775748 | 0.96853 | 0.181916 | 0.6889938 | 0.020858 | 0.9330551 | 0.432016 | 0.3176557 | -0.519037 | 0.04388564 | 0.292018 | 0.9565287 |
| ENSG00000185896.10 | LAMP1 | 0.230681 | 0.6124876 | 0.051195 | 0.6527736 | -0.76789 | 8.53E-06 | 1.150452 | 0.01554383 | 0.633672 | 0.8525311 | 0.146913 | 0.6885065 |
| ENSG00000185920.15 | PTCH1 | -0.935711 | 0.6214382 | 0.750138 | 0.7079982 | -0.472195 | 0.5290845 | 2.480366 | 6.15E-05 | -2.019141 | 0.8147074 | -3.089224 | 0.7346273 |
| ENSG00000185950.8 | IRS2 | 0.714392 | 0.6214382 | 0.002483 | 0.8703522 | 0.44331 | 0.09602319 | 2.266795 | 0.005375179 | 1.418131 | 0.04355409 | 0.335505 | 1 |
| ENSG00000185963.13 | BICD2 | -4.63085 | 0.3286597 | -0.352644 | 0.7449708 | 0.633843 | 0.01151805 | 1.610998 | 2.58E-05 | 0.928853 | 0.05836207 | 0.88011 | 0.9147227 |
| ENSG00000185985.8 | SLITRK2 | 2.295811 | 0.4406957 | -3.525325 | 0.03683482 | 0.339479 | 0.09799031 | 2.871989 | 0.2626326 | 0.406724 | 0.961805 | -0.072684 | 0.7767294 |
| ENSG00000186010.18 | NDUFA13 | 0.54862 | 0.004419341 | 0.278863 | 0.4215149 | -0.839421 | 1.58E-19 | -0.080465 | 0.1126321 | 1.422169 | 0.002076913 | -0.007197 | 0.6647029 |
| ENSG00000186017.14 | ZNF566 | 1.243485 | 0.6214382 | -0.336868 | 0.5240506 | 0.563903 | 0.06786536 | 1.99513 | 0.01507461 | 2.049037 | 0.6517097 | -1.682141 | 0.9352001 |
| ENSG00000186020.12 | ZNF529 | 1.886512 | 0.714717 | -0.744551 | 0.5984855 | 0.299702 | 0.536247 | 0.644897 | 0.0219796 | 1.24225 | 0.2570214 | 1.123463 | 0.7767294 |
| ENSG00000186106.11 | ANKRD46 | 0.809712 | 0.8006286 | 0.339496 | 0.7383998 | 0.703067 | 0.3491842 | 0.770715 | 0.2607967 | 1.53395 | 0.04558554 | 0.176446 | 0.6885065 |
| ENSG00000186111.8 | PIP5K1C | -3.658273 | 0.02815464 | 0.613506 | 0.6527736 | -0.778446 | 0.6717367 | 2.151726 | 0.000456725 | 1.615789 | 0.000752147 | -2.225528 | 0.6885065 |
| ENSG00000186132.14 | C2orf76 | 0.888772 | 0.6039189 | 0.342745 | 1 | -0.79464 | 0.1585172 | -0.887834 | 0.01034653 | -1.019218 | 0.3567865 | -0.499359 | 0.9147227 |
| ENSG00000186166.8 | CCDC84 | 0.091441 | 0.714717 | 0.174107 | 0.8703522 | 0.229917 | 0.5290845 | 1.152217 | 0.02421461 | 1.388613 | 0.9622018 | -0.718546 | 0.6829816 |
| ENSG00000186174.12 | BCL9L | -0.17963 | 0.9282145 | -0.659143 | 0.6527736 | -0.905064 | 0.3037092 | 1.130416 | 0.00066495 | 2.135246 | 0.03586689 | -0.13915 | 0.9147227 |
| ENSG00000186187.11 | ZNRF1 | -1.820059 | 0.529676 | -0.370823 | 1 | 0.897181 | 0.5455771 | 0.815416 | 0.08200106 | 1.764164 | 0.02488244 | 0.421715 | 0.9589332 |
| ENSG00000186193.8 | SAPCD2 | -0.145259 | 0.825472 | -1.127793 | 0.3522748 | -0.799296 | 0.08184338 | 1.223581 | 0.007728367 | 2.318131 | 1.53E-06 | -1.813783 | 0.7323674 |
| ENSG00000186222.4 | BLOC1S4 | -0.161455 | 0.6214382 | -0.224083 | 0.5953732 | -1.834869 | 0.01785955 | 0.99438 | 0.07141399 | 1.06773 | 0.4890731 | -0.006086 | 0.6885065 |
| ENSG00000186298.11 | PPP1CC | 0.387903 | 0.714717 | 0.220818 | 0.6527736 | 0.592035 | 0.1497049 | 0.897993 | 0.005620275 | 0.315561 | 0.766562 | 1.322062 | 0.6885065 |
| ENSG00000186395.6 | KRT10 | -1.579673 | 0.00903383 | -0.199555 | 0.07156425 | -1.509938 | 6.32E-25 | 0.420526 | 0.506159 | 0.49445 | 0.4826781 | -1.2267 | 0.6829816 |
| ENSG00000186432.8 | KPNA4 | -1.082844 | 0.6776314 | 0.075875 | 0.6527736 | 0.313683 | 0.9967097 | 0.990538 | 0.008373978 | 0.882808 | 0.1431053 | -0.735391 | 0.8562379 |
| ENSG00000186468.12 | RPS23 | -1.099908 | 0.3287298 | 0.205008 | 0.1313103 | -0.068463 | 0.6266498 | -0.189539 | 3.84E-05 | -0.138204 | 0.001277525 | -0.001259 | 0.2411252 |
| ENSG00000186532.11 | SMYD4 | 3.932062 | 0.7736255 | 0.228155 | 0.7281803 | -1.006814 | 0.8454005 | 0.88452 | 0.02468518 | 1.616592 | 0.1723311 | -0.501536 | 0.8479431 |
| ENSG00000186566.12 | GPATCH8 | -0.488081 | 0.6214382 | 0.742486 | 0.5953732 | 0.087372 | 0.3365506 | 1.009408 | 0.1956859 | 1.714022 | 0.0109684 | 1.237878 | 0.9276623 |
| ENSG00000186575.17 | NF2 | -0.434986 | 0.6214382 | -0.166462 | 0.6527736 | 0.588726 | 0.4055474 | 1.3096 | 0.04645593 | 0.805154 | 0.5196843 | 1.516152 | 0.6829816 |
| ENSG00000186577.11 | C6orf1 | 0.144865 | 0.6214382 | 0.235444 | 0.7013468 | -0.840931 | 1.49E-06 | -0.064572 | 0.7219547 | 0.723943 | 0.8381224 | -0.855197 | 0.5104122 |
| ENSG00000186591.11 | UBE2H | -2.303957 | 0.1448298 | -0.147288 | 0.5395105 | 0.477736 | 0.1991342 | -0.186282 | 0.7399696 | 0.905145 | 0.03937989 | 0.079674 | 0.6885065 |
| ENSG00000186648.14 | LRRC16B | 1.250732 | 1 | 0.737766 | 0.9329105 | 3.304563 | 0.5290845 | 4.005728 | 0.0434285 | -0.602808 | 0.2880317 | 4.10022 | 1 |
| ENSG00000186660.14 | ZFP91 | 0.606789 | 0.5579177 | 0.386253 | 0.6527736 | 1.916668 | 0.01263308 | 0.660587 | 0.02029956 | 0.743532 | 0.8181648 | 0.517278 | 0.9577321 |
| ENSG00000186687.15 | LYRM7 | -0.167745 | 1 | 0.341789 | 1 | -0.517001 | 0.8389806 | 1.376713 | 0.04691967 | 0.500196 | 0.7377262 | -0.663239 | 0.9119423 |
| ENSG00000186812.12 | ZNF397 | 0.536765 | 0.6324988 | 0.786438 | 0.9061805 | 1.569716 | 1.77E-05 | 1.765084 | 0.000549468 | 1.923659 | 0.0352973 | -1.14559 | 1 |
| ENSG00000186868.15 | MAPT | -1.187027 | 0.6014343 | 0.032279 | 0.6737951 | 0.290801 | 0.9700569 | 0.91563 | 0.002936174 | 0.3247 | 0.9599276 | -0.772882 | 0.5344421 |
| ENSG00000186918.13 | ZNF395 | 0.881276 | 1 | 0.898052 | 0.008839116 | 0.557182 | 0.6010661 | 1.010071 | 1 | 1.578498 | 0.3108583 | 0.534598 | 0.6829816 |
| ENSG00000187024.13 | PTRH1 | -2.373574 | 0.005603822 | -0.668685 | 0.6527736 | -0.814957 | 0.5290845 | -0.542365 | 0.01105541 | 0.21122 | 0.576218 | 1.415507 | 0.8479431 |
| ENSG00000187051.8 | RPS19BP1 | 0.29178 | 0.7940298 | 0.076381 | 1 | -0.57358 | 3.79E-07 | -0.16889 | 0.02598489 | 0.03697 | 0.1686314 | 2.515819 | 0.6829816 |
| ENSG00000187079.14 | TEAD1 | -1.335692 | 0.1204153 | -0.897849 | 0.000424311 | 0.793656 | 9.91E-05 | 0.453562 | 0.000217365 | -0.154382 | 0.3756673 | 0.60774 | 0.7571519 |
| ENSG00000187109.13 | NAP1L1 | -2.008322 | 0.714717 | -0.094065 | 0.6527736 | 0.597097 | 1.20E-06 | 0.63968 | 0.1781539 | 0.475737 | 0.3607078 | -0.209428 | 0.2159565 |
| ENSG00000187118.12 | CMC1 | -0.435007 | 0.7736255 | 0.041968 | 0.8703522 | -0.248539 | 0.000607495 | 0.168663 | 0.2928264 | 0.431774 | 0.8994036 | 0.784187 | 0.8479431 |
| ENSG00000187147.17 | RNF220 | 2.453668 | 0.8006286 | -0.165242 | 0.7349695 | 0.470297 | 0.5506935 | 1.035819 | 0.4572705 | 2.205819 | 0.03291688 | -0.204036 | 0.9067845 |
| ENSG00000187189.10 | TSPYL4 | 0.964918 | 0.5139066 | -0.825544 | 0.5940759 | 0.487962 | 0.0142718 | 2.198029 | 0.001147056 | 0.893285 | 0.7546247 | -0.249351 | 0.6885065 |
| ENSG00000187193.8 | MT1X | -1.207629 | 0.6792949 | 0.479118 | 0.0511244 | -0.458528 | 0.01408099 | -0.496767 | 0.008181357 | 0.514079 | 0.5196843 | 0.465521 | 0.821798 |
| ENSG00000187231.13 | SESTD1 | -0.482488 | 0.9351938 | 0.131223 | 0.8703522 | 1.271745 | 0.1372475 | 1.860991 | 0.02535905 | 1.042541 | 0.04125865 | 1.342028 | 0.6885065 |
| ENSG00000187239.16 | FNBP1 | -1.103931 | 0.8672448 | 0.173344 | 0.5953732 | 0.912576 | 3.59E-05 | -0.186204 | 1 | -0.014949 | 0.7035016 | 1.847994 | 0.6829816 |
| ENSG00000187257.14 | RSBN1L | -0.466054 | 0.6214382 | -0.494982 | 0.05201621 | 0.9447 | 8.83E-08 | 0.525332 | 0.8338473 | 0.55078 | 0.1363141 | -0.794876 | 0.6138783 |
| ENSG00000187479.5 | C11orf96 | -0.883167 | 0.5692871 | -1.385329 | 0.01966733 | -1.295955 | 0.121786 | 1.563919 | 0.1254664 | 2.410149 | 0.000427519 | 0.770499 | 0.8724921 |
| ENSG00000187514.14 | PTMA | -0.450195 | 0.8006286 | -0.033026 | 0.6527736 | 0.532381 | 3.09E-15 | -0.026409 | 0.03271749 | -0.429419 | 5.43E-06 | -0.032493 | 0.426223 |
| ENSG00000187531.13 | SIRT7 | -1.056285 | 0.7693197 | 0.711581 | 0.5010139 | -1.047319 | 0.02719861 | 0.821603 | 0.08918292 | 2.016105 | 0.3941931 | -2.997998 | 0.8956025 |
| ENSG00000187555.14 | USP7 | -3.640307 | 0.2990001 | 0.336667 | 0.9723966 | 0.874209 | 7.92E-05 | 0.910999 | 0.006275933 | 1.001936 | 0.01127024 | -0.025578 | 1 |
| ENSG00000187605.15 | TET3 | -0.371157 | 0.6214382 | 0.58507 | 0.6015027 | 0.650648 | 0.003206137 | 1.052701 | 0.04439064 | 1.409081 | 0.02149852 | 1.348412 | 0.3762381 |
| ENSG00000187607.15 | ZNF286A | 0.385787 | 0.8787256 | 0.73066 | 0.9730349 | 1.267237 | 0.1663608 | 0.50987 | 0.0028842 | 0.85913 | 0.5188602 | -1.021559 | 0.6885065 |
| ENSG00000187608.8 | ISG15 | 1.19112 | 0.6214382 | 0.143076 | 0.8717486 | -1.276418 | 2.77E-18 | 0.674482 | 0.4919368 | -0.169183 | 0.4531181 | 1.321088 | 0.936179 |
| ENSG00000187634.10 | SAMD11 | 0.096063 | 1 | -0.341706 | 0.6527736 | -6.488654 | 0.002447825 | 1.106277 | 0.09294199 | 0.140713 | 0.3371433 | -1.31571 | 1 |
| ENSG00000187735.12 | TCEA1 | 0.472256 | 0.6014343 | 0.532495 | 0.1886054 | 0.411897 | 0.02450741 | -0.717021 | 0.01433292 | -0.588968 | 5.77E-07 | 0.216724 | 0.6885065 |
| ENSG00000187742.14 | SECISBP2 | 0.057202 | 0.5711449 | 0.290347 | 0.9296542 | 1.096028 | 0.03915157 | 0.811482 | 0.7055936 | 1.200712 | 0.2317816 | 1.354486 | 0.8309897 |
| ENSG00000187801.14 | ZFP69B | 2.173237 | 0.8587992 | -0.321591 | 0.5953732 | 1.87038 | 0.5095769 | 0.737898 | 0.7864362 | 1.091175 | 0.8942969 |  |  |
| ENSG00000187837.3 | HIST1H1C | -1.61325 | 1 | -0.602574 | 0.804846 | 0.934167 | 0.009257526 | -0.234765 | 0.3357935 | 0.785103 | 0.9297852 | 0.918549 | 1 |
| ENSG00000187840.4 | EIF4EBP1 | 0.575551 | 0.01833589 | 0.364432 | 0.2578431 | -0.018363 | 0.02838024 | -0.373842 | 0.06877857 | -0.055054 | 0.08636113 | 0.249201 | 0.5411083 |
| ENSG00000187866.7 | FAM122A | -0.676402 | 0.4364395 | 0.315309 | 0.83982 | -1.19686 | 0.4848276 | 1.608135 | 0.03814667 | 1.121826 | 0.3387021 | -0.568739 | 0.9638718 |
| ENSG00000188010.12 | MORN2 | -0.438659 | 0.6214382 | -0.210886 | 0.7881421 | -0.238317 | 0.04480826 | -0.729996 | 0.02173865 | -0.010416 | 0.4224614 | -0.891632 | 0.6829816 |
| ENSG00000188021.8 | UBQLN2 | 0.767244 | 0.6214382 | -0.269425 | 0.6542448 | 0.929564 | 0.04392145 | 2.819451 | 0.009420281 | 1.375607 | 0.2707324 | -1.05529 | 0.8479431 |
| ENSG00000188042.7 | ARL4C | 0.120148 | 0.6014343 | 0.221645 | 0.1054706 | -0.017226 | 0.04587351 | 0.466745 | 0.00692364 | 0.720751 | 0.008830335 | -0.059404 | 0.4578379 |
| ENSG00000188070.9 | C11orf95 | -0.065955 | 0.6214382 | -0.336127 | 1 | 0.261696 | 0.7437308 | 1.750035 | 1.16E-05 | 0.949195 | 0.1835921 | -1.248269 | 1 |
| ENSG00000188163.7 | FAM166A | -1.839396 | 0.8006286 | 0.302201 | 0.8703522 | -1.517414 | 0.7437308 | 4.156067 | 0.02991167 | -2.73292 | 0.5196843 | 2.524442 | 0.8479431 |
| ENSG00000188186.10 | LAMTOR4 | 0.455381 | 0.6014343 | 0.062134 | 0.8263112 | -0.744142 | 3.09E-05 | -0.092275 | 0.03614793 | -0.148627 | 0.2100584 | -0.8445 | 0.5344421 |
| ENSG00000188227.12 | ZNF793 | 0.319324 | 0.714717 | -0.060517 | 0.6527736 | 0.950637 | 0.4929745 | 2.367654 | 7.51E-06 | 0.381326 | 0.4285976 | -2.117374 | 1 |
| ENSG00000188243.12 | COMMD6 | 0.222987 | 0.563505 | 0.098305 | 0.9984248 | -0.121697 | 0.2455464 | 0.174157 | 0.7401798 | 0.060958 | 0.04995173 | -0.074966 | 0.6185816 |
| ENSG00000188290.10 | HES4 | -0.933443 | 1 | 0.015011 | 0.8667612 | -3.648909 | 0.1138222 | 0.874743 | 0.2879598 | 2.608217 | 0.002280492 | 1.019697 | 0.8479431 |
| ENSG00000188315.7 | C3orf62 | 2.213127 | 1 | -0.56236 | 0.8263112 | 0.118687 | 0.2784089 | 2.530969 | 0.04179318 | 1.066472 | 0.2147756 | -0.105895 | 0.9626957 |
| ENSG00000188322.4 | SBK1 | 1.116753 | 0.6214382 | -0.425324 | 0.6514723 | -0.49715 | 0.711025 | 1.372842 | 0.06169181 | 1.3437 | 0.000426935 | -1.846111 | 0.6885065 |
| ENSG00000188338.14 | SLC38A3 | -3.910728 | 1 | 2.181729 | 0.6527736 | 2.828541 | 0.4382561 | 3.581116 | 0.657739 | 2.156391 | 0.3125964 |  |  |
| ENSG00000188375.4 | H3F3C | -0.278068 | 0.563505 | 2.035554 | 0.04367331 | -1.881674 | 0.01634596 | 1.718858 | 0.931053 | 1.611256 | 0.5196843 | -0.647769 | 1 |
| ENSG00000188483.7 | IER5L | 0.11946 | 0.9512403 | -0.610765 | 0.08356534 | -1.000967 | 0.003012856 | 1.292862 | 0.000710238 | 3.152879 | 1.06E-05 | -0.384981 | 0.6829816 |
| ENSG00000188486.3 | H2AFX | -0.777668 | 0.5671641 | -0.175923 | 0.7674398 | -2.163222 | 0.000155995 | 1.109261 | 0.1032024 | 1.305533 | 0.004899682 | 0.29179 | 0.7243489 |
| ENSG00000188529.14 | SRSF10 | -0.118374 | 1 | 0.096521 | 0.7383998 | 0.744497 | 0.001088591 | 1.024249 | 0.08941774 | 0.619139 | 0.06555897 | 0.643618 | 0.951232 |
| ENSG00000188554.13 | NBR1 | 0.1137 | 0.9649704 | -0.299505 | 0.5953732 | 0.147682 | 1 | 1.585756 | 0.01476691 | 0.604891 | 0.9738386 | 0.576858 | 0.826754 |
| ENSG00000188559.13 | RALGAPA2 | 1.572322 | 0.6214382 | 0.392557 | 0.7820217 | -1.366668 | 0.8862589 | 2.28638 | 0.02445851 | 2.017513 | 0.03309984 | -0.098179 | 0.9569765 |
| ENSG00000188573.7 | FBLL1 | -0.087194 | 0.6854446 | -1.03153 | 0.8263112 | -0.627931 | 0.789139 | 1.635587 | 0.29158 | 0.466048 | 0.04241657 | -0.428403 | 0.7417751 |
| ENSG00000188612.11 | SUMO2 | -0.529168 | 0.6214382 | -0.161677 | 0.3121527 | 0.084889 | 0.112253 | -0.148607 | 0.02193743 | -0.157259 | 0.001092187 | -0.451111 | 0.2276299 |
| ENSG00000188636.3 | LDOC1L | 0.457427 | 0.6214382 | -0.224539 | 0.846655 | -0.358954 | 0.5290845 | 1.180435 | 0.000454752 | 1.495032 | 0.7406781 | -2.440711 | 0.3284129 |
| ENSG00000188643.10 | S100A16 | -1.671245 | 0.07604642 | -0.286111 | 0.1276613 | -1.06145 | 2.88E-11 | 0.286385 | 0.01816429 | 2.019788 | 1 | 0.450564 | 0.9270789 |
| ENSG00000188647.12 | PTAR1 | 0.655316 | 0.959823 | -0.737783 | 0.08640355 | 1.499352 | 9.07E-05 | 0.661171 | 0.000542607 | 0.619439 | 0.004459026 | 1.220944 | 0.4304069 |
| ENSG00000188735.12 | TMEM120B | -0.269393 | 0.6214382 | 0.271021 | 1 | 0.461472 | 0.8772157 | 1.192485 | 0.2031747 | 1.066898 | 0.03956112 | -0.630547 | 0.913718 |
| ENSG00000188760.10 | TMEM198 | -1.942657 | 1 | 0.699886 | 0.6527736 | 0.874935 | 0.6081028 | 0.510589 | 0.03418824 | -0.371481 | 0.5196843 | 0.103113 | 0.6885065 |
| ENSG00000188766.12 | SPRED3 | 0.753476 | 0.714717 | 0.662103 | 0.7912196 | -1.354246 | 0.04623787 | 1.20513 | 3.96E-05 | 1.891599 | 0.8621012 | 1.87092 | 0.6885065 |
| ENSG00000188786.9 | MTF1 | -0.954969 | 0.6214382 | -0.407658 | 0.6527736 | 0.617603 | 0.08784542 | 2.270889 | 0.004513611 | 2.247684 | 0.1063758 | 2.392494 | 1 |
| ENSG00000188807.12 | TMEM201 | -0.366061 | 1 | -2.277967 | 0.04079597 | 0.23247 | 1 | 1.281112 | 0.154327 | 1.962766 | 0.1524869 | -0.526032 | 0.9147227 |
| ENSG00000188846.13 | RPL14 | -0.936115 | 2.48E-05 | -0.01227 | 0.7335559 | 0.042154 | 0.70474 | -0.215082 | 0.000212025 | -0.063059 | 0.006171639 | 0.035047 | 0.5344421 |
| ENSG00000188895.11 | MSL1 | -0.786031 | 0.05571134 | -0.185909 | 0.658906 | 0.761794 | 0.3577337 | 1.0747 | 0.003130059 | 1.133081 | 0.3450415 | 0.270322 | 0.7167272 |
| ENSG00000188916.8 | FAM196A | -0.89745 | 0.9282145 | 0.43307 | 0.6527736 | 0.56285 | 0.935949 | 3.815016 | 0.001477636 | -0.074513 | 0.5196843 | 0.954958 | 0.7950024 |
| ENSG00000188994.12 | ZNF292 | 0.396507 | 0.8240785 | 0.074604 | 0.8928139 | 0.835877 | 2.30E-05 | 2.676476 | 0.000494195 | 2.266383 | 1.46E-06 | 2.221487 | 0.6796295 |
| ENSG00000189043.9 | NDUFA4 | 0.326373 | 1.78E-06 | -0.017094 | 0.8953649 | -0.362322 | 1.54E-12 | -0.351457 | 1.90E-07 | -0.442077 | 3.64E-11 | -0.276187 | 0.09921197 |
| ENSG00000189058.8 | APOD | -1.318715 | 0.226923 | -0.364272 | 0.385697 | 0.570097 | 0.4339302 | -0.758443 | 3.03E-06 | -0.625114 | 0.4921378 | 1.31637 | 0.9539924 |
| ENSG00000189091.12 | SF3B3 | -0.062751 | 0.9710176 | 0.137366 | 0.979665 | 1.065123 | 0.09929095 | 1.724027 | 2.56E-07 | 1.117415 | 0.08617161 | -1.674078 | 0.6829816 |
| ENSG00000189159.15 | HN1 | -0.006208 | 0.5428806 | -0.533523 | 0.001546986 | -0.178156 | 0.07352297 | -0.003388 | 0.1435467 | 0.098433 | 0.03224226 | 0.488532 | 0.5193439 |
| ENSG00000189227.5 | C15orf61 | -0.752624 | 0.6214382 | -0.37126 | 0.8703522 | -1.531456 | 6.32E-07 | 0.535048 | 1 | 1.414084 | 0.9847639 | 1.346733 | 0.9130454 |
| ENSG00000189308.10 | LIN54 | 1.756771 | 1 | 1.307137 | 0.8910308 | -0.111945 | 1 | 2.740018 | 0.03287638 | 1.591973 | 0.8520331 | -0.925852 | 0.7728044 |
| ENSG00000189339.11 | SLC35E2B | 0.641564 | 0.7736255 | -0.283665 | 0.8374851 | -0.95923 | 0.08400047 | 1.038336 | 0.002259593 | 2.429686 | 9.25E-05 | -0.342578 | 0.9562147 |
| ENSG00000189403.14 | HMGB1 | -0.382013 | 0.563505 | -0.010066 | 0.6748674 | 0.142425 | 0.000785673 | -0.117982 | 0.001835662 | -0.164701 | 0.006496374 | -0.025617 | 0.06642752 |
| ENSG00000196081.9 | ZNF724P | 2.221787 | 0.6014343 | -0.258945 | 0.9330407 | 1.149718 | 0.01410444 | -0.210959 | 0.6486242 | -2.662904 | 0.4801191 | -2.221949 | 0.8479431 |
| ENSG00000196083.9 | IL1RAP | -0.086311 | 0.9444548 | 0.267643 | 0.6527736 | 0.090909 | 0.8479917 | 0.751124 | 0.02029956 | 0.720512 | 0.5196843 | 1.177785 | 1 |
| ENSG00000196123.12 | KIAA0895L | 0.640391 | 0.9451106 | 0.887464 | 0.4487482 | -0.079537 | 0.6157029 | 0.739153 | 0.03307286 | 1.718263 | 0.02184139 | 1.134726 | 0.6829816 |
| ENSG00000196132.11 | MYT1 | -2.510058 | 0.6214382 | 0.037173 | 1 | -1.064228 | 0.2350968 | 1.094551 | 0.1146382 | 1.781234 | 0.00105812 | -2.917825 | 0.5733051 |
| ENSG00000196172.9 | ZNF681 | 0.595014 | 0.6214382 | -0.095807 | 0.6527736 | 0.564681 | 0.7371191 | 6.864286 | 0.02694428 | -0.199111 | 0.4265932 | #NAME? | 0.5344421 |
| ENSG00000196182.10 | STK40 | -1.557452 | 0.6214382 | -0.040182 | 0.9551669 | 0.524156 | 0.6581709 | 1.620063 | 3.49E-05 | 1.18912 | 0.05654004 | 0.391079 | 1 |
| ENSG00000196199.13 | MPHOSPH8 | -1.803504 | 0.000108719 | 0.019384 | 0.8232846 | 0.36955 | 0.001139666 | -0.504041 | 1 | -0.625418 | 0.02847942 | -0.786567 | 0.6829816 |
| ENSG00000196208.13 | GREB1 | -3.507727 | 0.6214382 | 0.140719 | 0.7049761 | -0.110402 | 0.9063164 | 1.683586 | 0.007120083 | 1.674111 | 0.3255801 | -0.159771 | 0.9403971 |
| ENSG00000196230.12 | TUBB | -0.266361 | 0.7381604 | -0.315663 | 0.02021046 | 0.386915 | 0.02096472 | 0.219175 | 0.7007056 | 0.181374 | 0.7377262 | -0.070132 | 0.05841333 |
| ENSG00000196262.13 | PPIA | -1.446706 | 0.007523103 | -0.096373 | 0.2356743 | -0.09641 | 0.08884657 | 0.126485 | 0.9461242 | -0.069324 | 0.04278906 | -0.256555 | 0.3643965 |
| ENSG00000196268.11 | ZNF493 | -0.629255 | 0.714717 | 0.194322 | 0.6527736 | 0.930819 | 0.6310059 | 1.448222 | 0.04304509 | 0.864317 | 0.2344752 | -3.249009 | 0.7354313 |
| ENSG00000196275.13 | GTF2IRD2 | -0.274635 | 0.9193148 | -1.330514 | 0.9856582 | 0.398157 | 0.2552655 | 1.26681 | 0.02801458 | 2.882739 | 9.70E-05 | -1.267339 | 0.9738331 |
| ENSG00000196305.17 | IARS | 0.934262 | 0.6362998 | 0.119503 | 0.8263112 | 0.968034 | 0.01439134 | 1.629285 | 0.01246554 | 0.9819 | 0.1407934 | 1.053394 | 0.6185816 |
| ENSG00000196313.11 | POM121 | 0.27887 | 0.789027 | 0.356086 | 0.7049761 | -0.094296 | 0.4337744 | 0.696919 | 0.0402944 | 1.150755 | 0.6141465 | -0.462587 | 0.9589332 |
| ENSG00000196323.11 | ZBTB44 | 0.10912 | 0.9183025 | -0.881769 | 0.4433091 | -0.120477 | 0.3019061 | 1.350085 | 0.001345975 | 1.790659 | 0.008863234 | -1.302242 | 0.5733051 |
| ENSG00000196338.12 | NLGN3 | 0.318313 | 0.8006286 | -0.677133 | 0.04589273 | 0.489449 | 0.8523164 | 0.189076 | 0.2754358 | 0.391587 | 0.8718069 | 1.858266 | 0.9618031 |
| ENSG00000196361.9 | ELAVL3 | -0.984302 | 0.1345325 | 0.302982 | 0.6090372 | 0.548942 | 0.01806716 | 1.201392 | 1.69E-11 | 1.660773 | 9.75E-12 | -0.79842 | 0.5344421 |
| ENSG00000196363.9 | WDR5 | 0.110107 | 0.8006286 | -0.308551 | 1 | -0.518383 | 0.5188138 | 0.684262 | 0.0312972 | 1.411606 | 0.001075709 | 0.051081 | 0.8479431 |
| ENSG00000196367.12 | TRRAP | -0.946359 | 0.6014343 | -0.213735 | 0.9795629 | 0.673887 | 0.1309478 | 2.779391 | 0.002846276 | 1.707664 | 0.0698616 | 3.185528 | 0.926878 |
| ENSG00000196369.10 | SRGAP2B | -0.097286 | 0.6214382 | -0.595322 | 0.5010139 | 1.266308 | 0.001839621 | 1.391684 | 0.08669712 | 1.433371 | 0.2956461 | 0.080225 | 0.7812339 |
| ENSG00000196372.12 | ASB13 | -0.033521 | 0.7107729 | -1.305521 | 0.7813551 | 2.079069 | 0.5884118 | 3.243914 | 0.01849898 | 0.188997 | 0.8786084 | -1.3565 | 0.926878 |
| ENSG00000196376.10 | SLC35F1 | -0.733172 | 0.9281656 | -0.145945 | 0.2185295 | 0.653418 | 0.01904784 | 0.852269 | 0.1272439 | 0.177113 | 0.8520676 | -0.520094 | 0.6885065 |
| ENSG00000196378.11 | ZNF34 | 0.75018 | 0.8810808 | -0.023482 | 0.9754542 | 1.232607 | 0.1761889 | 2.98671 | 1.16E-05 | 1.191072 | 0.06213265 | 0.715298 | 0.8949489 |
| ENSG00000196396.9 | PTPN1 | -0.85869 | 0.3291921 | -0.025067 | 0.8922633 | 0.152052 | 0.5687544 | 1.672623 | 0.001831479 | -0.152478 | 0.4493174 | -1.109542 | 0.5336372 |
| ENSG00000196405.12 | EVL | -1.456741 | 0.2233335 | 0.003704 | 0.8263112 | -0.453249 | 0.8357355 | 1.282434 | 0.01831499 | 1.111606 | 0.2019818 | 0.345622 | 0.6885065 |
| ENSG00000196422.10 | PPP1R26 | -0.443606 | 1 | 0.031849 | 0.8353216 | -0.342946 | 0.5591443 | 1.667007 | 0.004826448 | 0.621569 | 0.280178 | 1.962564 | 0.6885065 |
| ENSG00000196428.12 | TSC22D2 | 0.569972 | 1 | 0.387323 | 0.86041 | -0.068563 | 0.8764224 | 1.244784 | 0.000776635 | 1.018031 | 0.03266311 | -0.948735 | 0.9147227 |
| ENSG00000196440.11 | ARMCX4 | 0.347635 | 0.6214382 | -1.053126 | 0.5953732 | 0.979936 | 0.4130427 | 0.827669 | 0.001105852 | 0.991576 | 0.5430328 | 4.979124 | 0.6885065 |
| ENSG00000196459.13 | TRAPPC2 | 0.50664 | 0.9883587 | 0.514875 | 0.6414243 | -0.584251 | 0.04887835 | -0.132829 | 0.2185751 | -0.466932 | 0.3810728 | -0.125492 | 0.8185117 |
| ENSG00000196498.13 | NCOR2 | -0.430808 | 0.8006286 | -0.726824 | 0.0524234 | 0.070036 | 0.7467564 | 0.852081 | 5.35E-05 | 0.807152 | 0.1206053 | 0.419006 | 0.9944282 |
| ENSG00000196504.15 | PRPF40A | -0.140125 | 0.6214382 | 0.229253 | 0.8542264 | 1.242487 | 7.90E-08 | 0.593003 | 0.7893808 | 0.157997 | 0.1115046 | 0.091781 | 0.9064846 |
| ENSG00000196507.10 | TCEAL3 | -1.707697 | 0.004540752 | -0.567154 | 0.07313683 | -0.174654 | 0.007549214 | 0.466807 | 0.02192208 | -0.7225 | 0.02130307 | 1.360369 | 1 |
| ENSG00000196531.10 | NACA | -0.704013 | 0.2643278 | 0.007122 | 0.9036408 | 0.115928 | 0.2277134 | -0.181632 | 0.0008221 | -0.3308 | 2.32E-05 | -0.140795 | 0.3746505 |
| ENSG00000196547.14 | MAN2A2 | -0.11537 | 0.8689227 | -0.00996 | 0.6527736 | 0.286886 | 0.5290845 | 1.964102 | 0.00052036 | 1.034505 | 0.3974318 | -1.96126 | 0.6167027 |
| ENSG00000196576.14 | PLXNB2 | -0.609942 | 0.8787256 | 0.312753 | 0.8703522 | -0.722135 | 0.4972329 | 1.905673 | 0.002967725 | 2.013441 | 0.1060143 | 0.299779 | 0.6885065 |
| ENSG00000196586.13 | MYO6 | -0.867432 | 0.8006286 | 0.321879 | 0.7383998 | 0.829526 | 0.008774211 | 0.768931 | 0.5493541 | 1.452564 | 0.04196275 | 2.097227 | 0.8829971 |
| ENSG00000196588.14 | MKL1 | -0.24899 | 0.9183025 | -1.338602 | 0.2185295 | 1.183587 | 0.3260512 | 1.766306 | 0.009724107 | 2.719432 | 5.66E-05 | -1.289755 | 0.9304225 |
| ENSG00000196591.11 | HDAC2 | 0.348698 | 0.4782461 | -0.152312 | 0.3898048 | 0.583092 | 0.000254909 | 0.247462 | 0.1860003 | -0.088707 | 0.4877978 | 0.563385 | 1 |
| ENSG00000196628.13 | TCF4 | -0.786754 | 0.3184862 | -0.213844 | 0.2864367 | 0.992294 | 5.93E-11 | 1.978093 | 1.05E-29 | 0.677977 | 0.01905099 | -1.142812 | 0.5026528 |
| ENSG00000196632.10 | WNK3 | -1.404483 | 0.6014343 | -0.180416 | 0.5028401 | 1.634409 | 0.000254115 | 0.778191 | 0.05931811 | 1.44953 | 0.3941931 | -0.701719 | 0.6885065 |
| ENSG00000196642.16 | RABL6 | -0.971646 | 0.7099868 | 0.137956 | 0.7336348 | 0.113765 | 0.5741718 | 0.619461 | 0.0465356 | 1.328536 | 0.001637036 | -1.200916 | 0.7920121 |
| ENSG00000196646.11 | ZNF136 | 4.164832 | 0.6214382 | 0.187019 | 0.8885132 | 0.411529 | 0.1933616 | 3.391252 | 0.01471784 | 1.074252 | 0.5196843 | 0.567125 | 0.7767294 |
| ENSG00000196652.11 | ZKSCAN5 | 0.026091 | 0.935955 | 0.415569 | 0.9547032 | -0.060372 | 1 | 1.665424 | 0.04817963 | 0.599615 | 0.8121319 | -0.826902 | 0.9545079 |
| ENSG00000196655.9 | TRAPPC4 | 0.429395 | 0.8006286 | 0.462919 | 0.5394427 | -0.039613 | 0.05510724 | 0.255014 | 0.3760083 | 0.274859 | 0.001624447 | -1.114316 | 0.6885065 |
| ENSG00000196683.10 | TOMM7 | -0.53627 | 0.6214382 | -0.173579 | 0.1881619 | -0.495275 | 2.93E-09 | 0.251169 | 0.2571656 | -0.066589 | 0.1691344 | 0.340293 | 0.6185816 |
| ENSG00000196700.7 | ZNF512B | -3.852916 | 0.4764974 | -0.140687 | 0.6869133 | -0.835428 | 0.001680264 | 2.623834 | 0.0135473 | 2.68772 | 0.02251042 | 1.234374 | 0.7915993 |
| ENSG00000196705.8 | ZNF431 | -0.121158 | 0.8867323 | 0.109637 | 0.5010139 | 0.586189 | 0.1682909 | 0.77864 | 0.000549468 | 0.251488 | 0.3941931 | -1.669741 | 0.7767294 |
| ENSG00000196712.16 | NF1 | 0.110729 | 1 | -0.240526 | 0.4130975 | 1.220202 | 5.04E-05 | 0.722964 | 0.02141258 | 2.321558 | 0.006214714 | 2.421921 | 0.6885065 |
| ENSG00000196754.10 | S100A2 | -0.946022 | 0.8006286 | -0.730287 | 0.6692821 | -0.302497 | 0.03684288 | 0.86586 | 0.8859784 | -4.550812 | 0.5196843 | -2.088757 | 0.3846916 |
| ENSG00000196776.14 | CD47 | 0.390085 | 0.6214382 | 0.354657 | 0.2128045 | -0.428065 | 0.022069 | 0.638565 | 0.5876718 | 0.409637 | 0.6418821 | 0.053535 | 0.6885065 |
| ENSG00000196792.11 | STRN3 | -0.146743 | 0.9708867 | 0.334334 | 0.9164212 | 0.198934 | 0.9314284 | 0.946432 | 0.002046134 | -0.038611 | 1 | -1.786349 | 0.7110407 |
| ENSG00000196814.14 | MVB12B | 0.260021 | 0.6798783 | 0.222192 | 0.6527736 | -0.037511 | 0.826493 | 0.319056 | 0.3467508 | 2.561633 | 0.03283007 | -1.986833 | 1 |
| ENSG00000196867.7 | ZFP28 | -0.654023 | 0.6214382 | 0.877569 | 0.6527736 | -1.538461 | 0.8100781 | 3.382336 | 0.02591905 | 0.061017 | 0.4917259 | -1.371255 | 1 |
| ENSG00000196876.13 | SCN8A | -1.761345 | 0.6214382 | -1.05376 | 0.4961426 | 1.127781 | 0.9582428 | 2.033286 | 0.03242844 | 2.220525 | 0.005808237 | -3.062577 | 0.8479431 |
| ENSG00000196912.12 | ANKRD36B | -0.704479 | 0.714717 | 0.88227 | 0.6527736 | 0.676948 | 0.2879524 | 2.052814 | 0.02172405 | 2.427563 | 0.001178701 | -2.127218 | 0.6885065 |
| ENSG00000196924.14 | FLNA | -0.812479 | 0.4562592 | 0.079633 | 0.9768713 | -0.492645 | 0.005011579 | 1.367004 | 0.000351591 | 0.930683 | 0.03738376 | -0.21981 | 0.9375673 |
| ENSG00000196935.8 | SRGAP1 | -1.870893 | 0.06581042 | -0.025202 | 0.9676809 | 0.250889 | 1 | 1.508913 | 1.25E-05 | 1.438853 | 0.1754266 | 0.115182 | 0.7241869 |
| ENSG00000196950.13 | SLC39A10 | -0.402658 | 0.9506557 | -0.411859 | 0.5953732 | 1.1898 | 0.02096472 | 1.355032 | 0.1903798 | -0.06396 | 0.3132498 | 1.016363 | 0.9147227 |
| ENSG00000196961.12 | AP2A1 | 0.690763 | 0.8860308 | -0.055856 | 0.8203105 | -0.163404 | 0.5290845 | 0.854837 | 0.042941 | 2.248651 | 0.2960559 | -0.140562 | 1 |
| ENSG00000196976.6 | LAGE3 | 0.714499 | 0.2744257 | -0.402299 | 0.5350015 | -1.120335 | 1.93E-08 | -0.498017 | 0.2096041 | 0.242562 | 0.95261 | -0.147768 | 0.8949489 |
| ENSG00000197013.9 | ZNF429 | 0.662025 | 0.8648814 | 0.683717 | 0.6527736 | 1.297791 | 0.04360461 | -0.300049 | 0.4576547 | 0.418756 | 0.9383375 | -3.448601 | 0.6829816 |
| ENSG00000197050.10 | ZNF420 | 1.574696 | 0.8867323 | -0.414058 | 0.7785668 | -0.262868 | 0.9115611 | 3.30837 | 0.003770254 | -1.039403 | 0.9537862 | -3.647254 | 0.5733051 |
| ENSG00000197061.4 | HIST1H4C | 0.91143 | 1.65E-07 | -0.61699 | 0.1245072 | 1.210591 | 1.94E-12 | -1.342115 | 0.02485579 | 0.935994 | 1 | -2.966053 | 0.736021 |
| ENSG00000197063.10 | MAFG | -1.281281 | 0.02363633 | 0.201617 | 0.7720357 | 0.435636 | 0.1188688 | 0.204299 | 0.4015684 | 0.914554 | 0.04196275 | 0.637932 | 1 |
| ENSG00000197102.10 | DYNC1H1 | -0.411315 | 0.6214382 | -0.04658 | 0.02727976 | -0.064898 | 0.09046443 | 1.182459 | 2.28E-05 | 1.337502 | 0.2126543 | 0.406764 | 0.6885065 |
| ENSG00000197111.15 | PCBP2 | 0.454196 | 0.2588298 | -0.069368 | 0.6527736 | -0.370015 | 5.95E-07 | 0.748855 | 0.000431472 | 0.627949 | 0.5360904 | -0.064683 | 0.6185816 |
| ENSG00000197119.12 | SLC25A29 | -1.186482 | 0.224833 | -0.044311 | 1 | 0.07905 | 0.1610041 | 1.312459 | 0.00015806 | 1.087753 | 0.2322731 | 0.702215 | 0.826754 |
| ENSG00000197121.14 | PGAP1 | -0.354231 | 0.9864272 | 0.276883 | 0.704767 | 0.928855 | 0.5290845 | 3.059648 | 3.09E-09 | 0.943537 | 0.01947854 | 2.024999 | 0.8213017 |
| ENSG00000197122.11 | SRC | 0.406639 | 1 | 0.413467 | 0.9407618 | -0.6397 | 0.5290845 | 1.921636 | 0.000110251 | 1.532786 | 0.02059149 | -0.040119 | 1 |
| ENSG00000197147.12 | LRRC8B | 0.221283 | 0.8006286 | -0.240905 | 0.6173703 | -0.336198 | 0.6028266 | 1.40859 | 0.000567377 | 3.155919 | 0.004562732 | 0.647071 | 0.9147227 |
| ENSG00000197157.10 | SND1 | -0.799333 | 0.9653624 | 0.521458 | 0.8528295 | 0.535876 | 0.01188286 | 0.162417 | 0.4315465 | 0.471446 | 0.6138391 | -0.645257 | 1 |
| ENSG00000197170.9 | PSMD12 | -0.191818 | 0.6214382 | 0.278566 | 0.9439293 | 0.630542 | 0.000251736 | 1.187398 | 0.03367078 | -0.033277 | 0.5196843 | 0.31441 | 0.2353643 |
| ENSG00000197181.11 | PIWIL2 | 0.59793 | 1 | 2.653655 | 0.6527736 | 0.69665 | 0.8825298 | 1.216258 | 0.1292522 | 3.00626 | 0.04336877 |  |  |
| ENSG00000197183.12 | NOL4L | -0.374628 | 0.8006286 | -1.030536 | 0.5671096 | -0.264586 | 0.919502 | 0.933903 | 0.004578293 | 2.386357 | 0.000954018 | -1.995941 | 0.7724671 |
| ENSG00000197238.4 | HIST1H4J | -1.067788 | 0.6014343 | 0.173478 | 0.8217443 | 1.639706 | 0.02823473 | 0.46026 | 0.8395525 | -2.115352 | 0.5196843 | -1.846102 | 0.9428375 |
| ENSG00000197299.10 | BLM | -0.372248 | 0.955833 | -0.876103 | 0.01624684 | 0.492157 | 0.03176686 | -0.383251 | 0.8223245 | 1.487657 | 0.3382129 | -1.422315 | 0.6885065 |
| ENSG00000197312.11 | DDI2 | -1.340143 | 0.6169321 | 1.001551 | 0.6514723 | 0.52978 | 0.6163624 | 0.376697 | 0.000444582 | 1.688608 | 0.6812141 | -0.819696 | 0.9374883 |
| ENSG00000197323.10 | TRIM33 | -1.459091 | 0.6214382 | -0.566543 | 0.5395105 | 2.882899 | 0.000735511 | 2.427231 | 6.42E-05 | 1.972973 | 0.01955383 | -1.023668 | 0.6200022 |
| ENSG00000197343.10 | ZNF655 | -0.255237 | 1 | 0.014658 | 0.7647276 | 0.501628 | 0.3995137 | 2.017158 | 1.03E-05 | 2.204293 | 0.000491145 | 1.067132 | 0.9453262 |
| ENSG00000197345.12 | MRPL21 | -0.766792 | 0.6559753 | -0.053395 | 0.8701081 | -0.747654 | 3.20E-07 | 0.155298 | 0.4191775 | -0.533507 | 0.003658915 | 0.620112 | 0.5160522 |
| ENSG00000197372.9 | ZNF675 | -2.140622 | 0.4764974 | -0.140143 | 0.9004336 | 2.273394 | 0.01483275 | 1.014043 | 0.6331786 | 0.173593 | 0.5192962 | 1.736298 | 0.9147227 |
| ENSG00000197380.10 | DACT3 | -0.161412 | 0.8587992 | -0.571 | 0.8515885 | 0.226082 | 0.9724248 | 2.210088 | 0.01475099 | 0.805715 | 0.06448943 | 0.150024 | 0.9663203 |
| ENSG00000197451.10 | HNRNPAB | -0.63065 | 0.2821482 | 0.020573 | 0.6527736 | 0.264259 | 0.5290845 | 0.748348 | 0.001894308 | 1.523431 | 2.83E-07 | -0.329547 | 0.1000281 |
| ENSG00000197457.9 | STMN3 | 0.625534 | 0.021504 | 0.188204 | 0.8263112 | -0.210003 | 0.5322465 | 0.869573 | 0.03040355 | 1.019963 | 0.07366152 | 0.454042 | 0.9353551 |
| ENSG00000197498.12 | RPF2 | -0.173234 | 0.6956464 | 0.331591 | 0.8628908 | 0.127408 | 0.4064599 | -0.137218 | 0.05070586 | -0.150116 | 9.77E-05 | -0.737385 | 0.8949489 |
| ENSG00000197530.12 | MIB2 | -2.083831 | 0.6169802 | -0.057284 | 0.9069847 | -2.014458 | 0.001019573 | -0.576676 | 1 | 0.583569 | 0.6367341 | 1.482353 | 1 |
| ENSG00000197535.14 | MYO5A | 0.608798 | 0.6002902 | 0.449557 | 0.7894251 | 0.418323 | 0.9974923 | 1.319566 | 0.000352765 | -0.282712 | 0.1742426 | 0.011708 | 0.9022264 |
| ENSG00000197601.12 | FAR1 | 0.224889 | 1 | 0.11974 | 0.8703522 | 0.706751 | 0.1604151 | 0.467568 | 0.02097249 | 1.908305 | 0.02084092 | -0.611147 | 1 |
| ENSG00000197603.13 | C5orf42 | 0.123485 | 0.3439898 | 0.019426 | 0.5953732 | 1.087908 | 0.0332374 | 0.657386 | 0.1825384 | 0.822973 | 0.4309406 | 2.381467 | 1 |
| ENSG00000197622.12 | CDC42SE1 | 0.085929 | 1 | 0.013851 | 0.7695736 | -0.251318 | 0.1790766 | 1.19562 | 4.07E-06 | 1.828296 | 0.001555174 | -0.21202 | 0.6885065 |
| ENSG00000197694.13 | SPTAN1 | 0.384228 | 1 | -0.037824 | 0.5647148 | -0.153526 | 0.7872586 | 1.636991 | 5.99E-05 | 0.429582 | 0.4607525 | 0.384809 | 0.8479431 |
| ENSG00000197696.9 | NMB | -0.17209 | 0.7411968 | -0.033759 | 0.9791082 | -1.965749 | 2.05E-06 | -1.158692 | 3.13E-05 | 1.130883 | 0.5440486 | 1.180576 | 0.7240243 |
| ENSG00000197728.9 | RPS26 | -0.654804 | 7.45E-05 | 0.386996 | 0.121014 | -0.689891 | 1.87E-29 | 0.015135 | 0.3760083 | -0.513609 | 3.74E-14 | -0.496279 | 0.006923841 |
| ENSG00000197746.13 | PSAP | -0.076416 | 0.02602047 | 0.273525 | 0.05028446 | 0.373528 | 0.0390447 | 0.321006 | 0.5299916 | -0.329934 | 0.001613143 | 0.588112 | 0.5122384 |
| ENSG00000197747.8 | S100A10 | -0.511996 | 0.8689227 | 0.022798 | 0.8895089 | -0.639776 | 4.82E-05 | -0.501167 | 1.82E-07 | 0.43687 | 0.008053024 | -0.341386 | 0.2959482 |
| ENSG00000197756.9 | RPL37A | -0.415119 | 0.2593412 | -0.019187 | 0.8732621 | -0.310564 | 3.12E-09 | 0.300582 | 0.005348797 | 0.143014 | 0.9815603 | -0.028114 | 0.5417688 |
| ENSG00000197780.9 | TAF13 | 1.304894 | 0.3372448 | 0.092902 | 0.681763 | -0.531307 | 0.0173006 | 0.894733 | 0.2623467 | 0.571111 | 1 | 0.458963 | 0.9068578 |
| ENSG00000197822.10 | OCLN | -1.513752 | 0.8006286 | 0.300928 | 0.6498315 | 0.454498 | 0.8454005 | 0.591987 | 0.2335397 | 3.366133 | 0.02162527 | inf | 0.6885065 |
| ENSG00000197852.9 | FAM212B | 0.565694 | 0.8806818 | 0.052625 | 0.6532996 | 0.782751 | 0.8578018 | 1.938288 | 0.004185528 | 0.803085 | 0.3573222 | 0.080105 | 0.8479431 |
| ENSG00000197858.10 | GPAA1 | 1.997962 | 0.01793817 | -0.505307 | 0.4519375 | 0.298335 | 0.6784963 | 1.302887 | 0.2383056 | -0.23436 | 0.5243319 | 0.047977 | 0.3746505 |
| ENSG00000197860.9 | SGTB | 0.192273 | 0.9492727 | 0.286511 | 0.8485366 | 1.219468 | 0.1605261 | 1.615966 | 0.000687068 | 1.575985 | 0.3978836 | -0.652603 | 0.9011704 |
| ENSG00000197892.12 | KIF13B | -0.898122 | 1 | -0.325212 | 0.6077765 | -0.602553 | 0.02395534 | 1.111387 | 0.3194824 | -0.876093 | 0.7377262 | 2.219803 | 1 |
| ENSG00000197912.13 | SPG7 | -1.671084 | 0.07871787 | -0.287032 | 1 | -0.193976 | 0.5353066 | 0.630515 | 6.38E-14 | 1.434494 | 0.000696277 | 0.029782 | 0.6885065 |
| ENSG00000197930.12 | ERO1A | 0.018911 | 0.8006286 | 0.527115 | 0.03750986 | -0.042178 | 0.7437308 | 0.948379 | 0.3110267 | 1.865251 | 0.578483 | 1.330049 | 0.5653062 |
| ENSG00000197937.12 | ZNF347 | 2.51327 | 0.6214382 | 0.229987 | 0.6353314 | 0.360529 | 0.6229203 | 2.174569 | 0.009766286 | -0.224573 | 0.8692148 | -0.445952 | 0.6885065 |
| ENSG00000197956.9 | S100A6 | 0.240072 | 0.1899814 | -0.630514 | 1.09E-25 | -0.592691 | 1.72E-19 | 0.116719 | 0.4202431 | -0.362989 | 0.0100413 | -0.379288 | 0.01006513 |
| ENSG00000197958.12 | RPL12 | -0.089293 | 0.563505 | -0.14751 | 0.3903174 | 0.313629 | 0.01128174 | -0.117141 | 0.03796887 | -0.424137 | 8.81E-08 | 0.092011 | 0.5344421 |
| ENSG00000197959.13 | DNM3 | -1.748544 | 0.6014343 | -1.42195 | 0.5512166 | 2.037968 | 0.000736286 | -0.031601 | 0.978347 | 0.371024 | 0.9571155 | -2.890525 | 1 |
| ENSG00000197961.11 | ZNF121 | 0.099725 | 1 | -0.167829 | 0.9631017 | 0.331651 | 0.06996973 | 1.428884 | 0.000134283 | -0.016029 | 0.1401461 | 1.453102 | 0.6829816 |
| ENSG00000197965.11 | MPZL1 | 1.582869 | 0.01740848 | -0.623268 | 0.1919448 | -0.009371 | 0.8647023 | -0.416033 | 0.9608791 | 1.911776 | 0.003433527 | 1.358709 | 0.7651872 |
| ENSG00000197982.13 | C1orf122 | -0.682139 | 0.9183025 | -0.177966 | 0.2863181 | -1.064769 | 6.41E-10 | -0.089331 | 0.412612 | -0.207061 | 0.311879 | 0.860403 | 0.6885065 |
| ENSG00000198000.11 | NOL8 | -0.363408 | 0.6616751 | 0.125584 | 0.9449335 | 1.300152 | 0.005837178 | 1.446353 | 0.001827364 | -0.046827 | 0.363108 | -2.233719 | 0.6885065 |
| ENSG00000198026.7 | ZNF335 | -0.771234 | 0.8587992 | -0.786201 | 0.6527736 | -1.455048 | 0.4302127 | 0.25412 | 0.5441302 | 2.312532 | 0.01104711 | 2.159877 | 0.4578379 |
| ENSG00000198034.10 | RPS4X | -0.532777 | 0.7736255 | 0.047781 | 0.9973179 | 0.251248 | 0.002479257 | 0.439626 | 0.1876965 | 0.340924 | 0.9308614 | 0.071538 | 0.5733051 |
| ENSG00000198089.14 | SFI1 | -0.001613 | 0.8006286 | -0.152024 | 0.8314425 | -0.955428 | 0.8220688 | 1.046552 | 0.01968523 | 1.44299 | 0.05854415 | 0.632913 | 0.8681734 |
| ENSG00000198146.4 | ZNF770 | -0.114981 | 0.563505 | 0.008778 | 0.829338 | 0.685442 | 0.4626116 | 0.761709 | 0.01717445 | 0.274796 | 0.8768974 | 0.139086 | 0.97072 |
| ENSG00000198160.14 | MIER1 | -0.577686 | 0.8587992 | -0.272315 | 0.3444625 | 1.286964 | 0.000897894 | 0.479 | 0.797431 | 1.035606 | 0.9069399 | -0.594208 | 0.6885065 |
| ENSG00000198162.12 | MAN1A2 | 1.028814 | 0.714717 | -0.564136 | 0.6527736 | 1.631064 | 0.000121848 | 0.506074 | 0.1059339 | 0.212632 | 0.9629001 | -0.759222 | 0.7963477 |
| ENSG00000198168.8 | SVIP | 1.510299 | 0.001807704 | -0.193565 | 0.8263112 | -0.431295 | 0.2240475 | 0.379744 | 0.3565431 | -0.507057 | 0.05183808 | -1.047168 | 0.744192 |
| ENSG00000198171.12 | DDRGK1 | -0.882165 | 0.4562592 | -0.054482 | 0.960561 | -0.514659 | 0.002035014 | 0.149833 | 0.07588336 | 0.665838 | 0.9638747 | 1.388935 | 0.8479431 |
| ENSG00000198189.10 | HSD17B11 | 2.632157 | 0.563505 | 0.177297 | 0.8846103 | 0.082912 | 0.7437308 | -0.507907 | 0.001374448 | 0.060696 | 0.6107287 | -0.871744 | 0.6885065 |
| ENSG00000198208.11 | RPS6KL1 | 0.060006 | 1 | -1.158445 | 0.8263112 | 1.500129 | 0.8966068 | 3.234829 | 0.002163125 | 0.568614 | 0.4108092 | 3.330192 | 0.6517664 |
| ENSG00000198218.10 | QRICH1 | 0.008116 | 0.8006286 | 0.106809 | 0.7669248 | -0.049452 | 0.9372061 | 0.567515 | 0.004297744 | 1.674598 | 0.04238114 | -0.063739 | 0.9147227 |
| ENSG00000198225.5 | FKBP1C | 0.916817 | 0.1633478 | 0.210496 | 0.8528295 | -1.042324 | 1.02E-05 | -0.514072 | 0.009744773 | 0.936294 | 0.8536581 | -0.310733 | 0.6885065 |
| ENSG00000198231.12 | DDX42 | -0.701706 | 0.3667121 | -0.004096 | 0.6146331 | 1.097636 | 0.000261407 | 0.751189 | 0.1638109 | -0.189646 | 0.7377262 | -0.59384 | 1 |
| ENSG00000198242.13 | RPL23A | -0.507137 | 0.4764974 | 0.094737 | 0.9726088 | -0.178023 | 0.003502914 | -0.00898 | 0.01243118 | 0.10487 | 0.2047158 | -0.362192 | 0.0620743 |
| ENSG00000198258.10 | UBL5 | -1.53534 | 0.3927055 | 0.158138 | 0.5905303 | -0.422451 | 1.46E-17 | -0.084874 | 3.88E-05 | -0.114925 | 0.008561424 | -0.337436 | 0.2159565 |
| ENSG00000198265.11 | HELZ | 1.663184 | 0.6214382 | -0.049371 | 0.9481419 | 0.4175 | 0.001286635 | 1.109175 | 0.001843879 | 1.302213 | 0.1401461 | -1.878019 | 0.6885065 |
| ENSG00000198301.11 | SDAD1 | -0.248314 | 0.7031212 | 0.109728 | 0.9408634 | 0.608292 | 0.03328468 | 1.930086 | 0.08403524 | -0.393647 | 0.0597997 | -0.296721 | 0.6829816 |
| ENSG00000198355.4 | PIM3 | 0.872255 | 0.529676 | -0.073998 | 0.7881421 | -0.658933 | 4.76E-05 | 0.575964 | 1.70E-06 | 0.984027 | 0.001277525 | 0.33047 | 0.9704101 |
| ENSG00000198356.11 | ASNA1 | 0.541983 | 0.6014343 | 0.079499 | 0.6527736 | -0.284881 | 9.67E-06 | 0.832029 | 0.05976993 | 0.046019 | 0.7377262 | -0.056072 | 0.5733051 |
| ENSG00000198373.12 | WWP2 | -0.904985 | 0.7736255 | -0.190603 | 0.8468859 | 0.018734 | 0.9333796 | 0.182686 | 0.04121412 | 3.113493 | 0.1415112 | -0.583504 | 0.8479431 |
| ENSG00000198393.7 | ZNF26 | -0.222435 | 0.8689227 | 0.438649 | 0.8532785 | 0.399792 | 0.3871766 | 1.256756 | 0.03362496 | 1.54264 | 0.3630456 | 3.653806 | 0.5733051 |
| ENSG00000198399.14 | ITSN2 | -0.806882 | 0.6014343 | 0.122636 | 0.9343836 | 0.836963 | 0.3636414 | 2.462753 | 0.000170983 | 0.845386 | 0.3812741 | -3.731464 | 0.5193439 |
| ENSG00000198408.13 | MGEA5 | -0.981468 | 0.01653545 | -0.166288 | 0.2810191 | 0.292413 | 0.000112394 | 0.8676 | 0.001147056 | 1.001511 | 0.0317483 | 0.066842 | 0.6885065 |
| ENSG00000198420.9 | TCAF1 | -0.138578 | 0.8006902 | -0.127054 | 0.7105285 | 0.802012 | 9.09E-08 | 1.001649 | 1.70E-06 | 0.534226 | 0.1401461 | 0.73203 | 0.8917781 |
| ENSG00000198431.15 | TXNRD1 | -0.153119 | 0.8650222 | 0.746849 | 0.01515307 | 0.584829 | 0.1167096 | 0.96484 | 0.01272241 | 0.950332 | 0.963799 | 0.741024 | 0.9822287 |
| ENSG00000198440.9 | ZNF583 | -1.120135 | 0.3946735 | 0.242018 | 0.6527736 | 1.154114 | 0.6928639 | 2.674101 | 0.04221781 | 0.360534 | 1 | -3.20504 | 1 |
| ENSG00000198467.13 | TPM2 | -2.703978 | 0.6214382 | 0.462606 | 0.3644419 | -1.836655 | 0.8369428 | 0.540205 | 0.8034388 | 0.071906 | 0.003204401 | 1.359031 | 0.7788913 |
| ENSG00000198478.7 | SH3BGRL2 | 1.002454 | 0.6214382 | 2.067111 | 0.8703522 | -1.844564 | 0.5290845 | 1.86205 | 0.009224418 | 0.583401 | 0.8176292 | -4.592836 | 0.6885065 |
| ENSG00000198492.14 | YTHDF2 | -1.316854 | 0.7117094 | -0.125538 | 0.9296311 | 0.52181 | 0.00365647 | 0.310834 | 0.7225458 | 0.202002 | 0.790426 | 0.296744 | 0.9973062 |
| ENSG00000198498.9 | TMA16 | 0.307856 | 0.6214382 | -0.057296 | 0.5517607 | 0.162534 | 0.7100718 | -0.107651 | 0.4515519 | -1.30333 | 0.02609751 | 0.040543 | 0.7918118 |
| ENSG00000198561.13 | CTNND1 | -0.358744 | 0.8006286 | -0.631135 | 0.8703522 | 0.249089 | 0.2408782 | 0.100263 | 0.5088602 | 0.714757 | 0.03387033 | 1.900014 | 0.6886945 |
| ENSG00000198586.13 | TLK1 | -4.3715 | 0.9183025 | -0.134672 | 0.2801475 | 0.966824 | 0.01029158 | 2.337896 | 0.02552731 | -0.569624 | 0.7114696 | 1.723104 | 0.6284961 |
| ENSG00000198604.10 | BAZ1A | -0.427193 | 0.6214382 | -0.488571 | 0.6417538 | 0.325289 | 0.3180431 | 0.94609 | 0.002939922 | 0.447038 | 0.7910583 | 0.43013 | 0.9467893 |
| ENSG00000198612.10 | COPS8 | 0.821752 | 0.3694036 | -0.006474 | 0.9028136 | -0.176085 | 0.5775267 | -0.334926 | 0.02929256 | 0.639035 | 0.301317 | -0.324774 | 0.8345878 |
| ENSG00000198625.12 | MDM4 | -1.052832 | 0.6214382 | 0.090994 | 0.8113877 | 0.604829 | 0.007843297 | 1.609053 | 0.000688376 | 1.056332 | 0.0169801 | -1.054778 | 0.6185816 |
| ENSG00000198646.13 | NCOA6 | -0.848601 | 0.5711449 | 0.311472 | 0.8953649 | -0.183698 | 0.5837381 | 0.974597 | 0.001133042 | 0.949429 | 0.008061688 | -0.1529 | 0.6185816 |
| ENSG00000198648.10 | STK39 | -0.436431 | 0.8006286 | -0.384183 | 0.401251 | 0.15301 | 0.0869859 | 1.267849 | 0.02028292 | 2.745058 | 0.000121003 | 1.117481 | 0.7595705 |
| ENSG00000198663.16 | C6orf89 | -2.786101 | 0.2012287 | -0.355358 | 0.6527736 | 0.560967 | 0.5131304 | 0.289091 | 0.007776291 | -1.587453 | 0.5196843 | -0.626849 | 0.9147227 |
| ENSG00000198668.10 | CALM1 | -1.615255 | 0.08456749 | -0.32446 | 9.19E-06 | 0.396382 | 4.51E-05 | 0.18978 | 0.7279279 | -0.206109 | 0.000556679 | -0.314603 | 0.6885065 |
| ENSG00000198677.10 | TTC37 | -0.275547 | 0.8867323 | 0.210096 | 0.9451258 | 0.603494 | 0.004020273 | 0.399709 | 0.8537667 | -0.344401 | 0.1617117 | -2.178061 | 0.5736188 |
| ENSG00000198680.4 | TUSC1 | -2.302976 | 0.8587992 | 0.168666 | 0.9832381 | 0.023376 | 0.5290845 | 2.349343 | 0.02034791 | 2.281988 | 0.2065601 | -2.386575 | 0.5008376 |
| ENSG00000198690.9 | FAN1 | 0.590544 | 0.8006286 | -0.098485 | 1 | 0.563569 | 0.8100781 | 2.269911 | 0.0272166 | 1.694729 | 0.2799994 | 3.15095 | 0.7260077 |
| ENSG00000198700.9 | IPO9 | 0.679415 | 1 | -0.083653 | 0.7958166 | 0.255249 | 0.5449348 | 0.586642 | 0.02536093 | 0.740482 | 0.4988048 | 0.618197 | 1 |
| ENSG00000198707.14 | CEP290 | 0.09765 | 0.529676 | -0.176065 | 0.2185295 | 1.469556 | 0.04239198 | 1.640474 | 0.1146644 | 0.973883 | 0.8387831 | 0.602869 | 0.9147227 |
| ENSG00000198719.8 | DLL1 | 2.349607 | 0.6214382 | -0.316251 | 0.8872857 | 0.212663 | 0.8322087 | -0.37309 | 0.9013275 | 1.454823 | 0.03056663 | -4.212593 | 0.6829816 |
| ENSG00000198720.12 | ANKRD13B | -1.610855 | 0.6214382 | 1.024634 | 0.5953732 | -0.396388 | 0.3276184 | 1.817734 | 0.002986579 | 2.735174 | 0.004711842 | -0.810511 | 0.9062221 |
| ENSG00000198721.12 | ECI2 | 0.088447 | 0.8006286 | -0.03603 | 0.9081785 | -0.080461 | 0.9428466 | 1.054237 | 0.02950643 | 0.31363 | 0.4014141 | 1.03415 | 0.9147227 |
| ENSG00000198740.8 | ZNF652 | -0.433904 | 1 | 0.094586 | 0.5010139 | 0.176796 | 0.5290845 | 1.917113 | 0.0412511 | 1.528589 | 0.006137746 | -0.464139 | 0.6885065 |
| ENSG00000198742.9 | SMURF1 | -0.714526 | 0.6837569 | 0.052577 | 0.6527736 | -0.368099 | 0.5376848 | 0.317501 | 0.04939234 | 1.721288 | 0.08496185 | 1.440775 | 0.9565287 |
| ENSG00000198743.5 | SLC5A3 | -0.453135 | 0.934454 | -0.456453 | 0.6030937 | 1.630545 | 0.03807631 | 1.356047 | 0.2835711 | 0.42408 | 0.01087007 | 0.949583 | 0.8949489 |
| ENSG00000198752.10 | CDC42BPB | 0.313897 | 0.9599095 | 0.329327 | 0.7335559 | 0.254845 | 0.5681115 | 1.171215 | 0.03106038 | 0.955075 | 0.2601785 | -1.598433 | 0.8949489 |
| ENSG00000198755.10 | RPL10A | -0.609115 | 0.09114542 | 0.005485 | 0.6688577 | -0.100101 | 0.870773 | -0.072863 | 0.001116302 | 0.22582 | 0.3250692 | -0.159076 | 0.2768456 |
| ENSG00000198792.12 | TMEM184B | -0.632849 | 0.8006286 | -0.479687 | 0.9452791 | 0.577836 | 0.9140472 | 2.350517 | 0.001921065 | 1.295213 | 0.006124632 | 1.004961 | 0.7767294 |
| ENSG00000198793.12 | MTOR | 0.249544 | 1 | 0.360041 | 0.9476887 | 0.404327 | 0.724574 | 1.859597 | 0.007265777 | 1.480465 | 0.01508558 | 0.064099 | 0.6829816 |
| ENSG00000198794.11 | SCAMP5 | -0.652831 | 0.6214382 | 0.368015 | 0.6200225 | -1.122326 | 0.6992619 | 0.876299 | 0.04743104 | 1.804082 | 0.04614451 | -0.104729 | 0.6185816 |
| ENSG00000198799.11 | LRIG2 | 0.022925 | 0.8006286 | 0.419303 | 0.6428284 | 1.2083 | 0.03715969 | 1.196481 | 0.2521743 | 1.473389 | 0.4242927 | 0.680306 | 0.8649436 |
| ENSG00000198805.11 | PNP | 1.39082 | 0.6214382 | -0.675594 | 0.7881421 | -0.806615 | 0.7437308 | 3.028466 | 0.008249854 | 1.820852 | 0.7573677 | -1.825504 | 1 |
| ENSG00000198815.8 | FOXJ3 | -0.272455 | 0.9346111 | -0.460233 | 0.434287 | 0.578897 | 0.01826961 | 1.499007 | 0.01506498 | 0.017775 | 0.5196843 | 1.48825 | 0.9568643 |
| ENSG00000198816.5 | ZNF358 | -1.55413 | 0.6014343 | 0.176119 | 0.7472055 | -1.107838 | 0.03042647 | 1.195973 | 0.003891416 | 1.549231 | 0.2510112 | 1.416081 | 0.782216 |
| ENSG00000198818.9 | SFT2D1 | -1.651276 | 0.422462 | 0.32469 | 0.5956071 | -0.31853 | 0.1666996 | 0.561705 | 0.5037188 | -0.281397 | 0.04889758 | -1.414966 | 0.7550414 |
| ENSG00000198832.10 | SELM | -0.535297 | 1 | -1.171685 | 0.6828476 | -1.688093 | 0.004999539 | -0.688537 | 0.02918812 | 0.778396 | 0.6883566 | 1.923884 | 0.8479431 |
| ENSG00000198836.8 | OPA1 | -0.886666 | 0.6214382 | 0.248734 | 0.872873 | 0.905415 | 0.03605421 | 1.871029 | 0.01252251 | 0.536609 | 1 | 1.394427 | 0.7385207 |
| ENSG00000198846.5 | TOX | -0.057123 | 0.9311232 | -0.063176 | 0.8587199 | -0.042682 | 1 | 0.667859 | 0.002813711 | 0.910033 | 0.1462636 | 1.196556 | 0.6885065 |
| ENSG00000198862.13 | LTN1 | -1.347435 | 0.8006286 | 0.071217 | 1 | 0.388465 | 0.01649772 | 1.545846 | 0.6498745 | -0.393991 | 0.3170597 | -0.814448 | 0.6885065 |
| ENSG00000198863.7 | RUNDC1 | -0.733219 | 0.9282145 | 0.923632 | 0.6527736 | 0.554254 | 0.4034134 | 3.073 | 0.001887814 | 1.6727 | 0.3291949 | -0.798474 | 0.8479431 |
| ENSG00000198876.12 | DCAF12 | -1.274863 | 0.6014343 | -0.677486 | 0.6527736 | 0.097876 | 0.67274 | 1.452134 | 0.0307211 | 0.604497 | 0.9782894 | -1.304559 | 0.736021 |
| ENSG00000198879.11 | SFMBT2 | 1.78227 | 1 | -0.101847 | 0.9329105 | -1.064703 | 0.6538987 | 1.755315 | 0.007660097 | -1.046696 | 0.4818826 | 2.8958 | 0.8479431 |
| ENSG00000198887.8 | SMC5 | -0.36097 | 0.9749561 | -0.226546 | 0.6436071 | 1.246282 | 1.77E-05 | 0.521228 | 0.09855916 | -0.00287 | 0.8700639 | 2.52291 | 0.7496444 |
| ENSG00000198892.6 | SHISA4 | -0.517104 | 0.9126213 | -0.883914 | 0.5010139 | -1.094834 | 0.000277782 | -0.312463 | 0.3362087 | 0.673558 | 0.8638047 | -0.657262 | 0.826754 |
| ENSG00000198898.12 | CAPZA2 | 1.737083 | 0.002536564 | -0.182779 | 0.4626812 | 0.193241 | 0.6987919 | 0.576823 | 0.9618755 | -0.815871 | 1.84E-06 | 1.087057 | 0.9638718 |
| ENSG00000198900.5 | TOP1 | -0.111896 | 0.6214382 | -0.175728 | 0.7371871 | 0.867035 | 0.001125165 | 0.300093 | 0.1148418 | 0.243639 | 0.4286052 | -0.532389 | 0.7468809 |
| ENSG00000198909.7 | MAP3K3 | -1.228641 | 0.8587992 | -1.241079 | 0.6527736 | 2.867201 | 0.123539 | 6.562763 | 0.008309395 | 3.547879 | 0.06813193 | 0.829551 | 0.6552711 |
| ENSG00000198911.11 | SREBF2 | 1.502646 | 0.08844455 | 0.737534 | 0.7517008 | 0.16928 | 0.8308365 | 0.717473 | 0.0100786 | 0.049306 | 0.03327729 | 1.295566 | 0.6185816 |
| ENSG00000198914.2 | POU3F3 | -1.824793 | 0.01499139 | -0.099011 | 0.4833764 | 0.115007 | 0.3276184 | 0.976227 | 0.001340844 | 0.533514 | 0.9990698 | -0.232583 | 0.9169962 |
| ENSG00000198918.7 | RPL39 | 0.303146 | 0.001829282 | -0.112199 | 0.4324917 | -0.048575 | 0.2150986 | -0.193001 | 5.51E-05 | -0.117322 | 0.00189447 | -0.1055 | 0.2123159 |
| ENSG00000198919.12 | DZIP3 | 0.964282 | 0.8726783 | 0.543048 | 0.906244 | 0.474819 | 0.2652894 | -0.028557 | 0.01859259 | 1.963385 | 0.09136799 | -1.534368 | 0.7315846 |
| ENSG00000198931.10 | APRT | 1.02613 | 0.01499139 | 0.11706 | 0.8307191 | -0.278817 | 0.1182794 | -0.386883 | 0.01846326 | 0.263321 | 0.576218 | 0.960278 | 0.6885065 |
| ENSG00000198933.9 | TBKBP1 | 1.048731 | 0.9198482 | 0.251248 | 0.9711003 | -1.136612 | 0.6545047 | 1.541541 | 0.002860427 | 0.949578 | 0.621723 | -0.696855 | 0.9368176 |
| ENSG00000198937.8 | CCDC167 | 0.164599 | 0.529676 | 0.060187 | 0.6527736 | -0.358333 | 0.000190189 | 0.026073 | 0.07032724 | -0.181615 | 0.01187671 | -1.216455 | 0.09856287 |
| ENSG00000198952.8 | SMG5 | -0.850502 | 0.6014343 | 0.172072 | 0.8322287 | 0.61672 | 0.7354314 | 0.768448 | 0.02326622 | 1.106376 | 0.1305498 | -0.589968 | 0.8757124 |
| ENSG00000198961.9 | PJA2 | 0.474256 | 0.6214382 | 0.100628 | 0.8654277 | 1.0739 | 1.27E-10 | 0.560656 | 0.03519633 | 0.521997 | 0.9831817 | 0.107539 | 0.6885065 |
| ENSG00000198963.10 | RORB | 0.241378 | 0.8006286 | -0.202835 | 0.727891 | 1.580592 | 0.004020273 | 2.763282 | 0.1274788 | 1.291627 | 0.8888294 | 0.3244 | 0.6885065 |
| ENSG00000203667.9 | COX20 | 0.798325 | 0.4406957 | -0.199308 | 0.5395105 | -0.121992 | 0.002274442 | 0.039915 | 0.7037023 | 0.926322 | 0.5196843 | -0.744115 | 0.348294 |
| ENSG00000203778.7 | FAM229B | 0.22702 | 0.6214382 | 0.07027 | 1 | 0.358018 | 0.4064599 | 0.611948 | 0.4919368 | -1.878075 | 0.005245519 | 0.190502 | 0.6829816 |
| ENSG00000203879.11 | GDI1 | 0.4999 | 0.5657211 | -0.095143 | 0.5314455 | -0.354513 | 0.2049228 | 0.4826 | 0.007197486 | 1.001756 | 0.001459778 | -0.55008 | 0.6885065 |
| ENSG00000203880.11 | PCMTD2 | 0.549328 | 0.7979044 | 0.139774 | 0.6929602 | 0.235137 | 0.3449792 | 1.376142 | 0.02485579 | 1.136631 | 0.03327974 | 0.058669 | 0.5200188 |
| ENSG00000204120.14 | GIGYF2 | 0.162042 | 0.7391347 | -0.080063 | 0.6527736 | 1.455179 | 2.89E-09 | 1.446789 | 0.2410327 | -0.41583 | 0.5196843 | 6.273734 | 0.426223 |
| ENSG00000204178.9 | TMEM57 | 0.707248 | 0.9538934 | 0.665413 | 0.4011918 | 0.88657 | 0.1964422 | 1.224805 | 0.000516363 | 0.826289 | 0.1644677 | 1.42995 | 0.9204395 |
| ENSG00000204186.7 | ZDBF2 | -2.303714 | 0.563505 | 1.719542 | 0.5108497 | -0.407889 | 0.4382561 | 1.378 | 3.01E-05 | 0.343911 | 0.5196843 | 1.756585 | 0.6185816 |
| ENSG00000204209.10 | DAXX | 0.596447 | 0.6214382 | 0.096041 | 1 | 0.810961 | 0.1771136 | 1.667578 | 0.1779476 | 0.900026 | 0.01487653 | -0.709901 | 0.6885065 |
| ENSG00000204217.12 | BMPR2 | -0.53045 | 0.6214382 | -0.203558 | 0.4536333 | 0.985467 | 0.06625098 | 0.902805 | 0.02137679 | 0.419292 | 0.17967 | -0.453268 | 0.9127491 |
| ENSG00000204220.9 | PFDN6 | -1.286488 | 0.4764974 | 0.276503 | 0.7927491 | -0.712334 | 4.74E-08 | 0.62172 | 0.1090508 | 0.29859 | 0.3974318 | -0.494839 | 0.5066477 |
| ENSG00000204237.4 | OXLD1 | -0.107516 | 0.7568783 | 0.478661 | 0.8263112 | -0.375141 | 0.01022277 | -0.213726 | 0.4880359 | 1.016222 | 0.7494748 | 0.809573 | 0.9077431 |
| ENSG00000204256.12 | BRD2 | 0.320568 | 0.8006286 | 0.12334 | 0.8485564 | 0.411645 | 0.03852063 | 0.267661 | 0.08714575 | 1.074756 | 0.002138492 | 0.519985 | 0.6885065 |
| ENSG00000204257.14 | HLA-DMA | 0.200085 | 0.8587992 | 2.511164 | 0.02674978 | -1.583968 | 0.7437308 | -3.568667 | 0.5930297 | 1.133819 | 1 | 0.613851 | 0.6885065 |
| ENSG00000204264.8 | PSMB8 | -0.797408 | 0.6084645 | 0.039905 | 0.8528295 | -0.185616 | 0.4064599 | -0.378473 | 0.3467508 | -0.004328 | 0.01251397 | 0.667336 | 0.8479431 |
| ENSG00000204356.11 | NELFE | 0.297496 | 0.8632491 | -0.030285 | 0.9793662 | -0.437838 | 0.00354988 | 0.588437 | 0.7245061 | -0.630942 | 0.03455371 | 0.686004 | 0.6829816 |
| ENSG00000204370.8 | SDHD | -0.001098 | 0.178542 | 0.09756 | 0.6996366 | 0.359242 | 0.1991342 | -0.572083 | 0.000383314 | 0.926949 | 0.0310413 | 0.760508 | 0.9733982 |
| ENSG00000204388.6 | HSPA1B | -0.688662 | 0.6214382 | -0.751804 | 0.9766083 | 1.189931 | 7.06E-15 | 1.074873 | 0.1850495 | 2.063627 | 1.33E-11 | 0.862635 | 0.9147227 |
| ENSG00000204389.9 | HSPA1A | -0.162785 | 0.7155003 | -0.641705 | 0.8703522 | 1.481487 | 2.67E-16 | 0.202151 | 0.4953678 | 2.151817 | 2.92E-09 | -0.067862 | 1 |
| ENSG00000204392.10 | LSM2 | 0.17706 | 0.9332642 | -0.369909 | 0.2792759 | -0.255254 | 0.2337559 | -0.518077 | 0.01572454 | -0.213679 | 0.01508043 | 0.016395 | 0.6490419 |
| ENSG00000204469.12 | PRRC2A | -1.400246 | 0.4801473 | 0.561191 | 0.7508235 | -0.496263 | 0.4436285 | 1.902545 | 1.39E-05 | 2.677541 | 2.79E-13 | -0.404494 | 0.8654874 |
| ENSG00000204525.15 | HLA-C | -0.423817 | 0.3526445 | 0.536639 | 4.18E-22 | -0.67836 | 2.07E-17 | -0.037965 | 0.02741529 | -0.499437 | 1.05E-05 | 0.319207 | 0.7408601 |
| ENSG00000204569.9 | PPP1R10 | -2.824093 | 0.3563101 | 0.062317 | 1 | 1.492356 | 0.000927901 | 2.54085 | 0.002616538 | 1.485722 | 0.583236 | -0.319046 | 0.7810316 |
| ENSG00000204592.8 | HLA-E | -1.452053 | 0.162295 | 0.17604 | 0.1889019 | -0.506634 | 0.02184901 | 0.540533 | 0.7144938 | 0.332513 | 0.7242087 | 1.382832 | 0.2440556 |
| ENSG00000204628.11 | GNB2L1 | -0.821961 | 0.8006286 | -0.060432 | 0.561939 | 0.024783 | 0.8371054 | 0.035053 | 0.5156115 | -0.194229 | 3.20E-05 | 0.148747 | 0.9451588 |
| ENSG00000204642.13 | HLA-F | -3.454218 | 0.5695633 | 0.979305 | 0.2713328 | 2.920962 | 0.000312839 | -0.475472 | 7.38E-06 | -0.492399 | 0.8161922 | 0.704862 | 0.5344421 |
| ENSG00000204673.10 | AKT1S1 | 0.571962 | 0.9282145 | 0.802823 | 0.3972124 | -0.500366 | 0.000353312 | 0.502922 | 0.3541189 | 1.219072 | 0.08626379 | -0.716794 | 0.6885065 |
| ENSG00000204681.10 | GABBR1 | -1.314116 | 0.6214382 | -1.645717 | 0.1547783 | 0.746556 | 0.1181265 | 0.830588 | 0.01046955 | 0.563236 | 0.5344037 | -0.728572 | 0.6441404 |
| ENSG00000204711.8 | C9orf135 | -1.105163 | 0.8006286 | 0.469773 | 0.8863478 | -3.201672 | 0.01667615 | -2.015242 | 0.04775009 | -1.476248 | 0.5196843 | 2.959744 | 1 |
| ENSG00000204842.14 | ATXN2 | -0.468701 | 0.4218563 | 0.322128 | 0.8854415 | 0.928993 | 0.02578415 | 1.188308 | 5.88E-10 | 1.77571 | 5.84E-05 | 0.496917 | 0.9425055 |
| ENSG00000204922.4 | UQCC3 | -0.62875 | 0.6214382 | -0.701419 | 0.04220726 | 0.101343 | 0.16214 | 0.513205 | 0.2506565 | -0.366513 | 0.3809839 | -0.542564 | 0.8654874 |
| ENSG00000204946.9 | ZNF783 | -2.126017 | 0.8006286 | 0.222226 | 0.8893429 | -1.2727 | 0.3853208 | 1.563758 | 0.03242844 | 0.670635 | 0.8223699 | -2.868507 | 0.6885065 |
| ENSG00000204991.10 | SPIRE2 | -1.756693 | 0.8006286 | -1.743896 | 0.5953732 | 0.64789 | 0.7554924 | 0.833485 | 0.1414521 | 1.774365 | 0.03477713 | 3.002942 | 0.9380942 |
| ENSG00000205060.10 | SLC35B4 | 0.656436 | 0.8006286 | -0.065725 | 0.6527736 | 0.245011 | 0.7268105 | 1.333048 | 4.81E-09 | 0.561141 | 0.512206 | -0.783778 | 0.4998763 |
| ENSG00000205084.10 | TMEM231 | -2.533289 | 0.8587992 | -0.132778 | 0.9787499 | -1.124124 | 0.04447082 | 1.136905 | 0.1267073 | 0.5089 | 0.9068628 | -0.775768 | 0.9270338 |
| ENSG00000205138.3 | SDHAF1 | 0.512966 | 0.739831 | -0.302047 | 0.486097 | -1.040775 | 6.06E-05 | -0.656354 | 0.1147845 | 2.63803 | 0.8515144 | 0.848733 | 0.819756 |
| ENSG00000205155.7 | PSENEN | 1.768432 | 0.1832812 | 0.075414 | 1 | -0.50004 | 7.07E-05 | 0.145272 | 0.2752129 | 0.308724 | 0.4918689 | 0.089951 | 0.926878 |
| ENSG00000205189.11 | ZBTB10 | 0.160082 | 0.8867323 | -0.344007 | 0.5456228 | 1.024224 | 0.0582643 | 1.062996 | 0.04746704 | 0.978357 | 0.2880317 | -0.234237 | 0.9539154 |
| ENSG00000205208.4 | C4orf46 | 1.982515 | 0.8006286 | -0.249585 | 0.8953649 | -0.959721 | 0.84154 | 0.90845 | 0.03982882 | 0.419508 | 0.7377262 | -0.204064 | 0.9439641 |
| ENSG00000205336.11 | ADGRG1 | -0.7435 | 0.000617602 | -0.586063 | 0.001778822 | 0.471446 | 0.01260924 | 0.80619 | 0.04615682 | 0.906045 | 0.1816663 | 0.573988 | 0.7767294 |
| ENSG00000205339.9 | IPO7 | 0.748113 | 0.1204153 | -0.237468 | 0.7971937 | 0.719972 | 0.000174662 | 1.174932 | 2.91E-05 | 1.06433 | 0.5196843 | 1.161933 | 0.9067556 |
| ENSG00000205362.10 | MT1A | -1.2951 | 1 | -0.550552 | 0.7426112 | -0.202696 | 0.02207811 | -1.482334 | 0.2574191 | 2.338468 | 1 | #NAME? | 0.6885065 |
| ENSG00000205476.8 | CCDC85C | -1.341691 | 0.6214382 | 0.883143 | 0.6527736 | -0.205709 | 0.8454005 | 1.979253 | 5.35E-05 | 2.406254 | 0.03104635 | -3.463454 | 0.6829816 |
| ENSG00000205542.10 | TMSB4X | -0.418038 | 0.9018396 | -0.112009 | 0.1076617 | -0.309575 | 1.89E-05 | -0.390551 | 1.57E-07 | -0.772019 | 6.02E-12 | -0.07051 | 0.9914916 |
| ENSG00000205581.10 | HMGN1 | -0.12655 | 0.9335818 | -0.123624 | 0.5375948 | -0.846646 | 9.84E-07 | 0.34883 | 0.6951585 | 0.072747 | 0.3974318 | -0.117141 | 0.6829816 |
| ENSG00000205639.9 | MFSD2B | 2.111323 | 0.1835496 | 0.508467 | 0.7063369 | -2.939482 | 9.49E-07 | 0.267969 | 0.2806819 | -0.357918 | 0.6217705 |  |  |
| ENSG00000205903.6 | ZNF316 | -1.397886 | 0.6214382 | -0.367281 | 0.7882366 | -0.4973 | 0.4064599 | 1.126447 | 0.007350873 | 0.528295 | 0.03147391 | -1.294292 | 0.8118172 |
| ENSG00000205929.9 | C21orf62 | -0.807954 | 0.6014343 | -0.465527 | 0.2886194 | 1.301859 | 0.018079 | -0.220565 | 0.4919368 | 0.90771 | 0.1688763 | #NAME? | 0.6829816 |
| ENSG00000205937.11 | RNPS1 | 0.772131 | 0.005603822 | -0.118744 | 0.5736784 | -0.01687 | 0.9137126 | 0.839124 | 0.009778502 | 0.335444 | 1 | -0.098548 | 0.3746505 |
| ENSG00000205981.6 | DNAJC19 | -0.622474 | 0.6209259 | 0.193011 | 0.8515885 | -0.441895 | 0.01022277 | 0.048385 | 0.1852027 | 0.007252 | 0.0527459 | 0.947535 | 0.9351719 |
| ENSG00000206053.12 | HN1L | -1.839275 | 1 | 0.001701 | 0.5818005 | -0.029504 | 0.7668614 | 1.26274 | 0.008452094 | 1.427591 | 0.03484403 | 0.347124 | 0.9641669 |
| ENSG00000206077.10 | ZDHHC11B | -2.84415 | 0.3946735 | -0.018659 | 0.6487262 | -0.528757 | 0.578623 | -0.494824 | 0.947988 | 1.28048 | 0.001827878 | 0.933723 | 0.7427692 |
| ENSG00000206418.3 | RAB12 | -0.376796 | 0.9282145 | 0.186845 | 0.8776804 | 0.386532 | 0.6380246 | 1.723714 | 0.01631939 | -0.579267 | 0.7833673 | -0.760106 | 0.6885065 |
| ENSG00000206503.11 | HLA-A | -1.169686 | 0.000247117 | 0.403501 | 3.34E-15 | -0.345235 | 1.47E-05 | 0.018216 | 0.1490863 | -0.601564 | 6.72E-07 | 0.336741 | 0.8479431 |
| ENSG00000206527.9 | HACD2 | -1.211488 | 0.8006286 | -0.272581 | 0.7601699 | 0.249239 | 0.6025382 | 1.244761 | 0.04188543 | -0.088824 | 0.9662339 | -0.430231 | 0.9822287 |
| ENSG00000206559.7 | ZCWPW2 | 1.170388 | 0.8006286 | -0.111982 | 1 | 0.042061 | 0.5290845 | -0.806092 | 1 | -0.792968 | 0.7377262 |  |  |
| ENSG00000206560.10 | ANKRD28 | 0.901263 | 0.8567944 | -0.677039 | 0.8703522 | 0.980475 | 0.2977668 | 1.891466 | 0.007351647 | 1.346113 | 0.05231516 | -2.476449 | 0.8845948 |
| ENSG00000211450.9 | C11orf31 | -0.497505 | 1 | 0.007303 | 0.6218841 | -0.121063 | 0.01443464 | 0.499522 | 0.03788242 | -1.344546 | 0.002398345 | 0.53434 | 0.6885065 |
| ENSG00000213015.8 | ZNF580 | -1.50452 | 0.3009675 | -0.346876 | 0.5010139 | -1.543939 | 5.36E-06 | 0.947255 | 0.7144938 | 0.624937 | 0.08724925 | -1.202889 | 0.6885065 |
| ENSG00000213066.11 | FGFR1OP | -0.633618 | 0.6014343 | 0.130592 | 0.979665 | -0.179169 | 0.8089907 | 1.768387 | 0.02462981 | 1.040041 | 0.217049 | 2.017266 | 0.9581926 |
| ENSG00000213123.10 | TCTEX1D2 | 0.415558 | 0.9575547 | 0.093193 | 0.6208241 | -0.522043 | 0.001839621 | -0.317867 | 0.4242944 | -0.09456 | 0.09177453 | -2.040531 | 0.4331698 |
| ENSG00000213190.3 | MLLT11 | 0.137712 | 0.6014343 | -0.260382 | 0.6527736 | 0.218921 | 0.5290845 | 0.310197 | 0.4919368 | 0.195639 | 0.205875 | -0.450349 | 0.04053673 |
| ENSG00000213380.13 | COG8 | -0.294671 | 0.8006286 | 0.637881 | 1 | 0.23286 | 0.9508895 | 1.528763 | 0.0540036 | 3.026674 | 0.007470624 | -1.012416 | 0.9618031 |
| ENSG00000213397.10 | HAUS7 | 2.611261 | 0.714717 | -0.187143 | 0.859273 | -2.051544 | 0.008474587 | 1.974669 | 0.006245438 | 1.039776 | 0.9361679 | 1.714609 | 0.7117761 |
| ENSG00000213420.7 | GPC2 | -1.087823 | 0.6409583 | -1.149913 | 0.7383998 | -0.965146 | 0.04371205 | 0.530107 | 0.4919368 | 0.619438 | 0.6950751 | -0.2434 | 0.9536525 |
| ENSG00000213462.4 | ERV3-1 | 0.048256 | 0.8006286 | -0.82133 | 0.8127993 | -0.168528 | 0.1618397 | 3.219632 | 0.0315178 | 0.774112 | 0.01124439 | 3.061109 | 0.9264343 |
| ENSG00000213563.6 | C8orf82 | 1.170667 | 1 | -0.126179 | 0.6527736 | -0.553514 | 0.3346772 | 3.719683 | 0.03573762 | -0.259565 | 0.4278631 | 1.413506 | 0.5733051 |
| ENSG00000213585.10 | VDAC1 | -0.594657 | 0.2006576 | -0.16727 | 0.04760656 | 0.222043 | 0.2879524 | 0.254338 | 0.4264406 | -0.017366 | 0.002340489 | 0.143181 | 0.4627089 |
| ENSG00000213619.9 | NDUFS3 | -0.688608 | 0.8006286 | -0.276082 | 0.4543215 | -0.300463 | 0.01488258 | 0.274472 | 0.2484437 | -0.380881 | 0.002398345 | -0.690437 | 0.6885065 |
| ENSG00000213625.8 | LEPROT | 0.653319 | 0.6214382 | 0.516671 | 0.5394427 | 0.713789 | 1 | -0.115947 | 0.7600554 | -2.293752 | 0.0411428 | 0.295395 | 1 |
| ENSG00000213639.9 | PPP1CB | 0.889976 | 0.01711078 | 0.010907 | 0.8472233 | 0.105979 | 0.8100781 | 0.508773 | 0.4919368 | -0.350931 | 0.5196843 | 0.696428 | 1 |
| ENSG00000213699.8 | SLC35F6 | 0.044299 | 0.3698143 | -0.88565 | 0.4961426 | -0.120266 | 0.8231116 | 0.645795 | 0.02477329 | 1.296551 | 1 | -0.870212 | 0.6037129 |
| ENSG00000213722.8 | DDAH2 | -0.043608 | 1 | 0.108747 | 0.6274147 | -0.067735 | 0.000242068 | 0.697927 | 0.03668755 | 0.298168 | 0.8255932 | 0.471737 | 0.8479431 |
| ENSG00000213741.8 | RPS29 | -1.533124 | 7.45E-05 | 0.075483 | 0.7529887 | -0.685712 | 5.67E-17 | -0.008684 | 0.3825038 | 0.182392 | 0.1477952 | 0.148256 | 0.6185816 |
| ENSG00000213760.10 | ATP6V1G2 | -0.374365 | 0.6214382 | 0.722735 | 1 | 0.150522 | 0.7437308 | 0.084135 | 1 | 1.360812 | 0.02084092 | 0.934891 | 1 |
| ENSG00000213762.10 | ZNF134 | 0.817083 | 0.6214382 | -0.391748 | 1 | 0.683957 | 0.01576963 | 0.032553 | 0.02251262 | 0.517125 | 0.8899273 | 0.745512 | 0.8479431 |
| ENSG00000213918.10 | DNASE1 | -3.225632 | 0.6214382 | -0.18908 | 0.9019145 | -0.4786 | 0.8100781 | 2.227628 | 0.008498161 | 2.045685 | 0.005460428 | 0.356353 | 0.796792 |
| ENSG00000213923.10 | CSNK1E | -1.086969 | 0.4672906 | -0.471446 | 0.1186917 | 1.488701 | 2.82E-06 | 1.216653 | 7.09E-07 | 1.211423 | 1.06E-05 | 0.022395 | 0.6885065 |
| ENSG00000214022.11 | REPIN1 | 0.222936 | 0.8006286 | 0.100037 | 0.9834088 | -0.042464 | 0.5937705 | 0.432431 | 0.09710095 | 1.668792 | 0.02926165 | 0.252025 | 1 |
| ENSG00000214026.10 | MRPL23 | -1.149164 | 0.4764974 | -0.040753 | 0.8661807 | -2.315798 | 6.13E-13 | -0.181017 | 0.04577302 | 1.428633 | 0.8577035 | -1.783472 | 0.5151181 |
| ENSG00000214087.8 | ARL16 | -1.067217 | 0.682906 | 0.112659 | 0.9277629 | -0.329739 | 0.001995716 | -0.19557 | 0.2832438 | 0.557403 | 0.4399626 | 0.618112 | 0.8479431 |
| ENSG00000214160.9 | ALG3 | -0.059984 | 0.9667286 | 0.610237 | 0.6527736 | -0.187746 | 0.09251875 | -0.604555 | 0.04795421 | 0.369469 | 0.6662427 | -0.442473 | 0.7767294 |
| ENSG00000214253.8 | FIS1 | -0.731345 | 0.4764974 | -0.48519 | 0.02922882 | -0.381623 | 0.005985965 | -0.336909 | 0.4919368 | 0.380689 | 0.1226567 | 0.52518 | 0.8629701 |
| ENSG00000214357.8 | NEURL1B | -1.843037 | 0.6214382 | -0.943124 | 0.6688577 | -0.033432 | 0.6828329 | -0.756756 | 0.4919368 | 1.557017 | 0.02162527 | 1.744227 | 0.9264343 |
| ENSG00000214517.8 | PPME1 | 0.086618 | 0.8587992 | 0.375068 | 0.9668554 | 0.629732 | 0.002073404 | 0.306655 | 0.4919368 | 1.100725 | 0.3978836 | 0.526487 | 0.9663203 |
| ENSG00000214655.10 | ZSWIM8 | -1.722951 | 0.6792949 | 0.154218 | 0.6527736 | -0.725906 | 0.03170797 | 1.245638 | 0.3901988 | 1.968277 | 0.3005108 | 1.163196 | 0.9147227 |
| ENSG00000214706.10 | IFRD2 | 1.516463 | 0.2821482 | 0.495338 | 0.6294699 | -1.228101 | 0.007036002 | 0.447265 | 0.9534362 | -0.21217 | 0.7867303 | -0.339175 | 0.841154 |
| ENSG00000214753.2 | HNRNPUL2 | -1.361215 | 0.6136506 | -0.697739 | 0.05691035 | -0.718284 | 0.000107016 | 1.17757 | 3.30E-11 | 0.686136 | 0.01690028 | -0.60948 | 0.3058061 |
| ENSG00000214756.7 | METTL12 | -0.014 | 0.6214382 | -0.126616 | 0.7526603 | -0.216871 | 0.000205713 | 0.09342 | 0.8519291 | 1.315653 | 0.06557358 | -0.210376 | 0.8479431 |
| ENSG00000214941.7 | ZSWIM7 | -0.642605 | 0.6214382 | -0.337605 | 0.5953732 | -0.287048 | 0.2616845 | -0.231986 | 0.2948808 | -1.027997 | 0.02125501 | 0.146001 | 0.6829816 |
| ENSG00000215193.12 | PEX26 | -0.311666 | 0.7703009 | -0.726912 | 0.3454639 | 0.100103 | 0.8758996 | 1.560578 | 2.25E-05 | 1.164345 | 0.2550605 | 0.039678 | 0.9014975 |
| ENSG00000215252.11 | GOLGA8B | -0.058668 | 0.9047689 | -0.210696 | 0.8703522 | 0.314314 | 0.8095917 | 2.701748 | 0.02147017 | 0.407588 | 0.7944207 | -0.114705 | 1 |
| ENSG00000215301.9 | DDX3X | 0.248855 | 0.6616751 | -0.08464 | 0.714583 | 0.874435 | 0.2171616 | 0.610178 | 0.003677139 | 1.358989 | 0.05508623 | 0.94806 | 1 |
| ENSG00000217555.12 | CKLF | 0.475031 | 0.7817523 | 0.735435 | 0.6527736 | -0.889316 | 4.12E-06 | 0.680169 | 0.6371865 | -0.062817 | 0.1336832 | 1.738784 | 0.9377765 |
| ENSG00000218891.4 | ZNF579 | 1.870444 | 0.6014343 | -0.336324 | 0.7211571 | -1.278771 | 0.4599581 | 1.07207 | 0.04997317 | 2.467656 | 0.04490897 | 0.929804 | 0.6885065 |
| ENSG00000220201.7 | ZGLP1 | 0.630026 | 1 | 1.722602 | 0.3227388 | 0.550896 | 0.8106155 | 1.623398 | 0.1097309 | 1.597447 | 0.01491143 | 1.802348 | 0.6885065 |
| ENSG00000220205.8 | VAMP2 | -1.21597 | 0.00311427 | 0.220805 | 0.5813882 | -0.246956 | 0.1392205 | 0.48453 | 0.8121812 | -0.017461 | 0.5196843 | -0.662659 | 0.8564155 |
| ENSG00000221823.10 | PPP3R1 | 0.499287 | 0.6214382 | 0.344785 | 0.8703522 | 1.599655 | 0.1306428 | 1.122387 | 0.000911272 | 1.42968 | 0.04251491 | -0.379243 | 0.7478109 |
| ENSG00000221866.9 | PLXNA4 | -1.420259 | 0.6214382 | -0.496373 | 0.6527736 | 0.28453 | 0.9734049 | 1.431668 | 0.003680048 | -0.627731 | 0.8172994 | -0.648288 | 0.8949489 |
| ENSG00000221909.2 | FAM200A | 1.656184 | 0.8006286 | 0.060152 | 0.7881421 | 0.350141 | 0.8100781 | 0.556121 | 0.0484318 | 1.21743 | 0.6418821 | -0.586026 | 1 |
| ENSG00000221914.8 | PPP2R2A | -0.911254 | 0.6589361 | -0.251415 | 0.1276613 | 0.542551 | 7.50E-05 | 0.601636 | 0.4962484 | 0.133305 | 0.5196843 | 0.392182 | 0.6784855 |
| ENSG00000221946.7 | FXYD7 | -0.585374 | 0.8132872 | -1.403592 | 0.7881421 | -0.861756 | 0.5290845 | 2.477338 | 0.009744773 | 0.33169 | 0.9403512 | -1.754748 | 0.7990826 |
| ENSG00000221978.11 | CCNL2 | 0.929623 | 0.563505 | 0.01563 | 0.7273234 | 0.149279 | 0.7437308 | 1.08759 | 1.65E-05 | 2.23071 | 1.31E-10 | 1.495798 | 0.8973038 |
| ENSG00000221983.7 | UBA52 | 0.142166 | 0.6214382 | 0.002552 | 0.8695649 | -0.176968 | 0.001997044 | 0.268089 | 0.5860824 | 0.160623 | 0.2226201 | -0.117832 | 0.4362234 |
| ENSG00000223496.2 | EXOSC6 | 1.079021 | 0.2523209 | 0.290516 | 0.8961834 | 0.077395 | 0.8063725 | 0.07468 | 0.002454634 | 1.504028 | 0.1193335 | -0.582886 | 0.2936114 |
| ENSG00000223547.9 | ZNF844 | -1.618926 | 0.6214382 | -0.377469 | 0.9798987 | 0.943157 | 0.1595644 | 1.083168 | 0.003246221 | 0.971147 | 0.5921002 | -1.298679 | 0.9539154 |
| ENSG00000224877.3 | C17orf89 | 0.10782 | 1 | -0.162312 | 0.3368845 | -2.669495 | 2.02E-29 | -0.491821 | 0.311428 | 0.277838 | 0.6705976 | 0.629492 | 0.3094676 |
| ENSG00000225921.6 | NOL7 | 0.069135 | 0.7833919 | 0.178295 | 1 | -0.197704 | 0.001268807 | 0.195779 | 0.5141053 | -0.266554 | 0.02228536 | 0.196481 | 0.348294 |
| ENSG00000227500.9 | SCAMP4 | 0.903991 | 0.6214382 | 0.452862 | 0.8263112 | -0.908222 | 0.7887553 | 0.732775 | 0.001643361 | 2.559046 | 0.7319903 | 1.09707 | 0.9725802 |
| ENSG00000228300.13 | C19orf24 | 0.836554 | 0.6214382 | 0.147303 | 0.9597479 | -1.938204 | 1.73E-12 | -0.284661 | 0.003493666 | 0.370172 | 0.9110564 | -1.916995 | 0.5066477 |
| ENSG00000228474.5 | OST4 | 0.544018 | 0.2374782 | 0.11956 | 0.8263112 | -0.625585 | 2.25E-12 | -0.192522 | 0.001450825 | -0.261564 | 0.003546663 | 0.826202 | 0.8479431 |
| ENSG00000228696.8 | ARL17B | 2.769628 | 1 | 0.126193 | 0.9104976 | 2.321485 | 0.01722079 | -2.60785 | 0.3215575 | -0.355472 | 0.621723 | inf | 0.6185816 |
| ENSG00000228716.6 | DHFR | 0.845562 | 0.6793885 | -0.458436 | 0.4433091 | 0.868269 | 0.6608038 | 0.962604 | 0.001150126 | 0.30584 | 0.9592424 | -0.684426 | 0.6885065 |
| ENSG00000229117.8 | RPL41 | 0.070086 | 0.3077195 | 0.096414 | 0.48118 | -0.127916 | 0.01437159 | 0.062001 | 0.6556769 | -0.107248 | 0.03626519 | -0.097192 | 0.5733051 |
| ENSG00000229833.9 | PET100 | 1.08982 | 0.04692741 | 0.300276 | 0.6527736 | -1.571207 | 2.00E-19 | 0.024211 | 0.320553 | 0.091485 | 0.6614237 | 0.473653 | 0.8479431 |
| ENSG00000230124.6 | ACBD6 | -0.609073 | 0.6014343 | -0.121925 | 0.5010139 | -0.324651 | 0.3533539 | -0.45103 | 0.000187941 | 0.067078 | 0.2744651 | 1.162521 | 0.7997982 |
| ENSG00000230989.6 | HSBP1 | 0.801814 | 0.04862915 | 0.00406 | 0.8263112 | -0.022987 | 0.8100781 | -0.09358 | 0.5630962 | 0.199585 | 0.2589228 | 0.292863 | 1 |
| ENSG00000231500.6 | RPS18 | -0.62949 | 0.6744243 | 0.145001 | 0.3231727 | 0.068831 | 0.1052211 | -0.152663 | 0.004513611 | -0.433374 | 5.33E-10 | -0.169942 | 0.1567333 |
| ENSG00000231925.11 | TAPBP | 0.642395 | 0.5788332 | -0.472712 | 0.01315349 | -1.094818 | 1 | 0.474882 | 0.2335397 | 0.034721 | 0.9790948 | 0.140009 | 0.8739809 |
| ENSG00000232112.3 | TMA7 | -0.157696 | 0.9723935 | 0.050948 | 0.9771402 | -0.507894 | 4.16E-22 | -0.214508 | 0.000353134 | -0.048419 | 0.01464417 | -0.231908 | 0.03724323 |
| ENSG00000232388.4 | LINC00493 | 0.140136 | 0.6214382 | 0.18609 | 0.7487049 | -0.809581 | 1.03E-10 | -0.721744 | 1.06E-05 | -0.058996 | 0.01717712 | 0.374614 | 0.6885065 |
| ENSG00000232838.3 | PET117 | -0.668853 | 0.8738443 | -0.307544 | 0.7791094 | -0.368506 | 0.01610187 | 1.016734 | 0.2141464 | 0.579002 | 0.6816292 | 2.488466 | 0.6885065 |
| ENSG00000233276.3 | GPX1 | 0.636752 | 0.2716673 | 0.045595 | 0.8853411 | -0.289699 | 0.000142765 | -0.096591 | 0.01642572 | -0.233257 | 0.03099545 | 0.369943 | 0.863256 |
| ENSG00000233927.4 | RPS28 | -0.275263 | 0.624814 | -0.003809 | 0.9691339 | -0.197472 | 0.001345929 | -0.101836 | 0.03036606 | -0.178679 | 0.000719629 | -0.13929 | 0.3532606 |
| ENSG00000233954.6 | UQCRHL | -0.343299 | 1 | 1.196915 | 0.6527736 | -0.935835 | 4.84E-12 | -0.188603 | 0.6934365 | 2.013793 | 0.000814998 | 1.112334 | 0.6885065 |
| ENSG00000234127.8 | TRIM26 | 0.21257 | 0.9453571 | 1.167121 | 0.6527736 | 0.253893 | 0.8683055 | 1.020383 | 0.01947567 | 1.891512 | 0.4874815 | 0.324523 | 0.5915825 |
| ENSG00000234545.7 | FAM133B | 0.233198 | 0.7611292 | -0.187867 | 0.6200225 | 1.104306 | 2.12E-06 | 0.85484 | 0.06036412 | 1.027955 | 0.3368835 | 0.002595 | 0.4706214 |
| ENSG00000234745.9 | HLA-B | -0.902058 | 0.7736255 | 0.362628 | 2.45E-15 | -0.479039 | 3.06E-10 | 0.245655 | 0.8872299 | -0.699422 | 0.002564532 | 0.805252 | 0.4589582 |
| ENSG00000235162.8 | C12orf75 | -0.212799 | 0.9047689 | -0.853843 | 0.04760656 | -0.02677 | 0.03507147 | 1.608606 | 0.1926059 | 0.721229 | 0.1037881 | 1.697273 | 0.9617345 |
| ENSG00000237440.8 | ZNF737 | -0.845886 | 1 | -0.355056 | 0.9104976 | 3.230049 | 0.002372301 | -0.084774 | 0.6996942 | -1.567533 | 0.6134399 | inf | 0.6885065 |
| ENSG00000238083.7 | LRRC37A2 | -4.073768 | 0.1341967 | -2.622499 | 0.4487482 | 1.822117 | 0.05638606 | 0.393662 | 0.04730658 | 1.182904 | 0.5196843 | -4.958225 | 0.5733051 |
| ENSG00000239521.7 | GATS | -1.18138 | 0.7736255 | -0.506997 | 0.9517299 | 0.196671 | 0.6215927 | 0.102632 | 0.04970658 | 0.868949 | 0.01506509 | -0.152449 | 0.6829816 |
| ENSG00000239672.7 | NME1 | -0.460121 | 0.6214382 | -0.007561 | 0.6527736 | -0.318455 | 6.30E-08 | 0.324998 | 0.5057416 | -0.469157 | 0.000281589 | -0.36004 | 0.01138072 |
| ENSG00000240065.7 | PSMB9 | -1.587093 | 1 | -0.108974 | 0.853835 | -0.724625 | 1.88E-06 | -0.801609 | 0.06536764 | 0.48486 | 0.5778808 | 1.426103 | 0.7603387 |
| ENSG00000240184.6 | PCDHGC3 | -1.090846 | 0.529676 | -0.318268 | 0.821929 | 0.807769 | 0.000484397 | -1.506997 | 0.02946863 | -1.991901 | 0.08783911 | 2.000265 | 0.8479431 |
| ENSG00000240230.5 | COX19 | -2.309913 | 0.2423804 | -0.289877 | 0.5394427 | -0.938814 | 4.40E-05 | 1.130766 | 0.1313631 | 0.325563 | 0.3931837 | -1.652687 | 0.8479431 |
| ENSG00000240694.8 | PNMA2 | 0.297771 | 0.9309704 | 0.446505 | 0.8824535 | 0.539782 | 0.000320027 | 1.031677 | 0.305163 | 0.099186 | 0.738195 | 0.769213 | 0.9699068 |
| ENSG00000240972.1 | MIF | 1.155507 | 3.41E-14 | 0.626192 | 0.008910742 | -1.176863 | 3.80E-19 | -0.450331 | 3.14E-09 | 0.039472 | 0.2221967 | 0.270444 | 0.6885065 |
| ENSG00000241343.9 | RPL36A | -0.838908 | 0.005603822 | 0.38576 | 0.121014 | -0.866128 | 1.51E-27 | 1.064011 | 1.35E-15 | -0.350495 | 3.95E-07 | -0.377117 | 0.01514169 |
| ENSG00000241370.5 | RPP21 | 2.706818 | 8.74E-05 | -0.578552 | 0.8468859 | -1.357193 | 2.40E-05 | 0.77378 | 0.7051646 | -0.674704 | 0.621723 | -4.722409 | 0.3532606 |
| ENSG00000241468.7 | ATP5J2 | 0.910616 | 0.03568534 | -0.016466 | 0.9848758 | -0.911805 | 6.06E-30 | 0.065046 | 0.3467508 | -0.630607 | 1.23E-07 | 0.352731 | 0.6572299 |
| ENSG00000241837.6 | ATP5O | 0.14885 | 0.6214382 | -0.140025 | 0.5010139 | 0.271505 | 0.07116515 | 0.129546 | 0.2906903 | -0.447116 | 0.0001059 | 0.619151 | 0.5296752 |
| ENSG00000241878.9 | PISD | 0.511518 | 0.5527656 | -0.342408 | 0.6527736 | -0.022883 | 0.9408856 | -0.202707 | 0.04803716 | 0.467924 | 1 | 0.647238 | 0.7303122 |
| ENSG00000242265.5 | PEG10 | -0.646465 | 0.6214382 | -0.367499 | 0.4518214 | 0.692437 | 0.00092849 | 1.305717 | 0.003202048 | -0.611373 | 3.68E-05 | 0.718837 | 0.4301971 |
| ENSG00000242372.6 | EIF6 | 0.311079 | 0.6792949 | -0.273012 | 0.6527736 | 0.02299 | 0.0821772 | 0.239367 | 0.1208866 | -0.325681 | 0.003550296 | 0.703925 | 0.8479431 |
| ENSG00000242485.5 | MRPL20 | -0.176352 | 0.5791794 | -0.055796 | 0.784167 | -0.145832 | 0.000217556 | -0.023757 | 0.0163289 | 0.010316 | 0.2913839 | -0.437615 | 0.3082307 |
| ENSG00000242802.6 | AP5Z1 | -0.407824 | 0.6214382 | 0.152864 | 0.91418 | -0.249761 | 0.002274839 | 1.468854 | 0.1783688 | -0.118488 | 0.963643 | 1.237612 | 0.8317638 |
| ENSG00000243147.7 | MRPL33 | -0.379248 | 0.6214382 | 0.006962 | 0.5222411 | -0.244317 | 0.002039559 | -0.096116 | 0.01594454 | -0.067138 | 0.3403243 | -0.300595 | 0.5601856 |
| ENSG00000243156.7 | MICAL3 | -1.205576 | 0.6214382 | -0.210413 | 0.8217443 | 0.563639 | 0.5563493 | 2.389633 | 0.3467508 | 1.181944 | 0.01432977 | -0.861426 | 0.9264343 |
| ENSG00000243279.3 | PRAF2 | -0.532243 | 0.3132333 | 0.063835 | 0.7156572 | -0.097377 | 0.003793729 | 1.20837 | 0.05317647 | 0.335113 | 0.1206129 | -0.100694 | 0.6885065 |
| ENSG00000243317.7 | C7orf73 | -0.128303 | 0.9289591 | 0.063598 | 0.9801261 | -0.111517 | 0.03909694 | 0.812477 | 0.9689579 | 0.142275 | 1 | 0.959415 | 0.9517616 |
| ENSG00000243449.6 | C4orf48 | 0.396339 | 0.000741602 | -0.476198 | 0.04799695 | -4.863919 | 1.61E-38 | -0.380665 | 0.000861715 | 0.881085 | 0.6489484 | -0.632718 | 0.1598778 |
| ENSG00000243477.5 | NAT6 | -0.790718 | 0.8006286 | 0.236583 | 0.6527736 | 0.543109 | 1 | -2.008131 | 0.03968708 | 3.593938 | 0.3255801 | -0.48635 | 0.9147227 |
| ENSG00000243678.11 | NME2 | -0.229787 | 0.6214382 | -0.028196 | 0.7274222 | 0.540341 | 0.01424104 | -0.336783 | 8.97E-05 | 1.37643 | 0.001624447 | -0.547283 | 0.4623015 |
| ENSG00000243716.10 | NPIPB5 | -0.845712 | 0.6214382 | 0.172573 | 1 | 0.886126 | 0.000920343 | 2.185205 | 5.55E-07 | 3.428756 | 0.05001113 | 1.320846 | 0.9531804 |
| ENSG00000243943.9 | ZNF512 | 0.550954 | 0.8587992 | -0.50604 | 0.6527736 | -1.199839 | 0.6928639 | 2.051443 | 0.001320529 | 0.52059 | 0.6965089 | 1.495454 | 0.7767294 |
| ENSG00000244005.12 | NFS1 | -1.714139 | 0.714717 | 0.881272 | 0.6527736 | -0.471942 | 0.03164635 | -0.218234 | 0.3467508 | 0.301108 | 0.6689424 | -1.793048 | 0.6885065 |
| ENSG00000244165.1 | P2RY11 | -2.602533 | 0.09877095 | 0.707966 | 0.1334054 | -0.884484 | 0.2802304 | 1.626238 | 0.00025815 | 2.728819 | 0.0619892 | -1.264298 | 0.5814532 |
| ENSG00000244405.7 | ETV5 | -0.126744 | 0.8237387 | -0.209256 | 0.8653057 | -0.217767 | 0.03849615 | 0.254214 | 0.9567398 | 1.539359 | 0.2880317 | 0.54398 | 0.6885065 |
| ENSG00000244462.7 | RBM12 | -0.396496 | 0.6214382 | 0.067594 | 0.7213677 | 0.426374 | 0.2431375 | 2.485687 | 0.005400628 | 1.474331 | 0.08980341 | -0.879892 | 0.868568 |
| ENSG00000244486.7 | SCARF2 | -0.794772 | 0.714717 | -0.182464 | 0.6527736 | -0.567076 | 0.4183778 | 0.820768 | 0.1166342 | 1.858842 | 0.00014889 | -0.605758 | 0.9205794 |
| ENSG00000244754.8 | N4BP2L2 | -0.239843 | 0.7640133 | 0.889566 | 0.5953732 | 0.844778 | 1.07E-05 | 0.588974 | 0.2499795 | 0.77378 | 0.8021966 | -0.456583 | 0.8479431 |
| ENSG00000247077.6 | PGAM5 | 0.444554 | 0.59287 | -0.047018 | 0.2802741 | -0.429802 | 0.3276184 | 1.659331 | 0.01850972 | 2.054784 | 0.006149094 | -0.736771 | 0.6767422 |
| ENSG00000247315.3 | ZCCHC3 | -1.547649 | 0.5711449 | 0.063967 | 0.8899439 | 0.008729 | 0.4424178 | 0.458896 | 0.02312534 | 1.092554 | 0.1661214 | 2.284199 | 0.6829816 |
| ENSG00000248487.8 | ABHD14A | -0.917834 | 0.9473277 | -0.020883 | 0.95708 | -0.68903 | 0.04239485 | 0.02298 | 0.8224647 | 0.019174 | 0.6816292 | -2.745023 | 0.3746505 |
| ENSG00000249141.1 | RP11-514O12.4 | 0.576355 | 1 | -0.412997 | 0.9329105 | -1.578994 | 1 | -0.785451 | 0.4919368 | 1.667423 | 1 |  |  |
| ENSG00000249437.7 | NAIP | -1.227713 | 0.7736255 | 0.064754 | 0.8703522 | 0.769788 | 0.3983857 | 0.969564 | 0.01023944 | 1.171739 | 0.005274811 | -0.416222 | 1 |
| ENSG00000249459.8 | ZNF286B | -0.179893 | 0.8783629 | 1.09335 | 0.5953732 | 1.175929 | 0.01401688 | -0.120994 | 0.5792659 | 1.595806 | 0.234333 | -0.833515 | 0.6829816 |
| ENSG00000249915.7 | PDCD6 | -0.665235 | 0.5084093 | 0.745455 | 0.04943722 | -0.056556 | 0.005574707 | 0.352359 | 1 | 1.087434 | 0.9968301 | 0.454738 | 0.8800291 |
| ENSG00000249992.1 | TMEM158 | -0.382381 | 0.4764974 | 0.142051 | 0.5010139 | -0.56886 | 0.000206498 | 1.084554 | 2.50E-07 | 1.042507 | 0.4862573 | -0.526744 | 0.03721295 |
| ENSG00000250479.8 | CHCHD10 | -1.160667 | 0.529676 | -0.251788 | 0.1505221 | -0.75304 | 0.001103149 | -0.020314 | 0.4294044 | 0.385923 | 0.7115116 | 1.811511 | 0.7100152 |
| ENSG00000253506.2 | NACA2 | 0.560013 | 0.02326877 | 1.440912 | 0.004626623 | -1.799853 | 1.29E-19 | -0.522745 | 0.8028893 | -0.739851 | 0.2573878 | -0.413745 | 0.9016395 |
| ENSG00000253626.3 | EIF5AL1 | 2.212802 | 0.001448711 | 1.442347 | 0.001504246 | -2.242409 | 5.45E-28 | -0.033188 | 0.6051264 | 1.722836 | 0.2880317 | -0.579282 | 0.8479431 |
| ENSG00000253729.7 | PRKDC | -1.012997 | 0.6214382 | -0.381423 | 0.08888279 | 0.93018 | 5.42E-07 | 1.843877 | 7.08E-07 | 0.343138 | 0.8942969 | 0.005733 | 0.6833517 |
| ENSG00000254505.9 | CHMP4A | -2.499643 | 0.4692975 | 0.313968 | 1 | -0.689802 | 0.03337556 | 2.520709 | 0.3099929 | 0.76858 | 0.9278532 | 2.655101 | 0.926878 |
| ENSG00000254598.2 | CSNK2A3 | -1.198671 | 0.8265276 | 0.374747 | 0.6527736 | -0.760705 | 0.4638867 | -0.130786 | 0.160631 | -2.363664 | 0.03043237 | 1.839795 | 0.9597711 |
| ENSG00000254726.2 | MEX3A | 0.600379 | 0.5711449 | -0.176361 | 0.6527736 | -0.091535 | 0.5290845 | 1.703514 | 8.93E-07 | 2.232922 | 2.79E-13 | 0.00466 | 0.8949489 |
| ENSG00000254999.3 | BRK1 | 0.041432 | 0.7736255 | 0.017653 | 0.6527736 | 0.131823 | 0.4370871 | -0.162826 | 0.000207128 | -0.505085 | 0.1209675 | -0.107405 | 0.5344421 |
| ENSG00000255302.3 | EID1 | -2.427358 | 0.1253248 | 0.090238 | 0.6527736 | 0.331371 | 0.003248071 | 0.041033 | 0.01791766 | -0.247554 | 3.58E-07 | -0.299982 | 0.7767294 |
| ENSG00000255823.2 | MTRNR2L8 | 2.378973 | 0.4764974 | 1.272902 | 0.4011918 | 3.837618 | 0.009429051 | 3.616317 | 0.00187277 | 1.030327 | 0.2364543 | 3.190079 | 0.8949489 |
| ENSG00000256060.2 | TRAPPC2P1 | 1.724554 | 0.5692871 | -0.742982 | 0.7483725 | -0.696387 | 0.002416155 | -0.680416 | 0.2982077 | -1.348044 | 0.0963637 | 0.681761 | 0.7767294 |
| ENSG00000256087.6 | ZNF432 | -2.817766 | 0.6214382 | 0.346116 | 0.6527736 | 0.827425 | 0.3993016 | 0.974952 | 0.02412601 | 1.937236 | 0.09667266 | -0.532191 | 0.9147227 |
| ENSG00000256463.8 | SALL3 | -2.674548 | 0.6214382 | 1.09286 | 0.9640934 | -0.31843 | 0.6275552 | 2.143888 | 0.01631219 | 1.256697 | 0.5161226 | 1.794108 | 0.960853 |
| ENSG00000256618.2 | MTRNR2L1 | 1.399491 | 3.60E-20 | 1.725073 | 0.001800034 | -0.290384 | 0.6470059 | 2.940387 | 1.45E-08 | 5.366265 | 0.01007426 | -2.893795 | 0.6885065 |
| ENSG00000257093.6 | KIAA1147 | 0.444944 | 1 | -0.246271 | 0.5394427 | 1.548138 | 0.01775551 | -0.114381 | 0.3147727 | 1.973277 | 0.180165 | 0.254651 | 0.9539154 |
| ENSG00000257218.5 | GATC | 0.697674 | 0.9470141 | -0.334129 | 0.415753 | 0.78387 | 0.8443322 | 1.212844 | 4.14E-07 | 1.402333 | 0.02424787 | -0.871452 | 0.4964698 |
| ENSG00000257704.3 | INAFM1 | -0.722746 | 0.714717 | -0.146774 | 0.1256222 | -2.006494 | 3.33E-10 | -0.106857 | 0.1052874 | 0.131026 | 0.9440398 | -0.083688 | 0.8363382 |
| ENSG00000257727.5 | CNPY2 | -0.033607 | 0.8006286 | 0.355232 | 0.2957998 | -0.704399 | 0.00015264 | 0.525589 | 0.6185149 | 0.008435 | 0.130167 | -1.034263 | 0.4981184 |
| ENSG00000257923.9 | CUX1 | 0.122596 | 0.8587992 | -0.103805 | 0.6775766 | 0.861796 | 0.6718295 | 1.277806 | 0.04890934 | 0.985168 | 0.1197239 | -1.065778 | 0.4028895 |
| ENSG00000258102.3 | MAP1LC3B2 | -0.601919 | 0.9282145 | 0.522799 | 0.5427944 | -1.930714 | 4.16E-10 | 1.949426 | 0.4919368 | -0.4638 | 1 | 4.397599 | 0.6885065 |
| ENSG00000258472.8 | RP11-192H23.4 | 0.620056 | 1 | -1.412253 | 0.7383998 | -2.919771 | 0.3144442 | 0.79069 | 0.03032508 | 0.888482 | 0.5196843 | -2.881603 | 0.6885065 |
| ENSG00000258839.2 | MC1R | -0.796947 | 0.714717 | 4.78076 | 0.8217443 | -1.458789 | 0.5290845 | 1.415508 | 0.000911448 | 3.055266 | 0.01406336 | -3.48238 | 0.6155125 |
| ENSG00000259956.1 | RBM15B | -1.181523 | 0.714717 | -0.532002 | 0.354537 | 0.118768 | 0.4721767 | 2.006488 | 0.01471784 | 0.726884 | 0.004902851 | -0.923395 | 1 |
| ENSG00000261115.5 | TMEM178B | -0.597423 | 1 | 0.772602 | 0.6527736 | 0.435339 | 0.1045708 | 1.305933 | 0.002024012 | 0.79079 | 0.9039452 | -0.476555 | 0.6885065 |
| ENSG00000261609.3 | GAN | -0.073106 | 1 | 0.314092 | 0.9341021 | 2.180168 | 0.002028948 | 1.441796 | 0.00045627 | 1.457901 | 0.06634352 | -0.94258 | 0.7767294 |
| ENSG00000262246.5 | CORO7 | 4.604129 | 0.3946735 | 1.009961 | 0.5953732 | -1.438987 | 0.4064599 | -0.373508 | 0.5435298 | 0.793484 | 0.004919348 | 3.59647 | 0.5733051 |
| ENSG00000262814.6 | MRPL12 | 0.160954 | 0.7598375 | -0.222799 | 0.4083382 | -0.839975 | 7.84E-06 | -0.179404 | 0.571042 | -0.360957 | 0.006387737 | -1.121884 | 0.6885065 |
| ENSG00000263001.5 | GTF2I | -0.980945 | 0.6014343 | -0.091289 | 0.8374851 | -0.130736 | 0.883719 | 3.158221 | 0.8308849 | 2.366777 | 0.003978748 | 0.866648 | 0.6885065 |
| ENSG00000263956.5 | NBPF11 | 0.860564 | 1 | 0.014274 | 0.6527736 | -0.323937 | 0.6603698 | 2.386863 | 0.009270402 | 1.342076 | 0.01766597 | -2.697917 | 0.6517664 |
| ENSG00000264364.2 | DYNLL2 | -3.70916 | 0.05012058 | -0.313638 | 0.6527736 | 0.491722 | 0.6628486 | 0.407267 | 0.01084756 | 0.183383 | 0.38864 | 0.38804 | 0.8654874 |
| ENSG00000265241.6 | RBM8A | -0.154624 | 0.7687347 | 0.117356 | 0.7647276 | 0.315898 | 0.4721767 | 0.215511 | 0.03449997 | 0.368076 | 0.1568461 | -0.385137 | 0.2321124 |
| ENSG00000265491.4 | RNF115 | -0.132077 | 0.8144419 | -0.095704 | 0.9720618 | 0.726165 | 0.6381738 | 1.297072 | 0.02260971 | 0.953984 | 0.08626379 | 1.75283 | 0.9738331 |
| ENSG00000265972.5 | TXNIP | 0.309534 | 0.714717 | -0.096541 | 0.6527736 | 1.377557 | 0.000596577 | -0.494036 | 0.009744773 | 1.13944 | 0.02599105 | 0.226781 | 1 |
| ENSG00000266028.7 | SRGAP2 | -0.644251 | 0.7736255 | -0.824765 | 0.1688282 | 0.511427 | 0.1992941 | -0.218286 | 0.1932203 | 0.74742 | 0.01243218 | 1.220682 | 0.7152543 |
| ENSG00000266074.8 | BAHCC1 | 1.56081 | 0.8006286 | 1.467299 | 0.8079089 | 0.816768 | 0.8145878 | 2.468546 | 0.1148418 | 2.624244 | 0.005747048 | -3.163392 | 0.5066477 |
| ENSG00000266472.5 | MRPS21 | 0.192822 | 0.4728584 | 0.014161 | 1 | -0.342601 | 1.86E-05 | -0.021875 | 0.03676124 | -0.493187 | 0.001807581 | 0.252811 | 0.278077 |
| ENSG00000267855.5 | NDUFA7 | -0.584674 | 0.7410745 | 0.561436 | 0.1476286 | -1.166611 | 2.13E-16 | 0.644724 | 0.1248679 | -0.966589 | 0.2640175 | -0.431925 | 0.9638718 |
| ENSG00000268043.7 | NBPF12 | -0.617395 | 0.9047689 | 0.781948 | 0.5738452 | 1.07277 | 0.2879524 | 2.107477 | 0.01245367 | 1.765499 | 0.1291534 | -3.258572 | 0.8479431 |
| ENSG00000269713.7 | NBPF9 | -0.329979 | 0.714717 | 0.292435 | 0.9676809 | 0.850267 | 0.0327941 | 0.382515 | 0.8512427 | 1.009511 | 0.6554785 | -2.493251 | 0.8931285 |
| ENSG00000269858.5 | EGLN2 | 1.656364 | 0.8006286 | 0.054977 | 0.8953649 | 0.274719 | 0.9124169 | 1.224317 | 0.0168188 | 0.425388 | 0.6003632 | -0.530679 | 0.9270338 |
| ENSG00000270170.1 | NCBP2-AS2 | 0.37912 | 0.529676 | 0.121659 | 0.7519994 | -0.828762 | 6.54E-07 | 0.342759 | 0.7211844 | 1.472433 | 0.5900158 | -0.13882 | 0.9671638 |
| ENSG00000271425.6 | NBPF10 | -0.554965 | 1 | 0.622794 | 0.6527736 | 0.477278 | 0.725917 | 0.897845 | 0.03364882 | 3.482416 | 0.02707314 | 3.022587 | 0.8479431 |
| ENSG00000272047.1 | GTF2H5 | -0.767336 | 0.6214382 | 0.032948 | 0.8070253 | -0.448979 | 0.001027852 | -0.035333 | 0.062815 | -0.409942 | 0.04251491 | 0.792088 | 0.8479431 |
| ENSG00000272196.2 | HIST2H2AA4 | -1.907053 | 0.9252683 | 0.014392 | 0.9222921 | -0.427608 | 0.01326601 | -0.63101 | 0.1357963 | -0.321376 | 0.2880317 | -0.09689 | 0.9147227 |
| ENSG00000272325.1 | NUDT3 | -0.667162 | 0.6214382 | -0.192256 | 0.3736565 | 0.599606 | 5.05E-05 | 1.286636 | 1.41E-05 | 0.567439 | 0.2648054 | 0.01578 | 0.6885065 |
| ENSG00000272333.5 | KMT2B | -1.61159 | 0.6169321 | 0.107575 | 0.9126089 | 0.091651 | 0.7446597 | 0.217039 | 0.01087494 | 1.059727 | 0.3514917 | -2.001773 | 0.9690114 |
| ENSG00000272391.5 | POM121C | -1.765802 | 0.1201494 | 0.244234 | 0.6527736 | 0.500175 | 0.7164952 | 1.075806 | 0.03362496 | 0.890651 | 0.5196843 | -0.884388 | 0.5553892 |
| ENSG00000272886.5 | DCP1A | 0.231753 | 0.6214382 | 0.825676 | 0.6340237 | 1.409996 | 0.000397816 | 1.837614 | 0.001466045 | 0.208183 | 0.3552415 | 1.213056 | 1 |
| ENSG00000273749.4 | CYFIP1 | 0.323274 | 0.7321786 | 0.206619 | 0.6200225 | 0.255361 | 0.01419177 | 1.330224 | 0.1143494 | 0.05644 | 0.5196843 | -0.643192 | 0.6885065 |
| ENSG00000273841.4 | TAF9 | 0.46071 | 0.8006286 | 0.4552 | 0.592336 | -0.217542 | 0.02841787 | 0.521473 | 0.5027003 | 0.758473 | 0.03081869 | -0.180887 | 0.9506884 |
| ENSG00000273899.4 | NOL12 | -0.772729 | 0.6214382 | 0.817695 | 0.8217443 | -0.51023 | 0.002068226 | 0.209305 | 0.4791525 | 0.837618 | 1 | -0.395835 | 1 |
| ENSG00000274211.4 | SOCS7 | 0.073513 | 0.8006286 | 0.388629 | 0.8593459 | -0.172837 | 0.7437308 | 0.95996 | 0.04764465 | 0.813949 | 0.2019818 | -0.755684 | 0.8178576 |
| ENSG00000275004.3 | ZNF280B | -2.722968 | 0.529676 | -3.529405 | 0.7881421 | 0.167252 | 0.4626116 | 0.747905 | 0.9069344 | -0.148917 | 0.920782 |  |  |
| ENSG00000275023.4 | MLLT6 | -0.632278 | 0.7823641 | 0.728373 | 0.6527736 | 0.083603 | 0.991425 | 1.309217 | 0.1857339 | 1.245788 | 0.00018356 | 1.570069 | 0.8704806 |
| ENSG00000275052.4 | PPP4R3B | -1.437405 | 0.682906 | -0.110564 | 0.9104976 | 0.716338 | 9.95E-05 | 0.277006 | 0.01579705 | 1.29103 | 0.2019818 | -1.799649 | 0.7494298 |
| ENSG00000275066.4 | SYNRG | -0.135929 | 0.8650222 | 0.099297 | 0.8677969 | 0.137575 | 0.238902 | 1.938646 | 0.03007978 | 0.489936 | 0.6481815 | 0.744658 | 0.834639 |
| ENSG00000275464.3 | CH507-9B2.5 | -0.276797 | 0.9348915 | 0.502592 | 0.4691942 | -1.613257 | 0.9139328 | 1.684184 | 0.9478836 | 0.342013 | 0.506562 |  |  |
| ENSG00000275832.4 | ARHGAP23 | 0.177848 | 0.6214382 | -1.536654 | 0.4719032 | 0.000488 | 0.3219588 | 0.443592 | 0.2450967 | 1.480649 | 0.006623006 | -1.620756 | 0.8986153 |
| ENSG00000276043.4 | UHRF1 | -1.917368 | 0.8587992 | -0.1334 | 0.7201934 | -0.343314 | 0.7510691 | 1.568625 | 0.001009941 | 0.676112 | 0.5781199 | -0.517999 | 0.7153537 |
| ENSG00000276293.4 | PIP4K2B | -1.150068 | 0.2006576 | -0.610754 | 0.2751093 | -0.587401 | 0.8271566 | 1.057276 | 0.01454436 | 0.949842 | 0.1119361 | 0.486073 | 1 |
| ENSG00000277149.4 | TYW1B | -1.088072 | 0.8006286 | 0.591354 | 1 | 4.218445 | 0.000140916 | -0.602045 | 1 | 1.508214 | 0.787129 | -0.83928 | 0.8949489 |
| ENSG00000277443.1 | MARCKS | -0.439157 | 0.6214382 | 0.097507 | 0.6527736 | 0.422719 | 0.00141721 | 1.127849 | 6.53E-11 | 1.071968 | 4.01E-08 | -0.279122 | 0.6885065 |
| ENSG00000277791.4 | PSMB3 | 0.51795 | 0.000114999 | 0.072542 | 0.8379413 | -0.125576 | 0.01671207 | 0.057561 | 0.04936461 | -0.077798 | 0.002410666 | -0.786589 | 0.03975463 |
| ENSG00000277972.1 | CISD3 | -0.278356 | 0.8006286 | 0.001699 | 0.6527736 | -0.900011 | 0.01776045 | -0.263795 | 1 | 1.21855 | 0.7491998 | 0.779122 | 1 |
| ENSG00000278053.4 | DDX52 | 0.620824 | 0.8294538 | 0.484808 | 0.8117341 | 0.705908 | 0.000614971 | 1.401902 | 0.02131841 | 1.591909 | 0.321864 | -0.94589 | 0.6559143 |
| ENSG00000278129.1 | ZNF8 | -1.522519 | 0.6792949 | 1.379761 | 0.5010139 | 0.391825 | 0.5279962 | 0.084481 | 0.92164 | 0.644011 | 0.04017399 | 6.886042 | 0.6517664 |
| ENSG00000278535.4 | DHRS11 | 1.014525 | 1 | 0.202592 | 0.690756 | 3.813887 | 0.6295125 | 0.93184 | 0.171072 | 2.936098 | 0.01243193 | 0.184732 | 0.9147227 |
| ENSG00000278540.4 | ACACA | -1.940865 | 0.5453725 | -0.86351 | 0.1957576 | 0.605705 | 0.1992941 | 1.881431 | 0.00133523 | 1.189732 | 0.007444469 | 0.435737 | 0.6885065 |
